# Cost-effectively dissecting the genetic architecture of complex wool traits in rabbits by low-coverage sequencing

**DOI:** 10.1101/2022.03.09.483689

**Authors:** Dan Wang, Kerui Xie, Yanyan Wang, Jiaqing Hu, Wenqiang Li, Qin Zhang, Chao Ning, Xinzhong Fan

## Abstract

Wool traits of rabbits are important in fiber production and model organism research on hair growth, while the genetic architecture remains obscure. In this study, we focused on wool characteristics in Angora rabbits, a well-known fiber breed. Balancing genotyping cost and variant detection, we proposed low-coverage whole genome sequencing (LCS) followed by genotype imputation for genotyping. Different genotype imputation strategies, sequencing coverages and sample sizes were compared, and we found by BaseVar + STITCH, genotyping reached high accuracy (>0.97) at a depth of 1.0X and a sample size > 300. Multivariate GWAS followed by conditional GWAS and confidence interval estimation of QTLs were used to reveal the genetic architecture of wool traits. Six QTLs were detected with phenotypic variation contribution ranging from 0.42% to 7.50%. Gene-level mapping implicated *FGF10* associated with fiber growth and diameter, which supported previous function research on fibroblast growth factor family in other species and provided genetic information for wool rabbit breeding. We suggest LCS as a cost-effective alternative for assessing common variants. GWAS combined with LCS can excavate QTLs and fine-map genes associated with quantitative traits. This study provides a powerful analysis mentality for investigating complex traits, which lays the foundation for genomic breeding.

## INTRODUCTION

Genome-wide association studies (GWAS) have delivered new insights into the biology and genetic architecture of complex traits. In the past decades, GWAS accelerated the rate of gene discovery to an unprecedented scale, identifying many replicated genetic variants associated with complex diseases and quantitative traits in livestock, plants, humans and model organisms (da Silva Xavier et al., 2013, Huang et al., 2017, Freebern et al., 2020, Qin et al., 2021). Phenotypic variations of complex traits are always caused by the cumulative effect of numerous common variants, *i*.*e*., polygenic, so high marker density GWAS could provide novel insights into the genomic architecture (Kainer et al., 2019). The traditional approach for high marker density requires two distinct genetic testing technologies: high coverage sequencing of whole genome and a genome-wide genotyping array followed by imputation. Considering the cost of population sequencing and the case of lacking in chip array, low-coverage whole genome sequencing (LCS) followed by imputation is a much more affordable alternative for assessing common genetic variants and testing the association of millions of variants (Loos, 2020). Furthermore, it has been proposed to increase the discovery power of trait-associated and/or causative genetic variants (Ros-Freixedes et al., 2017, Loos, 2020). At the present stage, LCS has been widely used to accurately assess common variants in GWAS. Studies showed that 0.5-1X LCS performed comparably to commonly used low-density GWAS arrays (Martin et al., 2021). LCS at a depth of 1X was able to find signals missed by standard imputation of SNP arrays (Gilly et al., 2016). A more systematic examination of the power of GWAS suggested that 1X LCS sequencing allows discovering up to twice as many associations as standard SNP array imputation (Gilly et al., 2019). LCS at a depth of ≥4X captured variants of all frequencies more accurately than all commonly used GWAS arrays investigated at a comparable cost (Martin et al., 2021).

The LCS approach (LCS followed by imputation) exploits the fact that individuals in the same cohort are sufficiently related to share large genome segments (Ros-Freixedes et al., 2017). Missing genotypes in LCS data are imputed using local linkage patterns to infer unknown genotypes in target samples from known genotypes. Current available tools for imputation of LCS data include STITCH (Davies et al., 2016), Beagle (Browning S.R. and B.L., 2007), GeneImp (Spiliopoulou et al., 2017), GLIMPSE (Rubinacci et al., 2021) and loimpute (Wasik et al., 2021). The methods are two typical ways to obtain imputed genotypes: with a haplotype reference panel and without reference panels. STITCH (Davies et al., 2016) imputes genotype based only on sequencing read data, without requiring additional reference panels or array data, and is applicable in settings of extremely low sequencing coverage (Liu et al., 2018a, Meier et al., 2021). The others are imputation tools based on reference panel information, for example, GLIMPSE phases and imputes LCS data using large reference panels (Rubinacci et al., 2021). In addition, Beagle is developed for genotype imputation tailored to work both with and without reference panels (Browning S.R. and B.L., 2007).

Since both library and sequencing costs decrease, LCS has become increasingly attractive for obtaining genotyping information of farm animals (Meier et al., 2021). Angora rabbits are well known farm animals for wool production. The economic value of Angora wool depends mainly on the texture of rabbit hair including fiber diameter, length and so on. In this study, we generated the accurate and dense genotypes of Angora rabbits with a cost-efficient LCS approach by demonstrating the imputation performance across five levels of sequencing coverages and six levels of sample sizes using three imputation strategies (BaseVar + STITCH, Bcftools + Beagle4 and GATK + Beagle5). To reveal the genetic architecture of complex wool traits in Angora rabbits, we performed GWAS of six important economic traits at various time points with high resolution. Furthermore, we developed a conditional GWAS and the drop (Δ) in log-transformed *P* values in multivariate linear mixed model to confirm confidence intervals of QTLs which aided in candidate genes identification in high-resolution.

## RESULTS

### LCS imputation pipeline

In order to accurately capture variants in the rabbit genome, we compared three genotyping pipelines using LCS data, and used the high-depth sequencing data results on chromosome 11 (chr11) as the gold standard for accuracy evaluation (Fig. 1). Chr11 was used because of its similar LD extent to the whole genome (shown in the results of genetic architecture below) and the medium length. Genotypic imputation of genetic variants was performed in the 600 rabbits with a down-sampled sequencing depth of 2X. Among the reference-panel-free methods, highly accurate genotypes were obtained using the pipeline BaseVar + STITCH with an average GC of 99.08% and an average GA of 0.98, while by neither Bcftools + Beagle4 nor GATK + Beagle5, GC didn’t exceed 95.73%, and GA didn’t exceed 0.88 (Fig. 2A-B, Table S1).

**Fig. 1.**
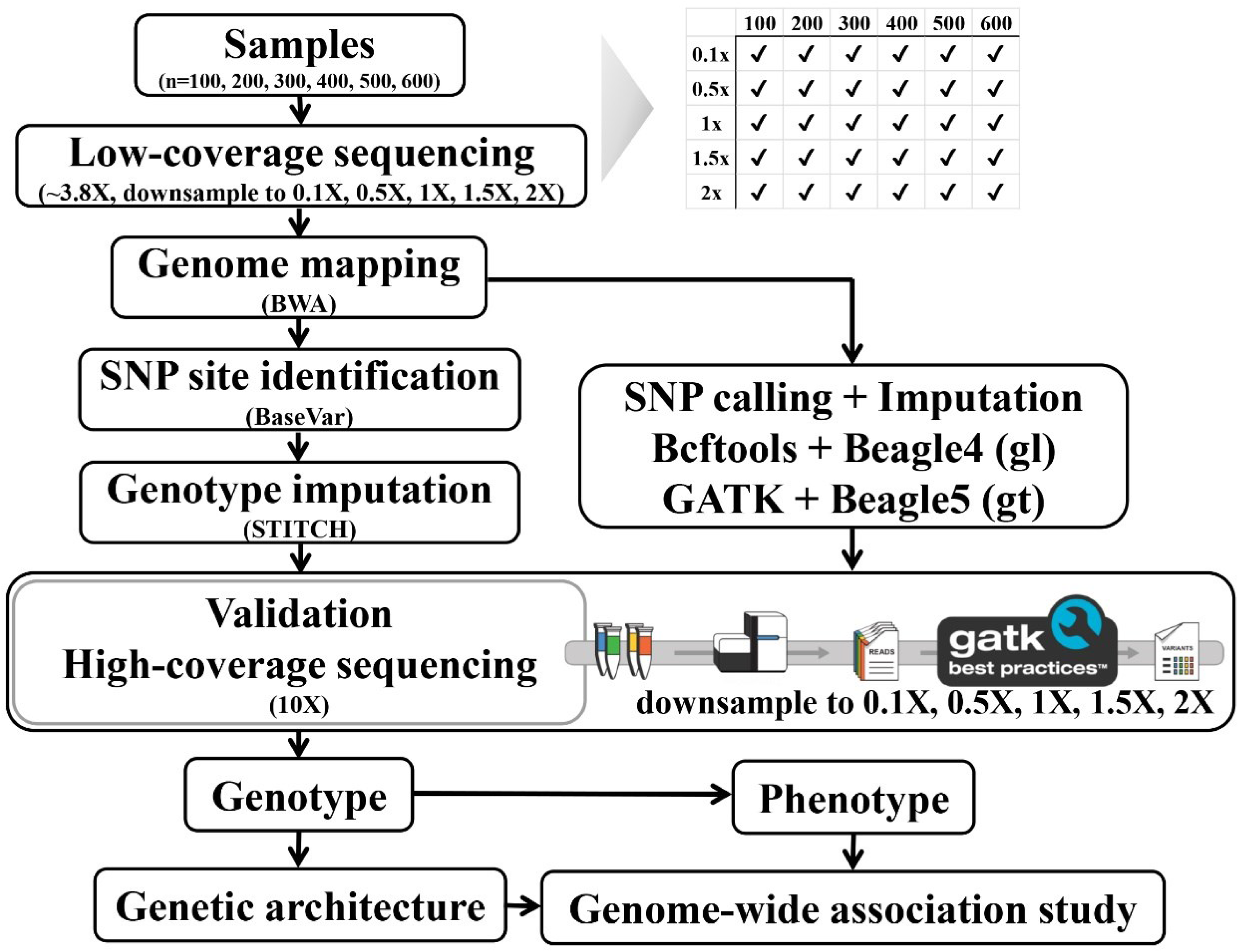
Analysis pipeline for low coverage sequence data and genetic architecture.

**Fig. 2.**
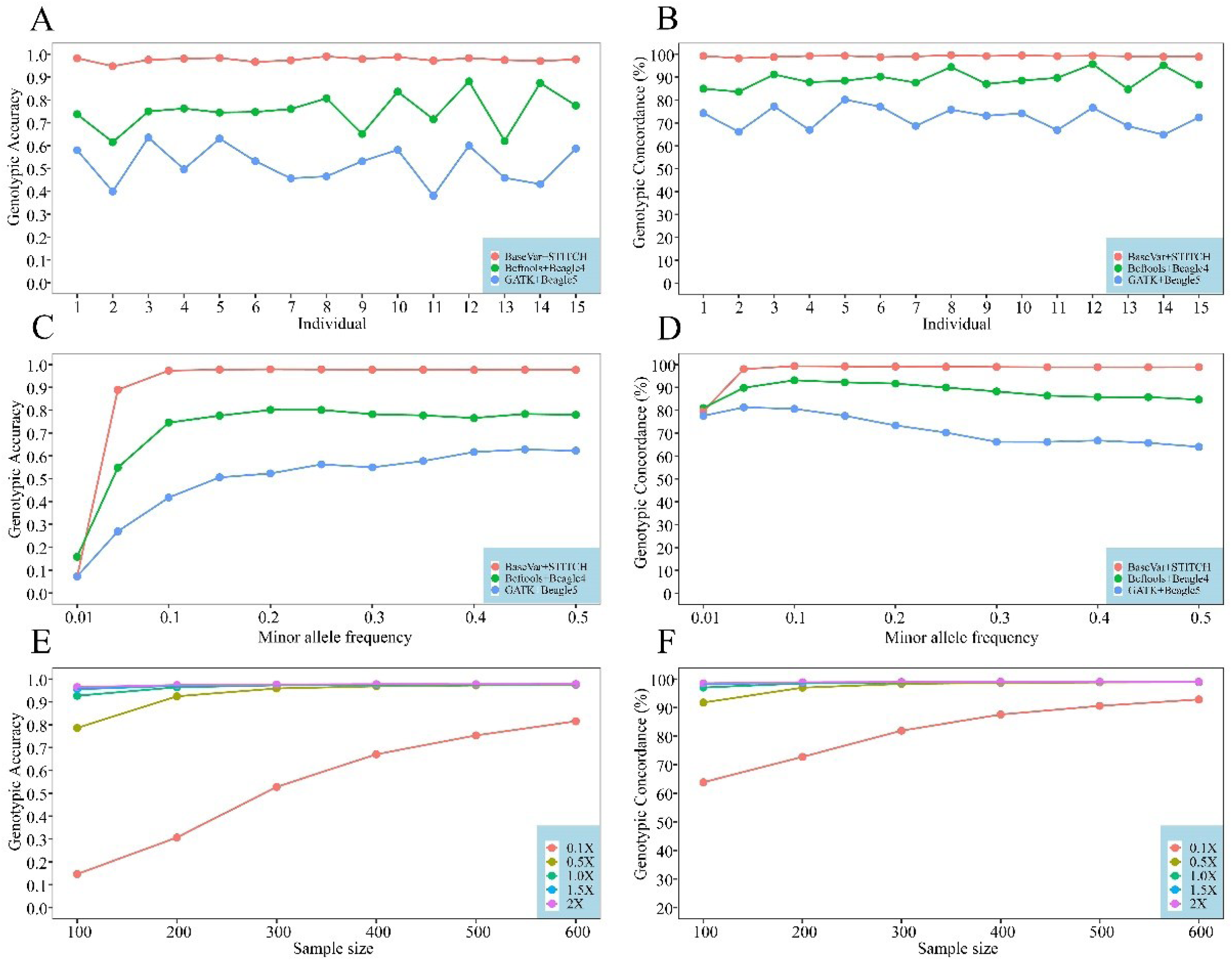
Genotype imputation performance compared among the three pipelines (red: BaseVar + STITCH, green: Bcftools + Beagle4 and blue: GATK + Beagle5) and different numbers of samples (100, 200, 300, 400, 500 and 600) and sequencing depths (0.1×, 0.5×, 1.0×, 1.5×, 2.0×) by genotypic concordance (the right ones) and genotypic accuracy (the left ones). A and B were imputation performance among the 15 individuals, C and D were among different minor allele frequency, and E and F were among different numbers of samples and sequencing depths.

The pattern of imputation performance in relation to minor allele frequency (MAF) was investigated among the pipelines. Using the pipeline BaseVar + STITCH, high and steady imputation accuracy (an average GC of 98.98% and GC ranging from 98.82% to 99.31%, an average GA of 0.98 and GA ranging from 0.97 to 0.98) was obtained for common variants with MAF ranging from 0.05 to 0.5. However, in the MAF range, imputation accuracy was a bit poor and greatly affected by MAF with GC waving from 84.66% to 93.10% and GA waving from 0.75 to 0.80 by Bcftools + Beagle4, and it was worse and more fluctuant by GATK + Beagle5 with GC waving from 64.02% to 80.60% and GA waving from 0.42 to 0.63. For SNPs with MAF lower than 0.05, both the genotypic accuracy and the genotypic concordance tended to decrease and were hugely affected by MAF (Fig. 2C-D, Table S2), showing the imputation accuracy of rare variants could be highly influenced by MAF. Based on the above results, the pipeline BaseVar + STITCH performed best and was used for the subsequent analyses.

### Effect of sample size and sequencing depth on imputation

In order to examine the influence of sample size and sequence coverage to imputation accuracy, we performed genotypic imputation by the pipeline BaseVar + STITCH with different numbers of samples (100, 200, 300, 400, 500 and 600) and sequencing depths (0.1X, 0.5X, 1.0X, 1.5X, 2.0X) in this population. As expected, genotypic concordance and genotypic accuracy generally increased as sample size and sequencing depth increased. Especially, when sample size increased from 100 to 300 and sequence coverage increased from 0.1X to 1.0X, the imputation accuracy was hugely improved. For the > 1X coverage, a sample size >300 had little effect on imputation performance, and showed to guarantee the credibility of genotyping (Fig. 2E-F, Table S3).

### Tagging SNPs

We retained 18,577,154 high-quality imputed SNPs by a two-step imputation using STITCH followed by Beagle and stringent quality control. The SNP density corresponded to 1 SNP per 150 bp in the rabbit genome. The variants were distributed uniformly across the genome (Fig. 3A). The majority of the identified SNPs were located in intergenic regions (57.78%) and intronic regions (35.50%). The exonic regions contained 0.52% of the SNPs. A total of 72,552 synonymous SNPs and 23,328 nonsynonymous SNPs of exons were identified, for a nonsynonymous/synonymous ratio of 0.32 (Table S4).

**Fig. 3.**
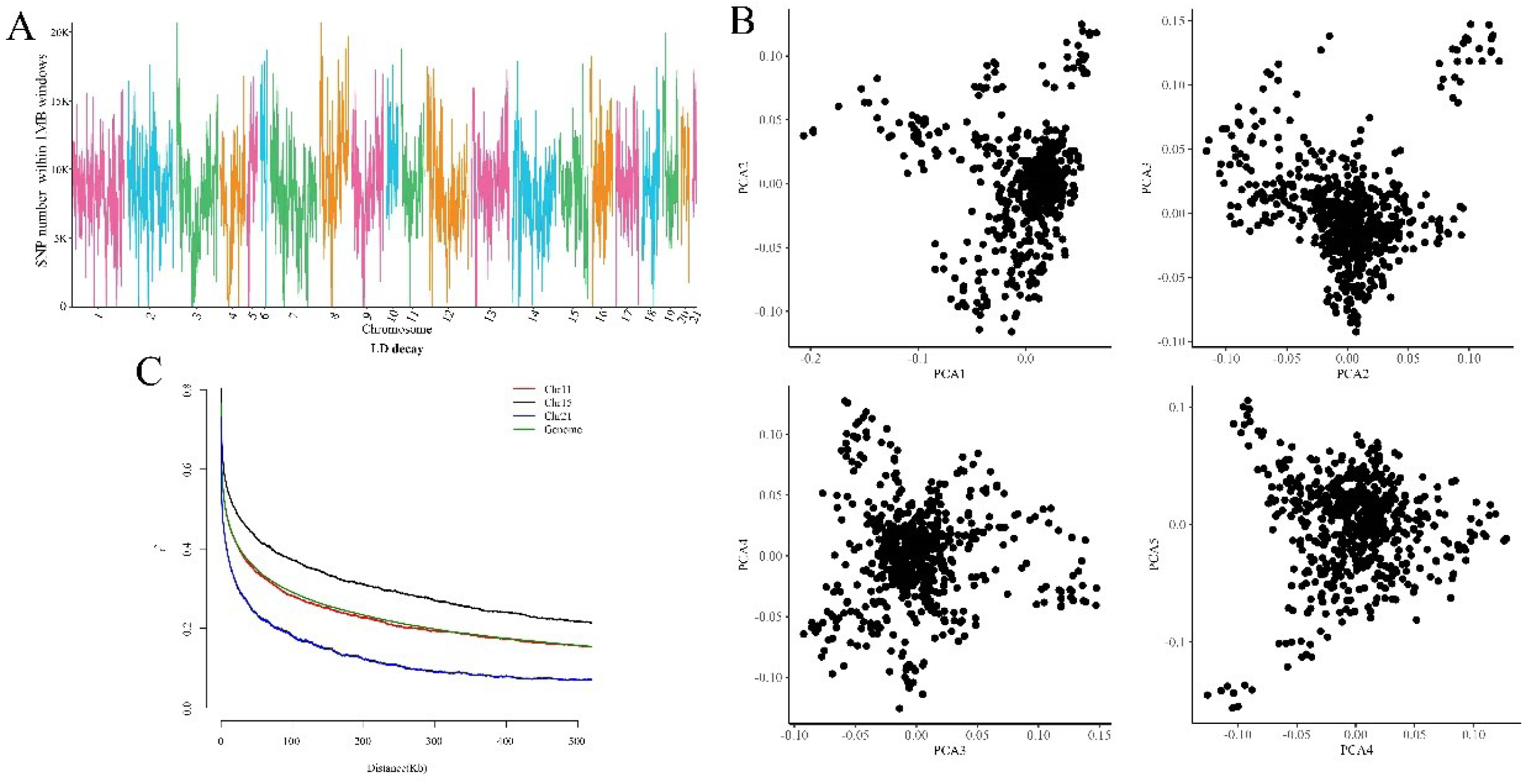
Genetic diversity of the Angora rabbit population. (A) Distribution of SNPs in 1-Mb windows across the genome. (B) Principal component analyses plotting the first to the fifth dimension. (C) The extent of linkage disequilibrium (LD), values were mean LD r2 values for all pairs of SNPs binned by distance. The slowest LD decay was observed for chr15, and the fastest was observed for chr21. Chr11 showed similar LD extent to the whole genome.

### Genetic architecture

The population structure of the 629 rabbits was assessed by performing PCA. The relationships between the first five principal components show no distinct evidence of population structure (Fig. 3B). LD analysis indicated that the physical distance between SNPs occurred at ∼6.5 kb (r^2^ = 0.50, Fig. 3C and Table S5). The average pairwise LD r2 values decreased to 0.16 at 500 kb and to 0.11 at 1 Mb. The distribution of r2 with respect to the physical distance for each chromosome was different. The slowest LD decay was observed for chr15, and the fastest was observed for chr21. Chr11 showed similar LD extent to the whole genome. Combining CLR and Pi analyses, we identified 151 potential selective-sweep regions overlapping with 309 candidate genes (Fig. S1, Table S6). The regions displayed significant overrepresentation of genes involved in immunity (*P* = 4.10E-12) and vitamin B6 metabolism (*P* = 1.30E-04) (Table S7). Immune system is one of the strongly targeted functions by natural selection during evolution because it serves as the backbone of defence against pathogens (Quintana-Murci, 2019, Barreiro and Quintana-Murci, 2020, Gerardo et al., 2020). Vitamin B6 is actively involved in protein metabolism as a catalyst in the body. It activates the enzymes and chemical reactions that start the metabolism of the hair proteins, keratin and melanin, in the hair follicles. This makes the hair follicles get enough keratin and melanin, which promotes hair growth and hair renewal. On the basis of clinical and trichological studies, vitamin B6 was revealed to induces improvement in the hair condition and reduce the hair loss (Brzezinska-Wcislo, 2001, D’Agostini et al., 2007). In addition, several regions involving in tryptophan, valine, leucine, isoleucine, nicotinate, nicotinamide, tyrosine and retinol metabolism showed selective signatures (Table S7).

In order to explore further detect genomic footprints of selection, 14 domesticated rabbits were sampled from the population to analyze genetic diversity and population structure by comparing to their wild progenitor, 14 wild rabbits (*Oryctolagus cuniculus*). A maximum-likelihood tree showed that the genotypes were classified into obvious two divergent groups (Fig. 4A). The PCA showed diversity among the rabbit genotypes with the first two principal components explaining 8.26% and 1.48% of the genetic variance, respectively (Fig. 4B). For the Angora population, all individuals were grouped together and showed a consistent genetic relationship, while for the wild population, the individuals were relative dispersive probably because of different geographical origins. What’s more, population structure was assessed for *K* values ranging from 1 to 5. the most significant change of likelihood occurred when K increased from 1 to 2 (Fig. 4C). Thus, the most likely value of *K* was 2. At *K* = 2, the two populations were separated from each other, and their genetic backgrounds are clearly significantly different. Such a partitioning of the population was consistent with significant delta *K* values (Fig. 4D). This was also in accordance with the maximum-likelihood tree (Fig. 4A). LD was calculated to provide information for population history. LD between markers decreased as physical distance between markers increased, and the degree of LD attenuation hugely differed between the two populations. From the current samples, the wild population exhibited an extremely rapid LD decay, indicating the high diversity of the wild ancestors. However, the Angora population showed a slow decay of LD, and markers separated by 350 kb showed r^2^ higher than 0.2, suggesting high inbreeding potentially due to artificial selection (Fig. 4E and Table S8). Furthermore, using the top 5% of *F*_ST_ values and *θ*_*π*_ ratio (cutoffs: *F*_ST_ > 1.87 and log_2_ (*θ*_*π*_ ratio (*θ*_*π* wild_/*θ*_*π* Angora_) ≥4.09), we identified 464 candidate domestication regions overlapping 775 genes under selection in the domestic rabbits (Fig. 4F, Table S9 and S10). The enrichment of genes was mainly involved in nervous system probably reflecting tameness and aggression during domestication (Carneiro et al., 2014).

**Fig. 4.**
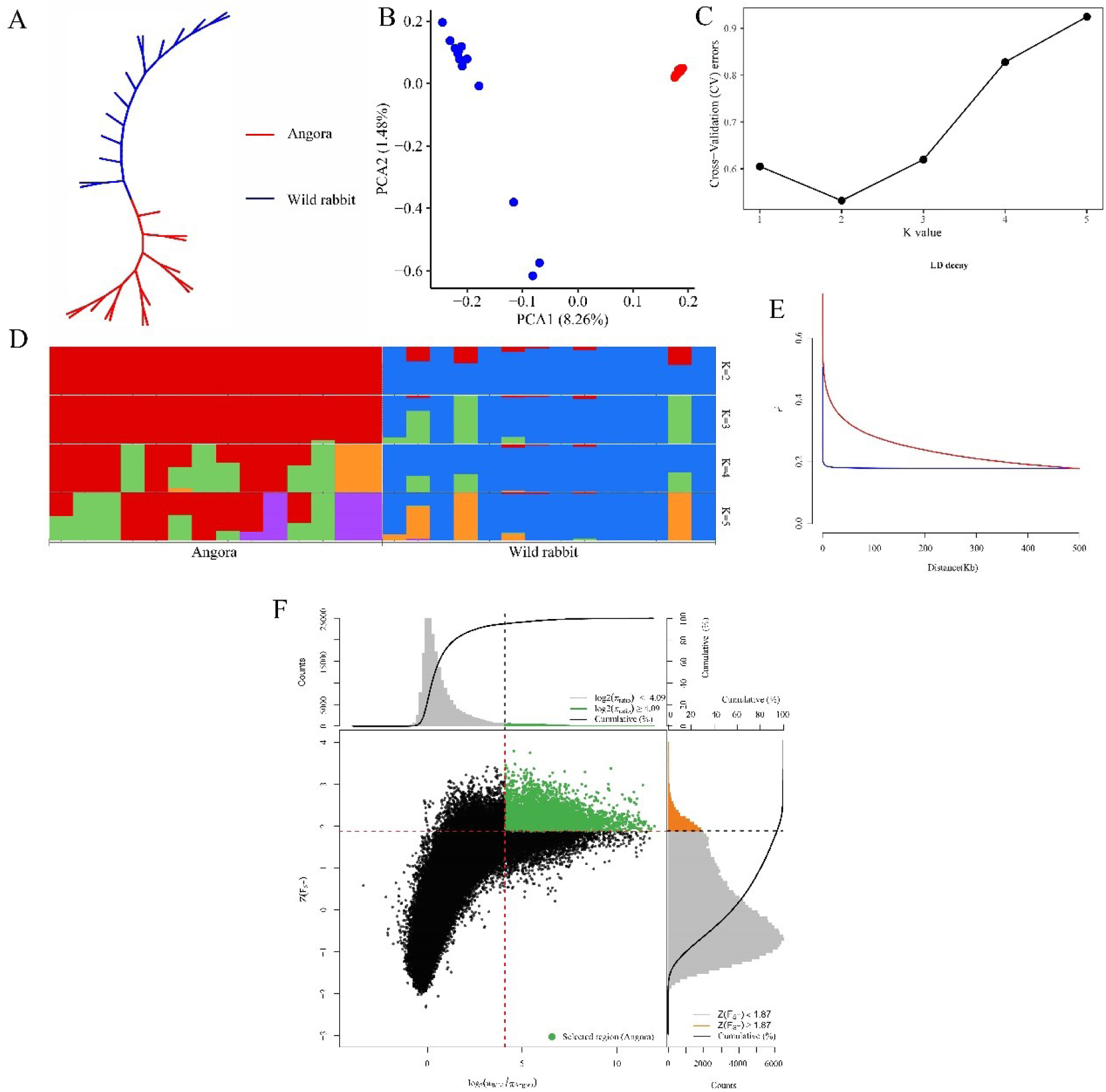
Genetic diversity between the Angora and wild rabbit populations. (A) Maximum-likelihood tree. (B) Principal component analysis revealing genetic differentiation of two populations. (C) Ancestral population analysis (K = 1-5). (D) Admixture plots based on different number of assumed ancestors. (E) The extent of LD, in which values were mean LD r2 values for all pairs of SNPs binned by distance. In all Fig.s, red referred to Angora rabbits and blue referred to wild rabbits. (F) Genomic regions with strong selective sweep signals in Angora and wild rabbits. Distributions of π ratios (wild/Angora) and Z(*F*_ST_) values were calculated by 50-kb windows with 10-kb steps. Genomic regions under selection during domestication were shown as green points located to the top-right regions correspond to the 5% right tails of empirical *log*_2_ (*π*_*wild*_/*π*_*Angora*_) ratio distribution and the top 5% empirical Z(*F*_ST_) distribution. The vertical and horizontal gray lines represent the top 5% value of *log*_2_ (*π*_*wild*_/*π*_*Angora*_) (4.09) and Z(*F*_ST_) (1.87), respectively.

### Genome-wide association analyses

Phenotypes of six traits (LFW, DFW, CVDFW, LCW, RCW and BW) and genotypes including up to 18,577,154 autosomal SNPs after imputation were available for 629 rabbits. For association testing, we used multivariate linear mixed model as implemented in our developed software GMAT (https://github.com/chaoning/GMAT). After LD-based pruning with PLINK, there were 391,976 SNPs in approximate linkage equilibrium with each other and the genome-wide significance level was 1.28e-8 after Bonferroni correction. We showed significant SNPs associated with DFW, CVDFW, LFW and BW in Table S11-S14. Circle Manhattan plot showing significant associations between SNPs and the three traits (DFW, CVDFW, LFW and BW) was given in Fig. 5. Quantile-quantile plot and Manhattan plot for each trait were given in <u>Fig.</u> S2-S3. To sum up, a total of six (five non-overlapping) quantitative trait loci (QTLs, Table 1) were identified for the traits of CVDFW, DFW and BW, and totally 6 independent top significant SNPs are located for the three traits. No QTLs were identified for LFW, LCW or RCW.

**Table 1.** QTL mappings and the 95% confidence interval (95%CI) of each QTL

| **Trait** | **QTL** | **Chr** | **95%CI** |  | **95%CI width** | **Top SNP** | **Allele0** | **Allele1** | **MAF** | **P** | **Gene <sup>1</sup>** |
| --- | --- | --- | --- | --- | --- | --- | --- | --- | --- | --- | --- |
| **DFW** | **QTL1** | 7 | 138,910,411 | 139,598,335 | 687,925 | 138,975,863 | T | C | 0.053963 | 1.36E-07 | 2 |
|  | **QTL2** | 11 | 64,944,841 | 64,952,273 | 7,433 | 64,951,174 | G | C | 0.071669 | 7.12E-23 | 2 |
| **CVDFW** | **QTL1** | 11 | 64,270,630 | 65,432,659 | 1,162,030 | 65,412,003 | A | T | 0.079723 | 1.42E-11 | 6 |
| **BW** | **QTL1** | 2 | 8,257,045 | 8,923,024 | 665,980 | 8,877,739 | G | A | 0.174952 | 7.33E-12 | 4 |
|  | **QTL2** | 4 | 36,704,911 | 38,161,753 | 1,456,843 | 38,127,528 | A | G | 0.083174 | 5.93E-07 | 61 |
|  | **QTL3** | 12 | 148,360,582 | 148,642,693 | 282,112 | 148,369,145 | G | A | 0.083174 | 4.61E-07 | 3 |

1. QTLs for DFW contained no genes, and the nearest genes in the flanking regions were annotated.

**Fig. 5.**
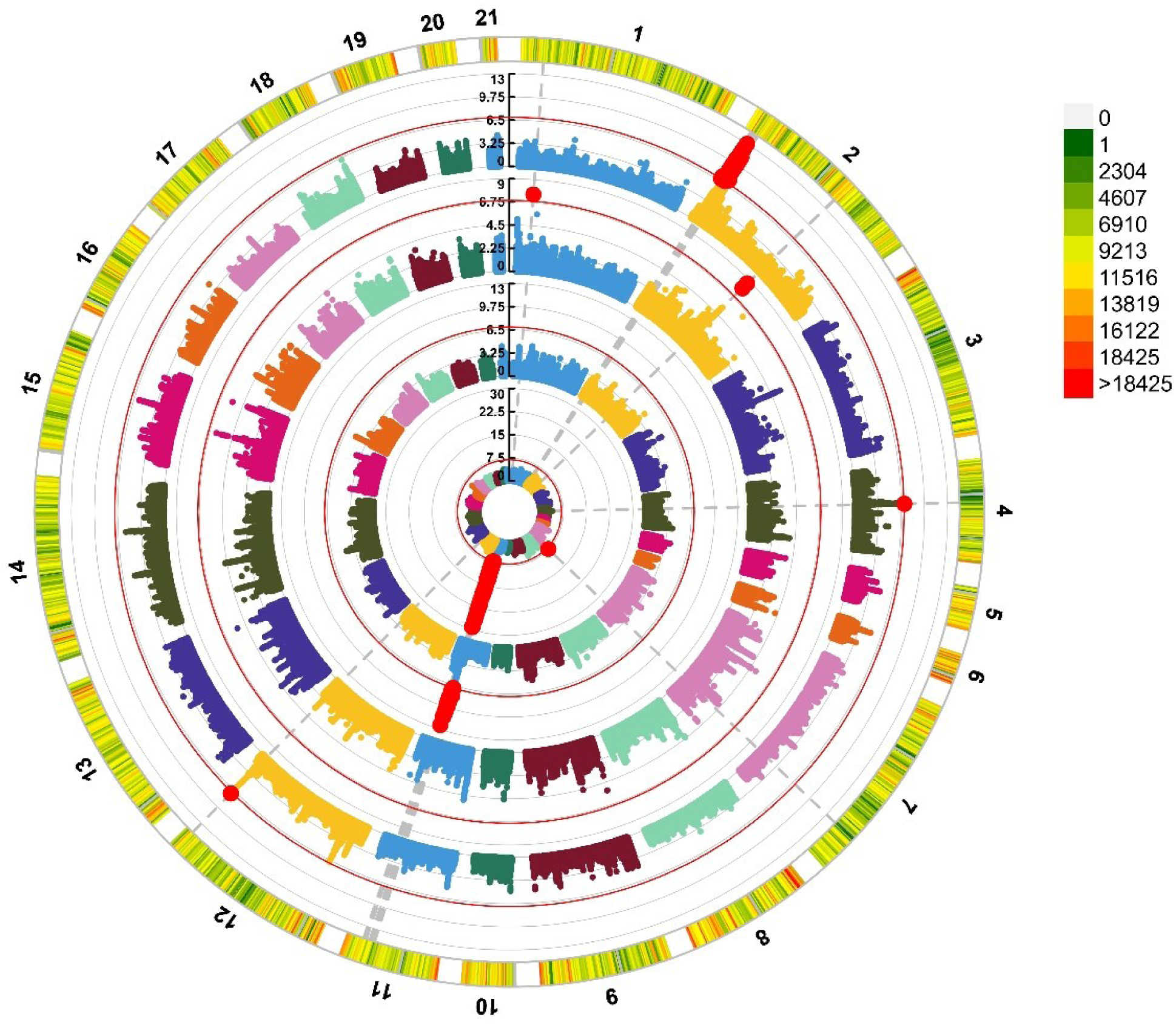
Circle Manhattan plot showing associations between single nucleotide polymorphisms and diameter of fine wool (DFW), coefficient of variation of diameter of fine wool (CVDFW), length of fine wool (LFW) and body weight (BW), respectively (from inside to outside circle), in the Angora rabbit population. The threshold lines indicated the genome-wide significance level (-log_10_(0.05/391,976)) after Bonferroni correction.

Conditional genome-wide association studies were applied with significant lead SNPs regarded as fixed regression. No significant SNPs were located after conditional analyses, which indicated all the significant SNPs in each QTL were in linkage disequilibrium with the most significant SNP and the causal SNP might be at or nearby it. To assist gene identification, the 95% confidence interval (95%CI) of each QTL was estimated using simulations based on the drop in logarithm of *P* values (Nicod et al., 2016). Among the QTLs, the width of 95%CI averaged 0.71 Mb (7.43 kb - 1.46 Mb) with 67% less than 1 Mb (Table 1).

### Heritability estimation based on genome-wide SNPs and by QTLs

SNP-based heritability was estimated for the six traits resulting in a range of 7.45-39.05% indicating a low to medium heritability, with a mean value of 18.98% (Table S15). Among them, the heritability was highest for BW, and lowest for CVDFW. In order to assess the heritability explained by detected QTLs, the effect size of the overall decreased proportion of heritability was analyzed by using the most significant SNPs distributed in these QTLs as fixed effects. The QTLs associated with the three traits explained 2.28-8.52% of the phenotypic variation (Table S16). Among them, DFW exhibited the highest QTL effect with a high phenotypic variation explained by QTL2 (7.50%). CVDFW showed a low QTL effect, similar to its SNP-based heritability.

### Candidate genes identification in high-resolution QTLs

The number of annotated genes covered by the QTLs (for 95%CI) ranged from 0 to 61, with a mean of 12 genes (<u>Table</u> 1 and Table S17). Among them, three contained less than 10 genes in the 95% confidence interval. Two QTLs for DFW did not overlap any gene because of small width, then we found the nearest upstream and downstream genes to the QTLs with distances of 243,754 bp and 279,302 bp away from QTL1 and 474,777 bp and 37,659 bp away from QTL2. QTLs containing smaller numbers of genes were focused on because they provided a starting point for functional investigations. For rabbit wool traits, the QTL for CVDFW on chr11 contained six genes. Among them, *FGF10* caught our attention. The most significant locus (chr11: 65,412,003 bp, *P* = 1.42E-11) associated with fine fiber was detected closest to the *FGF10* gene (chr11: 64,989,932-65,080,400) that was also the nearest gene to the QTL for DFW (chr11: 64,944,841-64,952,273). Fibroblast growth factor 10 (*FGF10*) is a member of the fibroblast growth factor (FGF) family possessing broad mitogenic and cell survival activities and famous for regulation of hair morphogenesis and cycle hair growth in human and mice (Greco et al., 2009, Kinoshita-Ise et al., 2020).

For body weight, QTLs located in chr2 and chr12 contained four and three genes, respectively. Among the genes, several have been announced the association in previous studies. *FAM184B* was associated with chicken body weight by genome-wide study (Zhang et al., 2015, Fan et al., 2017). The *DCAF16-NCAPG* region was susceptible for average daily gain in cattle from the multi-strategy GWASs (Zhang et al., 2016). *NCAPG-LCORL* is known as a locus for adult human height, and have been detected for association with body weight/height in cattle and horses and for selective sweep in dogs and pigs (Takasuga, 2016).

## DISCUSSION

Wool traits are important in rabbits, because the fur is one of the most preferred natural fibers among the textile industries. A well-known breed for fiber production is the Angora rabbit. The fibers obtained from its wool are usually chosen for the production of luxury textile materials. In addition, the rabbit was first and has long been used as models to improve the understanding of human maladies, and was crucial (Esteves et al., 2018), for example, a wool rabbit is considered as an extra model in terms of hair growth. However, there is not an abundance of research on rabbit fiber, with the most studies on hair growth being focused on humans, sheep and mice (Chai et al., 2019, Gur-Cohen et al., 2019, Plowman et al., 2020, Zhao et al., 2021a). In this study, we focus on wool characteristics such as diameter and length that are essential parameters of the wool trait applied in wool rabbit breeding, as well as important indicators of the spinning efficiency of the wool. To investigate the genetic foundation underlying wool traits of Angora rabbits, firstly we cost-effectively explore high-density SNP markers using ultra-low coverage whole genome sequencing combined with three genotype imputation strategies (BaseVar + STITCH, Bcftools + Beagle4 and GATK + Beagle5) and with several levels of sequencing coverages and sample sizes to guarantee imputation performance; then we perform GWAS and conditional GWAS in multivariate linear mixed model in sequence, and map QTLs into 95% confidence intervals for candidate genes of high-resolution to dissect the genetic architecture of complex wool traits in rabbits.

Genotype imputation carried out across the whole genome boosts the number of SNPs. It has been used widely in GWAS to provide a high-resolution view of an associated region, and to increases the chance that a causal SNP can be directly identified (Marchini J. and B., 2010). LCS combined with imputation plays out advantages in obtaining genotyping information since both DNA library and sequencing cost decreased (Nicod et al., 2016, Meier et al., 2021) especially when lacking in microarray (Davies et al., 2016, Davies et al., 2021). It shows to increase performance in a number of scenarios with different outcomes and study designs. We refined and compared genetic variant imputation pipelines using commonly used tools for sequence data and tools specially designed for LCS data based on non-reference panels. Beagle, as one of the most frequently used imputation tools, was attempted to determine whether suitable for LCS data following the conventional SNP calling software Bcftools/GATK. STITCH and its ally BaseVar, which were designed for genotype imputation in LCS data, were applied in this study. As expected, STITCH following BaseVar greatly outperformed the other pipelines, showing GATK is not suitable for LCS data. As a proof of principle, we imputed genotypes for five levels of LCS coverages in overlapping samples of six sizes to assess how sequencing depth and sample size influence power in imputation. We unraveled that sample size hugely affected imputation accuracy at an ultra-low sequencing coverage (<1.0X). At a sequencing depth of 1.0X, the high imputation accuracy by BaseVar + STITCH could be reached and remain stable with genotypic concordance >98.84% and genotypic accuracy >0.97 when a sample size was larger than 300. The patterns of imputation accuracy in different strategies and with several levels of sequencing coverages and sample sizes in this study are in line with previous studies (Davies et al., 2016, Yang et al., 2021, Zhao et al., 2021b).

Comparing to SNP chips, LCS surmounts the problem induced by the ascertainment of common arrays, accurately captures genetic variation in an unbiased manner, effectively identifies novel variation, and enhances variant discovery particularly in underrepresented populations (Martin et al., 2021). In terms of cost, LCS at the sequencing depth of 1X is comparable to and can even be lower than SNP array. In addition, despite the decrease in unit sequencing cost, the cost of high-depth sequencing of a large population remains substantial. Comparing to high-coverage sequencing, LCS can self-evidently save tens of times of genotyping cost per sample in the meantime providing enough allele information, which makes it practicable to sequencing larger samples. Therefore, we strongly corroborate that low coverage sequencing combining with genotype imputation provides a cost-effective and powerful alternative to SNP arrays and high-depth sequencing for more powerful genetic analyses.

With a large sample size and a high-resolution genome-screen, recombination events could be contained to accurately detect causative variants that underlie a quantitative trait(Ros-Freixedes et al., 2017, Loos, 2020). Using whole-genome SNPs explored by low coverage sequencing combining with genotype imputation in a population of 629 rabbits, six QTLs associated with growth and wool traits were recognized, with phenotypic variations ranging from 0.42% to 7.50%. After fine mapping, we focus on the *FGF10* gene associated with fiber growth and diameter. Several FGF members, including *FGF10*, are reported to regulate cycle hair growth mainly to promote the hair follicle (HF) telogen anagen transition *via* providing the stimulatory signals to the HF stem cells and/or their progenies residing in the HF bulge and secondary hair germ (Greco et al., 2009, Mardaryev et al., 2010, Wei-hong Lin, 2015). In addition, FGFs might play an important role in hair morphogenesis. For example, *FGF2* (Takabayashi et al., 2016), *FGF7* (Seo et al., 2016), *FGF9* (Kinoshita-Ise et al., 2020), *FGF10* (Schlake, 2005) and so on have been studied to contribute to a different fiber diameter in human and mice. What’s more, FGF7 and FGF10 efficiently and specifically bind to FGFR2-IIIb, one of several diverse protein variants with distinct binding characteristics encoded by the *FGFR2* gene. Transgenic mice deficient for *FGFR2-IIIb* suffer from abnormally thin hairs, characterized by single columns of medulla cells in all hair types (Schlake, 2005).

Response to strong artificial selection acts on standing genetic variation and completely fixed mutations across many genomic regions, which reflected the long-term directional selection history for wool traits of the Angora rabbit population and probably explained few QTLs detected for some fiber traits. The LD decay and quantile-quantile plots reflected the strong artificial selection. LD extended over a long distance in the Angora rabbit genome, in which markers were separated by 300 kb showing *r*^2^ higher than 0.2. Quantile-quantile plots showed deviation from the expected values extended over a large range of *P* values. What’s more, few QTLs detected for other fiber traits might be due to polygenic genetic architectures, *i*.*e*., small effect of individual mutations contributing to the total genetic variation but not reaching the significant level for a typical complex trait.

## CONCLUSIONS

Low coverage sequencing combined with genotype imputation allows accurate achievement of high-density genotypes even without a good reference panel. GWAS based on LCS data excavates QTLs and fine-maps genes associated with quantitative traits. This study provides a cost-effective analysis pipeline for facilitating our understanding of genetic architecture of complex traits, which lays the foundation for genomic breeding. As accumulation of sequence data, we wish the pipeline will contribute to comprehend genetic mechanism behind important economic traits and to increase genetic progress in livestock.

## MATERIALS AND METHODS

### Animals and phenotypes

A total of 629 Agora rabbits (298 males and 331 females) used for this study were from same batch. All rabbits were housed under the same conditions on a farm, including diet, water and temperature. The experimental procedures in this study were approved by the Animal Care and Use Committee of Shandong Agricultural University. The associated wool traits including length of fine wool (LFW), diameter of fine wool (DFW), coefficient of variation of diameter of fine wool (CVDFW), length of coarse wool (LCW) and rate of coarse wool (RCW) were measured at 70, 140 and 210 days of age. The wool samples from center of lateral body were shaved with clippers. In addition, body weight (BW) was measured including weaning weight at 35 days and body weight at 70, 140 and 210 days of age.

### Sequencing

Ear samples were collected from the individuals. Genomic DNA was isolated using the Qiagen MinElute Kit. Genomic DNA from each sample was used to construct a paired-end library (PE150) with an insert size of ∼ 350 bp. All libraries were sequenced on the BGISEQ-500 platform. An average of 3.84X genomic coverage for 627 samples was sequenced, with the read depth varying from 1.51X to 8.03X. In addition, 15 samples were deep-sequenced at 10X coverage for genotype validation. We generated a total of 7305 gigabases of genomic sequence data.

### Preprocessing of sequence data

Read quality was assessed using FastQC (https://www.bioinformatics.babraham.ac.uk/projects/fastqc/) with a focus on base quality scores, GC content, N content and sequence duplication levels. Adapters and low-quality bases were removed using Trimmomatic as “java -jar trimmomatic-0.38.jar PE sample_1.fq.gz sample_2.fq.gz 1_paired.fq.gz 1_unpaired.fq.gz 2_paired.fq.gz 2_unpaired.fq.gz ILLUMINACLIP:TruSeq3-PE.fa:2:30:10 SLIDINGWINDOW:5:20 LEADING:5 TRAILING:5 MINLEN:50” (Bolger et al., 2014). Sample reads were mapped to the rabbit reference sequence GCF_000003625.3 (*Oryctolagus cuniculus*) using BWA-mem (Li and Durbin, 2009). All PCR duplicates were removed using Picard tools (https://broadinstitute.github.io/picard/).

### Genotyping using high-depth sequencing

Variant calling was performed using GATK4 best practices (McKenna et al., 2010). Base quality score realignment and recalibration were applied to each sample, and haplotypecaller was used for variant discovery. Average coverage was estimated using Qualimap 2.2.1. To simulate low-pass sequencing, the 15 BAM files were downsampled to 0.1X, 0.5X, 1.0X, 1.5X and 2.0X coverage using Picard.

### Genotype imputation using low-coverage sequencing

Due to a lack of a reference panel, the imputation tools that can work without reference information, STITCH and Beagle, were employed to impute genotype of the rabbits using low-coverage sequencing data, combining with different variant calling tools including traditional methods (GATK and SAMtools followed by Bcftools) and the special method for low-pass WGS data (BaseVar). Three imputation pipelines were compared, including (1) BaseVar + STITCH: SNPs were called using BaseVar (Liu et al., 2018b) and imputed genotype dosages at missing sites using STITCH (Davies et al., 2016); (2) Bcftools + Beagle4 (genotype likelihoods): SNPs were called using Bcftools (Li, 2011) and then conducted genotype imputation (with probabilities, dosage genotype) using Beagle v4.1 (Browning and Browning, 2016); (3) GATK + Beagle5: SNPs were called using GATK (McKenna et al., 2010) and then conducted genotype imputation using Beagle v5.1 (Browning et al., 2018). The above two versions of Beagle impute genotypes by different information, Beagle v4.1 infers genotypes from genotype likelihood input data, while Beagle v5.1 does not have this capability but provides significantly fast genotype phasing and similar imputation accuracy (Browning et al., 2018).

### Assessing imputation accuracy

The genotypes directly called from high-coverage sequencing are *de facto* standard for validating the imputation of untyped SNPs (the measure of imputation quality). The imputation accuracy was assessed by comparing imputed genotypes to high-coverage genotypes, and was measured with genotypic concordance and genotypic accuracy by identifying sites shared across both datasets. Sites were considered shared if position, reference allele, and alternate allele were identical. Genotypic concordance (GC) is defined as the proportion of correctly imputed genotypes which was identical to the genotype determined using high-coverage sequencing. Genotypic accuracy (GA) is defined as Pearson correlation coefficient between imputed genotypes and typed genotypes by high-coverage sequencing. The performance of the three cost-effective genotype imputation strategies described above was evaluated by the two criteria. Furthermore, in order to examine the influence of the number of samples and sequence coverage to imputation representation of population genetic variations, we compared imputation accuracy of altering sample size and sequencing depth in this population. Genotypic concordance and genotypic accuracy across different coverages (0.1X, 0.5X, 1.0X, 1.5X, 2.0X) and populations (100, 200, 300, 400, 500 and 600) by down-sampling were calculated to evaluate the influence of sequencing depth and sample size on imputation accuracy.

### Selection of tagging SNPs and annotation

The SNPs (directly genotyped and imputed by STITCH) were filtered for an imputation info score >0.4 using Bcftools, and then with ‘MAF > 0.05, genotype missing rate < 0.1 and a Hardy-Weinberg equilibrium (HWE) *p*-value > 1E-6’ using PLINK (Chang et al., 2015). The sites which were missing in 10% of the individuals after STITCH imputation were then imputed by Beagle v5.1.

SNPs were annotated and categorized as variations in exonic regions, intronic regions and intergenic regions using ANNOVAR (Wang et al., 2010) based on the rabbit reference genome. Those in exons were further classified into synonymous or non-synonymous SNPs.

### Population genetics analysis

In the population of 629 rabbit, principal component analyses (PCA) were performed using the GCTA software (Yang et al., 2011). The first five principal components were extracted and visualized in R. Linkage disequilibrium (LD) decay was measured across the whole genome using r^2^ by PopLDdecay (Zhang et al., 2019). In order to analyze selective sweep regions, CLR were implemented using SweeD (Pavlidis et al., 2013), and nucleotide diversity (Pi) were implemented using Vcftools (Danecek et al., 2011) simultaneously with a window size set to 50 kb and a step size of 10 kb. Sliding windows with the top 1% of CLR and Pi values in the whole genome were regarded as putative selection regions.

Domestication and centuries of selective breeding have changed genomes of rabbit breeds to respond to environmental challenges and human needs. In order to explore further detect genomic footprints of selection, 14 domesticated rabbits were sampled from the population to analyze genetic diversity and population structure by comparing to their wild progenitor, 14 wild rabbits (*Oryctolagus cuniculus*). A maximum-likelihood tree was constructed using the phylogeny program IQ-TREE2. PCA of the first two principal components was visualized. Linkage disequilibrium (LD) decay was compared between two groups. The genetic structure of the two populations was analyzed with ADMIXTURE (Alexander et al., 2009) with *K* values ranging from 1 to 5. *K*-value (*K* = 2) had the lowest cross validation error (CV-error). Pi analysis was applied to estimate the degree of variability within each group, and the fixation statistic *F*_ST_ was applied to explain population differentiation on the basis of the variance of allele frequencies between two groups. Both Pi and *F*_ST_ were calculated using a sliding window analysis with a window size of 50 kb and a step size of 10 kb by Vcftools. The candidate selective sweeps discovered with the top 5% of Pi and *F*_**ST**_ were treated as highly divergent windows. Adjacent windows were merged into a single divergent region and annotated.

Functional enrichment analysis was performed by the Database for Annotation, Visualization and Integrated Discovery (DAVID) to analyze gene ontology (GO) and Kyoto Encyclopedia of Gene and Genome (KEGG) pathway (Huang d W et al., 2009). The P-value of the gene enrichment was corrected by Benjamini-Hochberg FDR (false discovery rate).

### Estimation of whole-genome SNP-based heritability

In our studies, we used three-trait model to analyze wool traits at three time points and four-trait model for body weight. The three-trait model for the heritability estimations of different traits is formulated as

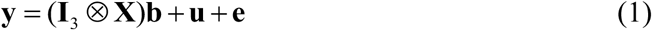

where

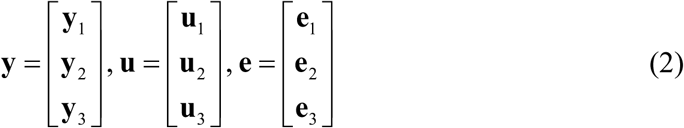

Here, **y** _*i*_ is a vector of phenotypic values for the trait at the *i*th time point; **b**_*i*_ is a vector of fixed effects (population mean, sex and rabbit house); **u**_*i*_ is a vector of additive polygenic genetic effects; **e**_*i*_ is a vector of residual errors. **X** is the design matrix for the fixed effects; ⊗ is the Kronecker product. The distributions of the random effects are

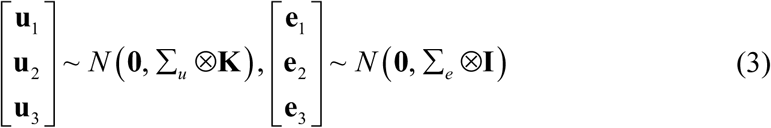

Here, Σ_*u*_ and Σ_*e*_ are a 3× 3 covariance matrix for the additive polygenic effects and residual errors; **K** is the genomic relationship matrix. The additive heritability for the trait at the *i*th time point is defined as

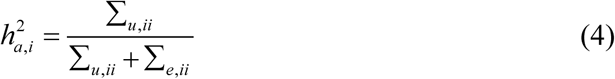

where Σ _*u* ,*ii*_ is the additive genetic variance for the *i*th time point, *i*.*e*., the *i*th diagonal element of Σ_*u*_ ; Σ _*u* ,*ii*_ is the residual variance for the *i*th time point, *i*.*e*., the *i*th diagonal element of Σ_*e*_.

### GWAS

We add the SNP as the fixed effects into the equation (1) to perform multivariate GWAS. The model is formulated as

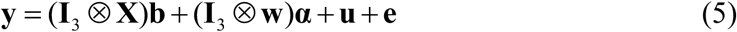

Where ***α*** = [*α*_1_ *α*_2_ *α*_3_]’. Here, *α*_*i*_ is the SNP effect for the trait at the *i*th time point and **w** is a vector of SNP genotypes assigned a value of 0, 1 or 2, respectively, for *aa, Aa* and *AA*. ⊗ is the Kronecker product. We constructed the Wald Chi-square test statistics to test the significance of the SNP effects

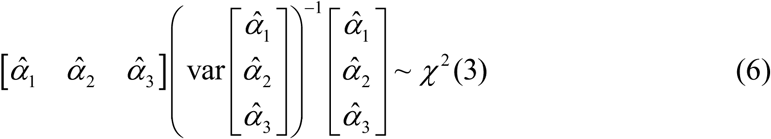

Bonferroni correction was adopted to adjust for multiple testing to control false-positive rates. The threshold for genome-wide significance was 0.05/N, where N was the number of effective SNP calculated by the PLINK “--indep-pairwise 50 5 0.2” command (Purcell et al., 2007).

### Conditional GWAS

To confirm whether the significant SNPs within clusters of loci are independent or linked due to high linkage disequilibrium, we also performed conditional GWAS with significant lead SNP regarded as fixed effect. The model is formulated as

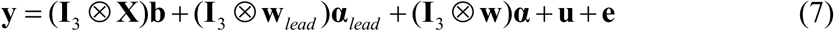

where ***α***_*lead*_ is the lead SNP effect and **w**_*lead*_ is a vector of SNP genotypes for the lead SNP.

### Confidence interval

Confidence intervals were estimated by drop log(*P*) method similarly to previous study (Nicod et al., 2016). In the study, we expanded it into multivariate linear mixed model. Using the SNP effects estimated with equation (5), we first removed the effect of lead SNP at each QTL from the phenotype vector. We random selected 1,000 SNPs within the candidate QTL region and assigned them with effect of lead SNP. The effects of simulated causal SNPs were added to the above residual phenotype vector, one at a time, to produce 1000 simulated datasets. A local association analysis of the region using the equation (5) with the simulated phenotype was performed, and the drop in log(*P*) value between top SNP and simulated causal SNP was recorded for each dataset. Across the 1,000 simulations, we estimated the distribution of these drops in log(*P*) and used the 95th percentile to determine confidence intervals for the original phenotype data.

### Heritability estimation of QTL

We use the following model to estimate as the QTL heritability

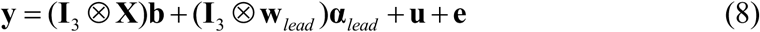

**y** = (**I**_3_ ⊗ **X**)**b** + (**I**_3_ ⊗ **w**_*lead*_)***α***_*lead*_ + **u** + **e** (8) With the lead SNP added into model, we re-calculated the whole-genome SNP-based heritability, which was defined as 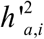. The QTL heritability was estimated using the following equation

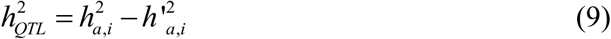

where 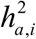 is the whole-genome SNP-based heritability from equation (4).

## Data availability

The sequencing data used for analysis is available at NCBI (PRJNA810279). SNP-based heritability, GWAS, conditional GWAS, confidence interval and QTL heritability estimation were analyzed with our self-written software available at https://github.com/chaoning/GMAT.

## Compliance and ethics

The authors declare that they have no conflict of interest.

## Acknowledgments

This work was supported by the Shandong Province Special Economic Animal Innovation Team (SDAIT-21-02), Agricultural Improved Seed Project of Shandong Province (2021LZGC002), National Natural Science Foundation of China (32102526 and 32002172), China Postdoctoral Science Foundation (2020M682217), Shandong Provincial Postdoctoral Program for Innovative Talent and Shandong Provincial Natural Science Foundation (ZR2020QC176 and ZR2020QC175).

## Author contributions

Xinzhong Fan, Chao Ning and Dan Wang conceived the idea. Yanyan Wang and Wenqiang Li collected the data. Chao Ning performed the theoretical study. Kerui Xie and Dan Wang analyzed the data. Dan Wang, Chao Ning and Xinzhong Fan wrote the manuscript with input from all authors. Xinzhong Fan, Qin Zhang and Jiaqing Hu supervised the research. All authors contributed to the discussions of the results.

## References

Alexander, D. H., J. Novembre, and K. Lange. (2009). Fast model-based estimation of ancestry in unrelated individuals. Genome Res 19, 1655–1664.

Barreiro, L. B. and L. Quintana-Murci. (2020). Evolutionary and population (epi)genetics of immunity to infection. Hum Genet 139, 723–732.

Bolger, A. M., M. Lohse, and B. Usadel. (2014). Trimmomatic: A flexible trimmer for Illumina sequence data. Bioinformatics 30, 2114–2120.

Browning, B. L. and S. R. Browning. (2016). Genotype imputation with millions of reference samples. Am J Hum Genet 98, 116–126.

Browning, B. L., Y. Zhou, and S. R. Browning. (2018). A one-penny imputed genome from next generation reference panels. Am J Hum Genet 103, 338–348.

Browning S.R. and B. B.L. (2007). Rapid and accurate haplotype phasing and missing-data inference for whole-genome association studies by use of localized haplotype clustering. The American Journal of Human Genetics 81, 1084–1097.

Brzezinska-Wcislo, L. (2001). Evaluation of vitamin B6 and calcium pantothenate effectiveness on hair growth from clinical and trichographic aspects for treatment of diffuse alopecia in women. Wiad Lek 54, 11–18.

Carneiro, M., C. J. Rubin, F. Di Palma, F. W. Albert, J. Alfoldi, A. Martinez Barrio, G. Pielberg, N. Rafati, S. Sayyab, J. Turner-Maier, S. Younis, S. Afonso, B. Aken, J. M. Alves, D. Barrell, G. Bolet, S. Boucher, H. A. Burbano, R. Campos, J. L. Chang, V. Duranthon, L. Fontanesi, H. Garreau, D. Heiman, J. Johnson, R. G. Mage, Z. Peng, G. Queney, C. Rogel-Gaillard, M. Ruffier, S. Searle, R. Villafuerte, A. Xiong, S. Young, K. Forsberg-Nilsson, J. M. Good, E. S. Lander, N. Ferrand, K. Lindblad-Toh, and L. Andersson. (2014). Rabbit genome analysis reveals a polygenic basis for phenotypic change during domestication. Science 345, 1074–1079.

Chai, M., M. S. Jiang, L. Vergnes, X. D. Fu, S. C. de Barros, N. B. Doan, W. Huang, J. Chu, J. Jiao, H. Herschman, G. M. Crooks, K. Reue, and J. Huang. (2019). Stimulation of hair growth by small molecules that activate autophagy. Cell Rep 27, 3413-+.

Chang, C. C., C. C. Chow, L. C. Tellier, S. Vattikuti, S. M. Purcell, and J. J. Lee. (2015). Second-generation PLINK: Rising to the challenge of larger and richer datasets. Gigascience 4, 7.

D’Agostini, F., P. Fiallo, T. M. Pennisi, and S. De Flora. (2007). Chemoprevention of smoke-induced alopecia in mice by oral administration of l-cystine and vitamin B6. J Dermatol Sci 46, 189–198.

da Silva Xavier, G., E. A. Bellomo, J. A. McGinty, P. M. French, and G. A. Rutter. (2013). Animal models of GWAS-identified type 2 diabetes genes. J Diabetes Res 2013, 906590.

Danecek, P., A. Auton, G. Abecasis, C. A. Albers, E. Banks, M. A. DePristo, R. E. Handsaker, G. Lunter, G. T. Marth, S. T. Sherry, G. McVean, R. Durbin, and G. Genomes Project Analysis. (2011). The variant call format and VCFtools. Bioinformatics 27, 2156–2158.

Davies, R. W., J. Flint, S. Myers, and R. Mott. (2016). Rapid genotype imputation from sequence without reference panels. Nature Genetics 48, 965-+.

Davies, R. W., M. Kucka, D. W. Su, S. N. Shi, M. Flanagan, C. M. Cunniff, Y. F. Chan, and S. Myers. (2021). Rapid genotype imputation from sequence with reference panels. Nature Genetics 53, 1104-+.

Esteves, P. J., J. Abrantes, and H. Baldauf. (2018). The wide utility of rabbits as models of human diseases. Experimental & Molecular Medicine 50, 1–10.

Fan, Q. C., P. F. Wu, G. J. Dai, G. X. Zhang, T. Zhang, Q. Xue, H. Q. Shi, and J. Y. Wang. (2017). Identification of 19 loci for reproductive traits in a local Chinese chicken by genome-wide study. Genet Mol Res 16.

Freebern, E., D. J. A. Santos, L. Fang, J. Jiang, K. L. Parker Gaddis, G. E. Liu, P. M. VanRaden, C. Maltecca, J. B. Cole, and L. Ma. (2020). GWAS and fine-mapping of livability and six disease traits in Holstein cattle. BMC Genomics 21, 41.

Gerardo, N. M., K. L. Hoang, and K. S. Stoy. (2020). Evolution of animal immunity in the light of beneficial symbioses. Philos Trans R Soc Lond B Biol Sci 375, 20190601.

Gilly, A., G. R. Ritchie, L. Southam, A. E. Farmaki, E. Tsafantakis, G. Dedoussis, and E. Zeggini. (2016). Very low-depth sequencing in a founder population identifies a cardioprotective APOC3 signal missed by genome-wide imputation. Hum Mol Genet 25, 2360–2365.

Gilly, S. A., L. Southam, D. Suveges, K. Kuchenbaecker, R. Moore, G. E. M. Melloni, K. Hatzikotoulas, A. E. Farmaki, G. Ritchie, J. Schwartzentruber, P. Danecek, B. Kilian, M. O. Pollard, X. Ge, E. Tsafantakis, G. Dedoussis, and E. Zeggini. (2019). Very low-depth whole-genome sequencing in complex trait association studies. Bioinformatics 35, 2555–2561.

Greco, V., T. Chen, M. Rendl, M. Schober, H. A. Pasolli, N. Stokes, J. Dela Cruz-Racelis, and E. Fuchs. (2009). A Two-Step Mechanism for Stem Cell Activation during Hair Regeneration. Cell Stem Cell 4, 155–169.

Gur-Cohen, S., H. Yang, S. C. Baksh, Y. Miao, J. Levorse, R. P. Kataru, X. Liu, J. de la Cruz-Racelis, B. J. Mehrara, and E. Fuchs. (2019). Stem cell-driven lymphatic remodeling coordinates tissue regeneration. Science 366, 1218–1225.

Huang d W, Sherman BT, and L. Ra. (2009). Systematic and integrative analysis of large gene lists using DAVID bioinformatics resources. Nature Protocols 4, 44–57.

Huang, H., M. Fang, L. Jostins, M. Umicevic Mirkov, G. Boucher, C. A. Anderson, V. Andersen, I. Cleynen, A. Cortes, F. Crins, M. D’Amato, V. Deffontaine, J. Dmitrieva, E. Docampo, M. Elansary, K. K. Farh, A. Franke, A. S. Gori, P. Goyette, J. Halfvarson, T. Haritunians, J. Knight, I. C. Lawrance, C. W. Lees, E. Louis, R. Mariman, T. Meuwissen, M. Mni, Y. Momozawa, M. Parkes, S. L. Spain, E. Theatre, G. Trynka, J. Satsangi, S. van Sommeren, S. Vermeire, R. J. Xavier, C. International Inflammatory Bowel Disease Genetics, R. K. Weersma, R. H. Duerr, C. G. Mathew, J. D. Rioux, D. P. B. McGovern, J. H. Cho, M. Georges, M. J. Daly, and J. C. Barrett. (2017). Fine-mapping inflammatory bowel disease loci to single-variant resolution. Nature 547, 173–178.

Kainer, D., A. Padovan, J. Degenhardt, S. Krause, P. Mondal, W. J. Foley, and C. Kulheim. (2019). High marker density GWAS provides novel insights into the genomic architecture of terpene oil yield in Eucalyptus. New Phytol 223, 1489–1504.

Kinoshita-Ise, M., A. Tsukashima, T. Kinoshita, Y. Yamazaki, and M. Ohyama. (2020). Altered FGF expression profile in human scalp-derived fibroblasts upon WNT activation: implication of their role to provide folliculogenetic microenvironment. Inflamm Regen 40.

Li, H. (2011). A statistical framework for SNP calling, mutation discovery, association mapping and population genetical parameter estimation from sequencing data. Bioinformatics 27, 2987–2993.

Li, H. and R. Durbin. (2009). Fast and accurate short read alignment with Burrows-Wheeler transform. Bioinformatics 25, 1754–1760.

Liu, S., S. Huang, F. Chen, L. Zhao, Y. Yuan, S. S. Francis, L. Fang, Z. Li, L. Lin, R. Liu, Y. Zhang, H. Xu, S. Li, Y. Zhou, R. W. Davies, Q. Liu, R. G. Walters, K. Lin, J. Ju, T. Korneliussen, M. A. Yang, Q. Fu, J. Wang, L. Zhou, A. Krogh, H. Zhang, W. Wang, Z. Chen, Z. Cai, Y. Yin, H. Yang, M. Mao, J. Shendure, J. Wang, A. Albrechtsen, X. Jin, R. Nielsen, and X. Xu. (2018a). Genomic analyses from non-invasive prenatal testing reveal genetic associations, patterns of viral infections, and Chinese population history. Cell 175, 347–359 e314.

Liu, S., S. Huang, F. Chen, L. Zhao, Y. Yuan, S. S. Francis, L. Fang, Z. Li, L. Lin, R. Liu, Y. Zhang, H. Xu, S. Li, Y. Zhou, R. W. Davies, Q. Liu, R. G. Walters, K. Lin, J. Ju, T. Korneliussen, M. A. Yang, Q. Fu, J. Wang, L. Zhou, A. Krogh, H. Zhang, W. Wang, Z. Chen, Z. Cai, Y. Yin, H. Yang, M. Mao, J. Shendure, J. Wang, A. Albrechtsen, X. Jin, R. Nielsen, and X. Xu. (2018b). Genomic analyses from non-invasive prenatal testing reveal Ggenetic associations, patterns of viral infections, and Chinese population history. Cell 175, 347–359 e314.

Loos, R. J. F. (2020). 15 years of genome-wide association studies and no signs of slowing down. Nat Commun 11.

Marchini J. and H. B. (2010). Genotype imputation for genome-wide association studies. Nature Reviews Genetics 11, 499–511.

Mardaryev, A. N., M. I. Ahmed, N. V. Vlahov, M. Y. Fessing, J. H. Gill, A. A. Sharov, and N. V. Botchkareva. (2010). Micro-RNA-31 controls hair cycle-associated changes in gene expression programs of the skin and hair follicle. FASEB J 24, 3869–3881.

Martin, A. R., E. G. Atkinson, S. B. Chapman, A. Stevenson, R. E. Stroud, T. Abebe, D. Akena, M. Alemayehu, F. K. Ashaba, L. Atwoli, T. Bowers, L. B. Chibnik, M. J. Daly, T. DeSmet, S. Dodge, A. Fekadu, S. Ferriera, B. Gelaye, S. Gichuru, W. E. Injera, R. James, S. M. Kariuki, G. Kigen, K. C. Koenen, E. Kwobah, J. Kyebuzibwa, L. Majara, H. Musinguzi, R. M. Mwema, B. M. Neale, C. P. Newman, C. Newton, J. K. Pickrell, R. Ramesar, W. Shiferaw, D. J. Stein, S. Teferra, C. van der Merwe, Z. Zingela, and G. A. P. P. S. T. Neuro. (2021). Low-coverage sequencing cost-effectively detects known and novel variation in underrepresented populations. Am J Hum Genet 108, 656–668.

McKenna, A., M. Hanna, E. Banks, A. Sivachenko, K. Cibulskis, A. Kernytsky, K. Garimella, D. Altshuler, S. Gabriel, M. Daly, and M. A. DePristo. (2010). The genome analysis toolkit: a MapReduce framework for analyzing next-generation DNA sequencing data. Genome Res 20, 1297–1303.

Meier, J. I., P. A. Salazar, M. Kucka, R. W. Davies, A. Dreau, I. Aldas, O. Box Power, N. J. Nadeau, J. R. Bridle, C. Rolian, N. H. Barton, W. O. McMillan, C. D. Jiggins, and Y. F. Chan. (2021). Haplotype tagging reveals parallel formation of hybrid races in two butterfly species. Proc Natl Acad Sci U S A 118.

Nicod, J., R. W. Davies, N. Cai, C. Hassett, L. Goodstadt, C. Cosgrove, B. K. Yee, V. Lionikaite, R. E. McIntyre, C. A. Remme, E. M. Lodder, J. S. Gregory, T. Hough, R. Joynson, H. Phelps, B. Nell, C. Rowe, J. Wood, A. Walling, N. Bopp, A. Bhomra, P. Hernandez-Pliego, J. Callebert, R. M. Aspden, N. P. Talbot, P. A. Robbins, M. Harrison, M. Fray, J. M. Launay, Y. M. Pinto, D. A. Blizard, C. R. Bezzina, D. J. Adams, P. Franken, T. Weaver, S. Wells, S. D. Brown, P. K. Potter, P. Klenerman, A. Lionikas, R. Mott, and J. Flint. (2016). Genome-wide association of multiple complex traits in outbred mice by ultra-low-coverage sequencing. Nat Genet 48, 912–918.

Pavlidis, P., D. Zivkovic, A. Stamatakis, and N. Alachiotis. (2013). SweeD: likelihood-based detection of selective sweeps in thousands of genomes. Mol Biol Evol 30, 2224–2234.

Plowman, J. E., D. P. Harland, A. M. O. Campos, E. S. S. Rocha, A. Thomas, J. A. Vernon, C. van Koten, C. Hefer, S. Clerens, and A. M. de Almeida. (2020). The wool proteome and fibre characteristics of three distinct genetic ovine breeds from Portugal. J Proteomics 225, 103853.

Purcell, S., B. Neale, K. Todd-Brown, L. Thomas, M. A. Ferreira, D. Bender, J. Maller, P. Sklar, P. I. de Bakker, M. J. Daly, and P. C. Sham. (2007). PLINK: A tool set for whole-genome association and population-based linkage analyses. Am J Hum Genet 81, 559–575.

Qin, P., H. Lu, H. Du, H. Wang, W. Chen, Z. Chen, Q. He, S. Ou, H. Zhang, X. Li, X. Li, Y. Li, Y. Liao, Q. Gao, B. Tu, H. Yuan, B. Ma, Y. Wang, Y. Qian, S. Fan, W. Li, J. Wang, M. He, J. Yin, T. Li, N. Jiang, X. Chen, C. Liang, and S. Li. (2021). Pan-genome analysis of 33 genetically diverse rice accessions reveals hidden genomic variations. Cell 184, 3542–3558 e3516.

Quintana-Murci, L. (2019). Human immunology through the lens of evolutionary genetics. Cell 177, 184–199.

Ros-Freixedes, R., S. Gonen, G. Gorjanc, and J. M. Hickey. (2017). A method for allocating low-coverage sequencing resources by targeting haplotypes rather than individuals. Genet Sel Evol 49.

Rubinacci, S., D. M. Ribeiro, R. J. Hofmeister, and O. Delaneau. (2021). Efficient phasing and imputation of low-coverage sequencing data using large reference panels. Nat Genet 53, 120–126.

Schlake, T. (2005). FGF signals specifically regulate the structure of hair shaft medulla via IGF-binding protein 5. Development 132, 2981–2990.

Seo, H. S., D. J. Lee, J. H. Chung, C. H. Lee, H. R. Kim, J. E. Kim, B. J. Kim, M. H. Jung, K. T. Ha, and H. S. Jeong. (2016). Hominis Placenta facilitates hair re-growth by upregulating cellular proliferation and expression of fibroblast growth factor-7. BMC Complement Altern Med 16, 187.

Spiliopoulou, A., M. Colombo, P. Orchard, F. Agakov, and P. McKeigue. (2017). GeneImp: Fast imputation to large reference panels using genotype likelihoods from ultralow coverage sequencing. Genetics 206, 91–104.

Takabayashi, Y., M. Nambu, M. Ishihara, M. Kuwabara, K. Fukuda, S. Nakamura, H. Hattori, and T. Kiyosawa. (2016). Enhanced effect of fibroblast growth factor-2-containing dalteparin/protamine nanoparticles on hair growth. Clin Cosmet Investig Dermatol 9, 127–134.

Takasuga, A. (2016). PLAG1 and NCAPG-LCORL in livestock. Anim Sci J 87, 159–167.

Wang, K., M. Li, and H. Hakonarson. (2010). ANNOVAR: functional annotation of genetic variants from high-throughput sequencing data. Nucleic Acids Res 38, e164.

Wasik, K., T. Berisa, J. K. Pickrell, J. H. Li, D. J. Fraser, K. King, and C. Cox. (2021). Comparing low-pass sequencing and genotyping for trait mapping in pharmacogenetics. BMC Genomics 22, 197.

Wei-hong Lin, L.-J. X., Hong-Xue Shi, Jian Zhang, Li-ping Jiang, Ping-tao Cai, Zhen-Lang Lin, Bei-Bei Lin, Yan Huang, Hai-Lin Zhang, Xiao-Bing Fu, Ding-Jiong Guo, Xiao-Kun Li, Xiao-Jie Wang, Jian Xiao. (2015). Fibroblast growth factors stimulate hair growth through β-catenin and Shh expression in C57BL/6 mice. BioMed Research International 2015, 730139.

Yang, J., S. H. Lee, M. E. Goddard, and P. M. Visscher. (2011). GCTA: a tool for Genome-wide Complex Trait Analysis. Am J Hum Genet 88, 76–82.

Yang, R., X. Guo, D. Zhu, C. Tan, C. Bian, J. Ren, Z. Huang, Y. Zhao, G. Cai, D. Liu, Z. Wu, Y. Wang, N. Li, and X. Hu. (2021). Accelerated deciphering of the genetic architecture of agricultural economic traits in pigs using a low-coverage whole-genome sequencing strategy. GigaScience 10.

Zhang, C., S. S. Dong, J. Y. Xu, W. M. He, and T. L. Yang. (2019). PopLDdecay: a fast and effective tool for linkage disequilibrium decay analysis based on variant call format files. Bioinformatics 35, 1786–1788.

Zhang, G. X., Q. C. Fan, J. Y. Wang, T. Zhang, Q. Xue, and H. Q. Shi. (2015). Genome-wide association study on reproductive traits in Jinghai Yellow Chicken. Anim Reprod Sci 163, 30–34.

Zhang, W. G., J. Y. Li, Y. Guo, L. P. Zhang, L. Y. Xu, X. Gao, B. Zhu, H. J. Gao, H. M. Ni, and Y. Chen. (2016). Multi-strategy genome-wide association studies identify the DCAF16-NCAPG region as a susceptibility locus for average daily gain in cattle. Sci Rep-Uk 6.

Zhao, B., H. Luo, J. He, X. Huang, S. Chen, X. Fu, W. Zeng, Y. Tian, S. Liu, C. J. Li, G. E. Liu, L. Fang, S. Zhang, and K. Tian. (2021a). Comprehensive transcriptome and methylome analysis delineates the biological basis of hair follicle development and wool-related traits in Merino sheep. BMC Biol 19, 197.

Zhao, C., J. Teng, X. Zhang, D. Wang, X. Zhang, S. Li, X. Jiang, H. Li, C. Ning, and Q. Zhang. (2021b). Towards a Cost-Effective Implementation of Genomic Prediction Based on Low Coverage Whole Genome Sequencing in Dezhou Donkey. Frontiers in Genetics 2.

