## supplement tables and figures for "Cost-effectively dissecting the genetic architecture of complex wool traits in rabbits by low-coverage sequencing": Supplemental Figures.docx

**Fig. S1.** CLR and Pi analyses in the Angora rabbit population

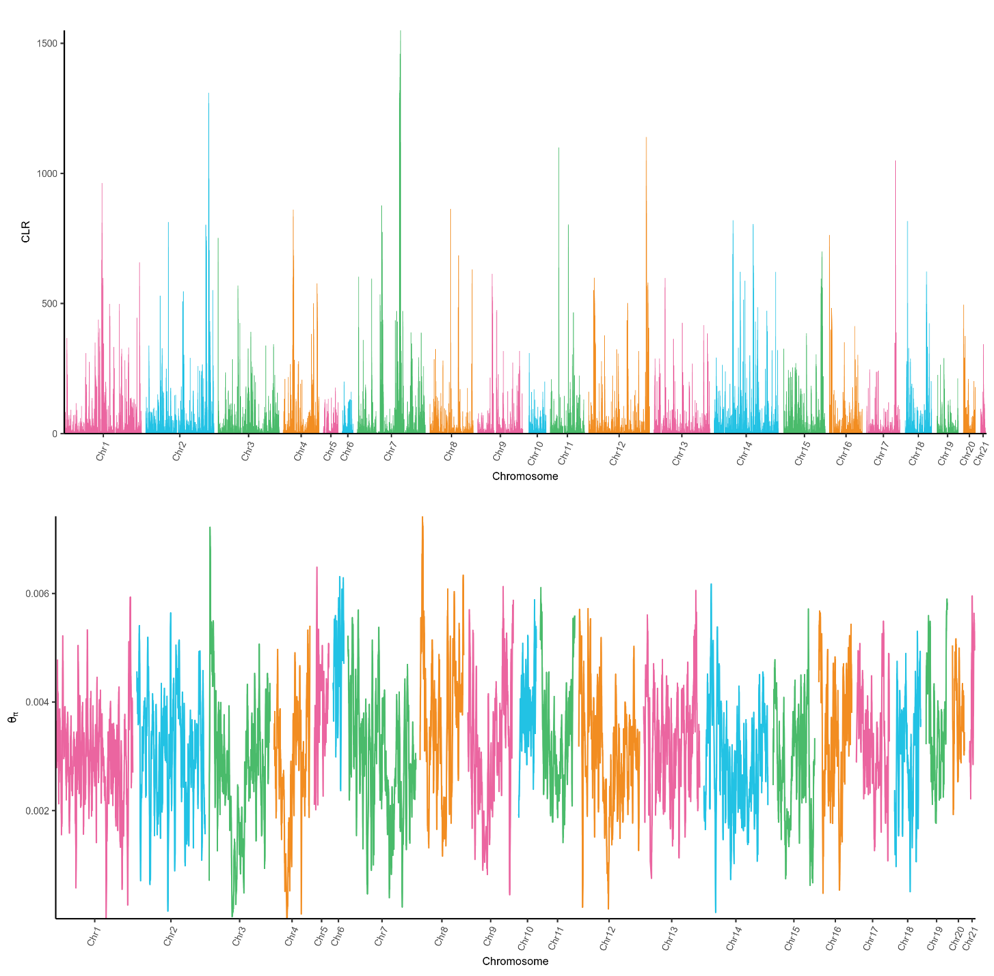

**Fig. S2.** Q-Q plots for the Angora rabbits (A: Body weight, B: DFW, C: CVDFW, D: LCW, E: LFW, F: RCW)

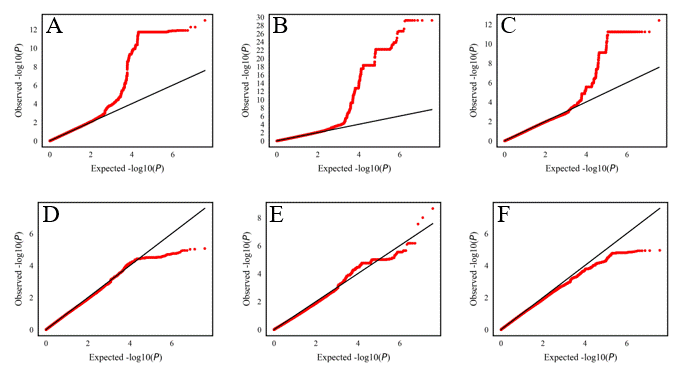

**Fig. S3.** Manhattan plots for the Angora rabbits (A: Body weight, B: DFW, C: CVDFW, D: LCW, E: LFW, F: RCW)

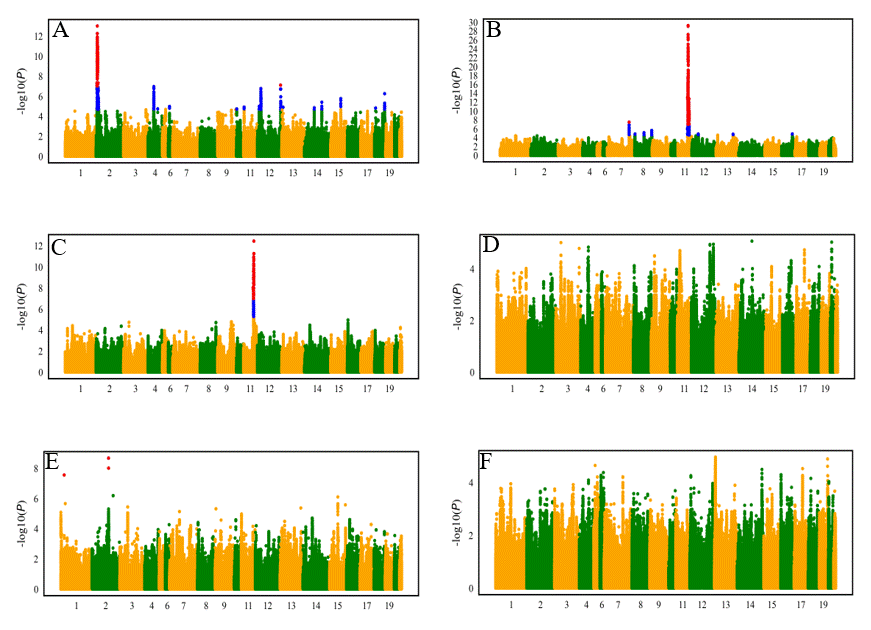
