## supplement tables and figures for "Cost-effectively dissecting the genetic architecture of complex wool traits in rabbits by low-coverage sequencing": Supplemental Tables .pdf

**Table S1. Genotypic accuracy and concordance of imputation by the three pipelines on 15 individuals**

| Pipeline | Individual | Genotypic_Accuracy | Genotypic_Concordance(%) |
|---|---|---|---|
| BaseVar+STITCH | 1 | 0.98 | 99.25 |
| BaseVar+STITCH | 2 | 0.95 | 98.24 |
| BaseVar+STITCH | 3 | 0.98 | 98.80 |
| BaseVar+STITCH | 4 | 0.98 | 99.24 |
| BaseVar+STITCH | 5 | 0.98 | 99.31 |
| BaseVar+STITCH | 6 | 0.97 | 98.74 |
| BaseVar+STITCH | 7 | 0.97 | 99.04 |
| BaseVar+STITCH | 8 | 0.99 | 99.58 |
| BaseVar+STITCH | 9 | 0.98 | 99.19 |
| BaseVar+STITCH | 10 | 0.99 | 99.47 |
| BaseVar+STITCH | 11 | 0.97 | 99.17 |
| BaseVar+STITCH | 12 | 0.98 | 99.30 |
| BaseVar+STITCH | 13 | 0.97 | 99.10 |
| BaseVar+STITCH | 14 | 0.97 | 98.90 |
| BaseVar+STITCH | 15 | 0.98 | 98.89 |
| Bcftools+Beagle4 | 1 | 0.74 | 84.99 |
| Bcftools+Beagle4 | 2 | 0.62 | 83.62 |
| Bcftools+Beagle4 | 3 | 0.75 | 91.14 |
| Bcftools+Beagle4 | 4 | 0.76 | 87.78 |
| Bcftools+Beagle4 | 5 | 0.74 | 88.43 |
| Bcftools+Beagle4 | 6 | 0.75 | 90.19 |
| Bcftools+Beagle4 | 7 | 0.76 | 87.58 |
| Bcftools+Beagle4 | 8 | 0.81 | 94.42 |
| Bcftools+Beagle4 | 9 | 0.65 | 87.00 |
| Bcftools+Beagle4 | 10 | 0.84 | 88.48 |
| Bcftools+Beagle4 | 11 | 0.72 | 89.67 |
| Bcftools+Beagle4 | 12 | 0.88 | 95.73 |
| Bcftools+Beagle4 | 13 | 0.62 | 84.69 |
| Bcftools+Beagle4 | 14 | 0.87 | 95.18 |
| Bcftools+Beagle4 | 15 | 0.78 | 86.68 |
| GATK+Beagle5 | 1 | 0.58 | 74.30 |
| GATK+Beagle5 | 2 | 0.40 | 66.11 |
| GATK+Beagle5 | 3 | 0.64 | 77.18 |
| GATK+Beagle5 | 4 | 0.50 | 66.93 |
| GATK+Beagle5 | 5 | 0.63 | 80.19 |
| GATK+Beagle5 | 6 | 0.53 | 77.07 |
| GATK+Beagle5 | 7 | 0.46 | 68.71 |
| GATK+Beagle5 | 8 | 0.47 | 75.77 |
| GATK+Beagle5 | 9 | 0.53 | 73.13 |
| GATK+Beagle5 | 10 | 0.58 | 74.18 |
| GATK+Beagle5 | 11 | 0.38 | 66.86 |
| GATK+Beagle5 | 12 | 0.60 | 76.62 |
| GATK+Beagle5 | 13 | 0.46 | 68.62 |
| GATK+Beagle5 | 14 | 0.43 | 64.81 |
| GATK+Beagle5 | 15 | 0.59 | 72.36 |

**Table S2. Genotypic accuracy and concordance of imputation by the three pipelines with the MAF ranges**

| Pipeline | MAF | Genotypic Accuracy | Genotypic Concordance(%) |
|---|---|---|---|
| BaseVar+STITCH | 0-0.01 | 0.07 | 79.55 |
| BaseVar+STITCH | 0.01-0.05 | 0.89 | 97.99 |
| BaseVar+STITCH | 0.05-0.1 | 0.97 | 99.31 |
| BaseVar+STITCH | 0.1-0.15 | 0.98 | 99.19 |
| BaseVar+STITCH | 0.15-0.2 | 0.98 | 99.11 |
| BaseVar+STITCH | 0.2-0.25 | 0.98 | 98.98 |
| BaseVar+STITCH | 0.25-0.3 | 0.98 | 98.91 |
| BaseVar+STITCH | 0.3-0.35 | 0.98 | 98.82 |
| BaseVar+STITCH | 0.35-0.4 | 0.98 | 98.82 |
| BaseVar+STITCH | 0.4-0.45 | 0.98 | 98.82 |
| BaseVar+STITCH | 0.45-0.5 | 0.98 | 98.85 |
| Bcftools+Beagle4 | 0-0.01 | 0.16 | 80.97 |
| Bcftools+Beagle4 | 0.01-0.05 | 0.55 | 89.79 |
| Bcftools+Beagle4 | 0.05-0.1 | 0.75 | 93.10 |
| Bcftools+Beagle4 | 0.1-0.15 | 0.78 | 92.17 |
| Bcftools+Beagle4 | 0.15-0.2 | 0.80 | 91.65 |
| Bcftools+Beagle4 | 0.2-0.25 | 0.80 | 89.90 |
| Bcftools+Beagle4 | 0.25-0.3 | 0.78 | 88.20 |
| Bcftools+Beagle4 | 0.3-0.35 | 0.78 | 86.40 |
| Bcftools+Beagle4 | 0.35-0.4 | 0.77 | 85.80 |
| Bcftools+Beagle4 | 0.4-0.45 | 0.78 | 85.77 |
| Bcftools+Beagle4 | 0.45-0.5 | 0.78 | 84.66 |
| GATK+Beagle5 | 0-0.01 | 0.07 | 77.53 |
| GATK+Beagle5 | 0.01-0.05 | 0.27 | 81.26 |
| GATK+Beagle5 | 0.05-0.1 | 0.42 | 80.60 |
| GATK+Beagle5 | 0.1-0.15 | 0.51 | 77.54 |
| GATK+Beagle5 | 0.15-0.2 | 0.52 | 73.38 |
| GATK+Beagle5 | 0.2-0.25 | 0.56 | 70.23 |
| GATK+Beagle5 | 0.25-0.3 | 0.55 | 66.18 |
| GATK+Beagle5 | 0.3-0.35 | 0.58 | 66.14 |
| GATK+Beagle5 | 0.35-0.4 | 0.62 | 66.75 |
| GATK+Beagle5 | 0.4-0.45 | 0.63 | 65.72 |
| GATK+Beagle5 | 0.45-0.5 | 0.62 | 64.02 |

**Table S3. Genotypic accuracy and concordance of imputation with different sample size and depth by BaseVar+STITCH**

| Depth | Sample size | Genotypic Accuracy | Genotypic Concordance(%) |
|---|---|---|---|
| 0.1× | 100 | 0.15 | 63.84 |
| 0.1× | 200 | 0.31 | 72.74 |
| 0.1× | 300 | 0.53 | 81.92 |
| 0.1× | 400 | 0.67 | 87.57 |
| 0.1× | 500 | 0.75 | 90.59 |
| 0.1× | 600 | 0.82 | 92.86 |
| 0.5× | 100 | 0.79 | 91.78 |
| 0.5× | 200 | 0.92 | 96.97 |
| 0.5× | 300 | 0.96 | 98.31 |
| 0.5× | 400 | 0.97 | 98.67 |
| 0.5× | 500 | 0.97 | 98.82 |
| 0.5× | 600 | 0.97 | 98.92 |
| 1.0× | 100 | 0.93 | 97.01 |
| 1.0× | 200 | 0.96 | 98.49 |
| 1.0× | 300 | 0.97 | 98.84 |
| 1.0× | 400 | 0.97 | 98.91 |
| 1.0× | 500 | 0.98 | 98.97 |
| 1.0× | 600 | 0.98 | 98.99 |
| 1.5× | 100 | 0.96 | 98.14 |
| 1.5× | 200 | 0.97 | 98.76 |
| 1.5× | 300 | 0.98 | 98.95 |
| 1.5× | 400 | 0.98 | 98.96 |
| 1.5× | 500 | 0.98 | 99.00 |
| 1.5× | 600 | 0.98 | 99.01 |
| 2.0× | 100 | 0.96 | 98.51 |
| 2.0× | 200 | 0.97 | 98.87 |
| 2.0× | 300 | 0.98 | 98.97 |
| 2.0× | 400 | 0.98 | 99.04 |
| 2.0× | 500 | 0.98 | 99.04 |
| 2.0× | 600 | 0.98 | 99.05 |

**Table S4. Variant analysis by regions and functions**

| Region | SNP number | Percentage |
|---|---|---|
| **intergenic** | 10,733,466 | 57.78% |
| **intronic** | 6,594,031 | 35.50% |
| **exonic** | 96,276 | 0.52% |
| **upstream** | 131,610 | 0.71% |
| **downstream** | 128,233 | 0.69% |
| **UTR5** | 11,730 | 0.06% |
| **UTR3** | 31,490 | 0.17% |
| **splicing** | 323 | 0.00% |
| **ncRNA** | 0 | 0.00% |

| Function | SNP Number | Nonsynonymous/synonymous |
|---|---|---|
| **synonymous** | 72,552 | 0.32 |
| **nonsynonymous** | 23,328 | |

**Table S5. LD decay in the Angora rabbit population**

| Distance | Mean_r^2 across the ge[illegible] | Mean_r^2 on chr15 | Mean_r^2 on chr21 | Mean_r^2 on chr11 |
|---|---|---|---|---|
| 10 | 0.764228903 | 0.803498026 | 0.731700904 | 0.757011428 |
| 20 | 0.723217377 | 0.772132579 | 0.675324519 | 0.71390568 |
| 30 | 0.708651517 | 0.761388571 | 0.652020306 | 0.698369492 |
| 40 | 0.700082494 | 0.752994011 | 0.63652428 | 0.691805937 |
| 50 | 0.693746703 | 0.746847047 | 0.631434934 | 0.684064068 |
| 60 | 0.68852647 | 0.744754903 | 0.623121266 | 0.678838331 |
| 70 | 0.685725401 | 0.742642411 | 0.61812611 | 0.678053631 |
| 80 | 0.681822696 | 0.739089324 | 0.611738456 | 0.673915531 |
| 90 | 0.680174339 | 0.737717566 | 0.61021329 | 0.671782213 |
| 100 | 0.677208182 | 0.734908081 | 0.606728367 | 0.670119545 |
| 200 | 0.668108722 | 0.726961692 | 0.5947081 | 0.660091686 |
| 300 | 0.655416358 | 0.716837824 | 0.578492157 | 0.647837843 |
| 400 | 0.645714061 | 0.708090814 | 0.566288357 | 0.639972733 |
| 500 | 0.637663972 | 0.701503578 | 0.556255665 | 0.631995384 |
| 600 | 0.630971975 | 0.695454069 | 0.549346957 | 0.625015552 |
| 700 | 0.625108743 | 0.689591855 | 0.543752531 | 0.619833024 |
| 800 | 0.619534542 | 0.68399733 | 0.539578376 | 0.613974678 |
| 900 | 0.6139436 | 0.678789844 | 0.531323587 | 0.608781181 |
| 1,000 | 0.609299893 | 0.675141272 | 0.530661268 | 0.604488626 |
| 1,100 | 0.60464404 | 0.671130895 | 0.523736585 | 0.599741725 |
| 1,200 | 0.600263823 | 0.666090041 | 0.521465824 | 0.59524276 |
| 1,300 | 0.596419135 | 0.662948691 | 0.515515646 | 0.592248168 |
| 1,400 | 0.592766208 | 0.658735552 | 0.513337698 | 0.589347425 |
| 1,500 | 0.589164835 | 0.655463356 | 0.510035182 | 0.585933074 |
| 1,600 | 0.585530577 | 0.652357104 | 0.505365295 | 0.581966277 |
| 1,700 | 0.582405504 | 0.649735449 | 0.502913726 | 0.578010963 |
| 1,800 | 0.579235646 | 0.646157471 | 0.49718939 | 0.576135852 |
| 1,900 | 0.576429738 | 0.643041974 | 0.4957792 | 0.57329301 |
| 2,000 | 0.57355748 | 0.640622495 | 0.490256451 | 0.569445613 |
| 2,100 | 0.570658457 | 0.638189498 | 0.487253869 | 0.566077189 |
| 2,200 | 0.56818623 | 0.636149931 | 0.484627371 | 0.564883233 |
| 2,300 | 0.565579999 | 0.632302653 | 0.482692048 | 0.562356924 |
| 2,400 | 0.563063316 | 0.631094027 | 0.481109486 | 0.559312711 |
| 2,500 | 0.56067848 | 0.628834828 | 0.478141857 | 0.556999496 |
| 2,600 | 0.55822993 | 0.625360625 | 0.475299107 | 0.55515209 |
| 2,700 | 0.555930306 | 0.62307993 | 0.471367409 | 0.552602234 |
| 2,800 | 0.553986432 | 0.621194793 | 0.468206524 | 0.551009314 |
| 2,900 | 0.551909566 | 0.619451992 | 0.467852893 | 0.548662713 |
| 3,000 | 0.549981518 | 0.616545893 | 0.465621997 | 0.547519836 |
| 3,100 | 0.548150636 | 0.615699086 | 0.466002248 | 0.544227067 |
| 3,200 | 0.546045828 | 0.612686644 | 0.463077516 | 0.543070429 |
| 3,300 | 0.544175432 | 0.611175964 | 0.462014504 | 0.540707067 |
| 3,400 | 0.542232639 | 0.608080563 | 0.460600796 | 0.538757733 |
| 3,500 | 0.540639619 | 0.6071765 | 0.460028125 | 0.537579751 |
| 3,600 | 0.538657181 | 0.60466349 | 0.459545961 | 0.537393825 |
| 3,700 | 0.537037075 | 0.603536163 | 0.455002872 | 0.535178487 |
| 3,800 | 0.535312766 | 0.601968808 | 0.45413955 | 0.534138332 |
| 3,900 | 0.533820671 | 0.60056044 | 0.451389024 | 0.531695814 |
| 4,000 | 0.532258523 | 0.599517155 | 0.449915629 | 0.530218098 |
| 4,100 | 0.530645159 | 0.597085134 | 0.446239402 | 0.528616714 |
| 4,200 | 0.529192616 | 0.595167827 | 0.446576544 | 0.526789954 |
| 4,300 | 0.527575407 | 0.595237776 | 0.445677476 | 0.525169792 |
| 4,400 | 0.526148502 | 0.593368062 | 0.445198888 | 0.523233139 |
| 4,500 | 0.524656491 | 0.591614975 | 0.442735447 | 0.522437052 |
| 4,600 | 0.523377806 | 0.590367766 | 0.440930887 | 0.520240254 |
| 4,700 | 0.522013795 | 0.588633305 | 0.440693546 | 0.520774943 |
| 4,800 | 0.52072517 | 0.587102315 | 0.438831094 | 0.51973956 |
| 4,900 | 0.519468698 | 0.586378581 | 0.438371743 | 0.517929539 |
| 5,000 | 0.518112561 | 0.584385775 | 0.436527018 | 0.516961045 |
| 5,100 | 0.516985038 | 0.584049569 | 0.433813473 | 0.514514077 |
| 5,200 | 0.515587011 | 0.581704376 | 0.431863092 | 0.513857136 |
| 5,300 | 0.514359918 | 0.58180496 | 0.430914884 | 0.513120647 |
| 5,400 | 0.51316337 | 0.581322011 | 0.429212459 | 0.510484341 |
| 5,500 | 0.511966988 | 0.57958385 | 0.428231868 | 0.510628891 |
| 5,600 | 0.510750171 | 0.578418662 | 0.424513159 | 0.510716512 |

| | | | | |
|---|---|---|---|---|
| 5,700 | 0.509605382 | 0.577263322 | 0.422519166 | 0.509189007 |
| 5,800 | 0.508343076 | 0.575959521 | 0.421906573 | 0.506191759 |
| 5,900 | 0.507368734 | 0.575775496 | 0.422627016 | 0.505718785 |
| 6,000 | 0.506167946 | 0.57573002 | 0.419833354 | 0.505341687 |
| 6,100 | 0.505293784 | 0.573381235 | 0.420876329 | 0.503072311 |
| 6,200 | 0.504201014 | 0.573090973 | 0.418217931 | 0.501913958 |
| 6,300 | 0.503095091 | 0.572105864 | 0.419405589 | 0.5005185 |
| 6,400 | 0.50195193 | 0.571370903 | 0.41898776 | 0.500287975 |
| 6,500 | 0.500970507 | 0.57049664 | 0.415331271 | 0.499571457 |
| 6,600 | 0.49979656 | 0.568345691 | 0.415722824 | 0.497866729 |
| 6,700 | 0.498943629 | 0.569014018 | 0.414239467 | 0.498171185 |
| 6,800 | 0.497796082 | 0.567859756 | 0.411828439 | 0.496106488 |
| 6,900 | 0.496945081 | 0.56727224 | 0.412461288 | 0.494385259 |
| 7,000 | 0.496074403 | 0.565914128 | 0.411464356 | 0.494012989 |
| 7,100 | 0.494874076 | 0.56471818 | 0.410780191 | 0.492515645 |
| 7,200 | 0.49417231 | 0.564243184 | 0.409201582 | 0.491935207 |
| 7,300 | 0.493142941 | 0.563198028 | 0.407401889 | 0.490554175 |
| 7,400 | 0.492386556 | 0.563102526 | 0.40643729 | 0.489844695 |
| 7,500 | 0.491513369 | 0.562231826 | 0.403834677 | 0.48827692 |
| 7,600 | 0.490549613 | 0.561243216 | 0.404956196 | 0.489341088 |
| 7,700 | 0.489873139 | 0.559775814 | 0.402338075 | 0.48774387 |
| 7,800 | 0.48902884 | 0.558531285 | 0.398792521 | 0.486467266 |
| 7,900 | 0.488032359 | 0.558309234 | 0.399199338 | 0.485990193 |
| 8,000 | 0.487126528 | 0.557302467 | 0.399366982 | 0.485859282 |
| 8,100 | 0.486380267 | 0.556187507 | 0.398831395 | 0.482525052 |
| 8,200 | 0.485404117 | 0.555957126 | 0.397331529 | 0.482426744 |
| 8,300 | 0.484334212 | 0.554992253 | 0.397497195 | 0.481464997 |
| 8,400 | 0.483463571 | 0.553330116 | 0.394844036 | 0.480704379 |
| 8,500 | 0.482722684 | 0.553023429 | 0.39662389 | 0.480330426 |
| 8,600 | 0.482259442 | 0.553021507 | 0.398242467 | 0.479910045 |
| 8,700 | 0.481211875 | 0.552293886 | 0.395418013 | 0.478838129 |
| 8,800 | 0.480478825 | 0.551429233 | 0.392652523 | 0.476794992 |
| 8,900 | 0.479788544 | 0.550647626 | 0.393269869 | 0.47700581 |
| 9,000 | 0.478971917 | 0.549958069 | 0.392344022 | 0.476183159 |
| 9,100 | 0.478417137 | 0.55023121 | 0.389840335 | 0.475956038 |
| 9,200 | 0.47752396 | 0.54928952 | 0.388574111 | 0.474295435 |
| 9,300 | 0.476613898 | 0.549031964 | 0.388836214 | 0.474122589 |
| 9,400 | 0.475945772 | 0.546965499 | 0.390214831 | 0.472386731 |
| 9,500 | 0.475328426 | 0.546095294 | 0.386600239 | 0.472742296 |
| 9,600 | 0.474724597 | 0.5465163 | 0.384285548 | 0.472225543 |
| 9,700 | 0.4739397 | 0.545236203 | 0.384573312 | 0.472247793 |
| 9,800 | 0.473299769 | 0.54508388 | 0.381689642 | 0.471233651 |
| 9,900 | 0.472720767 | 0.544380275 | 0.382383087 | 0.472450344 |
| 10,000 | 0.471739424 | 0.542826753 | 0.381477 | 0.470170984 |
| 10,100 | 0.471286712 | 0.542297638 | 0.380559108 | 0.469776562 |
| 10,200 | 0.470540535 | 0.54183399 | 0.381574162 | 0.468411335 |
| 10,300 | 0.469974247 | 0.54274377 | 0.37924564 | 0.469146831 |
| 10,400 | 0.469287049 | 0.541755734 | 0.377209156 | 0.467076841 |
| 10,500 | 0.468645783 | 0.54191335 | 0.37796587 | 0.466812333 |
| 10,600 | 0.467831909 | 0.539803411 | 0.376898867 | 0.466988767 |
| 10,700 | 0.467166075 | 0.54051383 | 0.375538307 | 0.465493486 |
| 10,800 | 0.466336335 | 0.539371504 | 0.374870196 | 0.465047182 |
| 10,900 | 0.465921246 | 0.538801536 | 0.373945047 | 0.465316461 |
| 11,000 | 0.465193474 | 0.538531801 | 0.37203611 | 0.463231375 |
| 11,100 | 0.464727393 | 0.538609467 | 0.373916035 | 0.463228938 |
| 11,200 | 0.463837628 | 0.53639708 | 0.370614199 | 0.462856602 |
| 11,300 | 0.463356765 | 0.5367124 | 0.371004577 | 0.461203683 |
| 11,400 | 0.462783404 | 0.536245888 | 0.372299636 | 0.461640369 |
| 11,500 | 0.461980826 | 0.534992302 | 0.370673297 | 0.459603699 |
| 11,600 | 0.461470842 | 0.536217759 | 0.369607031 | 0.459801869 |
| 11,700 | 0.46101283 | 0.534412702 | 0.368989015 | 0.458201974 |
| 11,800 | 0.460192664 | 0.534258102 | 0.36766946 | 0.458456966 |
| 11,900 | 0.4598391 | 0.534155918 | 0.36738749 | 0.457905165 |
| 12,000 | 0.459003116 | 0.532886682 | 0.365368069 | 0.456417957 |
| 12,100 | 0.458675301 | 0.53288993 | 0.366033858 | 0.45623867 |
| 12,200 | 0.457884043 | 0.532112887 | 0.366063252 | 0.45543666 |
| 12,300 | 0.457609224 | 0.530952014 | 0.366374679 | 0.455935914 |

| | | | | |
|---|---|---|---|---|
| 12,400 | 0.456932118 | 0.531108583 | 0.364994794 | 0.454695486 |
| 12,500 | 0.456379956 | 0.531728916 | 0.366485968 | 0.453508875 |
| 12,600 | 0.455779062 | 0.530338091 | 0.364420034 | 0.453131018 |
| 12,700 | 0.455178427 | 0.528882914 | 0.363720052 | 0.452362064 |
| 12,800 | 0.454632496 | 0.528911383 | 0.361976358 | 0.451985577 |
| 12,900 | 0.453942062 | 0.52769689 | 0.359995545 | 0.451883353 |
| 13,000 | 0.453377171 | 0.528234022 | 0.360653858 | 0.451761146 |
| 13,100 | 0.452860775 | 0.527630432 | 0.3597536 | 0.451949003 |
| 13,200 | 0.452468538 | 0.526709874 | 0.355491401 | 0.450193057 |
| 13,300 | 0.451703362 | 0.527347815 | 0.357108634 | 0.449722982 |
| 13,400 | 0.451066894 | 0.526284871 | 0.354403458 | 0.449610261 |
| 13,500 | 0.450713164 | 0.525130657 | 0.3542931 | 0.449143781 |
| 13,600 | 0.450134726 | 0.525398379 | 0.353218903 | 0.448554119 |
| 13,700 | 0.449402671 | 0.524369929 | 0.350800482 | 0.447871611 |
| 13,800 | 0.448893222 | 0.523637338 | 0.352888945 | 0.447747755 |
| 13,900 | 0.448596037 | 0.52399513 | 0.351244034 | 0.447563229 |
| 14,000 | 0.447976548 | 0.522815668 | 0.348085821 | 0.447216234 |
| 14,100 | 0.447374756 | 0.522189106 | 0.350759368 | 0.446351137 |
| 14,200 | 0.447031119 | 0.520482824 | 0.352675523 | 0.445014295 |
| 14,300 | 0.446266277 | 0.519318964 | 0.350471819 | 0.444854613 |
| 14,400 | 0.446232755 | 0.520134089 | 0.348786385 | 0.444999272 |
| 14,500 | 0.445404079 | 0.519096698 | 0.347897833 | 0.443558963 |
| 14,600 | 0.444640425 | 0.519227171 | 0.346691316 | 0.443491401 |
| 14,700 | 0.444287668 | 0.517951123 | 0.345264624 | 0.442432116 |
| 14,800 | 0.4438847 | 0.517250695 | 0.344541451 | 0.440838452 |
| 14,900 | 0.443111779 | 0.516901634 | 0.345695536 | 0.440431757 |
| 15,000 | 0.442643243 | 0.516641901 | 0.344119679 | 0.440768204 |
| 15,100 | 0.442243115 | 0.515574873 | 0.342922736 | 0.440491241 |
| 15,200 | 0.441754114 | 0.516383268 | 0.346325964 | 0.439472942 |
| 15,300 | 0.441145826 | 0.517067422 | 0.34353267 | 0.438329025 |
| 15,400 | 0.440972403 | 0.515353161 | 0.341452779 | 0.43791832 |
| 15,500 | 0.440313853 | 0.514833596 | 0.343547053 | 0.437670942 |
| 15,600 | 0.439893461 | 0.513614867 | 0.344283867 | 0.437778332 |
| 15,700 | 0.439491389 | 0.514476227 | 0.342689249 | 0.436283844 |
| 15,800 | 0.438924729 | 0.512771025 | 0.344303026 | 0.435861941 |
| 15,900 | 0.438758587 | 0.512338941 | 0.342290195 | 0.436245682 |
| 16,000 | 0.437823796 | 0.511900976 | 0.340432824 | 0.435929316 |
| 16,100 | 0.437400455 | 0.511289616 | 0.340373338 | 0.435712611 |
| 16,200 | 0.436914416 | 0.511951766 | 0.341380921 | 0.434426394 |
| 16,300 | 0.436705527 | 0.5111664 | 0.341942609 | 0.434773811 |
| 16,400 | 0.436046714 | 0.511099229 | 0.340208446 | 0.434292843 |
| 16,500 | 0.436062804 | 0.511354316 | 0.339518303 | 0.43395544 |
| 16,600 | 0.435212502 | 0.51036561 | 0.338637382 | 0.433410024 |
| 16,700 | 0.434805542 | 0.509562331 | 0.337194126 | 0.433331702 |
| 16,800 | 0.434359494 | 0.509714296 | 0.335444407 | 0.432540228 |
| 16,900 | 0.433759317 | 0.509280121 | 0.335305247 | 0.432159854 |
| 17,000 | 0.433530922 | 0.50811742 | 0.335072402 | 0.432144039 |
| 17,100 | 0.432787119 | 0.50766239 | 0.334560368 | 0.431277487 |
| 17,200 | 0.43227908 | 0.507369864 | 0.334146728 | 0.431243652 |
| 17,300 | 0.432103577 | 0.508042064 | 0.333917509 | 0.429833043 |
| 17,400 | 0.431958938 | 0.507201763 | 0.334338084 | 0.430817131 |
| 17,500 | 0.431585848 | 0.507741332 | 0.334426451 | 0.430715837 |
| 17,600 | 0.431009877 | 0.507328759 | 0.332851799 | 0.429080241 |
| 17,700 | 0.430604024 | 0.50689699 | 0.331842033 | 0.430150303 |
| 17,800 | 0.430036931 | 0.506641461 | 0.331312261 | 0.429624317 |
| 17,900 | 0.429546447 | 0.506266397 | 0.328339912 | 0.427235705 |
| 18,000 | 0.428933249 | 0.504869663 | 0.330212323 | 0.427323417 |
| 18,100 | 0.428668738 | 0.505025734 | 0.330052933 | 0.427356434 |
| 18,200 | 0.428129257 | 0.50420479 | 0.329641731 | 0.426947818 |
| 18,300 | 0.427689525 | 0.504261705 | 0.327507951 | 0.426121288 |
| 18,400 | 0.42757747 | 0.503425229 | 0.32827784 | 0.426222844 |
| 18,500 | 0.426926041 | 0.503780465 | 0.328209507 | 0.425759768 |
| 18,600 | 0.426677278 | 0.50392475 | 0.327560106 | 0.425477361 |
| 18,700 | 0.426409512 | 0.503388255 | 0.325736089 | 0.425113694 |
| 18,800 | 0.425984211 | 0.503139224 | 0.323832146 | 0.42434307 |
| 18,900 | 0.425218246 | 0.503241292 | 0.32517264 | 0.424051576 |
| 19,000 | 0.424941201 | 0.502324861 | 0.321835037 | 0.424166359 |

| | | | | |
|---|---|---|---|---|
| 19,100 | 0.424662241 | 0.502789262 | 0.324258489 | 0.423928949 |
| 19,200 | 0.424130044 | 0.502022988 | 0.322435358 | 0.423983887 |
| 19,300 | 0.423786626 | 0.50210511 | 0.321977604 | 0.423749077 |
| 19,400 | 0.423562319 | 0.501969859 | 0.322501735 | 0.423405654 |
| 19,500 | 0.423226188 | 0.501786471 | 0.325106479 | 0.42321558 |
| 19,600 | 0.422677731 | 0.50117448 | 0.324263619 | 0.422324336 |
| 19,700 | 0.422313864 | 0.500864348 | 0.320888975 | 0.422311053 |
| 19,800 | 0.421883461 | 0.501389291 | 0.321101538 | 0.420541665 |
| 19,900 | 0.421420043 | 0.499739675 | 0.31898772 | 0.420169586 |
| 20,000 | 0.421012454 | 0.499350323 | 0.317475577 | 0.419045277 |
| 20,100 | 0.420748877 | 0.499943806 | 0.319227637 | 0.41793681 |
| 20,200 | 0.420103735 | 0.498777997 | 0.318374082 | 0.418212804 |
| 20,300 | 0.419843876 | 0.498707797 | 0.316660247 | 0.419201995 |
| 20,400 | 0.419780879 | 0.499048305 | 0.315923936 | 0.419497075 |
| 20,500 | 0.419054327 | 0.497524374 | 0.315898202 | 0.417902386 |
| 20,600 | 0.4187836 | 0.497549146 | 0.315693991 | 0.417174795 |
| 20,700 | 0.418360868 | 0.496615242 | 0.317596007 | 0.418282872 |
| 20,800 | 0.417889656 | 0.497265899 | 0.315000253 | 0.416682352 |
| 20,900 | 0.41775109 | 0.495994501 | 0.316612698 | 0.416664062 |
| 21,000 | 0.417280896 | 0.495446264 | 0.314923249 | 0.417388632 |
| 21,100 | 0.416898161 | 0.495172314 | 0.312749428 | 0.416553981 |
| 21,200 | 0.416465998 | 0.495191818 | 0.31428622 | 0.41526982 |
| 21,300 | 0.416034457 | 0.494398493 | 0.314619433 | 0.414559786 |
| 21,400 | 0.41584257 | 0.493728847 | 0.313338672 | 0.414731436 |
| 21,500 | 0.415635367 | 0.493084824 | 0.313626095 | 0.414920361 |
| 21,600 | 0.415282037 | 0.49248604 | 0.313247036 | 0.414201328 |
| 21,700 | 0.414479509 | 0.491512828 | 0.312280431 | 0.41358671 |
| 21,800 | 0.414032469 | 0.489832231 | 0.312608563 | 0.413380564 |
| 21,900 | 0.413862063 | 0.490493245 | 0.307345838 | 0.412992917 |
| 22,000 | 0.413605268 | 0.490461282 | 0.308265904 | 0.412316258 |
| 22,100 | 0.413161326 | 0.489937833 | 0.308552108 | 0.411687092 |
| 22,200 | 0.412990325 | 0.489632822 | 0.309442091 | 0.411757917 |
| 22,300 | 0.412675176 | 0.489479766 | 0.307662655 | 0.411492089 |
| 22,400 | 0.412420252 | 0.489976977 | 0.308440794 | 0.410924675 |
| 22,500 | 0.412021452 | 0.490622017 | 0.306750328 | 0.410797184 |
| 22,600 | 0.411518191 | 0.489278076 | 0.305462437 | 0.40998951 |
| 22,700 | 0.411420714 | 0.488407102 | 0.305674973 | 0.410186133 |
| 22,800 | 0.410993356 | 0.488222828 | 0.305317237 | 0.409601794 |
| 22,900 | 0.410371269 | 0.488007233 | 0.304258291 | 0.409602633 |
| 23,000 | 0.410418404 | 0.487666278 | 0.305939546 | 0.409336164 |
| 23,100 | 0.409857543 | 0.48709039 | 0.306794715 | 0.408284535 |
| 23,200 | 0.409594454 | 0.487145908 | 0.30746891 | 0.408067983 |
| 23,300 | 0.409206312 | 0.487222542 | 0.304554145 | 0.407666089 |
| 23,400 | 0.408668191 | 0.4848712 | 0.305477866 | 0.407652253 |
| 23,500 | 0.408474064 | 0.484691493 | 0.304008183 | 0.406763225 |
| 23,600 | 0.407988583 | 0.484451553 | 0.300244711 | 0.4075388 |
| 23,700 | 0.407886048 | 0.484613768 | 0.299527023 | 0.406900757 |
| 23,800 | 0.407632257 | 0.484857843 | 0.302489493 | 0.406347716 |
| 23,900 | 0.40705708 | 0.484369641 | 0.303417146 | 0.405037398 |
| 24,000 | 0.406858415 | 0.48389024 | 0.300942445 | 0.405303897 |
| 24,100 | 0.406752136 | 0.482486204 | 0.302856939 | 0.406214328 |
| 24,200 | 0.406166559 | 0.480675813 | 0.300966675 | 0.406371054 |
| 24,300 | 0.405943885 | 0.481492305 | 0.30125333 | 0.405298144 |
| 24,400 | 0.405458451 | 0.480797529 | 0.29930489 | 0.403915218 |
| 24,500 | 0.405114374 | 0.481216746 | 0.297633906 | 0.402355565 |
| 24,600 | 0.405008005 | 0.481372341 | 0.298666714 | 0.402723091 |
| 24,700 | 0.404784475 | 0.481270649 | 0.298136474 | 0.402283028 |
| 24,800 | 0.404338337 | 0.480419286 | 0.299399334 | 0.403178847 |
| 24,900 | 0.403989975 | 0.480814222 | 0.29848857 | 0.401936338 |
| 25,000 | 0.403737936 | 0.481169104 | 0.300030286 | 0.401896253 |
| 25,100 | 0.403569765 | 0.480806392 | 0.296657534 | 0.402460376 |
| 25,200 | 0.403439095 | 0.480110403 | 0.297223398 | 0.40181173 |
| 25,300 | 0.403040802 | 0.480294706 | 0.296085309 | 0.401270659 |
| 25,400 | 0.40266315 | 0.480014556 | 0.295862734 | 0.400998936 |
| 25,500 | 0.402289391 | 0.480044545 | 0.297571022 | 0.398886526 |
| 25,600 | 0.401862258 | 0.479399162 | 0.296295722 | 0.400179491 |
| 25,700 | 0.401597552 | 0.479227119 | 0.295232744 | 0.399291411 |

| | | | | |
|---|---|---|---|---|
| 25,800 | 0.401274939 | 0.477891589 | 0.295087052 | 0.399230366 |
| 25,900 | 0.400773539 | 0.477287142 | 0.295028505 | 0.397751615 |
| 26,000 | 0.400659647 | 0.477526591 | 0.293903742 | 0.39770293 |
| 26,100 | 0.400351612 | 0.477480635 | 0.293649965 | 0.397456175 |
| 26,200 | 0.400083333 | 0.477928447 | 0.291480952 | 0.398039696 |
| 26,300 | 0.399845461 | 0.477187925 | 0.291697702 | 0.397699173 |
| 26,400 | 0.399175292 | 0.477347912 | 0.293013289 | 0.396044177 |
| 26,500 | 0.399023158 | 0.477718988 | 0.291552713 | 0.394880998 |
| 26,600 | 0.39877033 | 0.476486791 | 0.289425504 | 0.395442433 |
| 26,700 | 0.398364457 | 0.477503113 | 0.291648654 | 0.39580919 |
| 26,800 | 0.397831678 | 0.475628292 | 0.289419484 | 0.394983444 |
| 26,900 | 0.397774396 | 0.476911199 | 0.28882939 | 0.395093839 |
| 27,000 | 0.39754132 | 0.475719265 | 0.289348417 | 0.394478027 |
| 27,100 | 0.397181539 | 0.475213867 | 0.288084322 | 0.393400294 |
| 27,200 | 0.397056286 | 0.476862694 | 0.289176571 | 0.394404911 |
| 27,300 | 0.396626301 | 0.474256242 | 0.288442658 | 0.392564823 |
| 27,400 | 0.396534734 | 0.474328501 | 0.289786147 | 0.392090375 |
| 27,500 | 0.396368572 | 0.473844842 | 0.290464799 | 0.391730552 |
| 27,600 | 0.395848317 | 0.473709352 | 0.289562171 | 0.391481945 |
| 27,700 | 0.395537238 | 0.473724895 | 0.289996739 | 0.390736775 |
| 27,800 | 0.395453124 | 0.473513663 | 0.288116504 | 0.39177221 |
| 27,900 | 0.395145185 | 0.473003412 | 0.287044225 | 0.390514166 |
| 28,000 | 0.394777359 | 0.473098101 | 0.288672286 | 0.390330236 |
| 28,100 | 0.394370292 | 0.47271919 | 0.287234939 | 0.389472577 |
| 28,200 | 0.394166038 | 0.472287172 | 0.287679548 | 0.389343667 |
| 28,300 | 0.394067348 | 0.472004695 | 0.287080569 | 0.389658378 |
| 28,400 | 0.393802574 | 0.471834436 | 0.289116102 | 0.389083773 |
| 28,500 | 0.393176223 | 0.470853486 | 0.288842101 | 0.388330919 |
| 28,600 | 0.393073473 | 0.470982898 | 0.285448546 | 0.389164839 |
| 28,700 | 0.393022125 | 0.471808298 | 0.285574132 | 0.389228958 |
| 28,800 | 0.392842483 | 0.470417493 | 0.285581578 | 0.38992185 |
| 28,900 | 0.392429123 | 0.470832284 | 0.287347343 | 0.388466001 |
| 29,000 | 0.392116432 | 0.469301812 | 0.287149554 | 0.388582632 |
| 29,100 | 0.391849819 | 0.469275485 | 0.287054725 | 0.388240574 |
| 29,200 | 0.39139036 | 0.469414562 | 0.286016601 | 0.387636543 |
| 29,300 | 0.39122847 | 0.469141743 | 0.286033881 | 0.387970061 |
| 29,400 | 0.390958468 | 0.469548759 | 0.285728418 | 0.387473536 |
| 29,500 | 0.390791846 | 0.468085284 | 0.285228511 | 0.3863887 |
| 29,600 | 0.390462331 | 0.467380438 | 0.28338254 | 0.386119785 |
| 29,700 | 0.390242833 | 0.467306121 | 0.280289082 | 0.385809118 |
| 29,800 | 0.389743827 | 0.466755328 | 0.284000947 | 0.384976319 |
| 29,900 | 0.389704463 | 0.467016381 | 0.284130601 | 0.386327147 |
| 30,000 | 0.389320949 | 0.467986967 | 0.282858617 | 0.384325686 |
| 30,100 | 0.389262131 | 0.46809791 | 0.281215043 | 0.385713613 |
| 30,200 | 0.388789932 | 0.467262306 | 0.286046162 | 0.38493832 |
| 30,300 | 0.388703927 | 0.46785742 | 0.2844609 | 0.383893502 |
| 30,400 | 0.388490421 | 0.468171914 | 0.283814571 | 0.383748387 |
| 30,500 | 0.388149776 | 0.467887548 | 0.282667976 | 0.383610397 |
| 30,600 | 0.387829899 | 0.466767759 | 0.282571333 | 0.382713664 |
| 30,700 | 0.387875677 | 0.467272848 | 0.281476891 | 0.383409752 |
| 30,800 | 0.387469299 | 0.466942072 | 0.279331219 | 0.3817331 |
| 30,900 | 0.387003175 | 0.465826033 | 0.280362207 | 0.381182593 |
| 31,000 | 0.386608228 | 0.465859791 | 0.278143264 | 0.380651047 |
| 31,100 | 0.386277937 | 0.465231007 | 0.278717001 | 0.381169918 |
| 31,200 | 0.386115627 | 0.465025598 | 0.279243772 | 0.381143418 |
| 31,300 | 0.385799808 | 0.464699144 | 0.278362599 | 0.380186725 |
| 31,400 | 0.385757452 | 0.465048883 | 0.280375368 | 0.381342538 |
| 31,500 | 0.385514938 | 0.464858052 | 0.277685384 | 0.379743122 |
| 31,600 | 0.385306235 | 0.463912989 | 0.275744349 | 0.379703732 |
| 31,700 | 0.385129874 | 0.464120687 | 0.276329567 | 0.379141572 |
| 31,800 | 0.384661181 | 0.464083758 | 0.277358413 | 0.379619373 |
| 31,900 | 0.38459955 | 0.464101909 | 0.278183246 | 0.379662562 |
| 32,000 | 0.384165402 | 0.463289899 | 0.276184498 | 0.37863203 |
| 32,100 | 0.384017632 | 0.464451255 | 0.276466963 | 0.378541185 |
| 32,200 | 0.383941704 | 0.463339034 | 0.277474093 | 0.378180506 |
| 32,300 | 0.383697701 | 0.463359665 | 0.277458173 | 0.378141306 |
| 32,400 | 0.383431603 | 0.463747065 | 0.277669813 | 0.377570214 |

| | | | | |
|---|---|---|---|---|
| 32,500 | 0.383215041 | 0.463536081 | 0.279174652 | 0.377078177 |
| 32,600 | 0.382890124 | 0.462638444 | 0.27610131 | 0.377953982 |
| 32,700 | 0.382724222 | 0.461736279 | 0.277273373 | 0.377015755 |
| 32,800 | 0.382599733 | 0.461581707 | 0.278373708 | 0.375073073 |
| 32,900 | 0.382125982 | 0.461752652 | 0.276633177 | 0.375077852 |
| 33,000 | 0.381781025 | 0.460893412 | 0.274473227 | 0.374944598 |
| 33,100 | 0.381506577 | 0.460961141 | 0.27645943 | 0.374438776 |
| 33,200 | 0.381361692 | 0.460429613 | 0.276319059 | 0.37565153 |
| 33,300 | 0.381116553 | 0.460274354 | 0.277243096 | 0.374704318 |
| 33,400 | 0.381166833 | 0.460847872 | 0.274870333 | 0.374901606 |
| 33,500 | 0.380888301 | 0.461058072 | 0.276213251 | 0.373799268 |
| 33,600 | 0.380761716 | 0.460408957 | 0.275024673 | 0.374418924 |
| 33,700 | 0.380621534 | 0.46148705 | 0.274311465 | 0.373501175 |
| 33,800 | 0.380456123 | 0.460488002 | 0.27506534 | 0.373892406 |
| 33,900 | 0.380215481 | 0.459621064 | 0.273362118 | 0.372953283 |
| 34,000 | 0.379961625 | 0.459703546 | 0.276201664 | 0.373364331 |
| 34,100 | 0.379801992 | 0.459226989 | 0.274548646 | 0.37300904 |
| 34,200 | 0.379483002 | 0.458684362 | 0.274453283 | 0.372534182 |
| 34,300 | 0.379082644 | 0.457850672 | 0.273817817 | 0.37196436 |
| 34,400 | 0.379049755 | 0.458771981 | 0.273860309 | 0.372380495 |
| 34,500 | 0.378694071 | 0.457560278 | 0.275110969 | 0.371543657 |
| 34,600 | 0.378399891 | 0.458278475 | 0.273914477 | 0.372214537 |
| 34,700 | 0.378369 | 0.458111568 | 0.27050441 | 0.371494781 |
| 34,800 | 0.377898364 | 0.456975339 | 0.271236776 | 0.371261216 |
| 34,900 | 0.377732159 | 0.456935666 | 0.272314891 | 0.372537448 |
| 35,000 | 0.377356607 | 0.456428438 | 0.269514217 | 0.371824434 |
| 35,100 | 0.377263751 | 0.456631569 | 0.269929624 | 0.370666335 |
| 35,200 | 0.377119005 | 0.457287437 | 0.270819553 | 0.370615154 |
| 35,300 | 0.37696601 | 0.456228449 | 0.269623367 | 0.371008314 |
| 35,400 | 0.376732598 | 0.456282484 | 0.270484496 | 0.37082633 |
| 35,500 | 0.376583462 | 0.456674426 | 0.267975312 | 0.371428379 |
| 35,600 | 0.376239949 | 0.45548194 | 0.268593498 | 0.369485817 |
| 35,700 | 0.376344187 | 0.457164753 | 0.268892705 | 0.369466727 |
| 35,800 | 0.37577192 | 0.456086379 | 0.270477088 | 0.368446158 |
| 35,900 | 0.375773324 | 0.455828206 | 0.270568534 | 0.368238747 |
| 36,000 | 0.375402229 | 0.455490547 | 0.26939992 | 0.367802439 |
| 36,100 | 0.375126958 | 0.454312492 | 0.268896266 | 0.367992705 |
| 36,200 | 0.374910821 | 0.454192582 | 0.270435953 | 0.367119868 |
| 36,300 | 0.374676448 | 0.453583974 | 0.268947109 | 0.366908938 |
| 36,400 | 0.374444581 | 0.452876668 | 0.268151543 | 0.367985971 |
| 36,500 | 0.374321044 | 0.453836078 | 0.268076954 | 0.367582175 |
| 36,600 | 0.374097608 | 0.453571842 | 0.269348917 | 0.366616923 |
| 36,700 | 0.373896018 | 0.452918754 | 0.267741873 | 0.36642054 |
| 36,800 | 0.373766807 | 0.452512114 | 0.267167442 | 0.366570066 |
| 36,900 | 0.373129993 | 0.452476929 | 0.267565513 | 0.365551451 |
| 37,000 | 0.373082494 | 0.452110495 | 0.26616376 | 0.365973133 |
| 37,100 | 0.372721235 | 0.451782008 | 0.265783594 | 0.365944795 |
| 37,200 | 0.372600734 | 0.452493461 | 0.26719675 | 0.365397507 |
| 37,300 | 0.372410386 | 0.451775316 | 0.266634058 | 0.366244523 |
| 37,400 | 0.372175103 | 0.451257541 | 0.265450701 | 0.365021458 |
| 37,500 | 0.372023681 | 0.452140256 | 0.263393135 | 0.366368165 |
| 37,600 | 0.371763604 | 0.452167057 | 0.26582689 | 0.3651791 |
| 37,700 | 0.371524719 | 0.452532683 | 0.262762896 | 0.364395128 |
| 37,800 | 0.371279624 | 0.451617539 | 0.262576707 | 0.364467709 |
| 37,900 | 0.371184737 | 0.451705714 | 0.263121599 | 0.364152677 |
| 38,000 | 0.371049316 | 0.451698356 | 0.261677135 | 0.363792099 |
| 38,100 | 0.370760979 | 0.451931589 | 0.261896814 | 0.362596689 |
| 38,200 | 0.370347968 | 0.451351501 | 0.260965837 | 0.363745745 |
| 38,300 | 0.370262705 | 0.451223989 | 0.26199268 | 0.362896685 |
| 38,400 | 0.370096653 | 0.450996438 | 0.261140231 | 0.362418552 |
| 38,500 | 0.36996041 | 0.450023091 | 0.26029066 | 0.362712158 |
| 38,600 | 0.369705954 | 0.449840797 | 0.261747895 | 0.362649989 |
| 38,700 | 0.369782822 | 0.449465575 | 0.260048246 | 0.361843491 |
| 38,800 | 0.369453263 | 0.450070314 | 0.261617627 | 0.362244801 |
| 38,900 | 0.369070553 | 0.449696922 | 0.259027596 | 0.362049441 |
| 39,000 | 0.368807164 | 0.449004107 | 0.260493217 | 0.361792974 |
| 39,100 | 0.368622697 | 0.44982497 | 0.261397405 | 0.361151666 |

| | | | | |
|---|---|---|---|---|
| 39,200 | 0.368283824 | 0.449053958 | 0.258020423 | 0.362157799 |
| 39,300 | 0.3681064 | 0.448518428 | 0.260749935 | 0.361636632 |
| 39,400 | 0.367640269 | 0.4476913 | 0.260843862 | 0.360826457 |
| 39,500 | 0.367552206 | 0.447989107 | 0.260038604 | 0.360093827 |
| 39,600 | 0.367450718 | 0.447406022 | 0.25974857 | 0.361002428 |
| 39,700 | 0.367184919 | 0.447024909 | 0.259560075 | 0.359889194 |
| 39,800 | 0.367177223 | 0.447402669 | 0.258588293 | 0.359922084 |
| 39,900 | 0.36682685 | 0.446239705 | 0.258126706 | 0.360439744 |
| 40,000 | 0.366670908 | 0.445956166 | 0.257310129 | 0.358911249 |
| 40,100 | 0.366447432 | 0.445922828 | 0.258431064 | 0.358506419 |
| 40,200 | 0.366432246 | 0.44584811 | 0.257924441 | 0.359426369 |
| 40,300 | 0.366232668 | 0.445599265 | 0.257743019 | 0.359439381 |
| 40,400 | 0.365995069 | 0.444556792 | 0.256119183 | 0.360692375 |
| 40,500 | 0.365990407 | 0.44494007 | 0.257826694 | 0.358577735 |
| 40,600 | 0.365778625 | 0.445312392 | 0.257118862 | 0.358155136 |
| 40,700 | 0.365369048 | 0.4441237 | 0.255799762 | 0.357789755 |
| 40,800 | 0.365110574 | 0.44493446 | 0.257196972 | 0.358123276 |
| 40,900 | 0.364814071 | 0.444275976 | 0.256505538 | 0.358318453 |
| 41,000 | 0.364558741 | 0.443287904 | 0.253659817 | 0.356640382 |
| 41,100 | 0.364628782 | 0.444379003 | 0.25520468 | 0.357898465 |
| 41,200 | 0.364540412 | 0.444117784 | 0.254848754 | 0.357668777 |
| 41,300 | 0.364235345 | 0.44386613 | 0.25493924 | 0.358018302 |
| 41,400 | 0.363869801 | 0.443404636 | 0.254090958 | 0.35592153 |
| 41,500 | 0.363762944 | 0.443540187 | 0.252378874 | 0.356808847 |
| 41,600 | 0.363673787 | 0.443097222 | 0.253624954 | 0.356879674 |
| 41,700 | 0.363643739 | 0.442812952 | 0.254189827 | 0.357165328 |
| 41,800 | 0.363371413 | 0.443608693 | 0.251379967 | 0.356038014 |
| 41,900 | 0.362878382 | 0.442398575 | 0.252060126 | 0.355577568 |
| 42,000 | 0.36270239 | 0.442839888 | 0.25013431 | 0.355297354 |
| 42,100 | 0.362667853 | 0.442153709 | 0.252354838 | 0.355849744 |
| 42,200 | 0.362390116 | 0.442950511 | 0.253549723 | 0.354962347 |
| 42,300 | 0.362357106 | 0.44322198 | 0.253130392 | 0.355725994 |
| 42,400 | 0.36214649 | 0.443356993 | 0.251875083 | 0.354316536 |
| 42,500 | 0.361901359 | 0.443139407 | 0.249236029 | 0.354426294 |
| 42,600 | 0.361788839 | 0.442655073 | 0.251695401 | 0.354226515 |
| 42,700 | 0.361544765 | 0.443844803 | 0.249768432 | 0.354413302 |
| 42,800 | 0.361280063 | 0.443833129 | 0.249665191 | 0.354305388 |
| 42,900 | 0.361275669 | 0.442172899 | 0.246896525 | 0.35479296 |
| 43,000 | 0.361044785 | 0.442466127 | 0.249182074 | 0.35411018 |
| 43,100 | 0.360923927 | 0.442080175 | 0.249154701 | 0.354329742 |
| 43,200 | 0.360719659 | 0.440850484 | 0.248289514 | 0.353384406 |
| 43,300 | 0.360448791 | 0.441201813 | 0.247526965 | 0.352933818 |
| 43,400 | 0.360202912 | 0.441117568 | 0.247846923 | 0.35276833 |
| 43,500 | 0.360205488 | 0.440539665 | 0.249538713 | 0.353065522 |
| 43,600 | 0.360140744 | 0.440514975 | 0.2476953 | 0.353177192 |
| 43,700 | 0.359754014 | 0.440375485 | 0.247921302 | 0.353067758 |
| 43,800 | 0.359414902 | 0.439966341 | 0.247506508 | 0.352961184 |
| 43,900 | 0.359218212 | 0.439899699 | 0.246946738 | 0.353522609 |
| 44,000 | 0.358983401 | 0.439282941 | 0.246755128 | 0.352222743 |
| 44,100 | 0.359074708 | 0.440103783 | 0.246122001 | 0.354211163 |
| 44,200 | 0.358948431 | 0.439382467 | 0.246830354 | 0.35395473 |
| 44,300 | 0.358607851 | 0.438331711 | 0.245223304 | 0.353667536 |
| 44,400 | 0.358185386 | 0.43864507 | 0.244472145 | 0.352570563 |
| 44,500 | 0.358216773 | 0.438085825 | 0.245071663 | 0.352358879 |
| 44,600 | 0.358007218 | 0.437964391 | 0.242503419 | 0.353355318 |
| 44,700 | 0.357928638 | 0.438129357 | 0.245120183 | 0.351441581 |
| 44,800 | 0.357485856 | 0.438967287 | 0.242536134 | 0.349868338 |
| 44,900 | 0.357410062 | 0.438096846 | 0.24366047 | 0.350283789 |
| 45,000 | 0.357445976 | 0.437769464 | 0.244422263 | 0.350550782 |
| 45,100 | 0.357107926 | 0.437455115 | 0.24237504 | 0.349746022 |
| 45,200 | 0.357019344 | 0.437713823 | 0.242084711 | 0.350229028 |
| 45,300 | 0.356738256 | 0.437281392 | 0.240002913 | 0.349290086 |
| 45,400 | 0.356597031 | 0.43681996 | 0.241258111 | 0.348798556 |
| 45,500 | 0.356494961 | 0.435408963 | 0.241729258 | 0.3493762 |
| 45,600 | 0.356435796 | 0.436418109 | 0.241231921 | 0.34963616 |
| 45,700 | 0.356020473 | 0.435790618 | 0.241966587 | 0.349217654 |
| 45,800 | 0.355947053 | 0.435542833 | 0.244319758 | 0.349656025 |

| | | | | |
|---|---|---|---|---|
| 45,900 | 0.355749855 | 0.434639516 | 0.242843467 | 0.34986672 |
| 46,000 | 0.355390297 | 0.434781667 | 0.243307815 | 0.348509157 |
| 46,100 | 0.355353809 | 0.435987176 | 0.242137362 | 0.348932479 |
| 46,200 | 0.355114205 | 0.435438803 | 0.243913601 | 0.347768966 |
| 46,300 | 0.355099426 | 0.434735958 | 0.242163156 | 0.348128389 |
| 46,400 | 0.354995318 | 0.434974774 | 0.242795812 | 0.347905051 |
| 46,500 | 0.354749309 | 0.43521705 | 0.241415157 | 0.346273344 |
| 46,600 | 0.354478224 | 0.435964915 | 0.241580726 | 0.346375189 |
| 46,700 | 0.354376795 | 0.434307261 | 0.239896313 | 0.346373324 |
| 46,800 | 0.354114214 | 0.43391578 | 0.241387185 | 0.346593475 |
| 46,900 | 0.353905501 | 0.434219811 | 0.240387049 | 0.346352047 |
| 47,000 | 0.353887374 | 0.433718712 | 0.239644827 | 0.346018177 |
| 47,100 | 0.35353906 | 0.432897042 | 0.240421641 | 0.345590542 |
| 47,200 | 0.353239462 | 0.43213236 | 0.240512324 | 0.345657092 |
| 47,300 | 0.353121312 | 0.432930189 | 0.240712969 | 0.346306686 |
| 47,400 | 0.352843734 | 0.432654556 | 0.242910659 | 0.344884894 |
| 47,500 | 0.352723724 | 0.432898499 | 0.241366669 | 0.344443262 |
| 47,600 | 0.352542293 | 0.431374378 | 0.239156122 | 0.344293966 |
| 47,700 | 0.35258737 | 0.432411802 | 0.239233547 | 0.346214169 |
| 47,800 | 0.35226263 | 0.431448056 | 0.241231066 | 0.346027784 |
| 47,900 | 0.352016139 | 0.431732392 | 0.241323801 | 0.343968636 |
| 48,000 | 0.351907252 | 0.431073106 | 0.241053636 | 0.345168718 |
| 48,100 | 0.351810276 | 0.431350066 | 0.239346302 | 0.344026355 |
| 48,200 | 0.35131904 | 0.430518848 | 0.240285054 | 0.344051341 |
| 48,300 | 0.351241137 | 0.429357416 | 0.241402744 | 0.342553537 |
| 48,400 | 0.351145437 | 0.429646937 | 0.240263751 | 0.342959585 |
| 48,500 | 0.350847691 | 0.42930351 | 0.238329665 | 0.343151191 |
| 48,600 | 0.350695393 | 0.429275688 | 0.238083781 | 0.343144098 |
| 48,700 | 0.350640081 | 0.430794379 | 0.236972268 | 0.342007484 |
| 48,800 | 0.35032968 | 0.430019409 | 0.239171119 | 0.341316739 |
| 48,900 | 0.350352602 | 0.43021233 | 0.240108201 | 0.341513299 |
| 49,000 | 0.350084703 | 0.429131083 | 0.238552723 | 0.340852319 |
| 49,100 | 0.350034744 | 0.429163195 | 0.239617165 | 0.340874633 |
| 49,200 | 0.349427636 | 0.428815025 | 0.238954676 | 0.340387202 |
| 49,300 | 0.349441481 | 0.429746765 | 0.240435231 | 0.341846442 |
| 49,400 | 0.349224602 | 0.428834532 | 0.240200966 | 0.342280499 |
| 49,500 | 0.349396279 | 0.428058559 | 0.239640736 | 0.342010241 |
| 49,600 | 0.348956969 | 0.42698004 | 0.238925899 | 0.341893142 |
| 49,700 | 0.34898918 | 0.427087658 | 0.239389645 | 0.341173048 |
| 49,800 | 0.348775335 | 0.427718033 | 0.237699917 | 0.340560683 |
| 49,900 | 0.348331057 | 0.42683706 | 0.237209132 | 0.341235685 |
| 50,000 | 0.348146919 | 0.426159826 | 0.23768642 | 0.341197867 |
| 50,100 | 0.348179156 | 0.426195776 | 0.236343648 | 0.341217133 |
| 50,200 | 0.347810869 | 0.426361339 | 0.23814981 | 0.341220971 |
| 50,300 | 0.34790486 | 0.425234039 | 0.237621037 | 0.340654469 |
| 50,400 | 0.347648602 | 0.426366705 | 0.236465321 | 0.340054184 |
| 50,500 | 0.347301193 | 0.424676774 | 0.238110109 | 0.341123362 |
| 50,600 | 0.347194196 | 0.42584809 | 0.23378483 | 0.340391252 |
| 50,700 | 0.346764027 | 0.424956936 | 0.236566236 | 0.340396526 |
| 50,800 | 0.346807632 | 0.424789601 | 0.235246903 | 0.34011811 |
| 50,900 | 0.346621735 | 0.424772806 | 0.234338679 | 0.339466986 |
| 51,000 | 0.346672046 | 0.424982229 | 0.234326789 | 0.340990822 |
| 51,100 | 0.346262186 | 0.423936456 | 0.232951448 | 0.339292875 |
| 51,200 | 0.346280062 | 0.424116106 | 0.23374004 | 0.339038331 |
| 51,300 | 0.346173519 | 0.424397346 | 0.233143559 | 0.338979555 |
| 51,400 | 0.34604502 | 0.423964243 | 0.23340876 | 0.338418076 |
| 51,500 | 0.345656158 | 0.423854941 | 0.233005769 | 0.337999936 |
| 51,600 | 0.345618738 | 0.424791945 | 0.232131901 | 0.339098946 |
| 51,700 | 0.345458656 | 0.42420374 | 0.231624115 | 0.338946396 |
| 51,800 | 0.345129916 | 0.42290582 | 0.232312222 | 0.338354789 |
| 51,900 | 0.344898517 | 0.423559778 | 0.230907303 | 0.338253979 |
| 52,000 | 0.344772851 | 0.422630171 | 0.232102147 | 0.338604022 |
| 52,100 | 0.344703423 | 0.42240725 | 0.231746614 | 0.338987649 |
| 52,200 | 0.344615258 | 0.422443408 | 0.231028233 | 0.338690781 |
| 52,300 | 0.344225807 | 0.422302071 | 0.230611798 | 0.337722556 |
| 52,400 | 0.344295186 | 0.421344674 | 0.229596088 | 0.338968248 |
| 52,500 | 0.343992328 | 0.421401018 | 0.230639201 | 0.337008352 |

| | | | | |
|---|---|---|---|---|
| 52,600 | 0.344088553 | 0.421357185 | 0.230121009 | 0.337475597 |
| 52,700 | 0.343801593 | 0.421263823 | 0.229781265 | 0.337959613 |
| 52,800 | 0.34376397 | 0.42229697 | 0.229332811 | 0.338280886 |
| 52,900 | 0.34345929 | 0.423170418 | 0.228001342 | 0.336828538 |
| 53,000 | 0.343131617 | 0.421740353 | 0.228042158 | 0.337541161 |
| 53,100 | 0.342898867 | 0.4226365 | 0.230033358 | 0.337273323 |
| 53,200 | 0.342918691 | 0.42258893 | 0.228201868 | 0.335912822 |
| 53,300 | 0.342526402 | 0.422682343 | 0.227804771 | 0.336789904 |
| 53,400 | 0.342316691 | 0.421177158 | 0.229327451 | 0.336595631 |
| 53,500 | 0.342206176 | 0.420830591 | 0.228807463 | 0.334994476 |
| 53,600 | 0.341963575 | 0.421337858 | 0.22918371 | 0.335310641 |
| 53,700 | 0.341918576 | 0.421803856 | 0.228637597 | 0.335227865 |
| 53,800 | 0.341839965 | 0.420626717 | 0.231912995 | 0.334790247 |
| 53,900 | 0.341650952 | 0.419748607 | 0.231497878 | 0.334296375 |
| 54,000 | 0.341505289 | 0.420141832 | 0.228974945 | 0.33467067 |
| 54,100 | 0.341413212 | 0.419587391 | 0.227563262 | 0.334184055 |
| 54,200 | 0.341538689 | 0.419429216 | 0.229494611 | 0.334724084 |
| 54,300 | 0.341396701 | 0.420729692 | 0.228592112 | 0.334384234 |
| 54,400 | 0.340901147 | 0.419990358 | 0.226104273 | 0.33448154 |
| 54,500 | 0.341046686 | 0.41957875 | 0.22694206 | 0.333980644 |
| 54,600 | 0.340656919 | 0.419619859 | 0.228203403 | 0.334622072 |
| 54,700 | 0.340481383 | 0.419160396 | 0.226699896 | 0.334898751 |
| 54,800 | 0.340192053 | 0.419423146 | 0.227212334 | 0.33402219 |
| 54,900 | 0.34004923 | 0.418883265 | 0.228402493 | 0.334814175 |
| 55,000 | 0.339820815 | 0.418007007 | 0.227324442 | 0.332790924 |
| 55,100 | 0.339805109 | 0.417229939 | 0.227558353 | 0.333350445 |
| 55,200 | 0.339324439 | 0.417617454 | 0.226210297 | 0.332686401 |
| 55,300 | 0.339525589 | 0.418438297 | 0.225577583 | 0.332891168 |
| 55,400 | 0.339253034 | 0.418547263 | 0.2252124 | 0.333324411 |
| 55,500 | 0.339132089 | 0.417610002 | 0.225949517 | 0.332319916 |
| 55,600 | 0.338920176 | 0.417635689 | 0.224692059 | 0.332286073 |
| 55,700 | 0.33884133 | 0.416946698 | 0.225250223 | 0.333154244 |
| 55,800 | 0.338745087 | 0.416503584 | 0.225906678 | 0.332187734 |
| 55,900 | 0.338556372 | 0.417201115 | 0.226269789 | 0.33151721 |
| 56,000 | 0.33851361 | 0.416143219 | 0.226691314 | 0.331700127 |
| 56,100 | 0.338363502 | 0.41616045 | 0.223473412 | 0.331972165 |
| 56,200 | 0.338027827 | 0.41591724 | 0.223556845 | 0.331779354 |
| 56,300 | 0.338060866 | 0.415373536 | 0.222812055 | 0.332251236 |
| 56,400 | 0.337913628 | 0.416057267 | 0.225003834 | 0.33088309 |
| 56,500 | 0.337884213 | 0.416182343 | 0.223291192 | 0.330944966 |
| 56,600 | 0.337672049 | 0.415677718 | 0.222629202 | 0.331465493 |
| 56,700 | 0.337415689 | 0.41504629 | 0.223187582 | 0.331076623 |
| 56,800 | 0.337140974 | 0.415413621 | 0.220485552 | 0.331138346 |
| 56,900 | 0.336857498 | 0.413406771 | 0.221976447 | 0.330358391 |
| 57,000 | 0.336794741 | 0.414292328 | 0.224725581 | 0.33035702 |
| 57,100 | 0.336891436 | 0.415009252 | 0.223074337 | 0.330391624 |
| 57,200 | 0.336613772 | 0.413948577 | 0.222440688 | 0.331524604 |
| 57,300 | 0.336619646 | 0.413960797 | 0.223061543 | 0.332083806 |
| 57,400 | 0.33620105 | 0.414351452 | 0.22272791 | 0.330756726 |
| 57,500 | 0.336145528 | 0.414475981 | 0.221631356 | 0.330875228 |
| 57,600 | 0.336053894 | 0.415017941 | 0.221685472 | 0.331618408 |
| 57,700 | 0.336022198 | 0.41411762 | 0.220884413 | 0.330519697 |
| 57,800 | 0.335881343 | 0.413843035 | 0.222622211 | 0.33080708 |
| 57,900 | 0.335435096 | 0.413963303 | 0.223288162 | 0.329736967 |
| 58,000 | 0.33529397 | 0.413409711 | 0.221307736 | 0.330297294 |
| 58,100 | 0.335381262 | 0.41326038 | 0.221690736 | 0.330599973 |
| 58,200 | 0.335302516 | 0.413774291 | 0.221661744 | 0.330212675 |
| 58,300 | 0.335185836 | 0.411974602 | 0.222694089 | 0.330404916 |
| 58,400 | 0.33503727 | 0.412366192 | 0.223815759 | 0.331120916 |
| 58,500 | 0.3347423 | 0.412470356 | 0.221708938 | 0.329508013 |
| 58,600 | 0.334738161 | 0.413160093 | 0.222137941 | 0.330596283 |
| 58,700 | 0.334598527 | 0.411953726 | 0.222930269 | 0.328676705 |
| 58,800 | 0.334300196 | 0.412628531 | 0.222966649 | 0.329886966 |
| 58,900 | 0.33431162 | 0.412059508 | 0.222550754 | 0.328753722 |
| 59,000 | 0.334129072 | 0.413038126 | 0.221521974 | 0.329307808 |
| 59,100 | 0.33397759 | 0.412590974 | 0.221588408 | 0.328765048 |
| 59,200 | 0.333984202 | 0.411621336 | 0.21868107 | 0.328391776 |

| | | | | |
|---|---|---|---|---|
| 59,300 | 0.333941343 | 0.411781628 | 0.222573544 | 0.328942848 |
| 59,400 | 0.333766537 | 0.413159665 | 0.22200928 | 0.329391262 |
| 59,500 | 0.33350359 | 0.412624966 | 0.221952594 | 0.329568414 |
| 59,600 | 0.333413167 | 0.413099653 | 0.224063935 | 0.328816534 |
| 59,700 | 0.333079874 | 0.412466644 | 0.222152274 | 0.328671318 |
| 59,800 | 0.33302312 | 0.412695569 | 0.222915117 | 0.328149904 |
| 59,900 | 0.332875545 | 0.411729632 | 0.222433083 | 0.328203748 |
| 60,000 | 0.332750043 | 0.413117328 | 0.221505358 | 0.32799614 |
| 60,100 | 0.332761222 | 0.413226445 | 0.22062832 | 0.329041612 |
| 60,200 | 0.332622299 | 0.411793989 | 0.221465377 | 0.328244968 |
| 60,300 | 0.332360133 | 0.412602316 | 0.218606026 | 0.327668337 |
| 60,400 | 0.332171768 | 0.411586943 | 0.218448676 | 0.328063149 |
| 60,500 | 0.332068333 | 0.412304074 | 0.221008277 | 0.328069489 |
| 60,600 | 0.331935074 | 0.413229491 | 0.220240583 | 0.326771747 |
| 60,700 | 0.331836016 | 0.412359361 | 0.220422199 | 0.32706934 |
| 60,800 | 0.331800327 | 0.412027578 | 0.222132207 | 0.326983352 |
| 60,900 | 0.331622753 | 0.412132767 | 0.220915495 | 0.326865049 |
| 61,000 | 0.33163303 | 0.412440248 | 0.219224296 | 0.327197909 |
| 61,100 | 0.331234217 | 0.411560014 | 0.219475152 | 0.326633221 |
| 61,200 | 0.331198544 | 0.411448395 | 0.219166801 | 0.325768505 |
| 61,300 | 0.331139978 | 0.410756174 | 0.220587013 | 0.32547056 |
| 61,400 | 0.331051252 | 0.412100616 | 0.220287561 | 0.326126481 |
| 61,500 | 0.330810523 | 0.410559264 | 0.220672649 | 0.325681496 |
| 61,600 | 0.330642256 | 0.410137048 | 0.218344475 | 0.325523037 |
| 61,700 | 0.330789048 | 0.411706685 | 0.218945646 | 0.325359192 |
| 61,800 | 0.330484503 | 0.411299066 | 0.219816268 | 0.324528086 |
| 61,900 | 0.330645809 | 0.410117965 | 0.220638481 | 0.325368747 |
| 62,000 | 0.330487881 | 0.410138774 | 0.221538536 | 0.32470307 |
| 62,100 | 0.330250199 | 0.41011133 | 0.219351696 | 0.324003726 |
| 62,200 | 0.330263373 | 0.410176161 | 0.221868069 | 0.323714827 |
| 62,300 | 0.329949811 | 0.409310162 | 0.222027688 | 0.324199541 |
| 62,400 | 0.32975829 | 0.410138894 | 0.22191087 | 0.323873852 |
| 62,500 | 0.329772132 | 0.409975485 | 0.22155713 | 0.324483888 |
| 62,600 | 0.329567112 | 0.41057614 | 0.219796689 | 0.324167828 |
| 62,700 | 0.329520405 | 0.410861943 | 0.219981203 | 0.323688232 |
| 62,800 | 0.329457828 | 0.410828279 | 0.219680918 | 0.324275169 |
| 62,900 | 0.329474191 | 0.409974735 | 0.219928651 | 0.323498166 |
| 63,000 | 0.329218891 | 0.409725265 | 0.21693359 | 0.324117493 |
| 63,100 | 0.328960475 | 0.40934927 | 0.218527406 | 0.323581149 |
| 63,200 | 0.328874928 | 0.409682688 | 0.219411173 | 0.322992336 |
| 63,300 | 0.328761058 | 0.409013276 | 0.22138291 | 0.323593896 |
| 63,400 | 0.328629015 | 0.408981273 | 0.222362871 | 0.323424228 |
| 63,500 | 0.328651391 | 0.408619085 | 0.220592689 | 0.323925284 |
| 63,600 | 0.328581106 | 0.407872889 | 0.220083617 | 0.325059426 |
| 63,700 | 0.328497566 | 0.408352328 | 0.220326649 | 0.323660437 |
| 63,800 | 0.328479538 | 0.408264743 | 0.219286895 | 0.32411222 |
| 63,900 | 0.328397582 | 0.408081956 | 0.219440931 | 0.322154011 |
| 64,000 | 0.327998225 | 0.408197208 | 0.218266434 | 0.323276419 |
| 64,100 | 0.328024185 | 0.408348341 | 0.218065774 | 0.323860035 |
| 64,200 | 0.327862199 | 0.40791313 | 0.217977389 | 0.322932042 |
| 64,300 | 0.327740927 | 0.408090654 | 0.216133135 | 0.322711549 |
| 64,400 | 0.327667237 | 0.407371559 | 0.216409923 | 0.323664241 |
| 64,500 | 0.327226095 | 0.407742072 | 0.217854563 | 0.323205612 |
| 64,600 | 0.327112189 | 0.407394679 | 0.218072481 | 0.321870847 |
| 64,700 | 0.327004952 | 0.407063349 | 0.21975138 | 0.322641405 |
| 64,800 | 0.326934373 | 0.405979491 | 0.219009885 | 0.323082874 |
| 64,900 | 0.326821271 | 0.405942091 | 0.220364139 | 0.322148947 |
| 65,000 | 0.326559754 | 0.406426455 | 0.21880175 | 0.321236513 |
| 65,100 | 0.326546977 | 0.407157429 | 0.218140324 | 0.321924107 |
| 65,200 | 0.326365307 | 0.407450239 | 0.216810494 | 0.32228017 |
| 65,300 | 0.326354498 | 0.406727077 | 0.217718818 | 0.321747898 |
| 65,400 | 0.326163841 | 0.406389423 | 0.216847573 | 0.321910343 |
| 65,500 | 0.326064234 | 0.405884753 | 0.216297931 | 0.321922837 |
| 65,600 | 0.32597925 | 0.406334688 | 0.218577289 | 0.32224581 |
| 65,700 | 0.325835885 | 0.406018732 | 0.219368885 | 0.321939538 |
| 65,800 | 0.325809576 | 0.406266154 | 0.219444947 | 0.320972155 |
| 65,900 | 0.325709534 | 0.406180233 | 0.218898819 | 0.320288623 |

| | | | | |
|---|---|---|---|---|
| 66,000 | 0.325738762 | 0.405869461 | 0.218636186 | 0.320731415 |
| 66,100 | 0.325503193 | 0.40671979 | 0.219666259 | 0.320382225 |
| 66,200 | 0.325509797 | 0.407396469 | 0.218083146 | 0.319820453 |
| 66,300 | 0.32534383 | 0.40595972 | 0.217729737 | 0.320122746 |
| 66,400 | 0.325100652 | 0.406445966 | 0.216577058 | 0.320288367 |
| 66,500 | 0.325018669 | 0.40514663 | 0.215267553 | 0.319806884 |
| 66,600 | 0.324853082 | 0.406235679 | 0.215019089 | 0.319736856 |
| 66,700 | 0.324563559 | 0.406255996 | 0.215209034 | 0.319943721 |
| 66,800 | 0.324545433 | 0.406021419 | 0.214798406 | 0.319177922 |
| 66,900 | 0.324415231 | 0.406464948 | 0.215338117 | 0.318763613 |
| 67,000 | 0.324289875 | 0.405233923 | 0.214582855 | 0.31888067 |
| 67,100 | 0.324042568 | 0.405276271 | 0.214013211 | 0.318899557 |
| 67,200 | 0.324062311 | 0.405103831 | 0.214885143 | 0.318300079 |
| 67,300 | 0.323942704 | 0.40568692 | 0.212651031 | 0.318209076 |
| 67,400 | 0.323794783 | 0.405329672 | 0.212764146 | 0.318232076 |
| 67,500 | 0.323487146 | 0.406386853 | 0.213213222 | 0.31746523 |
| 67,600 | 0.32340843 | 0.4058422 | 0.213396592 | 0.316775518 |
| 67,700 | 0.323420065 | 0.407111401 | 0.214860665 | 0.316503011 |
| 67,800 | 0.323238748 | 0.40633874 | 0.213774454 | 0.316464698 |
| 67,900 | 0.323046358 | 0.405299268 | 0.214393239 | 0.317513801 |
| 68,000 | 0.323057448 | 0.405311583 | 0.213689589 | 0.317029809 |
| 68,100 | 0.32292994 | 0.40446724 | 0.213966383 | 0.317203663 |
| 68,200 | 0.32285048 | 0.404488359 | 0.212791849 | 0.31561239 |
| 68,300 | 0.322802339 | 0.404267489 | 0.212902002 | 0.316223443 |
| 68,400 | 0.322521499 | 0.404202775 | 0.211522431 | 0.315824553 |
| 68,500 | 0.322609779 | 0.403173785 | 0.212067515 | 0.315675496 |
| 68,600 | 0.322426715 | 0.404477358 | 0.211392655 | 0.315646961 |
| 68,700 | 0.322067879 | 0.40389594 | 0.213350963 | 0.315355878 |
| 68,800 | 0.321996731 | 0.4041832 | 0.214400244 | 0.315559264 |
| 68,900 | 0.322006569 | 0.404027009 | 0.210880894 | 0.316320922 |
| 69,000 | 0.321805441 | 0.40406608 | 0.211600776 | 0.315580141 |
| 69,100 | 0.321684039 | 0.402164495 | 0.212290435 | 0.314264445 |
| 69,200 | 0.321876782 | 0.403014453 | 0.21350398 | 0.315702611 |
| 69,300 | 0.321493356 | 0.40332521 | 0.211293864 | 0.315286642 |
| 69,400 | 0.321336722 | 0.403204749 | 0.211417257 | 0.314867179 |
| 69,500 | 0.321259778 | 0.404017229 | 0.207816434 | 0.314359328 |
| 69,600 | 0.321244836 | 0.403539412 | 0.208908573 | 0.315235693 |
| 69,700 | 0.321014893 | 0.403518432 | 0.211121515 | 0.315026022 |
| 69,800 | 0.32086774 | 0.403226793 | 0.210316019 | 0.315229189 |
| 69,900 | 0.320643392 | 0.402154202 | 0.209962146 | 0.315242322 |
| 70,000 | 0.320651997 | 0.402664394 | 0.210068472 | 0.314640462 |
| 70,100 | 0.320450576 | 0.40225887 | 0.211192745 | 0.314707084 |
| 70,200 | 0.320040664 | 0.40076068 | 0.208595721 | 0.313962998 |
| 70,300 | 0.320007697 | 0.401554716 | 0.209518692 | 0.31524664 |
| 70,400 | 0.31979684 | 0.401519076 | 0.20898659 | 0.314681865 |
| 70,500 | 0.319748909 | 0.400457907 | 0.209666601 | 0.313465755 |
| 70,600 | 0.319904444 | 0.39985525 | 0.209387393 | 0.313708608 |
| 70,700 | 0.319621172 | 0.400015651 | 0.207752715 | 0.313860019 |
| 70,800 | 0.319531338 | 0.400731342 | 0.207129547 | 0.314142941 |
| 70,900 | 0.319463152 | 0.399536934 | 0.206546182 | 0.314110162 |
| 71,000 | 0.319259593 | 0.399202108 | 0.207678227 | 0.313847598 |
| 71,100 | 0.319142724 | 0.399326492 | 0.2074553 | 0.31323801 |
| 71,200 | 0.318933427 | 0.398225337 | 0.20794263 | 0.313240201 |
| 71,300 | 0.318893687 | 0.399670601 | 0.209519346 | 0.313467106 |
| 71,400 | 0.318877077 | 0.399796852 | 0.206881945 | 0.312864265 |
| 71,500 | 0.31875392 | 0.398931447 | 0.207881311 | 0.312262009 |
| 71,600 | 0.318640191 | 0.397040362 | 0.206460635 | 0.312344588 |
| 71,700 | 0.318552014 | 0.39821276 | 0.208071165 | 0.312927458 |
| 71,800 | 0.318388295 | 0.398059159 | 0.206369976 | 0.312703594 |
| 71,900 | 0.318259291 | 0.397975471 | 0.205849193 | 0.31314943 |
| 72,000 | 0.318152415 | 0.399009835 | 0.206615703 | 0.312483363 |
| 72,100 | 0.318061399 | 0.399421439 | 0.205399435 | 0.312592656 |
| 72,200 | 0.318208517 | 0.399043952 | 0.208202312 | 0.312938282 |
| 72,300 | 0.31813009 | 0.397958419 | 0.205504088 | 0.313239827 |
| 72,400 | 0.318010212 | 0.398623836 | 0.205575638 | 0.312836231 |
| 72,500 | 0.317863672 | 0.398460898 | 0.204123285 | 0.312525769 |
| 72,600 | 0.317519776 | 0.397710128 | 0.206312793 | 0.311463099 |

| | | | | |
|---|---|---|---|---|
| 72,700 | 0.31752285 | 0.398232906 | 0.205638984 | 0.311420273 |
| 72,800 | 0.317567855 | 0.398941951 | 0.204212775 | 0.310698212 |
| 72,900 | 0.317206962 | 0.39878111 | 0.202748402 | 0.311013662 |
| 73,000 | 0.317099435 | 0.397316506 | 0.204796744 | 0.310467679 |
| 73,100 | 0.316982995 | 0.397850213 | 0.203089129 | 0.310647715 |
| 73,200 | 0.316819142 | 0.397234812 | 0.204019783 | 0.311255416 |
| 73,300 | 0.316825829 | 0.396826841 | 0.203268984 | 0.310848623 |
| 73,400 | 0.316882207 | 0.397984963 | 0.20456706 | 0.309844934 |
| 73,500 | 0.31652388 | 0.396233026 | 0.205047404 | 0.311373663 |
| 73,600 | 0.316425145 | 0.398164472 | 0.203689378 | 0.309911747 |
| 73,700 | 0.316240339 | 0.396532876 | 0.205397773 | 0.309819818 |
| 73,800 | 0.316076406 | 0.396240754 | 0.202621187 | 0.310046627 |
| 73,900 | 0.316136152 | 0.396863058 | 0.204215667 | 0.310066467 |
| 74,000 | 0.316000296 | 0.395492849 | 0.202547249 | 0.309942632 |
| 74,100 | 0.315629749 | 0.396674467 | 0.203615119 | 0.310358916 |
| 74,200 | 0.31562558 | 0.395636558 | 0.204024004 | 0.30947609 |
| 74,300 | 0.315629473 | 0.393923385 | 0.204590245 | 0.310569116 |
| 74,400 | 0.315399871 | 0.395567668 | 0.204578058 | 0.309535261 |
| 74,500 | 0.315337778 | 0.394874838 | 0.203750197 | 0.308540656 |
| 74,600 | 0.315371331 | 0.395315848 | 0.203319528 | 0.308617525 |
| 74,700 | 0.315199 | 0.394326779 | 0.202547165 | 0.308442068 |
| 74,800 | 0.315113603 | 0.394558842 | 0.204636211 | 0.308201203 |
| 74,900 | 0.315116421 | 0.394524264 | 0.202499022 | 0.30832557 |
| 75,000 | 0.315018318 | 0.394347539 | 0.203311151 | 0.309460874 |
| 75,100 | 0.314794459 | 0.394776371 | 0.201426136 | 0.308355316 |
| 75,200 | 0.314750003 | 0.393976449 | 0.200946804 | 0.307776441 |
| 75,300 | 0.314601344 | 0.394393469 | 0.203474273 | 0.307436404 |
| 75,400 | 0.314546776 | 0.394729478 | 0.204054322 | 0.308219351 |
| 75,500 | 0.314418477 | 0.39350885 | 0.203841324 | 0.307311552 |
| 75,600 | 0.314172593 | 0.393219576 | 0.201542864 | 0.308017659 |
| 75,700 | 0.314119505 | 0.392812589 | 0.20234103 | 0.308159416 |
| 75,800 | 0.313902908 | 0.392567995 | 0.200972168 | 0.307769403 |
| 75,900 | 0.31377328 | 0.39294014 | 0.201508376 | 0.307227676 |
| 76,000 | 0.313715655 | 0.392118213 | 0.200478761 | 0.307371807 |
| 76,100 | 0.313712766 | 0.392589572 | 0.201542836 | 0.307151456 |
| 76,200 | 0.313552597 | 0.393122166 | 0.200344251 | 0.307316196 |
| 76,300 | 0.313505872 | 0.391521763 | 0.202422437 | 0.3073475 |
| 76,400 | 0.313341965 | 0.391939495 | 0.202552651 | 0.306229602 |
| 76,500 | 0.313158249 | 0.390992158 | 0.20062132 | 0.306769864 |
| 76,600 | 0.313140021 | 0.391326441 | 0.200659762 | 0.306835737 |
| 76,700 | 0.313252788 | 0.391070391 | 0.201344555 | 0.306442918 |
| 76,800 | 0.313001716 | 0.390534188 | 0.202066958 | 0.305537328 |
| 76,900 | 0.31296212 | 0.391034576 | 0.201621686 | 0.305739578 |
| 77,000 | 0.312607874 | 0.390649261 | 0.201618519 | 0.30607537 |
| 77,100 | 0.312593911 | 0.390992699 | 0.201896245 | 0.306703481 |
| 77,200 | 0.312408438 | 0.390351992 | 0.201169981 | 0.306509692 |
| 77,300 | 0.312037399 | 0.389876324 | 0.201237187 | 0.305618491 |
| 77,400 | 0.312051965 | 0.390278103 | 0.200510249 | 0.305774467 |
| 77,500 | 0.311934198 | 0.391142109 | 0.200165481 | 0.304911374 |
| 77,600 | 0.312134916 | 0.391175342 | 0.201845692 | 0.304610653 |
| 77,700 | 0.311657312 | 0.390295761 | 0.201175625 | 0.30571437 |
| 77,800 | 0.311670354 | 0.390054685 | 0.202354469 | 0.30578571 |
| 77,900 | 0.311406018 | 0.390302454 | 0.200390803 | 0.305935075 |
| 78,000 | 0.311244956 | 0.389403582 | 0.200036539 | 0.305564903 |
| 78,100 | 0.311273327 | 0.388460824 | 0.198578809 | 0.304545848 |
| 78,200 | 0.311256001 | 0.388545893 | 0.200285344 | 0.304148173 |
| 78,300 | 0.311130386 | 0.390206015 | 0.200245272 | 0.303867183 |
| 78,400 | 0.311038504 | 0.390451156 | 0.199898752 | 0.3054219 |
| 78,500 | 0.310850142 | 0.389180698 | 0.200009816 | 0.304224733 |
| 78,600 | 0.310725756 | 0.390035476 | 0.201559618 | 0.303682062 |
| 78,700 | 0.310714212 | 0.389009976 | 0.199444294 | 0.303568075 |
| 78,800 | 0.310376804 | 0.389534009 | 0.200867305 | 0.30468806 |
| 78,900 | 0.310382596 | 0.389739024 | 0.200774079 | 0.304803445 |
| 79,000 | 0.310183178 | 0.389434562 | 0.200950106 | 0.304087517 |
| 79,100 | 0.309938344 | 0.389564824 | 0.200837421 | 0.304489449 |
| 79,200 | 0.309912573 | 0.388172973 | 0.199490941 | 0.304085777 |
| 79,300 | 0.309827438 | 0.38882018 | 0.199844518 | 0.303766109 |

| | | | | |
|---|---|---|---|---|
| 79,400 | 0.309667407 | 0.388872965 | 0.20211339 | 0.30233986 |
| 79,500 | 0.309593776 | 0.388942986 | 0.201226206 | 0.30236114 |
| 79,600 | 0.309435502 | 0.389042733 | 0.198077514 | 0.302482872 |
| 79,700 | 0.309416318 | 0.389188666 | 0.200105973 | 0.301995325 |
| 79,800 | 0.309311666 | 0.388430099 | 0.202046365 | 0.30173158 |
| 79,900 | 0.309378819 | 0.388806375 | 0.199088347 | 0.302314231 |
| 80,000 | 0.309028 | 0.388050311 | 0.200552808 | 0.301358378 |
| 80,100 | 0.308945437 | 0.388572093 | 0.200194985 | 0.301914515 |
| 80,200 | 0.309001134 | 0.389399809 | 0.200944515 | 0.302145446 |
| 80,300 | 0.30892621 | 0.388905545 | 0.200295207 | 0.302825452 |
| 80,400 | 0.308813807 | 0.388916387 | 0.197629248 | 0.30194285 |
| 80,500 | 0.308401537 | 0.387623067 | 0.199659865 | 0.301358025 |
| 80,600 | 0.308449498 | 0.388301418 | 0.198102318 | 0.301141619 |
| 80,700 | 0.308624881 | 0.38860531 | 0.197907793 | 0.302120738 |
| 80,800 | 0.3083673 | 0.387718283 | 0.198436355 | 0.301125442 |
| 80,900 | 0.308268503 | 0.388741478 | 0.198763938 | 0.300632222 |
| 81,000 | 0.308080681 | 0.388135614 | 0.197869912 | 0.301042441 |
| 81,100 | 0.308018473 | 0.387461978 | 0.199243757 | 0.300220428 |
| 81,200 | 0.308045888 | 0.388664478 | 0.197797762 | 0.299287677 |
| 81,300 | 0.307671657 | 0.387555996 | 0.198079006 | 0.299644904 |
| 81,400 | 0.307517742 | 0.387059166 | 0.199894101 | 0.300409741 |
| 81,500 | 0.307335059 | 0.387001814 | 0.197699596 | 0.300390503 |
| 81,600 | 0.307372107 | 0.387269077 | 0.199146961 | 0.299803046 |
| 81,700 | 0.307313197 | 0.387170244 | 0.198464048 | 0.29967651 |
| 81,800 | 0.307276342 | 0.387716603 | 0.19854293 | 0.299547929 |
| 81,900 | 0.307293317 | 0.387799582 | 0.197799277 | 0.299759018 |
| 82,000 | 0.307125803 | 0.387744997 | 0.197565886 | 0.298986103 |
| 82,100 | 0.307108554 | 0.387506712 | 0.199629882 | 0.298194931 |
| 82,200 | 0.306912865 | 0.387702813 | 0.199011666 | 0.298469544 |
| 82,300 | 0.306914199 | 0.386678043 | 0.198886012 | 0.299737961 |
| 82,400 | 0.306618701 | 0.387107099 | 0.198188233 | 0.299142625 |
| 82,500 | 0.306434099 | 0.386636253 | 0.196885555 | 0.298039731 |
| 82,600 | 0.306344937 | 0.386538813 | 0.198278216 | 0.297811975 |
| 82,700 | 0.306322038 | 0.386588989 | 0.19860998 | 0.298568755 |
| 82,800 | 0.306360579 | 0.386299019 | 0.198076402 | 0.298784283 |
| 82,900 | 0.305987299 | 0.385873256 | 0.19870212 | 0.297618236 |
| 83,000 | 0.305837265 | 0.386497264 | 0.197383365 | 0.296971004 |
| 83,100 | 0.305792101 | 0.384713117 | 0.195907303 | 0.297697462 |
| 83,200 | 0.305877123 | 0.386133002 | 0.197671239 | 0.297938062 |
| 83,300 | 0.30563071 | 0.386135003 | 0.19549596 | 0.297468387 |
| 83,400 | 0.305588673 | 0.386103308 | 0.196927339 | 0.29682447 |
| 83,500 | 0.305543275 | 0.385947721 | 0.198150782 | 0.29708742 |
| 83,600 | 0.305367847 | 0.385946753 | 0.198306852 | 0.297267546 |
| 83,700 | 0.305493419 | 0.384997654 | 0.197116146 | 0.297338279 |
| 83,800 | 0.30517586 | 0.385734204 | 0.198262155 | 0.295572449 |
| 83,900 | 0.304962511 | 0.385419915 | 0.19727057 | 0.296101165 |
| 84,000 | 0.305025026 | 0.385871679 | 0.197716695 | 0.296731072 |
| 84,100 | 0.30475579 | 0.384989865 | 0.198612979 | 0.295953417 |
| 84,200 | 0.30483939 | 0.384905833 | 0.197345156 | 0.296651577 |
| 84,300 | 0.304697676 | 0.385629408 | 0.195648648 | 0.296021838 |
| 84,400 | 0.304520234 | 0.385248927 | 0.196940723 | 0.29590213 |
| 84,500 | 0.304356668 | 0.384197055 | 0.196886086 | 0.295624809 |
| 84,600 | 0.304435379 | 0.384386693 | 0.19692734 | 0.294951527 |
| 84,700 | 0.304270875 | 0.385303014 | 0.196924629 | 0.294618133 |
| 84,800 | 0.304128174 | 0.384501869 | 0.197415727 | 0.294413186 |
| 84,900 | 0.304035608 | 0.384857843 | 0.198413038 | 0.295100491 |
| 85,000 | 0.304021937 | 0.384748115 | 0.199045128 | 0.29421123 |
| 85,100 | 0.303923916 | 0.384014551 | 0.197243137 | 0.29410569 |
| 85,200 | 0.303754151 | 0.383359909 | 0.197299702 | 0.29453927 |
| 85,300 | 0.303821135 | 0.38317016 | 0.197548299 | 0.294377661 |
| 85,400 | 0.303789295 | 0.38407083 | 0.19642553 | 0.294595983 |
| 85,500 | 0.303671539 | 0.382524529 | 0.197272845 | 0.294145472 |
| 85,600 | 0.303521708 | 0.382545213 | 0.198025947 | 0.294514786 |
| 85,700 | 0.303336126 | 0.382953793 | 0.195254782 | 0.29428179 |
| 85,800 | 0.303178062 | 0.382360472 | 0.195434522 | 0.293868915 |
| 85,900 | 0.303125355 | 0.383197266 | 0.194866148 | 0.294560234 |
| 86,000 | 0.302880096 | 0.382758593 | 0.196304794 | 0.293117412 |

| | | | | |
|---|---|---|---|---|
| 86,100 | 0.303030206 | 0.382257808 | 0.197052904 | 0.29384888 |
| 86,200 | 0.302982471 | 0.381722514 | 0.195195938 | 0.293961362 |
| 86,300 | 0.3025753 | 0.381171615 | 0.194985734 | 0.293876067 |
| 86,400 | 0.302778226 | 0.382475354 | 0.194607128 | 0.293823145 |
| 86,500 | 0.30258946 | 0.381467312 | 0.193519681 | 0.293579242 |
| 86,600 | 0.302814759 | 0.382534728 | 0.194701658 | 0.29404402 |
| 86,700 | 0.302446797 | 0.381875212 | 0.193438608 | 0.291777065 |
| 86,800 | 0.302268909 | 0.381105712 | 0.195237312 | 0.292070562 |
| 86,900 | 0.302347233 | 0.381646173 | 0.19356373 | 0.293181299 |
| 87,000 | 0.302086072 | 0.382560172 | 0.193453903 | 0.292966147 |
| 87,100 | 0.301859165 | 0.381030854 | 0.192811431 | 0.292225036 |
| 87,200 | 0.301625152 | 0.381725283 | 0.192700651 | 0.291995671 |
| 87,300 | 0.301744311 | 0.380968555 | 0.193044659 | 0.291943228 |
| 87,400 | 0.301692692 | 0.380511249 | 0.195011395 | 0.292333504 |
| 87,500 | 0.301446263 | 0.380666804 | 0.192992558 | 0.290938898 |
| 87,600 | 0.30150903 | 0.38249807 | 0.191999753 | 0.292545914 |
| 87,700 | 0.301426546 | 0.38187906 | 0.191319638 | 0.292300835 |
| 87,800 | 0.301332035 | 0.381157695 | 0.19273367 | 0.291394864 |
| 87,900 | 0.301318284 | 0.381189667 | 0.19511047 | 0.291989833 |
| 88,000 | 0.30133442 | 0.380049541 | 0.194931045 | 0.29281372 |
| 88,100 | 0.301077622 | 0.379590578 | 0.192530585 | 0.292241868 |
| 88,200 | 0.300972883 | 0.381768347 | 0.191841374 | 0.291845787 |
| 88,300 | 0.30100488 | 0.380871839 | 0.19093966 | 0.291938271 |
| 88,400 | 0.300892237 | 0.379900365 | 0.191842461 | 0.291959723 |
| 88,500 | 0.30062154 | 0.38024732 | 0.190554165 | 0.291042506 |
| 88,600 | 0.300506135 | 0.380919908 | 0.192440204 | 0.290886295 |
| 88,700 | 0.30063491 | 0.380833225 | 0.192309989 | 0.291196877 |
| 88,800 | 0.300632419 | 0.380356805 | 0.191424218 | 0.291765408 |
| 88,900 | 0.300644309 | 0.380885573 | 0.192373428 | 0.290772814 |
| 89,000 | 0.30044913 | 0.38084901 | 0.190903059 | 0.29083119 |
| 89,100 | 0.300478132 | 0.380224955 | 0.191185518 | 0.290093528 |
| 89,200 | 0.30026244 | 0.381297999 | 0.191255118 | 0.290908998 |
| 89,300 | 0.30016054 | 0.380444248 | 0.19280345 | 0.290147612 |
| 89,400 | 0.299975093 | 0.380761127 | 0.192123548 | 0.290457489 |
| 89,500 | 0.299816938 | 0.380393039 | 0.190312592 | 0.290494831 |
| 89,600 | 0.299847843 | 0.38096337 | 0.192032099 | 0.289966588 |
| 89,700 | 0.299705764 | 0.379600779 | 0.190412502 | 0.289968952 |
| 89,800 | 0.299668766 | 0.380012089 | 0.191178356 | 0.29043501 |
| 89,900 | 0.299433049 | 0.380036872 | 0.189185114 | 0.289794716 |
| 90,000 | 0.299287523 | 0.379291932 | 0.190104636 | 0.288572467 |
| 90,100 | 0.299362244 | 0.379660164 | 0.189670631 | 0.287846595 |
| 90,200 | 0.299192827 | 0.380192991 | 0.191556182 | 0.288048982 |
| 90,300 | 0.299152712 | 0.379943078 | 0.191312295 | 0.288437453 |
| 90,400 | 0.298979984 | 0.379897704 | 0.192467293 | 0.288190251 |
| 90,500 | 0.299157218 | 0.38093618 | 0.190599352 | 0.288551127 |
| 90,600 | 0.299054515 | 0.379766205 | 0.191878672 | 0.288471693 |
| 90,700 | 0.29888153 | 0.37969212 | 0.19126299 | 0.288569808 |
| 90,800 | 0.29878631 | 0.380179708 | 0.193534659 | 0.28887222 |
| 90,900 | 0.298775964 | 0.380434401 | 0.194305743 | 0.288287767 |
| 91,000 | 0.29861896 | 0.380488697 | 0.192714479 | 0.288531289 |
| 91,100 | 0.298565652 | 0.380144312 | 0.191075202 | 0.287797124 |
| 91,200 | 0.298471229 | 0.38018698 | 0.191038203 | 0.287227366 |
| 91,300 | 0.298297674 | 0.378722187 | 0.191407818 | 0.286791237 |
| 91,400 | 0.298437467 | 0.379865601 | 0.190048839 | 0.287355229 |
| 91,500 | 0.298178976 | 0.379704593 | 0.191957538 | 0.287447217 |
| 91,600 | 0.298206153 | 0.379356283 | 0.192077438 | 0.286662655 |
| 91,700 | 0.298348252 | 0.378732476 | 0.19322392 | 0.28631048 |
| 91,800 | 0.297932515 | 0.378603141 | 0.190282092 | 0.286955778 |
| 91,900 | 0.297916801 | 0.37901206 | 0.190698758 | 0.286329809 |
| 92,000 | 0.297870504 | 0.378409193 | 0.192821203 | 0.28757668 |
| 92,100 | 0.297842457 | 0.378689454 | 0.192389632 | 0.285536817 |
| 92,200 | 0.297474154 | 0.37869528 | 0.191704198 | 0.285942802 |
| 92,300 | 0.297634542 | 0.379329583 | 0.190379364 | 0.285695597 |
| 92,400 | 0.297378942 | 0.378584423 | 0.190958415 | 0.286024312 |
| 92,500 | 0.297320617 | 0.378330645 | 0.189580666 | 0.285559097 |
| 92,600 | 0.297237725 | 0.378326427 | 0.190123968 | 0.284885528 |
| 92,700 | 0.296921199 | 0.377804701 | 0.19086666 | 0.285523709 |

| | | | | |
|---|---|---|---|---|
| 92,800 | 0.296706266 | 0.377700155 | 0.189707253 | 0.284637411 |
| 92,900 | 0.296575992 | 0.377603663 | 0.190805158 | 0.284908059 |
| 93,000 | 0.296681986 | 0.377123898 | 0.190358782 | 0.285229074 |
| 93,100 | 0.296738869 | 0.376991926 | 0.190935302 | 0.285165311 |
| 93,200 | 0.296796028 | 0.377048232 | 0.190192963 | 0.286612557 |
| 93,300 | 0.296522613 | 0.378347357 | 0.190339422 | 0.28514607 |
| 93,400 | 0.296189583 | 0.377688049 | 0.188747232 | 0.285069286 |
| 93,500 | 0.296320257 | 0.377857733 | 0.191444768 | 0.284304102 |
| 93,600 | 0.296311119 | 0.377709285 | 0.190517748 | 0.284069539 |
| 93,700 | 0.29595699 | 0.378158135 | 0.188621922 | 0.283603458 |
| 93,800 | 0.295770611 | 0.37757524 | 0.189687934 | 0.283501552 |
| 93,900 | 0.295684377 | 0.377661108 | 0.189136786 | 0.283643058 |
| 94,000 | 0.295855794 | 0.377613635 | 0.189722292 | 0.283928003 |
| 94,100 | 0.29583715 | 0.37731893 | 0.189909518 | 0.285118297 |
| 94,200 | 0.295599262 | 0.376827124 | 0.189413477 | 0.284433651 |
| 94,300 | 0.295483091 | 0.377212173 | 0.189178056 | 0.284004802 |
| 94,400 | 0.295532351 | 0.376901557 | 0.188445922 | 0.284316949 |
| 94,500 | 0.295264363 | 0.376579207 | 0.189418741 | 0.28423994 |
| 94,600 | 0.295257814 | 0.376355612 | 0.192154775 | 0.284854328 |
| 94,700 | 0.295358192 | 0.375675073 | 0.187673189 | 0.284560412 |
| 94,800 | 0.295003695 | 0.374930139 | 0.188985528 | 0.284686109 |
| 94,900 | 0.295186343 | 0.375801309 | 0.187789945 | 0.284655131 |
| 95,000 | 0.29507348 | 0.376159823 | 0.187768828 | 0.284143719 |
| 95,100 | 0.294975527 | 0.375723429 | 0.189697819 | 0.283111541 |
| 95,200 | 0.294767233 | 0.376466658 | 0.188629664 | 0.284461165 |
| 95,300 | 0.294675975 | 0.376464205 | 0.188345469 | 0.284185274 |
| 95,400 | 0.294644615 | 0.376896035 | 0.188049867 | 0.28416628 |
| 95,500 | 0.294550557 | 0.37661932 | 0.188046518 | 0.284505197 |
| 95,600 | 0.294534918 | 0.375650504 | 0.189996765 | 0.284552287 |
| 95,700 | 0.294316284 | 0.377232661 | 0.18731469 | 0.283965079 |
| 95,800 | 0.294256851 | 0.375899841 | 0.186462173 | 0.284228788 |
| 95,900 | 0.294262221 | 0.377148646 | 0.18635287 | 0.284116055 |
| 96,000 | 0.294191045 | 0.375569426 | 0.187419123 | 0.283709715 |
| 96,100 | 0.294023188 | 0.375316046 | 0.18506089 | 0.284000227 |
| 96,200 | 0.293991017 | 0.375174544 | 0.186895221 | 0.283462528 |
| 96,300 | 0.29393433 | 0.374779603 | 0.18776425 | 0.284010938 |
| 96,400 | 0.294067667 | 0.37540032 | 0.188560661 | 0.28474373 |
| 96,500 | 0.29380386 | 0.374714419 | 0.18807419 | 0.284573915 |
| 96,600 | 0.293965868 | 0.376059448 | 0.187213361 | 0.28353177 |
| 96,700 | 0.293593623 | 0.375489119 | 0.187123069 | 0.283436718 |
| 96,800 | 0.293595223 | 0.375145561 | 0.185401265 | 0.283358241 |
| 96,900 | 0.293479806 | 0.374960834 | 0.186174408 | 0.283794373 |
| 97,000 | 0.293392754 | 0.374619238 | 0.185910428 | 0.284202259 |
| 97,100 | 0.29339986 | 0.375152364 | 0.18563909 | 0.282880148 |
| 97,200 | 0.293274141 | 0.375251069 | 0.184685716 | 0.28264785 |
| 97,300 | 0.293208942 | 0.374435144 | 0.182945622 | 0.282664607 |
| 97,400 | 0.292958192 | 0.374540026 | 0.184428026 | 0.283099662 |
| 97,500 | 0.292907755 | 0.373867587 | 0.186590546 | 0.282387378 |
| 97,600 | 0.293039228 | 0.374163858 | 0.18598224 | 0.28249271 |
| 97,700 | 0.29278244 | 0.373661805 | 0.183530883 | 0.281545 |
| 97,800 | 0.292703988 | 0.373317347 | 0.183785564 | 0.281061633 |
| 97,900 | 0.292596044 | 0.373691069 | 0.183543795 | 0.281804754 |
| 98,000 | 0.292783511 | 0.373314202 | 0.184226123 | 0.281686721 |
| 98,100 | 0.292652369 | 0.373255517 | 0.183373966 | 0.281848877 |
| 98,200 | 0.292306752 | 0.373072283 | 0.183688163 | 0.282231574 |
| 98,300 | 0.292335246 | 0.372875793 | 0.184976268 | 0.282279132 |
| 98,400 | 0.291978847 | 0.371948113 | 0.183245837 | 0.28258425 |
| 98,500 | 0.291947687 | 0.373734232 | 0.182972219 | 0.281526935 |
| 98,600 | 0.291856517 | 0.372861461 | 0.181901805 | 0.281781476 |
| 98,700 | 0.291953049 | 0.372422565 | 0.182943916 | 0.282088089 |
| 98,800 | 0.291804642 | 0.372927812 | 0.18075578 | 0.282622202 |
| 98,900 | 0.291772052 | 0.372404001 | 0.182424208 | 0.281931621 |
| 99,000 | 0.291733849 | 0.372148774 | 0.18214576 | 0.282091943 |
| 99,100 | 0.291598794 | 0.371978441 | 0.183051665 | 0.282908218 |
| 99,200 | 0.291576364 | 0.371531711 | 0.180831185 | 0.282254888 |
| 99,300 | 0.291496697 | 0.371139721 | 0.181375447 | 0.282204383 |
| 99,400 | 0.291313722 | 0.370855093 | 0.180787245 | 0.282156368 |

| | | | | |
|---|---|---|---|---|
| 99,500 | 0.291153168 | 0.371231135 | 0.181561715 | 0.282288599 |
| 99,600 | 0.291115764 | 0.371732765 | 0.180768597 | 0.282304888 |
| 99,700 | 0.291010259 | 0.371976326 | 0.181782548 | 0.281768855 |
| 99,800 | 0.290904074 | 0.372519793 | 0.180212619 | 0.281491861 |
| 99,900 | 0.290896597 | 0.372286797 | 0.180203489 | 0.282930041 |
| 100,000 | 0.290826342 | 0.371900482 | 0.180730513 | 0.282135957 |
| 100,100 | 0.290687432 | 0.370990323 | 0.180984764 | 0.281670585 |
| 100,200 | 0.290776801 | 0.3708404 | 0.179383918 | 0.281316 |
| 100,300 | 0.290535836 | 0.370758274 | 0.178934256 | 0.281291054 |
| 100,400 | 0.290476717 | 0.371704332 | 0.180057982 | 0.280553491 |
| 100,500 | 0.290505226 | 0.371643732 | 0.178526195 | 0.280127393 |
| 100,600 | 0.290486701 | 0.372254211 | 0.179810693 | 0.281469889 |
| 100,700 | 0.290468838 | 0.371565399 | 0.179440753 | 0.281509996 |
| 100,800 | 0.290027839 | 0.371711931 | 0.180007117 | 0.280548847 |
| 100,900 | 0.2903343 | 0.371805368 | 0.180588636 | 0.281202557 |
| 101,000 | 0.29009041 | 0.37121637 | 0.180346388 | 0.281083659 |
| 101,100 | 0.289993941 | 0.370952743 | 0.180886866 | 0.281580841 |
| 101,200 | 0.289808618 | 0.371027757 | 0.181624151 | 0.281113625 |
| 101,300 | 0.28988146 | 0.3721342 | 0.18049454 | 0.28148959 |
| 101,400 | 0.289681011 | 0.372172283 | 0.179274042 | 0.280093943 |
| 101,500 | 0.289649621 | 0.371248982 | 0.179359884 | 0.28063863 |
| 101,600 | 0.289396351 | 0.370971143 | 0.179430904 | 0.280887171 |
| 101,700 | 0.28936765 | 0.371291987 | 0.179254638 | 0.279661796 |
| 101,800 | 0.289310324 | 0.371353425 | 0.179851722 | 0.279446303 |
| 101,900 | 0.289131369 | 0.371089974 | 0.180111487 | 0.280120616 |
| 102,000 | 0.288997377 | 0.370993207 | 0.178466685 | 0.280357483 |
| 102,100 | 0.289187948 | 0.371604543 | 0.176568943 | 0.2806798 |
| 102,200 | 0.288846565 | 0.37123691 | 0.178254448 | 0.280271618 |
| 102,300 | 0.288728119 | 0.371093077 | 0.176585487 | 0.279332213 |
| 102,400 | 0.288920111 | 0.371404173 | 0.17722135 | 0.278832045 |
| 102,500 | 0.288773148 | 0.370879606 | 0.17667011 | 0.279028835 |
| 102,600 | 0.288659429 | 0.370918376 | 0.17687979 | 0.279190051 |
| 102,700 | 0.288654389 | 0.37117289 | 0.177366828 | 0.278859781 |
| 102,800 | 0.288349414 | 0.371046212 | 0.17618517 | 0.279140172 |
| 102,900 | 0.288430508 | 0.370948618 | 0.175909471 | 0.278958442 |
| 103,000 | 0.288344447 | 0.370922219 | 0.176117038 | 0.278578967 |
| 103,100 | 0.288183621 | 0.369944656 | 0.178728311 | 0.27976517 |
| 103,200 | 0.287940692 | 0.370653627 | 0.177197943 | 0.278358887 |
| 103,300 | 0.287915278 | 0.370081791 | 0.17878683 | 0.278053613 |
| 103,400 | 0.288147716 | 0.371231223 | 0.178544205 | 0.279708226 |
| 103,500 | 0.287744338 | 0.369628019 | 0.177196893 | 0.277792318 |
| 103,600 | 0.287654024 | 0.369189764 | 0.178016004 | 0.278516396 |
| 103,700 | 0.287505087 | 0.36891279 | 0.177361298 | 0.277895591 |
| 103,800 | 0.287557986 | 0.370074975 | 0.176622378 | 0.278851352 |
| 103,900 | 0.287433495 | 0.369945567 | 0.176809185 | 0.279619541 |
| 104,000 | 0.28754249 | 0.369239165 | 0.178331939 | 0.279339629 |
| 104,100 | 0.287335582 | 0.370763981 | 0.177057444 | 0.278019109 |
| 104,200 | 0.287184266 | 0.370480372 | 0.177104023 | 0.278505093 |
| 104,300 | 0.287200823 | 0.371672499 | 0.177617602 | 0.27768621 |
| 104,400 | 0.287217716 | 0.370016985 | 0.176190339 | 0.277274361 |
| 104,500 | 0.287231626 | 0.370949863 | 0.17766937 | 0.277685813 |
| 104,600 | 0.28699491 | 0.370902905 | 0.176403553 | 0.277189856 |
| 104,700 | 0.286915162 | 0.371519836 | 0.175961419 | 0.277404524 |
| 104,800 | 0.286905409 | 0.370979301 | 0.176539784 | 0.277260155 |
| 104,900 | 0.286928598 | 0.370646715 | 0.173662826 | 0.276553846 |
| 105,000 | 0.286867601 | 0.370082796 | 0.175765985 | 0.278542882 |
| 105,100 | 0.286675061 | 0.369952015 | 0.176237557 | 0.278033278 |
| 105,200 | 0.28646075 | 0.369496513 | 0.176445097 | 0.27756269 |
| 105,300 | 0.286623359 | 0.370045308 | 0.176261955 | 0.277254259 |
| 105,400 | 0.286548089 | 0.368803738 | 0.174988972 | 0.276502213 |
| 105,500 | 0.286460001 | 0.369646466 | 0.175808888 | 0.276534477 |
| 105,600 | 0.286252917 | 0.369275997 | 0.176383042 | 0.276612712 |
| 105,700 | 0.286281862 | 0.369372888 | 0.17478364 | 0.276857054 |
| 105,800 | 0.286157302 | 0.370337183 | 0.17634491 | 0.276791605 |
| 105,900 | 0.286230852 | 0.369803358 | 0.176921222 | 0.277629078 |
| 106,000 | 0.286183046 | 0.369395939 | 0.17461176 | 0.27740491 |
| 106,100 | 0.286169979 | 0.369666645 | 0.17224169 | 0.277301126 |

| | | | | |
|---|---|---|---|---|
| 106,200 | 0.286290172 | 0.370318548 | 0.174127426 | 0.277191203 |
| 106,300 | 0.286144438 | 0.370291044 | 0.174815267 | 0.276353244 |
| 106,400 | 0.286090315 | 0.36957056 | 0.175202544 | 0.276401433 |
| 106,500 | 0.285910445 | 0.369788614 | 0.174616479 | 0.276831047 |
| 106,600 | 0.285944405 | 0.368777125 | 0.173511278 | 0.277430562 |
| 106,700 | 0.285881859 | 0.368383288 | 0.172430177 | 0.277507305 |
| 106,800 | 0.285730232 | 0.36767835 | 0.173483649 | 0.277885726 |
| 106,900 | 0.285742499 | 0.367893891 | 0.171795448 | 0.277380206 |
| 107,000 | 0.285430672 | 0.366916048 | 0.172421731 | 0.276962506 |
| 107,100 | 0.285520702 | 0.366966475 | 0.170765761 | 0.277084878 |
| 107,200 | 0.285508601 | 0.366423543 | 0.170041503 | 0.276369255 |
| 107,300 | 0.285407347 | 0.366800237 | 0.171353071 | 0.276799696 |
| 107,400 | 0.285037727 | 0.365871162 | 0.170774424 | 0.276295334 |
| 107,500 | 0.284824342 | 0.366573842 | 0.170105913 | 0.276720378 |
| 107,600 | 0.2849574 | 0.366205422 | 0.170145472 | 0.275787343 |
| 107,700 | 0.284806869 | 0.365744195 | 0.170661556 | 0.275355391 |
| 107,800 | 0.284551166 | 0.365775313 | 0.171677177 | 0.275731993 |
| 107,900 | 0.284511148 | 0.366705961 | 0.17115006 | 0.275242029 |
| 108,000 | 0.284501988 | 0.365305574 | 0.171836154 | 0.276293986 |
| 108,100 | 0.28444349 | 0.366226294 | 0.172574752 | 0.275220494 |
| 108,200 | 0.284383708 | 0.365313877 | 0.173030219 | 0.275424865 |
| 108,300 | 0.284150998 | 0.365112542 | 0.170467409 | 0.274404849 |
| 108,400 | 0.284375549 | 0.364344195 | 0.171682754 | 0.275715908 |
| 108,500 | 0.284312985 | 0.365844411 | 0.171004148 | 0.275664449 |
| 108,600 | 0.284082695 | 0.36593762 | 0.171846579 | 0.275390554 |
| 108,700 | 0.283967852 | 0.365744341 | 0.170327544 | 0.275600602 |
| 108,800 | 0.284014515 | 0.365635386 | 0.168605333 | 0.275000436 |
| 108,900 | 0.283711342 | 0.36578023 | 0.169894246 | 0.275200136 |
| 109,000 | 0.283472268 | 0.365163187 | 0.169111097 | 0.275013601 |
| 109,100 | 0.283394344 | 0.36551344 | 0.171710666 | 0.275436197 |
| 109,200 | 0.283248728 | 0.365631333 | 0.171364035 | 0.275647005 |
| 109,300 | 0.283297631 | 0.365591547 | 0.171992106 | 0.275089192 |
| 109,400 | 0.283141598 | 0.36456929 | 0.172090185 | 0.274984266 |
| 109,500 | 0.28306346 | 0.364384772 | 0.170889937 | 0.273833383 |
| 109,600 | 0.282839428 | 0.364399554 | 0.171420011 | 0.274029056 |
| 109,700 | 0.283044039 | 0.365158305 | 0.170589365 | 0.27447105 |
| 109,800 | 0.282802444 | 0.364888369 | 0.170325694 | 0.274560504 |
| 109,900 | 0.283023361 | 0.365324854 | 0.171359692 | 0.274443268 |
| 110,000 | 0.282960308 | 0.365355277 | 0.170525168 | 0.274844757 |
| 110,100 | 0.282750409 | 0.364180925 | 0.171459687 | 0.275236495 |
| 110,200 | 0.282666623 | 0.363361451 | 0.169545225 | 0.274988031 |
| 110,300 | 0.28282764 | 0.36433248 | 0.173517829 | 0.274892836 |
| 110,400 | 0.282587203 | 0.364337525 | 0.172029659 | 0.273844111 |
| 110,500 | 0.282569824 | 0.364581109 | 0.17144448 | 0.273376134 |
| 110,600 | 0.282475076 | 0.363575193 | 0.172289229 | 0.273834315 |
| 110,700 | 0.282324764 | 0.363076782 | 0.171448558 | 0.273584057 |
| 110,800 | 0.282238884 | 0.362947728 | 0.173937989 | 0.273150994 |
| 110,900 | 0.2821372 | 0.361882008 | 0.172250805 | 0.272367968 |
| 111,000 | 0.282039295 | 0.362616157 | 0.17218436 | 0.272231928 |
| 111,100 | 0.282123835 | 0.363242371 | 0.171222306 | 0.27285399 |
| 111,200 | 0.281968424 | 0.362030125 | 0.172335886 | 0.273139165 |
| 111,300 | 0.281619047 | 0.36109243 | 0.172736136 | 0.272260184 |
| 111,400 | 0.281675058 | 0.361724144 | 0.171969854 | 0.272335789 |
| 111,500 | 0.2815105 | 0.361181442 | 0.170180737 | 0.271789596 |
| 111,600 | 0.281581503 | 0.362026177 | 0.171327258 | 0.272897267 |
| 111,700 | 0.281343004 | 0.361560265 | 0.169689356 | 0.27227168 |
| 111,800 | 0.281356451 | 0.362167719 | 0.169558928 | 0.273271873 |
| 111,900 | 0.281195662 | 0.360731738 | 0.171192274 | 0.271636842 |
| 112,000 | 0.281174045 | 0.361414593 | 0.17006761 | 0.272234085 |
| 112,100 | 0.281115221 | 0.361585592 | 0.170214105 | 0.272106665 |
| 112,200 | 0.280907825 | 0.361827003 | 0.171634735 | 0.272385766 |
| 112,300 | 0.280937687 | 0.361169516 | 0.170266907 | 0.272138238 |
| 112,400 | 0.280892396 | 0.361885937 | 0.169526964 | 0.271886533 |
| 112,500 | 0.280829866 | 0.36145447 | 0.171548708 | 0.272402474 |
| 112,600 | 0.280805401 | 0.361865176 | 0.171478713 | 0.271739661 |
| 112,700 | 0.28046376 | 0.361751118 | 0.170621976 | 0.271222836 |
| 112,800 | 0.280354875 | 0.361889399 | 0.168946619 | 0.271280752 |

| | | | | |
|---|---|---|---|---|
| 112,900 | 0.280367461 | 0.36171026 | 0.168054735 | 0.271274801 |
| 113,000 | 0.280254592 | 0.362057869 | 0.169027499 | 0.271409091 |
| 113,100 | 0.280255638 | 0.361775828 | 0.167732384 | 0.271847798 |
| 113,200 | 0.280176583 | 0.361367537 | 0.168922531 | 0.271858766 |
| 113,300 | 0.280377936 | 0.362031784 | 0.166973272 | 0.272181616 |
| 113,400 | 0.280098925 | 0.361525817 | 0.166039927 | 0.27136499 |
| 113,500 | 0.280123485 | 0.361123978 | 0.167156382 | 0.270764188 |
| 113,600 | 0.279970837 | 0.360886648 | 0.167725312 | 0.270634262 |
| 113,700 | 0.279919857 | 0.361482594 | 0.168428003 | 0.271031515 |
| 113,800 | 0.279912062 | 0.360490126 | 0.168491672 | 0.27182881 |
| 113,900 | 0.279776985 | 0.360842015 | 0.169657647 | 0.270958546 |
| 114,000 | 0.279673005 | 0.360249203 | 0.168492686 | 0.270889208 |
| 114,100 | 0.279624091 | 0.360359458 | 0.168255866 | 0.269921297 |
| 114,200 | 0.279495014 | 0.360947319 | 0.166792832 | 0.27130432 |
| 114,300 | 0.279353896 | 0.360188463 | 0.167681278 | 0.271369019 |
| 114,400 | 0.279172453 | 0.359180067 | 0.167672838 | 0.270519486 |
| 114,500 | 0.279125061 | 0.359284408 | 0.168510149 | 0.270675394 |
| 114,600 | 0.27912211 | 0.359458534 | 0.166354078 | 0.270402311 |
| 114,700 | 0.279115681 | 0.359613975 | 0.167434325 | 0.271134041 |
| 114,800 | 0.279187369 | 0.359432979 | 0.169860161 | 0.270654593 |
| 114,900 | 0.278967019 | 0.359042839 | 0.166290029 | 0.270705405 |
| 115,000 | 0.278814007 | 0.358123607 | 0.166786623 | 0.270035026 |
| 115,100 | 0.27895039 | 0.35927098 | 0.167223961 | 0.270752578 |
| 115,200 | 0.27891087 | 0.359462475 | 0.166311592 | 0.27144962 |
| 115,300 | 0.278830055 | 0.358568995 | 0.167026812 | 0.271039013 |
| 115,400 | 0.278820606 | 0.357749202 | 0.167352997 | 0.271274095 |
| 115,500 | 0.278670756 | 0.359107697 | 0.16712899 | 0.269429845 |
| 115,600 | 0.27867126 | 0.359781999 | 0.167208028 | 0.270696758 |
| 115,700 | 0.278619122 | 0.358448079 | 0.168215483 | 0.270535614 |
| 115,800 | 0.278458379 | 0.358555869 | 0.1672209 | 0.271062877 |
| 115,900 | 0.278305112 | 0.359140206 | 0.166444493 | 0.270113419 |
| 116,000 | 0.278216505 | 0.357914236 | 0.167196933 | 0.270894138 |
| 116,100 | 0.278250545 | 0.358747742 | 0.168839332 | 0.271112727 |
| 116,200 | 0.277939181 | 0.358203376 | 0.167903232 | 0.269240517 |
| 116,300 | 0.277874772 | 0.358150363 | 0.167487518 | 0.270475383 |
| 116,400 | 0.277890097 | 0.357483347 | 0.168271924 | 0.270629673 |
| 116,500 | 0.277923909 | 0.358452452 | 0.169543386 | 0.270530895 |
| 116,600 | 0.277835071 | 0.358223582 | 0.167720988 | 0.269514123 |
| 116,700 | 0.277748939 | 0.358249547 | 0.168039583 | 0.269925614 |
| 116,800 | 0.277462464 | 0.35765986 | 0.164913937 | 0.269295057 |
| 116,900 | 0.277792754 | 0.357520019 | 0.167112102 | 0.268187862 |
| 117,000 | 0.277524731 | 0.356945956 | 0.167690694 | 0.269213743 |
| 117,100 | 0.277659698 | 0.357230841 | 0.168902783 | 0.269223451 |
| 117,200 | 0.277479037 | 0.356160876 | 0.167613601 | 0.268654323 |
| 117,300 | 0.277389602 | 0.357371239 | 0.167412246 | 0.267810643 |
| 117,400 | 0.277071077 | 0.356663148 | 0.1670064 | 0.26779177 |
| 117,500 | 0.277154161 | 0.355937268 | 0.167159979 | 0.268181781 |
| 117,600 | 0.277112922 | 0.356770339 | 0.16738681 | 0.267559257 |
| 117,700 | 0.277136129 | 0.357296566 | 0.167787473 | 0.268240358 |
| 117,800 | 0.277138793 | 0.357856587 | 0.171453129 | 0.26798831 |
| 117,900 | 0.277146664 | 0.357375048 | 0.170918282 | 0.26851809 |
| 118,000 | 0.276887202 | 0.356989515 | 0.169237943 | 0.268325804 |
| 118,100 | 0.2766899 | 0.357064166 | 0.169700494 | 0.267404093 |
| 118,200 | 0.276544128 | 0.356741179 | 0.169243937 | 0.267386284 |
| 118,300 | 0.276647068 | 0.357360745 | 0.16971935 | 0.268476676 |
| 118,400 | 0.276665634 | 0.356436877 | 0.169012829 | 0.267760922 |
| 118,500 | 0.276494939 | 0.356795636 | 0.168776213 | 0.266737487 |
| 118,600 | 0.276395676 | 0.356168674 | 0.169252973 | 0.265860531 |
| 118,700 | 0.276537316 | 0.356950441 | 0.169271377 | 0.266335024 |
| 118,800 | 0.276285136 | 0.356556478 | 0.16986302 | 0.26682471 |
| 118,900 | 0.276071999 | 0.355523989 | 0.168745276 | 0.266472654 |
| 119,000 | 0.276200368 | 0.355353845 | 0.170008211 | 0.265957902 |
| 119,100 | 0.276025717 | 0.356146728 | 0.170034147 | 0.266291935 |
| 119,200 | 0.275820854 | 0.355195731 | 0.170996688 | 0.265598169 |
| 119,300 | 0.275995135 | 0.355099413 | 0.171109934 | 0.266206769 |
| 119,400 | 0.275728965 | 0.355084066 | 0.168770537 | 0.265256522 |
| 119,500 | 0.275606832 | 0.354510181 | 0.169782351 | 0.265525535 |

| | | | | |
|---|---|---|---|---|
| 119,600 | 0.275505016 | 0.353991387 | 0.16956738 | 0.265652946 |
| 119,700 | 0.275464918 | 0.353554427 | 0.169631866 | 0.264952375 |
| 119,800 | 0.275429898 | 0.352904034 | 0.168010444 | 0.26624714 |
| 119,900 | 0.275395679 | 0.354452468 | 0.169539639 | 0.266086366 |
| 120,000 | 0.275236436 | 0.353211409 | 0.16999195 | 0.266780937 |
| 120,100 | 0.275301706 | 0.354754367 | 0.167500427 | 0.266477736 |
| 120,200 | 0.275084972 | 0.354627147 | 0.167939725 | 0.266656725 |
| 120,300 | 0.27498917 | 0.353793807 | 0.167196296 | 0.265556471 |
| 120,400 | 0.274805877 | 0.354199361 | 0.167418569 | 0.264871468 |
| 120,500 | 0.274830592 | 0.353326618 | 0.16715804 | 0.265754374 |
| 120,600 | 0.274875249 | 0.35353367 | 0.16701127 | 0.265763168 |
| 120,700 | 0.274704623 | 0.353109281 | 0.166854206 | 0.265616115 |
| 120,800 | 0.274723364 | 0.352966369 | 0.167523711 | 0.265539482 |
| 120,900 | 0.274534178 | 0.353583667 | 0.166461086 | 0.264793718 |
| 121,000 | 0.274496086 | 0.352851751 | 0.168290701 | 0.264840576 |
| 121,100 | 0.27476349 | 0.353166064 | 0.168567757 | 0.264342516 |
| 121,200 | 0.274373249 | 0.353101977 | 0.167785618 | 0.264891923 |
| 121,300 | 0.274322038 | 0.353239036 | 0.168182228 | 0.264986897 |
| 121,400 | 0.274489988 | 0.352983576 | 0.168393733 | 0.265755862 |
| 121,500 | 0.274374677 | 0.353875701 | 0.168498032 | 0.265224681 |
| 121,600 | 0.274097777 | 0.35377978 | 0.168164465 | 0.26511954 |
| 121,700 | 0.274032807 | 0.35322701 | 0.16741596 | 0.265891244 |
| 121,800 | 0.273918956 | 0.351860748 | 0.167297137 | 0.265798084 |
| 121,900 | 0.27376365 | 0.353325156 | 0.167587851 | 0.264917011 |
| 122,000 | 0.273926635 | 0.352388695 | 0.166507169 | 0.265120782 |
| 122,100 | 0.273824134 | 0.352263187 | 0.167404617 | 0.264857812 |
| 122,200 | 0.273648246 | 0.352201923 | 0.166798469 | 0.265599761 |
| 122,300 | 0.27374003 | 0.353494075 | 0.165704918 | 0.265255666 |
| 122,400 | 0.273727429 | 0.352514404 | 0.165333205 | 0.265537916 |
| 122,500 | 0.27364182 | 0.352007108 | 0.165693058 | 0.265702382 |
| 122,600 | 0.273499877 | 0.351646864 | 0.166553097 | 0.265111489 |
| 122,700 | 0.273576312 | 0.3525027 | 0.166845608 | 0.26450555 |
| 122,800 | 0.27342617 | 0.351756002 | 0.167475249 | 0.264437319 |
| 122,900 | 0.273554059 | 0.35194679 | 0.16638281 | 0.264215858 |
| 123,000 | 0.273436122 | 0.351710085 | 0.167193716 | 0.263701256 |
| 123,100 | 0.273354829 | 0.351509063 | 0.1651842 | 0.263431663 |
| 123,200 | 0.273297928 | 0.351335347 | 0.164986147 | 0.2652584 |
| 123,300 | 0.27301074 | 0.351915477 | 0.165338269 | 0.264816289 |
| 123,400 | 0.272953289 | 0.351674563 | 0.165348497 | 0.264322449 |
| 123,500 | 0.272679802 | 0.352153902 | 0.165009291 | 0.264222297 |
| 123,600 | 0.272830265 | 0.352557209 | 0.165147617 | 0.264733474 |
| 123,700 | 0.272691613 | 0.353168163 | 0.16448334 | 0.265126752 |
| 123,800 | 0.272749159 | 0.353070749 | 0.165214282 | 0.26452021 |
| 123,900 | 0.272645855 | 0.353635265 | 0.162632356 | 0.263815096 |
| 124,000 | 0.272392392 | 0.352935131 | 0.16202238 | 0.264337156 |
| 124,100 | 0.272326255 | 0.35308748 | 0.162151394 | 0.264608022 |
| 124,200 | 0.272329455 | 0.352492342 | 0.163411217 | 0.264304593 |
| 124,300 | 0.272344757 | 0.353003405 | 0.163954517 | 0.264689189 |
| 124,400 | 0.272389629 | 0.352148464 | 0.164960327 | 0.264673125 |
| 124,500 | 0.27236575 | 0.351206309 | 0.164998941 | 0.263962797 |
| 124,600 | 0.272211956 | 0.35177508 | 0.165814931 | 0.264182389 |
| 124,700 | 0.272083474 | 0.351559443 | 0.165699361 | 0.26391078 |
| 124,800 | 0.272066613 | 0.351818788 | 0.164367099 | 0.263941103 |
| 124,900 | 0.271932737 | 0.351891181 | 0.16443952 | 0.263714722 |
| 125,000 | 0.271998758 | 0.351810729 | 0.164903108 | 0.263581739 |
| 125,100 | 0.271915385 | 0.352083702 | 0.164307848 | 0.263992086 |
| 125,200 | 0.271666994 | 0.351936264 | 0.163595777 | 0.263811371 |
| 125,300 | 0.271715918 | 0.351385858 | 0.165348564 | 0.264162131 |
| 125,400 | 0.271432593 | 0.350918926 | 0.164363685 | 0.264815415 |
| 125,500 | 0.271590184 | 0.351844965 | 0.164037279 | 0.264317776 |
| 125,600 | 0.271262286 | 0.351700348 | 0.164813268 | 0.264161492 |
| 125,700 | 0.271394491 | 0.350431487 | 0.166408385 | 0.265049138 |
| 125,800 | 0.271343864 | 0.350393683 | 0.165604528 | 0.264682633 |
| 125,900 | 0.27117329 | 0.349919339 | 0.164589398 | 0.264488141 |
| 126,000 | 0.271113358 | 0.351120198 | 0.164779325 | 0.264727543 |
| 126,100 | 0.271034883 | 0.350911116 | 0.164684269 | 0.264111096 |
| 126,200 | 0.2711211 | 0.351172922 | 0.164702515 | 0.264996272 |

| | | | | |
|---|---|---|---|---|
| 126,300 | 0.270773811 | 0.350581926 | 0.164663384 | 0.264655313 |
| 126,400 | 0.270827681 | 0.350988646 | 0.163019321 | 0.263396563 |
| 126,500 | 0.270625759 | 0.351388208 | 0.162436345 | 0.263800902 |
| 126,600 | 0.270701969 | 0.35085576 | 0.163361649 | 0.263179823 |
| 126,700 | 0.270765264 | 0.351443426 | 0.164201058 | 0.262919487 |
| 126,800 | 0.270626743 | 0.350927013 | 0.163861093 | 0.26351779 |
| 126,900 | 0.270742424 | 0.351431845 | 0.164445937 | 0.263660047 |
| 127,000 | 0.270668306 | 0.349963748 | 0.166149934 | 0.264043476 |
| 127,100 | 0.270573397 | 0.350747084 | 0.163869352 | 0.263814566 |
| 127,200 | 0.270581573 | 0.350622014 | 0.165436272 | 0.263111661 |
| 127,300 | 0.270524984 | 0.350796732 | 0.166321857 | 0.263370824 |
| 127,400 | 0.270302684 | 0.350959249 | 0.164762221 | 0.263179849 |
| 127,500 | 0.270228576 | 0.350535406 | 0.164329154 | 0.261953144 |
| 127,600 | 0.270211578 | 0.350791032 | 0.164934414 | 0.26243138 |
| 127,700 | 0.270202704 | 0.351231934 | 0.16564267 | 0.263284856 |
| 127,800 | 0.270324832 | 0.350180165 | 0.164017719 | 0.262180644 |
| 127,900 | 0.270352148 | 0.350288539 | 0.16269724 | 0.262429955 |
| 128,000 | 0.270173824 | 0.349521631 | 0.160678542 | 0.261968795 |
| 128,100 | 0.269905284 | 0.350107696 | 0.160219754 | 0.262646089 |
| 128,200 | 0.269947297 | 0.35027269 | 0.159485413 | 0.26240519 |
| 128,300 | 0.269769452 | 0.350215591 | 0.161917969 | 0.262061471 |
| 128,400 | 0.269952034 | 0.350163415 | 0.161571603 | 0.262524713 |
| 128,500 | 0.269724535 | 0.349650975 | 0.162842606 | 0.262722381 |
| 128,600 | 0.269480107 | 0.349903139 | 0.162779303 | 0.261143852 |
| 128,700 | 0.269498454 | 0.3498198 | 0.161882263 | 0.261652048 |
| 128,800 | 0.269300775 | 0.349556151 | 0.161729764 | 0.261475493 |
| 128,900 | 0.269410747 | 0.349624533 | 0.161200221 | 0.261267701 |
| 129,000 | 0.269307014 | 0.348871997 | 0.161139043 | 0.262311831 |
| 129,100 | 0.269331842 | 0.349968357 | 0.160334429 | 0.260602703 |
| 129,200 | 0.269191697 | 0.349698019 | 0.159815105 | 0.260933052 |
| 129,300 | 0.269038118 | 0.348586496 | 0.161774296 | 0.261285411 |
| 129,400 | 0.269138812 | 0.349292045 | 0.16067173 | 0.261306339 |
| 129,500 | 0.269151742 | 0.350165175 | 0.159854369 | 0.260883254 |
| 129,600 | 0.268977113 | 0.350211936 | 0.160882048 | 0.260978505 |
| 129,700 | 0.268938879 | 0.350651139 | 0.159028181 | 0.260663041 |
| 129,800 | 0.269043293 | 0.350173964 | 0.158583522 | 0.26115119 |
| 129,900 | 0.268838608 | 0.349231761 | 0.158780802 | 0.261543788 |
| 130,000 | 0.268927402 | 0.350106259 | 0.159733004 | 0.261644609 |
| 130,100 | 0.268652583 | 0.349343123 | 0.159911262 | 0.261803937 |
| 130,200 | 0.268901194 | 0.348397916 | 0.158776356 | 0.261529969 |
| 130,300 | 0.268467021 | 0.349501215 | 0.159179782 | 0.260992973 |
| 130,400 | 0.268521058 | 0.348749356 | 0.159324924 | 0.26131677 |
| 130,500 | 0.268350617 | 0.348855082 | 0.157943462 | 0.261354523 |
| 130,600 | 0.268377586 | 0.349526881 | 0.159061287 | 0.261890485 |
| 130,700 | 0.268338964 | 0.349522057 | 0.158760331 | 0.261650359 |
| 130,800 | 0.268333023 | 0.348197875 | 0.159510105 | 0.261971472 |
| 130,900 | 0.268182512 | 0.349226807 | 0.15857487 | 0.260724531 |
| 131,000 | 0.268192646 | 0.348917222 | 0.158728422 | 0.260966905 |
| 131,100 | 0.268102949 | 0.349916006 | 0.158452068 | 0.260945254 |
| 131,200 | 0.267838945 | 0.349621305 | 0.160357728 | 0.259427426 |
| 131,300 | 0.267810421 | 0.348657591 | 0.158941044 | 0.259726224 |
| 131,400 | 0.267699874 | 0.348084678 | 0.157479495 | 0.259540465 |
| 131,500 | 0.26767191 | 0.347984162 | 0.158196315 | 0.259723671 |
| 131,600 | 0.267329124 | 0.348237473 | 0.157539981 | 0.259266141 |
| 131,700 | 0.267327788 | 0.347275849 | 0.158684825 | 0.260224172 |
| 131,800 | 0.267341597 | 0.347102765 | 0.158440671 | 0.259403084 |
| 131,900 | 0.267347997 | 0.347537798 | 0.159087677 | 0.259384927 |
| 132,000 | 0.267234249 | 0.347423477 | 0.15785916 | 0.258932122 |
| 132,100 | 0.267115302 | 0.347631219 | 0.157070743 | 0.259418454 |
| 132,200 | 0.26712418 | 0.347937504 | 0.157528906 | 0.259848639 |
| 132,300 | 0.267104531 | 0.347292193 | 0.156668846 | 0.259556193 |
| 132,400 | 0.267011277 | 0.347577249 | 0.156968171 | 0.259066841 |
| 132,500 | 0.267001395 | 0.347858473 | 0.157680956 | 0.259606045 |
| 132,600 | 0.267064164 | 0.348470704 | 0.156898622 | 0.259972114 |
| 132,700 | 0.266772913 | 0.347831998 | 0.15742226 | 0.25940588 |
| 132,800 | 0.266773203 | 0.348701316 | 0.15712944 | 0.258857658 |
| 132,900 | 0.26679103 | 0.347604788 | 0.158236054 | 0.25848534 |

| | | | | |
|---|---|---|---|---|
| 133,000 | 0.266493667 | 0.348185956 | 0.157709529 | 0.257508198 |
| 133,100 | 0.266417601 | 0.348222155 | 0.157883417 | 0.257269274 |
| 133,200 | 0.266420228 | 0.348478916 | 0.156786039 | 0.258047322 |
| 133,300 | 0.266517824 | 0.347161135 | 0.157196846 | 0.259672207 |
| 133,400 | 0.26635671 | 0.347509738 | 0.157768394 | 0.259146931 |
| 133,500 | 0.266134191 | 0.34781029 | 0.158437479 | 0.258620581 |
| 133,600 | 0.266163797 | 0.347322782 | 0.15677355 | 0.258513059 |
| 133,700 | 0.266301307 | 0.348677641 | 0.157626374 | 0.258420584 |
| 133,800 | 0.266147011 | 0.347779067 | 0.156889351 | 0.25803214 |
| 133,900 | 0.266210945 | 0.348086231 | 0.156424571 | 0.258707793 |
| 134,000 | 0.266052893 | 0.347309864 | 0.156818152 | 0.258119337 |
| 134,100 | 0.26610141 | 0.347141702 | 0.157278985 | 0.258534724 |
| 134,200 | 0.265940811 | 0.346885549 | 0.156293221 | 0.257950581 |
| 134,300 | 0.266011572 | 0.346714638 | 0.156195 | 0.259063812 |
| 134,400 | 0.265887119 | 0.346516678 | 0.154914999 | 0.258532938 |
| 134,500 | 0.265980765 | 0.347548286 | 0.155349671 | 0.258992744 |
| 134,600 | 0.265780581 | 0.347100305 | 0.15632391 | 0.25780114 |
| 134,700 | 0.265739316 | 0.346415326 | 0.155747111 | 0.258613024 |
| 134,800 | 0.265569604 | 0.347422287 | 0.154388155 | 0.257761763 |
| 134,900 | 0.265403035 | 0.347239865 | 0.154769214 | 0.257772045 |
| 135,000 | 0.265613755 | 0.347278846 | 0.154729836 | 0.258597957 |
| 135,100 | 0.265346592 | 0.346789453 | 0.155844996 | 0.257473506 |
| 135,200 | 0.265361028 | 0.347428179 | 0.155693911 | 0.257070199 |
| 135,300 | 0.265305004 | 0.347577041 | 0.154111166 | 0.257487462 |
| 135,400 | 0.265134401 | 0.346429681 | 0.154346914 | 0.257693575 |
| 135,500 | 0.26517745 | 0.34667557 | 0.154828846 | 0.258180631 |
| 135,600 | 0.265014016 | 0.346587348 | 0.15508967 | 0.257526707 |
| 135,700 | 0.265077952 | 0.345702045 | 0.155789898 | 0.258037913 |
| 135,800 | 0.264879267 | 0.346051277 | 0.155581527 | 0.258560136 |
| 135,900 | 0.264826638 | 0.346343251 | 0.155018212 | 0.25836836 |
| 136,000 | 0.264918937 | 0.347203879 | 0.155704107 | 0.258050897 |
| 136,100 | 0.264921495 | 0.346760357 | 0.156618879 | 0.257410557 |
| 136,200 | 0.264716331 | 0.346918656 | 0.156860081 | 0.257725743 |
| 136,300 | 0.264829622 | 0.346682625 | 0.156090807 | 0.257525475 |
| 136,400 | 0.264752829 | 0.346502726 | 0.156746867 | 0.257703318 |
| 136,500 | 0.264454179 | 0.345781241 | 0.155414341 | 0.257337644 |
| 136,600 | 0.264571893 | 0.345900257 | 0.15596842 | 0.257790761 |
| 136,700 | 0.264614632 | 0.346550478 | 0.155205141 | 0.258091432 |
| 136,800 | 0.264399816 | 0.345275107 | 0.155314739 | 0.257179014 |
| 136,900 | 0.264352393 | 0.345785436 | 0.153074043 | 0.257737143 |
| 137,000 | 0.264386252 | 0.345656653 | 0.154255682 | 0.257192598 |
| 137,100 | 0.264455424 | 0.345969837 | 0.154346189 | 0.257114558 |
| 137,200 | 0.264405224 | 0.346462138 | 0.155229174 | 0.257580918 |
| 137,300 | 0.264117859 | 0.346294352 | 0.153919169 | 0.257439771 |
| 137,400 | 0.26402074 | 0.345812241 | 0.153310176 | 0.257598283 |
| 137,500 | 0.26400968 | 0.344744803 | 0.154214305 | 0.257674382 |
| 137,600 | 0.263922686 | 0.344664382 | 0.15597762 | 0.257452334 |
| 137,700 | 0.2640008 | 0.344985885 | 0.156083779 | 0.257590377 |
| 137,800 | 0.263791909 | 0.344710036 | 0.155100297 | 0.257734342 |
| 137,900 | 0.263809686 | 0.345346971 | 0.154621804 | 0.257332854 |
| 138,000 | 0.263796726 | 0.346146367 | 0.153610555 | 0.257767025 |
| 138,100 | 0.263791811 | 0.344313226 | 0.153174933 | 0.256355037 |
| 138,200 | 0.263493347 | 0.344031935 | 0.153949297 | 0.2566959 |
| 138,300 | 0.263455744 | 0.344215567 | 0.154251702 | 0.256610594 |
| 138,400 | 0.263603989 | 0.343834468 | 0.153231554 | 0.256567762 |
| 138,500 | 0.263184315 | 0.344478234 | 0.151671674 | 0.256869272 |
| 138,600 | 0.263233858 | 0.343623994 | 0.152938497 | 0.256657373 |
| 138,700 | 0.263243557 | 0.343664913 | 0.152583669 | 0.257291421 |
| 138,800 | 0.262971147 | 0.343467859 | 0.151453884 | 0.255884117 |
| 138,900 | 0.263131627 | 0.344453176 | 0.153008891 | 0.255767576 |
| 139,000 | 0.262896383 | 0.344654561 | 0.15194566 | 0.255439572 |
| 139,100 | 0.263025209 | 0.344695158 | 0.151918121 | 0.255642514 |
| 139,200 | 0.262894474 | 0.343824206 | 0.152615598 | 0.255365202 |
| 139,300 | 0.262741109 | 0.342314035 | 0.153217815 | 0.255216961 |
| 139,400 | 0.262723617 | 0.34232216 | 0.153326284 | 0.254898931 |
| 139,500 | 0.262561006 | 0.342290725 | 0.152906246 | 0.255027967 |
| 139,600 | 0.262722365 | 0.342755471 | 0.151297607 | 0.254805496 |

| | | | | |
|---|---|---|---|---|
| 139,700 | 0.26259636 | 0.342871252 | 0.152158268 | 0.254830026 |
| 139,800 | 0.262329829 | 0.34226411 | 0.152207909 | 0.254547837 |
| 139,900 | 0.262486262 | 0.342973381 | 0.151125239 | 0.254259519 |
| 140,000 | 0.262254376 | 0.342466697 | 0.152505652 | 0.254841489 |
| 140,100 | 0.262178841 | 0.342755847 | 0.153206344 | 0.254216813 |
| 140,200 | 0.262022291 | 0.341757241 | 0.153472243 | 0.255419644 |
| 140,300 | 0.262092433 | 0.34194775 | 0.153514008 | 0.255902625 |
| 140,400 | 0.262110056 | 0.342061869 | 0.152329369 | 0.255121749 |
| 140,500 | 0.261859712 | 0.342123932 | 0.152554506 | 0.254356199 |
| 140,600 | 0.26189235 | 0.341585562 | 0.152926697 | 0.254469037 |
| 140,700 | 0.2615129 | 0.34238168 | 0.15194334 | 0.253326934 |
| 140,800 | 0.261587738 | 0.342195456 | 0.153650079 | 0.253311471 |
| 140,900 | 0.261806345 | 0.34143157 | 0.15321508 | 0.253591918 |
| 141,000 | 0.26180705 | 0.342405996 | 0.155618277 | 0.252415091 |
| 141,100 | 0.261674815 | 0.341789634 | 0.154003478 | 0.253089827 |
| 141,200 | 0.261604765 | 0.341455183 | 0.15355732 | 0.253131919 |
| 141,300 | 0.261593164 | 0.341081127 | 0.151485155 | 0.253913409 |
| 141,400 | 0.261373186 | 0.341500189 | 0.151725726 | 0.252834552 |
| 141,500 | 0.261530937 | 0.342096432 | 0.153266834 | 0.25374172 |
| 141,600 | 0.261492453 | 0.340551642 | 0.151924235 | 0.252084661 |
| 141,700 | 0.26126535 | 0.341001 | 0.150637445 | 0.253896056 |
| 141,800 | 0.261392612 | 0.341263097 | 0.153198061 | 0.25359402 |
| 141,900 | 0.261252544 | 0.341167019 | 0.151141086 | 0.253552158 |
| 142,000 | 0.261081069 | 0.341207244 | 0.152465826 | 0.253656847 |
| 142,100 | 0.261284547 | 0.341408522 | 0.152861009 | 0.253699882 |
| 142,200 | 0.261047572 | 0.34211213 | 0.15364192 | 0.253410015 |
| 142,300 | 0.261095009 | 0.341207747 | 0.152682696 | 0.253453867 |
| 142,400 | 0.260926838 | 0.341056122 | 0.153595964 | 0.253808273 |
| 142,500 | 0.260973858 | 0.341558408 | 0.153300565 | 0.253278091 |
| 142,600 | 0.260791624 | 0.341191531 | 0.152915484 | 0.253347421 |
| 142,700 | 0.260946371 | 0.34195226 | 0.153794235 | 0.252271153 |
| 142,800 | 0.260764582 | 0.341478868 | 0.154851142 | 0.253249354 |
| 142,900 | 0.260512919 | 0.341315003 | 0.152608075 | 0.252342814 |
| 143,000 | 0.260462739 | 0.342841043 | 0.151480248 | 0.253431227 |
| 143,100 | 0.260577889 | 0.342340034 | 0.152164855 | 0.252813656 |
| 143,200 | 0.260290713 | 0.342075481 | 0.151710929 | 0.252564168 |
| 143,300 | 0.260187592 | 0.341972426 | 0.151737982 | 0.252149499 |
| 143,400 | 0.260040344 | 0.340616831 | 0.150804092 | 0.25218329 |
| 143,500 | 0.260185816 | 0.341259764 | 0.151194901 | 0.251911612 |
| 143,600 | 0.260192206 | 0.340789592 | 0.150084971 | 0.25220327 |
| 143,700 | 0.260165625 | 0.339247807 | 0.149177623 | 0.252448232 |
| 143,800 | 0.259900496 | 0.340035003 | 0.150628414 | 0.251277101 |
| 143,900 | 0.259889838 | 0.339499432 | 0.151604298 | 0.251878267 |
| 144,000 | 0.259833498 | 0.340100821 | 0.149914484 | 0.251834377 |
| 144,100 | 0.259745071 | 0.340650659 | 0.15013331 | 0.252453905 |
| 144,200 | 0.25978014 | 0.34025307 | 0.149713214 | 0.251633027 |
| 144,300 | 0.259812026 | 0.340306935 | 0.149021422 | 0.252063938 |
| 144,400 | 0.259814149 | 0.339901349 | 0.150399923 | 0.251803677 |
| 144,500 | 0.259758897 | 0.340445128 | 0.151531305 | 0.251086117 |
| 144,600 | 0.259602452 | 0.339804322 | 0.149328098 | 0.2509346 |
| 144,700 | 0.259544011 | 0.34016647 | 0.149837289 | 0.252041147 |
| 144,800 | 0.259308736 | 0.339031668 | 0.149265237 | 0.251743889 |
| 144,900 | 0.259299385 | 0.339452446 | 0.149651805 | 0.252046493 |
| 145,000 | 0.259254653 | 0.338779626 | 0.14953606 | 0.252021702 |
| 145,100 | 0.259484752 | 0.340039728 | 0.150499931 | 0.252431761 |
| 145,200 | 0.259303916 | 0.339043735 | 0.150260687 | 0.252747807 |
| 145,300 | 0.259323889 | 0.338406156 | 0.1503951 | 0.252814983 |
| 145,400 | 0.259271035 | 0.338391665 | 0.149452371 | 0.252529204 |
| 145,500 | 0.259253742 | 0.339100745 | 0.150253429 | 0.25243289 |
| 145,600 | 0.259218789 | 0.339617981 | 0.148415592 | 0.251718436 |
| 145,700 | 0.259271854 | 0.339143973 | 0.148212601 | 0.252984407 |
| 145,800 | 0.25907269 | 0.338827076 | 0.149944089 | 0.252432405 |
| 145,900 | 0.258737247 | 0.338125154 | 0.147840412 | 0.252133667 |
| 146,000 | 0.258941436 | 0.33790766 | 0.150134121 | 0.252028587 |
| 146,100 | 0.25883099 | 0.3389947 | 0.148268956 | 0.251513566 |
| 146,200 | 0.258785438 | 0.338211674 | 0.148790384 | 0.251703219 |
| 146,300 | 0.258814907 | 0.337028773 | 0.149831663 | 0.25184538 |

| | | | | |
|---|---|---|---|---|
| 146,400 | 0.258643984 | 0.337071132 | 0.148983996 | 0.251678665 |
| 146,500 | 0.258674511 | 0.337711852 | 0.149982433 | 0.252700874 |
| 146,600 | 0.258559259 | 0.337496394 | 0.149735101 | 0.252153552 |
| 146,700 | 0.258579083 | 0.337838637 | 0.146550872 | 0.253046225 |
| 146,800 | 0.258371279 | 0.337360975 | 0.148910675 | 0.252806296 |
| 146,900 | 0.258345812 | 0.337839956 | 0.148969451 | 0.253006792 |
| 147,000 | 0.258208233 | 0.338164674 | 0.151417133 | 0.252053474 |
| 147,100 | 0.258163307 | 0.336959094 | 0.149363001 | 0.251458366 |
| 147,200 | 0.258175721 | 0.337282366 | 0.150016992 | 0.251182118 |
| 147,300 | 0.257975675 | 0.336423751 | 0.150832533 | 0.251618054 |
| 147,400 | 0.257942944 | 0.336563903 | 0.150684714 | 0.251630627 |
| 147,500 | 0.257706913 | 0.336140893 | 0.149478951 | 0.251156814 |
| 147,600 | 0.257783613 | 0.336223056 | 0.148661635 | 0.251500988 |
| 147,700 | 0.257690303 | 0.336389536 | 0.147867834 | 0.251650066 |
| 147,800 | 0.257909948 | 0.335916356 | 0.14871183 | 0.251809231 |
| 147,900 | 0.257853009 | 0.33583789 | 0.148804433 | 0.251518866 |
| 148,000 | 0.257755906 | 0.335531169 | 0.148327192 | 0.252181174 |
| 148,100 | 0.257748183 | 0.335303899 | 0.149137232 | 0.252252175 |
| 148,200 | 0.257603599 | 0.336355308 | 0.149395974 | 0.25264013 |
| 148,300 | 0.257366628 | 0.335230588 | 0.149629209 | 0.252017939 |
| 148,400 | 0.257415126 | 0.335861348 | 0.149325726 | 0.251502238 |
| 148,500 | 0.257189554 | 0.336445934 | 0.147633437 | 0.251407283 |
| 148,600 | 0.257303536 | 0.336022201 | 0.148801314 | 0.251710417 |
| 148,700 | 0.25728383 | 0.33543139 | 0.14878545 | 0.251771592 |
| 148,800 | 0.257329491 | 0.335399555 | 0.148808099 | 0.252747307 |
| 148,900 | 0.257281982 | 0.336106504 | 0.148448081 | 0.252987288 |
| 149,000 | 0.257145844 | 0.335321582 | 0.149401382 | 0.251524808 |
| 149,100 | 0.256969391 | 0.334755736 | 0.148576073 | 0.252105734 |
| 149,200 | 0.256820841 | 0.334683635 | 0.148618532 | 0.251826839 |
| 149,300 | 0.25694391 | 0.33521074 | 0.148047269 | 0.252676966 |
| 149,400 | 0.256944421 | 0.335173606 | 0.147741421 | 0.25232612 |
| 149,500 | 0.256865763 | 0.334776665 | 0.148354962 | 0.25236086 |
| 149,600 | 0.256784214 | 0.334613017 | 0.148101467 | 0.252335178 |
| 149,700 | 0.256854452 | 0.334835276 | 0.147692568 | 0.251846036 |
| 149,800 | 0.256735682 | 0.334674913 | 0.149137156 | 0.251718631 |
| 149,900 | 0.256592708 | 0.333577927 | 0.147637348 | 0.251935345 |
| 150,000 | 0.256518974 | 0.333877746 | 0.148618506 | 0.251775706 |
| 150,100 | 0.256509499 | 0.333588078 | 0.149271442 | 0.25050084 |
| 150,200 | 0.25652127 | 0.335001485 | 0.149637409 | 0.251907534 |
| 150,300 | 0.256491644 | 0.334273498 | 0.149085199 | 0.251824018 |
| 150,400 | 0.256578342 | 0.334125611 | 0.146946304 | 0.252326547 |
| 150,500 | 0.25633158 | 0.334288153 | 0.147465281 | 0.250999106 |
| 150,600 | 0.256229593 | 0.334490435 | 0.148212194 | 0.251165885 |
| 150,700 | 0.256207487 | 0.333703865 | 0.146599186 | 0.250213655 |
| 150,800 | 0.255822083 | 0.333363821 | 0.148239642 | 0.24991441 |
| 150,900 | 0.255994544 | 0.333218016 | 0.149544299 | 0.24979423 |
| 151,000 | 0.256032156 | 0.33341087 | 0.148592624 | 0.249997422 |
| 151,100 | 0.256015301 | 0.333435068 | 0.147362158 | 0.249965396 |
| 151,200 | 0.256036081 | 0.332663685 | 0.149040977 | 0.249675753 |
| 151,300 | 0.255884854 | 0.334016106 | 0.146910325 | 0.249659285 |
| 151,400 | 0.255875306 | 0.333421553 | 0.147492936 | 0.250122482 |
| 151,500 | 0.255794011 | 0.33356286 | 0.148450606 | 0.250016553 |
| 151,600 | 0.255808233 | 0.332951362 | 0.149261185 | 0.249069207 |
| 151,700 | 0.255687795 | 0.333374269 | 0.147874736 | 0.249206035 |
| 151,800 | 0.255747263 | 0.333117772 | 0.147217434 | 0.249792621 |
| 151,900 | 0.255639473 | 0.333096192 | 0.147467675 | 0.249003746 |
| 152,000 | 0.255567143 | 0.332441189 | 0.147721262 | 0.250029158 |
| 152,100 | 0.25526766 | 0.332906089 | 0.146772243 | 0.248799962 |
| 152,200 | 0.255309178 | 0.333057331 | 0.147554952 | 0.248105997 |
| 152,300 | 0.255254237 | 0.333102028 | 0.148117977 | 0.248931473 |
| 152,400 | 0.255127694 | 0.334362691 | 0.147927709 | 0.248032092 |
| 152,500 | 0.255192076 | 0.3343686 | 0.148346472 | 0.24825954 |
| 152,600 | 0.255273861 | 0.33385414 | 0.150585516 | 0.248500434 |
| 152,700 | 0.255210068 | 0.334228189 | 0.149925296 | 0.24843148 |
| 152,800 | 0.255067558 | 0.333347226 | 0.148482029 | 0.247665317 |
| 152,900 | 0.254954855 | 0.333481686 | 0.148125341 | 0.248481969 |
| 153,000 | 0.254938598 | 0.333265013 | 0.148261978 | 0.247553176 |

| | | | | |
|---|---|---|---|---|
| 153,100 | 0.254643122 | 0.331622872 | 0.146602632 | 0.248381871 |
| 153,200 | 0.254847389 | 0.331961424 | 0.147517562 | 0.247801166 |
| 153,300 | 0.254958772 | 0.332070637 | 0.147846942 | 0.24751405 |
| 153,400 | 0.254933025 | 0.332279881 | 0.148033508 | 0.247350787 |
| 153,500 | 0.254935653 | 0.332757133 | 0.147483972 | 0.2489379 |
| 153,600 | 0.254831391 | 0.332534407 | 0.147741116 | 0.24784175 |
| 153,700 | 0.254688396 | 0.333079299 | 0.146569285 | 0.247712673 |
| 153,800 | 0.254668156 | 0.331968767 | 0.147183973 | 0.247502951 |
| 153,900 | 0.254513081 | 0.332552057 | 0.148455762 | 0.24836528 |
| 154,000 | 0.254678019 | 0.332839938 | 0.147811991 | 0.24801826 |
| 154,100 | 0.254402236 | 0.331696139 | 0.148296298 | 0.247123169 |
| 154,200 | 0.254399125 | 0.331761381 | 0.147586291 | 0.247379517 |
| 154,300 | 0.254352688 | 0.332263469 | 0.147966976 | 0.247757553 |
| 154,400 | 0.254306202 | 0.332174176 | 0.147944757 | 0.247016814 |
| 154,500 | 0.254187523 | 0.33195904 | 0.148138755 | 0.247422165 |
| 154,600 | 0.254152669 | 0.33211765 | 0.148509983 | 0.246648278 |
| 154,700 | 0.254149176 | 0.33137671 | 0.14854466 | 0.247224612 |
| 154,800 | 0.253990827 | 0.331728202 | 0.148502598 | 0.246774985 |
| 154,900 | 0.253976129 | 0.331063548 | 0.147407507 | 0.247011431 |
| 155,000 | 0.253956484 | 0.330588242 | 0.148327402 | 0.247290991 |
| 155,100 | 0.25387177 | 0.331875821 | 0.1465801 | 0.246909649 |
| 155,200 | 0.253892593 | 0.331764163 | 0.148756388 | 0.245268413 |
| 155,300 | 0.254101343 | 0.332139462 | 0.15012341 | 0.24657247 |
| 155,400 | 0.253914883 | 0.330620138 | 0.14848675 | 0.246443027 |
| 155,500 | 0.253855444 | 0.331115362 | 0.150026534 | 0.247259841 |
| 155,600 | 0.253806269 | 0.330427094 | 0.148648963 | 0.247192626 |
| 155,700 | 0.253721748 | 0.330740918 | 0.147069493 | 0.247643228 |
| 155,800 | 0.253674259 | 0.33196491 | 0.147572686 | 0.247106448 |
| 155,900 | 0.253513926 | 0.332163764 | 0.147670777 | 0.246840382 |
| 156,000 | 0.253528771 | 0.332265393 | 0.148838236 | 0.246815674 |
| 156,100 | 0.253576428 | 0.33226359 | 0.146864795 | 0.247646314 |
| 156,200 | 0.253765943 | 0.331895426 | 0.14772874 | 0.247225788 |
| 156,300 | 0.253660588 | 0.330892793 | 0.147330958 | 0.246224671 |
| 156,400 | 0.253583612 | 0.331388533 | 0.147714977 | 0.246909962 |
| 156,500 | 0.253526899 | 0.331955483 | 0.147286122 | 0.247022318 |
| 156,600 | 0.253588103 | 0.331689565 | 0.146608341 | 0.2470962 |
| 156,700 | 0.253307478 | 0.331469156 | 0.145358229 | 0.247548422 |
| 156,800 | 0.253262205 | 0.330219283 | 0.145604573 | 0.247501765 |
| 156,900 | 0.253206966 | 0.330486626 | 0.145907123 | 0.24700068 |
| 157,000 | 0.253141941 | 0.330704121 | 0.144864498 | 0.247105199 |
| 157,100 | 0.253376558 | 0.330372339 | 0.143408018 | 0.248063465 |
| 157,200 | 0.253045248 | 0.330905838 | 0.145433555 | 0.246775951 |
| 157,300 | 0.253106796 | 0.331319027 | 0.144158601 | 0.246780309 |
| 157,400 | 0.252918598 | 0.331567725 | 0.144493924 | 0.246794808 |
| 157,500 | 0.252779507 | 0.330704611 | 0.144607863 | 0.246581013 |
| 157,600 | 0.252858565 | 0.330556234 | 0.145501957 | 0.246411447 |
| 157,700 | 0.252537726 | 0.330501148 | 0.143729385 | 0.246349112 |
| 157,800 | 0.252719628 | 0.331080737 | 0.142522568 | 0.245852359 |
| 157,900 | 0.252726872 | 0.331018263 | 0.143239537 | 0.24597533 |
| 158,000 | 0.252448291 | 0.331294535 | 0.143147724 | 0.245208678 |
| 158,100 | 0.252540181 | 0.330599589 | 0.142734109 | 0.245901786 |
| 158,200 | 0.25249612 | 0.329496938 | 0.142273933 | 0.245253191 |
| 158,300 | 0.252597771 | 0.330877071 | 0.143062856 | 0.244972014 |
| 158,400 | 0.252396882 | 0.330677313 | 0.144861024 | 0.245873186 |
| 158,500 | 0.252229835 | 0.330193985 | 0.14268855 | 0.245651165 |
| 158,600 | 0.252355531 | 0.330761583 | 0.145118159 | 0.245237211 |
| 158,700 | 0.252187997 | 0.330296614 | 0.144488836 | 0.245019035 |
| 158,800 | 0.252287451 | 0.330609342 | 0.144402984 | 0.245471717 |
| 158,900 | 0.252015312 | 0.329550859 | 0.142957145 | 0.244912625 |
| 159,000 | 0.25202681 | 0.329862013 | 0.144385603 | 0.245149766 |
| 159,100 | 0.252140677 | 0.330032158 | 0.141306216 | 0.244436358 |
| 159,200 | 0.25232597 | 0.329790522 | 0.141834868 | 0.245063382 |
| 159,300 | 0.251986455 | 0.329118386 | 0.142410139 | 0.245927947 |
| 159,400 | 0.25229583 | 0.329284403 | 0.141730322 | 0.246358782 |
| 159,500 | 0.251966987 | 0.329950443 | 0.141910118 | 0.245602573 |
| 159,600 | 0.252146513 | 0.329688101 | 0.141493783 | 0.245124823 |
| 159,700 | 0.251972832 | 0.329521355 | 0.141500962 | 0.244843168 |

| | | | | |
|---|---|---|---|---|
| 159,800 | 0.251968309 | 0.330355399 | 0.14141139 | 0.244816833 |
| 159,900 | 0.251857666 | 0.330282751 | 0.140537045 | 0.244676589 |
| 160,000 | 0.252024486 | 0.330116142 | 0.140678873 | 0.245264109 |
| 160,100 | 0.251694676 | 0.330237716 | 0.141401435 | 0.244043357 |
| 160,200 | 0.251644423 | 0.329568964 | 0.141566928 | 0.244065117 |
| 160,300 | 0.251573152 | 0.330067488 | 0.141687337 | 0.244619346 |
| 160,400 | 0.251384054 | 0.330097737 | 0.140949224 | 0.244017546 |
| 160,500 | 0.251403436 | 0.328889175 | 0.139346556 | 0.244059315 |
| 160,600 | 0.251485874 | 0.32911806 | 0.141433021 | 0.243984379 |
| 160,700 | 0.251579941 | 0.328418103 | 0.141785605 | 0.244159331 |
| 160,800 | 0.251428581 | 0.329026411 | 0.141706669 | 0.244106271 |
| 160,900 | 0.25131205 | 0.328597444 | 0.140562948 | 0.243478252 |
| 161,000 | 0.251453113 | 0.329951592 | 0.140279559 | 0.242747452 |
| 161,100 | 0.25111088 | 0.32935406 | 0.139409262 | 0.24337038 |
| 161,200 | 0.251260541 | 0.328200853 | 0.140226136 | 0.244048438 |
| 161,300 | 0.251126807 | 0.329031164 | 0.139668126 | 0.244377508 |
| 161,400 | 0.251088175 | 0.327983434 | 0.138464777 | 0.243520329 |
| 161,500 | 0.250929841 | 0.328355718 | 0.139700703 | 0.243980017 |
| 161,600 | 0.251042659 | 0.328022924 | 0.140303195 | 0.244969507 |
| 161,700 | 0.250935617 | 0.327813817 | 0.137985493 | 0.243580447 |
| 161,800 | 0.250727463 | 0.327239027 | 0.138885513 | 0.241971465 |
| 161,900 | 0.250884742 | 0.327556438 | 0.13721391 | 0.243571578 |
| 162,000 | 0.250578961 | 0.326763828 | 0.137578364 | 0.243230957 |
| 162,100 | 0.250753626 | 0.326826371 | 0.138389402 | 0.243580354 |
| 162,200 | 0.250664303 | 0.327136333 | 0.13908082 | 0.244020373 |
| 162,300 | 0.250641486 | 0.326777925 | 0.137847838 | 0.243752599 |
| 162,400 | 0.250589777 | 0.327155861 | 0.138418577 | 0.243641081 |
| 162,500 | 0.25057563 | 0.327174943 | 0.137069126 | 0.243487615 |
| 162,600 | 0.250350778 | 0.326282553 | 0.136875032 | 0.243201912 |
| 162,700 | 0.250385187 | 0.326140588 | 0.138010925 | 0.242372988 |
| 162,800 | 0.250393567 | 0.327009904 | 0.137331053 | 0.243308631 |
| 162,900 | 0.250449317 | 0.326405492 | 0.136761808 | 0.242729536 |
| 163,000 | 0.250356369 | 0.325403715 | 0.139564345 | 0.24264065 |
| 163,100 | 0.250297345 | 0.325462255 | 0.138034159 | 0.242873201 |
| 163,200 | 0.250139212 | 0.325381199 | 0.137517163 | 0.242892886 |
| 163,300 | 0.250052201 | 0.325259596 | 0.13935931 | 0.242070354 |
| 163,400 | 0.249976229 | 0.325537616 | 0.139792341 | 0.24258237 |
| 163,500 | 0.249822787 | 0.324478474 | 0.139409043 | 0.242015879 |
| 163,600 | 0.249873724 | 0.325179585 | 0.140166268 | 0.242031831 |
| 163,700 | 0.24997308 | 0.325436989 | 0.13953734 | 0.241659441 |
| 163,800 | 0.249881581 | 0.324796179 | 0.138695706 | 0.242430169 |
| 163,900 | 0.249845329 | 0.324635798 | 0.137866303 | 0.242294737 |
| 164,000 | 0.249797083 | 0.325308802 | 0.137809309 | 0.242551367 |
| 164,100 | 0.249791374 | 0.32438526 | 0.1373191 | 0.242400015 |
| 164,200 | 0.249809569 | 0.32479608 | 0.137899311 | 0.242100382 |
| 164,300 | 0.24988383 | 0.324808981 | 0.136942419 | 0.243001713 |
| 164,400 | 0.249548157 | 0.32391651 | 0.13812011 | 0.241858184 |
| 164,500 | 0.249619922 | 0.324732094 | 0.137936046 | 0.241612027 |
| 164,600 | 0.249636095 | 0.325049828 | 0.138544431 | 0.241757225 |
| 164,700 | 0.249348242 | 0.325044985 | 0.137179817 | 0.241239952 |
| 164,800 | 0.2495206 | 0.324961069 | 0.136903204 | 0.241345493 |
| 164,900 | 0.249332946 | 0.324942551 | 0.13665022 | 0.242561235 |
| 165,000 | 0.249247064 | 0.324394758 | 0.137908789 | 0.241599651 |
| 165,100 | 0.249268218 | 0.324368618 | 0.13782741 | 0.240995811 |
| 165,200 | 0.249184118 | 0.324492229 | 0.136952967 | 0.240778395 |
| 165,300 | 0.249078852 | 0.324572234 | 0.136767128 | 0.241348291 |
| 165,400 | 0.248953352 | 0.325129836 | 0.136737133 | 0.240883715 |
| 165,500 | 0.248993768 | 0.325251782 | 0.136277007 | 0.241469514 |
| 165,600 | 0.248818495 | 0.324083395 | 0.135415548 | 0.240764665 |
| 165,700 | 0.248884769 | 0.324323496 | 0.13627352 | 0.241204509 |
| 165,800 | 0.248720651 | 0.324175382 | 0.135454162 | 0.241531464 |
| 165,900 | 0.248555656 | 0.323985882 | 0.134902859 | 0.240211827 |
| 166,000 | 0.248512291 | 0.324291198 | 0.13530717 | 0.24041812 |
| 166,100 | 0.248337743 | 0.324650291 | 0.134746114 | 0.239978003 |
| 166,200 | 0.248519077 | 0.325134736 | 0.134674845 | 0.240941993 |
| 166,300 | 0.24848736 | 0.3243999 | 0.135990364 | 0.240728396 |
| 166,400 | 0.248460892 | 0.324259296 | 0.136588662 | 0.240731496 |

| | | | | |
|---|---|---|---|---|
| 166,500 | 0.248413423 | 0.32438826 | 0.136028254 | 0.240612149 |
| 166,600 | 0.248112241 | 0.32329792 | 0.136558091 | 0.240846684 |
| 166,700 | 0.248130189 | 0.323397446 | 0.135221369 | 0.240882443 |
| 166,800 | 0.248207 | 0.32359393 | 0.134316347 | 0.241101217 |
| 166,900 | 0.248180537 | 0.322856743 | 0.135026069 | 0.240921122 |
| 167,000 | 0.247946386 | 0.322234997 | 0.136161896 | 0.239951655 |
| 167,100 | 0.247845796 | 0.323134676 | 0.13507793 | 0.240526986 |
| 167,200 | 0.247822317 | 0.322663689 | 0.13527291 | 0.240377723 |
| 167,300 | 0.247906752 | 0.322801844 | 0.134191397 | 0.240397881 |
| 167,400 | 0.247751766 | 0.321988522 | 0.134034267 | 0.24047185 |
| 167,500 | 0.24773259 | 0.322443639 | 0.133679376 | 0.240378564 |
| 167,600 | 0.247854286 | 0.322979183 | 0.133383542 | 0.240317812 |
| 167,700 | 0.247625909 | 0.322717544 | 0.134788122 | 0.240058022 |
| 167,800 | 0.247491178 | 0.323276503 | 0.134321645 | 0.240479802 |
| 167,900 | 0.247590862 | 0.324062189 | 0.133362135 | 0.240954664 |
| 168,000 | 0.247472295 | 0.323465008 | 0.136009678 | 0.24036836 |
| 168,100 | 0.247397149 | 0.323555921 | 0.135060271 | 0.23989791 |
| 168,200 | 0.247404991 | 0.323867784 | 0.133434099 | 0.24022782 |
| 168,300 | 0.247516876 | 0.323218479 | 0.134949215 | 0.239671752 |
| 168,400 | 0.247453289 | 0.323518612 | 0.134574952 | 0.239731325 |
| 168,500 | 0.247229307 | 0.323060604 | 0.135458443 | 0.23983284 |
| 168,600 | 0.247402371 | 0.32401348 | 0.133930937 | 0.239326839 |
| 168,700 | 0.24742419 | 0.323427049 | 0.133695931 | 0.239333629 |
| 168,800 | 0.247206144 | 0.322485929 | 0.132750234 | 0.239204649 |
| 168,900 | 0.247145085 | 0.322683835 | 0.132966111 | 0.238554632 |
| 169,000 | 0.247091154 | 0.322592589 | 0.13223014 | 0.238972779 |
| 169,100 | 0.247076914 | 0.322614824 | 0.130990383 | 0.23884247 |
| 169,200 | 0.247142883 | 0.321969852 | 0.130592732 | 0.238626541 |
| 169,300 | 0.247039517 | 0.322832086 | 0.131371972 | 0.238329371 |
| 169,400 | 0.246994601 | 0.32092157 | 0.130850571 | 0.238696267 |
| 169,500 | 0.246969863 | 0.322309253 | 0.13202391 | 0.238713493 |
| 169,600 | 0.246866166 | 0.32105374 | 0.131414389 | 0.237705803 |
| 169,700 | 0.24678536 | 0.32212036 | 0.130613879 | 0.23741538 |
| 169,800 | 0.24700057 | 0.321550917 | 0.131832316 | 0.238017506 |
| 169,900 | 0.246809478 | 0.321546334 | 0.132393506 | 0.237286287 |
| 170,000 | 0.246665612 | 0.321688947 | 0.13167127 | 0.238733721 |
| 170,100 | 0.246642524 | 0.322130942 | 0.131826351 | 0.238737039 |
| 170,200 | 0.246671542 | 0.320763457 | 0.13190469 | 0.238248242 |
| 170,300 | 0.246653647 | 0.321438939 | 0.131909405 | 0.238670082 |
| 170,400 | 0.246548201 | 0.321142463 | 0.132733518 | 0.238458879 |
| 170,500 | 0.246484213 | 0.321302571 | 0.132693348 | 0.238471457 |
| 170,600 | 0.246475428 | 0.322397371 | 0.132171597 | 0.238264664 |
| 170,700 | 0.246258205 | 0.320898629 | 0.133221232 | 0.238454426 |
| 170,800 | 0.246264867 | 0.320834329 | 0.13216816 | 0.238304263 |
| 170,900 | 0.24618582 | 0.320802527 | 0.131664844 | 0.238584514 |
| 171,000 | 0.24628502 | 0.321598904 | 0.132520696 | 0.238298924 |
| 171,100 | 0.24608021 | 0.320448729 | 0.132600072 | 0.237589993 |
| 171,200 | 0.245997328 | 0.319580333 | 0.131024364 | 0.23810615 |
| 171,300 | 0.246052271 | 0.319884674 | 0.131656806 | 0.238248853 |
| 171,400 | 0.245975417 | 0.320933183 | 0.131435342 | 0.238441337 |
| 171,500 | 0.245867514 | 0.320737417 | 0.131251965 | 0.238578967 |
| 171,600 | 0.245926485 | 0.320581538 | 0.132013856 | 0.23864356 |
| 171,700 | 0.245851169 | 0.31946534 | 0.132348152 | 0.239327145 |
| 171,800 | 0.245849248 | 0.32089626 | 0.132599367 | 0.23953113 |
| 171,900 | 0.245798368 | 0.320572467 | 0.131647221 | 0.238441357 |
| 172,000 | 0.245725129 | 0.320697885 | 0.13348605 | 0.238184506 |
| 172,100 | 0.245660191 | 0.320216199 | 0.13210059 | 0.238439407 |
| 172,200 | 0.245623856 | 0.319982493 | 0.132965023 | 0.238384904 |
| 172,300 | 0.245514903 | 0.319706621 | 0.132251675 | 0.23822005 |
| 172,400 | 0.245473504 | 0.320298739 | 0.13248514 | 0.238116314 |
| 172,500 | 0.245519001 | 0.319938959 | 0.132208657 | 0.238403606 |
| 172,600 | 0.245626406 | 0.320659583 | 0.13190352 | 0.238060952 |
| 172,700 | 0.245366751 | 0.319838073 | 0.130758214 | 0.238285927 |
| 172,800 | 0.245174797 | 0.320187748 | 0.131604506 | 0.237488147 |
| 172,900 | 0.245139928 | 0.32043865 | 0.130230394 | 0.238445686 |
| 173,000 | 0.245340218 | 0.319910426 | 0.133065307 | 0.238264848 |
| 173,100 | 0.245210569 | 0.319608446 | 0.131871566 | 0.238510526 |

| | | | | |
|---|---|---|---|---|
| 173,200 | 0.244977955 | 0.319556911 | 0.133366857 | 0.237192367 |
| 173,300 | 0.245094082 | 0.320192281 | 0.13225503 | 0.238614757 |
| 173,400 | 0.245090983 | 0.31958988 | 0.132393833 | 0.237974727 |
| 173,500 | 0.24510398 | 0.319932376 | 0.132965583 | 0.23839338 |
| 173,600 | 0.245056636 | 0.31961721 | 0.132753213 | 0.238211136 |
| 173,700 | 0.245256586 | 0.319737064 | 0.131662138 | 0.238813167 |
| 173,800 | 0.24514096 | 0.319075814 | 0.133097474 | 0.238502484 |
| 173,900 | 0.24500764 | 0.319498816 | 0.131740186 | 0.238261981 |
| 174,000 | 0.244795329 | 0.319942747 | 0.130699456 | 0.238132342 |
| 174,100 | 0.244831477 | 0.319146257 | 0.131316581 | 0.23786877 |
| 174,200 | 0.244788498 | 0.318957499 | 0.131727586 | 0.237755671 |
| 174,300 | 0.244816458 | 0.318599162 | 0.132553169 | 0.237882563 |
| 174,400 | 0.244798014 | 0.317622677 | 0.132790127 | 0.238149272 |
| 174,500 | 0.244680691 | 0.318515821 | 0.132488643 | 0.23733995 |
| 174,600 | 0.244513505 | 0.319039231 | 0.133216409 | 0.237800759 |
| 174,700 | 0.244672943 | 0.318851194 | 0.133236643 | 0.237799877 |
| 174,800 | 0.244555852 | 0.318515135 | 0.133312196 | 0.237949904 |
| 174,900 | 0.244619297 | 0.31893505 | 0.13364119 | 0.237287095 |
| 175,000 | 0.24455833 | 0.318291832 | 0.133073846 | 0.237821732 |
| 175,100 | 0.244311305 | 0.318761352 | 0.132015246 | 0.236907985 |
| 175,200 | 0.244482569 | 0.319108005 | 0.131551405 | 0.237725176 |
| 175,300 | 0.244577877 | 0.31892392 | 0.133890742 | 0.238095042 |
| 175,400 | 0.244508051 | 0.31824376 | 0.132679609 | 0.238048036 |
| 175,500 | 0.244520727 | 0.318349279 | 0.133106132 | 0.236779083 |
| 175,600 | 0.244329124 | 0.318798819 | 0.131674768 | 0.237351266 |
| 175,700 | 0.244421108 | 0.318122215 | 0.132757871 | 0.236952597 |
| 175,800 | 0.244417715 | 0.318174927 | 0.13242688 | 0.237405885 |
| 175,900 | 0.244339049 | 0.31727143 | 0.133245939 | 0.237119899 |
| 176,000 | 0.244330934 | 0.317536167 | 0.132996609 | 0.237111549 |
| 176,100 | 0.24412543 | 0.317763455 | 0.131195547 | 0.237766302 |
| 176,200 | 0.244149129 | 0.318321904 | 0.131553374 | 0.23749896 |
| 176,300 | 0.244168794 | 0.317927292 | 0.132137902 | 0.237079319 |
| 176,400 | 0.244091413 | 0.318806148 | 0.133668387 | 0.236472423 |
| 176,500 | 0.244188383 | 0.318382931 | 0.133535823 | 0.237392085 |
| 176,600 | 0.244014332 | 0.318636606 | 0.132667093 | 0.236502486 |
| 176,700 | 0.244203623 | 0.318439989 | 0.133108331 | 0.236237755 |
| 176,800 | 0.244097097 | 0.317813034 | 0.132722515 | 0.23634485 |
| 176,900 | 0.244027112 | 0.318273316 | 0.133418715 | 0.236178251 |
| 177,000 | 0.243992552 | 0.317968695 | 0.133010077 | 0.235693345 |
| 177,100 | 0.243938994 | 0.31788837 | 0.134168106 | 0.236646056 |
| 177,200 | 0.243762105 | 0.318171638 | 0.132299279 | 0.237147904 |
| 177,300 | 0.243833145 | 0.31805382 | 0.132020792 | 0.236254981 |
| 177,400 | 0.243574508 | 0.31733732 | 0.132936075 | 0.236169608 |
| 177,500 | 0.243703728 | 0.317689734 | 0.134590775 | 0.235725207 |
| 177,600 | 0.243568326 | 0.318522559 | 0.134192909 | 0.236288213 |
| 177,700 | 0.243483557 | 0.317186751 | 0.133208607 | 0.236173385 |
| 177,800 | 0.243431601 | 0.316342717 | 0.132481183 | 0.236614569 |
| 177,900 | 0.243506614 | 0.318177305 | 0.133114929 | 0.236004934 |
| 178,000 | 0.243550759 | 0.316493272 | 0.133638258 | 0.237112026 |
| 178,100 | 0.243467333 | 0.316402661 | 0.133179362 | 0.236051117 |
| 178,200 | 0.243038665 | 0.316832415 | 0.13288481 | 0.235500373 |
| 178,300 | 0.243219435 | 0.31675712 | 0.13273918 | 0.236545561 |
| 178,400 | 0.242929102 | 0.3171563 | 0.13232497 | 0.235641383 |
| 178,500 | 0.243078605 | 0.319035653 | 0.131718059 | 0.235520096 |
| 178,600 | 0.24284894 | 0.317064138 | 0.130903056 | 0.235181963 |
| 178,700 | 0.242949359 | 0.318421838 | 0.131493657 | 0.235094705 |
| 178,800 | 0.242940342 | 0.31750283 | 0.132631013 | 0.235020455 |
| 178,900 | 0.242781009 | 0.316859297 | 0.133182655 | 0.235057888 |
| 179,000 | 0.242817003 | 0.316937505 | 0.131606343 | 0.235305717 |
| 179,100 | 0.242600591 | 0.317533356 | 0.132936366 | 0.235422336 |
| 179,200 | 0.242523093 | 0.316348581 | 0.132320905 | 0.235264975 |
| 179,300 | 0.242391602 | 0.316937936 | 0.130972586 | 0.23475813 |
| 179,400 | 0.242565793 | 0.317076583 | 0.132240201 | 0.234850739 |
| 179,500 | 0.242424861 | 0.317113677 | 0.133011826 | 0.234876189 |
| 179,600 | 0.242209247 | 0.31601878 | 0.131135459 | 0.234746015 |
| 179,700 | 0.242246824 | 0.317413761 | 0.13286033 | 0.234233284 |
| 179,800 | 0.242246572 | 0.318630005 | 0.130657783 | 0.234740775 |

| | | | | |
|---|---|---|---|---|
| 179,900 | 0.24215002 | 0.317578884 | 0.132027195 | 0.234325455 |
| 180,000 | 0.241970182 | 0.317464878 | 0.131704208 | 0.234235237 |
| 180,100 | 0.241950887 | 0.317337153 | 0.131357787 | 0.234320239 |
| 180,200 | 0.241829668 | 0.317565217 | 0.131856117 | 0.23329027 |
| 180,300 | 0.241868729 | 0.317628329 | 0.129721559 | 0.234135164 |
| 180,400 | 0.241843782 | 0.317333796 | 0.129615294 | 0.234053904 |
| 180,500 | 0.242056122 | 0.31691968 | 0.129735447 | 0.233455298 |
| 180,600 | 0.241819317 | 0.317799516 | 0.128588946 | 0.23411158 |
| 180,700 | 0.241786424 | 0.317726391 | 0.129866856 | 0.233779493 |
| 180,800 | 0.241610336 | 0.317775054 | 0.129190427 | 0.233995325 |
| 180,900 | 0.24176801 | 0.31829973 | 0.130368479 | 0.234824892 |
| 181,000 | 0.24156424 | 0.318174802 | 0.129833331 | 0.234294393 |
| 181,100 | 0.241624177 | 0.317870473 | 0.129430713 | 0.233632135 |
| 181,200 | 0.241640003 | 0.317292003 | 0.128333101 | 0.233412787 |
| 181,300 | 0.241311909 | 0.317397861 | 0.128198682 | 0.234714344 |
| 181,400 | 0.241313912 | 0.316954795 | 0.129317006 | 0.234651278 |
| 181,500 | 0.241453949 | 0.317881834 | 0.129153458 | 0.234081058 |
| 181,600 | 0.241320648 | 0.317922149 | 0.129962033 | 0.233775347 |
| 181,700 | 0.241180538 | 0.317896101 | 0.128224946 | 0.232544542 |
| 181,800 | 0.241072469 | 0.317954513 | 0.128646977 | 0.233019208 |
| 181,900 | 0.241102108 | 0.317317958 | 0.12976965 | 0.233019867 |
| 182,000 | 0.24084075 | 0.318521302 | 0.130384591 | 0.233290708 |
| 182,100 | 0.241097411 | 0.318538128 | 0.129724619 | 0.233311883 |
| 182,200 | 0.240958932 | 0.31755876 | 0.129180289 | 0.232851718 |
| 182,300 | 0.240872283 | 0.317482286 | 0.130121212 | 0.233020835 |
| 182,400 | 0.240845499 | 0.317955056 | 0.128858933 | 0.232824681 |
| 182,500 | 0.240698844 | 0.318957 | 0.129941321 | 0.232338357 |
| 182,600 | 0.240872956 | 0.319193461 | 0.129389098 | 0.233642038 |
| 182,700 | 0.240893956 | 0.318312493 | 0.129633482 | 0.23288691 |
| 182,800 | 0.240909617 | 0.31769626 | 0.129189552 | 0.232883777 |
| 182,900 | 0.240767454 | 0.317381244 | 0.131192893 | 0.231515294 |
| 183,000 | 0.24066258 | 0.317782896 | 0.131778109 | 0.231791182 |
| 183,100 | 0.24059405 | 0.317725816 | 0.12967291 | 0.232139502 |
| 183,200 | 0.240550268 | 0.317586136 | 0.130396287 | 0.232938573 |
| 183,300 | 0.240220314 | 0.317776075 | 0.131337369 | 0.232645948 |
| 183,400 | 0.240481091 | 0.31839726 | 0.131517907 | 0.232557637 |
| 183,500 | 0.240349768 | 0.319095155 | 0.130445647 | 0.232879918 |
| 183,600 | 0.240370012 | 0.316987713 | 0.130186302 | 0.232812561 |
| 183,700 | 0.240331847 | 0.318164339 | 0.131607463 | 0.231659271 |
| 183,800 | 0.240208402 | 0.318171906 | 0.129517781 | 0.232151998 |
| 183,900 | 0.240207602 | 0.317778771 | 0.129263943 | 0.232728746 |
| 184,000 | 0.240199325 | 0.317290558 | 0.128725299 | 0.231686535 |
| 184,100 | 0.240197579 | 0.317486239 | 0.128116428 | 0.231957808 |
| 184,200 | 0.240193755 | 0.317644231 | 0.128689446 | 0.232492505 |
| 184,300 | 0.240039874 | 0.317362893 | 0.130478347 | 0.232456529 |
| 184,400 | 0.240066155 | 0.318086493 | 0.129850731 | 0.23216918 |
| 184,500 | 0.239922427 | 0.318247135 | 0.129271492 | 0.23202902 |
| 184,600 | 0.239995136 | 0.318189391 | 0.129559826 | 0.232627234 |
| 184,700 | 0.240085272 | 0.317393147 | 0.129866495 | 0.232419774 |
| 184,800 | 0.239911047 | 0.318198983 | 0.130867111 | 0.231884907 |
| 184,900 | 0.239957065 | 0.318589406 | 0.129953894 | 0.232732327 |
| 185,000 | 0.239713603 | 0.318005496 | 0.129739204 | 0.232324325 |
| 185,100 | 0.239639701 | 0.318311441 | 0.129661337 | 0.232554118 |
| 185,200 | 0.239596944 | 0.318196546 | 0.130052675 | 0.232554332 |
| 185,300 | 0.239707302 | 0.318891056 | 0.129806967 | 0.232732077 |
| 185,400 | 0.239710705 | 0.318902314 | 0.130825346 | 0.233087225 |
| 185,500 | 0.239565686 | 0.317681079 | 0.130624946 | 0.232848836 |
| 185,600 | 0.239437642 | 0.31671033 | 0.131277527 | 0.232533706 |
| 185,700 | 0.23958261 | 0.318564447 | 0.129206825 | 0.232792734 |
| 185,800 | 0.239640193 | 0.317768154 | 0.130803098 | 0.233543417 |
| 185,900 | 0.239491053 | 0.319031992 | 0.129875231 | 0.233422875 |
| 186,000 | 0.239457007 | 0.31879873 | 0.130084455 | 0.2327199 |
| 186,100 | 0.239493654 | 0.318465319 | 0.131910021 | 0.23271106 |
| 186,200 | 0.23924405 | 0.319223446 | 0.130565905 | 0.23253343 |
| 186,300 | 0.239238624 | 0.318703009 | 0.129563327 | 0.232870555 |
| 186,400 | 0.239212545 | 0.318134949 | 0.129635049 | 0.232301092 |
| 186,500 | 0.239353725 | 0.31851785 | 0.130598743 | 0.232540899 |

| | | | | |
|---|---|---|---|---|
| 186,600 | 0.239265406 | 0.318437005 | 0.130594954 | 0.232983363 |
| 186,700 | 0.23920798 | 0.317616948 | 0.130338869 | 0.232536264 |
| 186,800 | 0.239109605 | 0.318197175 | 0.13154637 | 0.232704417 |
| 186,900 | 0.239148651 | 0.317376299 | 0.130227459 | 0.232974318 |
| 187,000 | 0.239002214 | 0.317309307 | 0.130248132 | 0.233435035 |
| 187,100 | 0.239042921 | 0.317423869 | 0.131571458 | 0.233762882 |
| 187,200 | 0.238872351 | 0.317209173 | 0.13061577 | 0.233092297 |
| 187,300 | 0.238884788 | 0.317487034 | 0.129295503 | 0.233503808 |
| 187,400 | 0.238579508 | 0.315450397 | 0.130497889 | 0.23298328 |
| 187,500 | 0.238644579 | 0.316082407 | 0.12968461 | 0.23252855 |
| 187,600 | 0.238707222 | 0.315906434 | 0.128522591 | 0.233158331 |
| 187,700 | 0.238724482 | 0.315735699 | 0.130439299 | 0.232738305 |
| 187,800 | 0.238517829 | 0.315821697 | 0.130337442 | 0.232294192 |
| 187,900 | 0.238531759 | 0.31593981 | 0.131796269 | 0.231986361 |
| 188,000 | 0.238598444 | 0.31543598 | 0.13203991 | 0.232099336 |
| 188,100 | 0.238383151 | 0.315254887 | 0.131135554 | 0.232233972 |
| 188,200 | 0.238480864 | 0.31527941 | 0.131786068 | 0.231785791 |
| 188,300 | 0.238319794 | 0.315127094 | 0.132551138 | 0.232332075 |
| 188,400 | 0.238332754 | 0.315709377 | 0.132298812 | 0.232545852 |
| 188,500 | 0.23838907 | 0.315280499 | 0.133262546 | 0.23196485 |
| 188,600 | 0.238365521 | 0.314384912 | 0.13216776 | 0.232488799 |
| 188,700 | 0.238063924 | 0.314473491 | 0.130768445 | 0.232673696 |
| 188,800 | 0.238036167 | 0.313650933 | 0.131912005 | 0.232378166 |
| 188,900 | 0.237945486 | 0.313121332 | 0.131406333 | 0.232049855 |
| 189,000 | 0.237787821 | 0.313131374 | 0.130299737 | 0.231842115 |
| 189,100 | 0.237951897 | 0.31356609 | 0.129947669 | 0.233101496 |
| 189,200 | 0.237868524 | 0.314987982 | 0.129599914 | 0.232177636 |
| 189,300 | 0.237883394 | 0.314007896 | 0.130985544 | 0.232450478 |
| 189,400 | 0.237799686 | 0.314410984 | 0.128695112 | 0.231001088 |
| 189,500 | 0.23784938 | 0.3148393 | 0.130715419 | 0.231649972 |
| 189,600 | 0.237644434 | 0.314115814 | 0.12986968 | 0.232394392 |
| 189,700 | 0.237639427 | 0.31442025 | 0.130267571 | 0.230615514 |
| 189,800 | 0.237592178 | 0.313754709 | 0.130982617 | 0.232079231 |
| 189,900 | 0.23747883 | 0.313826836 | 0.130327205 | 0.232678869 |
| 190,000 | 0.237569258 | 0.314004578 | 0.130011364 | 0.232339345 |
| 190,100 | 0.237498072 | 0.313877054 | 0.130327048 | 0.232090383 |
| 190,200 | 0.237394323 | 0.314174724 | 0.129598853 | 0.232259338 |
| 190,300 | 0.237488667 | 0.31414504 | 0.130699594 | 0.231977246 |
| 190,400 | 0.237303428 | 0.313550812 | 0.129589911 | 0.231685896 |
| 190,500 | 0.237239884 | 0.31328455 | 0.130318579 | 0.231385379 |
| 190,600 | 0.23730575 | 0.313093957 | 0.129981528 | 0.231619266 |
| 190,700 | 0.23720578 | 0.313483041 | 0.12985115 | 0.230589977 |
| 190,800 | 0.237369696 | 0.313669747 | 0.128579909 | 0.230445331 |
| 190,900 | 0.237019665 | 0.313141665 | 0.130085378 | 0.230266794 |
| 191,000 | 0.236957688 | 0.313340721 | 0.12990219 | 0.231174565 |
| 191,100 | 0.237087929 | 0.312289538 | 0.130119682 | 0.230530652 |
| 191,200 | 0.237002635 | 0.312402312 | 0.129756226 | 0.230140667 |
| 191,300 | 0.236982155 | 0.313124414 | 0.128695904 | 0.230940394 |
| 191,400 | 0.237018872 | 0.311918414 | 0.130067211 | 0.231089187 |
| 191,500 | 0.236943071 | 0.312884384 | 0.129065922 | 0.230798227 |
| 191,600 | 0.236842948 | 0.313167016 | 0.129142948 | 0.229738229 |
| 191,700 | 0.236894399 | 0.312430132 | 0.129590249 | 0.229966658 |
| 191,800 | 0.236786745 | 0.313045485 | 0.129569907 | 0.230529506 |
| 191,900 | 0.236806286 | 0.312409425 | 0.129164211 | 0.230504024 |
| 192,000 | 0.236787622 | 0.312208788 | 0.13050796 | 0.230368848 |
| 192,100 | 0.236567918 | 0.311614403 | 0.129235504 | 0.230096698 |
| 192,200 | 0.236519902 | 0.311918653 | 0.129976913 | 0.22984299 |
| 192,300 | 0.236541422 | 0.31262397 | 0.131247753 | 0.229739046 |
| 192,400 | 0.236450767 | 0.312353684 | 0.130086701 | 0.229763813 |
| 192,500 | 0.236417559 | 0.312888761 | 0.130904404 | 0.230460957 |
| 192,600 | 0.236390528 | 0.312895537 | 0.12956852 | 0.228823154 |
| 192,700 | 0.23628128 | 0.312175305 | 0.129154215 | 0.229572989 |
| 192,800 | 0.236289337 | 0.312532052 | 0.129196139 | 0.228885996 |
| 192,900 | 0.236263532 | 0.312071898 | 0.127359405 | 0.229213259 |
| 193,000 | 0.236345513 | 0.31200209 | 0.128586719 | 0.229455771 |
| 193,100 | 0.2361265 | 0.313090916 | 0.130017115 | 0.228907175 |
| 193,200 | 0.236368768 | 0.313559147 | 0.129308375 | 0.229769735 |

| | | | | |
|---|---|---|---|---|
| 193,300 | 0.236357713 | 0.312953777 | 0.128995775 | 0.229944711 |
| 193,400 | 0.236121923 | 0.312981872 | 0.128713984 | 0.22874581 |
| 193,500 | 0.236143309 | 0.312531065 | 0.129651022 | 0.228771782 |
| 193,600 | 0.236014891 | 0.313510107 | 0.128407468 | 0.229219133 |
| 193,700 | 0.235898661 | 0.313360661 | 0.128321647 | 0.228094353 |
| 193,800 | 0.23595152 | 0.313605006 | 0.127634783 | 0.228539181 |
| 193,900 | 0.235942057 | 0.313127 | 0.129036407 | 0.228477409 |
| 194,000 | 0.235924469 | 0.312384455 | 0.128584521 | 0.228681347 |
| 194,100 | 0.235861554 | 0.312480797 | 0.128308153 | 0.228552585 |
| 194,200 | 0.235648492 | 0.31213331 | 0.127041858 | 0.227689781 |
| 194,300 | 0.235854345 | 0.312634662 | 0.127786436 | 0.22860845 |
| 194,400 | 0.235813088 | 0.31293892 | 0.127053518 | 0.22805466 |
| 194,500 | 0.235762181 | 0.312696834 | 0.126369467 | 0.227889302 |
| 194,600 | 0.235550628 | 0.312994212 | 0.127144317 | 0.227954642 |
| 194,700 | 0.23543098 | 0.313129896 | 0.127769326 | 0.228256577 |
| 194,800 | 0.235520107 | 0.313582369 | 0.127831264 | 0.22728995 |
| 194,900 | 0.235501223 | 0.313283083 | 0.127213081 | 0.227807207 |
| 195,000 | 0.23554003 | 0.312880456 | 0.126462528 | 0.227314258 |
| 195,100 | 0.235390791 | 0.313033758 | 0.125707463 | 0.22746957 |
| 195,200 | 0.235330669 | 0.313052091 | 0.127194427 | 0.227690333 |
| 195,300 | 0.235200353 | 0.313611283 | 0.126943324 | 0.227520115 |
| 195,400 | 0.235131516 | 0.31245885 | 0.126170416 | 0.227156478 |
| 195,500 | 0.235168432 | 0.312802207 | 0.126567852 | 0.227638914 |
| 195,600 | 0.235211195 | 0.31359634 | 0.127122687 | 0.22747287 |
| 195,700 | 0.235079594 | 0.312588665 | 0.126820967 | 0.227097405 |
| 195,800 | 0.235142783 | 0.311888278 | 0.12721133 | 0.226990413 |
| 195,900 | 0.234850778 | 0.311468931 | 0.127407213 | 0.227197901 |
| 196,000 | 0.23490211 | 0.311787822 | 0.126971154 | 0.227563461 |
| 196,100 | 0.234852177 | 0.311582596 | 0.126803353 | 0.226719843 |
| 196,200 | 0.234872286 | 0.311561013 | 0.126428554 | 0.226642681 |
| 196,300 | 0.234855517 | 0.311267207 | 0.127324923 | 0.226812541 |
| 196,400 | 0.234722316 | 0.311336924 | 0.126922583 | 0.227504673 |
| 196,500 | 0.234532849 | 0.311071101 | 0.1257171 | 0.227050185 |
| 196,600 | 0.234474587 | 0.312470593 | 0.126566606 | 0.22639619 |
| 196,700 | 0.234599887 | 0.311624444 | 0.124479815 | 0.227068137 |
| 196,800 | 0.234402801 | 0.311707279 | 0.125463782 | 0.226812116 |
| 196,900 | 0.234442251 | 0.312663982 | 0.124124362 | 0.227702739 |
| 197,000 | 0.234358585 | 0.311671414 | 0.125008326 | 0.227578456 |
| 197,100 | 0.234326321 | 0.31100044 | 0.125204579 | 0.226947745 |
| 197,200 | 0.234375054 | 0.31060546 | 0.126072174 | 0.22603967 |
| 197,300 | 0.234330627 | 0.311130836 | 0.124612798 | 0.226427902 |
| 197,400 | 0.234332258 | 0.31169222 | 0.125410789 | 0.226648672 |
| 197,500 | 0.234343618 | 0.311849349 | 0.125532581 | 0.227381504 |
| 197,600 | 0.234268105 | 0.31121371 | 0.125019885 | 0.226851866 |
| 197,700 | 0.234066574 | 0.311765235 | 0.126971765 | 0.227230392 |
| 197,800 | 0.233961394 | 0.311428186 | 0.125012982 | 0.226363858 |
| 197,900 | 0.23404484 | 0.311753098 | 0.124615102 | 0.226859079 |
| 198,000 | 0.23393626 | 0.312602167 | 0.123769971 | 0.226879557 |
| 198,100 | 0.233960537 | 0.310881964 | 0.124623736 | 0.22641744 |
| 198,200 | 0.233959838 | 0.311606355 | 0.123967822 | 0.225721129 |
| 198,300 | 0.233883923 | 0.312061621 | 0.122947199 | 0.226033875 |
| 198,400 | 0.233890448 | 0.312312675 | 0.122579355 | 0.225535448 |
| 198,500 | 0.233975632 | 0.31157706 | 0.122188302 | 0.22658632 |
| 198,600 | 0.233777935 | 0.312121784 | 0.12298149 | 0.227682864 |
| 198,700 | 0.233819881 | 0.312831308 | 0.123997452 | 0.226141998 |
| 198,800 | 0.233704004 | 0.312212282 | 0.122611388 | 0.225851234 |
| 198,900 | 0.233774271 | 0.311752641 | 0.122727235 | 0.227339671 |
| 199,000 | 0.233612002 | 0.312298573 | 0.123584178 | 0.226317504 |
| 199,100 | 0.233624237 | 0.311796454 | 0.123602521 | 0.22639991 |
| 199,200 | 0.233626621 | 0.311551945 | 0.123321025 | 0.22590805 |
| 199,300 | 0.233523982 | 0.31118083 | 0.122086441 | 0.226505603 |
| 199,400 | 0.23343413 | 0.311237738 | 0.122276709 | 0.226824877 |
| 199,500 | 0.233457103 | 0.310435472 | 0.122803426 | 0.227205388 |
| 199,600 | 0.233383552 | 0.309995554 | 0.122302769 | 0.227330565 |
| 199,700 | 0.233318849 | 0.310698139 | 0.122276847 | 0.227234635 |
| 199,800 | 0.233118861 | 0.309892068 | 0.123223524 | 0.227410683 |
| 199,900 | 0.233162806 | 0.310420548 | 0.122882891 | 0.227032255 |

| | | | | |
|---|---|---|---|---|
| 200,000 | 0.233062711 | 0.310687721 | 0.121738618 | 0.2277551 |
| 200,100 | 0.233115963 | 0.31115554 | 0.122460124 | 0.227092311 |
| 200,200 | 0.233156092 | 0.310954895 | 0.122328307 | 0.2274078 |
| 200,300 | 0.233102364 | 0.310872024 | 0.121185311 | 0.2277997 |
| 200,400 | 0.233020998 | 0.31119391 | 0.121363159 | 0.226856278 |
| 200,500 | 0.233005218 | 0.310948705 | 0.120751591 | 0.227740308 |
| 200,600 | 0.232884545 | 0.310160429 | 0.122048793 | 0.226920646 |
| 200,700 | 0.232734374 | 0.310227594 | 0.120538791 | 0.227438845 |
| 200,800 | 0.232843042 | 0.310191916 | 0.121029486 | 0.227145791 |
| 200,900 | 0.232717886 | 0.310127365 | 0.121552352 | 0.227208904 |
| 201,000 | 0.232627566 | 0.309137075 | 0.121682323 | 0.227372593 |
| 201,100 | 0.232478157 | 0.310421008 | 0.121830431 | 0.22661228 |
| 201,200 | 0.232434649 | 0.309291231 | 0.121709901 | 0.2264012 |
| 201,300 | 0.232563311 | 0.309876396 | 0.121728466 | 0.226889962 |
| 201,400 | 0.232537346 | 0.310821402 | 0.121761884 | 0.226398725 |
| 201,500 | 0.232561656 | 0.311049955 | 0.120721968 | 0.227200069 |
| 201,600 | 0.232549912 | 0.310597733 | 0.121868868 | 0.226469286 |
| 201,700 | 0.23241708 | 0.3110948 | 0.122357443 | 0.226730272 |
| 201,800 | 0.232549548 | 0.310599117 | 0.122359915 | 0.22640709 |
| 201,900 | 0.232318961 | 0.310355467 | 0.121224465 | 0.227188624 |
| 202,000 | 0.232256988 | 0.309892856 | 0.122633762 | 0.22784882 |
| 202,100 | 0.232280722 | 0.309209512 | 0.121508687 | 0.227303812 |
| 202,200 | 0.232219671 | 0.30877527 | 0.121274307 | 0.227423085 |
| 202,300 | 0.23217299 | 0.308608527 | 0.122368826 | 0.227678467 |
| 202,400 | 0.232245594 | 0.308988464 | 0.123462555 | 0.227663672 |
| 202,500 | 0.232154414 | 0.308233849 | 0.123191661 | 0.226599336 |
| 202,600 | 0.232072436 | 0.308933663 | 0.121891841 | 0.226943187 |
| 202,700 | 0.231808226 | 0.308499061 | 0.123409123 | 0.226779363 |
| 202,800 | 0.231761408 | 0.307960369 | 0.124177261 | 0.226466837 |
| 202,900 | 0.231859257 | 0.308833229 | 0.123383024 | 0.226789522 |
| 203,000 | 0.231803608 | 0.308010864 | 0.122411178 | 0.226998152 |
| 203,100 | 0.23173283 | 0.3081568 | 0.122070171 | 0.227147678 |
| 203,200 | 0.231810491 | 0.307860521 | 0.123498647 | 0.227327217 |
| 203,300 | 0.231607325 | 0.308201107 | 0.1218854 | 0.226550382 |
| 203,400 | 0.231674575 | 0.308099051 | 0.12238744 | 0.226817057 |
| 203,500 | 0.231814128 | 0.307850046 | 0.122963598 | 0.226714424 |
| 203,600 | 0.231672319 | 0.307834372 | 0.122550537 | 0.227956386 |
| 203,700 | 0.231778293 | 0.308209344 | 0.122433709 | 0.227183449 |
| 203,800 | 0.231470257 | 0.307263429 | 0.122495114 | 0.226343094 |
| 203,900 | 0.231431184 | 0.30737524 | 0.121651569 | 0.226625626 |
| 204,000 | 0.231266677 | 0.30657098 | 0.120458823 | 0.226934645 |
| 204,100 | 0.231334169 | 0.30728868 | 0.122912784 | 0.226851614 |
| 204,200 | 0.231308244 | 0.307257981 | 0.122003932 | 0.226400924 |
| 204,300 | 0.231239864 | 0.306693259 | 0.122898336 | 0.226326914 |
| 204,400 | 0.231422338 | 0.307533559 | 0.123367138 | 0.226995161 |
| 204,500 | 0.231328206 | 0.307154571 | 0.122318188 | 0.226152259 |
| 204,600 | 0.231331834 | 0.307236041 | 0.122353628 | 0.226597409 |
| 204,700 | 0.231051365 | 0.307671805 | 0.122081808 | 0.227388648 |
| 204,800 | 0.231235232 | 0.307993551 | 0.122547857 | 0.226738328 |
| 204,900 | 0.231087927 | 0.307693293 | 0.121176212 | 0.226334345 |
| 205,000 | 0.231028465 | 0.307931519 | 0.121211802 | 0.227075557 |
| 205,100 | 0.230980876 | 0.307964424 | 0.120942399 | 0.227269948 |
| 205,200 | 0.230830729 | 0.307846339 | 0.121668453 | 0.226204579 |
| 205,300 | 0.230961027 | 0.306851074 | 0.122087426 | 0.22643603 |
| 205,400 | 0.230785908 | 0.307065423 | 0.120812853 | 0.22636518 |
| 205,500 | 0.230917751 | 0.306482413 | 0.120585069 | 0.226134476 |
| 205,600 | 0.230720864 | 0.307331328 | 0.12207598 | 0.22648458 |
| 205,700 | 0.230645461 | 0.307909169 | 0.119936989 | 0.226527836 |
| 205,800 | 0.230473604 | 0.306363649 | 0.120031393 | 0.225704678 |
| 205,900 | 0.230656209 | 0.30717687 | 0.121182647 | 0.226062272 |
| 206,000 | 0.230699084 | 0.306210168 | 0.119357263 | 0.226081805 |
| 206,100 | 0.230462683 | 0.30598233 | 0.120606516 | 0.225836564 |
| 206,200 | 0.230532407 | 0.306574294 | 0.119922001 | 0.225334514 |
| 206,300 | 0.230523968 | 0.30723804 | 0.120327048 | 0.225222021 |
| 206,400 | 0.230600746 | 0.306311262 | 0.120000168 | 0.225803195 |
| 206,500 | 0.230461779 | 0.306782221 | 0.119238388 | 0.224858507 |
| 206,600 | 0.230344013 | 0.305902475 | 0.119130675 | 0.225175973 |

| | | | | |
|---|---|---|---|---|
| 206,700 | 0.230225338 | 0.306954308 | 0.120172689 | 0.225184053 |
| 206,800 | 0.230097767 | 0.306535568 | 0.120341562 | 0.225353253 |
| 206,900 | 0.230237949 | 0.306224743 | 0.119833625 | 0.225536876 |
| 207,000 | 0.230125969 | 0.306401903 | 0.119986847 | 0.225172209 |
| 207,100 | 0.23016207 | 0.30690677 | 0.11962238 | 0.224788827 |
| 207,200 | 0.230030571 | 0.307002193 | 0.120725413 | 0.225163232 |
| 207,300 | 0.229867284 | 0.306310199 | 0.120127298 | 0.224859455 |
| 207,400 | 0.230049054 | 0.307325219 | 0.119801638 | 0.224323066 |
| 207,500 | 0.230034474 | 0.306466288 | 0.119445192 | 0.223807458 |
| 207,600 | 0.230056717 | 0.306801147 | 0.120532031 | 0.224513632 |
| 207,700 | 0.229978974 | 0.306396718 | 0.120977729 | 0.224864403 |
| 207,800 | 0.230081255 | 0.306702275 | 0.119885399 | 0.224169291 |
| 207,900 | 0.229973096 | 0.306771837 | 0.118537448 | 0.224365669 |
| 208,000 | 0.229983471 | 0.306872253 | 0.118714565 | 0.224214564 |
| 208,100 | 0.229908926 | 0.306537644 | 0.117696425 | 0.224925311 |
| 208,200 | 0.230045862 | 0.306886826 | 0.117417179 | 0.224726664 |
| 208,300 | 0.229840125 | 0.306269746 | 0.118006934 | 0.223636046 |
| 208,400 | 0.230039794 | 0.30704911 | 0.119775633 | 0.224333582 |
| 208,500 | 0.229700943 | 0.305665368 | 0.117640179 | 0.224968143 |
| 208,600 | 0.229773681 | 0.305680463 | 0.11704661 | 0.224816246 |
| 208,700 | 0.229721181 | 0.305663465 | 0.118212795 | 0.225949872 |
| 208,800 | 0.229698732 | 0.306254268 | 0.118196311 | 0.225106096 |
| 208,900 | 0.229494452 | 0.306378066 | 0.11853744 | 0.224613249 |
| 209,000 | 0.229507203 | 0.306391773 | 0.118635065 | 0.224177004 |
| 209,100 | 0.229429516 | 0.305441992 | 0.118229269 | 0.224227475 |
| 209,200 | 0.229644318 | 0.306119718 | 0.117597088 | 0.225506972 |
| 209,300 | 0.229380308 | 0.305495502 | 0.117634643 | 0.225045381 |
| 209,400 | 0.229554182 | 0.306110449 | 0.117173511 | 0.22474336 |
| 209,500 | 0.22961572 | 0.30677252 | 0.117884758 | 0.224611708 |
| 209,600 | 0.229378248 | 0.305999461 | 0.118666581 | 0.225227155 |
| 209,700 | 0.229437708 | 0.306420685 | 0.118138225 | 0.224500292 |
| 209,800 | 0.229442829 | 0.306254822 | 0.11847372 | 0.224463036 |
| 209,900 | 0.229263856 | 0.306202973 | 0.11882603 | 0.224332709 |
| 210,000 | 0.22924849 | 0.305916791 | 0.118461423 | 0.224300072 |
| 210,100 | 0.229047016 | 0.305655702 | 0.117624513 | 0.223518366 |
| 210,200 | 0.229228097 | 0.305343488 | 0.117804279 | 0.224329826 |
| 210,300 | 0.229154011 | 0.305503581 | 0.117856654 | 0.224392398 |
| 210,400 | 0.228919236 | 0.305843794 | 0.117482876 | 0.223609104 |
| 210,500 | 0.229029388 | 0.306290926 | 0.117960128 | 0.223427092 |
| 210,600 | 0.228830244 | 0.305009698 | 0.117655518 | 0.223658728 |
| 210,700 | 0.228877016 | 0.305928001 | 0.117429028 | 0.223515441 |
| 210,800 | 0.228947333 | 0.305765475 | 0.117150689 | 0.222825841 |
| 210,900 | 0.228761379 | 0.306050083 | 0.116533918 | 0.223787755 |
| 211,000 | 0.22869151 | 0.306064186 | 0.118681001 | 0.223516992 |
| 211,100 | 0.2287592 | 0.30591411 | 0.117017645 | 0.223775319 |
| 211,200 | 0.228699659 | 0.305746651 | 0.117789963 | 0.222997732 |
| 211,300 | 0.228685337 | 0.306619398 | 0.117466099 | 0.222868363 |
| 211,400 | 0.22859207 | 0.306956779 | 0.118949545 | 0.22313087 |
| 211,500 | 0.228490608 | 0.306712248 | 0.118738967 | 0.224184417 |
| 211,600 | 0.228420121 | 0.305013039 | 0.119515563 | 0.222359702 |
| 211,700 | 0.228519132 | 0.306032737 | 0.118447068 | 0.223093631 |
| 211,800 | 0.228410067 | 0.305995269 | 0.120041567 | 0.223299202 |
| 211,900 | 0.228334272 | 0.305896847 | 0.120264617 | 0.223700829 |
| 212,000 | 0.228315716 | 0.305943873 | 0.119252333 | 0.222630664 |
| 212,100 | 0.228239726 | 0.305407347 | 0.11821329 | 0.222621904 |
| 212,200 | 0.228058929 | 0.30481232 | 0.119358443 | 0.223057758 |
| 212,300 | 0.228006936 | 0.30441584 | 0.118381683 | 0.224085719 |
| 212,400 | 0.228084651 | 0.30504563 | 0.117862307 | 0.223010392 |
| 212,500 | 0.227876525 | 0.305101714 | 0.119142483 | 0.223543074 |
| 212,600 | 0.227841275 | 0.304773984 | 0.12000872 | 0.222864667 |
| 212,700 | 0.227925996 | 0.305302423 | 0.119231632 | 0.223367671 |
| 212,800 | 0.227828447 | 0.30447743 | 0.117828344 | 0.223911055 |
| 212,900 | 0.227971594 | 0.305105286 | 0.119024394 | 0.222790671 |
| 213,000 | 0.227861443 | 0.30556824 | 0.117405051 | 0.222727501 |
| 213,100 | 0.227586862 | 0.304565637 | 0.119166586 | 0.223011458 |
| 213,200 | 0.227513147 | 0.303758277 | 0.117218892 | 0.222517594 |
| 213,300 | 0.227379519 | 0.304472452 | 0.11781429 | 0.221491158 |

| | | | | |
|---|---|---|---|---|
| 213,400 | 0.227452593 | 0.303806661 | 0.117513053 | 0.221868416 |
| 213,500 | 0.227611611 | 0.304189973 | 0.117325822 | 0.221974593 |
| 213,600 | 0.227414407 | 0.303817939 | 0.118158432 | 0.221641087 |
| 213,700 | 0.227517114 | 0.304108637 | 0.118319061 | 0.222154859 |
| 213,800 | 0.22738803 | 0.303489709 | 0.119018178 | 0.221780957 |
| 213,900 | 0.227227926 | 0.303442624 | 0.117807255 | 0.221495602 |
| 214,000 | 0.227020198 | 0.303220238 | 0.118647356 | 0.221001447 |
| 214,100 | 0.227163321 | 0.302663953 | 0.117215531 | 0.221374905 |
| 214,200 | 0.227137943 | 0.303220205 | 0.11914107 | 0.22120925 |
| 214,300 | 0.227081091 | 0.303386629 | 0.117595458 | 0.220625125 |
| 214,400 | 0.226920336 | 0.302479193 | 0.117616156 | 0.222115955 |
| 214,500 | 0.226931095 | 0.303184288 | 0.117944136 | 0.221579285 |
| 214,600 | 0.226821748 | 0.302763993 | 0.118505286 | 0.221389655 |
| 214,700 | 0.226632569 | 0.301627962 | 0.117211298 | 0.220695722 |
| 214,800 | 0.226684094 | 0.30291783 | 0.116917179 | 0.22130868 |
| 214,900 | 0.226747923 | 0.303151753 | 0.117558547 | 0.220326939 |
| 215,000 | 0.226636823 | 0.303008571 | 0.116953258 | 0.220736286 |
| 215,100 | 0.226690189 | 0.303456626 | 0.118418882 | 0.220827161 |
| 215,200 | 0.226645067 | 0.302455618 | 0.118195864 | 0.221105069 |
| 215,300 | 0.226681436 | 0.302145129 | 0.117868782 | 0.221220623 |
| 215,400 | 0.226411747 | 0.303388645 | 0.116176916 | 0.221269833 |
| 215,500 | 0.226466568 | 0.302851184 | 0.117962105 | 0.22142405 |
| 215,600 | 0.226209809 | 0.301980692 | 0.117711575 | 0.220791572 |
| 215,700 | 0.226461191 | 0.302656633 | 0.115320638 | 0.220365884 |
| 215,800 | 0.226339723 | 0.302965656 | 0.116336776 | 0.220186704 |
| 215,900 | 0.22630137 | 0.30357461 | 0.11719605 | 0.220156478 |
| 216,000 | 0.2262701 | 0.30319407 | 0.116252291 | 0.220905348 |
| 216,100 | 0.226200394 | 0.30299729 | 0.11804937 | 0.220573153 |
| 216,200 | 0.226184188 | 0.304239625 | 0.117152382 | 0.220459899 |
| 216,300 | 0.226101784 | 0.304044938 | 0.116692187 | 0.219783607 |
| 216,400 | 0.226083459 | 0.30348697 | 0.116016523 | 0.220416212 |
| 216,500 | 0.226117018 | 0.303668093 | 0.116508468 | 0.220871961 |
| 216,600 | 0.226172028 | 0.303754647 | 0.117892084 | 0.220026535 |
| 216,700 | 0.226159978 | 0.303873037 | 0.116675693 | 0.220388804 |
| 216,800 | 0.226064257 | 0.303925091 | 0.116419605 | 0.220265311 |
| 216,900 | 0.225962521 | 0.304159853 | 0.116617984 | 0.220285657 |
| 217,000 | 0.226070427 | 0.304094365 | 0.116349003 | 0.219971451 |
| 217,100 | 0.225680384 | 0.303282908 | 0.115541856 | 0.219511494 |
| 217,200 | 0.225627503 | 0.301803139 | 0.116740851 | 0.219764708 |
| 217,300 | 0.22579379 | 0.303076038 | 0.115723143 | 0.219720978 |
| 217,400 | 0.225613793 | 0.303362757 | 0.116127453 | 0.220643959 |
| 217,500 | 0.225686392 | 0.303701606 | 0.115171588 | 0.219959293 |
| 217,600 | 0.225828985 | 0.303218539 | 0.114299327 | 0.220499466 |
| 217,700 | 0.225688817 | 0.303341404 | 0.11479812 | 0.219257565 |
| 217,800 | 0.22573078 | 0.302945439 | 0.115309693 | 0.22036886 |
| 217,900 | 0.225588194 | 0.303762785 | 0.115214372 | 0.220232002 |
| 218,000 | 0.22559725 | 0.302981542 | 0.116106328 | 0.220344226 |
| 218,100 | 0.22576329 | 0.302811275 | 0.115575456 | 0.220972877 |
| 218,200 | 0.225444306 | 0.303155226 | 0.114505394 | 0.220019583 |
| 218,300 | 0.225479631 | 0.303042244 | 0.114839899 | 0.220136125 |
| 218,400 | 0.225460076 | 0.303227418 | 0.115434689 | 0.219776871 |
| 218,500 | 0.225250162 | 0.303463015 | 0.115430325 | 0.219726868 |
| 218,600 | 0.225291258 | 0.303703323 | 0.116311702 | 0.220386726 |
| 218,700 | 0.225117754 | 0.302376782 | 0.115738805 | 0.220194646 |
| 218,800 | 0.225367598 | 0.303402012 | 0.115673379 | 0.220454067 |
| 218,900 | 0.225404398 | 0.303722438 | 0.115144931 | 0.221347245 |
| 219,000 | 0.225259832 | 0.303183087 | 0.114364682 | 0.22042868 |
| 219,100 | 0.225139236 | 0.302473798 | 0.114385747 | 0.219890378 |
| 219,200 | 0.225175895 | 0.303002232 | 0.114976784 | 0.219945793 |
| 219,300 | 0.22486925 | 0.3030324 | 0.114829157 | 0.219684343 |
| 219,400 | 0.224975563 | 0.303474892 | 0.113855934 | 0.220173807 |
| 219,500 | 0.224836084 | 0.303051526 | 0.115351185 | 0.220288119 |
| 219,600 | 0.224916328 | 0.303736269 | 0.115494935 | 0.220032444 |
| 219,700 | 0.224923149 | 0.303788284 | 0.114649665 | 0.220120383 |
| 219,800 | 0.224795539 | 0.303073081 | 0.115096088 | 0.220342965 |
| 219,900 | 0.224825455 | 0.302980476 | 0.115676066 | 0.219685501 |
| 220,000 | 0.224775736 | 0.30278141 | 0.11583236 | 0.220199452 |

| | | | | |
|---|---|---|---|---|
| 220,100 | 0.224666258 | 0.303011552 | 0.115319029 | 0.219828827 |
| 220,200 | 0.224629256 | 0.304269721 | 0.115415819 | 0.219817877 |
| 220,300 | 0.224395621 | 0.303759408 | 0.114491123 | 0.218705363 |
| 220,400 | 0.2244592 | 0.303603913 | 0.115022025 | 0.220009388 |
| 220,500 | 0.224483428 | 0.303049993 | 0.114741386 | 0.218998011 |
| 220,600 | 0.224396628 | 0.303172926 | 0.114199589 | 0.219561432 |
| 220,700 | 0.224506684 | 0.303023163 | 0.114205645 | 0.219114317 |
| 220,800 | 0.224508198 | 0.303152315 | 0.114633396 | 0.219703223 |
| 220,900 | 0.224473769 | 0.302952228 | 0.113470607 | 0.219690123 |
| 221,000 | 0.224368394 | 0.302578786 | 0.113308267 | 0.219706406 |
| 221,100 | 0.224084713 | 0.302473932 | 0.113571574 | 0.219207302 |
| 221,200 | 0.22436437 | 0.303472721 | 0.113137573 | 0.219586267 |
| 221,300 | 0.224089428 | 0.302920666 | 0.113387454 | 0.219064174 |
| 221,400 | 0.224262733 | 0.302971269 | 0.1134021 | 0.219226775 |
| 221,500 | 0.224197668 | 0.302424567 | 0.113482601 | 0.218357421 |
| 221,600 | 0.224090411 | 0.302392555 | 0.113389683 | 0.218587737 |
| 221,700 | 0.224018672 | 0.301833872 | 0.113509657 | 0.218231313 |
| 221,800 | 0.224153732 | 0.302241583 | 0.114011334 | 0.217807767 |
| 221,900 | 0.224104776 | 0.301359832 | 0.113475531 | 0.218526791 |
| 222,000 | 0.224136728 | 0.301681124 | 0.113202428 | 0.217671653 |
| 222,100 | 0.224266055 | 0.302065594 | 0.114231221 | 0.218259863 |
| 222,200 | 0.224055817 | 0.302411387 | 0.114312006 | 0.217921097 |
| 222,300 | 0.223865241 | 0.301703591 | 0.111929731 | 0.217844897 |
| 222,400 | 0.224032259 | 0.301548884 | 0.11291915 | 0.2178779 |
| 222,500 | 0.223790857 | 0.300857455 | 0.113291498 | 0.217648446 |
| 222,600 | 0.223902006 | 0.300814783 | 0.113195829 | 0.218187509 |
| 222,700 | 0.223772331 | 0.30104716 | 0.11231577 | 0.218191051 |
| 222,800 | 0.223787956 | 0.300957035 | 0.11337509 | 0.218197464 |
| 222,900 | 0.223773229 | 0.300460188 | 0.114306139 | 0.218128918 |
| 223,000 | 0.223610758 | 0.301087042 | 0.112021497 | 0.218048261 |
| 223,100 | 0.223491153 | 0.300266128 | 0.112839102 | 0.217751677 |
| 223,200 | 0.22371147 | 0.301154057 | 0.113588407 | 0.218560921 |
| 223,300 | 0.223587737 | 0.299191655 | 0.112641719 | 0.218703743 |
| 223,400 | 0.223564494 | 0.300128461 | 0.114555439 | 0.217982784 |
| 223,500 | 0.223423468 | 0.299532351 | 0.11449815 | 0.217902901 |
| 223,600 | 0.22327186 | 0.300483336 | 0.114930249 | 0.217317672 |
| 223,700 | 0.223418882 | 0.299704424 | 0.114196651 | 0.218028476 |
| 223,800 | 0.223369719 | 0.29995686 | 0.112920621 | 0.217459223 |
| 223,900 | 0.223333336 | 0.299608879 | 0.11353589 | 0.218280603 |
| 224,000 | 0.223197248 | 0.299782722 | 0.113713168 | 0.218401927 |
| 224,100 | 0.223103903 | 0.300126599 | 0.113144865 | 0.218600754 |
| 224,200 | 0.222961529 | 0.299301451 | 0.112931905 | 0.217626919 |
| 224,300 | 0.223062107 | 0.299585761 | 0.112565928 | 0.217465456 |
| 224,400 | 0.222864478 | 0.299116614 | 0.112738412 | 0.217243692 |
| 224,500 | 0.223071476 | 0.299160144 | 0.112709385 | 0.217274282 |
| 224,600 | 0.222978105 | 0.299181693 | 0.112343027 | 0.21727805 |
| 224,700 | 0.222920566 | 0.299570512 | 0.111785627 | 0.217490664 |
| 224,800 | 0.222963177 | 0.300899904 | 0.112981331 | 0.217531189 |
| 224,900 | 0.222731104 | 0.299757844 | 0.113669142 | 0.21784069 |
| 225,000 | 0.22285628 | 0.299432623 | 0.1132147 | 0.217691067 |
| 225,100 | 0.222810511 | 0.298694181 | 0.113937108 | 0.218349515 |
| 225,200 | 0.222764823 | 0.298880882 | 0.112727258 | 0.217931971 |
| 225,300 | 0.222587729 | 0.29866653 | 0.113957225 | 0.217942681 |
| 225,400 | 0.222845409 | 0.298634991 | 0.113346587 | 0.217537745 |
| 225,500 | 0.222691061 | 0.298549927 | 0.112437011 | 0.21714886 |
| 225,600 | 0.222787789 | 0.29920833 | 0.112219385 | 0.217555388 |
| 225,700 | 0.222783907 | 0.297734861 | 0.114069583 | 0.218282745 |
| 225,800 | 0.222599287 | 0.29695095 | 0.114173043 | 0.217990448 |
| 225,900 | 0.222548348 | 0.297303985 | 0.114183273 | 0.21745789 |
| 226,000 | 0.222659867 | 0.297578955 | 0.11334244 | 0.21743836 |
| 226,100 | 0.222304415 | 0.296799038 | 0.112499832 | 0.217194234 |
| 226,200 | 0.222518013 | 0.296952075 | 0.112940859 | 0.217880768 |
| 226,300 | 0.222257726 | 0.297919767 | 0.112093782 | 0.218204452 |
| 226,400 | 0.222258822 | 0.296889982 | 0.111724527 | 0.218032137 |
| 226,500 | 0.222106479 | 0.297011324 | 0.111789718 | 0.216863993 |
| 226,600 | 0.222133244 | 0.297948815 | 0.111377744 | 0.217066179 |
| 226,700 | 0.222059527 | 0.296601485 | 0.111822441 | 0.217259697 |

| | | | | |
|---|---|---|---|---|
| 226,800 | 0.222002963 | 0.297555369 | 0.111136364 | 0.216799849 |
| 226,900 | 0.222043806 | 0.297628132 | 0.111487247 | 0.216984032 |
| 227,000 | 0.221940296 | 0.296971115 | 0.11055122 | 0.217077981 |
| 227,100 | 0.221930788 | 0.296970196 | 0.111428972 | 0.216714745 |
| 227,200 | 0.221899053 | 0.297009472 | 0.112608508 | 0.21613514 |
| 227,300 | 0.221994199 | 0.296495598 | 0.1110567 | 0.216672557 |
| 227,400 | 0.221884809 | 0.29623306 | 0.112165805 | 0.216593132 |
| 227,500 | 0.221809238 | 0.296291845 | 0.113167347 | 0.216141632 |
| 227,600 | 0.221887163 | 0.296729954 | 0.112266405 | 0.216211369 |
| 227,700 | 0.221620255 | 0.295799232 | 0.11251689 | 0.216369231 |
| 227,800 | 0.221762054 | 0.296314809 | 0.112890908 | 0.21607685 |
| 227,900 | 0.22160367 | 0.296983826 | 0.11195477 | 0.21588252 |
| 228,000 | 0.22166478 | 0.295371095 | 0.112996282 | 0.215628048 |
| 228,100 | 0.22160664 | 0.295686093 | 0.111426082 | 0.215819838 |
| 228,200 | 0.221742054 | 0.296306175 | 0.112766171 | 0.216194707 |
| 228,300 | 0.221730172 | 0.297001128 | 0.112617203 | 0.216095227 |
| 228,400 | 0.221490642 | 0.29512582 | 0.112785198 | 0.215406406 |
| 228,500 | 0.221457906 | 0.296269372 | 0.112599129 | 0.215365321 |
| 228,600 | 0.221406109 | 0.294828607 | 0.11342326 | 0.215550894 |
| 228,700 | 0.221479379 | 0.29446472 | 0.1121804 | 0.215792199 |
| 228,800 | 0.221399504 | 0.295211558 | 0.112050014 | 0.216090169 |
| 228,900 | 0.221310706 | 0.295415649 | 0.111529005 | 0.216139514 |
| 229,000 | 0.221395982 | 0.295684729 | 0.112027956 | 0.216473832 |
| 229,100 | 0.221165264 | 0.295298509 | 0.113081347 | 0.215693465 |
| 229,200 | 0.221159935 | 0.296494853 | 0.113913631 | 0.216009496 |
| 229,300 | 0.221311415 | 0.296406109 | 0.112349699 | 0.215847363 |
| 229,400 | 0.221292078 | 0.296533435 | 0.112444614 | 0.21519546 |
| 229,500 | 0.221168235 | 0.295170384 | 0.112658552 | 0.21503883 |
| 229,600 | 0.22097862 | 0.296145879 | 0.112395679 | 0.21494306 |
| 229,700 | 0.220920166 | 0.295687745 | 0.11213473 | 0.214365647 |
| 229,800 | 0.221065118 | 0.296012558 | 0.111185822 | 0.214615805 |
| 229,900 | 0.220918841 | 0.295588735 | 0.111629611 | 0.214279802 |
| 230,000 | 0.220953667 | 0.296408624 | 0.110933892 | 0.214741133 |
| 230,100 | 0.220710863 | 0.295535185 | 0.111888298 | 0.214969905 |
| 230,200 | 0.220866991 | 0.295926872 | 0.110847578 | 0.21507953 |
| 230,300 | 0.220724434 | 0.296259308 | 0.111332914 | 0.215034416 |
| 230,400 | 0.220743848 | 0.296185099 | 0.110859473 | 0.214783154 |
| 230,500 | 0.220721911 | 0.296444595 | 0.110091542 | 0.214570008 |
| 230,600 | 0.220611128 | 0.295665309 | 0.110032517 | 0.214670895 |
| 230,700 | 0.220647522 | 0.295017809 | 0.109850927 | 0.214144396 |
| 230,800 | 0.22068171 | 0.2952385 | 0.109487272 | 0.214205559 |
| 230,900 | 0.220524836 | 0.295233636 | 0.11053457 | 0.213818856 |
| 231,000 | 0.220459413 | 0.295789773 | 0.110286217 | 0.214318636 |
| 231,100 | 0.220573582 | 0.295015884 | 0.108628422 | 0.214330518 |
| 231,200 | 0.220469491 | 0.294941918 | 0.109344942 | 0.213324106 |
| 231,300 | 0.22042888 | 0.295246215 | 0.109265366 | 0.213538548 |
| 231,400 | 0.220434797 | 0.295584588 | 0.108116499 | 0.213482897 |
| 231,500 | 0.220337034 | 0.295853617 | 0.1094274 | 0.212927636 |
| 231,600 | 0.220194394 | 0.295843823 | 0.10924822 | 0.21377278 |
| 231,700 | 0.22011602 | 0.295342063 | 0.109640009 | 0.213609157 |
| 231,800 | 0.22010143 | 0.295225792 | 0.108849358 | 0.212904294 |
| 231,900 | 0.220070345 | 0.295052037 | 0.10852209 | 0.212943427 |
| 232,000 | 0.220000368 | 0.294474801 | 0.108547142 | 0.212463789 |
| 232,100 | 0.220066465 | 0.295489704 | 0.109214441 | 0.212353295 |
| 232,200 | 0.220130798 | 0.294593597 | 0.108568478 | 0.21310757 |
| 232,300 | 0.220086019 | 0.295671429 | 0.108449328 | 0.21279445 |
| 232,400 | 0.219888997 | 0.295755051 | 0.109563882 | 0.212741825 |
| 232,500 | 0.219966697 | 0.295267438 | 0.110717566 | 0.212098685 |
| 232,600 | 0.22005126 | 0.295886884 | 0.111067185 | 0.212371782 |
| 232,700 | 0.219882591 | 0.296133178 | 0.110720027 | 0.211446899 |
| 232,800 | 0.219854006 | 0.296174627 | 0.110654579 | 0.211514435 |
| 232,900 | 0.219717171 | 0.296024333 | 0.110008166 | 0.211389608 |
| 233,000 | 0.219654926 | 0.296222287 | 0.10959693 | 0.210147014 |
| 233,100 | 0.219444706 | 0.295747602 | 0.109624314 | 0.210626237 |
| 233,200 | 0.219632404 | 0.295289236 | 0.110666767 | 0.210371351 |
| 233,300 | 0.219504904 | 0.295974412 | 0.110457257 | 0.210428406 |
| 233,400 | 0.219415902 | 0.295625416 | 0.108815498 | 0.210615818 |

| | | | | |
|---|---|---|---|---|
| 233,500 | 0.219522802 | 0.294416743 | 0.11013091 | 0.211405375 |
| 233,600 | 0.219520813 | 0.295646251 | 0.110708156 | 0.210791429 |
| 233,700 | 0.219313665 | 0.29519518 | 0.110112496 | 0.211233958 |
| 233,800 | 0.219470128 | 0.294474369 | 0.109260627 | 0.211343811 |
| 233,900 | 0.219190183 | 0.295415621 | 0.109596316 | 0.210995803 |
| 234,000 | 0.219352129 | 0.29549371 | 0.110519545 | 0.210515862 |
| 234,100 | 0.219079679 | 0.295893675 | 0.109570757 | 0.210529461 |
| 234,200 | 0.219085162 | 0.296131649 | 0.108160983 | 0.210453167 |
| 234,300 | 0.219214459 | 0.296323027 | 0.108459852 | 0.210447895 |
| 234,400 | 0.219163136 | 0.297138167 | 0.109038138 | 0.210420003 |
| 234,500 | 0.219086351 | 0.29709951 | 0.108849749 | 0.210384675 |
| 234,600 | 0.219069908 | 0.296813901 | 0.108450124 | 0.210540906 |
| 234,700 | 0.218943526 | 0.295906899 | 0.108227891 | 0.210887848 |
| 234,800 | 0.218989431 | 0.296041299 | 0.107640738 | 0.21129567 |
| 234,900 | 0.219014415 | 0.295436025 | 0.108287179 | 0.211040719 |
| 235,000 | 0.219041934 | 0.295999464 | 0.108707696 | 0.210744627 |
| 235,100 | 0.218869082 | 0.295894804 | 0.108837803 | 0.210665181 |
| 235,200 | 0.219029008 | 0.295838189 | 0.109275135 | 0.210371565 |
| 235,300 | 0.218822368 | 0.295570521 | 0.109557167 | 0.210059682 |
| 235,400 | 0.218788649 | 0.295394031 | 0.109443755 | 0.209636202 |
| 235,500 | 0.218992575 | 0.296036107 | 0.109509559 | 0.210266215 |
| 235,600 | 0.218950445 | 0.295954728 | 0.108748161 | 0.209737778 |
| 235,700 | 0.21885612 | 0.295841146 | 0.108700252 | 0.209323586 |
| 235,800 | 0.218649358 | 0.295333985 | 0.109497396 | 0.209977669 |
| 235,900 | 0.21881412 | 0.295979736 | 0.109015425 | 0.209830654 |
| 236,000 | 0.218689426 | 0.29644774 | 0.108781871 | 0.209610457 |
| 236,100 | 0.218624841 | 0.296091011 | 0.10934805 | 0.209300692 |
| 236,200 | 0.218582438 | 0.295657604 | 0.108887296 | 0.20996585 |
| 236,300 | 0.218540053 | 0.295497612 | 0.108279605 | 0.21093414 |
| 236,400 | 0.218479728 | 0.296091004 | 0.108230638 | 0.210641687 |
| 236,500 | 0.218542605 | 0.295892087 | 0.108819057 | 0.210321372 |
| 236,600 | 0.218371407 | 0.294276206 | 0.109452933 | 0.209706961 |
| 236,700 | 0.218329104 | 0.293691972 | 0.108122219 | 0.210840295 |
| 236,800 | 0.218244292 | 0.294116314 | 0.108468494 | 0.209553643 |
| 236,900 | 0.218225596 | 0.294163932 | 0.107998412 | 0.210101288 |
| 237,000 | 0.218117435 | 0.294021982 | 0.108273507 | 0.209604734 |
| 237,100 | 0.218144918 | 0.294475437 | 0.107853428 | 0.210444558 |
| 237,200 | 0.218162224 | 0.294381709 | 0.108725556 | 0.210713615 |
| 237,300 | 0.218045124 | 0.295472168 | 0.10920873 | 0.210740768 |
| 237,400 | 0.218110414 | 0.29473704 | 0.108663549 | 0.210655806 |
| 237,500 | 0.217932893 | 0.294611663 | 0.108229228 | 0.210523871 |
| 237,600 | 0.217956282 | 0.294038749 | 0.107031273 | 0.210207784 |
| 237,700 | 0.217958526 | 0.294462194 | 0.109235306 | 0.210584741 |
| 237,800 | 0.217841092 | 0.293675125 | 0.109253788 | 0.210794647 |
| 237,900 | 0.217781051 | 0.294357565 | 0.10858767 | 0.210859607 |
| 238,000 | 0.217591243 | 0.293784954 | 0.108634997 | 0.21057709 |
| 238,100 | 0.21800737 | 0.293849463 | 0.109842588 | 0.210885074 |
| 238,200 | 0.217849986 | 0.29393633 | 0.109799572 | 0.211082652 |
| 238,300 | 0.217809107 | 0.294556205 | 0.108894036 | 0.211405869 |
| 238,400 | 0.217705844 | 0.293594256 | 0.107688186 | 0.210808464 |
| 238,500 | 0.217754835 | 0.293127442 | 0.108792556 | 0.210559017 |
| 238,600 | 0.217727639 | 0.294250461 | 0.109020636 | 0.211536701 |
| 238,700 | 0.217644798 | 0.2934446 | 0.108272034 | 0.210415019 |
| 238,800 | 0.217646088 | 0.29338519 | 0.109182885 | 0.210333341 |
| 238,900 | 0.217583231 | 0.293938731 | 0.108859435 | 0.209315414 |
| 239,000 | 0.217761075 | 0.294236664 | 0.108844317 | 0.2095647 |
| 239,100 | 0.217602517 | 0.293586314 | 0.108902036 | 0.209486982 |
| 239,200 | 0.217672378 | 0.292977507 | 0.108500112 | 0.209678962 |
| 239,300 | 0.217677502 | 0.293858674 | 0.108965541 | 0.210119795 |
| 239,400 | 0.217676743 | 0.29295681 | 0.109211312 | 0.209594003 |
| 239,500 | 0.217569397 | 0.29347 | 0.109868829 | 0.210007352 |
| 239,600 | 0.217620498 | 0.29368275 | 0.10962773 | 0.208915652 |
| 239,700 | 0.217685697 | 0.295015714 | 0.109408777 | 0.20932862 |
| 239,800 | 0.217512541 | 0.294193499 | 0.110969672 | 0.208845838 |
| 239,900 | 0.217313226 | 0.292748561 | 0.110356076 | 0.208883771 |
| 240,000 | 0.217484314 | 0.293620221 | 0.109996649 | 0.20855061 |
| 240,100 | 0.217547507 | 0.293055185 | 0.109881369 | 0.209176698 |

| | | | | |
|---|---|---|---|---|
| 240,200 | 0.217352306 | 0.292858591 | 0.109942987 | 0.208911974 |
| 240,300 | 0.217350945 | 0.292374987 | 0.108363366 | 0.208585539 |
| 240,400 | 0.217233152 | 0.293109664 | 0.108352907 | 0.208601087 |
| 240,500 | 0.217202165 | 0.292968426 | 0.108929834 | 0.208576984 |
| 240,600 | 0.217328333 | 0.293171247 | 0.108976917 | 0.208267874 |
| 240,700 | 0.217164543 | 0.292184063 | 0.109855168 | 0.208802543 |
| 240,800 | 0.217301181 | 0.29307865 | 0.109421791 | 0.207993455 |
| 240,900 | 0.217186439 | 0.292925313 | 0.108723743 | 0.207537116 |
| 241,000 | 0.217110028 | 0.293053223 | 0.110563603 | 0.208074932 |
| 241,100 | 0.21717166 | 0.29266676 | 0.109456684 | 0.208071433 |
| 241,200 | 0.217054203 | 0.293761907 | 0.109189327 | 0.207397683 |
| 241,300 | 0.217081806 | 0.293551403 | 0.109016857 | 0.207699462 |
| 241,400 | 0.216967852 | 0.29437992 | 0.108864062 | 0.207472792 |
| 241,500 | 0.216891639 | 0.293515803 | 0.109572345 | 0.207847748 |
| 241,600 | 0.216885631 | 0.29340505 | 0.110091793 | 0.207420475 |
| 241,700 | 0.216831129 | 0.292704372 | 0.109324427 | 0.207803678 |
| 241,800 | 0.216694552 | 0.293576019 | 0.109573171 | 0.20832567 |
| 241,900 | 0.216756368 | 0.29403129 | 0.110010033 | 0.207769434 |
| 242,000 | 0.216559477 | 0.29340883 | 0.109003492 | 0.207586858 |
| 242,100 | 0.216713081 | 0.294442955 | 0.110193052 | 0.207913664 |
| 242,200 | 0.21655863 | 0.293218416 | 0.109262506 | 0.207281299 |
| 242,300 | 0.216579684 | 0.294290849 | 0.109028107 | 0.207240205 |
| 242,400 | 0.216405495 | 0.293874488 | 0.108583306 | 0.206920375 |
| 242,500 | 0.216454366 | 0.292939494 | 0.109481827 | 0.207864419 |
| 242,600 | 0.216428913 | 0.293152332 | 0.10851318 | 0.207292001 |
| 242,700 | 0.216416097 | 0.293809059 | 0.109530413 | 0.207236953 |
| 242,800 | 0.216469316 | 0.293249388 | 0.107656702 | 0.207872696 |
| 242,900 | 0.216216718 | 0.293294901 | 0.108056916 | 0.206588751 |
| 243,000 | 0.216211787 | 0.292662818 | 0.107287769 | 0.206707908 |
| 243,100 | 0.216279385 | 0.293899457 | 0.107554264 | 0.207626102 |
| 243,200 | 0.216391265 | 0.295209468 | 0.107893658 | 0.206751582 |
| 243,300 | 0.216380094 | 0.294173928 | 0.108061384 | 0.207357451 |
| 243,400 | 0.216225448 | 0.293323326 | 0.10834716 | 0.207169049 |
| 243,500 | 0.216314078 | 0.294185432 | 0.10951566 | 0.208201566 |
| 243,600 | 0.216266695 | 0.294740796 | 0.108537 | 0.206656124 |
| 243,700 | 0.216164451 | 0.294593839 | 0.108705227 | 0.20700609 |
| 243,800 | 0.216160083 | 0.294409987 | 0.110185764 | 0.207043728 |
| 243,900 | 0.216006114 | 0.294607064 | 0.109396383 | 0.207233971 |
| 244,000 | 0.216121108 | 0.294608666 | 0.110039411 | 0.206706629 |
| 244,100 | 0.216158942 | 0.295176845 | 0.110286423 | 0.207013413 |
| 244,200 | 0.215961334 | 0.294337677 | 0.110942312 | 0.206642342 |
| 244,300 | 0.215903436 | 0.29537839 | 0.110447161 | 0.206503321 |
| 244,400 | 0.216009935 | 0.295477287 | 0.109907484 | 0.206493727 |
| 244,500 | 0.21604865 | 0.294774985 | 0.109678038 | 0.205825175 |
| 244,600 | 0.215897414 | 0.294277071 | 0.110005449 | 0.205799214 |
| 244,700 | 0.216005006 | 0.293966268 | 0.109756186 | 0.205985093 |
| 244,800 | 0.215759735 | 0.294045601 | 0.109585794 | 0.206617436 |
| 244,900 | 0.215801169 | 0.294038273 | 0.10984238 | 0.206970626 |
| 245,000 | 0.215610161 | 0.293416777 | 0.107285121 | 0.205716602 |
| 245,100 | 0.215585862 | 0.29336264 | 0.108575703 | 0.205340035 |
| 245,200 | 0.215648873 | 0.293611122 | 0.109030409 | 0.205472831 |
| 245,300 | 0.215605983 | 0.293404335 | 0.108047619 | 0.205670559 |
| 245,400 | 0.215547399 | 0.292751173 | 0.108676922 | 0.205342801 |
| 245,500 | 0.215544282 | 0.29327871 | 0.10919811 | 0.20514769 |
| 245,600 | 0.215411974 | 0.29316639 | 0.108089647 | 0.203988928 |
| 245,700 | 0.215557819 | 0.293059034 | 0.109667647 | 0.205498067 |
| 245,800 | 0.215430005 | 0.292611352 | 0.108124288 | 0.20537027 |
| 245,900 | 0.215412974 | 0.293597083 | 0.107926314 | 0.205550165 |
| 246,000 | 0.215374366 | 0.292471796 | 0.108025358 | 0.205355454 |
| 246,100 | 0.215305437 | 0.2918986 | 0.107135661 | 0.205958064 |
| 246,200 | 0.215261132 | 0.292331262 | 0.107659503 | 0.205667479 |
| 246,300 | 0.215189262 | 0.29129539 | 0.107582272 | 0.205024777 |
| 246,400 | 0.215052204 | 0.291648725 | 0.106528047 | 0.205075163 |
| 246,500 | 0.215033642 | 0.291721397 | 0.106581003 | 0.205271671 |
| 246,600 | 0.214907495 | 0.291076268 | 0.106978529 | 0.205553599 |
| 246,700 | 0.214856746 | 0.291422874 | 0.107200161 | 0.205168331 |
| 246,800 | 0.214829499 | 0.291753203 | 0.106947471 | 0.205527856 |

| | | | | |
|---|---|---|---|---|
| 246,900 | 0.214832893 | 0.291667993 | 0.10809524 | 0.205604199 |
| 247,000 | 0.214973277 | 0.292130959 | 0.107884517 | 0.20538058 |
| 247,100 | 0.214844587 | 0.291538287 | 0.108300983 | 0.206271905 |
| 247,200 | 0.214812154 | 0.292090292 | 0.10694127 | 0.205371051 |
| 247,300 | 0.21472773 | 0.29152437 | 0.107702325 | 0.205398897 |
| 247,400 | 0.214585889 | 0.291343513 | 0.10597224 | 0.205801128 |
| 247,500 | 0.214425133 | 0.291279143 | 0.107018665 | 0.205875409 |
| 247,600 | 0.214446887 | 0.291684727 | 0.106209833 | 0.206271697 |
| 247,700 | 0.214543208 | 0.291484053 | 0.107601207 | 0.205887648 |
| 247,800 | 0.21444395 | 0.290754721 | 0.107048538 | 0.205922235 |
| 247,900 | 0.214341085 | 0.291579924 | 0.107771502 | 0.20515015 |
| 248,000 | 0.214540975 | 0.291492022 | 0.105610959 | 0.205433192 |
| 248,100 | 0.214277592 | 0.290844663 | 0.106781152 | 0.205154635 |
| 248,200 | 0.21437205 | 0.29183503 | 0.106565157 | 0.205313595 |
| 248,300 | 0.214211477 | 0.29120498 | 0.106483294 | 0.205074381 |
| 248,400 | 0.214103418 | 0.291103492 | 0.106967333 | 0.205114639 |
| 248,500 | 0.214182432 | 0.290878219 | 0.106768854 | 0.206076066 |
| 248,600 | 0.21420183 | 0.29156619 | 0.106114005 | 0.206053498 |
| 248,700 | 0.214171798 | 0.290887991 | 0.106977779 | 0.206399293 |
| 248,800 | 0.213979468 | 0.291056755 | 0.106480675 | 0.204989034 |
| 248,900 | 0.213928442 | 0.290804754 | 0.106878706 | 0.204538497 |
| 249,000 | 0.213844181 | 0.291181943 | 0.106362603 | 0.204768828 |
| 249,100 | 0.213783989 | 0.290188566 | 0.106500024 | 0.204825123 |
| 249,200 | 0.213929066 | 0.290593947 | 0.106670752 | 0.205114182 |
| 249,300 | 0.21385878 | 0.291378277 | 0.105166679 | 0.205693612 |
| 249,400 | 0.213627039 | 0.290335081 | 0.105628512 | 0.205292175 |
| 249,500 | 0.213576042 | 0.290063274 | 0.104328588 | 0.205810507 |
| 249,600 | 0.213479188 | 0.289960396 | 0.105475878 | 0.205024959 |
| 249,700 | 0.213633787 | 0.290673003 | 0.106118679 | 0.205668027 |
| 249,800 | 0.21340363 | 0.290285317 | 0.105893295 | 0.204715144 |
| 249,900 | 0.213469155 | 0.289377322 | 0.106458508 | 0.204865606 |
| 250,000 | 0.213319284 | 0.29127356 | 0.106965782 | 0.205176863 |
| 250,100 | 0.213300383 | 0.289582257 | 0.106580732 | 0.204361373 |
| 250,200 | 0.213262557 | 0.28980844 | 0.105849591 | 0.204865987 |
| 250,300 | 0.213314687 | 0.290763412 | 0.107097789 | 0.205707004 |
| 250,400 | 0.213311321 | 0.29014697 | 0.106875489 | 0.204510042 |
| 250,500 | 0.213217916 | 0.289930106 | 0.105679436 | 0.204889371 |
| 250,600 | 0.213315142 | 0.289459341 | 0.106954519 | 0.20491936 |
| 250,700 | 0.213227092 | 0.289140652 | 0.107271192 | 0.205406736 |
| 250,800 | 0.212927737 | 0.28892075 | 0.105916194 | 0.205261501 |
| 250,900 | 0.212817549 | 0.289555262 | 0.106961158 | 0.20509653 |
| 251,000 | 0.212805753 | 0.289844411 | 0.106387988 | 0.20556433 |
| 251,100 | 0.213019791 | 0.289547547 | 0.106299585 | 0.205388267 |
| 251,200 | 0.21287694 | 0.289597226 | 0.105141457 | 0.205576751 |
| 251,300 | 0.212873256 | 0.288808066 | 0.105661149 | 0.205442843 |
| 251,400 | 0.212836607 | 0.289102808 | 0.104886559 | 0.205061013 |
| 251,500 | 0.212713554 | 0.289329751 | 0.105099298 | 0.205662193 |
| 251,600 | 0.212702032 | 0.287766171 | 0.103710974 | 0.205229734 |
| 251,700 | 0.212561658 | 0.288461968 | 0.104195503 | 0.20531658 |
| 251,800 | 0.212496003 | 0.289116545 | 0.104872438 | 0.205093114 |
| 251,900 | 0.21260789 | 0.288912675 | 0.104264741 | 0.205461356 |
| 252,000 | 0.212661978 | 0.289187246 | 0.104756185 | 0.205486643 |
| 252,100 | 0.212527921 | 0.288823205 | 0.10489169 | 0.205040783 |
| 252,200 | 0.212461192 | 0.288526603 | 0.105398801 | 0.204412899 |
| 252,300 | 0.212426055 | 0.288537368 | 0.104884775 | 0.204582692 |
| 252,400 | 0.212430096 | 0.288287702 | 0.105066759 | 0.20480647 |
| 252,500 | 0.212461139 | 0.287489461 | 0.105421678 | 0.205552209 |
| 252,600 | 0.212265942 | 0.287499104 | 0.104571239 | 0.205126061 |
| 252,700 | 0.212231581 | 0.288324645 | 0.104692086 | 0.205311553 |
| 252,800 | 0.212227507 | 0.287130661 | 0.103768879 | 0.205369865 |
| 252,900 | 0.212342185 | 0.287179802 | 0.104336899 | 0.205127608 |
| 253,000 | 0.212284079 | 0.28789176 | 0.104308569 | 0.205334882 |
| 253,100 | 0.212011618 | 0.287423615 | 0.104667034 | 0.204702025 |
| 253,200 | 0.212301866 | 0.288589923 | 0.104356424 | 0.204791397 |
| 253,300 | 0.212109356 | 0.288117466 | 0.104809138 | 0.205175054 |
| 253,400 | 0.212015072 | 0.287041919 | 0.104139021 | 0.204583502 |
| 253,500 | 0.211973277 | 0.287663965 | 0.104018235 | 0.204750836 |

| | | | | |
|---|---|---|---|---|
| 253,600 | 0.212057677 | 0.287800451 | 0.10427432 | 0.204388324 |
| 253,700 | 0.211935386 | 0.288294521 | 0.104643187 | 0.204202025 |
| 253,800 | 0.211748316 | 0.288165339 | 0.104293329 | 0.204159539 |
| 253,900 | 0.211817493 | 0.286968944 | 0.103623569 | 0.204673041 |
| 254,000 | 0.211823661 | 0.287474084 | 0.104974153 | 0.204513462 |
| 254,100 | 0.211982853 | 0.287882253 | 0.105122695 | 0.20411937 |
| 254,200 | 0.21195324 | 0.286917259 | 0.104340719 | 0.204239514 |
| 254,300 | 0.211939414 | 0.287079895 | 0.103746832 | 0.204389205 |
| 254,400 | 0.211711199 | 0.287249553 | 0.104834688 | 0.204540172 |
| 254,500 | 0.211810025 | 0.287481042 | 0.10329442 | 0.203820327 |
| 254,600 | 0.211772758 | 0.287598975 | 0.104055142 | 0.203354443 |
| 254,700 | 0.211636238 | 0.287932748 | 0.102935926 | 0.203932783 |
| 254,800 | 0.211677714 | 0.28674776 | 0.103292404 | 0.203752301 |
| 254,900 | 0.211709607 | 0.288077392 | 0.103368542 | 0.204016186 |
| 255,000 | 0.211560698 | 0.287821847 | 0.103030136 | 0.203067771 |
| 255,100 | 0.211529325 | 0.288018178 | 0.102455601 | 0.203650494 |
| 255,200 | 0.211564438 | 0.289162134 | 0.103325458 | 0.204046899 |
| 255,300 | 0.211449945 | 0.288794874 | 0.103378662 | 0.204232019 |
| 255,400 | 0.211419269 | 0.288434843 | 0.103086879 | 0.2038189 |
| 255,500 | 0.211548108 | 0.287785946 | 0.104052516 | 0.203573218 |
| 255,600 | 0.211335386 | 0.287818754 | 0.102758293 | 0.203221706 |
| 255,700 | 0.211343413 | 0.287770874 | 0.102444043 | 0.203005405 |
| 255,800 | 0.21137425 | 0.28728297 | 0.102178283 | 0.203039032 |
| 255,900 | 0.211113816 | 0.287005506 | 0.102561054 | 0.202496816 |
| 256,000 | 0.211379945 | 0.287010724 | 0.103593805 | 0.202579042 |
| 256,100 | 0.211236769 | 0.287174933 | 0.104438238 | 0.203240518 |
| 256,200 | 0.21137479 | 0.28796757 | 0.102066868 | 0.202274185 |
| 256,300 | 0.211194105 | 0.287528702 | 0.103336363 | 0.202809221 |
| 256,400 | 0.211352223 | 0.287715998 | 0.103148373 | 0.203354115 |
| 256,500 | 0.211094009 | 0.28632024 | 0.10481151 | 0.202490726 |
| 256,600 | 0.211174962 | 0.286531216 | 0.104421338 | 0.203033759 |
| 256,700 | 0.211034308 | 0.286697403 | 0.104137166 | 0.20259886 |
| 256,800 | 0.211005883 | 0.287374563 | 0.103259112 | 0.202172834 |
| 256,900 | 0.210880252 | 0.286688975 | 0.102690561 | 0.201526031 |
| 257,000 | 0.210991362 | 0.287307012 | 0.102611379 | 0.201951535 |
| 257,100 | 0.210894378 | 0.286935348 | 0.103976205 | 0.202132108 |
| 257,200 | 0.210904782 | 0.28841891 | 0.103739026 | 0.201623646 |
| 257,300 | 0.210897301 | 0.287086933 | 0.103260873 | 0.202238186 |
| 257,400 | 0.210793596 | 0.28714432 | 0.102765774 | 0.20149157 |
| 257,500 | 0.210991809 | 0.286819902 | 0.102224543 | 0.202005051 |
| 257,600 | 0.210856652 | 0.28651863 | 0.10303842 | 0.201878737 |
| 257,700 | 0.210847706 | 0.286052714 | 0.102962832 | 0.201897739 |
| 257,800 | 0.210895256 | 0.286297656 | 0.103088884 | 0.201946355 |
| 257,900 | 0.210757668 | 0.287640904 | 0.101750996 | 0.201200316 |
| 258,000 | 0.210892766 | 0.286372659 | 0.101193693 | 0.201573597 |
| 258,100 | 0.210616755 | 0.286275151 | 0.103081218 | 0.201217454 |
| 258,200 | 0.210729574 | 0.287091955 | 0.101447812 | 0.200550572 |
| 258,300 | 0.210751621 | 0.286692601 | 0.103190844 | 0.201109954 |
| 258,400 | 0.210488975 | 0.285488286 | 0.102327575 | 0.200262833 |
| 258,500 | 0.210697113 | 0.286546188 | 0.103021934 | 0.200247403 |
| 258,600 | 0.210480611 | 0.286646361 | 0.101368623 | 0.200120124 |
| 258,700 | 0.210466849 | 0.286980631 | 0.100242477 | 0.200910105 |
| 258,800 | 0.210508417 | 0.286325382 | 0.099284126 | 0.20100497 |
| 258,900 | 0.210433835 | 0.285528593 | 0.098005914 | 0.201158203 |
| 259,000 | 0.210364161 | 0.286027019 | 0.099088995 | 0.201184921 |
| 259,100 | 0.210408927 | 0.285820054 | 0.100662421 | 0.201106219 |
| 259,200 | 0.210282108 | 0.286421817 | 0.099997388 | 0.201174476 |
| 259,300 | 0.21039412 | 0.288251474 | 0.100500059 | 0.200662912 |
| 259,400 | 0.210342067 | 0.287184115 | 0.099588177 | 0.201180823 |
| 259,500 | 0.210098817 | 0.286441269 | 0.100489295 | 0.200935119 |
| 259,600 | 0.210292465 | 0.287464843 | 0.099633987 | 0.201282238 |
| 259,700 | 0.210328129 | 0.287150545 | 0.099456305 | 0.201630655 |
| 259,800 | 0.210437673 | 0.287151477 | 0.099419003 | 0.201849738 |
| 259,900 | 0.210274625 | 0.287139381 | 0.099127738 | 0.200933353 |
| 260,000 | 0.210220369 | 0.287280178 | 0.098485014 | 0.20132708 |
| 260,100 | 0.210047526 | 0.286871686 | 0.099330423 | 0.201684322 |
| 260,200 | 0.209942273 | 0.286544695 | 0.098563132 | 0.20164225 |

| | | | | |
|---|---|---|---|---|
| 260,300 | 0.209995417 | 0.28695301 | 0.098909733 | 0.20148906 |
| 260,400 | 0.209768057 | 0.285626328 | 0.098735002 | 0.201342988 |
| 260,500 | 0.209820023 | 0.285877136 | 0.099933164 | 0.200753801 |
| 260,600 | 0.20972403 | 0.285870923 | 0.098986349 | 0.201445043 |
| 260,700 | 0.209676891 | 0.285705641 | 0.099192564 | 0.200851059 |
| 260,800 | 0.209763372 | 0.285826009 | 0.099669148 | 0.201501255 |
| 260,900 | 0.209784787 | 0.286362488 | 0.100090857 | 0.201626221 |
| 261,000 | 0.209858446 | 0.286411715 | 0.099773726 | 0.201297024 |
| 261,100 | 0.209686496 | 0.285884372 | 0.100300112 | 0.201164929 |
| 261,200 | 0.209604686 | 0.284179866 | 0.102224797 | 0.201184214 |
| 261,300 | 0.209620677 | 0.28487402 | 0.100777825 | 0.200753796 |
| 261,400 | 0.209580747 | 0.284676484 | 0.099992796 | 0.201193404 |
| 261,500 | 0.209558326 | 0.284608176 | 0.09991138 | 0.201451692 |
| 261,600 | 0.209531425 | 0.284930098 | 0.099740767 | 0.201289475 |
| 261,700 | 0.209389988 | 0.284747088 | 0.099032878 | 0.201346942 |
| 261,800 | 0.209283324 | 0.285071849 | 0.098897358 | 0.201233744 |
| 261,900 | 0.20920464 | 0.284948734 | 0.098895858 | 0.200781458 |
| 262,000 | 0.20936221 | 0.285605263 | 0.099680133 | 0.201496322 |
| 262,100 | 0.209415387 | 0.284538573 | 0.100608852 | 0.201606655 |
| 262,200 | 0.209347865 | 0.284337927 | 0.099173504 | 0.201570126 |
| 262,300 | 0.209359506 | 0.284298237 | 0.100061089 | 0.201672043 |
| 262,400 | 0.20936646 | 0.284687264 | 0.100066678 | 0.201427653 |
| 262,500 | 0.209285098 | 0.284096691 | 0.098953146 | 0.201919943 |
| 262,600 | 0.209141574 | 0.283631101 | 0.099792014 | 0.20140427 |
| 262,700 | 0.209118537 | 0.283583984 | 0.09858253 | 0.201177772 |
| 262,800 | 0.209124743 | 0.284136647 | 0.099912668 | 0.201248691 |
| 262,900 | 0.209056742 | 0.28373187 | 0.098476744 | 0.202057893 |
| 263,000 | 0.209067612 | 0.283260357 | 0.099456771 | 0.201890257 |
| 263,100 | 0.209077263 | 0.28336176 | 0.097264794 | 0.201907652 |
| 263,200 | 0.208919546 | 0.283861773 | 0.098469688 | 0.201519564 |
| 263,300 | 0.208838197 | 0.282746641 | 0.098456655 | 0.202461683 |
| 263,400 | 0.208981547 | 0.283088152 | 0.099090087 | 0.201889157 |
| 263,500 | 0.208963997 | 0.283616964 | 0.099233674 | 0.202581861 |
| 263,600 | 0.208797421 | 0.283653067 | 0.099222429 | 0.202206241 |
| 263,700 | 0.208894337 | 0.284388798 | 0.099425103 | 0.201455014 |
| 263,800 | 0.208884688 | 0.283863559 | 0.098854458 | 0.202004416 |
| 263,900 | 0.208806045 | 0.284453523 | 0.098332334 | 0.201550385 |
| 264,000 | 0.208980207 | 0.284216059 | 0.099246629 | 0.202057561 |
| 264,100 | 0.208971342 | 0.28439507 | 0.098041282 | 0.202636019 |
| 264,200 | 0.209056134 | 0.284100403 | 0.098545935 | 0.20118171 |
| 264,300 | 0.208845533 | 0.284734479 | 0.097159409 | 0.20154376 |
| 264,400 | 0.208793137 | 0.285063678 | 0.098538857 | 0.201939843 |
| 264,500 | 0.208866841 | 0.284227285 | 0.098279164 | 0.201798165 |
| 264,600 | 0.208671776 | 0.282934897 | 0.09916421 | 0.201865663 |
| 264,700 | 0.20863662 | 0.284486207 | 0.099637627 | 0.202074246 |
| 264,800 | 0.208640565 | 0.28336364 | 0.099810508 | 0.201110851 |
| 264,900 | 0.208430492 | 0.284251201 | 0.09880014 | 0.201478148 |
| 265,000 | 0.208528785 | 0.284396102 | 0.098925476 | 0.202136454 |
| 265,100 | 0.208503847 | 0.284521152 | 0.09805369 | 0.201432859 |
| 265,200 | 0.208339817 | 0.283537316 | 0.099140988 | 0.200853276 |
| 265,300 | 0.208404016 | 0.284280752 | 0.09808349 | 0.201856071 |
| 265,400 | 0.208437344 | 0.283702652 | 0.098452847 | 0.201226135 |
| 265,500 | 0.208501091 | 0.284020831 | 0.097802188 | 0.201703442 |
| 265,600 | 0.208301875 | 0.28388945 | 0.097214406 | 0.201474028 |
| 265,700 | 0.208372452 | 0.284991497 | 0.097639882 | 0.201041929 |
| 265,800 | 0.208169336 | 0.284720592 | 0.09718044 | 0.201146222 |
| 265,900 | 0.208188463 | 0.284035183 | 0.098057745 | 0.200939856 |
| 266,000 | 0.208062972 | 0.284518079 | 0.097737361 | 0.201501279 |
| 266,100 | 0.208171391 | 0.284732106 | 0.098104601 | 0.200814823 |
| 266,200 | 0.208172285 | 0.284921201 | 0.098195718 | 0.200287206 |
| 266,300 | 0.208062927 | 0.284424321 | 0.098263008 | 0.200878361 |
| 266,400 | 0.208022125 | 0.28407309 | 0.097227445 | 0.200975356 |
| 266,500 | 0.207978143 | 0.283635829 | 0.096939784 | 0.201817317 |
| 266,600 | 0.207909934 | 0.284638767 | 0.097330533 | 0.202555841 |
| 266,700 | 0.207750672 | 0.284335088 | 0.09684665 | 0.20142228 |
| 266,800 | 0.207803815 | 0.285030781 | 0.096615275 | 0.201831594 |
| 266,900 | 0.207759595 | 0.283789018 | 0.096626997 | 0.201922834 |

| | | | | |
|---|---|---|---|---|
| 267,000 | 0.207798258 | 0.284239528 | 0.098124315 | 0.201379887 |
| 267,100 | 0.207693795 | 0.283416345 | 0.097676158 | 0.200899131 |
| 267,200 | 0.207920103 | 0.283629732 | 0.097891443 | 0.201747951 |
| 267,300 | 0.207704577 | 0.283913781 | 0.0985178 | 0.201322765 |
| 267,400 | 0.207922225 | 0.284720947 | 0.097349447 | 0.201880793 |
| 267,500 | 0.207828743 | 0.283956293 | 0.096894853 | 0.201844248 |
| 267,600 | 0.207785966 | 0.283834581 | 0.097802653 | 0.201849299 |
| 267,700 | 0.207578316 | 0.283125384 | 0.096850256 | 0.201201339 |
| 267,800 | 0.207576756 | 0.283484224 | 0.096464742 | 0.201223966 |
| 267,900 | 0.207673457 | 0.284491336 | 0.09799715 | 0.201426568 |
| 268,000 | 0.207599436 | 0.284160004 | 0.097369786 | 0.20052795 |
| 268,100 | 0.207535369 | 0.284065259 | 0.097440585 | 0.200482996 |
| 268,200 | 0.207567793 | 0.283772568 | 0.097942192 | 0.200930204 |
| 268,300 | 0.207614664 | 0.283051171 | 0.097006993 | 0.200455249 |
| 268,400 | 0.207588203 | 0.283830778 | 0.098628984 | 0.201192498 |
| 268,500 | 0.207724183 | 0.284029262 | 0.097705999 | 0.200445787 |
| 268,600 | 0.207589488 | 0.283323105 | 0.097553177 | 0.200996915 |
| 268,700 | 0.207362338 | 0.283247221 | 0.097432755 | 0.200186639 |
| 268,800 | 0.207616362 | 0.28282214 | 0.097877445 | 0.200151743 |
| 268,900 | 0.207422983 | 0.283063134 | 0.097917087 | 0.199617144 |
| 269,000 | 0.207546985 | 0.282814527 | 0.097177823 | 0.200208159 |
| 269,100 | 0.207529669 | 0.283126539 | 0.097676118 | 0.20010986 |
| 269,200 | 0.207400382 | 0.282888042 | 0.097803333 | 0.199710746 |
| 269,300 | 0.207526432 | 0.283051283 | 0.098199615 | 0.199507918 |
| 269,400 | 0.207210744 | 0.282587334 | 0.097295342 | 0.200230676 |
| 269,500 | 0.207371632 | 0.282973114 | 0.098105921 | 0.200102645 |
| 269,600 | 0.207267848 | 0.282325208 | 0.097442749 | 0.200616522 |
| 269,700 | 0.207373712 | 0.282341734 | 0.097254238 | 0.200788527 |
| 269,800 | 0.207211029 | 0.282198693 | 0.098782419 | 0.199826142 |
| 269,900 | 0.207110482 | 0.281880249 | 0.098482672 | 0.200453206 |
| 270,000 | 0.207090157 | 0.281723102 | 0.099111466 | 0.201052828 |
| 270,100 | 0.206995335 | 0.281698394 | 0.098612025 | 0.200258899 |
| 270,200 | 0.206919565 | 0.282165847 | 0.09826538 | 0.199820527 |
| 270,300 | 0.206959516 | 0.282226817 | 0.098760054 | 0.200101938 |
| 270,400 | 0.207027538 | 0.281614873 | 0.097795562 | 0.200243425 |
| 270,500 | 0.206907392 | 0.282245532 | 0.098938581 | 0.200589619 |
| 270,600 | 0.206817896 | 0.281431323 | 0.098468597 | 0.200216107 |
| 270,700 | 0.206738117 | 0.282024265 | 0.098546466 | 0.200450452 |
| 270,800 | 0.206825744 | 0.282627688 | 0.099247116 | 0.200966352 |
| 270,900 | 0.206843191 | 0.281970194 | 0.098413001 | 0.200911902 |
| 271,000 | 0.206695835 | 0.282261864 | 0.098131098 | 0.201748181 |
| 271,100 | 0.20697219 | 0.282552917 | 0.097758379 | 0.200679824 |
| 271,200 | 0.206850461 | 0.282184559 | 0.096951049 | 0.200919745 |
| 271,300 | 0.206720173 | 0.282487372 | 0.097367039 | 0.20076309 |
| 271,400 | 0.206623782 | 0.281312043 | 0.097608386 | 0.200885967 |
| 271,500 | 0.20662414 | 0.282188145 | 0.097301019 | 0.200646518 |
| 271,600 | 0.206568531 | 0.281345289 | 0.097457928 | 0.201071503 |
| 271,700 | 0.206632292 | 0.281341326 | 0.097384983 | 0.200949279 |
| 271,800 | 0.206832638 | 0.281649105 | 0.096924624 | 0.201250667 |
| 271,900 | 0.206895667 | 0.282103085 | 0.096628205 | 0.201158282 |
| 272,000 | 0.206625427 | 0.281706846 | 0.096478848 | 0.200734413 |
| 272,100 | 0.206483914 | 0.281592113 | 0.097329606 | 0.201260271 |
| 272,200 | 0.206442284 | 0.281434591 | 0.097245627 | 0.200727216 |
| 272,300 | 0.206356266 | 0.281366785 | 0.096883719 | 0.200542677 |
| 272,400 | 0.206356203 | 0.281368439 | 0.096404376 | 0.200488106 |
| 272,500 | 0.206263663 | 0.281232648 | 0.09764393 | 0.200829519 |
| 272,600 | 0.206356645 | 0.280477082 | 0.097630115 | 0.20139402 |
| 272,700 | 0.206303836 | 0.280370613 | 0.097543568 | 0.200168985 |
| 272,800 | 0.206273298 | 0.2806762 | 0.097983265 | 0.199744573 |
| 272,900 | 0.206033558 | 0.279543369 | 0.098369223 | 0.200653278 |
| 273,000 | 0.206259399 | 0.27930529 | 0.097456036 | 0.201022524 |
| 273,100 | 0.206239477 | 0.279646485 | 0.098388526 | 0.200492892 |
| 273,200 | 0.206232661 | 0.280542242 | 0.097119349 | 0.200269729 |
| 273,300 | 0.206345596 | 0.280298397 | 0.097466176 | 0.200378401 |
| 273,400 | 0.206167339 | 0.280436828 | 0.097770645 | 0.200108042 |
| 273,500 | 0.206149437 | 0.281534751 | 0.097622319 | 0.200437354 |
| 273,600 | 0.206107982 | 0.279910738 | 0.098479498 | 0.201115578 |

| | | | | |
|---|---|---|---|---|
| 273,700 | 0.20594792 | 0.280407293 | 0.097732184 | 0.200677388 |
| 273,800 | 0.206110274 | 0.280627124 | 0.096069857 | 0.200582904 |
| 273,900 | 0.206101018 | 0.281073377 | 0.097234609 | 0.199468777 |
| 274,000 | 0.206208686 | 0.280552694 | 0.097906988 | 0.199741198 |
| 274,100 | 0.206094633 | 0.280799391 | 0.097660144 | 0.199607606 |
| 274,200 | 0.205888247 | 0.2801172 | 0.097089466 | 0.199679527 |
| 274,300 | 0.205853247 | 0.279875137 | 0.096682377 | 0.198952909 |
| 274,400 | 0.205858749 | 0.279647003 | 0.097387718 | 0.199211088 |
| 274,500 | 0.205638291 | 0.279655411 | 0.097428717 | 0.199032624 |
| 274,600 | 0.20577865 | 0.279683888 | 0.098301583 | 0.198967178 |
| 274,700 | 0.205697755 | 0.279024523 | 0.098158187 | 0.199374665 |
| 274,800 | 0.205825994 | 0.278949695 | 0.097646125 | 0.199210119 |
| 274,900 | 0.205741954 | 0.27900521 | 0.09712316 | 0.199412976 |
| 275,000 | 0.205609425 | 0.278763456 | 0.097478957 | 0.199576724 |
| 275,100 | 0.205816806 | 0.279512119 | 0.097920261 | 0.199151809 |
| 275,200 | 0.205797834 | 0.279373853 | 0.096628533 | 0.1995008 |
| 275,300 | 0.205887727 | 0.28012553 | 0.096578434 | 0.200413059 |
| 275,400 | 0.205783541 | 0.278633816 | 0.096456065 | 0.199735873 |
| 275,500 | 0.205690886 | 0.279384429 | 0.095681188 | 0.199723846 |
| 275,600 | 0.205454929 | 0.279734654 | 0.095958225 | 0.199274329 |
| 275,700 | 0.205547103 | 0.2794134 | 0.095586366 | 0.200299968 |
| 275,800 | 0.20546166 | 0.279064897 | 0.096546888 | 0.199360315 |
| 275,900 | 0.2053964 | 0.279703057 | 0.095979207 | 0.198798157 |
| 276,000 | 0.205419926 | 0.279702059 | 0.095213544 | 0.19943544 |
| 276,100 | 0.205338284 | 0.279158668 | 0.095554719 | 0.198547688 |
| 276,200 | 0.205370268 | 0.279307138 | 0.09499477 | 0.198870487 |
| 276,300 | 0.205237722 | 0.27874164 | 0.095131214 | 0.199190041 |
| 276,400 | 0.205410104 | 0.279256834 | 0.095830386 | 0.19949588 |
| 276,500 | 0.205335594 | 0.279793299 | 0.096211532 | 0.198692464 |
| 276,600 | 0.205155944 | 0.279523775 | 0.095524025 | 0.198393023 |
| 276,700 | 0.205023918 | 0.278730559 | 0.095297314 | 0.198576392 |
| 276,800 | 0.205246202 | 0.278852873 | 0.095711777 | 0.198818907 |
| 276,900 | 0.205135127 | 0.279139138 | 0.095137066 | 0.198682164 |
| 277,000 | 0.205327956 | 0.279845087 | 0.095231156 | 0.199184446 |
| 277,100 | 0.205175244 | 0.278536199 | 0.095916053 | 0.198934288 |
| 277,200 | 0.20516618 | 0.279096426 | 0.095492989 | 0.199170923 |
| 277,300 | 0.205012272 | 0.279522042 | 0.095866221 | 0.19876621 |
| 277,400 | 0.204946701 | 0.279947685 | 0.095944039 | 0.198205627 |
| 277,500 | 0.204892874 | 0.278523597 | 0.09608088 | 0.198785727 |
| 277,600 | 0.204812708 | 0.278754174 | 0.095526141 | 0.199282222 |
| 277,700 | 0.204779816 | 0.278386378 | 0.09604294 | 0.198127579 |
| 277,800 | 0.204792268 | 0.278005284 | 0.095427038 | 0.198620564 |
| 277,900 | 0.204566243 | 0.278472771 | 0.096125117 | 0.198382231 |
| 278,000 | 0.204763248 | 0.277737622 | 0.096069506 | 0.198683072 |
| 278,100 | 0.204689251 | 0.277705851 | 0.096357858 | 0.198225762 |
| 278,200 | 0.204582604 | 0.278159816 | 0.095434776 | 0.198059917 |
| 278,300 | 0.204811094 | 0.279534634 | 0.096905653 | 0.198274309 |
| 278,400 | 0.204649425 | 0.279029237 | 0.096558697 | 0.198639308 |
| 278,500 | 0.204516739 | 0.278748858 | 0.09666174 | 0.198178389 |
| 278,600 | 0.204493222 | 0.27881974 | 0.09631391 | 0.198654112 |
| 278,700 | 0.204412861 | 0.278334574 | 0.095668902 | 0.198010044 |
| 278,800 | 0.204652696 | 0.278581782 | 0.096943307 | 0.198843174 |
| 278,900 | 0.204459338 | 0.278024821 | 0.096465423 | 0.198897803 |
| 279,000 | 0.20436703 | 0.278092425 | 0.096945295 | 0.199438459 |
| 279,100 | 0.204334387 | 0.278224455 | 0.096181685 | 0.199340572 |
| 279,200 | 0.204285181 | 0.278770252 | 0.096570809 | 0.199125434 |
| 279,300 | 0.204334727 | 0.278661457 | 0.096036985 | 0.199148156 |
| 279,400 | 0.204403269 | 0.279088701 | 0.096364573 | 0.199503226 |
| 279,500 | 0.204359824 | 0.278802741 | 0.095811987 | 0.199589957 |
| 279,600 | 0.204270044 | 0.279609551 | 0.09615089 | 0.199389662 |
| 279,700 | 0.204237629 | 0.278235261 | 0.095853251 | 0.198806784 |
| 279,800 | 0.204268847 | 0.27890783 | 0.096623766 | 0.199181715 |
| 279,900 | 0.204128681 | 0.278764961 | 0.096400372 | 0.199146107 |
| 280,000 | 0.204199486 | 0.278396474 | 0.096918076 | 0.199504841 |
| 280,100 | 0.204035621 | 0.278743284 | 0.096020729 | 0.199148767 |
| 280,200 | 0.203982723 | 0.279756772 | 0.096122297 | 0.199316501 |
| 280,300 | 0.204023571 | 0.279375058 | 0.095291641 | 0.199295375 |

| | | | | |
|---|---|---|---|---|
| 280,400 | 0.20388162 | 0.278316714 | 0.095774096 | 0.199095988 |
| 280,500 | 0.203891044 | 0.277977275 | 0.095122413 | 0.198868394 |
| 280,600 | 0.203879355 | 0.277987235 | 0.094557272 | 0.198994289 |
| 280,700 | 0.203901587 | 0.278084752 | 0.094632082 | 0.199200325 |
| 280,800 | 0.203820341 | 0.277856747 | 0.094703153 | 0.198686988 |
| 280,900 | 0.203802704 | 0.277759348 | 0.09459831 | 0.198390417 |
| 281,000 | 0.20373755 | 0.277773146 | 0.094739113 | 0.19837392 |
| 281,100 | 0.203679767 | 0.277672002 | 0.095501261 | 0.198250882 |
| 281,200 | 0.203707818 | 0.278324226 | 0.096221137 | 0.198131372 |
| 281,300 | 0.203640018 | 0.277632307 | 0.095594019 | 0.19868068 |
| 281,400 | 0.203596556 | 0.277431375 | 0.096772929 | 0.198372362 |
| 281,500 | 0.203568099 | 0.278059267 | 0.09552067 | 0.198218069 |
| 281,600 | 0.203771911 | 0.27710682 | 0.094574722 | 0.198663041 |
| 281,700 | 0.203489383 | 0.276748909 | 0.095499969 | 0.197801599 |
| 281,800 | 0.203649735 | 0.278243885 | 0.096057334 | 0.199143847 |
| 281,900 | 0.203486081 | 0.277366548 | 0.095903159 | 0.198793723 |
| 282,000 | 0.203476542 | 0.277327425 | 0.094904926 | 0.19836845 |
| 282,100 | 0.203448162 | 0.277283514 | 0.09587673 | 0.198712957 |
| 282,200 | 0.203476814 | 0.277755316 | 0.09524363 | 0.198688006 |
| 282,300 | 0.203396194 | 0.277861285 | 0.096014313 | 0.198214109 |
| 282,400 | 0.203324207 | 0.277364433 | 0.095449651 | 0.1989066 |
| 282,500 | 0.203376209 | 0.278015266 | 0.095804946 | 0.198575549 |
| 282,600 | 0.203236065 | 0.278163603 | 0.095232039 | 0.198579337 |
| 282,700 | 0.203419662 | 0.277957869 | 0.095352494 | 0.198185579 |
| 282,800 | 0.203184934 | 0.277372661 | 0.094943293 | 0.198524433 |
| 282,900 | 0.203214751 | 0.276905807 | 0.094597169 | 0.198821817 |
| 283,000 | 0.203141712 | 0.277551554 | 0.09550929 | 0.19819208 |
| 283,100 | 0.203044181 | 0.277509377 | 0.09477563 | 0.198290011 |
| 283,200 | 0.202978111 | 0.277362601 | 0.094860234 | 0.198393274 |
| 283,300 | 0.202838151 | 0.277247524 | 0.094785575 | 0.198641608 |
| 283,400 | 0.202730116 | 0.277429533 | 0.094152472 | 0.198451215 |
| 283,500 | 0.20287996 | 0.277467121 | 0.094220296 | 0.198207245 |
| 283,600 | 0.202829705 | 0.278485384 | 0.094381315 | 0.198618282 |
| 283,700 | 0.202816661 | 0.277986562 | 0.093820974 | 0.19838422 |
| 283,800 | 0.202834956 | 0.278102605 | 0.095308437 | 0.198872229 |
| 283,900 | 0.202834704 | 0.277985715 | 0.094254625 | 0.198193234 |
| 284,000 | 0.202753877 | 0.277195278 | 0.093686841 | 0.199096611 |
| 284,100 | 0.202792351 | 0.278283235 | 0.094480577 | 0.198409071 |
| 284,200 | 0.202772814 | 0.278049348 | 0.094667843 | 0.198741757 |
| 284,300 | 0.202641188 | 0.278251163 | 0.09466484 | 0.198780354 |
| 284,400 | 0.202774971 | 0.277964172 | 0.094861152 | 0.197885791 |
| 284,500 | 0.202631256 | 0.278263935 | 0.093460834 | 0.19847679 |
| 284,600 | 0.202667206 | 0.278625349 | 0.093792672 | 0.198839844 |
| 284,700 | 0.202725272 | 0.278146396 | 0.094353103 | 0.198767509 |
| 284,800 | 0.202670978 | 0.278716959 | 0.093996066 | 0.198106714 |
| 284,900 | 0.202604447 | 0.277983255 | 0.095272876 | 0.197850029 |
| 285,000 | 0.202517594 | 0.278692433 | 0.095399886 | 0.198151356 |
| 285,100 | 0.202617059 | 0.278530421 | 0.094086518 | 0.198149172 |
| 285,200 | 0.202432705 | 0.277731848 | 0.094614444 | 0.198237558 |
| 285,300 | 0.202437409 | 0.277079244 | 0.094423216 | 0.198118131 |
| 285,400 | 0.202410028 | 0.276231072 | 0.095398257 | 0.197680627 |
| 285,500 | 0.202425315 | 0.278301926 | 0.095157802 | 0.197118174 |
| 285,600 | 0.202377816 | 0.278741896 | 0.09488387 | 0.197101718 |
| 285,700 | 0.202271048 | 0.278410639 | 0.09594647 | 0.197448038 |
| 285,800 | 0.202344179 | 0.279315462 | 0.095240088 | 0.197080804 |
| 285,900 | 0.202352283 | 0.278422694 | 0.094278553 | 0.196918484 |
| 286,000 | 0.20217789 | 0.278142757 | 0.095829113 | 0.197156366 |
| 286,100 | 0.202309889 | 0.278982521 | 0.095755633 | 0.197669813 |
| 286,200 | 0.202137232 | 0.278166951 | 0.094767939 | 0.197905433 |
| 286,300 | 0.202055411 | 0.278664115 | 0.095029948 | 0.198392636 |
| 286,400 | 0.202056117 | 0.278416076 | 0.095036367 | 0.197483892 |
| 286,500 | 0.201907267 | 0.278824469 | 0.094484755 | 0.197182028 |
| 286,600 | 0.201802709 | 0.278201987 | 0.09489693 | 0.19617128 |
| 286,700 | 0.201890078 | 0.279341526 | 0.095362217 | 0.196804707 |
| 286,800 | 0.201899626 | 0.278512881 | 0.094895772 | 0.196233419 |
| 286,900 | 0.201778124 | 0.278684992 | 0.094712994 | 0.197288667 |
| 287,000 | 0.201707809 | 0.278461885 | 0.09518622 | 0.196905653 |

| | | | | |
|---|---|---|---|---|
| 287,100 | 0.201640241 | 0.277516647 | 0.094469129 | 0.196621964 |
| 287,200 | 0.2018374 | 0.278678365 | 0.093421656 | 0.196742064 |
| 287,300 | 0.201664359 | 0.278022249 | 0.093538346 | 0.197329785 |
| 287,400 | 0.201726856 | 0.278736022 | 0.093961109 | 0.197386691 |
| 287,500 | 0.201747009 | 0.278247754 | 0.094313882 | 0.19716994 |
| 287,600 | 0.20169187 | 0.279241949 | 0.094531871 | 0.196970558 |
| 287,700 | 0.201487424 | 0.277551843 | 0.093393586 | 0.19619087 |
| 287,800 | 0.201406719 | 0.278130064 | 0.093546471 | 0.19610329 |
| 287,900 | 0.201395876 | 0.277452791 | 0.092409556 | 0.196864194 |
| 288,000 | 0.201491094 | 0.277635425 | 0.092589101 | 0.196648798 |
| 288,100 | 0.201454863 | 0.278023184 | 0.093626607 | 0.196980286 |
| 288,200 | 0.201443999 | 0.277233274 | 0.092375793 | 0.196663778 |
| 288,300 | 0.20117877 | 0.277861176 | 0.092789413 | 0.196447559 |
| 288,400 | 0.201022183 | 0.277789136 | 0.092702052 | 0.196100365 |
| 288,500 | 0.201156872 | 0.277370531 | 0.092497761 | 0.196226608 |
| 288,600 | 0.201206366 | 0.27701808 | 0.093321154 | 0.195449985 |
| 288,700 | 0.201167742 | 0.276894588 | 0.092814511 | 0.195558699 |
| 288,800 | 0.201128177 | 0.276473437 | 0.092317505 | 0.196187471 |
| 288,900 | 0.201048368 | 0.277984846 | 0.092264225 | 0.195930827 |
| 289,000 | 0.201045291 | 0.276693222 | 0.092333283 | 0.195020261 |
| 289,100 | 0.201125878 | 0.276578507 | 0.093231914 | 0.196039757 |
| 289,200 | 0.200845643 | 0.276585817 | 0.093342102 | 0.194809012 |
| 289,300 | 0.201053197 | 0.276874829 | 0.093706363 | 0.195646759 |
| 289,400 | 0.201034484 | 0.27684962 | 0.092649634 | 0.195939916 |
| 289,500 | 0.200988867 | 0.276061437 | 0.092306886 | 0.195257185 |
| 289,600 | 0.200920067 | 0.27592612 | 0.09368083 | 0.195585589 |
| 289,700 | 0.200839474 | 0.276365912 | 0.094609958 | 0.195580672 |
| 289,800 | 0.200889273 | 0.277287311 | 0.092630869 | 0.195860172 |
| 289,900 | 0.200844031 | 0.276914943 | 0.092387182 | 0.195910161 |
| 290,000 | 0.200941453 | 0.276305741 | 0.092394018 | 0.194761696 |
| 290,100 | 0.200733519 | 0.275850209 | 0.09260887 | 0.19548644 |
| 290,200 | 0.200779221 | 0.276234989 | 0.092390655 | 0.19527621 |
| 290,300 | 0.200655279 | 0.277062028 | 0.092743986 | 0.194680295 |
| 290,400 | 0.20083779 | 0.277226325 | 0.092014029 | 0.194693722 |
| 290,500 | 0.200582655 | 0.275844121 | 0.091776023 | 0.194938188 |
| 290,600 | 0.20047752 | 0.275914577 | 0.092942371 | 0.194592212 |
| 290,700 | 0.200567896 | 0.276222327 | 0.092771059 | 0.194549365 |
| 290,800 | 0.200501276 | 0.276402622 | 0.09309627 | 0.195169477 |
| 290,900 | 0.200513626 | 0.27587128 | 0.092859895 | 0.1942672 |
| 291,000 | 0.20041628 | 0.276343304 | 0.092593616 | 0.194304384 |
| 291,100 | 0.200532747 | 0.275448367 | 0.093617696 | 0.194090859 |
| 291,200 | 0.200511041 | 0.275784781 | 0.094312575 | 0.195026701 |
| 291,300 | 0.200377227 | 0.276372772 | 0.094090592 | 0.194437609 |
| 291,400 | 0.200415316 | 0.276763462 | 0.093530572 | 0.194251147 |
| 291,500 | 0.200274515 | 0.275816593 | 0.094353007 | 0.194660261 |
| 291,600 | 0.200257574 | 0.276169549 | 0.093169898 | 0.194051027 |
| 291,700 | 0.200290135 | 0.276297584 | 0.093726508 | 0.193743709 |
| 291,800 | 0.200221111 | 0.274675338 | 0.093406018 | 0.194214574 |
| 291,900 | 0.20028039 | 0.2759153 | 0.093994063 | 0.193390579 |
| 292,000 | 0.200172503 | 0.274940452 | 0.094477773 | 0.193899761 |
| 292,100 | 0.200243451 | 0.275594261 | 0.094924079 | 0.193784287 |
| 292,200 | 0.200334076 | 0.276055668 | 0.093900721 | 0.193547156 |
| 292,300 | 0.200309087 | 0.275750876 | 0.093730047 | 0.194724987 |
| 292,400 | 0.200312697 | 0.275251287 | 0.092935006 | 0.194245148 |
| 292,500 | 0.200258817 | 0.275722824 | 0.092057742 | 0.194109623 |
| 292,600 | 0.200123305 | 0.275704079 | 0.092799054 | 0.19379148 |
| 292,700 | 0.199977884 | 0.275112701 | 0.092779668 | 0.193849833 |
| 292,800 | 0.199970477 | 0.275027682 | 0.093093874 | 0.193953407 |
| 292,900 | 0.199897543 | 0.275265587 | 0.093110743 | 0.193737309 |
| 293,000 | 0.200123714 | 0.27474622 | 0.092768395 | 0.194289658 |
| 293,100 | 0.199777562 | 0.274396222 | 0.092559792 | 0.193498732 |
| 293,200 | 0.199823584 | 0.273969773 | 0.09209505 | 0.19368429 |
| 293,300 | 0.199796082 | 0.273755247 | 0.093173905 | 0.193740455 |
| 293,400 | 0.199836747 | 0.273932058 | 0.09348956 | 0.192086238 |
| 293,500 | 0.199818295 | 0.273976137 | 0.092418952 | 0.192494252 |
| 293,600 | 0.199675319 | 0.274203915 | 0.091813018 | 0.19281654 |
| 293,700 | 0.199598987 | 0.274164136 | 0.092389403 | 0.192363277 |

| | | | | |
|---|---|---|---|---|
| 293,800 | 0.199468983 | 0.273772392 | 0.092261018 | 0.192992705 |
| 293,900 | 0.199580932 | 0.274423746 | 0.093398389 | 0.193502461 |
| 294,000 | 0.199702331 | 0.274381716 | 0.093412576 | 0.193085877 |
| 294,100 | 0.199605605 | 0.274069957 | 0.093131889 | 0.192959655 |
| 294,200 | 0.199598999 | 0.273650304 | 0.093252202 | 0.192697782 |
| 294,300 | 0.19972827 | 0.274825861 | 0.093741591 | 0.193299604 |
| 294,400 | 0.19959424 | 0.274151835 | 0.092875718 | 0.19257055 |
| 294,500 | 0.199601226 | 0.273599265 | 0.094199395 | 0.193280286 |
| 294,600 | 0.19941745 | 0.273866038 | 0.093277517 | 0.192625325 |
| 294,700 | 0.199622961 | 0.273491936 | 0.093270497 | 0.192817209 |
| 294,800 | 0.199343175 | 0.272949489 | 0.092678967 | 0.192679448 |
| 294,900 | 0.199371661 | 0.27395173 | 0.094473849 | 0.193801868 |
| 295,000 | 0.199210252 | 0.272458351 | 0.094108494 | 0.192764332 |
| 295,100 | 0.199232408 | 0.27284607 | 0.094251399 | 0.192741934 |
| 295,200 | 0.199237158 | 0.273270328 | 0.094237356 | 0.192459644 |
| 295,300 | 0.199025696 | 0.27348511 | 0.093208076 | 0.192838994 |
| 295,400 | 0.199176653 | 0.272943769 | 0.093071081 | 0.193706818 |
| 295,500 | 0.19915959 | 0.272906593 | 0.092418042 | 0.193071696 |
| 295,600 | 0.199204196 | 0.272752361 | 0.093274764 | 0.193147964 |
| 295,700 | 0.199071317 | 0.27298506 | 0.094394338 | 0.192901638 |
| 295,800 | 0.199191303 | 0.273520652 | 0.093527138 | 0.193419544 |
| 295,900 | 0.199145612 | 0.273144849 | 0.093920181 | 0.193517915 |
| 296,000 | 0.199139364 | 0.273710813 | 0.093135249 | 0.193745143 |
| 296,100 | 0.199106089 | 0.273488917 | 0.093598825 | 0.19352998 |
| 296,200 | 0.198999971 | 0.273407795 | 0.092431302 | 0.194072449 |
| 296,300 | 0.199190173 | 0.272631491 | 0.093596155 | 0.193693404 |
| 296,400 | 0.199132502 | 0.272608201 | 0.093832018 | 0.19324747 |
| 296,500 | 0.198952968 | 0.271656279 | 0.093301843 | 0.193468987 |
| 296,600 | 0.199007115 | 0.272260102 | 0.093042505 | 0.193333223 |
| 296,700 | 0.198916584 | 0.272712119 | 0.093575362 | 0.193094839 |
| 296,800 | 0.198836115 | 0.272583284 | 0.093628601 | 0.193414446 |
| 296,900 | 0.198763207 | 0.27209271 | 0.092558167 | 0.193847097 |
| 297,000 | 0.198681721 | 0.272219822 | 0.092631961 | 0.193388453 |
| 297,100 | 0.198698406 | 0.271337171 | 0.09119117 | 0.193592144 |
| 297,200 | 0.198802476 | 0.271275413 | 0.092404484 | 0.193406948 |
| 297,300 | 0.198854921 | 0.271721336 | 0.092666118 | 0.193301903 |
| 297,400 | 0.198880989 | 0.273068323 | 0.092109548 | 0.192827869 |
| 297,500 | 0.198868966 | 0.27164531 | 0.092188004 | 0.193389696 |
| 297,600 | 0.198784384 | 0.271977158 | 0.092522467 | 0.193179451 |
| 297,700 | 0.198648727 | 0.27168106 | 0.0915042 | 0.193286578 |
| 297,800 | 0.198561372 | 0.272062883 | 0.091735979 | 0.194040322 |
| 297,900 | 0.198484947 | 0.271871863 | 0.090464582 | 0.193890557 |
| 298,000 | 0.19844022 | 0.270886489 | 0.090421062 | 0.193859394 |
| 298,100 | 0.198411516 | 0.270699385 | 0.090415246 | 0.194328358 |
| 298,200 | 0.198427622 | 0.2704506 | 0.089752105 | 0.194409964 |
| 298,300 | 0.198555361 | 0.27016778 | 0.089872136 | 0.1946294 |
| 298,400 | 0.198474325 | 0.270527448 | 0.090612315 | 0.193704183 |
| 298,500 | 0.198330764 | 0.270740217 | 0.090677977 | 0.193536946 |
| 298,600 | 0.198392117 | 0.270334198 | 0.091005691 | 0.194141206 |
| 298,700 | 0.198391079 | 0.271318526 | 0.090625715 | 0.193717105 |
| 298,800 | 0.198202187 | 0.270516737 | 0.090559058 | 0.193672613 |
| 298,900 | 0.198322681 | 0.270819332 | 0.090986272 | 0.193566544 |
| 299,000 | 0.198285719 | 0.270754695 | 0.090266638 | 0.193321118 |
| 299,100 | 0.198173212 | 0.26992871 | 0.089581498 | 0.19372175 |
| 299,200 | 0.198187449 | 0.270224615 | 0.090166424 | 0.192938651 |
| 299,300 | 0.198064094 | 0.269897108 | 0.090797123 | 0.19345442 |
| 299,400 | 0.198102537 | 0.269583593 | 0.090215043 | 0.193355552 |
| 299,500 | 0.197894281 | 0.27007243 | 0.089864955 | 0.193527483 |
| 299,600 | 0.197903323 | 0.269904487 | 0.090188075 | 0.194034435 |
| 299,700 | 0.197894158 | 0.269948653 | 0.090000971 | 0.193985339 |
| 299,800 | 0.197731786 | 0.269333859 | 0.090735094 | 0.194542827 |
| 299,900 | 0.197670128 | 0.26966925 | 0.089338377 | 0.194216585 |
| 300,000 | 0.197685882 | 0.269983104 | 0.090116366 | 0.193232168 |
| 300,100 | 0.197694358 | 0.269729552 | 0.089944586 | 0.193737502 |
| 300,200 | 0.197683568 | 0.268871077 | 0.09031553 | 0.194304143 |
| 300,300 | 0.197588769 | 0.269029994 | 0.090350652 | 0.194185838 |
| 300,400 | 0.197683174 | 0.26992741 | 0.090493503 | 0.193862278 |

| | | | | |
|---|---|---|---|---|
| 300,500 | 0.19762874 | 0.269943329 | 0.09079837 | 0.194070069 |
| 300,600 | 0.197620612 | 0.268598039 | 0.09099095 | 0.1934815 |
| 300,700 | 0.197480799 | 0.268259277 | 0.090267106 | 0.194003916 |
| 300,800 | 0.197489663 | 0.268991212 | 0.090584835 | 0.193589681 |
| 300,900 | 0.197589096 | 0.26977754 | 0.089768412 | 0.193545947 |
| 301,000 | 0.197414343 | 0.269416014 | 0.089008686 | 0.193637418 |
| 301,100 | 0.197366805 | 0.268857311 | 0.089491747 | 0.192652256 |
| 301,200 | 0.197306377 | 0.268928004 | 0.089330308 | 0.193214288 |
| 301,300 | 0.197454557 | 0.268697007 | 0.089496461 | 0.192941061 |
| 301,400 | 0.197418411 | 0.26803739 | 0.089559531 | 0.193411313 |
| 301,500 | 0.197296391 | 0.26821528 | 0.089765583 | 0.192581516 |
| 301,600 | 0.197385895 | 0.269076873 | 0.090157796 | 0.193615742 |
| 301,700 | 0.197227273 | 0.268595306 | 0.089526646 | 0.193375602 |
| 301,800 | 0.197341979 | 0.268329891 | 0.090534656 | 0.193195618 |
| 301,900 | 0.197130707 | 0.268857073 | 0.088266008 | 0.193294689 |
| 302,000 | 0.197067377 | 0.268831787 | 0.089014539 | 0.19280562 |
| 302,100 | 0.197150199 | 0.269011797 | 0.089754553 | 0.193249565 |
| 302,200 | 0.197110771 | 0.269052309 | 0.089544247 | 0.192351326 |
| 302,300 | 0.197326156 | 0.269605765 | 0.089280195 | 0.192954737 |
| 302,400 | 0.197115969 | 0.269376981 | 0.088825612 | 0.192364595 |
| 302,500 | 0.197150848 | 0.269175422 | 0.089751551 | 0.193360592 |
| 302,600 | 0.197185408 | 0.268645259 | 0.08966405 | 0.192974641 |
| 302,700 | 0.196859277 | 0.269136556 | 0.090081536 | 0.192870069 |
| 302,800 | 0.19695395 | 0.268656084 | 0.090255392 | 0.193153649 |
| 302,900 | 0.197026916 | 0.269468593 | 0.089002652 | 0.192436796 |
| 303,000 | 0.196846094 | 0.269470932 | 0.088317999 | 0.193086596 |
| 303,100 | 0.196884247 | 0.26992155 | 0.088208591 | 0.192857614 |
| 303,200 | 0.196805373 | 0.269358447 | 0.088912194 | 0.193108361 |
| 303,300 | 0.196862659 | 0.269736221 | 0.089576004 | 0.193334228 |
| 303,400 | 0.196823711 | 0.269329463 | 0.089561386 | 0.193779561 |
| 303,500 | 0.196785391 | 0.270148509 | 0.090091167 | 0.192905587 |
| 303,600 | 0.196805715 | 0.269236449 | 0.090200666 | 0.193003287 |
| 303,700 | 0.196817375 | 0.269499632 | 0.090364498 | 0.192551877 |
| 303,800 | 0.196778516 | 0.269414685 | 0.089695667 | 0.192138709 |
| 303,900 | 0.196670242 | 0.269690856 | 0.089129322 | 0.192311732 |
| 304,000 | 0.196607885 | 0.268557741 | 0.08973478 | 0.192790714 |
| 304,100 | 0.196671446 | 0.269098678 | 0.089528243 | 0.192656571 |
| 304,200 | 0.196704249 | 0.269502638 | 0.091330229 | 0.19286297 |
| 304,300 | 0.196598207 | 0.268097541 | 0.091302724 | 0.192981404 |
| 304,400 | 0.196599183 | 0.268823662 | 0.089128479 | 0.192774371 |
| 304,500 | 0.196520052 | 0.26832619 | 0.08968513 | 0.192848646 |
| 304,600 | 0.196533685 | 0.267393207 | 0.091280019 | 0.193036698 |
| 304,700 | 0.196451916 | 0.268145366 | 0.089666998 | 0.193168104 |
| 304,800 | 0.196482157 | 0.268726774 | 0.090202942 | 0.192909376 |
| 304,900 | 0.196329529 | 0.268308269 | 0.08995038 | 0.192825065 |
| 305,000 | 0.196220985 | 0.267084631 | 0.090164968 | 0.192195559 |
| 305,100 | 0.19619392 | 0.267402254 | 0.089855828 | 0.192539767 |
| 305,200 | 0.19612052 | 0.267231984 | 0.089444079 | 0.192734808 |
| 305,300 | 0.19630503 | 0.267961255 | 0.089497725 | 0.193262034 |
| 305,400 | 0.19624129 | 0.267318342 | 0.088834326 | 0.19253034 |
| 305,500 | 0.196094902 | 0.26770934 | 0.089546588 | 0.192477173 |
| 305,600 | 0.196036148 | 0.267635689 | 0.090154725 | 0.192388606 |
| 305,700 | 0.196160526 | 0.268480361 | 0.090591502 | 0.192105159 |
| 305,800 | 0.19612169 | 0.267889841 | 0.09161796 | 0.192373769 |
| 305,900 | 0.196089101 | 0.267431756 | 0.090796113 | 0.192842942 |
| 306,000 | 0.196038793 | 0.26721 | 0.091290488 | 0.192586863 |
| 306,100 | 0.195991892 | 0.267073056 | 0.090516561 | 0.1933068 |
| 306,200 | 0.196009126 | 0.266970673 | 0.091636318 | 0.192496757 |
| 306,300 | 0.195935514 | 0.267320915 | 0.090878461 | 0.193274334 |
| 306,400 | 0.195853743 | 0.267838238 | 0.092318268 | 0.192413644 |
| 306,500 | 0.195881958 | 0.267284819 | 0.090960484 | 0.192378928 |
| 306,600 | 0.195798154 | 0.266823437 | 0.090277778 | 0.19254222 |
| 306,700 | 0.195818242 | 0.266881225 | 0.090631024 | 0.192649187 |
| 306,800 | 0.19563555 | 0.267116035 | 0.091364448 | 0.192372869 |
| 306,900 | 0.195683911 | 0.267824379 | 0.090785201 | 0.192620128 |
| 307,000 | 0.19582401 | 0.267675943 | 0.090554017 | 0.192460782 |
| 307,100 | 0.195771437 | 0.267766809 | 0.091282499 | 0.193194856 |

| | | | | |
|---|---|---|---|---|
| 307,200 | 0.19576319 | 0.26728549 | 0.091338955 | 0.192830572 |
| 307,300 | 0.195618369 | 0.267073863 | 0.090610921 | 0.192749504 |
| 307,400 | 0.195890376 | 0.267845942 | 0.093653306 | 0.192527084 |
| 307,500 | 0.195655837 | 0.267146696 | 0.091351326 | 0.192256071 |
| 307,600 | 0.195680238 | 0.267485061 | 0.091104761 | 0.192431801 |
| 307,700 | 0.195647101 | 0.267853151 | 0.091647752 | 0.192334869 |
| 307,800 | 0.195603996 | 0.267906692 | 0.091271685 | 0.192079627 |
| 307,900 | 0.195693854 | 0.267926855 | 0.091563972 | 0.192249041 |
| 308,000 | 0.195559697 | 0.267373083 | 0.09135127 | 0.192189354 |
| 308,100 | 0.195696025 | 0.268619372 | 0.091167019 | 0.19255128 |
| 308,200 | 0.195667841 | 0.268413747 | 0.090683039 | 0.192249423 |
| 308,300 | 0.195558688 | 0.267272881 | 0.09022895 | 0.192701762 |
| 308,400 | 0.195564276 | 0.266684319 | 0.090929076 | 0.193104784 |
| 308,500 | 0.195778009 | 0.267864112 | 0.090623886 | 0.192914947 |
| 308,600 | 0.195627066 | 0.267741941 | 0.091057213 | 0.192663757 |
| 308,700 | 0.195740986 | 0.268270618 | 0.091198412 | 0.192257825 |
| 308,800 | 0.195563698 | 0.268050183 | 0.090608615 | 0.191391393 |
| 308,900 | 0.195456765 | 0.267076603 | 0.090010299 | 0.191457013 |
| 309,000 | 0.195325963 | 0.267845556 | 0.089651617 | 0.191485754 |
| 309,100 | 0.195268416 | 0.267520398 | 0.089594939 | 0.191656838 |
| 309,200 | 0.195427279 | 0.267938076 | 0.089514073 | 0.19242824 |
| 309,300 | 0.195217971 | 0.266748962 | 0.091889894 | 0.192157805 |
| 309,400 | 0.195133009 | 0.267110142 | 0.090080163 | 0.192547486 |
| 309,500 | 0.195166949 | 0.267316815 | 0.090312857 | 0.192415772 |
| 309,600 | 0.195262327 | 0.267388922 | 0.09015995 | 0.192308962 |
| 309,700 | 0.195354369 | 0.267674562 | 0.090699103 | 0.192312293 |
| 309,800 | 0.195204084 | 0.266589297 | 0.089031591 | 0.192125901 |
| 309,900 | 0.195115054 | 0.267215581 | 0.090114678 | 0.191802259 |
| 310,000 | 0.19517391 | 0.266677609 | 0.090726872 | 0.192264699 |
| 310,100 | 0.195126871 | 0.266851703 | 0.0917237 | 0.191582383 |
| 310,200 | 0.194926185 | 0.266685417 | 0.089507477 | 0.191830879 |
| 310,300 | 0.195082494 | 0.266927 | 0.090113634 | 0.192422072 |
| 310,400 | 0.194976202 | 0.26657537 | 0.089556194 | 0.19220975 |
| 310,500 | 0.194981437 | 0.266728069 | 0.089944923 | 0.192294823 |
| 310,600 | 0.194949867 | 0.266548925 | 0.090689904 | 0.192146937 |
| 310,700 | 0.194990151 | 0.26711504 | 0.088281585 | 0.192070387 |
| 310,800 | 0.194910389 | 0.26716574 | 0.090238646 | 0.19242286 |
| 310,900 | 0.194960923 | 0.267697122 | 0.090488809 | 0.192015924 |
| 311,000 | 0.194707624 | 0.267014638 | 0.089692526 | 0.192842769 |
| 311,100 | 0.194834117 | 0.266895102 | 0.089443882 | 0.191765262 |
| 311,200 | 0.194803062 | 0.266680988 | 0.089074059 | 0.193160261 |
| 311,300 | 0.19472024 | 0.266701313 | 0.090053177 | 0.192127795 |
| 311,400 | 0.194699292 | 0.268151227 | 0.089334087 | 0.192508259 |
| 311,500 | 0.194656929 | 0.267427166 | 0.089820127 | 0.192318394 |
| 311,600 | 0.194630519 | 0.268411226 | 0.08946475 | 0.192478314 |
| 311,700 | 0.194599715 | 0.268500304 | 0.088809442 | 0.192671485 |
| 311,800 | 0.194624481 | 0.267456115 | 0.089277817 | 0.192280227 |
| 311,900 | 0.194612553 | 0.266951422 | 0.089529362 | 0.191957077 |
| 312,000 | 0.194361866 | 0.267625619 | 0.088964465 | 0.192662181 |
| 312,100 | 0.194507452 | 0.266820766 | 0.089090323 | 0.192139754 |
| 312,200 | 0.194225643 | 0.266918302 | 0.088361683 | 0.191999501 |
| 312,300 | 0.194296319 | 0.266699488 | 0.088351787 | 0.191960649 |
| 312,400 | 0.19423646 | 0.26683345 | 0.087802657 | 0.192152795 |
| 312,500 | 0.194377687 | 0.267558221 | 0.088758362 | 0.192177297 |
| 312,600 | 0.194280124 | 0.268147212 | 0.087969329 | 0.192156889 |
| 312,700 | 0.194299265 | 0.267375911 | 0.087741588 | 0.191444202 |
| 312,800 | 0.194411906 | 0.267415343 | 0.088261173 | 0.192656706 |
| 312,900 | 0.194556731 | 0.268119229 | 0.088820887 | 0.192671374 |
| 313,000 | 0.194399285 | 0.267495877 | 0.089307837 | 0.191629295 |
| 313,100 | 0.194471621 | 0.268934122 | 0.088835097 | 0.192532501 |
| 313,200 | 0.194476916 | 0.268586399 | 0.08922713 | 0.192453635 |
| 313,300 | 0.194348598 | 0.267822094 | 0.089134681 | 0.192084905 |
| 313,400 | 0.194222449 | 0.267625065 | 0.089423537 | 0.191920469 |
| 313,500 | 0.194228579 | 0.267634737 | 0.090246889 | 0.191737057 |
| 313,600 | 0.194119253 | 0.267043883 | 0.089340237 | 0.191827009 |
| 313,700 | 0.193993776 | 0.266092832 | 0.089978052 | 0.192942853 |
| 313,800 | 0.193928443 | 0.266607463 | 0.089824772 | 0.192600004 |

| | | | | |
|---|---|---|---|---|
| 313,900 | 0.193899306 | 0.266970242 | 0.090156128 | 0.192899587 |
| 314,000 | 0.193828792 | 0.267181465 | 0.089299836 | 0.192325058 |
| 314,100 | 0.193839714 | 0.267025009 | 0.08968275 | 0.192137685 |
| 314,200 | 0.193890162 | 0.266530815 | 0.089518671 | 0.192431328 |
| 314,300 | 0.193966403 | 0.266997142 | 0.088607043 | 0.192482057 |
| 314,400 | 0.193842923 | 0.266310105 | 0.088714216 | 0.19222375 |
| 314,500 | 0.193792872 | 0.26711466 | 0.089600297 | 0.192707432 |
| 314,600 | 0.193699197 | 0.266877704 | 0.09011726 | 0.191738864 |
| 314,700 | 0.193695046 | 0.267136739 | 0.088619458 | 0.191801641 |
| 314,800 | 0.193715987 | 0.267077026 | 0.089264482 | 0.192104679 |
| 314,900 | 0.193452792 | 0.265915629 | 0.089598572 | 0.19155677 |
| 315,000 | 0.193349862 | 0.265882732 | 0.088676984 | 0.191316912 |
| 315,100 | 0.193502035 | 0.265878065 | 0.088656496 | 0.191647744 |
| 315,200 | 0.193478205 | 0.26621165 | 0.089158569 | 0.191049855 |
| 315,300 | 0.193451611 | 0.265769687 | 0.088928089 | 0.191620985 |
| 315,400 | 0.193367191 | 0.264931419 | 0.089254281 | 0.191243465 |
| 315,500 | 0.193191832 | 0.264950306 | 0.089210194 | 0.191275963 |
| 315,600 | 0.193161308 | 0.26475405 | 0.088751816 | 0.191196641 |
| 315,700 | 0.193164329 | 0.264336785 | 0.089025581 | 0.190918281 |
| 315,800 | 0.193341793 | 0.264433045 | 0.088852248 | 0.191520056 |
| 315,900 | 0.193276566 | 0.264937223 | 0.088502885 | 0.191394314 |
| 316,000 | 0.193222702 | 0.265229536 | 0.088908651 | 0.191259104 |
| 316,100 | 0.193235949 | 0.265422841 | 0.088583731 | 0.191072573 |
| 316,200 | 0.193201108 | 0.265609388 | 0.08876576 | 0.190758196 |
| 316,300 | 0.193197053 | 0.266440663 | 0.089174061 | 0.190974778 |
| 316,400 | 0.19323676 | 0.266033958 | 0.088042256 | 0.192124006 |
| 316,500 | 0.193201876 | 0.266078508 | 0.089721525 | 0.191575068 |
| 316,600 | 0.193176949 | 0.266200072 | 0.088232153 | 0.191284795 |
| 316,700 | 0.193286884 | 0.265620758 | 0.088519628 | 0.191406848 |
| 316,800 | 0.19306258 | 0.264586837 | 0.089645658 | 0.191725057 |
| 316,900 | 0.193070556 | 0.264998647 | 0.088448162 | 0.192145105 |
| 317,000 | 0.193185994 | 0.264636961 | 0.089267714 | 0.192108371 |
| 317,100 | 0.193101129 | 0.265256979 | 0.088939825 | 0.192559687 |
| 317,200 | 0.193016278 | 0.264941208 | 0.08882668 | 0.192188147 |
| 317,300 | 0.192984932 | 0.265009728 | 0.087784609 | 0.191852034 |
| 317,400 | 0.192808747 | 0.265537122 | 0.087405064 | 0.191839663 |
| 317,500 | 0.192833963 | 0.264799216 | 0.087452772 | 0.19183539 |
| 317,600 | 0.192754446 | 0.265930027 | 0.088623893 | 0.192276278 |
| 317,700 | 0.192798929 | 0.264846914 | 0.088643295 | 0.191717748 |
| 317,800 | 0.192782637 | 0.264993329 | 0.089326526 | 0.191953257 |
| 317,900 | 0.192681279 | 0.265073038 | 0.087771294 | 0.191693931 |
| 318,000 | 0.19278658 | 0.265000294 | 0.088119386 | 0.191523332 |
| 318,100 | 0.192534784 | 0.264558871 | 0.088047778 | 0.190490446 |
| 318,200 | 0.192639657 | 0.26479054 | 0.089003844 | 0.191202777 |
| 318,300 | 0.192565899 | 0.26511051 | 0.088682711 | 0.191536735 |
| 318,400 | 0.192524882 | 0.264709695 | 0.087175732 | 0.191599254 |
| 318,500 | 0.192428573 | 0.265133266 | 0.08811215 | 0.19170154 |
| 318,600 | 0.192315757 | 0.264619958 | 0.088232842 | 0.191021109 |
| 318,700 | 0.192402643 | 0.264863589 | 0.089167323 | 0.191843659 |
| 318,800 | 0.192290521 | 0.26455398 | 0.088438306 | 0.191614268 |
| 318,900 | 0.192464383 | 0.265028741 | 0.088643659 | 0.191132222 |
| 319,000 | 0.192372633 | 0.264260846 | 0.087703168 | 0.19082767 |
| 319,100 | 0.192435615 | 0.264686985 | 0.08730991 | 0.190678852 |
| 319,200 | 0.192349073 | 0.264900505 | 0.088024301 | 0.191708109 |
| 319,300 | 0.192258912 | 0.265219061 | 0.088698788 | 0.192275383 |
| 319,400 | 0.192274129 | 0.264841364 | 0.088313729 | 0.190888099 |
| 319,500 | 0.192365676 | 0.264167278 | 0.089326543 | 0.191801049 |
| 319,600 | 0.192139451 | 0.263851312 | 0.089031895 | 0.191631357 |
| 319,700 | 0.192109085 | 0.26313697 | 0.088429867 | 0.191052761 |
| 319,800 | 0.192135255 | 0.263368236 | 0.088257485 | 0.189903525 |
| 319,900 | 0.19214871 | 0.263277243 | 0.08792718 | 0.190703344 |
| 320,000 | 0.192264124 | 0.264063854 | 0.087919148 | 0.19076373 |
| 320,100 | 0.1921016 | 0.263571232 | 0.087955486 | 0.190662799 |
| 320,200 | 0.192056181 | 0.26291446 | 0.086712146 | 0.191792552 |
| 320,300 | 0.192190693 | 0.262961298 | 0.087085336 | 0.191546839 |
| 320,400 | 0.19200946 | 0.262985574 | 0.087779985 | 0.191729845 |
| 320,500 | 0.192042125 | 0.262154003 | 0.086313984 | 0.191389395 |

| | | | | |
|---|---|---|---|---|
| 320,600 | 0.191835337 | 0.262022642 | 0.087122021 | 0.190805072 |
| 320,700 | 0.191981653 | 0.262319775 | 0.086770789 | 0.191600389 |
| 320,800 | 0.191831484 | 0.261470285 | 0.087554788 | 0.190696802 |
| 320,900 | 0.191846391 | 0.261625554 | 0.087324646 | 0.191473177 |
| 321,000 | 0.191891259 | 0.262125521 | 0.088199947 | 0.190454649 |
| 321,100 | 0.192036672 | 0.262498176 | 0.087800123 | 0.190940084 |
| 321,200 | 0.192075877 | 0.261976186 | 0.087314484 | 0.191334369 |
| 321,300 | 0.191961388 | 0.261811167 | 0.086677421 | 0.191634777 |
| 321,400 | 0.191974427 | 0.260841679 | 0.087079949 | 0.191513172 |
| 321,500 | 0.191898884 | 0.260542853 | 0.087290582 | 0.191214302 |
| 321,600 | 0.191907643 | 0.26184671 | 0.087309672 | 0.191592526 |
| 321,700 | 0.191811675 | 0.261209467 | 0.086940878 | 0.190878663 |
| 321,800 | 0.191883459 | 0.262613259 | 0.087244511 | 0.190871173 |
| 321,900 | 0.19203624 | 0.261897101 | 0.088350478 | 0.191378539 |
| 322,000 | 0.191949497 | 0.26155926 | 0.088496926 | 0.191688211 |
| 322,100 | 0.191699308 | 0.261277618 | 0.087894165 | 0.190942842 |
| 322,200 | 0.191870214 | 0.261537931 | 0.087343013 | 0.19056156 |
| 322,300 | 0.19180465 | 0.261540729 | 0.087939339 | 0.190319163 |
| 322,400 | 0.19158625 | 0.261575013 | 0.087742249 | 0.190757288 |
| 322,500 | 0.191640357 | 0.261339942 | 0.088079039 | 0.190497582 |
| 322,600 | 0.191517548 | 0.261501658 | 0.087524599 | 0.190748706 |
| 322,700 | 0.191429063 | 0.261475161 | 0.088531951 | 0.189920972 |
| 322,800 | 0.191400159 | 0.260874605 | 0.088250412 | 0.19045385 |
| 322,900 | 0.191424345 | 0.261110893 | 0.087985465 | 0.190687318 |
| 323,000 | 0.191421055 | 0.260868572 | 0.087173507 | 0.190328607 |
| 323,100 | 0.191429054 | 0.26143022 | 0.087212783 | 0.190668148 |
| 323,200 | 0.191473881 | 0.262035283 | 0.086690831 | 0.19026606 |
| 323,300 | 0.191300407 | 0.261846677 | 0.088090243 | 0.19090897 |
| 323,400 | 0.191354475 | 0.261973626 | 0.088317727 | 0.190549018 |
| 323,500 | 0.191418583 | 0.261693537 | 0.087329907 | 0.189746716 |
| 323,600 | 0.191419817 | 0.262317698 | 0.087444538 | 0.18987392 |
| 323,700 | 0.191423349 | 0.262943335 | 0.087439963 | 0.190613095 |
| 323,800 | 0.191452147 | 0.262760203 | 0.087811643 | 0.19058548 |
| 323,900 | 0.191352481 | 0.262196764 | 0.087370031 | 0.190283688 |
| 324,000 | 0.191173706 | 0.261842957 | 0.088007638 | 0.19097482 |
| 324,100 | 0.191009455 | 0.261613778 | 0.088330212 | 0.18960742 |
| 324,200 | 0.191010293 | 0.26210574 | 0.087633967 | 0.189794065 |
| 324,300 | 0.191288058 | 0.26099747 | 0.087744159 | 0.190332069 |
| 324,400 | 0.191091235 | 0.26103285 | 0.08773953 | 0.190559507 |
| 324,500 | 0.191185622 | 0.260867614 | 0.088691584 | 0.19096707 |
| 324,600 | 0.191147603 | 0.261145594 | 0.089073154 | 0.18997336 |
| 324,700 | 0.191201806 | 0.261357036 | 0.087747801 | 0.190453569 |
| 324,800 | 0.19130874 | 0.261706647 | 0.088646252 | 0.190736455 |
| 324,900 | 0.191154757 | 0.261224164 | 0.087832598 | 0.191256891 |
| 325,000 | 0.191126372 | 0.261393775 | 0.088397008 | 0.191202911 |
| 325,100 | 0.191167742 | 0.261600094 | 0.088660074 | 0.190353561 |
| 325,200 | 0.191009678 | 0.260355551 | 0.088189017 | 0.190132501 |
| 325,300 | 0.190938485 | 0.261124161 | 0.087682843 | 0.190856596 |
| 325,400 | 0.191059454 | 0.260901883 | 0.088272989 | 0.190512809 |
| 325,500 | 0.191066507 | 0.260896942 | 0.088392752 | 0.190824042 |
| 325,600 | 0.1911381 | 0.26122649 | 0.08776849 | 0.19052352 |
| 325,700 | 0.191092985 | 0.261476987 | 0.08679615 | 0.191148351 |
| 325,800 | 0.191039223 | 0.261170179 | 0.088037171 | 0.191498586 |
| 325,900 | 0.191054387 | 0.261226272 | 0.087220337 | 0.19173077 |
| 326,000 | 0.191155982 | 0.261721756 | 0.087701411 | 0.191531711 |
| 326,100 | 0.190859175 | 0.261382712 | 0.087668461 | 0.190752688 |
| 326,200 | 0.191114101 | 0.26165611 | 0.085916518 | 0.191618889 |
| 326,300 | 0.191055366 | 0.260962921 | 0.087479555 | 0.190930977 |
| 326,400 | 0.191004949 | 0.260729587 | 0.087003102 | 0.190821341 |
| 326,500 | 0.190960363 | 0.260814685 | 0.087888449 | 0.190328937 |
| 326,600 | 0.190995233 | 0.260840365 | 0.08730056 | 0.190670513 |
| 326,700 | 0.191082238 | 0.260747627 | 0.08779683 | 0.190023886 |
| 326,800 | 0.190988558 | 0.260756805 | 0.087706615 | 0.189475057 |
| 326,900 | 0.191142185 | 0.26076905 | 0.087094004 | 0.190097468 |
| 327,000 | 0.190913879 | 0.259709885 | 0.087980377 | 0.18955827 |
| 327,100 | 0.190954716 | 0.259752306 | 0.088500114 | 0.190106273 |
| 327,200 | 0.190912434 | 0.260416552 | 0.088635591 | 0.190554684 |

| | | | | |
|---|---|---|---|---|
| 327,300 | 0.190849673 | 0.26078598 | 0.088623784 | 0.190528413 |
| 327,400 | 0.190757723 | 0.260351534 | 0.088348256 | 0.190642231 |
| 327,500 | 0.190884986 | 0.260274806 | 0.088134119 | 0.190482406 |
| 327,600 | 0.19092326 | 0.260547284 | 0.088321397 | 0.190587267 |
| 327,700 | 0.190892132 | 0.260626454 | 0.08757607 | 0.189662491 |
| 327,800 | 0.190737148 | 0.260031787 | 0.086998293 | 0.190080069 |
| 327,900 | 0.190544948 | 0.260637159 | 0.087210178 | 0.189539904 |
| 328,000 | 0.190643117 | 0.260145796 | 0.0876247 | 0.189675385 |
| 328,100 | 0.19066394 | 0.260504822 | 0.087171104 | 0.189542387 |
| 328,200 | 0.190607 | 0.260435034 | 0.087218053 | 0.189429078 |
| 328,300 | 0.190705402 | 0.261111418 | 0.086999663 | 0.189753461 |
| 328,400 | 0.190520604 | 0.260163675 | 0.087781415 | 0.189157552 |
| 328,500 | 0.190601776 | 0.260839104 | 0.087875782 | 0.188851249 |
| 328,600 | 0.190463785 | 0.261004035 | 0.087577237 | 0.18966312 |
| 328,700 | 0.190272887 | 0.260139807 | 0.087571751 | 0.188894794 |
| 328,800 | 0.190456612 | 0.261128387 | 0.089158316 | 0.188962155 |
| 328,900 | 0.190343804 | 0.259944729 | 0.089007278 | 0.188821141 |
| 329,000 | 0.190443984 | 0.260526415 | 0.089060909 | 0.189069856 |
| 329,100 | 0.190314132 | 0.261219266 | 0.088292839 | 0.189414053 |
| 329,200 | 0.190183275 | 0.260806297 | 0.088655717 | 0.188597977 |
| 329,300 | 0.190256076 | 0.260390821 | 0.08744959 | 0.188371887 |
| 329,400 | 0.190245729 | 0.260463446 | 0.089088486 | 0.188086804 |
| 329,500 | 0.190152797 | 0.26071123 | 0.088863312 | 0.18800272 |
| 329,600 | 0.190146818 | 0.261254165 | 0.089576406 | 0.18776945 |
| 329,700 | 0.190201306 | 0.260754363 | 0.088696958 | 0.187636379 |
| 329,800 | 0.19007302 | 0.259693799 | 0.088734783 | 0.187449227 |
| 329,900 | 0.190024178 | 0.259682404 | 0.088241861 | 0.187337003 |
| 330,000 | 0.189960819 | 0.260456024 | 0.087852111 | 0.188069966 |
| 330,100 | 0.189981188 | 0.259918238 | 0.089405235 | 0.187341216 |
| 330,200 | 0.189861701 | 0.259414901 | 0.088385788 | 0.187491897 |
| 330,300 | 0.190004988 | 0.259828904 | 0.088570066 | 0.187659109 |
| 330,400 | 0.189986546 | 0.260265359 | 0.088772287 | 0.187291907 |
| 330,500 | 0.189886516 | 0.259872819 | 0.088226809 | 0.18753094 |
| 330,600 | 0.189876319 | 0.259423435 | 0.088545741 | 0.187412286 |
| 330,700 | 0.189778565 | 0.258220042 | 0.089533034 | 0.188042029 |
| 330,800 | 0.189749797 | 0.258946495 | 0.089558839 | 0.187279952 |
| 330,900 | 0.189745594 | 0.259196022 | 0.088941976 | 0.187569158 |
| 331,000 | 0.189860592 | 0.259791577 | 0.090049125 | 0.188444452 |
| 331,100 | 0.189820524 | 0.259351524 | 0.089263717 | 0.188439012 |
| 331,200 | 0.189843584 | 0.260090904 | 0.089326687 | 0.188974123 |
| 331,300 | 0.189652616 | 0.259176072 | 0.089509283 | 0.188435412 |
| 331,400 | 0.189527487 | 0.258722565 | 0.088690641 | 0.188718756 |
| 331,500 | 0.189612952 | 0.258996207 | 0.088278343 | 0.188485132 |
| 331,600 | 0.189403559 | 0.259366174 | 0.088129811 | 0.187977955 |
| 331,700 | 0.189584965 | 0.258819406 | 0.088366344 | 0.188266012 |
| 331,800 | 0.189620014 | 0.258644244 | 0.089451847 | 0.188769246 |
| 331,900 | 0.189465598 | 0.258768716 | 0.088864042 | 0.188080574 |
| 332,000 | 0.189450628 | 0.258493587 | 0.089726024 | 0.18840857 |
| 332,100 | 0.18938639 | 0.258858881 | 0.088776074 | 0.188165968 |
| 332,200 | 0.189360593 | 0.258051715 | 0.089588534 | 0.187797606 |
| 332,300 | 0.189215553 | 0.257872826 | 0.089280942 | 0.187613566 |
| 332,400 | 0.189328101 | 0.258948861 | 0.089811636 | 0.187207223 |
| 332,500 | 0.189238078 | 0.259254637 | 0.089429025 | 0.188180965 |
| 332,600 | 0.189301942 | 0.258936981 | 0.089577872 | 0.18776589 |
| 332,700 | 0.189222743 | 0.259282097 | 0.089632002 | 0.187438292 |
| 332,800 | 0.18924695 | 0.258199164 | 0.089934972 | 0.18810572 |
| 332,900 | 0.18900499 | 0.258099168 | 0.089825202 | 0.187783594 |
| 333,000 | 0.18924192 | 0.258766145 | 0.089841636 | 0.187959356 |
| 333,100 | 0.189041584 | 0.258038483 | 0.09052898 | 0.187687935 |
| 333,200 | 0.189124728 | 0.257757022 | 0.089660031 | 0.187702103 |
| 333,300 | 0.18904106 | 0.258721439 | 0.089994099 | 0.188047927 |
| 333,400 | 0.189074407 | 0.257997668 | 0.089890495 | 0.187736797 |
| 333,500 | 0.189163284 | 0.257619356 | 0.089173391 | 0.187794028 |
| 333,600 | 0.18913601 | 0.257789646 | 0.089699884 | 0.188080205 |
| 333,700 | 0.189058548 | 0.257738532 | 0.090384167 | 0.187660295 |
| 333,800 | 0.18909851 | 0.257928177 | 0.089936043 | 0.188570595 |
| 333,900 | 0.189009433 | 0.258209899 | 0.090182917 | 0.188032727 |

| | | | | |
|---|---|---|---|---|
| 334,000 | 0.188931342 | 0.257581926 | 0.091021109 | 0.187448793 |
| 334,100 | 0.188956119 | 0.257857303 | 0.090723113 | 0.1873012 |
| 334,200 | 0.189052586 | 0.258951223 | 0.090239257 | 0.187331799 |
| 334,300 | 0.188837169 | 0.258431863 | 0.090527262 | 0.187799238 |
| 334,400 | 0.188940767 | 0.258307676 | 0.090103786 | 0.188133715 |
| 334,500 | 0.188776267 | 0.258626395 | 0.090113623 | 0.187090182 |
| 334,600 | 0.188728885 | 0.257330046 | 0.090776943 | 0.187995323 |
| 334,700 | 0.188686552 | 0.25730697 | 0.090158547 | 0.187482627 |
| 334,800 | 0.188880303 | 0.258101123 | 0.089477409 | 0.187704252 |
| 334,900 | 0.188867087 | 0.258074281 | 0.090567322 | 0.187932861 |
| 335,000 | 0.188738705 | 0.257421714 | 0.09008375 | 0.188264177 |
| 335,100 | 0.188651023 | 0.25736251 | 0.08939155 | 0.187785603 |
| 335,200 | 0.188616016 | 0.258027542 | 0.089307602 | 0.18678821 |
| 335,300 | 0.188741248 | 0.257508286 | 0.089552422 | 0.187445444 |
| 335,400 | 0.188722699 | 0.257793587 | 0.089081131 | 0.187015075 |
| 335,500 | 0.188637892 | 0.257402715 | 0.089343803 | 0.187827181 |
| 335,600 | 0.188781249 | 0.258900049 | 0.089387086 | 0.187053518 |
| 335,700 | 0.188831149 | 0.258148166 | 0.088581351 | 0.187284197 |
| 335,800 | 0.18843773 | 0.257120898 | 0.088199492 | 0.187269967 |
| 335,900 | 0.18869126 | 0.257269805 | 0.088018419 | 0.187111373 |
| 336,000 | 0.188511635 | 0.256947771 | 0.087185697 | 0.186250666 |
| 336,100 | 0.188463581 | 0.256464402 | 0.088338828 | 0.186700292 |
| 336,200 | 0.188540157 | 0.256385863 | 0.087667056 | 0.186248182 |
| 336,300 | 0.188534885 | 0.257298288 | 0.088180276 | 0.186183535 |
| 336,400 | 0.18834776 | 0.257140404 | 0.087394395 | 0.186388305 |
| 336,500 | 0.188464643 | 0.256603637 | 0.088274095 | 0.187076707 |
| 336,600 | 0.188371131 | 0.256381776 | 0.088745108 | 0.186360176 |
| 336,700 | 0.188459592 | 0.257358595 | 0.089282871 | 0.186857183 |
| 336,800 | 0.188546154 | 0.257775747 | 0.087926369 | 0.187430802 |
| 336,900 | 0.188601884 | 0.258484969 | 0.08832183 | 0.187717626 |
| 337,000 | 0.188458324 | 0.257639912 | 0.087506452 | 0.186142904 |
| 337,100 | 0.188354144 | 0.257601288 | 0.0873968 | 0.18633701 |
| 337,200 | 0.188268715 | 0.257513269 | 0.08770609 | 0.186553258 |
| 337,300 | 0.188445432 | 0.257935952 | 0.087868557 | 0.18686489 |
| 337,400 | 0.188450977 | 0.258083232 | 0.087286648 | 0.186166596 |
| 337,500 | 0.188298704 | 0.257248864 | 0.087897131 | 0.185626004 |
| 337,600 | 0.188360228 | 0.257290546 | 0.086432094 | 0.186551335 |
| 337,700 | 0.188212181 | 0.257051562 | 0.087147875 | 0.186424922 |
| 337,800 | 0.188220772 | 0.25669061 | 0.087278897 | 0.186051478 |
| 337,900 | 0.18833827 | 0.25703522 | 0.087728383 | 0.18580784 |
| 338,000 | 0.188345294 | 0.257039253 | 0.087221149 | 0.186192224 |
| 338,100 | 0.188247595 | 0.256579605 | 0.087378827 | 0.186925357 |
| 338,200 | 0.188241488 | 0.256646922 | 0.087513986 | 0.186993828 |
| 338,300 | 0.188150544 | 0.256404095 | 0.086692695 | 0.186343244 |
| 338,400 | 0.188144365 | 0.256617605 | 0.087277459 | 0.186481124 |
| 338,500 | 0.188230135 | 0.257162515 | 0.088130882 | 0.187502807 |
| 338,600 | 0.188176061 | 0.255568936 | 0.088496235 | 0.18794331 |
| 338,700 | 0.188056826 | 0.255865715 | 0.087251961 | 0.187539871 |
| 338,800 | 0.18805506 | 0.256042509 | 0.088432965 | 0.18666826 |
| 338,900 | 0.1880269 | 0.256371704 | 0.088807293 | 0.186923708 |
| 339,000 | 0.188062515 | 0.255965361 | 0.088330226 | 0.187003276 |
| 339,100 | 0.188018441 | 0.256413024 | 0.088720607 | 0.186835733 |
| 339,200 | 0.187959786 | 0.255702174 | 0.089554398 | 0.186493269 |
| 339,300 | 0.187992172 | 0.256316603 | 0.089253862 | 0.186454818 |
| 339,400 | 0.187978533 | 0.255335895 | 0.088687279 | 0.186671452 |
| 339,500 | 0.187921663 | 0.25573815 | 0.089578864 | 0.186273474 |
| 339,600 | 0.187941395 | 0.255909491 | 0.088293762 | 0.186502215 |
| 339,700 | 0.187754159 | 0.255675725 | 0.08869918 | 0.186507283 |
| 339,800 | 0.187871499 | 0.254945463 | 0.08771691 | 0.186513306 |
| 339,900 | 0.187828072 | 0.255771605 | 0.088834365 | 0.18622323 |
| 340,000 | 0.187695906 | 0.255028531 | 0.088724523 | 0.186590066 |
| 340,100 | 0.187734911 | 0.255097224 | 0.088397293 | 0.187076034 |
| 340,200 | 0.187721376 | 0.255362991 | 0.088166926 | 0.18665598 |
| 340,300 | 0.187735982 | 0.254834586 | 0.088862707 | 0.186049534 |
| 340,400 | 0.187743982 | 0.254772307 | 0.08824694 | 0.185978763 |
| 340,500 | 0.187679813 | 0.254172771 | 0.088353299 | 0.187657296 |
| 340,600 | 0.187644928 | 0.254404168 | 0.089108028 | 0.186123046 |

| | | | | |
|---|---|---|---|---|
| 340,700 | 0.187635162 | 0.255620685 | 0.088525737 | 0.186021299 |
| 340,800 | 0.187693349 | 0.255544528 | 0.087468586 | 0.186418611 |
| 340,900 | 0.187710326 | 0.255824631 | 0.087738143 | 0.186336254 |
| 341,000 | 0.187780263 | 0.254155484 | 0.089024782 | 0.186543996 |
| 341,100 | 0.187618677 | 0.254340592 | 0.087476737 | 0.185934384 |
| 341,200 | 0.187730204 | 0.254813622 | 0.087983276 | 0.18639351 |
| 341,300 | 0.187683286 | 0.255540941 | 0.089252428 | 0.185616683 |
| 341,400 | 0.187479319 | 0.254515209 | 0.088143159 | 0.185444243 |
| 341,500 | 0.187658302 | 0.254112697 | 0.088494688 | 0.184952873 |
| 341,600 | 0.187476041 | 0.254643212 | 0.088233528 | 0.185457135 |
| 341,700 | 0.187453752 | 0.254425969 | 0.087797657 | 0.185087236 |
| 341,800 | 0.18754291 | 0.254440876 | 0.087671762 | 0.18512764 |
| 341,900 | 0.187469046 | 0.254009867 | 0.087925513 | 0.185190649 |
| 342,000 | 0.187436482 | 0.253918684 | 0.088314609 | 0.184838416 |
| 342,100 | 0.187493373 | 0.254259073 | 0.088523687 | 0.184492518 |
| 342,200 | 0.187291402 | 0.254022642 | 0.088074471 | 0.184183006 |
| 342,300 | 0.187432361 | 0.254162787 | 0.08783206 | 0.18525094 |
| 342,400 | 0.187251368 | 0.254194049 | 0.08803231 | 0.18442762 |
| 342,500 | 0.187350304 | 0.254059736 | 0.087003813 | 0.18459957 |
| 342,600 | 0.187255198 | 0.254903742 | 0.087651169 | 0.185457816 |
| 342,700 | 0.187279787 | 0.254161732 | 0.087640809 | 0.185479458 |
| 342,800 | 0.187337941 | 0.254928887 | 0.087937581 | 0.184605766 |
| 342,900 | 0.187246736 | 0.255542977 | 0.087677792 | 0.184730974 |
| 343,000 | 0.187394317 | 0.254480316 | 0.088227841 | 0.184506243 |
| 343,100 | 0.187269836 | 0.254816875 | 0.088044462 | 0.184610854 |
| 343,200 | 0.18723518 | 0.254341796 | 0.08785074 | 0.185457011 |
| 343,300 | 0.187233995 | 0.254385098 | 0.087975757 | 0.185298356 |
| 343,400 | 0.187251726 | 0.254452337 | 0.08785001 | 0.185210595 |
| 343,500 | 0.187062725 | 0.25408558 | 0.087988931 | 0.184897885 |
| 343,600 | 0.187074561 | 0.253527222 | 0.087361663 | 0.185717485 |
| 343,700 | 0.186897408 | 0.254540954 | 0.087765259 | 0.18478649 |
| 343,800 | 0.18681603 | 0.254373199 | 0.087947331 | 0.184692395 |
| 343,900 | 0.187010812 | 0.254623707 | 0.088085287 | 0.18495482 |
| 344,000 | 0.186927097 | 0.253795808 | 0.087695622 | 0.185191597 |
| 344,100 | 0.186773219 | 0.254182203 | 0.087665454 | 0.185328832 |
| 344,200 | 0.186683126 | 0.253873415 | 0.086865418 | 0.185350045 |
| 344,300 | 0.186683086 | 0.25478816 | 0.087050642 | 0.184450249 |
| 344,400 | 0.186726518 | 0.254460712 | 0.087122051 | 0.184758019 |
| 344,500 | 0.186733645 | 0.254156532 | 0.087143889 | 0.184977947 |
| 344,600 | 0.186760974 | 0.254144832 | 0.086366379 | 0.185892288 |
| 344,700 | 0.18682549 | 0.255215043 | 0.086831107 | 0.185006399 |
| 344,800 | 0.186930382 | 0.255264709 | 0.086619923 | 0.185678486 |
| 344,900 | 0.186726375 | 0.254795411 | 0.087532075 | 0.185746576 |
| 345,000 | 0.18666306 | 0.255161314 | 0.086793025 | 0.185193925 |
| 345,100 | 0.186713065 | 0.25586881 | 0.086853106 | 0.185508839 |
| 345,200 | 0.186666837 | 0.255346269 | 0.086635895 | 0.185581594 |
| 345,300 | 0.186580778 | 0.255872559 | 0.087083028 | 0.185312006 |
| 345,400 | 0.186630917 | 0.255275637 | 0.086577373 | 0.185795263 |
| 345,500 | 0.186447867 | 0.254730303 | 0.086990569 | 0.185681559 |
| 345,600 | 0.186480882 | 0.255640856 | 0.086789802 | 0.18573021 |
| 345,700 | 0.186385444 | 0.254634882 | 0.086265177 | 0.18544422 |
| 345,800 | 0.18644367 | 0.254575588 | 0.08663418 | 0.184145101 |
| 345,900 | 0.186533009 | 0.254519748 | 0.086853866 | 0.185185678 |
| 346,000 | 0.186581167 | 0.253382092 | 0.086757005 | 0.18490136 |
| 346,100 | 0.186712841 | 0.254341542 | 0.086321212 | 0.184942319 |
| 346,200 | 0.186407969 | 0.253959797 | 0.086537419 | 0.184889921 |
| 346,300 | 0.186415761 | 0.254032483 | 0.086161038 | 0.184467947 |
| 346,400 | 0.186391499 | 0.254225547 | 0.085088612 | 0.185282478 |
| 346,500 | 0.186455969 | 0.254575497 | 0.085550198 | 0.184412521 |
| 346,600 | 0.186637807 | 0.255964778 | 0.085520951 | 0.184436101 |
| 346,700 | 0.186699167 | 0.254715229 | 0.08683641 | 0.185470004 |
| 346,800 | 0.186568246 | 0.255413708 | 0.086835564 | 0.185123668 |
| 346,900 | 0.186511557 | 0.254653401 | 0.085707996 | 0.185482953 |
| 347,000 | 0.186630295 | 0.255187887 | 0.085615646 | 0.184928812 |
| 347,100 | 0.186594202 | 0.255289017 | 0.085708717 | 0.184854277 |
| 347,200 | 0.186538678 | 0.254945415 | 0.085396136 | 0.185207677 |
| 347,300 | 0.186514571 | 0.255490814 | 0.085534135 | 0.185127197 |

| | | | | |
|---|---|---|---|---|
| 347,400 | 0.186404098 | 0.254965621 | 0.085920014 | 0.184794726 |
| 347,500 | 0.186452469 | 0.253868382 | 0.085806291 | 0.185274012 |
| 347,600 | 0.18636644 | 0.253845016 | 0.085139593 | 0.185777665 |
| 347,700 | 0.186359751 | 0.253752398 | 0.084486576 | 0.185576521 |
| 347,800 | 0.186236467 | 0.254203013 | 0.085071777 | 0.184667661 |
| 347,900 | 0.186311926 | 0.254877084 | 0.084833056 | 0.184471842 |
| 348,000 | 0.186223844 | 0.254218501 | 0.083877129 | 0.184466199 |
| 348,100 | 0.18615529 | 0.253900045 | 0.085039512 | 0.184777211 |
| 348,200 | 0.186129084 | 0.253511377 | 0.083586884 | 0.18482053 |
| 348,300 | 0.186146797 | 0.254211578 | 0.084371866 | 0.184141591 |
| 348,400 | 0.18606947 | 0.25422603 | 0.08430038 | 0.184270932 |
| 348,500 | 0.186138978 | 0.254606779 | 0.084792158 | 0.183947562 |
| 348,600 | 0.186147064 | 0.254435188 | 0.084266968 | 0.184325516 |
| 348,700 | 0.186065747 | 0.254863337 | 0.084676057 | 0.183987605 |
| 348,800 | 0.186006141 | 0.253374418 | 0.08518856 | 0.183917821 |
| 348,900 | 0.186075092 | 0.253980299 | 0.084881873 | 0.184254224 |
| 349,000 | 0.185796868 | 0.253492564 | 0.083325367 | 0.182821272 |
| 349,100 | 0.185978342 | 0.253536175 | 0.083392724 | 0.183349382 |
| 349,200 | 0.185840991 | 0.253213327 | 0.083266633 | 0.183259043 |
| 349,300 | 0.185692762 | 0.252612462 | 0.083873943 | 0.183426797 |
| 349,400 | 0.185726064 | 0.254102328 | 0.083283845 | 0.183841071 |
| 349,500 | 0.1857677 | 0.253470036 | 0.082890464 | 0.18338108 |
| 349,600 | 0.185563811 | 0.25311374 | 0.082384097 | 0.184066146 |
| 349,700 | 0.185474246 | 0.252987282 | 0.081822776 | 0.183370072 |
| 349,800 | 0.185729288 | 0.253605385 | 0.08240418 | 0.183696536 |
| 349,900 | 0.185698525 | 0.252757184 | 0.08241351 | 0.184099307 |
| 350,000 | 0.185597181 | 0.252595397 | 0.083780874 | 0.183613334 |
| 350,100 | 0.185512239 | 0.252851501 | 0.083881946 | 0.18236771 |
| 350,200 | 0.185280227 | 0.252728844 | 0.083172056 | 0.182245305 |
| 350,300 | 0.185486431 | 0.252355422 | 0.083288962 | 0.182534747 |
| 350,400 | 0.185549158 | 0.252755157 | 0.083259321 | 0.18310642 |
| 350,500 | 0.185614248 | 0.252290006 | 0.082484273 | 0.183537152 |
| 350,600 | 0.185662468 | 0.252641477 | 0.082388844 | 0.183255087 |
| 350,700 | 0.185568643 | 0.251804352 | 0.083453667 | 0.182832246 |
| 350,800 | 0.18551209 | 0.251252453 | 0.083932188 | 0.183052112 |
| 350,900 | 0.185492249 | 0.251525082 | 0.082781304 | 0.182695023 |
| 351,000 | 0.185294081 | 0.251118995 | 0.083124421 | 0.18314513 |
| 351,100 | 0.185272196 | 0.25153873 | 0.083553889 | 0.18266487 |
| 351,200 | 0.185265299 | 0.250624402 | 0.0828484 | 0.182160587 |
| 351,300 | 0.185182241 | 0.25148894 | 0.083940865 | 0.182338241 |
| 351,400 | 0.185154231 | 0.251243627 | 0.083406016 | 0.182781042 |
| 351,500 | 0.185119487 | 0.251922528 | 0.084593506 | 0.182836466 |
| 351,600 | 0.18518741 | 0.252102056 | 0.083403334 | 0.18323774 |
| 351,700 | 0.185143146 | 0.252244308 | 0.083923284 | 0.182724784 |
| 351,800 | 0.185160434 | 0.251220837 | 0.082185664 | 0.182626933 |
| 351,900 | 0.185164356 | 0.250831152 | 0.083208035 | 0.182873988 |
| 352,000 | 0.185318437 | 0.250846064 | 0.082469443 | 0.182569833 |
| 352,100 | 0.185252074 | 0.250006581 | 0.08285766 | 0.183084349 |
| 352,200 | 0.185128953 | 0.250320283 | 0.082050211 | 0.183016096 |
| 352,300 | 0.185194058 | 0.250485822 | 0.082640688 | 0.183203829 |
| 352,400 | 0.185279318 | 0.250453863 | 0.082852908 | 0.182766408 |
| 352,500 | 0.185107157 | 0.250004386 | 0.082176098 | 0.182557256 |
| 352,600 | 0.185074566 | 0.250496947 | 0.082255964 | 0.182700481 |
| 352,700 | 0.185079779 | 0.251201081 | 0.08226043 | 0.183264657 |
| 352,800 | 0.185030102 | 0.250182107 | 0.083136946 | 0.182577343 |
| 352,900 | 0.185037281 | 0.250627047 | 0.083005613 | 0.182747805 |
| 353,000 | 0.185118334 | 0.250436573 | 0.082906897 | 0.182302486 |
| 353,100 | 0.184944176 | 0.249998149 | 0.082712935 | 0.182496093 |
| 353,200 | 0.184900912 | 0.249465014 | 0.081676488 | 0.182762991 |
| 353,300 | 0.184893851 | 0.25010912 | 0.082535511 | 0.182963693 |
| 353,400 | 0.184893455 | 0.25041968 | 0.083050326 | 0.183196505 |
| 353,500 | 0.184693053 | 0.24988211 | 0.0824283 | 0.183065315 |
| 353,600 | 0.184710548 | 0.250174467 | 0.082342952 | 0.183030547 |
| 353,700 | 0.184645817 | 0.249548031 | 0.081988469 | 0.182760469 |
| 353,800 | 0.184647418 | 0.249790771 | 0.082038186 | 0.182922872 |
| 353,900 | 0.184775618 | 0.250602917 | 0.082496401 | 0.183000737 |
| 354,000 | 0.184560278 | 0.250612841 | 0.082244014 | 0.182976754 |

| | | | | |
|---|---|---|---|---|
| 354,100 | 0.184559426 | 0.251267296 | 0.082705726 | 0.182918755 |
| 354,200 | 0.184621652 | 0.251019507 | 0.082691165 | 0.182671595 |
| 354,300 | 0.18457046 | 0.251055252 | 0.08270166 | 0.183412025 |
| 354,400 | 0.184437565 | 0.250943946 | 0.082529967 | 0.182758125 |
| 354,500 | 0.184488759 | 0.25157409 | 0.082497638 | 0.182743758 |
| 354,600 | 0.184308419 | 0.250726104 | 0.082058695 | 0.182312008 |
| 354,700 | 0.184159425 | 0.250416425 | 0.082486404 | 0.182927072 |
| 354,800 | 0.184319644 | 0.250265135 | 0.083413835 | 0.182599797 |
| 354,900 | 0.184145663 | 0.25007075 | 0.083186179 | 0.182481965 |
| 355,000 | 0.184212055 | 0.25114408 | 0.083107068 | 0.182179067 |
| 355,100 | 0.184135749 | 0.250775114 | 0.082779041 | 0.182358952 |
| 355,200 | 0.184253011 | 0.249877317 | 0.083499969 | 0.182169015 |
| 355,300 | 0.184037196 | 0.249879121 | 0.08258821 | 0.182767356 |
| 355,400 | 0.184268185 | 0.250898988 | 0.083133881 | 0.182213058 |
| 355,500 | 0.184233925 | 0.250767394 | 0.082750623 | 0.182593783 |
| 355,600 | 0.184131233 | 0.250564454 | 0.082656358 | 0.182749001 |
| 355,700 | 0.183994515 | 0.25016345 | 0.08228242 | 0.182595819 |
| 355,800 | 0.184054148 | 0.251310363 | 0.082797998 | 0.182499265 |
| 355,900 | 0.184129734 | 0.251179899 | 0.082444375 | 0.182031264 |
| 356,000 | 0.184002727 | 0.25060131 | 0.081821943 | 0.182564808 |
| 356,100 | 0.184045256 | 0.250488862 | 0.08166413 | 0.18257797 |
| 356,200 | 0.184051949 | 0.250191904 | 0.081936308 | 0.182674851 |
| 356,300 | 0.184079461 | 0.250856381 | 0.082182362 | 0.182717552 |
| 356,400 | 0.184035583 | 0.250109814 | 0.081851924 | 0.1823962 |
| 356,500 | 0.184032621 | 0.249897818 | 0.082341113 | 0.182320464 |
| 356,600 | 0.183933715 | 0.249674266 | 0.081592592 | 0.182327426 |
| 356,700 | 0.183912714 | 0.249841854 | 0.082137294 | 0.182588888 |
| 356,800 | 0.183922816 | 0.249408659 | 0.082262542 | 0.182332964 |
| 356,900 | 0.183985831 | 0.250015394 | 0.081684643 | 0.182675078 |
| 357,000 | 0.18395155 | 0.249877229 | 0.08152966 | 0.182166884 |
| 357,100 | 0.183973746 | 0.249800362 | 0.081749871 | 0.182142307 |
| 357,200 | 0.183896678 | 0.250485715 | 0.081528225 | 0.182823795 |
| 357,300 | 0.183825494 | 0.250000084 | 0.082340834 | 0.182465145 |
| 357,400 | 0.183653224 | 0.25007188 | 0.081005385 | 0.181779945 |
| 357,500 | 0.183668664 | 0.250518572 | 0.08246697 | 0.182276247 |
| 357,600 | 0.183761357 | 0.250403294 | 0.081710512 | 0.182087373 |
| 357,700 | 0.183743624 | 0.250737875 | 0.081304034 | 0.182684931 |
| 357,800 | 0.18376993 | 0.251010314 | 0.08253485 | 0.182476815 |
| 357,900 | 0.183692949 | 0.250596083 | 0.08137086 | 0.181554109 |
| 358,000 | 0.18374258 | 0.250926341 | 0.082587906 | 0.182413583 |
| 358,100 | 0.183542236 | 0.250756462 | 0.081493091 | 0.181680421 |
| 358,200 | 0.183441552 | 0.249660999 | 0.081797556 | 0.181696628 |
| 358,300 | 0.183524072 | 0.250463309 | 0.08209169 | 0.181594288 |
| 358,400 | 0.18352148 | 0.250870403 | 0.081797634 | 0.181486157 |
| 358,500 | 0.183506059 | 0.251013727 | 0.08307457 | 0.180903483 |
| 358,600 | 0.183636364 | 0.251059076 | 0.081930108 | 0.181648306 |
| 358,700 | 0.183334447 | 0.24962436 | 0.082459014 | 0.180983624 |
| 358,800 | 0.183515868 | 0.250382938 | 0.081689485 | 0.181398113 |
| 358,900 | 0.183366289 | 0.249946406 | 0.082194763 | 0.181299691 |
| 359,000 | 0.183312331 | 0.250722564 | 0.082266842 | 0.181189722 |
| 359,100 | 0.183244345 | 0.250275183 | 0.081468399 | 0.180714699 |
| 359,200 | 0.183024042 | 0.250467211 | 0.081514948 | 0.180057094 |
| 359,300 | 0.18320013 | 0.249862255 | 0.08165856 | 0.181718013 |
| 359,400 | 0.183248136 | 0.249820596 | 0.081323096 | 0.181305754 |
| 359,500 | 0.183109126 | 0.250413653 | 0.082571866 | 0.181368586 |
| 359,600 | 0.183267741 | 0.250492853 | 0.081747109 | 0.181296702 |
| 359,700 | 0.183149535 | 0.249451034 | 0.082057176 | 0.181572528 |
| 359,800 | 0.183236003 | 0.250143509 | 0.082954 | 0.180898222 |
| 359,900 | 0.183170654 | 0.249711958 | 0.082363654 | 0.181486113 |
| 360,000 | 0.183203264 | 0.250426423 | 0.081506405 | 0.180586919 |
| 360,100 | 0.18332968 | 0.249886079 | 0.083020255 | 0.180690105 |
| 360,200 | 0.183294862 | 0.249703713 | 0.083380485 | 0.18026327 |
| 360,300 | 0.183252464 | 0.249748386 | 0.08361361 | 0.179872984 |
| 360,400 | 0.183185673 | 0.250225993 | 0.08332534 | 0.180248446 |
| 360,500 | 0.183297769 | 0.250380568 | 0.083892277 | 0.180331273 |
| 360,600 | 0.18331327 | 0.24943204 | 0.084125544 | 0.181020128 |
| 360,700 | 0.183355046 | 0.249324691 | 0.083345319 | 0.180403417 |

| | | | | |
|---|---|---|---|---|
| 360,800 | 0.183129644 | 0.249552074 | 0.08328997 | 0.180100257 |
| 360,900 | 0.183086492 | 0.249762988 | 0.084103358 | 0.180234394 |
| 361,000 | 0.183067154 | 0.249203963 | 0.083940455 | 0.179979466 |
| 361,100 | 0.182890986 | 0.248271006 | 0.084061492 | 0.18044152 |
| 361,200 | 0.182836817 | 0.248038898 | 0.083630581 | 0.180879884 |
| 361,300 | 0.182825354 | 0.248034517 | 0.083639542 | 0.180276171 |
| 361,400 | 0.182842182 | 0.248223189 | 0.083770238 | 0.180207335 |
| 361,500 | 0.182929614 | 0.24890029 | 0.083994618 | 0.179984859 |
| 361,600 | 0.182892169 | 0.248212933 | 0.083242453 | 0.180338226 |
| 361,700 | 0.182787938 | 0.247637265 | 0.082884906 | 0.180236361 |
| 361,800 | 0.182727564 | 0.24873276 | 0.082737047 | 0.180169734 |
| 361,900 | 0.182555434 | 0.247692509 | 0.083097159 | 0.179633338 |
| 362,000 | 0.182677794 | 0.248869568 | 0.083220245 | 0.179910726 |
| 362,100 | 0.182524288 | 0.248462362 | 0.082714409 | 0.179479449 |
| 362,200 | 0.182591109 | 0.248249266 | 0.081799912 | 0.179474617 |
| 362,300 | 0.18247465 | 0.248463517 | 0.082858239 | 0.180016754 |
| 362,400 | 0.182553597 | 0.248566727 | 0.083220378 | 0.180197197 |
| 362,500 | 0.182454018 | 0.249399692 | 0.082240706 | 0.180349548 |
| 362,600 | 0.182518481 | 0.248911773 | 0.083192771 | 0.180283283 |
| 362,700 | 0.182504758 | 0.249017984 | 0.082902009 | 0.18090417 |
| 362,800 | 0.182521147 | 0.249562231 | 0.083505976 | 0.179838244 |
| 362,900 | 0.182281554 | 0.248387239 | 0.082804787 | 0.179765357 |
| 363,000 | 0.182284207 | 0.247926746 | 0.082401837 | 0.180098092 |
| 363,100 | 0.182334692 | 0.2482382 | 0.082823707 | 0.179724578 |
| 363,200 | 0.182068354 | 0.247442452 | 0.082599383 | 0.17986076 |
| 363,300 | 0.182196898 | 0.247928005 | 0.083254026 | 0.179584489 |
| 363,400 | 0.182063757 | 0.248433974 | 0.08326215 | 0.179233571 |
| 363,500 | 0.182163581 | 0.247087276 | 0.084125335 | 0.179805566 |
| 363,600 | 0.182185293 | 0.247384487 | 0.083575391 | 0.179862089 |
| 363,700 | 0.182144458 | 0.247800882 | 0.082155173 | 0.179714327 |
| 363,800 | 0.18204435 | 0.247619625 | 0.083133989 | 0.179440573 |
| 363,900 | 0.182128759 | 0.246378996 | 0.082311426 | 0.179703711 |
| 364,000 | 0.182081011 | 0.247467646 | 0.08169673 | 0.17992046 |
| 364,100 | 0.18191506 | 0.247725805 | 0.081631104 | 0.179250076 |
| 364,200 | 0.18201407 | 0.247650908 | 0.081649666 | 0.179214731 |
| 364,300 | 0.181990934 | 0.247280845 | 0.083500522 | 0.178955023 |
| 364,400 | 0.181983405 | 0.247769873 | 0.083893881 | 0.179279004 |
| 364,500 | 0.181844259 | 0.247474369 | 0.082540519 | 0.179074131 |
| 364,600 | 0.182026984 | 0.246850581 | 0.083076588 | 0.179220668 |
| 364,700 | 0.182099164 | 0.246334043 | 0.083778569 | 0.179743782 |
| 364,800 | 0.1818515 | 0.24613207 | 0.082958938 | 0.179692425 |
| 364,900 | 0.181876448 | 0.246200297 | 0.08270492 | 0.179194016 |
| 365,000 | 0.181843512 | 0.246280826 | 0.082393725 | 0.17902098 |
| 365,100 | 0.181954987 | 0.246766362 | 0.082520188 | 0.179104918 |
| 365,200 | 0.18194444 | 0.247212216 | 0.083101424 | 0.178416839 |
| 365,300 | 0.181798194 | 0.246266793 | 0.082464528 | 0.177751119 |
| 365,400 | 0.181756336 | 0.24580027 | 0.0823291 | 0.178355361 |
| 365,500 | 0.181790666 | 0.245821743 | 0.081987829 | 0.17883266 |
| 365,600 | 0.181902368 | 0.246014796 | 0.082058129 | 0.178881746 |
| 365,700 | 0.18164234 | 0.245059956 | 0.081971126 | 0.17930076 |
| 365,800 | 0.181806814 | 0.245941101 | 0.082231326 | 0.177964745 |
| 365,900 | 0.181830379 | 0.245599589 | 0.082089094 | 0.179129027 |
| 366,000 | 0.181718121 | 0.245982285 | 0.082433017 | 0.177962805 |
| 366,100 | 0.181686424 | 0.246386796 | 0.083093217 | 0.177816644 |
| 366,200 | 0.181671313 | 0.245886311 | 0.081596003 | 0.178652711 |
| 366,300 | 0.181738543 | 0.246385805 | 0.082018314 | 0.17805147 |
| 366,400 | 0.181708069 | 0.245294729 | 0.082725082 | 0.17862753 |
| 366,500 | 0.181658832 | 0.244969897 | 0.083055151 | 0.178111782 |
| 366,600 | 0.181609658 | 0.244613049 | 0.083030663 | 0.178540132 |
| 366,700 | 0.181583715 | 0.244415279 | 0.082683295 | 0.178065455 |
| 366,800 | 0.181667915 | 0.24479704 | 0.083520646 | 0.17866859 |
| 366,900 | 0.18162886 | 0.245699978 | 0.08410035 | 0.17798177 |
| 367,000 | 0.181476277 | 0.245293433 | 0.082926627 | 0.178629108 |
| 367,100 | 0.181430596 | 0.245003129 | 0.081599711 | 0.178567195 |
| 367,200 | 0.181528904 | 0.244950079 | 0.081743474 | 0.178904983 |
| 367,300 | 0.181442197 | 0.244748285 | 0.081213303 | 0.178643826 |
| 367,400 | 0.181548164 | 0.244812728 | 0.081711038 | 0.178583945 |

| | | | | |
|---|---|---|---|---|
| 367,500 | 0.18137683 | 0.244390199 | 0.081593688 | 0.178000492 |
| 367,600 | 0.181389017 | 0.244677218 | 0.08157002 | 0.178082901 |
| 367,700 | 0.181365951 | 0.24461805 | 0.081102442 | 0.178682851 |
| 367,800 | 0.181392408 | 0.245129261 | 0.082442965 | 0.17876259 |
| 367,900 | 0.181241678 | 0.244747898 | 0.081731164 | 0.178405046 |
| 368,000 | 0.181300827 | 0.245386111 | 0.081770222 | 0.177865177 |
| 368,100 | 0.181198615 | 0.244622762 | 0.081363931 | 0.178664468 |
| 368,200 | 0.181250516 | 0.245332618 | 0.081678156 | 0.178123115 |
| 368,300 | 0.181198676 | 0.244182485 | 0.082857676 | 0.178368429 |
| 368,400 | 0.181257561 | 0.244738469 | 0.082567301 | 0.177883583 |
| 368,500 | 0.18120928 | 0.244043171 | 0.081756131 | 0.178237253 |
| 368,600 | 0.181133264 | 0.244281998 | 0.082884596 | 0.178053953 |
| 368,700 | 0.181187742 | 0.244640166 | 0.082106558 | 0.178194964 |
| 368,800 | 0.181017831 | 0.243735051 | 0.081923582 | 0.178226025 |
| 368,900 | 0.181034531 | 0.244848028 | 0.081764311 | 0.178372611 |
| 369,000 | 0.180968511 | 0.245036699 | 0.081457824 | 0.179157117 |
| 369,100 | 0.181020017 | 0.244947078 | 0.081996082 | 0.178963001 |
| 369,200 | 0.181206038 | 0.244878657 | 0.082596903 | 0.178693782 |
| 369,300 | 0.181148502 | 0.245224937 | 0.082539197 | 0.178830532 |
| 369,400 | 0.180935699 | 0.244245591 | 0.081973422 | 0.178847275 |
| 369,500 | 0.180981793 | 0.243960385 | 0.082404301 | 0.178638647 |
| 369,600 | 0.180881733 | 0.243352165 | 0.082183759 | 0.177929946 |
| 369,700 | 0.180902276 | 0.243421129 | 0.083136887 | 0.177802454 |
| 369,800 | 0.180918633 | 0.244590053 | 0.082232747 | 0.177620102 |
| 369,900 | 0.180870215 | 0.244698557 | 0.082108205 | 0.177510684 |
| 370,000 | 0.180863239 | 0.243832096 | 0.08166359 | 0.177670373 |
| 370,100 | 0.180865401 | 0.244086608 | 0.080654145 | 0.177089084 |
| 370,200 | 0.180700585 | 0.243677477 | 0.081428372 | 0.177222233 |
| 370,300 | 0.180846575 | 0.244408667 | 0.081667612 | 0.177977826 |
| 370,400 | 0.180907044 | 0.244587232 | 0.081273092 | 0.177964191 |
| 370,500 | 0.180698005 | 0.24449126 | 0.081655936 | 0.177334015 |
| 370,600 | 0.180696224 | 0.244727729 | 0.080833689 | 0.177058136 |
| 370,700 | 0.180706686 | 0.244442498 | 0.08143845 | 0.177328403 |
| 370,800 | 0.180682264 | 0.244181259 | 0.081556684 | 0.176706215 |
| 370,900 | 0.180566905 | 0.243727299 | 0.081412134 | 0.177942339 |
| 371,000 | 0.18060468 | 0.244032211 | 0.081191849 | 0.177583236 |
| 371,100 | 0.180464468 | 0.243281415 | 0.080640963 | 0.177087348 |
| 371,200 | 0.180401947 | 0.243280298 | 0.080437371 | 0.177276379 |
| 371,300 | 0.180504579 | 0.244458571 | 0.080408148 | 0.177406837 |
| 371,400 | 0.180302937 | 0.243903132 | 0.081492441 | 0.177491955 |
| 371,500 | 0.180425944 | 0.243794413 | 0.082018731 | 0.176968399 |
| 371,600 | 0.18025057 | 0.243461438 | 0.082380341 | 0.177149085 |
| 371,700 | 0.180386657 | 0.244060145 | 0.080907428 | 0.177238491 |
| 371,800 | 0.180391722 | 0.244030273 | 0.081266123 | 0.177928954 |
| 371,900 | 0.180225752 | 0.244122758 | 0.081644566 | 0.177201557 |
| 372,000 | 0.180237379 | 0.24351808 | 0.08048534 | 0.17764875 |
| 372,100 | 0.180284354 | 0.243430066 | 0.081151277 | 0.177217294 |
| 372,200 | 0.180380295 | 0.243522753 | 0.081408269 | 0.177316247 |
| 372,300 | 0.180116957 | 0.243688569 | 0.080743131 | 0.177184014 |
| 372,400 | 0.180220986 | 0.242945533 | 0.081658427 | 0.177980634 |
| 372,500 | 0.180103032 | 0.243489887 | 0.080989532 | 0.177334014 |
| 372,600 | 0.18008246 | 0.243529265 | 0.080372118 | 0.177793877 |
| 372,700 | 0.180109081 | 0.243770419 | 0.080387335 | 0.177542186 |
| 372,800 | 0.180196568 | 0.243386993 | 0.080630874 | 0.178090626 |
| 372,900 | 0.180025054 | 0.243645956 | 0.08079272 | 0.177429586 |
| 373,000 | 0.180143435 | 0.243629027 | 0.081354882 | 0.178134823 |
| 373,100 | 0.180020977 | 0.243060606 | 0.081478875 | 0.17749172 |
| 373,200 | 0.180065596 | 0.244354197 | 0.08229446 | 0.177231011 |
| 373,300 | 0.180001254 | 0.244200439 | 0.082842848 | 0.178013724 |
| 373,400 | 0.179967776 | 0.24450965 | 0.082691717 | 0.177142572 |
| 373,500 | 0.179908582 | 0.24396766 | 0.083400738 | 0.178148187 |
| 373,600 | 0.179974035 | 0.243685014 | 0.083169727 | 0.177795658 |
| 373,700 | 0.180131412 | 0.243227581 | 0.082374321 | 0.177249991 |
| 373,800 | 0.179874923 | 0.243300116 | 0.082816146 | 0.177610654 |
| 373,900 | 0.179704466 | 0.242979719 | 0.081712711 | 0.177615405 |
| 374,000 | 0.179710094 | 0.242406251 | 0.082866507 | 0.178067842 |
| 374,100 | 0.179737483 | 0.242498294 | 0.081993033 | 0.176418552 |

| | | | | |
|---|---|---|---|---|
| 374,200 | 0.179750018 | 0.242467232 | 0.082394592 | 0.177197071 |
| 374,300 | 0.179790694 | 0.242909147 | 0.082286377 | 0.176237054 |
| 374,400 | 0.17965101 | 0.243266335 | 0.081611145 | 0.176779887 |
| 374,500 | 0.17955804 | 0.242611511 | 0.081852435 | 0.177422264 |
| 374,600 | 0.179711639 | 0.242531175 | 0.081831753 | 0.177113262 |
| 374,700 | 0.17964886 | 0.242986991 | 0.082230107 | 0.176708255 |
| 374,800 | 0.179633795 | 0.243456605 | 0.082138095 | 0.17705819 |
| 374,900 | 0.179548528 | 0.242554619 | 0.081977953 | 0.176346743 |
| 375,000 | 0.179616731 | 0.242592678 | 0.081873003 | 0.176891173 |
| 375,100 | 0.179698025 | 0.242121524 | 0.081238872 | 0.176038875 |
| 375,200 | 0.179543765 | 0.243058927 | 0.081402127 | 0.176090954 |
| 375,300 | 0.179547985 | 0.243560773 | 0.081945159 | 0.176390685 |
| 375,400 | 0.179623557 | 0.244170147 | 0.082250596 | 0.176007258 |
| 375,500 | 0.179475754 | 0.242375622 | 0.081607378 | 0.176683599 |
| 375,600 | 0.179494524 | 0.242853692 | 0.082233359 | 0.176155547 |
| 375,700 | 0.17936977 | 0.242828552 | 0.083086315 | 0.175740711 |
| 375,800 | 0.179473064 | 0.243348338 | 0.082528791 | 0.176284919 |
| 375,900 | 0.179327599 | 0.243139451 | 0.082304476 | 0.175644469 |
| 376,000 | 0.179378737 | 0.243349539 | 0.082739244 | 0.175722511 |
| 376,100 | 0.179352061 | 0.243500757 | 0.082161128 | 0.17586802 |
| 376,200 | 0.179429037 | 0.243678123 | 0.081784931 | 0.176322538 |
| 376,300 | 0.179571094 | 0.243403235 | 0.082083971 | 0.176559302 |
| 376,400 | 0.179545633 | 0.243510886 | 0.08223146 | 0.176971996 |
| 376,500 | 0.179582451 | 0.243243253 | 0.082021269 | 0.176936395 |
| 376,600 | 0.179581572 | 0.243153396 | 0.081070033 | 0.176640597 |
| 376,700 | 0.179599708 | 0.243523999 | 0.080835869 | 0.176586864 |
| 376,800 | 0.179440146 | 0.242893653 | 0.080360633 | 0.176437605 |
| 376,900 | 0.179476045 | 0.242883136 | 0.081495934 | 0.177008131 |
| 377,000 | 0.179424069 | 0.24259432 | 0.081006807 | 0.17715254 |
| 377,100 | 0.179410881 | 0.243208721 | 0.080723686 | 0.177864116 |
| 377,200 | 0.179393687 | 0.243575456 | 0.080875585 | 0.177289875 |
| 377,300 | 0.179526071 | 0.243852378 | 0.080451487 | 0.17776634 |
| 377,400 | 0.179433056 | 0.243209815 | 0.081439128 | 0.177792616 |
| 377,500 | 0.179445099 | 0.243053216 | 0.08091743 | 0.177959223 |
| 377,600 | 0.179409344 | 0.243367002 | 0.080217364 | 0.177964572 |
| 377,700 | 0.179288769 | 0.243575258 | 0.080240331 | 0.176761293 |
| 377,800 | 0.179104864 | 0.24376372 | 0.080219644 | 0.176568879 |
| 377,900 | 0.179056388 | 0.243344436 | 0.0806727 | 0.177158067 |
| 378,000 | 0.179229954 | 0.242936361 | 0.080815993 | 0.17679073 |
| 378,100 | 0.17915331 | 0.243600497 | 0.080662174 | 0.176584806 |
| 378,200 | 0.179264955 | 0.244064891 | 0.080544707 | 0.176609937 |
| 378,300 | 0.179278244 | 0.244525919 | 0.080295179 | 0.177303129 |
| 378,400 | 0.179194042 | 0.243842898 | 0.080865315 | 0.177132948 |
| 378,500 | 0.179182625 | 0.243831513 | 0.080740859 | 0.176694215 |
| 378,600 | 0.179084563 | 0.243721329 | 0.079917739 | 0.176519411 |
| 378,700 | 0.179136817 | 0.243234163 | 0.080865179 | 0.176719978 |
| 378,800 | 0.178936642 | 0.243754594 | 0.080336568 | 0.176441912 |
| 378,900 | 0.17896262 | 0.243712388 | 0.080043502 | 0.176743001 |
| 379,000 | 0.178936465 | 0.243301971 | 0.079857526 | 0.17660146 |
| 379,100 | 0.178787372 | 0.243862793 | 0.080349822 | 0.17707951 |
| 379,200 | 0.178860871 | 0.243150286 | 0.080504384 | 0.176208053 |
| 379,300 | 0.178845053 | 0.243823199 | 0.080433591 | 0.176372892 |
| 379,400 | 0.178849627 | 0.244214452 | 0.079988928 | 0.177114837 |
| 379,500 | 0.178762467 | 0.243899294 | 0.079825945 | 0.177823581 |
| 379,600 | 0.178687635 | 0.243090255 | 0.08014252 | 0.177071452 |
| 379,700 | 0.178780734 | 0.243352383 | 0.079499763 | 0.177007087 |
| 379,800 | 0.178600062 | 0.243003758 | 0.079776513 | 0.17765572 |
| 379,900 | 0.178754457 | 0.243255262 | 0.080227604 | 0.177434653 |
| 380,000 | 0.178684495 | 0.243195114 | 0.079552995 | 0.177203686 |
| 380,100 | 0.178650074 | 0.243062748 | 0.078877001 | 0.177069426 |
| 380,200 | 0.178621433 | 0.243013188 | 0.079449024 | 0.176796367 |
| 380,300 | 0.178715904 | 0.242900769 | 0.079291756 | 0.177222484 |
| 380,400 | 0.178621522 | 0.242694068 | 0.078630633 | 0.177674769 |
| 380,500 | 0.178604477 | 0.242971889 | 0.080032003 | 0.177239842 |
| 380,600 | 0.178491597 | 0.242766977 | 0.07871273 | 0.17776808 |
| 380,700 | 0.178564562 | 0.243480376 | 0.079362335 | 0.1767274 |
| 380,800 | 0.178474748 | 0.241935207 | 0.079013039 | 0.176433332 |

| | | | | |
|---|---|---|---|---|
| 380,900 | 0.178436585 | 0.242717307 | 0.079058156 | 0.177092803 |
| 381,000 | 0.178503447 | 0.242498861 | 0.078478235 | 0.177278982 |
| 381,100 | 0.1783773 | 0.2426457 | 0.079895854 | 0.177012683 |
| 381,200 | 0.178390524 | 0.24195072 | 0.079914466 | 0.176847819 |
| 381,300 | 0.178323509 | 0.242719052 | 0.078754481 | 0.177311331 |
| 381,400 | 0.178318327 | 0.242800969 | 0.078960249 | 0.177616849 |
| 381,500 | 0.178364333 | 0.242455045 | 0.079397466 | 0.177174007 |
| 381,600 | 0.178451544 | 0.242716008 | 0.07951465 | 0.17732192 |
| 381,700 | 0.17836695 | 0.242276631 | 0.07907313 | 0.177256737 |
| 381,800 | 0.17844258 | 0.242959561 | 0.079196691 | 0.177522749 |
| 381,900 | 0.178312467 | 0.242103104 | 0.078908779 | 0.17806584 |
| 382,000 | 0.178309641 | 0.242551059 | 0.079455471 | 0.17742742 |
| 382,100 | 0.17835708 | 0.243165142 | 0.07857905 | 0.177028034 |
| 382,200 | 0.178373691 | 0.243986265 | 0.078263062 | 0.177160141 |
| 382,300 | 0.178376527 | 0.243365796 | 0.078268773 | 0.177400627 |
| 382,400 | 0.178254674 | 0.243757209 | 0.079198147 | 0.177663329 |
| 382,500 | 0.178330896 | 0.244002486 | 0.079158322 | 0.176855558 |
| 382,600 | 0.178242872 | 0.244785434 | 0.078766111 | 0.176716519 |
| 382,700 | 0.178156791 | 0.243511988 | 0.078999486 | 0.176437754 |
| 382,800 | 0.178083456 | 0.243456719 | 0.078588083 | 0.176741073 |
| 382,900 | 0.178037384 | 0.242623417 | 0.079219615 | 0.176390147 |
| 383,000 | 0.178040599 | 0.243395105 | 0.078154012 | 0.177053848 |
| 383,100 | 0.178036204 | 0.244073207 | 0.07735906 | 0.176874342 |
| 383,200 | 0.178056163 | 0.243033812 | 0.077219265 | 0.176533836 |
| 383,300 | 0.177884544 | 0.242862756 | 0.077538801 | 0.17658609 |
| 383,400 | 0.178130664 | 0.243718871 | 0.077971759 | 0.176796347 |
| 383,500 | 0.177833234 | 0.243100587 | 0.078403477 | 0.176228492 |
| 383,600 | 0.17792644 | 0.243771253 | 0.078429963 | 0.176479357 |
| 383,700 | 0.177897617 | 0.243745318 | 0.07822667 | 0.176443778 |
| 383,800 | 0.177860797 | 0.243985772 | 0.078214443 | 0.176442725 |
| 383,900 | 0.177819363 | 0.243528207 | 0.078636211 | 0.176920804 |
| 384,000 | 0.177919141 | 0.243156654 | 0.078121512 | 0.177277725 |
| 384,100 | 0.177724005 | 0.24380823 | 0.078224317 | 0.17695089 |
| 384,200 | 0.17769421 | 0.243870887 | 0.078655207 | 0.176637112 |
| 384,300 | 0.177617869 | 0.24399793 | 0.077973712 | 0.176150792 |
| 384,400 | 0.177612993 | 0.243714031 | 0.076928416 | 0.176729429 |
| 384,500 | 0.177709711 | 0.244010569 | 0.077988343 | 0.176184789 |
| 384,600 | 0.177529782 | 0.243070283 | 0.078141311 | 0.176645296 |
| 384,700 | 0.177578837 | 0.243678868 | 0.07884027 | 0.176521585 |
| 384,800 | 0.17757421 | 0.243399086 | 0.078470458 | 0.176503247 |
| 384,900 | 0.177592123 | 0.243251195 | 0.07777318 | 0.176008173 |
| 385,000 | 0.177545208 | 0.243511289 | 0.07737623 | 0.176793308 |
| 385,100 | 0.17753153 | 0.243883747 | 0.077505653 | 0.176728871 |
| 385,200 | 0.177662924 | 0.244000406 | 0.076697257 | 0.176358619 |
| 385,300 | 0.177575024 | 0.242510143 | 0.077902274 | 0.176552978 |
| 385,400 | 0.177503112 | 0.243813221 | 0.077610057 | 0.176018812 |
| 385,500 | 0.177400953 | 0.243832637 | 0.077683714 | 0.175720575 |
| 385,600 | 0.177505795 | 0.242992447 | 0.078029821 | 0.176256568 |
| 385,700 | 0.17746668 | 0.242761283 | 0.078225839 | 0.176113862 |
| 385,800 | 0.177468325 | 0.242941732 | 0.07828005 | 0.175917852 |
| 385,900 | 0.17741545 | 0.242932583 | 0.077780057 | 0.175643231 |
| 386,000 | 0.177367921 | 0.243632506 | 0.078356804 | 0.175413955 |
| 386,100 | 0.177553497 | 0.24302802 | 0.07822442 | 0.175528901 |
| 386,200 | 0.177323599 | 0.242746545 | 0.0776887 | 0.176072537 |
| 386,300 | 0.177273158 | 0.243462479 | 0.077536212 | 0.176238062 |
| 386,400 | 0.177418672 | 0.243183033 | 0.077611376 | 0.176364922 |
| 386,500 | 0.177367524 | 0.242427172 | 0.077602653 | 0.175508071 |
| 386,600 | 0.177264068 | 0.243236012 | 0.07751266 | 0.175935356 |
| 386,700 | 0.177200898 | 0.242508645 | 0.077922912 | 0.175731257 |
| 386,800 | 0.177263022 | 0.242545601 | 0.077289412 | 0.176263892 |
| 386,900 | 0.177163895 | 0.24214333 | 0.078010736 | 0.175265124 |
| 387,000 | 0.177085259 | 0.241864164 | 0.078083923 | 0.175744703 |
| 387,100 | 0.177061291 | 0.242851924 | 0.077443135 | 0.175443401 |
| 387,200 | 0.17689171 | 0.242674496 | 0.076542475 | 0.175179351 |
| 387,300 | 0.176926618 | 0.242557601 | 0.076787108 | 0.175708872 |
| 387,400 | 0.177066022 | 0.241678296 | 0.077609385 | 0.175305101 |
| 387,500 | 0.177054148 | 0.242457734 | 0.077689451 | 0.175524257 |

| | | | | |
|---|---|---|---|---|
| 387,600 | 0.177063209 | 0.242501825 | 0.077831581 | 0.175942835 |
| 387,700 | 0.177080695 | 0.243150542 | 0.078066867 | 0.175547145 |
| 387,800 | 0.177045785 | 0.241841423 | 0.078701385 | 0.175110064 |
| 387,900 | 0.177017159 | 0.241802431 | 0.078381772 | 0.174687318 |
| 388,000 | 0.176979134 | 0.240709743 | 0.077721922 | 0.175186333 |
| 388,100 | 0.176995849 | 0.241322258 | 0.078361987 | 0.175138799 |
| 388,200 | 0.176992274 | 0.242677741 | 0.077966061 | 0.175416101 |
| 388,300 | 0.176975296 | 0.242543946 | 0.078781682 | 0.174971723 |
| 388,400 | 0.17694166 | 0.241489999 | 0.07803723 | 0.175320332 |
| 388,500 | 0.176976418 | 0.241530732 | 0.078551433 | 0.175074079 |
| 388,600 | 0.176902789 | 0.24144628 | 0.077407751 | 0.17464085 |
| 388,700 | 0.176877956 | 0.241651138 | 0.078290117 | 0.174962883 |
| 388,800 | 0.176644023 | 0.241014359 | 0.0783399 | 0.175010205 |
| 388,900 | 0.176889068 | 0.241244627 | 0.077304021 | 0.175630642 |
| 389,000 | 0.176814266 | 0.241431455 | 0.077705449 | 0.175097267 |
| 389,100 | 0.176835252 | 0.241802818 | 0.078097005 | 0.175236105 |
| 389,200 | 0.176795005 | 0.240765183 | 0.07824203 | 0.174619069 |
| 389,300 | 0.176785756 | 0.240665279 | 0.079175673 | 0.174717549 |
| 389,400 | 0.176710847 | 0.240824465 | 0.0789054 | 0.174764114 |
| 389,500 | 0.176850047 | 0.240827039 | 0.078640222 | 0.174560377 |
| 389,600 | 0.176658766 | 0.241303657 | 0.077320254 | 0.174757569 |
| 389,700 | 0.176810392 | 0.240688266 | 0.07898313 | 0.17489832 |
| 389,800 | 0.176677457 | 0.24100276 | 0.078583448 | 0.174850157 |
| 389,900 | 0.176594198 | 0.240594744 | 0.078305635 | 0.175100326 |
| 390,000 | 0.176629374 | 0.240793153 | 0.077868803 | 0.174840611 |
| 390,100 | 0.176597478 | 0.240400946 | 0.077741217 | 0.175342967 |
| 390,200 | 0.176563999 | 0.240126588 | 0.078946294 | 0.175554901 |
| 390,300 | 0.176605629 | 0.241154216 | 0.078810102 | 0.175318273 |
| 390,400 | 0.176461542 | 0.241820672 | 0.078693992 | 0.175111794 |
| 390,500 | 0.176397772 | 0.241260732 | 0.078669256 | 0.175257374 |
| 390,600 | 0.176438812 | 0.240975342 | 0.079620095 | 0.174692758 |
| 390,700 | 0.176401737 | 0.24069583 | 0.079014803 | 0.174114405 |
| 390,800 | 0.176387186 | 0.242146993 | 0.080357804 | 0.175163685 |
| 390,900 | 0.176457582 | 0.242238358 | 0.079597972 | 0.174885106 |
| 391,000 | 0.176450281 | 0.241877604 | 0.079630846 | 0.175409853 |
| 391,100 | 0.176551927 | 0.242541414 | 0.079534069 | 0.175015909 |
| 391,200 | 0.176505967 | 0.242320449 | 0.079242949 | 0.174429769 |
| 391,300 | 0.1763358 | 0.241452483 | 0.080122494 | 0.174341523 |
| 391,400 | 0.176257571 | 0.241254534 | 0.079516382 | 0.174306558 |
| 391,500 | 0.176173152 | 0.241202117 | 0.080262528 | 0.173861942 |
| 391,600 | 0.176164103 | 0.240840174 | 0.080125747 | 0.17360387 |
| 391,700 | 0.176207341 | 0.240792932 | 0.080344501 | 0.17438439 |
| 391,800 | 0.176158443 | 0.241302753 | 0.07950711 | 0.174296389 |
| 391,900 | 0.176341698 | 0.240841023 | 0.079457586 | 0.174491763 |
| 392,000 | 0.176070998 | 0.240614495 | 0.079748203 | 0.174444202 |
| 392,100 | 0.176148758 | 0.241710632 | 0.080015251 | 0.174505033 |
| 392,200 | 0.176124945 | 0.24100204 | 0.079947471 | 0.175056795 |
| 392,300 | 0.176167584 | 0.241797237 | 0.08121477 | 0.175141678 |
| 392,400 | 0.176085252 | 0.24144744 | 0.080550577 | 0.174630171 |
| 392,500 | 0.176177698 | 0.241759913 | 0.079982745 | 0.175325653 |
| 392,600 | 0.176057559 | 0.241296678 | 0.079453084 | 0.174868137 |
| 392,700 | 0.176155515 | 0.24181424 | 0.078811754 | 0.175449543 |
| 392,800 | 0.176193092 | 0.241183102 | 0.07902116 | 0.175965994 |
| 392,900 | 0.176125808 | 0.241777175 | 0.0796669 | 0.175130268 |
| 393,000 | 0.176123815 | 0.241912988 | 0.079192696 | 0.175550096 |
| 393,100 | 0.175915937 | 0.240384077 | 0.079844478 | 0.175084391 |
| 393,200 | 0.176026289 | 0.24089732 | 0.078977311 | 0.175539662 |
| 393,300 | 0.175816473 | 0.241303916 | 0.080588797 | 0.174224478 |
| 393,400 | 0.175971781 | 0.240805954 | 0.079678617 | 0.174519973 |
| 393,500 | 0.175986083 | 0.241073694 | 0.079902795 | 0.17437929 |
| 393,600 | 0.175987192 | 0.241329927 | 0.080111819 | 0.174578963 |
| 393,700 | 0.175909446 | 0.241459328 | 0.079303056 | 0.173741331 |
| 393,800 | 0.176025706 | 0.241730677 | 0.079567691 | 0.17412086 |
| 393,900 | 0.175897981 | 0.241433898 | 0.079654773 | 0.173586842 |
| 394,000 | 0.175926906 | 0.241515542 | 0.078898302 | 0.173925705 |
| 394,100 | 0.175743624 | 0.241442914 | 0.079466284 | 0.174040545 |
| 394,200 | 0.17583067 | 0.241800208 | 0.079296127 | 0.174591701 |

| | | | | |
|---|---|---|---|---|
| 394,300 | 0.175834078 | 0.241746281 | 0.078577002 | 0.174293151 |
| 394,400 | 0.175706089 | 0.240602274 | 0.078822808 | 0.17393417 |
| 394,500 | 0.175683926 | 0.240913436 | 0.078970607 | 0.173923668 |
| 394,600 | 0.175649838 | 0.240920271 | 0.079820511 | 0.174035527 |
| 394,700 | 0.175618394 | 0.24108543 | 0.079406042 | 0.173736459 |
| 394,800 | 0.175655797 | 0.240728316 | 0.079125772 | 0.17344052 |
| 394,900 | 0.175575905 | 0.240744538 | 0.078066647 | 0.174040811 |
| 395,000 | 0.175638045 | 0.241123699 | 0.078715769 | 0.174468547 |
| 395,100 | 0.175570348 | 0.241771327 | 0.078119852 | 0.173156743 |
| 395,200 | 0.175556867 | 0.241373725 | 0.077461147 | 0.173852448 |
| 395,300 | 0.175667551 | 0.242105992 | 0.078452032 | 0.173760979 |
| 395,400 | 0.175532995 | 0.240974147 | 0.078010577 | 0.173919952 |
| 395,500 | 0.175436593 | 0.23998825 | 0.078897453 | 0.173631447 |
| 395,600 | 0.175440633 | 0.241095331 | 0.078653909 | 0.174034018 |
| 395,700 | 0.175393987 | 0.240648024 | 0.079489109 | 0.173744504 |
| 395,800 | 0.175327813 | 0.240617558 | 0.079129268 | 0.173758398 |
| 395,900 | 0.17523805 | 0.240151981 | 0.078829386 | 0.173133576 |
| 396,000 | 0.175439044 | 0.24087609 | 0.078902666 | 0.173479473 |
| 396,100 | 0.175411817 | 0.241647647 | 0.078549731 | 0.173258045 |
| 396,200 | 0.175241955 | 0.24115839 | 0.079390144 | 0.1732381 |
| 396,300 | 0.175169387 | 0.241180725 | 0.079340267 | 0.173489516 |
| 396,400 | 0.175173903 | 0.241390366 | 0.078922904 | 0.173109313 |
| 396,500 | 0.175187306 | 0.241065118 | 0.078412941 | 0.173563985 |
| 396,600 | 0.175176709 | 0.241391514 | 0.078545333 | 0.173665359 |
| 396,700 | 0.175142091 | 0.240824496 | 0.078377311 | 0.17255061 |
| 396,800 | 0.175206509 | 0.240791128 | 0.077817975 | 0.173559006 |
| 396,900 | 0.175079598 | 0.240721834 | 0.078021622 | 0.173696197 |
| 397,000 | 0.175029914 | 0.240357083 | 0.077651275 | 0.173612872 |
| 397,100 | 0.175111415 | 0.240414601 | 0.077515627 | 0.172737582 |
| 397,200 | 0.175088625 | 0.240154615 | 0.078367365 | 0.173525562 |
| 397,300 | 0.175194103 | 0.240337914 | 0.078556327 | 0.17347075 |
| 397,400 | 0.175223373 | 0.24053082 | 0.078940011 | 0.173732012 |
| 397,500 | 0.175178715 | 0.240729365 | 0.078497347 | 0.172754699 |
| 397,600 | 0.1751517 | 0.241007153 | 0.078809978 | 0.172868985 |
| 397,700 | 0.175093193 | 0.240485722 | 0.079354062 | 0.173029049 |
| 397,800 | 0.174960436 | 0.240245311 | 0.078299463 | 0.17340945 |
| 397,900 | 0.175012095 | 0.240725659 | 0.078308375 | 0.173891096 |
| 398,000 | 0.174896633 | 0.240169155 | 0.078497416 | 0.173443872 |
| 398,100 | 0.174887728 | 0.240607971 | 0.078046374 | 0.17341886 |
| 398,200 | 0.174898097 | 0.239635777 | 0.078923003 | 0.173753254 |
| 398,300 | 0.175062509 | 0.241482729 | 0.078937375 | 0.173638213 |
| 398,400 | 0.174916805 | 0.240454821 | 0.079244683 | 0.174064146 |
| 398,500 | 0.174978808 | 0.240939024 | 0.078561447 | 0.173351474 |
| 398,600 | 0.174970713 | 0.241698161 | 0.079178326 | 0.173578343 |
| 398,700 | 0.174876769 | 0.241001373 | 0.080022511 | 0.173117335 |
| 398,800 | 0.174934984 | 0.241161473 | 0.078802471 | 0.173372671 |
| 398,900 | 0.174892608 | 0.240657248 | 0.078438708 | 0.17331833 |
| 399,000 | 0.174853227 | 0.241168271 | 0.079334405 | 0.173539448 |
| 399,100 | 0.174892215 | 0.240749751 | 0.079482068 | 0.17319835 |
| 399,200 | 0.17494096 | 0.241554782 | 0.079849561 | 0.173855397 |
| 399,300 | 0.174600944 | 0.240599494 | 0.079434361 | 0.173862842 |
| 399,400 | 0.174623648 | 0.240987939 | 0.080182317 | 0.173445034 |
| 399,500 | 0.17464511 | 0.241090518 | 0.080323556 | 0.172662423 |
| 399,600 | 0.174797085 | 0.240995334 | 0.079764366 | 0.173167694 |
| 399,700 | 0.174806586 | 0.241316238 | 0.079583864 | 0.172771291 |
| 399,800 | 0.174868616 | 0.240570233 | 0.080954379 | 0.173722246 |
| 399,900 | 0.174681644 | 0.24157429 | 0.080177953 | 0.172789125 |
| 400,000 | 0.174650129 | 0.24141652 | 0.079269377 | 0.173072314 |
| 400,100 | 0.174566151 | 0.241204733 | 0.078763998 | 0.172887938 |
| 400,200 | 0.174693937 | 0.240964648 | 0.079375682 | 0.173110492 |
| 400,300 | 0.174675084 | 0.241589465 | 0.080099563 | 0.173596498 |
| 400,400 | 0.17463742 | 0.240971535 | 0.079802201 | 0.17314822 |
| 400,500 | 0.174722719 | 0.240709395 | 0.079985109 | 0.17331293 |
| 400,600 | 0.174617591 | 0.241073861 | 0.080263994 | 0.173832529 |
| 400,700 | 0.174629067 | 0.241139024 | 0.079261933 | 0.173663776 |
| 400,800 | 0.174453484 | 0.240936319 | 0.079363613 | 0.173814779 |
| 400,900 | 0.174458318 | 0.24070335 | 0.078463344 | 0.174118376 |

| | | | | |
|---|---|---|---|---|
| 401,000 | 0.17445899 | 0.240177805 | 0.079021824 | 0.173452269 |
| 401,100 | 0.174523051 | 0.240608089 | 0.079152446 | 0.174832789 |
| 401,200 | 0.174399628 | 0.240732333 | 0.080214997 | 0.174585089 |
| 401,300 | 0.174367476 | 0.241086579 | 0.079979974 | 0.17494183 |
| 401,400 | 0.174444822 | 0.240491212 | 0.079167131 | 0.17407692 |
| 401,500 | 0.174586621 | 0.241700172 | 0.079393391 | 0.174289743 |
| 401,600 | 0.174450893 | 0.240408926 | 0.079473401 | 0.174511353 |
| 401,700 | 0.174346763 | 0.240890238 | 0.078480703 | 0.173575258 |
| 401,800 | 0.174389283 | 0.240334584 | 0.078494739 | 0.173761597 |
| 401,900 | 0.17443535 | 0.240360512 | 0.079339041 | 0.17389469 |
| 402,000 | 0.174420038 | 0.240569821 | 0.078804808 | 0.173911518 |
| 402,100 | 0.174297587 | 0.239907517 | 0.078859862 | 0.174062282 |
| 402,200 | 0.174316077 | 0.239449355 | 0.079246932 | 0.174201068 |
| 402,300 | 0.174327321 | 0.239812744 | 0.079708318 | 0.173576532 |
| 402,400 | 0.174157726 | 0.239534314 | 0.079084244 | 0.173714581 |
| 402,500 | 0.174245768 | 0.239688902 | 0.080139286 | 0.173884062 |
| 402,600 | 0.174152457 | 0.23940516 | 0.079743986 | 0.172960957 |
| 402,700 | 0.17416167 | 0.238833923 | 0.07959064 | 0.173665852 |
| 402,800 | 0.174228694 | 0.239425635 | 0.079935838 | 0.173705025 |
| 402,900 | 0.173952058 | 0.238999275 | 0.079841159 | 0.173716813 |
| 403,000 | 0.174126804 | 0.239401915 | 0.079673582 | 0.173619541 |
| 403,100 | 0.174216252 | 0.239935219 | 0.080130595 | 0.173708266 |
| 403,200 | 0.174125394 | 0.240191524 | 0.079219968 | 0.174128746 |
| 403,300 | 0.174129987 | 0.240180578 | 0.07957854 | 0.174452737 |
| 403,400 | 0.174007717 | 0.240178501 | 0.079330058 | 0.174659512 |
| 403,500 | 0.173933861 | 0.240032579 | 0.080170433 | 0.17465466 |
| 403,600 | 0.173905372 | 0.239377915 | 0.079634219 | 0.17486518 |
| 403,700 | 0.174049262 | 0.23992876 | 0.079093227 | 0.174516248 |
| 403,800 | 0.174112356 | 0.239723559 | 0.0789071 | 0.174529743 |
| 403,900 | 0.173956257 | 0.239673258 | 0.079246716 | 0.174779067 |
| 404,000 | 0.173973229 | 0.239787508 | 0.078639653 | 0.174546021 |
| 404,100 | 0.173900091 | 0.239313002 | 0.079135681 | 0.174518986 |
| 404,200 | 0.17372497 | 0.238759313 | 0.080050154 | 0.17436589 |
| 404,300 | 0.173855482 | 0.239372062 | 0.079666413 | 0.17470468 |
| 404,400 | 0.173758818 | 0.239356187 | 0.079543172 | 0.173955777 |
| 404,500 | 0.17375447 | 0.238913471 | 0.079308177 | 0.17410656 |
| 404,600 | 0.17382445 | 0.238837069 | 0.078961615 | 0.173755946 |
| 404,700 | 0.173804419 | 0.239260569 | 0.078588014 | 0.174155956 |
| 404,800 | 0.173844771 | 0.239133026 | 0.07936288 | 0.174166374 |
| 404,900 | 0.173757072 | 0.23884549 | 0.079964347 | 0.173579293 |
| 405,000 | 0.173684215 | 0.238871688 | 0.079313304 | 0.173345809 |
| 405,100 | 0.173793141 | 0.239395102 | 0.07931737 | 0.173332689 |
| 405,200 | 0.173641887 | 0.239666467 | 0.078485771 | 0.173648646 |
| 405,300 | 0.173593624 | 0.239332426 | 0.078336612 | 0.174047314 |
| 405,400 | 0.173593411 | 0.239045648 | 0.078767764 | 0.173568301 |
| 405,500 | 0.173553439 | 0.240239553 | 0.077935962 | 0.173610948 |
| 405,600 | 0.173504647 | 0.23995212 | 0.078509406 | 0.174067626 |
| 405,700 | 0.173430709 | 0.239495506 | 0.079170871 | 0.173929069 |
| 405,800 | 0.173564219 | 0.239463331 | 0.079322125 | 0.173967779 |
| 405,900 | 0.173401755 | 0.239505362 | 0.079091303 | 0.172965622 |
| 406,000 | 0.173505997 | 0.239925292 | 0.078980235 | 0.173615017 |
| 406,100 | 0.173372732 | 0.240449823 | 0.078729031 | 0.172997456 |
| 406,200 | 0.173306508 | 0.240530815 | 0.079201312 | 0.173241354 |
| 406,300 | 0.173333657 | 0.239989325 | 0.078382682 | 0.173457465 |
| 406,400 | 0.173434593 | 0.239382118 | 0.078536155 | 0.173874384 |
| 406,500 | 0.173237676 | 0.239097908 | 0.079041312 | 0.173772457 |
| 406,600 | 0.173208948 | 0.238933336 | 0.078998989 | 0.172289222 |
| 406,700 | 0.173249626 | 0.23916924 | 0.078838575 | 0.172713822 |
| 406,800 | 0.173279445 | 0.2389877 | 0.078450946 | 0.173169731 |
| 406,900 | 0.173109996 | 0.238874031 | 0.078470263 | 0.17272051 |
| 407,000 | 0.173139831 | 0.238203533 | 0.079594656 | 0.173409316 |
| 407,100 | 0.173214937 | 0.238691095 | 0.078963681 | 0.173168362 |
| 407,200 | 0.173167605 | 0.239076931 | 0.078059947 | 0.172936522 |
| 407,300 | 0.173196998 | 0.23873992 | 0.077810969 | 0.173148944 |
| 407,400 | 0.173150609 | 0.239212981 | 0.077821419 | 0.172487865 |
| 407,500 | 0.172950044 | 0.238491168 | 0.077768057 | 0.172877114 |
| 407,600 | 0.173135088 | 0.238939814 | 0.078048036 | 0.172709068 |

| | | | | |
|---|---|---|---|---|
| 407,700 | 0.172974711 | 0.238283872 | 0.076681491 | 0.172606529 |
| 407,800 | 0.173113269 | 0.238757135 | 0.078157799 | 0.173586862 |
| 407,900 | 0.173081221 | 0.238959948 | 0.077164981 | 0.17287444 |
| 408,000 | 0.173098055 | 0.239060622 | 0.077904587 | 0.172924322 |
| 408,100 | 0.173108134 | 0.239440382 | 0.077319598 | 0.172748444 |
| 408,200 | 0.17303248 | 0.23924527 | 0.077465967 | 0.172809773 |
| 408,300 | 0.173177963 | 0.23925432 | 0.078465072 | 0.173415592 |
| 408,400 | 0.173035116 | 0.23857442 | 0.077904632 | 0.173261626 |
| 408,500 | 0.173025272 | 0.238890038 | 0.078027985 | 0.173164752 |
| 408,600 | 0.17310835 | 0.238565451 | 0.077559642 | 0.173452549 |
| 408,700 | 0.173062092 | 0.239749295 | 0.077616585 | 0.172609065 |
| 408,800 | 0.173085254 | 0.239721837 | 0.078210919 | 0.172758299 |
| 408,900 | 0.173013191 | 0.239083558 | 0.077352974 | 0.17307497 |
| 409,000 | 0.173053968 | 0.239269819 | 0.07823654 | 0.173278578 |
| 409,100 | 0.173019134 | 0.239212098 | 0.077939872 | 0.172837228 |
| 409,200 | 0.172873713 | 0.238228268 | 0.077809908 | 0.173049913 |
| 409,300 | 0.172924422 | 0.23884669 | 0.07840581 | 0.172192807 |
| 409,400 | 0.173062054 | 0.238837738 | 0.077481461 | 0.17263751 |
| 409,500 | 0.172929178 | 0.239068322 | 0.07843256 | 0.173728344 |
| 409,600 | 0.172753803 | 0.238806266 | 0.078107582 | 0.172810406 |
| 409,700 | 0.172824466 | 0.238408191 | 0.07753983 | 0.17234228 |
| 409,800 | 0.172734975 | 0.238646029 | 0.077317502 | 0.172957612 |
| 409,900 | 0.172689647 | 0.237903319 | 0.077737134 | 0.172531514 |
| 410,000 | 0.172677413 | 0.238379925 | 0.077212689 | 0.171761389 |
| 410,100 | 0.172658535 | 0.238297815 | 0.077174282 | 0.172087194 |
| 410,200 | 0.172698334 | 0.2379466 | 0.076803443 | 0.172600391 |
| 410,300 | 0.172636885 | 0.237720822 | 0.076984247 | 0.172008987 |
| 410,400 | 0.172541582 | 0.237357199 | 0.076828108 | 0.171697568 |
| 410,500 | 0.172590597 | 0.237865562 | 0.076943068 | 0.171822217 |
| 410,600 | 0.17241029 | 0.237838122 | 0.076942449 | 0.171891287 |
| 410,700 | 0.172424425 | 0.237586243 | 0.07707979 | 0.171450965 |
| 410,800 | 0.172551961 | 0.238015155 | 0.076969561 | 0.171370783 |
| 410,900 | 0.172397721 | 0.23782197 | 0.077309435 | 0.17158841 |
| 411,000 | 0.172523033 | 0.238957229 | 0.076269844 | 0.171861862 |
| 411,100 | 0.172386995 | 0.238213542 | 0.075747512 | 0.172162903 |
| 411,200 | 0.172357058 | 0.238052996 | 0.076384183 | 0.172264309 |
| 411,300 | 0.172448409 | 0.238581115 | 0.075936441 | 0.172118983 |
| 411,400 | 0.172350002 | 0.238863777 | 0.076079488 | 0.171759441 |
| 411,500 | 0.172460719 | 0.239180994 | 0.07614159 | 0.172456817 |
| 411,600 | 0.17223717 | 0.238532349 | 0.075849747 | 0.171583608 |
| 411,700 | 0.17232216 | 0.238834977 | 0.076413113 | 0.171681066 |
| 411,800 | 0.17242179 | 0.238688783 | 0.075863532 | 0.172341451 |
| 411,900 | 0.172347065 | 0.238308486 | 0.076685102 | 0.17176568 |
| 412,000 | 0.172179944 | 0.238247027 | 0.076114309 | 0.17113181 |
| 412,100 | 0.17213884 | 0.238227765 | 0.076140118 | 0.170787295 |
| 412,200 | 0.172197164 | 0.238717233 | 0.076288293 | 0.17118927 |
| 412,300 | 0.172201889 | 0.238336165 | 0.07542358 | 0.170939007 |
| 412,400 | 0.172307648 | 0.238597731 | 0.07665146 | 0.171174957 |
| 412,500 | 0.172173201 | 0.238380161 | 0.076779879 | 0.171156208 |
| 412,600 | 0.172046378 | 0.238476309 | 0.076657015 | 0.170789323 |
| 412,700 | 0.172105637 | 0.238669559 | 0.076410845 | 0.170489025 |
| 412,800 | 0.171943259 | 0.238046791 | 0.076044186 | 0.170621853 |
| 412,900 | 0.172131544 | 0.238106021 | 0.076466026 | 0.170543378 |
| 413,000 | 0.172127372 | 0.238829123 | 0.076056355 | 0.170912004 |
| 413,100 | 0.172127106 | 0.238638734 | 0.075903721 | 0.170822623 |
| 413,200 | 0.172172559 | 0.23863351 | 0.0750945 | 0.171333675 |
| 413,300 | 0.17203684 | 0.238480859 | 0.075494034 | 0.171079719 |
| 413,400 | 0.172115625 | 0.23884514 | 0.074970894 | 0.170776348 |
| 413,500 | 0.172093352 | 0.238580113 | 0.075320373 | 0.170867846 |
| 413,600 | 0.172106288 | 0.238647073 | 0.076262699 | 0.170541779 |
| 413,700 | 0.172028119 | 0.238116724 | 0.075435705 | 0.170683664 |
| 413,800 | 0.171988103 | 0.237896533 | 0.075075628 | 0.170716581 |
| 413,900 | 0.172023796 | 0.236931231 | 0.075721009 | 0.170735529 |
| 414,000 | 0.171921509 | 0.238095886 | 0.075418863 | 0.170492792 |
| 414,100 | 0.171971374 | 0.237646867 | 0.074676647 | 0.170823741 |
| 414,200 | 0.172004335 | 0.238122507 | 0.07520492 | 0.170431634 |
| 414,300 | 0.171911789 | 0.238388931 | 0.075100052 | 0.170403239 |

| | | | | |
|---|---|---|---|---|
| 414,400 | 0.171936675 | 0.237440193 | 0.074434677 | 0.170388844 |
| 414,500 | 0.171937823 | 0.237825093 | 0.074870334 | 0.171141615 |
| 414,600 | 0.171759236 | 0.237918393 | 0.074738835 | 0.169993252 |
| 414,700 | 0.171902559 | 0.237031187 | 0.074849054 | 0.170460557 |
| 414,800 | 0.171942085 | 0.237609753 | 0.075491984 | 0.169402224 |
| 414,900 | 0.171847953 | 0.236227329 | 0.0750343 | 0.169954103 |
| 415,000 | 0.171914945 | 0.236168654 | 0.074078296 | 0.169872487 |
| 415,100 | 0.171879187 | 0.237119901 | 0.074452511 | 0.169742543 |
| 415,200 | 0.171889315 | 0.236792864 | 0.074577542 | 0.169798111 |
| 415,300 | 0.171787985 | 0.236949593 | 0.074739989 | 0.170091785 |
| 415,400 | 0.171833294 | 0.237772381 | 0.074255151 | 0.169446769 |
| 415,500 | 0.171843581 | 0.23805732 | 0.074010889 | 0.169173208 |
| 415,600 | 0.171683683 | 0.237142309 | 0.073681199 | 0.169532536 |
| 415,700 | 0.171658236 | 0.237862897 | 0.073392271 | 0.169107285 |
| 415,800 | 0.171761757 | 0.237610166 | 0.074048668 | 0.16890921 |
| 415,900 | 0.171650746 | 0.238332679 | 0.073342649 | 0.1694413 |
| 416,000 | 0.171560868 | 0.237755638 | 0.073926147 | 0.168967974 |
| 416,100 | 0.171619133 | 0.237484134 | 0.074304411 | 0.168595321 |
| 416,200 | 0.171655527 | 0.237292318 | 0.074975053 | 0.169243489 |
| 416,300 | 0.17165812 | 0.237131982 | 0.074789204 | 0.169442781 |
| 416,400 | 0.171682589 | 0.237900709 | 0.075076295 | 0.169467654 |
| 416,500 | 0.171777045 | 0.237804793 | 0.074755755 | 0.169853926 |
| 416,600 | 0.171608376 | 0.237344686 | 0.074145936 | 0.169632006 |
| 416,700 | 0.171573363 | 0.23684162 | 0.075245454 | 0.169615625 |
| 416,800 | 0.171584959 | 0.237383127 | 0.074887361 | 0.16883039 |
| 416,900 | 0.171465637 | 0.237528582 | 0.075038884 | 0.169164387 |
| 417,000 | 0.171493922 | 0.237088249 | 0.074745735 | 0.16909132 |
| 417,100 | 0.171275985 | 0.23771191 | 0.074423929 | 0.168324601 |
| 417,200 | 0.171291681 | 0.236808622 | 0.075696727 | 0.169515116 |
| 417,300 | 0.171410401 | 0.236592248 | 0.075142745 | 0.169076865 |
| 417,400 | 0.171321436 | 0.236668948 | 0.074934981 | 0.169215912 |
| 417,500 | 0.171322427 | 0.236943511 | 0.074318016 | 0.168868792 |
| 417,600 | 0.171368236 | 0.237371506 | 0.074586523 | 0.16911373 |
| 417,700 | 0.171478376 | 0.237559327 | 0.075360507 | 0.169630726 |
| 417,800 | 0.171308282 | 0.236553066 | 0.075453901 | 0.168814212 |
| 417,900 | 0.171334551 | 0.236236373 | 0.075196306 | 0.169243569 |
| 418,000 | 0.17128697 | 0.237434334 | 0.07555303 | 0.168528208 |
| 418,100 | 0.171296995 | 0.236798143 | 0.074850829 | 0.168533397 |
| 418,200 | 0.171190845 | 0.237019358 | 0.074559252 | 0.168645588 |
| 418,300 | 0.171255649 | 0.236431843 | 0.074527222 | 0.168645365 |
| 418,400 | 0.171156977 | 0.236869443 | 0.074763513 | 0.168995922 |
| 418,500 | 0.171101462 | 0.236126641 | 0.074321892 | 0.168460868 |
| 418,600 | 0.171149179 | 0.236624395 | 0.075070017 | 0.168478297 |
| 418,700 | 0.171077069 | 0.2358467 | 0.075257136 | 0.168229446 |
| 418,800 | 0.171188835 | 0.236267786 | 0.07544555 | 0.167843044 |
| 418,900 | 0.171146648 | 0.236695063 | 0.07482544 | 0.168234248 |
| 419,000 | 0.171115805 | 0.237230642 | 0.075488435 | 0.168069985 |
| 419,100 | 0.17100206 | 0.237047158 | 0.075845071 | 0.168601818 |
| 419,200 | 0.171081556 | 0.236732593 | 0.074752471 | 0.168117253 |
| 419,300 | 0.170979174 | 0.235883645 | 0.074366492 | 0.16841042 |
| 419,400 | 0.171086069 | 0.236513391 | 0.074326654 | 0.169091857 |
| 419,500 | 0.170910418 | 0.236768787 | 0.074322503 | 0.168601031 |
| 419,600 | 0.170967477 | 0.236326495 | 0.074274529 | 0.168709742 |
| 419,700 | 0.170907377 | 0.236541219 | 0.073836643 | 0.168175931 |
| 419,800 | 0.170865821 | 0.236543899 | 0.074746046 | 0.168113296 |
| 419,900 | 0.171001298 | 0.237012509 | 0.074346005 | 0.168049601 |
| 420,000 | 0.170801844 | 0.236397179 | 0.075141439 | 0.167620702 |
| 420,100 | 0.170793343 | 0.235863181 | 0.075011453 | 0.167486193 |
| 420,200 | 0.170808116 | 0.236893316 | 0.074792044 | 0.167594633 |
| 420,300 | 0.170692761 | 0.236555303 | 0.074466597 | 0.168011029 |
| 420,400 | 0.170710967 | 0.237011033 | 0.074148074 | 0.16801055 |
| 420,500 | 0.170785427 | 0.236265603 | 0.074105569 | 0.167498198 |
| 420,600 | 0.170754846 | 0.236386663 | 0.074548104 | 0.167630398 |
| 420,700 | 0.170802496 | 0.236967963 | 0.074181148 | 0.167935445 |
| 420,800 | 0.17088177 | 0.236944706 | 0.074878833 | 0.167535562 |
| 420,900 | 0.170821623 | 0.23660222 | 0.074407201 | 0.16742675 |
| 421,000 | 0.170827871 | 0.237099147 | 0.074142992 | 0.167874457 |

| | | | | |
|---|---|---|---|---|
| 421,100 | 0.170896061 | 0.236098732 | 0.074904379 | 0.168684214 |
| 421,200 | 0.170813981 | 0.235632634 | 0.074969353 | 0.168815079 |
| 421,300 | 0.170750162 | 0.236447472 | 0.074491322 | 0.168668646 |
| 421,400 | 0.17058988 | 0.236793257 | 0.074693468 | 0.168565923 |
| 421,500 | 0.170558775 | 0.235759609 | 0.074640744 | 0.168133 |
| 421,600 | 0.170517907 | 0.23595522 | 0.074241759 | 0.168328097 |
| 421,700 | 0.17055085 | 0.234983911 | 0.073899222 | 0.168100613 |
| 421,800 | 0.170513307 | 0.234381916 | 0.074268169 | 0.168738953 |
| 421,900 | 0.170498246 | 0.233895001 | 0.073768501 | 0.168496291 |
| 422,000 | 0.170505194 | 0.234620128 | 0.073849606 | 0.167596676 |
| 422,100 | 0.170430088 | 0.234608971 | 0.073436077 | 0.168084851 |
| 422,200 | 0.170412912 | 0.235497628 | 0.074052599 | 0.168044112 |
| 422,300 | 0.170471931 | 0.234553636 | 0.073370513 | 0.168367761 |
| 422,400 | 0.170430436 | 0.23489561 | 0.073503516 | 0.168277155 |
| 422,500 | 0.170271955 | 0.234895468 | 0.073053182 | 0.168847402 |
| 422,600 | 0.17041593 | 0.235316418 | 0.073457976 | 0.168339358 |
| 422,700 | 0.170398687 | 0.235181901 | 0.073794325 | 0.168297118 |
| 422,800 | 0.170365732 | 0.234929861 | 0.074606353 | 0.168807829 |
| 422,900 | 0.170411939 | 0.235135883 | 0.073443965 | 0.168631689 |
| 423,000 | 0.170392206 | 0.235103398 | 0.074507612 | 0.168361873 |
| 423,100 | 0.170374024 | 0.234475837 | 0.073342907 | 0.168983166 |
| 423,200 | 0.17034672 | 0.234529494 | 0.073483992 | 0.168125549 |
| 423,300 | 0.170358025 | 0.234396005 | 0.072764582 | 0.167691926 |
| 423,400 | 0.170354568 | 0.235384892 | 0.072497093 | 0.167764218 |
| 423,500 | 0.170251323 | 0.234783756 | 0.072852101 | 0.167544729 |
| 423,600 | 0.170454424 | 0.235301746 | 0.07314217 | 0.167345281 |
| 423,700 | 0.17037553 | 0.234629327 | 0.073733175 | 0.167911617 |
| 423,800 | 0.170426182 | 0.235009067 | 0.07384881 | 0.168251611 |
| 423,900 | 0.170254254 | 0.234041826 | 0.074020997 | 0.168078056 |
| 424,000 | 0.170378142 | 0.23446914 | 0.074410616 | 0.167974225 |
| 424,100 | 0.170177239 | 0.234077716 | 0.073545317 | 0.167871198 |
| 424,200 | 0.170239383 | 0.234511239 | 0.073785762 | 0.167515576 |
| 424,300 | 0.170016817 | 0.233737284 | 0.073792575 | 0.167388774 |
| 424,400 | 0.170069878 | 0.233688964 | 0.073443488 | 0.166899441 |
| 424,500 | 0.17014489 | 0.234019301 | 0.074249063 | 0.167206227 |
| 424,600 | 0.170072833 | 0.23359421 | 0.073177565 | 0.167351553 |
| 424,700 | 0.170158134 | 0.233788296 | 0.073018187 | 0.166925085 |
| 424,800 | 0.17016118 | 0.234725942 | 0.07308524 | 0.166993475 |
| 424,900 | 0.170036088 | 0.234262574 | 0.07351725 | 0.166727204 |
| 425,000 | 0.170093847 | 0.234318253 | 0.072668705 | 0.166732562 |
| 425,100 | 0.169974148 | 0.23398051 | 0.074063564 | 0.166801524 |
| 425,200 | 0.170059327 | 0.235521016 | 0.073415955 | 0.166384273 |
| 425,300 | 0.169905969 | 0.233938349 | 0.073041403 | 0.166678785 |
| 425,400 | 0.169781283 | 0.234169865 | 0.073472663 | 0.16631842 |
| 425,500 | 0.169949037 | 0.23437186 | 0.074153983 | 0.166923376 |
| 425,600 | 0.16997189 | 0.234268244 | 0.073529991 | 0.167328081 |
| 425,700 | 0.169817538 | 0.234313182 | 0.073172436 | 0.166335117 |
| 425,800 | 0.169901122 | 0.234893814 | 0.073055932 | 0.166794547 |
| 425,900 | 0.169956237 | 0.233893624 | 0.073150458 | 0.166430766 |
| 426,000 | 0.169736221 | 0.23345158 | 0.072486354 | 0.166525955 |
| 426,100 | 0.169714775 | 0.2335588 | 0.07318337 | 0.166294112 |
| 426,200 | 0.169881711 | 0.233681575 | 0.073288417 | 0.166649034 |
| 426,300 | 0.169777392 | 0.233285591 | 0.073817615 | 0.166591635 |
| 426,400 | 0.169845464 | 0.233644509 | 0.072897017 | 0.166396584 |
| 426,500 | 0.169560026 | 0.233464969 | 0.072315787 | 0.166353231 |
| 426,600 | 0.16972472 | 0.2330076 | 0.072636514 | 0.166024685 |
| 426,700 | 0.169657135 | 0.233129083 | 0.072859679 | 0.166005028 |
| 426,800 | 0.169605512 | 0.232697461 | 0.072398109 | 0.16602214 |
| 426,900 | 0.169658702 | 0.232714099 | 0.072563278 | 0.166244753 |
| 427,000 | 0.169558762 | 0.232649758 | 0.072392842 | 0.166235832 |
| 427,100 | 0.169516705 | 0.233555564 | 0.07357946 | 0.164887453 |
| 427,200 | 0.16962995 | 0.23341119 | 0.072972939 | 0.165602263 |
| 427,300 | 0.169618683 | 0.232758153 | 0.073734411 | 0.165924536 |
| 427,400 | 0.169725895 | 0.233568551 | 0.073771571 | 0.166334377 |
| 427,500 | 0.169649884 | 0.23361808 | 0.073827114 | 0.165923704 |
| 427,600 | 0.169624321 | 0.233002444 | 0.074469641 | 0.165565666 |
| 427,700 | 0.16956963 | 0.232514431 | 0.07338707 | 0.165953019 |

| | | | | |
|---|---|---|---|---|
| 427,800 | 0.169658037 | 0.232593645 | 0.073484141 | 0.166069994 |
| 427,900 | 0.169560586 | 0.232284456 | 0.07368704 | 0.165185834 |
| 428,000 | 0.16957503 | 0.232223653 | 0.073305676 | 0.166283921 |
| 428,100 | 0.169513218 | 0.231817652 | 0.073409043 | 0.165744078 |
| 428,200 | 0.169340884 | 0.232261282 | 0.073357833 | 0.165793095 |
| 428,300 | 0.169481161 | 0.232250509 | 0.072873137 | 0.165694058 |
| 428,400 | 0.169362322 | 0.231963201 | 0.073831963 | 0.165949111 |
| 428,500 | 0.169471344 | 0.232663998 | 0.073944963 | 0.166389723 |
| 428,600 | 0.169467573 | 0.231985952 | 0.073858893 | 0.166762328 |
| 428,700 | 0.169347988 | 0.231920848 | 0.073167312 | 0.166103477 |
| 428,800 | 0.169451259 | 0.232338706 | 0.074037419 | 0.165607411 |
| 428,900 | 0.169252339 | 0.23127447 | 0.074075021 | 0.165683817 |
| 429,000 | 0.169472911 | 0.231720907 | 0.074481641 | 0.165479876 |
| 429,100 | 0.169330496 | 0.232354558 | 0.074086802 | 0.166649507 |
| 429,200 | 0.169374305 | 0.231978802 | 0.074372551 | 0.16620886 |
| 429,300 | 0.169406984 | 0.231999439 | 0.073533054 | 0.166371063 |
| 429,400 | 0.169436676 | 0.231810979 | 0.07307798 | 0.166730411 |
| 429,500 | 0.169344835 | 0.231563297 | 0.072759712 | 0.166539042 |
| 429,600 | 0.169285608 | 0.232344465 | 0.072854791 | 0.16606105 |
| 429,700 | 0.169262058 | 0.232090164 | 0.073420766 | 0.166573754 |
| 429,800 | 0.169391496 | 0.231355966 | 0.073930223 | 0.16599418 |
| 429,900 | 0.169215418 | 0.23171807 | 0.074301414 | 0.165781698 |
| 430,000 | 0.169187356 | 0.231590559 | 0.073969694 | 0.165613484 |
| 430,100 | 0.169094227 | 0.230963871 | 0.07351087 | 0.166747469 |
| 430,200 | 0.169246383 | 0.232134284 | 0.074009292 | 0.166142713 |
| 430,300 | 0.169170017 | 0.232109287 | 0.073768325 | 0.166246059 |
| 430,400 | 0.169306307 | 0.231959773 | 0.074592063 | 0.166099983 |
| 430,500 | 0.169055635 | 0.230368233 | 0.074225044 | 0.165605769 |
| 430,600 | 0.169001972 | 0.231235518 | 0.073682452 | 0.165497264 |
| 430,700 | 0.168888361 | 0.230630326 | 0.073512989 | 0.165542965 |
| 430,800 | 0.168987919 | 0.231102731 | 0.073083564 | 0.16597816 |
| 430,900 | 0.168864469 | 0.23047963 | 0.074043095 | 0.165630772 |
| 431,000 | 0.168961021 | 0.231253617 | 0.073751725 | 0.165877737 |
| 431,100 | 0.168848213 | 0.230683434 | 0.072790225 | 0.165851072 |
| 431,200 | 0.168922343 | 0.230789637 | 0.072609471 | 0.165999375 |
| 431,300 | 0.168766944 | 0.230634454 | 0.073154102 | 0.165009306 |
| 431,400 | 0.16883122 | 0.231392029 | 0.072792074 | 0.165721383 |
| 431,500 | 0.168925855 | 0.230679352 | 0.072645966 | 0.165149934 |
| 431,600 | 0.168929646 | 0.230460781 | 0.073510156 | 0.165651344 |
| 431,700 | 0.168984504 | 0.230697565 | 0.072618411 | 0.165534653 |
| 431,800 | 0.168913023 | 0.230815476 | 0.073941842 | 0.16543444 |
| 431,900 | 0.168798546 | 0.230423406 | 0.073207642 | 0.165217901 |
| 432,000 | 0.168795878 | 0.230896189 | 0.07302393 | 0.165783639 |
| 432,100 | 0.169058974 | 0.230453157 | 0.07304407 | 0.166136535 |
| 432,200 | 0.168912603 | 0.23090159 | 0.072719549 | 0.165740736 |
| 432,300 | 0.168858909 | 0.230045359 | 0.073075582 | 0.166360969 |
| 432,400 | 0.168874991 | 0.230530602 | 0.072758743 | 0.165917396 |
| 432,500 | 0.168741431 | 0.230319849 | 0.072455632 | 0.16594753 |
| 432,600 | 0.168775957 | 0.230390604 | 0.072662522 | 0.166188138 |
| 432,700 | 0.168579059 | 0.229379882 | 0.073569232 | 0.164989307 |
| 432,800 | 0.168614403 | 0.229718311 | 0.07331876 | 0.165377264 |
| 432,900 | 0.168499696 | 0.229571924 | 0.074256537 | 0.165755128 |
| 433,000 | 0.168446619 | 0.228705406 | 0.07343891 | 0.165752923 |
| 433,100 | 0.168362704 | 0.229212506 | 0.073297611 | 0.165542808 |
| 433,200 | 0.16853012 | 0.229032019 | 0.074348545 | 0.165897243 |
| 433,300 | 0.168499891 | 0.228980152 | 0.073696894 | 0.16636104 |
| 433,400 | 0.168374904 | 0.228990476 | 0.073978318 | 0.165776643 |
| 433,500 | 0.168377963 | 0.228265587 | 0.074643845 | 0.16579461 |
| 433,600 | 0.168311467 | 0.228336934 | 0.073656401 | 0.165454835 |
| 433,700 | 0.168328289 | 0.228380164 | 0.073343901 | 0.166126252 |
| 433,800 | 0.168156602 | 0.227875698 | 0.07371656 | 0.165173568 |
| 433,900 | 0.168293101 | 0.229142418 | 0.07349293 | 0.165798238 |
| 434,000 | 0.16832325 | 0.228911405 | 0.073726305 | 0.166203168 |
| 434,100 | 0.168177549 | 0.228255574 | 0.073090345 | 0.16574599 |
| 434,200 | 0.168349825 | 0.22895616 | 0.073012392 | 0.165674398 |
| 434,300 | 0.168361681 | 0.228759383 | 0.07292897 | 0.165886996 |
| 434,400 | 0.168282406 | 0.228370219 | 0.072500689 | 0.165296342 |

| | | | | |
|---|---|---|---|---|
| 434,500 | 0.168210583 | 0.228482573 | 0.072419194 | 0.165948538 |
| 434,600 | 0.168180227 | 0.228625597 | 0.073513617 | 0.165248601 |
| 434,700 | 0.168109853 | 0.229151344 | 0.073727272 | 0.165267227 |
| 434,800 | 0.168191794 | 0.228586677 | 0.073283887 | 0.165446718 |
| 434,900 | 0.168129082 | 0.227770427 | 0.073032122 | 0.165717909 |
| 435,000 | 0.168153027 | 0.228346629 | 0.072449567 | 0.165453423 |
| 435,100 | 0.168157924 | 0.228176944 | 0.072103093 | 0.165595301 |
| 435,200 | 0.168205417 | 0.228891383 | 0.07279819 | 0.16536447 |
| 435,300 | 0.16813733 | 0.228877421 | 0.072920997 | 0.164998817 |
| 435,400 | 0.167987792 | 0.229028813 | 0.072678475 | 0.164833023 |
| 435,500 | 0.168071938 | 0.229309572 | 0.072829103 | 0.164337507 |
| 435,600 | 0.168063012 | 0.22814525 | 0.072974178 | 0.164480829 |
| 435,700 | 0.168051445 | 0.227976111 | 0.072419856 | 0.164668957 |
| 435,800 | 0.168021024 | 0.228206488 | 0.072558973 | 0.165173862 |
| 435,900 | 0.167991686 | 0.228807628 | 0.072749664 | 0.164804425 |
| 436,000 | 0.167861341 | 0.228528913 | 0.073710545 | 0.164423564 |
| 436,100 | 0.167861878 | 0.228708228 | 0.073312161 | 0.16480899 |
| 436,200 | 0.167899796 | 0.22839416 | 0.072386981 | 0.164345729 |
| 436,300 | 0.167967189 | 0.228862109 | 0.072382751 | 0.164316061 |
| 436,400 | 0.167892951 | 0.227972119 | 0.072875517 | 0.163935171 |
| 436,500 | 0.167635437 | 0.227615966 | 0.071734187 | 0.164426484 |
| 436,600 | 0.167693914 | 0.22799543 | 0.072899102 | 0.163302122 |
| 436,700 | 0.167768242 | 0.227987228 | 0.073374249 | 0.164165747 |
| 436,800 | 0.167886923 | 0.228542649 | 0.072741399 | 0.164488711 |
| 436,900 | 0.167730028 | 0.227027678 | 0.073435977 | 0.163742231 |
| 437,000 | 0.167896734 | 0.228347009 | 0.07317267 | 0.164251039 |
| 437,100 | 0.167901423 | 0.228633619 | 0.073664764 | 0.16433585 |
| 437,200 | 0.16788776 | 0.229490296 | 0.072972796 | 0.164756178 |
| 437,300 | 0.167832819 | 0.229500286 | 0.073413374 | 0.164310256 |
| 437,400 | 0.167823414 | 0.228571305 | 0.07436625 | 0.163809842 |
| 437,500 | 0.167848657 | 0.228980494 | 0.073560081 | 0.164504735 |
| 437,600 | 0.167933856 | 0.229325345 | 0.074137759 | 0.164194705 |
| 437,700 | 0.167753548 | 0.229012457 | 0.07369573 | 0.163770282 |
| 437,800 | 0.167720087 | 0.229473624 | 0.074155019 | 0.164380582 |
| 437,900 | 0.16785687 | 0.229421818 | 0.073112258 | 0.16434978 |
| 438,000 | 0.167803842 | 0.22852208 | 0.073637064 | 0.164230685 |
| 438,100 | 0.167715497 | 0.228749827 | 0.07418005 | 0.164159491 |
| 438,200 | 0.167739862 | 0.229737844 | 0.073725656 | 0.163899536 |
| 438,300 | 0.167732447 | 0.228339201 | 0.073790506 | 0.164068849 |
| 438,400 | 0.167680826 | 0.228758162 | 0.074082545 | 0.164693762 |
| 438,500 | 0.167651558 | 0.228750766 | 0.074176047 | 0.164554013 |
| 438,600 | 0.167645117 | 0.228377169 | 0.074709901 | 0.164112843 |
| 438,700 | 0.16751052 | 0.228009843 | 0.074019029 | 0.16409592 |
| 438,800 | 0.167618065 | 0.228858251 | 0.073576597 | 0.163967176 |
| 438,900 | 0.167486182 | 0.228818508 | 0.07428959 | 0.163641468 |
| 439,000 | 0.167523766 | 0.228939465 | 0.074226713 | 0.163636519 |
| 439,100 | 0.167502488 | 0.229506258 | 0.074164588 | 0.163300012 |
| 439,200 | 0.167499788 | 0.22900992 | 0.074728805 | 0.162786244 |
| 439,300 | 0.167599986 | 0.229534656 | 0.074329434 | 0.163561752 |
| 439,400 | 0.16762816 | 0.228836631 | 0.074134522 | 0.164172802 |
| 439,500 | 0.167588582 | 0.229207578 | 0.073193184 | 0.164089091 |
| 439,600 | 0.16754224 | 0.22928529 | 0.072959961 | 0.164157231 |
| 439,700 | 0.167619371 | 0.228958981 | 0.073402753 | 0.163731993 |
| 439,800 | 0.167542393 | 0.22932163 | 0.074129577 | 0.163765453 |
| 439,900 | 0.167694133 | 0.229572945 | 0.074360695 | 0.163858409 |
| 440,000 | 0.167540338 | 0.228987073 | 0.073954981 | 0.163760064 |
| 440,100 | 0.16741414 | 0.22922501 | 0.074259935 | 0.162999059 |
| 440,200 | 0.16739802 | 0.228669852 | 0.073521056 | 0.163199307 |
| 440,300 | 0.16745789 | 0.229002979 | 0.07356274 | 0.16319239 |
| 440,400 | 0.16735566 | 0.228199923 | 0.074301829 | 0.16384503 |
| 440,500 | 0.167334291 | 0.228408976 | 0.074053359 | 0.163517803 |
| 440,600 | 0.167445373 | 0.228199462 | 0.073620304 | 0.163461843 |
| 440,700 | 0.167371021 | 0.22878435 | 0.074365457 | 0.163828735 |
| 440,800 | 0.167301921 | 0.228033943 | 0.074236177 | 0.163521744 |
| 440,900 | 0.167126219 | 0.228329071 | 0.072479216 | 0.163503241 |
| 441,000 | 0.167232944 | 0.22763189 | 0.072918785 | 0.163403077 |
| 441,100 | 0.167109952 | 0.227110032 | 0.073523799 | 0.164091163 |

| | | | | |
|---|---|---|---|---|
| 441,200 | 0.167183281 | 0.227574655 | 0.0729362 | 0.164094151 |
| 441,300 | 0.167196169 | 0.227844148 | 0.07316229 | 0.164563632 |
| 441,400 | 0.167240306 | 0.227688754 | 0.073853742 | 0.164038157 |
| 441,500 | 0.167078279 | 0.227606434 | 0.073398319 | 0.164604139 |
| 441,600 | 0.167112992 | 0.227346215 | 0.073793343 | 0.164441239 |
| 441,700 | 0.166986815 | 0.227817801 | 0.074359524 | 0.163715035 |
| 441,800 | 0.16709189 | 0.227593795 | 0.074257558 | 0.164095304 |
| 441,900 | 0.167076965 | 0.227514254 | 0.074339337 | 0.164305209 |
| 442,000 | 0.167025019 | 0.227587505 | 0.074684373 | 0.164414906 |
| 442,100 | 0.166854955 | 0.226740719 | 0.074284441 | 0.164610618 |
| 442,200 | 0.166837756 | 0.227179612 | 0.073489195 | 0.16403912 |
| 442,300 | 0.166919258 | 0.227039086 | 0.073826995 | 0.164249915 |
| 442,400 | 0.166685314 | 0.226546545 | 0.072990995 | 0.163934921 |
| 442,500 | 0.16686767 | 0.227460456 | 0.073900367 | 0.164051654 |
| 442,600 | 0.166690692 | 0.226532976 | 0.073721555 | 0.163813601 |
| 442,700 | 0.166674386 | 0.226797718 | 0.074196062 | 0.163067673 |
| 442,800 | 0.166677991 | 0.226333574 | 0.074553446 | 0.16397399 |
| 442,900 | 0.166517366 | 0.225480502 | 0.073503413 | 0.16351096 |
| 443,000 | 0.16640194 | 0.225279075 | 0.07467106 | 0.162807472 |
| 443,100 | 0.166427567 | 0.225509308 | 0.073439243 | 0.163351146 |
| 443,200 | 0.166527186 | 0.224989114 | 0.07367779 | 0.16322582 |
| 443,300 | 0.166589985 | 0.224837971 | 0.073915572 | 0.163369614 |
| 443,400 | 0.166493759 | 0.224962331 | 0.073898439 | 0.163207389 |
| 443,500 | 0.166450518 | 0.2248926 | 0.074907551 | 0.162740741 |
| 443,600 | 0.166545727 | 0.225193136 | 0.0743194 | 0.163449043 |
| 443,700 | 0.166583731 | 0.225290341 | 0.074638333 | 0.163409685 |
| 443,800 | 0.166491784 | 0.225146147 | 0.074319672 | 0.163479324 |
| 443,900 | 0.166341321 | 0.22468416 | 0.074210579 | 0.163129023 |
| 444,000 | 0.166440059 | 0.224607466 | 0.075221549 | 0.163213633 |
| 444,100 | 0.166338041 | 0.225282246 | 0.074520938 | 0.163933401 |
| 444,200 | 0.166336771 | 0.225231807 | 0.074745285 | 0.16372176 |
| 444,300 | 0.16659341 | 0.225286237 | 0.074609122 | 0.164413236 |
| 444,400 | 0.166404059 | 0.225031618 | 0.074261624 | 0.164391802 |
| 444,500 | 0.166396258 | 0.225386261 | 0.075220062 | 0.1638981 |
| 444,600 | 0.166419286 | 0.225099034 | 0.075015447 | 0.162960875 |
| 444,700 | 0.166408739 | 0.224658193 | 0.075168622 | 0.163761371 |
| 444,800 | 0.166367879 | 0.225339254 | 0.074333508 | 0.163303671 |
| 444,900 | 0.166233383 | 0.224949342 | 0.074215845 | 0.163555189 |
| 445,000 | 0.166115905 | 0.225337052 | 0.074556045 | 0.163422802 |
| 445,100 | 0.16609472 | 0.224857256 | 0.075185711 | 0.163648616 |
| 445,200 | 0.1661807 | 0.225970753 | 0.075288255 | 0.163570608 |
| 445,300 | 0.166215025 | 0.225114622 | 0.07557914 | 0.163359626 |
| 445,400 | 0.166222985 | 0.226019387 | 0.073842477 | 0.162994392 |
| 445,500 | 0.166271406 | 0.225447036 | 0.073906567 | 0.16353276 |
| 445,600 | 0.166045552 | 0.224726676 | 0.074502902 | 0.163577006 |
| 445,700 | 0.166089693 | 0.224559195 | 0.074237142 | 0.163592581 |
| 445,800 | 0.166217194 | 0.22589592 | 0.074303733 | 0.163691459 |
| 445,900 | 0.16616879 | 0.225792367 | 0.074437708 | 0.164074025 |
| 446,000 | 0.166168021 | 0.225264112 | 0.074838496 | 0.163514273 |
| 446,100 | 0.166030161 | 0.225750753 | 0.075003634 | 0.163499275 |
| 446,200 | 0.166072382 | 0.225129216 | 0.074574272 | 0.164058605 |
| 446,300 | 0.165890022 | 0.225603724 | 0.074698588 | 0.164171766 |
| 446,400 | 0.166101325 | 0.225398197 | 0.0752374 | 0.163346989 |
| 446,500 | 0.165918728 | 0.226199648 | 0.075303332 | 0.163430263 |
| 446,600 | 0.165830498 | 0.225627204 | 0.074740809 | 0.163997777 |
| 446,700 | 0.165841752 | 0.225559497 | 0.074296558 | 0.164440308 |
| 446,800 | 0.165920703 | 0.225895957 | 0.074119024 | 0.163934511 |
| 446,900 | 0.165834394 | 0.226037583 | 0.074286715 | 0.163870646 |
| 447,000 | 0.165828925 | 0.225767072 | 0.075059259 | 0.164691694 |
| 447,100 | 0.165935622 | 0.226638099 | 0.07440519 | 0.165223506 |
| 447,200 | 0.165681109 | 0.226382467 | 0.073817231 | 0.164147455 |
| 447,300 | 0.165745071 | 0.224823248 | 0.074955411 | 0.164308726 |
| 447,400 | 0.165478189 | 0.225387552 | 0.074331586 | 0.164076769 |
| 447,500 | 0.16558912 | 0.224782178 | 0.07477189 | 0.1640461 |
| 447,600 | 0.165386957 | 0.225917209 | 0.074854107 | 0.163757318 |
| 447,700 | 0.165508521 | 0.224914273 | 0.074378112 | 0.163221826 |
| 447,800 | 0.165492205 | 0.2253507 | 0.074536177 | 0.163862007 |

| | | | | |
|---|---|---|---|---|
| 447,900 | 0.165673048 | 0.225153797 | 0.074510447 | 0.163706789 |
| 448,000 | 0.165460302 | 0.225099857 | 0.074565478 | 0.163705664 |
| 448,100 | 0.165436064 | 0.2254634 | 0.074662045 | 0.163537769 |
| 448,200 | 0.165493084 | 0.224853955 | 0.073989821 | 0.163303171 |
| 448,300 | 0.165369619 | 0.224912243 | 0.074599352 | 0.163605792 |
| 448,400 | 0.165405084 | 0.225631926 | 0.074448203 | 0.163427525 |
| 448,500 | 0.165373101 | 0.225259331 | 0.07397817 | 0.163505949 |
| 448,600 | 0.165572004 | 0.225398269 | 0.075093683 | 0.164215417 |
| 448,700 | 0.165402274 | 0.225534279 | 0.074268214 | 0.162749093 |
| 448,800 | 0.165407565 | 0.224973462 | 0.0747025 | 0.163558767 |
| 448,900 | 0.16549515 | 0.225920525 | 0.074924351 | 0.163543657 |
| 449,000 | 0.165398596 | 0.225905794 | 0.073931325 | 0.16382441 |
| 449,100 | 0.165406381 | 0.225507723 | 0.075015038 | 0.163703044 |
| 449,200 | 0.165616583 | 0.225867324 | 0.074972086 | 0.163121294 |
| 449,300 | 0.165474896 | 0.224951676 | 0.074339431 | 0.163706866 |
| 449,400 | 0.165405892 | 0.225581707 | 0.074347924 | 0.163463688 |
| 449,500 | 0.165438485 | 0.225786005 | 0.07423985 | 0.163735445 |
| 449,600 | 0.165486705 | 0.226023311 | 0.073968626 | 0.163790487 |
| 449,700 | 0.165231187 | 0.225655939 | 0.074473663 | 0.163722974 |
| 449,800 | 0.165428163 | 0.226416801 | 0.074234558 | 0.163540657 |
| 449,900 | 0.165410462 | 0.226595008 | 0.074511436 | 0.163970478 |
| 450,000 | 0.16537692 | 0.226289816 | 0.074223952 | 0.16315267 |
| 450,100 | 0.165331009 | 0.225604621 | 0.074293287 | 0.163029984 |
| 450,200 | 0.165264704 | 0.225611689 | 0.074107197 | 0.162719899 |
| 450,300 | 0.165261632 | 0.226308709 | 0.074588915 | 0.163244848 |
| 450,400 | 0.165379827 | 0.227238472 | 0.074480671 | 0.162977639 |
| 450,500 | 0.165307282 | 0.226393603 | 0.074047441 | 0.163587312 |
| 450,600 | 0.165522997 | 0.22719462 | 0.074407575 | 0.162717834 |
| 450,700 | 0.165438191 | 0.226710234 | 0.074216038 | 0.163128041 |
| 450,800 | 0.165290195 | 0.225960613 | 0.074437325 | 0.163126612 |
| 450,900 | 0.165337552 | 0.226547658 | 0.073894682 | 0.163994709 |
| 451,000 | 0.1652583 | 0.226049089 | 0.074195401 | 0.164158913 |
| 451,100 | 0.165363858 | 0.225709454 | 0.075188577 | 0.16386729 |
| 451,200 | 0.165468234 | 0.226174422 | 0.074485745 | 0.163647296 |
| 451,300 | 0.165359886 | 0.226352832 | 0.074302732 | 0.163848656 |
| 451,400 | 0.165290595 | 0.226119464 | 0.074509217 | 0.163361138 |
| 451,500 | 0.165259115 | 0.225491602 | 0.074666579 | 0.164474537 |
| 451,600 | 0.165135471 | 0.225434974 | 0.074281932 | 0.163704896 |
| 451,700 | 0.16514108 | 0.225825224 | 0.075464528 | 0.164348333 |
| 451,800 | 0.16528743 | 0.2255663 | 0.07496136 | 0.163546012 |
| 451,900 | 0.165174584 | 0.224756397 | 0.074418878 | 0.163008936 |
| 452,000 | 0.165285829 | 0.226076313 | 0.074358616 | 0.163675456 |
| 452,100 | 0.165113719 | 0.225344026 | 0.074663271 | 0.164269425 |
| 452,200 | 0.165281304 | 0.224805444 | 0.075168424 | 0.164124777 |
| 452,300 | 0.165072542 | 0.225111041 | 0.074752696 | 0.164126531 |
| 452,400 | 0.16507961 | 0.225627417 | 0.075597448 | 0.163793563 |
| 452,500 | 0.165278654 | 0.225580615 | 0.075250115 | 0.165041702 |
| 452,600 | 0.165031666 | 0.225035151 | 0.074807289 | 0.163760785 |
| 452,700 | 0.164931191 | 0.225610134 | 0.074830291 | 0.164050722 |
| 452,800 | 0.164982528 | 0.226482666 | 0.075036944 | 0.16404497 |
| 452,900 | 0.165053603 | 0.225598684 | 0.074356343 | 0.163783286 |
| 453,000 | 0.16513697 | 0.226161967 | 0.074812303 | 0.163741645 |
| 453,100 | 0.165165142 | 0.226290369 | 0.074514617 | 0.164410409 |
| 453,200 | 0.16510634 | 0.225362744 | 0.074303973 | 0.163416227 |
| 453,300 | 0.165036418 | 0.2259477 | 0.074860979 | 0.163445444 |
| 453,400 | 0.164988933 | 0.225675787 | 0.07420808 | 0.163576094 |
| 453,500 | 0.164917938 | 0.225339406 | 0.074884943 | 0.163388147 |
| 453,600 | 0.165063697 | 0.226179478 | 0.074593368 | 0.163684735 |
| 453,700 | 0.164948489 | 0.225513544 | 0.074674231 | 0.163447686 |
| 453,800 | 0.164991202 | 0.225832323 | 0.074350794 | 0.162804754 |
| 453,900 | 0.164921933 | 0.225461613 | 0.074633814 | 0.163101698 |
| 454,000 | 0.164947194 | 0.226223258 | 0.076071271 | 0.163471174 |
| 454,100 | 0.164824509 | 0.22612016 | 0.076326076 | 0.163106755 |
| 454,200 | 0.164779793 | 0.225914537 | 0.075089618 | 0.163200607 |
| 454,300 | 0.164992136 | 0.226249589 | 0.075269368 | 0.163022406 |
| 454,400 | 0.164738113 | 0.224114816 | 0.075068377 | 0.162918933 |
| 454,500 | 0.164764184 | 0.224572731 | 0.075469775 | 0.163342986 |

| | | | | |
|---|---|---|---|---|
| 454,600 | 0.164863544 | 0.224968291 | 0.075827639 | 0.163493171 |
| 454,700 | 0.164838066 | 0.225037173 | 0.074185393 | 0.162438346 |
| 454,800 | 0.164806478 | 0.224767657 | 0.074739651 | 0.162509649 |
| 454,900 | 0.164686389 | 0.224614991 | 0.075039115 | 0.162242949 |
| 455,000 | 0.164659711 | 0.224623477 | 0.075342264 | 0.16263881 |
| 455,100 | 0.164507799 | 0.223988156 | 0.074651588 | 0.162385872 |
| 455,200 | 0.164549444 | 0.224261128 | 0.074535389 | 0.162248637 |
| 455,300 | 0.164515797 | 0.22505649 | 0.074087149 | 0.162719653 |
| 455,400 | 0.16471972 | 0.224373445 | 0.075221643 | 0.162782202 |
| 455,500 | 0.164516754 | 0.224068253 | 0.074946165 | 0.163029047 |
| 455,600 | 0.164516529 | 0.224287407 | 0.075258823 | 0.162196968 |
| 455,700 | 0.164423585 | 0.223688502 | 0.074249612 | 0.161561539 |
| 455,800 | 0.164378277 | 0.223983901 | 0.07557369 | 0.161934925 |
| 455,900 | 0.16440625 | 0.223556036 | 0.074127652 | 0.16112816 |
| 456,000 | 0.164335693 | 0.224145044 | 0.074968434 | 0.160881031 |
| 456,100 | 0.16443329 | 0.223570353 | 0.075042475 | 0.161412219 |
| 456,200 | 0.164499083 | 0.223833825 | 0.075278818 | 0.161523271 |
| 456,300 | 0.164336883 | 0.223119264 | 0.075396151 | 0.161990377 |
| 456,400 | 0.16436519 | 0.224426952 | 0.075572234 | 0.162298136 |
| 456,500 | 0.16419966 | 0.224975507 | 0.074862374 | 0.160902201 |
| 456,600 | 0.164358024 | 0.224364778 | 0.075064757 | 0.161069697 |
| 456,700 | 0.164416288 | 0.224406898 | 0.074834869 | 0.161634294 |
| 456,800 | 0.16418205 | 0.224277776 | 0.074885733 | 0.161066207 |
| 456,900 | 0.16420588 | 0.224783123 | 0.074610468 | 0.161218208 |
| 457,000 | 0.164359442 | 0.225771257 | 0.07448705 | 0.161276263 |
| 457,100 | 0.164229123 | 0.224581371 | 0.075201636 | 0.161119681 |
| 457,200 | 0.164207073 | 0.224271309 | 0.075663007 | 0.161551358 |
| 457,300 | 0.164173007 | 0.224211269 | 0.074874837 | 0.160835898 |
| 457,400 | 0.163991373 | 0.224684489 | 0.075416773 | 0.161023767 |
| 457,500 | 0.164085001 | 0.223293058 | 0.074802884 | 0.161567584 |
| 457,600 | 0.164119571 | 0.224336928 | 0.075266417 | 0.161363925 |
| 457,700 | 0.164045671 | 0.223993289 | 0.075187859 | 0.162122323 |
| 457,800 | 0.163945768 | 0.223787198 | 0.074179372 | 0.161228756 |
| 457,900 | 0.164059298 | 0.224292257 | 0.075622484 | 0.161241907 |
| 458,000 | 0.164105419 | 0.22423025 | 0.074912643 | 0.160665643 |
| 458,100 | 0.163961945 | 0.223934908 | 0.074903144 | 0.161086729 |
| 458,200 | 0.164037774 | 0.224671883 | 0.075293976 | 0.160689879 |
| 458,300 | 0.163848843 | 0.224713251 | 0.075527465 | 0.160285338 |
| 458,400 | 0.163777711 | 0.225085022 | 0.074768649 | 0.161199481 |
| 458,500 | 0.16384043 | 0.224844976 | 0.074814085 | 0.160827892 |
| 458,600 | 0.163850214 | 0.224787014 | 0.074660795 | 0.160896426 |
| 458,700 | 0.163835306 | 0.224961056 | 0.074714087 | 0.160654767 |
| 458,800 | 0.163848383 | 0.224151146 | 0.074708027 | 0.161344136 |
| 458,900 | 0.16370987 | 0.223784959 | 0.07597521 | 0.160941198 |
| 459,000 | 0.163865089 | 0.223815226 | 0.075798663 | 0.161643327 |
| 459,100 | 0.163919549 | 0.224581316 | 0.07563238 | 0.160953771 |
| 459,200 | 0.16397535 | 0.224475851 | 0.075446716 | 0.160892027 |
| 459,300 | 0.163813255 | 0.224130831 | 0.075171868 | 0.160594696 |
| 459,400 | 0.163858577 | 0.224326412 | 0.075769365 | 0.161344843 |
| 459,500 | 0.163762587 | 0.223685198 | 0.075952769 | 0.160943893 |
| 459,600 | 0.163798815 | 0.223282117 | 0.075688149 | 0.160794695 |
| 459,700 | 0.163720965 | 0.2243789 | 0.075658409 | 0.160803555 |
| 459,800 | 0.163784728 | 0.224044081 | 0.075212934 | 0.160737897 |
| 459,900 | 0.163736448 | 0.224381043 | 0.07455427 | 0.16057833 |
| 460,000 | 0.163809631 | 0.224344069 | 0.074653882 | 0.160781693 |
| 460,100 | 0.163684218 | 0.22402914 | 0.075582212 | 0.160360001 |
| 460,200 | 0.163789871 | 0.224254843 | 0.075913721 | 0.16061981 |
| 460,300 | 0.163660935 | 0.223631061 | 0.076087567 | 0.160321106 |
| 460,400 | 0.163704831 | 0.224167707 | 0.075392931 | 0.160021831 |
| 460,500 | 0.163643694 | 0.22437222 | 0.075109661 | 0.160047955 |
| 460,600 | 0.163625455 | 0.223738764 | 0.075162697 | 0.16080088 |
| 460,700 | 0.163734652 | 0.225171804 | 0.075441398 | 0.160879498 |
| 460,800 | 0.163715678 | 0.224475882 | 0.074982816 | 0.161360439 |
| 460,900 | 0.163731846 | 0.224505675 | 0.075832854 | 0.160788165 |
| 461,000 | 0.163550946 | 0.224536058 | 0.074621311 | 0.160060947 |
| 461,100 | 0.163603399 | 0.224771149 | 0.07523863 | 0.160543241 |
| 461,200 | 0.163548094 | 0.225333758 | 0.075598773 | 0.159741331 |

| | | | | |
|---|---|---|---|---|
| 461,300 | 0.163635668 | 0.224678904 | 0.076310215 | 0.159990853 |
| 461,400 | 0.163622986 | 0.225292736 | 0.075926167 | 0.160047317 |
| 461,500 | 0.163628967 | 0.226174406 | 0.075360436 | 0.159595053 |
| 461,600 | 0.163656686 | 0.225528189 | 0.075277467 | 0.160193374 |
| 461,700 | 0.163593407 | 0.225568333 | 0.07413967 | 0.159689773 |
| 461,800 | 0.16358036 | 0.225119473 | 0.074445097 | 0.159776137 |
| 461,900 | 0.163417107 | 0.225568097 | 0.074143632 | 0.159730633 |
| 462,000 | 0.163450061 | 0.225264999 | 0.074041547 | 0.159276809 |
| 462,100 | 0.163545087 | 0.226154563 | 0.07441849 | 0.159352837 |
| 462,200 | 0.163575794 | 0.22623523 | 0.074745888 | 0.159592932 |
| 462,300 | 0.163547552 | 0.225517606 | 0.074412051 | 0.159809418 |
| 462,400 | 0.163399579 | 0.225163544 | 0.074166208 | 0.159545034 |
| 462,500 | 0.163602858 | 0.225289596 | 0.074713044 | 0.160243747 |
| 462,600 | 0.163422752 | 0.224860013 | 0.0740086 | 0.159883102 |
| 462,700 | 0.163425973 | 0.224454975 | 0.074390081 | 0.160064073 |
| 462,800 | 0.163246396 | 0.224684695 | 0.074199466 | 0.160295616 |
| 462,900 | 0.163260733 | 0.225169759 | 0.074101562 | 0.15987922 |
| 463,000 | 0.163349258 | 0.224574935 | 0.074999065 | 0.15996158 |
| 463,100 | 0.163306419 | 0.22413936 | 0.07333001 | 0.159328204 |
| 463,200 | 0.163228982 | 0.224481884 | 0.07416744 | 0.159495876 |
| 463,300 | 0.163298715 | 0.225200212 | 0.07363268 | 0.159651429 |
| 463,400 | 0.163164159 | 0.224290818 | 0.073460728 | 0.15926282 |
| 463,500 | 0.163258124 | 0.22431924 | 0.074268176 | 0.159423925 |
| 463,600 | 0.163276457 | 0.224103703 | 0.073620732 | 0.16014502 |
| 463,700 | 0.163392734 | 0.224468018 | 0.073843757 | 0.160143048 |
| 463,800 | 0.163256497 | 0.224275337 | 0.073763782 | 0.159669787 |
| 463,900 | 0.163120465 | 0.223930289 | 0.074590664 | 0.159658106 |
| 464,000 | 0.16309676 | 0.224601743 | 0.07434998 | 0.159806397 |
| 464,100 | 0.163096944 | 0.223690172 | 0.073890925 | 0.16001166 |
| 464,200 | 0.163012687 | 0.223720353 | 0.074039201 | 0.16019672 |
| 464,300 | 0.163012327 | 0.224209379 | 0.074465951 | 0.160390604 |
| 464,400 | 0.162977611 | 0.224176628 | 0.073840067 | 0.159745737 |
| 464,500 | 0.163052191 | 0.224259325 | 0.073079508 | 0.159212017 |
| 464,600 | 0.162947055 | 0.223938409 | 0.07321176 | 0.159167475 |
| 464,700 | 0.162811764 | 0.223729571 | 0.07280151 | 0.159075809 |
| 464,800 | 0.162839891 | 0.223487417 | 0.072762453 | 0.159106754 |
| 464,900 | 0.162896708 | 0.223320444 | 0.072779047 | 0.159326553 |
| 465,000 | 0.16287131 | 0.223458744 | 0.073347436 | 0.159685274 |
| 465,100 | 0.162816305 | 0.223867437 | 0.074525815 | 0.159661018 |
| 465,200 | 0.162797566 | 0.222545814 | 0.073382482 | 0.15937994 |
| 465,300 | 0.162738052 | 0.222442641 | 0.073572046 | 0.160019102 |
| 465,400 | 0.162921806 | 0.223527476 | 0.07334881 | 0.159473214 |
| 465,500 | 0.162816502 | 0.22352641 | 0.073498966 | 0.159165472 |
| 465,600 | 0.162675256 | 0.222959152 | 0.073634289 | 0.159429615 |
| 465,700 | 0.162872412 | 0.223947073 | 0.073328007 | 0.159143961 |
| 465,800 | 0.162878596 | 0.222617675 | 0.073587395 | 0.159542809 |
| 465,900 | 0.162756971 | 0.222708796 | 0.072850057 | 0.159863196 |
| 466,000 | 0.162818181 | 0.222776107 | 0.073004999 | 0.159826133 |
| 466,100 | 0.162811343 | 0.222725712 | 0.073567742 | 0.160144328 |
| 466,200 | 0.162693988 | 0.221798787 | 0.073886544 | 0.159767811 |
| 466,300 | 0.162800667 | 0.222618379 | 0.073512672 | 0.159868607 |
| 466,400 | 0.16272816 | 0.222559023 | 0.073043959 | 0.159576603 |
| 466,500 | 0.16275761 | 0.223060335 | 0.073634443 | 0.159952164 |
| 466,600 | 0.162761618 | 0.222823044 | 0.073727341 | 0.1602979 |
| 466,700 | 0.16262504 | 0.222669576 | 0.073161346 | 0.160172387 |
| 466,800 | 0.162636716 | 0.222699586 | 0.072718945 | 0.160705803 |
| 466,900 | 0.162678682 | 0.222326749 | 0.073503679 | 0.160532003 |
| 467,000 | 0.162521542 | 0.222556694 | 0.073137319 | 0.16088551 |
| 467,100 | 0.162489561 | 0.222923382 | 0.072716784 | 0.160428057 |
| 467,200 | 0.16255032 | 0.222137865 | 0.073304525 | 0.160403772 |
| 467,300 | 0.162518012 | 0.22272815 | 0.073317738 | 0.160272187 |
| 467,400 | 0.162446123 | 0.222310332 | 0.07252035 | 0.160437968 |
| 467,500 | 0.162574762 | 0.222953488 | 0.073012011 | 0.161268626 |
| 467,600 | 0.162442409 | 0.222536737 | 0.072354398 | 0.160594307 |
| 467,700 | 0.162510628 | 0.222257589 | 0.072511689 | 0.160550883 |
| 467,800 | 0.162398113 | 0.221430074 | 0.072880996 | 0.16053007 |
| 467,900 | 0.162364249 | 0.221549335 | 0.072692864 | 0.160369873 |

| | | | | |
|---|---|---|---|---|
| 468,000 | 0.162414441 | 0.221776538 | 0.072353434 | 0.160503168 |
| 468,100 | 0.162430177 | 0.221889388 | 0.072054438 | 0.160948541 |
| 468,200 | 0.162259709 | 0.222129634 | 0.071660105 | 0.160540844 |
| 468,300 | 0.162293659 | 0.221521338 | 0.072142785 | 0.16039906 |
| 468,400 | 0.162280581 | 0.221599117 | 0.072831446 | 0.160338838 |
| 468,500 | 0.162366878 | 0.222352987 | 0.072287591 | 0.160722096 |
| 468,600 | 0.162255485 | 0.222262086 | 0.072246046 | 0.160568504 |
| 468,700 | 0.162091588 | 0.221450949 | 0.073133388 | 0.160419163 |
| 468,800 | 0.162237539 | 0.221887424 | 0.072870877 | 0.160470682 |
| 468,900 | 0.162127759 | 0.222225613 | 0.072648727 | 0.160065111 |
| 469,000 | 0.162159497 | 0.222070716 | 0.072251417 | 0.161113745 |
| 469,100 | 0.162024738 | 0.22203151 | 0.073255425 | 0.159802687 |
| 469,200 | 0.16216177 | 0.222308841 | 0.073567491 | 0.160503537 |
| 469,300 | 0.162081025 | 0.22233949 | 0.072371622 | 0.160329072 |
| 469,400 | 0.161989196 | 0.222026143 | 0.07231827 | 0.160766339 |
| 469,500 | 0.161971058 | 0.221698054 | 0.072822523 | 0.15972584 |
| 469,600 | 0.161876853 | 0.221920405 | 0.073264637 | 0.160596809 |
| 469,700 | 0.161864557 | 0.222023525 | 0.072929355 | 0.160951146 |
| 469,800 | 0.161918966 | 0.221654193 | 0.072432479 | 0.160629205 |
| 469,900 | 0.16194115 | 0.221430176 | 0.073253728 | 0.161161493 |
| 470,000 | 0.161888878 | 0.221999943 | 0.072472831 | 0.160332343 |
| 470,100 | 0.161799806 | 0.221953721 | 0.072616992 | 0.160776291 |
| 470,200 | 0.161711935 | 0.221674685 | 0.072848176 | 0.160887682 |
| 470,300 | 0.161886122 | 0.221163916 | 0.072391898 | 0.160503955 |
| 470,400 | 0.161889923 | 0.221994436 | 0.072369831 | 0.161062802 |
| 470,500 | 0.16190658 | 0.22162276 | 0.072757012 | 0.1614723 |
| 470,600 | 0.161837541 | 0.221065061 | 0.072939274 | 0.160549848 |
| 470,700 | 0.161814933 | 0.221444191 | 0.072970613 | 0.160510645 |
| 470,800 | 0.161773923 | 0.220976536 | 0.072282345 | 0.160427638 |
| 470,900 | 0.161635572 | 0.22141508 | 0.072012031 | 0.160851693 |
| 471,000 | 0.161640912 | 0.220737389 | 0.072352379 | 0.160571937 |
| 471,100 | 0.16160546 | 0.22129913 | 0.072495184 | 0.160199194 |
| 471,200 | 0.161730463 | 0.222236931 | 0.07279849 | 0.160894397 |
| 471,300 | 0.161572277 | 0.222505885 | 0.072908347 | 0.160520399 |
| 471,400 | 0.161551188 | 0.22185988 | 0.072828962 | 0.159813562 |
| 471,500 | 0.161665624 | 0.222415667 | 0.072515861 | 0.160204546 |
| 471,600 | 0.16166076 | 0.222322083 | 0.072263957 | 0.160915213 |
| 471,700 | 0.161596015 | 0.222313695 | 0.072388918 | 0.160587955 |
| 471,800 | 0.1617013 | 0.222185107 | 0.07286917 | 0.160821081 |
| 471,900 | 0.161667733 | 0.22148834 | 0.072252443 | 0.161108077 |
| 472,000 | 0.161593359 | 0.222612551 | 0.072598042 | 0.160424689 |
| 472,100 | 0.161580546 | 0.222475951 | 0.071486211 | 0.160379108 |
| 472,200 | 0.161550684 | 0.222027847 | 0.072608831 | 0.159932657 |
| 472,300 | 0.161511341 | 0.222617242 | 0.071494763 | 0.160369996 |
| 472,400 | 0.161515732 | 0.222363792 | 0.071717839 | 0.161300403 |
| 472,500 | 0.161550473 | 0.222291959 | 0.071938017 | 0.160962195 |
| 472,600 | 0.161524416 | 0.222693436 | 0.072435109 | 0.160935317 |
| 472,700 | 0.161527363 | 0.222970331 | 0.073278418 | 0.160645801 |
| 472,800 | 0.161552452 | 0.223148252 | 0.072812608 | 0.160539755 |
| 472,900 | 0.161511984 | 0.222607371 | 0.072050622 | 0.160565877 |
| 473,000 | 0.161608378 | 0.223279416 | 0.071585249 | 0.160756133 |
| 473,100 | 0.161469494 | 0.221494642 | 0.071435117 | 0.160297208 |
| 473,200 | 0.161596446 | 0.222075888 | 0.07177738 | 0.160738856 |
| 473,300 | 0.161324865 | 0.222320573 | 0.071497053 | 0.160647874 |
| 473,400 | 0.161290846 | 0.222368756 | 0.07226766 | 0.160718816 |
| 473,500 | 0.161300477 | 0.222963154 | 0.071893937 | 0.160394535 |
| 473,600 | 0.16128289 | 0.222543744 | 0.071530883 | 0.160064203 |
| 473,700 | 0.161244083 | 0.221848827 | 0.071182305 | 0.160704458 |
| 473,800 | 0.161196682 | 0.222415562 | 0.071298185 | 0.160469438 |
| 473,900 | 0.16128799 | 0.222436864 | 0.071150018 | 0.160755285 |
| 474,000 | 0.161242366 | 0.221788682 | 0.070607827 | 0.160832986 |
| 474,100 | 0.16136905 | 0.222982623 | 0.070828892 | 0.161098467 |
| 474,200 | 0.161313372 | 0.223468962 | 0.071244202 | 0.16080459 |
| 474,300 | 0.161354315 | 0.222895371 | 0.071316584 | 0.160441979 |
| 474,400 | 0.161112148 | 0.222743589 | 0.071927998 | 0.160743886 |
| 474,500 | 0.16121157 | 0.22273482 | 0.071590435 | 0.160075174 |
| 474,600 | 0.161224927 | 0.222892093 | 0.072449024 | 0.16007393 |

| | | | | |
|---|---|---|---|---|
| 474,700 | 0.161232022 | 0.221896559 | 0.071153414 | 0.160117778 |
| 474,800 | 0.161153944 | 0.222479229 | 0.07142382 | 0.159875914 |
| 474,900 | 0.161312971 | 0.222331947 | 0.071747203 | 0.160133198 |
| 475,000 | 0.161207556 | 0.222949264 | 0.072187492 | 0.159290368 |
| 475,100 | 0.161142269 | 0.223381221 | 0.072133019 | 0.158880313 |
| 475,200 | 0.161111628 | 0.222564628 | 0.072017643 | 0.159328487 |
| 475,300 | 0.161167553 | 0.22273349 | 0.071919277 | 0.159913224 |
| 475,400 | 0.161060595 | 0.222080014 | 0.071740431 | 0.160057387 |
| 475,500 | 0.161201564 | 0.223568319 | 0.071896902 | 0.159700855 |
| 475,600 | 0.161221838 | 0.223585877 | 0.071917424 | 0.160607801 |
| 475,700 | 0.161352493 | 0.222950621 | 0.072025679 | 0.1600426 |
| 475,800 | 0.161174189 | 0.223383569 | 0.072019972 | 0.160257924 |
| 475,900 | 0.161105314 | 0.223020249 | 0.070909363 | 0.159819702 |
| 476,000 | 0.161084818 | 0.223448531 | 0.071331267 | 0.160838108 |
| 476,100 | 0.161030048 | 0.222777644 | 0.072366481 | 0.160350662 |
| 476,200 | 0.161095814 | 0.222683955 | 0.071890617 | 0.159894953 |
| 476,300 | 0.161033583 | 0.223332105 | 0.071079221 | 0.160280661 |
| 476,400 | 0.161008947 | 0.222921591 | 0.071149114 | 0.159853452 |
| 476,500 | 0.161129609 | 0.223890656 | 0.070772175 | 0.160009927 |
| 476,600 | 0.160961851 | 0.223008366 | 0.071659001 | 0.159923872 |
| 476,700 | 0.160922986 | 0.22276579 | 0.071738907 | 0.160022716 |
| 476,800 | 0.161025073 | 0.222820158 | 0.072027767 | 0.160466291 |
| 476,900 | 0.161074783 | 0.222767115 | 0.072086286 | 0.159959596 |
| 477,000 | 0.160988432 | 0.222974567 | 0.070438304 | 0.160136877 |
| 477,100 | 0.160879029 | 0.222917663 | 0.0718881 | 0.159991185 |
| 477,200 | 0.160831141 | 0.222305649 | 0.071731044 | 0.159835186 |
| 477,300 | 0.160932209 | 0.22201091 | 0.071754577 | 0.15977176 |
| 477,400 | 0.160854343 | 0.222511099 | 0.071515291 | 0.1599443 |
| 477,500 | 0.160950927 | 0.222682103 | 0.071878488 | 0.15963577 |
| 477,600 | 0.1608031 | 0.222687755 | 0.071932514 | 0.159991028 |
| 477,700 | 0.160918724 | 0.222865912 | 0.071904207 | 0.159963493 |
| 477,800 | 0.160737035 | 0.222808266 | 0.071651106 | 0.159840855 |
| 477,900 | 0.160731571 | 0.222645014 | 0.07239723 | 0.159333007 |
| 478,000 | 0.160713663 | 0.222399257 | 0.072142441 | 0.159760945 |
| 478,100 | 0.160642056 | 0.2220436 | 0.072052407 | 0.159996208 |
| 478,200 | 0.160609469 | 0.222120397 | 0.071007999 | 0.159933752 |
| 478,300 | 0.160598111 | 0.222067855 | 0.071734103 | 0.160012975 |
| 478,400 | 0.160663869 | 0.222852932 | 0.071752173 | 0.159369633 |
| 478,500 | 0.160535042 | 0.223104939 | 0.071670983 | 0.159011376 |
| 478,600 | 0.16046849 | 0.222776143 | 0.07160936 | 0.158903189 |
| 478,700 | 0.160552368 | 0.223203717 | 0.071627871 | 0.159451206 |
| 478,800 | 0.160535656 | 0.222089883 | 0.071317508 | 0.159471299 |
| 478,900 | 0.160410264 | 0.221855652 | 0.071629021 | 0.159011871 |
| 479,000 | 0.160354758 | 0.22199273 | 0.071491624 | 0.158564263 |
| 479,100 | 0.160427917 | 0.2224089 | 0.07152725 | 0.159085248 |
| 479,200 | 0.160398018 | 0.221704381 | 0.07143065 | 0.159213178 |
| 479,300 | 0.160425709 | 0.221995705 | 0.071291498 | 0.15954807 |
| 479,400 | 0.160161641 | 0.221332143 | 0.070874638 | 0.159301488 |
| 479,500 | 0.160204288 | 0.221458598 | 0.071000301 | 0.159183187 |
| 479,600 | 0.160243575 | 0.221513495 | 0.070955436 | 0.159120688 |
| 479,700 | 0.160363249 | 0.221561474 | 0.070708217 | 0.158796967 |
| 479,800 | 0.160203354 | 0.22083056 | 0.071116466 | 0.159478017 |
| 479,900 | 0.160272705 | 0.221502175 | 0.070465914 | 0.159487931 |
| 480,000 | 0.160390089 | 0.221762125 | 0.07139338 | 0.159144521 |
| 480,100 | 0.160258778 | 0.221704259 | 0.071706049 | 0.159263573 |
| 480,200 | 0.160263545 | 0.221924342 | 0.071096822 | 0.159174774 |
| 480,300 | 0.160163519 | 0.221132674 | 0.071690812 | 0.1586084 |
| 480,400 | 0.160209381 | 0.221653167 | 0.071769618 | 0.159253282 |
| 480,500 | 0.160199023 | 0.22115258 | 0.071185924 | 0.158791665 |
| 480,600 | 0.160036194 | 0.221207879 | 0.070477893 | 0.15849858 |
| 480,700 | 0.160107357 | 0.220812663 | 0.070248089 | 0.158782526 |
| 480,800 | 0.160078616 | 0.220490554 | 0.070886755 | 0.158650131 |
| 480,900 | 0.16005804 | 0.221299086 | 0.070731538 | 0.158207751 |
| 481,000 | 0.160040781 | 0.221025482 | 0.070387825 | 0.158789391 |
| 481,100 | 0.160128999 | 0.21988982 | 0.070925368 | 0.158712891 |
| 481,200 | 0.159945063 | 0.22082535 | 0.070807846 | 0.158527158 |
| 481,300 | 0.159887339 | 0.220296036 | 0.070445481 | 0.158485543 |

| | | | | |
|---|---|---|---|---|
| 481,400 | 0.15998906 | 0.220617506 | 0.070213076 | 0.15847862 |
| 481,500 | 0.15991593 | 0.220698314 | 0.070491523 | 0.158792148 |
| 481,600 | 0.159835778 | 0.220232913 | 0.070230559 | 0.158220065 |
| 481,700 | 0.159963582 | 0.219982694 | 0.069849995 | 0.159040724 |
| 481,800 | 0.159896406 | 0.219966234 | 0.06967085 | 0.159036765 |
| 481,900 | 0.159999523 | 0.219540477 | 0.070527193 | 0.1592625 |
| 482,000 | 0.159847552 | 0.220693472 | 0.070613608 | 0.158813814 |
| 482,100 | 0.15987262 | 0.220023107 | 0.070439729 | 0.15893206 |
| 482,200 | 0.159766054 | 0.220007889 | 0.071305963 | 0.159924942 |
| 482,300 | 0.159855202 | 0.220820167 | 0.070503607 | 0.158864248 |
| 482,400 | 0.159872091 | 0.220238402 | 0.071170021 | 0.159598833 |
| 482,500 | 0.159678704 | 0.219570182 | 0.07055548 | 0.159420478 |
| 482,600 | 0.159755655 | 0.219974215 | 0.071544565 | 0.159265462 |
| 482,700 | 0.159758001 | 0.220164491 | 0.071392046 | 0.159302618 |
| 482,800 | 0.159742084 | 0.220269977 | 0.070757615 | 0.159472666 |
| 482,900 | 0.15972718 | 0.220767235 | 0.071056851 | 0.158771713 |
| 483,000 | 0.159751163 | 0.221098862 | 0.070878339 | 0.159257902 |
| 483,100 | 0.159667147 | 0.221182641 | 0.070574049 | 0.159609504 |
| 483,200 | 0.159769273 | 0.22068547 | 0.070912127 | 0.159355092 |
| 483,300 | 0.159754322 | 0.221329939 | 0.070884106 | 0.159267022 |
| 483,400 | 0.159576457 | 0.220819316 | 0.070361048 | 0.158889843 |
| 483,500 | 0.159757685 | 0.220730119 | 0.070055784 | 0.159604924 |
| 483,600 | 0.159595732 | 0.220093021 | 0.070252637 | 0.159763829 |
| 483,700 | 0.159551775 | 0.220269579 | 0.06978129 | 0.159476939 |
| 483,800 | 0.159540308 | 0.220069851 | 0.069700825 | 0.159096702 |
| 483,900 | 0.159614157 | 0.220318455 | 0.069824161 | 0.159233902 |
| 484,000 | 0.159455716 | 0.219859294 | 0.069243603 | 0.159746095 |
| 484,100 | 0.159556423 | 0.221012299 | 0.069421432 | 0.159897084 |
| 484,200 | 0.159585503 | 0.221363935 | 0.06973755 | 0.15974663 |
| 484,300 | 0.159408721 | 0.220588858 | 0.07014813 | 0.159944729 |
| 484,400 | 0.159365899 | 0.220545894 | 0.069929418 | 0.159906143 |
| 484,500 | 0.159462283 | 0.220827769 | 0.069415553 | 0.160176386 |
| 484,600 | 0.159399937 | 0.219899799 | 0.069180372 | 0.160007037 |
| 484,700 | 0.159421559 | 0.219865738 | 0.069836224 | 0.159131253 |
| 484,800 | 0.159382846 | 0.219809168 | 0.069969074 | 0.159397208 |
| 484,900 | 0.159433892 | 0.220547697 | 0.070110875 | 0.159657405 |
| 485,000 | 0.159302521 | 0.220446276 | 0.070076574 | 0.158826813 |
| 485,100 | 0.15923177 | 0.220211971 | 0.0699834 | 0.159369268 |
| 485,200 | 0.159323005 | 0.220413052 | 0.069201547 | 0.159488841 |
| 485,300 | 0.159448018 | 0.220433814 | 0.070259664 | 0.158962661 |
| 485,400 | 0.159412972 | 0.220759828 | 0.070809454 | 0.158778876 |
| 485,500 | 0.159480214 | 0.220223882 | 0.069969969 | 0.158980871 |
| 485,600 | 0.159314572 | 0.220479291 | 0.069973632 | 0.158961548 |
| 485,700 | 0.159540216 | 0.220846644 | 0.070148225 | 0.158851577 |
| 485,800 | 0.159561207 | 0.220503847 | 0.069616013 | 0.158897097 |
| 485,900 | 0.159421954 | 0.220151801 | 0.069883976 | 0.159402548 |
| 486,000 | 0.159313773 | 0.22008046 | 0.069395419 | 0.1584281 |
| 486,100 | 0.159416237 | 0.22034816 | 0.069999903 | 0.15922159 |
| 486,200 | 0.159230332 | 0.220679833 | 0.068957374 | 0.158353237 |
| 486,300 | 0.159355669 | 0.220447399 | 0.069703036 | 0.158800931 |
| 486,400 | 0.159292291 | 0.219276691 | 0.07057483 | 0.159373839 |
| 486,500 | 0.159224532 | 0.219605749 | 0.069905516 | 0.159131428 |
| 486,600 | 0.159233566 | 0.219400107 | 0.069621504 | 0.158535847 |
| 486,700 | 0.159229022 | 0.219316943 | 0.070510903 | 0.159215148 |
| 486,800 | 0.159323207 | 0.220124673 | 0.070216219 | 0.15917692 |
| 486,900 | 0.159224563 | 0.219167577 | 0.070561356 | 0.158718596 |
| 487,000 | 0.15919767 | 0.219213178 | 0.070076599 | 0.158784224 |
| 487,100 | 0.159157802 | 0.219038642 | 0.070362402 | 0.158527356 |
| 487,200 | 0.159140678 | 0.219060259 | 0.070005717 | 0.158818735 |
| 487,300 | 0.159227626 | 0.219188538 | 0.070619986 | 0.158908155 |
| 487,400 | 0.159346917 | 0.219050415 | 0.071337967 | 0.158330761 |
| 487,500 | 0.158995999 | 0.219404746 | 0.070972033 | 0.158552112 |
| 487,600 | 0.159162747 | 0.21879652 | 0.070358917 | 0.158965875 |
| 487,700 | 0.159090028 | 0.218580873 | 0.070704569 | 0.159101666 |
| 487,800 | 0.158966939 | 0.219434762 | 0.071003328 | 0.158284297 |
| 487,900 | 0.159018849 | 0.219118204 | 0.070254328 | 0.159048136 |
| 488,000 | 0.159168782 | 0.219332563 | 0.070528956 | 0.15853556 |

| | | | | |
|---|---|---|---|---|
| 488,100 | 0.159209662 | 0.218522577 | 0.070513639 | 0.158156698 |
| 488,200 | 0.158959911 | 0.218368077 | 0.071493613 | 0.158089642 |
| 488,300 | 0.158866117 | 0.219046677 | 0.070766523 | 0.158683575 |
| 488,400 | 0.158834408 | 0.219301614 | 0.071043943 | 0.157832146 |
| 488,500 | 0.158968022 | 0.219772093 | 0.071081036 | 0.158376003 |
| 488,600 | 0.15894276 | 0.21971288 | 0.071298292 | 0.158279095 |
| 488,700 | 0.158919486 | 0.218931215 | 0.071284282 | 0.158263481 |
| 488,800 | 0.158879581 | 0.219273333 | 0.071241173 | 0.158549982 |
| 488,900 | 0.158863229 | 0.218660115 | 0.071430541 | 0.158364083 |
| 489,000 | 0.158767379 | 0.218556658 | 0.071717087 | 0.158587159 |
| 489,100 | 0.158886349 | 0.218456988 | 0.071104982 | 0.158258527 |
| 489,200 | 0.158786056 | 0.21885284 | 0.071344826 | 0.157838668 |
| 489,300 | 0.1589644 | 0.218063578 | 0.070889071 | 0.158604192 |
| 489,400 | 0.158980876 | 0.218436593 | 0.071763164 | 0.158489165 |
| 489,500 | 0.158914048 | 0.218093631 | 0.070747286 | 0.158796876 |
| 489,600 | 0.158913579 | 0.21820246 | 0.071309239 | 0.158154555 |
| 489,700 | 0.158985381 | 0.218562422 | 0.071405281 | 0.158368114 |
| 489,800 | 0.158984716 | 0.217667927 | 0.070876425 | 0.158243671 |
| 489,900 | 0.159003417 | 0.218090714 | 0.071134802 | 0.159236388 |
| 490,000 | 0.158890883 | 0.218546219 | 0.071772528 | 0.158596691 |
| 490,100 | 0.158990621 | 0.218937129 | 0.071024264 | 0.158058924 |
| 490,200 | 0.15888337 | 0.218326334 | 0.071718843 | 0.158144855 |
| 490,300 | 0.159018162 | 0.219124811 | 0.070800876 | 0.158117983 |
| 490,400 | 0.158785073 | 0.218656803 | 0.071948888 | 0.15830683 |
| 490,500 | 0.158790346 | 0.218555275 | 0.071644107 | 0.158788767 |
| 490,600 | 0.158888149 | 0.21842891 | 0.071035292 | 0.158605617 |
| 490,700 | 0.1588387 | 0.21887942 | 0.072462724 | 0.158109142 |
| 490,800 | 0.158603899 | 0.21864404 | 0.071199213 | 0.158216066 |
| 490,900 | 0.158617103 | 0.219352075 | 0.072580258 | 0.158549226 |
| 491,000 | 0.158667367 | 0.218483754 | 0.072250378 | 0.158123532 |
| 491,100 | 0.158701603 | 0.218894403 | 0.071865886 | 0.158905704 |
| 491,200 | 0.158669828 | 0.218438196 | 0.072089856 | 0.15866593 |
| 491,300 | 0.158689653 | 0.218526747 | 0.071896586 | 0.158453031 |
| 491,400 | 0.15877414 | 0.21942768 | 0.072158282 | 0.157676983 |
| 491,500 | 0.158760494 | 0.218665182 | 0.072363772 | 0.158026934 |
| 491,600 | 0.158596019 | 0.218878249 | 0.072463175 | 0.158020287 |
| 491,700 | 0.158572749 | 0.219359756 | 0.071591373 | 0.158103771 |
| 491,800 | 0.15855815 | 0.219019352 | 0.072142526 | 0.157811381 |
| 491,900 | 0.158495594 | 0.218772645 | 0.07174009 | 0.157568707 |
| 492,000 | 0.158658463 | 0.218967193 | 0.072040858 | 0.157699437 |
| 492,100 | 0.158542916 | 0.218636699 | 0.07207211 | 0.157582037 |
| 492,200 | 0.15847514 | 0.218383208 | 0.071711176 | 0.158253549 |
| 492,300 | 0.158377706 | 0.218074378 | 0.072503908 | 0.157421522 |
| 492,400 | 0.158384771 | 0.218114541 | 0.071457083 | 0.158088476 |
| 492,500 | 0.158438988 | 0.218002356 | 0.071877785 | 0.157876652 |
| 492,600 | 0.158390988 | 0.218055932 | 0.072242528 | 0.157752922 |
| 492,700 | 0.158353915 | 0.217741493 | 0.071085349 | 0.157494625 |
| 492,800 | 0.158416025 | 0.217941962 | 0.071574549 | 0.157872422 |
| 492,900 | 0.158350328 | 0.218684261 | 0.071034905 | 0.157642686 |
| 493,000 | 0.158447643 | 0.218724147 | 0.071817118 | 0.15794881 |
| 493,100 | 0.158472121 | 0.218418447 | 0.070948055 | 0.157348435 |
| 493,200 | 0.158245965 | 0.218181472 | 0.071031331 | 0.157181393 |
| 493,300 | 0.158275498 | 0.218011917 | 0.0707455 | 0.157981087 |
| 493,400 | 0.158373487 | 0.218319888 | 0.070662419 | 0.157689463 |
| 493,500 | 0.15836177 | 0.218049576 | 0.070719792 | 0.157386396 |
| 493,600 | 0.158383411 | 0.217959888 | 0.069974058 | 0.15800794 |
| 493,700 | 0.158247553 | 0.218787711 | 0.070313493 | 0.157614444 |
| 493,800 | 0.158213774 | 0.218703524 | 0.070736391 | 0.157891952 |
| 493,900 | 0.158231492 | 0.21863982 | 0.070361402 | 0.158291462 |
| 494,000 | 0.158209951 | 0.217924835 | 0.070417145 | 0.158349039 |
| 494,100 | 0.158235966 | 0.217543744 | 0.070353115 | 0.158273942 |
| 494,200 | 0.158307486 | 0.218463784 | 0.07055723 | 0.158384721 |
| 494,300 | 0.158203051 | 0.217971505 | 0.071033655 | 0.158101309 |
| 494,400 | 0.158273704 | 0.218302005 | 0.070947076 | 0.157648275 |
| 494,500 | 0.158267717 | 0.218464239 | 0.071236671 | 0.158047576 |
| 494,600 | 0.15822121 | 0.217037501 | 0.071594624 | 0.15807828 |
| 494,700 | 0.158157795 | 0.217370489 | 0.071221858 | 0.158099303 |

| | | | | |
|---|---|---|---|---|
| 494,800 | 0.158122717 | 0.217358476 | 0.07179563 | 0.15794595 |
| 494,900 | 0.157987743 | 0.216651669 | 0.071219257 | 0.157888187 |
| 495,000 | 0.158057565 | 0.217634035 | 0.071772023 | 0.157733773 |
| 495,100 | 0.157948767 | 0.217018573 | 0.071638273 | 0.157871738 |
| 495,200 | 0.158060856 | 0.217833879 | 0.071212239 | 0.158062086 |
| 495,300 | 0.157915412 | 0.218825911 | 0.070764462 | 0.157194675 |
| 495,400 | 0.157935193 | 0.218499781 | 0.070454926 | 0.158059483 |
| 495,500 | 0.157912423 | 0.218022804 | 0.070618812 | 0.157997687 |
| 495,600 | 0.157929471 | 0.217228025 | 0.070986376 | 0.157798685 |
| 495,700 | 0.157714872 | 0.217643648 | 0.071055933 | 0.157292387 |
| 495,800 | 0.1577258 | 0.21758338 | 0.069716147 | 0.157995621 |
| 495,900 | 0.157610902 | 0.217246553 | 0.069294826 | 0.156930347 |
| 496,000 | 0.157679125 | 0.217328316 | 0.070621822 | 0.157512791 |
| 496,100 | 0.157600998 | 0.216790128 | 0.069737457 | 0.157531211 |
| 496,200 | 0.157679448 | 0.216884489 | 0.069945117 | 0.15712724 |
| 496,300 | 0.157832068 | 0.216421457 | 0.070214376 | 0.157855834 |
| 496,400 | 0.157614131 | 0.217194018 | 0.070958823 | 0.15732603 |
| 496,500 | 0.157613338 | 0.21715199 | 0.071184864 | 0.157296188 |
| 496,600 | 0.157691677 | 0.2174304 | 0.071358865 | 0.157329848 |
| 496,700 | 0.157649872 | 0.216859862 | 0.071808153 | 0.157154981 |
| 496,800 | 0.157759277 | 0.217204707 | 0.070874439 | 0.157389684 |
| 496,900 | 0.15762684 | 0.217419894 | 0.071547985 | 0.157113667 |
| 497,000 | 0.157605866 | 0.217369659 | 0.071644445 | 0.156211155 |
| 497,100 | 0.157537999 | 0.217451377 | 0.071621399 | 0.156378135 |
| 497,200 | 0.157493066 | 0.216331932 | 0.07071801 | 0.157117242 |
| 497,300 | 0.157535343 | 0.217037666 | 0.0709719 | 0.157197217 |
| 497,400 | 0.157488238 | 0.216885253 | 0.070623503 | 0.156760558 |
| 497,500 | 0.157532518 | 0.217455905 | 0.070276594 | 0.157322014 |
| 497,600 | 0.157376128 | 0.216893364 | 0.070227428 | 0.156674265 |
| 497,700 | 0.157431183 | 0.216934694 | 0.070738207 | 0.15636131 |
| 497,800 | 0.15728454 | 0.217112705 | 0.070497218 | 0.156742978 |
| 497,900 | 0.157276966 | 0.21704838 | 0.071032623 | 0.157257591 |
| 498,000 | 0.157318571 | 0.217264619 | 0.070823983 | 0.157555904 |
| 498,100 | 0.157393703 | 0.21852192 | 0.070680409 | 0.157190423 |
| 498,200 | 0.157428887 | 0.218249654 | 0.070777856 | 0.157602755 |
| 498,300 | 0.157249926 | 0.217121737 | 0.070878504 | 0.157150821 |
| 498,400 | 0.157356194 | 0.217557271 | 0.070971783 | 0.156640651 |
| 498,500 | 0.157258082 | 0.216786905 | 0.071372501 | 0.157216423 |
| 498,600 | 0.157338491 | 0.217380501 | 0.070707519 | 0.156493667 |
| 498,700 | 0.157335855 | 0.217520789 | 0.070181872 | 0.156645635 |
| 498,800 | 0.157213117 | 0.217621347 | 0.070097677 | 0.156618982 |
| 498,900 | 0.157172847 | 0.217119137 | 0.069619217 | 0.156217266 |
| 499,000 | 0.157266029 | 0.217595429 | 0.070759639 | 0.156197442 |
| 499,100 | 0.157189971 | 0.217239133 | 0.070524233 | 0.156552414 |
| 499,200 | 0.157144318 | 0.216791896 | 0.070483816 | 0.157001998 |
| 499,300 | 0.157045167 | 0.217359337 | 0.070403205 | 0.156675987 |
| 499,400 | 0.157094793 | 0.217379487 | 0.070891984 | 0.157335545 |
| 499,500 | 0.156904448 | 0.217438577 | 0.069962779 | 0.15715158 |
| 499,600 | 0.157007825 | 0.218109296 | 0.070552609 | 0.157229933 |
| 499,700 | 0.157040873 | 0.218561279 | 0.0703111 | 0.156951557 |
| 499,800 | 0.157181849 | 0.218188082 | 0.069941632 | 0.157154616 |
| 499,900 | 0.157020189 | 0.218328075 | 0.070711741 | 0.156916187 |
| 500,000 | 0.15698669 | 0.217076092 | 0.070777917 | 0.157024113 |
| 500,100 | 0.157043832 | 0.21814132 | 0.07113592 | 0.157008826 |
| 500,200 | 0.156918949 | 0.217335253 | 0.070289064 | 0.156859335 |
| 500,300 | 0.157010509 | 0.217169599 | 0.070546438 | 0.156885421 |
| 500,400 | 0.156885443 | 0.217498527 | 0.070873303 | 0.156096664 |
| 500,500 | 0.156999837 | 0.217618933 | 0.070484747 | 0.15622858 |
| 500,600 | 0.156880456 | 0.217415802 | 0.070935106 | 0.156682953 |
| 500,700 | 0.156921928 | 0.216971853 | 0.07072587 | 0.156453307 |
| 500,800 | 0.156934312 | 0.21711479 | 0.070746898 | 0.1568207 |
| 500,900 | 0.156891947 | 0.216618678 | 0.070994034 | 0.156019995 |
| 501,000 | 0.156764144 | 0.217089939 | 0.070706079 | 0.156416222 |
| 501,100 | 0.156812334 | 0.217223337 | 0.070300763 | 0.156126125 |
| 501,200 | 0.156861595 | 0.217514017 | 0.070786477 | 0.156562483 |
| 501,300 | 0.156927278 | 0.217500578 | 0.070453286 | 0.156305144 |
| 501,400 | 0.156942972 | 0.218056499 | 0.069991872 | 0.156347345 |

| | | | | |
|---|---|---|---|---|
| 501,500 | 0.156976365 | 0.217583228 | 0.070390747 | 0.15639582 |
| 501,600 | 0.156879948 | 0.2172944 | 0.070316542 | 0.156249487 |
| 501,700 | 0.156808704 | 0.217425342 | 0.070464795 | 0.156092186 |
| 501,800 | 0.156907174 | 0.217247383 | 0.070482977 | 0.155943045 |
| 501,900 | 0.156891062 | 0.217680594 | 0.071356294 | 0.155423233 |
| 502,000 | 0.156784699 | 0.21834311 | 0.070621427 | 0.155858042 |
| 502,100 | 0.156740464 | 0.21799776 | 0.070012559 | 0.155877927 |
| 502,200 | 0.156761475 | 0.2183164 | 0.069277044 | 0.155330133 |
| 502,300 | 0.156956216 | 0.218141826 | 0.0707122 | 0.156524532 |
| 502,400 | 0.156792344 | 0.217974034 | 0.071010254 | 0.156129857 |
| 502,500 | 0.156840992 | 0.21805311 | 0.071268119 | 0.155870567 |
| 502,600 | 0.156734603 | 0.21804335 | 0.070381964 | 0.15532815 |
| 502,700 | 0.1566162 | 0.217609432 | 0.070540112 | 0.155607302 |
| 502,800 | 0.156601838 | 0.217665492 | 0.070264941 | 0.155974464 |
| 502,900 | 0.15661662 | 0.217724921 | 0.070773352 | 0.156341734 |
| 503,000 | 0.156616601 | 0.217764498 | 0.069761106 | 0.156204301 |
| 503,100 | 0.156555529 | 0.216985586 | 0.070846277 | 0.1555771 |
| 503,200 | 0.156580279 | 0.217417906 | 0.069845555 | 0.156165471 |
| 503,300 | 0.156514037 | 0.217989425 | 0.07032624 | 0.155460592 |
| 503,400 | 0.156635717 | 0.217302465 | 0.069965621 | 0.155978944 |
| 503,500 | 0.156411358 | 0.217069793 | 0.07071513 | 0.156306777 |
| 503,600 | 0.156494625 | 0.217863294 | 0.070630659 | 0.155510535 |
| 503,700 | 0.156554782 | 0.217834863 | 0.070149302 | 0.156026779 |
| 503,800 | 0.156584427 | 0.217507683 | 0.070981825 | 0.156341888 |
| 503,900 | 0.156593673 | 0.216666188 | 0.070436172 | 0.156774429 |
| 504,000 | 0.15641916 | 0.21718349 | 0.070833252 | 0.156231752 |
| 504,100 | 0.156486453 | 0.217205672 | 0.070977871 | 0.156520559 |
| 504,200 | 0.156596886 | 0.217237432 | 0.070786033 | 0.156095702 |
| 504,300 | 0.156418951 | 0.217292151 | 0.070608598 | 0.155849324 |
| 504,400 | 0.156344795 | 0.216808442 | 0.070209889 | 0.155537753 |
| 504,500 | 0.156430548 | 0.217083801 | 0.069137708 | 0.156235055 |
| 504,600 | 0.156373435 | 0.21681091 | 0.069469515 | 0.155707099 |
| 504,700 | 0.156319953 | 0.216848059 | 0.070281079 | 0.155505483 |
| 504,800 | 0.156400581 | 0.216924129 | 0.069582599 | 0.155282264 |
| 504,900 | 0.156282188 | 0.216688615 | 0.069909656 | 0.1550225 |
| 505,000 | 0.156392218 | 0.216722979 | 0.070119336 | 0.155945736 |
| 505,100 | 0.156279036 | 0.216481137 | 0.070070648 | 0.155420552 |
| 505,200 | 0.156208165 | 0.216234724 | 0.070389156 | 0.155195071 |
| 505,300 | 0.156233661 | 0.215789888 | 0.070833572 | 0.155760386 |
| 505,400 | 0.156277957 | 0.215999008 | 0.069632381 | 0.155211622 |
| 505,500 | 0.156145457 | 0.216343558 | 0.069638259 | 0.155946279 |
| 505,600 | 0.15616736 | 0.216263033 | 0.070375541 | 0.155106747 |
| 505,700 | 0.156194227 | 0.216884383 | 0.07067306 | 0.155060847 |
| 505,800 | 0.156145994 | 0.216411455 | 0.070688412 | 0.154721403 |
| 505,900 | 0.156305866 | 0.21743957 | 0.070463276 | 0.155178745 |
| 506,000 | 0.156198748 | 0.217032756 | 0.070208758 | 0.15449513 |
| 506,100 | 0.156187356 | 0.217296479 | 0.069834952 | 0.154608395 |
| 506,200 | 0.156374083 | 0.216727232 | 0.070020544 | 0.155357872 |
| 506,300 | 0.156191617 | 0.217204884 | 0.070238998 | 0.155272653 |
| 506,400 | 0.156214668 | 0.217164082 | 0.07036971 | 0.155823825 |
| 506,500 | 0.156223812 | 0.216949506 | 0.070278603 | 0.1553117 |
| 506,600 | 0.156250173 | 0.217287307 | 0.070827749 | 0.155051729 |
| 506,700 | 0.156060568 | 0.217117092 | 0.070107075 | 0.154851103 |
| 506,800 | 0.156064194 | 0.216519369 | 0.070308915 | 0.154915601 |
| 506,900 | 0.156083319 | 0.216701694 | 0.070427322 | 0.154546879 |
| 507,000 | 0.156111546 | 0.216522326 | 0.071137247 | 0.1545558 |
| 507,100 | 0.156033177 | 0.216698589 | 0.071231518 | 0.154384606 |
| 507,200 | 0.15604249 | 0.216970525 | 0.070779727 | 0.154011347 |
| 507,300 | 0.155963493 | 0.216714282 | 0.070293442 | 0.154422343 |
| 507,400 | 0.155968378 | 0.216477454 | 0.071053507 | 0.154146476 |
| 507,500 | 0.155805849 | 0.21565619 | 0.070768226 | 0.153656324 |
| 507,600 | 0.155975954 | 0.215729722 | 0.0712603 | 0.154576487 |
| 507,700 | 0.156054022 | 0.216692005 | 0.070575765 | 0.154601818 |
| 507,800 | 0.15608835 | 0.216764312 | 0.070860873 | 0.154413152 |
| 507,900 | 0.155993304 | 0.216886117 | 0.070546221 | 0.154184943 |
| 508,000 | 0.156067598 | 0.216006047 | 0.0707073 | 0.154758719 |
| 508,100 | 0.156038473 | 0.21631889 | 0.070634477 | 0.153979881 |

| | | | | |
|---|---|---|---|---|
| 508,200 | 0.155854923 | 0.216138367 | 0.070445393 | 0.154277502 |
| 508,300 | 0.155897779 | 0.215516542 | 0.070428874 | 0.154483431 |
| 508,400 | 0.155812821 | 0.216353767 | 0.070531088 | 0.155252606 |
| 508,500 | 0.155986679 | 0.215959324 | 0.070637436 | 0.154834343 |
| 508,600 | 0.155871586 | 0.215861441 | 0.07029359 | 0.155220345 |
| 508,700 | 0.155853942 | 0.216673103 | 0.071583548 | 0.154664387 |
| 508,800 | 0.15578645 | 0.216644861 | 0.070405866 | 0.154460456 |
| 508,900 | 0.155745356 | 0.216127429 | 0.071109618 | 0.154125422 |
| 509,000 | 0.15585474 | 0.216180474 | 0.07164434 | 0.154690219 |
| 509,100 | 0.155863722 | 0.215794838 | 0.071295571 | 0.154100596 |
| 509,200 | 0.155897354 | 0.215733164 | 0.070922641 | 0.155366697 |
| 509,300 | 0.155855795 | 0.215899882 | 0.070970991 | 0.154776259 |
| 509,400 | 0.155691679 | 0.216113757 | 0.070251922 | 0.153560911 |
| 509,500 | 0.155867174 | 0.215985902 | 0.070137134 | 0.154302279 |
| 509,600 | 0.155879436 | 0.216883171 | 0.070738986 | 0.15406992 |
| 509,700 | 0.155716178 | 0.215942926 | 0.071386775 | 0.153761327 |
| 509,800 | 0.155747782 | 0.215586472 | 0.071239121 | 0.153768483 |
| 509,900 | 0.155652297 | 0.215948716 | 0.070497019 | 0.153675753 |
| 510,000 | 0.15574499 | 0.215794871 | 0.070861066 | 0.154375429 |
| 510,100 | 0.155682359 | 0.216485303 | 0.070062038 | 0.154148647 |
| 510,200 | 0.155571425 | 0.215554608 | 0.070909558 | 0.154218783 |
| 510,300 | 0.155731094 | 0.21622522 | 0.070636232 | 0.154372604 |
| 510,400 | 0.155632033 | 0.21629074 | 0.071147285 | 0.154271195 |
| 510,500 | 0.155651119 | 0.215730129 | 0.070898694 | 0.153975127 |
| 510,600 | 0.155714586 | 0.215954966 | 0.071142416 | 0.154221808 |
| 510,700 | 0.155713229 | 0.216184451 | 0.071627233 | 0.154404858 |
| 510,800 | 0.155734242 | 0.215763735 | 0.070735966 | 0.154360164 |
| 510,900 | 0.155560138 | 0.21467288 | 0.071093484 | 0.154366577 |
| 511,000 | 0.155465817 | 0.215834044 | 0.070726139 | 0.153503338 |
| 511,100 | 0.155522044 | 0.215633736 | 0.071392326 | 0.153983425 |
| 511,200 | 0.155497753 | 0.216096436 | 0.071125027 | 0.15431263 |
| 511,300 | 0.155497046 | 0.215485379 | 0.070094122 | 0.153695332 |
| 511,400 | 0.155579609 | 0.216264825 | 0.070552603 | 0.154589475 |
| 511,500 | 0.155650002 | 0.216549504 | 0.070694437 | 0.154338225 |
| 511,600 | 0.155465273 | 0.216056111 | 0.070779844 | 0.153860795 |
| 511,700 | 0.155568468 | 0.216595796 | 0.070775817 | 0.154221784 |
| 511,800 | 0.155500032 | 0.216190251 | 0.07081236 | 0.153964806 |
| 511,900 | 0.155466992 | 0.215975171 | 0.070548112 | 0.154079258 |
| 512,000 | 0.155467569 | 0.216462368 | 0.069975483 | 0.153912215 |
| 512,100 | 0.15552081 | 0.216438587 | 0.071208376 | 0.154680787 |
| 512,200 | 0.155340378 | 0.215758407 | 0.070565452 | 0.154463547 |
| 512,300 | 0.155642793 | 0.216933193 | 0.07049328 | 0.154754079 |
| 512,400 | 0.155517248 | 0.215872638 | 0.069965752 | 0.154801026 |
| 512,500 | 0.155549799 | 0.21571429 | 0.071041335 | 0.154887011 |
| 512,600 | 0.155388449 | 0.215629007 | 0.070839898 | 0.154921463 |
| 512,700 | 0.15530158 | 0.214961688 | 0.070442727 | 0.155476547 |
| 512,800 | 0.155355314 | 0.215067358 | 0.071044837 | 0.154423515 |
| 512,900 | 0.155380683 | 0.215930498 | 0.070519675 | 0.154543374 |
| 513,000 | 0.155351907 | 0.216112072 | 0.071466732 | 0.155168635 |
| 513,100 | 0.155422684 | 0.216773118 | 0.071286515 | 0.155607673 |
| 513,200 | 0.155332125 | 0.216490792 | 0.070813289 | 0.155489271 |
| 513,300 | 0.155153348 | 0.215966461 | 0.070832077 | 0.154215459 |
| 513,400 | 0.155174875 | 0.215698897 | 0.070661337 | 0.154099093 |
| 513,500 | 0.155290853 | 0.216506613 | 0.070761886 | 0.154594575 |
| 513,600 | 0.155286299 | 0.216232123 | 0.070436929 | 0.154190629 |
| 513,700 | 0.155317571 | 0.216238506 | 0.071621221 | 0.154504023 |
| 513,800 | 0.155319546 | 0.216351951 | 0.072078254 | 0.154798298 |
| 513,900 | 0.155252486 | 0.216712418 | 0.071816212 | 0.154247797 |
| 514,000 | 0.155245852 | 0.216033893 | 0.071587656 | 0.154132868 |
| 514,100 | 0.155094101 | 0.215581913 | 0.071376682 | 0.153878038 |
| 514,200 | 0.155105809 | 0.21552369 | 0.070920632 | 0.154459508 |
| 514,300 | 0.155047162 | 0.215858999 | 0.070918942 | 0.153936575 |
| 514,400 | 0.1551597 | 0.215268661 | 0.071132951 | 0.154188213 |
| 514,500 | 0.155104184 | 0.2151326 | 0.070823179 | 0.154181633 |
| 514,600 | 0.155091731 | 0.215847427 | 0.071498622 | 0.154037384 |
| 514,700 | 0.155065074 | 0.214894429 | 0.070961193 | 0.154509954 |
| 514,800 | 0.155090205 | 0.215288235 | 0.071675611 | 0.154251047 |

| | | | | |
|---|---|---|---|---|
| 514,900 | 0.155094057 | 0.21503096 | 0.072306285 | 0.154148269 |
| 515,000 | 0.155087743 | 0.21542913 | 0.071651613 | 0.154135521 |
| 515,100 | 0.155002426 | 0.215195366 | 0.071543416 | 0.153817165 |
| 515,200 | 0.154878574 | 0.215462673 | 0.071768089 | 0.154028516 |
| 515,300 | 0.154975788 | 0.215634285 | 0.070738819 | 0.154426683 |
| 515,400 | 0.154962688 | 0.215122482 | 0.070951725 | 0.154313529 |
| 515,500 | 0.155041425 | 0.215118327 | 0.071517559 | 0.154596038 |
| 515,600 | 0.154959013 | 0.214987549 | 0.070968353 | 0.154447849 |
| 515,700 | 0.155010456 | 0.21564777 | 0.071344036 | 0.15456076 |
| 515,800 | 0.154923432 | 0.215584242 | 0.070920564 | 0.154274703 |
| 515,900 | 0.155020492 | 0.216103521 | 0.071001741 | 0.154090012 |
| 516,000 | 0.154796507 | 0.215939822 | 0.071234111 | 0.153617097 |
| 516,100 | 0.154973602 | 0.215812384 | 0.070935834 | 0.153777906 |
| 516,200 | 0.154777503 | 0.215042473 | 0.071178179 | 0.154070314 |
| 516,300 | 0.154816281 | 0.215349232 | 0.071749377 | 0.153951634 |
| 516,400 | 0.154659039 | 0.214991122 | 0.071215176 | 0.154285644 |
| 516,500 | 0.154608907 | 0.215275095 | 0.071758854 | 0.153833539 |
| 516,600 | 0.154671145 | 0.215203242 | 0.071120047 | 0.152947503 |
| 516,700 | 0.154592006 | 0.215702339 | 0.070480548 | 0.15325387 |
| 516,800 | 0.154524243 | 0.215021763 | 0.071737482 | 0.153346094 |
| 516,900 | 0.154488245 | 0.214505359 | 0.071657722 | 0.154298613 |
| 517,000 | 0.154781308 | 0.215177459 | 0.07191969 | 0.153893346 |
| 517,100 | 0.154548714 | 0.214620014 | 0.072358699 | 0.153728508 |
| 517,200 | 0.15453764 | 0.215004312 | 0.071226998 | 0.153951114 |
| 517,300 | 0.15437784 | 0.214842203 | 0.07231644 | 0.153606198 |
| 517,400 | 0.154661184 | 0.214324333 | 0.071431479 | 0.153439029 |
| 517,500 | 0.154607295 | 0.21437645 | 0.071774195 | 0.153987066 |
| 517,600 | 0.154518893 | 0.213949412 | 0.07187264 | 0.153474685 |
| 517,700 | 0.154493572 | 0.213463516 | 0.07206891 | 0.153504573 |
| 517,800 | 0.154456682 | 0.213461383 | 0.071113236 | 0.153735594 |
| 517,900 | 0.154345013 | 0.213829994 | 0.071568064 | 0.154056547 |
| 518,000 | 0.154429537 | 0.214187767 | 0.071107823 | 0.153730281 |
| 518,100 | 0.154374075 | 0.214538621 | 0.070489817 | 0.154214307 |
| 518,200 | 0.154301854 | 0.21295926 | 0.071801591 | 0.154056929 |
| 518,300 | 0.154195754 | 0.213305574 | 0.071140846 | 0.154082533 |
| 518,400 | 0.154274042 | 0.213841673 | 0.072284606 | 0.153850593 |
| 518,500 | 0.154179511 | 0.213474981 | 0.071537525 | 0.154708308 |
| 518,600 | 0.154142817 | 0.213605594 | 0.071012309 | 0.154045644 |
| 518,700 | 0.154262415 | 0.213671275 | 0.070285548 | 0.153987772 |
| 518,800 | 0.154306435 | 0.214148355 | 0.071127705 | 0.154026701 |
| 518,900 | 0.154230089 | 0.213339962 | 0.070501696 | 0.154316369 |
| 519,000 | 0.154206268 | 0.213432762 | 0.071128192 | 0.154044388 |
| 519,100 | 0.154199417 | 0.214047543 | 0.071218016 | 0.153702623 |
| 519,200 | 0.154190087 | 0.214125433 | 0.070594278 | 0.153597068 |
| 519,300 | 0.154006353 | 0.213441868 | 0.070561916 | 0.153522912 |
| 519,400 | 0.154072231 | 0.214279347 | 0.071110081 | 0.153387468 |
| 519,500 | 0.154223714 | 0.213438707 | 0.071150091 | 0.153648066 |
| 519,600 | 0.154229206 | 0.214026088 | 0.070999076 | 0.153878508 |
| 519,700 | 0.154192535 | 0.213325561 | 0.07115737 | 0.153799528 |
| 519,800 | 0.154084965 | 0.21360153 | 0.071502554 | 0.154172634 |
| 519,900 | 0.15407125 | 0.213613405 | 0.07084204 | 0.153821764 |
| 520,000 | 0.154181777 | 0.213061478 | 0.071301869 | 0.154382728 |
| 520,100 | 0.154174793 | 0.214030685 | 0.070908549 | 0.154121083 |
| 520,200 | 0.154052593 | 0.213877016 | 0.070340437 | 0.15351463 |
| 520,300 | 0.154143405 | 0.214078897 | 0.0701439 | 0.153626341 |
| 520,400 | 0.154020478 | 0.213991284 | 0.071424087 | 0.153372765 |
| 520,500 | 0.154046188 | 0.214286944 | 0.071060353 | 0.153773422 |
| 520,600 | 0.154046921 | 0.213422243 | 0.070194507 | 0.153896389 |
| 520,700 | 0.154005709 | 0.213142783 | 0.070911758 | 0.152959201 |
| 520,800 | 0.154028592 | 0.213738744 | 0.070229979 | 0.152874502 |
| 520,900 | 0.154027066 | 0.213322445 | 0.070036688 | 0.153074009 |
| 521,000 | 0.154020795 | 0.213448513 | 0.070989306 | 0.15338919 |
| 521,100 | 0.154016506 | 0.213589157 | 0.070183697 | 0.153434904 |
| 521,200 | 0.153994928 | 0.212903398 | 0.070218764 | 0.15455337 |
| 521,300 | 0.153934842 | 0.21292641 | 0.070915064 | 0.153574424 |
| 521,400 | 0.153890073 | 0.212506746 | 0.070202073 | 0.15369639 |
| 521,500 | 0.153852633 | 0.212619672 | 0.070121619 | 0.153612703 |

| | | | | |
|---|---|---|---|---|
| 521,600 | 0.153825667 | 0.212797736 | 0.07009256 | 0.153973641 |
| 521,700 | 0.153854115 | 0.213297352 | 0.070780608 | 0.153048377 |
| 521,800 | 0.153796295 | 0.212280769 | 0.070028896 | 0.15325629 |
| 521,900 | 0.153688895 | 0.212840354 | 0.07033698 | 0.153009366 |
| 522,000 | 0.153761869 | 0.212630484 | 0.070375363 | 0.153803482 |
| 522,100 | 0.153653225 | 0.213434775 | 0.070948013 | 0.153311571 |
| 522,200 | 0.153815739 | 0.213188639 | 0.070861397 | 0.153609736 |
| 522,300 | 0.153711937 | 0.212788255 | 0.070660653 | 0.153629271 |
| 522,400 | 0.153691648 | 0.213116396 | 0.070988496 | 0.153428795 |
| 522,500 | 0.1535915 | 0.213489629 | 0.070911389 | 0.15308648 |
| 522,600 | 0.15350417 | 0.212842839 | 0.070741573 | 0.152813753 |
| 522,700 | 0.153619587 | 0.212100352 | 0.070306554 | 0.153613955 |
| 522,800 | 0.153593214 | 0.212500769 | 0.07051037 | 0.153169124 |
| 522,900 | 0.153555039 | 0.212205608 | 0.070594515 | 0.15381042 |
| 523,000 | 0.153582813 | 0.21234995 | 0.069664618 | 0.153579564 |
| 523,100 | 0.153558505 | 0.212083829 | 0.070902675 | 0.153735444 |
| 523,200 | 0.153435629 | 0.21192282 | 0.06899612 | 0.153744599 |
| 523,300 | 0.153534941 | 0.211978293 | 0.069505573 | 0.153848093 |
| 523,400 | 0.153482829 | 0.212108658 | 0.070147255 | 0.153187415 |
| 523,500 | 0.153603745 | 0.212248368 | 0.070968865 | 0.15320618 |
| 523,600 | 0.153523199 | 0.213022625 | 0.071334133 | 0.153384222 |
| 523,700 | 0.15357649 | 0.213977782 | 0.070800359 | 0.153563707 |
| 523,800 | 0.153541271 | 0.213826172 | 0.071141878 | 0.153415588 |
| 523,900 | 0.153552742 | 0.212985733 | 0.071002887 | 0.153785765 |
| 524,000 | 0.153345341 | 0.212693597 | 0.071053296 | 0.152872061 |
| 524,100 | 0.153539254 | 0.213729361 | 0.070944811 | 0.153317768 |
| 524,200 | 0.153435161 | 0.213221887 | 0.071936582 | 0.153625436 |
| 524,300 | 0.153484106 | 0.213558082 | 0.071298952 | 0.153337067 |
| 524,400 | 0.153439782 | 0.212741561 | 0.071717233 | 0.153605553 |
| 524,500 | 0.153477011 | 0.212771402 | 0.072131145 | 0.153227573 |
| 524,600 | 0.153522474 | 0.213526847 | 0.071493548 | 0.15377643 |
| 524,700 | 0.153450879 | 0.213271033 | 0.072008866 | 0.153318986 |
| 524,800 | 0.153306277 | 0.212887297 | 0.071791281 | 0.153434203 |
| 524,900 | 0.153252022 | 0.212678989 | 0.070724194 | 0.152952406 |
| 525,000 | 0.153419766 | 0.213643523 | 0.070664169 | 0.153674348 |
| 525,100 | 0.153241525 | 0.212933699 | 0.07073075 | 0.153200014 |
| 525,200 | 0.153191417 | 0.213485973 | 0.070488539 | 0.152892872 |
| 525,300 | 0.153308757 | 0.21296171 | 0.071112488 | 0.153037243 |
| 525,400 | 0.153275767 | 0.212616036 | 0.071067085 | 0.152895146 |
| 525,500 | 0.153072948 | 0.212169465 | 0.069740221 | 0.153121754 |
| 525,600 | 0.153068114 | 0.212530132 | 0.069786019 | 0.152838637 |
| 525,700 | 0.153157105 | 0.212592155 | 0.070171775 | 0.152972425 |
| 525,800 | 0.153138025 | 0.211833398 | 0.069768196 | 0.152949117 |
| 525,900 | 0.153175538 | 0.211655935 | 0.070356507 | 0.153781622 |
| 526,000 | 0.153189989 | 0.211823499 | 0.070437277 | 0.15337617 |
| 526,100 | 0.153174564 | 0.21093917 | 0.069679836 | 0.153602696 |
| 526,200 | 0.153129168 | 0.211831139 | 0.069341027 | 0.152113213 |
| 526,300 | 0.153116672 | 0.211984948 | 0.069265432 | 0.152322723 |
| 526,400 | 0.152997421 | 0.211986135 | 0.069267519 | 0.152781675 |
| 526,500 | 0.152992253 | 0.211326226 | 0.069122068 | 0.153701782 |
| 526,600 | 0.153132587 | 0.211653683 | 0.069093775 | 0.153564048 |
| 526,700 | 0.153071671 | 0.211865562 | 0.068776222 | 0.152626732 |
| 526,800 | 0.153029559 | 0.212336127 | 0.069000168 | 0.152483248 |
| 526,900 | 0.152847153 | 0.212144039 | 0.069083104 | 0.152683942 |
| 527,000 | 0.152984473 | 0.212772076 | 0.069608673 | 0.152545653 |
| 527,100 | 0.15297204 | 0.212703976 | 0.069456965 | 0.153327473 |
| 527,200 | 0.153036572 | 0.212785516 | 0.070676673 | 0.153376152 |
| 527,300 | 0.153026031 | 0.212834081 | 0.070334316 | 0.152649137 |
| 527,400 | 0.152974173 | 0.212411889 | 0.069707362 | 0.153065487 |
| 527,500 | 0.153028003 | 0.212896772 | 0.07046004 | 0.153405075 |
| 527,600 | 0.152975307 | 0.212364738 | 0.069835453 | 0.152713237 |
| 527,700 | 0.152963615 | 0.212120235 | 0.069223089 | 0.152816032 |
| 527,800 | 0.153096726 | 0.213005417 | 0.069928863 | 0.153190373 |
| 527,900 | 0.152868668 | 0.212443708 | 0.069233527 | 0.152801161 |
| 528,000 | 0.152923939 | 0.212505476 | 0.068848029 | 0.152859532 |
| 528,100 | 0.153014198 | 0.212316136 | 0.069664514 | 0.152982182 |
| 528,200 | 0.152932239 | 0.211894644 | 0.069486793 | 0.152780507 |

| | | | | |
|---|---|---|---|---|
| 528,300 | 0.152817024 | 0.21229156 | 0.069408444 | 0.152496694 |
| 528,400 | 0.152804553 | 0.212120386 | 0.069147965 | 0.152987203 |
| 528,500 | 0.152911845 | 0.212220557 | 0.06866008 | 0.152529781 |
| 528,600 | 0.152868539 | 0.212564288 | 0.068840665 | 0.152631717 |
| 528,700 | 0.152914283 | 0.212224351 | 0.068720513 | 0.15281115 |
| 528,800 | 0.152790192 | 0.211470986 | 0.069278426 | 0.152813754 |
| 528,900 | 0.152787102 | 0.212023996 | 0.068452242 | 0.152171361 |
| 529,000 | 0.152756126 | 0.211820386 | 0.068280266 | 0.15235507 |
| 529,100 | 0.152846345 | 0.212179879 | 0.069100672 | 0.152636752 |
| 529,200 | 0.152686384 | 0.211519606 | 0.068900739 | 0.152206166 |
| 529,300 | 0.152882379 | 0.21253126 | 0.069098296 | 0.153187631 |
| 529,400 | 0.152796467 | 0.211997537 | 0.069566767 | 0.153127527 |
| 529,500 | 0.152752338 | 0.212412739 | 0.068874054 | 0.152622714 |
| 529,600 | 0.15269203 | 0.211173258 | 0.068948544 | 0.153051243 |
| 529,700 | 0.152857481 | 0.211595184 | 0.069080191 | 0.15301906 |
| 529,800 | 0.152821984 | 0.210991294 | 0.069199929 | 0.153322549 |
| 529,900 | 0.152842429 | 0.210285192 | 0.069265507 | 0.153291167 |
| 530,000 | 0.15283229 | 0.210887809 | 0.069522515 | 0.152627551 |
| 530,100 | 0.152713968 | 0.21147539 | 0.069107311 | 0.152420678 |
| 530,200 | 0.152673839 | 0.210473359 | 0.069241489 | 0.153055966 |
| 530,300 | 0.152534187 | 0.210759426 | 0.068726193 | 0.152933229 |
| 530,400 | 0.152617614 | 0.21135438 | 0.069671753 | 0.153833555 |
| 530,500 | 0.15264821 | 0.210473569 | 0.069505213 | 0.153073307 |
| 530,600 | 0.152639028 | 0.210627376 | 0.069555848 | 0.152734756 |
| 530,700 | 0.152563191 | 0.210742216 | 0.069224239 | 0.152688333 |
| 530,800 | 0.152744682 | 0.211140982 | 0.069560653 | 0.152619726 |
| 530,900 | 0.152594727 | 0.210743732 | 0.069133965 | 0.152421833 |
| 531,000 | 0.152658567 | 0.211689255 | 0.069276156 | 0.152335758 |
| 531,100 | 0.15254467 | 0.211174454 | 0.068979243 | 0.152686717 |
| 531,200 | 0.152415 | 0.210659918 | 0.0688954 | 0.152194338 |
| 531,300 | 0.15257704 | 0.211381606 | 0.068981419 | 0.152303757 |
| 531,400 | 0.152528179 | 0.211625172 | 0.068607126 | 0.15283445 |
| 531,500 | 0.152406531 | 0.211899836 | 0.068202128 | 0.153569613 |
| 531,600 | 0.152515527 | 0.210998536 | 0.068218176 | 0.152778654 |
| 531,700 | 0.152419538 | 0.210463291 | 0.068545902 | 0.152780861 |
| 531,800 | 0.152451989 | 0.210629184 | 0.068442219 | 0.153481891 |
| 531,900 | 0.152406545 | 0.211283788 | 0.06818226 | 0.153769379 |
| 532,000 | 0.152420667 | 0.211235476 | 0.068289241 | 0.152477547 |
| 532,100 | 0.152281247 | 0.210790694 | 0.068057817 | 0.152613973 |
| 532,200 | 0.15232194 | 0.210875713 | 0.067713895 | 0.153543784 |
| 532,300 | 0.152328124 | 0.210618546 | 0.068081519 | 0.153355681 |
| 532,400 | 0.15238178 | 0.210473138 | 0.068070068 | 0.153004674 |
| 532,500 | 0.152347678 | 0.210368675 | 0.067180662 | 0.153192884 |
| 532,600 | 0.152278368 | 0.210313806 | 0.067104707 | 0.152723512 |
| 532,700 | 0.152382828 | 0.210460789 | 0.067689152 | 0.153298144 |
| 532,800 | 0.152326615 | 0.209854638 | 0.067798348 | 0.153236033 |
| 532,900 | 0.152252106 | 0.210699991 | 0.067757381 | 0.153236562 |
| 533,000 | 0.152209395 | 0.21002409 | 0.068594623 | 0.152855929 |
| 533,100 | 0.152321982 | 0.210051476 | 0.0669406 | 0.153705259 |
| 533,200 | 0.152251835 | 0.209774099 | 0.067345897 | 0.153099968 |
| 533,300 | 0.152208767 | 0.209607415 | 0.067742889 | 0.152919914 |
| 533,400 | 0.152238684 | 0.209592332 | 0.066849146 | 0.153434269 |
| 533,500 | 0.15210915 | 0.209966771 | 0.066913577 | 0.153002296 |
| 533,600 | 0.152258431 | 0.210089468 | 0.067774116 | 0.152813123 |
| 533,700 | 0.152285692 | 0.21046185 | 0.067908126 | 0.152857389 |
| 533,800 | 0.152372015 | 0.210317847 | 0.067359948 | 0.15248789 |
| 533,900 | 0.152158163 | 0.210099294 | 0.066788928 | 0.152921039 |
| 534,000 | 0.152295068 | 0.210122863 | 0.066969437 | 0.153216922 |
| 534,100 | 0.152160088 | 0.209886165 | 0.067285144 | 0.152818096 |
| 534,200 | 0.152013552 | 0.210241896 | 0.067484038 | 0.152833376 |
| 534,300 | 0.152112142 | 0.20988015 | 0.067484124 | 0.152535286 |
| 534,400 | 0.152132207 | 0.20986152 | 0.066899659 | 0.153083435 |
| 534,500 | 0.152046849 | 0.209736836 | 0.06773272 | 0.152609112 |
| 534,600 | 0.152036732 | 0.210074623 | 0.066832958 | 0.152528395 |
| 534,700 | 0.152076648 | 0.210171649 | 0.067006593 | 0.153099236 |
| 534,800 | 0.152062228 | 0.210062765 | 0.066934749 | 0.152207027 |
| 534,900 | 0.151900714 | 0.210297277 | 0.067173914 | 0.152474556 |

| | | | | |
|---|---|---|---|---|
| 535,000 | 0.152128462 | 0.210305139 | 0.067010726 | 0.152589396 |
| 535,100 | 0.151972233 | 0.209238779 | 0.067102478 | 0.152479013 |
| 535,200 | 0.151842727 | 0.209890461 | 0.066770188 | 0.152343154 |
| 535,300 | 0.151839658 | 0.209849983 | 0.066373087 | 0.152620865 |
| 535,400 | 0.151994984 | 0.209165779 | 0.067249258 | 0.152554467 |
| 535,500 | 0.151847368 | 0.209217213 | 0.067267854 | 0.152102503 |
| 535,600 | 0.15207105 | 0.209966913 | 0.067696317 | 0.153301224 |
| 535,700 | 0.151826405 | 0.209254402 | 0.067102053 | 0.152830872 |
| 535,800 | 0.15185344 | 0.208789208 | 0.066884116 | 0.152616374 |
| 535,900 | 0.1518135 | 0.208936765 | 0.066869477 | 0.152443177 |
| 536,000 | 0.151737642 | 0.208724767 | 0.066569072 | 0.152493682 |
| 536,100 | 0.151814081 | 0.209081228 | 0.066723857 | 0.153165469 |
| 536,200 | 0.151692654 | 0.208522647 | 0.066472426 | 0.152533731 |
| 536,300 | 0.151690758 | 0.208269091 | 0.066685675 | 0.152472482 |
| 536,400 | 0.151821644 | 0.207974945 | 0.066176885 | 0.152774184 |
| 536,500 | 0.151730908 | 0.20836863 | 0.066968292 | 0.152210309 |
| 536,600 | 0.151724416 | 0.208380545 | 0.066528227 | 0.152874149 |
| 536,700 | 0.151834504 | 0.208450501 | 0.066250373 | 0.152459647 |
| 536,800 | 0.151688802 | 0.208116944 | 0.06589832 | 0.151762391 |
| 536,900 | 0.151584963 | 0.208854567 | 0.066322646 | 0.151805644 |
| 537,000 | 0.151592197 | 0.208123595 | 0.066502567 | 0.151836567 |
| 537,100 | 0.15165236 | 0.208453706 | 0.066702742 | 0.152808609 |
| 537,200 | 0.151558336 | 0.208288088 | 0.066360943 | 0.152153237 |
| 537,300 | 0.151351914 | 0.20807583 | 0.066223856 | 0.152744435 |
| 537,400 | 0.151464377 | 0.207759745 | 0.066370695 | 0.153029424 |
| 537,500 | 0.151390684 | 0.207420477 | 0.06616525 | 0.152632094 |
| 537,600 | 0.151395235 | 0.207971298 | 0.066343193 | 0.152594407 |
| 537,700 | 0.151413081 | 0.207579176 | 0.066240192 | 0.152120209 |
| 537,800 | 0.151397697 | 0.207963937 | 0.065551535 | 0.152747994 |
| 537,900 | 0.151349091 | 0.207578135 | 0.065701087 | 0.15217843 |
| 538,000 | 0.151250388 | 0.206830476 | 0.065776328 | 0.151909611 |
| 538,100 | 0.15134346 | 0.207172288 | 0.06584145 | 0.152540643 |
| 538,200 | 0.151424013 | 0.207022046 | 0.065885873 | 0.152934668 |
| 538,300 | 0.151240768 | 0.206620871 | 0.066805483 | 0.152235421 |
| 538,400 | 0.151350542 | 0.207822834 | 0.066528573 | 0.152813715 |
| 538,500 | 0.151320582 | 0.207264555 | 0.066906969 | 0.1529684 |
| 538,600 | 0.15120282 | 0.206955837 | 0.066188836 | 0.15246803 |
| 538,700 | 0.151180342 | 0.207938509 | 0.06599437 | 0.152784131 |
| 538,800 | 0.151207895 | 0.208290305 | 0.066375959 | 0.152910805 |
| 538,900 | 0.151128696 | 0.207796749 | 0.066034758 | 0.152759976 |
| 539,000 | 0.150970932 | 0.207406667 | 0.066490154 | 0.152938357 |
| 539,100 | 0.15105365 | 0.207742796 | 0.066621114 | 0.152595622 |
| 539,200 | 0.151126149 | 0.208120527 | 0.066849955 | 0.15290133 |
| 539,300 | 0.151012119 | 0.207802483 | 0.066284403 | 0.152873067 |
| 539,400 | 0.151088924 | 0.208023575 | 0.06588852 | 0.1525702 |
| 539,500 | 0.151059305 | 0.207083699 | 0.066028995 | 0.152307925 |
| 539,600 | 0.151220325 | 0.208066559 | 0.06681927 | 0.152583979 |
| 539,700 | 0.151036252 | 0.207473304 | 0.065861125 | 0.152258153 |
| 539,800 | 0.151010173 | 0.208055766 | 0.066567424 | 0.151531363 |
| 539,900 | 0.150978232 | 0.207703235 | 0.065600586 | 0.152033937 |
| 540,000 | 0.151058807 | 0.207288172 | 0.065973194 | 0.152283506 |
| 540,100 | 0.151142583 | 0.207231698 | 0.065001809 | 0.152150557 |
| 540,200 | 0.150991834 | 0.206660438 | 0.065142537 | 0.152315248 |
| 540,300 | 0.151127252 | 0.207352408 | 0.065520238 | 0.152078142 |
| 540,400 | 0.151044083 | 0.208350545 | 0.065636623 | 0.151884512 |
| 540,500 | 0.150965035 | 0.208096155 | 0.06562789 | 0.151911359 |
| 540,600 | 0.150887054 | 0.207758568 | 0.066068286 | 0.151760949 |
| 540,700 | 0.150888588 | 0.208245397 | 0.065775332 | 0.151389969 |
| 540,800 | 0.151009191 | 0.208286889 | 0.065477759 | 0.152375901 |
| 540,900 | 0.150911003 | 0.208023959 | 0.064826044 | 0.151704208 |
| 541,000 | 0.150884869 | 0.20864008 | 0.065283545 | 0.151714368 |
| 541,100 | 0.150747192 | 0.208180283 | 0.064769626 | 0.15198897 |
| 541,200 | 0.150796188 | 0.208215616 | 0.065227566 | 0.152188085 |
| 541,300 | 0.150853757 | 0.209317411 | 0.064822924 | 0.151840759 |
| 541,400 | 0.150792562 | 0.209163591 | 0.06512566 | 0.152063613 |
| 541,500 | 0.150839166 | 0.209057816 | 0.065239516 | 0.151734447 |
| 541,600 | 0.150796964 | 0.209265833 | 0.064937596 | 0.151704694 |

| | | | | |
|---|---|---|---|---|
| 541,700 | 0.150767059 | 0.209020385 | 0.065418509 | 0.152190666 |
| 541,800 | 0.150640562 | 0.208498036 | 0.06546682 | 0.151396778 |
| 541,900 | 0.150719706 | 0.209016098 | 0.065690158 | 0.151355334 |
| 542,000 | 0.150732297 | 0.207939441 | 0.064873343 | 0.151612616 |
| 542,100 | 0.150741641 | 0.207931898 | 0.065586659 | 0.151440675 |
| 542,200 | 0.150616541 | 0.207266069 | 0.065273508 | 0.151643474 |
| 542,300 | 0.15063111 | 0.208263214 | 0.065098388 | 0.150788618 |
| 542,400 | 0.150702104 | 0.207946289 | 0.065092061 | 0.15157806 |
| 542,500 | 0.150624166 | 0.207353175 | 0.065076309 | 0.151463364 |
| 542,600 | 0.150551781 | 0.207651532 | 0.064464256 | 0.150871519 |
| 542,700 | 0.150509948 | 0.20689788 | 0.064252529 | 0.151363964 |
| 542,800 | 0.150526258 | 0.206554251 | 0.064860475 | 0.15053902 |
| 542,900 | 0.150596062 | 0.20725527 | 0.064662081 | 0.151340915 |
| 543,000 | 0.150554916 | 0.207039327 | 0.06480823 | 0.151394161 |
| 543,100 | 0.150567575 | 0.207260412 | 0.064374252 | 0.150614868 |
| 543,200 | 0.150540207 | 0.206530473 | 0.064931726 | 0.151928139 |
| 543,300 | 0.150561862 | 0.206616923 | 0.064440808 | 0.15101086 |
| 543,400 | 0.150632214 | 0.206608029 | 0.065224288 | 0.151481311 |
| 543,500 | 0.15040113 | 0.207257533 | 0.065010725 | 0.151263693 |
| 543,600 | 0.150530212 | 0.207287043 | 0.064785229 | 0.151389075 |
| 543,700 | 0.150501283 | 0.20763886 | 0.06491694 | 0.150809644 |
| 543,800 | 0.150561554 | 0.207882047 | 0.064861616 | 0.150945286 |
| 543,900 | 0.150373411 | 0.207406456 | 0.064526322 | 0.151287896 |
| 544,000 | 0.150419301 | 0.20687086 | 0.063996323 | 0.151375036 |
| 544,100 | 0.15049201 | 0.207485871 | 0.065037395 | 0.150752871 |
| 544,200 | 0.150430696 | 0.207050541 | 0.065006755 | 0.151048814 |
| 544,300 | 0.150495674 | 0.207241264 | 0.064287342 | 0.151251238 |
| 544,400 | 0.150345694 | 0.207208317 | 0.064879895 | 0.151334943 |
| 544,500 | 0.150383001 | 0.207095387 | 0.06405289 | 0.151460269 |
| 544,600 | 0.150528799 | 0.207654641 | 0.064688574 | 0.151531112 |
| 544,700 | 0.150533097 | 0.207331817 | 0.064597226 | 0.151088923 |
| 544,800 | 0.150403972 | 0.207343135 | 0.064371155 | 0.150996943 |
| 544,900 | 0.150381511 | 0.207496121 | 0.064738624 | 0.150875858 |
| 545,000 | 0.150411351 | 0.207740023 | 0.064951377 | 0.150619349 |
| 545,100 | 0.150326479 | 0.206879757 | 0.064314754 | 0.151148567 |
| 545,200 | 0.150347199 | 0.207002312 | 0.064922098 | 0.151614318 |
| 545,300 | 0.150311143 | 0.207484103 | 0.064705911 | 0.15074892 |
| 545,400 | 0.150309681 | 0.207817436 | 0.064075726 | 0.150901867 |
| 545,500 | 0.150438421 | 0.207311933 | 0.064871823 | 0.15163736 |
| 545,600 | 0.150232941 | 0.206848558 | 0.064504139 | 0.150607507 |
| 545,700 | 0.150230997 | 0.206854422 | 0.065236415 | 0.15073084 |
| 545,800 | 0.150448219 | 0.206828852 | 0.064942153 | 0.150897622 |
| 545,900 | 0.150281747 | 0.206433295 | 0.064646864 | 0.150988707 |
| 546,000 | 0.15017859 | 0.206512374 | 0.064908431 | 0.151844472 |
| 546,100 | 0.1502118 | 0.206880272 | 0.063675392 | 0.151000303 |
| 546,200 | 0.150115704 | 0.206178876 | 0.064572276 | 0.151772955 |
| 546,300 | 0.150180706 | 0.206626229 | 0.064902868 | 0.151663387 |
| 546,400 | 0.150214585 | 0.20631291 | 0.064241245 | 0.15147731 |
| 546,500 | 0.150066123 | 0.206591735 | 0.06466573 | 0.151661483 |
| 546,600 | 0.150015167 | 0.205716362 | 0.064424718 | 0.151063285 |
| 546,700 | 0.150025241 | 0.206164106 | 0.064303356 | 0.151179685 |
| 546,800 | 0.149961537 | 0.20592811 | 0.064271483 | 0.15094676 |
| 546,900 | 0.150109023 | 0.205897607 | 0.064717099 | 0.151212135 |
| 547,000 | 0.15014107 | 0.205742205 | 0.064530476 | 0.150875623 |
| 547,100 | 0.150023954 | 0.205976055 | 0.064326232 | 0.150913067 |
| 547,200 | 0.150065859 | 0.205840653 | 0.06465223 | 0.150564447 |
| 547,300 | 0.149969995 | 0.206158624 | 0.064103679 | 0.15073893 |
| 547,400 | 0.150167921 | 0.20643016 | 0.064785238 | 0.151070898 |
| 547,500 | 0.15001217 | 0.205788221 | 0.064225144 | 0.150532492 |
| 547,600 | 0.149948163 | 0.206706951 | 0.064804371 | 0.150011394 |
| 547,700 | 0.150007272 | 0.206375909 | 0.064733473 | 0.150873514 |
| 547,800 | 0.14999663 | 0.205656355 | 0.064527132 | 0.150669505 |
| 547,900 | 0.149986098 | 0.20621354 | 0.064065791 | 0.150755459 |
| 548,000 | 0.149968542 | 0.205383361 | 0.063878282 | 0.150697321 |
| 548,100 | 0.150028759 | 0.206550964 | 0.063859932 | 0.150882871 |
| 548,200 | 0.149854722 | 0.205508432 | 0.064160257 | 0.150662109 |
| 548,300 | 0.149779359 | 0.205199966 | 0.064292009 | 0.150887403 |

| | | | | |
|---|---|---|---|---|
| 548,400 | 0.149872721 | 0.205554743 | 0.064190493 | 0.150862533 |
| 548,500 | 0.149706405 | 0.205537533 | 0.063382699 | 0.150582101 |
| 548,600 | 0.14973019 | 0.206169029 | 0.063745565 | 0.151094613 |
| 548,700 | 0.149717532 | 0.205568488 | 0.063609031 | 0.150368531 |
| 548,800 | 0.149793192 | 0.205390224 | 0.063899214 | 0.150726466 |
| 548,900 | 0.149743804 | 0.205114178 | 0.063367182 | 0.150907015 |
| 549,000 | 0.149766758 | 0.205471265 | 0.064230067 | 0.151206547 |
| 549,100 | 0.149584113 | 0.204655263 | 0.064409787 | 0.150713688 |
| 549,200 | 0.149647514 | 0.205095325 | 0.064711057 | 0.150762877 |
| 549,300 | 0.149717861 | 0.205144664 | 0.064092382 | 0.150647961 |
| 549,400 | 0.149705815 | 0.206145588 | 0.064772123 | 0.15079119 |
| 549,500 | 0.14964951 | 0.205989208 | 0.06443319 | 0.151062084 |
| 549,600 | 0.149736779 | 0.205374841 | 0.064508282 | 0.150784755 |
| 549,700 | 0.149698263 | 0.205943143 | 0.064477464 | 0.150934904 |
| 549,800 | 0.149677174 | 0.205189229 | 0.064826663 | 0.150576254 |
| 549,900 | 0.14958483 | 0.205485385 | 0.064318383 | 0.150997335 |
| 550,000 | 0.149644569 | 0.205424971 | 0.064770123 | 0.150695589 |
| 550,100 | 0.149464452 | 0.205471955 | 0.064873428 | 0.150337548 |
| 550,200 | 0.149521665 | 0.205322333 | 0.064379241 | 0.150525294 |
| 550,300 | 0.149559046 | 0.205316005 | 0.064838407 | 0.150991728 |
| 550,400 | 0.149494249 | 0.205093219 | 0.063953449 | 0.149746613 |
| 550,500 | 0.149591524 | 0.204876067 | 0.064465253 | 0.15059663 |
| 550,600 | 0.149531259 | 0.205432587 | 0.064543986 | 0.150523036 |
| 550,700 | 0.149612281 | 0.2049584 | 0.064223218 | 0.15049151 |
| 550,800 | 0.149392918 | 0.205138467 | 0.064279325 | 0.150159098 |
| 550,900 | 0.149389296 | 0.204922598 | 0.063790012 | 0.14942188 |
| 551,000 | 0.149365873 | 0.20486595 | 0.064449225 | 0.149710479 |
| 551,100 | 0.149559024 | 0.205220922 | 0.063961439 | 0.149963315 |
| 551,200 | 0.149432779 | 0.204485816 | 0.063160701 | 0.150106252 |
| 551,300 | 0.149483928 | 0.204071872 | 0.064134675 | 0.150176778 |
| 551,400 | 0.149396956 | 0.205038219 | 0.06425201 | 0.150046786 |
| 551,500 | 0.149478039 | 0.205298718 | 0.06413662 | 0.14988302 |
| 551,600 | 0.149241604 | 0.203809807 | 0.06409897 | 0.149929168 |
| 551,700 | 0.149285695 | 0.204371794 | 0.064084421 | 0.149977567 |
| 551,800 | 0.149349538 | 0.203567843 | 0.064520508 | 0.1498406 |
| 551,900 | 0.149311935 | 0.204067059 | 0.06415694 | 0.150310042 |
| 552,000 | 0.149397362 | 0.204184967 | 0.06456486 | 0.149984927 |
| 552,100 | 0.149323118 | 0.204858974 | 0.064747622 | 0.150270098 |
| 552,200 | 0.149230591 | 0.204206234 | 0.064468555 | 0.149855758 |
| 552,300 | 0.149304141 | 0.204706096 | 0.064267798 | 0.149601054 |
| 552,400 | 0.149240637 | 0.204638588 | 0.064329175 | 0.149064979 |
| 552,500 | 0.149256577 | 0.204369062 | 0.064559188 | 0.149586143 |
| 552,600 | 0.149230796 | 0.205087612 | 0.064589322 | 0.149627655 |
| 552,700 | 0.149338121 | 0.205449673 | 0.064708648 | 0.149218292 |
| 552,800 | 0.149192793 | 0.204639529 | 0.06460584 | 0.148848465 |
| 552,900 | 0.149128273 | 0.204641253 | 0.064255348 | 0.149339191 |
| 553,000 | 0.149099371 | 0.204025263 | 0.064544142 | 0.149241254 |
| 553,100 | 0.149046606 | 0.204272852 | 0.064311747 | 0.149302518 |
| 553,200 | 0.14910474 | 0.204491095 | 0.064289126 | 0.149059036 |
| 553,300 | 0.149085451 | 0.204493879 | 0.064036124 | 0.149480139 |
| 553,400 | 0.149123824 | 0.204109831 | 0.063883193 | 0.149390805 |
| 553,500 | 0.14906379 | 0.204608347 | 0.063704775 | 0.148864891 |
| 553,600 | 0.148924928 | 0.203951663 | 0.063465647 | 0.148069986 |
| 553,700 | 0.149089895 | 0.203715283 | 0.063547702 | 0.148961269 |
| 553,800 | 0.149142312 | 0.203866387 | 0.063932653 | 0.148543384 |
| 553,900 | 0.148933679 | 0.203908463 | 0.06422778 | 0.149255955 |
| 554,000 | 0.149038555 | 0.204152355 | 0.063843511 | 0.148804765 |
| 554,100 | 0.148981512 | 0.204429416 | 0.064189515 | 0.148850466 |
| 554,200 | 0.148892677 | 0.204716589 | 0.064562467 | 0.148697397 |
| 554,300 | 0.14878722 | 0.204800649 | 0.064398354 | 0.148282846 |
| 554,400 | 0.148978089 | 0.204955286 | 0.063863812 | 0.148969651 |
| 554,500 | 0.148898846 | 0.204671833 | 0.064066045 | 0.149202027 |
| 554,600 | 0.148887726 | 0.204211641 | 0.064719533 | 0.148330835 |
| 554,700 | 0.148762456 | 0.204448484 | 0.064153084 | 0.148720194 |
| 554,800 | 0.14894515 | 0.204126547 | 0.064153618 | 0.148923866 |
| 554,900 | 0.148799752 | 0.20405281 | 0.064449332 | 0.148395642 |
| 555,000 | 0.14879805 | 0.203955986 | 0.064452743 | 0.148728834 |

| | | | | |
|---|---|---|---|---|
| 555,100 | 0.148768803 | 0.20467997 | 0.064155733 | 0.147912985 |
| 555,200 | 0.148777884 | 0.204694968 | 0.064165721 | 0.148063682 |
| 555,300 | 0.148771488 | 0.204086756 | 0.063328253 | 0.148510852 |
| 555,400 | 0.148724214 | 0.204024644 | 0.063826429 | 0.147872944 |
| 555,500 | 0.148739708 | 0.203957812 | 0.06413746 | 0.148083616 |
| 555,600 | 0.148714006 | 0.202956506 | 0.063788953 | 0.147581772 |
| 555,700 | 0.148747948 | 0.204305673 | 0.063742657 | 0.147881461 |
| 555,800 | 0.148738019 | 0.203877879 | 0.064166404 | 0.148244908 |
| 555,900 | 0.148753457 | 0.203966524 | 0.064115749 | 0.148220403 |
| 556,000 | 0.148679274 | 0.203050914 | 0.063753903 | 0.148192862 |
| 556,100 | 0.148624189 | 0.204436526 | 0.064191663 | 0.147845557 |
| 556,200 | 0.148660869 | 0.204055524 | 0.063962326 | 0.148180633 |
| 556,300 | 0.148665027 | 0.204059177 | 0.064292879 | 0.148304334 |
| 556,400 | 0.148623792 | 0.204191032 | 0.064470405 | 0.148125472 |
| 556,500 | 0.148581422 | 0.204770806 | 0.064232401 | 0.148397628 |
| 556,600 | 0.148467547 | 0.205076994 | 0.064187642 | 0.148077061 |
| 556,700 | 0.148595912 | 0.204643244 | 0.063849388 | 0.148024247 |
| 556,800 | 0.148474698 | 0.204343487 | 0.063590979 | 0.148448691 |
| 556,900 | 0.148501158 | 0.205018858 | 0.0641021 | 0.147989109 |
| 557,000 | 0.148305479 | 0.204185924 | 0.064602737 | 0.147957816 |
| 557,100 | 0.148338538 | 0.204271852 | 0.064478931 | 0.148170674 |
| 557,200 | 0.148330032 | 0.204560559 | 0.064530935 | 0.148642862 |
| 557,300 | 0.148387552 | 0.203671901 | 0.064217357 | 0.147959913 |
| 557,400 | 0.148251287 | 0.20364571 | 0.064608965 | 0.147753277 |
| 557,500 | 0.148401325 | 0.204134727 | 0.064581418 | 0.147974475 |
| 557,600 | 0.148321435 | 0.204632107 | 0.064163468 | 0.147683643 |
| 557,700 | 0.148374304 | 0.203929691 | 0.064719495 | 0.147782155 |
| 557,800 | 0.148370078 | 0.204138089 | 0.064314303 | 0.147840278 |
| 557,900 | 0.148409793 | 0.203953016 | 0.064428195 | 0.147870249 |
| 558,000 | 0.148390869 | 0.203732902 | 0.064967136 | 0.147688984 |
| 558,100 | 0.148364905 | 0.203695899 | 0.064685409 | 0.147433541 |
| 558,200 | 0.148376984 | 0.203231048 | 0.064493556 | 0.147849761 |
| 558,300 | 0.148210012 | 0.202863605 | 0.064307955 | 0.147903274 |
| 558,400 | 0.14821442 | 0.203214257 | 0.063921497 | 0.147890318 |
| 558,500 | 0.148459924 | 0.203352335 | 0.064559602 | 0.148141375 |
| 558,600 | 0.148374051 | 0.204091819 | 0.06433061 | 0.148614535 |
| 558,700 | 0.148484368 | 0.204076162 | 0.064528713 | 0.148002349 |
| 558,800 | 0.148282469 | 0.203202344 | 0.063865224 | 0.147753172 |
| 558,900 | 0.148409315 | 0.203368408 | 0.063861466 | 0.147956402 |
| 559,000 | 0.148584108 | 0.203421189 | 0.064311628 | 0.148438429 |
| 559,100 | 0.14836232 | 0.203377497 | 0.063890463 | 0.1482419 |
| 559,200 | 0.148321653 | 0.20307338 | 0.06342837 | 0.14822505 |
| 559,300 | 0.14838466 | 0.203813388 | 0.063450592 | 0.148204927 |
| 559,400 | 0.148361849 | 0.203900122 | 0.063801253 | 0.147686948 |
| 559,500 | 0.148303088 | 0.202929962 | 0.063123072 | 0.147793045 |
| 559,600 | 0.148210518 | 0.203335046 | 0.063964227 | 0.147706172 |
| 559,700 | 0.148346673 | 0.202990272 | 0.06396122 | 0.148078821 |
| 559,800 | 0.148192272 | 0.202968378 | 0.064559994 | 0.147447458 |
| 559,900 | 0.148221251 | 0.202490009 | 0.063229443 | 0.147459339 |
| 560,000 | 0.148341127 | 0.203362532 | 0.063908205 | 0.147497755 |
| 560,100 | 0.148281166 | 0.203035101 | 0.063845411 | 0.147722328 |
| 560,200 | 0.14825365 | 0.203070099 | 0.064282208 | 0.147680476 |
| 560,300 | 0.148166459 | 0.203227971 | 0.063480505 | 0.147703414 |
| 560,400 | 0.14836503 | 0.203116787 | 0.063847197 | 0.14860545 |
| 560,500 | 0.148292728 | 0.203797147 | 0.0637482 | 0.148253566 |
| 560,600 | 0.148179514 | 0.203501594 | 0.063949732 | 0.147644683 |
| 560,700 | 0.148103767 | 0.203583354 | 0.064221043 | 0.147504021 |
| 560,800 | 0.148217798 | 0.202976463 | 0.064551293 | 0.147617376 |
| 560,900 | 0.148093125 | 0.203380688 | 0.063931525 | 0.147822174 |
| 561,000 | 0.148069356 | 0.203852382 | 0.06470738 | 0.147783666 |
| 561,100 | 0.148124018 | 0.203781283 | 0.06417701 | 0.147663904 |
| 561,200 | 0.148281154 | 0.20314785 | 0.063253113 | 0.146947317 |
| 561,300 | 0.148178978 | 0.203041705 | 0.063791666 | 0.147217399 |
| 561,400 | 0.147960947 | 0.202939964 | 0.064071428 | 0.147806708 |
| 561,500 | 0.147881682 | 0.202838955 | 0.063576009 | 0.147324912 |
| 561,600 | 0.147932381 | 0.203522813 | 0.063971772 | 0.147168246 |
| 561,700 | 0.147822392 | 0.20233962 | 0.064301158 | 0.147709652 |

| | | | | |
|---|---|---|---|---|
| 561,800 | 0.14790504 | 0.202727488 | 0.064281449 | 0.147320971 |
| 561,900 | 0.147897039 | 0.203662571 | 0.063789031 | 0.147897948 |
| 562,000 | 0.147833801 | 0.20355892 | 0.063618338 | 0.14719306 |
| 562,100 | 0.147816526 | 0.203363561 | 0.063105804 | 0.147303413 |
| 562,200 | 0.147840306 | 0.203380946 | 0.063964415 | 0.147525879 |
| 562,300 | 0.147964648 | 0.202447707 | 0.063898412 | 0.147563159 |
| 562,400 | 0.147859989 | 0.202684744 | 0.063383176 | 0.147477031 |
| 562,500 | 0.14783919 | 0.202698853 | 0.063557371 | 0.147379722 |
| 562,600 | 0.147736788 | 0.202896326 | 0.064012833 | 0.147378479 |
| 562,700 | 0.147837109 | 0.202957188 | 0.064099 | 0.1465066 |
| 562,800 | 0.147668925 | 0.202938445 | 0.064156447 | 0.147091108 |
| 562,900 | 0.147781001 | 0.202865152 | 0.06314136 | 0.146937159 |
| 563,000 | 0.147715793 | 0.203566351 | 0.063330818 | 0.146667726 |
| 563,100 | 0.147750738 | 0.202958797 | 0.063560693 | 0.146895603 |
| 563,200 | 0.147794093 | 0.20231865 | 0.063572865 | 0.14713848 |
| 563,300 | 0.147861852 | 0.20258364 | 0.06310594 | 0.147404622 |
| 563,400 | 0.14775953 | 0.202656828 | 0.063381981 | 0.14699311 |
| 563,500 | 0.147748637 | 0.202374615 | 0.063266756 | 0.146960615 |
| 563,600 | 0.147569199 | 0.202317226 | 0.063519062 | 0.146906732 |
| 563,700 | 0.147610875 | 0.202320474 | 0.062956124 | 0.146583073 |
| 563,800 | 0.147499444 | 0.202654022 | 0.063429561 | 0.146114735 |
| 563,900 | 0.147465426 | 0.202000753 | 0.063674602 | 0.146696327 |
| 564,000 | 0.147421228 | 0.201827973 | 0.063907668 | 0.146392844 |
| 564,100 | 0.14742304 | 0.202408823 | 0.063848981 | 0.147171512 |
| 564,200 | 0.147485982 | 0.202627196 | 0.063113618 | 0.146302976 |
| 564,300 | 0.147450559 | 0.201565733 | 0.063491263 | 0.146265918 |
| 564,400 | 0.147422438 | 0.201913582 | 0.063909742 | 0.146338803 |
| 564,500 | 0.147320257 | 0.201897263 | 0.0634894 | 0.146517511 |
| 564,600 | 0.147406279 | 0.202638962 | 0.063701053 | 0.14618783 |
| 564,700 | 0.147232282 | 0.20237056 | 0.063372349 | 0.146080617 |
| 564,800 | 0.147348292 | 0.202389049 | 0.063525938 | 0.145754958 |
| 564,900 | 0.14727897 | 0.202262057 | 0.063087046 | 0.146696609 |
| 565,000 | 0.147262157 | 0.202819766 | 0.063049701 | 0.146439686 |
| 565,100 | 0.147260079 | 0.202269257 | 0.063553052 | 0.146113411 |
| 565,200 | 0.147328699 | 0.203137384 | 0.063258168 | 0.146001251 |
| 565,300 | 0.147139692 | 0.201937639 | 0.063849795 | 0.145573616 |
| 565,400 | 0.147275404 | 0.201763728 | 0.063273663 | 0.145480015 |
| 565,500 | 0.147286521 | 0.202544845 | 0.063473533 | 0.145471005 |
| 565,600 | 0.147350645 | 0.202985116 | 0.063091641 | 0.146104095 |
| 565,700 | 0.147221837 | 0.202743707 | 0.063248539 | 0.145649806 |
| 565,800 | 0.147165455 | 0.201951035 | 0.063792172 | 0.146151214 |
| 565,900 | 0.147211091 | 0.201997182 | 0.063293598 | 0.146072424 |
| 566,000 | 0.14731741 | 0.201844841 | 0.063613764 | 0.145697425 |
| 566,100 | 0.147210213 | 0.202133556 | 0.063006466 | 0.145750333 |
| 566,200 | 0.147189576 | 0.20149036 | 0.063811627 | 0.145830839 |
| 566,300 | 0.147157683 | 0.202106475 | 0.064103193 | 0.145650432 |
| 566,400 | 0.147190535 | 0.202388593 | 0.064258489 | 0.146058683 |
| 566,500 | 0.147114853 | 0.20267085 | 0.064007594 | 0.145088695 |
| 566,600 | 0.147140876 | 0.202837102 | 0.064016615 | 0.145308643 |
| 566,700 | 0.147154651 | 0.202992898 | 0.064283554 | 0.145207708 |
| 566,800 | 0.147137767 | 0.202730219 | 0.063836483 | 0.145278664 |
| 566,900 | 0.147011482 | 0.201671482 | 0.063362056 | 0.145702939 |
| 567,000 | 0.14708227 | 0.202106671 | 0.064035612 | 0.144925684 |
| 567,100 | 0.147039971 | 0.201835862 | 0.063953189 | 0.145645098 |
| 567,200 | 0.147003652 | 0.202291496 | 0.063798656 | 0.145490976 |
| 567,300 | 0.14698792 | 0.202160854 | 0.064395397 | 0.145490582 |
| 567,400 | 0.147072844 | 0.202038078 | 0.064053775 | 0.145871991 |
| 567,500 | 0.14710731 | 0.202791792 | 0.063849907 | 0.145840668 |
| 567,600 | 0.146955257 | 0.202288894 | 0.06388682 | 0.145680758 |
| 567,700 | 0.14694884 | 0.201596384 | 0.063686036 | 0.145736533 |
| 567,800 | 0.146868725 | 0.202021605 | 0.063204946 | 0.145792851 |
| 567,900 | 0.146991718 | 0.201636474 | 0.062999965 | 0.144804705 |
| 568,000 | 0.146937533 | 0.201336329 | 0.063928237 | 0.145550075 |
| 568,100 | 0.146864196 | 0.201421145 | 0.063858198 | 0.14542327 |
| 568,200 | 0.146851578 | 0.201796745 | 0.063700979 | 0.144985191 |
| 568,300 | 0.146830123 | 0.201647235 | 0.063114768 | 0.145035716 |
| 568,400 | 0.146816712 | 0.201677287 | 0.062925896 | 0.145328706 |

| | | | | |
|---|---|---|---|---|
| 568,500 | 0.146764579 | 0.202479186 | 0.063839832 | 0.145026865 |
| 568,600 | 0.146825852 | 0.202288409 | 0.063495127 | 0.145446871 |
| 568,700 | 0.146752035 | 0.202582136 | 0.063306638 | 0.145194142 |
| 568,800 | 0.146746372 | 0.20201391 | 0.062902212 | 0.145249924 |
| 568,900 | 0.14671442 | 0.202561811 | 0.063128536 | 0.144882006 |
| 569,000 | 0.146669325 | 0.201856624 | 0.063184741 | 0.144909048 |
| 569,100 | 0.146715241 | 0.201593398 | 0.062808288 | 0.144892977 |
| 569,200 | 0.146682921 | 0.201696143 | 0.063272195 | 0.145173861 |
| 569,300 | 0.146608613 | 0.201281134 | 0.062760517 | 0.145162478 |
| 569,400 | 0.146652168 | 0.201816561 | 0.062883598 | 0.14533127 |
| 569,500 | 0.146558418 | 0.201670734 | 0.062942137 | 0.145247182 |
| 569,600 | 0.146576917 | 0.201189001 | 0.06301282 | 0.145059791 |
| 569,700 | 0.146567248 | 0.202012057 | 0.063577261 | 0.144973188 |
| 569,800 | 0.146472764 | 0.201949233 | 0.063152196 | 0.144506174 |
| 569,900 | 0.146483895 | 0.202061915 | 0.062841248 | 0.145115097 |
| 570,000 | 0.146528727 | 0.202252493 | 0.062209779 | 0.145056935 |
| 570,100 | 0.146521742 | 0.201970064 | 0.062586813 | 0.144773107 |
| 570,200 | 0.146438597 | 0.201598483 | 0.062737214 | 0.145704851 |
| 570,300 | 0.14647716 | 0.201636874 | 0.062490207 | 0.145520956 |
| 570,400 | 0.146494939 | 0.201232477 | 0.062855047 | 0.145073385 |
| 570,500 | 0.146569986 | 0.201919965 | 0.062757551 | 0.144894777 |
| 570,600 | 0.146510847 | 0.202131103 | 0.062037174 | 0.144364182 |
| 570,700 | 0.146426143 | 0.20194908 | 0.062831311 | 0.144647322 |
| 570,800 | 0.146481365 | 0.201781896 | 0.061866898 | 0.144626308 |
| 570,900 | 0.146449705 | 0.201654363 | 0.0627337 | 0.144640682 |
| 571,000 | 0.146396414 | 0.202068196 | 0.06225207 | 0.145013341 |
| 571,100 | 0.14635266 | 0.201690445 | 0.062021779 | 0.144685941 |
| 571,200 | 0.146272051 | 0.201429953 | 0.061796496 | 0.144705749 |
| 571,300 | 0.146418547 | 0.201655535 | 0.061979138 | 0.144735776 |
| 571,400 | 0.146311016 | 0.201221892 | 0.061948266 | 0.145070212 |
| 571,500 | 0.146259232 | 0.200517707 | 0.062749732 | 0.144565664 |
| 571,600 | 0.146249813 | 0.201602152 | 0.062748375 | 0.144447132 |
| 571,700 | 0.146331901 | 0.201887071 | 0.062905282 | 0.145429504 |
| 571,800 | 0.146180233 | 0.201760166 | 0.061820488 | 0.144785234 |
| 571,900 | 0.146163857 | 0.202004204 | 0.062335095 | 0.144424516 |
| 572,000 | 0.146144687 | 0.201979889 | 0.061735056 | 0.144159479 |
| 572,100 | 0.146176764 | 0.201132888 | 0.062216543 | 0.144728602 |
| 572,200 | 0.146250876 | 0.200930527 | 0.062173739 | 0.144961915 |
| 572,300 | 0.146102106 | 0.201449753 | 0.062224892 | 0.144693642 |
| 572,400 | 0.146104901 | 0.201245766 | 0.062691524 | 0.144467182 |
| 572,500 | 0.14624031 | 0.20190181 | 0.062588565 | 0.144660443 |
| 572,600 | 0.146093385 | 0.20098586 | 0.062617211 | 0.14518173 |
| 572,700 | 0.146025447 | 0.201876099 | 0.062047877 | 0.144740044 |
| 572,800 | 0.145882591 | 0.200723835 | 0.062319719 | 0.14514316 |
| 572,900 | 0.146017971 | 0.201531846 | 0.061980518 | 0.144644664 |
| 573,000 | 0.146060167 | 0.200720287 | 0.062454743 | 0.145050777 |
| 573,100 | 0.145990134 | 0.200752901 | 0.062362217 | 0.14494031 |
| 573,200 | 0.14609937 | 0.201211119 | 0.062164118 | 0.145113377 |
| 573,300 | 0.145920105 | 0.201562962 | 0.062174348 | 0.144944438 |
| 573,400 | 0.146040896 | 0.201544334 | 0.062333001 | 0.144415429 |
| 573,500 | 0.145907389 | 0.201438663 | 0.062162609 | 0.144551756 |
| 573,600 | 0.146039294 | 0.200670357 | 0.062747343 | 0.144745373 |
| 573,700 | 0.14587023 | 0.20113009 | 0.062089332 | 0.144602987 |
| 573,800 | 0.145916975 | 0.201236505 | 0.062224005 | 0.144764868 |
| 573,900 | 0.145781022 | 0.200970479 | 0.062237083 | 0.144659943 |
| 574,000 | 0.145938511 | 0.200490275 | 0.061881868 | 0.144926145 |
| 574,100 | 0.145881215 | 0.201288132 | 0.062689168 | 0.144477395 |
| 574,200 | 0.145882587 | 0.201569248 | 0.063082368 | 0.144548997 |
| 574,300 | 0.145990471 | 0.201669609 | 0.06312925 | 0.145117005 |
| 574,400 | 0.145903138 | 0.201925649 | 0.062880529 | 0.144824422 |
| 574,500 | 0.145910948 | 0.201411849 | 0.06235834 | 0.144445139 |
| 574,600 | 0.14586631 | 0.201652668 | 0.061320864 | 0.14436339 |
| 574,700 | 0.145941697 | 0.20109324 | 0.06161426 | 0.14465316 |
| 574,800 | 0.145980578 | 0.201868858 | 0.061596946 | 0.144384266 |
| 574,900 | 0.145917515 | 0.202337861 | 0.062469925 | 0.144394012 |
| 575,000 | 0.145882484 | 0.201580948 | 0.062393781 | 0.14384461 |
| 575,100 | 0.145788797 | 0.201145315 | 0.062568071 | 0.143972823 |

| | | | | |
|---|---|---|---|---|
| 575,200 | 0.145819008 | 0.200617255 | 0.062448419 | 0.143760251 |
| 575,300 | 0.14582894 | 0.202167621 | 0.061985791 | 0.144467625 |
| 575,400 | 0.14585849 | 0.202173917 | 0.062235936 | 0.144340589 |
| 575,500 | 0.145956971 | 0.202006837 | 0.062446629 | 0.144347151 |
| 575,600 | 0.145733192 | 0.201304659 | 0.062891787 | 0.14423705 |
| 575,700 | 0.145753428 | 0.201564901 | 0.062603943 | 0.144232124 |
| 575,800 | 0.145840008 | 0.201523129 | 0.06166146 | 0.144263689 |
| 575,900 | 0.145835937 | 0.201822901 | 0.061850592 | 0.144473671 |
| 576,000 | 0.145673674 | 0.201607383 | 0.062421979 | 0.143555768 |
| 576,100 | 0.145827134 | 0.201724881 | 0.061982401 | 0.144133911 |
| 576,200 | 0.145768922 | 0.202020067 | 0.062384631 | 0.143560793 |
| 576,300 | 0.145663748 | 0.202271586 | 0.063119274 | 0.143358294 |
| 576,400 | 0.14561169 | 0.201445666 | 0.062664232 | 0.143905697 |
| 576,500 | 0.145619245 | 0.201457696 | 0.062585987 | 0.143523587 |
| 576,600 | 0.145737782 | 0.20180906 | 0.06218903 | 0.14366504 |
| 576,700 | 0.145717323 | 0.201702678 | 0.062587292 | 0.143800307 |
| 576,800 | 0.145690516 | 0.202570893 | 0.062815147 | 0.143649 |
| 576,900 | 0.145690294 | 0.202141323 | 0.06244483 | 0.143083874 |
| 577,000 | 0.145757178 | 0.201461192 | 0.062940563 | 0.143491251 |
| 577,100 | 0.145623078 | 0.201563201 | 0.063429134 | 0.143011538 |
| 577,200 | 0.145584371 | 0.201315421 | 0.063110986 | 0.143510874 |
| 577,300 | 0.145534826 | 0.201169905 | 0.062829103 | 0.142919599 |
| 577,400 | 0.145620954 | 0.201489832 | 0.06328119 | 0.143678058 |
| 577,500 | 0.14560583 | 0.200950044 | 0.063397216 | 0.143353841 |
| 577,600 | 0.145603456 | 0.201233089 | 0.063544021 | 0.143217405 |
| 577,700 | 0.14546498 | 0.200517828 | 0.062859048 | 0.143562389 |
| 577,800 | 0.145474685 | 0.20033791 | 0.063715171 | 0.143079269 |
| 577,900 | 0.145427518 | 0.20064204 | 0.063888142 | 0.143649756 |
| 578,000 | 0.145497706 | 0.201256051 | 0.06315637 | 0.143331142 |
| 578,100 | 0.145447387 | 0.20073591 | 0.06312909 | 0.143529843 |
| 578,200 | 0.145480798 | 0.201463732 | 0.062505633 | 0.143642108 |
| 578,300 | 0.145501449 | 0.200679518 | 0.062953586 | 0.14379536 |
| 578,400 | 0.145311173 | 0.200507729 | 0.063005847 | 0.14276748 |
| 578,500 | 0.145249801 | 0.199794955 | 0.063004646 | 0.143405334 |
| 578,600 | 0.145286886 | 0.200597559 | 0.062668829 | 0.143562047 |
| 578,700 | 0.145395339 | 0.200518107 | 0.062214339 | 0.143756742 |
| 578,800 | 0.145258298 | 0.200324743 | 0.062076115 | 0.143409591 |
| 578,900 | 0.145242916 | 0.200261788 | 0.062442578 | 0.143438399 |
| 579,000 | 0.145294177 | 0.200670221 | 0.062569957 | 0.142990636 |
| 579,100 | 0.145361636 | 0.201092748 | 0.062452702 | 0.142616425 |
| 579,200 | 0.145385859 | 0.200877895 | 0.062928193 | 0.143016122 |
| 579,300 | 0.145334079 | 0.200424261 | 0.062783972 | 0.14308734 |
| 579,400 | 0.145287855 | 0.200537836 | 0.062639296 | 0.143346366 |
| 579,500 | 0.145416852 | 0.200467341 | 0.063422549 | 0.143555545 |
| 579,600 | 0.145360227 | 0.200214913 | 0.06374508 | 0.143768205 |
| 579,700 | 0.145238346 | 0.199820027 | 0.063459631 | 0.143823884 |
| 579,800 | 0.145266892 | 0.200235235 | 0.063327502 | 0.143781278 |
| 579,900 | 0.145253543 | 0.199360869 | 0.0636248 | 0.143342188 |
| 580,000 | 0.145187372 | 0.200153871 | 0.063268061 | 0.143462178 |
| 580,100 | 0.145241966 | 0.200232786 | 0.063731584 | 0.14380911 |
| 580,200 | 0.145257058 | 0.200521481 | 0.063792502 | 0.143696675 |
| 580,300 | 0.145231069 | 0.200361176 | 0.062803883 | 0.143696717 |
| 580,400 | 0.145299193 | 0.201106934 | 0.06328257 | 0.143792935 |
| 580,500 | 0.145281988 | 0.200654258 | 0.063241978 | 0.142986568 |
| 580,600 | 0.145283666 | 0.200598182 | 0.064038675 | 0.143339425 |
| 580,700 | 0.145244302 | 0.20145656 | 0.063659832 | 0.143153265 |
| 580,800 | 0.145184292 | 0.200926252 | 0.063549132 | 0.143311628 |
| 580,900 | 0.145173398 | 0.200797122 | 0.062894315 | 0.143136382 |
| 581,000 | 0.145083258 | 0.200163875 | 0.063368722 | 0.14319121 |
| 581,100 | 0.145161756 | 0.200699167 | 0.06304548 | 0.142875637 |
| 581,200 | 0.145077312 | 0.200019961 | 0.063407992 | 0.142395691 |
| 581,300 | 0.144988372 | 0.200670351 | 0.06243223 | 0.143090149 |
| 581,400 | 0.145173974 | 0.201140404 | 0.062726091 | 0.143117427 |
| 581,500 | 0.145172617 | 0.201438056 | 0.062684944 | 0.142669544 |
| 581,600 | 0.145181526 | 0.200461798 | 0.063016707 | 0.142997965 |
| 581,700 | 0.145114438 | 0.200929937 | 0.061961211 | 0.142868655 |
| 581,800 | 0.145262158 | 0.200610433 | 0.062848372 | 0.142767015 |

| | | | | |
|---|---|---|---|---|
| 581,900 | 0.145204704 | 0.201196157 | 0.06224355 | 0.142975109 |
| 582,000 | 0.145255995 | 0.200326916 | 0.062271734 | 0.143000643 |
| 582,100 | 0.145106972 | 0.200411274 | 0.062030842 | 0.143234678 |
| 582,200 | 0.144994119 | 0.200050851 | 0.062393309 | 0.1429601 |
| 582,300 | 0.144970744 | 0.200216335 | 0.062293753 | 0.142984348 |
| 582,400 | 0.145120797 | 0.200219037 | 0.062167758 | 0.143251369 |
| 582,500 | 0.145075367 | 0.200305644 | 0.062105679 | 0.143841443 |
| 582,600 | 0.145013434 | 0.200155575 | 0.061742453 | 0.143734729 |
| 582,700 | 0.1450375 | 0.199733355 | 0.06165942 | 0.142985879 |
| 582,800 | 0.144926699 | 0.19995954 | 0.061979545 | 0.14294098 |
| 582,900 | 0.144922966 | 0.199413313 | 0.062224285 | 0.143355662 |
| 583,000 | 0.144823283 | 0.200060292 | 0.062449496 | 0.142440327 |
| 583,100 | 0.144915141 | 0.199889783 | 0.062032637 | 0.142975145 |
| 583,200 | 0.144838394 | 0.199276756 | 0.062480041 | 0.142837994 |
| 583,300 | 0.144904054 | 0.199374285 | 0.061889775 | 0.142910696 |
| 583,400 | 0.144960894 | 0.200157074 | 0.062567686 | 0.142889857 |
| 583,500 | 0.145012768 | 0.200681189 | 0.062845966 | 0.142494619 |
| 583,600 | 0.145127156 | 0.200325114 | 0.062949471 | 0.143375371 |
| 583,700 | 0.144996633 | 0.19994591 | 0.062753867 | 0.142790221 |
| 583,800 | 0.145115296 | 0.200054259 | 0.063221422 | 0.142851984 |
| 583,900 | 0.144864977 | 0.200370326 | 0.062426711 | 0.142870397 |
| 584,000 | 0.144980236 | 0.200779952 | 0.06316771 | 0.143355559 |
| 584,100 | 0.144880006 | 0.201066522 | 0.06276758 | 0.142470097 |
| 584,200 | 0.144946054 | 0.200722286 | 0.063387603 | 0.142422039 |
| 584,300 | 0.144803297 | 0.200595343 | 0.063000217 | 0.142322769 |
| 584,400 | 0.144889367 | 0.201563245 | 0.063057195 | 0.143079627 |
| 584,500 | 0.144850161 | 0.200843079 | 0.063321564 | 0.142379184 |
| 584,600 | 0.144895188 | 0.201076367 | 0.063403116 | 0.141371197 |
| 584,700 | 0.144873813 | 0.200474869 | 0.063172164 | 0.141679146 |
| 584,800 | 0.144832409 | 0.200093159 | 0.063393642 | 0.142613197 |
| 584,900 | 0.144618132 | 0.199658748 | 0.063123451 | 0.142407447 |
| 585,000 | 0.144798546 | 0.200681616 | 0.062560527 | 0.141985893 |
| 585,100 | 0.144827443 | 0.199900416 | 0.06329722 | 0.14211648 |
| 585,200 | 0.144762057 | 0.199855783 | 0.062633721 | 0.141868351 |
| 585,300 | 0.14480685 | 0.200170326 | 0.062341731 | 0.14157703 |
| 585,400 | 0.144797299 | 0.200121408 | 0.062931333 | 0.142031923 |
| 585,500 | 0.144881113 | 0.200196365 | 0.062494554 | 0.142190388 |
| 585,600 | 0.144867775 | 0.200367562 | 0.062427488 | 0.142295933 |
| 585,700 | 0.144910628 | 0.200283369 | 0.06287023 | 0.14241824 |
| 585,800 | 0.14473338 | 0.199865468 | 0.062052244 | 0.142140548 |
| 585,900 | 0.144673646 | 0.20042948 | 0.062977704 | 0.141793526 |
| 586,000 | 0.144725026 | 0.200200985 | 0.062543274 | 0.141523277 |
| 586,100 | 0.144615166 | 0.200154966 | 0.063173778 | 0.142477885 |
| 586,200 | 0.144594164 | 0.19994757 | 0.063650597 | 0.142448522 |
| 586,300 | 0.144601252 | 0.200099235 | 0.06353821 | 0.142249215 |
| 586,400 | 0.144696306 | 0.19974888 | 0.062988626 | 0.14227084 |
| 586,500 | 0.144855461 | 0.200537032 | 0.063992535 | 0.142735297 |
| 586,600 | 0.144755719 | 0.199441421 | 0.064200256 | 0.142134283 |
| 586,700 | 0.144843188 | 0.20052663 | 0.063252647 | 0.142987531 |
| 586,800 | 0.144708655 | 0.199291164 | 0.06315736 | 0.142708433 |
| 586,900 | 0.144730783 | 0.199670757 | 0.063822588 | 0.141974915 |
| 587,000 | 0.144765316 | 0.199709719 | 0.063378679 | 0.142019777 |
| 587,100 | 0.144637561 | 0.199331177 | 0.062846943 | 0.141772553 |
| 587,200 | 0.144560285 | 0.199294749 | 0.062759987 | 0.141764065 |
| 587,300 | 0.144521313 | 0.19829373 | 0.062439233 | 0.141564956 |
| 587,400 | 0.14451916 | 0.198628054 | 0.062915378 | 0.141083263 |
| 587,500 | 0.14450022 | 0.199373196 | 0.062901316 | 0.141054863 |
| 587,600 | 0.14450356 | 0.198250597 | 0.063302077 | 0.141474848 |
| 587,700 | 0.144445466 | 0.198281916 | 0.062792208 | 0.141090519 |
| 587,800 | 0.144333592 | 0.198261513 | 0.062533831 | 0.140756137 |
| 587,900 | 0.144370322 | 0.197914678 | 0.063220281 | 0.141498853 |
| 588,000 | 0.144555424 | 0.198239405 | 0.063204488 | 0.141964821 |
| 588,100 | 0.144351539 | 0.198348261 | 0.063034507 | 0.141744693 |
| 588,200 | 0.144221799 | 0.198143041 | 0.062671128 | 0.140986991 |
| 588,300 | 0.144313751 | 0.198861721 | 0.063151117 | 0.14166698 |
| 588,400 | 0.144278718 | 0.1978795 | 0.062404045 | 0.141831601 |
| 588,500 | 0.144156734 | 0.198073657 | 0.062761404 | 0.141703981 |

| | | | | |
|---|---|---|---|---|
| 588,600 | 0.14431804 | 0.197957379 | 0.062368683 | 0.141560706 |
| 588,700 | 0.144271352 | 0.198317453 | 0.062615895 | 0.141492004 |
| 588,800 | 0.144376214 | 0.198125424 | 0.062217843 | 0.141326599 |
| 588,900 | 0.144317043 | 0.198715146 | 0.062123188 | 0.141378047 |
| 589,000 | 0.14437841 | 0.19879015 | 0.061971059 | 0.141385182 |
| 589,100 | 0.144296419 | 0.198436459 | 0.062685023 | 0.141014162 |
| 589,200 | 0.144356851 | 0.198079849 | 0.062494698 | 0.14146935 |
| 589,300 | 0.144311129 | 0.197013407 | 0.06220482 | 0.141748655 |
| 589,400 | 0.144260822 | 0.197757497 | 0.062436285 | 0.141529873 |
| 589,500 | 0.144262861 | 0.197345192 | 0.062090187 | 0.141319778 |
| 589,600 | 0.144359248 | 0.198241423 | 0.062322817 | 0.141594564 |
| 589,700 | 0.144168004 | 0.198011742 | 0.062082792 | 0.141215658 |
| 589,800 | 0.144126186 | 0.197263251 | 0.062653958 | 0.141390134 |
| 589,900 | 0.144331369 | 0.197907944 | 0.062219833 | 0.141683307 |
| 590,000 | 0.144226181 | 0.197973205 | 0.062097699 | 0.141529508 |
| 590,100 | 0.144231025 | 0.198774125 | 0.061661803 | 0.141894791 |
| 590,200 | 0.144154897 | 0.198371958 | 0.061963774 | 0.141814071 |
| 590,300 | 0.144141327 | 0.19778997 | 0.061910162 | 0.141105311 |
| 590,400 | 0.144172223 | 0.198333586 | 0.062044595 | 0.141428749 |
| 590,500 | 0.144253718 | 0.198285126 | 0.062170822 | 0.141809175 |
| 590,600 | 0.144135268 | 0.198099556 | 0.06235493 | 0.141649083 |
| 590,700 | 0.144247728 | 0.197747335 | 0.061711811 | 0.141572447 |
| 590,800 | 0.144257368 | 0.197989738 | 0.061822166 | 0.141779201 |
| 590,900 | 0.144118743 | 0.197362733 | 0.062312876 | 0.141017865 |
| 591,000 | 0.144102692 | 0.197383787 | 0.061004349 | 0.141898037 |
| 591,100 | 0.144107812 | 0.197811824 | 0.061201823 | 0.141900982 |
| 591,200 | 0.144047256 | 0.197587056 | 0.061627569 | 0.142004975 |
| 591,300 | 0.144095896 | 0.198018086 | 0.06158403 | 0.14212052 |
| 591,400 | 0.14410039 | 0.197255384 | 0.061888315 | 0.141647128 |
| 591,500 | 0.144232818 | 0.197441434 | 0.06250689 | 0.141920974 |
| 591,600 | 0.144061039 | 0.197952225 | 0.062368916 | 0.141719579 |
| 591,700 | 0.143991822 | 0.197688279 | 0.062537501 | 0.141365618 |
| 591,800 | 0.144226777 | 0.198549849 | 0.061824169 | 0.141484864 |
| 591,900 | 0.144204401 | 0.197948806 | 0.061949104 | 0.141565245 |
| 592,000 | 0.144069498 | 0.197726478 | 0.06233219 | 0.141536467 |
| 592,100 | 0.144099236 | 0.197773848 | 0.06114013 | 0.141232849 |
| 592,200 | 0.143995904 | 0.198056529 | 0.0616759 | 0.141013777 |
| 592,300 | 0.143977009 | 0.197429465 | 0.06155955 | 0.141568605 |
| 592,400 | 0.143877051 | 0.198492127 | 0.061290676 | 0.141350751 |
| 592,500 | 0.143973178 | 0.197269283 | 0.062166132 | 0.142050232 |
| 592,600 | 0.144004485 | 0.197051725 | 0.062421791 | 0.1418823 |
| 592,700 | 0.144042253 | 0.197761506 | 0.061598454 | 0.141354401 |
| 592,800 | 0.14383279 | 0.197299869 | 0.062288093 | 0.141390027 |
| 592,900 | 0.143869951 | 0.197621516 | 0.062289432 | 0.142129286 |
| 593,000 | 0.143715114 | 0.197697397 | 0.062075693 | 0.141312401 |
| 593,100 | 0.143938478 | 0.197325812 | 0.06202693 | 0.141286363 |
| 593,200 | 0.143834928 | 0.19755363 | 0.062366428 | 0.141464835 |
| 593,300 | 0.144003899 | 0.197882105 | 0.062189425 | 0.140865219 |
| 593,400 | 0.143922682 | 0.197288358 | 0.062286232 | 0.140742303 |
| 593,500 | 0.143888962 | 0.197979958 | 0.062543413 | 0.1408262 |
| 593,600 | 0.143925363 | 0.197903144 | 0.062526336 | 0.14092531 |
| 593,700 | 0.143867875 | 0.197150404 | 0.062005056 | 0.140859199 |
| 593,800 | 0.143758821 | 0.19740893 | 0.062290343 | 0.14130995 |
| 593,900 | 0.143764138 | 0.19727515 | 0.062227984 | 0.141573872 |
| 594,000 | 0.143761961 | 0.197760409 | 0.061961972 | 0.141044613 |
| 594,100 | 0.143713666 | 0.198257264 | 0.062734178 | 0.140907213 |
| 594,200 | 0.143694464 | 0.197795348 | 0.062568637 | 0.141544224 |
| 594,300 | 0.143713857 | 0.197684697 | 0.062684428 | 0.141802363 |
| 594,400 | 0.143562665 | 0.197595944 | 0.062218054 | 0.140931075 |
| 594,500 | 0.143665044 | 0.197627113 | 0.062499908 | 0.141396195 |
| 594,600 | 0.143808532 | 0.197886006 | 0.06187589 | 0.141355495 |
| 594,700 | 0.143703807 | 0.197756314 | 0.06203464 | 0.140694831 |
| 594,800 | 0.143710905 | 0.197841695 | 0.062157984 | 0.141160348 |
| 594,900 | 0.143599064 | 0.198336437 | 0.062306505 | 0.140692839 |
| 595,000 | 0.143517959 | 0.197836608 | 0.062364304 | 0.14051044 |
| 595,100 | 0.143510145 | 0.198558839 | 0.06245184 | 0.14071705 |
| 595,200 | 0.143531237 | 0.198424116 | 0.062241668 | 0.14111592 |

| | | | | |
|---|---|---|---|---|
| 595,300 | 0.143466295 | 0.197419332 | 0.062118336 | 0.141023002 |
| 595,400 | 0.14350372 | 0.196894924 | 0.062632839 | 0.141473903 |
| 595,500 | 0.143529188 | 0.19753584 | 0.062681685 | 0.14154527 |
| 595,600 | 0.143533696 | 0.197411089 | 0.062751704 | 0.141169192 |
| 595,700 | 0.143539779 | 0.196755376 | 0.062267364 | 0.141721023 |
| 595,800 | 0.143534156 | 0.197192558 | 0.062509496 | 0.141805964 |
| 595,900 | 0.143532514 | 0.197915235 | 0.062607589 | 0.141885906 |
| 596,000 | 0.143621207 | 0.19798265 | 0.062394707 | 0.14210553 |
| 596,100 | 0.14359809 | 0.197584024 | 0.062624584 | 0.14226846 |
| 596,200 | 0.143492567 | 0.19736953 | 0.06259807 | 0.141536173 |
| 596,300 | 0.143489159 | 0.197307761 | 0.062768462 | 0.142249535 |
| 596,400 | 0.143463414 | 0.196574484 | 0.061908242 | 0.142404121 |
| 596,500 | 0.143471186 | 0.197141372 | 0.063086409 | 0.142447989 |
| 596,600 | 0.143446479 | 0.196482764 | 0.062860289 | 0.142065084 |
| 596,700 | 0.143355047 | 0.197237202 | 0.062543005 | 0.142220769 |
| 596,800 | 0.143281585 | 0.196853342 | 0.062601387 | 0.141760122 |
| 596,900 | 0.143181346 | 0.1971426 | 0.063315534 | 0.14162696 |
| 597,000 | 0.143242727 | 0.197375637 | 0.062532771 | 0.142074361 |
| 597,100 | 0.143202974 | 0.196303912 | 0.062152963 | 0.142604833 |
| 597,200 | 0.143236881 | 0.197013376 | 0.06172275 | 0.141853551 |
| 597,300 | 0.143283983 | 0.19645465 | 0.062279193 | 0.141762935 |
| 597,400 | 0.14330823 | 0.19680055 | 0.061778461 | 0.141670345 |
| 597,500 | 0.143274633 | 0.197082978 | 0.062030733 | 0.141061063 |
| 597,600 | 0.143255296 | 0.197787708 | 0.062490705 | 0.140536454 |
| 597,700 | 0.143305358 | 0.197461385 | 0.062538508 | 0.140587155 |
| 597,800 | 0.143226155 | 0.196426245 | 0.062527491 | 0.141231428 |
| 597,900 | 0.143172661 | 0.195871248 | 0.062225041 | 0.141856333 |
| 598,000 | 0.143122682 | 0.196084349 | 0.061434422 | 0.140869336 |
| 598,100 | 0.143061778 | 0.195584156 | 0.061958998 | 0.140842852 |
| 598,200 | 0.143013234 | 0.196215736 | 0.061917547 | 0.140703337 |
| 598,300 | 0.142945707 | 0.196054666 | 0.061651444 | 0.140819696 |
| 598,400 | 0.143042061 | 0.196490626 | 0.060811822 | 0.140648885 |
| 598,500 | 0.142885145 | 0.195880875 | 0.061854582 | 0.140823971 |
| 598,600 | 0.142913346 | 0.195715026 | 0.061452327 | 0.140883381 |
| 598,700 | 0.142870519 | 0.196183936 | 0.061170129 | 0.140890583 |
| 598,800 | 0.142887846 | 0.196219168 | 0.06092164 | 0.141022837 |
| 598,900 | 0.142824518 | 0.196328504 | 0.061495005 | 0.140839 |
| 599,000 | 0.142763779 | 0.195969734 | 0.061057218 | 0.14033912 |
| 599,100 | 0.14273325 | 0.196350433 | 0.061101456 | 0.140416523 |
| 599,200 | 0.142680574 | 0.196490779 | 0.061627999 | 0.140803259 |
| 599,300 | 0.142715431 | 0.196472379 | 0.060889896 | 0.141001452 |
| 599,400 | 0.14269764 | 0.196003885 | 0.061011924 | 0.141255807 |
| 599,500 | 0.14261271 | 0.195661569 | 0.061041369 | 0.140196356 |
| 599,600 | 0.142759221 | 0.196645733 | 0.061229383 | 0.140464662 |
| 599,700 | 0.142721213 | 0.196357588 | 0.060687203 | 0.140810676 |
| 599,800 | 0.142756938 | 0.196787136 | 0.060804989 | 0.140216719 |
| 599,900 | 0.142819089 | 0.195973559 | 0.060745943 | 0.140275062 |
| 600,000 | 0.14268332 | 0.197101327 | 0.060494091 | 0.140145673 |
| 600,100 | 0.142691721 | 0.196852786 | 0.059770598 | 0.14042432 |
| 600,200 | 0.142541277 | 0.196300634 | 0.060140913 | 0.140224826 |
| 600,300 | 0.142623876 | 0.19658305 | 0.060411635 | 0.140440463 |
| 600,400 | 0.142701075 | 0.196591089 | 0.061090351 | 0.140382824 |
| 600,500 | 0.142729479 | 0.195894556 | 0.060478887 | 0.140239725 |
| 600,600 | 0.142648879 | 0.196634583 | 0.060732864 | 0.139953226 |
| 600,700 | 0.142667273 | 0.196016207 | 0.061367006 | 0.140269423 |
| 600,800 | 0.142756408 | 0.19672204 | 0.06009223 | 0.140843637 |
| 600,900 | 0.142714876 | 0.196202741 | 0.059907408 | 0.140192598 |
| 601,000 | 0.142806999 | 0.196304375 | 0.061165009 | 0.140753168 |
| 601,100 | 0.142766044 | 0.196302051 | 0.060685167 | 0.141634331 |
| 601,200 | 0.14275891 | 0.196195381 | 0.061025012 | 0.139862959 |
| 601,300 | 0.142698713 | 0.196074491 | 0.061377841 | 0.1406871 |
| 601,400 | 0.142628345 | 0.195984322 | 0.061598581 | 0.140831653 |
| 601,500 | 0.142598646 | 0.195837999 | 0.061382896 | 0.140482335 |
| 601,600 | 0.142626823 | 0.196087762 | 0.061078983 | 0.140476505 |
| 601,700 | 0.142627379 | 0.195952864 | 0.06109132 | 0.140608478 |
| 601,800 | 0.142668942 | 0.196312525 | 0.060622528 | 0.140448718 |
| 601,900 | 0.142508851 | 0.195278436 | 0.060107787 | 0.140101299 |

| | | | | |
|---|---|---|---|---|
| 602,000 | 0.142626923 | 0.195812257 | 0.060756079 | 0.1410081 |
| 602,100 | 0.142396026 | 0.195499746 | 0.060839017 | 0.140169861 |
| 602,200 | 0.142447169 | 0.195138383 | 0.061119396 | 0.140211945 |
| 602,300 | 0.142418585 | 0.194940775 | 0.061525465 | 0.140059713 |
| 602,400 | 0.142579684 | 0.194752962 | 0.061149364 | 0.140388004 |
| 602,500 | 0.142341213 | 0.194460695 | 0.060933642 | 0.13990949 |
| 602,600 | 0.142407579 | 0.193773428 | 0.061003483 | 0.140696361 |
| 602,700 | 0.14234805 | 0.194045136 | 0.060810296 | 0.140857319 |
| 602,800 | 0.142501584 | 0.193884484 | 0.060401419 | 0.141020744 |
| 602,900 | 0.142396262 | 0.193977865 | 0.06047474 | 0.140701319 |
| 603,000 | 0.142509475 | 0.194437769 | 0.061011451 | 0.140627007 |
| 603,100 | 0.1425452 | 0.193460618 | 0.060780589 | 0.14081655 |
| 603,200 | 0.142448015 | 0.19403609 | 0.060258261 | 0.140520125 |
| 603,300 | 0.142440112 | 0.194045511 | 0.060860892 | 0.140166077 |
| 603,400 | 0.142387554 | 0.194307774 | 0.060547529 | 0.140388752 |
| 603,500 | 0.142353215 | 0.194385787 | 0.061432549 | 0.140866929 |
| 603,600 | 0.142398458 | 0.194122514 | 0.061205539 | 0.140753918 |
| 603,700 | 0.142318378 | 0.194010414 | 0.061220233 | 0.140431315 |
| 603,800 | 0.142376494 | 0.194215725 | 0.061036888 | 0.140694202 |
| 603,900 | 0.142358826 | 0.19458417 | 0.060728517 | 0.140605656 |
| 604,000 | 0.142225089 | 0.194398555 | 0.061314081 | 0.140874894 |
| 604,100 | 0.142302268 | 0.194473926 | 0.061053178 | 0.140725721 |
| 604,200 | 0.14221475 | 0.193922711 | 0.061243943 | 0.140693537 |
| 604,300 | 0.142279459 | 0.195018696 | 0.060653541 | 0.141210542 |
| 604,400 | 0.142266605 | 0.194683183 | 0.061088631 | 0.141919953 |
| 604,500 | 0.142111176 | 0.193895617 | 0.061288778 | 0.140888012 |
| 604,600 | 0.142160179 | 0.194155143 | 0.06092574 | 0.140673208 |
| 604,700 | 0.142320167 | 0.194306029 | 0.060595332 | 0.140281817 |
| 604,800 | 0.142235507 | 0.194380991 | 0.0607666 | 0.140645442 |
| 604,900 | 0.142289193 | 0.194679154 | 0.061197881 | 0.140535729 |
| 605,000 | 0.14218587 | 0.194443363 | 0.061109677 | 0.140104006 |
| 605,100 | 0.1421266 | 0.194689843 | 0.060778616 | 0.14024201 |
| 605,200 | 0.142247808 | 0.194182502 | 0.060540106 | 0.140666121 |
| 605,300 | 0.14217722 | 0.194771084 | 0.061146522 | 0.140315973 |
| 605,400 | 0.142138034 | 0.194508679 | 0.06081967 | 0.140863535 |
| 605,500 | 0.142166867 | 0.193978566 | 0.06058889 | 0.140837754 |
| 605,600 | 0.142270068 | 0.194794637 | 0.060461842 | 0.140663918 |
| 605,700 | 0.142279001 | 0.195315953 | 0.060808408 | 0.140978672 |
| 605,800 | 0.142104039 | 0.194685407 | 0.061131899 | 0.140711138 |
| 605,900 | 0.14205284 | 0.194868943 | 0.061145521 | 0.140283848 |
| 606,000 | 0.142057489 | 0.195283977 | 0.061149088 | 0.140566824 |
| 606,100 | 0.142090504 | 0.194879352 | 0.060943466 | 0.140695541 |
| 606,200 | 0.14201998 | 0.194973426 | 0.060810259 | 0.14089503 |
| 606,300 | 0.142175307 | 0.194741117 | 0.06079692 | 0.140583091 |
| 606,400 | 0.142138286 | 0.194712733 | 0.061182596 | 0.140439529 |
| 606,500 | 0.142203893 | 0.195035818 | 0.06056895 | 0.140672852 |
| 606,600 | 0.14206809 | 0.19382922 | 0.060557725 | 0.140926627 |
| 606,700 | 0.142026872 | 0.194853461 | 0.061092567 | 0.14097772 |
| 606,800 | 0.141974854 | 0.194067068 | 0.060641368 | 0.140782611 |
| 606,900 | 0.142027865 | 0.194479124 | 0.061339623 | 0.140991457 |
| 607,000 | 0.1419341 | 0.194886026 | 0.060648644 | 0.14066176 |
| 607,100 | 0.141907372 | 0.194621028 | 0.060720212 | 0.141025973 |
| 607,200 | 0.141946341 | 0.194424996 | 0.060832421 | 0.140821241 |
| 607,300 | 0.141904223 | 0.194857013 | 0.061235792 | 0.141098075 |
| 607,400 | 0.141836081 | 0.194314961 | 0.061307335 | 0.141878655 |
| 607,500 | 0.141735009 | 0.19436737 | 0.061025614 | 0.141350334 |
| 607,600 | 0.141812612 | 0.194946871 | 0.061012644 | 0.141236009 |
| 607,700 | 0.141985207 | 0.195250282 | 0.060669915 | 0.142018216 |
| 607,800 | 0.141882247 | 0.194892457 | 0.060082176 | 0.141824129 |
| 607,900 | 0.141944824 | 0.195153488 | 0.06098591 | 0.141455655 |
| 608,000 | 0.141674553 | 0.193931097 | 0.061046302 | 0.14154491 |
| 608,100 | 0.141912131 | 0.1946943 | 0.061077299 | 0.142010237 |
| 608,200 | 0.141862355 | 0.194518543 | 0.060666804 | 0.141696329 |
| 608,300 | 0.141816217 | 0.194577559 | 0.060872004 | 0.141978627 |
| 608,400 | 0.141731589 | 0.194225432 | 0.059925396 | 0.141604589 |
| 608,500 | 0.141716889 | 0.193988961 | 0.06058616 | 0.142227892 |
| 608,600 | 0.141751528 | 0.194144786 | 0.060821756 | 0.141668229 |

| | | | | |
|---|---|---|---|---|
| 608,700 | 0.141701398 | 0.193739699 | 0.060633997 | 0.141901815 |
| 608,800 | 0.141702486 | 0.194169731 | 0.060155709 | 0.141913965 |
| 608,900 | 0.141671463 | 0.194399994 | 0.059653457 | 0.141633026 |
| 609,000 | 0.141663003 | 0.193435764 | 0.059873358 | 0.141537505 |
| 609,100 | 0.141783117 | 0.194173409 | 0.05980078 | 0.141249104 |
| 609,200 | 0.141786913 | 0.194002803 | 0.060092626 | 0.141535536 |
| 609,300 | 0.14169658 | 0.194378011 | 0.060235848 | 0.141088473 |
| 609,400 | 0.141704947 | 0.194507062 | 0.060200391 | 0.141271972 |
| 609,500 | 0.14165152 | 0.194609914 | 0.059894648 | 0.141807072 |
| 609,600 | 0.141616578 | 0.194547422 | 0.059908183 | 0.141783932 |
| 609,700 | 0.141636359 | 0.194646912 | 0.06040494 | 0.14215131 |
| 609,800 | 0.141702826 | 0.193839891 | 0.060126444 | 0.141480467 |
| 609,900 | 0.141692 | 0.194562987 | 0.059852308 | 0.141394267 |
| 610,000 | 0.141628726 | 0.193891709 | 0.060021822 | 0.141906277 |
| 610,100 | 0.141505409 | 0.192767236 | 0.060168267 | 0.141855675 |
| 610,200 | 0.141445633 | 0.193233959 | 0.059624459 | 0.141267872 |
| 610,300 | 0.141510017 | 0.193383135 | 0.059601068 | 0.141651967 |
| 610,400 | 0.141392473 | 0.193189544 | 0.060135606 | 0.141575679 |
| 610,500 | 0.141360417 | 0.192755313 | 0.06014778 | 0.141543473 |
| 610,600 | 0.141399933 | 0.193492837 | 0.05967317 | 0.141430044 |
| 610,700 | 0.141323541 | 0.193601349 | 0.059873523 | 0.141458045 |
| 610,800 | 0.141259465 | 0.193072079 | 0.060043114 | 0.141440446 |
| 610,900 | 0.141282502 | 0.193928788 | 0.060230543 | 0.141430099 |
| 611,000 | 0.141289568 | 0.193931741 | 0.060321136 | 0.141244587 |
| 611,100 | 0.141312874 | 0.194816037 | 0.060896117 | 0.141155605 |
| 611,200 | 0.141330461 | 0.195135791 | 0.060511174 | 0.141221076 |
| 611,300 | 0.141080932 | 0.19402476 | 0.06010387 | 0.141052408 |
| 611,400 | 0.141116824 | 0.194331175 | 0.060444433 | 0.141477015 |
| 611,500 | 0.141083355 | 0.193720556 | 0.05997159 | 0.141222499 |
| 611,600 | 0.141106476 | 0.194076256 | 0.059764396 | 0.141604259 |
| 611,700 | 0.141222359 | 0.194334802 | 0.0595876 | 0.141875907 |
| 611,800 | 0.141304179 | 0.194390775 | 0.059959934 | 0.141700192 |
| 611,900 | 0.141111801 | 0.193895908 | 0.059963812 | 0.141518196 |
| 612,000 | 0.141255804 | 0.193568653 | 0.06059675 | 0.141225697 |
| 612,100 | 0.141225703 | 0.193678901 | 0.060054003 | 0.141535801 |
| 612,200 | 0.141219165 | 0.193233841 | 0.060270186 | 0.141822338 |
| 612,300 | 0.141126846 | 0.193066065 | 0.060123223 | 0.141688544 |
| 612,400 | 0.141139576 | 0.192720582 | 0.060793426 | 0.141496362 |
| 612,500 | 0.141159459 | 0.19354121 | 0.061197527 | 0.141823545 |
| 612,600 | 0.141145802 | 0.193084119 | 0.060979129 | 0.141900306 |
| 612,700 | 0.141052927 | 0.19321246 | 0.060231861 | 0.141710347 |
| 612,800 | 0.141228197 | 0.193770451 | 0.060657176 | 0.1422959 |
| 612,900 | 0.141176834 | 0.19425824 | 0.060952035 | 0.141820791 |
| 613,000 | 0.141134957 | 0.193155824 | 0.060978589 | 0.1423909 |
| 613,100 | 0.141221665 | 0.19328093 | 0.060600707 | 0.142035633 |
| 613,200 | 0.141077326 | 0.193554888 | 0.06073226 | 0.141434747 |
| 613,300 | 0.141040274 | 0.193664434 | 0.060810702 | 0.142025535 |
| 613,400 | 0.141088352 | 0.192710461 | 0.060547436 | 0.141088231 |
| 613,500 | 0.141067752 | 0.192426897 | 0.059929129 | 0.1414367 |
| 613,600 | 0.141083484 | 0.192847931 | 0.059745681 | 0.142100485 |
| 613,700 | 0.141011519 | 0.192854312 | 0.059667439 | 0.141836639 |
| 613,800 | 0.141100199 | 0.193186624 | 0.060016275 | 0.141395878 |
| 613,900 | 0.140986517 | 0.192850141 | 0.060134076 | 0.142191584 |
| 614,000 | 0.141053357 | 0.192907172 | 0.059735175 | 0.141289221 |
| 614,100 | 0.140968264 | 0.193010839 | 0.060599704 | 0.141638782 |
| 614,200 | 0.141030956 | 0.193400911 | 0.061035832 | 0.141302958 |
| 614,300 | 0.141066243 | 0.193105306 | 0.06061398 | 0.141835043 |
| 614,400 | 0.141024606 | 0.193204463 | 0.060517406 | 0.14176071 |
| 614,500 | 0.141098256 | 0.193798483 | 0.060930799 | 0.141528361 |
| 614,600 | 0.141010178 | 0.193527231 | 0.061042567 | 0.141449346 |
| 614,700 | 0.140993664 | 0.193753733 | 0.061155543 | 0.141551048 |
| 614,800 | 0.140885514 | 0.193460754 | 0.06098773 | 0.141630257 |
| 614,900 | 0.140930447 | 0.193609751 | 0.060698145 | 0.141685002 |
| 615,000 | 0.140974882 | 0.192969984 | 0.059996484 | 0.141654005 |
| 615,100 | 0.14091602 | 0.193469248 | 0.060486558 | 0.141687515 |
| 615,200 | 0.14091898 | 0.193397655 | 0.060516654 | 0.141185239 |
| 615,300 | 0.140781913 | 0.193084796 | 0.06072341 | 0.1413725 |

| | | | | |
|---|---|---|---|---|
| 615,400 | 0.140932533 | 0.193494693 | 0.060380313 | 0.141235534 |
| 615,500 | 0.140848375 | 0.192667935 | 0.060712458 | 0.141557274 |
| 615,600 | 0.140842829 | 0.193397958 | 0.061029048 | 0.140662316 |
| 615,700 | 0.140883801 | 0.193398642 | 0.061117875 | 0.140998176 |
| 615,800 | 0.140830795 | 0.193085702 | 0.060998694 | 0.141022092 |
| 615,900 | 0.140912249 | 0.192984704 | 0.060935377 | 0.140722608 |
| 616,000 | 0.140972614 | 0.192940598 | 0.061250813 | 0.141139341 |
| 616,100 | 0.140997156 | 0.193105409 | 0.061104682 | 0.141154855 |
| 616,200 | 0.140979846 | 0.193157592 | 0.060469569 | 0.141570956 |
| 616,300 | 0.140924807 | 0.192912863 | 0.061251173 | 0.14150967 |
| 616,400 | 0.140958577 | 0.193066387 | 0.061442777 | 0.141320266 |
| 616,500 | 0.140961018 | 0.192574527 | 0.06209444 | 0.141414388 |
| 616,600 | 0.141056361 | 0.192681881 | 0.061066427 | 0.141357611 |
| 616,700 | 0.140910774 | 0.192427051 | 0.061000257 | 0.141668372 |
| 616,800 | 0.140987656 | 0.192664524 | 0.060724952 | 0.141439953 |
| 616,900 | 0.140899556 | 0.192570079 | 0.060819202 | 0.141876345 |
| 617,000 | 0.140998206 | 0.192796296 | 0.061175328 | 0.141524208 |
| 617,100 | 0.140981322 | 0.192648455 | 0.061705233 | 0.142040362 |
| 617,200 | 0.14087805 | 0.19223728 | 0.061182943 | 0.14165258 |
| 617,300 | 0.140912016 | 0.19187136 | 0.061451658 | 0.141391458 |
| 617,400 | 0.140836151 | 0.192715829 | 0.0615085 | 0.14117636 |
| 617,500 | 0.140889378 | 0.192975275 | 0.06131519 | 0.141771261 |
| 617,600 | 0.140871978 | 0.192700137 | 0.061376018 | 0.140953255 |
| 617,700 | 0.140844927 | 0.192623994 | 0.061743816 | 0.140770206 |
| 617,800 | 0.140938343 | 0.193080709 | 0.061009609 | 0.140880189 |
| 617,900 | 0.140882501 | 0.1931038 | 0.061375027 | 0.141541855 |
| 618,000 | 0.140765756 | 0.192272467 | 0.061335187 | 0.141523235 |
| 618,100 | 0.140788081 | 0.192114271 | 0.06156936 | 0.141686701 |
| 618,200 | 0.140761072 | 0.192541825 | 0.060898354 | 0.141430738 |
| 618,300 | 0.14073713 | 0.192377744 | 0.061424691 | 0.141555874 |
| 618,400 | 0.14079031 | 0.192021412 | 0.061243575 | 0.141887066 |
| 618,500 | 0.140767268 | 0.191907434 | 0.061414565 | 0.141526132 |
| 618,600 | 0.140824794 | 0.19271124 | 0.061117137 | 0.142256742 |
| 618,700 | 0.140732139 | 0.192140133 | 0.060951465 | 0.141747419 |
| 618,800 | 0.1407051 | 0.192071757 | 0.061251358 | 0.142305838 |
| 618,900 | 0.140475299 | 0.191724533 | 0.060835197 | 0.141986262 |
| 619,000 | 0.140619249 | 0.192471565 | 0.061603032 | 0.142095744 |
| 619,100 | 0.140657861 | 0.192221859 | 0.06160462 | 0.141932166 |
| 619,200 | 0.140775566 | 0.192620509 | 0.060942771 | 0.141984122 |
| 619,300 | 0.140735728 | 0.192448971 | 0.061011698 | 0.142122472 |
| 619,400 | 0.140575189 | 0.192060208 | 0.061129296 | 0.141769409 |
| 619,500 | 0.140599767 | 0.192460489 | 0.060905142 | 0.142545998 |
| 619,600 | 0.140631569 | 0.19187623 | 0.06150352 | 0.141311614 |
| 619,700 | 0.140647825 | 0.192135479 | 0.060780234 | 0.141769288 |
| 619,800 | 0.1406297 | 0.192101799 | 0.060997011 | 0.141203159 |
| 619,900 | 0.140525056 | 0.192625823 | 0.061086799 | 0.140985609 |
| 620,000 | 0.14063728 | 0.192451358 | 0.060955532 | 0.141294782 |
| 620,100 | 0.140648795 | 0.192817951 | 0.060984026 | 0.141794349 |
| 620,200 | 0.140626734 | 0.193048859 | 0.060933115 | 0.14145426 |
| 620,300 | 0.140540583 | 0.192868093 | 0.060965154 | 0.142052183 |
| 620,400 | 0.140497582 | 0.192732339 | 0.060819751 | 0.141576099 |
| 620,500 | 0.14053534 | 0.191627157 | 0.060657726 | 0.142393608 |
| 620,600 | 0.140494969 | 0.192104247 | 0.060926943 | 0.141327228 |
| 620,700 | 0.140381611 | 0.191695347 | 0.060724557 | 0.141814645 |
| 620,800 | 0.140467827 | 0.191971518 | 0.061424478 | 0.141491915 |
| 620,900 | 0.140415826 | 0.192443262 | 0.06073177 | 0.141070256 |
| 621,000 | 0.140335492 | 0.192488211 | 0.061191363 | 0.141320231 |
| 621,100 | 0.140449214 | 0.192355854 | 0.061505742 | 0.140999542 |
| 621,200 | 0.140494893 | 0.192545912 | 0.060569423 | 0.14154123 |
| 621,300 | 0.140577369 | 0.193481785 | 0.060574667 | 0.141486674 |
| 621,400 | 0.140480915 | 0.193305618 | 0.060406543 | 0.141413035 |
| 621,500 | 0.140510528 | 0.193262015 | 0.060487153 | 0.141703968 |
| 621,600 | 0.14043166 | 0.192766462 | 0.060771328 | 0.141535795 |
| 621,700 | 0.140544582 | 0.192695044 | 0.060799051 | 0.141247441 |
| 621,800 | 0.14049222 | 0.192574849 | 0.060605736 | 0.141431842 |
| 621,900 | 0.140369427 | 0.192572069 | 0.060668665 | 0.141272943 |
| 622,000 | 0.140389945 | 0.192478009 | 0.060653701 | 0.141254172 |

| | | | | |
|---|---|---|---|---|
| 622,100 | 0.140318216 | 0.19281647 | 0.060775143 | 0.140516754 |
| 622,200 | 0.140214282 | 0.192355453 | 0.061440889 | 0.141310696 |
| 622,300 | 0.140214114 | 0.192830833 | 0.061724713 | 0.14103532 |
| 622,400 | 0.140284913 | 0.192964882 | 0.060961014 | 0.141174705 |
| 622,500 | 0.140222072 | 0.192973066 | 0.060810416 | 0.141222392 |
| 622,600 | 0.140274385 | 0.193414987 | 0.061243725 | 0.141066176 |
| 622,700 | 0.140102673 | 0.19279215 | 0.060490291 | 0.140688775 |
| 622,800 | 0.140242693 | 0.193179177 | 0.061108747 | 0.141131478 |
| 622,900 | 0.140187354 | 0.192982236 | 0.061057973 | 0.141008162 |
| 623,000 | 0.140102371 | 0.192816434 | 0.060100979 | 0.140725233 |
| 623,100 | 0.140088532 | 0.193210726 | 0.060995465 | 0.140682253 |
| 623,200 | 0.140016954 | 0.193378557 | 0.060823086 | 0.140874752 |
| 623,300 | 0.140047753 | 0.192591696 | 0.060742092 | 0.141074924 |
| 623,400 | 0.13996803 | 0.193485451 | 0.061155617 | 0.140950406 |
| 623,500 | 0.139851696 | 0.192549798 | 0.060844834 | 0.140558647 |
| 623,600 | 0.14006778 | 0.193011042 | 0.061018411 | 0.141037206 |
| 623,700 | 0.139892952 | 0.193163216 | 0.061149921 | 0.140452564 |
| 623,800 | 0.140011548 | 0.192748062 | 0.060680421 | 0.140366146 |
| 623,900 | 0.139868351 | 0.192527811 | 0.061223525 | 0.139775219 |
| 624,000 | 0.139965463 | 0.192949152 | 0.061024682 | 0.140100095 |
| 624,100 | 0.140018826 | 0.192957639 | 0.061214073 | 0.140734834 |
| 624,200 | 0.140022758 | 0.193025635 | 0.060999873 | 0.141280765 |
| 624,300 | 0.139892858 | 0.192531356 | 0.061536279 | 0.140386248 |
| 624,400 | 0.139915523 | 0.193371137 | 0.06169058 | 0.140812086 |
| 624,500 | 0.140056903 | 0.192893048 | 0.061559006 | 0.140727386 |
| 624,600 | 0.14004904 | 0.193707854 | 0.062050918 | 0.140590144 |
| 624,700 | 0.140030373 | 0.192967876 | 0.061981284 | 0.140751901 |
| 624,800 | 0.140034634 | 0.193324659 | 0.061807114 | 0.140448628 |
| 624,900 | 0.140115319 | 0.193625303 | 0.062300385 | 0.140739117 |
| 625,000 | 0.140015449 | 0.192836929 | 0.061405289 | 0.141271121 |
| 625,100 | 0.140176638 | 0.193513011 | 0.062349254 | 0.141185543 |
| 625,200 | 0.140011319 | 0.193384328 | 0.062173989 | 0.141023453 |
| 625,300 | 0.139976321 | 0.19256247 | 0.062027728 | 0.140786601 |
| 625,400 | 0.139878136 | 0.192796525 | 0.061512911 | 0.141147463 |
| 625,500 | 0.140031978 | 0.193301119 | 0.0612937 | 0.140915101 |
| 625,600 | 0.139872176 | 0.192927641 | 0.06083509 | 0.141346412 |
| 625,700 | 0.139898881 | 0.192258691 | 0.061599962 | 0.141410368 |
| 625,800 | 0.139810052 | 0.192415981 | 0.061092596 | 0.141139605 |
| 625,900 | 0.139883141 | 0.191812111 | 0.061329311 | 0.14139359 |
| 626,000 | 0.139888917 | 0.19262745 | 0.06124855 | 0.140933133 |
| 626,100 | 0.139913858 | 0.192386431 | 0.061556372 | 0.141004824 |
| 626,200 | 0.139874891 | 0.192262823 | 0.061395425 | 0.141256442 |
| 626,300 | 0.139835212 | 0.192396212 | 0.061541379 | 0.141090826 |
| 626,400 | 0.139783213 | 0.192306964 | 0.061671961 | 0.14137654 |
| 626,500 | 0.139757316 | 0.191749463 | 0.061731343 | 0.141216023 |
| 626,600 | 0.139909643 | 0.192237689 | 0.062157485 | 0.141645288 |
| 626,700 | 0.139772376 | 0.192576128 | 0.061724804 | 0.141276633 |
| 626,800 | 0.13976414 | 0.192509132 | 0.061255623 | 0.141132869 |
| 626,900 | 0.139813809 | 0.192396353 | 0.060945508 | 0.141640531 |
| 627,000 | 0.139824086 | 0.192359394 | 0.061074498 | 0.141851821 |
| 627,100 | 0.139728422 | 0.192093735 | 0.061564295 | 0.141118837 |
| 627,200 | 0.139783095 | 0.192048499 | 0.06110566 | 0.141377662 |
| 627,300 | 0.139785746 | 0.191697865 | 0.0610797 | 0.141349235 |
| 627,400 | 0.139707057 | 0.191113475 | 0.061264204 | 0.141089215 |
| 627,500 | 0.139725342 | 0.191272442 | 0.061411526 | 0.141347578 |
| 627,600 | 0.139769374 | 0.191530085 | 0.061522381 | 0.141785864 |
| 627,700 | 0.139776416 | 0.191240978 | 0.06038683 | 0.141455234 |
| 627,800 | 0.139756481 | 0.191272253 | 0.060435274 | 0.140791507 |
| 627,900 | 0.139605256 | 0.191313407 | 0.060574108 | 0.140781307 |
| 628,000 | 0.139682482 | 0.191630138 | 0.060658774 | 0.141030473 |
| 628,100 | 0.139713504 | 0.191351065 | 0.060982488 | 0.140818553 |
| 628,200 | 0.139697614 | 0.191717652 | 0.060749097 | 0.140976381 |
| 628,300 | 0.139636746 | 0.191747347 | 0.061223683 | 0.140530129 |
| 628,400 | 0.139640012 | 0.192420274 | 0.061230165 | 0.140413293 |
| 628,500 | 0.139645268 | 0.191678054 | 0.061454483 | 0.140737844 |
| 628,600 | 0.139711163 | 0.191405351 | 0.061479609 | 0.140500039 |
| 628,700 | 0.139562308 | 0.191504624 | 0.061720178 | 0.140649841 |

| | | | | |
|---|---|---|---|---|
| 628,800 | 0.139578555 | 0.191436109 | 0.061920386 | 0.14098267 |
| 628,900 | 0.139684278 | 0.191810918 | 0.061239512 | 0.14146743 |
| 629,000 | 0.13968354 | 0.191477111 | 0.061499343 | 0.14050892 |
| 629,100 | 0.139628877 | 0.191476834 | 0.061418747 | 0.140373196 |
| 629,200 | 0.139613566 | 0.192005886 | 0.06084178 | 0.140386729 |
| 629,300 | 0.139559714 | 0.191828465 | 0.060803816 | 0.140767153 |
| 629,400 | 0.139491211 | 0.191764293 | 0.060939753 | 0.141081648 |
| 629,500 | 0.13951793 | 0.192220311 | 0.060843594 | 0.140689053 |
| 629,600 | 0.139618511 | 0.192011288 | 0.061087336 | 0.140851826 |
| 629,700 | 0.139465885 | 0.192001702 | 0.060658865 | 0.140593364 |
| 629,800 | 0.139463672 | 0.191964106 | 0.060859793 | 0.140151551 |
| 629,900 | 0.13953974 | 0.192164204 | 0.0609107 | 0.140505216 |
| 630,000 | 0.13946965 | 0.191687029 | 0.061235365 | 0.140960961 |
| 630,100 | 0.139578197 | 0.192657947 | 0.060950198 | 0.139957838 |
| 630,200 | 0.139468943 | 0.192993324 | 0.061353267 | 0.139941449 |
| 630,300 | 0.139458861 | 0.192254513 | 0.06182465 | 0.140337613 |
| 630,400 | 0.139459922 | 0.192359619 | 0.061049631 | 0.140816562 |
| 630,500 | 0.139462004 | 0.191941919 | 0.061864216 | 0.140736368 |
| 630,600 | 0.139563186 | 0.191949781 | 0.062198805 | 0.140894826 |
| 630,700 | 0.139492387 | 0.191809466 | 0.062125016 | 0.140317813 |
| 630,800 | 0.139416384 | 0.19183837 | 0.061926827 | 0.140555647 |
| 630,900 | 0.139523661 | 0.19136128 | 0.062044635 | 0.140435222 |
| 631,000 | 0.139401569 | 0.191488894 | 0.061697966 | 0.140621996 |
| 631,100 | 0.139373273 | 0.191530265 | 0.061809704 | 0.140359381 |
| 631,200 | 0.139378195 | 0.191898701 | 0.061995004 | 0.141318892 |
| 631,300 | 0.139314709 | 0.191668563 | 0.062014645 | 0.141555502 |
| 631,400 | 0.139429704 | 0.191454722 | 0.061782867 | 0.1411432 |
| 631,500 | 0.139304058 | 0.191428731 | 0.062153888 | 0.140888132 |
| 631,600 | 0.139273606 | 0.190355208 | 0.061704347 | 0.140284778 |
| 631,700 | 0.139309695 | 0.190997978 | 0.061143105 | 0.140435061 |
| 631,800 | 0.139352114 | 0.191551664 | 0.061517959 | 0.139896796 |
| 631,900 | 0.139377294 | 0.190420186 | 0.061535193 | 0.140183662 |
| 632,000 | 0.139225195 | 0.190763137 | 0.061763498 | 0.140383878 |
| 632,100 | 0.139187638 | 0.190631966 | 0.061640596 | 0.140788733 |
| 632,200 | 0.139330438 | 0.191470339 | 0.061321191 | 0.139969377 |
| 632,300 | 0.13928294 | 0.190844319 | 0.061261437 | 0.139855548 |
| 632,400 | 0.139254642 | 0.191019427 | 0.061011774 | 0.140142627 |
| 632,500 | 0.139079596 | 0.19103397 | 0.061214828 | 0.140282585 |
| 632,600 | 0.139305252 | 0.191052376 | 0.061631639 | 0.140002258 |
| 632,700 | 0.139223267 | 0.191000394 | 0.062351486 | 0.140002947 |
| 632,800 | 0.139136943 | 0.191268032 | 0.062191288 | 0.139682869 |
| 632,900 | 0.139172091 | 0.190905672 | 0.061884466 | 0.139895148 |
| 633,000 | 0.139072066 | 0.191214812 | 0.062137625 | 0.139734001 |
| 633,100 | 0.139116048 | 0.191204793 | 0.06224197 | 0.139633497 |
| 633,200 | 0.138970651 | 0.190494481 | 0.061793504 | 0.139697268 |
| 633,300 | 0.1390826 | 0.191248472 | 0.061907696 | 0.139819932 |
| 633,400 | 0.138942404 | 0.191139594 | 0.061845596 | 0.139759441 |
| 633,500 | 0.139036277 | 0.191121374 | 0.061546201 | 0.139999962 |
| 633,600 | 0.139057687 | 0.191208856 | 0.06139535 | 0.13920675 |
| 633,700 | 0.138977198 | 0.190614323 | 0.061578621 | 0.1394534 |
| 633,800 | 0.139019498 | 0.190483967 | 0.061727443 | 0.139756388 |
| 633,900 | 0.139082516 | 0.190615783 | 0.06201369 | 0.139993513 |
| 634,000 | 0.139050563 | 0.19048 | 0.06211531 | 0.139574973 |
| 634,100 | 0.139069646 | 0.190153126 | 0.061512628 | 0.139607264 |
| 634,200 | 0.138940285 | 0.189314271 | 0.061389787 | 0.139360365 |
| 634,300 | 0.139018006 | 0.190338484 | 0.061346258 | 0.139071751 |
| 634,400 | 0.139041113 | 0.190327811 | 0.062103287 | 0.139516381 |
| 634,500 | 0.138995047 | 0.190009435 | 0.061836454 | 0.14003242 |
| 634,600 | 0.138943604 | 0.189619573 | 0.061094906 | 0.139407846 |
| 634,700 | 0.138887868 | 0.189782599 | 0.061271406 | 0.1403075 |
| 634,800 | 0.138954637 | 0.189637919 | 0.06120538 | 0.139741417 |
| 634,900 | 0.138921812 | 0.189850918 | 0.060840414 | 0.139758656 |
| 635,000 | 0.138923639 | 0.189735235 | 0.060801214 | 0.139517759 |
| 635,100 | 0.138801843 | 0.188787355 | 0.060551829 | 0.139659367 |
| 635,200 | 0.138885218 | 0.189189425 | 0.060583907 | 0.139582747 |
| 635,300 | 0.138855473 | 0.189116547 | 0.060851038 | 0.138724276 |
| 635,400 | 0.138879934 | 0.188790581 | 0.061069779 | 0.139639217 |

| | | | | |
|---|---|---|---|---|
| 635,500 | 0.138830032 | 0.18900072 | 0.060813283 | 0.139643537 |
| 635,600 | 0.138760381 | 0.189250148 | 0.061823373 | 0.139305562 |
| 635,700 | 0.138795054 | 0.188717802 | 0.061527943 | 0.139810613 |
| 635,800 | 0.138847781 | 0.189839225 | 0.061750012 | 0.139652691 |
| 635,900 | 0.13888378 | 0.189986058 | 0.061386267 | 0.139944615 |
| 636,000 | 0.138735419 | 0.189693551 | 0.060742308 | 0.139684279 |
| 636,100 | 0.138770406 | 0.189187248 | 0.061569564 | 0.139670178 |
| 636,200 | 0.138750264 | 0.189669924 | 0.060989762 | 0.139446653 |
| 636,300 | 0.138702809 | 0.189272988 | 0.061346577 | 0.13956022 |
| 636,400 | 0.138854488 | 0.189600847 | 0.060951408 | 0.139803833 |
| 636,500 | 0.13875974 | 0.189834397 | 0.061343355 | 0.139767363 |
| 636,600 | 0.138775036 | 0.189765834 | 0.060843318 | 0.139573382 |
| 636,700 | 0.138776569 | 0.189181055 | 0.060821557 | 0.139655492 |
| 636,800 | 0.138649367 | 0.18860886 | 0.060023288 | 0.139873399 |
| 636,900 | 0.13874255 | 0.189474417 | 0.060700319 | 0.139470175 |
| 637,000 | 0.138746874 | 0.188612147 | 0.0602555 | 0.140039131 |
| 637,100 | 0.138734184 | 0.189218519 | 0.060729191 | 0.13898088 |
| 637,200 | 0.138708763 | 0.189437735 | 0.060855494 | 0.139727382 |
| 637,300 | 0.13870806 | 0.188560689 | 0.060686167 | 0.139694307 |
| 637,400 | 0.138706745 | 0.188556135 | 0.060645864 | 0.140103002 |
| 637,500 | 0.138688645 | 0.189380134 | 0.060184685 | 0.140156157 |
| 637,600 | 0.138706888 | 0.189512209 | 0.060316321 | 0.139825311 |
| 637,700 | 0.138790814 | 0.189448263 | 0.060463753 | 0.13969001 |
| 637,800 | 0.138780699 | 0.189427799 | 0.060599521 | 0.140055436 |
| 637,900 | 0.138719159 | 0.189409762 | 0.060549025 | 0.140050745 |
| 638,000 | 0.138811039 | 0.190015289 | 0.060410129 | 0.139909728 |
| 638,100 | 0.138778662 | 0.189682764 | 0.060832566 | 0.139390829 |
| 638,200 | 0.138761489 | 0.190070993 | 0.060840696 | 0.139551416 |
| 638,300 | 0.138648943 | 0.18930722 | 0.060612333 | 0.13982173 |
| 638,400 | 0.138729755 | 0.189298455 | 0.060284224 | 0.139836292 |
| 638,500 | 0.138468455 | 0.188971638 | 0.060242242 | 0.13981846 |
| 638,600 | 0.138528614 | 0.189335412 | 0.060581602 | 0.139872975 |
| 638,700 | 0.138578646 | 0.189490395 | 0.060624702 | 0.139646061 |
| 638,800 | 0.138532134 | 0.189437916 | 0.059792435 | 0.139669726 |
| 638,900 | 0.138459988 | 0.189453943 | 0.060592327 | 0.139398819 |
| 639,000 | 0.138469631 | 0.189224384 | 0.060693363 | 0.139965041 |
| 639,100 | 0.138470721 | 0.189575801 | 0.060383373 | 0.139702118 |
| 639,200 | 0.138520996 | 0.189975591 | 0.060018837 | 0.139305547 |
| 639,300 | 0.138495933 | 0.18957192 | 0.059939387 | 0.139508437 |
| 639,400 | 0.138510029 | 0.189933791 | 0.059612031 | 0.13940046 |
| 639,500 | 0.138574503 | 0.189916063 | 0.06018472 | 0.140326673 |
| 639,600 | 0.138484733 | 0.18992875 | 0.060009603 | 0.140311904 |
| 639,700 | 0.138488563 | 0.190485441 | 0.059632473 | 0.139670739 |
| 639,800 | 0.138384777 | 0.189671714 | 0.059777856 | 0.139774422 |
| 639,900 | 0.138450529 | 0.189774432 | 0.059492068 | 0.139519864 |
| 640,000 | 0.138497492 | 0.190057739 | 0.059749599 | 0.139893428 |
| 640,100 | 0.138514685 | 0.190194511 | 0.059405822 | 0.139545903 |
| 640,200 | 0.138431345 | 0.190197143 | 0.059727422 | 0.139513682 |
| 640,300 | 0.138458223 | 0.189496545 | 0.059384088 | 0.139360523 |
| 640,400 | 0.138380899 | 0.1897596 | 0.059616626 | 0.139663487 |
| 640,500 | 0.138409255 | 0.189054684 | 0.05976468 | 0.139366618 |
| 640,600 | 0.138365765 | 0.188352225 | 0.059762382 | 0.139601587 |
| 640,700 | 0.138428141 | 0.189516251 | 0.060266508 | 0.139737906 |
| 640,800 | 0.138448084 | 0.189303083 | 0.059391705 | 0.139273954 |
| 640,900 | 0.138266578 | 0.189340287 | 0.059796626 | 0.139260175 |
| 641,000 | 0.1383982 | 0.189289788 | 0.05975955 | 0.139596203 |
| 641,100 | 0.138336531 | 0.189594599 | 0.059784711 | 0.139430067 |
| 641,200 | 0.13822765 | 0.189247322 | 0.059608938 | 0.139382998 |
| 641,300 | 0.138250589 | 0.189398048 | 0.05986933 | 0.139313592 |
| 641,400 | 0.138292474 | 0.189846637 | 0.059084547 | 0.139195472 |
| 641,500 | 0.138164963 | 0.189011384 | 0.058913103 | 0.139321538 |
| 641,600 | 0.13822409 | 0.188644906 | 0.059298317 | 0.139051564 |
| 641,700 | 0.138219913 | 0.188720854 | 0.05949051 | 0.139019002 |
| 641,800 | 0.138401583 | 0.188884286 | 0.059442941 | 0.139280488 |
| 641,900 | 0.138398042 | 0.188984034 | 0.05927301 | 0.139144835 |
| 642,000 | 0.138352817 | 0.188370253 | 0.059691556 | 0.13906058 |
| 642,100 | 0.138308283 | 0.189057213 | 0.05957385 | 0.138538293 |

| | | | | |
|---|---|---|---|---|
| 642,200 | 0.138247808 | 0.188893307 | 0.059821579 | 0.139035182 |
| 642,300 | 0.138263086 | 0.189365878 | 0.060114756 | 0.139037754 |
| 642,400 | 0.138289323 | 0.189879926 | 0.060186767 | 0.139247246 |
| 642,500 | 0.138205144 | 0.189629416 | 0.059569411 | 0.138700297 |
| 642,600 | 0.13829991 | 0.190091981 | 0.059813507 | 0.138980035 |
| 642,700 | 0.138348088 | 0.189488857 | 0.059780353 | 0.139269684 |
| 642,800 | 0.138372745 | 0.189025459 | 0.059570899 | 0.138602843 |
| 642,900 | 0.138523002 | 0.189537849 | 0.060104353 | 0.138885934 |
| 643,000 | 0.138291421 | 0.189546788 | 0.059820324 | 0.13908898 |
| 643,100 | 0.138296152 | 0.188911862 | 0.05995873 | 0.139040845 |
| 643,200 | 0.138141081 | 0.189031467 | 0.059920833 | 0.138713614 |
| 643,300 | 0.138213883 | 0.189022963 | 0.059882991 | 0.138749233 |
| 643,400 | 0.138270103 | 0.189299693 | 0.060104344 | 0.138927374 |
| 643,500 | 0.138254379 | 0.188573173 | 0.059502448 | 0.139053625 |
| 643,600 | 0.138287356 | 0.189604109 | 0.059535421 | 0.138608156 |
| 643,700 | 0.138175838 | 0.189052728 | 0.060024341 | 0.139052945 |
| 643,800 | 0.138231468 | 0.189278459 | 0.059923775 | 0.139379194 |
| 643,900 | 0.138239488 | 0.189120379 | 0.059822132 | 0.138987699 |
| 644,000 | 0.138182379 | 0.189180103 | 0.059286517 | 0.139446633 |
| 644,100 | 0.138214517 | 0.189006166 | 0.059537075 | 0.139707619 |
| 644,200 | 0.13821197 | 0.188964795 | 0.059074499 | 0.139533984 |
| 644,300 | 0.138174982 | 0.189073242 | 0.059970625 | 0.138951267 |
| 644,400 | 0.138119493 | 0.188272323 | 0.059603829 | 0.139447847 |
| 644,500 | 0.138096644 | 0.18899864 | 0.05934967 | 0.139435516 |
| 644,600 | 0.138063949 | 0.188861191 | 0.059423954 | 0.139219043 |
| 644,700 | 0.138166119 | 0.188783586 | 0.059914477 | 0.138761013 |
| 644,800 | 0.138044966 | 0.188875096 | 0.059595357 | 0.138738699 |
| 644,900 | 0.138119825 | 0.189262649 | 0.059640797 | 0.139127479 |
| 645,000 | 0.138101325 | 0.187869841 | 0.059732387 | 0.139222039 |
| 645,100 | 0.138031167 | 0.188846409 | 0.058874786 | 0.13885281 |
| 645,200 | 0.137993901 | 0.189021515 | 0.059400844 | 0.139181444 |
| 645,300 | 0.137937405 | 0.189380996 | 0.059634427 | 0.139086667 |
| 645,400 | 0.137945023 | 0.188437191 | 0.059268873 | 0.139279944 |
| 645,500 | 0.137812042 | 0.188400432 | 0.05936949 | 0.138965072 |
| 645,600 | 0.137827803 | 0.188633356 | 0.059422958 | 0.138979296 |
| 645,700 | 0.137913067 | 0.189045059 | 0.058691593 | 0.139217143 |
| 645,800 | 0.137869262 | 0.188032327 | 0.059300685 | 0.139331555 |
| 645,900 | 0.137962282 | 0.18845941 | 0.059626792 | 0.138710307 |
| 646,000 | 0.137884201 | 0.188280342 | 0.059463119 | 0.138636779 |
| 646,100 | 0.137877112 | 0.189151464 | 0.0593959 | 0.139108917 |
| 646,200 | 0.13788276 | 0.188982666 | 0.059514239 | 0.139271156 |
| 646,300 | 0.137873308 | 0.189521186 | 0.059516772 | 0.139066651 |
| 646,400 | 0.137846604 | 0.188058283 | 0.05949197 | 0.139616042 |
| 646,500 | 0.137834963 | 0.188501913 | 0.059874465 | 0.139174618 |
| 646,600 | 0.137725411 | 0.188772392 | 0.059828677 | 0.139305462 |
| 646,700 | 0.13785559 | 0.188656197 | 0.058991923 | 0.13953798 |
| 646,800 | 0.137812882 | 0.189186014 | 0.059129854 | 0.138707663 |
| 646,900 | 0.137791959 | 0.18895114 | 0.059653578 | 0.139497892 |
| 647,000 | 0.137771708 | 0.188642218 | 0.059290137 | 0.13969051 |
| 647,100 | 0.137713306 | 0.188811532 | 0.059323535 | 0.140103146 |
| 647,200 | 0.137830786 | 0.188436018 | 0.059827711 | 0.139894355 |
| 647,300 | 0.137727767 | 0.188879897 | 0.05909805 | 0.139772948 |
| 647,400 | 0.137693339 | 0.188674957 | 0.059350185 | 0.139204686 |
| 647,500 | 0.137743374 | 0.18863781 | 0.059940583 | 0.139695841 |
| 647,600 | 0.137716005 | 0.188828725 | 0.059020365 | 0.139445682 |
| 647,700 | 0.137597645 | 0.188260297 | 0.059935454 | 0.139627795 |
| 647,800 | 0.137672188 | 0.188718761 | 0.059741501 | 0.14019528 |
| 647,900 | 0.137664174 | 0.188375863 | 0.059578661 | 0.140558805 |
| 648,000 | 0.137699667 | 0.188523965 | 0.059901809 | 0.140437969 |
| 648,100 | 0.137624699 | 0.188135623 | 0.060329542 | 0.139811624 |
| 648,200 | 0.137779464 | 0.188602429 | 0.06007552 | 0.140461434 |
| 648,300 | 0.137663193 | 0.187964378 | 0.059761075 | 0.140406643 |
| 648,400 | 0.137643042 | 0.18865506 | 0.060240387 | 0.140363841 |
| 648,500 | 0.137645793 | 0.18849579 | 0.059618219 | 0.140830629 |
| 648,600 | 0.137592831 | 0.188643003 | 0.059222604 | 0.140176938 |
| 648,700 | 0.137669143 | 0.188450708 | 0.059387056 | 0.140154019 |
| 648,800 | 0.137523297 | 0.188577802 | 0.059775451 | 0.140251065 |

| | | | | |
|---|---|---|---|---|
| 648,900 | 0.137454205 | 0.188658484 | 0.059798485 | 0.140029409 |
| 649,000 | 0.137509732 | 0.188976781 | 0.05946741 | 0.13988966 |
| 649,100 | 0.13745722 | 0.188987183 | 0.058927027 | 0.139948131 |
| 649,200 | 0.137559135 | 0.189378535 | 0.059189608 | 0.14025841 |
| 649,300 | 0.137584511 | 0.189445289 | 0.059699645 | 0.1399476 |
| 649,400 | 0.137533273 | 0.189937634 | 0.059017585 | 0.13981264 |
| 649,500 | 0.137563723 | 0.189841607 | 0.059028374 | 0.13967152 |
| 649,600 | 0.137434834 | 0.1890954 | 0.058778897 | 0.138982598 |
| 649,700 | 0.137322716 | 0.189492652 | 0.058665097 | 0.13964809 |
| 649,800 | 0.137342093 | 0.189166715 | 0.059356076 | 0.139321555 |
| 649,900 | 0.137490537 | 0.189226028 | 0.060060337 | 0.13909532 |
| 650,000 | 0.137414288 | 0.189266807 | 0.059638296 | 0.139928484 |
| 650,100 | 0.137434922 | 0.18940054 | 0.059646916 | 0.139727932 |
| 650,200 | 0.137374031 | 0.189314528 | 0.059894186 | 0.139387186 |
| 650,300 | 0.137410561 | 0.189909047 | 0.059476134 | 0.139535436 |
| 650,400 | 0.137372908 | 0.189762709 | 0.059543371 | 0.139857548 |
| 650,500 | 0.137165914 | 0.189073482 | 0.059475393 | 0.139748229 |
| 650,600 | 0.137240099 | 0.189730938 | 0.059770101 | 0.139136951 |
| 650,700 | 0.137264427 | 0.189779479 | 0.059562089 | 0.139075404 |
| 650,800 | 0.137265322 | 0.189471515 | 0.059321156 | 0.138726115 |
| 650,900 | 0.137201006 | 0.1891025 | 0.059098593 | 0.139473477 |
| 651,000 | 0.137168393 | 0.189545672 | 0.05918444 | 0.139326905 |
| 651,100 | 0.137168677 | 0.188641502 | 0.059764879 | 0.139289241 |
| 651,200 | 0.137192253 | 0.188874115 | 0.059588214 | 0.139375127 |
| 651,300 | 0.137103006 | 0.188756376 | 0.05923932 | 0.139958918 |
| 651,400 | 0.137209723 | 0.189338911 | 0.05940589 | 0.138966047 |
| 651,500 | 0.137178989 | 0.18824782 | 0.059550169 | 0.139373393 |
| 651,600 | 0.137151562 | 0.187960034 | 0.059629655 | 0.139722763 |
| 651,700 | 0.137151062 | 0.188184015 | 0.06002134 | 0.139154928 |
| 651,800 | 0.137111389 | 0.187841849 | 0.060124201 | 0.139274522 |
| 651,900 | 0.137079296 | 0.18810477 | 0.059715994 | 0.138794266 |
| 652,000 | 0.137127108 | 0.187600841 | 0.059480173 | 0.139302515 |
| 652,100 | 0.137107762 | 0.187644128 | 0.059752287 | 0.139378727 |
| 652,200 | 0.136942995 | 0.188145519 | 0.059693205 | 0.139036138 |
| 652,300 | 0.137037457 | 0.187278864 | 0.059649184 | 0.139309139 |
| 652,400 | 0.137015297 | 0.187348161 | 0.058862758 | 0.138626217 |
| 652,500 | 0.137077353 | 0.18761813 | 0.059384146 | 0.139693208 |
| 652,600 | 0.137019998 | 0.187008001 | 0.059390177 | 0.139030809 |
| 652,700 | 0.136920531 | 0.187038479 | 0.059642113 | 0.139238754 |
| 652,800 | 0.137019396 | 0.187674735 | 0.05989891 | 0.139328557 |
| 652,900 | 0.137083438 | 0.187610372 | 0.059563782 | 0.139084274 |
| 653,000 | 0.137038327 | 0.187554683 | 0.06029385 | 0.138916398 |
| 653,100 | 0.136969046 | 0.187528951 | 0.059555912 | 0.138982167 |
| 653,200 | 0.137014871 | 0.187696146 | 0.059740079 | 0.138618527 |
| 653,300 | 0.136874698 | 0.187474074 | 0.059785773 | 0.139232286 |
| 653,400 | 0.137011695 | 0.187507083 | 0.060166362 | 0.138996597 |
| 653,500 | 0.136969732 | 0.187716647 | 0.059956121 | 0.13884159 |
| 653,600 | 0.136797129 | 0.187210369 | 0.059464433 | 0.139063843 |
| 653,700 | 0.136895643 | 0.187505284 | 0.059482304 | 0.138775126 |
| 653,800 | 0.13686087 | 0.187343572 | 0.059828778 | 0.138512714 |
| 653,900 | 0.137027628 | 0.187906884 | 0.06001167 | 0.139195644 |
| 654,000 | 0.136937674 | 0.187684798 | 0.059500693 | 0.139381802 |
| 654,100 | 0.136991079 | 0.187776618 | 0.059544426 | 0.139432741 |
| 654,200 | 0.137015551 | 0.188696374 | 0.059057985 | 0.139559013 |
| 654,300 | 0.137024243 | 0.188645161 | 0.059214139 | 0.139454673 |
| 654,400 | 0.136892592 | 0.187694034 | 0.058888469 | 0.139814423 |
| 654,500 | 0.136930239 | 0.187735895 | 0.058584622 | 0.13963385 |
| 654,600 | 0.136796768 | 0.187702257 | 0.058673869 | 0.138999959 |
| 654,700 | 0.136879018 | 0.18799037 | 0.058440871 | 0.139439634 |
| 654,800 | 0.136814546 | 0.187598247 | 0.059652577 | 0.139185855 |
| 654,900 | 0.136849561 | 0.188282337 | 0.058932653 | 0.13948114 |
| 655,000 | 0.136879167 | 0.187971806 | 0.059191059 | 0.139177218 |
| 655,100 | 0.136805426 | 0.187671947 | 0.059083433 | 0.138895368 |
| 655,200 | 0.136848964 | 0.188664564 | 0.059243022 | 0.139090782 |
| 655,300 | 0.136766296 | 0.187645515 | 0.059338387 | 0.138566743 |
| 655,400 | 0.136806184 | 0.188328101 | 0.058805379 | 0.138554405 |
| 655,500 | 0.136778014 | 0.188456094 | 0.059075205 | 0.138550543 |

| | | | | |
|---|---|---|---|---|
| 655,600 | 0.136634265 | 0.187646492 | 0.058999774 | 0.138726123 |
| 655,700 | 0.136682159 | 0.188393153 | 0.058703937 | 0.138717313 |
| 655,800 | 0.136694005 | 0.187946416 | 0.059511432 | 0.139047636 |
| 655,900 | 0.136647186 | 0.187754959 | 0.059166426 | 0.139159695 |
| 656,000 | 0.136624255 | 0.187976283 | 0.059085557 | 0.138872865 |
| 656,100 | 0.136630109 | 0.187642709 | 0.059144099 | 0.139466357 |
| 656,200 | 0.13666187 | 0.187286219 | 0.05915103 | 0.139560691 |
| 656,300 | 0.136624948 | 0.187154471 | 0.059743927 | 0.139179959 |
| 656,400 | 0.136734808 | 0.186821466 | 0.059353538 | 0.139094366 |
| 656,500 | 0.136582262 | 0.187085112 | 0.059117394 | 0.138921675 |
| 656,600 | 0.136662934 | 0.187006555 | 0.059291943 | 0.139766577 |
| 656,700 | 0.136673264 | 0.187313121 | 0.059692397 | 0.139413055 |
| 656,800 | 0.136617419 | 0.187454985 | 0.059593131 | 0.138853314 |
| 656,900 | 0.136553429 | 0.18729761 | 0.059363792 | 0.139250778 |
| 657,000 | 0.136571116 | 0.187511754 | 0.060029687 | 0.138968576 |
| 657,100 | 0.136597537 | 0.187800499 | 0.060080845 | 0.138539135 |
| 657,200 | 0.136521734 | 0.187702683 | 0.060075646 | 0.139259946 |
| 657,300 | 0.136582743 | 0.187665093 | 0.059836534 | 0.139120755 |
| 657,400 | 0.136616772 | 0.187838581 | 0.060169145 | 0.138787121 |
| 657,500 | 0.136459094 | 0.187282481 | 0.060338546 | 0.138824705 |
| 657,600 | 0.136488145 | 0.187429883 | 0.059853784 | 0.139445346 |
| 657,700 | 0.1365834 | 0.187444509 | 0.060028789 | 0.138978432 |
| 657,800 | 0.136506518 | 0.187201342 | 0.05991253 | 0.138893352 |
| 657,900 | 0.136581783 | 0.187099465 | 0.060294223 | 0.139167588 |
| 658,000 | 0.13641012 | 0.186810666 | 0.060388386 | 0.13871722 |
| 658,100 | 0.1362814 | 0.187011313 | 0.05984126 | 0.138397071 |
| 658,200 | 0.136404322 | 0.187204159 | 0.059596581 | 0.138952749 |
| 658,300 | 0.136391853 | 0.186688254 | 0.059877675 | 0.139148621 |
| 658,400 | 0.136328608 | 0.187030856 | 0.059554219 | 0.138834059 |
| 658,500 | 0.136191004 | 0.186875018 | 0.059942649 | 0.138796358 |
| 658,600 | 0.136325457 | 0.186684582 | 0.059249392 | 0.139142581 |
| 658,700 | 0.136284191 | 0.187402667 | 0.060347026 | 0.138311491 |
| 658,800 | 0.136301746 | 0.187235107 | 0.060062854 | 0.138840812 |
| 658,900 | 0.136271165 | 0.187388783 | 0.059422284 | 0.138815859 |
| 659,000 | 0.136366951 | 0.187350686 | 0.059627367 | 0.138474515 |
| 659,100 | 0.136254143 | 0.186978111 | 0.058855473 | 0.138457051 |
| 659,200 | 0.136420147 | 0.18756642 | 0.059515514 | 0.138989855 |
| 659,300 | 0.136331551 | 0.186903587 | 0.05913653 | 0.138620522 |
| 659,400 | 0.136356404 | 0.187200081 | 0.059689106 | 0.138414539 |
| 659,500 | 0.136347957 | 0.187880319 | 0.059408395 | 0.139052961 |
| 659,600 | 0.136308523 | 0.187405991 | 0.058750531 | 0.138890775 |
| 659,700 | 0.136261097 | 0.186526511 | 0.059107199 | 0.139127753 |
| 659,800 | 0.136296928 | 0.187440283 | 0.058672286 | 0.138986925 |
| 659,900 | 0.136223749 | 0.187715089 | 0.058902346 | 0.138684153 |
| 660,000 | 0.136277853 | 0.187402209 | 0.058759232 | 0.138846329 |
| 660,100 | 0.136362822 | 0.186965902 | 0.058624763 | 0.139393986 |
| 660,200 | 0.136277889 | 0.18736332 | 0.058957471 | 0.139235655 |
| 660,300 | 0.136290304 | 0.187196828 | 0.0584406 | 0.139192453 |
| 660,400 | 0.136329455 | 0.18680858 | 0.05813193 | 0.139290095 |
| 660,500 | 0.136196341 | 0.18760499 | 0.058240588 | 0.139421632 |
| 660,600 | 0.136316548 | 0.187335 | 0.058823651 | 0.139072148 |
| 660,700 | 0.13619451 | 0.187640987 | 0.058387595 | 0.139127873 |
| 660,800 | 0.13626835 | 0.187464683 | 0.057824657 | 0.13904923 |
| 660,900 | 0.136320489 | 0.18799211 | 0.058014378 | 0.139006509 |
| 661,000 | 0.136331077 | 0.18760144 | 0.058438266 | 0.138705107 |
| 661,100 | 0.136284209 | 0.18734368 | 0.058430276 | 0.138748571 |
| 661,200 | 0.136314213 | 0.187073127 | 0.058291469 | 0.138094585 |
| 661,300 | 0.13631017 | 0.187175809 | 0.058948399 | 0.139198774 |
| 661,400 | 0.136306488 | 0.186850884 | 0.058427603 | 0.138304489 |
| 661,500 | 0.136270777 | 0.187230419 | 0.058193537 | 0.138410649 |
| 661,600 | 0.136137045 | 0.186378405 | 0.058891613 | 0.139017416 |
| 661,700 | 0.136144268 | 0.18633855 | 0.058599439 | 0.139113107 |
| 661,800 | 0.136113827 | 0.187220889 | 0.058800276 | 0.138953048 |
| 661,900 | 0.136087628 | 0.186922118 | 0.058857976 | 0.139387652 |
| 662,000 | 0.136075445 | 0.187318186 | 0.058919349 | 0.139247498 |
| 662,100 | 0.1360225 | 0.186207236 | 0.058473379 | 0.138883763 |
| 662,200 | 0.136060016 | 0.186494875 | 0.058852297 | 0.139709778 |

| | | | | |
|---|---|---|---|---|
| 662,300 | 0.136092274 | 0.187112784 | 0.058236827 | 0.13846545 |
| 662,400 | 0.136021104 | 0.187748126 | 0.058831776 | 0.139074333 |
| 662,500 | 0.136089717 | 0.186894162 | 0.058963749 | 0.139458075 |
| 662,600 | 0.135958807 | 0.186758227 | 0.058608966 | 0.139113983 |
| 662,700 | 0.136003004 | 0.186687069 | 0.058417158 | 0.139192986 |
| 662,800 | 0.135827028 | 0.186982927 | 0.058963349 | 0.138603374 |
| 662,900 | 0.135876111 | 0.187352587 | 0.0585512 | 0.138628564 |
| 663,000 | 0.135861127 | 0.186316593 | 0.058497469 | 0.138660094 |
| 663,100 | 0.135871737 | 0.186983755 | 0.058274636 | 0.138659335 |
| 663,200 | 0.135918761 | 0.186557519 | 0.059052042 | 0.139105236 |
| 663,300 | 0.135955868 | 0.186710167 | 0.059415675 | 0.138939909 |
| 663,400 | 0.135964039 | 0.18657171 | 0.058952286 | 0.138987425 |
| 663,500 | 0.13587803 | 0.186210453 | 0.058213182 | 0.138242303 |
| 663,600 | 0.135839337 | 0.186685269 | 0.058884032 | 0.138601804 |
| 663,700 | 0.135796131 | 0.186977925 | 0.058372617 | 0.138135141 |
| 663,800 | 0.135764935 | 0.186566285 | 0.057843307 | 0.137822447 |
| 663,900 | 0.135819212 | 0.186638545 | 0.058511682 | 0.138184885 |
| 664,000 | 0.13567613 | 0.186731986 | 0.058302696 | 0.13808174 |
| 664,100 | 0.135781667 | 0.186095159 | 0.058028912 | 0.138658406 |
| 664,200 | 0.135799173 | 0.187164828 | 0.058615769 | 0.138549639 |
| 664,300 | 0.135812247 | 0.187097472 | 0.057899756 | 0.138517396 |
| 664,400 | 0.135824513 | 0.18681551 | 0.058002861 | 0.138645523 |
| 664,500 | 0.135823082 | 0.186255224 | 0.058142106 | 0.138361071 |
| 664,600 | 0.13580026 | 0.185578369 | 0.057932672 | 0.138616481 |
| 664,700 | 0.135936342 | 0.186445362 | 0.057752675 | 0.138591082 |
| 664,800 | 0.13582352 | 0.186551809 | 0.057802659 | 0.138419042 |
| 664,900 | 0.13572782 | 0.186531089 | 0.05801964 | 0.138662932 |
| 665,000 | 0.135754174 | 0.186964697 | 0.057825245 | 0.138557069 |
| 665,100 | 0.135925994 | 0.187264463 | 0.057852208 | 0.138362521 |
| 665,200 | 0.135890117 | 0.186959912 | 0.057754956 | 0.13861161 |
| 665,300 | 0.135900396 | 0.186809568 | 0.057808216 | 0.138711422 |
| 665,400 | 0.13584412 | 0.186681961 | 0.057498666 | 0.138698334 |
| 665,500 | 0.135772902 | 0.185948479 | 0.0572952 | 0.138629526 |
| 665,600 | 0.135651875 | 0.186691329 | 0.057373639 | 0.138169431 |
| 665,700 | 0.135559609 | 0.186424209 | 0.057238819 | 0.138168723 |
| 665,800 | 0.135742563 | 0.18701229 | 0.057643847 | 0.13835778 |
| 665,900 | 0.135729916 | 0.186972651 | 0.058099938 | 0.137864369 |
| 666,000 | 0.13563739 | 0.186247798 | 0.057897947 | 0.138089175 |
| 666,100 | 0.135698855 | 0.186907614 | 0.058505839 | 0.138194223 |
| 666,200 | 0.135699973 | 0.187017667 | 0.05826959 | 0.138067398 |
| 666,300 | 0.135647039 | 0.187427532 | 0.058231361 | 0.138170653 |
| 666,400 | 0.13564091 | 0.187658592 | 0.058020832 | 0.137305892 |
| 666,500 | 0.135783916 | 0.187151887 | 0.058213484 | 0.13793484 |
| 666,600 | 0.135628689 | 0.187420426 | 0.05831953 | 0.137262798 |
| 666,700 | 0.135557607 | 0.187619554 | 0.058676703 | 0.137626821 |
| 666,800 | 0.135639661 | 0.187689449 | 0.058399872 | 0.137261245 |
| 666,900 | 0.135611559 | 0.187940814 | 0.058862035 | 0.137850008 |
| 667,000 | 0.135505678 | 0.187627309 | 0.058076578 | 0.137641874 |
| 667,100 | 0.135574057 | 0.187648823 | 0.058489697 | 0.13754628 |
| 667,200 | 0.135601274 | 0.18830442 | 0.058331693 | 0.138313828 |
| 667,300 | 0.135621396 | 0.187502968 | 0.058419505 | 0.138061137 |
| 667,400 | 0.135619962 | 0.186978225 | 0.057870498 | 0.138258701 |
| 667,500 | 0.13564599 | 0.187596834 | 0.057362467 | 0.138184431 |
| 667,600 | 0.135653874 | 0.187640701 | 0.057900604 | 0.138681585 |
| 667,700 | 0.135497 | 0.188199029 | 0.058033641 | 0.137948582 |
| 667,800 | 0.135376643 | 0.187403156 | 0.05775783 | 0.137852438 |
| 667,900 | 0.135481529 | 0.18709027 | 0.058156895 | 0.13789686 |
| 668,000 | 0.135395107 | 0.187147459 | 0.057942889 | 0.137653848 |
| 668,100 | 0.135474712 | 0.187197616 | 0.057358327 | 0.138013575 |
| 668,200 | 0.135372705 | 0.187108504 | 0.058226172 | 0.137845818 |
| 668,300 | 0.135335567 | 0.187241699 | 0.057845555 | 0.138024526 |
| 668,400 | 0.135372996 | 0.18689234 | 0.058241947 | 0.139016457 |
| 668,500 | 0.135257783 | 0.187515929 | 0.058123134 | 0.1382975 |
| 668,600 | 0.135422309 | 0.187331362 | 0.057702383 | 0.138208408 |
| 668,700 | 0.135349815 | 0.187595149 | 0.057933705 | 0.138059108 |
| 668,800 | 0.135278813 | 0.186952249 | 0.058618466 | 0.138281167 |
| 668,900 | 0.135420687 | 0.187392299 | 0.057837423 | 0.138423245 |

| | | | | |
|---|---|---|---|---|
| 669,000 | 0.135366808 | 0.18740019 | 0.058232392 | 0.137874807 |
| 669,100 | 0.135375131 | 0.187156733 | 0.058310086 | 0.138090899 |
| 669,200 | 0.135367361 | 0.187526671 | 0.05771404 | 0.137910335 |
| 669,300 | 0.135370608 | 0.187783511 | 0.05840308 | 0.138150734 |
| 669,400 | 0.135236999 | 0.187086321 | 0.058064746 | 0.137534211 |
| 669,500 | 0.135352511 | 0.187268052 | 0.057724232 | 0.137555781 |
| 669,600 | 0.135307525 | 0.187418133 | 0.058438372 | 0.137648272 |
| 669,700 | 0.135158444 | 0.187000686 | 0.058135752 | 0.138012885 |
| 669,800 | 0.135138285 | 0.187117313 | 0.058087396 | 0.13844046 |
| 669,900 | 0.135118428 | 0.187035269 | 0.058144669 | 0.138683123 |
| 670,000 | 0.135246508 | 0.187451817 | 0.057808878 | 0.138086333 |
| 670,100 | 0.135161251 | 0.187073812 | 0.058185744 | 0.138144293 |
| 670,200 | 0.135140252 | 0.186928026 | 0.057838286 | 0.138295379 |
| 670,300 | 0.135155316 | 0.186355814 | 0.057938177 | 0.13849211 |
| 670,400 | 0.135149649 | 0.187282973 | 0.058429464 | 0.137704722 |
| 670,500 | 0.135086299 | 0.186993862 | 0.058516366 | 0.13799535 |
| 670,600 | 0.134946416 | 0.186382867 | 0.0579471 | 0.138131834 |
| 670,700 | 0.134969002 | 0.186957997 | 0.058081552 | 0.138513468 |
| 670,800 | 0.134973702 | 0.186049297 | 0.058238983 | 0.138189605 |
| 670,900 | 0.135028446 | 0.18553616 | 0.058243583 | 0.138167559 |
| 671,000 | 0.134922214 | 0.18632885 | 0.057932787 | 0.138790927 |
| 671,100 | 0.135044448 | 0.186238097 | 0.057965883 | 0.137988474 |
| 671,200 | 0.135009103 | 0.186015324 | 0.05808391 | 0.137398333 |
| 671,300 | 0.135085753 | 0.185839287 | 0.057861079 | 0.137944615 |
| 671,400 | 0.134954967 | 0.185761664 | 0.058690685 | 0.13808178 |
| 671,500 | 0.134950461 | 0.185645622 | 0.058295086 | 0.138124692 |
| 671,600 | 0.134852864 | 0.185888057 | 0.057777242 | 0.138426533 |
| 671,700 | 0.134797078 | 0.185943303 | 0.058473156 | 0.138053525 |
| 671,800 | 0.13479661 | 0.18558329 | 0.05786296 | 0.137859312 |
| 671,900 | 0.134790888 | 0.186443405 | 0.057531512 | 0.138117509 |
| 672,000 | 0.134712892 | 0.185316681 | 0.057749886 | 0.137619766 |
| 672,100 | 0.134621678 | 0.185832529 | 0.057852569 | 0.137312715 |
| 672,200 | 0.13480549 | 0.18554105 | 0.05714817 | 0.13781711 |
| 672,300 | 0.134866924 | 0.185988626 | 0.057658346 | 0.137803185 |
| 672,400 | 0.134850885 | 0.185872609 | 0.057433855 | 0.137233727 |
| 672,500 | 0.134901734 | 0.185489418 | 0.058293483 | 0.13748275 |
| 672,600 | 0.134819617 | 0.185683179 | 0.057485479 | 0.137924917 |
| 672,700 | 0.134937643 | 0.186005542 | 0.058113107 | 0.136842133 |
| 672,800 | 0.134913634 | 0.186287159 | 0.05753978 | 0.137409843 |
| 672,900 | 0.134860926 | 0.185386809 | 0.057794129 | 0.136808501 |
| 673,000 | 0.134820124 | 0.186093951 | 0.05747996 | 0.137053022 |
| 673,100 | 0.134903601 | 0.185161494 | 0.057725789 | 0.136847436 |
| 673,200 | 0.134770364 | 0.185218435 | 0.057604714 | 0.136931504 |
| 673,300 | 0.134914609 | 0.185746855 | 0.057453191 | 0.137505065 |
| 673,400 | 0.13487711 | 0.185592469 | 0.057395205 | 0.136924566 |
| 673,500 | 0.134799246 | 0.185269628 | 0.057961198 | 0.137623251 |
| 673,600 | 0.13495745 | 0.185585426 | 0.057726535 | 0.137663579 |
| 673,700 | 0.134742454 | 0.185622779 | 0.057356414 | 0.137239173 |
| 673,800 | 0.134756378 | 0.185546782 | 0.057586242 | 0.137100276 |
| 673,900 | 0.134703955 | 0.185586403 | 0.058414779 | 0.136817649 |
| 674,000 | 0.13477698 | 0.185149058 | 0.057621781 | 0.13666603 |
| 674,100 | 0.134727706 | 0.185094998 | 0.058249962 | 0.136649883 |
| 674,200 | 0.134884774 | 0.185331226 | 0.058088351 | 0.137191071 |
| 674,300 | 0.134688778 | 0.185066771 | 0.058313676 | 0.136805703 |
| 674,400 | 0.134794868 | 0.185323474 | 0.058469082 | 0.13703293 |
| 674,500 | 0.13468139 | 0.184829475 | 0.058597697 | 0.13709122 |
| 674,600 | 0.134663756 | 0.185416919 | 0.058135995 | 0.136903659 |
| 674,700 | 0.134664632 | 0.184935027 | 0.05832616 | 0.136375117 |
| 674,800 | 0.134669061 | 0.185513313 | 0.05818554 | 0.136804193 |
| 674,900 | 0.134696769 | 0.185305326 | 0.058078002 | 0.137394363 |
| 675,000 | 0.134660197 | 0.184704718 | 0.057894824 | 0.137144799 |
| 675,100 | 0.134732442 | 0.185283017 | 0.057879673 | 0.137302078 |
| 675,200 | 0.134588194 | 0.184441556 | 0.057659609 | 0.137271674 |
| 675,300 | 0.134472327 | 0.184731747 | 0.058330869 | 0.137452646 |
| 675,400 | 0.134521258 | 0.184037032 | 0.058231127 | 0.136825373 |
| 675,500 | 0.134610578 | 0.183546171 | 0.057971439 | 0.137035298 |
| 675,600 | 0.134595568 | 0.184177164 | 0.058108882 | 0.137421592 |

| | | | | |
|---|---|---|---|---|
| 675,700 | 0.134559731 | 0.18370227 | 0.058032083 | 0.137226258 |
| 675,800 | 0.134715832 | 0.184244014 | 0.058003802 | 0.136905841 |
| 675,900 | 0.134627031 | 0.184564063 | 0.057961551 | 0.137453958 |
| 676,000 | 0.134663163 | 0.184506848 | 0.058266317 | 0.137285295 |
| 676,100 | 0.134648829 | 0.184421187 | 0.058231779 | 0.137433825 |
| 676,200 | 0.134622428 | 0.184935959 | 0.058113898 | 0.137287845 |
| 676,300 | 0.134582711 | 0.184688701 | 0.058125782 | 0.137987465 |
| 676,400 | 0.134633852 | 0.184906073 | 0.057645093 | 0.138012897 |
| 676,500 | 0.13456375 | 0.18483937 | 0.05792033 | 0.137614506 |
| 676,600 | 0.134601006 | 0.184588539 | 0.056760393 | 0.137379116 |
| 676,700 | 0.134628068 | 0.18475293 | 0.057236728 | 0.137704562 |
| 676,800 | 0.13458169 | 0.185013815 | 0.057283436 | 0.137412324 |
| 676,900 | 0.134443644 | 0.184780596 | 0.057344689 | 0.137440924 |
| 677,000 | 0.134471034 | 0.185347061 | 0.05786862 | 0.137336189 |
| 677,100 | 0.134440179 | 0.185267376 | 0.057739702 | 0.137486803 |
| 677,200 | 0.13445122 | 0.184980445 | 0.057463481 | 0.137214055 |
| 677,300 | 0.13434537 | 0.18519401 | 0.057862416 | 0.13726256 |
| 677,400 | 0.134331652 | 0.184322149 | 0.058304963 | 0.137003287 |
| 677,500 | 0.134423076 | 0.184872896 | 0.057904543 | 0.137342757 |
| 677,600 | 0.134511524 | 0.185471998 | 0.057948247 | 0.137077596 |
| 677,700 | 0.134423261 | 0.184273811 | 0.058330853 | 0.137502921 |
| 677,800 | 0.13437971 | 0.184703024 | 0.057603842 | 0.13768118 |
| 677,900 | 0.134331505 | 0.18419941 | 0.058086211 | 0.137389022 |
| 678,000 | 0.13435271 | 0.184829509 | 0.058432868 | 0.137782188 |
| 678,100 | 0.134338101 | 0.1843566 | 0.058537356 | 0.137492767 |
| 678,200 | 0.134336467 | 0.184998841 | 0.058281873 | 0.137114923 |
| 678,300 | 0.134254449 | 0.184477049 | 0.058243583 | 0.136938617 |
| 678,400 | 0.134226658 | 0.184358779 | 0.058464297 | 0.137346621 |
| 678,500 | 0.13412945 | 0.184664203 | 0.057741536 | 0.136609996 |
| 678,600 | 0.134225498 | 0.184745154 | 0.058386679 | 0.137792667 |
| 678,700 | 0.134208156 | 0.184576888 | 0.057694391 | 0.13772593 |
| 678,800 | 0.134102314 | 0.183889611 | 0.057827281 | 0.137480614 |
| 678,900 | 0.134136693 | 0.184183136 | 0.058295189 | 0.13726244 |
| 679,000 | 0.134095277 | 0.183702372 | 0.058188448 | 0.137630708 |
| 679,100 | 0.134107128 | 0.184123755 | 0.05783248 | 0.137161621 |
| 679,200 | 0.134088774 | 0.18447268 | 0.058100516 | 0.137355034 |
| 679,300 | 0.134064942 | 0.184028781 | 0.058604966 | 0.137331388 |
| 679,400 | 0.134038548 | 0.183770022 | 0.058138107 | 0.137657814 |
| 679,500 | 0.133865059 | 0.183690168 | 0.05801097 | 0.136994971 |
| 679,600 | 0.13400477 | 0.184066487 | 0.05871832 | 0.136573233 |
| 679,700 | 0.133885312 | 0.184109851 | 0.058535325 | 0.136273752 |
| 679,800 | 0.133947668 | 0.184388478 | 0.058951571 | 0.136890307 |
| 679,900 | 0.133896793 | 0.184784151 | 0.059016876 | 0.136960019 |
| 680,000 | 0.133791702 | 0.184412369 | 0.059022495 | 0.137093073 |
| 680,100 | 0.133826758 | 0.184089553 | 0.05901097 | 0.136581065 |
| 680,200 | 0.133871685 | 0.184035322 | 0.059058716 | 0.136850667 |
| 680,300 | 0.133892997 | 0.183629981 | 0.058905618 | 0.136936436 |
| 680,400 | 0.133844351 | 0.183591828 | 0.059194611 | 0.136999273 |
| 680,500 | 0.133793078 | 0.184035823 | 0.059087197 | 0.137216531 |
| 680,600 | 0.133814965 | 0.183455492 | 0.059038826 | 0.136989865 |
| 680,700 | 0.133878838 | 0.183133535 | 0.059392123 | 0.136993976 |
| 680,800 | 0.133853665 | 0.183983911 | 0.059423565 | 0.137526926 |
| 680,900 | 0.133770379 | 0.183901242 | 0.059480432 | 0.137270146 |
| 681,000 | 0.133741139 | 0.184122542 | 0.059750238 | 0.137355846 |
| 681,100 | 0.133763305 | 0.184198501 | 0.06005209 | 0.137029806 |
| 681,200 | 0.13373167 | 0.183610285 | 0.05981561 | 0.136443168 |
| 681,300 | 0.133673497 | 0.183655513 | 0.060018387 | 0.137169437 |
| 681,400 | 0.133829526 | 0.183801841 | 0.060192709 | 0.137067606 |
| 681,500 | 0.133786181 | 0.183468546 | 0.05980755 | 0.137057161 |
| 681,600 | 0.133657592 | 0.1842718 | 0.059537573 | 0.136668716 |
| 681,700 | 0.133641939 | 0.183740602 | 0.05955382 | 0.137000005 |
| 681,800 | 0.133568343 | 0.183071737 | 0.059796373 | 0.136482115 |
| 681,900 | 0.133643316 | 0.183319714 | 0.059730429 | 0.137231738 |
| 682,000 | 0.133594939 | 0.183931418 | 0.059157928 | 0.136548158 |
| 682,100 | 0.133605006 | 0.182709575 | 0.059025856 | 0.137172028 |
| 682,200 | 0.133682645 | 0.182744231 | 0.060070391 | 0.137278001 |
| 682,300 | 0.133748813 | 0.182880138 | 0.059398276 | 0.137252242 |

| | | | | |
|---|---|---|---|---|
| 682,400 | 0.133678981 | 0.182600382 | 0.059632638 | 0.137001902 |
| 682,500 | 0.133615008 | 0.183288451 | 0.059580797 | 0.136900489 |
| 682,600 | 0.13367268 | 0.182798293 | 0.059765126 | 0.1365052 |
| 682,700 | 0.133629387 | 0.183006627 | 0.058934262 | 0.136822029 |
| 682,800 | 0.133545029 | 0.182812006 | 0.060510531 | 0.136707114 |
| 682,900 | 0.133615843 | 0.183330943 | 0.059963531 | 0.13722764 |
| 683,000 | 0.133467355 | 0.18295761 | 0.060200798 | 0.137023998 |
| 683,100 | 0.13349605 | 0.183017733 | 0.059867379 | 0.137077397 |
| 683,200 | 0.133570237 | 0.183521436 | 0.059913512 | 0.137275212 |
| 683,300 | 0.133439785 | 0.18308126 | 0.060372653 | 0.137888031 |
| 683,400 | 0.133459532 | 0.183368096 | 0.0599724 | 0.137309268 |
| 683,500 | 0.133350688 | 0.182803206 | 0.060513964 | 0.137348473 |
| 683,600 | 0.133360782 | 0.183148158 | 0.060188642 | 0.137598295 |
| 683,700 | 0.133319116 | 0.183341319 | 0.060658345 | 0.137287449 |
| 683,800 | 0.13336768 | 0.183806284 | 0.060124417 | 0.137250217 |
| 683,900 | 0.133447731 | 0.183915369 | 0.059953839 | 0.137738218 |
| 684,000 | 0.133347461 | 0.183468658 | 0.059771336 | 0.137600712 |
| 684,100 | 0.133395219 | 0.183795311 | 0.059494732 | 0.138119606 |
| 684,200 | 0.13330386 | 0.183722407 | 0.059541086 | 0.137849494 |
| 684,300 | 0.133367885 | 0.183477336 | 0.059954163 | 0.137889944 |
| 684,400 | 0.133352285 | 0.183993349 | 0.059576928 | 0.137642357 |
| 684,500 | 0.133247664 | 0.182893675 | 0.060076014 | 0.137710767 |
| 684,600 | 0.133199662 | 0.183334032 | 0.059690343 | 0.137540487 |
| 684,700 | 0.133219418 | 0.183569789 | 0.060361505 | 0.137431562 |
| 684,800 | 0.133352577 | 0.184162017 | 0.059971652 | 0.137850789 |
| 684,900 | 0.133294537 | 0.183306516 | 0.059586517 | 0.137856374 |
| 685,000 | 0.133279351 | 0.18323583 | 0.059019475 | 0.13769886 |
| 685,100 | 0.133169341 | 0.18361285 | 0.059652725 | 0.137735054 |
| 685,200 | 0.133266685 | 0.183476064 | 0.059098724 | 0.137604523 |
| 685,300 | 0.133237549 | 0.183228939 | 0.059507656 | 0.137471821 |
| 685,400 | 0.133242916 | 0.183705885 | 0.059826492 | 0.137931667 |
| 685,500 | 0.133270202 | 0.183491261 | 0.060018616 | 0.137457338 |
| 685,600 | 0.133038337 | 0.182955767 | 0.059623043 | 0.137424066 |
| 685,700 | 0.133064589 | 0.18376831 | 0.059888725 | 0.137037093 |
| 685,800 | 0.133012545 | 0.183094663 | 0.060418006 | 0.137518182 |
| 685,900 | 0.133029724 | 0.183515975 | 0.059957704 | 0.137282802 |
| 686,000 | 0.132919849 | 0.183352256 | 0.060100348 | 0.13698127 |
| 686,100 | 0.132885052 | 0.183388358 | 0.059654225 | 0.136896053 |
| 686,200 | 0.132868779 | 0.182867465 | 0.059549924 | 0.137463802 |
| 686,300 | 0.132923475 | 0.182780235 | 0.060021795 | 0.1371126 |
| 686,400 | 0.132972967 | 0.183406337 | 0.060123288 | 0.137014501 |
| 686,500 | 0.132922141 | 0.183455386 | 0.059981785 | 0.137893015 |
| 686,600 | 0.133042853 | 0.182939161 | 0.060299475 | 0.137643348 |
| 686,700 | 0.132985239 | 0.18307995 | 0.060555828 | 0.137401899 |
| 686,800 | 0.133072403 | 0.183034123 | 0.060389157 | 0.137646244 |
| 686,900 | 0.133002655 | 0.183927325 | 0.060806174 | 0.137311522 |
| 687,000 | 0.132808607 | 0.183017077 | 0.06091379 | 0.137461184 |
| 687,100 | 0.132905888 | 0.183570436 | 0.060450315 | 0.13766932 |
| 687,200 | 0.133006153 | 0.183852237 | 0.060311264 | 0.138140237 |
| 687,300 | 0.132881892 | 0.183240685 | 0.059852879 | 0.137948075 |
| 687,400 | 0.13289127 | 0.183675793 | 0.060167372 | 0.13770124 |
| 687,500 | 0.132925781 | 0.182413541 | 0.060008929 | 0.137571109 |
| 687,600 | 0.132832212 | 0.182703364 | 0.059700694 | 0.137992952 |
| 687,700 | 0.132940538 | 0.183084464 | 0.060449602 | 0.137470607 |
| 687,800 | 0.132853236 | 0.182519235 | 0.060533692 | 0.137394308 |
| 687,900 | 0.13270233 | 0.182384738 | 0.060253214 | 0.137603848 |
| 688,000 | 0.132701159 | 0.182447331 | 0.059998551 | 0.137000602 |
| 688,100 | 0.132666968 | 0.183056467 | 0.060464965 | 0.137034125 |
| 688,200 | 0.132606788 | 0.182743961 | 0.060336625 | 0.137112424 |
| 688,300 | 0.132672361 | 0.182710114 | 0.06010276 | 0.137366915 |
| 688,400 | 0.132625588 | 0.182858394 | 0.06061596 | 0.137106764 |
| 688,500 | 0.132790521 | 0.182088049 | 0.060495532 | 0.137373582 |
| 688,600 | 0.132682538 | 0.182209027 | 0.060720009 | 0.13734575 |
| 688,700 | 0.132634748 | 0.182142592 | 0.060268526 | 0.137478837 |
| 688,800 | 0.132566448 | 0.182464401 | 0.060300915 | 0.137448962 |
| 688,900 | 0.132510609 | 0.18221309 | 0.059965651 | 0.137637357 |
| 689,000 | 0.132477058 | 0.182417086 | 0.060090645 | 0.136771462 |

| | | | | |
|---|---|---|---|---|
| 689,100 | 0.13254689 | 0.182820683 | 0.059672888 | 0.137086451 |
| 689,200 | 0.132595488 | 0.182989126 | 0.060555136 | 0.137013577 |
| 689,300 | 0.132597454 | 0.182605373 | 0.060044134 | 0.13709548 |
| 689,400 | 0.132635709 | 0.182752125 | 0.060048333 | 0.137519735 |
| 689,500 | 0.132496889 | 0.182939024 | 0.059859044 | 0.137032193 |
| 689,600 | 0.132365934 | 0.182087644 | 0.059691461 | 0.137049615 |
| 689,700 | 0.132383799 | 0.182807317 | 0.05992445 | 0.137317063 |
| 689,800 | 0.132517097 | 0.183593343 | 0.059929691 | 0.1376074 |
| 689,900 | 0.132458223 | 0.18353337 | 0.059562163 | 0.137274471 |
| 690,000 | 0.132458673 | 0.183778818 | 0.059975454 | 0.137273383 |
| 690,100 | 0.132464993 | 0.18330752 | 0.05985232 | 0.137347964 |
| 690,200 | 0.132460878 | 0.183351431 | 0.059962087 | 0.137127396 |
| 690,300 | 0.132562922 | 0.183782812 | 0.059702075 | 0.137058842 |
| 690,400 | 0.132510305 | 0.18375873 | 0.059937562 | 0.136967704 |
| 690,500 | 0.132388306 | 0.183243929 | 0.059829847 | 0.136410804 |
| 690,600 | 0.132458616 | 0.183141735 | 0.060198243 | 0.137062542 |
| 690,700 | 0.132511418 | 0.183771014 | 0.060024588 | 0.13681188 |
| 690,800 | 0.132376578 | 0.184086465 | 0.059916573 | 0.1370612 |
| 690,900 | 0.132455563 | 0.18376456 | 0.060463956 | 0.137278959 |
| 691,000 | 0.132422295 | 0.183060209 | 0.059986101 | 0.137115107 |
| 691,100 | 0.132468257 | 0.183657534 | 0.060188109 | 0.137177541 |
| 691,200 | 0.132371196 | 0.183058406 | 0.059884825 | 0.136959835 |
| 691,300 | 0.132400177 | 0.183207181 | 0.059899083 | 0.137033954 |
| 691,400 | 0.132417029 | 0.18266459 | 0.0600317 | 0.137480239 |
| 691,500 | 0.13237222 | 0.183356801 | 0.059760663 | 0.137683381 |
| 691,600 | 0.132589418 | 0.183311749 | 0.059890169 | 0.137409776 |
| 691,700 | 0.132359673 | 0.183018757 | 0.059857685 | 0.137404556 |
| 691,800 | 0.132385808 | 0.182577952 | 0.060263822 | 0.136905974 |
| 691,900 | 0.13239065 | 0.183224613 | 0.060244047 | 0.136820755 |
| 692,000 | 0.132390734 | 0.183410526 | 0.060409481 | 0.136886775 |
| 692,100 | 0.132257079 | 0.183114495 | 0.060762983 | 0.136820918 |
| 692,200 | 0.132407436 | 0.183995114 | 0.059667142 | 0.136785223 |
| 692,300 | 0.132322108 | 0.183640721 | 0.060667931 | 0.136232812 |
| 692,400 | 0.132516284 | 0.183875452 | 0.060043241 | 0.136630299 |
| 692,500 | 0.132442939 | 0.183877209 | 0.060061893 | 0.136123975 |
| 692,600 | 0.132533979 | 0.183946341 | 0.059930584 | 0.135871647 |
| 692,700 | 0.132289955 | 0.183180771 | 0.059265297 | 0.13569305 |
| 692,800 | 0.132353056 | 0.182858242 | 0.059974407 | 0.135883271 |
| 692,900 | 0.132437566 | 0.183319698 | 0.060128677 | 0.136192693 |
| 693,000 | 0.132410345 | 0.183190712 | 0.060253108 | 0.135789401 |
| 693,100 | 0.132348071 | 0.183455117 | 0.06000281 | 0.136282813 |
| 693,200 | 0.132415147 | 0.183438583 | 0.060339856 | 0.136421721 |
| 693,300 | 0.132476917 | 0.183167448 | 0.060583376 | 0.136221005 |
| 693,400 | 0.132473975 | 0.183323057 | 0.06101506 | 0.136248171 |
| 693,500 | 0.132439863 | 0.18344572 | 0.060446847 | 0.135734678 |
| 693,600 | 0.132269956 | 0.18290342 | 0.060064532 | 0.136050801 |
| 693,700 | 0.132236668 | 0.183435982 | 0.060625758 | 0.136333314 |
| 693,800 | 0.13222421 | 0.183291447 | 0.060476963 | 0.136412373 |
| 693,900 | 0.132174551 | 0.182961801 | 0.060504624 | 0.136356487 |
| 694,000 | 0.132190859 | 0.183439514 | 0.060297499 | 0.135914719 |
| 694,100 | 0.132168162 | 0.183415683 | 0.060439451 | 0.135892348 |
| 694,200 | 0.132215229 | 0.18393147 | 0.059889881 | 0.136405859 |
| 694,300 | 0.131969322 | 0.183407412 | 0.059629928 | 0.136576089 |
| 694,400 | 0.132160116 | 0.1833131 | 0.060588406 | 0.136269833 |
| 694,500 | 0.132005971 | 0.183334503 | 0.060390595 | 0.135801185 |
| 694,600 | 0.132014301 | 0.18343845 | 0.060603456 | 0.135482389 |
| 694,700 | 0.131927672 | 0.182753468 | 0.059979118 | 0.136229132 |
| 694,800 | 0.132064342 | 0.183018535 | 0.060216238 | 0.136645298 |
| 694,900 | 0.132000711 | 0.183195966 | 0.060366299 | 0.136392101 |
| 695,000 | 0.132006581 | 0.182704312 | 0.059947536 | 0.136562731 |
| 695,100 | 0.132091227 | 0.182567131 | 0.0597322 | 0.135975303 |
| 695,200 | 0.13199835 | 0.182707548 | 0.060043453 | 0.136246885 |
| 695,300 | 0.132001861 | 0.182139452 | 0.059841526 | 0.136361266 |
| 695,400 | 0.132071883 | 0.183143998 | 0.060112662 | 0.13653326 |
| 695,500 | 0.132051915 | 0.182998121 | 0.059707658 | 0.136380368 |
| 695,600 | 0.132108744 | 0.182872099 | 0.060029732 | 0.136211398 |
| 695,700 | 0.132033947 | 0.182120549 | 0.059414136 | 0.136058553 |

| | | | | |
|---|---|---|---|---|
| 695,800 | 0.132083604 | 0.182460375 | 0.060088082 | 0.135995173 |
| 695,900 | 0.132038315 | 0.18241921 | 0.059759421 | 0.136168073 |
| 696,000 | 0.132133868 | 0.183077766 | 0.059521394 | 0.135899535 |
| 696,100 | 0.13195584 | 0.182154548 | 0.059343058 | 0.135473826 |
| 696,200 | 0.132132477 | 0.182723778 | 0.060001521 | 0.136357613 |
| 696,300 | 0.1320685 | 0.182709188 | 0.059767737 | 0.136008451 |
| 696,400 | 0.132038918 | 0.182440643 | 0.059165223 | 0.136466903 |
| 696,500 | 0.132004192 | 0.182823071 | 0.059254193 | 0.13658386 |
| 696,600 | 0.131873616 | 0.182720947 | 0.059945519 | 0.135868723 |
| 696,700 | 0.131859483 | 0.182808355 | 0.059970481 | 0.135876523 |
| 696,800 | 0.131932135 | 0.182661383 | 0.05973034 | 0.136172949 |
| 696,900 | 0.131954787 | 0.182312903 | 0.059707808 | 0.135840788 |
| 697,000 | 0.131837868 | 0.182231985 | 0.059972426 | 0.135265368 |
| 697,100 | 0.131914826 | 0.182543946 | 0.060792237 | 0.13563966 |
| 697,200 | 0.131840373 | 0.182569359 | 0.060141747 | 0.135469845 |
| 697,300 | 0.131842379 | 0.182083716 | 0.060361918 | 0.135359703 |
| 697,400 | 0.131843298 | 0.181869473 | 0.05994903 | 0.135598503 |
| 697,500 | 0.131963793 | 0.182835974 | 0.060217126 | 0.135797522 |
| 697,600 | 0.132004188 | 0.182375211 | 0.060136227 | 0.136191558 |
| 697,700 | 0.131860292 | 0.182104032 | 0.059741668 | 0.136201972 |
| 697,800 | 0.131970355 | 0.181710408 | 0.059796956 | 0.136121875 |
| 697,900 | 0.131940933 | 0.181884716 | 0.059436409 | 0.136366529 |
| 698,000 | 0.13193895 | 0.182012601 | 0.059980081 | 0.135688468 |
| 698,100 | 0.13172127 | 0.181782888 | 0.060160207 | 0.135888079 |
| 698,200 | 0.131843258 | 0.181093048 | 0.059870886 | 0.135499807 |
| 698,300 | 0.13178182 | 0.181953598 | 0.059580044 | 0.135472846 |
| 698,400 | 0.131799696 | 0.181777489 | 0.059971637 | 0.13607769 |
| 698,500 | 0.131913786 | 0.181770431 | 0.059970609 | 0.135986576 |
| 698,600 | 0.131827244 | 0.182011598 | 0.059724572 | 0.135720054 |
| 698,700 | 0.131912641 | 0.182078693 | 0.059951238 | 0.135506015 |
| 698,800 | 0.131918675 | 0.181697573 | 0.060100491 | 0.135107458 |
| 698,900 | 0.131887568 | 0.182093724 | 0.05985246 | 0.135575421 |
| 699,000 | 0.131832773 | 0.182290757 | 0.059791683 | 0.135510743 |
| 699,100 | 0.131819647 | 0.181967522 | 0.059660144 | 0.135311228 |
| 699,200 | 0.131885208 | 0.182457103 | 0.05989532 | 0.135708918 |
| 699,300 | 0.131873886 | 0.182509962 | 0.059400888 | 0.134604816 |
| 699,400 | 0.131741978 | 0.181808063 | 0.058797086 | 0.135194901 |
| 699,500 | 0.131865725 | 0.182227204 | 0.059335662 | 0.134975262 |
| 699,600 | 0.131798734 | 0.182369012 | 0.059098395 | 0.135118639 |
| 699,700 | 0.131824627 | 0.181926072 | 0.05873511 | 0.134798374 |
| 699,800 | 0.131805916 | 0.182422737 | 0.059459454 | 0.134841437 |
| 699,900 | 0.131602682 | 0.182358053 | 0.059614053 | 0.134733601 |
| 700,000 | 0.131711414 | 0.182144098 | 0.05971485 | 0.134467806 |
| 700,100 | 0.131774959 | 0.181765273 | 0.059325586 | 0.134908961 |
| 700,200 | 0.131729309 | 0.181834024 | 0.059274055 | 0.135660815 |
| 700,300 | 0.13162008 | 0.181774931 | 0.059000978 | 0.135073304 |
| 700,400 | 0.13166123 | 0.181913589 | 0.05905296 | 0.134749357 |
| 700,500 | 0.131710268 | 0.181545706 | 0.058850657 | 0.134839003 |
| 700,600 | 0.131548716 | 0.182104314 | 0.059143015 | 0.134941981 |
| 700,700 | 0.131596581 | 0.181576403 | 0.059796521 | 0.134814905 |
| 700,800 | 0.131634225 | 0.181527005 | 0.059748891 | 0.135023451 |
| 700,900 | 0.131615227 | 0.182241961 | 0.059778165 | 0.134399191 |
| 701,000 | 0.131702027 | 0.181900033 | 0.05996211 | 0.134442982 |
| 701,100 | 0.131658513 | 0.181861222 | 0.058961507 | 0.134836751 |
| 701,200 | 0.131721315 | 0.18192843 | 0.059578991 | 0.134678369 |
| 701,300 | 0.131680189 | 0.181758986 | 0.059184483 | 0.134877965 |
| 701,400 | 0.131663848 | 0.182250278 | 0.059615978 | 0.134804249 |
| 701,500 | 0.131570021 | 0.181952396 | 0.059396045 | 0.134649564 |
| 701,600 | 0.131769842 | 0.181599562 | 0.05903629 | 0.135354427 |
| 701,700 | 0.131605738 | 0.181859611 | 0.059244418 | 0.135168574 |
| 701,800 | 0.131563252 | 0.181853638 | 0.060003738 | 0.135223996 |
| 701,900 | 0.131624848 | 0.18143014 | 0.059134017 | 0.135090065 |
| 702,000 | 0.131561672 | 0.181559208 | 0.059452923 | 0.134838026 |
| 702,100 | 0.131461312 | 0.181524988 | 0.059332646 | 0.135295559 |
| 702,200 | 0.131525543 | 0.1816538 | 0.059181955 | 0.135242053 |
| 702,300 | 0.131679414 | 0.181214211 | 0.059003669 | 0.135295557 |
| 702,400 | 0.131627047 | 0.181105267 | 0.059319791 | 0.135108751 |

| | | | | |
|---|---|---|---|---|
| 702,500 | 0.131579069 | 0.181012796 | 0.058324369 | 0.135418172 |
| 702,600 | 0.131525449 | 0.181044066 | 0.058861015 | 0.134803695 |
| 702,700 | 0.131445747 | 0.181313157 | 0.059192207 | 0.13515662 |
| 702,800 | 0.131666283 | 0.181586418 | 0.058730061 | 0.135144429 |
| 702,900 | 0.131560954 | 0.181102342 | 0.059042543 | 0.134812649 |
| 703,000 | 0.131609083 | 0.181184675 | 0.05918217 | 0.134755583 |
| 703,100 | 0.131537635 | 0.1816371 | 0.059214533 | 0.134768227 |
| 703,200 | 0.131522146 | 0.180896566 | 0.059329547 | 0.134396769 |
| 703,300 | 0.13141013 | 0.181339997 | 0.058834415 | 0.135241451 |
| 703,400 | 0.131457731 | 0.181336451 | 0.058209082 | 0.134980374 |
| 703,500 | 0.131475028 | 0.181503625 | 0.058412731 | 0.134653677 |
| 703,600 | 0.131439531 | 0.180866293 | 0.059340744 | 0.134928351 |
| 703,700 | 0.131372335 | 0.181077979 | 0.059373227 | 0.134824549 |
| 703,800 | 0.131454636 | 0.181469444 | 0.058308956 | 0.13472775 |
| 703,900 | 0.131285387 | 0.180770314 | 0.059157129 | 0.135012595 |
| 704,000 | 0.131354234 | 0.180892978 | 0.058658266 | 0.134942017 |
| 704,100 | 0.131303567 | 0.181520294 | 0.058896806 | 0.134929138 |
| 704,200 | 0.131187498 | 0.181341391 | 0.058917567 | 0.134731465 |
| 704,300 | 0.131309568 | 0.181173303 | 0.058742934 | 0.135157747 |
| 704,400 | 0.131343523 | 0.181419066 | 0.059371822 | 0.13528225 |
| 704,500 | 0.131284242 | 0.181452827 | 0.058516768 | 0.135151281 |
| 704,600 | 0.131235103 | 0.18145735 | 0.05868605 | 0.135179381 |
| 704,700 | 0.131219731 | 0.181998885 | 0.058448738 | 0.135219523 |
| 704,800 | 0.131213018 | 0.181549766 | 0.058403108 | 0.134717108 |
| 704,900 | 0.13122402 | 0.181836467 | 0.058519503 | 0.134881079 |
| 705,000 | 0.131234728 | 0.181683056 | 0.058335492 | 0.135039427 |
| 705,100 | 0.131188061 | 0.181393368 | 0.058390668 | 0.135179082 |
| 705,200 | 0.131148833 | 0.181238959 | 0.057713544 | 0.135391259 |
| 705,300 | 0.131191289 | 0.181431633 | 0.057932126 | 0.135556839 |
| 705,400 | 0.13107959 | 0.181550316 | 0.058199674 | 0.135634853 |
| 705,500 | 0.131009739 | 0.181713842 | 0.058174818 | 0.13580941 |
| 705,600 | 0.130948841 | 0.180918558 | 0.057949904 | 0.135753583 |
| 705,700 | 0.131154804 | 0.181652702 | 0.058212861 | 0.136036552 |
| 705,800 | 0.130976891 | 0.181509787 | 0.058301968 | 0.135708664 |
| 705,900 | 0.130996082 | 0.181240535 | 0.058365196 | 0.135163467 |
| 706,000 | 0.130963673 | 0.181940442 | 0.057518073 | 0.135800776 |
| 706,100 | 0.131013617 | 0.181667917 | 0.057847263 | 0.135919028 |
| 706,200 | 0.130985937 | 0.181751254 | 0.058167307 | 0.13631247 |
| 706,300 | 0.131053518 | 0.182006577 | 0.058002596 | 0.136086081 |
| 706,400 | 0.131114041 | 0.181980993 | 0.058042738 | 0.136829522 |
| 706,500 | 0.130982296 | 0.181733103 | 0.0580213 | 0.136111144 |
| 706,600 | 0.13102233 | 0.181868961 | 0.057957012 | 0.135708171 |
| 706,700 | 0.130899298 | 0.181246937 | 0.058062663 | 0.135669451 |
| 706,800 | 0.13093692 | 0.181030319 | 0.0581768 | 0.135905415 |
| 706,900 | 0.13084053 | 0.18078026 | 0.057635682 | 0.13609354 |
| 707,000 | 0.130929539 | 0.181064723 | 0.057419083 | 0.136080884 |
| 707,100 | 0.130787143 | 0.180536674 | 0.057703488 | 0.135868584 |
| 707,200 | 0.130824078 | 0.180826805 | 0.05800287 | 0.135772568 |
| 707,300 | 0.131017135 | 0.181748087 | 0.057965602 | 0.135861336 |
| 707,400 | 0.130875087 | 0.181226392 | 0.057648286 | 0.135527604 |
| 707,500 | 0.130908146 | 0.181140224 | 0.057902985 | 0.135941231 |
| 707,600 | 0.130805728 | 0.180846862 | 0.058045891 | 0.135749265 |
| 707,700 | 0.130895438 | 0.180950239 | 0.058835107 | 0.135911231 |
| 707,800 | 0.130875355 | 0.180743882 | 0.05838064 | 0.135754517 |
| 707,900 | 0.13083318 | 0.180792383 | 0.057924792 | 0.136055945 |
| 708,000 | 0.130899273 | 0.180956805 | 0.057996607 | 0.136204915 |
| 708,100 | 0.130809269 | 0.180830258 | 0.057936561 | 0.136083679 |
| 708,200 | 0.13081045 | 0.181384847 | 0.058082842 | 0.136210882 |
| 708,300 | 0.130914002 | 0.180986596 | 0.057768675 | 0.136137484 |
| 708,400 | 0.130850957 | 0.180542092 | 0.057696528 | 0.136143781 |
| 708,500 | 0.130833917 | 0.18058612 | 0.058020263 | 0.136353975 |
| 708,600 | 0.130770639 | 0.180506758 | 0.057682007 | 0.1363873 |
| 708,700 | 0.130712314 | 0.180257481 | 0.057551954 | 0.135805518 |
| 708,800 | 0.130869151 | 0.181058174 | 0.057707832 | 0.136113614 |
| 708,900 | 0.130950085 | 0.1814372 | 0.057512099 | 0.136399593 |
| 709,000 | 0.130696678 | 0.181082244 | 0.05805799 | 0.136614916 |
| 709,100 | 0.130775922 | 0.181522426 | 0.057433685 | 0.136526119 |

| | | | | |
|---|---|---|---|---|
| 709,200 | 0.130830054 | 0.181324258 | 0.057179145 | 0.136072589 |
| 709,300 | 0.13071075 | 0.181699235 | 0.057349433 | 0.136469342 |
| 709,400 | 0.130585198 | 0.181120127 | 0.057201137 | 0.136376496 |
| 709,500 | 0.13077104 | 0.180948139 | 0.05814871 | 0.135497273 |
| 709,600 | 0.130651053 | 0.180504944 | 0.057649754 | 0.135532107 |
| 709,700 | 0.130617484 | 0.181154304 | 0.058248505 | 0.134808144 |
| 709,800 | 0.130612051 | 0.181026279 | 0.058104938 | 0.13490822 |
| 709,900 | 0.13062087 | 0.18098135 | 0.05779851 | 0.135510799 |
| 710,000 | 0.130566204 | 0.181158231 | 0.057595252 | 0.135323833 |
| 710,100 | 0.130525897 | 0.180571639 | 0.057488176 | 0.135372766 |
| 710,200 | 0.130575158 | 0.18128805 | 0.057718663 | 0.135722436 |
| 710,300 | 0.130541932 | 0.181592718 | 0.057758863 | 0.135317379 |
| 710,400 | 0.130600078 | 0.181691084 | 0.058239732 | 0.135899435 |
| 710,500 | 0.130574626 | 0.181391411 | 0.057650997 | 0.135301132 |
| 710,600 | 0.130577581 | 0.181615842 | 0.057110004 | 0.135569386 |
| 710,700 | 0.13047166 | 0.181659213 | 0.05745272 | 0.135416323 |
| 710,800 | 0.130481483 | 0.181277594 | 0.057570375 | 0.134783596 |
| 710,900 | 0.130510437 | 0.181858637 | 0.057781254 | 0.135055429 |
| 711,000 | 0.130526759 | 0.182108811 | 0.057401398 | 0.135692648 |
| 711,100 | 0.130410336 | 0.182121927 | 0.057634238 | 0.134981275 |
| 711,200 | 0.130486592 | 0.181999282 | 0.057553479 | 0.134709623 |
| 711,300 | 0.130400839 | 0.181681707 | 0.057289191 | 0.135001499 |
| 711,400 | 0.130411103 | 0.181125928 | 0.057549255 | 0.134905273 |
| 711,500 | 0.130484223 | 0.180890149 | 0.057865219 | 0.13483092 |
| 711,600 | 0.130434523 | 0.181090873 | 0.057371418 | 0.135241677 |
| 711,700 | 0.130333823 | 0.180357503 | 0.057986211 | 0.134846908 |
| 711,800 | 0.13029023 | 0.180618213 | 0.057975169 | 0.135492759 |
| 711,900 | 0.130383593 | 0.180912451 | 0.058099884 | 0.13522405 |
| 712,000 | 0.130213036 | 0.180443752 | 0.057415117 | 0.135381564 |
| 712,100 | 0.130206233 | 0.181106098 | 0.057395531 | 0.135029908 |
| 712,200 | 0.130196979 | 0.18071928 | 0.058174875 | 0.135235906 |
| 712,300 | 0.130178144 | 0.180459379 | 0.057934515 | 0.135134065 |
| 712,400 | 0.130271549 | 0.181421411 | 0.057599026 | 0.135854346 |
| 712,500 | 0.13016002 | 0.180830461 | 0.057232138 | 0.135473011 |
| 712,600 | 0.130180306 | 0.181487304 | 0.05718212 | 0.134429607 |
| 712,700 | 0.130226869 | 0.18123239 | 0.056877447 | 0.135450398 |
| 712,800 | 0.130108949 | 0.180971224 | 0.057601174 | 0.135182968 |
| 712,900 | 0.13017701 | 0.180743414 | 0.057098168 | 0.135127861 |
| 713,000 | 0.130074494 | 0.18026592 | 0.057159953 | 0.135200117 |
| 713,100 | 0.130028677 | 0.180435811 | 0.057180159 | 0.135133851 |
| 713,200 | 0.13009714 | 0.180318317 | 0.057207284 | 0.135514099 |
| 713,300 | 0.130080291 | 0.180194071 | 0.057466935 | 0.135329319 |
| 713,400 | 0.1300129 | 0.180684355 | 0.057270926 | 0.13532366 |
| 713,500 | 0.130130352 | 0.180279582 | 0.05740591 | 0.135743485 |
| 713,600 | 0.13004332 | 0.180457591 | 0.057654573 | 0.135672886 |
| 713,700 | 0.130038385 | 0.180287695 | 0.05753264 | 0.135549989 |
| 713,800 | 0.130136635 | 0.179858352 | 0.057057467 | 0.135775828 |
| 713,900 | 0.130125468 | 0.180285563 | 0.057239143 | 0.136086 |
| 714,000 | 0.130233155 | 0.180509206 | 0.057325453 | 0.135402481 |
| 714,100 | 0.130178875 | 0.18059036 | 0.05772826 | 0.135454634 |
| 714,200 | 0.130101033 | 0.180524209 | 0.058088913 | 0.136065156 |
| 714,300 | 0.130125933 | 0.180644094 | 0.058161183 | 0.136141007 |
| 714,400 | 0.130045864 | 0.180783266 | 0.057787329 | 0.136135527 |
| 714,500 | 0.130042115 | 0.180133531 | 0.058117974 | 0.136336559 |
| 714,600 | 0.129965349 | 0.180308059 | 0.057530107 | 0.135688245 |
| 714,700 | 0.130008835 | 0.180623625 | 0.057394795 | 0.135982019 |
| 714,800 | 0.129976101 | 0.180424561 | 0.057653879 | 0.135936003 |
| 714,900 | 0.130005752 | 0.180149815 | 0.05756119 | 0.135767852 |
| 715,000 | 0.129918719 | 0.180336097 | 0.057881748 | 0.135563679 |
| 715,100 | 0.129946662 | 0.180883501 | 0.057899086 | 0.135909593 |
| 715,200 | 0.130073021 | 0.180901535 | 0.057898912 | 0.135670122 |
| 715,300 | 0.130049818 | 0.180249682 | 0.058271317 | 0.13541262 |
| 715,400 | 0.130109266 | 0.180461783 | 0.0581074 | 0.135531684 |
| 715,500 | 0.130095568 | 0.180741907 | 0.057681903 | 0.135211813 |
| 715,600 | 0.130122321 | 0.180386462 | 0.058018828 | 0.135174758 |
| 715,700 | 0.130215809 | 0.180288955 | 0.058209291 | 0.135216797 |
| 715,800 | 0.130110917 | 0.180476791 | 0.058163094 | 0.135273456 |

| | | | | |
|---|---|---|---|---|
| 715,900 | 0.130099724 | 0.180375556 | 0.057941024 | 0.135518966 |
| 716,000 | 0.130051842 | 0.179956574 | 0.057769743 | 0.135346703 |
| 716,100 | 0.130131015 | 0.180629888 | 0.058091004 | 0.135077254 |
| 716,200 | 0.130043612 | 0.179431572 | 0.058165268 | 0.13519046 |
| 716,300 | 0.130051342 | 0.18072767 | 0.057761068 | 0.135509232 |
| 716,400 | 0.130069314 | 0.181079167 | 0.058722128 | 0.135211121 |
| 716,500 | 0.129977732 | 0.180552435 | 0.058105474 | 0.135805015 |
| 716,600 | 0.129976284 | 0.180330461 | 0.057985693 | 0.135024237 |
| 716,700 | 0.130023501 | 0.180320663 | 0.057810826 | 0.135362657 |
| 716,800 | 0.130019945 | 0.180922943 | 0.057475819 | 0.135346685 |
| 716,900 | 0.129951578 | 0.180564533 | 0.057810076 | 0.135604367 |
| 717,000 | 0.130045841 | 0.180346398 | 0.058014849 | 0.135298026 |
| 717,100 | 0.129891797 | 0.181065091 | 0.057871574 | 0.135574309 |
| 717,200 | 0.129965004 | 0.181291816 | 0.058181772 | 0.134840688 |
| 717,300 | 0.130028513 | 0.180801092 | 0.058384658 | 0.135828504 |
| 717,400 | 0.129986292 | 0.18057982 | 0.058409878 | 0.136134607 |
| 717,500 | 0.130008574 | 0.180354827 | 0.058029677 | 0.136186017 |
| 717,600 | 0.12998277 | 0.18044242 | 0.058108343 | 0.135620558 |
| 717,700 | 0.130051998 | 0.180055313 | 0.058152189 | 0.135541104 |
| 717,800 | 0.129865941 | 0.180130676 | 0.057992773 | 0.135396186 |
| 717,900 | 0.129884737 | 0.179866787 | 0.058263387 | 0.136328175 |
| 718,000 | 0.129839897 | 0.179500015 | 0.057698203 | 0.135782385 |
| 718,100 | 0.129780154 | 0.178958086 | 0.058017232 | 0.135543202 |
| 718,200 | 0.129781379 | 0.178932557 | 0.057591687 | 0.135755252 |
| 718,300 | 0.129789427 | 0.179455387 | 0.057543605 | 0.135447846 |
| 718,400 | 0.129832268 | 0.179831147 | 0.058047894 | 0.135063799 |
| 718,500 | 0.129695436 | 0.179697659 | 0.05783368 | 0.134913285 |
| 718,600 | 0.1298006 | 0.179469376 | 0.057651231 | 0.135408254 |
| 718,700 | 0.129654515 | 0.179651919 | 0.058113552 | 0.135197386 |
| 718,800 | 0.129767974 | 0.17958882 | 0.057999827 | 0.135519277 |
| 718,900 | 0.129855466 | 0.180286074 | 0.057843925 | 0.135875789 |
| 719,000 | 0.129769408 | 0.179471144 | 0.058233597 | 0.135923692 |
| 719,100 | 0.129804252 | 0.179140175 | 0.058078479 | 0.135414604 |
| 719,200 | 0.129670926 | 0.179456665 | 0.057647774 | 0.135231779 |
| 719,300 | 0.129689795 | 0.17952737 | 0.057416192 | 0.13526118 |
| 719,400 | 0.12972137 | 0.179548139 | 0.058134246 | 0.135375421 |
| 719,500 | 0.129688883 | 0.17943084 | 0.057770424 | 0.135253683 |
| 719,600 | 0.129582345 | 0.179801102 | 0.058001232 | 0.134875685 |
| 719,700 | 0.129554466 | 0.17963624 | 0.057654292 | 0.134802234 |
| 719,800 | 0.129730364 | 0.180107147 | 0.057444244 | 0.134737272 |
| 719,900 | 0.129665078 | 0.179512208 | 0.057643424 | 0.134593844 |
| 720,000 | 0.129606894 | 0.17924862 | 0.057999067 | 0.13455818 |
| 720,100 | 0.129481055 | 0.178120248 | 0.057523795 | 0.13457208 |
| 720,200 | 0.129610311 | 0.179561841 | 0.057855239 | 0.13492692 |
| 720,300 | 0.129564939 | 0.1792635 | 0.05780696 | 0.134271072 |
| 720,400 | 0.129579995 | 0.179542291 | 0.057075374 | 0.135386427 |
| 720,500 | 0.129668074 | 0.179627881 | 0.057940814 | 0.135926473 |
| 720,600 | 0.129596339 | 0.179072325 | 0.05785487 | 0.134941489 |
| 720,700 | 0.12941059 | 0.179056283 | 0.05814672 | 0.134349021 |
| 720,800 | 0.12951135 | 0.178846558 | 0.0578557 | 0.13452105 |
| 720,900 | 0.129558131 | 0.179233932 | 0.0572227 | 0.134861128 |
| 721,000 | 0.129462686 | 0.178963157 | 0.057723838 | 0.134974108 |
| 721,100 | 0.129356443 | 0.178935745 | 0.058101497 | 0.134948582 |
| 721,200 | 0.129375132 | 0.178040578 | 0.057417047 | 0.134948016 |
| 721,300 | 0.129348697 | 0.17841707 | 0.057665407 | 0.134228271 |
| 721,400 | 0.129435815 | 0.178645719 | 0.057394102 | 0.134132624 |
| 721,500 | 0.129400217 | 0.179171094 | 0.056811343 | 0.134089945 |
| 721,600 | 0.129371559 | 0.178390891 | 0.057305087 | 0.134015919 |
| 721,700 | 0.129298805 | 0.178193991 | 0.05701342 | 0.13419396 |
| 721,800 | 0.129282967 | 0.177946251 | 0.057145459 | 0.133993954 |
| 721,900 | 0.129162548 | 0.177530899 | 0.057460218 | 0.133825883 |
| 722,000 | 0.129240286 | 0.177623082 | 0.057618651 | 0.134468098 |
| 722,100 | 0.129262694 | 0.178660228 | 0.057822917 | 0.134841698 |
| 722,200 | 0.129388035 | 0.178301335 | 0.056974957 | 0.134498452 |
| 722,300 | 0.129323532 | 0.178509708 | 0.057589055 | 0.134634615 |
| 722,400 | 0.129253157 | 0.177941716 | 0.05717091 | 0.134243412 |
| 722,500 | 0.129254192 | 0.178343805 | 0.057094613 | 0.133869649 |

| | | | | |
|---|---|---|---|---|
| 722,600 | 0.129270337 | 0.178459805 | 0.056971726 | 0.134276302 |
| 722,700 | 0.129242029 | 0.178516287 | 0.057047371 | 0.134479585 |
| 722,800 | 0.129217904 | 0.178026122 | 0.056918235 | 0.133834024 |
| 722,900 | 0.129316249 | 0.177946001 | 0.056845385 | 0.133774532 |
| 723,000 | 0.129259623 | 0.178146347 | 0.057116033 | 0.134076756 |
| 723,100 | 0.129258671 | 0.178093427 | 0.056820632 | 0.133650359 |
| 723,200 | 0.129291041 | 0.177697196 | 0.056738233 | 0.134059262 |
| 723,300 | 0.129258156 | 0.177427032 | 0.056033269 | 0.134120202 |
| 723,400 | 0.129180509 | 0.177615072 | 0.056553519 | 0.134120421 |
| 723,500 | 0.129135682 | 0.177272249 | 0.056363911 | 0.134344746 |
| 723,600 | 0.129191655 | 0.177820517 | 0.056718252 | 0.133768883 |
| 723,700 | 0.129256784 | 0.177292692 | 0.056384823 | 0.133823584 |
| 723,800 | 0.129222699 | 0.177855465 | 0.056817098 | 0.133833363 |
| 723,900 | 0.129139969 | 0.176961744 | 0.056640501 | 0.132837213 |
| 724,000 | 0.12911592 | 0.1772593 | 0.057079147 | 0.133492433 |
| 724,100 | 0.12908275 | 0.177037113 | 0.056927576 | 0.13345157 |
| 724,200 | 0.129149011 | 0.177313112 | 0.056636592 | 0.133280532 |
| 724,300 | 0.129205648 | 0.17784325 | 0.056921945 | 0.133195641 |
| 724,400 | 0.129113916 | 0.177873293 | 0.056978509 | 0.13284835 |
| 724,500 | 0.129258437 | 0.177957749 | 0.057192701 | 0.133773951 |
| 724,600 | 0.129009103 | 0.177026043 | 0.056835242 | 0.133384524 |
| 724,700 | 0.129145844 | 0.177738402 | 0.057335885 | 0.133725978 |
| 724,800 | 0.129121206 | 0.177938904 | 0.057062774 | 0.133521475 |
| 724,900 | 0.129206767 | 0.178477288 | 0.056699056 | 0.133742666 |
| 725,000 | 0.129055324 | 0.178093315 | 0.057595348 | 0.133113715 |
| 725,100 | 0.129041774 | 0.178426153 | 0.05709251 | 0.133151909 |
| 725,200 | 0.129052556 | 0.177767587 | 0.057580711 | 0.133351329 |
| 725,300 | 0.129011093 | 0.177830295 | 0.05769986 | 0.133335938 |
| 725,400 | 0.129081372 | 0.17793481 | 0.057483408 | 0.133241037 |
| 725,500 | 0.128990061 | 0.177748539 | 0.057470512 | 0.133343855 |
| 725,600 | 0.12898575 | 0.177333087 | 0.057762234 | 0.133875906 |
| 725,700 | 0.128961918 | 0.17739026 | 0.057875476 | 0.1330641 |
| 725,800 | 0.12908041 | 0.178240998 | 0.057755274 | 0.133655237 |
| 725,900 | 0.128975474 | 0.177961764 | 0.057070451 | 0.133279825 |
| 726,000 | 0.129052519 | 0.17820933 | 0.057399325 | 0.133352647 |
| 726,100 | 0.128972745 | 0.178069846 | 0.057626004 | 0.133063806 |
| 726,200 | 0.128999352 | 0.17850135 | 0.057643847 | 0.133107076 |
| 726,300 | 0.129071295 | 0.178749698 | 0.057648921 | 0.132491182 |
| 726,400 | 0.128953023 | 0.178084693 | 0.057614265 | 0.133277234 |
| 726,500 | 0.128902079 | 0.178653004 | 0.057498543 | 0.133468769 |
| 726,600 | 0.128846386 | 0.178157409 | 0.057337498 | 0.13334023 |
| 726,700 | 0.128894592 | 0.177786477 | 0.057855194 | 0.133070849 |
| 726,800 | 0.128743663 | 0.178117786 | 0.057685533 | 0.132862716 |
| 726,900 | 0.128752766 | 0.177666814 | 0.057667098 | 0.133356183 |
| 727,000 | 0.128797571 | 0.178268972 | 0.057322867 | 0.132680996 |
| 727,100 | 0.12878301 | 0.17839212 | 0.056933745 | 0.132173987 |
| 727,200 | 0.128769505 | 0.178491755 | 0.056996503 | 0.132980671 |
| 727,300 | 0.128865971 | 0.177903518 | 0.057374787 | 0.13339432 |
| 727,400 | 0.128767187 | 0.178121817 | 0.057626623 | 0.132619681 |
| 727,500 | 0.128864851 | 0.178379325 | 0.05710298 | 0.132272651 |
| 727,600 | 0.128840898 | 0.1785287 | 0.056702301 | 0.1332762 |
| 727,700 | 0.128739875 | 0.177404496 | 0.056739999 | 0.132666126 |
| 727,800 | 0.128669292 | 0.178459356 | 0.056525978 | 0.132711087 |
| 727,900 | 0.128720221 | 0.178002671 | 0.056441276 | 0.132607573 |
| 728,000 | 0.128647825 | 0.178091301 | 0.056269267 | 0.133014009 |
| 728,100 | 0.128665028 | 0.178719238 | 0.056958264 | 0.13268798 |
| 728,200 | 0.128783783 | 0.178901641 | 0.056737737 | 0.132982083 |
| 728,300 | 0.128739718 | 0.17758333 | 0.056728895 | 0.133026362 |
| 728,400 | 0.128795256 | 0.178441472 | 0.056545375 | 0.132857843 |
| 728,500 | 0.128658096 | 0.178227785 | 0.05694326 | 0.132834141 |
| 728,600 | 0.128729293 | 0.178358453 | 0.057011887 | 0.133219924 |
| 728,700 | 0.128621072 | 0.178305579 | 0.056704945 | 0.132686136 |
| 728,800 | 0.128690117 | 0.178483699 | 0.057175111 | 0.132240752 |
| 728,900 | 0.128603448 | 0.17878652 | 0.056286689 | 0.132694858 |
| 729,000 | 0.128672401 | 0.178745649 | 0.057166039 | 0.132945033 |
| 729,100 | 0.128696286 | 0.178844313 | 0.056533873 | 0.132724767 |
| 729,200 | 0.128666746 | 0.179192672 | 0.057001565 | 0.133196492 |

| | | | | |
|---|---|---|---|---|
| 729,300 | 0.128729286 | 0.178660608 | 0.056533654 | 0.1334267 |
| 729,400 | 0.128822168 | 0.179368976 | 0.056766814 | 0.132922488 |
| 729,500 | 0.128713962 | 0.179071969 | 0.05667853 | 0.132787802 |
| 729,600 | 0.128835878 | 0.179743937 | 0.05628552 | 0.132830861 |
| 729,700 | 0.128633649 | 0.179054784 | 0.056371565 | 0.132434867 |
| 729,800 | 0.128685687 | 0.178601149 | 0.056882201 | 0.132654954 |
| 729,900 | 0.128671685 | 0.178710669 | 0.057001737 | 0.132714749 |
| 730,000 | 0.128700594 | 0.178730423 | 0.057197596 | 0.132484478 |
| 730,100 | 0.128761777 | 0.179050403 | 0.05675669 | 0.132038349 |
| 730,200 | 0.128768522 | 0.178794269 | 0.056956907 | 0.132342886 |
| 730,300 | 0.128655481 | 0.179077633 | 0.057406019 | 0.132396203 |
| 730,400 | 0.128574041 | 0.178728559 | 0.056430686 | 0.132463188 |
| 730,500 | 0.128686497 | 0.178665392 | 0.05650191 | 0.132567079 |
| 730,600 | 0.128673267 | 0.179516241 | 0.056299652 | 0.132207889 |
| 730,700 | 0.128621201 | 0.178576475 | 0.056523663 | 0.132673926 |
| 730,800 | 0.128703259 | 0.179108357 | 0.05707022 | 0.132250725 |
| 730,900 | 0.128556117 | 0.17887951 | 0.05737848 | 0.132241392 |
| 731,000 | 0.128596432 | 0.17874681 | 0.057171305 | 0.131994148 |
| 731,100 | 0.128643854 | 0.178648805 | 0.057003409 | 0.132234442 |
| 731,200 | 0.128553586 | 0.178936403 | 0.057228977 | 0.13183547 |
| 731,300 | 0.128619793 | 0.178762278 | 0.057138281 | 0.131428513 |
| 731,400 | 0.128507267 | 0.178259036 | 0.056901145 | 0.13163173 |
| 731,500 | 0.128485728 | 0.178051307 | 0.057750505 | 0.131562631 |
| 731,600 | 0.128475012 | 0.177904841 | 0.057649592 | 0.13224955 |
| 731,700 | 0.128440222 | 0.178354626 | 0.05704661 | 0.131781289 |
| 731,800 | 0.12845715 | 0.178412461 | 0.056995127 | 0.132029407 |
| 731,900 | 0.128476786 | 0.178026685 | 0.0571171 | 0.132312454 |
| 732,000 | 0.128409421 | 0.17826846 | 0.05720224 | 0.131889377 |
| 732,100 | 0.128470603 | 0.178254825 | 0.056986685 | 0.132406126 |
| 732,200 | 0.128523112 | 0.178352431 | 0.057584014 | 0.132015875 |
| 732,300 | 0.128615658 | 0.178090617 | 0.056855882 | 0.131926206 |
| 732,400 | 0.128643239 | 0.178659817 | 0.056824997 | 0.131946266 |
| 732,500 | 0.128512075 | 0.178609084 | 0.056525074 | 0.131825423 |
| 732,600 | 0.128443732 | 0.178258104 | 0.057214922 | 0.131555277 |
| 732,700 | 0.128494402 | 0.178275382 | 0.057124241 | 0.131791681 |
| 732,800 | 0.128493607 | 0.178478722 | 0.057751529 | 0.131705306 |
| 732,900 | 0.128434047 | 0.178313196 | 0.057513595 | 0.131648106 |
| 733,000 | 0.128470303 | 0.17829249 | 0.05766574 | 0.131492434 |
| 733,100 | 0.128443934 | 0.178339386 | 0.057559875 | 0.131182241 |
| 733,200 | 0.128512563 | 0.178526208 | 0.057501645 | 0.132106444 |
| 733,300 | 0.128582376 | 0.178656237 | 0.057839897 | 0.132101025 |
| 733,400 | 0.128512782 | 0.17857713 | 0.057773 | 0.132212135 |
| 733,500 | 0.128510721 | 0.178153653 | 0.058064829 | 0.132054971 |
| 733,600 | 0.128520018 | 0.178630339 | 0.057783988 | 0.132203075 |
| 733,700 | 0.128493463 | 0.179083377 | 0.058263195 | 0.132316405 |
| 733,800 | 0.128504544 | 0.178644629 | 0.058215762 | 0.132146588 |
| 733,900 | 0.128585742 | 0.178779364 | 0.057365399 | 0.132035735 |
| 734,000 | 0.128588337 | 0.178949122 | 0.058021719 | 0.132086471 |
| 734,100 | 0.128471934 | 0.177767493 | 0.058028202 | 0.131465312 |
| 734,200 | 0.128455563 | 0.177062802 | 0.057669061 | 0.132072061 |
| 734,300 | 0.128654547 | 0.178496053 | 0.058025826 | 0.131744134 |
| 734,400 | 0.128520566 | 0.177687281 | 0.058083499 | 0.131726977 |
| 734,500 | 0.128498197 | 0.178207144 | 0.058546934 | 0.131398734 |
| 734,600 | 0.12845036 | 0.178158858 | 0.058450862 | 0.131400127 |
| 734,700 | 0.128400397 | 0.178444942 | 0.057678924 | 0.131516697 |
| 734,800 | 0.128313744 | 0.178543371 | 0.058387644 | 0.131916157 |
| 734,900 | 0.12837246 | 0.178887278 | 0.058224259 | 0.131257643 |
| 735,000 | 0.128397893 | 0.178300463 | 0.058029445 | 0.1315385 |
| 735,100 | 0.128399583 | 0.178860849 | 0.058722915 | 0.131359762 |
| 735,200 | 0.128427539 | 0.179119558 | 0.058352268 | 0.131695579 |
| 735,300 | 0.128372116 | 0.178907896 | 0.058228388 | 0.131740716 |
| 735,400 | 0.128470377 | 0.178869426 | 0.058133048 | 0.131737888 |
| 735,500 | 0.12838573 | 0.179098953 | 0.058390579 | 0.131922909 |
| 735,600 | 0.12837091 | 0.178710611 | 0.058266038 | 0.131708527 |
| 735,700 | 0.128399857 | 0.178073029 | 0.058433279 | 0.131781061 |
| 735,800 | 0.128396286 | 0.178530674 | 0.057876767 | 0.131456366 |
| 735,900 | 0.12838731 | 0.178902674 | 0.057498284 | 0.131795414 |

| | | | | |
|---|---|---|---|---|
| 736,000 | 0.128377306 | 0.178822677 | 0.057360386 | 0.1321189 |
| 736,100 | 0.128414812 | 0.179230904 | 0.058056132 | 0.131925321 |
| 736,200 | 0.128379856 | 0.178481225 | 0.057416019 | 0.131757963 |
| 736,300 | 0.128337926 | 0.178268476 | 0.057537233 | 0.13209619 |
| 736,400 | 0.128391168 | 0.177950143 | 0.057315365 | 0.131935405 |
| 736,500 | 0.128309571 | 0.178381313 | 0.057638949 | 0.131491846 |
| 736,600 | 0.12841563 | 0.177961556 | 0.057591966 | 0.131954453 |
| 736,700 | 0.128281626 | 0.177464194 | 0.057836611 | 0.132686184 |
| 736,800 | 0.128289146 | 0.178424536 | 0.057241957 | 0.132007617 |
| 736,900 | 0.128404321 | 0.178193616 | 0.057256553 | 0.131688056 |
| 737,000 | 0.128226646 | 0.178361238 | 0.057318128 | 0.131690581 |
| 737,100 | 0.128231584 | 0.17788474 | 0.05757477 | 0.131857851 |
| 737,200 | 0.128152848 | 0.177458312 | 0.057792366 | 0.131791354 |
| 737,300 | 0.128298608 | 0.177956111 | 0.057901223 | 0.132370069 |
| 737,400 | 0.128171913 | 0.178027285 | 0.057647737 | 0.132123218 |
| 737,500 | 0.128276427 | 0.178563423 | 0.057927662 | 0.132272526 |
| 737,600 | 0.128223191 | 0.177972253 | 0.05732805 | 0.132428955 |
| 737,700 | 0.128220694 | 0.17805036 | 0.057199787 | 0.132498663 |
| 737,800 | 0.128306908 | 0.178047906 | 0.058187853 | 0.132189934 |
| 737,900 | 0.128187166 | 0.177389667 | 0.057539066 | 0.132573028 |
| 738,000 | 0.128169298 | 0.178040703 | 0.057443211 | 0.132878532 |
| 738,100 | 0.128236102 | 0.177292615 | 0.05729911 | 0.132424598 |
| 738,200 | 0.12822136 | 0.177673561 | 0.057594341 | 0.132068709 |
| 738,300 | 0.128140822 | 0.17682272 | 0.057533804 | 0.132267438 |
| 738,400 | 0.128074134 | 0.177209581 | 0.05747622 | 0.132228295 |
| 738,500 | 0.128140148 | 0.177042281 | 0.057830929 | 0.132631846 |
| 738,600 | 0.128147835 | 0.177125331 | 0.057710303 | 0.132011077 |
| 738,700 | 0.128185345 | 0.177459034 | 0.056874677 | 0.132297183 |
| 738,800 | 0.128120233 | 0.1766236 | 0.057351211 | 0.131954709 |
| 738,900 | 0.128188828 | 0.176947448 | 0.057098169 | 0.132308686 |
| 739,000 | 0.128164021 | 0.176982981 | 0.057234008 | 0.132188162 |
| 739,100 | 0.128271862 | 0.17755263 | 0.057208962 | 0.132329198 |
| 739,200 | 0.128112836 | 0.17793814 | 0.057069968 | 0.132238172 |
| 739,300 | 0.128117518 | 0.177690155 | 0.057677095 | 0.132595568 |
| 739,400 | 0.128016348 | 0.177490077 | 0.056580298 | 0.132266648 |
| 739,500 | 0.128066705 | 0.177780635 | 0.057125919 | 0.132431549 |
| 739,600 | 0.127927988 | 0.177369033 | 0.056695698 | 0.132123459 |
| 739,700 | 0.12803844 | 0.177178538 | 0.057174562 | 0.132180903 |
| 739,800 | 0.128059007 | 0.177858685 | 0.057338644 | 0.131457364 |
| 739,900 | 0.128091013 | 0.177399827 | 0.057231658 | 0.132284836 |
| 740,000 | 0.12816219 | 0.177564854 | 0.057265172 | 0.131741312 |
| 740,100 | 0.128048901 | 0.177622778 | 0.056752288 | 0.13233835 |
| 740,200 | 0.128052561 | 0.177972456 | 0.056984762 | 0.131993798 |
| 740,300 | 0.128034461 | 0.177938565 | 0.056954972 | 0.132497993 |
| 740,400 | 0.128003229 | 0.177083089 | 0.057396327 | 0.132624412 |
| 740,500 | 0.128094839 | 0.177732008 | 0.057239958 | 0.132176421 |
| 740,600 | 0.128003773 | 0.177885399 | 0.057075784 | 0.131945638 |
| 740,700 | 0.127939151 | 0.177843196 | 0.056946911 | 0.132362896 |
| 740,800 | 0.127995507 | 0.178047419 | 0.056938534 | 0.13232817 |
| 740,900 | 0.127828631 | 0.178013463 | 0.056621255 | 0.132117668 |
| 741,000 | 0.127862057 | 0.177473375 | 0.056792708 | 0.132058657 |
| 741,100 | 0.127976688 | 0.177869029 | 0.056866236 | 0.132735717 |
| 741,200 | 0.127844564 | 0.176946712 | 0.056811513 | 0.131869939 |
| 741,300 | 0.127905139 | 0.177133763 | 0.057083086 | 0.132195427 |
| 741,400 | 0.127995425 | 0.176828538 | 0.057238107 | 0.132142004 |
| 741,500 | 0.127854987 | 0.177484014 | 0.056506157 | 0.132612828 |
| 741,600 | 0.127707912 | 0.177396211 | 0.056008592 | 0.131841999 |
| 741,700 | 0.127916518 | 0.177511818 | 0.056605287 | 0.132058512 |
| 741,800 | 0.127798747 | 0.177110773 | 0.05663885 | 0.131713669 |
| 741,900 | 0.127845254 | 0.177419419 | 0.056834867 | 0.132192095 |
| 742,000 | 0.127989996 | 0.1778217 | 0.057318873 | 0.131989893 |
| 742,100 | 0.127923006 | 0.177923529 | 0.056865204 | 0.132361985 |
| 742,200 | 0.127866089 | 0.177920186 | 0.057842809 | 0.13212999 |
| 742,300 | 0.127834782 | 0.177585186 | 0.057543963 | 0.13179595 |
| 742,400 | 0.127895411 | 0.177602904 | 0.057109951 | 0.132097599 |
| 742,500 | 0.127901056 | 0.178108981 | 0.056723043 | 0.131905204 |
| 742,600 | 0.128031814 | 0.177935587 | 0.057076558 | 0.132174209 |

| | | | | |
|---|---|---|---|---|
| 742,700 | 0.127911402 | 0.177808096 | 0.057374568 | 0.131978366 |
| 742,800 | 0.127926267 | 0.177218167 | 0.058066483 | 0.132321444 |
| 742,900 | 0.127890299 | 0.176531163 | 0.057633082 | 0.13244396 |
| 743,000 | 0.12781632 | 0.177108439 | 0.057655809 | 0.132349332 |
| 743,100 | 0.12789731 | 0.17731998 | 0.057645686 | 0.132032154 |
| 743,200 | 0.12801032 | 0.176775263 | 0.057875914 | 0.132087788 |
| 743,300 | 0.127884378 | 0.176954967 | 0.058461671 | 0.132232742 |
| 743,400 | 0.127890928 | 0.17694073 | 0.058097059 | 0.132366016 |
| 743,500 | 0.127770911 | 0.177039802 | 0.057996513 | 0.13165391 |
| 743,600 | 0.127832226 | 0.176778153 | 0.05809642 | 0.131662139 |
| 743,700 | 0.127872745 | 0.17684307 | 0.057809505 | 0.131868006 |
| 743,800 | 0.127846292 | 0.176576184 | 0.057485384 | 0.132244962 |
| 743,900 | 0.127752333 | 0.176847627 | 0.057599462 | 0.131564806 |
| 744,000 | 0.127739349 | 0.176774234 | 0.058134714 | 0.132008953 |
| 744,100 | 0.127739823 | 0.176636088 | 0.058105 | 0.13220616 |
| 744,200 | 0.12774236 | 0.176967039 | 0.057712064 | 0.131627251 |
| 744,300 | 0.127669872 | 0.177206046 | 0.058214657 | 0.131842545 |
| 744,400 | 0.127690076 | 0.176637019 | 0.058080127 | 0.131527738 |
| 744,500 | 0.127745615 | 0.177006393 | 0.058124703 | 0.131458753 |
| 744,600 | 0.127693905 | 0.176807853 | 0.057978431 | 0.131264814 |
| 744,700 | 0.127718355 | 0.176824034 | 0.058086512 | 0.131279965 |
| 744,800 | 0.127724126 | 0.176663982 | 0.058072889 | 0.13159469 |
| 744,900 | 0.127656579 | 0.176414206 | 0.05746513 | 0.131997197 |
| 745,000 | 0.127630775 | 0.176380386 | 0.058164404 | 0.131543476 |
| 745,100 | 0.127617199 | 0.175590571 | 0.057744502 | 0.131554812 |
| 745,200 | 0.127663526 | 0.176468499 | 0.057892918 | 0.131748654 |
| 745,300 | 0.127651519 | 0.176636371 | 0.058189938 | 0.131553231 |
| 745,400 | 0.127791572 | 0.176317085 | 0.057916278 | 0.132262817 |
| 745,500 | 0.127672775 | 0.175956852 | 0.05789794 | 0.13189762 |
| 745,600 | 0.127809178 | 0.177167308 | 0.057250782 | 0.132455286 |
| 745,700 | 0.127732268 | 0.175984991 | 0.057381879 | 0.132096163 |
| 745,800 | 0.127695531 | 0.176686203 | 0.057078531 | 0.131961056 |
| 745,900 | 0.127740598 | 0.176900957 | 0.057259544 | 0.132426731 |
| 746,000 | 0.127677761 | 0.176498594 | 0.056873613 | 0.132571286 |
| 746,100 | 0.127672066 | 0.176665934 | 0.056721442 | 0.132477683 |
| 746,200 | 0.127669444 | 0.176994177 | 0.056325851 | 0.132674821 |
| 746,300 | 0.127567119 | 0.17650439 | 0.057270859 | 0.132203582 |
| 746,400 | 0.127608947 | 0.176235113 | 0.056815673 | 0.132883742 |
| 746,500 | 0.127629866 | 0.176533277 | 0.056875612 | 0.132206127 |
| 746,600 | 0.127578664 | 0.176425458 | 0.056731302 | 0.132069772 |
| 746,700 | 0.127467947 | 0.176125394 | 0.057219261 | 0.13215941 |
| 746,800 | 0.12751532 | 0.176340441 | 0.056606976 | 0.132691486 |
| 746,900 | 0.127481923 | 0.176286609 | 0.057135703 | 0.132826898 |
| 747,000 | 0.127425652 | 0.176010104 | 0.056934106 | 0.132534864 |
| 747,100 | 0.127452072 | 0.175665596 | 0.056833743 | 0.132912927 |
| 747,200 | 0.12742969 | 0.17571938 | 0.057052615 | 0.13194485 |
| 747,300 | 0.127445402 | 0.176393784 | 0.056333753 | 0.132355037 |
| 747,400 | 0.127346648 | 0.175968827 | 0.05670835 | 0.132181097 |
| 747,500 | 0.127413746 | 0.175997251 | 0.056523519 | 0.132282428 |
| 747,600 | 0.127267229 | 0.175673059 | 0.056369477 | 0.13207094 |
| 747,700 | 0.127379342 | 0.175693877 | 0.056606934 | 0.132475267 |
| 747,800 | 0.127482146 | 0.175495072 | 0.056677854 | 0.132444991 |
| 747,900 | 0.127257552 | 0.175395348 | 0.056847371 | 0.132520482 |
| 748,000 | 0.127300187 | 0.176081512 | 0.056236811 | 0.131978162 |
| 748,100 | 0.127395988 | 0.176032345 | 0.056241348 | 0.132372081 |
| 748,200 | 0.127327138 | 0.175872604 | 0.056075293 | 0.131951213 |
| 748,300 | 0.127290017 | 0.175707547 | 0.056192474 | 0.132573502 |
| 748,400 | 0.127288437 | 0.176595383 | 0.055647073 | 0.132477791 |
| 748,500 | 0.127386907 | 0.176287976 | 0.055985947 | 0.131955546 |
| 748,600 | 0.12734209 | 0.176564267 | 0.056276543 | 0.132596837 |
| 748,700 | 0.127374521 | 0.176631943 | 0.056047194 | 0.132373983 |
| 748,800 | 0.12744918 | 0.17702969 | 0.056395051 | 0.132156003 |
| 748,900 | 0.127464355 | 0.176382946 | 0.056419254 | 0.132339053 |
| 749,000 | 0.127427358 | 0.176221969 | 0.056696875 | 0.13223766 |
| 749,100 | 0.127455165 | 0.176983073 | 0.056593977 | 0.133074362 |
| 749,200 | 0.127418094 | 0.176197569 | 0.055934636 | 0.132833889 |
| 749,300 | 0.127466582 | 0.176877545 | 0.056818743 | 0.132464583 |

| | | | | |
|---|---|---|---|---|
| 749,400 | 0.127351246 | 0.17656172 | 0.056097304 | 0.132326466 |
| 749,500 | 0.127386826 | 0.176938919 | 0.055726806 | 0.132693257 |
| 749,600 | 0.127361949 | 0.17700251 | 0.055979607 | 0.133044924 |
| 749,700 | 0.127329745 | 0.177047418 | 0.056436506 | 0.132397223 |
| 749,800 | 0.127357136 | 0.176666269 | 0.055937149 | 0.132199351 |
| 749,900 | 0.127306665 | 0.176587049 | 0.056167923 | 0.132700205 |
| 750,000 | 0.127267294 | 0.177048302 | 0.055794951 | 0.132733906 |
| 750,100 | 0.12731917 | 0.176823461 | 0.055535445 | 0.133178482 |
| 750,200 | 0.127352449 | 0.177014339 | 0.056219651 | 0.133041519 |
| 750,300 | 0.127418363 | 0.176896625 | 0.056460387 | 0.133572834 |
| 750,400 | 0.127315078 | 0.176351054 | 0.055883699 | 0.133281651 |
| 750,500 | 0.127254895 | 0.176520558 | 0.055874865 | 0.132822078 |
| 750,600 | 0.127293795 | 0.176866922 | 0.055803335 | 0.133242058 |
| 750,700 | 0.127224423 | 0.177183941 | 0.056309711 | 0.133579814 |
| 750,800 | 0.127222087 | 0.176813352 | 0.05594174 | 0.133230249 |
| 750,900 | 0.127194237 | 0.176767537 | 0.055387097 | 0.133064416 |
| 751,000 | 0.127151702 | 0.176223779 | 0.055395424 | 0.13345218 |
| 751,100 | 0.127166845 | 0.176465252 | 0.056015556 | 0.134194426 |
| 751,200 | 0.127162118 | 0.175880386 | 0.055514701 | 0.133610067 |
| 751,300 | 0.127089747 | 0.176449132 | 0.055352095 | 0.132929998 |
| 751,400 | 0.12710046 | 0.176161185 | 0.055177019 | 0.132876289 |
| 751,500 | 0.127154015 | 0.176514512 | 0.055193411 | 0.13248571 |
| 751,600 | 0.127188077 | 0.176590885 | 0.055756272 | 0.133255377 |
| 751,700 | 0.12719723 | 0.176999174 | 0.055935111 | 0.133560871 |
| 751,800 | 0.127193099 | 0.176543417 | 0.055616974 | 0.133618264 |
| 751,900 | 0.127141505 | 0.176335321 | 0.05561871 | 0.133506282 |
| 752,000 | 0.127210157 | 0.176602341 | 0.055684797 | 0.133620914 |
| 752,100 | 0.127149708 | 0.176530757 | 0.055011612 | 0.13397451 |
| 752,200 | 0.127138079 | 0.176942422 | 0.054921465 | 0.133158022 |
| 752,300 | 0.127275145 | 0.177444762 | 0.05569726 | 0.133551665 |
| 752,400 | 0.127216599 | 0.177695269 | 0.055462639 | 0.13360965 |
| 752,500 | 0.127113206 | 0.177121873 | 0.055284384 | 0.132836962 |
| 752,600 | 0.127165136 | 0.177065647 | 0.055377618 | 0.132835886 |
| 752,700 | 0.127124276 | 0.177067975 | 0.054985664 | 0.133023383 |
| 752,800 | 0.126961621 | 0.176058271 | 0.055273724 | 0.133104374 |
| 752,900 | 0.127005387 | 0.177065694 | 0.055922699 | 0.132831574 |
| 753,000 | 0.126982486 | 0.176155824 | 0.055906561 | 0.133167167 |
| 753,100 | 0.126974228 | 0.176450902 | 0.054822649 | 0.133350259 |
| 753,200 | 0.126987328 | 0.176669986 | 0.054672118 | 0.133062156 |
| 753,300 | 0.127070504 | 0.176687317 | 0.054901782 | 0.133394527 |
| 753,400 | 0.1271211 | 0.177556764 | 0.054633781 | 0.133461604 |
| 753,500 | 0.127075787 | 0.177461447 | 0.054870114 | 0.133564419 |
| 753,600 | 0.1268965 | 0.176498119 | 0.055144689 | 0.133188416 |
| 753,700 | 0.126967475 | 0.17691529 | 0.054725722 | 0.132634044 |
| 753,800 | 0.12699771 | 0.176425049 | 0.05513929 | 0.132769187 |
| 753,900 | 0.126874959 | 0.176803297 | 0.054363507 | 0.133411648 |
| 754,000 | 0.127022228 | 0.176063964 | 0.054358818 | 0.133419977 |
| 754,100 | 0.126894629 | 0.176073796 | 0.054407852 | 0.133183452 |
| 754,200 | 0.126954346 | 0.176582595 | 0.05480713 | 0.13297601 |
| 754,300 | 0.126801585 | 0.176516753 | 0.054944387 | 0.132616925 |
| 754,400 | 0.126824481 | 0.175213242 | 0.054598766 | 0.133476473 |
| 754,500 | 0.126758039 | 0.175910822 | 0.054468745 | 0.13269983 |
| 754,600 | 0.126833088 | 0.176201478 | 0.055275389 | 0.132817537 |
| 754,700 | 0.126759488 | 0.17616515 | 0.054180849 | 0.132532366 |
| 754,800 | 0.12679679 | 0.176009121 | 0.054000045 | 0.132719559 |
| 754,900 | 0.126921986 | 0.175683451 | 0.054092486 | 0.132633583 |
| 755,000 | 0.126869468 | 0.17581181 | 0.054317946 | 0.132263844 |
| 755,100 | 0.126869102 | 0.175681222 | 0.054277826 | 0.132389263 |
| 755,200 | 0.126892069 | 0.175891367 | 0.05417707 | 0.132437769 |
| 755,300 | 0.12675768 | 0.176397095 | 0.054490791 | 0.131702854 |
| 755,400 | 0.1268172 | 0.176256939 | 0.054349056 | 0.132229718 |
| 755,500 | 0.126779459 | 0.175670847 | 0.053982917 | 0.132467549 |
| 755,600 | 0.126829427 | 0.176055833 | 0.053983173 | 0.132387017 |
| 755,700 | 0.126746051 | 0.175663261 | 0.053549769 | 0.131583879 |
| 755,800 | 0.126739107 | 0.176333871 | 0.054534868 | 0.131894251 |
| 755,900 | 0.126684417 | 0.175483767 | 0.054468195 | 0.132422327 |
| 756,000 | 0.126714091 | 0.175711825 | 0.05374245 | 0.132201738 |

| | | | | |
|---|---|---|---|---|
| 756,100 | 0.126684157 | 0.175211004 | 0.053786124 | 0.131764256 |
| 756,200 | 0.126579414 | 0.175385753 | 0.053610823 | 0.131788652 |
| 756,300 | 0.12666444 | 0.175157801 | 0.053902489 | 0.132019793 |
| 756,400 | 0.126602891 | 0.175682975 | 0.053843183 | 0.132396297 |
| 756,500 | 0.126689639 | 0.176248944 | 0.053973432 | 0.132130551 |
| 756,600 | 0.126565256 | 0.175605845 | 0.053873348 | 0.132116925 |
| 756,700 | 0.12664361 | 0.174672969 | 0.054309028 | 0.132243819 |
| 756,800 | 0.126631385 | 0.174645436 | 0.054741159 | 0.132250201 |
| 756,900 | 0.126561641 | 0.175325849 | 0.054153479 | 0.132002609 |
| 757,000 | 0.126554302 | 0.175158269 | 0.053555621 | 0.132216437 |
| 757,100 | 0.126614614 | 0.175309248 | 0.053752048 | 0.131649832 |
| 757,200 | 0.126609305 | 0.175347977 | 0.053424821 | 0.131764603 |
| 757,300 | 0.126561253 | 0.174993595 | 0.053792592 | 0.131483295 |
| 757,400 | 0.126547955 | 0.17525008 | 0.053916205 | 0.13159351 |
| 757,500 | 0.12649801 | 0.17531226 | 0.054139878 | 0.131888913 |
| 757,600 | 0.126497924 | 0.17536575 | 0.053897538 | 0.13201689 |
| 757,700 | 0.12648022 | 0.175370386 | 0.05368242 | 0.131653997 |
| 757,800 | 0.126466243 | 0.175318146 | 0.054042192 | 0.131280108 |
| 757,900 | 0.126521769 | 0.174918627 | 0.054407321 | 0.131399449 |
| 758,000 | 0.126367556 | 0.174539265 | 0.053916604 | 0.131457255 |
| 758,100 | 0.126478813 | 0.174949375 | 0.054106363 | 0.131622917 |
| 758,200 | 0.126283385 | 0.174347717 | 0.05409026 | 0.131143907 |
| 758,300 | 0.126348138 | 0.174784794 | 0.053685437 | 0.131489208 |
| 758,400 | 0.126165776 | 0.173887112 | 0.053793312 | 0.131324187 |
| 758,500 | 0.126342877 | 0.175146529 | 0.054183991 | 0.131182813 |
| 758,600 | 0.126313009 | 0.174303626 | 0.054013983 | 0.131309507 |
| 758,700 | 0.126424978 | 0.17472221 | 0.053953791 | 0.130963465 |
| 758,800 | 0.126321273 | 0.174669787 | 0.053513283 | 0.132017438 |
| 758,900 | 0.126328432 | 0.174788603 | 0.053542784 | 0.131928676 |
| 759,000 | 0.126415252 | 0.175465551 | 0.053557435 | 0.131517667 |
| 759,100 | 0.126360575 | 0.174692351 | 0.053627715 | 0.131792753 |
| 759,200 | 0.126255307 | 0.17405101 | 0.053819027 | 0.131677771 |
| 759,300 | 0.126391342 | 0.175266328 | 0.053855571 | 0.131410013 |
| 759,400 | 0.126441757 | 0.174936968 | 0.053642089 | 0.131740702 |
| 759,500 | 0.126456071 | 0.175011139 | 0.053335867 | 0.131400803 |
| 759,600 | 0.12637843 | 0.17416572 | 0.0536519 | 0.131444546 |
| 759,700 | 0.126318215 | 0.173870876 | 0.05312247 | 0.131293651 |
| 759,800 | 0.126328181 | 0.174265568 | 0.053160478 | 0.13089084 |
| 759,900 | 0.126289212 | 0.174238784 | 0.053561979 | 0.130660583 |
| 760,000 | 0.126342493 | 0.174144965 | 0.053473368 | 0.130958599 |
| 760,100 | 0.126302075 | 0.174300878 | 0.052842583 | 0.130939939 |
| 760,200 | 0.126204091 | 0.173528776 | 0.053026437 | 0.131384513 |
| 760,300 | 0.126258424 | 0.174042035 | 0.053396295 | 0.13154135 |
| 760,400 | 0.126227232 | 0.173741063 | 0.053233011 | 0.131716522 |
| 760,500 | 0.126203025 | 0.173094896 | 0.053375232 | 0.13225885 |
| 760,600 | 0.12615588 | 0.173046575 | 0.053054033 | 0.131510822 |
| 760,700 | 0.12614918 | 0.173172099 | 0.053076176 | 0.131812787 |
| 760,800 | 0.126279526 | 0.173818268 | 0.053203283 | 0.131465095 |
| 760,900 | 0.126285381 | 0.173502798 | 0.053681464 | 0.131402827 |
| 761,000 | 0.126260348 | 0.173127274 | 0.053214912 | 0.131370245 |
| 761,100 | 0.126281811 | 0.173552303 | 0.053049852 | 0.131409504 |
| 761,200 | 0.126279625 | 0.173295895 | 0.052682549 | 0.131891464 |
| 761,300 | 0.126480491 | 0.173145099 | 0.053435606 | 0.131773409 |
| 761,400 | 0.12637292 | 0.173626511 | 0.053714212 | 0.131856431 |
| 761,500 | 0.126273013 | 0.173961872 | 0.053158388 | 0.13204248 |
| 761,600 | 0.126355666 | 0.174087451 | 0.053234525 | 0.132733704 |
| 761,700 | 0.126234681 | 0.173976135 | 0.05273096 | 0.132356679 |
| 761,800 | 0.126194856 | 0.173500341 | 0.053567687 | 0.132318147 |
| 761,900 | 0.126124663 | 0.173453133 | 0.053189837 | 0.131967994 |
| 762,000 | 0.126279996 | 0.174121116 | 0.053275496 | 0.131634562 |
| 762,100 | 0.126175555 | 0.173940572 | 0.053453511 | 0.131611157 |
| 762,200 | 0.126194988 | 0.173539573 | 0.053260337 | 0.131515069 |
| 762,300 | 0.125940724 | 0.173485844 | 0.053574666 | 0.131521185 |
| 762,400 | 0.126092441 | 0.173977989 | 0.053381247 | 0.131450546 |
| 762,500 | 0.126132095 | 0.173436498 | 0.053294275 | 0.1315629 |
| 762,600 | 0.126096546 | 0.17364175 | 0.052953393 | 0.131670799 |
| 762,700 | 0.126057365 | 0.172806501 | 0.052943816 | 0.131254418 |

| | | | | |
|---|---|---|---|---|
| 762,800 | 0.126190008 | 0.173224039 | 0.052997796 | 0.131366247 |
| 762,900 | 0.126050992 | 0.173106927 | 0.052883674 | 0.131066548 |
| 763,000 | 0.126148268 | 0.173213187 | 0.052447922 | 0.13108558 |
| 763,100 | 0.126060162 | 0.173189537 | 0.052509051 | 0.131579668 |
| 763,200 | 0.125961064 | 0.173394232 | 0.053058784 | 0.131539618 |
| 763,300 | 0.12606635 | 0.173431818 | 0.05273132 | 0.131284985 |
| 763,400 | 0.126053484 | 0.173081774 | 0.052231984 | 0.13108991 |
| 763,500 | 0.126055134 | 0.173367173 | 0.052284098 | 0.131306175 |
| 763,600 | 0.126038319 | 0.173546339 | 0.052722132 | 0.130826082 |
| 763,700 | 0.126068331 | 0.173222781 | 0.052498005 | 0.131193919 |
| 763,800 | 0.125954463 | 0.173292692 | 0.052439565 | 0.130590168 |
| 763,900 | 0.126161212 | 0.173841737 | 0.052523376 | 0.131064041 |
| 764,000 | 0.126165325 | 0.173111276 | 0.052437797 | 0.131319486 |
| 764,100 | 0.126152565 | 0.173796243 | 0.052524746 | 0.131071745 |
| 764,200 | 0.126086192 | 0.173930373 | 0.052687536 | 0.131312016 |
| 764,300 | 0.126092387 | 0.173524147 | 0.053207021 | 0.131244428 |
| 764,400 | 0.126125894 | 0.173121579 | 0.052648283 | 0.131447602 |
| 764,500 | 0.125997641 | 0.173510325 | 0.052193743 | 0.130841013 |
| 764,600 | 0.126061195 | 0.173621831 | 0.052720914 | 0.131181115 |
| 764,700 | 0.126091936 | 0.173267516 | 0.052708415 | 0.131424895 |
| 764,800 | 0.126058425 | 0.173090569 | 0.052353912 | 0.131775524 |
| 764,900 | 0.126077841 | 0.173405193 | 0.052388125 | 0.131280308 |
| 765,000 | 0.126083464 | 0.17371467 | 0.052503689 | 0.131662388 |
| 765,100 | 0.126078205 | 0.173609759 | 0.052252907 | 0.13145653 |
| 765,200 | 0.125989752 | 0.172998754 | 0.052636896 | 0.131531226 |
| 765,300 | 0.125930331 | 0.173563233 | 0.052495659 | 0.130825552 |
| 765,400 | 0.126034567 | 0.173258235 | 0.052448915 | 0.131447804 |
| 765,500 | 0.125947896 | 0.173481883 | 0.05247571 | 0.131203456 |
| 765,600 | 0.12587428 | 0.172761841 | 0.052875036 | 0.131575415 |
| 765,700 | 0.125925955 | 0.173309186 | 0.052777255 | 0.131150747 |
| 765,800 | 0.125946301 | 0.17337793 | 0.052426501 | 0.130921234 |
| 765,900 | 0.125844361 | 0.173136584 | 0.052694565 | 0.130970721 |
| 766,000 | 0.125843649 | 0.173161808 | 0.052492206 | 0.131657713 |
| 766,100 | 0.125860528 | 0.173062958 | 0.052660163 | 0.131436911 |
| 766,200 | 0.125722591 | 0.172880122 | 0.052449184 | 0.131336016 |
| 766,300 | 0.125712242 | 0.172965935 | 0.052804994 | 0.130718647 |
| 766,400 | 0.125810995 | 0.173381491 | 0.052488548 | 0.130928512 |
| 766,500 | 0.125766668 | 0.173451706 | 0.052164473 | 0.131021562 |
| 766,600 | 0.125713956 | 0.173344724 | 0.052102519 | 0.131028711 |
| 766,700 | 0.12579221 | 0.172708538 | 0.052262561 | 0.130928888 |
| 766,800 | 0.1256854 | 0.172350277 | 0.05265846 | 0.131259503 |
| 766,900 | 0.125663071 | 0.172486752 | 0.051452892 | 0.131086635 |
| 767,000 | 0.125694948 | 0.172322182 | 0.051581556 | 0.131497251 |
| 767,100 | 0.125711722 | 0.172063533 | 0.05203611 | 0.131130283 |
| 767,200 | 0.125751213 | 0.172295826 | 0.052606061 | 0.131639959 |
| 767,300 | 0.125773879 | 0.17267526 | 0.052138602 | 0.130971617 |
| 767,400 | 0.125686072 | 0.172554217 | 0.052259041 | 0.131446529 |
| 767,500 | 0.125748243 | 0.172870424 | 0.052300132 | 0.131311583 |
| 767,600 | 0.125773183 | 0.172512268 | 0.051720246 | 0.131017157 |
| 767,700 | 0.125543911 | 0.172104576 | 0.052544702 | 0.130774998 |
| 767,800 | 0.125704606 | 0.172223947 | 0.052409915 | 0.131232645 |
| 767,900 | 0.125794675 | 0.172436595 | 0.052972533 | 0.131725959 |
| 768,000 | 0.125749745 | 0.172123916 | 0.052073861 | 0.131236376 |
| 768,100 | 0.125621281 | 0.172290349 | 0.051938854 | 0.130512527 |
| 768,200 | 0.125727134 | 0.172644807 | 0.052302281 | 0.130252729 |
| 768,300 | 0.125681721 | 0.172727701 | 0.052392145 | 0.130749673 |
| 768,400 | 0.125529641 | 0.171999988 | 0.052736434 | 0.130426582 |
| 768,500 | 0.125694548 | 0.172689078 | 0.052255226 | 0.130317319 |
| 768,600 | 0.125568633 | 0.172212597 | 0.052000995 | 0.130905945 |
| 768,700 | 0.125659341 | 0.172527031 | 0.052232584 | 0.130697283 |
| 768,800 | 0.12564109 | 0.172037132 | 0.052367445 | 0.130982602 |
| 768,900 | 0.125583 | 0.172177113 | 0.052203771 | 0.130441427 |
| 769,000 | 0.125574676 | 0.172067767 | 0.052347647 | 0.13057738 |
| 769,100 | 0.125455054 | 0.172586852 | 0.052393977 | 0.130028047 |
| 769,200 | 0.125474176 | 0.172249855 | 0.052372019 | 0.130147647 |
| 769,300 | 0.125530695 | 0.172210271 | 0.052355463 | 0.130548214 |
| 769,400 | 0.125446548 | 0.171927791 | 0.052542026 | 0.130113915 |

| | | | | |
|---|---|---|---|---|
| 769,500 | 0.125402682 | 0.172682657 | 0.052768911 | 0.12986633 |
| 769,600 | 0.125360273 | 0.171555817 | 0.052490235 | 0.129972229 |
| 769,700 | 0.125506388 | 0.17195013 | 0.051984644 | 0.130408088 |
| 769,800 | 0.125364333 | 0.172284201 | 0.052643892 | 0.13013385 |
| 769,900 | 0.125327144 | 0.171606036 | 0.052400765 | 0.129430466 |
| 770,000 | 0.125403061 | 0.17189227 | 0.052033548 | 0.129723186 |
| 770,100 | 0.125355391 | 0.171898102 | 0.05214009 | 0.130034671 |
| 770,200 | 0.125409374 | 0.171461518 | 0.052170021 | 0.130182809 |
| 770,300 | 0.125349254 | 0.172441917 | 0.052224554 | 0.129599409 |
| 770,400 | 0.125328711 | 0.17206303 | 0.052175672 | 0.12994086 |
| 770,500 | 0.125235481 | 0.172162867 | 0.05236572 | 0.129486003 |
| 770,600 | 0.125191739 | 0.172230092 | 0.052296565 | 0.129366167 |
| 770,700 | 0.125148071 | 0.171420854 | 0.051857226 | 0.129386619 |
| 770,800 | 0.12526501 | 0.172044141 | 0.052526599 | 0.129492144 |
| 770,900 | 0.125215146 | 0.171829348 | 0.05289786 | 0.129512739 |
| 771,000 | 0.125269794 | 0.171506103 | 0.052957143 | 0.129884629 |
| 771,100 | 0.125235788 | 0.171948365 | 0.052829129 | 0.129363447 |
| 771,200 | 0.125108942 | 0.171537318 | 0.053113296 | 0.129299378 |
| 771,300 | 0.12511153 | 0.171770936 | 0.053126031 | 0.129680757 |
| 771,400 | 0.125322598 | 0.171774509 | 0.052675202 | 0.129862496 |
| 771,500 | 0.12516641 | 0.171485803 | 0.052881306 | 0.129951156 |
| 771,600 | 0.125135624 | 0.172014463 | 0.053532647 | 0.129751615 |
| 771,700 | 0.125072974 | 0.171960996 | 0.053087027 | 0.129531597 |
| 771,800 | 0.125088853 | 0.172208315 | 0.052614177 | 0.129283808 |
| 771,900 | 0.125143909 | 0.171779163 | 0.052510216 | 0.129543648 |
| 772,000 | 0.12515314 | 0.171864431 | 0.05249171 | 0.1296142 |
| 772,100 | 0.125113525 | 0.172232068 | 0.053004937 | 0.12977167 |
| 772,200 | 0.125113229 | 0.171311733 | 0.053364599 | 0.129531268 |
| 772,300 | 0.124975533 | 0.172288624 | 0.053309472 | 0.129212785 |
| 772,400 | 0.125015683 | 0.171014752 | 0.052459026 | 0.129724519 |
| 772,500 | 0.125074437 | 0.170685267 | 0.052892302 | 0.129768794 |
| 772,600 | 0.125008241 | 0.17103299 | 0.053024952 | 0.129299343 |
| 772,700 | 0.125122691 | 0.171465867 | 0.053042054 | 0.129546307 |
| 772,800 | 0.125024581 | 0.171721267 | 0.053001557 | 0.12970459 |
| 772,900 | 0.124982345 | 0.171627026 | 0.0527984 | 0.129359515 |
| 773,000 | 0.125058442 | 0.171447342 | 0.053183244 | 0.129205271 |
| 773,100 | 0.124977632 | 0.171534147 | 0.052852975 | 0.129849266 |
| 773,200 | 0.124952119 | 0.171762993 | 0.053146238 | 0.128855596 |
| 773,300 | 0.125052113 | 0.171016277 | 0.052955568 | 0.129423442 |
| 773,400 | 0.124973031 | 0.170458334 | 0.053085805 | 0.129373226 |
| 773,500 | 0.124968098 | 0.170832871 | 0.052962707 | 0.130090571 |
| 773,600 | 0.124958639 | 0.170690121 | 0.052806845 | 0.129431741 |
| 773,700 | 0.12497028 | 0.170769071 | 0.052718696 | 0.129469155 |
| 773,800 | 0.124809442 | 0.170619088 | 0.053035024 | 0.129853364 |
| 773,900 | 0.124688149 | 0.170095669 | 0.052734741 | 0.129393167 |
| 774,000 | 0.124761655 | 0.17022884 | 0.052929515 | 0.129505273 |
| 774,100 | 0.124697835 | 0.170423682 | 0.052511329 | 0.12916533 |
| 774,200 | 0.124803345 | 0.170239697 | 0.052618976 | 0.129387233 |
| 774,300 | 0.124842814 | 0.17027357 | 0.052140164 | 0.129524175 |
| 774,400 | 0.124821362 | 0.170277783 | 0.052522345 | 0.12903448 |
| 774,500 | 0.124790439 | 0.170270222 | 0.052553448 | 0.128839901 |
| 774,600 | 0.124608121 | 0.169931037 | 0.052467361 | 0.129104298 |
| 774,700 | 0.124834116 | 0.170247794 | 0.052573667 | 0.129208885 |
| 774,800 | 0.124716678 | 0.169524572 | 0.052572365 | 0.129300998 |
| 774,900 | 0.124589735 | 0.169806896 | 0.052774739 | 0.129455851 |
| 775,000 | 0.124625815 | 0.169739597 | 0.052134492 | 0.129010185 |
| 775,100 | 0.124628451 | 0.169834573 | 0.052328686 | 0.129842332 |
| 775,200 | 0.124699272 | 0.170208893 | 0.052471832 | 0.129853145 |
| 775,300 | 0.124735743 | 0.170879494 | 0.05241225 | 0.129386048 |
| 775,400 | 0.124696294 | 0.170229093 | 0.052950811 | 0.129200407 |
| 775,500 | 0.124627647 | 0.170374635 | 0.052206982 | 0.129279019 |
| 775,600 | 0.124590426 | 0.170736164 | 0.05204977 | 0.128548815 |
| 775,700 | 0.124701221 | 0.170641299 | 0.051992525 | 0.129472304 |
| 775,800 | 0.124651848 | 0.170379323 | 0.05195726 | 0.129267834 |
| 775,900 | 0.124649586 | 0.17107053 | 0.051922521 | 0.128820702 |
| 776,000 | 0.124595897 | 0.171083411 | 0.051848546 | 0.129656557 |
| 776,100 | 0.124564591 | 0.170835431 | 0.051431836 | 0.128930711 |

| | | | | |
|---|---|---|---|---|
| 776,200 | 0.124569893 | 0.170289885 | 0.051859034 | 0.129049252 |
| 776,300 | 0.124575245 | 0.170212897 | 0.051701405 | 0.129191096 |
| 776,400 | 0.124674393 | 0.170916157 | 0.05187457 | 0.129490087 |
| 776,500 | 0.12466404 | 0.170637579 | 0.052150552 | 0.129638101 |
| 776,600 | 0.124615777 | 0.170837777 | 0.052241361 | 0.129567774 |
| 776,700 | 0.124643063 | 0.170317272 | 0.052173267 | 0.129927858 |
| 776,800 | 0.124545566 | 0.17152323 | 0.052001746 | 0.129268462 |
| 776,900 | 0.124646285 | 0.171111006 | 0.051740026 | 0.129451888 |
| 777,000 | 0.124597882 | 0.170454629 | 0.052556804 | 0.129547685 |
| 777,100 | 0.124730465 | 0.171055459 | 0.052764458 | 0.129632008 |
| 777,200 | 0.12470766 | 0.170593637 | 0.052721677 | 0.130298462 |
| 777,300 | 0.124687151 | 0.170807606 | 0.05269482 | 0.13025893 |
| 777,400 | 0.124749068 | 0.170783842 | 0.052290313 | 0.130158442 |
| 777,500 | 0.124711227 | 0.170498101 | 0.051955177 | 0.129860848 |
| 777,600 | 0.124530983 | 0.169751903 | 0.052680666 | 0.129211888 |
| 777,700 | 0.12455289 | 0.170022917 | 0.052213446 | 0.129545846 |
| 777,800 | 0.124579199 | 0.170758474 | 0.053167427 | 0.129290148 |
| 777,900 | 0.124558545 | 0.170223999 | 0.052207775 | 0.129258831 |
| 778,000 | 0.124535108 | 0.170297297 | 0.052584687 | 0.129227433 |
| 778,100 | 0.12458615 | 0.170547803 | 0.053101158 | 0.12947002 |
| 778,200 | 0.124481463 | 0.170496305 | 0.052796129 | 0.129443185 |
| 778,300 | 0.124718574 | 0.171050378 | 0.053252859 | 0.12955223 |
| 778,400 | 0.124560733 | 0.170034666 | 0.053200894 | 0.129276956 |
| 778,500 | 0.124584767 | 0.17001611 | 0.053012129 | 0.129134282 |
| 778,600 | 0.124641654 | 0.169444619 | 0.053137982 | 0.12938665 |
| 778,700 | 0.124577938 | 0.170078689 | 0.052879652 | 0.129144583 |
| 778,800 | 0.12442191 | 0.169380887 | 0.053086563 | 0.129495034 |
| 778,900 | 0.124480321 | 0.169382348 | 0.05300869 | 0.129684196 |
| 779,000 | 0.124449393 | 0.170075639 | 0.053362121 | 0.130076225 |
| 779,100 | 0.124459289 | 0.169749494 | 0.052836449 | 0.129265189 |
| 779,200 | 0.124560372 | 0.16940765 | 0.05259736 | 0.129611137 |
| 779,300 | 0.124481073 | 0.169178726 | 0.052306841 | 0.129669418 |
| 779,400 | 0.124565176 | 0.169384248 | 0.052362067 | 0.129536464 |
| 779,500 | 0.124523735 | 0.169025812 | 0.052854124 | 0.130034823 |
| 779,600 | 0.124502989 | 0.169861339 | 0.05278585 | 0.129848224 |
| 779,700 | 0.124484615 | 0.16966868 | 0.052462894 | 0.129816114 |
| 779,800 | 0.124703592 | 0.170200725 | 0.052640604 | 0.129841968 |
| 779,900 | 0.124632697 | 0.169632169 | 0.05247415 | 0.129755737 |
| 780,000 | 0.124499893 | 0.169400282 | 0.052382437 | 0.129469651 |
| 780,100 | 0.124456279 | 0.169489612 | 0.052435649 | 0.129203602 |
| 780,200 | 0.124515041 | 0.169482224 | 0.052221627 | 0.129391056 |
| 780,300 | 0.124526652 | 0.169087068 | 0.052653368 | 0.129455261 |
| 780,400 | 0.124618457 | 0.169281798 | 0.052192242 | 0.128929803 |
| 780,500 | 0.124552012 | 0.169570997 | 0.052187999 | 0.128707865 |
| 780,600 | 0.124399202 | 0.1687864 | 0.052649473 | 0.129638543 |
| 780,700 | 0.12465643 | 0.168854407 | 0.052528894 | 0.129543634 |
| 780,800 | 0.124483285 | 0.169046016 | 0.052504559 | 0.129327704 |
| 780,900 | 0.124528893 | 0.169109081 | 0.052910573 | 0.130171762 |
| 781,000 | 0.124517818 | 0.169398052 | 0.053351729 | 0.129587866 |
| 781,100 | 0.124606596 | 0.170148066 | 0.05288844 | 0.129747401 |
| 781,200 | 0.124581302 | 0.169502902 | 0.053462229 | 0.129941521 |
| 781,300 | 0.124513674 | 0.169369068 | 0.052996211 | 0.129962624 |
| 781,400 | 0.124471741 | 0.169936356 | 0.05234014 | 0.129564694 |
| 781,500 | 0.124453702 | 0.169653552 | 0.052788236 | 0.129418758 |
| 781,600 | 0.124422542 | 0.170254936 | 0.052547371 | 0.129451296 |
| 781,700 | 0.124390204 | 0.170002456 | 0.053110874 | 0.130026601 |
| 781,800 | 0.124478734 | 0.169938176 | 0.052577984 | 0.13013528 |
| 781,900 | 0.124550481 | 0.170321251 | 0.052895338 | 0.13042144 |
| 782,000 | 0.124500997 | 0.169847107 | 0.053109935 | 0.129574728 |
| 782,100 | 0.124494545 | 0.169639586 | 0.053136439 | 0.130031285 |
| 782,200 | 0.124393454 | 0.16925355 | 0.0526472 | 0.130012622 |
| 782,300 | 0.124441112 | 0.169912225 | 0.052628452 | 0.130036611 |
| 782,400 | 0.124438513 | 0.16955437 | 0.052715338 | 0.129672792 |
| 782,500 | 0.124353638 | 0.169621421 | 0.05259499 | 0.129971477 |
| 782,600 | 0.124308633 | 0.169783581 | 0.052372283 | 0.13015482 |
| 782,700 | 0.124342509 | 0.170138149 | 0.05210512 | 0.129793229 |
| 782,800 | 0.124348161 | 0.170997326 | 0.052542253 | 0.129794018 |

| | | | | |
|---|---|---|---|---|
| 782,900 | 0.124271703 | 0.169750465 | 0.05244859 | 0.129806665 |
| 783,000 | 0.124328443 | 0.170238215 | 0.052055183 | 0.129887225 |
| 783,100 | 0.124244959 | 0.169589218 | 0.05243786 | 0.129522475 |
| 783,200 | 0.124257011 | 0.169703576 | 0.051616282 | 0.12983912 |
| 783,300 | 0.124229897 | 0.169996562 | 0.052176327 | 0.129920955 |
| 783,400 | 0.124281271 | 0.169743193 | 0.052582534 | 0.129805515 |
| 783,500 | 0.124255349 | 0.170161942 | 0.052356129 | 0.129613255 |
| 783,600 | 0.124096282 | 0.170128329 | 0.052450172 | 0.129560336 |
| 783,700 | 0.124140704 | 0.170411084 | 0.053099799 | 0.12965223 |
| 783,800 | 0.124105155 | 0.17004768 | 0.052433815 | 0.130533922 |
| 783,900 | 0.124057155 | 0.169262945 | 0.052829392 | 0.129634228 |
| 784,000 | 0.12416969 | 0.170331881 | 0.052875334 | 0.130338103 |
| 784,100 | 0.124109707 | 0.170149588 | 0.052707794 | 0.130014066 |
| 784,200 | 0.124017476 | 0.169642803 | 0.052659563 | 0.129479381 |
| 784,300 | 0.124091974 | 0.169249388 | 0.052380948 | 0.130276422 |
| 784,400 | 0.12402065 | 0.169172841 | 0.052807386 | 0.130200311 |
| 784,500 | 0.124033505 | 0.16962249 | 0.052057301 | 0.12978676 |
| 784,600 | 0.124090042 | 0.169505292 | 0.052503933 | 0.13024286 |
| 784,700 | 0.123962613 | 0.169813244 | 0.052864952 | 0.12990704 |
| 784,800 | 0.123982905 | 0.170354088 | 0.053140401 | 0.130166184 |
| 784,900 | 0.124011391 | 0.170311627 | 0.052663228 | 0.129907361 |
| 785,000 | 0.124036377 | 0.170625175 | 0.052361563 | 0.129910189 |
| 785,100 | 0.124066477 | 0.170162229 | 0.052076119 | 0.12988407 |
| 785,200 | 0.124002602 | 0.170728261 | 0.051745934 | 0.129666435 |
| 785,300 | 0.123982377 | 0.170589147 | 0.051977846 | 0.129405362 |
| 785,400 | 0.124028917 | 0.170614342 | 0.05286978 | 0.129591124 |
| 785,500 | 0.123974495 | 0.170609834 | 0.052936906 | 0.129601947 |
| 785,600 | 0.123995431 | 0.170104316 | 0.053002376 | 0.129869493 |
| 785,700 | 0.123961648 | 0.169970501 | 0.05260625 | 0.129510567 |
| 785,800 | 0.12392494 | 0.169654943 | 0.052457925 | 0.128678703 |
| 785,900 | 0.123918836 | 0.170259687 | 0.052620177 | 0.12919646 |
| 786,000 | 0.123909814 | 0.169883224 | 0.052658147 | 0.128848955 |
| 786,100 | 0.123979695 | 0.169717762 | 0.052557735 | 0.12944895 |
| 786,200 | 0.123885092 | 0.169462138 | 0.051988354 | 0.128736241 |
| 786,300 | 0.123916421 | 0.170386549 | 0.051950149 | 0.129245483 |
| 786,400 | 0.123905317 | 0.170558599 | 0.05255789 | 0.12865479 |
| 786,500 | 0.12382016 | 0.169792392 | 0.052160899 | 0.128512498 |
| 786,600 | 0.123798614 | 0.169698411 | 0.052454888 | 0.129123294 |
| 786,700 | 0.1238044 | 0.17054399 | 0.052949308 | 0.128562805 |
| 786,800 | 0.123833992 | 0.169567882 | 0.052347082 | 0.128932014 |
| 786,900 | 0.123959404 | 0.169809007 | 0.052149071 | 0.129013034 |
| 787,000 | 0.123873646 | 0.169250279 | 0.05238197 | 0.12903623 |
| 787,100 | 0.123722579 | 0.16944218 | 0.052105767 | 0.129214318 |
| 787,200 | 0.123741644 | 0.169520362 | 0.051912091 | 0.129237702 |
| 787,300 | 0.123804246 | 0.169856759 | 0.052171617 | 0.129600354 |
| 787,400 | 0.12374232 | 0.169517989 | 0.0525832 | 0.129237374 |
| 787,500 | 0.123723132 | 0.169321192 | 0.052194822 | 0.129645312 |
| 787,600 | 0.123742301 | 0.169908445 | 0.052473363 | 0.129534252 |
| 787,700 | 0.123696726 | 0.169444758 | 0.052792593 | 0.129948519 |
| 787,800 | 0.123711513 | 0.170252696 | 0.052762886 | 0.129329685 |
| 787,900 | 0.123633771 | 0.169530683 | 0.052765634 | 0.129632497 |
| 788,000 | 0.123713072 | 0.169069663 | 0.052935732 | 0.130153434 |
| 788,100 | 0.123722633 | 0.169612852 | 0.052358368 | 0.129688486 |
| 788,200 | 0.123647747 | 0.168908497 | 0.052697987 | 0.129243091 |
| 788,300 | 0.123722409 | 0.169458822 | 0.052826439 | 0.129612627 |
| 788,400 | 0.123761098 | 0.169798146 | 0.052792825 | 0.129616098 |
| 788,500 | 0.123696556 | 0.169091566 | 0.052390198 | 0.130217419 |
| 788,600 | 0.123605305 | 0.169546137 | 0.052660348 | 0.129739565 |
| 788,700 | 0.123652471 | 0.169613322 | 0.052687529 | 0.129534 |
| 788,800 | 0.123627058 | 0.16917296 | 0.053005366 | 0.129050103 |
| 788,900 | 0.123723265 | 0.170163856 | 0.053223948 | 0.12945319 |
| 789,000 | 0.123831483 | 0.169933731 | 0.053227469 | 0.129330868 |
| 789,100 | 0.123722671 | 0.169771552 | 0.052352862 | 0.129450298 |
| 789,200 | 0.123738063 | 0.170329815 | 0.052749663 | 0.129330257 |
| 789,300 | 0.12362453 | 0.169924578 | 0.053243471 | 0.129105643 |
| 789,400 | 0.123619151 | 0.170075765 | 0.053015999 | 0.129025752 |
| 789,500 | 0.123712073 | 0.170357654 | 0.053385753 | 0.129347334 |

| | | | | |
|---|---|---|---|---|
| 789,600 | 0.123698594 | 0.170051841 | 0.053192385 | 0.129155098 |
| 789,700 | 0.12377036 | 0.170359636 | 0.052700841 | 0.128970049 |
| 789,800 | 0.123725083 | 0.170120558 | 0.052864529 | 0.128933601 |
| 789,900 | 0.123736427 | 0.170420625 | 0.052891765 | 0.129102272 |
| 790,000 | 0.123613982 | 0.169910801 | 0.053046763 | 0.129608433 |
| 790,100 | 0.123602912 | 0.170079191 | 0.052939016 | 0.128961977 |
| 790,200 | 0.123677453 | 0.170648531 | 0.053169385 | 0.129276906 |
| 790,300 | 0.123553894 | 0.170207652 | 0.052800262 | 0.129065935 |
| 790,400 | 0.123531472 | 0.170130783 | 0.053332007 | 0.129295762 |
| 790,500 | 0.123592211 | 0.170260337 | 0.053003934 | 0.128970198 |
| 790,600 | 0.123514063 | 0.169913298 | 0.053177167 | 0.129028139 |
| 790,700 | 0.1235424 | 0.170274144 | 0.053445917 | 0.129288111 |
| 790,800 | 0.12354257 | 0.169875069 | 0.052834489 | 0.129207606 |
| 790,900 | 0.123505121 | 0.169432031 | 0.052993321 | 0.129113766 |
| 791,000 | 0.123381116 | 0.169393355 | 0.052860012 | 0.128723739 |
| 791,100 | 0.123410903 | 0.169582586 | 0.052805806 | 0.128404704 |
| 791,200 | 0.123296445 | 0.169462456 | 0.052838704 | 0.129141266 |
| 791,300 | 0.123382033 | 0.169307027 | 0.052592382 | 0.128790028 |
| 791,400 | 0.123426204 | 0.170207758 | 0.05302316 | 0.129017417 |
| 791,500 | 0.123388173 | 0.169756353 | 0.05300256 | 0.129003755 |
| 791,600 | 0.123507484 | 0.169453211 | 0.05266697 | 0.129379856 |
| 791,700 | 0.123501776 | 0.16966931 | 0.052435957 | 0.129304461 |
| 791,800 | 0.123429055 | 0.170145294 | 0.052471027 | 0.128893023 |
| 791,900 | 0.12345639 | 0.169809689 | 0.052894986 | 0.129746511 |
| 792,000 | 0.123511681 | 0.169881707 | 0.052767328 | 0.129396524 |
| 792,100 | 0.123410565 | 0.170192196 | 0.052950642 | 0.12905935 |
| 792,200 | 0.123477574 | 0.17032608 | 0.053215139 | 0.128983339 |
| 792,300 | 0.123362951 | 0.170596101 | 0.05301727 | 0.129032089 |
| 792,400 | 0.12338866 | 0.17028106 | 0.052950418 | 0.129385787 |
| 792,500 | 0.123419759 | 0.169409558 | 0.053068751 | 0.128989448 |
| 792,600 | 0.123342813 | 0.170459617 | 0.053147889 | 0.128833786 |
| 792,700 | 0.123424094 | 0.170037082 | 0.053175241 | 0.129077768 |
| 792,800 | 0.123380238 | 0.170385102 | 0.05339819 | 0.129228023 |
| 792,900 | 0.123347716 | 0.170392484 | 0.053490187 | 0.129278462 |
| 793,000 | 0.123256097 | 0.169801778 | 0.053308271 | 0.129012152 |
| 793,100 | 0.12337802 | 0.16957209 | 0.053786388 | 0.129014421 |
| 793,200 | 0.123378274 | 0.170327787 | 0.053363316 | 0.129202556 |
| 793,300 | 0.123373456 | 0.170275928 | 0.053223902 | 0.129248834 |
| 793,400 | 0.123367748 | 0.170841756 | 0.054083007 | 0.129258625 |
| 793,500 | 0.12329668 | 0.170478182 | 0.053702146 | 0.129591813 |
| 793,600 | 0.123422314 | 0.170781067 | 0.053773745 | 0.129016178 |
| 793,700 | 0.123251952 | 0.169723068 | 0.053575941 | 0.129510109 |
| 793,800 | 0.123347482 | 0.169757515 | 0.05403873 | 0.129184349 |
| 793,900 | 0.123240419 | 0.169529813 | 0.054018755 | 0.129009364 |
| 794,000 | 0.123282053 | 0.169426822 | 0.053622591 | 0.129226894 |
| 794,100 | 0.12321722 | 0.169923717 | 0.053627313 | 0.129309522 |
| 794,200 | 0.12326554 | 0.170025258 | 0.053766541 | 0.12878816 |
| 794,300 | 0.123433922 | 0.170295851 | 0.05391813 | 0.129040171 |
| 794,400 | 0.123309865 | 0.169958692 | 0.053597037 | 0.12865973 |
| 794,500 | 0.123220177 | 0.17066396 | 0.054292132 | 0.128648208 |
| 794,600 | 0.123239742 | 0.170357113 | 0.053812374 | 0.12932169 |
| 794,700 | 0.1232489 | 0.170352398 | 0.053673499 | 0.129345162 |
| 794,800 | 0.123220172 | 0.170069215 | 0.053598693 | 0.128399167 |
| 794,900 | 0.12319756 | 0.169716769 | 0.053731894 | 0.128014372 |
| 795,000 | 0.123277022 | 0.170665149 | 0.053735497 | 0.128766853 |
| 795,100 | 0.123181073 | 0.170247796 | 0.054085539 | 0.128304694 |
| 795,200 | 0.123211175 | 0.169565547 | 0.053492329 | 0.128865948 |
| 795,300 | 0.123111026 | 0.17021548 | 0.053395262 | 0.128528565 |
| 795,400 | 0.123138342 | 0.170391924 | 0.052944698 | 0.128727415 |
| 795,500 | 0.123133789 | 0.169552477 | 0.053423662 | 0.128483941 |
| 795,600 | 0.123049429 | 0.169524262 | 0.05418125 | 0.128197438 |
| 795,700 | 0.123023365 | 0.170117372 | 0.053948996 | 0.12852888 |
| 795,800 | 0.12308532 | 0.16980748 | 0.054065451 | 0.127996443 |
| 795,900 | 0.123143999 | 0.170043111 | 0.054659773 | 0.128329555 |
| 796,000 | 0.123192083 | 0.170111351 | 0.054189426 | 0.128644623 |
| 796,100 | 0.123139696 | 0.171200842 | 0.054027168 | 0.128632264 |
| 796,200 | 0.123077329 | 0.17045753 | 0.05399118 | 0.128403264 |

| | | | | |
|---|---|---|---|---|
| 796,300 | 0.123013371 | 0.171539044 | 0.053539224 | 0.128433803 |
| 796,400 | 0.12297648 | 0.171040543 | 0.053553299 | 0.127948174 |
| 796,500 | 0.123098921 | 0.170707895 | 0.053705873 | 0.128328618 |
| 796,600 | 0.123107787 | 0.170782405 | 0.053772253 | 0.127946274 |
| 796,700 | 0.123079672 | 0.170811023 | 0.053396121 | 0.127925101 |
| 796,800 | 0.123036714 | 0.170319926 | 0.053793591 | 0.12817223 |
| 796,900 | 0.123040133 | 0.1700353 | 0.053919963 | 0.128022477 |
| 797,000 | 0.123108287 | 0.170414898 | 0.054094376 | 0.127979682 |
| 797,100 | 0.123032704 | 0.169958864 | 0.054339482 | 0.127987926 |
| 797,200 | 0.123074514 | 0.170203909 | 0.054551055 | 0.128416265 |
| 797,300 | 0.12296876 | 0.169956405 | 0.054021109 | 0.128108926 |
| 797,400 | 0.123022676 | 0.170266288 | 0.054507588 | 0.128133427 |
| 797,500 | 0.123194074 | 0.170524079 | 0.05398924 | 0.1279121 |
| 797,600 | 0.123045198 | 0.169890071 | 0.054135088 | 0.127688136 |
| 797,700 | 0.122952679 | 0.169944762 | 0.054627572 | 0.128234687 |
| 797,800 | 0.122913415 | 0.169883248 | 0.054515872 | 0.127690502 |
| 797,900 | 0.123014414 | 0.170470797 | 0.054039574 | 0.127757671 |
| 798,000 | 0.12296902 | 0.170332497 | 0.0542541 | 0.127518661 |
| 798,100 | 0.122905586 | 0.170392896 | 0.054597181 | 0.127629614 |
| 798,200 | 0.123039515 | 0.170377217 | 0.054822313 | 0.128520918 |
| 798,300 | 0.122980945 | 0.170590279 | 0.054443258 | 0.127732239 |
| 798,400 | 0.122988245 | 0.170023931 | 0.053508921 | 0.128323874 |
| 798,500 | 0.123027047 | 0.170637376 | 0.053961097 | 0.128717584 |
| 798,600 | 0.122882903 | 0.170170052 | 0.054186779 | 0.128191903 |
| 798,700 | 0.122986451 | 0.170610454 | 0.054059901 | 0.128353157 |
| 798,800 | 0.122981985 | 0.170765773 | 0.053595193 | 0.128327619 |
| 798,900 | 0.123001377 | 0.170097456 | 0.054171378 | 0.12812145 |
| 799,000 | 0.123098697 | 0.170355888 | 0.054531816 | 0.128467771 |
| 799,100 | 0.122941207 | 0.170454571 | 0.054486015 | 0.128155976 |
| 799,200 | 0.122926522 | 0.169951778 | 0.054393552 | 0.127920249 |
| 799,300 | 0.122821471 | 0.170508439 | 0.054513784 | 0.128562611 |
| 799,400 | 0.122822486 | 0.170956365 | 0.054398308 | 0.128566686 |
| 799,500 | 0.123022881 | 0.171281796 | 0.054227175 | 0.128426017 |
| 799,600 | 0.122883038 | 0.170160244 | 0.054093677 | 0.129081115 |
| 799,700 | 0.122893568 | 0.170516017 | 0.054684674 | 0.128155946 |
| 799,800 | 0.122755091 | 0.170501279 | 0.054345359 | 0.128125067 |
| 799,900 | 0.122742402 | 0.170275316 | 0.054026998 | 0.12791086 |
| 800,000 | 0.122858883 | 0.170514903 | 0.053897684 | 0.128006025 |
| 800,100 | 0.122832763 | 0.170716048 | 0.053856378 | 0.128021171 |
| 800,200 | 0.122833991 | 0.170410194 | 0.05366273 | 0.128195448 |
| 800,300 | 0.122864835 | 0.170366695 | 0.054110418 | 0.128258136 |
| 800,400 | 0.122823154 | 0.170225738 | 0.053971276 | 0.127528315 |
| 800,500 | 0.122867373 | 0.170542503 | 0.053989789 | 0.127976275 |
| 800,600 | 0.12274944 | 0.170260628 | 0.053593393 | 0.128064477 |
| 800,700 | 0.122867838 | 0.169945882 | 0.054340772 | 0.127412352 |
| 800,800 | 0.122800086 | 0.170149315 | 0.053710753 | 0.127600802 |
| 800,900 | 0.12286502 | 0.170399669 | 0.053821601 | 0.128176592 |
| 801,000 | 0.122733698 | 0.169367121 | 0.054058941 | 0.127308207 |
| 801,100 | 0.122742135 | 0.170153913 | 0.054138662 | 0.128034428 |
| 801,200 | 0.122830316 | 0.170003462 | 0.053727813 | 0.128321829 |
| 801,300 | 0.122796488 | 0.169296029 | 0.054046784 | 0.127977969 |
| 801,400 | 0.122716033 | 0.169537292 | 0.053732362 | 0.128214267 |
| 801,500 | 0.122621162 | 0.169150578 | 0.053484287 | 0.127870994 |
| 801,600 | 0.122676243 | 0.169618515 | 0.053900483 | 0.127427022 |
| 801,700 | 0.122591279 | 0.169853317 | 0.054144389 | 0.127490619 |
| 801,800 | 0.122597202 | 0.169635537 | 0.053818228 | 0.128190777 |
| 801,900 | 0.122590006 | 0.169485868 | 0.054088577 | 0.127529228 |
| 802,000 | 0.122593318 | 0.169554159 | 0.054352866 | 0.127858501 |
| 802,100 | 0.122555943 | 0.169496444 | 0.054286133 | 0.127164923 |
| 802,200 | 0.122637825 | 0.169085127 | 0.054319566 | 0.12775196 |
| 802,300 | 0.122625578 | 0.169369777 | 0.05453559 | 0.128085148 |
| 802,400 | 0.122659636 | 0.16928596 | 0.054308083 | 0.127906566 |
| 802,500 | 0.122488936 | 0.168718053 | 0.054216723 | 0.127494236 |
| 802,600 | 0.122544028 | 0.168716314 | 0.054515121 | 0.12758524 |
| 802,700 | 0.122579454 | 0.168586403 | 0.054073632 | 0.127881323 |
| 802,800 | 0.122549335 | 0.168599759 | 0.054497258 | 0.127799236 |
| 802,900 | 0.122461558 | 0.168673937 | 0.054138522 | 0.128197131 |

| | | | | |
|---|---|---|---|---|
| 803,000 | 0.122532102 | 0.168984118 | 0.054064509 | 0.127412653 |
| 803,100 | 0.122494739 | 0.168441989 | 0.054105448 | 0.127463446 |
| 803,200 | 0.122488469 | 0.168832588 | 0.054724469 | 0.127605217 |
| 803,300 | 0.122440426 | 0.168832 | 0.054556143 | 0.127724811 |
| 803,400 | 0.122459289 | 0.168550999 | 0.053952544 | 0.127545764 |
| 803,500 | 0.122393704 | 0.16900344 | 0.053897026 | 0.127593992 |
| 803,600 | 0.122509981 | 0.168382357 | 0.054775 | 0.127163816 |
| 803,700 | 0.12247913 | 0.168697467 | 0.054691562 | 0.127475674 |
| 803,800 | 0.122487861 | 0.16816904 | 0.054284642 | 0.126910291 |
| 803,900 | 0.1224936 | 0.167984076 | 0.05385265 | 0.127004428 |
| 804,000 | 0.122557492 | 0.168445383 | 0.053264328 | 0.127450219 |
| 804,100 | 0.122446868 | 0.16894583 | 0.053810917 | 0.127236705 |
| 804,200 | 0.122517001 | 0.168818752 | 0.054148394 | 0.127438183 |
| 804,300 | 0.122498359 | 0.169014896 | 0.054202053 | 0.12702227 |
| 804,400 | 0.122564216 | 0.168647434 | 0.054046345 | 0.127528563 |
| 804,500 | 0.122533198 | 0.168683603 | 0.054615687 | 0.127381299 |
| 804,600 | 0.122537566 | 0.1688938 | 0.053941911 | 0.127361792 |
| 804,700 | 0.122474567 | 0.168948309 | 0.053519738 | 0.127105375 |
| 804,800 | 0.122471173 | 0.169132743 | 0.054126007 | 0.127326853 |
| 804,900 | 0.122453483 | 0.16920331 | 0.053874947 | 0.127348016 |
| 805,000 | 0.122282549 | 0.169539942 | 0.054239817 | 0.12734079 |
| 805,100 | 0.122344557 | 0.168839258 | 0.054317931 | 0.127466134 |
| 805,200 | 0.122353464 | 0.168877257 | 0.054265211 | 0.127724867 |
| 805,300 | 0.122446943 | 0.168984515 | 0.054771605 | 0.128098359 |
| 805,400 | 0.122345393 | 0.168569982 | 0.054354984 | 0.127551264 |
| 805,500 | 0.122222309 | 0.168373181 | 0.053974161 | 0.127136853 |
| 805,600 | 0.122290785 | 0.168594833 | 0.054148332 | 0.12690956 |
| 805,700 | 0.122329948 | 0.168925374 | 0.054107504 | 0.127272451 |
| 805,800 | 0.122415227 | 0.169115644 | 0.053676051 | 0.127373869 |
| 805,900 | 0.122369554 | 0.168696587 | 0.054444861 | 0.127924888 |
| 806,000 | 0.122416724 | 0.168472515 | 0.053884847 | 0.127710335 |
| 806,100 | 0.122366128 | 0.168542922 | 0.054150396 | 0.127542239 |
| 806,200 | 0.122405914 | 0.16900874 | 0.053576536 | 0.127439946 |
| 806,300 | 0.122312869 | 0.168619257 | 0.054082554 | 0.127405753 |
| 806,400 | 0.122203434 | 0.168384932 | 0.053997559 | 0.126609374 |
| 806,500 | 0.122251835 | 0.168217696 | 0.05377197 | 0.127130819 |
| 806,600 | 0.122316534 | 0.168102661 | 0.053683702 | 0.127508398 |
| 806,700 | 0.122248525 | 0.168018697 | 0.054653896 | 0.126993308 |
| 806,800 | 0.12229339 | 0.16820444 | 0.054484688 | 0.127212602 |
| 806,900 | 0.122043297 | 0.167104439 | 0.054219386 | 0.127278229 |
| 807,000 | 0.122113017 | 0.166981656 | 0.05469338 | 0.126683793 |
| 807,100 | 0.122098534 | 0.16762473 | 0.053956102 | 0.126796896 |
| 807,200 | 0.122160996 | 0.167367036 | 0.054151617 | 0.127356749 |
| 807,300 | 0.122158199 | 0.167672443 | 0.053639983 | 0.12746757 |
| 807,400 | 0.122210846 | 0.167378901 | 0.0543671 | 0.127281116 |
| 807,500 | 0.12208654 | 0.167736311 | 0.053868591 | 0.127141419 |
| 807,600 | 0.122116931 | 0.167291221 | 0.053920175 | 0.127348326 |
| 807,700 | 0.122245354 | 0.168429758 | 0.054172757 | 0.127473277 |
| 807,800 | 0.122197038 | 0.168565703 | 0.054057245 | 0.126908452 |
| 807,900 | 0.122107301 | 0.168042684 | 0.054657384 | 0.127375227 |
| 808,000 | 0.122104735 | 0.168626806 | 0.054888466 | 0.12725327 |
| 808,100 | 0.122197849 | 0.168616755 | 0.054683405 | 0.127290919 |
| 808,200 | 0.122029787 | 0.167741224 | 0.054810851 | 0.127180049 |
| 808,300 | 0.121968085 | 0.167383424 | 0.054981415 | 0.126455654 |
| 808,400 | 0.122107447 | 0.167958148 | 0.054682974 | 0.127206459 |
| 808,500 | 0.122017941 | 0.167436688 | 0.054823958 | 0.126335513 |
| 808,600 | 0.122122417 | 0.168074015 | 0.054978052 | 0.127045221 |
| 808,700 | 0.122101323 | 0.168613817 | 0.054448776 | 0.126626814 |
| 808,800 | 0.12213662 | 0.167890723 | 0.054725971 | 0.126476014 |
| 808,900 | 0.122192305 | 0.168203353 | 0.055070385 | 0.126724231 |
| 809,000 | 0.122230636 | 0.167996776 | 0.054733169 | 0.127267643 |
| 809,100 | 0.122250624 | 0.168023807 | 0.054869506 | 0.12750461 |
| 809,200 | 0.122172574 | 0.167838465 | 0.055328807 | 0.126963306 |
| 809,300 | 0.122182335 | 0.168023549 | 0.055162228 | 0.127357815 |
| 809,400 | 0.122076717 | 0.168196225 | 0.054589578 | 0.127101309 |
| 809,500 | 0.122184214 | 0.168147856 | 0.054905638 | 0.126874632 |
| 809,600 | 0.122143657 | 0.168831725 | 0.054691563 | 0.126916347 |

| | | | | |
|---|---|---|---|---|
| 809,700 | 0.122213904 | 0.168535059 | 0.054047506 | 0.127062483 |
| 809,800 | 0.122065745 | 0.168067891 | 0.054945033 | 0.126530472 |
| 809,900 | 0.122098322 | 0.167648844 | 0.054507536 | 0.127346596 |
| 810,000 | 0.122110866 | 0.167887394 | 0.054456633 | 0.127010165 |
| 810,100 | 0.122139421 | 0.167601619 | 0.054057789 | 0.127230691 |
| 810,200 | 0.122101605 | 0.167974801 | 0.054865242 | 0.127504107 |
| 810,300 | 0.12210036 | 0.16744086 | 0.054725855 | 0.127647349 |
| 810,400 | 0.122004242 | 0.167217566 | 0.054494986 | 0.127608456 |
| 810,500 | 0.122015073 | 0.166892832 | 0.054664774 | 0.127909322 |
| 810,600 | 0.12199514 | 0.166927904 | 0.054415489 | 0.127202933 |
| 810,700 | 0.122173164 | 0.168063465 | 0.054171958 | 0.127954369 |
| 810,800 | 0.121965869 | 0.16749036 | 0.054490786 | 0.127362792 |
| 810,900 | 0.121985647 | 0.167664257 | 0.054171648 | 0.12714404 |
| 811,000 | 0.121999223 | 0.167450445 | 0.054596432 | 0.127463332 |
| 811,100 | 0.122077428 | 0.167919502 | 0.054425947 | 0.127362478 |
| 811,200 | 0.122042395 | 0.168278091 | 0.054088885 | 0.126997227 |
| 811,300 | 0.121958156 | 0.167473271 | 0.053566365 | 0.126603574 |
| 811,400 | 0.121913158 | 0.167483448 | 0.053598412 | 0.126698478 |
| 811,500 | 0.121951905 | 0.167892962 | 0.053784306 | 0.127163498 |
| 811,600 | 0.121853851 | 0.168276154 | 0.053800504 | 0.126736393 |
| 811,700 | 0.121852902 | 0.167511976 | 0.054035939 | 0.126586057 |
| 811,800 | 0.121962296 | 0.167238955 | 0.05372271 | 0.127267357 |
| 811,900 | 0.121966063 | 0.167848145 | 0.053655804 | 0.127111441 |
| 812,000 | 0.121924948 | 0.167297033 | 0.053530182 | 0.126857508 |
| 812,100 | 0.121934324 | 0.167693631 | 0.054010923 | 0.1269891 |
| 812,200 | 0.121904499 | 0.166912564 | 0.053448038 | 0.127106069 |
| 812,300 | 0.121936465 | 0.166828165 | 0.053710146 | 0.127559458 |
| 812,400 | 0.121842224 | 0.166811415 | 0.053338354 | 0.126397829 |
| 812,500 | 0.122055067 | 0.16730693 | 0.053582772 | 0.125942335 |
| 812,600 | 0.121828754 | 0.167231419 | 0.054020388 | 0.126015529 |
| 812,700 | 0.121935815 | 0.167546716 | 0.053762948 | 0.126278046 |
| 812,800 | 0.121960472 | 0.168037321 | 0.053615225 | 0.125945195 |
| 812,900 | 0.121976219 | 0.167269785 | 0.05375547 | 0.126314738 |
| 813,000 | 0.121994765 | 0.167285997 | 0.054071483 | 0.126418975 |
| 813,100 | 0.121985646 | 0.167268461 | 0.054088463 | 0.126223045 |
| 813,200 | 0.12184989 | 0.166399259 | 0.053555305 | 0.126293161 |
| 813,300 | 0.121923043 | 0.16694379 | 0.05402074 | 0.12584444 |
| 813,400 | 0.121968675 | 0.166838979 | 0.053633044 | 0.12614064 |
| 813,500 | 0.121927709 | 0.166838202 | 0.053861363 | 0.125808616 |
| 813,600 | 0.121910596 | 0.167466062 | 0.053994892 | 0.125969942 |
| 813,700 | 0.121908215 | 0.16694622 | 0.054116393 | 0.126284538 |
| 813,800 | 0.12189191 | 0.167022967 | 0.054022413 | 0.125805665 |
| 813,900 | 0.121896038 | 0.166864189 | 0.054272269 | 0.125406672 |
| 814,000 | 0.121939178 | 0.167818375 | 0.054174113 | 0.125994077 |
| 814,100 | 0.121836902 | 0.167822684 | 0.054544162 | 0.125245687 |
| 814,200 | 0.121852569 | 0.167157418 | 0.054446139 | 0.125548759 |
| 814,300 | 0.121883684 | 0.166889933 | 0.053892696 | 0.125851042 |
| 814,400 | 0.121836203 | 0.167639158 | 0.054242396 | 0.125495597 |
| 814,500 | 0.121835956 | 0.167442339 | 0.054367104 | 0.125187457 |
| 814,600 | 0.121844747 | 0.167493195 | 0.054215523 | 0.125799685 |
| 814,700 | 0.12183185 | 0.167522265 | 0.054196487 | 0.125916478 |
| 814,800 | 0.121755286 | 0.166597564 | 0.053897305 | 0.125727331 |
| 814,900 | 0.121727758 | 0.166694479 | 0.054107253 | 0.125687309 |
| 815,000 | 0.121689669 | 0.166314691 | 0.054688271 | 0.125870014 |
| 815,100 | 0.12177865 | 0.167077754 | 0.054498859 | 0.125661246 |
| 815,200 | 0.121666497 | 0.166304557 | 0.054708394 | 0.126012707 |
| 815,300 | 0.121688771 | 0.166881754 | 0.05424698 | 0.125439132 |
| 815,400 | 0.121601823 | 0.166568185 | 0.054084331 | 0.125639482 |
| 815,500 | 0.121708816 | 0.166508021 | 0.0545458 | 0.125610528 |
| 815,600 | 0.121711141 | 0.167003516 | 0.053686378 | 0.125829534 |
| 815,700 | 0.121740578 | 0.166873337 | 0.053993518 | 0.125934574 |
| 815,800 | 0.121726279 | 0.16677658 | 0.053775132 | 0.126075684 |
| 815,900 | 0.121737758 | 0.166766459 | 0.053678237 | 0.125951293 |
| 816,000 | 0.121716729 | 0.167051014 | 0.053737863 | 0.125501953 |
| 816,100 | 0.121650737 | 0.166680095 | 0.053275917 | 0.125502367 |
| 816,200 | 0.121615686 | 0.166715156 | 0.053272379 | 0.125472671 |
| 816,300 | 0.121658567 | 0.166709103 | 0.053175559 | 0.125571457 |

| | | | | |
|---|---|---|---|---|
| 816,400 | 0.121584512 | 0.166466532 | 0.053224107 | 0.125942896 |
| 816,500 | 0.121514076 | 0.166555569 | 0.05346155 | 0.126046896 |
| 816,600 | 0.121542327 | 0.167075674 | 0.054104702 | 0.125997744 |
| 816,700 | 0.121518806 | 0.166788168 | 0.053644719 | 0.126607059 |
| 816,800 | 0.1215805 | 0.166842074 | 0.05351421 | 0.125749492 |
| 816,900 | 0.12149074 | 0.16697672 | 0.05339367 | 0.126308968 |
| 817,000 | 0.121562882 | 0.167060843 | 0.053926619 | 0.125777251 |
| 817,100 | 0.121564846 | 0.166832695 | 0.054097773 | 0.126026464 |
| 817,200 | 0.121516777 | 0.167111206 | 0.053831029 | 0.125863904 |
| 817,300 | 0.121566352 | 0.166554101 | 0.053700355 | 0.125763377 |
| 817,400 | 0.121544263 | 0.166957297 | 0.053724791 | 0.125397249 |
| 817,500 | 0.121581547 | 0.167140044 | 0.053519748 | 0.124876714 |
| 817,600 | 0.121467485 | 0.166858031 | 0.054018363 | 0.125490196 |
| 817,700 | 0.121622718 | 0.166245102 | 0.054061836 | 0.125725154 |
| 817,800 | 0.121534084 | 0.166371766 | 0.053590204 | 0.125386354 |
| 817,900 | 0.121533445 | 0.166749152 | 0.05337205 | 0.125593018 |
| 818,000 | 0.121453958 | 0.166619956 | 0.053802613 | 0.126081101 |
| 818,100 | 0.121377715 | 0.166221265 | 0.053569458 | 0.125809163 |
| 818,200 | 0.121485995 | 0.167243488 | 0.053202839 | 0.125331122 |
| 818,300 | 0.121327938 | 0.166557474 | 0.053548659 | 0.125269994 |
| 818,400 | 0.121503243 | 0.166856445 | 0.053306765 | 0.125274153 |
| 818,500 | 0.121500864 | 0.166502457 | 0.05330541 | 0.125188463 |
| 818,600 | 0.121455246 | 0.166728643 | 0.053629152 | 0.125280555 |
| 818,700 | 0.121337961 | 0.165303414 | 0.053808171 | 0.125340386 |
| 818,800 | 0.121533365 | 0.165551458 | 0.053734938 | 0.125560799 |
| 818,900 | 0.12145666 | 0.165739155 | 0.05411338 | 0.125712922 |
| 819,000 | 0.121453161 | 0.166565789 | 0.053088028 | 0.125198505 |
| 819,100 | 0.121394692 | 0.16593516 | 0.053760646 | 0.125205491 |
| 819,200 | 0.121354367 | 0.166091496 | 0.053848146 | 0.125462565 |
| 819,300 | 0.121356382 | 0.166110852 | 0.053734838 | 0.125333184 |
| 819,400 | 0.121306479 | 0.165267697 | 0.053946276 | 0.125068866 |
| 819,500 | 0.121272325 | 0.16502424 | 0.053506346 | 0.125122417 |
| 819,600 | 0.12140209 | 0.165631991 | 0.053898309 | 0.125307321 |
| 819,700 | 0.121309263 | 0.165031511 | 0.053327352 | 0.124549397 |
| 819,800 | 0.121290358 | 0.165569847 | 0.05331045 | 0.124926617 |
| 819,900 | 0.121330463 | 0.16543384 | 0.053678396 | 0.125179638 |
| 820,000 | 0.121308036 | 0.164951191 | 0.05357448 | 0.124942529 |
| 820,100 | 0.121231448 | 0.164930562 | 0.052522548 | 0.125129059 |
| 820,200 | 0.12123634 | 0.164799818 | 0.053266252 | 0.125174804 |
| 820,300 | 0.121230879 | 0.165187533 | 0.053161398 | 0.124886278 |
| 820,400 | 0.121044545 | 0.16511894 | 0.053364068 | 0.125120402 |
| 820,500 | 0.121120889 | 0.165608835 | 0.05336648 | 0.125157969 |
| 820,600 | 0.121228712 | 0.165420963 | 0.053706374 | 0.125038889 |
| 820,700 | 0.121162172 | 0.165337646 | 0.053343821 | 0.124633187 |
| 820,800 | 0.121286034 | 0.165346016 | 0.053743812 | 0.124930713 |
| 820,900 | 0.121122324 | 0.166396835 | 0.05368306 | 0.124716389 |
| 821,000 | 0.121220439 | 0.166010733 | 0.053313475 | 0.124617284 |
| 821,100 | 0.121177783 | 0.165530398 | 0.053147683 | 0.125114378 |
| 821,200 | 0.121078387 | 0.165895772 | 0.053124737 | 0.124345595 |
| 821,300 | 0.121093419 | 0.166046167 | 0.053864571 | 0.124623551 |
| 821,400 | 0.121137833 | 0.166012536 | 0.053161771 | 0.124238239 |
| 821,500 | 0.121058827 | 0.165517474 | 0.053116999 | 0.124632918 |
| 821,600 | 0.120980694 | 0.165239241 | 0.052993681 | 0.124602251 |
| 821,700 | 0.121076437 | 0.165014275 | 0.053213886 | 0.124768858 |
| 821,800 | 0.120980204 | 0.165056878 | 0.053165548 | 0.125196742 |
| 821,900 | 0.121098272 | 0.165505816 | 0.053221878 | 0.124729364 |
| 822,000 | 0.120976183 | 0.164882995 | 0.053043647 | 0.124708392 |
| 822,100 | 0.121009034 | 0.165262921 | 0.052955052 | 0.124240308 |
| 822,200 | 0.12104433 | 0.166173534 | 0.053039481 | 0.124387684 |
| 822,300 | 0.120921237 | 0.165318174 | 0.052808812 | 0.124612475 |
| 822,400 | 0.120974855 | 0.165269185 | 0.053235912 | 0.124655385 |
| 822,500 | 0.120839507 | 0.164245197 | 0.053157656 | 0.124686089 |
| 822,600 | 0.120907752 | 0.164911233 | 0.053201061 | 0.124520915 |
| 822,700 | 0.120843957 | 0.164576415 | 0.053515976 | 0.124604563 |
| 822,800 | 0.120938843 | 0.165111186 | 0.053058791 | 0.124651815 |
| 822,900 | 0.120820265 | 0.164669246 | 0.053425818 | 0.124393357 |
| 823,000 | 0.120966098 | 0.165609885 | 0.053326852 | 0.124563941 |

| | | | | |
|---|---|---|---|---|
| 823,100 | 0.12090102 | 0.165483444 | 0.05337342 | 0.124365747 |
| 823,200 | 0.120908646 | 0.164849849 | 0.053631888 | 0.124837998 |
| 823,300 | 0.120698409 | 0.165869906 | 0.053166101 | 0.124365663 |
| 823,400 | 0.120738917 | 0.164195388 | 0.053455524 | 0.125158004 |
| 823,500 | 0.120653112 | 0.165332404 | 0.05299548 | 0.125063445 |
| 823,600 | 0.120779256 | 0.16479188 | 0.053051455 | 0.124874428 |
| 823,700 | 0.120628522 | 0.164096653 | 0.052777772 | 0.124706305 |
| 823,800 | 0.120832165 | 0.164770261 | 0.052875811 | 0.124837245 |
| 823,900 | 0.120805761 | 0.164704863 | 0.052942981 | 0.124772179 |
| 824,000 | 0.120770985 | 0.164703446 | 0.052696088 | 0.124796205 |
| 824,100 | 0.120797143 | 0.164428383 | 0.053350785 | 0.124284331 |
| 824,200 | 0.120733621 | 0.164610446 | 0.053081488 | 0.123978511 |
| 824,300 | 0.120706804 | 0.164461005 | 0.053501103 | 0.124362554 |
| 824,400 | 0.120699264 | 0.164575017 | 0.05278175 | 0.124228905 |
| 824,500 | 0.120737354 | 0.164508854 | 0.052623464 | 0.124544179 |
| 824,600 | 0.120647366 | 0.164807943 | 0.052920552 | 0.124296376 |
| 824,700 | 0.120643573 | 0.164503793 | 0.05245857 | 0.124115604 |
| 824,800 | 0.120680131 | 0.164402958 | 0.05260434 | 0.124375956 |
| 824,900 | 0.120611476 | 0.164699905 | 0.052976117 | 0.123958437 |
| 825,000 | 0.120645628 | 0.164168196 | 0.052658404 | 0.124027823 |
| 825,100 | 0.120643217 | 0.164322504 | 0.052479346 | 0.124240633 |
| 825,200 | 0.120745544 | 0.164160153 | 0.053291464 | 0.124170234 |
| 825,300 | 0.120561159 | 0.164507186 | 0.052789893 | 0.123820076 |
| 825,400 | 0.120584875 | 0.164856316 | 0.053092778 | 0.124037421 |
| 825,500 | 0.120639275 | 0.165181335 | 0.052278382 | 0.124445652 |
| 825,600 | 0.120653824 | 0.164359418 | 0.053002505 | 0.124441004 |
| 825,700 | 0.120638867 | 0.164792421 | 0.052352117 | 0.124303819 |
| 825,800 | 0.120612265 | 0.164363153 | 0.052545519 | 0.124056035 |
| 825,900 | 0.120568993 | 0.164726551 | 0.052570004 | 0.123634353 |
| 826,000 | 0.120605561 | 0.164743188 | 0.052587314 | 0.124101516 |
| 826,100 | 0.120538835 | 0.164709073 | 0.052617227 | 0.123851977 |
| 826,200 | 0.120526161 | 0.165000952 | 0.053165555 | 0.123622566 |
| 826,300 | 0.1204025 | 0.164400463 | 0.052418385 | 0.12393376 |
| 826,400 | 0.120448293 | 0.164670746 | 0.052395946 | 0.124180769 |
| 826,500 | 0.120406062 | 0.164262205 | 0.052156246 | 0.123692444 |
| 826,600 | 0.120366616 | 0.163551751 | 0.052300854 | 0.123486932 |
| 826,700 | 0.120407152 | 0.163685719 | 0.052297054 | 0.123352175 |
| 826,800 | 0.120383043 | 0.16336488 | 0.052844188 | 0.12346347 |
| 826,900 | 0.120298565 | 0.163940864 | 0.052310919 | 0.123062559 |
| 827,000 | 0.120361823 | 0.164390294 | 0.052638862 | 0.123306268 |
| 827,100 | 0.120284814 | 0.163818555 | 0.05274075 | 0.123081709 |
| 827,200 | 0.120398798 | 0.164382145 | 0.053366141 | 0.123772908 |
| 827,300 | 0.120459793 | 0.164556256 | 0.052401094 | 0.124072621 |
| 827,400 | 0.120511849 | 0.164447306 | 0.052615686 | 0.123667762 |
| 827,500 | 0.120359981 | 0.164402062 | 0.052610623 | 0.123143036 |
| 827,600 | 0.120401457 | 0.164264702 | 0.05277808 | 0.123205463 |
| 827,700 | 0.12050356 | 0.164331795 | 0.052431738 | 0.122997688 |
| 827,800 | 0.12045682 | 0.164110469 | 0.053135721 | 0.123150208 |
| 827,900 | 0.120500833 | 0.164326453 | 0.052215976 | 0.123697456 |
| 828,000 | 0.120424596 | 0.164940401 | 0.053058903 | 0.123511744 |
| 828,100 | 0.120375243 | 0.16413704 | 0.053154023 | 0.123753324 |
| 828,200 | 0.120311889 | 0.163542854 | 0.052665645 | 0.123361148 |
| 828,300 | 0.120297081 | 0.164295103 | 0.052430039 | 0.123323376 |
| 828,400 | 0.120318164 | 0.163900786 | 0.052061549 | 0.123544607 |
| 828,500 | 0.120339856 | 0.164456596 | 0.052445586 | 0.123067899 |
| 828,600 | 0.120289307 | 0.164632672 | 0.051953501 | 0.123240063 |
| 828,700 | 0.120333776 | 0.164073856 | 0.052141489 | 0.123187288 |
| 828,800 | 0.120300237 | 0.163610162 | 0.051928085 | 0.122904156 |
| 828,900 | 0.120274505 | 0.164213224 | 0.052133063 | 0.122548989 |
| 829,000 | 0.120364865 | 0.163727035 | 0.052232152 | 0.123229065 |
| 829,100 | 0.120207468 | 0.163162846 | 0.05223804 | 0.122767284 |
| 829,200 | 0.120277049 | 0.163457379 | 0.052272442 | 0.122489959 |
| 829,300 | 0.120248059 | 0.163848185 | 0.052520307 | 0.122980114 |
| 829,400 | 0.120175238 | 0.163575342 | 0.052694302 | 0.122583987 |
| 829,500 | 0.120222982 | 0.163757934 | 0.052280888 | 0.123368803 |
| 829,600 | 0.120164137 | 0.163720145 | 0.052424214 | 0.122851648 |
| 829,700 | 0.120134273 | 0.163460403 | 0.052449583 | 0.122730832 |

| | | | | |
|---|---|---|---|---|
| 829,800 | 0.120075083 | 0.16296637 | 0.052856437 | 0.122728642 |
| 829,900 | 0.120209795 | 0.163089532 | 0.053176681 | 0.122769897 |
| 830,000 | 0.120081611 | 0.164211325 | 0.052669915 | 0.122690063 |
| 830,100 | 0.120115548 | 0.163991334 | 0.052855473 | 0.122357428 |
| 830,200 | 0.120090178 | 0.164214502 | 0.052463243 | 0.122302359 |
| 830,300 | 0.120084546 | 0.164391834 | 0.052787518 | 0.122748469 |
| 830,400 | 0.120113278 | 0.1643431 | 0.052663216 | 0.12246459 |
| 830,500 | 0.120025805 | 0.163656685 | 0.052461241 | 0.122800783 |
| 830,600 | 0.120066729 | 0.16375168 | 0.052696662 | 0.122523687 |
| 830,700 | 0.120150951 | 0.164320279 | 0.053099986 | 0.122707037 |
| 830,800 | 0.120056412 | 0.164678407 | 0.053185158 | 0.1228007 |
| 830,900 | 0.120061841 | 0.163163733 | 0.052638727 | 0.122603173 |
| 831,000 | 0.119970777 | 0.163701813 | 0.052794879 | 0.122662027 |
| 831,100 | 0.120074687 | 0.163876852 | 0.052339044 | 0.122604092 |
| 831,200 | 0.119944801 | 0.16359211 | 0.052832141 | 0.121790562 |
| 831,300 | 0.120051305 | 0.164098945 | 0.052957006 | 0.122274913 |
| 831,400 | 0.120148893 | 0.164251998 | 0.052514323 | 0.122985827 |
| 831,500 | 0.120132729 | 0.164596963 | 0.052563338 | 0.122642585 |
| 831,600 | 0.120089579 | 0.163860649 | 0.052183796 | 0.122581917 |
| 831,700 | 0.12007936 | 0.164957411 | 0.052855061 | 0.122356582 |
| 831,800 | 0.120011977 | 0.16441187 | 0.052275498 | 0.121784625 |
| 831,900 | 0.120044015 | 0.164699113 | 0.052798341 | 0.122119933 |
| 832,000 | 0.120019657 | 0.164633141 | 0.052133913 | 0.122015211 |
| 832,100 | 0.120018882 | 0.164520354 | 0.052832579 | 0.122279164 |
| 832,200 | 0.120047825 | 0.164335562 | 0.051876473 | 0.122338106 |
| 832,300 | 0.120007098 | 0.164709866 | 0.051831876 | 0.122313773 |
| 832,400 | 0.120094281 | 0.164990888 | 0.051525733 | 0.122715293 |
| 832,500 | 0.120019612 | 0.16474805 | 0.052304789 | 0.122475086 |
| 832,600 | 0.120032712 | 0.164705834 | 0.05171948 | 0.122730138 |
| 832,700 | 0.120067668 | 0.163728947 | 0.052138965 | 0.122803681 |
| 832,800 | 0.120003307 | 0.164680239 | 0.052060302 | 0.122417889 |
| 832,900 | 0.119936855 | 0.164665892 | 0.052340147 | 0.122393713 |
| 833,000 | 0.120055996 | 0.164885345 | 0.051798367 | 0.122962943 |
| 833,100 | 0.119854311 | 0.16446061 | 0.052552905 | 0.122453543 |
| 833,200 | 0.119895308 | 0.164268181 | 0.052207673 | 0.122074192 |
| 833,300 | 0.119946899 | 0.165183347 | 0.051707108 | 0.122261977 |
| 833,400 | 0.119996251 | 0.165035053 | 0.05259739 | 0.121930553 |
| 833,500 | 0.119919734 | 0.165336179 | 0.052713529 | 0.122722483 |
| 833,600 | 0.119879301 | 0.164918382 | 0.052125719 | 0.12244653 |
| 833,700 | 0.119937911 | 0.165144654 | 0.05234396 | 0.122546266 |
| 833,800 | 0.119839646 | 0.164473774 | 0.052101966 | 0.122673889 |
| 833,900 | 0.119917042 | 0.164502209 | 0.051697091 | 0.122752054 |
| 834,000 | 0.119818602 | 0.164681641 | 0.051828776 | 0.122966689 |
| 834,100 | 0.119826449 | 0.164952485 | 0.052454758 | 0.123153524 |
| 834,200 | 0.119828959 | 0.164227246 | 0.052082769 | 0.123302915 |
| 834,300 | 0.119750556 | 0.164033831 | 0.052135105 | 0.123197796 |
| 834,400 | 0.119781454 | 0.164808124 | 0.052094505 | 0.12261669 |
| 834,500 | 0.119704499 | 0.164148171 | 0.052182006 | 0.122760132 |
| 834,600 | 0.11978467 | 0.164155555 | 0.052055036 | 0.123015139 |
| 834,700 | 0.119873159 | 0.164438092 | 0.0527587 | 0.123058379 |
| 834,800 | 0.11981806 | 0.164455583 | 0.052425014 | 0.123138361 |
| 834,900 | 0.119821105 | 0.164277026 | 0.052682362 | 0.123036947 |
| 835,000 | 0.119884766 | 0.164694078 | 0.053046253 | 0.123559021 |
| 835,100 | 0.119946638 | 0.164687317 | 0.052430451 | 0.123173699 |
| 835,200 | 0.119872147 | 0.164204568 | 0.052586627 | 0.123235011 |
| 835,300 | 0.119829316 | 0.164593476 | 0.052460242 | 0.122562678 |
| 835,400 | 0.11991394 | 0.165055217 | 0.05242043 | 0.122723023 |
| 835,500 | 0.11979064 | 0.163948332 | 0.052494567 | 0.122530429 |
| 835,600 | 0.119795719 | 0.164433317 | 0.052480855 | 0.123086034 |
| 835,700 | 0.119779796 | 0.164014117 | 0.052521376 | 0.123139308 |
| 835,800 | 0.119629714 | 0.163932002 | 0.052409336 | 0.122519413 |
| 835,900 | 0.119742946 | 0.16447768 | 0.051941736 | 0.122647569 |
| 836,000 | 0.119730813 | 0.163724491 | 0.053189458 | 0.122370066 |
| 836,100 | 0.119710875 | 0.163837339 | 0.053169002 | 0.122135948 |
| 836,200 | 0.119765001 | 0.163938008 | 0.053134545 | 0.122466849 |
| 836,300 | 0.11970568 | 0.163776973 | 0.053545386 | 0.121906616 |
| 836,400 | 0.119578487 | 0.164329688 | 0.053253224 | 0.121463135 |

| | | | | |
|---|---|---|---|---|
| 836,500 | 0.119648202 | 0.163840607 | 0.052639949 | 0.122121411 |
| 836,600 | 0.119631737 | 0.163335238 | 0.052277099 | 0.122440679 |
| 836,700 | 0.119582948 | 0.163769384 | 0.052529075 | 0.122763773 |
| 836,800 | 0.119600141 | 0.163511337 | 0.052916513 | 0.122760531 |
| 836,900 | 0.119517294 | 0.163705 | 0.052496057 | 0.122634147 |
| 837,000 | 0.119549773 | 0.163508864 | 0.053068116 | 0.122133948 |
| 837,100 | 0.119485394 | 0.163514517 | 0.05260911 | 0.122742248 |
| 837,200 | 0.119557168 | 0.162873887 | 0.0529915 | 0.122451174 |
| 837,300 | 0.119537688 | 0.163844441 | 0.052389014 | 0.122466475 |
| 837,400 | 0.119462403 | 0.163747776 | 0.052396543 | 0.122252043 |
| 837,500 | 0.119481285 | 0.163894846 | 0.052426842 | 0.122215769 |
| 837,600 | 0.119493281 | 0.163321869 | 0.052341469 | 0.122351241 |
| 837,700 | 0.119600778 | 0.162981467 | 0.052695952 | 0.121882926 |
| 837,800 | 0.119378554 | 0.162897613 | 0.052766008 | 0.122040343 |
| 837,900 | 0.119460007 | 0.163666565 | 0.05275548 | 0.122512113 |
| 838,000 | 0.11940195 | 0.163933206 | 0.052715041 | 0.122092046 |
| 838,100 | 0.119531617 | 0.163895825 | 0.052656702 | 0.122193945 |
| 838,200 | 0.11950359 | 0.163380997 | 0.052658393 | 0.122068541 |
| 838,300 | 0.119420288 | 0.163652272 | 0.052436256 | 0.12202984 |
| 838,400 | 0.119388714 | 0.163740063 | 0.052624897 | 0.121609166 |
| 838,500 | 0.119463219 | 0.163030247 | 0.052660398 | 0.122346307 |
| 838,600 | 0.119558106 | 0.163893943 | 0.052646393 | 0.122309443 |
| 838,700 | 0.119437148 | 0.16433324 | 0.052194778 | 0.122054654 |
| 838,800 | 0.119495685 | 0.163719635 | 0.052070795 | 0.122490683 |
| 838,900 | 0.119529291 | 0.164271609 | 0.052414783 | 0.122469977 |
| 839,000 | 0.119413129 | 0.164069308 | 0.052788062 | 0.121928889 |
| 839,100 | 0.119500457 | 0.164293535 | 0.052801938 | 0.122589418 |
| 839,200 | 0.119520297 | 0.164008369 | 0.052544619 | 0.122731073 |
| 839,300 | 0.119533442 | 0.163980186 | 0.053357637 | 0.122632756 |
| 839,400 | 0.119418402 | 0.16343063 | 0.052962888 | 0.122595353 |
| 839,500 | 0.119401462 | 0.162696778 | 0.052691127 | 0.122656878 |
| 839,600 | 0.119439382 | 0.163419963 | 0.053142457 | 0.122416947 |
| 839,700 | 0.119331336 | 0.163283862 | 0.052888041 | 0.122638235 |
| 839,800 | 0.11916623 | 0.162668318 | 0.052516219 | 0.122406711 |
| 839,900 | 0.119265238 | 0.162440458 | 0.052118206 | 0.122326553 |
| 840,000 | 0.119339563 | 0.162674116 | 0.052599353 | 0.122621102 |
| 840,100 | 0.119389854 | 0.162931502 | 0.052815613 | 0.122250065 |
| 840,200 | 0.119315659 | 0.163146866 | 0.053348292 | 0.122332822 |
| 840,300 | 0.119304635 | 0.162756107 | 0.053165289 | 0.121877245 |
| 840,400 | 0.119327675 | 0.16314389 | 0.053038106 | 0.122063636 |
| 840,500 | 0.119382529 | 0.162932645 | 0.053001874 | 0.122162834 |
| 840,600 | 0.119350319 | 0.162664508 | 0.053459361 | 0.121901671 |
| 840,700 | 0.119316257 | 0.163005177 | 0.053071854 | 0.121583866 |
| 840,800 | 0.119446795 | 0.163705994 | 0.053363572 | 0.1219511 |
| 840,900 | 0.119233495 | 0.162768987 | 0.052797759 | 0.122082573 |
| 841,000 | 0.119436011 | 0.163231014 | 0.052616853 | 0.122441204 |
| 841,100 | 0.119387142 | 0.163162066 | 0.05301603 | 0.12237881 |
| 841,200 | 0.119507697 | 0.163073006 | 0.053198724 | 0.122177829 |
| 841,300 | 0.119372373 | 0.162435603 | 0.052913886 | 0.122570526 |
| 841,400 | 0.119435746 | 0.162648356 | 0.053089238 | 0.122583261 |
| 841,500 | 0.119353405 | 0.162908431 | 0.052959629 | 0.122342342 |
| 841,600 | 0.119327499 | 0.163312087 | 0.053280672 | 0.122843145 |
| 841,700 | 0.11936862 | 0.163224723 | 0.05322487 | 0.123029087 |
| 841,800 | 0.119326092 | 0.163117677 | 0.053230385 | 0.122753045 |
| 841,900 | 0.119401968 | 0.163764245 | 0.053335259 | 0.123208808 |
| 842,000 | 0.119303539 | 0.162678119 | 0.053129443 | 0.122667194 |
| 842,100 | 0.119351281 | 0.162810958 | 0.052648689 | 0.123468903 |
| 842,200 | 0.119317204 | 0.162599889 | 0.0530381 | 0.12290156 |
| 842,300 | 0.11935712 | 0.163297291 | 0.052698359 | 0.12332628 |
| 842,400 | 0.119323684 | 0.163142651 | 0.053258288 | 0.122906063 |
| 842,500 | 0.119217908 | 0.162874125 | 0.052476447 | 0.12282132 |
| 842,600 | 0.119261912 | 0.162632495 | 0.053276603 | 0.123327017 |
| 842,700 | 0.119287797 | 0.162671672 | 0.053638313 | 0.123957849 |
| 842,800 | 0.119245238 | 0.163238129 | 0.05325518 | 0.12349006 |
| 842,900 | 0.119205969 | 0.162341867 | 0.053397109 | 0.123138509 |
| 843,000 | 0.119219635 | 0.162326602 | 0.052240374 | 0.12385246 |
| 843,100 | 0.119284822 | 0.162558599 | 0.052493277 | 0.123491404 |

| | | | | |
|---|---|---|---|---|
| 843,200 | 0.119224985 | 0.163161358 | 0.052813893 | 0.122889612 |
| 843,300 | 0.11918503 | 0.16271623 | 0.052359555 | 0.122888713 |
| 843,400 | 0.119051866 | 0.16310192 | 0.052720815 | 0.123388787 |
| 843,500 | 0.119134328 | 0.163318377 | 0.053097877 | 0.123143197 |
| 843,600 | 0.119129252 | 0.163993659 | 0.052590997 | 0.123451126 |
| 843,700 | 0.119192181 | 0.16386342 | 0.052551777 | 0.123101218 |
| 843,800 | 0.119246052 | 0.163664743 | 0.052797425 | 0.123365652 |
| 843,900 | 0.119246524 | 0.163367473 | 0.052658266 | 0.123059116 |
| 844,000 | 0.119267716 | 0.163943849 | 0.053098351 | 0.122889422 |
| 844,100 | 0.119188421 | 0.164266768 | 0.052916231 | 0.123385403 |
| 844,200 | 0.119246857 | 0.164365674 | 0.052667836 | 0.122946728 |
| 844,300 | 0.119231988 | 0.163630944 | 0.052783345 | 0.123256843 |
| 844,400 | 0.119221936 | 0.163622805 | 0.052416702 | 0.123428281 |
| 844,500 | 0.119098858 | 0.163243323 | 0.053077111 | 0.12357648 |
| 844,600 | 0.119113651 | 0.163351474 | 0.052789909 | 0.12325768 |
| 844,700 | 0.119154824 | 0.163747463 | 0.052488592 | 0.123379047 |
| 844,800 | 0.119149127 | 0.16435388 | 0.052733763 | 0.123038657 |
| 844,900 | 0.119167472 | 0.163498519 | 0.052602286 | 0.122906034 |
| 845,000 | 0.119032113 | 0.163532456 | 0.05240137 | 0.122781787 |
| 845,100 | 0.118998145 | 0.163412249 | 0.052388245 | 0.122668236 |
| 845,200 | 0.119092991 | 0.164143241 | 0.052859386 | 0.123123889 |
| 845,300 | 0.119113505 | 0.164096728 | 0.052584219 | 0.122965142 |
| 845,400 | 0.119023075 | 0.163920613 | 0.052585443 | 0.123407444 |
| 845,500 | 0.11897744 | 0.163228805 | 0.053088801 | 0.122753297 |
| 845,600 | 0.118874695 | 0.163547218 | 0.053351384 | 0.123135775 |
| 845,700 | 0.118904645 | 0.16373635 | 0.053044993 | 0.122937482 |
| 845,800 | 0.118888803 | 0.163248656 | 0.053160354 | 0.123158753 |
| 845,900 | 0.119060631 | 0.163544582 | 0.052660679 | 0.123505008 |
| 846,000 | 0.118976708 | 0.163555375 | 0.052593194 | 0.123432959 |
| 846,100 | 0.118968483 | 0.163704112 | 0.052481287 | 0.123020605 |
| 846,200 | 0.118976772 | 0.164365207 | 0.05276103 | 0.12319487 |
| 846,300 | 0.11897698 | 0.163986555 | 0.053659419 | 0.123037853 |
| 846,400 | 0.119067575 | 0.164235832 | 0.053298524 | 0.123896075 |
| 846,500 | 0.119054286 | 0.163952508 | 0.053645605 | 0.123190496 |
| 846,600 | 0.119081963 | 0.164265495 | 0.053114025 | 0.123347744 |
| 846,700 | 0.119018304 | 0.16365373 | 0.054025563 | 0.122898732 |
| 846,800 | 0.118990742 | 0.164427353 | 0.052911579 | 0.123249828 |
| 846,900 | 0.119023267 | 0.164683015 | 0.053268298 | 0.12368899 |
| 847,000 | 0.118959139 | 0.164029153 | 0.053099601 | 0.123757379 |
| 847,100 | 0.118787397 | 0.164256106 | 0.053139695 | 0.123637187 |
| 847,200 | 0.118864991 | 0.164234945 | 0.054005834 | 0.123937935 |
| 847,300 | 0.118951287 | 0.164372154 | 0.053472362 | 0.124489955 |
| 847,400 | 0.11888873 | 0.164685698 | 0.052661934 | 0.124228254 |
| 847,500 | 0.118943169 | 0.164908761 | 0.052542634 | 0.124376482 |
| 847,600 | 0.118952674 | 0.164255948 | 0.052468767 | 0.123710485 |
| 847,700 | 0.118823391 | 0.163580808 | 0.052785198 | 0.123498393 |
| 847,800 | 0.118918986 | 0.163673636 | 0.052776232 | 0.123847233 |
| 847,900 | 0.118900721 | 0.163504956 | 0.05271007 | 0.123952063 |
| 848,000 | 0.118889873 | 0.163381602 | 0.052434531 | 0.123833591 |
| 848,100 | 0.118840555 | 0.163020959 | 0.052489001 | 0.123596837 |
| 848,200 | 0.118783154 | 0.163265853 | 0.052654956 | 0.123665176 |
| 848,300 | 0.118758103 | 0.164088285 | 0.052747452 | 0.123565687 |
| 848,400 | 0.118730379 | 0.16314837 | 0.052054223 | 0.12311949 |
| 848,500 | 0.11872311 | 0.162985892 | 0.052261025 | 0.123266877 |
| 848,600 | 0.118782691 | 0.163258386 | 0.051714094 | 0.123086172 |
| 848,700 | 0.1188303 | 0.163521246 | 0.051986597 | 0.123698439 |
| 848,800 | 0.118849621 | 0.163180348 | 0.052467905 | 0.123861506 |
| 848,900 | 0.118691403 | 0.162946789 | 0.052119261 | 0.123515312 |
| 849,000 | 0.118782641 | 0.163288291 | 0.052837285 | 0.123142256 |
| 849,100 | 0.118812731 | 0.163377872 | 0.052584811 | 0.12365454 |
| 849,200 | 0.118716024 | 0.163046435 | 0.052307971 | 0.12348301 |
| 849,300 | 0.118754568 | 0.163199704 | 0.052536182 | 0.123665621 |
| 849,400 | 0.118730477 | 0.163466202 | 0.052499105 | 0.124626187 |
| 849,500 | 0.118664119 | 0.163262807 | 0.052950462 | 0.123811053 |
| 849,600 | 0.118740723 | 0.163137505 | 0.052895894 | 0.123957765 |
| 849,700 | 0.118574449 | 0.163212027 | 0.052988284 | 0.123655087 |
| 849,800 | 0.118656832 | 0.162398641 | 0.052298698 | 0.124117591 |

| | | | | |
|---|---|---|---|---|
| 849,900 | 0.118796972 | 0.162519788 | 0.052294643 | 0.123965423 |
| 850,000 | 0.118696329 | 0.163301767 | 0.052214101 | 0.124494521 |
| 850,100 | 0.118667614 | 0.162946883 | 0.052039096 | 0.124296096 |
| 850,200 | 0.118646926 | 0.16299328 | 0.052408218 | 0.123864561 |
| 850,300 | 0.118606264 | 0.162523775 | 0.052127653 | 0.123734959 |
| 850,400 | 0.118579595 | 0.162550327 | 0.052554708 | 0.124069252 |
| 850,500 | 0.118595753 | 0.163188366 | 0.052649229 | 0.123727826 |
| 850,600 | 0.118470043 | 0.162230874 | 0.052236722 | 0.123764825 |
| 850,700 | 0.118506821 | 0.162618737 | 0.052660496 | 0.123592946 |
| 850,800 | 0.118468737 | 0.162115679 | 0.053051423 | 0.123753732 |
| 850,900 | 0.118424105 | 0.162274945 | 0.052669453 | 0.123741454 |
| 851,000 | 0.118556974 | 0.162544521 | 0.052931616 | 0.124180533 |
| 851,100 | 0.118461271 | 0.162668641 | 0.052617934 | 0.124046127 |
| 851,200 | 0.118420231 | 0.16243432 | 0.053011675 | 0.123627232 |
| 851,300 | 0.118464146 | 0.162587894 | 0.052820735 | 0.123504746 |
| 851,400 | 0.118468842 | 0.162661503 | 0.053377987 | 0.123964879 |
| 851,500 | 0.11850953 | 0.163098339 | 0.053568481 | 0.124184429 |
| 851,600 | 0.118617013 | 0.162735382 | 0.05377653 | 0.124141916 |
| 851,700 | 0.118502096 | 0.162702875 | 0.05322254 | 0.123872934 |
| 851,800 | 0.118436589 | 0.162334552 | 0.053202989 | 0.123998194 |
| 851,900 | 0.118509391 | 0.162715711 | 0.053221964 | 0.124548739 |
| 852,000 | 0.118532067 | 0.162932562 | 0.053466921 | 0.124022293 |
| 852,100 | 0.118475193 | 0.162614853 | 0.05307353 | 0.124333912 |
| 852,200 | 0.118409337 | 0.16309012 | 0.053635139 | 0.124243354 |
| 852,300 | 0.118332893 | 0.162823685 | 0.052873757 | 0.124127191 |
| 852,400 | 0.118404324 | 0.16221041 | 0.053920543 | 0.124100434 |
| 852,500 | 0.118458393 | 0.162833121 | 0.053202904 | 0.124147739 |
| 852,600 | 0.118448369 | 0.163071459 | 0.053627619 | 0.124486337 |
| 852,700 | 0.118481042 | 0.162884517 | 0.053693133 | 0.124027666 |
| 852,800 | 0.118402909 | 0.162845802 | 0.053163303 | 0.124154979 |
| 852,900 | 0.11838026 | 0.162585649 | 0.053603527 | 0.124199944 |
| 853,000 | 0.118440355 | 0.162792899 | 0.053741264 | 0.124680487 |
| 853,100 | 0.118503106 | 0.163154237 | 0.053596228 | 0.124856376 |
| 853,200 | 0.118502486 | 0.163341158 | 0.053742769 | 0.124921349 |
| 853,300 | 0.118388058 | 0.163106156 | 0.053216089 | 0.124581536 |
| 853,400 | 0.118438089 | 0.163223366 | 0.052777665 | 0.124478958 |
| 853,500 | 0.118384507 | 0.163235258 | 0.052986394 | 0.124475448 |
| 853,600 | 0.11846079 | 0.163350594 | 0.052715528 | 0.124985459 |
| 853,700 | 0.118371634 | 0.162298419 | 0.05262273 | 0.124727251 |
| 853,800 | 0.118394576 | 0.162487492 | 0.052117648 | 0.124784172 |
| 853,900 | 0.118301496 | 0.16261098 | 0.052565634 | 0.124503374 |
| 854,000 | 0.118309624 | 0.162053026 | 0.052490242 | 0.124468762 |
| 854,100 | 0.118361487 | 0.16249852 | 0.052671126 | 0.124903536 |
| 854,200 | 0.118242764 | 0.162170383 | 0.053483685 | 0.124500146 |
| 854,300 | 0.118260234 | 0.162467187 | 0.052848222 | 0.1242958 |
| 854,400 | 0.118307644 | 0.162063455 | 0.052553981 | 0.124502625 |
| 854,500 | 0.118307106 | 0.162471169 | 0.052356041 | 0.125144237 |
| 854,600 | 0.118358459 | 0.162478251 | 0.053666549 | 0.124282379 |
| 854,700 | 0.118235463 | 0.161864788 | 0.052845979 | 0.124453198 |
| 854,800 | 0.118189157 | 0.162907422 | 0.053504915 | 0.124500424 |
| 854,900 | 0.118145853 | 0.162775167 | 0.053262714 | 0.124216449 |
| 855,000 | 0.118145531 | 0.16274637 | 0.053084911 | 0.125166353 |
| 855,100 | 0.118228075 | 0.163389189 | 0.053148362 | 0.124640289 |
| 855,200 | 0.118127537 | 0.163399145 | 0.053504601 | 0.124771886 |
| 855,300 | 0.118145116 | 0.163380726 | 0.053146354 | 0.124534027 |
| 855,400 | 0.11821025 | 0.162188877 | 0.052801143 | 0.124802075 |
| 855,500 | 0.11817126 | 0.162928761 | 0.053224377 | 0.124735251 |
| 855,600 | 0.118093473 | 0.162403435 | 0.053240085 | 0.124435396 |
| 855,700 | 0.118012502 | 0.163025877 | 0.053406103 | 0.124622066 |
| 855,800 | 0.118088089 | 0.162697677 | 0.052562105 | 0.124482293 |
| 855,900 | 0.118037785 | 0.16275138 | 0.052999837 | 0.12446579 |
| 856,000 | 0.118075992 | 0.161903389 | 0.052883106 | 0.124858979 |
| 856,100 | 0.117972195 | 0.161819643 | 0.052683301 | 0.124600254 |
| 856,200 | 0.117911993 | 0.162349894 | 0.052449342 | 0.125032476 |
| 856,300 | 0.117917863 | 0.161790058 | 0.053220373 | 0.124597002 |
| 856,400 | 0.117965777 | 0.162188954 | 0.053994381 | 0.124886347 |
| 856,500 | 0.117933767 | 0.161895967 | 0.053069865 | 0.124551084 |

| | | | | |
|---|---|---|---|---|
| 856,600 | 0.118038684 | 0.16227421 | 0.052717834 | 0.124807181 |
| 856,700 | 0.118030859 | 0.161878787 | 0.053024898 | 0.124802817 |
| 856,800 | 0.117998437 | 0.162670906 | 0.053108391 | 0.124503438 |
| 856,900 | 0.117966189 | 0.162103649 | 0.053064066 | 0.125166106 |
| 857,000 | 0.11796772 | 0.162052685 | 0.053312852 | 0.125127607 |
| 857,100 | 0.118013562 | 0.162349223 | 0.053037472 | 0.125576851 |
| 857,200 | 0.117983395 | 0.162241499 | 0.053391759 | 0.125157324 |
| 857,300 | 0.117935466 | 0.161742564 | 0.053360614 | 0.124952714 |
| 857,400 | 0.117969492 | 0.161296431 | 0.053524124 | 0.124362752 |
| 857,500 | 0.118035099 | 0.161393428 | 0.053403807 | 0.124425205 |
| 857,600 | 0.117957396 | 0.161262642 | 0.053155475 | 0.124695816 |
| 857,700 | 0.117922541 | 0.161497785 | 0.053412846 | 0.12496549 |
| 857,800 | 0.117937038 | 0.161345524 | 0.052928889 | 0.124300804 |
| 857,900 | 0.117884346 | 0.161072871 | 0.053051873 | 0.124406626 |
| 858,000 | 0.117947723 | 0.161751343 | 0.053009249 | 0.124752858 |
| 858,100 | 0.117867206 | 0.161614809 | 0.053326679 | 0.124563196 |
| 858,200 | 0.117831317 | 0.16171706 | 0.053113538 | 0.12484571 |
| 858,300 | 0.117774268 | 0.161137352 | 0.053006817 | 0.124458579 |
| 858,400 | 0.117804004 | 0.16080184 | 0.053825825 | 0.1238884 |
| 858,500 | 0.117694735 | 0.161862802 | 0.05385338 | 0.124107723 |
| 858,600 | 0.117704649 | 0.161010903 | 0.053522794 | 0.123748176 |
| 858,700 | 0.117666345 | 0.160745248 | 0.05340368 | 0.123942841 |
| 858,800 | 0.117641144 | 0.16119283 | 0.053568981 | 0.124041652 |
| 858,900 | 0.117682282 | 0.161002645 | 0.053200363 | 0.124083335 |
| 859,000 | 0.117720531 | 0.161381744 | 0.053744171 | 0.124175382 |
| 859,100 | 0.117635929 | 0.160982552 | 0.053281543 | 0.124379989 |
| 859,200 | 0.117665856 | 0.161009606 | 0.053395317 | 0.12400074 |
| 859,300 | 0.117690206 | 0.161398697 | 0.052557886 | 0.12399786 |
| 859,400 | 0.117767757 | 0.161401164 | 0.05262595 | 0.124101035 |
| 859,500 | 0.117865654 | 0.161237313 | 0.053152578 | 0.123884926 |
| 859,600 | 0.117751024 | 0.160884557 | 0.052611195 | 0.124102281 |
| 859,700 | 0.117762899 | 0.161005534 | 0.052629977 | 0.1241008 |
| 859,800 | 0.117775197 | 0.161247213 | 0.052757228 | 0.124761413 |
| 859,900 | 0.117697916 | 0.161513624 | 0.052673039 | 0.124366763 |
| 860,000 | 0.117712938 | 0.161071236 | 0.053059126 | 0.123708232 |
| 860,100 | 0.117773908 | 0.16132407 | 0.053181468 | 0.124625742 |
| 860,200 | 0.117761813 | 0.160945642 | 0.053853245 | 0.123958869 |
| 860,300 | 0.117725027 | 0.161392883 | 0.053409561 | 0.124343213 |
| 860,400 | 0.117689627 | 0.161407519 | 0.05358123 | 0.124897297 |
| 860,500 | 0.117700401 | 0.161177241 | 0.053401145 | 0.124282118 |
| 860,600 | 0.117553024 | 0.160884282 | 0.053294086 | 0.12376057 |
| 860,700 | 0.117574857 | 0.160658059 | 0.053326925 | 0.124286458 |
| 860,800 | 0.117545507 | 0.160734784 | 0.053718184 | 0.123954463 |
| 860,900 | 0.117507983 | 0.161339795 | 0.053223048 | 0.124307101 |
| 861,000 | 0.117496005 | 0.161049994 | 0.053228746 | 0.124023209 |
| 861,100 | 0.117475207 | 0.160962765 | 0.053215502 | 0.12465352 |
| 861,200 | 0.117506469 | 0.161137088 | 0.053606504 | 0.124062412 |
| 861,300 | 0.117471648 | 0.161478445 | 0.053635183 | 0.123751078 |
| 861,400 | 0.117451683 | 0.160919307 | 0.053927303 | 0.123843411 |
| 861,500 | 0.117555519 | 0.161498478 | 0.053887806 | 0.123910097 |
| 861,600 | 0.117696217 | 0.161257329 | 0.053547832 | 0.124725002 |
| 861,700 | 0.117611282 | 0.161603294 | 0.054814094 | 0.124112802 |
| 861,800 | 0.117537036 | 0.161365937 | 0.054706405 | 0.124131377 |
| 861,900 | 0.117595316 | 0.161290053 | 0.054552624 | 0.124837945 |
| 862,000 | 0.117656842 | 0.161782352 | 0.054258907 | 0.124450711 |
| 862,100 | 0.117564884 | 0.161402393 | 0.053596107 | 0.124471789 |
| 862,200 | 0.11755216 | 0.161659529 | 0.053720696 | 0.124668859 |
| 862,300 | 0.117555199 | 0.161338461 | 0.053967029 | 0.124427415 |
| 862,400 | 0.117638066 | 0.160954397 | 0.053879333 | 0.124647516 |
| 862,500 | 0.117491866 | 0.160883461 | 0.054145965 | 0.124436704 |
| 862,600 | 0.117437986 | 0.161407212 | 0.054340721 | 0.124583727 |
| 862,700 | 0.117528496 | 0.161083959 | 0.054400767 | 0.124687204 |
| 862,800 | 0.117584335 | 0.161658825 | 0.053993028 | 0.124567972 |
| 862,900 | 0.117603489 | 0.161255166 | 0.053945067 | 0.124553613 |
| 863,000 | 0.117511288 | 0.160551795 | 0.053777051 | 0.124483107 |
| 863,100 | 0.117425713 | 0.160540902 | 0.053517024 | 0.124171688 |
| 863,200 | 0.117337371 | 0.1608401 | 0.053846182 | 0.123715086 |

| | | | | |
|---|---|---|---|---|
| 863,300 | 0.117417507 | 0.160260509 | 0.053532002 | 0.123948332 |
| 863,400 | 0.117417734 | 0.160536873 | 0.05377439 | 0.124448458 |
| 863,500 | 0.117468831 | 0.160638783 | 0.05380022 | 0.123629097 |
| 863,600 | 0.117462332 | 0.160155404 | 0.053506554 | 0.124103906 |
| 863,700 | 0.117476989 | 0.160187825 | 0.053945171 | 0.124315157 |
| 863,800 | 0.117433365 | 0.160374952 | 0.053293522 | 0.123935706 |
| 863,900 | 0.117422069 | 0.160246153 | 0.053842956 | 0.124059017 |
| 864,000 | 0.117430802 | 0.160077273 | 0.053611391 | 0.124149839 |
| 864,100 | 0.117377878 | 0.160230068 | 0.053521871 | 0.123769695 |
| 864,200 | 0.117237829 | 0.159228422 | 0.053573125 | 0.123982703 |
| 864,300 | 0.117332004 | 0.1600349 | 0.053803746 | 0.124104509 |
| 864,400 | 0.117237658 | 0.160424487 | 0.05365223 | 0.123873546 |
| 864,500 | 0.117388985 | 0.160351409 | 0.053338495 | 0.124164528 |
| 864,600 | 0.11732411 | 0.160274208 | 0.053221101 | 0.124026965 |
| 864,700 | 0.117166715 | 0.16022623 | 0.053367211 | 0.124190857 |
| 864,800 | 0.117244664 | 0.159701844 | 0.053471487 | 0.124054301 |
| 864,900 | 0.117264412 | 0.160298854 | 0.053813673 | 0.124277191 |
| 865,000 | 0.11732542 | 0.159911154 | 0.053701058 | 0.124014463 |
| 865,100 | 0.117151292 | 0.160141644 | 0.053622704 | 0.123935058 |
| 865,200 | 0.117199348 | 0.160317004 | 0.053742516 | 0.123948587 |
| 865,300 | 0.117102719 | 0.160606299 | 0.053537107 | 0.123790885 |
| 865,400 | 0.117181694 | 0.160544038 | 0.053153608 | 0.124491083 |
| 865,500 | 0.117082165 | 0.160384529 | 0.053241227 | 0.124203826 |
| 865,600 | 0.117212158 | 0.161303309 | 0.05341121 | 0.124224723 |
| 865,700 | 0.117183444 | 0.160232388 | 0.053351989 | 0.124780223 |
| 865,800 | 0.117177113 | 0.16056341 | 0.053416584 | 0.124533876 |
| 865,900 | 0.117175405 | 0.160925406 | 0.053627562 | 0.124695881 |
| 866,000 | 0.117176913 | 0.159670969 | 0.052972095 | 0.124621034 |
| 866,100 | 0.117178434 | 0.160554135 | 0.053760098 | 0.124905249 |
| 866,200 | 0.117123573 | 0.160442264 | 0.05381232 | 0.125102433 |
| 866,300 | 0.1171765 | 0.160834126 | 0.053892216 | 0.124822021 |
| 866,400 | 0.117192866 | 0.160239124 | 0.053890529 | 0.12499018 |
| 866,500 | 0.117203813 | 0.160864791 | 0.054104338 | 0.125007724 |
| 866,600 | 0.117167914 | 0.160545065 | 0.053940789 | 0.124621417 |
| 866,700 | 0.117212111 | 0.161055015 | 0.05361983 | 0.124509126 |
| 866,800 | 0.117117431 | 0.160425345 | 0.053992512 | 0.124897177 |
| 866,900 | 0.117158798 | 0.160722032 | 0.053569246 | 0.124880681 |
| 867,000 | 0.117149634 | 0.160713039 | 0.054049815 | 0.125110663 |
| 867,100 | 0.117140001 | 0.160913377 | 0.053142597 | 0.124987864 |
| 867,200 | 0.117138904 | 0.160871347 | 0.053758196 | 0.124672785 |
| 867,300 | 0.117192446 | 0.161445918 | 0.053147437 | 0.124949675 |
| 867,400 | 0.117082269 | 0.16094622 | 0.053477933 | 0.125121771 |
| 867,500 | 0.117113233 | 0.160876734 | 0.053733494 | 0.124902355 |
| 867,600 | 0.117032805 | 0.160349532 | 0.053472287 | 0.124248813 |
| 867,700 | 0.11707453 | 0.16068294 | 0.054077809 | 0.124764605 |
| 867,800 | 0.117033168 | 0.160492737 | 0.05334276 | 0.125075607 |
| 867,900 | 0.11695954 | 0.160005994 | 0.053986552 | 0.124391064 |
| 868,000 | 0.116845518 | 0.159845027 | 0.054143531 | 0.124374612 |
| 868,100 | 0.116888095 | 0.160485496 | 0.053791465 | 0.124545276 |
| 868,200 | 0.117061427 | 0.160875244 | 0.053806527 | 0.124686224 |
| 868,300 | 0.117054701 | 0.161125076 | 0.05456829 | 0.124673517 |
| 868,400 | 0.117099167 | 0.160853593 | 0.053568681 | 0.124611788 |
| 868,500 | 0.11697127 | 0.160858056 | 0.053731091 | 0.124407203 |
| 868,600 | 0.117012319 | 0.160945291 | 0.053626695 | 0.124387077 |
| 868,700 | 0.117048092 | 0.160925146 | 0.053672982 | 0.124457685 |
| 868,800 | 0.11712956 | 0.160923972 | 0.053259994 | 0.125189475 |
| 868,900 | 0.117041406 | 0.161117388 | 0.053092604 | 0.124846811 |
| 869,000 | 0.116974494 | 0.160990607 | 0.053234138 | 0.124371482 |
| 869,100 | 0.116923036 | 0.161442685 | 0.053034334 | 0.12475264 |
| 869,200 | 0.116996455 | 0.161461695 | 0.052720903 | 0.125021643 |
| 869,300 | 0.117054844 | 0.162320594 | 0.053723157 | 0.124596742 |
| 869,400 | 0.116969819 | 0.161498093 | 0.053396394 | 0.124581488 |
| 869,500 | 0.116918713 | 0.161049462 | 0.053587968 | 0.124809004 |
| 869,600 | 0.116821291 | 0.16132998 | 0.053435537 | 0.124601976 |
| 869,700 | 0.116998284 | 0.161345919 | 0.053262692 | 0.124955085 |
| 869,800 | 0.116909946 | 0.1610039 | 0.053644202 | 0.124416425 |
| 869,900 | 0.116953229 | 0.161086115 | 0.053344774 | 0.124624686 |

| | | | | |
|---|---|---|---|---|
| 870,000 | 0.117024937 | 0.161227664 | 0.053317098 | 0.124606482 |
| 870,100 | 0.11705575 | 0.160939346 | 0.05351043 | 0.124506965 |
| 870,200 | 0.116887686 | 0.16098208 | 0.053000422 | 0.124284204 |
| 870,300 | 0.116796569 | 0.160748553 | 0.052826066 | 0.124893663 |
| 870,400 | 0.117070663 | 0.160725048 | 0.052725722 | 0.124648789 |
| 870,500 | 0.117026534 | 0.160712192 | 0.052906912 | 0.125154039 |
| 870,600 | 0.117070661 | 0.161105734 | 0.053229068 | 0.124556214 |
| 870,700 | 0.117187953 | 0.161328922 | 0.053267248 | 0.125016854 |
| 870,800 | 0.117099497 | 0.161233677 | 0.053038274 | 0.124969607 |
| 870,900 | 0.117083399 | 0.160752429 | 0.053014394 | 0.125338994 |
| 871,000 | 0.117078058 | 0.161285502 | 0.053271096 | 0.124992694 |
| 871,100 | 0.117181251 | 0.161137431 | 0.053351916 | 0.124996632 |
| 871,200 | 0.117170637 | 0.161277021 | 0.053217766 | 0.125335422 |
| 871,300 | 0.117105347 | 0.161179337 | 0.053445999 | 0.125390067 |
| 871,400 | 0.117008252 | 0.160894302 | 0.053135056 | 0.125034463 |
| 871,500 | 0.117129064 | 0.16145171 | 0.052891499 | 0.125441858 |
| 871,600 | 0.11716093 | 0.161812322 | 0.053376826 | 0.125219962 |
| 871,700 | 0.117124386 | 0.161455071 | 0.053141037 | 0.124842601 |
| 871,800 | 0.117032857 | 0.161824176 | 0.054191627 | 0.125138971 |
| 871,900 | 0.117118769 | 0.161785536 | 0.053288102 | 0.124640748 |
| 872,000 | 0.117104828 | 0.161743017 | 0.053694112 | 0.124670012 |
| 872,100 | 0.117040767 | 0.161935475 | 0.053729506 | 0.12501509 |
| 872,200 | 0.116902079 | 0.161299496 | 0.053899017 | 0.125118224 |
| 872,300 | 0.117053218 | 0.161316051 | 0.053958595 | 0.124734763 |
| 872,400 | 0.117016156 | 0.161394356 | 0.053542497 | 0.124353082 |
| 872,500 | 0.116980062 | 0.160986846 | 0.053442633 | 0.124405221 |
| 872,600 | 0.116958361 | 0.161905597 | 0.053726825 | 0.124615431 |
| 872,700 | 0.116914564 | 0.162380162 | 0.0535342 | 0.124043246 |
| 872,800 | 0.116906732 | 0.161417618 | 0.054102957 | 0.124008748 |
| 872,900 | 0.116993395 | 0.162176569 | 0.053918861 | 0.124214874 |
| 873,000 | 0.116925371 | 0.161283512 | 0.053734939 | 0.124539615 |
| 873,100 | 0.11682655 | 0.161607339 | 0.054182856 | 0.124471276 |
| 873,200 | 0.116903988 | 0.161319948 | 0.053964284 | 0.124493317 |
| 873,300 | 0.116927821 | 0.161356063 | 0.053237503 | 0.124484525 |
| 873,400 | 0.11693911 | 0.161538837 | 0.053194019 | 0.124732784 |
| 873,500 | 0.116855133 | 0.161401767 | 0.053253479 | 0.124766152 |
| 873,600 | 0.116681522 | 0.160829877 | 0.05313086 | 0.123976438 |
| 873,700 | 0.11680948 | 0.161344483 | 0.05281953 | 0.124367059 |
| 873,800 | 0.116852577 | 0.160916016 | 0.053275862 | 0.124561125 |
| 873,900 | 0.116758422 | 0.161055437 | 0.052793231 | 0.124370104 |
| 874,000 | 0.116834687 | 0.161657271 | 0.052752509 | 0.124702744 |
| 874,100 | 0.116839583 | 0.161661 | 0.053216005 | 0.12456483 |
| 874,200 | 0.116733522 | 0.161560471 | 0.053503642 | 0.12420928 |
| 874,300 | 0.116842449 | 0.161325388 | 0.05389009 | 0.12477757 |
| 874,400 | 0.116790733 | 0.161282188 | 0.053384622 | 0.124698475 |
| 874,500 | 0.116781992 | 0.161422785 | 0.05314633 | 0.12464373 |
| 874,600 | 0.116834458 | 0.161629516 | 0.053039896 | 0.124859511 |
| 874,700 | 0.116763269 | 0.160713113 | 0.053056117 | 0.124264642 |
| 874,800 | 0.116724388 | 0.160818437 | 0.053443665 | 0.124300787 |
| 874,900 | 0.116688992 | 0.16125573 | 0.052916354 | 0.124791927 |
| 875,000 | 0.116590483 | 0.16076446 | 0.053057996 | 0.124249708 |
| 875,100 | 0.116646429 | 0.160976931 | 0.053392264 | 0.125023363 |
| 875,200 | 0.116653543 | 0.160773177 | 0.053286458 | 0.124819502 |
| 875,300 | 0.116485055 | 0.160719495 | 0.053242805 | 0.125280098 |
| 875,400 | 0.116554535 | 0.1607429 | 0.053586626 | 0.125355945 |
| 875,500 | 0.116605131 | 0.160675164 | 0.053468224 | 0.125353156 |
| 875,600 | 0.116583252 | 0.161017253 | 0.053831267 | 0.124520074 |
| 875,700 | 0.116627889 | 0.160899592 | 0.053739436 | 0.12459103 |
| 875,800 | 0.116565079 | 0.160607944 | 0.053119753 | 0.124807103 |
| 875,900 | 0.116538308 | 0.160723781 | 0.053249458 | 0.124463134 |
| 876,000 | 0.11652625 | 0.160927872 | 0.052976359 | 0.124849604 |
| 876,100 | 0.116446085 | 0.160607419 | 0.052764109 | 0.124740722 |
| 876,200 | 0.116568935 | 0.161546877 | 0.052548268 | 0.124768633 |
| 876,300 | 0.116598634 | 0.16155844 | 0.054034221 | 0.124782859 |
| 876,400 | 0.11646779 | 0.161228275 | 0.053344771 | 0.124767404 |
| 876,500 | 0.11655175 | 0.161215941 | 0.053227925 | 0.124571948 |
| 876,600 | 0.116469323 | 0.160884572 | 0.053584054 | 0.124366323 |

| | | | | |
|---|---|---|---|---|
| 876,700 | 0.116565717 | 0.160871962 | 0.054091719 | 0.124786053 |
| 876,800 | 0.116479056 | 0.160311924 | 0.053582243 | 0.125063547 |
| 876,900 | 0.116403984 | 0.16127895 | 0.052850992 | 0.124400936 |
| 877,000 | 0.116513087 | 0.160909177 | 0.052962543 | 0.125001818 |
| 877,100 | 0.116420399 | 0.160836485 | 0.052397071 | 0.124256086 |
| 877,200 | 0.116401844 | 0.160670418 | 0.053748359 | 0.124493894 |
| 877,300 | 0.116476184 | 0.160824611 | 0.053017633 | 0.124625085 |
| 877,400 | 0.116395658 | 0.160413727 | 0.053630024 | 0.124486261 |
| 877,500 | 0.116283219 | 0.160706363 | 0.053213386 | 0.124294481 |
| 877,600 | 0.116293408 | 0.160181848 | 0.052777284 | 0.124233171 |
| 877,700 | 0.116321611 | 0.159803662 | 0.05258922 | 0.124418863 |
| 877,800 | 0.116317956 | 0.160110125 | 0.05300903 | 0.124535732 |
| 877,900 | 0.116323657 | 0.159977202 | 0.052870762 | 0.124185436 |
| 878,000 | 0.116239432 | 0.159734097 | 0.053434489 | 0.124390772 |
| 878,100 | 0.116275746 | 0.159970358 | 0.052564912 | 0.124533181 |
| 878,200 | 0.116364169 | 0.159969318 | 0.052519343 | 0.124541723 |
| 878,300 | 0.116179224 | 0.160088862 | 0.053103543 | 0.124739374 |
| 878,400 | 0.116261866 | 0.160293338 | 0.053430286 | 0.124844459 |
| 878,500 | 0.1163129 | 0.160424298 | 0.053482115 | 0.124556133 |
| 878,600 | 0.11628218 | 0.159999038 | 0.052869048 | 0.125238983 |
| 878,700 | 0.116320815 | 0.160882191 | 0.053108163 | 0.124793802 |
| 878,800 | 0.116218354 | 0.160400163 | 0.053316861 | 0.124654999 |
| 878,900 | 0.116333688 | 0.160293872 | 0.052934162 | 0.124713255 |
| 879,000 | 0.116172739 | 0.160027037 | 0.052785105 | 0.124948605 |
| 879,100 | 0.116279178 | 0.160411703 | 0.052761553 | 0.124563575 |
| 879,200 | 0.116195139 | 0.159630414 | 0.052274086 | 0.125016411 |
| 879,300 | 0.116177349 | 0.159536738 | 0.052821017 | 0.124943406 |
| 879,400 | 0.116234829 | 0.159614144 | 0.053039978 | 0.124701943 |
| 879,500 | 0.116172711 | 0.159692886 | 0.053139543 | 0.124229627 |
| 879,600 | 0.116176221 | 0.159744899 | 0.052904187 | 0.124171083 |
| 879,700 | 0.116158182 | 0.160558089 | 0.052465814 | 0.124399668 |
| 879,800 | 0.116131559 | 0.160156216 | 0.05223936 | 0.124023327 |
| 879,900 | 0.116165372 | 0.15996767 | 0.052485916 | 0.124248782 |
| 880,000 | 0.116243593 | 0.160300803 | 0.052567346 | 0.123879472 |
| 880,100 | 0.116083391 | 0.159664016 | 0.051959507 | 0.124459499 |
| 880,200 | 0.116059357 | 0.16019293 | 0.052386657 | 0.123986442 |
| 880,300 | 0.116061991 | 0.159124504 | 0.05232204 | 0.124217919 |
| 880,400 | 0.116070716 | 0.160317239 | 0.052129883 | 0.12412259 |
| 880,500 | 0.116023771 | 0.15945726 | 0.051942048 | 0.123741324 |
| 880,600 | 0.116074349 | 0.159518312 | 0.052210228 | 0.124174971 |
| 880,700 | 0.115984858 | 0.159338202 | 0.052114239 | 0.123546917 |
| 880,800 | 0.11595957 | 0.159750928 | 0.052663563 | 0.123299181 |
| 880,900 | 0.115886195 | 0.159570492 | 0.052215823 | 0.123500539 |
| 881,000 | 0.115988554 | 0.159821909 | 0.052146886 | 0.123685146 |
| 881,100 | 0.115877833 | 0.159090048 | 0.052600495 | 0.123212968 |
| 881,200 | 0.115920261 | 0.158957538 | 0.052538755 | 0.123455472 |
| 881,300 | 0.115843499 | 0.15902554 | 0.052241309 | 0.123247249 |
| 881,400 | 0.115914678 | 0.159293233 | 0.051693547 | 0.123694764 |
| 881,500 | 0.115956415 | 0.158766755 | 0.051916152 | 0.123447432 |
| 881,600 | 0.115953849 | 0.158581965 | 0.052100137 | 0.123516801 |
| 881,700 | 0.116033492 | 0.159052547 | 0.052329713 | 0.123807645 |
| 881,800 | 0.115993146 | 0.158509865 | 0.051799767 | 0.123327863 |
| 881,900 | 0.11598765 | 0.158182331 | 0.051817278 | 0.123768685 |
| 882,000 | 0.115927733 | 0.159020031 | 0.051803751 | 0.123542151 |
| 882,100 | 0.116000523 | 0.158891424 | 0.051350539 | 0.123480045 |
| 882,200 | 0.116121107 | 0.159019738 | 0.051132132 | 0.123678195 |
| 882,300 | 0.116085172 | 0.159127323 | 0.051338436 | 0.123321568 |
| 882,400 | 0.116022309 | 0.15959257 | 0.051040817 | 0.122986658 |
| 882,500 | 0.116062476 | 0.158897565 | 0.051577176 | 0.123181404 |
| 882,600 | 0.116085335 | 0.158917891 | 0.05145213 | 0.123516565 |
| 882,700 | 0.115919704 | 0.158582926 | 0.051433202 | 0.123671294 |
| 882,800 | 0.116012585 | 0.159137958 | 0.051174775 | 0.123512232 |
| 882,900 | 0.11596066 | 0.159063282 | 0.051336504 | 0.123173979 |
| 883,000 | 0.115942706 | 0.159031929 | 0.050900866 | 0.123215583 |
| 883,100 | 0.115954706 | 0.159054649 | 0.051017448 | 0.123176762 |
| 883,200 | 0.115881959 | 0.158599262 | 0.051090758 | 0.123321297 |
| 883,300 | 0.115806125 | 0.158575097 | 0.050474461 | 0.123319922 |

| | | | | |
|---|---|---|---|---|
| 883,400 | 0.115789389 | 0.158747624 | 0.050790576 | 0.122972089 |
| 883,500 | 0.115779525 | 0.158902552 | 0.050999172 | 0.123169087 |
| 883,600 | 0.115809234 | 0.159322997 | 0.051195453 | 0.122520645 |
| 883,700 | 0.115785867 | 0.158702869 | 0.050408337 | 0.12273763 |
| 883,800 | 0.115810973 | 0.158459842 | 0.051115183 | 0.123616609 |
| 883,900 | 0.115663016 | 0.157901281 | 0.050495676 | 0.122966124 |
| 884,000 | 0.11583383 | 0.158994668 | 0.050250566 | 0.123514651 |
| 884,100 | 0.115720416 | 0.158262088 | 0.050435806 | 0.123389221 |
| 884,200 | 0.115926955 | 0.158483547 | 0.051146919 | 0.12347518 |
| 884,300 | 0.115909209 | 0.158644899 | 0.050869764 | 0.123237049 |
| 884,400 | 0.115817306 | 0.158530954 | 0.050777367 | 0.123460576 |
| 884,500 | 0.115825611 | 0.158622544 | 0.05093715 | 0.123256145 |
| 884,600 | 0.115821059 | 0.158550774 | 0.050773333 | 0.12351048 |
| 884,700 | 0.115860936 | 0.158609575 | 0.050031522 | 0.123820545 |
| 884,800 | 0.115855616 | 0.158673231 | 0.050410258 | 0.123310035 |
| 884,900 | 0.115851238 | 0.158415902 | 0.050583696 | 0.123562192 |
| 885,000 | 0.115804129 | 0.159273132 | 0.05067321 | 0.123571751 |
| 885,100 | 0.115775269 | 0.158481556 | 0.050697288 | 0.123813072 |
| 885,200 | 0.115823366 | 0.158362399 | 0.050561052 | 0.124134897 |
| 885,300 | 0.115876308 | 0.158761583 | 0.050518521 | 0.123710654 |
| 885,400 | 0.115756255 | 0.158130614 | 0.050458135 | 0.123792141 |
| 885,500 | 0.115824128 | 0.15804601 | 0.049844419 | 0.123826177 |
| 885,600 | 0.115803786 | 0.158023939 | 0.050065157 | 0.123706123 |
| 885,700 | 0.115845483 | 0.158045011 | 0.050311364 | 0.123528836 |
| 885,800 | 0.11570529 | 0.157378446 | 0.050647583 | 0.124109227 |
| 885,900 | 0.115758905 | 0.157717666 | 0.050072905 | 0.12348714 |
| 886,000 | 0.115849572 | 0.158338954 | 0.050139312 | 0.1236751 |
| 886,100 | 0.115782934 | 0.158078157 | 0.050359015 | 0.124028849 |
| 886,200 | 0.115756762 | 0.158033228 | 0.050211228 | 0.123644972 |
| 886,300 | 0.115676667 | 0.158474493 | 0.050368588 | 0.123815354 |
| 886,400 | 0.115732568 | 0.158231418 | 0.050046335 | 0.123577654 |
| 886,500 | 0.115789252 | 0.158907989 | 0.050479515 | 0.123606023 |
| 886,600 | 0.115756282 | 0.158289629 | 0.050246234 | 0.124315993 |
| 886,700 | 0.115693941 | 0.15738349 | 0.049893464 | 0.123532656 |
| 886,800 | 0.115746791 | 0.158277493 | 0.050544802 | 0.124091958 |
| 886,900 | 0.115618617 | 0.158788686 | 0.050050865 | 0.123860069 |
| 887,000 | 0.115635461 | 0.157957005 | 0.050264351 | 0.123459053 |
| 887,100 | 0.115675878 | 0.158660292 | 0.050153072 | 0.123760301 |
| 887,200 | 0.115668586 | 0.158274788 | 0.049810028 | 0.123838791 |
| 887,300 | 0.115692886 | 0.158238768 | 0.050207153 | 0.123394088 |
| 887,400 | 0.115559524 | 0.157688232 | 0.049784758 | 0.123751576 |
| 887,500 | 0.115532564 | 0.158176725 | 0.049761102 | 0.124050782 |
| 887,600 | 0.115590741 | 0.157800088 | 0.0497135 | 0.123528196 |
| 887,700 | 0.115590699 | 0.157923328 | 0.049909894 | 0.123696909 |
| 887,800 | 0.115589899 | 0.157557085 | 0.049580366 | 0.123620772 |
| 887,900 | 0.115639408 | 0.157963852 | 0.049966885 | 0.124046277 |
| 888,000 | 0.115624587 | 0.157035963 | 0.049806315 | 0.123357518 |
| 888,100 | 0.11570994 | 0.157542266 | 0.049758625 | 0.123523289 |
| 888,200 | 0.115581226 | 0.157036209 | 0.04921088 | 0.1231641 |
| 888,300 | 0.115498723 | 0.157233083 | 0.049574205 | 0.123143824 |
| 888,400 | 0.115637631 | 0.157608478 | 0.049708281 | 0.123967367 |
| 888,500 | 0.115441359 | 0.157645429 | 0.049620593 | 0.123257099 |
| 888,600 | 0.115585025 | 0.158438749 | 0.050153266 | 0.123020114 |
| 888,700 | 0.115544623 | 0.157670216 | 0.049802003 | 0.123100409 |
| 888,800 | 0.115482967 | 0.157901523 | 0.05029223 | 0.123135208 |
| 888,900 | 0.115690999 | 0.15795825 | 0.050249405 | 0.123164247 |
| 889,000 | 0.115652814 | 0.157918646 | 0.050168367 | 0.123310076 |
| 889,100 | 0.115620216 | 0.157974406 | 0.050295742 | 0.123262266 |
| 889,200 | 0.115607221 | 0.158232475 | 0.050307527 | 0.123389654 |
| 889,300 | 0.115518472 | 0.158311833 | 0.050526766 | 0.123137238 |
| 889,400 | 0.115517929 | 0.157879292 | 0.050213533 | 0.123331093 |
| 889,500 | 0.115439545 | 0.157652595 | 0.050097056 | 0.123294336 |
| 889,600 | 0.115531586 | 0.158914195 | 0.05032501 | 0.123744219 |
| 889,700 | 0.115512535 | 0.158082636 | 0.05039474 | 0.123882438 |
| 889,800 | 0.115532079 | 0.158513994 | 0.050256462 | 0.123161161 |
| 889,900 | 0.115484208 | 0.157895191 | 0.050189758 | 0.123310068 |
| 890,000 | 0.115448755 | 0.158214896 | 0.04975138 | 0.12276244 |

| | | | | |
|---|---|---|---|---|
| 890,100 | 0.115489711 | 0.158559006 | 0.050144318 | 0.122881433 |
| 890,200 | 0.115399404 | 0.158036317 | 0.050053536 | 0.122813676 |
| 890,300 | 0.115500188 | 0.158632788 | 0.049974059 | 0.12274855 |
| 890,400 | 0.115377495 | 0.158676918 | 0.050162343 | 0.123304667 |
| 890,500 | 0.115364005 | 0.158359668 | 0.050030956 | 0.123333871 |
| 890,600 | 0.11537831 | 0.158296347 | 0.049770617 | 0.122483136 |
| 890,700 | 0.11542625 | 0.158486305 | 0.049844433 | 0.122623964 |
| 890,800 | 0.115312751 | 0.157834373 | 0.050287487 | 0.122566007 |
| 890,900 | 0.11538959 | 0.15787321 | 0.050099931 | 0.122695765 |
| 891,000 | 0.115362964 | 0.157430481 | 0.050336123 | 0.122719299 |
| 891,100 | 0.115239431 | 0.157354866 | 0.049830981 | 0.123488893 |
| 891,200 | 0.115329339 | 0.157789878 | 0.049914968 | 0.123082094 |
| 891,300 | 0.115334003 | 0.157165491 | 0.050270788 | 0.123222546 |
| 891,400 | 0.115464887 | 0.158231021 | 0.050063986 | 0.123676857 |
| 891,500 | 0.115479297 | 0.15774259 | 0.049968466 | 0.12361583 |
| 891,600 | 0.11536524 | 0.158010261 | 0.049999235 | 0.122902451 |
| 891,700 | 0.115339799 | 0.158476908 | 0.04979289 | 0.122675693 |
| 891,800 | 0.115370641 | 0.158390688 | 0.050541812 | 0.123034455 |
| 891,900 | 0.115288945 | 0.157144847 | 0.049802047 | 0.123426075 |
| 892,000 | 0.115325118 | 0.158213146 | 0.050193814 | 0.122700741 |
| 892,100 | 0.115300268 | 0.158422845 | 0.050386415 | 0.122540227 |
| 892,200 | 0.115260774 | 0.15842176 | 0.049914252 | 0.122818565 |
| 892,300 | 0.115435568 | 0.158291383 | 0.049593374 | 0.122781542 |
| 892,400 | 0.115352471 | 0.158457943 | 0.04970694 | 0.12325435 |
| 892,500 | 0.115413897 | 0.157997679 | 0.049599076 | 0.123303572 |
| 892,600 | 0.115271429 | 0.15810092 | 0.050351696 | 0.123130656 |
| 892,700 | 0.115280913 | 0.158088224 | 0.050024399 | 0.123154304 |
| 892,800 | 0.115413228 | 0.158608044 | 0.050177378 | 0.122724624 |
| 892,900 | 0.115362096 | 0.158343714 | 0.049865958 | 0.122972342 |
| 893,000 | 0.115332388 | 0.158045332 | 0.050028472 | 0.122993299 |
| 893,100 | 0.115359992 | 0.158300084 | 0.049801351 | 0.122853312 |
| 893,200 | 0.115402382 | 0.158373083 | 0.050062631 | 0.123013413 |
| 893,300 | 0.115347146 | 0.1584298 | 0.049778194 | 0.122873868 |
| 893,400 | 0.115347959 | 0.158523696 | 0.049068091 | 0.123167014 |
| 893,500 | 0.115241931 | 0.157801998 | 0.049067191 | 0.122709449 |
| 893,600 | 0.115221088 | 0.158032305 | 0.048965526 | 0.122821032 |
| 893,700 | 0.115192327 | 0.157524107 | 0.049386534 | 0.122779382 |
| 893,800 | 0.115155216 | 0.157895253 | 0.049743986 | 0.122429939 |
| 893,900 | 0.115164534 | 0.157933336 | 0.049332841 | 0.122955999 |
| 894,000 | 0.115139598 | 0.157954535 | 0.049968239 | 0.123218491 |
| 894,100 | 0.115183627 | 0.15793596 | 0.049453757 | 0.123350341 |
| 894,200 | 0.115058047 | 0.157956007 | 0.050093751 | 0.122943714 |
| 894,300 | 0.115107354 | 0.158255605 | 0.05001158 | 0.122464641 |
| 894,400 | 0.115013622 | 0.158480277 | 0.049758701 | 0.122484469 |
| 894,500 | 0.115037912 | 0.158103805 | 0.049743521 | 0.122678828 |
| 894,600 | 0.115138095 | 0.158359858 | 0.049976375 | 0.122524152 |
| 894,700 | 0.115049918 | 0.158403858 | 0.049237452 | 0.122141643 |
| 894,800 | 0.115018259 | 0.158491494 | 0.049520964 | 0.122018837 |
| 894,900 | 0.115048453 | 0.158279139 | 0.049725204 | 0.121736087 |
| 895,000 | 0.115074517 | 0.158603901 | 0.049678773 | 0.122016755 |
| 895,100 | 0.115018462 | 0.15784665 | 0.049995813 | 0.121977118 |
| 895,200 | 0.114974463 | 0.15763069 | 0.050393428 | 0.122199388 |
| 895,300 | 0.115004906 | 0.157877543 | 0.049765966 | 0.122388433 |
| 895,400 | 0.11501283 | 0.157979843 | 0.050293255 | 0.122093607 |
| 895,500 | 0.115089743 | 0.159193243 | 0.050153879 | 0.122393583 |
| 895,600 | 0.115008421 | 0.158712606 | 0.050132995 | 0.121885557 |
| 895,700 | 0.115001328 | 0.15846768 | 0.050241939 | 0.122232225 |
| 895,800 | 0.115111718 | 0.158311361 | 0.050076698 | 0.122264258 |
| 895,900 | 0.115003018 | 0.157615176 | 0.050089247 | 0.122088012 |
| 896,000 | 0.114969423 | 0.158537523 | 0.049660801 | 0.122016854 |
| 896,100 | 0.114896812 | 0.157899113 | 0.05010078 | 0.12190861 |
| 896,200 | 0.115035967 | 0.158016376 | 0.050624353 | 0.122507114 |
| 896,300 | 0.115054201 | 0.157390916 | 0.050798206 | 0.122136111 |
| 896,400 | 0.114994932 | 0.157703796 | 0.050517468 | 0.122356223 |
| 896,500 | 0.114959928 | 0.157628207 | 0.050739998 | 0.12238897 |
| 896,600 | 0.114906862 | 0.157795964 | 0.050521128 | 0.122367266 |
| 896,700 | 0.115055179 | 0.158427796 | 0.050169744 | 0.122162853 |

| | | | | |
|---|---|---|---|---|
| 896,800 | 0.11494206 | 0.15793077 | 0.050367277 | 0.122196147 |
| 896,900 | 0.114927936 | 0.157560507 | 0.050525591 | 0.122229745 |
| 897,000 | 0.114871468 | 0.157432577 | 0.050632431 | 0.122248835 |
| 897,100 | 0.115047904 | 0.157938919 | 0.050218435 | 0.12275181 |
| 897,200 | 0.114962427 | 0.157579795 | 0.050719546 | 0.123110146 |
| 897,300 | 0.114993651 | 0.158233494 | 0.050836609 | 0.122336162 |
| 897,400 | 0.114999985 | 0.15815827 | 0.050548668 | 0.122632309 |
| 897,500 | 0.114956169 | 0.158219407 | 0.050351101 | 0.122511411 |
| 897,600 | 0.114938713 | 0.15828514 | 0.050434996 | 0.122638948 |
| 897,700 | 0.114834054 | 0.158501784 | 0.050244941 | 0.12249423 |
| 897,800 | 0.1148105 | 0.158161484 | 0.04991693 | 0.122292882 |
| 897,900 | 0.114785533 | 0.15887549 | 0.049890128 | 0.122160713 |
| 898,000 | 0.114806506 | 0.158645747 | 0.049818298 | 0.121681387 |
| 898,100 | 0.114915514 | 0.158998001 | 0.049997424 | 0.122636246 |
| 898,200 | 0.114784251 | 0.158540769 | 0.050217662 | 0.121653834 |
| 898,300 | 0.114730545 | 0.158684473 | 0.049935645 | 0.121894481 |
| 898,400 | 0.114791453 | 0.158853676 | 0.050066212 | 0.122484906 |
| 898,500 | 0.114823764 | 0.158472687 | 0.049836729 | 0.122372714 |
| 898,600 | 0.114823861 | 0.158442565 | 0.049870186 | 0.122429775 |
| 898,700 | 0.114777886 | 0.159033299 | 0.050126773 | 0.122673021 |
| 898,800 | 0.114736637 | 0.158921547 | 0.04990419 | 0.122513024 |
| 898,900 | 0.114660911 | 0.158761721 | 0.049921767 | 0.122387473 |
| 899,000 | 0.114681256 | 0.158786212 | 0.049753529 | 0.12155857 |
| 899,100 | 0.114750162 | 0.158453124 | 0.050157165 | 0.12158494 |
| 899,200 | 0.114707101 | 0.158711283 | 0.050001158 | 0.122078079 |
| 899,300 | 0.114645189 | 0.158592436 | 0.049271087 | 0.121746167 |
| 899,400 | 0.114704761 | 0.158661444 | 0.05005297 | 0.121566805 |
| 899,500 | 0.114704781 | 0.158316294 | 0.0498179 | 0.121818869 |
| 899,600 | 0.114758452 | 0.15882597 | 0.049484769 | 0.121466755 |
| 899,700 | 0.114887894 | 0.159064016 | 0.049805891 | 0.121963638 |
| 899,800 | 0.114839614 | 0.158552384 | 0.049638175 | 0.121523887 |
| 899,900 | 0.114828463 | 0.158170606 | 0.050195642 | 0.121675311 |
| 900,000 | 0.114790615 | 0.158147262 | 0.050167269 | 0.121943847 |
| 900,100 | 0.114722566 | 0.157618866 | 0.049896964 | 0.121790408 |
| 900,200 | 0.114782266 | 0.158245202 | 0.050486983 | 0.121719234 |
| 900,300 | 0.114726501 | 0.157392464 | 0.050346267 | 0.121568672 |
| 900,400 | 0.114793224 | 0.157729988 | 0.050149719 | 0.121280975 |
| 900,500 | 0.114757543 | 0.157771508 | 0.050011765 | 0.121254075 |
| 900,600 | 0.11471661 | 0.157386962 | 0.050136141 | 0.121670014 |
| 900,700 | 0.114729513 | 0.157385498 | 0.050325207 | 0.121887305 |
| 900,800 | 0.114690663 | 0.156898187 | 0.050195967 | 0.122020625 |
| 900,900 | 0.114747014 | 0.157831329 | 0.049946784 | 0.121861203 |
| 901,000 | 0.114698041 | 0.15782749 | 0.050018792 | 0.121740924 |
| 901,100 | 0.114578622 | 0.157633683 | 0.05014176 | 0.121395978 |
| 901,200 | 0.114610276 | 0.158117838 | 0.050637342 | 0.121088918 |
| 901,300 | 0.114564803 | 0.158156276 | 0.050384016 | 0.121085833 |
| 901,400 | 0.114621114 | 0.157441361 | 0.049945016 | 0.121420789 |
| 901,500 | 0.114558922 | 0.157623105 | 0.049969399 | 0.121161651 |
| 901,600 | 0.114572618 | 0.15690523 | 0.049737312 | 0.121338732 |
| 901,700 | 0.11461588 | 0.157264529 | 0.049892183 | 0.121388618 |
| 901,800 | 0.114541308 | 0.156909074 | 0.04991377 | 0.121175154 |
| 901,900 | 0.114649046 | 0.156954984 | 0.050813708 | 0.121551637 |
| 902,000 | 0.114612107 | 0.157193758 | 0.049878671 | 0.121561555 |
| 902,100 | 0.114711109 | 0.15697144 | 0.049779266 | 0.121642198 |
| 902,200 | 0.114581852 | 0.156838994 | 0.050429034 | 0.121751588 |
| 902,300 | 0.114620064 | 0.157093583 | 0.050173317 | 0.121690471 |
| 902,400 | 0.114635547 | 0.157056731 | 0.05026521 | 0.121688394 |
| 902,500 | 0.114617115 | 0.157430614 | 0.050026944 | 0.121309534 |
| 902,600 | 0.114597182 | 0.157138065 | 0.049706349 | 0.121174333 |
| 902,700 | 0.114529104 | 0.157685316 | 0.049877706 | 0.121266013 |
| 902,800 | 0.114532751 | 0.157113911 | 0.049688975 | 0.121393132 |
| 902,900 | 0.114432642 | 0.157329557 | 0.049709272 | 0.121048503 |
| 903,000 | 0.114517997 | 0.157262448 | 0.049733016 | 0.121260806 |
| 903,100 | 0.114633726 | 0.157617795 | 0.050456647 | 0.1215642 |
| 903,200 | 0.114668282 | 0.157704096 | 0.050130713 | 0.121443314 |
| 903,300 | 0.11457783 | 0.157976134 | 0.050171404 | 0.12157805 |
| 903,400 | 0.114545395 | 0.157828308 | 0.050047239 | 0.121458003 |

| | | | | |
|---|---|---|---|---|
| 903,500 | 0.114578803 | 0.15731743 | 0.049718293 | 0.121932699 |
| 903,600 | 0.114563863 | 0.157235621 | 0.050351163 | 0.121818154 |
| 903,700 | 0.114462527 | 0.157017774 | 0.049969386 | 0.121594271 |
| 903,800 | 0.114514848 | 0.156986973 | 0.050222282 | 0.121842586 |
| 903,900 | 0.114494634 | 0.156985235 | 0.050161181 | 0.121907471 |
| 904,000 | 0.114633319 | 0.157372678 | 0.050441457 | 0.121562665 |
| 904,100 | 0.11454424 | 0.157441392 | 0.050052165 | 0.121462865 |
| 904,200 | 0.114481691 | 0.157781704 | 0.049972471 | 0.121103981 |
| 904,300 | 0.114479211 | 0.157402433 | 0.050291757 | 0.120918742 |
| 904,400 | 0.114479034 | 0.157589857 | 0.050482256 | 0.121384865 |
| 904,500 | 0.114409302 | 0.156897482 | 0.050186227 | 0.121116502 |
| 904,600 | 0.114372682 | 0.15735676 | 0.049998334 | 0.120995223 |
| 904,700 | 0.11431002 | 0.156713096 | 0.050632313 | 0.121005858 |
| 904,800 | 0.114334732 | 0.156699725 | 0.050179508 | 0.121479201 |
| 904,900 | 0.11439098 | 0.157239828 | 0.049876943 | 0.121983158 |
| 905,000 | 0.114398256 | 0.156971902 | 0.049355416 | 0.121595879 |
| 905,100 | 0.11439568 | 0.157097553 | 0.050047631 | 0.12104232 |
| 905,200 | 0.11447753 | 0.158121247 | 0.050619845 | 0.121211191 |
| 905,300 | 0.114479011 | 0.158111349 | 0.050354791 | 0.121294958 |
| 905,400 | 0.114317773 | 0.158024104 | 0.049695097 | 0.121249124 |
| 905,500 | 0.114243674 | 0.158200347 | 0.049738686 | 0.120741537 |
| 905,600 | 0.114356557 | 0.158319836 | 0.049655937 | 0.121174214 |
| 905,700 | 0.11435303 | 0.159140408 | 0.049646528 | 0.121036652 |
| 905,800 | 0.114398354 | 0.158524956 | 0.049392028 | 0.120627779 |
| 905,900 | 0.114343029 | 0.158008218 | 0.049511222 | 0.121009055 |
| 906,000 | 0.114435335 | 0.158305128 | 0.050031361 | 0.121261238 |
| 906,100 | 0.114448896 | 0.159018084 | 0.049582964 | 0.121379514 |
| 906,200 | 0.114425265 | 0.158296643 | 0.050027905 | 0.120956732 |
| 906,300 | 0.114457896 | 0.158553152 | 0.050044526 | 0.121039225 |
| 906,400 | 0.114321949 | 0.15779559 | 0.049400648 | 0.121023259 |
| 906,500 | 0.114331147 | 0.157628424 | 0.050302098 | 0.120899663 |
| 906,600 | 0.114172411 | 0.158075581 | 0.049891807 | 0.120579135 |
| 906,700 | 0.114249883 | 0.157965114 | 0.049837798 | 0.1210737 |
| 906,800 | 0.114179079 | 0.158086879 | 0.050357541 | 0.120909698 |
| 906,900 | 0.114270325 | 0.158045087 | 0.050391118 | 0.120930362 |
| 907,000 | 0.114227853 | 0.157914601 | 0.050233793 | 0.121110722 |
| 907,100 | 0.114173429 | 0.158395162 | 0.050817684 | 0.12026572 |
| 907,200 | 0.114232609 | 0.157625862 | 0.050360779 | 0.120647399 |
| 907,300 | 0.114194967 | 0.157467305 | 0.050748684 | 0.120557304 |
| 907,400 | 0.114239804 | 0.158487961 | 0.050050036 | 0.120304876 |
| 907,500 | 0.114257843 | 0.158276421 | 0.050099005 | 0.120353138 |
| 907,600 | 0.114292117 | 0.158013603 | 0.050082872 | 0.120797652 |
| 907,700 | 0.114297641 | 0.15828794 | 0.04980431 | 0.120486222 |
| 907,800 | 0.114261167 | 0.15763505 | 0.050074873 | 0.120713393 |
| 907,900 | 0.114244706 | 0.158118684 | 0.04999984 | 0.120265872 |
| 908,000 | 0.114206755 | 0.157981298 | 0.050130854 | 0.12082937 |
| 908,100 | 0.114235783 | 0.158220325 | 0.049847002 | 0.120765173 |
| 908,200 | 0.114220232 | 0.158025919 | 0.049910667 | 0.120539244 |
| 908,300 | 0.114233296 | 0.158125132 | 0.050245585 | 0.120741593 |
| 908,400 | 0.114248146 | 0.157713798 | 0.049920096 | 0.120920116 |
| 908,500 | 0.114226241 | 0.15775834 | 0.050208351 | 0.12050283 |
| 908,600 | 0.114261273 | 0.158548839 | 0.049777591 | 0.12082118 |
| 908,700 | 0.114162022 | 0.157735807 | 0.050088287 | 0.121317729 |
| 908,800 | 0.114166285 | 0.157473438 | 0.049981505 | 0.120988954 |
| 908,900 | 0.114250202 | 0.15821163 | 0.049709944 | 0.121480651 |
| 909,000 | 0.114221076 | 0.157756273 | 0.04991326 | 0.121072946 |
| 909,100 | 0.114164154 | 0.157612683 | 0.049697856 | 0.120960721 |
| 909,200 | 0.114078077 | 0.157180375 | 0.05044663 | 0.120702047 |
| 909,300 | 0.114068607 | 0.157442494 | 0.049397951 | 0.121133805 |
| 909,400 | 0.114044968 | 0.156967206 | 0.050043313 | 0.120926101 |
| 909,500 | 0.114110434 | 0.15732947 | 0.049845967 | 0.121026546 |
| 909,600 | 0.114022977 | 0.157353204 | 0.04937668 | 0.120494913 |
| 909,700 | 0.114074362 | 0.157008362 | 0.050235676 | 0.120749086 |
| 909,800 | 0.114037849 | 0.157168312 | 0.049846019 | 0.120375118 |
| 909,900 | 0.114107558 | 0.156630047 | 0.049371065 | 0.120774733 |
| 910,000 | 0.11407423 | 0.157235016 | 0.049700622 | 0.120824504 |
| 910,100 | 0.11411455 | 0.157728693 | 0.049526406 | 0.119937622 |

| | | | | |
|---|---|---|---|---|
| 910,200 | 0.114212542 | 0.157210891 | 0.050086429 | 0.120312186 |
| 910,300 | 0.114160844 | 0.157890862 | 0.049933715 | 0.120296427 |
| 910,400 | 0.114125187 | 0.158058424 | 0.049961584 | 0.120013525 |
| 910,500 | 0.114065772 | 0.15738786 | 0.050347066 | 0.120372056 |
| 910,600 | 0.11410695 | 0.158270861 | 0.049228386 | 0.120033648 |
| 910,700 | 0.114133183 | 0.158245462 | 0.049493449 | 0.120346506 |
| 910,800 | 0.11415019 | 0.158071798 | 0.049755999 | 0.120452263 |
| 910,900 | 0.114178524 | 0.158325105 | 0.049493687 | 0.120536524 |
| 911,000 | 0.114162098 | 0.159031753 | 0.049831555 | 0.120129617 |
| 911,100 | 0.114072432 | 0.158306777 | 0.049814054 | 0.121021426 |
| 911,200 | 0.114059487 | 0.158195817 | 0.04983478 | 0.120755436 |
| 911,300 | 0.114146875 | 0.158260416 | 0.049721641 | 0.120660971 |
| 911,400 | 0.114222381 | 0.158482676 | 0.049745368 | 0.120351104 |
| 911,500 | 0.114154056 | 0.15830103 | 0.050174279 | 0.120469446 |
| 911,600 | 0.11406738 | 0.157664456 | 0.050359135 | 0.12039535 |
| 911,700 | 0.114142727 | 0.157912172 | 0.050330863 | 0.120027019 |
| 911,800 | 0.11412621 | 0.158285914 | 0.049868539 | 0.120165762 |
| 911,900 | 0.114129625 | 0.158125183 | 0.049702621 | 0.120020791 |
| 912,000 | 0.114049271 | 0.158033925 | 0.050154995 | 0.119831922 |
| 912,100 | 0.114097467 | 0.157817279 | 0.049783449 | 0.119331833 |
| 912,200 | 0.114070175 | 0.157942108 | 0.049966764 | 0.119791687 |
| 912,300 | 0.114023474 | 0.157876392 | 0.05024345 | 0.119667298 |
| 912,400 | 0.114138449 | 0.157763086 | 0.050468096 | 0.119782608 |
| 912,500 | 0.11414223 | 0.157327808 | 0.049892145 | 0.119648199 |
| 912,600 | 0.114188107 | 0.157383753 | 0.050353771 | 0.120020065 |
| 912,700 | 0.114103077 | 0.157091464 | 0.049523287 | 0.119916555 |
| 912,800 | 0.114040446 | 0.157536842 | 0.049553968 | 0.119713735 |
| 912,900 | 0.114091003 | 0.157079758 | 0.049704438 | 0.119907738 |
| 913,000 | 0.114097499 | 0.157745261 | 0.04956592 | 0.120154894 |
| 913,100 | 0.114121135 | 0.157906224 | 0.049812666 | 0.119524964 |
| 913,200 | 0.114036594 | 0.157474587 | 0.049856728 | 0.1194121 |
| 913,300 | 0.114057575 | 0.157459004 | 0.049714376 | 0.119600622 |
| 913,400 | 0.114041591 | 0.157702691 | 0.049891551 | 0.120060147 |
| 913,500 | 0.114051558 | 0.158580663 | 0.050055546 | 0.119732131 |
| 913,600 | 0.113966802 | 0.157297408 | 0.049888154 | 0.119713158 |
| 913,700 | 0.11404769 | 0.15807228 | 0.04917641 | 0.119750903 |
| 913,800 | 0.114009205 | 0.158141661 | 0.049339674 | 0.11993326 |
| 913,900 | 0.114030252 | 0.158825354 | 0.049756036 | 0.11969742 |
| 914,000 | 0.113993957 | 0.158476749 | 0.049633133 | 0.119681629 |
| 914,100 | 0.114071658 | 0.15855031 | 0.04963777 | 0.119717882 |
| 914,200 | 0.114026856 | 0.158040073 | 0.04958638 | 0.119869904 |
| 914,300 | 0.114016024 | 0.157820101 | 0.049190299 | 0.119914891 |
| 914,400 | 0.114139505 | 0.158138927 | 0.048806833 | 0.120277269 |
| 914,500 | 0.114047311 | 0.158148535 | 0.049880877 | 0.119854509 |
| 914,600 | 0.114060503 | 0.158051299 | 0.049416247 | 0.119728344 |
| 914,700 | 0.113916014 | 0.158071182 | 0.049719397 | 0.120022567 |
| 914,800 | 0.11403376 | 0.15875732 | 0.049203883 | 0.120053834 |
| 914,900 | 0.114090547 | 0.158491704 | 0.049022238 | 0.120385228 |
| 915,000 | 0.113987889 | 0.158027556 | 0.049412532 | 0.119859702 |
| 915,100 | 0.114029088 | 0.158439032 | 0.049579869 | 0.119965121 |
| 915,200 | 0.114018691 | 0.157646548 | 0.049542185 | 0.119816037 |
| 915,300 | 0.114043272 | 0.157566076 | 0.049101008 | 0.119691768 |
| 915,400 | 0.113936206 | 0.157360014 | 0.049354905 | 0.119440134 |
| 915,500 | 0.11395059 | 0.157546619 | 0.049887469 | 0.11944721 |
| 915,600 | 0.113983283 | 0.157661392 | 0.049391381 | 0.119808064 |
| 915,700 | 0.113912127 | 0.157691451 | 0.049593973 | 0.119932453 |
| 915,800 | 0.114032913 | 0.157525486 | 0.049370183 | 0.1198067 |
| 915,900 | 0.114044564 | 0.157632677 | 0.050046186 | 0.119981219 |
| 916,000 | 0.114055887 | 0.157470134 | 0.049302397 | 0.120299756 |
| 916,100 | 0.114122701 | 0.157741205 | 0.049430099 | 0.120515527 |
| 916,200 | 0.114061237 | 0.158208543 | 0.049274554 | 0.119860986 |
| 916,300 | 0.114006167 | 0.15752242 | 0.049450709 | 0.12005395 |
| 916,400 | 0.114096495 | 0.157848435 | 0.04892008 | 0.119800065 |
| 916,500 | 0.114078442 | 0.15839517 | 0.049237187 | 0.120005332 |
| 916,600 | 0.114049444 | 0.157226742 | 0.048938372 | 0.119538155 |
| 916,700 | 0.11400543 | 0.157641156 | 0.0490393 | 0.119884846 |
| 916,800 | 0.113983168 | 0.157669144 | 0.049065703 | 0.119801247 |

| | | | | |
|---|---|---|---|---|
| 916,900 | 0.114018519 | 0.157586174 | 0.048655033 | 0.119827414 |
| 917,000 | 0.113998741 | 0.157298738 | 0.048963318 | 0.119267024 |
| 917,100 | 0.113995875 | 0.157256431 | 0.049400974 | 0.119291551 |
| 917,200 | 0.113931207 | 0.157612632 | 0.049366036 | 0.119826234 |
| 917,300 | 0.113908018 | 0.157466618 | 0.049200135 | 0.119008633 |
| 917,400 | 0.114029339 | 0.158090734 | 0.049612017 | 0.119261012 |
| 917,500 | 0.113880531 | 0.157355209 | 0.049330852 | 0.119581903 |
| 917,600 | 0.113835014 | 0.156822632 | 0.049430711 | 0.119283753 |
| 917,700 | 0.11376545 | 0.157446018 | 0.050111116 | 0.119319891 |
| 917,800 | 0.113923199 | 0.157022477 | 0.048908474 | 0.119515621 |
| 917,900 | 0.113919215 | 0.15726415 | 0.049297942 | 0.119690421 |
| 918,000 | 0.113879653 | 0.157841621 | 0.049202194 | 0.119469375 |
| 918,100 | 0.113936791 | 0.157765903 | 0.048761509 | 0.119926932 |
| 918,200 | 0.113766133 | 0.157253789 | 0.049000271 | 0.119765162 |
| 918,300 | 0.113829991 | 0.156955866 | 0.048877445 | 0.119774644 |
| 918,400 | 0.113786014 | 0.157566119 | 0.048656931 | 0.119226296 |
| 918,500 | 0.113702954 | 0.157281969 | 0.048797359 | 0.119064098 |
| 918,600 | 0.113677312 | 0.157011717 | 0.048853405 | 0.11909976 |
| 918,700 | 0.113641477 | 0.157511548 | 0.048504767 | 0.119006852 |
| 918,800 | 0.113659397 | 0.158002436 | 0.049052412 | 0.119315811 |
| 918,900 | 0.113808937 | 0.1578218 | 0.048458642 | 0.11956619 |
| 919,000 | 0.113654183 | 0.157125577 | 0.048069726 | 0.119573666 |
| 919,100 | 0.113760823 | 0.157464436 | 0.048707909 | 0.119230095 |
| 919,200 | 0.113757746 | 0.157346283 | 0.048374344 | 0.119597287 |
| 919,300 | 0.113720518 | 0.157206072 | 0.048513904 | 0.119754523 |
| 919,400 | 0.113742811 | 0.157282996 | 0.048699296 | 0.119484055 |
| 919,500 | 0.113677151 | 0.156484511 | 0.048879059 | 0.119287086 |
| 919,600 | 0.113721434 | 0.157027972 | 0.048556558 | 0.119200059 |
| 919,700 | 0.11369535 | 0.156692565 | 0.048737356 | 0.118811417 |
| 919,800 | 0.113635511 | 0.156823884 | 0.048483861 | 0.119363522 |
| 919,900 | 0.113590739 | 0.157174714 | 0.048654878 | 0.118877234 |
| 920,000 | 0.113696499 | 0.156713842 | 0.048640728 | 0.11932116 |
| 920,100 | 0.113558269 | 0.157363683 | 0.048684283 | 0.1192719 |
| 920,200 | 0.113549532 | 0.157493697 | 0.048880667 | 0.119361524 |
| 920,300 | 0.1135955 | 0.15692717 | 0.04903984 | 0.119063199 |
| 920,400 | 0.113622216 | 0.157274609 | 0.048401872 | 0.118710716 |
| 920,500 | 0.113658326 | 0.157607849 | 0.048762381 | 0.119374952 |
| 920,600 | 0.113550535 | 0.156951422 | 0.048650754 | 0.118804466 |
| 920,700 | 0.113485352 | 0.15690728 | 0.048839757 | 0.119687055 |
| 920,800 | 0.113586858 | 0.156879237 | 0.048747384 | 0.119324766 |
| 920,900 | 0.113664786 | 0.15681554 | 0.048580592 | 0.120042456 |
| 921,000 | 0.113674891 | 0.156785234 | 0.048574022 | 0.120013391 |
| 921,100 | 0.113670987 | 0.15598844 | 0.049155128 | 0.120110059 |
| 921,200 | 0.113660071 | 0.155803665 | 0.048587514 | 0.120263747 |
| 921,300 | 0.113671452 | 0.156361245 | 0.048852379 | 0.119901468 |
| 921,400 | 0.113701404 | 0.155744841 | 0.049550149 | 0.120066133 |
| 921,500 | 0.113607506 | 0.156753029 | 0.048853967 | 0.120012612 |
| 921,600 | 0.113676578 | 0.15598834 | 0.048874134 | 0.120067339 |
| 921,700 | 0.113655485 | 0.156032066 | 0.049364828 | 0.120200191 |
| 921,800 | 0.113631268 | 0.155980879 | 0.049119805 | 0.120298817 |
| 921,900 | 0.113616492 | 0.155956008 | 0.049214172 | 0.120172598 |
| 922,000 | 0.113611844 | 0.155724685 | 0.049224401 | 0.120410207 |
| 922,100 | 0.113545916 | 0.156633403 | 0.048783743 | 0.120468059 |
| 922,200 | 0.113487001 | 0.156714133 | 0.048895205 | 0.119827578 |
| 922,300 | 0.113616224 | 0.156479697 | 0.048900776 | 0.120734987 |
| 922,400 | 0.113428166 | 0.155733377 | 0.048611658 | 0.120144261 |
| 922,500 | 0.11339281 | 0.155930031 | 0.048873477 | 0.120109452 |
| 922,600 | 0.113412291 | 0.155783626 | 0.048563355 | 0.119991485 |
| 922,700 | 0.11337125 | 0.155254786 | 0.048585749 | 0.120329663 |
| 922,800 | 0.113379229 | 0.155841972 | 0.048604215 | 0.119985108 |
| 922,900 | 0.113396772 | 0.156295829 | 0.048427263 | 0.120309546 |
| 923,000 | 0.113413854 | 0.15610199 | 0.048423981 | 0.120126542 |
| 923,100 | 0.113374365 | 0.155675694 | 0.048779634 | 0.1200234 |
| 923,200 | 0.11347179 | 0.156259671 | 0.049149189 | 0.120018923 |
| 923,300 | 0.113442281 | 0.156982149 | 0.048961617 | 0.12043183 |
| 923,400 | 0.113607018 | 0.156782021 | 0.048824318 | 0.120374888 |
| 923,500 | 0.113363287 | 0.156153004 | 0.048096817 | 0.120063711 |

| | | | | |
|---|---|---|---|---|
| 923,600 | 0.113463896 | 0.156523649 | 0.048858207 | 0.119539634 |
| 923,700 | 0.113423012 | 0.156506829 | 0.048399512 | 0.119748293 |
| 923,800 | 0.113351831 | 0.15668338 | 0.049015658 | 0.119500317 |
| 923,900 | 0.11329772 | 0.156838145 | 0.048747075 | 0.119271576 |
| 924,000 | 0.113247055 | 0.156541667 | 0.048565326 | 0.119182104 |
| 924,100 | 0.113287933 | 0.156225668 | 0.048826804 | 0.119774214 |
| 924,200 | 0.113387188 | 0.156776112 | 0.048353186 | 0.119726241 |
| 924,300 | 0.113332383 | 0.156409009 | 0.048641419 | 0.119859099 |
| 924,400 | 0.113284695 | 0.156116586 | 0.048504654 | 0.119249512 |
| 924,500 | 0.113194839 | 0.156190792 | 0.048530546 | 0.119119803 |
| 924,600 | 0.113206788 | 0.156367975 | 0.048354338 | 0.119849718 |
| 924,700 | 0.113139749 | 0.155983822 | 0.048712899 | 0.119574138 |
| 924,800 | 0.113071537 | 0.155762559 | 0.048889346 | 0.11954377 |
| 924,900 | 0.113196185 | 0.155909287 | 0.048837945 | 0.119512365 |
| 925,000 | 0.11325045 | 0.156169595 | 0.049122711 | 0.11933055 |
| 925,100 | 0.113261071 | 0.156325152 | 0.048904519 | 0.119619078 |
| 925,200 | 0.113166515 | 0.156043932 | 0.048784784 | 0.119017552 |
| 925,300 | 0.113195003 | 0.155787161 | 0.049385011 | 0.118947805 |
| 925,400 | 0.113134895 | 0.156105235 | 0.049118397 | 0.119018662 |
| 925,500 | 0.113212279 | 0.156226673 | 0.049223941 | 0.118721149 |
| 925,600 | 0.113126702 | 0.155960297 | 0.049243652 | 0.119088681 |
| 925,700 | 0.113053331 | 0.156102603 | 0.049422726 | 0.11819329 |
| 925,800 | 0.113200623 | 0.156195799 | 0.049387451 | 0.118701222 |
| 925,900 | 0.113211999 | 0.156922868 | 0.049521662 | 0.11946859 |
| 926,000 | 0.113037641 | 0.156377176 | 0.049653933 | 0.118801674 |
| 926,100 | 0.113028051 | 0.156160365 | 0.049541377 | 0.119286354 |
| 926,200 | 0.113042173 | 0.155838537 | 0.049041011 | 0.11920929 |
| 926,300 | 0.113002024 | 0.155940158 | 0.049302423 | 0.118512108 |
| 926,400 | 0.113037266 | 0.155879035 | 0.049534961 | 0.118197052 |
| 926,500 | 0.113039033 | 0.155964424 | 0.049843571 | 0.118456478 |
| 926,600 | 0.112972486 | 0.155774244 | 0.049280118 | 0.117854669 |
| 926,700 | 0.113050052 | 0.155654499 | 0.049351343 | 0.11842678 |
| 926,800 | 0.113037138 | 0.156208696 | 0.049587408 | 0.118847191 |
| 926,900 | 0.113035869 | 0.155729334 | 0.049778002 | 0.118078971 |
| 927,000 | 0.113067091 | 0.155879611 | 0.049587447 | 0.118247386 |
| 927,100 | 0.113032515 | 0.155904673 | 0.049520553 | 0.118116302 |
| 927,200 | 0.112928117 | 0.155574589 | 0.049248637 | 0.118531852 |
| 927,300 | 0.112961062 | 0.155771414 | 0.049469716 | 0.118524999 |
| 927,400 | 0.112939799 | 0.155774538 | 0.048981172 | 0.118423036 |
| 927,500 | 0.11290669 | 0.155285813 | 0.048842728 | 0.118900385 |
| 927,600 | 0.11301355 | 0.15534757 | 0.049799383 | 0.118746677 |
| 927,700 | 0.113026258 | 0.155494383 | 0.049282863 | 0.118644925 |
| 927,800 | 0.112950946 | 0.155539508 | 0.049246052 | 0.118766068 |
| 927,900 | 0.113090091 | 0.155943807 | 0.049422649 | 0.119034972 |
| 928,000 | 0.112892458 | 0.155735612 | 0.049170574 | 0.118888175 |
| 928,100 | 0.112862076 | 0.154503085 | 0.049254903 | 0.118840883 |
| 928,200 | 0.112915311 | 0.155514063 | 0.049710716 | 0.118799798 |
| 928,300 | 0.112910885 | 0.155444425 | 0.049619136 | 0.118677504 |
| 928,400 | 0.112859917 | 0.154997573 | 0.049581789 | 0.118297521 |
| 928,500 | 0.112851334 | 0.155276215 | 0.049078996 | 0.118254645 |
| 928,600 | 0.112974716 | 0.154802897 | 0.049403302 | 0.118145729 |
| 928,700 | 0.11294937 | 0.154801543 | 0.049496988 | 0.118176576 |
| 928,800 | 0.1128757 | 0.154715775 | 0.04978648 | 0.117894785 |
| 928,900 | 0.112870198 | 0.154682264 | 0.04949319 | 0.118085621 |
| 929,000 | 0.1128513 | 0.154924071 | 0.049735255 | 0.117686483 |
| 929,100 | 0.112934347 | 0.155026924 | 0.049227084 | 0.118090347 |
| 929,200 | 0.112887481 | 0.155508294 | 0.04965444 | 0.118502628 |
| 929,300 | 0.112879075 | 0.155000905 | 0.049476037 | 0.118513387 |
| 929,400 | 0.112870261 | 0.154340729 | 0.049940332 | 0.118468833 |
| 929,500 | 0.112965478 | 0.155238666 | 0.049449513 | 0.11850041 |
| 929,600 | 0.112996724 | 0.154560739 | 0.049133462 | 0.118664886 |
| 929,700 | 0.113099569 | 0.155585844 | 0.049600707 | 0.118064468 |
| 929,800 | 0.112993736 | 0.155189672 | 0.05009678 | 0.117764545 |
| 929,900 | 0.112991837 | 0.154878644 | 0.05018321 | 0.117755119 |
| 930,000 | 0.112973699 | 0.155375961 | 0.050398258 | 0.118396862 |
| 930,100 | 0.112968644 | 0.155638324 | 0.050503179 | 0.118611506 |
| 930,200 | 0.112899246 | 0.155267611 | 0.050142006 | 0.118461572 |

| | | | | |
|---|---|---|---|---|
| 930,300 | 0.112854831 | 0.154880136 | 0.050393549 | 0.118526612 |
| 930,400 | 0.112838835 | 0.154614195 | 0.050301805 | 0.118366272 |
| 930,500 | 0.112853734 | 0.1550628 | 0.050348924 | 0.118247705 |
| 930,600 | 0.112755402 | 0.154892645 | 0.050082987 | 0.118315527 |
| 930,700 | 0.112921577 | 0.154209383 | 0.05038416 | 0.118396929 |
| 930,800 | 0.112956567 | 0.154433752 | 0.050074247 | 0.11799223 |
| 930,900 | 0.112881474 | 0.154035117 | 0.049846619 | 0.118381171 |
| 931,000 | 0.112878367 | 0.154765972 | 0.049879399 | 0.118778756 |
| 931,100 | 0.112847419 | 0.154067882 | 0.049841619 | 0.118519912 |
| 931,200 | 0.112825685 | 0.155048487 | 0.049636469 | 0.118602671 |
| 931,300 | 0.112855307 | 0.154884323 | 0.049329809 | 0.118249328 |
| 931,400 | 0.112886758 | 0.154434278 | 0.049497422 | 0.118137774 |
| 931,500 | 0.112818016 | 0.154749949 | 0.049546832 | 0.118414353 |
| 931,600 | 0.112832458 | 0.154542446 | 0.048877263 | 0.118757664 |
| 931,700 | 0.112914434 | 0.154826636 | 0.049175479 | 0.118614811 |
| 931,800 | 0.112858286 | 0.154643875 | 0.049401337 | 0.118629527 |
| 931,900 | 0.112881408 | 0.155159858 | 0.049152847 | 0.118663773 |
| 932,000 | 0.11294513 | 0.15556656 | 0.048960798 | 0.11842885 |
| 932,100 | 0.112951633 | 0.155153842 | 0.048601654 | 0.119005051 |
| 932,200 | 0.112816028 | 0.154823501 | 0.04824775 | 0.118455078 |
| 932,300 | 0.112810378 | 0.155493333 | 0.048526464 | 0.118497494 |
| 932,400 | 0.112796317 | 0.155564093 | 0.048480305 | 0.118319374 |
| 932,500 | 0.112727314 | 0.154799378 | 0.048245012 | 0.118110266 |
| 932,600 | 0.112646476 | 0.154895398 | 0.048494105 | 0.118084068 |
| 932,700 | 0.112766178 | 0.155053386 | 0.048555267 | 0.118109099 |
| 932,800 | 0.112762954 | 0.15546321 | 0.048416523 | 0.117951729 |
| 932,900 | 0.112792202 | 0.155578651 | 0.0485577 | 0.118114689 |
| 933,000 | 0.112665042 | 0.155081343 | 0.048482546 | 0.11815886 |
| 933,100 | 0.112665357 | 0.154660341 | 0.048318222 | 0.118180243 |
| 933,200 | 0.112619663 | 0.154543671 | 0.048359735 | 0.118342395 |
| 933,300 | 0.112681977 | 0.154730458 | 0.04873432 | 0.117966607 |
| 933,400 | 0.112718067 | 0.154908705 | 0.047905425 | 0.117874061 |
| 933,500 | 0.112785783 | 0.154975382 | 0.048076266 | 0.118066021 |
| 933,600 | 0.112655076 | 0.154943982 | 0.048240345 | 0.117958646 |
| 933,700 | 0.112606114 | 0.155189528 | 0.047856235 | 0.117755171 |
| 933,800 | 0.112624616 | 0.155559321 | 0.048102362 | 0.117927509 |
| 933,900 | 0.112574661 | 0.155541075 | 0.047748072 | 0.117565112 |
| 934,000 | 0.112628807 | 0.155657048 | 0.048479021 | 0.117664764 |
| 934,100 | 0.112570394 | 0.155887858 | 0.04821114 | 0.117946763 |
| 934,200 | 0.112555137 | 0.155495804 | 0.048494512 | 0.117427068 |
| 934,300 | 0.112598611 | 0.155101551 | 0.048460381 | 0.117386268 |
| 934,400 | 0.112521329 | 0.154966939 | 0.048459832 | 0.117686735 |
| 934,500 | 0.112531481 | 0.155143058 | 0.048703253 | 0.117572619 |
| 934,600 | 0.112569638 | 0.155612062 | 0.048730566 | 0.117639011 |
| 934,700 | 0.112496441 | 0.155302316 | 0.048635816 | 0.117465422 |
| 934,800 | 0.112491071 | 0.155183008 | 0.048605258 | 0.11707762 |
| 934,900 | 0.112518613 | 0.155143524 | 0.048464245 | 0.117566724 |
| 935,000 | 0.11246434 | 0.155070504 | 0.048217278 | 0.117163368 |
| 935,100 | 0.112513923 | 0.155370344 | 0.047858349 | 0.117224228 |
| 935,200 | 0.112479292 | 0.155853101 | 0.048226078 | 0.116792259 |
| 935,300 | 0.11239819 | 0.156023142 | 0.048713911 | 0.116659872 |
| 935,400 | 0.112529877 | 0.155760243 | 0.048608821 | 0.117054333 |
| 935,500 | 0.112506213 | 0.156120137 | 0.048331823 | 0.117278505 |
| 935,600 | 0.112346651 | 0.155215068 | 0.048253497 | 0.116829149 |
| 935,700 | 0.112331813 | 0.155645971 | 0.048884478 | 0.117179423 |
| 935,800 | 0.112424296 | 0.156067779 | 0.048541228 | 0.117601723 |
| 935,900 | 0.112371505 | 0.156177321 | 0.048703183 | 0.117721975 |
| 936,000 | 0.112387177 | 0.15582702 | 0.049092436 | 0.117270516 |
| 936,100 | 0.112375315 | 0.156002145 | 0.048844341 | 0.117467842 |
| 936,200 | 0.112334684 | 0.156355414 | 0.04858369 | 0.117484182 |
| 936,300 | 0.112424149 | 0.155854853 | 0.048982562 | 0.117059413 |
| 936,400 | 0.112403697 | 0.156094123 | 0.048543176 | 0.117294975 |
| 936,500 | 0.112460199 | 0.155642799 | 0.048481519 | 0.117083703 |
| 936,600 | 0.112374667 | 0.155938921 | 0.048381626 | 0.117174293 |
| 936,700 | 0.112323617 | 0.155761925 | 0.048382639 | 0.117147109 |
| 936,800 | 0.112310882 | 0.155796899 | 0.048661883 | 0.117563417 |
| 936,900 | 0.112385457 | 0.15590955 | 0.048905825 | 0.117104732 |

| | | | | |
|---|---|---|---|---|
| 937,000 | 0.112319677 | 0.155807676 | 0.048880503 | 0.11759423 |
| 937,100 | 0.112424485 | 0.155841324 | 0.04862009 | 0.117128583 |
| 937,200 | 0.112383491 | 0.155633276 | 0.048608326 | 0.117355779 |
| 937,300 | 0.112304075 | 0.155638367 | 0.048893651 | 0.117266388 |
| 937,400 | 0.112310908 | 0.155708322 | 0.048887404 | 0.117436446 |
| 937,500 | 0.112255166 | 0.155748657 | 0.048546957 | 0.117609801 |
| 937,600 | 0.112262246 | 0.155764714 | 0.048787244 | 0.117907575 |
| 937,700 | 0.112310666 | 0.155810925 | 0.048804863 | 0.117783518 |
| 937,800 | 0.112373098 | 0.156225993 | 0.048563312 | 0.11763958 |
| 937,900 | 0.112377185 | 0.156071497 | 0.048359665 | 0.117920683 |
| 938,000 | 0.11222048 | 0.155733696 | 0.048650471 | 0.117570397 |
| 938,100 | 0.112218011 | 0.155320904 | 0.048664109 | 0.117310482 |
| 938,200 | 0.112225568 | 0.155770866 | 0.048396701 | 0.117054051 |
| 938,300 | 0.112184329 | 0.155293566 | 0.048404604 | 0.117319907 |
| 938,400 | 0.112216145 | 0.155015429 | 0.048671729 | 0.117347413 |
| 938,500 | 0.112208247 | 0.154869914 | 0.048860228 | 0.116886069 |
| 938,600 | 0.112226165 | 0.155277835 | 0.048031278 | 0.117833089 |
| 938,700 | 0.112152773 | 0.154984239 | 0.048379888 | 0.117083643 |
| 938,800 | 0.112104656 | 0.155237601 | 0.048415678 | 0.117134402 |
| 938,900 | 0.112123415 | 0.154965643 | 0.048590317 | 0.117599391 |
| 939,000 | 0.112113622 | 0.155336075 | 0.048485118 | 0.117158133 |
| 939,100 | 0.112076034 | 0.155425098 | 0.048465897 | 0.117371868 |
| 939,200 | 0.112096334 | 0.155115569 | 0.048408976 | 0.117559565 |
| 939,300 | 0.112051827 | 0.15506852 | 0.048628685 | 0.117223628 |
| 939,400 | 0.111987539 | 0.155488093 | 0.048376003 | 0.11739957 |
| 939,500 | 0.112062178 | 0.154843961 | 0.048457978 | 0.117222219 |
| 939,600 | 0.111962726 | 0.155148487 | 0.048526862 | 0.117085069 |
| 939,700 | 0.112009625 | 0.155371754 | 0.048196986 | 0.116933336 |
| 939,800 | 0.112002169 | 0.155836453 | 0.04813382 | 0.116990908 |
| 939,900 | 0.112043758 | 0.154932395 | 0.04798516 | 0.117542217 |
| 940,000 | 0.11202627 | 0.155203809 | 0.048165636 | 0.117431173 |
| 940,100 | 0.112106993 | 0.155312134 | 0.048218234 | 0.117741352 |
| 940,200 | 0.112070992 | 0.155605592 | 0.048533228 | 0.116998299 |
| 940,300 | 0.111986787 | 0.155113495 | 0.048695412 | 0.117117218 |
| 940,400 | 0.111886717 | 0.155320934 | 0.048246214 | 0.116556722 |
| 940,500 | 0.111880272 | 0.155326317 | 0.048527135 | 0.116802484 |
| 940,600 | 0.111888591 | 0.154991946 | 0.048549324 | 0.116624961 |
| 940,700 | 0.111969368 | 0.155224912 | 0.048568994 | 0.116945255 |
| 940,800 | 0.11198084 | 0.155326746 | 0.048448679 | 0.117280629 |
| 940,900 | 0.11188016 | 0.155141874 | 0.048843475 | 0.117001053 |
| 941,000 | 0.111914487 | 0.155393256 | 0.048601707 | 0.116586368 |
| 941,100 | 0.111941376 | 0.155708707 | 0.049071548 | 0.11694248 |
| 941,200 | 0.111896983 | 0.155086709 | 0.048561349 | 0.116991232 |
| 941,300 | 0.111829766 | 0.155172244 | 0.049216403 | 0.116791434 |
| 941,400 | 0.111884869 | 0.154825135 | 0.04887169 | 0.116620223 |
| 941,500 | 0.111916024 | 0.154694755 | 0.048787057 | 0.116939515 |
| 941,600 | 0.111960147 | 0.154677981 | 0.04834458 | 0.117072647 |
| 941,700 | 0.111884174 | 0.154257378 | 0.048415355 | 0.116849234 |
| 941,800 | 0.111953066 | 0.154703184 | 0.047814595 | 0.116985257 |
| 941,900 | 0.111806656 | 0.154661729 | 0.048038106 | 0.117076366 |
| 942,000 | 0.111846395 | 0.155125947 | 0.047797249 | 0.117202311 |
| 942,100 | 0.111813986 | 0.15439318 | 0.047919878 | 0.116806291 |
| 942,200 | 0.11176874 | 0.154487577 | 0.047620743 | 0.116774458 |
| 942,300 | 0.11184451 | 0.155103225 | 0.047911096 | 0.117219697 |
| 942,400 | 0.111816907 | 0.154910375 | 0.047876078 | 0.116919609 |
| 942,500 | 0.111790198 | 0.155066343 | 0.047605687 | 0.11686475 |
| 942,600 | 0.111890619 | 0.155350908 | 0.047548999 | 0.117220397 |
| 942,700 | 0.111872458 | 0.155488173 | 0.047321254 | 0.117041168 |
| 942,800 | 0.111795107 | 0.154536777 | 0.047711774 | 0.116734982 |
| 942,900 | 0.111845686 | 0.154497445 | 0.047742322 | 0.116871207 |
| 943,000 | 0.111845937 | 0.155089389 | 0.04783621 | 0.117535707 |
| 943,100 | 0.111812921 | 0.155303006 | 0.047599345 | 0.117027184 |
| 943,200 | 0.111769859 | 0.154489101 | 0.047230694 | 0.117333247 |
| 943,300 | 0.111692681 | 0.154807966 | 0.047544509 | 0.116844839 |
| 943,400 | 0.111752294 | 0.154465 | 0.047302394 | 0.116901094 |
| 943,500 | 0.111696999 | 0.154184387 | 0.047043664 | 0.116929873 |
| 943,600 | 0.111760863 | 0.154230558 | 0.046905334 | 0.116692448 |

| | | | | |
|---|---|---|---|---|
| 943,700 | 0.111756348 | 0.154134138 | 0.047164132 | 0.11688913 |
| 943,800 | 0.111788223 | 0.154352161 | 0.047644869 | 0.11748059 |
| 943,900 | 0.111687079 | 0.154249285 | 0.047781246 | 0.116985639 |
| 944,000 | 0.111809538 | 0.154394503 | 0.047433057 | 0.117604911 |
| 944,100 | 0.111681558 | 0.153986978 | 0.047370631 | 0.117571928 |
| 944,200 | 0.111726997 | 0.154088676 | 0.047399526 | 0.117561223 |
| 944,300 | 0.111683306 | 0.154280777 | 0.046746658 | 0.117679926 |
| 944,400 | 0.111727083 | 0.154193963 | 0.047246 | 0.117433777 |
| 944,500 | 0.111679322 | 0.154235532 | 0.047676236 | 0.11749938 |
| 944,600 | 0.111763319 | 0.154295981 | 0.047593788 | 0.117657982 |
| 944,700 | 0.11172214 | 0.154323235 | 0.047725766 | 0.117659344 |
| 944,800 | 0.111723817 | 0.154159666 | 0.047801305 | 0.117576501 |
| 944,900 | 0.111828259 | 0.154719094 | 0.048329185 | 0.117533566 |
| 945,000 | 0.111761068 | 0.154295043 | 0.047676595 | 0.117893307 |
| 945,100 | 0.111728859 | 0.154561189 | 0.047483007 | 0.117378047 |
| 945,200 | 0.111834976 | 0.154842894 | 0.04756239 | 0.117484103 |
| 945,300 | 0.11179742 | 0.154171805 | 0.047663354 | 0.117668875 |
| 945,400 | 0.111684652 | 0.154170285 | 0.048070097 | 0.117218312 |
| 945,500 | 0.111692064 | 0.154331299 | 0.047482362 | 0.11656777 |
| 945,600 | 0.111723294 | 0.154299249 | 0.047614333 | 0.117139921 |
| 945,700 | 0.111831972 | 0.154596013 | 0.047090797 | 0.117359535 |
| 945,800 | 0.111686494 | 0.153866742 | 0.047463834 | 0.117487852 |
| 945,900 | 0.111610375 | 0.153102017 | 0.047830339 | 0.116805557 |
| 946,000 | 0.11165722 | 0.153729467 | 0.047820108 | 0.117472842 |
| 946,100 | 0.111595923 | 0.153721738 | 0.047567689 | 0.117278069 |
| 946,200 | 0.111563623 | 0.154386288 | 0.047570709 | 0.117351819 |
| 946,300 | 0.111587399 | 0.153883367 | 0.047765313 | 0.117686355 |
| 946,400 | 0.111604779 | 0.154158174 | 0.047557404 | 0.117726558 |
| 946,500 | 0.111631312 | 0.154674836 | 0.047147286 | 0.117574019 |
| 946,600 | 0.111569123 | 0.153255249 | 0.047358537 | 0.117572764 |
| 946,700 | 0.111628163 | 0.153949931 | 0.047747764 | 0.118169429 |
| 946,800 | 0.111560395 | 0.153773802 | 0.047617679 | 0.117127264 |
| 946,900 | 0.111422531 | 0.15370307 | 0.047627936 | 0.117360535 |
| 947,000 | 0.111383014 | 0.153510056 | 0.047972847 | 0.117361685 |
| 947,100 | 0.111473767 | 0.153285675 | 0.047591505 | 0.11715762 |
| 947,200 | 0.111596146 | 0.153462314 | 0.047828602 | 0.117157029 |
| 947,300 | 0.111501657 | 0.153547044 | 0.047789285 | 0.117054098 |
| 947,400 | 0.111465419 | 0.153436904 | 0.047395323 | 0.117392266 |
| 947,500 | 0.111487356 | 0.153358502 | 0.047499393 | 0.117209597 |
| 947,600 | 0.111547878 | 0.152716378 | 0.04767319 | 0.117189741 |
| 947,700 | 0.111518286 | 0.153082079 | 0.047700269 | 0.116591648 |
| 947,800 | 0.111487346 | 0.153373759 | 0.047671815 | 0.116664224 |
| 947,900 | 0.111415853 | 0.152948832 | 0.047911694 | 0.117176389 |
| 948,000 | 0.111472061 | 0.153214754 | 0.047754626 | 0.116483688 |
| 948,100 | 0.111431634 | 0.153060397 | 0.048117676 | 0.116624244 |
| 948,200 | 0.111497822 | 0.153025025 | 0.047524581 | 0.116980164 |
| 948,300 | 0.111395819 | 0.153246837 | 0.047771461 | 0.116555109 |
| 948,400 | 0.1114208 | 0.153212882 | 0.047702287 | 0.116781517 |
| 948,500 | 0.111383566 | 0.153224677 | 0.047573429 | 0.116526917 |
| 948,600 | 0.111432174 | 0.153690655 | 0.047825976 | 0.116283314 |
| 948,700 | 0.111345051 | 0.153563121 | 0.047512178 | 0.116526988 |
| 948,800 | 0.111426959 | 0.152957414 | 0.047738458 | 0.116822922 |
| 948,900 | 0.111393711 | 0.152858019 | 0.047625273 | 0.11737862 |
| 949,000 | 0.111385649 | 0.153381206 | 0.048106412 | 0.116750275 |
| 949,100 | 0.111379426 | 0.153213565 | 0.047448631 | 0.116443646 |
| 949,200 | 0.111390581 | 0.153409773 | 0.046807045 | 0.116822609 |
| 949,300 | 0.111377155 | 0.153283692 | 0.047142012 | 0.116086284 |
| 949,400 | 0.111371032 | 0.153717049 | 0.047478414 | 0.116205005 |
| 949,500 | 0.111358274 | 0.153546067 | 0.048078118 | 0.116532635 |
| 949,600 | 0.11140226 | 0.153215901 | 0.047633625 | 0.116182232 |
| 949,700 | 0.111411355 | 0.153914479 | 0.047321534 | 0.116426948 |
| 949,800 | 0.111409856 | 0.153443706 | 0.04777809 | 0.116505634 |
| 949,900 | 0.11140035 | 0.153599329 | 0.047687869 | 0.116580203 |
| 950,000 | 0.111340546 | 0.15380669 | 0.047284179 | 0.11622277 |
| 950,100 | 0.111627183 | 0.154368605 | 0.047533818 | 0.116775052 |
| 950,200 | 0.111467863 | 0.153958098 | 0.047392574 | 0.116332771 |
| 950,300 | 0.111391616 | 0.154540865 | 0.047457163 | 0.116267515 |

| | | | | |
|---|---|---|---|---|
| 950,400 | 0.111415465 | 0.154439883 | 0.047369235 | 0.116306626 |
| 950,500 | 0.111359472 | 0.153943583 | 0.04684915 | 0.116371249 |
| 950,600 | 0.111381771 | 0.153745478 | 0.046966766 | 0.116372363 |
| 950,700 | 0.111447574 | 0.153693037 | 0.046864525 | 0.1167694 |
| 950,800 | 0.111332587 | 0.153405898 | 0.047004268 | 0.116694801 |
| 950,900 | 0.111406755 | 0.153389345 | 0.047525453 | 0.116615893 |
| 951,000 | 0.111463811 | 0.153610286 | 0.047560927 | 0.116558152 |
| 951,100 | 0.111413421 | 0.153573669 | 0.046839847 | 0.116113091 |
| 951,200 | 0.111306004 | 0.15389848 | 0.047235158 | 0.116633028 |
| 951,300 | 0.111285104 | 0.153576685 | 0.046902858 | 0.116250493 |
| 951,400 | 0.111309601 | 0.153666923 | 0.046400262 | 0.11675194 |
| 951,500 | 0.111282918 | 0.153528708 | 0.046951813 | 0.116696171 |
| 951,600 | 0.111296765 | 0.15357434 | 0.046672684 | 0.116391721 |
| 951,700 | 0.111248191 | 0.153749747 | 0.047065821 | 0.11620998 |
| 951,800 | 0.111242735 | 0.153493725 | 0.046578387 | 0.116557088 |
| 951,900 | 0.111269798 | 0.153938518 | 0.046265225 | 0.116021094 |
| 952,000 | 0.111340709 | 0.153887829 | 0.046799375 | 0.115881229 |
| 952,100 | 0.111247898 | 0.154515414 | 0.046641605 | 0.115935153 |
| 952,200 | 0.11126477 | 0.15379792 | 0.046106823 | 0.116420609 |
| 952,300 | 0.111213396 | 0.153490557 | 0.046518253 | 0.115441897 |
| 952,400 | 0.111245815 | 0.153614284 | 0.046649139 | 0.116170985 |
| 952,500 | 0.11125241 | 0.15378185 | 0.04666103 | 0.115749362 |
| 952,600 | 0.111199586 | 0.153625759 | 0.046198619 | 0.115659825 |
| 952,700 | 0.111187299 | 0.153357216 | 0.046228434 | 0.116250849 |
| 952,800 | 0.111275503 | 0.153418993 | 0.046017849 | 0.116378654 |
| 952,900 | 0.111222462 | 0.15341369 | 0.046395126 | 0.115676111 |
| 953,000 | 0.1112302 | 0.154157274 | 0.046792953 | 0.116378127 |
| 953,100 | 0.111241569 | 0.154068124 | 0.046596393 | 0.116571467 |
| 953,200 | 0.111070503 | 0.154077585 | 0.046231185 | 0.115695645 |
| 953,300 | 0.111113993 | 0.153627507 | 0.046269741 | 0.116048975 |
| 953,400 | 0.111141557 | 0.153632878 | 0.04661162 | 0.116013135 |
| 953,500 | 0.111157094 | 0.154509173 | 0.046583989 | 0.11580367 |
| 953,600 | 0.111194686 | 0.153802032 | 0.046703721 | 0.116474377 |
| 953,700 | 0.111129603 | 0.15430179 | 0.047134883 | 0.116024051 |
| 953,800 | 0.111187719 | 0.154126548 | 0.047388458 | 0.116036597 |
| 953,900 | 0.111083179 | 0.153932929 | 0.046975501 | 0.116057721 |
| 954,000 | 0.111067367 | 0.154143311 | 0.047326983 | 0.115353982 |
| 954,100 | 0.111147521 | 0.15395961 | 0.046572917 | 0.116220955 |
| 954,200 | 0.11106125 | 0.154605734 | 0.047030484 | 0.116351045 |
| 954,300 | 0.111085788 | 0.155336584 | 0.046835886 | 0.115999054 |
| 954,400 | 0.1111697 | 0.154955642 | 0.046894901 | 0.115843746 |
| 954,500 | 0.111053932 | 0.154143096 | 0.046670825 | 0.115736455 |
| 954,600 | 0.110952554 | 0.154306825 | 0.046651097 | 0.116064883 |
| 954,700 | 0.110958686 | 0.154107529 | 0.047121041 | 0.116153236 |
| 954,800 | 0.110977366 | 0.153355501 | 0.046565617 | 0.116301914 |
| 954,900 | 0.110987573 | 0.154164414 | 0.046890838 | 0.116315313 |
| 955,000 | 0.110907388 | 0.154037548 | 0.04656648 | 0.11612051 |
| 955,100 | 0.110941708 | 0.154002711 | 0.046578856 | 0.116075573 |
| 955,200 | 0.110910651 | 0.153911294 | 0.046968035 | 0.116510293 |
| 955,300 | 0.110990911 | 0.154342778 | 0.046950859 | 0.11642652 |
| 955,400 | 0.111046824 | 0.153969996 | 0.046511257 | 0.116142622 |
| 955,500 | 0.111044182 | 0.153998622 | 0.046473281 | 0.116380835 |
| 955,600 | 0.110977082 | 0.154526372 | 0.046330632 | 0.116443722 |
| 955,700 | 0.111029952 | 0.154104954 | 0.046852218 | 0.116141065 |
| 955,800 | 0.111019949 | 0.153741678 | 0.046591051 | 0.116223632 |
| 955,900 | 0.111003037 | 0.154287602 | 0.046809316 | 0.116569387 |
| 956,000 | 0.111036226 | 0.154313158 | 0.046712898 | 0.116744842 |
| 956,100 | 0.111008292 | 0.154209868 | 0.046692372 | 0.11642379 |
| 956,200 | 0.111040582 | 0.154382991 | 0.046555275 | 0.116588126 |
| 956,300 | 0.110968241 | 0.154105145 | 0.047295798 | 0.116373993 |
| 956,400 | 0.110959185 | 0.154668641 | 0.047059251 | 0.116308454 |
| 956,500 | 0.11104844 | 0.154626029 | 0.047169343 | 0.11600029 |
| 956,600 | 0.111060664 | 0.154403851 | 0.047267005 | 0.116075634 |
| 956,700 | 0.111144484 | 0.154718443 | 0.046852575 | 0.11638118 |
| 956,800 | 0.11112791 | 0.154749905 | 0.046934994 | 0.116098295 |
| 956,900 | 0.111024969 | 0.153926351 | 0.047085269 | 0.115867737 |
| 957,000 | 0.111047837 | 0.15386566 | 0.047382678 | 0.115730268 |

| | | | | |
|---|---|---|---|---|
| 957,100 | 0.111129906 | 0.154318765 | 0.047167768 | 0.116036076 |
| 957,200 | 0.111170448 | 0.154069319 | 0.04746742 | 0.116508955 |
| 957,300 | 0.111125167 | 0.154407991 | 0.047536215 | 0.116119586 |
| 957,400 | 0.111151626 | 0.155104768 | 0.047340934 | 0.115885317 |
| 957,500 | 0.111070793 | 0.154915816 | 0.047308231 | 0.115804758 |
| 957,600 | 0.111129523 | 0.154829767 | 0.047116256 | 0.116230714 |
| 957,700 | 0.111076017 | 0.154666147 | 0.047379046 | 0.115944252 |
| 957,800 | 0.11111373 | 0.155091107 | 0.047427137 | 0.116398619 |
| 957,900 | 0.111059393 | 0.154922604 | 0.047470165 | 0.116630365 |
| 958,000 | 0.11093691 | 0.154746792 | 0.047484811 | 0.116242784 |
| 958,100 | 0.110951724 | 0.154777925 | 0.047179195 | 0.115916276 |
| 958,200 | 0.11091858 | 0.154107472 | 0.047331159 | 0.116147837 |
| 958,300 | 0.110973512 | 0.154596257 | 0.047007931 | 0.116445027 |
| 958,400 | 0.111012768 | 0.154783948 | 0.046956197 | 0.116491098 |
| 958,500 | 0.111102142 | 0.154158404 | 0.04708767 | 0.11634645 |
| 958,600 | 0.110902734 | 0.153994482 | 0.047454054 | 0.115977462 |
| 958,700 | 0.110868882 | 0.153892941 | 0.047010421 | 0.116224546 |
| 958,800 | 0.111021594 | 0.154893974 | 0.047113945 | 0.11602546 |
| 958,900 | 0.111000787 | 0.154970011 | 0.047087079 | 0.116192455 |
| 959,000 | 0.110948062 | 0.154509979 | 0.047632183 | 0.116224391 |
| 959,100 | 0.110937112 | 0.154541422 | 0.047306921 | 0.116592485 |
| 959,200 | 0.110929721 | 0.154530111 | 0.047534103 | 0.116254482 |
| 959,300 | 0.111015395 | 0.155431663 | 0.047120819 | 0.116997859 |
| 959,400 | 0.110994124 | 0.155398029 | 0.046948904 | 0.116921058 |
| 959,500 | 0.110921074 | 0.154798087 | 0.046843381 | 0.117074331 |
| 959,600 | 0.110897816 | 0.155347578 | 0.046974914 | 0.116711871 |
| 959,700 | 0.110854014 | 0.154485954 | 0.046794723 | 0.116673071 |
| 959,800 | 0.11076403 | 0.154381188 | 0.047220352 | 0.116854752 |
| 959,900 | 0.110845004 | 0.15461649 | 0.04746261 | 0.116764147 |
| 960,000 | 0.110863626 | 0.154288336 | 0.048028931 | 0.116891004 |
| 960,100 | 0.110848945 | 0.154650771 | 0.048125975 | 0.11675255 |
| 960,200 | 0.110800434 | 0.154582737 | 0.04741638 | 0.116983381 |
| 960,300 | 0.110796225 | 0.15451756 | 0.047432766 | 0.11719177 |
| 960,400 | 0.110758934 | 0.154749286 | 0.047774395 | 0.116629011 |
| 960,500 | 0.110804567 | 0.154228698 | 0.047420347 | 0.11681977 |
| 960,600 | 0.110908375 | 0.155240249 | 0.047398831 | 0.117232548 |
| 960,700 | 0.110816374 | 0.155175754 | 0.047565372 | 0.116650185 |
| 960,800 | 0.110810795 | 0.154829419 | 0.047639791 | 0.116161637 |
| 960,900 | 0.11075953 | 0.154660274 | 0.047695063 | 0.116083659 |
| 961,000 | 0.110840817 | 0.15516724 | 0.047538368 | 0.116372313 |
| 961,100 | 0.11085575 | 0.15387393 | 0.047604303 | 0.116346989 |
| 961,200 | 0.110825118 | 0.155045795 | 0.047260879 | 0.115958546 |
| 961,300 | 0.110751527 | 0.154891209 | 0.047635811 | 0.116369513 |
| 961,400 | 0.110812289 | 0.154708514 | 0.047148639 | 0.116477251 |
| 961,500 | 0.110776181 | 0.155159439 | 0.046981181 | 0.116547305 |
| 961,600 | 0.110897649 | 0.155419181 | 0.046984903 | 0.116072503 |
| 961,700 | 0.11096 | 0.154798303 | 0.04702274 | 0.116034757 |
| 961,800 | 0.110838305 | 0.155060796 | 0.046804188 | 0.115806131 |
| 961,900 | 0.110762222 | 0.155221151 | 0.04720594 | 0.11598136 |
| 962,000 | 0.110828253 | 0.155194651 | 0.046989483 | 0.115836772 |
| 962,100 | 0.110790474 | 0.154525708 | 0.047020461 | 0.116040363 |
| 962,200 | 0.110762553 | 0.154290708 | 0.046865542 | 0.116023191 |
| 962,300 | 0.110834384 | 0.155247449 | 0.046753411 | 0.11601847 |
| 962,400 | 0.110758804 | 0.155007691 | 0.047054079 | 0.115801673 |
| 962,500 | 0.110738352 | 0.154638646 | 0.046921731 | 0.115871773 |
| 962,600 | 0.110768941 | 0.154814174 | 0.046928833 | 0.116218297 |
| 962,700 | 0.110721606 | 0.154718134 | 0.047461539 | 0.116291675 |
| 962,800 | 0.110820618 | 0.154390787 | 0.047302397 | 0.116091361 |
| 962,900 | 0.11074855 | 0.154119068 | 0.047529939 | 0.116160008 |
| 963,000 | 0.110691993 | 0.153591339 | 0.047769581 | 0.116369603 |
| 963,100 | 0.110660102 | 0.154148876 | 0.047208494 | 0.116395015 |
| 963,200 | 0.110608434 | 0.153477194 | 0.046880372 | 0.116237368 |
| 963,300 | 0.110609051 | 0.153694221 | 0.046955319 | 0.116371872 |
| 963,400 | 0.110703031 | 0.154144867 | 0.046699224 | 0.116105879 |
| 963,500 | 0.110612406 | 0.153939094 | 0.047265606 | 0.11654013 |
| 963,600 | 0.110666294 | 0.15398538 | 0.047311474 | 0.116610763 |
| 963,700 | 0.11070017 | 0.154151143 | 0.04765388 | 0.116286486 |

| | | | | |
|---|---|---|---|---|
| 963,800 | 0.110703159 | 0.153895425 | 0.047535848 | 0.116136691 |
| 963,900 | 0.110710856 | 0.153291861 | 0.047894539 | 0.116585919 |
| 964,000 | 0.110678964 | 0.153686909 | 0.048157656 | 0.115805086 |
| 964,100 | 0.11075372 | 0.154299053 | 0.047831986 | 0.115809225 |
| 964,200 | 0.110726697 | 0.153734543 | 0.047731959 | 0.116493345 |
| 964,300 | 0.110913888 | 0.154250769 | 0.047710472 | 0.116398249 |
| 964,400 | 0.110856188 | 0.153973165 | 0.04760459 | 0.116080591 |
| 964,500 | 0.11068632 | 0.153459416 | 0.047500889 | 0.116346805 |
| 964,600 | 0.110665461 | 0.153326391 | 0.047303037 | 0.116060607 |
| 964,700 | 0.110717215 | 0.153643262 | 0.047284324 | 0.115913915 |
| 964,800 | 0.110592244 | 0.15271587 | 0.047269008 | 0.115930346 |
| 964,900 | 0.110619431 | 0.153717092 | 0.046468307 | 0.116189321 |
| 965,000 | 0.110718819 | 0.153707707 | 0.046926331 | 0.116515753 |
| 965,100 | 0.11073842 | 0.154813321 | 0.046996059 | 0.116424351 |
| 965,200 | 0.11061199 | 0.154213243 | 0.0472224 | 0.116272072 |
| 965,300 | 0.110591064 | 0.153379921 | 0.047518157 | 0.116494155 |
| 965,400 | 0.110594766 | 0.153623399 | 0.047309162 | 0.116742703 |
| 965,500 | 0.110580726 | 0.153237994 | 0.047416058 | 0.116501255 |
| 965,600 | 0.110606002 | 0.153696212 | 0.046811245 | 0.116639379 |
| 965,700 | 0.110667015 | 0.154010729 | 0.04657648 | 0.117214342 |
| 965,800 | 0.110572043 | 0.153345982 | 0.046550128 | 0.117281504 |
| 965,900 | 0.110500678 | 0.153538431 | 0.046941857 | 0.11698164 |
| 966,000 | 0.110500068 | 0.153469938 | 0.046895771 | 0.116691943 |
| 966,100 | 0.110473249 | 0.153238175 | 0.046859005 | 0.116564884 |
| 966,200 | 0.110531471 | 0.153599306 | 0.046772499 | 0.11648033 |
| 966,300 | 0.110441623 | 0.15354139 | 0.046642417 | 0.116898526 |
| 966,400 | 0.110473439 | 0.153882051 | 0.046786295 | 0.11673204 |
| 966,500 | 0.110436279 | 0.153507403 | 0.046749525 | 0.11660532 |
| 966,600 | 0.110526079 | 0.153425464 | 0.046623033 | 0.116331679 |
| 966,700 | 0.110539007 | 0.153846768 | 0.046351565 | 0.116684268 |
| 966,800 | 0.110511359 | 0.153353419 | 0.045976838 | 0.116437722 |
| 966,900 | 0.110493043 | 0.153083607 | 0.046165452 | 0.116407442 |
| 967,000 | 0.110443567 | 0.153273206 | 0.046454436 | 0.116230436 |
| 967,100 | 0.110459722 | 0.152962256 | 0.046133315 | 0.116748598 |
| 967,200 | 0.110425395 | 0.153188683 | 0.046146239 | 0.11730121 |
| 967,300 | 0.110440127 | 0.153364568 | 0.046334352 | 0.117264893 |
| 967,400 | 0.11053036 | 0.15303098 | 0.046377817 | 0.117355543 |
| 967,500 | 0.110467106 | 0.152903603 | 0.046458427 | 0.116838683 |
| 967,600 | 0.11041018 | 0.152876798 | 0.0465974 | 0.117503444 |
| 967,700 | 0.11039766 | 0.153569053 | 0.04664766 | 0.117150591 |
| 967,800 | 0.110380721 | 0.153395292 | 0.046457377 | 0.117308287 |
| 967,900 | 0.110381329 | 0.153052686 | 0.046533722 | 0.117457829 |
| 968,000 | 0.110316828 | 0.153833599 | 0.046694494 | 0.117831631 |
| 968,100 | 0.110248653 | 0.153165112 | 0.046616879 | 0.117798426 |
| 968,200 | 0.110305622 | 0.153144851 | 0.046756217 | 0.117554574 |
| 968,300 | 0.110221061 | 0.152637323 | 0.046671115 | 0.117374512 |
| 968,400 | 0.110306358 | 0.153294011 | 0.046435139 | 0.117555239 |
| 968,500 | 0.110270006 | 0.153455814 | 0.046021201 | 0.117323017 |
| 968,600 | 0.110338111 | 0.153644354 | 0.046127579 | 0.117507734 |
| 968,700 | 0.110322601 | 0.153507668 | 0.046375255 | 0.117343637 |
| 968,800 | 0.110308442 | 0.153419081 | 0.046562634 | 0.11815676 |
| 968,900 | 0.11036472 | 0.153516394 | 0.046599675 | 0.117637871 |
| 969,000 | 0.110300938 | 0.153830348 | 0.046381345 | 0.117688494 |
| 969,100 | 0.110238749 | 0.153362564 | 0.04678831 | 0.118054078 |
| 969,200 | 0.110192709 | 0.153464933 | 0.046489363 | 0.117813477 |
| 969,300 | 0.110182519 | 0.153719442 | 0.046637524 | 0.117514164 |
| 969,400 | 0.110226963 | 0.153452836 | 0.046752093 | 0.117717027 |
| 969,500 | 0.110171819 | 0.153423029 | 0.046557933 | 0.116931696 |
| 969,600 | 0.11008867 | 0.153448472 | 0.046267261 | 0.116986774 |
| 969,700 | 0.110215001 | 0.153829609 | 0.046620424 | 0.117193647 |
| 969,800 | 0.110195869 | 0.153627628 | 0.046527878 | 0.11732956 |
| 969,900 | 0.110327183 | 0.154042867 | 0.046452828 | 0.117822739 |
| 970,000 | 0.110235847 | 0.153581101 | 0.046361499 | 0.117258202 |
| 970,100 | 0.110243708 | 0.15295376 | 0.046641762 | 0.117320992 |
| 970,200 | 0.110206661 | 0.152973469 | 0.046499565 | 0.117302018 |
| 970,300 | 0.110255464 | 0.153071212 | 0.046347338 | 0.117580677 |
| 970,400 | 0.110206546 | 0.152556666 | 0.046694319 | 0.117577095 |

| | | | | |
|---|---|---|---|---|
| 970,500 | 0.110143829 | 0.152880902 | 0.046865913 | 0.117318756 |
| 970,600 | 0.110171405 | 0.15332337 | 0.046675614 | 0.116958489 |
| 970,700 | 0.110078602 | 0.153308326 | 0.046082091 | 0.117052762 |
| 970,800 | 0.110223411 | 0.153373072 | 0.046451015 | 0.117040499 |
| 970,900 | 0.110121869 | 0.15407916 | 0.046933354 | 0.11767055 |
| 971,000 | 0.110162394 | 0.153435945 | 0.046652437 | 0.117367828 |
| 971,100 | 0.110068494 | 0.153284277 | 0.046683354 | 0.117602583 |
| 971,200 | 0.110161997 | 0.152912575 | 0.04670539 | 0.116804207 |
| 971,300 | 0.110181263 | 0.153217077 | 0.046157894 | 0.117166702 |
| 971,400 | 0.110088016 | 0.153583845 | 0.046399065 | 0.11706563 |
| 971,500 | 0.110059667 | 0.153551903 | 0.046251239 | 0.116656944 |
| 971,600 | 0.110204438 | 0.153449018 | 0.046788444 | 0.116843737 |
| 971,700 | 0.110139332 | 0.152887891 | 0.046288726 | 0.116657796 |
| 971,800 | 0.110099753 | 0.153018702 | 0.046985515 | 0.116540727 |
| 971,900 | 0.110089508 | 0.153418471 | 0.046645477 | 0.116832073 |
| 972,000 | 0.110121486 | 0.153127889 | 0.046208961 | 0.116933124 |
| 972,100 | 0.110139875 | 0.153197608 | 0.046101151 | 0.116560112 |
| 972,200 | 0.110062764 | 0.153348263 | 0.046607597 | 0.117298123 |
| 972,300 | 0.110062239 | 0.152705119 | 0.04714113 | 0.11698163 |
| 972,400 | 0.109981731 | 0.152755286 | 0.04676911 | 0.117435431 |
| 972,500 | 0.11006997 | 0.153095047 | 0.047160775 | 0.117400846 |
| 972,600 | 0.11008985 | 0.152974535 | 0.047077922 | 0.117299251 |
| 972,700 | 0.110049539 | 0.152850038 | 0.046631413 | 0.117551894 |
| 972,800 | 0.110101754 | 0.152885419 | 0.046711423 | 0.117490065 |
| 972,900 | 0.110005046 | 0.152591947 | 0.046590383 | 0.117531757 |
| 973,000 | 0.110061607 | 0.152866034 | 0.046472201 | 0.116926041 |
| 973,100 | 0.110031542 | 0.152345158 | 0.046860951 | 0.116757753 |
| 973,200 | 0.110016626 | 0.152102754 | 0.046986038 | 0.116591822 |
| 973,300 | 0.109938123 | 0.152492257 | 0.046893219 | 0.116914955 |
| 973,400 | 0.109944554 | 0.152146152 | 0.046607036 | 0.117196928 |
| 973,500 | 0.109996328 | 0.15225272 | 0.047208194 | 0.117155513 |
| 973,600 | 0.110080849 | 0.152107704 | 0.046989129 | 0.117113474 |
| 973,700 | 0.109982642 | 0.151227776 | 0.04703418 | 0.11717089 |
| 973,800 | 0.10998327 | 0.151983813 | 0.046328313 | 0.117558676 |
| 973,900 | 0.109950457 | 0.152478802 | 0.047325759 | 0.117681718 |
| 974,000 | 0.109892735 | 0.151388807 | 0.046756718 | 0.117137463 |
| 974,100 | 0.109943298 | 0.151586194 | 0.046801217 | 0.117477906 |
| 974,200 | 0.110035254 | 0.151703471 | 0.046997861 | 0.116914559 |
| 974,300 | 0.109912637 | 0.152069567 | 0.047531777 | 0.117084845 |
| 974,400 | 0.11002682 | 0.152445083 | 0.047243211 | 0.116888291 |
| 974,500 | 0.109883118 | 0.152087607 | 0.046983093 | 0.116432801 |
| 974,600 | 0.10993716 | 0.152550173 | 0.047023576 | 0.116840342 |
| 974,700 | 0.110050747 | 0.152989227 | 0.047197226 | 0.11671345 |
| 974,800 | 0.109980202 | 0.151635945 | 0.047172426 | 0.116669278 |
| 974,900 | 0.109902567 | 0.151810019 | 0.047039481 | 0.116666339 |
| 975,000 | 0.109939022 | 0.152330913 | 0.046962488 | 0.116738413 |
| 975,100 | 0.109910583 | 0.151759912 | 0.046721084 | 0.116796757 |
| 975,200 | 0.10988744 | 0.151948044 | 0.047342092 | 0.116710973 |
| 975,300 | 0.109853765 | 0.1518457 | 0.046950193 | 0.116668673 |
| 975,400 | 0.109965194 | 0.152429387 | 0.047059757 | 0.116199004 |
| 975,500 | 0.110022663 | 0.151823452 | 0.047121495 | 0.116786175 |
| 975,600 | 0.110011456 | 0.151661868 | 0.046315573 | 0.116688352 |
| 975,700 | 0.110034278 | 0.151716282 | 0.047417101 | 0.116901639 |
| 975,800 | 0.109968013 | 0.152088578 | 0.046674633 | 0.116839636 |
| 975,900 | 0.109928933 | 0.151705716 | 0.046679211 | 0.116696434 |
| 976,000 | 0.109914796 | 0.151347342 | 0.046488979 | 0.116465542 |
| 976,100 | 0.109892912 | 0.150940264 | 0.046402641 | 0.116443013 |
| 976,200 | 0.10999439 | 0.151387247 | 0.046066174 | 0.116481045 |
| 976,300 | 0.109837737 | 0.151130033 | 0.046244874 | 0.116068882 |
| 976,400 | 0.109918854 | 0.151211924 | 0.04605239 | 0.116756546 |
| 976,500 | 0.109888516 | 0.1513075 | 0.04614556 | 0.116722906 |
| 976,600 | 0.109810688 | 0.151316918 | 0.04656167 | 0.116461539 |
| 976,700 | 0.109705777 | 0.150716871 | 0.046442438 | 0.116416336 |
| 976,800 | 0.109731772 | 0.150647027 | 0.04617361 | 0.116370055 |
| 976,900 | 0.109722789 | 0.150986155 | 0.046437949 | 0.116259903 |
| 977,000 | 0.109736236 | 0.150563011 | 0.046862988 | 0.116475919 |
| 977,100 | 0.109734738 | 0.150979668 | 0.046569273 | 0.116780799 |

| | | | | |
|---|---|---|---|---|
| 977,200 | 0.109804013 | 0.150591896 | 0.0465786 | 0.116424272 |
| 977,300 | 0.109751057 | 0.151454957 | 0.046801995 | 0.116428855 |
| 977,400 | 0.109663024 | 0.151229404 | 0.046705535 | 0.116660571 |
| 977,500 | 0.109702924 | 0.151198682 | 0.046988006 | 0.116413763 |
| 977,600 | 0.109658 | 0.150939394 | 0.046954475 | 0.117024151 |
| 977,700 | 0.109637965 | 0.151097658 | 0.047039233 | 0.116837345 |
| 977,800 | 0.109560132 | 0.150260668 | 0.046770421 | 0.116708787 |
| 977,900 | 0.109738917 | 0.150442018 | 0.046962513 | 0.116948885 |
| 978,000 | 0.10966792 | 0.150297161 | 0.046836367 | 0.116832898 |
| 978,100 | 0.109668265 | 0.15049645 | 0.046622081 | 0.116925139 |
| 978,200 | 0.109688781 | 0.150856457 | 0.046498174 | 0.116755654 |
| 978,300 | 0.1096536 | 0.150722991 | 0.046470684 | 0.116248934 |
| 978,400 | 0.10965917 | 0.15105762 | 0.046670879 | 0.1159189 |
| 978,500 | 0.109580586 | 0.150712271 | 0.046278382 | 0.116223785 |
| 978,600 | 0.109569964 | 0.150936179 | 0.047134727 | 0.115961858 |
| 978,700 | 0.109662558 | 0.150753426 | 0.047420112 | 0.116374755 |
| 978,800 | 0.109690517 | 0.15076546 | 0.04756394 | 0.116097496 |
| 978,900 | 0.109648966 | 0.150700471 | 0.04702663 | 0.116237632 |
| 979,000 | 0.109479261 | 0.150277803 | 0.047297595 | 0.115839995 |
| 979,100 | 0.109623887 | 0.150469028 | 0.047338423 | 0.116050976 |
| 979,200 | 0.109667313 | 0.150995257 | 0.04736023 | 0.116011724 |
| 979,300 | 0.109697484 | 0.150754322 | 0.047278244 | 0.115840867 |
| 979,400 | 0.109664462 | 0.15036293 | 0.047395274 | 0.1160097 |
| 979,500 | 0.109581593 | 0.150832911 | 0.047184439 | 0.115973085 |
| 979,600 | 0.109605188 | 0.150478535 | 0.047101381 | 0.115620966 |
| 979,700 | 0.109484219 | 0.150709756 | 0.047145575 | 0.115830287 |
| 979,800 | 0.109545786 | 0.150962637 | 0.047079714 | 0.115713752 |
| 979,900 | 0.109544381 | 0.150740867 | 0.047572605 | 0.115519939 |
| 980,000 | 0.109510229 | 0.150326374 | 0.047403788 | 0.115552068 |
| 980,100 | 0.109543251 | 0.150919947 | 0.046794493 | 0.115782638 |
| 980,200 | 0.109578813 | 0.150736602 | 0.046287174 | 0.115650607 |
| 980,300 | 0.109451114 | 0.150164156 | 0.046576214 | 0.115434474 |
| 980,400 | 0.10952599 | 0.15075034 | 0.046432938 | 0.115714032 |
| 980,500 | 0.109470033 | 0.150368939 | 0.04671935 | 0.116053044 |
| 980,600 | 0.109493713 | 0.150465675 | 0.046180703 | 0.115619657 |
| 980,700 | 0.109570953 | 0.150678133 | 0.046506826 | 0.115832771 |
| 980,800 | 0.109500185 | 0.151185204 | 0.04676135 | 0.115777514 |
| 980,900 | 0.109540632 | 0.151205034 | 0.046532938 | 0.115965658 |
| 981,000 | 0.109532814 | 0.151393343 | 0.046858967 | 0.115505661 |
| 981,100 | 0.109573546 | 0.151844659 | 0.046535605 | 0.116071091 |
| 981,200 | 0.109555345 | 0.151426546 | 0.046708463 | 0.11565311 |
| 981,300 | 0.109552095 | 0.150898793 | 0.047076081 | 0.115724779 |
| 981,400 | 0.109493009 | 0.151449466 | 0.047040657 | 0.115491991 |
| 981,500 | 0.109425024 | 0.151877711 | 0.046737704 | 0.115110817 |
| 981,600 | 0.109476712 | 0.151844848 | 0.046770923 | 0.115227226 |
| 981,700 | 0.10942124 | 0.151368664 | 0.046540035 | 0.115140052 |
| 981,800 | 0.10949067 | 0.151932297 | 0.046940974 | 0.115401142 |
| 981,900 | 0.109482156 | 0.151283184 | 0.046860178 | 0.115862414 |
| 982,000 | 0.109435559 | 0.150785385 | 0.046882571 | 0.115587913 |
| 982,100 | 0.109524399 | 0.151325026 | 0.046806674 | 0.115646933 |
| 982,200 | 0.109391466 | 0.151532164 | 0.046654505 | 0.115546296 |
| 982,300 | 0.109439209 | 0.151177022 | 0.047072227 | 0.114977559 |
| 982,400 | 0.109333212 | 0.15130982 | 0.047428289 | 0.115428471 |
| 982,500 | 0.109364677 | 0.151073423 | 0.047226527 | 0.115489855 |
| 982,600 | 0.109374386 | 0.150971332 | 0.04726664 | 0.11571413 |
| 982,700 | 0.109458776 | 0.151314699 | 0.047464649 | 0.11569386 |
| 982,800 | 0.109365421 | 0.151056442 | 0.047174805 | 0.115278004 |
| 982,900 | 0.109357952 | 0.151162433 | 0.046724223 | 0.115354049 |
| 983,000 | 0.109323717 | 0.150920504 | 0.047262938 | 0.115405415 |
| 983,100 | 0.109287295 | 0.151340013 | 0.047268624 | 0.114779499 |
| 983,200 | 0.10945525 | 0.151832157 | 0.046506439 | 0.115190704 |
| 983,300 | 0.109417908 | 0.152210356 | 0.046740954 | 0.114827164 |
| 983,400 | 0.109359896 | 0.151677846 | 0.046662388 | 0.115214353 |
| 983,500 | 0.109234917 | 0.151507532 | 0.046380357 | 0.114526881 |
| 983,600 | 0.109358796 | 0.15155302 | 0.046492026 | 0.114567046 |
| 983,700 | 0.109267139 | 0.151478581 | 0.046595167 | 0.114958097 |
| 983,800 | 0.109327547 | 0.151264969 | 0.046352091 | 0.115285306 |

| | | | | |
|---|---|---|---|---|
| 983,900 | 0.109308559 | 0.151372981 | 0.046994977 | 0.114547882 |
| 984,000 | 0.109261223 | 0.151584469 | 0.047046156 | 0.114505201 |
| 984,100 | 0.109271801 | 0.151421471 | 0.047079468 | 0.11485625 |
| 984,200 | 0.109392002 | 0.151327264 | 0.046832714 | 0.114380946 |
| 984,300 | 0.109250376 | 0.151236829 | 0.047236994 | 0.114821993 |
| 984,400 | 0.109191542 | 0.151467249 | 0.047249501 | 0.114345973 |
| 984,500 | 0.109351978 | 0.151538604 | 0.047273731 | 0.114764983 |
| 984,600 | 0.109349487 | 0.151868098 | 0.04709682 | 0.11447594 |
| 984,700 | 0.109199702 | 0.151456178 | 0.04711471 | 0.114624867 |
| 984,800 | 0.109207752 | 0.151144244 | 0.047069414 | 0.114797396 |
| 984,900 | 0.109198396 | 0.151022052 | 0.047158731 | 0.114662206 |
| 985,000 | 0.109248435 | 0.151581632 | 0.047711928 | 0.1145519 |
| 985,100 | 0.109163248 | 0.151070397 | 0.04740436 | 0.114181713 |
| 985,200 | 0.109198633 | 0.150028477 | 0.047096284 | 0.114367181 |
| 985,300 | 0.109201123 | 0.150844512 | 0.047168333 | 0.114453435 |
| 985,400 | 0.109294862 | 0.150828687 | 0.047599489 | 0.114526098 |
| 985,500 | 0.109193471 | 0.150983038 | 0.047307488 | 0.114707741 |
| 985,600 | 0.109237702 | 0.150791904 | 0.047555619 | 0.1143828 |
| 985,700 | 0.109277787 | 0.151022262 | 0.047595693 | 0.114390319 |
| 985,800 | 0.109269726 | 0.151308827 | 0.047766173 | 0.114193717 |
| 985,900 | 0.109095221 | 0.150362931 | 0.04776904 | 0.113955856 |
| 986,000 | 0.109211104 | 0.150751554 | 0.047511578 | 0.114230327 |
| 986,100 | 0.109142245 | 0.150628965 | 0.047255631 | 0.114063161 |
| 986,200 | 0.109222945 | 0.150906019 | 0.047314947 | 0.113906841 |
| 986,300 | 0.109103318 | 0.150909835 | 0.04752748 | 0.113845401 |
| 986,400 | 0.109165601 | 0.150038946 | 0.047127641 | 0.114324072 |
| 986,500 | 0.109041364 | 0.149961032 | 0.047282931 | 0.113997204 |
| 986,600 | 0.108985873 | 0.150288005 | 0.047148997 | 0.113794309 |
| 986,700 | 0.109094883 | 0.150650794 | 0.047701115 | 0.114271411 |
| 986,800 | 0.109074005 | 0.150905159 | 0.047313684 | 0.113904666 |
| 986,900 | 0.109163097 | 0.150911568 | 0.046875533 | 0.114222241 |
| 987,000 | 0.109211212 | 0.15070565 | 0.047014708 | 0.114565429 |
| 987,100 | 0.109193342 | 0.150052346 | 0.047278967 | 0.114413243 |
| 987,200 | 0.109259083 | 0.150641088 | 0.047410879 | 0.114125046 |
| 987,300 | 0.109137549 | 0.150488717 | 0.047512166 | 0.113440381 |
| 987,400 | 0.109143397 | 0.151242666 | 0.047865714 | 0.113933616 |
| 987,500 | 0.109107934 | 0.150835562 | 0.04775126 | 0.114002845 |
| 987,600 | 0.109119579 | 0.151005511 | 0.047636245 | 0.114111026 |
| 987,700 | 0.109065121 | 0.149977824 | 0.047036674 | 0.114177861 |
| 987,800 | 0.109153937 | 0.150473251 | 0.047654935 | 0.114130131 |
| 987,900 | 0.10911342 | 0.150354544 | 0.047648602 | 0.114498824 |
| 988,000 | 0.109124168 | 0.150493381 | 0.047440571 | 0.114081426 |
| 988,100 | 0.109198291 | 0.150651438 | 0.047980706 | 0.113776615 |
| 988,200 | 0.109098123 | 0.150010263 | 0.047935029 | 0.113853407 |
| 988,300 | 0.109015223 | 0.150147948 | 0.047628369 | 0.113765944 |
| 988,400 | 0.109051924 | 0.150230255 | 0.047731337 | 0.113867823 |
| 988,500 | 0.109025141 | 0.150367011 | 0.0470847 | 0.113742914 |
| 988,600 | 0.108980396 | 0.150421248 | 0.047411709 | 0.113805107 |
| 988,700 | 0.109018394 | 0.150836274 | 0.048077455 | 0.113742284 |
| 988,800 | 0.109039345 | 0.150781989 | 0.047658774 | 0.113907343 |
| 988,900 | 0.108949266 | 0.15056373 | 0.047554475 | 0.113496539 |
| 989,000 | 0.109037567 | 0.150261514 | 0.047638516 | 0.113635199 |
| 989,100 | 0.108937039 | 0.150685087 | 0.047768934 | 0.1138138 |
| 989,200 | 0.10898253 | 0.150647738 | 0.047851991 | 0.113972283 |
| 989,300 | 0.109046973 | 0.150739317 | 0.048172538 | 0.113629304 |
| 989,400 | 0.108849108 | 0.150628937 | 0.048202996 | 0.113617582 |
| 989,500 | 0.10891941 | 0.15032827 | 0.047993896 | 0.113459464 |
| 989,600 | 0.108921587 | 0.150727225 | 0.048170304 | 0.113570297 |
| 989,700 | 0.108896891 | 0.150122401 | 0.048151533 | 0.113531911 |
| 989,800 | 0.108873495 | 0.150315727 | 0.048170291 | 0.113422021 |
| 989,900 | 0.108996532 | 0.150139043 | 0.048415838 | 0.11365087 |
| 990,000 | 0.108925476 | 0.149694783 | 0.047969841 | 0.113823099 |
| 990,100 | 0.109003761 | 0.150527781 | 0.047737825 | 0.113945409 |
| 990,200 | 0.108909078 | 0.150181247 | 0.047583598 | 0.113663485 |
| 990,300 | 0.108858026 | 0.149919676 | 0.048037469 | 0.11407978 |
| 990,400 | 0.108871498 | 0.150149362 | 0.048214247 | 0.113748903 |
| 990,500 | 0.108945731 | 0.150245757 | 0.047625156 | 0.114054854 |

| | | | | |
|---|---|---|---|---|
| 990,600 | 0.108786815 | 0.150105054 | 0.047862964 | 0.113765687 |
| 990,700 | 0.108870819 | 0.150334884 | 0.048315295 | 0.113796542 |
| 990,800 | 0.108900344 | 0.150062502 | 0.048043939 | 0.113425284 |
| 990,900 | 0.108958271 | 0.149730067 | 0.047835358 | 0.113512691 |
| 991,000 | 0.108885753 | 0.149942584 | 0.048068963 | 0.113528364 |
| 991,100 | 0.108917756 | 0.150117558 | 0.04794002 | 0.113164557 |
| 991,200 | 0.108959763 | 0.150056918 | 0.04866471 | 0.113357098 |
| 991,300 | 0.108867674 | 0.149859333 | 0.048271785 | 0.113414606 |
| 991,400 | 0.108856983 | 0.148959339 | 0.048080046 | 0.113610355 |
| 991,500 | 0.108855931 | 0.149928317 | 0.048437851 | 0.113389863 |
| 991,600 | 0.108766346 | 0.149734741 | 0.04784718 | 0.113279006 |
| 991,700 | 0.108873989 | 0.149379963 | 0.048363089 | 0.113144608 |
| 991,800 | 0.108780815 | 0.149935152 | 0.048462845 | 0.113207998 |
| 991,900 | 0.108854252 | 0.149529284 | 0.048275148 | 0.113306812 |
| 992,000 | 0.108818896 | 0.149964156 | 0.047997116 | 0.11310859 |
| 992,100 | 0.108751638 | 0.149696019 | 0.047487455 | 0.114118412 |
| 992,200 | 0.10874389 | 0.149861796 | 0.047926207 | 0.114060423 |
| 992,300 | 0.108817382 | 0.150057035 | 0.048736982 | 0.114034877 |
| 992,400 | 0.108794943 | 0.149130492 | 0.048513691 | 0.113717753 |
| 992,500 | 0.10880894 | 0.150156151 | 0.048053806 | 0.113886945 |
| 992,600 | 0.108876852 | 0.149899084 | 0.048528373 | 0.113464567 |
| 992,700 | 0.108790726 | 0.149886885 | 0.048415794 | 0.113643008 |
| 992,800 | 0.108753233 | 0.149865854 | 0.048663361 | 0.11349443 |
| 992,900 | 0.108752459 | 0.149760458 | 0.048613049 | 0.11411385 |
| 993,000 | 0.108787991 | 0.149870051 | 0.048118794 | 0.114115942 |
| 993,100 | 0.108762621 | 0.149138199 | 0.048565144 | 0.114143912 |
| 993,200 | 0.108866055 | 0.149571107 | 0.048575581 | 0.114581639 |
| 993,300 | 0.108796713 | 0.148994299 | 0.04850157 | 0.113947612 |
| 993,400 | 0.108716073 | 0.148710877 | 0.048342457 | 0.113893037 |
| 993,500 | 0.108719179 | 0.148437788 | 0.048409042 | 0.114015137 |
| 993,600 | 0.108852818 | 0.149107851 | 0.048909079 | 0.114849816 |
| 993,700 | 0.108861558 | 0.149107115 | 0.048280098 | 0.114043027 |
| 993,800 | 0.108896329 | 0.148882954 | 0.048262621 | 0.114610808 |
| 993,900 | 0.108670536 | 0.148513759 | 0.048485482 | 0.114250031 |
| 994,000 | 0.108836162 | 0.148699773 | 0.048563236 | 0.114593169 |
| 994,100 | 0.10874386 | 0.148628072 | 0.048133875 | 0.114317852 |
| 994,200 | 0.108761847 | 0.148461262 | 0.048332835 | 0.114406568 |
| 994,300 | 0.108848705 | 0.148680329 | 0.048299104 | 0.114531268 |
| 994,400 | 0.108683411 | 0.148310359 | 0.047712707 | 0.113851128 |
| 994,500 | 0.10874805 | 0.148826775 | 0.048322821 | 0.113963949 |
| 994,600 | 0.108753613 | 0.148931482 | 0.048206427 | 0.114387247 |
| 994,700 | 0.108670691 | 0.148925405 | 0.048095765 | 0.113641686 |
| 994,800 | 0.108748801 | 0.148862788 | 0.04830391 | 0.114396017 |
| 994,900 | 0.108734509 | 0.148745731 | 0.048010626 | 0.11446908 |
| 995,000 | 0.108769247 | 0.149013933 | 0.048594909 | 0.114546685 |
| 995,100 | 0.108646529 | 0.148989087 | 0.047746623 | 0.114366508 |
| 995,200 | 0.108661099 | 0.148573715 | 0.047966868 | 0.114101435 |
| 995,300 | 0.108719181 | 0.148749895 | 0.048348508 | 0.113837886 |
| 995,400 | 0.10865993 | 0.149072325 | 0.047850492 | 0.114032434 |
| 995,500 | 0.108726358 | 0.148671662 | 0.047771222 | 0.114382537 |
| 995,600 | 0.108624456 | 0.14932063 | 0.048623493 | 0.114271063 |
| 995,700 | 0.108761978 | 0.148636874 | 0.048231252 | 0.114197606 |
| 995,800 | 0.108630824 | 0.148769065 | 0.048015983 | 0.113755426 |
| 995,900 | 0.108516793 | 0.14839705 | 0.047978082 | 0.113789918 |
| 996,000 | 0.108498311 | 0.148489318 | 0.047905548 | 0.113805607 |
| 996,100 | 0.108571335 | 0.148671948 | 0.047816983 | 0.11414206 |
| 996,200 | 0.10849075 | 0.148627823 | 0.04802868 | 0.113891175 |
| 996,300 | 0.108550167 | 0.148716521 | 0.047973241 | 0.113456722 |
| 996,400 | 0.108584838 | 0.14886963 | 0.047607745 | 0.114075418 |
| 996,500 | 0.108457899 | 0.149133824 | 0.047307172 | 0.113993637 |
| 996,600 | 0.108574045 | 0.149283991 | 0.047458038 | 0.114185538 |
| 996,700 | 0.108409278 | 0.148146852 | 0.047834124 | 0.11365896 |
| 996,800 | 0.108616269 | 0.149261664 | 0.047839043 | 0.114106877 |
| 996,900 | 0.108506476 | 0.149011741 | 0.047704581 | 0.113629806 |
| 997,000 | 0.108490664 | 0.149226172 | 0.0479136 | 0.114013381 |
| 997,100 | 0.108586861 | 0.14957776 | 0.047367046 | 0.114278023 |
| 997,200 | 0.108463265 | 0.149458708 | 0.047924285 | 0.114037646 |

| | | | | |
|---|---|---|---|---|
| 997,300 | 0.108478854 | 0.148856682 | 0.047938974 | 0.114000067 |
| 997,400 | 0.108503783 | 0.148917054 | 0.047722192 | 0.113921319 |
| 997,500 | 0.108448412 | 0.148524066 | 0.047649194 | 0.113917038 |
| 997,600 | 0.108502637 | 0.148889989 | 0.047573664 | 0.11389444 |
| 997,700 | 0.108448102 | 0.148630077 | 0.047267877 | 0.113765086 |
| 997,800 | 0.108415473 | 0.149258285 | 0.047364277 | 0.113988354 |
| 997,900 | 0.108515769 | 0.149572322 | 0.047443046 | 0.114434875 |
| 998,000 | 0.108450619 | 0.149008796 | 0.047773477 | 0.113867361 |
| 998,100 | 0.108382109 | 0.149548198 | 0.047636811 | 0.113658608 |
| 998,200 | 0.108390376 | 0.148877622 | 0.047606518 | 0.113834689 |
| 998,300 | 0.10840187 | 0.149567797 | 0.047020528 | 0.113508335 |
| 998,400 | 0.108523241 | 0.149830364 | 0.047514934 | 0.113790673 |
| 998,500 | 0.108411241 | 0.149013452 | 0.047121406 | 0.114180889 |
| 998,600 | 0.108426976 | 0.149198031 | 0.047231961 | 0.113837452 |
| 998,700 | 0.108431738 | 0.149558688 | 0.04703419 | 0.114090667 |
| 998,800 | 0.108416027 | 0.149730806 | 0.047371887 | 0.114128172 |
| 998,900 | 0.108390384 | 0.149632349 | 0.046925572 | 0.11382314 |
| 999,000 | 0.108278316 | 0.149341938 | 0.046811435 | 0.114109214 |
| 999,100 | 0.108354587 | 0.149572045 | 0.04702767 | 0.114119589 |
| 999,200 | 0.108316129 | 0.149268581 | 0.047103102 | 0.113977741 |
| 999,300 | 0.108396583 | 0.149870182 | 0.047289621 | 0.113782642 |
| 999,400 | 0.108338959 | 0.149821404 | 0.047384975 | 0.113626893 |
| 999,500 | 0.108304435 | 0.149888994 | 0.046887552 | 0.113892095 |
| 999,600 | 0.10829143 | 0.149425958 | 0.046962299 | 0.11415621 |
| 999,700 | 0.108295252 | 0.14942953 | 0.047124029 | 0.11398248 |
| 999,800 | 0.108253525 | 0.149108339 | 0.047027461 | 0.114221888 |
| 999,900 | 0.108239377 | 0.149206192 | 0.046973284 | 0.11440797 |
| 1,000,000 | 0.108277078 | 0.148983197 | 0.047054606 | 0.114099052 |
| 1,000,100 | 0.108481433 | 0.147698363 | 0.046348796 | 0.117149516 |

**Table S6. Genes located in the selected regions in the Angora rabbit population**

| Genes |
|---|
| MARCH4 |
| ABHD18 |
| ACAP2 |
| ACAT2 |
| ACCSL |
| ACOXL |
| ACTR6 |
| ADAM8 |
| ALCAM |
| ALG9 |
| ALS2 |
| AMDHD1 |
| ANK2 |
| ANKRD6 |
| ANKS1B |
| ANO4 |
| AOX1 |
| AOX2 |
| AOX3 |
| AOX4 |
| APOD |
| ARHGEF12 |
| ARSJ |
| ATF6 |
| AUNIP |
| BAZ2A |
| BAZ2B |
| BBIP1 |
| BCL2L11 |
| BDH1 |
| BIRC8 |
| BMP2 |
| BRD8 |
| BZW1 |
| C15H4orf33 |
| C2H2orf61 |
| CADM2 |
| CADPS2 |
| CALY |
| CARD6 |
| CASP10 |
| CCDC115 |
| CCDC38 |
| CDH7 |
| CDK15 |
| CERS6 |
| CFLAR |
| CHCHD5 |
| CHCHD7 |
| CKAP2L |
| CLGN |
| CLK1 |
| COQ10B |
| CPA6 |
| CRY1 |
| DEPDC4 |

DLG1
DMP1
DPP10
DSPP
DTX4
DUSP12
DUSP5
DYNC1I2
DZIP3
EBF2
ECHS1
ECT2
EIF4E3
EML4
FAM111A
FAM126B
FAM13B
FAM168A
FAM71C
FARSB
FAT4
FCRLA
FCRLB
FUOM
FUT8
GABRR1
GABRR2
GAS2L3
GBE1
GLI2
GLYAT
GLYATL1
GPM6A
GRID1
GRIK4
GTSF1
HERC1
HHLA2
HSPA4L
HSPD1
HSPE1
IL1A
IL1B
IL1F10
IL1RN
IL36A
IL36G
IL36RN
IL37
IMP4
INTU
KCNB2
KCTD18
KIAA1217
KIF20A
LAMP5
LARGE1
LARP1B

LOC100009591
LOC100339265
LOC100340013
LOC100340271
LOC100342427
LOC100342572
LOC100344217
LOC100344466
LOC100344894
LOC100344990
LOC100345055
LOC100345584
LOC100346950
LOC100353150
LOC100356310
LOC100356443
LOC100356569
LOC100357577
LOC100358083
LOC100358341
LOC100358862
LOC100359039
LOC100359291
LOC100359306
LOC103347497
LOC103347695
LOC103348047
LOC103348254
LOC103348668
LOC103348731
LOC103348788
LOC103348854
LOC103348916
LOC103349813
LOC103349946
LOC103350128
LOC103350491
LOC103350856
LOC103350884
LOC103351320
LOC103351321
LOC103351426
LOC103351431
LOC103351445
LOC103351450
LOC103351453
LOC103351749
LOC108176366
LOC108176375
LOC108176501
LOC108176680
LOC108176829
LOC108176864
LOC108176874
LOC108176882
LOC108176929
LOC108176950
LOC108177388

LOC108177389
LOC108177393
LOC108177496
LOC108177497
LOC108177995
LOC108178089
LOC108178131
LOC108178132
LOC108178319
LOC108178452
LOC108178467
LOC108178674
LYRM2
MAGI2
MAGI3
MAN1C1
MCRIP2
MDN1
MFSD1
MFSD8
MGA
MGAT4D
MME
MOS
MPEG1
MTERF2
MTFR1L
MTG1
NCEH1
NCKAP1L
NDST4
NDUFB3
NF1
NHEJ1
NIF3L1
NME5
NPFFR1
NR1H4
NT5DC4
NUP98
OAF
OLFML2B
OLIG3
ORC2
PACRG
PAK5
PAOX
PAPPA2
PAQR7
PARK2
PAX8
PDCD4
PDE11A
PDK1
PGAP1
PGAP2
PKD2L2
PLAG1

PLCB1
PLCB4
PLCXD3
PLK4
PM20D2
PNRC1
POLR1B
POU2F3
PPA1
PPIL3
PPP1R2
PPP2R1B
PRAP1
PRKCQ
PROK2
PSD3
PSD4
PTPN12
PTPN18
PUM2
QKI
RBM20
RBMS2
RFTN2
RHOG
RNF133
RNF148
RNGTT
RPL37A
RPS20
RRAGD
SAR1A
SCLT1
SCN7A
SCN9A
SCYL2
SEPN1
SFMBT2
SGO2
SHOC2
SIK2
SLC17A8
SLC20A1
SLC25A12
SLC25A31
SLC4A10
SLCO5A1
SMARCAL1
SMC3
SNX9
SOX2
SPATA16
SPATA5
SPATS2L
SPRN
SRSF12
SULF1
TAS2R16

| |
|---|
| THADA |
| TMEM136 |
| TMEM263 |
| TNFAIP3 |
| TRAM1L1 |
| TRNAE-UUC |
| TRNAI-AAU-2 |
| TTC21B |
| TTC7A |
| TTLL5 |
| TUBGCP2 |
| UBE2J1 |
| UGT8 |
| UHRF1BP1L |
| WASF1 |
| WNT8A |
| WTAP |
| XKR4 |
| XRCC5 |
| ZBTB16 |
| ZNF511 |

Table S7. Enrichment of genes located in the selected regions in the Angora rabbit population

| Annotation Cluster 1 | Enrichment Score: 4.86 | G | | Count | P_Value | Benjamini |
|---|---|---|---|---|---|---|
| INTERPRO | Interleukin-1 | RT | | 8 | 4.10E-12 | 2.10E-09 |
| INTERPRO | Interleukin-1 receptor antagonist / Interleukin-36 | RT | | 6 | 3.50E-09 | 9.00E-07 |
| INTERPRO | Cytokine, IL-1-like | RT | | 8 | 1.10E-07 | 1.80E-05 |
| INTERPRO | Interleukin-1 conserved site | RT | | 4 | 1.20E-05 | 1.50E-03 |
| UP_KEYWORDS | Cytokine | RT | | 8 | 4.90E-05 | 6.50E-03 |
| UP_KEYWORDS | Secreted | RT | | 12 | 2.90E-02 | 3.80E-01 |
| GOTERM_BP_DIREC | immune response | RT | | 5 | 1.60E-01 | 1.00E+00 |
| GOTERM_CC_DIREC | extracellular space | RT | | 12 | 3.30E-01 | 1.00E+00 |
| Annotation Cluster 2 | Enrichment Score: 2.13 | G | | Count | P_Value | Benjamini |
| KEGG_PATHWAY | Vitamin B6 metabolism | RT | | 4 | 1.30E-04 | 2.40E-02 |
| INTERPRO | Aldehyde oxidase | RT | | 3 | 6.10E-04 | 5.70E-02 |
| KEGG_PATHWAY | Tryptophan metabolism | RT | | 6 | 9.30E-04 | 8.40E-02 |
| INTERPRO | Molybdopterin dehydrogenase, FAD-binding | RT | | 3 | 1.20E-03 | 5.70E-02 |
| INTERPRO | [2Fe-2S]-binding | RT | | 3 | 1.20E-03 | 5.70E-02 |
| INTERPRO | CO dehydrogenase flavoprotein, C-terminal | RT | | 3 | 1.20E-03 | 5.70E-02 |
| INTERPRO | Aldehyde oxidase/xanthine dehydrogenase, molybdopterin binding | RT | | 3 | 1.20E-03 | 5.70E-02 |
| INTERPRO | Aldehyde oxidase/xanthine dehydrogenase | RT | | 3 | 1.20E-03 | 5.70E-02 |
| INTERPRO | Aldehyde oxidase/xanthine dehydrogenase, a/b hammerhead | RT | | 3 | 1.20E-03 | 5.70E-02 |
| SMART | SM01008 | RT | | 3 | 1.20E-03 | 7.20E-02 |
| SMART | SM01092 | RT | | 3 | 1.20E-03 | 7.20E-02 |
| UP_KEYWORDS | Molybdenum | RT | | 3 | 2.00E-03 | 1.30E-01 |
| INTERPRO | 2Fe-2S ferredoxin, iron-sulphur binding site | RT | | 3 | 2.00E-03 | 8.60E-02 |
| KEGG_PATHWAY | Valine, leucine and isoleucine degradation | RT | | 6 | 2.20E-03 | 1.30E-01 |
| PIR_SUPERFAMILY | xanthine dehydrogenase | RT | | 3 | 3.20E-03 | 8.80E-02 |
| INTERPRO | 2Fe-2S ferredoxin-type domain | RT | | 3 | 4.10E-03 | 1.60E-01 |
| UP_KEYWORDS | 2Fe-2S | RT | | 3 | 5.30E-03 | 2.30E-01 |
| GOTERM_MF_DIREC | oxidoreductase activity, acting on CH-OH group of donors | RT | | 3 | 5.40E-03 | 9.60E-01 |
| INTERPRO | FAD-binding, type 2, subdomain 1 | RT | | 3 | 5.40E-03 | 1.80E-01 |
| INTERPRO | FAD-binding, type 2 | RT | | 3 | 6.90E-03 | 2.00E-01 |
| INTERPRO | CO dehydrogenase flavoprotein-like, FAD-binding, subdomain 2 | RT | | 3 | 6.90E-03 | 2.00E-01 |
| INTERPRO | Beta-grasp domain | RT | | 3 | 1.20E-02 | 3.40E-01 |
| KEGG_PATHWAY | Nicotinate and nicotinamide metabolism | RT | | 4 | 1.30E-02 | 5.70E-01 |
| GOTERM_MF_DIREC | 2 iron, 2 sulfur cluster binding | RT | | 3 | 1.70E-02 | 9.90E-01 |
| UP_KEYWORDS | Iron-sulfur | RT | | 3 | 1.90E-02 | 3.80E-01 |
| KEGG_PATHWAY | Tyrosine metabolism | RT | | 4 | 2.10E-02 | 7.40E-01 |
| KEGG_PATHWAY | Drug metabolism - cytochrome P450 | RT | | 5 | 4.00E-02 | 1.00E+00 |
| GOTERM_MF_DIREC | NAD binding | RT | | 3 | 6.10E-02 | 1.00E+00 |
| GOTERM_MF_DIREC | electron carrier activity | RT | | 3 | 8.10E-02 | 1.00E+00 |
| KEGG_PATHWAY | Retinol metabolism | RT | | 4 | 1.30E-01 | 1.00E+00 |
| UP_KEYWORDS | FAD | RT | | 3 | 1.50E-01 | 1.00E+00 |
| GOTERM_MF_DIREC | flavin adenine dinucleotide binding | RT | | 3 | 1.70E-01 | 1.00E+00 |
| UP_KEYWORDS | Flavoprotein | RT | | 3 | 1.70E-01 | 1.00E+00 |
| GOTERM_MF_DIREC | iron ion binding | RT | | 4 | 3.00E-01 | 1.00E+00 |
| UP_KEYWORDS | Iron | RT | | 4 | 3.20E-01 | 1.00E+00 |
| Annotation Cluster 3 | Enrichment Score: 1.59 | G | | Count | P_Value | Benjamini |
| SMART | PLCXc | RT | | 3 | 2.00E-02 | 5.70E-01 |
| INTERPRO | Phospholipase C, phosphatidylinositol-specific , X domain | RT | | 3 | 2.20E-02 | 4.80E-01 |

| | | | | | | |
|---|---|---|---|---|---|---|
| INTERPRO | PLC-like phosphodiesterase, TIM beta/alpha-barrel domain | RT | | 3 | 3.90E-02 | 6.80E-01 |
| Annotation Cluster 4 | Enrichment Score: 1.46 | G | | Count | P_Value | Benjamini |
| INTERPRO | Voltage gated sodium channel, alpha subunit | RT | | 3 | 5.40E-03 | 1.80E-01 |
| INTERPRO | Sodium ion transport-associated | RT | | 3 | 5.40E-03 | 1.80E-01 |
| GOTERM_MF_DIREC | voltage-gated sodium channel activity | RT | | 3 | 1.20E-02 | 9.90E-01 |
| GOTERM_CC_DIREC | voltage-gated sodium channel complex | RT | | 3 | 1.30E-02 | 1.00E+00 |
| UP_KEYWORDS | Ion channel | RT | | 8 | 1.90E-02 | 3.80E-01 |
| UP_KEYWORDS | Sodium channel | RT | | 3 | 2.90E-02 | 3.80E-01 |
| UP_KEYWORDS | Voltage-gated channel | RT | | 4 | 1.00E-01 | 8.90E-01 |
| UP_KEYWORDS | Sodium | RT | | 3 | 1.30E-01 | 1.00E+00 |
| UP_KEYWORDS | Sodium transport | RT | | 3 | 1.30E-01 | 1.00E+00 |
| INTERPRO | Ion transport domain | RT | | 4 | 1.40E-01 | 1.00E+00 |
| INTERPRO | Voltage-dependent potassium channel, four helix bundle domain | RT | | 3 | 1.50E-01 | 1.00E+00 |
| Annotation Cluster 5 | Enrichment Score: 1.37 | G | | Count | P_Value | Benjamini |
| INTERPRO | Guanylate kinase, conserved site | RT | | 3 | 1.70E-02 | 4.30E-01 |
| INTERPRO | Guanylate kinase | RT | | 3 | 3.00E-02 | 5.50E-01 |
| INTERPRO | Guanylate kinase/L-type calcium channel | RT | | 3 | 3.90E-02 | 6.80E-01 |
| SMART | GuKc | RT | | 3 | 4.00E-02 | 9.40E-01 |
| SMART | PDZ | RT | | 5 | 8.30E-02 | 1.00E+00 |
| INTERPRO | PDZ domain | RT | | 5 | 9.30E-02 | 1.00E+00 |
| Annotation Cluster 6 | Enrichment Score: 1.12 | G | | Count | P_Value | Benjamini |
| UP_KEYWORDS | Ion channel | RT | | 8 | 1.90E-02 | 3.80E-01 |
| UP_KEYWORDS | Ion transport | RT | | 10 | 2.00E-02 | 3.80E-01 |
| UP_KEYWORDS | Transport | RT | | 16 | 2.00E-02 | 3.80E-01 |
| UP_KEYWORDS | Synapse | RT | | 4 | 6.30E-02 | 7.50E-01 |
| UP_KEYWORDS | Cell junction | RT | | 4 | 2.00E-01 | 1.00E+00 |
| GOTERM_CC_DIREC | cell junction | RT | | 4 | 2.60E-01 | 1.00E+00 |
| KEGG_PATHWAY | Neuroactive ligand-receptor interaction | RT | | 5 | 6.20E-01 | 1.00E+00 |
| Annotation Cluster 7 | Enrichment Score: 0.92 | G | | Count | P_Value | Benjamini |
| INTERPRO | Protein-tyrosine/Dual specificity phosphatase | RT | | 5 | 2.10E-02 | 4.80E-01 |
| INTERPRO | Protein-tyrosine phosphatase, active site | RT | | 4 | 7.20E-02 | 1.00E+00 |
| INTERPRO | Protein-tyrosine phosphatase, catalytic | RT | | 3 | 1.80E-01 | 1.00E+00 |
| UP_KEYWORDS | Protein phosphatase | RT | | 3 | 1.80E-01 | 1.00E+00 |
| SMART | PTPc_motif | RT | | 3 | 1.90E-01 | 1.00E+00 |
| GOTERM_MF_DIREC | protein tyrosine phosphatase activity | RT | | 3 | 3.20E-01 | 1.00E+00 |
| Annotation Cluster 8 | Enrichment Score: 0.76 | G | | Count | P_Value | Benjamini |
| KEGG_PATHWAY | African trypanosomiasis | RT | | 3 | 8.70E-02 | 1.00E+00 |
| KEGG_PATHWAY | Chagas disease (American trypanosomiasis) | RT | | 5 | 9.60E-02 | 1.00E+00 |
| KEGG_PATHWAY | Alzheimer's disease | RT | | 6 | 1.60E-01 | 1.00E+00 |
| KEGG_PATHWAY | Inflammatory mediator regulation of TRP channels | RT | | 4 | 2.20E-01 | 1.00E+00 |
| KEGG_PATHWAY | Amoebiasis | RT | | 3 | 5.20E-01 | 1.00E+00 |
| Annotation Cluster 9 | Enrichment Score: 0.71 | G | | Count | P_Value | Benjamini |
| KEGG_PATHWAY | Chagas disease (American trypanosomiasis) | RT | | 5 | 9.60E-02 | 1.00E+00 |
| KEGG_PATHWAY | Dopaminergic synapse | RT | | 5 | 1.20E-01 | 1.00E+00 |
| KEGG_PATHWAY | Long-term depression | RT | | 3 | 2.30E-01 | 1.00E+00 |
| KEGG_PATHWAY | Sphingolipid signaling pathway | RT | | 4 | 2.60E-01 | 1.00E+00 |
| KEGG_PATHWAY | Adrenergic signaling in cardiomyocytes | RT | | 4 | 3.90E-01 | 1.00E+00 |
| Annotation Cluster 10 | Enrichment Score: 0.64 | G | | Count | P_Value | Benjamini |
| UP_KEYWORDS | Lipid degradation | RT | | 3 | 8.80E-02 | 8.90E-01 |
| GOTERM_BP_DIREC | lipid catabolic process | RT | | 3 | 1.60E-01 | 1.00E+00 |
| UP_KEYWORDS | Lipid metabolism | RT | | 3 | 3.60E-01 | 1.00E+00 |

| | | | | | |
|---|---|---|---|---|---|
| KEGG_PATHWAY | Pancreatic secretion | RT | 3 | 5.60E-01 | 1.00E+00 |
| Annotation Cluster 11 | Enrichment Score: 0.6 | | Count | P_Value | Benjamini |
| KEGG_PATHWAY | Measles | RT | 5 | 1.50E-01 | 1.00E+00 |
| KEGG_PATHWAY | Tuberculosis | RT | 6 | 1.70E-01 | 1.00E+00 |
| KEGG_PATHWAY | Osteoclast differentiation | RT | 3 | 6.30E-01 | 1.00E+00 |
| Annotation Cluster 12 | Enrichment Score: 0.47 | | Count | P_Value | Benjamini |
| GOTERM_BP_DIREC | intracellular signal transduction | RT | 6 | 2.00E-01 | 1.00E+00 |
| KEGG_PATHWAY | Inflammatory mediator regulation of TRP channels | RT | 4 | 2.20E-01 | 1.00E+00 |
| KEGG_PATHWAY | Vascular smooth muscle contraction | RT | 4 | 3.10E-01 | 1.00E+00 |
| INTERPRO | C2 calcium-dependent membrane targeting | RT | 3 | 5.60E-01 | 1.00E+00 |
| KEGG_PATHWAY | Platelet activation | RT | 3 | 6.00E-01 | 1.00E+00 |
| Annotation Cluster 13 | Enrichment Score: 0.47 | | Count | P_Value | Benjamini |
| INTERPRO | Pleckstrin homology-like domain | RT | 9 | 1.20E-01 | 1.00E+00 |
| SMART | PH | RT | 5 | 3.50E-01 | 1.00E+00 |
| INTERPRO | Pleckstrin homology domain | RT | 5 | 3.70E-01 | 1.00E+00 |
| KEGG_PATHWAY | Endocytosis | RT | 3 | 9.10E-01 | 1.00E+00 |
| Annotation Cluster 14 | Enrichment Score: 0.42 | | Count | P_Value | Benjamini |
| UP_KEYWORDS | Metal-binding | RT | 18 | 3.20E-01 | 1.00E+00 |
| UP_KEYWORDS | Zinc-finger | RT | 9 | 3.50E-01 | 1.00E+00 |
| UP_KEYWORDS | Zinc | RT | 11 | 5.00E-01 | 1.00E+00 |
| Annotation Cluster 15 | Enrichment Score: 0.41 | | Count | P_Value | Benjamini |
| UP_SEQ_FEATURE | glycosylation site:N-linked (GlcNAc...) | RT | 7 | 1.20E-01 | 1.00E+00 |
| UP_KEYWORDS | Glycoprotein | RT | 8 | 1.70E-01 | 1.00E+00 |
| UP_KEYWORDS | Phosphoprotein | RT | 8 | 3.80E-01 | 1.00E+00 |
| UP_SEQ_FEATURE | disulfide bond | RT | 4 | 5.70E-01 | 1.00E+00 |
| UP_SEQ_FEATURE | signal peptide | RT | 3 | 8.40E-01 | 1.00E+00 |
| UP_SEQ_FEATURE | transmembrane region | RT | 3 | 9.10E-01 | 1.00E+00 |
| Annotation Cluster 16 | Enrichment Score: 0.37 | | Count | P_Value | Benjamini |
| GOTERM_MF_DIREC | protein serine/threonine kinase activity | RT | 6 | 9.50E-02 | 1.00E+00 |
| INTERPRO | Protein kinase, ATP binding site | RT | 7 | 2.90E-01 | 1.00E+00 |
| UP_KEYWORDS | Transferase | RT | 12 | 3.10E-01 | 1.00E+00 |
| UP_KEYWORDS | Serine/threonine-protein kinase | RT | 4 | 3.90E-01 | 1.00E+00 |
| INTERPRO | Protein kinase, catalytic domain | RT | 8 | 3.90E-01 | 1.00E+00 |
| SMART | S_TKc | RT | 6 | 4.50E-01 | 1.00E+00 |
| INTERPRO | Protein kinase-like domain | RT | 8 | 4.80E-01 | 1.00E+00 |
| GOTERM_MF_DIREC | ATP binding | RT | 17 | 4.80E-01 | 1.00E+00 |
| INTERPRO | Serine/threonine-protein kinase, active site | RT | 5 | 5.30E-01 | 1.00E+00 |
| UP_KEYWORDS | Kinase | RT | 5 | 6.20E-01 | 1.00E+00 |
| UP_KEYWORDS | Nucleotide-binding | RT | 10 | 8.20E-01 | 1.00E+00 |
| UP_KEYWORDS | ATP-binding | RT | 7 | 9.00E-01 | 1.00E+00 |
| Annotation Cluster 17 | Enrichment Score: 0.35 | | Count | P_Value | Benjamini |
| GOTERM_MF_DIREC | transcriptional activator activity, RNA polymerase II core promoter proximal region sequence-specific binding | RT | 5 | 1.90E-01 | 1.00E+00 |
| UP_KEYWORDS | Transcription | RT | 6 | 2.30E-01 | 1.00E+00 |
| UP_KEYWORDS | Transcription regulation | RT | 4 | 5.30E-01 | 1.00E+00 |
| UP_KEYWORDS | Nucleus | RT | 11 | 6.20E-01 | 1.00E+00 |
| GOTERM_BP_DIREC | transcription, DNA-templated | RT | 4 | 7.10E-01 | 1.00E+00 |
| UP_KEYWORDS | DNA-binding | RT | 6 | 7.80E-01 | 1.00E+00 |
| Annotation Cluster 18 | Enrichment Score: 0.29 | | Count | P_Value | Benjamini |
| KEGG_PATHWAY | Melanogenesis | RT | 3 | 4.50E-01 | 1.00E+00 |
| KEGG_PATHWAY | Pathways in cancer | RT | 8 | 4.70E-01 | 1.00E+00 |
| KEGG_PATHWAY | Wnt signaling pathway | RT | 3 | 6.30E-01 | 1.00E+00 |
| Annotation Cluster 19 | Enrichment Score: 0.15 | | Count | P_Value | Benjamini |
| UP_KEYWORDS | ANK repeat | RT | 3 | 6.70E-01 | 1.00E+00 |
| SMART | ANK | RT | 3 | 7.00E-01 | 1.00E+00 |
| INTERPRO | Ankyrin repeat | RT | 3 | 7.10E-01 | 1.00E+00 |

| | | | | | | |
|---|---|---|---|---|---|---|
| INTERPRO | Ankyrin repeat-containing domain | RT | | 3 | 7.30E-01 | 1.00E+00 |
| Annotation Cluster 20 | Enrichment Score: 0.15 | G | | Count | P_Value | Benjamini |
| INTERPRO | Zinc finger, RING/FYVE/PHD-type | RT | | 6 | 5.70E-01 | 1.00E+00 |
| SMART | RING | RT | | 3 | 7.40E-01 | 1.00E+00 |
| INTERPRO | Zinc finger, RING-type | RT | | 3 | 8.20E-01 | 1.00E+00 |
| Annotation Cluster 21 | Enrichment Score: 0.14 | G | | Count | P_Value | Benjamini |
| KEGG_PATHWAY | Huntington's disease | RT | | 4 | 6.40E-01 | 1.00E+00 |
| KEGG_PATHWAY | cGMP-PKG signaling pathway | RT | | 3 | 7.40E-01 | 1.00E+00 |
| KEGG_PATHWAY | Calcium signaling pathway | RT | | 3 | 7.90E-01 | 1.00E+00 |
| Annotation Cluster 22 | Enrichment Score: 0.11 | G | | Count | P_Value | Benjamini |
| SMART | RRM | RT | | 3 | 7.00E-01 | 1.00E+00 |
| INTERPRO | RNA recognition motif domain | RT | | 3 | 7.20E-01 | 1.00E+00 |
| INTERPRO | Nucleotide-binding, alpha-beta plait | RT | | 3 | 8.20E-01 | 1.00E+00 |
| GOTERM_MF_DIREC | nucleotide binding | RT | | 3 | 8.80E-01 | 1.00E+00 |
| Annotation Cluster 23 | Enrichment Score: 0.11 | G | | Count | P_Value | Benjamini |
| INTERPRO | Immunoglobulin subtype 2 | RT | | 3 | 7.20E-01 | 1.00E+00 |
| SMART | IGc2 | RT | | 3 | 7.30E-01 | 1.00E+00 |
| INTERPRO | Immunoglobulin subtype | RT | | 4 | 7.40E-01 | 1.00E+00 |
| SMART | IG | RT | | 4 | 7.60E-01 | 1.00E+00 |
| INTERPRO | Immunoglobulin-like fold | RT | | 6 | 8.50E-01 | 1.00E+00 |
| INTERPRO | Immunoglobulin-like domain | RT | | 4 | 8.80E-01 | 1.00E+00 |
| Annotation Cluster 24 | Enrichment Score: 0.06 | G | | Count | P_Value | Benjamini |
| INTERPRO | WD40 repeat | RT | | 3 | 8.40E-01 | 1.00E+00 |
| SMART | WD40 | RT | | 3 | 8.40E-01 | 1.00E+00 |
| INTERPRO | WD40-repeat-containing domain | RT | | 3 | 8.80E-01 | 1.00E+00 |
| INTERPRO | WD40/YVTN repeat-like-containing domain | RT | | 3 | 9.20E-01 | 1.00E+00 |
| Annotation Cluster 25 | Enrichment Score: 0.03 | G | | Count | P_Value | Benjamini |
| UP_KEYWORDS | Membrane | RT | | 55 | 9.10E-01 | 1.00E+00 |
| UP_KEYWORDS | Transmembrane helix | RT | | 51 | 9.40E-01 | 1.00E+00 |
| UP_KEYWORDS | Transmembrane | RT | | 51 | 9.40E-01 | 1.00E+00 |
| GOTERM_CC_DIREC | integral component of membrane | RT | | 45 | 9.80E-01 | 1.00E+00 |
| Annotation Cluster 26 | Enrichment Score: 0.03 | G | | Count | P_Value | Benjamini |
| UP_KEYWORDS | Cell membrane | RT | | 14 | 6.50E-01 | 1.00E+00 |
| UP_KEYWORDS | Transducer | RT | | 9 | 9.40E-01 | 1.00E+00 |
| UP_KEYWORDS | Sensory transduction | RT | | 6 | 9.50E-01 | 1.00E+00 |
| UP_KEYWORDS | Receptor | RT | | 10 | 9.70E-01 | 1.00E+00 |
| UP_KEYWORDS | Olfaction | RT | | 5 | 9.70E-01 | 1.00E+00 |
| KEGG_PATHWAY | Olfactory transduction | RT | | 9 | 9.80E-01 | 1.00E+00 |
| INTERPRO | Olfactory receptor | RT | | 5 | 9.80E-01 | 1.00E+00 |
| GOTERM_MF_DIREC | olfactory receptor activity | RT | | 5 | 9.80E-01 | 1.00E+00 |
| UP_KEYWORDS | G-protein coupled receptor | RT | | 7 | 9.90E-01 | 1.00E+00 |
| GOTERM_MF_DIREC | G-protein coupled receptor activity | RT | | 6 | 9.90E-01 | 1.00E+00 |
| INTERPRO | G protein-coupled receptor, rhodopsin-like | RT | | 6 | 9.90E-01 | 1.00E+00 |
| INTERPRO | GPCR, rhodopsin-like, 7TM | RT | | 6 | 1.00E+00 | 1.00E+00 |
| Annotation Cluster 27 | Enrichment Score: 0.02 | G | | Count | P_Value | Benjamini |
| INTERPRO | Zinc finger, C2H2 | RT | | 4 | 9.20E-01 | 1.00E+00 |
| INTERPRO | Zinc finger C2H2-type/integrase DNA-binding domain | RT | | 3 | 9.40E-01 | 1.00E+00 |
| INTERPRO | Zinc finger, C2H2-like | RT | | 3 | 9.70E-01 | 1.00E+00 |
| SMART | ZnF_C2H2 | RT | | 3 | 9.70E-01 | 1.00E+00 |

**Table S8. LD decay in the Angora and wild rabbit populations**

| Distance | Mean_r^2 of Ango | Mean_r^2 of wild rabbits |
|---|---|---|
| 10 | 0.743611655 | 0.50682634 |
| 20 | 0.692571625 | 0.39128698 |
| 30 | 0.665479069 | 0.3331901 |
| 40 | 0.643360062 | 0.28984015 |
| 50 | 0.630016601 | 0.26748072 |
| 60 | 0.619609069 | 0.25199509 |
| 70 | 0.612139647 | 0.24099891 |
| 80 | 0.605729566 | 0.23360797 |
| 90 | 0.600405219 | 0.22978141 |
| 100 | 0.596058253 | 0.22765199 |
| 200 | 0.579923758 | 0.21931972 |
| 300 | 0.565864033 | 0.21181427 |
| 400 | 0.556155585 | 0.20859467 |
| 500 | 0.549360011 | 0.20677943 |
| 600 | 0.5441067 | 0.20529369 |
| 700 | 0.539581187 | 0.2042451 |
| 800 | 0.535444634 | 0.20314872 |
| 900 | 0.531199396 | 0.20219323 |
| 1,000 | 0.52768963 | 0.20131013 |
| 1,100 | 0.524447929 | 0.20058087 |
| 1,200 | 0.521148829 | 0.19980995 |
| 1,300 | 0.518135944 | 0.19914542 |
| 1,400 | 0.515321294 | 0.19859045 |
| 1,500 | 0.512655026 | 0.19799451 |
| 1,600 | 0.509888297 | 0.19745996 |
| 1,700 | 0.507196526 | 0.19692078 |
| 1,800 | 0.504767117 | 0.19635834 |
| 1,900 | 0.50243946 | 0.19593242 |
| 2,000 | 0.500447435 | 0.19548674 |
| 2,100 | 0.498152376 | 0.19500578 |
| 2,200 | 0.496252443 | 0.19468412 |
| 2,300 | 0.494246217 | 0.19439821 |
| 2,400 | 0.492423426 | 0.19407362 |
| 2,500 | 0.490393644 | 0.19367899 |
| 2,600 | 0.48845158 | 0.19336851 |
| 2,700 | 0.486726043 | 0.19308817 |
| 2,800 | 0.485095191 | 0.1927858 |
| 2,900 | 0.48343014 | 0.19246164 |
| 3,000 | 0.482141745 | 0.19222758 |
| 3,100 | 0.480708498 | 0.19188541 |
| 3,200 | 0.479062201 | 0.19171538 |
| 3,300 | 0.477543773 | 0.1915189 |
| 3,400 | 0.476030947 | 0.1913341 |
| 3,500 | 0.474681094 | 0.19114406 |
| 3,600 | 0.473267159 | 0.19086203 |
| 3,700 | 0.472061481 | 0.19067097 |
| 3,800 | 0.470654014 | 0.19042558 |
| 3,900 | 0.46955169 | 0.1904208 |
| 4,000 | 0.468192573 | 0.19015964 |
| 4,100 | 0.467078845 | 0.18995541 |
| 4,200 | 0.465897702 | 0.18981139 |
| 4,300 | 0.464494623 | 0.18972022 |
| 4,400 | 0.463280766 | 0.18936573 |
| 4,500 | 0.462232556 | 0.18924068 |
| 4,600 | 0.461403017 | 0.18911419 |
| 4,700 | 0.460317331 | 0.18893598 |
| 4,800 | 0.459276213 | 0.18887481 |
| 4,900 | 0.458371329 | 0.18869738 |
| 5,000 | 0.457072724 | 0.18864235 |
| 5,100 | 0.456325249 | 0.18852701 |
| 5,200 | 0.455166175 | 0.18843889 |
| 5,300 | 0.454242381 | 0.18817217 |
| 5,400 | 0.453382402 | 0.18813699 |
| 5,500 | 0.452387243 | 0.18795726 |
| 5,600 | 0.45137214 | 0.18769904 |

| | | |
|---|---|---|
| 5,700 | 0.450344937 | 0.1877457 |
| 5,800 | 0.449432721 | 0.1876368 |
| 5,900 | 0.448634285 | 0.18754853 |
| 6,000 | 0.447780744 | 0.18756221 |
| 6,100 | 0.446947018 | 0.18733774 |
| 6,200 | 0.446141384 | 0.18730055 |
| 6,300 | 0.445275358 | 0.18718133 |
| 6,400 | 0.444497698 | 0.1870502 |
| 6,500 | 0.443714695 | 0.18699756 |
| 6,600 | 0.442711331 | 0.18683136 |
| 6,700 | 0.441996774 | 0.18684595 |
| 6,800 | 0.441154667 | 0.18672467 |
| 6,900 | 0.440373888 | 0.18675476 |
| 7,000 | 0.43989469 | 0.1865516 |
| 7,100 | 0.43888283 | 0.18645657 |
| 7,200 | 0.438275655 | 0.18644307 |
| 7,300 | 0.437479027 | 0.18638683 |
| 7,400 | 0.436891387 | 0.18628074 |
| 7,500 | 0.436340615 | 0.18619992 |
| 7,600 | 0.435682664 | 0.18616803 |
| 7,700 | 0.435022568 | 0.18612153 |
| 7,800 | 0.434260731 | 0.18601351 |
| 7,900 | 0.433658845 | 0.18595893 |
| 8,000 | 0.432889318 | 0.18589382 |
| 8,100 | 0.432374258 | 0.1858161 |
| 8,200 | 0.431483267 | 0.18572364 |
| 8,300 | 0.430963795 | 0.18579342 |
| 8,400 | 0.430134655 | 0.18568377 |
| 8,500 | 0.429617679 | 0.18561632 |
| 8,600 | 0.429189963 | 0.18555699 |
| 8,700 | 0.428284959 | 0.1855289 |
| 8,800 | 0.428072455 | 0.18545835 |
| 8,900 | 0.427411171 | 0.18536728 |
| 9,000 | 0.42675314 | 0.18535525 |
| 9,100 | 0.426279184 | 0.18532654 |
| 9,200 | 0.425572948 | 0.18529511 |
| 9,300 | 0.424874723 | 0.18529364 |
| 9,400 | 0.424430327 | 0.18515575 |
| 9,500 | 0.42408216 | 0.18506464 |
| 9,600 | 0.423492428 | 0.18501104 |
| 9,700 | 0.422872297 | 0.18501018 |
| 9,800 | 0.422484893 | 0.18492303 |
| 9,900 | 0.421769254 | 0.18484097 |
| 10,000 | 0.421162547 | 0.18484032 |
| 10,100 | 0.420659664 | 0.18475431 |
| 10,200 | 0.420153405 | 0.18470284 |
| 10,300 | 0.419819446 | 0.18480183 |
| 10,400 | 0.419358728 | 0.18485028 |
| 10,500 | 0.418699387 | 0.18474074 |
| 10,600 | 0.418190883 | 0.18465241 |
| 10,700 | 0.417503085 | 0.18459622 |
| 10,800 | 0.416882045 | 0.1845888 |
| 10,900 | 0.416802801 | 0.18453078 |
| 11,000 | 0.41614665 | 0.18449065 |
| 11,100 | 0.415662755 | 0.18460311 |
| 11,200 | 0.415033555 | 0.18451627 |
| 11,300 | 0.414719356 | 0.18449517 |
| 11,400 | 0.414074006 | 0.18444268 |
| 11,500 | 0.413757889 | 0.18438753 |
| 11,600 | 0.413176702 | 0.18437193 |
| 11,700 | 0.412890555 | 0.18432258 |
| 11,800 | 0.412228573 | 0.18421968 |
| 11,900 | 0.411978983 | 0.18426607 |
| 12,000 | 0.411440759 | 0.18423475 |
| 12,100 | 0.411265403 | 0.1841862 |
| 12,200 | 0.410579091 | 0.18416858 |
| 12,300 | 0.410464884 | 0.18404906 |

| | | |
|---|---|---|
| 12,400 | 0.409937417 | 0.1839938 |
| 12,500 | 0.409313521 | 0.1840449 |
| 12,600 | 0.408797209 | 0.18397646 |
| 12,700 | 0.408440293 | 0.18396175 |
| 12,800 | 0.408031004 | 0.18391333 |
| 12,900 | 0.407299959 | 0.18400705 |
| 13,000 | 0.406847401 | 0.18390465 |
| 13,100 | 0.406695819 | 0.18383082 |
| 13,200 | 0.406225816 | 0.1837736 |
| 13,300 | 0.405679001 | 0.18379242 |
| 13,400 | 0.405139486 | 0.18380129 |
| 13,500 | 0.404863815 | 0.18375321 |
| 13,600 | 0.404638921 | 0.18374536 |
| 13,700 | 0.4039087 | 0.1836533 |
| 13,800 | 0.403513715 | 0.18361999 |
| 13,900 | 0.40309037 | 0.18357666 |
| 14,000 | 0.402626547 | 0.18355973 |
| 14,100 | 0.402289505 | 0.18360123 |
| 14,200 | 0.401914929 | 0.18354374 |
| 14,300 | 0.40149496 | 0.1834418 |
| 14,400 | 0.401482885 | 0.18352698 |
| 14,500 | 0.400856249 | 0.18344525 |
| 14,600 | 0.400355608 | 0.18347324 |
| 14,700 | 0.399895381 | 0.18344281 |
| 14,800 | 0.399592552 | 0.18343489 |
| 14,900 | 0.398934052 | 0.18342949 |
| 15,000 | 0.398669926 | 0.18330635 |
| 15,100 | 0.398319511 | 0.18328115 |
| 15,200 | 0.397789831 | 0.18336585 |
| 15,300 | 0.397358259 | 0.18333352 |
| 15,400 | 0.397053906 | 0.18331029 |
| 15,500 | 0.396743814 | 0.18326157 |
| 15,600 | 0.396237957 | 0.18314363 |
| 15,700 | 0.39615237 | 0.18326626 |
| 15,800 | 0.395625009 | 0.18320228 |
| 15,900 | 0.395337937 | 0.18313584 |
| 16,000 | 0.394927224 | 0.18321622 |
| 16,100 | 0.394462724 | 0.18317682 |
| 16,200 | 0.394294539 | 0.18313915 |
| 16,300 | 0.393853282 | 0.18314679 |
| 16,400 | 0.39355145 | 0.18309509 |
| 16,500 | 0.393423258 | 0.18313481 |
| 16,600 | 0.392640473 | 0.18310698 |
| 16,700 | 0.39254734 | 0.18309345 |
| 16,800 | 0.392046504 | 0.18304081 |
| 16,900 | 0.391706604 | 0.18296972 |
| 17,000 | 0.391414083 | 0.18294575 |
| 17,100 | 0.390998297 | 0.18286068 |
| 17,200 | 0.390597744 | 0.18290347 |
| 17,300 | 0.390505354 | 0.18281034 |
| 17,400 | 0.390444503 | 0.1828764 |
| 17,500 | 0.390086544 | 0.18277634 |
| 17,600 | 0.389429054 | 0.1828449 |
| 17,700 | 0.389364621 | 0.18285524 |
| 17,800 | 0.388928947 | 0.18292169 |
| 17,900 | 0.388459202 | 0.18272487 |
| 18,000 | 0.388077304 | 0.18275485 |
| 18,100 | 0.38767187 | 0.18277034 |
| 18,200 | 0.387506672 | 0.18278426 |
| 18,300 | 0.387334356 | 0.18272415 |
| 18,400 | 0.387048171 | 0.18280521 |
| 18,500 | 0.386572106 | 0.1827187 |
| 18,600 | 0.386233144 | 0.18271 |
| 18,700 | 0.386074446 | 0.18268069 |
| 18,800 | 0.385579707 | 0.18255767 |
| 18,900 | 0.385044036 | 0.182625 |
| 19,000 | 0.384814497 | 0.18255711 |

| | | |
|---|---|---|
| 19,100 | 0.384568159 | 0.18256474 |
| 19,200 | 0.384325716 | 0.18262265 |
| 19,300 | 0.383948204 | 0.18254894 |
| 19,400 | 0.383499362 | 0.18269951 |
| 19,500 | 0.38323987 | 0.18255725 |
| 19,600 | 0.383047735 | 0.18244832 |
| 19,700 | 0.382841652 | 0.18248197 |
| 19,800 | 0.382587816 | 0.18257442 |
| 19,900 | 0.382309682 | 0.18253198 |
| 20,000 | 0.381901623 | 0.18237795 |
| 20,100 | 0.381539372 | 0.1825634 |
| 20,200 | 0.381083016 | 0.18245181 |
| 20,300 | 0.380902757 | 0.18251188 |
| 20,400 | 0.380762799 | 0.18248832 |
| 20,500 | 0.380166179 | 0.18236838 |
| 20,600 | 0.380029477 | 0.18236674 |
| 20,700 | 0.379987094 | 0.18244552 |
| 20,800 | 0.379478245 | 0.18237392 |
| 20,900 | 0.379312255 | 0.18232858 |
| 21,000 | 0.378891788 | 0.18230761 |
| 21,100 | 0.378747967 | 0.1823121 |
| 21,200 | 0.378401525 | 0.18226895 |
| 21,300 | 0.378066719 | 0.18228336 |
| 21,400 | 0.377786448 | 0.18227278 |
| 21,500 | 0.377651784 | 0.18220564 |
| 21,600 | 0.37751663 | 0.18222109 |
| 21,700 | 0.376921306 | 0.18220117 |
| 21,800 | 0.37656719 | 0.18223759 |
| 21,900 | 0.376466362 | 0.18222397 |
| 22,000 | 0.376178186 | 0.18217597 |
| 22,100 | 0.375922072 | 0.18210659 |
| 22,200 | 0.37574452 | 0.18215659 |
| 22,300 | 0.375491048 | 0.18219022 |
| 22,400 | 0.375269933 | 0.18219823 |
| 22,500 | 0.374976132 | 0.18219718 |
| 22,600 | 0.374616646 | 0.18222585 |
| 22,700 | 0.374694764 | 0.18211754 |
| 22,800 | 0.374084915 | 0.18219057 |
| 22,900 | 0.373624899 | 0.18207204 |
| 23,000 | 0.37367785 | 0.18206711 |
| 23,100 | 0.373459909 | 0.18211472 |
| 23,200 | 0.373328967 | 0.18210995 |
| 23,300 | 0.373081521 | 0.18204061 |
| 23,400 | 0.372546859 | 0.181946 |
| 23,500 | 0.372507632 | 0.18197993 |
| 23,600 | 0.372129948 | 0.18207309 |
| 23,700 | 0.371812164 | 0.18193541 |
| 23,800 | 0.371700791 | 0.18190941 |
| 23,900 | 0.37150787 | 0.18195333 |
| 24,000 | 0.371191657 | 0.18205404 |
| 24,100 | 0.371097526 | 0.18212149 |
| 24,200 | 0.370621121 | 0.18206529 |
| 24,300 | 0.370418469 | 0.18195879 |
| 24,400 | 0.37032501 | 0.18193227 |
| 24,500 | 0.370119006 | 0.18194004 |
| 24,600 | 0.369518759 | 0.18191734 |
| 24,700 | 0.369477869 | 0.18189348 |
| 24,800 | 0.369295645 | 0.18199258 |
| 24,900 | 0.368900692 | 0.18196612 |
| 25,000 | 0.368903626 | 0.18197329 |
| 25,100 | 0.368648309 | 0.18186754 |
| 25,200 | 0.36829216 | 0.18190234 |
| 25,300 | 0.368245229 | 0.18186206 |
| 25,400 | 0.36782171 | 0.18188036 |
| 25,500 | 0.36757708 | 0.18177082 |
| 25,600 | 0.367364569 | 0.18183916 |
| 25,700 | 0.366926129 | 0.18180482 |

| | | |
|---|---|---|
| 25,800 | 0.366831803 | 0.18196056 |
| 25,900 | 0.366438051 | 0.18190927 |
| 26,000 | 0.366511331 | 0.1819017 |
| 26,100 | 0.366120459 | 0.18180781 |
| 26,200 | 0.366063585 | 0.18184826 |
| 26,300 | 0.365613244 | 0.18182715 |
| 26,400 | 0.365256464 | 0.18174395 |
| 26,500 | 0.365047796 | 0.18182901 |
| 26,600 | 0.36487475 | 0.18185636 |
| 26,700 | 0.364575153 | 0.18184961 |
| 26,800 | 0.36424842 | 0.18183495 |
| 26,900 | 0.36415335 | 0.18179007 |
| 27,000 | 0.363758191 | 0.18171264 |
| 27,100 | 0.363455007 | 0.18170644 |
| 27,200 | 0.363523893 | 0.18167517 |
| 27,300 | 0.363376625 | 0.181652 |
| 27,400 | 0.363233367 | 0.18164312 |
| 27,500 | 0.363023218 | 0.18174616 |
| 27,600 | 0.362659949 | 0.18167265 |
| 27,700 | 0.362297695 | 0.18171643 |
| 27,800 | 0.36227908 | 0.18171817 |
| 27,900 | 0.362202822 | 0.18171868 |
| 28,000 | 0.361801083 | 0.18177661 |
| 28,100 | 0.361408603 | 0.18165638 |
| 28,200 | 0.361412047 | 0.18162992 |
| 28,300 | 0.361318406 | 0.18165997 |
| 28,400 | 0.361161945 | 0.18161368 |
| 28,500 | 0.360599245 | 0.18163881 |
| 28,600 | 0.36049845 | 0.18157639 |
| 28,700 | 0.360354351 | 0.18160611 |
| 28,800 | 0.360248268 | 0.18156127 |
| 28,900 | 0.360105591 | 0.18167908 |
| 29,000 | 0.359750657 | 0.18155798 |
| 29,100 | 0.359519508 | 0.18156576 |
| 29,200 | 0.359285786 | 0.18152368 |
| 29,300 | 0.359282957 | 0.18149633 |
| 29,400 | 0.359085099 | 0.18146818 |
| 29,500 | 0.359042033 | 0.1815418 |
| 29,600 | 0.358598289 | 0.18152125 |
| 29,700 | 0.358242478 | 0.1814668 |
| 29,800 | 0.358111503 | 0.1814501 |
| 29,900 | 0.357862049 | 0.18147783 |
| 30,000 | 0.357779979 | 0.18144425 |
| 30,100 | 0.357765558 | 0.18146626 |
| 30,200 | 0.357219723 | 0.18145274 |
| 30,300 | 0.357060555 | 0.18151087 |
| 30,400 | 0.356938966 | 0.18150804 |
| 30,500 | 0.356679571 | 0.18140706 |
| 30,600 | 0.35658042 | 0.18145861 |
| 30,700 | 0.356646734 | 0.18145568 |
| 30,800 | 0.356456957 | 0.18138836 |
| 30,900 | 0.35610353 | 0.18146358 |
| 31,000 | 0.355519646 | 0.18145998 |
| 31,100 | 0.355303942 | 0.18146103 |
| 31,200 | 0.355197065 | 0.18143899 |
| 31,300 | 0.354946192 | 0.18140795 |
| 31,400 | 0.354837241 | 0.1813107 |
| 31,500 | 0.354668774 | 0.18140697 |
| 31,600 | 0.354693134 | 0.18139248 |
| 31,700 | 0.354588972 | 0.18139889 |
| 31,800 | 0.354275849 | 0.18133534 |
| 31,900 | 0.3540154 | 0.18135497 |
| 32,000 | 0.353864018 | 0.18139821 |
| 32,100 | 0.353699598 | 0.18138352 |
| 32,200 | 0.353524093 | 0.18138071 |
| 32,300 | 0.353553849 | 0.18145078 |
| 32,400 | 0.353374875 | 0.18134413 |

| | | |
|---|---|---|
| 32,500 | 0.353100244 | 0.18139278 |
| 32,600 | 0.352692653 | 0.18138034 |
| 32,700 | 0.352806692 | 0.18135401 |
| 32,800 | 0.352483037 | 0.18132406 |
| 32,900 | 0.352337741 | 0.18130031 |
| 33,000 | 0.352127811 | 0.18125106 |
| 33,100 | 0.351800126 | 0.18137085 |
| 33,200 | 0.351650977 | 0.18137018 |
| 33,300 | 0.351366138 | 0.18135454 |
| 33,400 | 0.351247182 | 0.1812839 |
| 33,500 | 0.351013939 | 0.18129813 |
| 33,600 | 0.350784181 | 0.18121475 |
| 33,700 | 0.350715375 | 0.18126569 |
| 33,800 | 0.350502638 | 0.18127943 |
| 33,900 | 0.350165639 | 0.18124405 |
| 34,000 | 0.350084757 | 0.18130982 |
| 34,100 | 0.349962567 | 0.18115257 |
| 34,200 | 0.349986739 | 0.1812265 |
| 34,300 | 0.34948506 | 0.18125201 |
| 34,400 | 0.349670603 | 0.18124267 |
| 34,500 | 0.349410965 | 0.18124616 |
| 34,600 | 0.349114264 | 0.18125331 |
| 34,700 | 0.34882982 | 0.18123689 |
| 34,800 | 0.348657111 | 0.18120703 |
| 34,900 | 0.348599938 | 0.18126329 |
| 35,000 | 0.348322788 | 0.18118335 |
| 35,100 | 0.348405304 | 0.18133404 |
| 35,200 | 0.348157277 | 0.18114223 |
| 35,300 | 0.348126098 | 0.18119583 |
| 35,400 | 0.347988802 | 0.18122216 |
| 35,500 | 0.347815222 | 0.18117046 |
| 35,600 | 0.347598139 | 0.18116657 |
| 35,700 | 0.347481065 | 0.18114647 |
| 35,800 | 0.347175273 | 0.18121475 |
| 35,900 | 0.346913336 | 0.18126312 |
| 36,000 | 0.346854494 | 0.18129191 |
| 36,100 | 0.346718374 | 0.18116101 |
| 36,200 | 0.346496281 | 0.18121124 |
| 36,300 | 0.346334952 | 0.18103174 |
| 36,400 | 0.346066343 | 0.18108946 |
| 36,500 | 0.346063873 | 0.18110312 |
| 36,600 | 0.345846928 | 0.18110933 |
| 36,700 | 0.345754567 | 0.18106368 |
| 36,800 | 0.345653028 | 0.18096742 |
| 36,900 | 0.345250751 | 0.18101509 |
| 37,000 | 0.345196983 | 0.18105576 |
| 37,100 | 0.344807603 | 0.18106381 |
| 37,200 | 0.344788578 | 0.18109867 |
| 37,300 | 0.344468896 | 0.18101769 |
| 37,400 | 0.344330915 | 0.18102404 |
| 37,500 | 0.344145764 | 0.1810248 |
| 37,600 | 0.344112339 | 0.18106075 |
| 37,700 | 0.343940101 | 0.18105328 |
| 37,800 | 0.343725996 | 0.18103945 |
| 37,900 | 0.343639252 | 0.18107979 |
| 38,000 | 0.343671487 | 0.18099094 |
| 38,100 | 0.343423812 | 0.18098954 |
| 38,200 | 0.343187921 | 0.18103717 |
| 38,300 | 0.343044259 | 0.18093096 |
| 38,400 | 0.342914557 | 0.18093271 |
| 38,500 | 0.342811475 | 0.18101235 |
| 38,600 | 0.342729537 | 0.18106423 |
| 38,700 | 0.342451715 | 0.18102309 |
| 38,800 | 0.342451767 | 0.18093741 |
| 38,900 | 0.34211656 | 0.1809011 |
| 39,000 | 0.341889661 | 0.18095288 |
| 39,100 | 0.341759874 | 0.18095966 |

| | | |
|---|---|---|
| 39,200 | 0.341520207 | 0.18093571 |
| 39,300 | 0.34156645 | 0.1809015 |
| 39,400 | 0.341069328 | 0.18091203 |
| 39,500 | 0.341089657 | 0.18092036 |
| 39,600 | 0.340950931 | 0.18089802 |
| 39,700 | 0.340653431 | 0.18082917 |
| 39,800 | 0.340732839 | 0.18089641 |
| 39,900 | 0.340623155 | 0.1807866 |
| 40,000 | 0.340240188 | 0.18094705 |
| 40,100 | 0.340102594 | 0.18095051 |
| 40,200 | 0.340179521 | 0.18091575 |
| 40,300 | 0.340023933 | 0.18085133 |
| 40,400 | 0.339839494 | 0.18091336 |
| 40,500 | 0.339810094 | 0.180891 |
| 40,600 | 0.339699732 | 0.18078927 |
| 40,700 | 0.339211875 | 0.18080788 |
| 40,800 | 0.339215707 | 0.18079845 |
| 40,900 | 0.339105321 | 0.18081348 |
| 41,000 | 0.338880292 | 0.18087446 |
| 41,100 | 0.338902302 | 0.18092926 |
| 41,200 | 0.338699269 | 0.18084448 |
| 41,300 | 0.338301381 | 0.18073759 |
| 41,400 | 0.338241747 | 0.18096081 |
| 41,500 | 0.338063215 | 0.18088534 |
| 41,600 | 0.337947534 | 0.18090226 |
| 41,700 | 0.337923876 | 0.18091082 |
| 41,800 | 0.337624862 | 0.18082742 |
| 41,900 | 0.337282488 | 0.18080615 |
| 42,000 | 0.337329402 | 0.18086613 |
| 42,100 | 0.337194995 | 0.18084472 |
| 42,200 | 0.336875659 | 0.18087723 |
| 42,300 | 0.336921897 | 0.18084323 |
| 42,400 | 0.336876911 | 0.18085086 |
| 42,500 | 0.336610893 | 0.18084108 |
| 42,600 | 0.336567907 | 0.18088341 |
| 42,700 | 0.336260981 | 0.18080377 |
| 42,800 | 0.336262197 | 0.18080298 |
| 42,900 | 0.336170698 | 0.1808258 |
| 43,000 | 0.336054982 | 0.18079933 |
| 43,100 | 0.335845222 | 0.18074427 |
| 43,200 | 0.335586387 | 0.18077984 |
| 43,300 | 0.33557135 | 0.1807617 |
| 43,400 | 0.335533342 | 0.18087907 |
| 43,500 | 0.335341207 | 0.18072035 |
| 43,600 | 0.335171194 | 0.18067071 |
| 43,700 | 0.335059499 | 0.18077057 |
| 43,800 | 0.334807583 | 0.18080616 |
| 43,900 | 0.334971623 | 0.18078614 |
| 44,000 | 0.334681379 | 0.18073799 |
| 44,100 | 0.334688883 | 0.18075409 |
| 44,200 | 0.33446411 | 0.18070419 |
| 44,300 | 0.334143419 | 0.18067669 |
| 44,400 | 0.334065165 | 0.18067587 |
| 44,500 | 0.333837193 | 0.18076108 |
| 44,600 | 0.333602921 | 0.18070018 |
| 44,700 | 0.333646011 | 0.18073319 |
| 44,800 | 0.333147517 | 0.18071798 |
| 44,900 | 0.333201279 | 0.18068079 |
| 45,000 | 0.3331987 | 0.1807905 |
| 45,100 | 0.332982071 | 0.18066023 |
| 45,200 | 0.332949402 | 0.18065592 |
| 45,300 | 0.332841842 | 0.18066579 |
| 45,400 | 0.332665622 | 0.18065201 |
| 45,500 | 0.332669225 | 0.18063866 |
| 45,600 | 0.332659486 | 0.18063521 |
| 45,700 | 0.332351173 | 0.18067349 |
| 45,800 | 0.332236372 | 0.18065579 |

| | | |
|---|---|---|
| 45,900 | 0.331984518 | 0.18066935 |
| 46,000 | 0.331888266 | 0.18070484 |
| 46,100 | 0.331702674 | 0.18062001 |
| 46,200 | 0.33153516 | 0.18055311 |
| 46,300 | 0.331315585 | 0.18056557 |
| 46,400 | 0.331282765 | 0.18074551 |
| 46,500 | 0.331028149 | 0.18062344 |
| 46,600 | 0.330986672 | 0.18060145 |
| 46,700 | 0.330766187 | 0.18059186 |
| 46,800 | 0.330728614 | 0.18057919 |
| 46,900 | 0.330422816 | 0.18070926 |
| 47,000 | 0.33027923 | 0.18062774 |
| 47,100 | 0.330253867 | 0.18059693 |
| 47,200 | 0.330275872 | 0.18065216 |
| 47,300 | 0.329973845 | 0.18070521 |
| 47,400 | 0.329697018 | 0.18057896 |
| 47,500 | 0.329785168 | 0.18062031 |
| 47,600 | 0.329583454 | 0.18064432 |
| 47,700 | 0.329603106 | 0.18062753 |
| 47,800 | 0.329316452 | 0.18063625 |
| 47,900 | 0.329255929 | 0.18062033 |
| 48,000 | 0.329173138 | 0.18060128 |
| 48,100 | 0.329108557 | 0.18058548 |
| 48,200 | 0.328539217 | 0.18060487 |
| 48,300 | 0.328537355 | 0.18063529 |
| 48,400 | 0.328627562 | 0.18065942 |
| 48,500 | 0.328335045 | 0.18056788 |
| 48,600 | 0.328257349 | 0.18063133 |
| 48,700 | 0.328243205 | 0.18048465 |
| 48,800 | 0.327976256 | 0.18061015 |
| 48,900 | 0.327883651 | 0.18052059 |
| 49,000 | 0.327864743 | 0.18055444 |
| 49,100 | 0.327832035 | 0.18055974 |
| 49,200 | 0.327409886 | 0.18066031 |
| 49,300 | 0.327436859 | 0.18056415 |
| 49,400 | 0.327353656 | 0.1805754 |
| 49,500 | 0.327182292 | 0.18057557 |
| 49,600 | 0.326922471 | 0.18048524 |
| 49,700 | 0.327044299 | 0.18058572 |
| 49,800 | 0.326609424 | 0.18054674 |
| 49,900 | 0.326331935 | 0.18054124 |
| 50,000 | 0.326369902 | 0.18059759 |
| 50,100 | 0.326412272 | 0.1804607 |
| 50,200 | 0.326111984 | 0.18055598 |
| 50,300 | 0.326265687 | 0.18056057 |
| 50,400 | 0.325929255 | 0.18048656 |
| 50,500 | 0.325806261 | 0.18054922 |
| 50,600 | 0.325696087 | 0.18056886 |
| 50,700 | 0.32549771 | 0.18055533 |
| 50,800 | 0.32561134 | 0.18051888 |
| 50,900 | 0.325362156 | 0.18052992 |
| 51,000 | 0.325304584 | 0.18056392 |
| 51,100 | 0.324917225 | 0.18051609 |
| 51,200 | 0.324635492 | 0.18046108 |
| 51,300 | 0.324870056 | 0.18054419 |
| 51,400 | 0.324449296 | 0.1804933 |
| 51,500 | 0.324415698 | 0.18048193 |
| 51,600 | 0.324282188 | 0.18045153 |
| 51,700 | 0.324340589 | 0.1804441 |
| 51,800 | 0.324230866 | 0.18058049 |
| 51,900 | 0.323873138 | 0.18053809 |
| 52,000 | 0.32386124 | 0.18051701 |
| 52,100 | 0.323840981 | 0.18048069 |
| 52,200 | 0.323820723 | 0.18046764 |
| 52,300 | 0.323491509 | 0.18054615 |
| 52,400 | 0.32335217 | 0.18060845 |
| 52,500 | 0.32323857 | 0.18056273 |

| | | |
|---|---|---|
| 52,600 | 0.323255126 | 0.18043742 |
| 52,700 | 0.322992192 | 0.18055084 |
| 52,800 | 0.323032625 | 0.18057373 |
| 52,900 | 0.322662864 | 0.18043536 |
| 53,000 | 0.322470632 | 0.18052444 |
| 53,100 | 0.322177442 | 0.18049366 |
| 53,200 | 0.322323698 | 0.1805717 |
| 53,300 | 0.322311252 | 0.18046188 |
| 53,400 | 0.321929733 | 0.18046201 |
| 53,500 | 0.321643387 | 0.18050853 |
| 53,600 | 0.321399573 | 0.18056332 |
| 53,700 | 0.321340302 | 0.18043948 |
| 53,800 | 0.321368915 | 0.18055882 |
| 53,900 | 0.321427909 | 0.18044998 |
| 54,000 | 0.32132802 | 0.18051563 |
| 54,100 | 0.321137255 | 0.18044524 |
| 54,200 | 0.321065099 | 0.18046033 |
| 54,300 | 0.321087434 | 0.18039925 |
| 54,400 | 0.320612729 | 0.18039727 |
| 54,500 | 0.320748981 | 0.18046229 |
| 54,600 | 0.320549823 | 0.18040012 |
| 54,700 | 0.320435359 | 0.1803292 |
| 54,800 | 0.320287335 | 0.18040028 |
| 54,900 | 0.320147284 | 0.1804201 |
| 55,000 | 0.319971592 | 0.18030654 |
| 55,100 | 0.319792504 | 0.18037813 |
| 55,200 | 0.319424978 | 0.18035291 |
| 55,300 | 0.319599809 | 0.18039112 |
| 55,400 | 0.319457283 | 0.18037657 |
| 55,500 | 0.319283477 | 0.18035574 |
| 55,600 | 0.318900388 | 0.1803047 |
| 55,700 | 0.318925998 | 0.18036032 |
| 55,800 | 0.319061203 | 0.18038156 |
| 55,900 | 0.319052689 | 0.18044268 |
| 56,000 | 0.318883371 | 0.18034102 |
| 56,100 | 0.318974054 | 0.18034688 |
| 56,200 | 0.318382192 | 0.18041034 |
| 56,300 | 0.318630205 | 0.18040253 |
| 56,400 | 0.318426933 | 0.18038938 |
| 56,500 | 0.318256939 | 0.18039182 |
| 56,600 | 0.318142949 | 0.18035477 |
| 56,700 | 0.31801387 | 0.18037281 |
| 56,800 | 0.317679569 | 0.18036377 |
| 56,900 | 0.317435594 | 0.18034941 |
| 57,000 | 0.317648348 | 0.18032271 |
| 57,100 | 0.317581286 | 0.18033287 |
| 57,200 | 0.317484389 | 0.18036297 |
| 57,300 | 0.317401669 | 0.18036106 |
| 57,400 | 0.317161462 | 0.1803683 |
| 57,500 | 0.316988095 | 0.18025487 |
| 57,600 | 0.316959302 | 0.18029513 |
| 57,700 | 0.316860036 | 0.18030852 |
| 57,800 | 0.316780031 | 0.1803683 |
| 57,900 | 0.316704403 | 0.18037262 |
| 58,000 | 0.316345246 | 0.18027571 |
| 58,100 | 0.31648447 | 0.18030656 |
| 58,200 | 0.316356493 | 0.18024924 |
| 58,300 | 0.316463403 | 0.18031739 |
| 58,400 | 0.316206715 | 0.18024362 |
| 58,500 | 0.31603468 | 0.18033617 |
| 58,600 | 0.316157939 | 0.18027142 |
| 58,700 | 0.315884267 | 0.18035531 |
| 58,800 | 0.31570344 | 0.1802795 |
| 58,900 | 0.315648453 | 0.18021093 |
| 59,000 | 0.315584419 | 0.18025051 |
| 59,100 | 0.315363895 | 0.18020535 |
| 59,200 | 0.315534913 | 0.18024185 |

| | | |
|---|---|---|
| 59,300 | 0.31536146 | 0.18019848 |
| 59,400 | 0.315323898 | 0.18018685 |
| 59,500 | 0.31517295 | 0.18018644 |
| 59,600 | 0.315026638 | 0.18019536 |
| 59,700 | 0.314701275 | 0.18015571 |
| 59,800 | 0.314705631 | 0.18024636 |
| 59,900 | 0.314584304 | 0.18019748 |
| 60,000 | 0.314516733 | 0.18025082 |
| 60,100 | 0.314413723 | 0.18016544 |
| 60,200 | 0.31433929 | 0.18023138 |
| 60,300 | 0.314179303 | 0.18021014 |
| 60,400 | 0.314240185 | 0.18013346 |
| 60,500 | 0.313966622 | 0.18018839 |
| 60,600 | 0.314131408 | 0.18019573 |
| 60,700 | 0.313944523 | 0.18026517 |
| 60,800 | 0.313971429 | 0.18017392 |
| 60,900 | 0.313821704 | 0.18022114 |
| 61,000 | 0.31376751 | 0.18021015 |
| 61,100 | 0.313528912 | 0.18025915 |
| 61,200 | 0.313298506 | 0.18021987 |
| 61,300 | 0.313416963 | 0.1802078 |
| 61,400 | 0.313423379 | 0.1801675 |
| 61,500 | 0.313192643 | 0.1801649 |
| 61,600 | 0.312930797 | 0.1802073 |
| 61,700 | 0.313207934 | 0.18021202 |
| 61,800 | 0.312900285 | 0.18020522 |
| 61,900 | 0.312985224 | 0.18016032 |
| 62,000 | 0.312928266 | 0.18014959 |
| 62,100 | 0.312763226 | 0.18028406 |
| 62,200 | 0.312757603 | 0.18016131 |
| 62,300 | 0.312488995 | 0.18017995 |
| 62,400 | 0.312540178 | 0.18017701 |
| 62,500 | 0.312512722 | 0.18015752 |
| 62,600 | 0.312507358 | 0.18013232 |
| 62,700 | 0.312320994 | 0.18029812 |
| 62,800 | 0.312085989 | 0.18016912 |
| 62,900 | 0.312133239 | 0.18014108 |
| 63,000 | 0.312004867 | 0.18014295 |
| 63,100 | 0.311780353 | 0.18030862 |
| 63,200 | 0.311518141 | 0.18019322 |
| 63,300 | 0.31153313 | 0.1801475 |
| 63,400 | 0.311419837 | 0.18007743 |
| 63,500 | 0.311483358 | 0.18020456 |
| 63,600 | 0.311380108 | 0.18017575 |
| 63,700 | 0.311298137 | 0.1802018 |
| 63,800 | 0.311270525 | 0.18019879 |
| 63,900 | 0.311210511 | 0.18017404 |
| 64,000 | 0.310642501 | 0.18021961 |
| 64,100 | 0.310890919 | 0.18022739 |
| 64,200 | 0.310784548 | 0.18019073 |
| 64,300 | 0.310514303 | 0.18020003 |
| 64,400 | 0.310588656 | 0.18010172 |
| 64,500 | 0.310439602 | 0.18021782 |
| 64,600 | 0.310219201 | 0.18008656 |
| 64,700 | 0.310245549 | 0.18017597 |
| 64,800 | 0.310096681 | 0.18008433 |
| 64,900 | 0.310113419 | 0.18011501 |
| 65,000 | 0.309792868 | 0.18014289 |
| 65,100 | 0.309886435 | 0.18008049 |
| 65,200 | 0.309732026 | 0.17999488 |
| 65,300 | 0.309691103 | 0.18007281 |
| 65,400 | 0.309575875 | 0.18006236 |
| 65,500 | 0.309536861 | 0.18008157 |
| 65,600 | 0.30951105 | 0.18016838 |
| 65,700 | 0.309419252 | 0.18018103 |
| 65,800 | 0.3091667 | 0.18011239 |
| 65,900 | 0.309037475 | 0.18012173 |

| | | |
|---|---|---|
| 66,000 | 0.309028416 | 0.18007226 |
| 66,100 | 0.308906612 | 0.18013542 |
| 66,200 | 0.308907107 | 0.18014666 |
| 66,300 | 0.308767021 | 0.18015501 |
| 66,400 | 0.308688785 | 0.1801499 |
| 66,500 | 0.308543839 | 0.18009976 |
| 66,600 | 0.308623798 | 0.18012808 |
| 66,700 | 0.308333338 | 0.18009224 |
| 66,800 | 0.308354783 | 0.18004996 |
| 66,900 | 0.308103609 | 0.18009873 |
| 67,000 | 0.308181083 | 0.18012871 |
| 67,100 | 0.308072197 | 0.18008633 |
| 67,200 | 0.307939798 | 0.18010988 |
| 67,300 | 0.307770598 | 0.1800559 |
| 67,400 | 0.307574526 | 0.18006926 |
| 67,500 | 0.307334883 | 0.1800679 |
| 67,600 | 0.307112508 | 0.18003759 |
| 67,700 | 0.30724054 | 0.18005715 |
| 67,800 | 0.307158977 | 0.18005264 |
| 67,900 | 0.307177754 | 0.18004032 |
| 68,000 | 0.307127081 | 0.17999137 |
| 68,100 | 0.307073258 | 0.1800714 |
| 68,200 | 0.307013112 | 0.18008899 |
| 68,300 | 0.306884832 | 0.18010612 |
| 68,400 | 0.30685804 | 0.18002204 |
| 68,500 | 0.306962791 | 0.17999558 |
| 68,600 | 0.306634241 | 0.18002622 |
| 68,700 | 0.306448838 | 0.18005601 |
| 68,800 | 0.306364773 | 0.17997041 |
| 68,900 | 0.306342864 | 0.18002879 |
| 69,000 | 0.306264293 | 0.18005141 |
| 69,100 | 0.306140394 | 0.18001099 |
| 69,200 | 0.306293682 | 0.18012126 |
| 69,300 | 0.306166518 | 0.18000583 |
| 69,400 | 0.305878836 | 0.1799402 |
| 69,500 | 0.30595971 | 0.18002024 |
| 69,600 | 0.305809997 | 0.17999103 |
| 69,700 | 0.305586035 | 0.17998675 |
| 69,800 | 0.305585442 | 0.17991685 |
| 69,900 | 0.305272611 | 0.1799691 |
| 70,000 | 0.305251595 | 0.17996083 |
| 70,100 | 0.305188717 | 0.1799996 |
| 70,200 | 0.305055197 | 0.17989644 |
| 70,300 | 0.304876782 | 0.17994581 |
| 70,400 | 0.30482286 | 0.17995842 |
| 70,500 | 0.304580693 | 0.18003411 |
| 70,600 | 0.304750527 | 0.18003897 |
| 70,700 | 0.304380445 | 0.17994609 |
| 70,800 | 0.304317067 | 0.17993522 |
| 70,900 | 0.304173256 | 0.18001681 |
| 71,000 | 0.304192386 | 0.1799204 |
| 71,100 | 0.304119943 | 0.17992062 |
| 71,200 | 0.304126201 | 0.17989398 |
| 71,300 | 0.304023155 | 0.17997002 |
| 71,400 | 0.303936743 | 0.1798807 |
| 71,500 | 0.303798431 | 0.1799553 |
| 71,600 | 0.303572044 | 0.17997727 |
| 71,700 | 0.303392166 | 0.17994372 |
| 71,800 | 0.303473832 | 0.18001666 |
| 71,900 | 0.303530524 | 0.17995058 |
| 72,000 | 0.303281937 | 0.17998147 |
| 72,100 | 0.303333347 | 0.17988483 |
| 72,200 | 0.3032139 | 0.18000234 |
| 72,300 | 0.303056099 | 0.17994065 |
| 72,400 | 0.303035015 | 0.17990755 |
| 72,500 | 0.302927928 | 0.18006363 |
| 72,600 | 0.302842039 | 0.17986585 |

| | | |
|---|---|---|
| 72,700 | 0.302729847 | 0.18001118 |
| 72,800 | 0.302699349 | 0.18003544 |
| 72,900 | 0.302318078 | 0.1799984 |
| 73,000 | 0.302318696 | 0.17991005 |
| 73,100 | 0.302303634 | 0.17994704 |
| 73,200 | 0.302033749 | 0.17989719 |
| 73,300 | 0.30226604 | 0.17993997 |
| 73,400 | 0.302095929 | 0.18004026 |
| 73,500 | 0.301950977 | 0.18001951 |
| 73,600 | 0.301825332 | 0.17998538 |
| 73,700 | 0.301733786 | 0.18003684 |
| 73,800 | 0.30155923 | 0.18006339 |
| 73,900 | 0.301554049 | 0.17997866 |
| 74,000 | 0.301453066 | 0.18003231 |
| 74,100 | 0.301146853 | 0.17998682 |
| 74,200 | 0.301317758 | 0.18004501 |
| 74,300 | 0.301097154 | 0.18000034 |
| 74,400 | 0.300825502 | 0.17998262 |
| 74,500 | 0.300527684 | 0.17992314 |
| 74,600 | 0.300972997 | 0.17993376 |
| 74,700 | 0.300916908 | 0.18000107 |
| 74,800 | 0.300822193 | 0.17993422 |
| 74,900 | 0.300742712 | 0.17989771 |
| 75,000 | 0.300711349 | 0.17986712 |
| 75,100 | 0.30042767 | 0.1799294 |
| 75,200 | 0.300291411 | 0.17985461 |
| 75,300 | 0.300244508 | 0.17996878 |
| 75,400 | 0.300299961 | 0.1798907 |
| 75,500 | 0.300405198 | 0.1798972 |
| 75,600 | 0.300091301 | 0.17992197 |
| 75,700 | 0.300162941 | 0.17993615 |
| 75,800 | 0.299775385 | 0.17999702 |
| 75,900 | 0.299887372 | 0.17993659 |
| 76,000 | 0.299880438 | 0.18002076 |
| 76,100 | 0.299875267 | 0.1799422 |
| 76,200 | 0.299682658 | 0.17990089 |
| 76,300 | 0.299750424 | 0.17995442 |
| 76,400 | 0.299554939 | 0.1799323 |
| 76,500 | 0.299384166 | 0.17988555 |
| 76,600 | 0.299384172 | 0.1798818 |
| 76,700 | 0.299468377 | 0.17986363 |
| 76,800 | 0.299313166 | 0.17988093 |
| 76,900 | 0.299187398 | 0.17994954 |
| 77,000 | 0.29910958 | 0.17989471 |
| 77,100 | 0.298886251 | 0.17990191 |
| 77,200 | 0.29876439 | 0.17974505 |
| 77,300 | 0.298579466 | 0.17993163 |
| 77,400 | 0.298635121 | 0.179814 |
| 77,500 | 0.298634406 | 0.17983207 |
| 77,600 | 0.298723771 | 0.17977284 |
| 77,700 | 0.29838209 | 0.17982529 |
| 77,800 | 0.298327187 | 0.17980606 |
| 77,900 | 0.29817262 | 0.17986889 |
| 78,000 | 0.298116836 | 0.17976193 |
| 78,100 | 0.298041867 | 0.17987643 |
| 78,200 | 0.298006133 | 0.17987734 |
| 78,300 | 0.297707374 | 0.17981681 |
| 78,400 | 0.297461114 | 0.17991146 |
| 78,500 | 0.297478991 | 0.1798553 |
| 78,600 | 0.297522564 | 0.17987438 |
| 78,700 | 0.297396229 | 0.179795 |
| 78,800 | 0.297359088 | 0.17990015 |
| 78,900 | 0.297272794 | 0.17990086 |
| 79,000 | 0.297061929 | 0.17982281 |
| 79,100 | 0.297039884 | 0.17980441 |
| 79,200 | 0.296901488 | 0.17976879 |
| 79,300 | 0.29670262 | 0.17976356 |

| | | |
|---|---|---|
| 79,400 | 0.29681836 | 0.17985642 |
| 79,500 | 0.296709365 | 0.17973476 |
| 79,600 | 0.296677637 | 0.17973426 |
| 79,700 | 0.296468458 | 0.17987358 |
| 79,800 | 0.296297223 | 0.17982365 |
| 79,900 | 0.296414299 | 0.17986803 |
| 80,000 | 0.296161573 | 0.17984601 |
| 80,100 | 0.296169844 | 0.17987679 |
| 80,200 | 0.296112311 | 0.17991023 |
| 80,300 | 0.296124668 | 0.17979935 |
| 80,400 | 0.296108914 | 0.17981835 |
| 80,500 | 0.295837581 | 0.17975637 |
| 80,600 | 0.295868449 | 0.17989046 |
| 80,700 | 0.295972834 | 0.17986082 |
| 80,800 | 0.295829325 | 0.1798256 |
| 80,900 | 0.295689238 | 0.17975338 |
| 81,000 | 0.295657036 | 0.17983304 |
| 81,100 | 0.295631507 | 0.17977337 |
| 81,200 | 0.295595345 | 0.17983812 |
| 81,300 | 0.295333775 | 0.17986236 |
| 81,400 | 0.295182013 | 0.17981087 |
| 81,500 | 0.295120923 | 0.17980229 |
| 81,600 | 0.295123571 | 0.17983498 |
| 81,700 | 0.295172396 | 0.17982184 |
| 81,800 | 0.295114083 | 0.17968798 |
| 81,900 | 0.295074068 | 0.17980284 |
| 82,000 | 0.295040548 | 0.17981326 |
| 82,100 | 0.2949155 | 0.17982355 |
| 82,200 | 0.294849217 | 0.17979402 |
| 82,300 | 0.294477813 | 0.17983103 |
| 82,400 | 0.294676901 | 0.17985318 |
| 82,500 | 0.294326359 | 0.17977107 |
| 82,600 | 0.294275811 | 0.1797833 |
| 82,700 | 0.294255686 | 0.17982515 |
| 82,800 | 0.294471625 | 0.1797961 |
| 82,900 | 0.293950777 | 0.17983244 |
| 83,000 | 0.293935068 | 0.17979862 |
| 83,100 | 0.293834374 | 0.17978954 |
| 83,200 | 0.293833374 | 0.17978493 |
| 83,300 | 0.293833289 | 0.17977811 |
| 83,400 | 0.29355535 | 0.17984079 |
| 83,500 | 0.293482614 | 0.17978655 |
| 83,600 | 0.293195898 | 0.17982193 |
| 83,700 | 0.293476637 | 0.17978796 |
| 83,800 | 0.293159546 | 0.17980449 |
| 83,900 | 0.293018291 | 0.17979386 |
| 84,000 | 0.293274841 | 0.17979363 |
| 84,100 | 0.293125597 | 0.1797251 |
| 84,200 | 0.293034478 | 0.17982076 |
| 84,300 | 0.293056184 | 0.17979508 |
| 84,400 | 0.292890275 | 0.17985871 |
| 84,500 | 0.292774066 | 0.179749 |
| 84,600 | 0.292708899 | 0.17977433 |
| 84,700 | 0.292522867 | 0.17972215 |
| 84,800 | 0.292578642 | 0.17984625 |
| 84,900 | 0.292376569 | 0.17977241 |
| 85,000 | 0.292415089 | 0.17981959 |
| 85,100 | 0.292269293 | 0.17990661 |
| 85,200 | 0.292261103 | 0.17981625 |
| 85,300 | 0.292296147 | 0.17983291 |
| 85,400 | 0.292147336 | 0.1798931 |
| 85,500 | 0.292146345 | 0.1798082 |
| 85,600 | 0.291954895 | 0.1798114 |
| 85,700 | 0.291986022 | 0.17982448 |
| 85,800 | 0.291764981 | 0.1798006 |
| 85,900 | 0.291856291 | 0.17976528 |
| 86,000 | 0.291748611 | 0.17981234 |

| | | |
|---|---|---|
| 86,100 | 0.291708963 | 0.17985137 |
| 86,200 | 0.291559243 | 0.17987924 |
| 86,300 | 0.291342535 | 0.17995003 |
| 86,400 | 0.291517307 | 0.17986457 |
| 86,500 | 0.291462651 | 0.17987128 |
| 86,600 | 0.291372438 | 0.17987706 |
| 86,700 | 0.291208928 | 0.1798313 |
| 86,800 | 0.291119295 | 0.17976323 |
| 86,900 | 0.291101021 | 0.17982035 |
| 87,000 | 0.290923528 | 0.17987638 |
| 87,100 | 0.290777444 | 0.17979572 |
| 87,200 | 0.290464656 | 0.17980792 |
| 87,300 | 0.290645679 | 0.17981719 |
| 87,400 | 0.290586638 | 0.17976607 |
| 87,500 | 0.290501609 | 0.17978455 |
| 87,600 | 0.29064319 | 0.1797584 |
| 87,700 | 0.290418198 | 0.17980544 |
| 87,800 | 0.290463887 | 0.17984329 |
| 87,900 | 0.290498996 | 0.17985688 |
| 88,000 | 0.290485689 | 0.17987001 |
| 88,100 | 0.290200695 | 0.17984187 |
| 88,200 | 0.290101707 | 0.17985458 |
| 88,300 | 0.290077342 | 0.17990776 |
| 88,400 | 0.290129872 | 0.17983927 |
| 88,500 | 0.289895278 | 0.17982189 |
| 88,600 | 0.289852745 | 0.17980204 |
| 88,700 | 0.289928101 | 0.17986911 |
| 88,800 | 0.289715032 | 0.17979382 |
| 88,900 | 0.289744523 | 0.17981264 |
| 89,000 | 0.289476294 | 0.17976735 |
| 89,100 | 0.289582948 | 0.17971721 |
| 89,200 | 0.289522687 | 0.17975337 |
| 89,300 | 0.289326337 | 0.17976406 |
| 89,400 | 0.289116847 | 0.17975119 |
| 89,500 | 0.289092097 | 0.17977654 |
| 89,600 | 0.289027527 | 0.17976913 |
| 89,700 | 0.288836541 | 0.17972619 |
| 89,800 | 0.289136958 | 0.1797602 |
| 89,900 | 0.288717533 | 0.17981614 |
| 90,000 | 0.288754242 | 0.17981311 |
| 90,100 | 0.28870679 | 0.17978619 |
| 90,200 | 0.288988766 | 0.17983564 |
| 90,300 | 0.28866991 | 0.1797323 |
| 90,400 | 0.288622532 | 0.17971371 |
| 90,500 | 0.288737061 | 0.17982329 |
| 90,600 | 0.288535377 | 0.1797847 |
| 90,700 | 0.288401605 | 0.17982744 |
| 90,800 | 0.288539866 | 0.17979977 |
| 90,900 | 0.28843614 | 0.17973915 |
| 91,000 | 0.288279626 | 0.17968766 |
| 91,100 | 0.288172658 | 0.17974568 |
| 91,200 | 0.288110443 | 0.17982331 |
| 91,300 | 0.288077015 | 0.17971989 |
| 91,400 | 0.288129959 | 0.17985703 |
| 91,500 | 0.287912327 | 0.17978246 |
| 91,600 | 0.288069184 | 0.17977217 |
| 91,700 | 0.287962061 | 0.17970599 |
| 91,800 | 0.287811557 | 0.17974326 |
| 91,900 | 0.287874867 | 0.1797853 |
| 92,000 | 0.28769335 | 0.17974958 |
| 92,100 | 0.287696196 | 0.17974632 |
| 92,200 | 0.287458644 | 0.17977488 |
| 92,300 | 0.287552611 | 0.17975348 |
| 92,400 | 0.287315428 | 0.17971819 |
| 92,500 | 0.287237837 | 0.17975249 |
| 92,600 | 0.287139519 | 0.17973698 |
| 92,700 | 0.28696136 | 0.17967968 |

| | | |
|---|---|---|
| 92,800 | 0.286805646 | 0.17967048 |
| 92,900 | 0.286637155 | 0.17961449 |
| 93,000 | 0.286837867 | 0.17965359 |
| 93,100 | 0.286799961 | 0.17976124 |
| 93,200 | 0.286711469 | 0.17962754 |
| 93,300 | 0.286558134 | 0.17980232 |
| 93,400 | 0.286569369 | 0.17971879 |
| 93,500 | 0.286777833 | 0.17970663 |
| 93,600 | 0.28665541 | 0.17962817 |
| 93,700 | 0.286260348 | 0.17962195 |
| 93,800 | 0.286144416 | 0.17959168 |
| 93,900 | 0.286232045 | 0.17964661 |
| 94,000 | 0.286162957 | 0.1797067 |
| 94,100 | 0.286022211 | 0.17961528 |
| 94,200 | 0.285965715 | 0.17972844 |
| 94,300 | 0.285881742 | 0.17963406 |
| 94,400 | 0.285972184 | 0.17969158 |
| 94,500 | 0.285739709 | 0.17969717 |
| 94,600 | 0.285720746 | 0.17965854 |
| 94,700 | 0.285900515 | 0.17973801 |
| 94,800 | 0.285767311 | 0.17960721 |
| 94,900 | 0.285652695 | 0.17960638 |
| 95,000 | 0.285765538 | 0.17967919 |
| 95,100 | 0.285638539 | 0.17972283 |
| 95,200 | 0.285537053 | 0.17969726 |
| 95,300 | 0.285492005 | 0.17963039 |
| 95,400 | 0.285358179 | 0.17970519 |
| 95,500 | 0.285381498 | 0.17971891 |
| 95,600 | 0.285252315 | 0.17970942 |
| 95,700 | 0.285330224 | 0.17968614 |
| 95,800 | 0.285040547 | 0.17964657 |
| 95,900 | 0.285163943 | 0.17964002 |
| 96,000 | 0.284983536 | 0.17964043 |
| 96,100 | 0.284928567 | 0.1797394 |
| 96,200 | 0.284955373 | 0.17968253 |
| 96,300 | 0.284852736 | 0.17966217 |
| 96,400 | 0.284969594 | 0.17974559 |
| 96,500 | 0.284849894 | 0.17970105 |
| 96,600 | 0.284784623 | 0.17959851 |
| 96,700 | 0.284648378 | 0.17970654 |
| 96,800 | 0.284511835 | 0.17968869 |
| 96,900 | 0.28445576 | 0.1795925 |
| 97,000 | 0.284377167 | 0.17972871 |
| 97,100 | 0.284375792 | 0.17971774 |
| 97,200 | 0.28439223 | 0.17968254 |
| 97,300 | 0.284183762 | 0.17965775 |
| 97,400 | 0.284111324 | 0.17966688 |
| 97,500 | 0.284041422 | 0.17968852 |
| 97,600 | 0.283996283 | 0.17963526 |
| 97,700 | 0.283881815 | 0.17973874 |
| 97,800 | 0.283802595 | 0.17972603 |
| 97,900 | 0.28376538 | 0.17975178 |
| 98,000 | 0.283929063 | 0.17962703 |
| 98,100 | 0.283609243 | 0.17970966 |
| 98,200 | 0.283552505 | 0.17963671 |
| 98,300 | 0.283525665 | 0.17960645 |
| 98,400 | 0.283574097 | 0.17966109 |
| 98,500 | 0.283470091 | 0.17968621 |
| 98,600 | 0.283363696 | 0.1796562 |
| 98,700 | 0.283390988 | 0.17969037 |
| 98,800 | 0.283259952 | 0.17967662 |
| 98,900 | 0.283353557 | 0.17962526 |
| 99,000 | 0.283143424 | 0.17971444 |
| 99,100 | 0.28291781 | 0.17963093 |
| 99,200 | 0.282846264 | 0.1796556 |
| 99,300 | 0.283046052 | 0.17967371 |
| 99,400 | 0.282815052 | 0.17975811 |

| | | |
|---|---|---|
| 99,500 | 0.282837652 | 0.17966857 |
| 99,600 | 0.282936696 | 0.17973282 |
| 99,700 | 0.282810571 | 0.17963454 |
| 99,800 | 0.282548247 | 0.17965178 |
| 99,900 | 0.282480436 | 0.17959628 |
| 100,000 | 0.282621877 | 0.17956779 |
| 100,100 | 0.28235414 | 0.17958516 |
| 100,200 | 0.282435436 | 0.17967165 |
| 100,300 | 0.282299084 | 0.17969378 |
| 100,400 | 0.282264307 | 0.17971175 |
| 100,500 | 0.282228958 | 0.17971853 |
| 100,600 | 0.282262897 | 0.17968403 |
| 100,700 | 0.282230913 | 0.17964063 |
| 100,800 | 0.282005344 | 0.17955154 |
| 100,900 | 0.282150207 | 0.17973605 |
| 101,000 | 0.281872569 | 0.17970377 |
| 101,100 | 0.281850845 | 0.17966211 |
| 101,200 | 0.281613103 | 0.17962083 |
| 101,300 | 0.281808914 | 0.17963475 |
| 101,400 | 0.281665786 | 0.17962762 |
| 101,500 | 0.281682823 | 0.17971767 |
| 101,600 | 0.2815039 | 0.17963293 |
| 101,700 | 0.28117595 | 0.1796007 |
| 101,800 | 0.281325476 | 0.17956465 |
| 101,900 | 0.280903067 | 0.17956099 |
| 102,000 | 0.281157869 | 0.17963528 |
| 102,100 | 0.281211859 | 0.17964921 |
| 102,200 | 0.280882924 | 0.17962693 |
| 102,300 | 0.280703275 | 0.17959963 |
| 102,400 | 0.28076827 | 0.17959037 |
| 102,500 | 0.280670144 | 0.17961085 |
| 102,600 | 0.280601209 | 0.17957488 |
| 102,700 | 0.280718058 | 0.17967202 |
| 102,800 | 0.280505121 | 0.179575 |
| 102,900 | 0.28055499 | 0.17972953 |
| 103,000 | 0.280663316 | 0.17961722 |
| 103,100 | 0.280375111 | 0.17960226 |
| 103,200 | 0.280254982 | 0.17975951 |
| 103,300 | 0.280381685 | 0.17967208 |
| 103,400 | 0.280552241 | 0.17969994 |
| 103,500 | 0.280186571 | 0.17960045 |
| 103,600 | 0.280159824 | 0.17963961 |
| 103,700 | 0.280062983 | 0.17956729 |
| 103,800 | 0.280055474 | 0.17958327 |
| 103,900 | 0.279980133 | 0.17962694 |
| 104,000 | 0.28001224 | 0.17952145 |
| 104,100 | 0.279834534 | 0.17958939 |
| 104,200 | 0.279764809 | 0.1795758 |
| 104,300 | 0.279752414 | 0.17969273 |
| 104,400 | 0.279822827 | 0.17958045 |
| 104,500 | 0.279892589 | 0.1795468 |
| 104,600 | 0.279424982 | 0.17947316 |
| 104,700 | 0.279441129 | 0.17953111 |
| 104,800 | 0.279505719 | 0.17958747 |
| 104,900 | 0.279378118 | 0.17963852 |
| 105,000 | 0.279332067 | 0.17958791 |
| 105,100 | 0.279215774 | 0.17955408 |
| 105,200 | 0.279180147 | 0.17963155 |
| 105,300 | 0.279244269 | 0.17960359 |
| 105,400 | 0.279135969 | 0.17958794 |
| 105,500 | 0.279204393 | 0.17954214 |
| 105,600 | 0.2790522 | 0.17955592 |
| 105,700 | 0.278993244 | 0.17962824 |
| 105,800 | 0.278793885 | 0.17951418 |
| 105,900 | 0.278851672 | 0.17958723 |
| 106,000 | 0.278803796 | 0.17950305 |
| 106,100 | 0.278614955 | 0.17954964 |

| | | |
|---|---|---|
| 106,200 | 0.278779191 | 0.17961904 |
| 106,300 | 0.278583819 | 0.17948078 |
| 106,400 | 0.278563551 | 0.17949658 |
| 106,500 | 0.278455142 | 0.1794225 |
| 106,600 | 0.278530174 | 0.17946927 |
| 106,700 | 0.278353481 | 0.17952428 |
| 106,800 | 0.278356734 | 0.17948905 |
| 106,900 | 0.278393301 | 0.17966961 |
| 107,000 | 0.278279358 | 0.17962226 |
| 107,100 | 0.278236358 | 0.17956017 |
| 107,200 | 0.278303775 | 0.17959624 |
| 107,300 | 0.278199708 | 0.17955399 |
| 107,400 | 0.278103613 | 0.17956684 |
| 107,500 | 0.277988791 | 0.17952278 |
| 107,600 | 0.277887827 | 0.17958101 |
| 107,700 | 0.27781401 | 0.17959131 |
| 107,800 | 0.277532761 | 0.17960064 |
| 107,900 | 0.277618809 | 0.1795836 |
| 108,000 | 0.277641308 | 0.17952775 |
| 108,100 | 0.277713859 | 0.17948358 |
| 108,200 | 0.277737768 | 0.17956042 |
| 108,300 | 0.277367391 | 0.17955717 |
| 108,400 | 0.277697241 | 0.17960357 |
| 108,500 | 0.277576331 | 0.1795621 |
| 108,600 | 0.277387709 | 0.17955103 |
| 108,700 | 0.277278546 | 0.17967532 |
| 108,800 | 0.277436108 | 0.17957762 |
| 108,900 | 0.277270673 | 0.17943289 |
| 109,000 | 0.276991756 | 0.1794993 |
| 109,100 | 0.276863126 | 0.17953298 |
| 109,200 | 0.276706783 | 0.17954733 |
| 109,300 | 0.276752498 | 0.17959229 |
| 109,400 | 0.276700439 | 0.17955271 |
| 109,500 | 0.276676546 | 0.17948628 |
| 109,600 | 0.276435998 | 0.17941498 |
| 109,700 | 0.276619422 | 0.17947596 |
| 109,800 | 0.276548522 | 0.17951707 |
| 109,900 | 0.276551685 | 0.17951765 |
| 110,000 | 0.276530614 | 0.17951708 |
| 110,100 | 0.27642781 | 0.17948457 |
| 110,200 | 0.276258717 | 0.17958327 |
| 110,300 | 0.276534989 | 0.17948325 |
| 110,400 | 0.27620323 | 0.17947287 |
| 110,500 | 0.276093822 | 0.17947619 |
| 110,600 | 0.276049227 | 0.17942165 |
| 110,700 | 0.276124939 | 0.17947261 |
| 110,800 | 0.275948331 | 0.17959676 |
| 110,900 | 0.275828734 | 0.17953719 |
| 111,000 | 0.275649094 | 0.17944068 |
| 111,100 | 0.275825778 | 0.17946246 |
| 111,200 | 0.275646396 | 0.17952992 |
| 111,300 | 0.275691491 | 0.17948525 |
| 111,400 | 0.275662237 | 0.17939482 |
| 111,500 | 0.275350588 | 0.1795775 |
| 111,600 | 0.275431182 | 0.17953076 |
| 111,700 | 0.275218912 | 0.17947779 |
| 111,800 | 0.275352527 | 0.17950038 |
| 111,900 | 0.275219718 | 0.17957955 |
| 112,000 | 0.275366462 | 0.17951061 |
| 112,100 | 0.275226619 | 0.17951972 |
| 112,200 | 0.274974683 | 0.1794205 |
| 112,300 | 0.274994485 | 0.17946219 |
| 112,400 | 0.275053248 | 0.17943206 |
| 112,500 | 0.275026543 | 0.17954688 |
| 112,600 | 0.275096829 | 0.17945184 |
| 112,700 | 0.274794549 | 0.17948804 |
| 112,800 | 0.274756696 | 0.17943028 |

| | | |
|---|---|---|
| 112,900 | 0.274819422 | 0.17951408 |
| 113,000 | 0.274693339 | 0.17956261 |
| 113,100 | 0.274650751 | 0.17955582 |
| 113,200 | 0.274674999 | 0.17945413 |
| 113,300 | 0.274458473 | 0.17951222 |
| 113,400 | 0.274378327 | 0.17948516 |
| 113,500 | 0.274336808 | 0.17951439 |
| 113,600 | 0.274221801 | 0.17963089 |
| 113,700 | 0.274268171 | 0.17952879 |
| 113,800 | 0.274337631 | 0.17967358 |
| 113,900 | 0.274230107 | 0.17960278 |
| 114,000 | 0.27406054 | 0.17956952 |
| 114,100 | 0.274007649 | 0.17952104 |
| 114,200 | 0.273978402 | 0.17947757 |
| 114,300 | 0.273823762 | 0.17949084 |
| 114,400 | 0.273859518 | 0.17954034 |
| 114,500 | 0.273719682 | 0.17954166 |
| 114,600 | 0.273741351 | 0.17952833 |
| 114,700 | 0.273849016 | 0.17958171 |
| 114,800 | 0.273893427 | 0.17953506 |
| 114,900 | 0.273817114 | 0.17954094 |
| 115,000 | 0.273682003 | 0.17954961 |
| 115,100 | 0.273805873 | 0.17943737 |
| 115,200 | 0.273796977 | 0.17940372 |
| 115,300 | 0.273632898 | 0.1795104 |
| 115,400 | 0.273654882 | 0.17944357 |
| 115,500 | 0.273554895 | 0.17952238 |
| 115,600 | 0.273492521 | 0.1794593 |
| 115,700 | 0.273609423 | 0.17955682 |
| 115,800 | 0.273410617 | 0.17957104 |
| 115,900 | 0.27310636 | 0.17944831 |
| 116,000 | 0.273197867 | 0.17950729 |
| 116,100 | 0.273133364 | 0.17949282 |
| 116,200 | 0.272933202 | 0.17957918 |
| 116,300 | 0.272818827 | 0.17940853 |
| 116,400 | 0.272967817 | 0.17944453 |
| 116,500 | 0.273003073 | 0.17944869 |
| 116,600 | 0.273014096 | 0.17943507 |
| 116,700 | 0.27287112 | 0.17955261 |
| 116,800 | 0.272463115 | 0.17955724 |
| 116,900 | 0.272647205 | 0.17943863 |
| 117,000 | 0.272430784 | 0.17952001 |
| 117,100 | 0.272593271 | 0.17943821 |
| 117,200 | 0.27243774 | 0.17941877 |
| 117,300 | 0.272351004 | 0.17952106 |
| 117,400 | 0.27219861 | 0.17943422 |
| 117,500 | 0.272192321 | 0.17941519 |
| 117,600 | 0.272403149 | 0.17962735 |
| 117,700 | 0.272381052 | 0.17947356 |
| 117,800 | 0.272316559 | 0.17935343 |
| 117,900 | 0.272287754 | 0.17941741 |
| 118,000 | 0.272270505 | 0.17954437 |
| 118,100 | 0.272115729 | 0.17960923 |
| 118,200 | 0.271810829 | 0.17943155 |
| 118,300 | 0.272026513 | 0.17950171 |
| 118,400 | 0.272061167 | 0.17941734 |
| 118,500 | 0.272092553 | 0.1795057 |
| 118,600 | 0.271859673 | 0.17944745 |
| 118,700 | 0.271846093 | 0.17953542 |
| 118,800 | 0.27154843 | 0.17942441 |
| 118,900 | 0.271664437 | 0.17942021 |
| 119,000 | 0.271558937 | 0.17943358 |
| 119,100 | 0.271352049 | 0.17941775 |
| 119,200 | 0.271329646 | 0.17940926 |
| 119,300 | 0.271374725 | 0.17938842 |
| 119,400 | 0.271406359 | 0.17945542 |
| 119,500 | 0.271218901 | 0.17940486 |

| | | |
|---|---|---|
| 119,600 | 0.271246499 | 0.17944946 |
| 119,700 | 0.271250025 | 0.17939912 |
| 119,800 | 0.271246111 | 0.17941481 |
| 119,900 | 0.271032624 | 0.17949573 |
| 120,000 | 0.270842398 | 0.17950135 |
| 120,100 | 0.27098227 | 0.17947166 |
| 120,200 | 0.271019186 | 0.17944676 |
| 120,300 | 0.270967827 | 0.17943072 |
| 120,400 | 0.270795519 | 0.17937858 |
| 120,500 | 0.27069718 | 0.17953888 |
| 120,600 | 0.270836799 | 0.17944228 |
| 120,700 | 0.270596543 | 0.17940671 |
| 120,800 | 0.270581249 | 0.17940356 |
| 120,900 | 0.270632218 | 0.17932603 |
| 121,000 | 0.270609617 | 0.17941877 |
| 121,100 | 0.270820617 | 0.17932871 |
| 121,200 | 0.270392787 | 0.17945197 |
| 121,300 | 0.270336653 | 0.17940842 |
| 121,400 | 0.270325313 | 0.17947038 |
| 121,500 | 0.270355402 | 0.17947578 |
| 121,600 | 0.270169468 | 0.17943409 |
| 121,700 | 0.27010674 | 0.17950835 |
| 121,800 | 0.270096223 | 0.17933401 |
| 121,900 | 0.269897491 | 0.17937653 |
| 122,000 | 0.269814114 | 0.17940484 |
| 122,100 | 0.269669879 | 0.17939021 |
| 122,200 | 0.269774046 | 0.17939313 |
| 122,300 | 0.26976488 | 0.17935152 |
| 122,400 | 0.269800824 | 0.17938761 |
| 122,500 | 0.269809718 | 0.17942901 |
| 122,600 | 0.269730323 | 0.17947707 |
| 122,700 | 0.269798112 | 0.17936457 |
| 122,800 | 0.269583372 | 0.17941086 |
| 122,900 | 0.269547739 | 0.17942604 |
| 123,000 | 0.269595317 | 0.1794557 |
| 123,100 | 0.269228394 | 0.17938316 |
| 123,200 | 0.269344169 | 0.17943893 |
| 123,300 | 0.269212959 | 0.17938017 |
| 123,400 | 0.269305487 | 0.17954834 |
| 123,500 | 0.269028053 | 0.17949958 |
| 123,600 | 0.269132559 | 0.17946687 |
| 123,700 | 0.26918693 | 0.17944659 |
| 123,800 | 0.269078452 | 0.17938236 |
| 123,900 | 0.268981098 | 0.17942832 |
| 124,000 | 0.26884121 | 0.17945778 |
| 124,100 | 0.268762735 | 0.17940042 |
| 124,200 | 0.268704779 | 0.17938144 |
| 124,300 | 0.268651529 | 0.17946078 |
| 124,400 | 0.268697649 | 0.17949371 |
| 124,500 | 0.268706224 | 0.17949352 |
| 124,600 | 0.268585466 | 0.17950142 |
| 124,700 | 0.268517302 | 0.17949858 |
| 124,800 | 0.268473415 | 0.17937021 |
| 124,900 | 0.268422896 | 0.17942488 |
| 125,000 | 0.268312907 | 0.17941433 |
| 125,100 | 0.268428993 | 0.1794368 |
| 125,200 | 0.268230411 | 0.1794355 |
| 125,300 | 0.268211259 | 0.17942585 |
| 125,400 | 0.268045038 | 0.17943248 |
| 125,500 | 0.268185615 | 0.17940912 |
| 125,600 | 0.267784955 | 0.17946836 |
| 125,700 | 0.268097281 | 0.17941328 |
| 125,800 | 0.267938591 | 0.1793887 |
| 125,900 | 0.267902094 | 0.17943186 |
| 126,000 | 0.267683472 | 0.17940773 |
| 126,100 | 0.267605865 | 0.17942942 |
| 126,200 | 0.267810997 | 0.17943709 |

| | | |
|---|---|---|
| 126,300 | 0.267731779 | 0.17940943 |
| 126,400 | 0.267573258 | 0.17947237 |
| 126,500 | 0.267345469 | 0.17947679 |
| 126,600 | 0.267252262 | 0.1793692 |
| 126,700 | 0.267366922 | 0.17945 |
| 126,800 | 0.267216295 | 0.17936397 |
| 126,900 | 0.267402371 | 0.1794464 |
| 127,000 | 0.267441078 | 0.17939298 |
| 127,100 | 0.267464003 | 0.17935955 |
| 127,200 | 0.267210856 | 0.17941373 |
| 127,300 | 0.267242302 | 0.17941665 |
| 127,400 | 0.266976252 | 0.17943951 |
| 127,500 | 0.267074053 | 0.17939593 |
| 127,600 | 0.266970231 | 0.17937421 |
| 127,700 | 0.267072592 | 0.17934498 |
| 127,800 | 0.267159014 | 0.17947039 |
| 127,900 | 0.267231517 | 0.17939755 |
| 128,000 | 0.267009973 | 0.17941809 |
| 128,100 | 0.266706661 | 0.17940468 |
| 128,200 | 0.266953759 | 0.17949278 |
| 128,300 | 0.266585816 | 0.17944604 |
| 128,400 | 0.266703969 | 0.17936994 |
| 128,500 | 0.266754633 | 0.17937673 |
| 128,600 | 0.266316052 | 0.17941697 |
| 128,700 | 0.266450641 | 0.17940941 |
| 128,800 | 0.266427036 | 0.17939113 |
| 128,900 | 0.266420315 | 0.17942193 |
| 129,000 | 0.266343915 | 0.17941012 |
| 129,100 | 0.266205293 | 0.17937412 |
| 129,200 | 0.266324483 | 0.17944954 |
| 129,300 | 0.266237573 | 0.17936292 |
| 129,400 | 0.266303912 | 0.17940013 |
| 129,500 | 0.266259134 | 0.17939732 |
| 129,600 | 0.266457839 | 0.17941697 |
| 129,700 | 0.266244062 | 0.179476 |
| 129,800 | 0.266298607 | 0.17944165 |
| 129,900 | 0.266114705 | 0.179373 |
| 130,000 | 0.266310641 | 0.17932596 |
| 130,100 | 0.265988188 | 0.17937492 |
| 130,200 | 0.266144996 | 0.17942722 |
| 130,300 | 0.266075005 | 0.17944762 |
| 130,400 | 0.266026467 | 0.17945768 |
| 130,500 | 0.2657694 | 0.17936864 |
| 130,600 | 0.265629759 | 0.17947106 |
| 130,700 | 0.265734577 | 0.17946599 |
| 130,800 | 0.265564228 | 0.17945169 |
| 130,900 | 0.26552688 | 0.17941071 |
| 131,000 | 0.265642898 | 0.1794741 |
| 131,100 | 0.265509111 | 0.17941421 |
| 131,200 | 0.265502906 | 0.17936804 |
| 131,300 | 0.265421194 | 0.17948822 |
| 131,400 | 0.26523841 | 0.17940213 |
| 131,500 | 0.265295505 | 0.17939306 |
| 131,600 | 0.26518471 | 0.17939734 |
| 131,700 | 0.265132574 | 0.17941927 |
| 131,800 | 0.265083539 | 0.17935847 |
| 131,900 | 0.265212605 | 0.17934287 |
| 132,000 | 0.265185754 | 0.1792992 |
| 132,100 | 0.26506491 | 0.17931749 |
| 132,200 | 0.265132293 | 0.17936006 |
| 132,300 | 0.265059191 | 0.17933122 |
| 132,400 | 0.264840119 | 0.17939774 |
| 132,500 | 0.264757838 | 0.17941468 |
| 132,600 | 0.264820709 | 0.17942845 |
| 132,700 | 0.264490335 | 0.17921618 |
| 132,800 | 0.264448424 | 0.1793638 |
| 132,900 | 0.264584801 | 0.17935791 |

| | | |
|---|---|---|
| 133,000 | 0.26439676 | 0.1793447 |
| 133,100 | 0.264314146 | 0.1794418 |
| 133,200 | 0.264263423 | 0.17932298 |
| 133,300 | 0.264417116 | 0.17950575 |
| 133,400 | 0.26411162 | 0.17937431 |
| 133,500 | 0.264093949 | 0.17937105 |
| 133,600 | 0.264022125 | 0.17946054 |
| 133,700 | 0.264196038 | 0.17940272 |
| 133,800 | 0.264021945 | 0.17937151 |
| 133,900 | 0.264043648 | 0.17939379 |
| 134,000 | 0.264035997 | 0.17928563 |
| 134,100 | 0.263925089 | 0.17936864 |
| 134,200 | 0.263845763 | 0.17936041 |
| 134,300 | 0.263785958 | 0.17929258 |
| 134,400 | 0.263702309 | 0.17935368 |
| 134,500 | 0.263857473 | 0.17938102 |
| 134,600 | 0.263790122 | 0.17941121 |
| 134,700 | 0.263721771 | 0.17941546 |
| 134,800 | 0.263721431 | 0.1793989 |
| 134,900 | 0.263517334 | 0.179371 |
| 135,000 | 0.263451329 | 0.17937988 |
| 135,100 | 0.263287073 | 0.17943081 |
| 135,200 | 0.263242142 | 0.17939733 |
| 135,300 | 0.263434379 | 0.1793762 |
| 135,400 | 0.263219714 | 0.17941925 |
| 135,500 | 0.263257704 | 0.17937853 |
| 135,600 | 0.26313152 | 0.17936904 |
| 135,700 | 0.263049082 | 0.17936471 |
| 135,800 | 0.262946231 | 0.1794623 |
| 135,900 | 0.26296545 | 0.17946089 |
| 136,000 | 0.263109486 | 0.17940634 |
| 136,100 | 0.262974144 | 0.17943618 |
| 136,200 | 0.263090041 | 0.17935218 |
| 136,300 | 0.263057167 | 0.17934111 |
| 136,400 | 0.262921155 | 0.17935263 |
| 136,500 | 0.262893841 | 0.17933472 |
| 136,600 | 0.262691447 | 0.179342 |
| 136,700 | 0.262704844 | 0.17932548 |
| 136,800 | 0.262566397 | 0.17943503 |
| 136,900 | 0.26248138 | 0.17936126 |
| 137,000 | 0.262419443 | 0.17941378 |
| 137,100 | 0.262445322 | 0.17937365 |
| 137,200 | 0.262530889 | 0.1793832 |
| 137,300 | 0.262119158 | 0.17935199 |
| 137,400 | 0.262225795 | 0.17940442 |
| 137,500 | 0.262166859 | 0.17942741 |
| 137,600 | 0.262255564 | 0.17938845 |
| 137,700 | 0.262192752 | 0.17936347 |
| 137,800 | 0.262120396 | 0.17935232 |
| 137,900 | 0.262034032 | 0.17934413 |
| 138,000 | 0.262062624 | 0.17934348 |
| 138,100 | 0.261976641 | 0.17943298 |
| 138,200 | 0.261787263 | 0.17938185 |
| 138,300 | 0.261795038 | 0.17938482 |
| 138,400 | 0.261836788 | 0.17941858 |
| 138,500 | 0.26172425 | 0.1793745 |
| 138,600 | 0.261914261 | 0.17939237 |
| 138,700 | 0.261774852 | 0.17927423 |
| 138,800 | 0.261632721 | 0.1792715 |
| 138,900 | 0.261558425 | 0.17927021 |
| 139,000 | 0.261446478 | 0.17928537 |
| 139,100 | 0.261546199 | 0.17931026 |
| 139,200 | 0.261464762 | 0.17939036 |
| 139,300 | 0.261399413 | 0.17930653 |
| 139,400 | 0.261245214 | 0.17931591 |
| 139,500 | 0.261294294 | 0.17935544 |
| 139,600 | 0.26130408 | 0.1794011 |

| | | |
|---|---|---|
| 139,700 | 0.2610076 | 0.17939843 |
| 139,800 | 0.26092795 | 0.17927145 |
| 139,900 | 0.261166032 | 0.17929467 |
| 140,000 | 0.260851789 | 0.17930079 |
| 140,100 | 0.261021217 | 0.17942607 |
| 140,200 | 0.260875712 | 0.17933023 |
| 140,300 | 0.260743453 | 0.17934145 |
| 140,400 | 0.260852147 | 0.17930424 |
| 140,500 | 0.260628399 | 0.17925355 |
| 140,600 | 0.260785934 | 0.17925057 |
| 140,700 | 0.260377338 | 0.17929655 |
| 140,800 | 0.260474635 | 0.17924665 |
| 140,900 | 0.26063076 | 0.17931686 |
| 141,000 | 0.260551237 | 0.17932651 |
| 141,100 | 0.260427296 | 0.17925856 |
| 141,200 | 0.26037882 | 0.17919968 |
| 141,300 | 0.260388794 | 0.17926652 |
| 141,400 | 0.260162467 | 0.17927172 |
| 141,500 | 0.260268555 | 0.17924104 |
| 141,600 | 0.260199565 | 0.17928902 |
| 141,700 | 0.260148831 | 0.17921457 |
| 141,800 | 0.260075118 | 0.17923465 |
| 141,900 | 0.260098764 | 0.17935076 |
| 142,000 | 0.259905976 | 0.17931334 |
| 142,100 | 0.260076558 | 0.17925589 |
| 142,200 | 0.259915544 | 0.17927535 |
| 142,300 | 0.260001321 | 0.17938178 |
| 142,400 | 0.259808548 | 0.17924729 |
| 142,500 | 0.259954826 | 0.17931244 |
| 142,600 | 0.259784914 | 0.17934031 |
| 142,700 | 0.259672168 | 0.17927435 |
| 142,800 | 0.259555573 | 0.1792956 |
| 142,900 | 0.259513796 | 0.1793122 |
| 143,000 | 0.259366032 | 0.1793185 |
| 143,100 | 0.259465631 | 0.17929876 |
| 143,200 | 0.259246891 | 0.17928331 |
| 143,300 | 0.259314336 | 0.17921137 |
| 143,400 | 0.259148388 | 0.17935025 |
| 143,500 | 0.259123282 | 0.17920742 |
| 143,600 | 0.25919289 | 0.17926442 |
| 143,700 | 0.259036581 | 0.17929374 |
| 143,800 | 0.259019533 | 0.17928342 |
| 143,900 | 0.258908977 | 0.17924451 |
| 144,000 | 0.25891324 | 0.17917163 |
| 144,100 | 0.258815383 | 0.1792105 |
| 144,200 | 0.258746302 | 0.1792531 |
| 144,300 | 0.258981291 | 0.17919043 |
| 144,400 | 0.258854464 | 0.17923278 |
| 144,500 | 0.258830651 | 0.17935771 |
| 144,600 | 0.258744017 | 0.17920789 |
| 144,700 | 0.258501088 | 0.17924525 |
| 144,800 | 0.258559876 | 0.17927947 |
| 144,900 | 0.258437998 | 0.17931011 |
| 145,000 | 0.258461049 | 0.17920086 |
| 145,100 | 0.25868906 | 0.17928953 |
| 145,200 | 0.258488973 | 0.17921215 |
| 145,300 | 0.258618593 | 0.17941799 |
| 145,400 | 0.258459171 | 0.17931233 |
| 145,500 | 0.258327498 | 0.17926734 |
| 145,600 | 0.258349208 | 0.17931412 |
| 145,700 | 0.258507641 | 0.17936556 |
| 145,800 | 0.258344518 | 0.17931231 |
| 145,900 | 0.258086096 | 0.17932565 |
| 146,000 | 0.258299961 | 0.1792651 |
| 146,100 | 0.25822326 | 0.17932204 |
| 146,200 | 0.258278004 | 0.17933106 |
| 146,300 | 0.258322312 | 0.17933151 |

| | | |
|---|---|---|
| 146,400 | 0.258257997 | 0.17922619 |
| 146,500 | 0.258034636 | 0.17925635 |
| 146,600 | 0.258036423 | 0.17921677 |
| 146,700 | 0.258056528 | 0.17922611 |
| 146,800 | 0.257857596 | 0.17919025 |
| 146,900 | 0.257851016 | 0.17927367 |
| 147,000 | 0.257799295 | 0.17919762 |
| 147,100 | 0.257622284 | 0.17928696 |
| 147,200 | 0.257639453 | 0.17928278 |
| 147,300 | 0.257638858 | 0.17929472 |
| 147,400 | 0.257557961 | 0.17930906 |
| 147,500 | 0.257438818 | 0.17924349 |
| 147,600 | 0.257454901 | 0.17929032 |
| 147,700 | 0.257480148 | 0.1793259 |
| 147,800 | 0.257494054 | 0.17918439 |
| 147,900 | 0.257406811 | 0.17927258 |
| 148,000 | 0.257362629 | 0.17927902 |
| 148,100 | 0.257318321 | 0.17928152 |
| 148,200 | 0.257148984 | 0.17921666 |
| 148,300 | 0.257250911 | 0.17927577 |
| 148,400 | 0.257181788 | 0.17927359 |
| 148,500 | 0.256901313 | 0.17929822 |
| 148,600 | 0.25702449 | 0.17923001 |
| 148,700 | 0.256832992 | 0.17922783 |
| 148,800 | 0.256986582 | 0.17928593 |
| 148,900 | 0.256747403 | 0.17922277 |
| 149,000 | 0.256755139 | 0.17920949 |
| 149,100 | 0.256599774 | 0.17922496 |
| 149,200 | 0.25664699 | 0.17921364 |
| 149,300 | 0.256561646 | 0.179225 |
| 149,400 | 0.256571441 | 0.1791954 |
| 149,500 | 0.256631379 | 0.17925159 |
| 149,600 | 0.256508106 | 0.17930728 |
| 149,700 | 0.256615147 | 0.17918734 |
| 149,800 | 0.256552262 | 0.17927097 |
| 149,900 | 0.256311688 | 0.1792977 |
| 150,000 | 0.256077032 | 0.17923274 |
| 150,100 | 0.256104631 | 0.17921436 |
| 150,200 | 0.256196587 | 0.17929299 |
| 150,300 | 0.256170629 | 0.17925317 |
| 150,400 | 0.256209641 | 0.17929269 |
| 150,500 | 0.25610418 | 0.17918794 |
| 150,600 | 0.256127963 | 0.17922964 |
| 150,700 | 0.255959013 | 0.17928945 |
| 150,800 | 0.255782961 | 0.17929887 |
| 150,900 | 0.255770383 | 0.17918733 |
| 151,000 | 0.255955227 | 0.17936055 |
| 151,100 | 0.255833117 | 0.17924722 |
| 151,200 | 0.255671144 | 0.1792548 |
| 151,300 | 0.255641979 | 0.17926331 |
| 151,400 | 0.255638575 | 0.17922893 |
| 151,500 | 0.255692478 | 0.17927836 |
| 151,600 | 0.255714989 | 0.17917251 |
| 151,700 | 0.25554489 | 0.17926076 |
| 151,800 | 0.255605376 | 0.17923518 |
| 151,900 | 0.255665618 | 0.17932024 |
| 152,000 | 0.255460048 | 0.17925039 |
| 152,100 | 0.255188884 | 0.17922534 |
| 152,200 | 0.255359735 | 0.17921039 |
| 152,300 | 0.255263908 | 0.17923725 |
| 152,400 | 0.255224803 | 0.17914696 |
| 152,500 | 0.255224447 | 0.17919214 |
| 152,600 | 0.255172737 | 0.17912613 |
| 152,700 | 0.255128305 | 0.17914948 |
| 152,800 | 0.255070704 | 0.17917464 |
| 152,900 | 0.254987977 | 0.17926592 |
| 153,000 | 0.254880539 | 0.17922812 |

| | | |
|---|---|---|
| 153,100 | 0.254858251 | 0.17930176 |
| 153,200 | 0.255105374 | 0.1792443 |
| 153,300 | 0.255031588 | 0.17925375 |
| 153,400 | 0.254918518 | 0.17931409 |
| 153,500 | 0.255041372 | 0.17912358 |
| 153,600 | 0.254852646 | 0.1793406 |
| 153,700 | 0.254728199 | 0.1792407 |
| 153,800 | 0.254765749 | 0.17931973 |
| 153,900 | 0.254727831 | 0.17925748 |
| 154,000 | 0.254791455 | 0.17922137 |
| 154,100 | 0.254665629 | 0.17923498 |
| 154,200 | 0.254707228 | 0.17920948 |
| 154,300 | 0.254633443 | 0.17917447 |
| 154,400 | 0.254496845 | 0.17921365 |
| 154,500 | 0.254440225 | 0.17932544 |
| 154,600 | 0.254436771 | 0.1792645 |
| 154,700 | 0.254342064 | 0.1792273 |
| 154,800 | 0.254343819 | 0.17928083 |
| 154,900 | 0.254156765 | 0.17925383 |
| 155,000 | 0.254315438 | 0.17920031 |
| 155,100 | 0.254020498 | 0.17916365 |
| 155,200 | 0.254144725 | 0.17926317 |
| 155,300 | 0.254346747 | 0.17919768 |
| 155,400 | 0.254267178 | 0.1792217 |
| 155,500 | 0.254237661 | 0.17914766 |
| 155,600 | 0.254108569 | 0.17919779 |
| 155,700 | 0.254090681 | 0.17926827 |
| 155,800 | 0.254199116 | 0.17920605 |
| 155,900 | 0.25409883 | 0.17922063 |
| 156,000 | 0.254137803 | 0.17915517 |
| 156,100 | 0.254091854 | 0.17916491 |
| 156,200 | 0.254054399 | 0.17916215 |
| 156,300 | 0.254062858 | 0.1791073 |
| 156,400 | 0.253900653 | 0.17919901 |
| 156,500 | 0.253936943 | 0.17910981 |
| 156,600 | 0.253939502 | 0.17919389 |
| 156,700 | 0.253622536 | 0.17907807 |
| 156,800 | 0.253596959 | 0.17917799 |
| 156,900 | 0.253476596 | 0.17917266 |
| 157,000 | 0.253610313 | 0.17912256 |
| 157,100 | 0.253683788 | 0.17912835 |
| 157,200 | 0.253651216 | 0.17918047 |
| 157,300 | 0.253574898 | 0.17919335 |
| 157,400 | 0.253399352 | 0.17925378 |
| 157,500 | 0.253385574 | 0.17923904 |
| 157,600 | 0.253367501 | 0.17926857 |
| 157,700 | 0.253083342 | 0.17913262 |
| 157,800 | 0.25328187 | 0.17915813 |
| 157,900 | 0.253255854 | 0.17914145 |
| 158,000 | 0.25309868 | 0.17919319 |
| 158,100 | 0.253233003 | 0.17912837 |
| 158,200 | 0.253148124 | 0.17909032 |
| 158,300 | 0.253261747 | 0.17917132 |
| 158,400 | 0.253127766 | 0.17913673 |
| 158,500 | 0.252867022 | 0.17915959 |
| 158,600 | 0.253006178 | 0.17916089 |
| 158,700 | 0.252885078 | 0.17914591 |
| 158,800 | 0.252892381 | 0.17908552 |
| 158,900 | 0.252745436 | 0.17915559 |
| 159,000 | 0.252931799 | 0.17919133 |
| 159,100 | 0.252921073 | 0.17916604 |
| 159,200 | 0.25289423 | 0.17924356 |
| 159,300 | 0.252809863 | 0.17921783 |
| 159,400 | 0.252904883 | 0.17925147 |
| 159,500 | 0.252871899 | 0.17915567 |
| 159,600 | 0.252913338 | 0.17914459 |
| 159,700 | 0.252646633 | 0.17911597 |

| | | |
|---|---|---|
| 159,800 | 0.252764254 | 0.17908804 |
| 159,900 | 0.252724738 | 0.17916365 |
| 160,000 | 0.252678696 | 0.17915244 |
| 160,100 | 0.252556508 | 0.17925254 |
| 160,200 | 0.252740997 | 0.17923903 |
| 160,300 | 0.252582832 | 0.17918833 |
| 160,400 | 0.252393251 | 0.17918152 |
| 160,500 | 0.252521037 | 0.17915093 |
| 160,600 | 0.252536369 | 0.17911698 |
| 160,700 | 0.25237573 | 0.17917843 |
| 160,800 | 0.252432778 | 0.1792173 |
| 160,900 | 0.252232153 | 0.17925107 |
| 161,000 | 0.252305638 | 0.17921637 |
| 161,100 | 0.252204427 | 0.1792351 |
| 161,200 | 0.25218715 | 0.17924525 |
| 161,300 | 0.252009441 | 0.17915909 |
| 161,400 | 0.252061225 | 0.17912729 |
| 161,500 | 0.25191128 | 0.17924033 |
| 161,600 | 0.252086124 | 0.17924152 |
| 161,700 | 0.252142495 | 0.17918271 |
| 161,800 | 0.251873661 | 0.17917638 |
| 161,900 | 0.251723128 | 0.17915716 |
| 162,000 | 0.251762155 | 0.17921606 |
| 162,100 | 0.251750492 | 0.17920526 |
| 162,200 | 0.251514711 | 0.17921172 |
| 162,300 | 0.251863433 | 0.17914908 |
| 162,400 | 0.251679757 | 0.17923695 |
| 162,500 | 0.25161884 | 0.17916167 |
| 162,600 | 0.251584773 | 0.1791928 |
| 162,700 | 0.251596228 | 0.17924864 |
| 162,800 | 0.251631519 | 0.17913155 |
| 162,900 | 0.251574844 | 0.17914609 |
| 163,000 | 0.251530783 | 0.17917303 |
| 163,100 | 0.251529271 | 0.17916724 |
| 163,200 | 0.25138253 | 0.179201 |
| 163,300 | 0.25125395 | 0.17918636 |
| 163,400 | 0.251287142 | 0.17922329 |
| 163,500 | 0.251213105 | 0.17913216 |
| 163,600 | 0.251219766 | 0.17921072 |
| 163,700 | 0.251110595 | 0.17921081 |
| 163,800 | 0.251107193 | 0.17917931 |
| 163,900 | 0.251167969 | 0.17915807 |
| 164,000 | 0.251091599 | 0.17921707 |
| 164,100 | 0.251238776 | 0.17920103 |
| 164,200 | 0.251187876 | 0.17914829 |
| 164,300 | 0.251126669 | 0.1792018 |
| 164,400 | 0.250884104 | 0.17915878 |
| 164,500 | 0.250870085 | 0.17917511 |
| 164,600 | 0.250930854 | 0.17930493 |
| 164,700 | 0.250811469 | 0.17915773 |
| 164,800 | 0.250899453 | 0.17911056 |
| 164,900 | 0.25075236 | 0.1791961 |
| 165,000 | 0.250717057 | 0.17908283 |
| 165,100 | 0.250751697 | 0.17920593 |
| 165,200 | 0.250587678 | 0.17917104 |
| 165,300 | 0.250561679 | 0.17918773 |
| 165,400 | 0.25052269 | 0.17916785 |
| 165,500 | 0.250658196 | 0.17924418 |
| 165,600 | 0.250532568 | 0.1791891 |
| 165,700 | 0.250521901 | 0.17908789 |
| 165,800 | 0.250343497 | 0.17917867 |
| 165,900 | 0.250236976 | 0.1791445 |
| 166,000 | 0.250216376 | 0.17917338 |
| 166,100 | 0.249997176 | 0.1791985 |
| 166,200 | 0.250008946 | 0.17917497 |
| 166,300 | 0.24998838 | 0.17915484 |
| 166,400 | 0.250191788 | 0.17908552 |

| | | |
|---|---|---|
| 166,500 | 0.249917737 | 0.17908183 |
| 166,600 | 0.24983783 | 0.17918659 |
| 166,700 | 0.249715667 | 0.17921039 |
| 166,800 | 0.249736288 | 0.17918525 |
| 166,900 | 0.24985824 | 0.1791492 |
| 167,000 | 0.249634838 | 0.17914623 |
| 167,100 | 0.249517884 | 0.17919461 |
| 167,200 | 0.249589773 | 0.17919132 |
| 167,300 | 0.249661581 | 0.17911104 |
| 167,400 | 0.249573776 | 0.17906559 |
| 167,500 | 0.249520573 | 0.17911325 |
| 167,600 | 0.24954063 | 0.17914117 |
| 167,700 | 0.249500596 | 0.17918821 |
| 167,800 | 0.249319665 | 0.17911312 |
| 167,900 | 0.24957855 | 0.17907356 |
| 168,000 | 0.249387213 | 0.17898646 |
| 168,100 | 0.249425832 | 0.17908138 |
| 168,200 | 0.249328886 | 0.17910445 |
| 168,300 | 0.249351619 | 0.17904215 |
| 168,400 | 0.249308979 | 0.17908069 |
| 168,500 | 0.249016449 | 0.17911152 |
| 168,600 | 0.249231292 | 0.17913353 |
| 168,700 | 0.249236473 | 0.17910839 |
| 168,800 | 0.249162741 | 0.17909555 |
| 168,900 | 0.248919164 | 0.17914018 |
| 169,000 | 0.249045283 | 0.17911784 |
| 169,100 | 0.248876623 | 0.17903497 |
| 169,200 | 0.249208545 | 0.17903732 |
| 169,300 | 0.248985015 | 0.17905701 |
| 169,400 | 0.249045505 | 0.17905961 |
| 169,500 | 0.248939607 | 0.1790449 |
| 169,600 | 0.248667417 | 0.1790597 |
| 169,700 | 0.248741161 | 0.17908983 |
| 169,800 | 0.248895112 | 0.17907775 |
| 169,900 | 0.248812785 | 0.17912818 |
| 170,000 | 0.248648406 | 0.17901844 |
| 170,100 | 0.248552654 | 0.17909152 |
| 170,200 | 0.248564277 | 0.17913535 |
| 170,300 | 0.248712731 | 0.17907802 |
| 170,400 | 0.248504295 | 0.17910685 |
| 170,500 | 0.248542133 | 0.17908915 |
| 170,600 | 0.248507652 | 0.17914781 |
| 170,700 | 0.24842374 | 0.17910011 |
| 170,800 | 0.248271389 | 0.17905616 |
| 170,900 | 0.248317524 | 0.1791834 |
| 171,000 | 0.248344711 | 0.17907379 |
| 171,100 | 0.248172111 | 0.17908233 |
| 171,200 | 0.248178349 | 0.17904867 |
| 171,300 | 0.248296 | 0.17912088 |
| 171,400 | 0.248244214 | 0.17903996 |
| 171,500 | 0.2482955 | 0.17917743 |
| 171,600 | 0.248187642 | 0.1790389 |
| 171,700 | 0.248228377 | 0.17906128 |
| 171,800 | 0.248178702 | 0.17907316 |
| 171,900 | 0.248255541 | 0.17904221 |
| 172,000 | 0.248069769 | 0.1790445 |
| 172,100 | 0.248118148 | 0.17910846 |
| 172,200 | 0.247952258 | 0.17903492 |
| 172,300 | 0.247899406 | 0.17912474 |
| 172,400 | 0.247933551 | 0.17905827 |
| 172,500 | 0.247851278 | 0.17906677 |
| 172,600 | 0.247974288 | 0.17905312 |
| 172,700 | 0.247799833 | 0.17909755 |
| 172,800 | 0.247593693 | 0.1790342 |
| 172,900 | 0.247594868 | 0.17910826 |
| 173,000 | 0.247663318 | 0.17904553 |
| 173,100 | 0.247756657 | 0.17902697 |

| | | |
|---|---|---|
| 173,200 | 0.247682441 | 0.17904049 |
| 173,300 | 0.247548331 | 0.17904602 |
| 173,400 | 0.247669758 | 0.17902406 |
| 173,500 | 0.247479709 | 0.17903172 |
| 173,600 | 0.247697644 | 0.17912223 |
| 173,700 | 0.247650249 | 0.17909531 |
| 173,800 | 0.247568943 | 0.1790445 |
| 173,900 | 0.247430523 | 0.17906031 |
| 174,000 | 0.24731787 | 0.17912974 |
| 174,100 | 0.247346246 | 0.17915736 |
| 174,200 | 0.247272909 | 0.17902133 |
| 174,300 | 0.247344738 | 0.17909065 |
| 174,400 | 0.247471671 | 0.17918774 |
| 174,500 | 0.247243104 | 0.17914921 |
| 174,600 | 0.247173382 | 0.17907172 |
| 174,700 | 0.247224331 | 0.17909048 |
| 174,800 | 0.247077586 | 0.17904012 |
| 174,900 | 0.247143897 | 0.1791123 |
| 175,000 | 0.247099632 | 0.17907726 |
| 175,100 | 0.246874577 | 0.17908308 |
| 175,200 | 0.247040737 | 0.17917348 |
| 175,300 | 0.247204083 | 0.1791235 |
| 175,400 | 0.24710278 | 0.17911038 |
| 175,500 | 0.247156614 | 0.17908887 |
| 175,600 | 0.24696829 | 0.17911056 |
| 175,700 | 0.247063108 | 0.17909732 |
| 175,800 | 0.246983362 | 0.17907827 |
| 175,900 | 0.246987342 | 0.17908894 |
| 176,000 | 0.246790298 | 0.17910808 |
| 176,100 | 0.246769118 | 0.17906808 |
| 176,200 | 0.246627486 | 0.17907236 |
| 176,300 | 0.24675559 | 0.17900986 |
| 176,400 | 0.246779771 | 0.17907367 |
| 176,500 | 0.246679669 | 0.17900953 |
| 176,600 | 0.246590719 | 0.17905049 |
| 176,700 | 0.246668717 | 0.17900544 |
| 176,800 | 0.246710082 | 0.17892099 |
| 176,900 | 0.246630325 | 0.17910206 |
| 177,000 | 0.246567054 | 0.17908053 |
| 177,100 | 0.246581765 | 0.17907619 |
| 177,200 | 0.246418186 | 0.17906512 |
| 177,300 | 0.246352684 | 0.1790317 |
| 177,400 | 0.246486443 | 0.17898991 |
| 177,500 | 0.246259237 | 0.17898338 |
| 177,600 | 0.246234476 | 0.17902325 |
| 177,700 | 0.246095997 | 0.17910898 |
| 177,800 | 0.246054957 | 0.17905542 |
| 177,900 | 0.245961831 | 0.17900511 |
| 178,000 | 0.246073785 | 0.17906644 |
| 178,100 | 0.246182219 | 0.17906651 |
| 178,200 | 0.24583637 | 0.17898201 |
| 178,300 | 0.245872729 | 0.17915235 |
| 178,400 | 0.245787812 | 0.17905936 |
| 178,500 | 0.245695894 | 0.17900869 |
| 178,600 | 0.245518972 | 0.17892792 |
| 178,700 | 0.245632565 | 0.1790639 |
| 178,800 | 0.245720969 | 0.17902942 |
| 178,900 | 0.245529126 | 0.17896622 |
| 179,000 | 0.24556904 | 0.17907158 |
| 179,100 | 0.24544122 | 0.17898904 |
| 179,200 | 0.245456247 | 0.17915208 |
| 179,300 | 0.245244019 | 0.17900264 |
| 179,400 | 0.245385636 | 0.17901694 |
| 179,500 | 0.245408168 | 0.17901729 |
| 179,600 | 0.245183689 | 0.17902822 |
| 179,700 | 0.245107067 | 0.17895513 |
| 179,800 | 0.245095306 | 0.17907251 |

| | | |
|---|---|---|
| 179,900 | 0.244959874 | 0.17899664 |
| 180,000 | 0.244924712 | 0.17908082 |
| 180,100 | 0.24489204 | 0.17905255 |
| 180,200 | 0.244931835 | 0.1790577 |
| 180,300 | 0.244777025 | 0.17906546 |
| 180,400 | 0.244780994 | 0.17902738 |
| 180,500 | 0.244859845 | 0.17907714 |
| 180,600 | 0.244912094 | 0.17906366 |
| 180,700 | 0.244865505 | 0.17906406 |
| 180,800 | 0.24469594 | 0.17904417 |
| 180,900 | 0.24464223 | 0.17900794 |
| 181,000 | 0.244577167 | 0.17901323 |
| 181,100 | 0.244536872 | 0.17896196 |
| 181,200 | 0.244513692 | 0.17903054 |
| 181,300 | 0.24418545 | 0.17894622 |
| 181,400 | 0.244346363 | 0.17901947 |
| 181,500 | 0.244384057 | 0.17893445 |
| 181,600 | 0.244262944 | 0.17894201 |
| 181,700 | 0.244272148 | 0.17894676 |
| 181,800 | 0.244185376 | 0.17896263 |
| 181,900 | 0.244129455 | 0.179036 |
| 182,000 | 0.244065526 | 0.17900418 |
| 182,100 | 0.244122208 | 0.17911237 |
| 182,200 | 0.244028849 | 0.1790518 |
| 182,300 | 0.243965425 | 0.17909423 |
| 182,400 | 0.244005891 | 0.17892417 |
| 182,500 | 0.243892 | 0.17896274 |
| 182,600 | 0.243911707 | 0.17901924 |
| 182,700 | 0.24414499 | 0.17907034 |
| 182,800 | 0.244101468 | 0.17900418 |
| 182,900 | 0.243977579 | 0.17910287 |
| 183,000 | 0.243782414 | 0.17894344 |
| 183,100 | 0.243801092 | 0.17902432 |
| 183,200 | 0.243775685 | 0.1790147 |
| 183,300 | 0.243517806 | 0.17905613 |
| 183,400 | 0.243729656 | 0.17904944 |
| 183,500 | 0.24360355 | 0.17905765 |
| 183,600 | 0.243627626 | 0.17909244 |
| 183,700 | 0.243470129 | 0.17907306 |
| 183,800 | 0.24341248 | 0.17904342 |
| 183,900 | 0.243455262 | 0.17907523 |
| 184,000 | 0.2433315 | 0.17902066 |
| 184,100 | 0.243439735 | 0.17899049 |
| 184,200 | 0.243338682 | 0.17913969 |
| 184,300 | 0.243179401 | 0.17903217 |
| 184,400 | 0.243111882 | 0.17908183 |
| 184,500 | 0.243232416 | 0.1790602 |
| 184,600 | 0.243162346 | 0.17910119 |
| 184,700 | 0.243199517 | 0.17905948 |
| 184,800 | 0.243179488 | 0.17905325 |
| 184,900 | 0.243163172 | 0.17902255 |
| 185,000 | 0.24301027 | 0.17905047 |
| 185,100 | 0.242914243 | 0.17896791 |
| 185,200 | 0.242834182 | 0.1790973 |
| 185,300 | 0.242841208 | 0.17902041 |
| 185,400 | 0.2427654 | 0.17909744 |
| 185,500 | 0.242843486 | 0.17900344 |
| 185,600 | 0.242759409 | 0.17903415 |
| 185,700 | 0.242739454 | 0.17898917 |
| 185,800 | 0.242849961 | 0.17904611 |
| 185,900 | 0.242755227 | 0.17900422 |
| 186,000 | 0.242705554 | 0.17902448 |
| 186,100 | 0.242856064 | 0.17893196 |
| 186,200 | 0.242612626 | 0.17895418 |
| 186,300 | 0.242565338 | 0.17906906 |
| 186,400 | 0.242528073 | 0.17899393 |
| 186,500 | 0.242598231 | 0.17892753 |

| | | |
|---|---|---|
| 186,600 | 0.242622809 | 0.17902243 |
| 186,700 | 0.242478365 | 0.17904907 |
| 186,800 | 0.242386857 | 0.17901674 |
| 186,900 | 0.24261302 | 0.17897055 |
| 187,000 | 0.242610961 | 0.1789628 |
| 187,100 | 0.242501541 | 0.17910632 |
| 187,200 | 0.242292958 | 0.17905914 |
| 187,300 | 0.242228852 | 0.17903937 |
| 187,400 | 0.242186032 | 0.17905382 |
| 187,500 | 0.2421338 | 0.17907448 |
| 187,600 | 0.242249016 | 0.17899228 |
| 187,700 | 0.242244445 | 0.17906956 |
| 187,800 | 0.242020793 | 0.17909908 |
| 187,900 | 0.24207938 | 0.17903502 |
| 188,000 | 0.242081873 | 0.17901456 |
| 188,100 | 0.241940101 | 0.17896028 |
| 188,200 | 0.242150868 | 0.17900059 |
| 188,300 | 0.241927278 | 0.17899911 |
| 188,400 | 0.242008649 | 0.17906324 |
| 188,500 | 0.241926326 | 0.17907529 |
| 188,600 | 0.241992212 | 0.17905271 |
| 188,700 | 0.241843893 | 0.17905205 |
| 188,800 | 0.241793995 | 0.17904425 |
| 188,900 | 0.241620626 | 0.1790259 |
| 189,000 | 0.241637472 | 0.179065 |
| 189,100 | 0.241772304 | 0.17903322 |
| 189,200 | 0.241750465 | 0.17900643 |
| 189,300 | 0.241772177 | 0.179061 |
| 189,400 | 0.241654921 | 0.17903372 |
| 189,500 | 0.24156607 | 0.17906465 |
| 189,600 | 0.241538246 | 0.17908601 |
| 189,700 | 0.24128903 | 0.17904828 |
| 189,800 | 0.241400281 | 0.17901346 |
| 189,900 | 0.241446956 | 0.17906063 |
| 190,000 | 0.241552624 | 0.17908471 |
| 190,100 | 0.241403234 | 0.17898449 |
| 190,200 | 0.241430732 | 0.17902335 |
| 190,300 | 0.241417405 | 0.17894952 |
| 190,400 | 0.241241435 | 0.17899784 |
| 190,500 | 0.241206516 | 0.17893458 |
| 190,600 | 0.241167636 | 0.17897751 |
| 190,700 | 0.241276249 | 0.17905567 |
| 190,800 | 0.241438804 | 0.17890133 |
| 190,900 | 0.241191212 | 0.17904022 |
| 191,000 | 0.241031408 | 0.17905243 |
| 191,100 | 0.241191716 | 0.17901706 |
| 191,200 | 0.241101259 | 0.17898102 |
| 191,300 | 0.241048887 | 0.17895246 |
| 191,400 | 0.241130434 | 0.17902796 |
| 191,500 | 0.241189327 | 0.17909265 |
| 191,600 | 0.240990051 | 0.17902263 |
| 191,700 | 0.240787426 | 0.17899958 |
| 191,800 | 0.240938847 | 0.17900892 |
| 191,900 | 0.240941705 | 0.17894168 |
| 192,000 | 0.240769364 | 0.17907138 |
| 192,100 | 0.240691086 | 0.17903532 |
| 192,200 | 0.240714374 | 0.1789445 |
| 192,300 | 0.240622191 | 0.17897359 |
| 192,400 | 0.240730329 | 0.1790061 |
| 192,500 | 0.240766074 | 0.17904378 |
| 192,600 | 0.24067839 | 0.17895419 |
| 192,700 | 0.240678362 | 0.17893274 |
| 192,800 | 0.24057718 | 0.17892168 |
| 192,900 | 0.240375345 | 0.17896511 |
| 193,000 | 0.24061051 | 0.1789871 |
| 193,100 | 0.24044151 | 0.17892459 |
| 193,200 | 0.240618421 | 0.17904595 |

| | | |
|---|---|---|
| 193,300 | 0.240438448 | 0.17902339 |
| 193,400 | 0.240146974 | 0.17906375 |
| 193,500 | 0.240440996 | 0.17900977 |
| 193,600 | 0.240159318 | 0.17892927 |
| 193,700 | 0.240226301 | 0.17891181 |
| 193,800 | 0.240295852 | 0.17896805 |
| 193,900 | 0.2402843 | 0.17898544 |
| 194,000 | 0.240228962 | 0.17893809 |
| 194,100 | 0.240229227 | 0.17908204 |
| 194,200 | 0.239903267 | 0.17899019 |
| 194,300 | 0.240036035 | 0.17902374 |
| 194,400 | 0.240054434 | 0.17888687 |
| 194,500 | 0.239921938 | 0.17890103 |
| 194,600 | 0.239865032 | 0.17899987 |
| 194,700 | 0.239952581 | 0.1789664 |
| 194,800 | 0.240102155 | 0.1790156 |
| 194,900 | 0.23999246 | 0.17899992 |
| 195,000 | 0.240020112 | 0.17894322 |
| 195,100 | 0.239797753 | 0.17893079 |
| 195,200 | 0.239816684 | 0.17898107 |
| 195,300 | 0.239674014 | 0.1790818 |
| 195,400 | 0.239646164 | 0.17897093 |
| 195,500 | 0.23976675 | 0.17894855 |
| 195,600 | 0.239889908 | 0.17891345 |
| 195,700 | 0.239722674 | 0.1789241 |
| 195,800 | 0.239822146 | 0.17888549 |
| 195,900 | 0.239373447 | 0.17907502 |
| 196,000 | 0.239482655 | 0.17900737 |
| 196,100 | 0.23954466 | 0.17897932 |
| 196,200 | 0.23961128 | 0.17899176 |
| 196,300 | 0.239429589 | 0.17901329 |
| 196,400 | 0.239463215 | 0.17898922 |
| 196,500 | 0.23931073 | 0.17892598 |
| 196,600 | 0.239200159 | 0.17895652 |
| 196,700 | 0.23922229 | 0.17896895 |
| 196,800 | 0.23894352 | 0.17898089 |
| 196,900 | 0.239097721 | 0.17894763 |
| 197,000 | 0.239030099 | 0.17903655 |
| 197,100 | 0.239192743 | 0.17896528 |
| 197,200 | 0.239191088 | 0.17897844 |
| 197,300 | 0.239078991 | 0.17904455 |
| 197,400 | 0.238990181 | 0.1790162 |
| 197,500 | 0.238950807 | 0.17897244 |
| 197,600 | 0.239126542 | 0.17904688 |
| 197,700 | 0.238960987 | 0.17897403 |
| 197,800 | 0.238795236 | 0.1790309 |
| 197,900 | 0.23876072 | 0.17900998 |
| 198,000 | 0.238564883 | 0.17905196 |
| 198,100 | 0.238676952 | 0.17894735 |
| 198,200 | 0.238706519 | 0.17900192 |
| 198,300 | 0.238460616 | 0.1788894 |
| 198,400 | 0.238394259 | 0.1789394 |
| 198,500 | 0.238658889 | 0.17894612 |
| 198,600 | 0.238586949 | 0.17889211 |
| 198,700 | 0.238694792 | 0.17901632 |
| 198,800 | 0.238504541 | 0.17895742 |
| 198,900 | 0.238454681 | 0.17896219 |
| 199,000 | 0.23855536 | 0.17892914 |
| 199,100 | 0.238687225 | 0.17904289 |
| 199,200 | 0.238664282 | 0.17899074 |
| 199,300 | 0.238571271 | 0.17893456 |
| 199,400 | 0.238602582 | 0.17895438 |
| 199,500 | 0.238406269 | 0.17905098 |
| 199,600 | 0.238494187 | 0.17898852 |
| 199,700 | 0.238485734 | 0.17899323 |
| 199,800 | 0.238279934 | 0.17896259 |
| 199,900 | 0.238243205 | 0.17891805 |

| | | |
|---|---|---|
| 200,000 | 0.238209021 | 0.17892024 |
| 200,100 | 0.238215424 | 0.17898021 |
| 200,200 | 0.238328178 | 0.17896658 |
| 200,300 | 0.238266962 | 0.17893707 |
| 200,400 | 0.238224405 | 0.17899985 |
| 200,500 | 0.238221587 | 0.17908522 |
| 200,600 | 0.238160091 | 0.17905738 |
| 200,700 | 0.238060048 | 0.17904474 |
| 200,800 | 0.238059216 | 0.17900908 |
| 200,900 | 0.237941914 | 0.17893344 |
| 201,000 | 0.237872922 | 0.17899982 |
| 201,100 | 0.237783606 | 0.17901574 |
| 201,200 | 0.237661711 | 0.17895678 |
| 201,300 | 0.237824677 | 0.17898964 |
| 201,400 | 0.237871535 | 0.17887824 |
| 201,500 | 0.237974923 | 0.17896346 |
| 201,600 | 0.237843071 | 0.17892669 |
| 201,700 | 0.237767394 | 0.17885961 |
| 201,800 | 0.237858166 | 0.17893567 |
| 201,900 | 0.23769328 | 0.17885076 |
| 202,000 | 0.237579937 | 0.17891327 |
| 202,100 | 0.237543852 | 0.17894606 |
| 202,200 | 0.237590591 | 0.17892662 |
| 202,300 | 0.237505966 | 0.17894663 |
| 202,400 | 0.237683122 | 0.17885653 |
| 202,500 | 0.237501524 | 0.17892942 |
| 202,600 | 0.237416122 | 0.17896471 |
| 202,700 | 0.237191799 | 0.17899965 |
| 202,800 | 0.237046674 | 0.17891819 |
| 202,900 | 0.237297424 | 0.17898956 |
| 203,000 | 0.237080628 | 0.17895617 |
| 203,100 | 0.237081113 | 0.17900396 |
| 203,200 | 0.237121438 | 0.17895191 |
| 203,300 | 0.236963129 | 0.17889112 |
| 203,400 | 0.237009427 | 0.17898252 |
| 203,500 | 0.237176828 | 0.17898229 |
| 203,600 | 0.237233753 | 0.17902658 |
| 203,700 | 0.237197147 | 0.1789336 |
| 203,800 | 0.236990698 | 0.17899989 |
| 203,900 | 0.236888844 | 0.17896787 |
| 204,000 | 0.236749515 | 0.1789655 |
| 204,100 | 0.236864417 | 0.17902012 |
| 204,200 | 0.236746159 | 0.17895142 |
| 204,300 | 0.236817775 | 0.17899479 |
| 204,400 | 0.236785648 | 0.17894085 |
| 204,500 | 0.236721876 | 0.17899368 |
| 204,600 | 0.236695513 | 0.17898663 |
| 204,700 | 0.236451934 | 0.17895391 |
| 204,800 | 0.236598212 | 0.17894632 |
| 204,900 | 0.236581947 | 0.17887133 |
| 205,000 | 0.236476505 | 0.17883509 |
| 205,100 | 0.236489027 | 0.17888178 |
| 205,200 | 0.236521882 | 0.17895711 |
| 205,300 | 0.236406285 | 0.17885847 |
| 205,400 | 0.236342375 | 0.17880517 |
| 205,500 | 0.236260577 | 0.17892293 |
| 205,600 | 0.236296746 | 0.17890911 |
| 205,700 | 0.236193349 | 0.17898988 |
| 205,800 | 0.236097426 | 0.17890951 |
| 205,900 | 0.236106032 | 0.17889796 |
| 206,000 | 0.236311781 | 0.17894568 |
| 206,100 | 0.235948417 | 0.17905245 |
| 206,200 | 0.236125848 | 0.17899829 |
| 206,300 | 0.23626107 | 0.17901305 |
| 206,400 | 0.235951064 | 0.17900036 |
| 206,500 | 0.236118368 | 0.17900867 |
| 206,600 | 0.236022124 | 0.17897051 |

| | | |
|---|---|---|
| 206,700 | 0.235967759 | 0.17888984 |
| 206,800 | 0.235818672 | 0.17901459 |
| 206,900 | 0.235954603 | 0.17890951 |
| 207,000 | 0.235870385 | 0.17900483 |
| 207,100 | 0.235668863 | 0.17887917 |
| 207,200 | 0.235668369 | 0.17899089 |
| 207,300 | 0.235469081 | 0.17890221 |
| 207,400 | 0.235560191 | 0.17890193 |
| 207,500 | 0.235580901 | 0.17893476 |
| 207,600 | 0.23583702 | 0.17882058 |
| 207,700 | 0.235667519 | 0.17892983 |
| 207,800 | 0.235670782 | 0.17887608 |
| 207,900 | 0.235700449 | 0.17886507 |
| 208,000 | 0.235574305 | 0.17882691 |
| 208,100 | 0.235570153 | 0.17878162 |
| 208,200 | 0.23570063 | 0.17877976 |
| 208,300 | 0.235449078 | 0.17890257 |
| 208,400 | 0.235572643 | 0.17894695 |
| 208,500 | 0.235408573 | 0.17879755 |
| 208,600 | 0.235494735 | 0.1789284 |
| 208,700 | 0.235512371 | 0.17893183 |
| 208,800 | 0.235427811 | 0.17891465 |
| 208,900 | 0.235174747 | 0.17888717 |
| 209,000 | 0.235321733 | 0.17897354 |
| 209,100 | 0.235269618 | 0.17892018 |
| 209,200 | 0.235262215 | 0.17895183 |
| 209,300 | 0.235066948 | 0.17893673 |
| 209,400 | 0.235235542 | 0.17896996 |
| 209,500 | 0.235300318 | 0.17896939 |
| 209,600 | 0.235114709 | 0.17888864 |
| 209,700 | 0.235266766 | 0.17891673 |
| 209,800 | 0.235142685 | 0.17894631 |
| 209,900 | 0.235114612 | 0.17904631 |
| 210,000 | 0.23506022 | 0.17894737 |
| 210,100 | 0.234974781 | 0.17880255 |
| 210,200 | 0.234915369 | 0.17879641 |
| 210,300 | 0.234850704 | 0.17894907 |
| 210,400 | 0.234835521 | 0.17890013 |
| 210,500 | 0.234835701 | 0.17896383 |
| 210,600 | 0.234750955 | 0.17891387 |
| 210,700 | 0.234719197 | 0.17875868 |
| 210,800 | 0.234704037 | 0.1788726 |
| 210,900 | 0.234545958 | 0.17883725 |
| 211,000 | 0.234610972 | 0.17880613 |
| 211,100 | 0.234609584 | 0.17897387 |
| 211,200 | 0.234528092 | 0.17888912 |
| 211,300 | 0.2345992 | 0.17881842 |
| 211,400 | 0.234516173 | 0.17880176 |
| 211,500 | 0.234560645 | 0.17890645 |
| 211,600 | 0.234373706 | 0.17892212 |
| 211,700 | 0.234443242 | 0.1788787 |
| 211,800 | 0.234455397 | 0.17885086 |
| 211,900 | 0.234329954 | 0.17889851 |
| 212,000 | 0.234369078 | 0.17887952 |
| 212,100 | 0.234100735 | 0.17890862 |
| 212,200 | 0.234103246 | 0.178863 |
| 212,300 | 0.233871697 | 0.17890065 |
| 212,400 | 0.234006668 | 0.17886188 |
| 212,500 | 0.234096214 | 0.17872589 |
| 212,600 | 0.233923688 | 0.17882045 |
| 212,700 | 0.234025969 | 0.1788633 |
| 212,800 | 0.233885026 | 0.17890632 |
| 212,900 | 0.233870735 | 0.17893007 |
| 213,000 | 0.233998742 | 0.17883879 |
| 213,100 | 0.233850808 | 0.17891918 |
| 213,200 | 0.233669584 | 0.17882619 |
| 213,300 | 0.233645032 | 0.17897783 |

| | | |
|---|---|---|
| 213,400 | 0.2337489 | 0.17888478 |
| 213,500 | 0.233854323 | 0.17883606 |
| 213,600 | 0.233527071 | 0.17893076 |
| 213,700 | 0.233535381 | 0.17893381 |
| 213,800 | 0.233462678 | 0.17891932 |
| 213,900 | 0.233500712 | 0.17890836 |
| 214,000 | 0.233249253 | 0.17888998 |
| 214,100 | 0.233398994 | 0.17894964 |
| 214,200 | 0.23339063 | 0.17884922 |
| 214,300 | 0.233375385 | 0.17887786 |
| 214,400 | 0.233155777 | 0.17892823 |
| 214,500 | 0.233234646 | 0.17886174 |
| 214,600 | 0.233241433 | 0.17896007 |
| 214,700 | 0.233178856 | 0.17901189 |
| 214,800 | 0.233003589 | 0.17895677 |
| 214,900 | 0.233065712 | 0.17887556 |
| 215,000 | 0.232942125 | 0.17886683 |
| 215,100 | 0.233063439 | 0.17890354 |
| 215,200 | 0.233084717 | 0.17894788 |
| 215,300 | 0.232992437 | 0.17883815 |
| 215,400 | 0.232882639 | 0.17878156 |
| 215,500 | 0.232829126 | 0.17882808 |
| 215,600 | 0.23255989 | 0.17887188 |
| 215,700 | 0.232836867 | 0.17889854 |
| 215,800 | 0.232774751 | 0.17889401 |
| 215,900 | 0.232622548 | 0.1788378 |
| 216,000 | 0.232532376 | 0.17888885 |
| 216,100 | 0.232602937 | 0.17888549 |
| 216,200 | 0.23267037 | 0.17888602 |
| 216,300 | 0.232572637 | 0.17885544 |
| 216,400 | 0.232505384 | 0.17892363 |
| 216,500 | 0.23255234 | 0.17888282 |
| 216,600 | 0.232589779 | 0.17889973 |
| 216,700 | 0.232544916 | 0.17894002 |
| 216,800 | 0.232506797 | 0.17890402 |
| 216,900 | 0.232422803 | 0.17890781 |
| 217,000 | 0.23263561 | 0.17899853 |
| 217,100 | 0.232177395 | 0.17892246 |
| 217,200 | 0.232221341 | 0.17904962 |
| 217,300 | 0.232267016 | 0.17896248 |
| 217,400 | 0.232246599 | 0.17890475 |
| 217,500 | 0.232266539 | 0.17900115 |
| 217,600 | 0.232451629 | 0.17896486 |
| 217,700 | 0.232188574 | 0.178969 |
| 217,800 | 0.232176172 | 0.1790268 |
| 217,900 | 0.232081346 | 0.1789481 |
| 218,000 | 0.232203335 | 0.17888243 |
| 218,100 | 0.232352975 | 0.17889525 |
| 218,200 | 0.232071702 | 0.17896172 |
| 218,300 | 0.232087176 | 0.17891829 |
| 218,400 | 0.232129398 | 0.17892235 |
| 218,500 | 0.231950916 | 0.17896726 |
| 218,600 | 0.232064509 | 0.17896999 |
| 218,700 | 0.231877475 | 0.17896317 |
| 218,800 | 0.232067741 | 0.17884356 |
| 218,900 | 0.231965597 | 0.1788944 |
| 219,000 | 0.232076786 | 0.17890252 |
| 219,100 | 0.231892151 | 0.17902408 |
| 219,200 | 0.231969185 | 0.1789996 |
| 219,300 | 0.231911267 | 0.17896193 |
| 219,400 | 0.231764104 | 0.17898776 |
| 219,500 | 0.231640787 | 0.17892096 |
| 219,600 | 0.231662006 | 0.17888109 |
| 219,700 | 0.231683347 | 0.1789778 |
| 219,800 | 0.23160764 | 0.17888636 |
| 219,900 | 0.231591258 | 0.17884818 |
| 220,000 | 0.231540568 | 0.17883039 |

| | | |
|---|---|---|
| 220,100 | 0.23151076 | 0.17873533 |
| 220,200 | 0.231437209 | 0.17892106 |
| 220,300 | 0.231205782 | 0.17888948 |
| 220,400 | 0.231425379 | 0.17886499 |
| 220,500 | 0.231426087 | 0.17896613 |
| 220,600 | 0.231344262 | 0.17888167 |
| 220,700 | 0.231344762 | 0.17890542 |
| 220,800 | 0.231319084 | 0.17892546 |
| 220,900 | 0.231231445 | 0.17890055 |
| 221,000 | 0.231306413 | 0.17895865 |
| 221,100 | 0.231127683 | 0.17896259 |
| 221,200 | 0.23122822 | 0.1789163 |
| 221,300 | 0.230998058 | 0.17892442 |
| 221,400 | 0.231307358 | 0.1788461 |
| 221,500 | 0.231150283 | 0.17884972 |
| 221,600 | 0.231031849 | 0.17893937 |
| 221,700 | 0.23113755 | 0.17900547 |
| 221,800 | 0.231119916 | 0.17891207 |
| 221,900 | 0.230984504 | 0.17889394 |
| 222,000 | 0.231024353 | 0.17891274 |
| 222,100 | 0.231132552 | 0.17894522 |
| 222,200 | 0.231066583 | 0.17898476 |
| 222,300 | 0.230830353 | 0.17893299 |
| 222,400 | 0.230938894 | 0.17894237 |
| 222,500 | 0.230823731 | 0.17895685 |
| 222,600 | 0.230789435 | 0.17894595 |
| 222,700 | 0.230593537 | 0.17890581 |
| 222,800 | 0.230763137 | 0.17900557 |
| 222,900 | 0.230813257 | 0.17887965 |
| 223,000 | 0.23088433 | 0.17892852 |
| 223,100 | 0.230932524 | 0.17899063 |
| 223,200 | 0.230962127 | 0.1788743 |
| 223,300 | 0.230746529 | 0.17907462 |
| 223,400 | 0.23074389 | 0.17892496 |
| 223,500 | 0.230791565 | 0.17891314 |
| 223,600 | 0.230808437 | 0.17899661 |
| 223,700 | 0.230595719 | 0.17891129 |
| 223,800 | 0.230538614 | 0.17889688 |
| 223,900 | 0.23035272 | 0.17899602 |
| 224,000 | 0.230410088 | 0.17896052 |
| 224,100 | 0.23034342 | 0.17888387 |
| 224,200 | 0.230185472 | 0.17897511 |
| 224,300 | 0.230208612 | 0.1789309 |
| 224,400 | 0.230126647 | 0.17903844 |
| 224,500 | 0.230173574 | 0.17897135 |
| 224,600 | 0.230126509 | 0.17885092 |
| 224,700 | 0.230126465 | 0.17891877 |
| 224,800 | 0.230037557 | 0.17885565 |
| 224,900 | 0.230122849 | 0.17886076 |
| 225,000 | 0.230077854 | 0.17886853 |
| 225,100 | 0.230101452 | 0.1789016 |
| 225,200 | 0.230073158 | 0.17894541 |
| 225,300 | 0.22980269 | 0.17892619 |
| 225,400 | 0.229870714 | 0.17896254 |
| 225,500 | 0.230015345 | 0.17895023 |
| 225,600 | 0.230048347 | 0.17888933 |
| 225,700 | 0.229917013 | 0.17889301 |
| 225,800 | 0.229718037 | 0.17895256 |
| 225,900 | 0.22977247 | 0.17898036 |
| 226,000 | 0.230000071 | 0.17893907 |
| 226,100 | 0.229660292 | 0.17880223 |
| 226,200 | 0.229826643 | 0.17887556 |
| 226,300 | 0.229478314 | 0.17895082 |
| 226,400 | 0.229708028 | 0.17894377 |
| 226,500 | 0.229559151 | 0.17888667 |
| 226,600 | 0.229507394 | 0.17888262 |
| 226,700 | 0.229426982 | 0.17887605 |

| | | |
|---|---|---|
| 226,800 | 0.229426356 | 0.17893014 |
| 226,900 | 0.229528564 | 0.17895152 |
| 227,000 | 0.229439638 | 0.17882624 |
| 227,100 | 0.229392719 | 0.17895344 |
| 227,200 | 0.229201718 | 0.17889569 |
| 227,300 | 0.22933989 | 0.1788566 |
| 227,400 | 0.229537267 | 0.17886406 |
| 227,500 | 0.2292176 | 0.1787529 |
| 227,600 | 0.229208786 | 0.17883931 |
| 227,700 | 0.229200528 | 0.17881624 |
| 227,800 | 0.229265147 | 0.17891305 |
| 227,900 | 0.229087051 | 0.17890686 |
| 228,000 | 0.229276841 | 0.17883891 |
| 228,100 | 0.229100845 | 0.17890701 |
| 228,200 | 0.229027189 | 0.17885651 |
| 228,300 | 0.229186396 | 0.17892273 |
| 228,400 | 0.229054712 | 0.17890767 |
| 228,500 | 0.229051598 | 0.17888275 |
| 228,600 | 0.229009065 | 0.17886125 |
| 228,700 | 0.228937482 | 0.17882416 |
| 228,800 | 0.22886892 | 0.17886493 |
| 228,900 | 0.228844771 | 0.17887433 |
| 229,000 | 0.228816856 | 0.17887324 |
| 229,100 | 0.22884276 | 0.17892809 |
| 229,200 | 0.22874125 | 0.17882504 |
| 229,300 | 0.228827016 | 0.17885337 |
| 229,400 | 0.229022635 | 0.17886071 |
| 229,500 | 0.228612938 | 0.17886208 |
| 229,600 | 0.228650447 | 0.17882427 |
| 229,700 | 0.228522366 | 0.17887648 |
| 229,800 | 0.22847874 | 0.17894932 |
| 229,900 | 0.228480993 | 0.17895626 |
| 230,000 | 0.228637823 | 0.17894905 |
| 230,100 | 0.228481898 | 0.17894978 |
| 230,200 | 0.228596445 | 0.17881206 |
| 230,300 | 0.228430902 | 0.17882043 |
| 230,400 | 0.22840442 | 0.17890427 |
| 230,500 | 0.228408239 | 0.17884294 |
| 230,600 | 0.228342217 | 0.17875852 |
| 230,700 | 0.228331858 | 0.17882664 |
| 230,800 | 0.228391442 | 0.17883858 |
| 230,900 | 0.228352995 | 0.17883491 |
| 231,000 | 0.228152971 | 0.17890762 |
| 231,100 | 0.228373607 | 0.17897317 |
| 231,200 | 0.228361295 | 0.17901364 |
| 231,300 | 0.228032314 | 0.17880194 |
| 231,400 | 0.22819907 | 0.17891095 |
| 231,500 | 0.228089965 | 0.17888728 |
| 231,600 | 0.228034014 | 0.17894754 |
| 231,700 | 0.228107771 | 0.17887592 |
| 231,800 | 0.228024121 | 0.17889324 |
| 231,900 | 0.22785704 | 0.17887285 |
| 232,000 | 0.227790617 | 0.17883848 |
| 232,100 | 0.227913741 | 0.17895516 |
| 232,200 | 0.227905354 | 0.1789424 |
| 232,300 | 0.227839007 | 0.17886919 |
| 232,400 | 0.22771127 | 0.17889357 |
| 232,500 | 0.227753251 | 0.17887282 |
| 232,600 | 0.227796934 | 0.17882977 |
| 232,700 | 0.22781183 | 0.17883062 |
| 232,800 | 0.227826193 | 0.17890956 |
| 232,900 | 0.227713129 | 0.17879518 |
| 233,000 | 0.227722361 | 0.17890418 |
| 233,100 | 0.227486471 | 0.17882815 |
| 233,200 | 0.227696587 | 0.17887213 |
| 233,300 | 0.227501619 | 0.17888918 |
| 233,400 | 0.227624485 | 0.17884968 |

| | | |
|---|---|---|
| 233,500 | 0.227545696 | 0.17885144 |
| 233,600 | 0.227627287 | 0.17885041 |
| 233,700 | 0.227512636 | 0.17890532 |
| 233,800 | 0.227603388 | 0.17887274 |
| 233,900 | 0.227318306 | 0.17882409 |
| 234,000 | 0.227478933 | 0.178926 |
| 234,100 | 0.227320798 | 0.17888837 |
| 234,200 | 0.227237894 | 0.1789422 |
| 234,300 | 0.227274949 | 0.17876567 |
| 234,400 | 0.227197057 | 0.17888567 |
| 234,500 | 0.227231592 | 0.178892 |
| 234,600 | 0.227142496 | 0.17887923 |
| 234,700 | 0.227066074 | 0.17883329 |
| 234,800 | 0.22723548 | 0.17891708 |
| 234,900 | 0.227173839 | 0.17894188 |
| 235,000 | 0.227126386 | 0.17891552 |
| 235,100 | 0.227175982 | 0.17894252 |
| 235,200 | 0.227277374 | 0.17888614 |
| 235,300 | 0.226853589 | 0.1788223 |
| 235,400 | 0.227011478 | 0.17883705 |
| 235,500 | 0.227176809 | 0.17887396 |
| 235,600 | 0.227078223 | 0.1787808 |
| 235,700 | 0.227026753 | 0.17880014 |
| 235,800 | 0.226785422 | 0.17881148 |
| 235,900 | 0.226907083 | 0.17892443 |
| 236,000 | 0.226869023 | 0.17882774 |
| 236,100 | 0.22681801 | 0.17890553 |
| 236,200 | 0.226939321 | 0.17880173 |
| 236,300 | 0.226662003 | 0.17891289 |
| 236,400 | 0.226731059 | 0.17889845 |
| 236,500 | 0.226766483 | 0.17885645 |
| 236,600 | 0.226592118 | 0.17892015 |
| 236,700 | 0.226553937 | 0.17886927 |
| 236,800 | 0.226662697 | 0.1789418 |
| 236,900 | 0.226447798 | 0.1788903 |
| 237,000 | 0.226433608 | 0.17882724 |
| 237,100 | 0.226478179 | 0.178872 |
| 237,200 | 0.226493202 | 0.17894874 |
| 237,300 | 0.226269749 | 0.17879107 |
| 237,400 | 0.226347554 | 0.17886746 |
| 237,500 | 0.226282745 | 0.17894406 |
| 237,600 | 0.226128583 | 0.17886105 |
| 237,700 | 0.226215441 | 0.17878511 |
| 237,800 | 0.226080052 | 0.17881546 |
| 237,900 | 0.226162943 | 0.17881689 |
| 238,000 | 0.226130386 | 0.17882774 |
| 238,100 | 0.226428186 | 0.17896412 |
| 238,200 | 0.226196301 | 0.17883758 |
| 238,300 | 0.226093428 | 0.17887054 |
| 238,400 | 0.226038201 | 0.17884799 |
| 238,500 | 0.226086639 | 0.17890158 |
| 238,600 | 0.226125835 | 0.17891399 |
| 238,700 | 0.22613209 | 0.17884438 |
| 238,800 | 0.225994108 | 0.17885609 |
| 238,900 | 0.226004489 | 0.17888343 |
| 239,000 | 0.225969344 | 0.1788524 |
| 239,100 | 0.225951175 | 0.17892447 |
| 239,200 | 0.225958314 | 0.1789569 |
| 239,300 | 0.226096202 | 0.17889567 |
| 239,400 | 0.22605951 | 0.17886098 |
| 239,500 | 0.225889636 | 0.17892838 |
| 239,600 | 0.225875729 | 0.17898302 |
| 239,700 | 0.225913648 | 0.1789724 |
| 239,800 | 0.225895369 | 0.17885147 |
| 239,900 | 0.225739828 | 0.17888854 |
| 240,000 | 0.225630373 | 0.17881419 |
| 240,100 | 0.225889498 | 0.17881443 |

| | | |
|---|---|---|
| 240,200 | 0.225633938 | 0.17881888 |
| 240,300 | 0.225730387 | 0.17877367 |
| 240,400 | 0.225634214 | 0.17873909 |
| 240,500 | 0.225594958 | 0.17884666 |
| 240,600 | 0.22569959 | 0.17888548 |
| 240,700 | 0.22554018 | 0.17889637 |
| 240,800 | 0.225524611 | 0.17889362 |
| 240,900 | 0.225569169 | 0.17886977 |
| 241,000 | 0.225458862 | 0.17888049 |
| 241,100 | 0.225530835 | 0.17880516 |
| 241,200 | 0.225471147 | 0.17882795 |
| 241,300 | 0.225369069 | 0.17888654 |
| 241,400 | 0.225312427 | 0.17889533 |
| 241,500 | 0.225234208 | 0.17891706 |
| 241,600 | 0.2251917 | 0.17888977 |
| 241,700 | 0.225382375 | 0.17896381 |
| 241,800 | 0.225152952 | 0.17879621 |
| 241,900 | 0.225064035 | 0.1788814 |
| 242,000 | 0.22516691 | 0.17888868 |
| 242,100 | 0.225331657 | 0.1788974 |
| 242,200 | 0.225156675 | 0.17886178 |
| 242,300 | 0.225026775 | 0.17878348 |
| 242,400 | 0.224876962 | 0.17880472 |
| 242,500 | 0.225023329 | 0.17885652 |
| 242,600 | 0.225066833 | 0.17879928 |
| 242,700 | 0.225129457 | 0.17895123 |
| 242,800 | 0.22514967 | 0.17882481 |
| 242,900 | 0.224840988 | 0.17881861 |
| 243,000 | 0.225071286 | 0.17882972 |
| 243,100 | 0.225086245 | 0.17892338 |
| 243,200 | 0.225068402 | 0.17886713 |
| 243,300 | 0.225045646 | 0.17891447 |
| 243,400 | 0.224943178 | 0.17874256 |
| 243,500 | 0.224970549 | 0.17874792 |
| 243,600 | 0.224983053 | 0.17886212 |
| 243,700 | 0.225025706 | 0.1787832 |
| 243,800 | 0.225061498 | 0.17888241 |
| 243,900 | 0.224846507 | 0.17888094 |
| 244,000 | 0.224775508 | 0.17886155 |
| 244,100 | 0.224885495 | 0.17884283 |
| 244,200 | 0.22499351 | 0.17883869 |
| 244,300 | 0.224793929 | 0.17887935 |
| 244,400 | 0.224863886 | 0.17886982 |
| 244,500 | 0.224895442 | 0.17888595 |
| 244,600 | 0.224839636 | 0.17894796 |
| 244,700 | 0.224905008 | 0.17889216 |
| 244,800 | 0.224631178 | 0.17883104 |
| 244,900 | 0.224926983 | 0.17881218 |
| 245,000 | 0.224767096 | 0.1789003 |
| 245,100 | 0.2246869 | 0.17885617 |
| 245,200 | 0.224631904 | 0.17888435 |
| 245,300 | 0.224663978 | 0.17891198 |
| 245,400 | 0.224505249 | 0.17884817 |
| 245,500 | 0.224411573 | 0.17891252 |
| 245,600 | 0.224325119 | 0.17887508 |
| 245,700 | 0.224350454 | 0.17897016 |
| 245,800 | 0.224301546 | 0.17891854 |
| 245,900 | 0.224374359 | 0.17882367 |
| 246,000 | 0.224372723 | 0.17889373 |
| 246,100 | 0.224424217 | 0.17889253 |
| 246,200 | 0.22427521 | 0.17889857 |
| 246,300 | 0.224272773 | 0.17881101 |
| 246,400 | 0.224425875 | 0.17880471 |
| 246,500 | 0.22419324 | 0.17885195 |
| 246,600 | 0.224092888 | 0.17881394 |
| 246,700 | 0.22417485 | 0.1788988 |
| 246,800 | 0.224244211 | 0.17891062 |

| | | |
|---|---|---|
| 246,900 | 0.224062582 | 0.17887262 |
| 247,000 | 0.224181675 | 0.1789199 |
| 247,100 | 0.224010525 | 0.178935 |
| 247,200 | 0.223851234 | 0.17883044 |
| 247,300 | 0.223856452 | 0.17896076 |
| 247,400 | 0.223815432 | 0.17887021 |
| 247,500 | 0.22355981 | 0.17886332 |
| 247,600 | 0.223700987 | 0.17892193 |
| 247,700 | 0.223819904 | 0.17886066 |
| 247,800 | 0.223792813 | 0.17894318 |
| 247,900 | 0.2237443 | 0.17892599 |
| 248,000 | 0.223816764 | 0.17884665 |
| 248,100 | 0.223531688 | 0.17893418 |
| 248,200 | 0.223755164 | 0.1788791 |
| 248,300 | 0.223562262 | 0.17889541 |
| 248,400 | 0.223504845 | 0.17884341 |
| 248,500 | 0.22368305 | 0.17877825 |
| 248,600 | 0.223594277 | 0.17886528 |
| 248,700 | 0.223451172 | 0.17899733 |
| 248,800 | 0.223391078 | 0.17883655 |
| 248,900 | 0.223437374 | 0.17877357 |
| 249,000 | 0.223183289 | 0.17880451 |
| 249,100 | 0.223187425 | 0.17890926 |
| 249,200 | 0.223224303 | 0.17889078 |
| 249,300 | 0.223275722 | 0.17886882 |
| 249,400 | 0.223002622 | 0.17891613 |
| 249,500 | 0.223033103 | 0.17889505 |
| 249,600 | 0.222965387 | 0.17883861 |
| 249,700 | 0.223116098 | 0.17889993 |
| 249,800 | 0.222900318 | 0.17889348 |
| 249,900 | 0.222918381 | 0.17891747 |
| 250,000 | 0.222890506 | 0.17890276 |
| 250,100 | 0.222892243 | 0.17883745 |
| 250,200 | 0.222813265 | 0.17896564 |
| 250,300 | 0.222764449 | 0.17891599 |
| 250,400 | 0.222471704 | 0.17881304 |
| 250,500 | 0.222546369 | 0.17889949 |
| 250,600 | 0.222505076 | 0.17885982 |
| 250,700 | 0.222612565 | 0.1788434 |
| 250,800 | 0.222381623 | 0.17882744 |
| 250,900 | 0.22243634 | 0.17885376 |
| 251,000 | 0.222527929 | 0.17887582 |
| 251,100 | 0.222646647 | 0.17887696 |
| 251,200 | 0.22254908 | 0.17895096 |
| 251,300 | 0.222564947 | 0.17886613 |
| 251,400 | 0.222580491 | 0.17894428 |
| 251,500 | 0.222356793 | 0.17889942 |
| 251,600 | 0.222397398 | 0.17891242 |
| 251,700 | 0.222367173 | 0.17884402 |
| 251,800 | 0.222254742 | 0.17890351 |
| 251,900 | 0.222483178 | 0.17885382 |
| 252,000 | 0.22245028 | 0.17884793 |
| 252,100 | 0.222220029 | 0.17888023 |
| 252,200 | 0.221972626 | 0.17878574 |
| 252,300 | 0.222047872 | 0.17880532 |
| 252,400 | 0.222100662 | 0.17874379 |
| 252,500 | 0.222208734 | 0.17867903 |
| 252,600 | 0.22213295 | 0.17879797 |
| 252,700 | 0.222166074 | 0.17885324 |
| 252,800 | 0.222103898 | 0.17875332 |
| 252,900 | 0.222183082 | 0.17884562 |
| 253,000 | 0.222027039 | 0.17879856 |
| 253,100 | 0.221918275 | 0.17880225 |
| 253,200 | 0.222189378 | 0.1788279 |
| 253,300 | 0.221931125 | 0.17884941 |
| 253,400 | 0.221828304 | 0.178943 |
| 253,500 | 0.221872422 | 0.17881281 |

| | | |
|---|---|---|
| 253,600 | 0.222043906 | 0.17885852 |
| 253,700 | 0.221674075 | 0.17888805 |
| 253,800 | 0.221721386 | 0.17887278 |
| 253,900 | 0.221682239 | 0.17876384 |
| 254,000 | 0.221557596 | 0.17882442 |
| 254,100 | 0.221653205 | 0.17874549 |
| 254,200 | 0.221701027 | 0.17882019 |
| 254,300 | 0.221874562 | 0.17876971 |
| 254,400 | 0.22160596 | 0.17889617 |
| 254,500 | 0.221638697 | 0.17879428 |
| 254,600 | 0.221604744 | 0.17885586 |
| 254,700 | 0.221523588 | 0.17885682 |
| 254,800 | 0.221587899 | 0.17876081 |
| 254,900 | 0.22154007 | 0.17881594 |
| 255,000 | 0.221441262 | 0.17883603 |
| 255,100 | 0.221372977 | 0.17876228 |
| 255,200 | 0.221446212 | 0.17878037 |
| 255,300 | 0.221447657 | 0.17873534 |
| 255,400 | 0.221314499 | 0.17875016 |
| 255,500 | 0.221329153 | 0.17887371 |
| 255,600 | 0.221273582 | 0.17891376 |
| 255,700 | 0.221213303 | 0.17880929 |
| 255,800 | 0.221281006 | 0.17874381 |
| 255,900 | 0.221106187 | 0.17879478 |
| 256,000 | 0.221264105 | 0.17883796 |
| 256,100 | 0.221156039 | 0.17879839 |
| 256,200 | 0.221281715 | 0.17877525 |
| 256,300 | 0.221192258 | 0.17881124 |
| 256,400 | 0.221177033 | 0.17881223 |
| 256,500 | 0.221133203 | 0.17874451 |
| 256,600 | 0.221135181 | 0.1787583 |
| 256,700 | 0.220763093 | 0.17877629 |
| 256,800 | 0.220881143 | 0.17879661 |
| 256,900 | 0.220760219 | 0.1788098 |
| 257,000 | 0.220958404 | 0.17883568 |
| 257,100 | 0.220902736 | 0.17886161 |
| 257,200 | 0.22067932 | 0.17879625 |
| 257,300 | 0.22073258 | 0.1787816 |
| 257,400 | 0.22077252 | 0.17885671 |
| 257,500 | 0.220791251 | 0.17888499 |
| 257,600 | 0.220738548 | 0.17878581 |
| 257,700 | 0.220802484 | 0.17870176 |
| 257,800 | 0.220696075 | 0.17870604 |
| 257,900 | 0.22068764 | 0.17876614 |
| 258,000 | 0.220763857 | 0.17871918 |
| 258,100 | 0.220682441 | 0.17878608 |
| 258,200 | 0.220824268 | 0.17884147 |
| 258,300 | 0.220590927 | 0.17873918 |
| 258,400 | 0.220637607 | 0.17866971 |
| 258,500 | 0.220509017 | 0.17870235 |
| 258,600 | 0.220371149 | 0.17875553 |
| 258,700 | 0.220452173 | 0.17868454 |
| 258,800 | 0.220320271 | 0.17863942 |
| 258,900 | 0.220343704 | 0.17878618 |
| 259,000 | 0.220315187 | 0.17873944 |
| 259,100 | 0.22036198 | 0.17874785 |
| 259,200 | 0.220197853 | 0.17877953 |
| 259,300 | 0.220310835 | 0.17875816 |
| 259,400 | 0.220126084 | 0.17875636 |
| 259,500 | 0.22007049 | 0.17879835 |
| 259,600 | 0.22033009 | 0.17879185 |
| 259,700 | 0.220222096 | 0.17872984 |
| 259,800 | 0.220155586 | 0.17874292 |
| 259,900 | 0.220233396 | 0.17874135 |
| 260,000 | 0.220170602 | 0.17869492 |
| 260,100 | 0.21992359 | 0.17863736 |
| 260,200 | 0.219913228 | 0.17865844 |

| | | |
|---|---|---|
| 260,300 | 0.219724951 | 0.17874377 |
| 260,400 | 0.219808306 | 0.17868285 |
| 260,500 | 0.21990355 | 0.17871985 |
| 260,600 | 0.219792469 | 0.17875869 |
| 260,700 | 0.219632946 | 0.17883819 |
| 260,800 | 0.219590351 | 0.1787699 |
| 260,900 | 0.219662465 | 0.17875806 |
| 261,000 | 0.219708211 | 0.17873648 |
| 261,100 | 0.219598462 | 0.17883539 |
| 261,200 | 0.219658403 | 0.17880614 |
| 261,300 | 0.219621868 | 0.17878266 |
| 261,400 | 0.219493499 | 0.17877281 |
| 261,500 | 0.219541777 | 0.178832 |
| 261,600 | 0.219516104 | 0.17878492 |
| 261,700 | 0.219432331 | 0.17874214 |
| 261,800 | 0.219419529 | 0.17868331 |
| 261,900 | 0.219302188 | 0.17870451 |
| 262,000 | 0.219460719 | 0.17873701 |
| 262,100 | 0.219539276 | 0.17865437 |
| 262,200 | 0.219410252 | 0.17865955 |
| 262,300 | 0.219330542 | 0.17865405 |
| 262,400 | 0.219374918 | 0.1786774 |
| 262,500 | 0.219360143 | 0.17873118 |
| 262,600 | 0.219268603 | 0.17874696 |
| 262,700 | 0.219214134 | 0.17881748 |
| 262,800 | 0.21909317 | 0.17869677 |
| 262,900 | 0.219104952 | 0.17875151 |
| 263,000 | 0.219096566 | 0.17872707 |
| 263,100 | 0.218983883 | 0.17879292 |
| 263,200 | 0.219010281 | 0.17873267 |
| 263,300 | 0.218948545 | 0.17871895 |
| 263,400 | 0.219036323 | 0.17879264 |
| 263,500 | 0.219024219 | 0.17881559 |
| 263,600 | 0.219035047 | 0.17880545 |
| 263,700 | 0.219021361 | 0.17868996 |
| 263,800 | 0.219132747 | 0.17878983 |
| 263,900 | 0.218920532 | 0.1787399 |
| 264,000 | 0.219020414 | 0.17871008 |
| 264,100 | 0.219059547 | 0.17880735 |
| 264,200 | 0.218902474 | 0.17877598 |
| 264,300 | 0.218972538 | 0.17879309 |
| 264,400 | 0.218788506 | 0.17879023 |
| 264,500 | 0.21887765 | 0.17875714 |
| 264,600 | 0.218798881 | 0.17882289 |
| 264,700 | 0.21876167 | 0.17874323 |
| 264,800 | 0.218771545 | 0.17883399 |
| 264,900 | 0.218496154 | 0.1787616 |
| 265,000 | 0.218376069 | 0.17875551 |
| 265,100 | 0.218622903 | 0.17869491 |
| 265,200 | 0.218390043 | 0.17868766 |
| 265,300 | 0.218436696 | 0.17883306 |
| 265,400 | 0.21840214 | 0.17875288 |
| 265,500 | 0.21847705 | 0.17888285 |
| 265,600 | 0.218450692 | 0.17868368 |
| 265,700 | 0.218401414 | 0.17874389 |
| 265,800 | 0.218236861 | 0.17873663 |
| 265,900 | 0.218091952 | 0.17884243 |
| 266,000 | 0.218106535 | 0.17885491 |
| 266,100 | 0.218320905 | 0.1787954 |
| 266,200 | 0.218226865 | 0.17884524 |
| 266,300 | 0.218132479 | 0.17884481 |
| 266,400 | 0.218180358 | 0.17880138 |
| 266,500 | 0.218114085 | 0.17871932 |
| 266,600 | 0.217972135 | 0.17873651 |
| 266,700 | 0.217851846 | 0.17872374 |
| 266,800 | 0.217952469 | 0.17880053 |
| 266,900 | 0.218041553 | 0.17878143 |

| | | |
|---|---|---|
| 267,000 | 0.217902466 | 0.17869971 |
| 267,100 | 0.217825467 | 0.17873371 |
| 267,200 | 0.217953307 | 0.17874138 |
| 267,300 | 0.217862927 | 0.17878316 |
| 267,400 | 0.217947954 | 0.17872295 |
| 267,500 | 0.218035497 | 0.17874969 |
| 267,600 | 0.217823923 | 0.17878 |
| 267,700 | 0.21782393 | 0.17874754 |
| 267,800 | 0.21770765 | 0.17870758 |
| 267,900 | 0.217797483 | 0.178803 |
| 268,000 | 0.21770406 | 0.17884286 |
| 268,100 | 0.217714474 | 0.17879524 |
| 268,200 | 0.217603083 | 0.17875556 |
| 268,300 | 0.217739577 | 0.17870177 |
| 268,400 | 0.217598685 | 0.17882902 |
| 268,500 | 0.217824624 | 0.17873184 |
| 268,600 | 0.217938309 | 0.17871687 |
| 268,700 | 0.217624968 | 0.17864813 |
| 268,800 | 0.217896996 | 0.178788 |
| 268,900 | 0.217584809 | 0.1788534 |
| 269,000 | 0.217765372 | 0.17876953 |
| 269,100 | 0.217685184 | 0.17870254 |
| 269,200 | 0.217568114 | 0.17874212 |
| 269,300 | 0.217729074 | 0.17876762 |
| 269,400 | 0.21749852 | 0.17872924 |
| 269,500 | 0.217592843 | 0.17876824 |
| 269,600 | 0.217557161 | 0.17879165 |
| 269,700 | 0.217743201 | 0.17870453 |
| 269,800 | 0.217478318 | 0.17870726 |
| 269,900 | 0.217464914 | 0.17864272 |
| 270,000 | 0.217366431 | 0.1787241 |
| 270,100 | 0.217235487 | 0.1787107 |
| 270,200 | 0.217381765 | 0.17861357 |
| 270,300 | 0.217373605 | 0.17872181 |
| 270,400 | 0.217277016 | 0.17866261 |
| 270,500 | 0.217178844 | 0.17867286 |
| 270,600 | 0.217062765 | 0.17879373 |
| 270,700 | 0.21693677 | 0.17869235 |
| 270,800 | 0.217224111 | 0.17869317 |
| 270,900 | 0.217156088 | 0.17870489 |
| 271,000 | 0.217094303 | 0.17877886 |
| 271,100 | 0.217231678 | 0.1786969 |
| 271,200 | 0.217153237 | 0.17873742 |
| 271,300 | 0.217120531 | 0.1787284 |
| 271,400 | 0.217065695 | 0.17869364 |
| 271,500 | 0.217146507 | 0.17869729 |
| 271,600 | 0.217171882 | 0.17872385 |
| 271,700 | 0.217017543 | 0.17873944 |
| 271,800 | 0.217080293 | 0.17884662 |
| 271,900 | 0.217218116 | 0.17877028 |
| 272,000 | 0.217158864 | 0.17885758 |
| 272,100 | 0.216916893 | 0.17875825 |
| 272,200 | 0.216806942 | 0.17877432 |
| 272,300 | 0.216822181 | 0.1787481 |
| 272,400 | 0.216927789 | 0.17869946 |
| 272,500 | 0.216893314 | 0.17871848 |
| 272,600 | 0.216892244 | 0.1786814 |
| 272,700 | 0.216860019 | 0.17865063 |
| 272,800 | 0.216933481 | 0.17873905 |
| 272,900 | 0.21677317 | 0.17867117 |
| 273,000 | 0.216874698 | 0.17870514 |
| 273,100 | 0.216753728 | 0.17869979 |
| 273,200 | 0.216864652 | 0.1786559 |
| 273,300 | 0.216911938 | 0.17872199 |
| 273,400 | 0.216917638 | 0.17865121 |
| 273,500 | 0.216759214 | 0.17873726 |
| 273,600 | 0.216743826 | 0.17868677 |

| | | |
|---|---|---|
| 273,700 | 0.216600123 | 0.17867985 |
| 273,800 | 0.216686832 | 0.17873386 |
| 273,900 | 0.216670395 | 0.17875484 |
| 274,000 | 0.216577738 | 0.17863965 |
| 274,100 | 0.216630913 | 0.17876874 |
| 274,200 | 0.21656677 | 0.178633 |
| 274,300 | 0.216510477 | 0.17873123 |
| 274,400 | 0.216636339 | 0.17875716 |
| 274,500 | 0.216474505 | 0.1786152 |
| 274,600 | 0.216659966 | 0.17870195 |
| 274,700 | 0.216463671 | 0.17867559 |
| 274,800 | 0.216516504 | 0.17871307 |
| 274,900 | 0.216461897 | 0.17867488 |
| 275,000 | 0.216570016 | 0.17869882 |
| 275,100 | 0.216572148 | 0.17864753 |
| 275,200 | 0.216613344 | 0.1786162 |
| 275,300 | 0.216612513 | 0.17865801 |
| 275,400 | 0.21646909 | 0.17869441 |
| 275,500 | 0.216371362 | 0.17865046 |
| 275,600 | 0.216337357 | 0.17867319 |
| 275,700 | 0.216485577 | 0.17869943 |
| 275,800 | 0.216251895 | 0.1786122 |
| 275,900 | 0.216141579 | 0.17870658 |
| 276,000 | 0.216217195 | 0.17870978 |
| 276,100 | 0.216161522 | 0.17874326 |
| 276,200 | 0.216159924 | 0.17869164 |
| 276,300 | 0.216007061 | 0.17869098 |
| 276,400 | 0.216213833 | 0.17866209 |
| 276,500 | 0.216039731 | 0.17855416 |
| 276,600 | 0.216093561 | 0.17863727 |
| 276,700 | 0.215992143 | 0.17858646 |
| 276,800 | 0.21609579 | 0.17863159 |
| 276,900 | 0.216027377 | 0.17865067 |
| 277,000 | 0.216171144 | 0.17873031 |
| 277,100 | 0.215973302 | 0.17864865 |
| 277,200 | 0.216112842 | 0.17866669 |
| 277,300 | 0.216070659 | 0.17861714 |
| 277,400 | 0.215924842 | 0.17866953 |
| 277,500 | 0.215943037 | 0.17861439 |
| 277,600 | 0.215754524 | 0.17867234 |
| 277,700 | 0.215782141 | 0.17875666 |
| 277,800 | 0.215891046 | 0.1787166 |
| 277,900 | 0.215705447 | 0.17868511 |
| 278,000 | 0.215891775 | 0.17867238 |
| 278,100 | 0.215777862 | 0.17864863 |
| 278,200 | 0.215776832 | 0.17864191 |
| 278,300 | 0.215712626 | 0.17866163 |
| 278,400 | 0.215621023 | 0.178704 |
| 278,500 | 0.21555511 | 0.17880326 |
| 278,600 | 0.215489057 | 0.17872498 |
| 278,700 | 0.215482134 | 0.17873281 |
| 278,800 | 0.215664799 | 0.17861054 |
| 278,900 | 0.215396449 | 0.17869163 |
| 279,000 | 0.215431565 | 0.17865821 |
| 279,100 | 0.21540871 | 0.17875504 |
| 279,200 | 0.215282055 | 0.17871652 |
| 279,300 | 0.215416405 | 0.178716 |
| 279,400 | 0.215379199 | 0.17866625 |
| 279,500 | 0.215313965 | 0.1787439 |
| 279,600 | 0.215316487 | 0.17871207 |
| 279,700 | 0.215280661 | 0.17879147 |
| 279,800 | 0.215179507 | 0.17878633 |
| 279,900 | 0.21519531 | 0.17864337 |
| 280,000 | 0.215257171 | 0.1787744 |
| 280,100 | 0.215217478 | 0.17880129 |
| 280,200 | 0.215278736 | 0.17876827 |
| 280,300 | 0.215207302 | 0.17873998 |

| | | |
|---|---|---|
| 280,400 | 0.215050363 | 0.17872581 |
| 280,500 | 0.215033777 | 0.17874614 |
| 280,600 | 0.214985149 | 0.1787701 |
| 280,700 | 0.215090283 | 0.17861288 |
| 280,800 | 0.214937794 | 0.1787139 |
| 280,900 | 0.214775621 | 0.178723 |
| 281,000 | 0.214940155 | 0.17872251 |
| 281,100 | 0.214834146 | 0.17861365 |
| 281,200 | 0.214779322 | 0.17867727 |
| 281,300 | 0.214816004 | 0.1787325 |
| 281,400 | 0.214817214 | 0.17872177 |
| 281,500 | 0.214905401 | 0.17868121 |
| 281,600 | 0.214963373 | 0.17868891 |
| 281,700 | 0.214667704 | 0.17864064 |
| 281,800 | 0.214815029 | 0.17873484 |
| 281,900 | 0.21463118 | 0.17869223 |
| 282,000 | 0.214730636 | 0.17871056 |
| 282,100 | 0.214654512 | 0.17870342 |
| 282,200 | 0.214728119 | 0.17875961 |
| 282,300 | 0.214611426 | 0.17868982 |
| 282,400 | 0.214809415 | 0.17873163 |
| 282,500 | 0.214742409 | 0.17861853 |
| 282,600 | 0.214604052 | 0.17864496 |
| 282,700 | 0.214818123 | 0.17863791 |
| 282,800 | 0.214576211 | 0.1786217 |
| 282,900 | 0.21458528 | 0.17868769 |
| 283,000 | 0.214668153 | 0.17863093 |
| 283,100 | 0.214496221 | 0.17866812 |
| 283,200 | 0.214340814 | 0.17861778 |
| 283,300 | 0.21452333 | 0.1787079 |
| 283,400 | 0.214366201 | 0.17867871 |
| 283,500 | 0.214455322 | 0.17873077 |
| 283,600 | 0.214307147 | 0.17864791 |
| 283,700 | 0.214339246 | 0.17868906 |
| 283,800 | 0.214233371 | 0.17861915 |
| 283,900 | 0.214192153 | 0.17857115 |
| 284,000 | 0.214311331 | 0.17861948 |
| 284,100 | 0.214256781 | 0.17854154 |
| 284,200 | 0.214180664 | 0.17864204 |
| 284,300 | 0.214262209 | 0.17868338 |
| 284,400 | 0.214395014 | 0.17859941 |
| 284,500 | 0.214259289 | 0.17861822 |
| 284,600 | 0.214212415 | 0.17854804 |
| 284,700 | 0.214191407 | 0.17865136 |
| 284,800 | 0.214159355 | 0.17866306 |
| 284,900 | 0.214084931 | 0.17861994 |
| 285,000 | 0.214051949 | 0.17870215 |
| 285,100 | 0.214002691 | 0.17863478 |
| 285,200 | 0.21400624 | 0.17867905 |
| 285,300 | 0.213925464 | 0.17862208 |
| 285,400 | 0.21387401 | 0.17867974 |
| 285,500 | 0.213907831 | 0.17864992 |
| 285,600 | 0.213922249 | 0.17874262 |
| 285,700 | 0.213819819 | 0.17866524 |
| 285,800 | 0.213963968 | 0.17866236 |
| 285,900 | 0.213802173 | 0.17862632 |
| 286,000 | 0.21369998 | 0.17860664 |
| 286,100 | 0.213773216 | 0.17864445 |
| 286,200 | 0.213716229 | 0.17864449 |
| 286,300 | 0.213682621 | 0.17858116 |
| 286,400 | 0.213632913 | 0.17867798 |
| 286,500 | 0.21365092 | 0.17856871 |
| 286,600 | 0.213508781 | 0.17865356 |
| 286,700 | 0.213620049 | 0.17870126 |
| 286,800 | 0.21369668 | 0.17860937 |
| 286,900 | 0.213553196 | 0.17857016 |
| 287,000 | 0.21352293 | 0.17862789 |

| | | |
|---|---|---|
| 287,100 | 0.213387499 | 0.17865415 |
| 287,200 | 0.213534359 | 0.17857364 |
| 287,300 | 0.213519541 | 0.17860993 |
| 287,400 | 0.213484843 | 0.17856818 |
| 287,500 | 0.213367698 | 0.17869273 |
| 287,600 | 0.213177049 | 0.17869377 |
| 287,700 | 0.213181191 | 0.17862026 |
| 287,800 | 0.212928391 | 0.17858222 |
| 287,900 | 0.213083883 | 0.17867182 |
| 288,000 | 0.213135394 | 0.17868073 |
| 288,100 | 0.213088169 | 0.17873005 |
| 288,200 | 0.212962991 | 0.1785223 |
| 288,300 | 0.212870925 | 0.17863043 |
| 288,400 | 0.2127386 | 0.17857317 |
| 288,500 | 0.212828468 | 0.17860827 |
| 288,600 | 0.212861965 | 0.17864458 |
| 288,700 | 0.212836938 | 0.17855434 |
| 288,800 | 0.212820549 | 0.17861355 |
| 288,900 | 0.212749755 | 0.17860312 |
| 289,000 | 0.212832169 | 0.1786338 |
| 289,100 | 0.212759897 | 0.17853941 |
| 289,200 | 0.212616289 | 0.17869111 |
| 289,300 | 0.212733118 | 0.17863289 |
| 289,400 | 0.212763074 | 0.17866993 |
| 289,500 | 0.212788289 | 0.17864716 |
| 289,600 | 0.212655137 | 0.17865351 |
| 289,700 | 0.2125509 | 0.17862248 |
| 289,800 | 0.212679316 | 0.17862733 |
| 289,900 | 0.212631737 | 0.17860179 |
| 290,000 | 0.212612418 | 0.17849681 |
| 290,100 | 0.212528721 | 0.17870398 |
| 290,200 | 0.212705641 | 0.17870568 |
| 290,300 | 0.212442213 | 0.17871292 |
| 290,400 | 0.212527455 | 0.17868058 |
| 290,500 | 0.212427719 | 0.17870168 |
| 290,600 | 0.212426114 | 0.17866007 |
| 290,700 | 0.212456942 | 0.17871526 |
| 290,800 | 0.212362986 | 0.17864889 |
| 290,900 | 0.212321641 | 0.17866374 |
| 291,000 | 0.212390398 | 0.17862487 |
| 291,100 | 0.212470389 | 0.17863149 |
| 291,200 | 0.212456586 | 0.17872234 |
| 291,300 | 0.212463704 | 0.1786936 |
| 291,400 | 0.212408965 | 0.17871078 |
| 291,500 | 0.212290024 | 0.17862967 |
| 291,600 | 0.212249831 | 0.17864138 |
| 291,700 | 0.212333393 | 0.17862962 |
| 291,800 | 0.212344029 | 0.17865215 |
| 291,900 | 0.212268274 | 0.17871912 |
| 292,000 | 0.212239977 | 0.17868727 |
| 292,100 | 0.212107041 | 0.17860465 |
| 292,200 | 0.212324691 | 0.17865545 |
| 292,300 | 0.212238879 | 0.17858886 |
| 292,400 | 0.212241965 | 0.17853012 |
| 292,500 | 0.211982382 | 0.17857644 |
| 292,600 | 0.212067785 | 0.17853647 |
| 292,700 | 0.21193535 | 0.17857273 |
| 292,800 | 0.211964155 | 0.17863671 |
| 292,900 | 0.211921243 | 0.17858427 |
| 293,000 | 0.211942697 | 0.17864795 |
| 293,100 | 0.211701738 | 0.17858811 |
| 293,200 | 0.211832711 | 0.17858041 |
| 293,300 | 0.211867346 | 0.17860303 |
| 293,400 | 0.211811757 | 0.17859296 |
| 293,500 | 0.211854624 | 0.17854517 |
| 293,600 | 0.211752818 | 0.17867558 |
| 293,700 | 0.211831388 | 0.17858915 |

| | | |
|---|---|---|
| 293,800 | 0.211612786 | 0.17868809 |
| 293,900 | 0.211698028 | 0.17865333 |
| 294,000 | 0.211737253 | 0.178562 |
| 294,100 | 0.211623662 | 0.17854443 |
| 294,200 | 0.211713276 | 0.17860883 |
| 294,300 | 0.211768755 | 0.1786119 |
| 294,400 | 0.211593079 | 0.17858348 |
| 294,500 | 0.211749891 | 0.17855979 |
| 294,600 | 0.211494005 | 0.17870002 |
| 294,700 | 0.211562865 | 0.17865611 |
| 294,800 | 0.211411877 | 0.17860341 |
| 294,900 | 0.211425509 | 0.17863259 |
| 295,000 | 0.211385015 | 0.17858106 |
| 295,100 | 0.211557403 | 0.17862514 |
| 295,200 | 0.21140986 | 0.17851208 |
| 295,300 | 0.21146595 | 0.17861451 |
| 295,400 | 0.211447025 | 0.17857712 |
| 295,500 | 0.211304962 | 0.17862038 |
| 295,600 | 0.211372893 | 0.17862424 |
| 295,700 | 0.211236251 | 0.1785619 |
| 295,800 | 0.211306224 | 0.17859078 |
| 295,900 | 0.211349849 | 0.17854462 |
| 296,000 | 0.211234808 | 0.17850701 |
| 296,100 | 0.211287043 | 0.17850878 |
| 296,200 | 0.211129614 | 0.17852376 |
| 296,300 | 0.211280442 | 0.17852778 |
| 296,400 | 0.211212158 | 0.17857561 |
| 296,500 | 0.211056224 | 0.17852377 |
| 296,600 | 0.211098474 | 0.178561 |
| 296,700 | 0.210990364 | 0.17858985 |
| 296,800 | 0.211156124 | 0.17852036 |
| 296,900 | 0.211114513 | 0.1785583 |
| 297,000 | 0.211031323 | 0.17851066 |
| 297,100 | 0.210978148 | 0.17865042 |
| 297,200 | 0.211133858 | 0.17862373 |
| 297,300 | 0.211043342 | 0.17854831 |
| 297,400 | 0.210957687 | 0.17859279 |
| 297,500 | 0.210954545 | 0.17851946 |
| 297,600 | 0.211077894 | 0.17860439 |
| 297,700 | 0.210916646 | 0.17858836 |
| 297,800 | 0.210883677 | 0.17857443 |
| 297,900 | 0.210795915 | 0.1785419 |
| 298,000 | 0.210794217 | 0.17852608 |
| 298,100 | 0.210810191 | 0.17850751 |
| 298,200 | 0.210766203 | 0.17861527 |
| 298,300 | 0.210771361 | 0.17862482 |
| 298,400 | 0.21076352 | 0.17858857 |
| 298,500 | 0.210682191 | 0.17857962 |
| 298,600 | 0.210717115 | 0.17866473 |
| 298,700 | 0.21074013 | 0.17865217 |
| 298,800 | 0.21055945 | 0.17861741 |
| 298,900 | 0.210691394 | 0.17867575 |
| 299,000 | 0.210500245 | 0.17862928 |
| 299,100 | 0.210535685 | 0.17858358 |
| 299,200 | 0.210426035 | 0.17867329 |
| 299,300 | 0.21040601 | 0.17865635 |
| 299,400 | 0.210442906 | 0.17860973 |
| 299,500 | 0.210307345 | 0.17858715 |
| 299,600 | 0.210188485 | 0.1786368 |
| 299,700 | 0.210155578 | 0.178675 |
| 299,800 | 0.210177054 | 0.17866808 |
| 299,900 | 0.210113502 | 0.17861273 |
| 300,000 | 0.210086791 | 0.17863182 |
| 300,100 | 0.210251107 | 0.17861054 |
| 300,200 | 0.210038547 | 0.17862486 |
| 300,300 | 0.21004792 | 0.17861498 |
| 300,400 | 0.209998884 | 0.17863353 |

| | | |
|---|---|---|
| 300,500 | 0.210015785 | 0.17859892 |
| 300,600 | 0.21001677 | 0.17852307 |
| 300,700 | 0.210038855 | 0.17856894 |
| 300,800 | 0.209891869 | 0.17859162 |
| 300,900 | 0.209980118 | 0.1786074 |
| 301,000 | 0.209923667 | 0.1786047 |
| 301,100 | 0.209890004 | 0.17866603 |
| 301,200 | 0.209613891 | 0.17873114 |
| 301,300 | 0.209870452 | 0.17860742 |
| 301,400 | 0.209888343 | 0.17861922 |
| 301,500 | 0.209839737 | 0.17864646 |
| 301,600 | 0.209794275 | 0.17852723 |
| 301,700 | 0.209638943 | 0.17859418 |
| 301,800 | 0.209656439 | 0.17854031 |
| 301,900 | 0.209764047 | 0.17863875 |
| 302,000 | 0.209593784 | 0.17869077 |
| 302,100 | 0.209588913 | 0.17861211 |
| 302,200 | 0.209518768 | 0.17863511 |
| 302,300 | 0.209539543 | 0.17855831 |
| 302,400 | 0.209521699 | 0.17862838 |
| 302,500 | 0.209563343 | 0.17862879 |
| 302,600 | 0.209490929 | 0.17856823 |
| 302,700 | 0.209335167 | 0.17850699 |
| 302,800 | 0.209376081 | 0.17853183 |
| 302,900 | 0.20938402 | 0.17855959 |
| 303,000 | 0.209334382 | 0.17857414 |
| 303,100 | 0.209370197 | 0.17869325 |
| 303,200 | 0.209377515 | 0.17856744 |
| 303,300 | 0.209460404 | 0.17857032 |
| 303,400 | 0.209284393 | 0.17852583 |
| 303,500 | 0.209409901 | 0.17851531 |
| 303,600 | 0.209279846 | 0.17856839 |
| 303,700 | 0.209529211 | 0.17853207 |
| 303,800 | 0.209383562 | 0.1785835 |
| 303,900 | 0.209485187 | 0.1786063 |
| 304,000 | 0.209258951 | 0.1784965 |
| 304,100 | 0.209362232 | 0.17854802 |
| 304,200 | 0.209439702 | 0.17855089 |
| 304,300 | 0.209201646 | 0.17859337 |
| 304,400 | 0.209358992 | 0.1785432 |
| 304,500 | 0.209213771 | 0.17851734 |
| 304,600 | 0.209312664 | 0.17852995 |
| 304,700 | 0.209153373 | 0.17858242 |
| 304,800 | 0.209184257 | 0.17850767 |
| 304,900 | 0.209136365 | 0.17849962 |
| 305,000 | 0.208945952 | 0.17858294 |
| 305,100 | 0.208960639 | 0.17856955 |
| 305,200 | 0.209071565 | 0.17852905 |
| 305,300 | 0.208926173 | 0.17858205 |
| 305,400 | 0.209023619 | 0.17854544 |
| 305,500 | 0.20879935 | 0.17850692 |
| 305,600 | 0.208823752 | 0.1785114 |
| 305,700 | 0.208917956 | 0.17851335 |
| 305,800 | 0.208953481 | 0.17865745 |
| 305,900 | 0.209093249 | 0.17860442 |
| 306,000 | 0.208871889 | 0.17855437 |
| 306,100 | 0.208898803 | 0.17867292 |
| 306,200 | 0.208932218 | 0.17851239 |
| 306,300 | 0.20876944 | 0.17851798 |
| 306,400 | 0.208860396 | 0.17856976 |
| 306,500 | 0.208840945 | 0.17849752 |
| 306,600 | 0.208838555 | 0.1785742 |
| 306,700 | 0.208837286 | 0.17854902 |
| 306,800 | 0.208739902 | 0.1785641 |
| 306,900 | 0.208777038 | 0.17862 |
| 307,000 | 0.208690471 | 0.17857801 |
| 307,100 | 0.208928088 | 0.17864374 |

| | | |
|---|---|---|
| 307,200 | 0.208782748 | 0.17860859 |
| 307,300 | 0.208608982 | 0.178607 |
| 307,400 | 0.208880202 | 0.1785457 |
| 307,500 | 0.208762137 | 0.17864449 |
| 307,600 | 0.208774234 | 0.17858779 |
| 307,700 | 0.208998338 | 0.17864475 |
| 307,800 | 0.208697969 | 0.17859227 |
| 307,900 | 0.208744978 | 0.17862325 |
| 308,000 | 0.208774943 | 0.17861456 |
| 308,100 | 0.208714364 | 0.17861694 |
| 308,200 | 0.20878931 | 0.17857862 |
| 308,300 | 0.208717108 | 0.17866213 |
| 308,400 | 0.208694017 | 0.17859688 |
| 308,500 | 0.208868111 | 0.17857267 |
| 308,600 | 0.208761185 | 0.17858037 |
| 308,700 | 0.208779482 | 0.1785527 |
| 308,800 | 0.208838085 | 0.17854495 |
| 308,900 | 0.208657481 | 0.17849677 |
| 309,000 | 0.208524577 | 0.17843635 |
| 309,100 | 0.208641995 | 0.17862911 |
| 309,200 | 0.20877076 | 0.17854899 |
| 309,300 | 0.208629804 | 0.17858583 |
| 309,400 | 0.208463743 | 0.17844166 |
| 309,500 | 0.208497997 | 0.1785732 |
| 309,600 | 0.208539206 | 0.17857518 |
| 309,700 | 0.208504281 | 0.17853919 |
| 309,800 | 0.208409768 | 0.17861068 |
| 309,900 | 0.208446931 | 0.17857606 |
| 310,000 | 0.208373442 | 0.17851959 |
| 310,100 | 0.20829935 | 0.17862788 |
| 310,200 | 0.208194706 | 0.17855022 |
| 310,300 | 0.208298578 | 0.17865498 |
| 310,400 | 0.208244087 | 0.17859083 |
| 310,500 | 0.208248976 | 0.17855146 |
| 310,600 | 0.208182724 | 0.17868071 |
| 310,700 | 0.20835388 | 0.17858711 |
| 310,800 | 0.208250524 | 0.17853087 |
| 310,900 | 0.208261369 | 0.17851695 |
| 311,000 | 0.208184187 | 0.17861471 |
| 311,100 | 0.208292255 | 0.17855562 |
| 311,200 | 0.208077622 | 0.17861141 |
| 311,300 | 0.208113441 | 0.1785743 |
| 311,400 | 0.208069136 | 0.17850633 |
| 311,500 | 0.208085901 | 0.17859223 |
| 311,600 | 0.208033769 | 0.1785738 |
| 311,700 | 0.208075039 | 0.17858579 |
| 311,800 | 0.208154382 | 0.17853789 |
| 311,900 | 0.208109514 | 0.17855513 |
| 312,000 | 0.207832524 | 0.17864141 |
| 312,100 | 0.208006706 | 0.17867869 |
| 312,200 | 0.207851227 | 0.1786163 |
| 312,300 | 0.207821723 | 0.17861901 |
| 312,400 | 0.207610158 | 0.17860902 |
| 312,500 | 0.207656479 | 0.17856881 |
| 312,600 | 0.20757887 | 0.17856326 |
| 312,700 | 0.207542007 | 0.17853927 |
| 312,800 | 0.207711233 | 0.17852182 |
| 312,900 | 0.207800134 | 0.17864582 |
| 313,000 | 0.20754646 | 0.17849708 |
| 313,100 | 0.20777243 | 0.17857838 |
| 313,200 | 0.207702953 | 0.17852917 |
| 313,300 | 0.207643762 | 0.17856808 |
| 313,400 | 0.207696262 | 0.17856537 |
| 313,500 | 0.207567197 | 0.17854604 |
| 313,600 | 0.207620006 | 0.1785716 |
| 313,700 | 0.207666206 | 0.17861182 |
| 313,800 | 0.207312461 | 0.17859601 |

| | | |
|---|---|---|
| 313,900 | 0.207409284 | 0.1785791 |
| 314,000 | 0.207268445 | 0.17864874 |
| 314,100 | 0.207363136 | 0.17866682 |
| 314,200 | 0.207310901 | 0.17861516 |
| 314,300 | 0.207288193 | 0.17853545 |
| 314,400 | 0.207309165 | 0.17863259 |
| 314,500 | 0.207265009 | 0.17855053 |
| 314,600 | 0.207133413 | 0.17855849 |
| 314,700 | 0.207219109 | 0.17860416 |
| 314,800 | 0.207226228 | 0.17851476 |
| 314,900 | 0.207221363 | 0.17852807 |
| 315,000 | 0.206982502 | 0.17852271 |
| 315,100 | 0.207037607 | 0.17847364 |
| 315,200 | 0.20713063 | 0.17856316 |
| 315,300 | 0.206982427 | 0.17857008 |
| 315,400 | 0.206859484 | 0.17855225 |
| 315,500 | 0.206807347 | 0.1786301 |
| 315,600 | 0.206810949 | 0.17861744 |
| 315,700 | 0.206807631 | 0.17858313 |
| 315,800 | 0.20678082 | 0.17866595 |
| 315,900 | 0.206694305 | 0.17862451 |
| 316,000 | 0.206672397 | 0.17859214 |
| 316,100 | 0.206866088 | 0.17858326 |
| 316,200 | 0.206786588 | 0.17858129 |
| 316,300 | 0.206737754 | 0.17856292 |
| 316,400 | 0.206813805 | 0.17863527 |
| 316,500 | 0.206844286 | 0.17853073 |
| 316,600 | 0.206755467 | 0.17854167 |
| 316,700 | 0.206802891 | 0.17852253 |
| 316,800 | 0.206687661 | 0.17853665 |
| 316,900 | 0.206696938 | 0.17848925 |
| 317,000 | 0.206809524 | 0.17859567 |
| 317,100 | 0.206623712 | 0.1785225 |
| 317,200 | 0.206594575 | 0.17854313 |
| 317,300 | 0.206585425 | 0.17854685 |
| 317,400 | 0.206382619 | 0.17864893 |
| 317,500 | 0.206564331 | 0.17849844 |
| 317,600 | 0.20651502 | 0.17857768 |
| 317,700 | 0.206599622 | 0.17868757 |
| 317,800 | 0.206548003 | 0.17859908 |
| 317,900 | 0.206457133 | 0.17864552 |
| 318,000 | 0.206376409 | 0.17860479 |
| 318,100 | 0.206292839 | 0.17864374 |
| 318,200 | 0.206415502 | 0.17862134 |
| 318,300 | 0.20623864 | 0.17857106 |
| 318,400 | 0.206314607 | 0.17859603 |
| 318,500 | 0.206102359 | 0.17864094 |
| 318,600 | 0.206168446 | 0.17857679 |
| 318,700 | 0.206192931 | 0.17870342 |
| 318,800 | 0.206249736 | 0.17863701 |
| 318,900 | 0.206312558 | 0.17855076 |
| 319,000 | 0.206205356 | 0.17865333 |
| 319,100 | 0.206270874 | 0.17853398 |
| 319,200 | 0.206220074 | 0.17850683 |
| 319,300 | 0.20616908 | 0.17859142 |
| 319,400 | 0.206095183 | 0.17852543 |
| 319,500 | 0.206145975 | 0.17867736 |
| 319,600 | 0.205934644 | 0.17860174 |
| 319,700 | 0.206037766 | 0.1785914 |
| 319,800 | 0.206134828 | 0.17847565 |
| 319,900 | 0.206056229 | 0.17849799 |
| 320,000 | 0.20608006 | 0.17858815 |
| 320,100 | 0.205871857 | 0.17866262 |
| 320,200 | 0.205894825 | 0.17854651 |
| 320,300 | 0.205987989 | 0.17863849 |
| 320,400 | 0.20594766 | 0.17869507 |
| 320,500 | 0.205900557 | 0.1786511 |

| | | |
|---|---|---|
| 320,600 | 0.205735745 | 0.17866511 |
| 320,700 | 0.20594385 | 0.17858258 |
| 320,800 | 0.205686654 | 0.1786404 |
| 320,900 | 0.205768771 | 0.17863049 |
| 321,000 | 0.205828645 | 0.17865635 |
| 321,100 | 0.20586764 | 0.17861586 |
| 321,200 | 0.206010338 | 0.17859692 |
| 321,300 | 0.205839828 | 0.17864634 |
| 321,400 | 0.205744637 | 0.17855198 |
| 321,500 | 0.20577102 | 0.17859504 |
| 321,600 | 0.205772236 | 0.17864655 |
| 321,700 | 0.205614759 | 0.17858587 |
| 321,800 | 0.205717184 | 0.17856795 |
| 321,900 | 0.205748766 | 0.17862283 |
| 322,000 | 0.205701858 | 0.17865065 |
| 322,100 | 0.205556931 | 0.1786674 |
| 322,200 | 0.205685054 | 0.17866498 |
| 322,300 | 0.205651816 | 0.17855763 |
| 322,400 | 0.20554043 | 0.17861794 |
| 322,500 | 0.205624535 | 0.17851578 |
| 322,600 | 0.205549045 | 0.17851181 |
| 322,700 | 0.205432633 | 0.1785227 |
| 322,800 | 0.205464724 | 0.17858166 |
| 322,900 | 0.205551769 | 0.17857993 |
| 323,000 | 0.205493254 | 0.17858739 |
| 323,100 | 0.205622543 | 0.17860123 |
| 323,200 | 0.205552358 | 0.17856413 |
| 323,300 | 0.205357027 | 0.1786302 |
| 323,400 | 0.205395395 | 0.17874659 |
| 323,500 | 0.205477297 | 0.17867925 |
| 323,600 | 0.20538506 | 0.1786712 |
| 323,700 | 0.205196115 | 0.17866536 |
| 323,800 | 0.205369974 | 0.17866557 |
| 323,900 | 0.205279171 | 0.17861978 |
| 324,000 | 0.205121057 | 0.17856424 |
| 324,100 | 0.205138438 | 0.17861568 |
| 324,200 | 0.205116757 | 0.17861928 |
| 324,300 | 0.205200298 | 0.17859166 |
| 324,400 | 0.20504204 | 0.17859233 |
| 324,500 | 0.205146885 | 0.17867551 |
| 324,600 | 0.205076761 | 0.1785723 |
| 324,700 | 0.205065107 | 0.17856981 |
| 324,800 | 0.20522576 | 0.17855167 |
| 324,900 | 0.205102492 | 0.17853734 |
| 325,000 | 0.205108731 | 0.17847752 |
| 325,100 | 0.205046609 | 0.1785367 |
| 325,200 | 0.204992731 | 0.17849628 |
| 325,300 | 0.204915012 | 0.17853915 |
| 325,400 | 0.204953617 | 0.17847293 |
| 325,500 | 0.204953858 | 0.17868315 |
| 325,600 | 0.205113805 | 0.17861052 |
| 325,700 | 0.205019022 | 0.17864654 |
| 325,800 | 0.204951746 | 0.17859762 |
| 325,900 | 0.204916268 | 0.17861093 |
| 326,000 | 0.204867873 | 0.17864327 |
| 326,100 | 0.204800631 | 0.17864913 |
| 326,200 | 0.204964553 | 0.17863483 |
| 326,300 | 0.204842795 | 0.17854217 |
| 326,400 | 0.204735168 | 0.17866162 |
| 326,500 | 0.204767713 | 0.1786953 |
| 326,600 | 0.204830621 | 0.17856373 |
| 326,700 | 0.204742631 | 0.17861432 |
| 326,800 | 0.20481974 | 0.17860452 |
| 326,900 | 0.204808049 | 0.17859124 |
| 327,000 | 0.204789476 | 0.17860653 |
| 327,100 | 0.204734908 | 0.17859654 |
| 327,200 | 0.204664558 | 0.17868518 |

| | | |
|---|---|---|
| 327,300 | 0.20452402 | 0.17860741 |
| 327,400 | 0.204490137 | 0.17862533 |
| 327,500 | 0.204572055 | 0.17858305 |
| 327,600 | 0.20474189 | 0.17865853 |
| 327,700 | 0.204726681 | 0.1785909 |
| 327,800 | 0.204694053 | 0.1785845 |
| 327,900 | 0.204465772 | 0.1787083 |
| 328,000 | 0.204427667 | 0.17858555 |
| 328,100 | 0.204560908 | 0.1786478 |
| 328,200 | 0.204453368 | 0.17859159 |
| 328,300 | 0.204537586 | 0.17865461 |
| 328,400 | 0.20437053 | 0.17871616 |
| 328,500 | 0.20435702 | 0.17867846 |
| 328,600 | 0.204299967 | 0.17847678 |
| 328,700 | 0.20417091 | 0.17856175 |
| 328,800 | 0.204447895 | 0.17858074 |
| 328,900 | 0.204347333 | 0.17858844 |
| 329,000 | 0.204250321 | 0.17860169 |
| 329,100 | 0.204180271 | 0.17859211 |
| 329,200 | 0.204098471 | 0.1786579 |
| 329,300 | 0.20424645 | 0.17864814 |
| 329,400 | 0.204087753 | 0.17876523 |
| 329,500 | 0.203983197 | 0.17867623 |
| 329,600 | 0.204071836 | 0.17864328 |
| 329,700 | 0.204059277 | 0.17867088 |
| 329,800 | 0.203928296 | 0.17857241 |
| 329,900 | 0.204041208 | 0.17865615 |
| 330,000 | 0.203953227 | 0.17864553 |
| 330,100 | 0.203736745 | 0.17860912 |
| 330,200 | 0.203853084 | 0.17864322 |
| 330,300 | 0.203926917 | 0.17862941 |
| 330,400 | 0.20388437 | 0.17858816 |
| 330,500 | 0.203743241 | 0.17858473 |
| 330,600 | 0.203879701 | 0.17860838 |
| 330,700 | 0.20380798 | 0.17856671 |
| 330,800 | 0.203772564 | 0.17861619 |
| 330,900 | 0.203763993 | 0.17865438 |
| 331,000 | 0.203807417 | 0.17866872 |
| 331,100 | 0.203774856 | 0.17865634 |
| 331,200 | 0.203705694 | 0.17855122 |
| 331,300 | 0.203739478 | 0.17855766 |
| 331,400 | 0.203651933 | 0.17856581 |
| 331,500 | 0.203804147 | 0.17865496 |
| 331,600 | 0.203656874 | 0.17863728 |
| 331,700 | 0.203612293 | 0.1785616 |
| 331,800 | 0.203651285 | 0.17853245 |
| 331,900 | 0.203589501 | 0.17844995 |
| 332,000 | 0.203495271 | 0.17844477 |
| 332,100 | 0.203430628 | 0.17853719 |
| 332,200 | 0.203533189 | 0.17858479 |
| 332,300 | 0.203375559 | 0.17851235 |
| 332,400 | 0.203494952 | 0.1785034 |
| 332,500 | 0.203377387 | 0.1785128 |
| 332,600 | 0.203488379 | 0.17850976 |
| 332,700 | 0.203304023 | 0.17853832 |
| 332,800 | 0.203442189 | 0.17851793 |
| 332,900 | 0.203343459 | 0.17851742 |
| 333,000 | 0.203430708 | 0.17858249 |
| 333,100 | 0.203360369 | 0.17849896 |
| 333,200 | 0.203469395 | 0.17866113 |
| 333,300 | 0.203317031 | 0.17853646 |
| 333,400 | 0.20335156 | 0.17853751 |
| 333,500 | 0.203433143 | 0.1784259 |
| 333,600 | 0.203340046 | 0.17858047 |
| 333,700 | 0.203280516 | 0.17848428 |
| 333,800 | 0.203291108 | 0.17868141 |
| 333,900 | 0.203317354 | 0.17858818 |

| | | |
|---|---|---|
| 334,000 | 0.203230368 | 0.1785322 |
| 334,100 | 0.203168879 | 0.17850628 |
| 334,200 | 0.203255284 | 0.17849671 |
| 334,300 | 0.203298234 | 0.17852296 |
| 334,400 | 0.202990914 | 0.17844611 |
| 334,500 | 0.20292755 | 0.17857425 |
| 334,600 | 0.203116072 | 0.17849586 |
| 334,700 | 0.203215346 | 0.17849044 |
| 334,800 | 0.203295641 | 0.17847508 |
| 334,900 | 0.203149379 | 0.17853201 |
| 335,000 | 0.203124802 | 0.17851206 |
| 335,100 | 0.203037198 | 0.17851378 |
| 335,200 | 0.203054398 | 0.17862088 |
| 335,300 | 0.203173545 | 0.17861734 |
| 335,400 | 0.203103974 | 0.17846118 |
| 335,500 | 0.20299249 | 0.17855428 |
| 335,600 | 0.203031252 | 0.17858215 |
| 335,700 | 0.203247387 | 0.17849425 |
| 335,800 | 0.202860471 | 0.17850539 |
| 335,900 | 0.20308953 | 0.17840398 |
| 336,000 | 0.202809909 | 0.17854311 |
| 336,100 | 0.203006507 | 0.17852053 |
| 336,200 | 0.202958638 | 0.17847863 |
| 336,300 | 0.202918056 | 0.17864193 |
| 336,400 | 0.202839315 | 0.17850251 |
| 336,500 | 0.202902224 | 0.17847744 |
| 336,600 | 0.202847952 | 0.17848652 |
| 336,700 | 0.20278367 | 0.17852372 |
| 336,800 | 0.202871345 | 0.178535 |
| 336,900 | 0.202899033 | 0.17851801 |
| 337,000 | 0.202735262 | 0.17855606 |
| 337,100 | 0.202630546 | 0.17853818 |
| 337,200 | 0.202591973 | 0.178612 |
| 337,300 | 0.202758913 | 0.17853272 |
| 337,400 | 0.202752307 | 0.17863111 |
| 337,500 | 0.20270516 | 0.17849216 |
| 337,600 | 0.202688674 | 0.17845424 |
| 337,700 | 0.202659436 | 0.17848235 |
| 337,800 | 0.202607985 | 0.1785136 |
| 337,900 | 0.202770613 | 0.1784351 |
| 338,000 | 0.202741769 | 0.17847733 |
| 338,100 | 0.202706705 | 0.17848946 |
| 338,200 | 0.202596706 | 0.1785263 |
| 338,300 | 0.202676042 | 0.17853789 |
| 338,400 | 0.202732922 | 0.17843438 |
| 338,500 | 0.202660414 | 0.17845819 |
| 338,600 | 0.202827142 | 0.17856469 |
| 338,700 | 0.202615712 | 0.17854138 |
| 338,800 | 0.202726127 | 0.17862162 |
| 338,900 | 0.202590849 | 0.17850822 |
| 339,000 | 0.202566513 | 0.17853302 |
| 339,100 | 0.202329646 | 0.17854765 |
| 339,200 | 0.202532264 | 0.17849965 |
| 339,300 | 0.202522229 | 0.17848422 |
| 339,400 | 0.202621689 | 0.17855584 |
| 339,500 | 0.202520503 | 0.17849491 |
| 339,600 | 0.202398292 | 0.17844428 |
| 339,700 | 0.202325683 | 0.17858787 |
| 339,800 | 0.202367303 | 0.17861764 |
| 339,900 | 0.20228025 | 0.17846594 |
| 340,000 | 0.202120903 | 0.17848924 |
| 340,100 | 0.202287321 | 0.17850564 |
| 340,200 | 0.202324431 | 0.17854406 |
| 340,300 | 0.202264916 | 0.17851334 |
| 340,400 | 0.202206159 | 0.17853227 |
| 340,500 | 0.202246955 | 0.17852747 |
| 340,600 | 0.20224597 | 0.17852976 |

| | | |
|---|---|---|
| 340,700 | 0.202174294 | 0.17840356 |
| 340,800 | 0.202290051 | 0.17851137 |
| 340,900 | 0.202103358 | 0.1784249 |
| 341,000 | 0.202266211 | 0.17841973 |
| 341,100 | 0.202101158 | 0.17845366 |
| 341,200 | 0.202152025 | 0.17847477 |
| 341,300 | 0.202177346 | 0.1784865 |
| 341,400 | 0.202144119 | 0.1784771 |
| 341,500 | 0.202174052 | 0.17845318 |
| 341,600 | 0.202009741 | 0.17843023 |
| 341,700 | 0.202047418 | 0.17840893 |
| 341,800 | 0.202094835 | 0.17847946 |
| 341,900 | 0.20206661 | 0.17846078 |
| 342,000 | 0.202110484 | 0.17850748 |
| 342,100 | 0.201947775 | 0.17845596 |
| 342,200 | 0.201667738 | 0.1784479 |
| 342,300 | 0.201731556 | 0.17849036 |
| 342,400 | 0.201799761 | 0.17855049 |
| 342,500 | 0.201918826 | 0.17849136 |
| 342,600 | 0.201779306 | 0.17852921 |
| 342,700 | 0.20180209 | 0.17853027 |
| 342,800 | 0.201749267 | 0.17855308 |
| 342,900 | 0.201737689 | 0.17854093 |
| 343,000 | 0.201863668 | 0.17848198 |
| 343,100 | 0.201849779 | 0.17848042 |
| 343,200 | 0.201685811 | 0.17858682 |
| 343,300 | 0.201809052 | 0.17857948 |
| 343,400 | 0.201789405 | 0.17845955 |
| 343,500 | 0.201703326 | 0.17847439 |
| 343,600 | 0.201668405 | 0.17842551 |
| 343,700 | 0.201774128 | 0.1785512 |
| 343,800 | 0.201518412 | 0.17855225 |
| 343,900 | 0.201697235 | 0.17854365 |
| 344,000 | 0.201573606 | 0.1784429 |
| 344,100 | 0.201578867 | 0.17861401 |
| 344,200 | 0.201448579 | 0.17847352 |
| 344,300 | 0.201468832 | 0.17841563 |
| 344,400 | 0.201452004 | 0.17850271 |
| 344,500 | 0.2015277 | 0.17842865 |
| 344,600 | 0.201417816 | 0.17847574 |
| 344,700 | 0.201435024 | 0.17846224 |
| 344,800 | 0.201592501 | 0.17843613 |
| 344,900 | 0.201425057 | 0.17856072 |
| 345,000 | 0.201470805 | 0.17847808 |
| 345,100 | 0.20135328 | 0.17848611 |
| 345,200 | 0.201190888 | 0.17846673 |
| 345,300 | 0.201326505 | 0.17852388 |
| 345,400 | 0.201236985 | 0.17845943 |
| 345,500 | 0.201290102 | 0.17856619 |
| 345,600 | 0.201204135 | 0.17851928 |
| 345,700 | 0.201053985 | 0.17861458 |
| 345,800 | 0.201067673 | 0.17862784 |
| 345,900 | 0.201171076 | 0.17851167 |
| 346,000 | 0.20130781 | 0.17854159 |
| 346,100 | 0.201283461 | 0.17849943 |
| 346,200 | 0.201183238 | 0.17855952 |
| 346,300 | 0.201236091 | 0.1786059 |
| 346,400 | 0.201163142 | 0.17853366 |
| 346,500 | 0.20117778 | 0.17857202 |
| 346,600 | 0.201181037 | 0.17854335 |
| 346,700 | 0.201176359 | 0.17855429 |
| 346,800 | 0.201112057 | 0.17854812 |
| 346,900 | 0.201026116 | 0.17858653 |
| 347,000 | 0.201155512 | 0.17850343 |
| 347,100 | 0.201066102 | 0.17851014 |
| 347,200 | 0.201160939 | 0.17852627 |
| 347,300 | 0.201082426 | 0.17850415 |

| | | |
|---|---|---|
| 347,400 | 0.201038263 | 0.17848959 |
| 347,500 | 0.201001083 | 0.17845882 |
| 347,600 | 0.201018025 | 0.17855677 |
| 347,700 | 0.200931352 | 0.17855171 |
| 347,800 | 0.200911816 | 0.17854979 |
| 347,900 | 0.200925171 | 0.17852955 |
| 348,000 | 0.200920364 | 0.17856885 |
| 348,100 | 0.20083409 | 0.1785125 |
| 348,200 | 0.200780528 | 0.17848866 |
| 348,300 | 0.200835723 | 0.17845038 |
| 348,400 | 0.200648097 | 0.17855568 |
| 348,500 | 0.200583419 | 0.17854162 |
| 348,600 | 0.200581595 | 0.17853941 |
| 348,700 | 0.200641362 | 0.17849008 |
| 348,800 | 0.200605239 | 0.17844509 |
| 348,900 | 0.200694121 | 0.17851309 |
| 349,000 | 0.200548563 | 0.17841264 |
| 349,100 | 0.200637629 | 0.17844482 |
| 349,200 | 0.200624384 | 0.17845257 |
| 349,300 | 0.200382805 | 0.17848793 |
| 349,400 | 0.200429413 | 0.17849474 |
| 349,500 | 0.200522719 | 0.17847353 |
| 349,600 | 0.200347345 | 0.1785881 |
| 349,700 | 0.200345558 | 0.17853399 |
| 349,800 | 0.200360624 | 0.17846759 |
| 349,900 | 0.200352627 | 0.17852163 |
| 350,000 | 0.20040878 | 0.1784671 |
| 350,100 | 0.200322346 | 0.17848588 |
| 350,200 | 0.200151386 | 0.17857365 |
| 350,300 | 0.200225595 | 0.1786349 |
| 350,400 | 0.200235291 | 0.17854653 |
| 350,500 | 0.200319768 | 0.17856168 |
| 350,600 | 0.200245059 | 0.17854242 |
| 350,700 | 0.200236426 | 0.17860478 |
| 350,800 | 0.200260093 | 0.17865518 |
| 350,900 | 0.200187786 | 0.17856526 |
| 351,000 | 0.200006305 | 0.17843579 |
| 351,100 | 0.199998135 | 0.17850782 |
| 351,200 | 0.199921919 | 0.17847723 |
| 351,300 | 0.199983893 | 0.17849842 |
| 351,400 | 0.199975045 | 0.17850547 |
| 351,500 | 0.199882313 | 0.17852303 |
| 351,600 | 0.200030867 | 0.17855755 |
| 351,700 | 0.199942662 | 0.17862974 |
| 351,800 | 0.200016486 | 0.17857502 |
| 351,900 | 0.200032564 | 0.17847522 |
| 352,000 | 0.20003155 | 0.17857783 |
| 352,100 | 0.199933925 | 0.17864362 |
| 352,200 | 0.199806842 | 0.17860061 |
| 352,300 | 0.199967377 | 0.17858571 |
| 352,400 | 0.199947593 | 0.17860149 |
| 352,500 | 0.199801404 | 0.17852181 |
| 352,600 | 0.199888918 | 0.17853486 |
| 352,700 | 0.199834342 | 0.17849856 |
| 352,800 | 0.199753676 | 0.17847255 |
| 352,900 | 0.19977851 | 0.17851325 |
| 353,000 | 0.199874368 | 0.1785168 |
| 353,100 | 0.199702103 | 0.17849813 |
| 353,200 | 0.199624891 | 0.17851906 |
| 353,300 | 0.199757723 | 0.17851694 |
| 353,400 | 0.199572599 | 0.17844068 |
| 353,500 | 0.19955062 | 0.17851833 |
| 353,600 | 0.199560976 | 0.17845921 |
| 353,700 | 0.199497364 | 0.17848819 |
| 353,800 | 0.199624353 | 0.17851223 |
| 353,900 | 0.19969353 | 0.17850346 |
| 354,000 | 0.199633494 | 0.1784973 |

| | | |
|---|---|---|
| 354,100 | 0.199612541 | 0.1784198 |
| 354,200 | 0.199543557 | 0.17852769 |
| 354,300 | 0.19946054 | 0.17849021 |
| 354,400 | 0.199496467 | 0.17850823 |
| 354,500 | 0.199506983 | 0.17851407 |
| 354,600 | 0.199442128 | 0.17846143 |
| 354,700 | 0.199385807 | 0.17844831 |
| 354,800 | 0.199465471 | 0.17855409 |
| 354,900 | 0.199303113 | 0.17853534 |
| 355,000 | 0.199207391 | 0.1786115 |
| 355,100 | 0.199164213 | 0.17844891 |
| 355,200 | 0.19933038 | 0.17847214 |
| 355,300 | 0.199186482 | 0.1784666 |
| 355,400 | 0.199468477 | 0.17858737 |
| 355,500 | 0.199325115 | 0.17863402 |
| 355,600 | 0.199108359 | 0.17857775 |
| 355,700 | 0.198920695 | 0.17846859 |
| 355,800 | 0.199162449 | 0.17852672 |
| 355,900 | 0.199135004 | 0.17842936 |
| 356,000 | 0.198988551 | 0.17852472 |
| 356,100 | 0.199126482 | 0.17852047 |
| 356,200 | 0.199061228 | 0.1784703 |
| 356,300 | 0.199012553 | 0.17855 |
| 356,400 | 0.198988177 | 0.17847954 |
| 356,500 | 0.199113565 | 0.17846413 |
| 356,600 | 0.199099501 | 0.17863634 |
| 356,700 | 0.199049499 | 0.17856254 |
| 356,800 | 0.19884569 | 0.17859016 |
| 356,900 | 0.19904893 | 0.17857343 |
| 357,000 | 0.198927955 | 0.17855227 |
| 357,100 | 0.198817159 | 0.17849719 |
| 357,200 | 0.198742455 | 0.1785411 |
| 357,300 | 0.198817257 | 0.17856916 |
| 357,400 | 0.198768281 | 0.17857517 |
| 357,500 | 0.198723197 | 0.17846366 |
| 357,600 | 0.198675512 | 0.1784959 |
| 357,700 | 0.198779865 | 0.17845829 |
| 357,800 | 0.198786159 | 0.17839135 |
| 357,900 | 0.198684358 | 0.17854101 |
| 358,000 | 0.19866118 | 0.17858152 |
| 358,100 | 0.198513945 | 0.17855653 |
| 358,200 | 0.198368021 | 0.17854043 |
| 358,300 | 0.198506099 | 0.17855393 |
| 358,400 | 0.198473358 | 0.17855929 |
| 358,500 | 0.198435114 | 0.17852685 |
| 358,600 | 0.198538291 | 0.17846748 |
| 358,700 | 0.19832275 | 0.17853226 |
| 358,800 | 0.198485716 | 0.17853191 |
| 358,900 | 0.198416739 | 0.17854525 |
| 359,000 | 0.198479935 | 0.17851793 |
| 359,100 | 0.19836012 | 0.17849754 |
| 359,200 | 0.198207417 | 0.17849299 |
| 359,300 | 0.198277153 | 0.17853872 |
| 359,400 | 0.198405132 | 0.1785423 |
| 359,500 | 0.198306216 | 0.17853029 |
| 359,600 | 0.198418867 | 0.17847185 |
| 359,700 | 0.198309627 | 0.17851051 |
| 359,800 | 0.198251829 | 0.17852293 |
| 359,900 | 0.198190871 | 0.17852335 |
| 360,000 | 0.198084014 | 0.17858066 |
| 360,100 | 0.198286009 | 0.1785124 |
| 360,200 | 0.198161278 | 0.17841027 |
| 360,300 | 0.198197521 | 0.17853395 |
| 360,400 | 0.198152146 | 0.17849829 |
| 360,500 | 0.198241712 | 0.1784809 |
| 360,600 | 0.198313078 | 0.17843869 |
| 360,700 | 0.198331874 | 0.17843222 |

| | | |
|---|---|---|
| 360,800 | 0.198287353 | 0.17848732 |
| 360,900 | 0.19812485 | 0.17845796 |
| 361,000 | 0.198095819 | 0.17849072 |
| 361,100 | 0.197989317 | 0.17852495 |
| 361,200 | 0.198046682 | 0.17856808 |
| 361,300 | 0.197813058 | 0.1785412 |
| 361,400 | 0.197938578 | 0.17848567 |
| 361,500 | 0.198027766 | 0.17843513 |
| 361,600 | 0.198109348 | 0.17855785 |
| 361,700 | 0.198005587 | 0.1785263 |
| 361,800 | 0.197825085 | 0.17843072 |
| 361,900 | 0.197688985 | 0.17849706 |
| 362,000 | 0.197850498 | 0.17858027 |
| 362,100 | 0.197694267 | 0.17844288 |
| 362,200 | 0.19776466 | 0.17847818 |
| 362,300 | 0.197767624 | 0.17841891 |
| 362,400 | 0.197620926 | 0.17848792 |
| 362,500 | 0.197561404 | 0.1785316 |
| 362,600 | 0.19762929 | 0.17844209 |
| 362,700 | 0.197642938 | 0.17853753 |
| 362,800 | 0.19758814 | 0.17848107 |
| 362,900 | 0.19749517 | 0.17846755 |
| 363,000 | 0.19742479 | 0.17846343 |
| 363,100 | 0.197466553 | 0.17847076 |
| 363,200 | 0.197387386 | 0.17836204 |
| 363,300 | 0.197408892 | 0.17843855 |
| 363,400 | 0.197262971 | 0.17847743 |
| 363,500 | 0.19727409 | 0.17846218 |
| 363,600 | 0.197169684 | 0.17853919 |
| 363,700 | 0.197374391 | 0.17857368 |
| 363,800 | 0.197404553 | 0.17850654 |
| 363,900 | 0.197558014 | 0.17846672 |
| 364,000 | 0.197333175 | 0.17845309 |
| 364,100 | 0.19727251 | 0.17840392 |
| 364,200 | 0.197479835 | 0.17833546 |
| 364,300 | 0.197443604 | 0.17837362 |
| 364,400 | 0.197262418 | 0.17836468 |
| 364,500 | 0.197260285 | 0.1784487 |
| 364,600 | 0.197371996 | 0.17846843 |
| 364,700 | 0.197354114 | 0.1783498 |
| 364,800 | 0.197279862 | 0.17835958 |
| 364,900 | 0.197406267 | 0.17850503 |
| 365,000 | 0.197099151 | 0.17845143 |
| 365,100 | 0.197339895 | 0.17851268 |
| 365,200 | 0.197284126 | 0.17848046 |
| 365,300 | 0.197161616 | 0.17844651 |
| 365,400 | 0.197303992 | 0.17846236 |
| 365,500 | 0.197277235 | 0.17855638 |
| 365,600 | 0.197312847 | 0.17842646 |
| 365,700 | 0.197131769 | 0.17846642 |
| 365,800 | 0.197193399 | 0.17841468 |
| 365,900 | 0.197232306 | 0.17850208 |
| 366,000 | 0.197206129 | 0.17851797 |
| 366,100 | 0.196999952 | 0.17848879 |
| 366,200 | 0.197007081 | 0.17848165 |
| 366,300 | 0.197044881 | 0.17848338 |
| 366,400 | 0.197203728 | 0.17850203 |
| 366,500 | 0.196975891 | 0.17857948 |
| 366,600 | 0.196919711 | 0.1785078 |
| 366,700 | 0.196927267 | 0.17844936 |
| 366,800 | 0.196849543 | 0.17843602 |
| 366,900 | 0.196979568 | 0.17846831 |
| 367,000 | 0.196799908 | 0.17843809 |
| 367,100 | 0.196713338 | 0.17834748 |
| 367,200 | 0.196824409 | 0.17846718 |
| 367,300 | 0.196831075 | 0.17849916 |
| 367,400 | 0.196942952 | 0.17838661 |

| | | |
|---|---|---|
| 367,500 | 0.196955634 | 0.17849383 |
| 367,600 | 0.196997956 | 0.17856121 |
| 367,700 | 0.196864065 | 0.17850162 |
| 367,800 | 0.197007849 | 0.17846465 |
| 367,900 | 0.196854832 | 0.17850512 |
| 368,000 | 0.19672857 | 0.17849405 |
| 368,100 | 0.196916569 | 0.17843633 |
| 368,200 | 0.19666301 | 0.17851237 |
| 368,300 | 0.196406765 | 0.17841986 |
| 368,400 | 0.196578015 | 0.17849513 |
| 368,500 | 0.196474701 | 0.17852926 |
| 368,600 | 0.196534663 | 0.17847543 |
| 368,700 | 0.196552627 | 0.17845733 |
| 368,800 | 0.196541997 | 0.17850241 |
| 368,900 | 0.196697969 | 0.17834544 |
| 369,000 | 0.196498759 | 0.17844909 |
| 369,100 | 0.196461033 | 0.17847187 |
| 369,200 | 0.196632837 | 0.17842056 |
| 369,300 | 0.19659712 | 0.17838423 |
| 369,400 | 0.196471454 | 0.17853801 |
| 369,500 | 0.196330302 | 0.17844492 |
| 369,600 | 0.196265697 | 0.17844516 |
| 369,700 | 0.196228615 | 0.17851583 |
| 369,800 | 0.196186148 | 0.17850553 |
| 369,900 | 0.19619742 | 0.17839817 |
| 370,000 | 0.196204806 | 0.1783745 |
| 370,100 | 0.196170644 | 0.17847589 |
| 370,200 | 0.196171682 | 0.17844788 |
| 370,300 | 0.196317306 | 0.17850093 |
| 370,400 | 0.196330349 | 0.17834663 |
| 370,500 | 0.196343952 | 0.17840408 |
| 370,600 | 0.196116437 | 0.17839224 |
| 370,700 | 0.196117427 | 0.17844687 |
| 370,800 | 0.196037936 | 0.17846537 |
| 370,900 | 0.195897797 | 0.1784292 |
| 371,000 | 0.196118579 | 0.17840383 |
| 371,100 | 0.195938889 | 0.17845083 |
| 371,200 | 0.195895943 | 0.17842752 |
| 371,300 | 0.195917801 | 0.17839981 |
| 371,400 | 0.195931186 | 0.17836233 |
| 371,500 | 0.195856239 | 0.17848861 |
| 371,600 | 0.195834483 | 0.1785382 |
| 371,700 | 0.19599258 | 0.17836908 |
| 371,800 | 0.19589794 | 0.17846733 |
| 371,900 | 0.195849101 | 0.17850492 |
| 372,000 | 0.195869375 | 0.17846281 |
| 372,100 | 0.195965885 | 0.17839994 |
| 372,200 | 0.195889429 | 0.17848201 |
| 372,300 | 0.195926566 | 0.17847429 |
| 372,400 | 0.195954315 | 0.1784535 |
| 372,500 | 0.195804428 | 0.17858694 |
| 372,600 | 0.195902208 | 0.17842895 |
| 372,700 | 0.195926976 | 0.17842377 |
| 372,800 | 0.195735231 | 0.17837807 |
| 372,900 | 0.195725321 | 0.1784571 |
| 373,000 | 0.195864596 | 0.17842119 |
| 373,100 | 0.195631696 | 0.17840449 |
| 373,200 | 0.195788036 | 0.17845232 |
| 373,300 | 0.195827468 | 0.17850343 |
| 373,400 | 0.195672033 | 0.17849815 |
| 373,500 | 0.195729039 | 0.17842969 |
| 373,600 | 0.195872959 | 0.17842881 |
| 373,700 | 0.19575563 | 0.17842106 |
| 373,800 | 0.195745958 | 0.17847688 |
| 373,900 | 0.195560188 | 0.17852616 |
| 374,000 | 0.195689803 | 0.17848464 |
| 374,100 | 0.195590441 | 0.17854847 |

| | | |
|---|---|---|
| 374,200 | 0.195565995 | 0.17845543 |
| 374,300 | 0.195677579 | 0.17857716 |
| 374,400 | 0.195523648 | 0.17845765 |
| 374,500 | 0.195311022 | 0.17848848 |
| 374,600 | 0.195408843 | 0.17857848 |
| 374,700 | 0.195291942 | 0.17843772 |
| 374,800 | 0.195360524 | 0.17832558 |
| 374,900 | 0.195321973 | 0.17831927 |
| 375,000 | 0.195403113 | 0.17837749 |
| 375,100 | 0.195403948 | 0.17835735 |
| 375,200 | 0.195274452 | 0.17844648 |
| 375,300 | 0.19532503 | 0.17838327 |
| 375,400 | 0.195323749 | 0.17836758 |
| 375,500 | 0.195298455 | 0.17844053 |
| 375,600 | 0.195317779 | 0.17845186 |
| 375,700 | 0.195363162 | 0.17847824 |
| 375,800 | 0.195243834 | 0.17842761 |
| 375,900 | 0.195060093 | 0.17843 |
| 376,000 | 0.195192261 | 0.17839991 |
| 376,100 | 0.19520741 | 0.17843324 |
| 376,200 | 0.195266734 | 0.1785144 |
| 376,300 | 0.195410745 | 0.17839052 |
| 376,400 | 0.195397962 | 0.17843973 |
| 376,500 | 0.195491842 | 0.17843603 |
| 376,600 | 0.195375975 | 0.17849227 |
| 376,700 | 0.195550593 | 0.17848517 |
| 376,800 | 0.195546445 | 0.17845937 |
| 376,900 | 0.19545502 | 0.17847117 |
| 377,000 | 0.195201665 | 0.17850902 |
| 377,100 | 0.195316373 | 0.17849365 |
| 377,200 | 0.195107655 | 0.17843884 |
| 377,300 | 0.195310383 | 0.17843391 |
| 377,400 | 0.195294088 | 0.17846129 |
| 377,500 | 0.19532996 | 0.17845424 |
| 377,600 | 0.195278728 | 0.17844577 |
| 377,700 | 0.19518999 | 0.17847179 |
| 377,800 | 0.195144356 | 0.17845291 |
| 377,900 | 0.195053771 | 0.17842704 |
| 378,000 | 0.19509635 | 0.17852042 |
| 378,100 | 0.195068327 | 0.17844087 |
| 378,200 | 0.195053059 | 0.17846107 |
| 378,300 | 0.195182439 | 0.17842617 |
| 378,400 | 0.195200364 | 0.17850533 |
| 378,500 | 0.195177528 | 0.17842483 |
| 378,600 | 0.195152084 | 0.17840809 |
| 378,700 | 0.195102061 | 0.17839999 |
| 378,800 | 0.195081456 | 0.17841316 |
| 378,900 | 0.195036043 | 0.17846774 |
| 379,000 | 0.195071921 | 0.17838819 |
| 379,100 | 0.19492159 | 0.17844712 |
| 379,200 | 0.195048864 | 0.17842403 |
| 379,300 | 0.195089758 | 0.17846534 |
| 379,400 | 0.195180534 | 0.17846043 |
| 379,500 | 0.195074532 | 0.17838966 |
| 379,600 | 0.194956467 | 0.17847166 |
| 379,700 | 0.195087576 | 0.17848915 |
| 379,800 | 0.194907571 | 0.17843443 |
| 379,900 | 0.19497623 | 0.17846849 |
| 380,000 | 0.194862606 | 0.17849675 |
| 380,100 | 0.194839774 | 0.17839408 |
| 380,200 | 0.194797082 | 0.1785116 |
| 380,300 | 0.194865628 | 0.17850076 |
| 380,400 | 0.194813203 | 0.17839231 |
| 380,500 | 0.194862767 | 0.17835987 |
| 380,600 | 0.194632899 | 0.17841445 |
| 380,700 | 0.194741891 | 0.17847464 |
| 380,800 | 0.194686313 | 0.17847018 |

| | | |
|---|---|---|
| 380,900 | 0.194573326 | 0.17839812 |
| 381,000 | 0.194637396 | 0.1784808 |
| 381,100 | 0.194452735 | 0.1783758 |
| 381,200 | 0.194452449 | 0.17844842 |
| 381,300 | 0.194503508 | 0.17846386 |
| 381,400 | 0.194530657 | 0.17837413 |
| 381,500 | 0.194430256 | 0.17840875 |
| 381,600 | 0.194431378 | 0.17841855 |
| 381,700 | 0.194388127 | 0.17832682 |
| 381,800 | 0.194442664 | 0.17844738 |
| 381,900 | 0.194390549 | 0.17834644 |
| 382,000 | 0.194366573 | 0.17839759 |
| 382,100 | 0.194262918 | 0.17843081 |
| 382,200 | 0.194329188 | 0.17836064 |
| 382,300 | 0.19434823 | 0.17828898 |
| 382,400 | 0.194332734 | 0.17838031 |
| 382,500 | 0.194425916 | 0.17837689 |
| 382,600 | 0.194396097 | 0.17842332 |
| 382,700 | 0.194212057 | 0.17839787 |
| 382,800 | 0.194245435 | 0.1784215 |
| 382,900 | 0.193922606 | 0.17848117 |
| 383,000 | 0.193980133 | 0.17841566 |
| 383,100 | 0.194057406 | 0.17836038 |
| 383,200 | 0.194199217 | 0.17834324 |
| 383,300 | 0.193975392 | 0.17828589 |
| 383,400 | 0.194237241 | 0.1783663 |
| 383,500 | 0.19386113 | 0.17833057 |
| 383,600 | 0.194079991 | 0.17848664 |
| 383,700 | 0.193935975 | 0.17840305 |
| 383,800 | 0.194036124 | 0.17840165 |
| 383,900 | 0.194023177 | 0.17835712 |
| 384,000 | 0.194127395 | 0.1783086 |
| 384,100 | 0.193857127 | 0.17842816 |
| 384,200 | 0.193971456 | 0.17843164 |
| 384,300 | 0.193908709 | 0.17837789 |
| 384,400 | 0.193927675 | 0.17839735 |
| 384,500 | 0.193945937 | 0.17840819 |
| 384,600 | 0.193765815 | 0.17844075 |
| 384,700 | 0.19387134 | 0.17838776 |
| 384,800 | 0.193794486 | 0.17839043 |
| 384,900 | 0.193947785 | 0.17824089 |
| 385,000 | 0.193773635 | 0.17832428 |
| 385,100 | 0.193772011 | 0.17833311 |
| 385,200 | 0.193977313 | 0.17835162 |
| 385,300 | 0.194081305 | 0.17836681 |
| 385,400 | 0.193800056 | 0.17835111 |
| 385,500 | 0.193811772 | 0.17842033 |
| 385,600 | 0.193741855 | 0.17840753 |
| 385,700 | 0.193685874 | 0.17839759 |
| 385,800 | 0.193750097 | 0.17850028 |
| 385,900 | 0.193760334 | 0.17837526 |
| 386,000 | 0.193811182 | 0.17835935 |
| 386,100 | 0.193676156 | 0.17833607 |
| 386,200 | 0.193516099 | 0.17838695 |
| 386,300 | 0.193547764 | 0.17835384 |
| 386,400 | 0.193557397 | 0.17839134 |
| 386,500 | 0.193493585 | 0.17836555 |
| 386,600 | 0.193475385 | 0.17836458 |
| 386,700 | 0.193514199 | 0.17837078 |
| 386,800 | 0.193442959 | 0.17841362 |
| 386,900 | 0.193438592 | 0.17843113 |
| 387,000 | 0.193240868 | 0.17831398 |
| 387,100 | 0.19329613 | 0.17843566 |
| 387,200 | 0.193301231 | 0.17847054 |
| 387,300 | 0.193257438 | 0.17844766 |
| 387,400 | 0.193447229 | 0.17840531 |
| 387,500 | 0.19338567 | 0.17837416 |

| | | |
|---|---|---|
| 387,600 | 0.193468489 | 0.17839612 |
| 387,700 | 0.193391594 | 0.17836363 |
| 387,800 | 0.193277194 | 0.17851248 |
| 387,900 | 0.193300826 | 0.178389 |
| 388,000 | 0.193236739 | 0.17836378 |
| 388,100 | 0.19322926 | 0.1783309 |
| 388,200 | 0.193226326 | 0.1783945 |
| 388,300 | 0.193353946 | 0.17841124 |
| 388,400 | 0.193317467 | 0.17841293 |
| 388,500 | 0.193158568 | 0.17843964 |
| 388,600 | 0.193105255 | 0.17849832 |
| 388,700 | 0.193192998 | 0.17849599 |
| 388,800 | 0.192971126 | 0.17834218 |
| 388,900 | 0.193216548 | 0.17842349 |
| 389,000 | 0.193127516 | 0.17839459 |
| 389,100 | 0.193057979 | 0.17841761 |
| 389,200 | 0.193149201 | 0.17836955 |
| 389,300 | 0.192886091 | 0.17832822 |
| 389,400 | 0.193041093 | 0.17839846 |
| 389,500 | 0.193106515 | 0.17840247 |
| 389,600 | 0.192881558 | 0.17843517 |
| 389,700 | 0.193236938 | 0.17836687 |
| 389,800 | 0.192984443 | 0.17836331 |
| 389,900 | 0.192871158 | 0.17834245 |
| 390,000 | 0.192930958 | 0.17833185 |
| 390,100 | 0.192982955 | 0.17831078 |
| 390,200 | 0.192857599 | 0.17835025 |
| 390,300 | 0.192835764 | 0.17833715 |
| 390,400 | 0.192692966 | 0.17833922 |
| 390,500 | 0.192770139 | 0.17837504 |
| 390,600 | 0.192782276 | 0.17839382 |
| 390,700 | 0.192710075 | 0.17837571 |
| 390,800 | 0.192772697 | 0.17828222 |
| 390,900 | 0.192886554 | 0.17836128 |
| 391,000 | 0.192941066 | 0.17838616 |
| 391,100 | 0.192947225 | 0.17838473 |
| 391,200 | 0.193009632 | 0.17837589 |
| 391,300 | 0.192754594 | 0.17840432 |
| 391,400 | 0.192766803 | 0.17839358 |
| 391,500 | 0.192612973 | 0.17847046 |
| 391,600 | 0.192830353 | 0.1783377 |
| 391,700 | 0.192741631 | 0.17833153 |
| 391,800 | 0.192592695 | 0.1783508 |
| 391,900 | 0.192770812 | 0.17835313 |
| 392,000 | 0.192436015 | 0.17826971 |
| 392,100 | 0.192553824 | 0.17834081 |
| 392,200 | 0.192601188 | 0.17832193 |
| 392,300 | 0.192654259 | 0.17838011 |
| 392,400 | 0.192562553 | 0.17837894 |
| 392,500 | 0.192580436 | 0.17834822 |
| 392,600 | 0.192420518 | 0.17845706 |
| 392,700 | 0.192705927 | 0.17841733 |
| 392,800 | 0.192616693 | 0.17830886 |
| 392,900 | 0.192676313 | 0.17835687 |
| 393,000 | 0.192683044 | 0.17834968 |
| 393,100 | 0.192595983 | 0.17842255 |
| 393,200 | 0.192480381 | 0.17842381 |
| 393,300 | 0.192346297 | 0.17838557 |
| 393,400 | 0.192422335 | 0.17839417 |
| 393,500 | 0.192534378 | 0.17837336 |
| 393,600 | 0.192593158 | 0.17832756 |
| 393,700 | 0.192501899 | 0.17825405 |
| 393,800 | 0.192562155 | 0.17827168 |
| 393,900 | 0.192442782 | 0.17832945 |
| 394,000 | 0.192622435 | 0.1783118 |
| 394,100 | 0.19241315 | 0.17834046 |
| 394,200 | 0.192482733 | 0.17829101 |

| | | |
|---|---|---|
| 394,300 | 0.192552177 | 0.17836348 |
| 394,400 | 0.19240981 | 0.17842834 |
| 394,500 | 0.192352858 | 0.17842479 |
| 394,600 | 0.192194086 | 0.17833575 |
| 394,700 | 0.192265375 | 0.17833395 |
| 394,800 | 0.192153161 | 0.17825406 |
| 394,900 | 0.19224681 | 0.17827675 |
| 395,000 | 0.192150383 | 0.17836205 |
| 395,100 | 0.192133256 | 0.17827019 |
| 395,200 | 0.192136583 | 0.17831016 |
| 395,300 | 0.192172424 | 0.17834897 |
| 395,400 | 0.192129033 | 0.17829995 |
| 395,500 | 0.191995223 | 0.17831014 |
| 395,600 | 0.192016364 | 0.17828231 |
| 395,700 | 0.192080149 | 0.17835547 |
| 395,800 | 0.191911582 | 0.17824393 |
| 395,900 | 0.1918598 | 0.17825613 |
| 396,000 | 0.192022415 | 0.1782427 |
| 396,100 | 0.192050605 | 0.17838652 |
| 396,200 | 0.191874284 | 0.17836044 |
| 396,300 | 0.191926347 | 0.17839862 |
| 396,400 | 0.191895784 | 0.17833438 |
| 396,500 | 0.191891978 | 0.17832859 |
| 396,600 | 0.191777533 | 0.1784016 |
| 396,700 | 0.191834034 | 0.17836773 |
| 396,800 | 0.19200705 | 0.17833181 |
| 396,900 | 0.191861812 | 0.17826284 |
| 397,000 | 0.191771107 | 0.17821244 |
| 397,100 | 0.191752345 | 0.17831742 |
| 397,200 | 0.19164856 | 0.1783322 |
| 397,300 | 0.191795702 | 0.17838989 |
| 397,400 | 0.191792205 | 0.17823183 |
| 397,500 | 0.191709472 | 0.17834185 |
| 397,600 | 0.191769225 | 0.17835281 |
| 397,700 | 0.191852667 | 0.17839497 |
| 397,800 | 0.191652365 | 0.17842152 |
| 397,900 | 0.191747479 | 0.17842133 |
| 398,000 | 0.191558912 | 0.17838523 |
| 398,100 | 0.191520101 | 0.17836095 |
| 398,200 | 0.191545041 | 0.17838683 |
| 398,300 | 0.191601698 | 0.17831087 |
| 398,400 | 0.191684751 | 0.17840561 |
| 398,500 | 0.191594276 | 0.17841725 |
| 398,600 | 0.191457242 | 0.17836303 |
| 398,700 | 0.191634817 | 0.17839186 |
| 398,800 | 0.191781745 | 0.17838541 |
| 398,900 | 0.19184765 | 0.17836724 |
| 399,000 | 0.191570528 | 0.17839678 |
| 399,100 | 0.191626018 | 0.17838038 |
| 399,200 | 0.191679303 | 0.1783256 |
| 399,300 | 0.19135093 | 0.17833076 |
| 399,400 | 0.191395302 | 0.17849734 |
| 399,500 | 0.191488097 | 0.17844303 |
| 399,600 | 0.191564935 | 0.17837802 |
| 399,700 | 0.191439244 | 0.17846803 |
| 399,800 | 0.191567404 | 0.17842711 |
| 399,900 | 0.19152382 | 0.17834961 |
| 400,000 | 0.191491797 | 0.17834717 |
| 400,100 | 0.191457283 | 0.17841896 |
| 400,200 | 0.191268664 | 0.17834273 |
| 400,300 | 0.191278869 | 0.17830107 |
| 400,400 | 0.191341477 | 0.17846086 |
| 400,500 | 0.191383685 | 0.17848693 |
| 400,600 | 0.191436169 | 0.17833685 |
| 400,700 | 0.19136989 | 0.17843414 |
| 400,800 | 0.191449464 | 0.17840889 |
| 400,900 | 0.191273597 | 0.17830994 |

| | | |
|---|---|---|
| 401,000 | 0.191332048 | 0.1783769 |
| 401,100 | 0.191293851 | 0.17844538 |
| 401,200 | 0.191314238 | 0.17835714 |
| 401,300 | 0.191199591 | 0.17835162 |
| 401,400 | 0.191322737 | 0.178424 |
| 401,500 | 0.191294793 | 0.17833787 |
| 401,600 | 0.191280858 | 0.17831074 |
| 401,700 | 0.191296637 | 0.17841462 |
| 401,800 | 0.191341908 | 0.17842486 |
| 401,900 | 0.191250478 | 0.17843017 |
| 402,000 | 0.191218061 | 0.17834116 |
| 402,100 | 0.191089316 | 0.1783302 |
| 402,200 | 0.19118354 | 0.17832998 |
| 402,300 | 0.191198075 | 0.17826465 |
| 402,400 | 0.191180145 | 0.17838325 |
| 402,500 | 0.191301362 | 0.17837679 |
| 402,600 | 0.191181619 | 0.17846218 |
| 402,700 | 0.191121615 | 0.17837768 |
| 402,800 | 0.191182966 | 0.17834595 |
| 402,900 | 0.190941521 | 0.17830758 |
| 403,000 | 0.191049156 | 0.17840725 |
| 403,100 | 0.191083122 | 0.17836295 |
| 403,200 | 0.191110319 | 0.17841249 |
| 403,300 | 0.191134913 | 0.17842894 |
| 403,400 | 0.191037679 | 0.17836933 |
| 403,500 | 0.190897834 | 0.17830518 |
| 403,600 | 0.19090224 | 0.17832533 |
| 403,700 | 0.190900343 | 0.17833997 |
| 403,800 | 0.190888849 | 0.17851244 |
| 403,900 | 0.190925306 | 0.17843276 |
| 404,000 | 0.190780219 | 0.17847234 |
| 404,100 | 0.190889901 | 0.178328 |
| 404,200 | 0.190744662 | 0.17845115 |
| 404,300 | 0.19080272 | 0.17851371 |
| 404,400 | 0.190857276 | 0.17842636 |
| 404,500 | 0.190897289 | 0.17850014 |
| 404,600 | 0.191012541 | 0.17837794 |
| 404,700 | 0.191090767 | 0.17843433 |
| 404,800 | 0.190968671 | 0.17839906 |
| 404,900 | 0.190957126 | 0.17841752 |
| 405,000 | 0.1909433 | 0.17837644 |
| 405,100 | 0.190844621 | 0.17831574 |
| 405,200 | 0.190866573 | 0.17837772 |
| 405,300 | 0.190720446 | 0.17837211 |
| 405,400 | 0.190728569 | 0.17835423 |
| 405,500 | 0.190678132 | 0.1783405 |
| 405,600 | 0.190723289 | 0.17825527 |
| 405,700 | 0.190656249 | 0.17841834 |
| 405,800 | 0.190781309 | 0.17837829 |
| 405,900 | 0.190491592 | 0.17841726 |
| 406,000 | 0.19075564 | 0.17837966 |
| 406,100 | 0.19069845 | 0.17841383 |
| 406,200 | 0.190631444 | 0.17838243 |
| 406,300 | 0.190611226 | 0.17840688 |
| 406,400 | 0.190578111 | 0.17838074 |
| 406,500 | 0.190530795 | 0.17840725 |
| 406,600 | 0.190590454 | 0.17835653 |
| 406,700 | 0.19050411 | 0.17837225 |
| 406,800 | 0.190353155 | 0.17826606 |
| 406,900 | 0.190294445 | 0.17833447 |
| 407,000 | 0.190396625 | 0.17836653 |
| 407,100 | 0.190425936 | 0.17834844 |
| 407,200 | 0.190451267 | 0.17836427 |
| 407,300 | 0.190520015 | 0.17839849 |
| 407,400 | 0.190395837 | 0.17838205 |
| 407,500 | 0.190212465 | 0.17834148 |
| 407,600 | 0.19033115 | 0.17836284 |

| | | |
|---|---|---|
| 407,700 | 0.190297873 | 0.17837625 |
| 407,800 | 0.190358689 | 0.17846557 |
| 407,900 | 0.190313313 | 0.17836538 |
| 408,000 | 0.190268684 | 0.17834861 |
| 408,100 | 0.190358419 | 0.17829196 |
| 408,200 | 0.19019485 | 0.17835566 |
| 408,300 | 0.190288284 | 0.17831783 |
| 408,400 | 0.190216028 | 0.17834948 |
| 408,500 | 0.190189329 | 0.17840228 |
| 408,600 | 0.1902237 | 0.17832116 |
| 408,700 | 0.190207609 | 0.17834055 |
| 408,800 | 0.190289786 | 0.17842463 |
| 408,900 | 0.190282222 | 0.17840067 |
| 409,000 | 0.190155666 | 0.17831858 |
| 409,100 | 0.190163066 | 0.17846272 |
| 409,200 | 0.19011049 | 0.17839693 |
| 409,300 | 0.190114124 | 0.17842088 |
| 409,400 | 0.189992938 | 0.17835407 |
| 409,500 | 0.190098358 | 0.17839118 |
| 409,600 | 0.19003841 | 0.17835264 |
| 409,700 | 0.19014369 | 0.1783269 |
| 409,800 | 0.190026083 | 0.17844452 |
| 409,900 | 0.190023227 | 0.17845135 |
| 410,000 | 0.190039935 | 0.17840646 |
| 410,100 | 0.190127752 | 0.17837042 |
| 410,200 | 0.189971246 | 0.17840592 |
| 410,300 | 0.190007275 | 0.17837236 |
| 410,400 | 0.189909096 | 0.17835781 |
| 410,500 | 0.189866841 | 0.17841954 |
| 410,600 | 0.189862881 | 0.17845723 |
| 410,700 | 0.189825476 | 0.1783571 |
| 410,800 | 0.189848952 | 0.17834129 |
| 410,900 | 0.189766017 | 0.1783113 |
| 411,000 | 0.189902536 | 0.17835761 |
| 411,100 | 0.189845479 | 0.17829433 |
| 411,200 | 0.189796449 | 0.17833684 |
| 411,300 | 0.189938441 | 0.17839524 |
| 411,400 | 0.189875385 | 0.17830624 |
| 411,500 | 0.189888212 | 0.17844527 |
| 411,600 | 0.189777335 | 0.17831317 |
| 411,700 | 0.189532069 | 0.17835953 |
| 411,800 | 0.18982929 | 0.17832644 |
| 411,900 | 0.189826585 | 0.17830595 |
| 412,000 | 0.18966182 | 0.17839019 |
| 412,100 | 0.189633624 | 0.17835078 |
| 412,200 | 0.189634634 | 0.1782353 |
| 412,300 | 0.189641609 | 0.17825584 |
| 412,400 | 0.189921259 | 0.17831643 |
| 412,500 | 0.189865024 | 0.17841352 |
| 412,600 | 0.189529846 | 0.17829432 |
| 412,700 | 0.1895985 | 0.1783365 |
| 412,800 | 0.189459074 | 0.17841409 |
| 412,900 | 0.18964386 | 0.17837246 |
| 413,000 | 0.189675776 | 0.17836455 |
| 413,100 | 0.189648126 | 0.1783217 |
| 413,200 | 0.189748663 | 0.17841621 |
| 413,300 | 0.189656927 | 0.17836768 |
| 413,400 | 0.189585938 | 0.17833654 |
| 413,500 | 0.189602493 | 0.17839516 |
| 413,600 | 0.189548802 | 0.17840703 |
| 413,700 | 0.18952229 | 0.17829575 |
| 413,800 | 0.189451637 | 0.17832821 |
| 413,900 | 0.189572154 | 0.17830959 |
| 414,000 | 0.189401152 | 0.1783083 |
| 414,100 | 0.18952072 | 0.17834638 |
| 414,200 | 0.189513966 | 0.17832765 |
| 414,300 | 0.189557113 | 0.17832231 |

| | | |
|---|---|---|
| 414,400 | 0.189407484 | 0.1782833 |
| 414,500 | 0.18948973 | 0.17834328 |
| 414,600 | 0.189382089 | 0.17830016 |
| 414,700 | 0.189559968 | 0.17826672 |
| 414,800 | 0.189618504 | 0.17831026 |
| 414,900 | 0.189557467 | 0.17841685 |
| 415,000 | 0.189421064 | 0.17838188 |
| 415,100 | 0.189547553 | 0.17836198 |
| 415,200 | 0.189469722 | 0.17827219 |
| 415,300 | 0.18940047 | 0.17841244 |
| 415,400 | 0.189259817 | 0.17844079 |
| 415,500 | 0.189371028 | 0.17835024 |
| 415,600 | 0.189274192 | 0.17835925 |
| 415,700 | 0.18927015 | 0.17827954 |
| 415,800 | 0.189269352 | 0.17822232 |
| 415,900 | 0.18912023 | 0.17823539 |
| 416,000 | 0.189173049 | 0.17831829 |
| 416,100 | 0.189227188 | 0.17826062 |
| 416,200 | 0.189262564 | 0.17832712 |
| 416,300 | 0.189421686 | 0.17832906 |
| 416,400 | 0.189300368 | 0.17831075 |
| 416,500 | 0.189457017 | 0.17829565 |
| 416,600 | 0.189276896 | 0.17837919 |
| 416,700 | 0.189427743 | 0.17835493 |
| 416,800 | 0.18928173 | 0.17830913 |
| 416,900 | 0.189267679 | 0.17834136 |
| 417,000 | 0.189317145 | 0.17835766 |
| 417,100 | 0.189114961 | 0.17839966 |
| 417,200 | 0.18914628 | 0.17839025 |
| 417,300 | 0.189098306 | 0.17841338 |
| 417,400 | 0.189137401 | 0.1783255 |
| 417,500 | 0.189006307 | 0.17839693 |
| 417,600 | 0.189007097 | 0.17839391 |
| 417,700 | 0.189139712 | 0.17819928 |
| 417,800 | 0.189096576 | 0.17830109 |
| 417,900 | 0.18907533 | 0.17828616 |
| 418,000 | 0.189044539 | 0.17826863 |
| 418,100 | 0.188947435 | 0.17823647 |
| 418,200 | 0.188941836 | 0.17829799 |
| 418,300 | 0.188892015 | 0.17831366 |
| 418,400 | 0.188941332 | 0.17833858 |
| 418,500 | 0.188890638 | 0.17827212 |
| 418,600 | 0.188925024 | 0.17837269 |
| 418,700 | 0.188928277 | 0.17827053 |
| 418,800 | 0.188973923 | 0.1783066 |
| 418,900 | 0.188929644 | 0.17828186 |
| 419,000 | 0.188898686 | 0.17841205 |
| 419,100 | 0.188765315 | 0.17827043 |
| 419,200 | 0.18870392 | 0.17836471 |
| 419,300 | 0.188855019 | 0.17832078 |
| 419,400 | 0.188936667 | 0.17829953 |
| 419,500 | 0.188698656 | 0.17825723 |
| 419,600 | 0.188792686 | 0.17824732 |
| 419,700 | 0.188859518 | 0.17831601 |
| 419,800 | 0.188882313 | 0.17840376 |
| 419,900 | 0.188864904 | 0.17825142 |
| 420,000 | 0.188771816 | 0.17834584 |
| 420,100 | 0.188711688 | 0.17823329 |
| 420,200 | 0.188580998 | 0.17824009 |
| 420,300 | 0.188526291 | 0.17834024 |
| 420,400 | 0.18863587 | 0.17825826 |
| 420,500 | 0.1887442 | 0.17832256 |
| 420,600 | 0.188829415 | 0.17831104 |
| 420,700 | 0.188785885 | 0.17828907 |
| 420,800 | 0.188761376 | 0.17829183 |
| 420,900 | 0.188717567 | 0.17829087 |
| 421,000 | 0.188742016 | 0.17833819 |

| | | |
|---|---|---|
| 421,100 | 0.188872464 | 0.17828774 |
| 421,200 | 0.188863972 | 0.17835889 |
| 421,300 | 0.188684983 | 0.17822757 |
| 421,400 | 0.188609099 | 0.17827599 |
| 421,500 | 0.188686102 | 0.17833305 |
| 421,600 | 0.188540882 | 0.17829609 |
| 421,700 | 0.188616119 | 0.17830262 |
| 421,800 | 0.188528409 | 0.17836135 |
| 421,900 | 0.188640145 | 0.17830715 |
| 422,000 | 0.18862219 | 0.17822714 |
| 422,100 | 0.18847414 | 0.17836719 |
| 422,200 | 0.18855462 | 0.17830681 |
| 422,300 | 0.188544203 | 0.17827559 |
| 422,400 | 0.188557886 | 0.17840871 |
| 422,500 | 0.188398286 | 0.17830233 |
| 422,600 | 0.188547537 | 0.17832649 |
| 422,700 | 0.188495773 | 0.17830335 |
| 422,800 | 0.188490403 | 0.17833761 |
| 422,900 | 0.188434765 | 0.17825018 |
| 423,000 | 0.188373141 | 0.17834847 |
| 423,100 | 0.188498926 | 0.17819349 |
| 423,200 | 0.18832323 | 0.17830814 |
| 423,300 | 0.188487514 | 0.17831442 |
| 423,400 | 0.188478961 | 0.17832277 |
| 423,500 | 0.188452887 | 0.17830524 |
| 423,600 | 0.188598175 | 0.17832906 |
| 423,700 | 0.188367967 | 0.17832531 |
| 423,800 | 0.18856302 | 0.17832932 |
| 423,900 | 0.188288837 | 0.17829402 |
| 424,000 | 0.188439091 | 0.17827616 |
| 424,100 | 0.18844937 | 0.17836606 |
| 424,200 | 0.188285176 | 0.17834724 |
| 424,300 | 0.188190242 | 0.1783344 |
| 424,400 | 0.188307607 | 0.1782978 |
| 424,500 | 0.188386479 | 0.17831756 |
| 424,600 | 0.188295234 | 0.17828673 |
| 424,700 | 0.188439076 | 0.17833464 |
| 424,800 | 0.188514067 | 0.1783633 |
| 424,900 | 0.188334033 | 0.17835968 |
| 425,000 | 0.188320445 | 0.17840261 |
| 425,100 | 0.188400183 | 0.17829109 |
| 425,200 | 0.188345351 | 0.17838843 |
| 425,300 | 0.188271793 | 0.17839126 |
| 425,400 | 0.188271007 | 0.17843632 |
| 425,500 | 0.188399751 | 0.17829365 |
| 425,600 | 0.188300966 | 0.17833039 |
| 425,700 | 0.188127013 | 0.17841686 |
| 425,800 | 0.188284511 | 0.17835264 |
| 425,900 | 0.188389098 | 0.1784341 |
| 426,000 | 0.188256313 | 0.17832691 |
| 426,100 | 0.188044365 | 0.17839591 |
| 426,200 | 0.188065754 | 0.17841695 |
| 426,300 | 0.188057312 | 0.17833806 |
| 426,400 | 0.188082677 | 0.17835754 |
| 426,500 | 0.187886121 | 0.17838995 |
| 426,600 | 0.187999518 | 0.17832003 |
| 426,700 | 0.187916237 | 0.17832169 |
| 426,800 | 0.187814568 | 0.17836532 |
| 426,900 | 0.187806543 | 0.17842884 |
| 427,000 | 0.187730173 | 0.17837533 |
| 427,100 | 0.187829345 | 0.17835006 |
| 427,200 | 0.187945057 | 0.17833249 |
| 427,300 | 0.188055315 | 0.17840907 |
| 427,400 | 0.187935718 | 0.17842403 |
| 427,500 | 0.188001058 | 0.17853979 |
| 427,600 | 0.188056062 | 0.17830531 |
| 427,700 | 0.187873759 | 0.17839988 |

| | | |
|---|---|---|
| 427,800 | 0.187923702 | 0.17836162 |
| 427,900 | 0.187943404 | 0.17839468 |
| 428,000 | 0.187900441 | 0.17836882 |
| 428,100 | 0.187845958 | 0.17855202 |
| 428,200 | 0.187644341 | 0.17843301 |
| 428,300 | 0.187738945 | 0.17846477 |
| 428,400 | 0.187684517 | 0.17838037 |
| 428,500 | 0.187790619 | 0.17833673 |
| 428,600 | 0.18785605 | 0.17840049 |
| 428,700 | 0.187650998 | 0.17842965 |
| 428,800 | 0.187601412 | 0.17841618 |
| 428,900 | 0.187598458 | 0.17839181 |
| 429,000 | 0.187762345 | 0.17835784 |
| 429,100 | 0.187663965 | 0.17842969 |
| 429,200 | 0.187559525 | 0.17840804 |
| 429,300 | 0.18755217 | 0.17832003 |
| 429,400 | 0.187644504 | 0.17833532 |
| 429,500 | 0.18767888 | 0.17835907 |
| 429,600 | 0.187553001 | 0.17840645 |
| 429,700 | 0.187571526 | 0.17840392 |
| 429,800 | 0.187630636 | 0.17849182 |
| 429,900 | 0.187607506 | 0.17846095 |
| 430,000 | 0.187529622 | 0.17835118 |
| 430,100 | 0.18748971 | 0.17833119 |
| 430,200 | 0.1875748 | 0.17833459 |
| 430,300 | 0.187501818 | 0.17840177 |
| 430,400 | 0.187465559 | 0.1784326 |
| 430,500 | 0.18738834 | 0.17838936 |
| 430,600 | 0.187169646 | 0.17837358 |
| 430,700 | 0.187362038 | 0.1783657 |
| 430,800 | 0.187346756 | 0.17835227 |
| 430,900 | 0.187374221 | 0.17835281 |
| 431,000 | 0.187229634 | 0.17837505 |
| 431,100 | 0.187302946 | 0.17833435 |
| 431,200 | 0.187216315 | 0.17845717 |
| 431,300 | 0.187146705 | 0.1784045 |
| 431,400 | 0.187073076 | 0.17837855 |
| 431,500 | 0.187103624 | 0.17836168 |
| 431,600 | 0.187366875 | 0.17834743 |
| 431,700 | 0.187325735 | 0.17844627 |
| 431,800 | 0.187282061 | 0.17843703 |
| 431,900 | 0.187251212 | 0.17843625 |
| 432,000 | 0.187066833 | 0.17839022 |
| 432,100 | 0.187406242 | 0.1784068 |
| 432,200 | 0.187125065 | 0.17847131 |
| 432,300 | 0.187282725 | 0.17837016 |
| 432,400 | 0.187144745 | 0.1784125 |
| 432,500 | 0.187033745 | 0.17836261 |
| 432,600 | 0.187022354 | 0.1784452 |
| 432,700 | 0.186945646 | 0.1783557 |
| 432,800 | 0.186954158 | 0.17830241 |
| 432,900 | 0.186948869 | 0.17837099 |
| 433,000 | 0.186906717 | 0.17824917 |
| 433,100 | 0.186767921 | 0.17832815 |
| 433,200 | 0.186892794 | 0.17830382 |
| 433,300 | 0.186837558 | 0.17835722 |
| 433,400 | 0.186803144 | 0.17824119 |
| 433,500 | 0.186807454 | 0.17832474 |
| 433,600 | 0.186773299 | 0.17839184 |
| 433,700 | 0.186764162 | 0.17836219 |
| 433,800 | 0.186741591 | 0.17833021 |
| 433,900 | 0.186915114 | 0.17844037 |
| 434,000 | 0.186836622 | 0.17839977 |
| 434,100 | 0.18678039 | 0.17839798 |
| 434,200 | 0.186769374 | 0.17831898 |
| 434,300 | 0.186882086 | 0.17835394 |
| 434,400 | 0.186784937 | 0.17837299 |

| | | |
|---|---|---|
| 434,500 | 0.186556194 | 0.17829231 |
| 434,600 | 0.186820614 | 0.17834158 |
| 434,700 | 0.186913208 | 0.17835957 |
| 434,800 | 0.186882967 | 0.17837014 |
| 434,900 | 0.186767965 | 0.17838981 |
| 435,000 | 0.186666211 | 0.17833855 |
| 435,100 | 0.186739401 | 0.1784256 |
| 435,200 | 0.186732379 | 0.17840146 |
| 435,300 | 0.186640807 | 0.17834817 |
| 435,400 | 0.186637415 | 0.1784325 |
| 435,500 | 0.186679819 | 0.17834067 |
| 435,600 | 0.186718672 | 0.17833906 |
| 435,700 | 0.186689435 | 0.17836264 |
| 435,800 | 0.186661375 | 0.17837033 |
| 435,900 | 0.186632151 | 0.17831652 |
| 436,000 | 0.186497886 | 0.17838005 |
| 436,100 | 0.186467987 | 0.17835567 |
| 436,200 | 0.18642749 | 0.17844385 |
| 436,300 | 0.186568326 | 0.17844187 |
| 436,400 | 0.186539223 | 0.17838409 |
| 436,500 | 0.186241309 | 0.1784109 |
| 436,600 | 0.186327563 | 0.17835229 |
| 436,700 | 0.18648665 | 0.17839552 |
| 436,800 | 0.186353707 | 0.17837738 |
| 436,900 | 0.186428327 | 0.17837083 |
| 437,000 | 0.186455443 | 0.17837622 |
| 437,100 | 0.18646498 | 0.17835852 |
| 437,200 | 0.186423663 | 0.17833469 |
| 437,300 | 0.186452621 | 0.17840562 |
| 437,400 | 0.186476939 | 0.17835824 |
| 437,500 | 0.186575627 | 0.17838881 |
| 437,600 | 0.186631807 | 0.17836393 |
| 437,700 | 0.18642516 | 0.17839369 |
| 437,800 | 0.186403712 | 0.17836215 |
| 437,900 | 0.186410216 | 0.1784058 |
| 438,000 | 0.186481419 | 0.17839714 |
| 438,100 | 0.18638268 | 0.17835251 |
| 438,200 | 0.186417645 | 0.17837445 |
| 438,300 | 0.186423378 | 0.17834583 |
| 438,400 | 0.186293716 | 0.17837589 |
| 438,500 | 0.186332206 | 0.17836104 |
| 438,600 | 0.186270618 | 0.178393 |
| 438,700 | 0.186129894 | 0.17834691 |
| 438,800 | 0.186195611 | 0.17838806 |
| 438,900 | 0.186311342 | 0.17833636 |
| 439,000 | 0.186138861 | 0.17840725 |
| 439,100 | 0.18632235 | 0.17838361 |
| 439,200 | 0.186253341 | 0.17836124 |
| 439,300 | 0.186433145 | 0.17842827 |
| 439,400 | 0.186265421 | 0.17837946 |
| 439,500 | 0.18629832 | 0.17842363 |
| 439,600 | 0.186189978 | 0.17841113 |
| 439,700 | 0.186214373 | 0.17837355 |
| 439,800 | 0.186099586 | 0.17836289 |
| 439,900 | 0.18630642 | 0.17839647 |
| 440,000 | 0.186259197 | 0.17840375 |
| 440,100 | 0.186044767 | 0.17832668 |
| 440,200 | 0.186003298 | 0.17829223 |
| 440,300 | 0.186222925 | 0.17837806 |
| 440,400 | 0.186082655 | 0.1784228 |
| 440,500 | 0.186116229 | 0.17842939 |
| 440,600 | 0.186217647 | 0.17840723 |
| 440,700 | 0.186110635 | 0.17846205 |
| 440,800 | 0.186083261 | 0.17846882 |
| 440,900 | 0.185793806 | 0.17851326 |
| 441,000 | 0.185898215 | 0.17834305 |
| 441,100 | 0.185764785 | 0.17832567 |

| | | |
|---|---|---|
| 441,200 | 0.185964094 | 0.1783791 |
| 441,300 | 0.186014811 | 0.17824132 |
| 441,400 | 0.186127201 | 0.17832188 |
| 441,500 | 0.185989921 | 0.17833366 |
| 441,600 | 0.18605016 | 0.17835894 |
| 441,700 | 0.185801384 | 0.17824624 |
| 441,800 | 0.185913204 | 0.17836539 |
| 441,900 | 0.185900615 | 0.17833615 |
| 442,000 | 0.185855306 | 0.17829099 |
| 442,100 | 0.185865315 | 0.17832244 |
| 442,200 | 0.185727672 | 0.17840746 |
| 442,300 | 0.185856695 | 0.17832287 |
| 442,400 | 0.185687235 | 0.1782555 |
| 442,500 | 0.185797902 | 0.17834646 |
| 442,600 | 0.185801883 | 0.17827493 |
| 442,700 | 0.185695832 | 0.17838803 |
| 442,800 | 0.185466463 | 0.17829569 |
| 442,900 | 0.185595892 | 0.17830255 |
| 443,000 | 0.185515434 | 0.17836119 |
| 443,100 | 0.185423322 | 0.17838013 |
| 443,200 | 0.185637522 | 0.17833431 |
| 443,300 | 0.185491019 | 0.17829335 |
| 443,400 | 0.185551824 | 0.1783241 |
| 443,500 | 0.185373279 | 0.17837905 |
| 443,600 | 0.185598861 | 0.17834011 |
| 443,700 | 0.185508283 | 0.17836905 |
| 443,800 | 0.185399202 | 0.17831993 |
| 443,900 | 0.185185486 | 0.17836181 |
| 444,000 | 0.185408808 | 0.17832744 |
| 444,100 | 0.185210837 | 0.17830839 |
| 444,200 | 0.185210119 | 0.17834883 |
| 444,300 | 0.185396301 | 0.17845976 |
| 444,400 | 0.185299874 | 0.17830966 |
| 444,500 | 0.185368317 | 0.1783912 |
| 444,600 | 0.185260802 | 0.17835519 |
| 444,700 | 0.185259767 | 0.17835393 |
| 444,800 | 0.185234087 | 0.17837162 |
| 444,900 | 0.185129601 | 0.17827521 |
| 445,000 | 0.185025275 | 0.17829977 |
| 445,100 | 0.185082962 | 0.17838054 |
| 445,200 | 0.185147799 | 0.17836359 |
| 445,300 | 0.185042023 | 0.17837144 |
| 445,400 | 0.185111825 | 0.17834641 |
| 445,500 | 0.185276996 | 0.17832239 |
| 445,600 | 0.1852118 | 0.17836709 |
| 445,700 | 0.185122861 | 0.17835043 |
| 445,800 | 0.185167156 | 0.17841637 |
| 445,900 | 0.185103916 | 0.17835205 |
| 446,000 | 0.185025753 | 0.17835428 |
| 446,100 | 0.185065375 | 0.17839685 |
| 446,200 | 0.185003086 | 0.17841694 |
| 446,300 | 0.184973264 | 0.17835812 |
| 446,400 | 0.185071291 | 0.17827829 |
| 446,500 | 0.184933463 | 0.17830523 |
| 446,600 | 0.184840306 | 0.17837066 |
| 446,700 | 0.184926902 | 0.17832006 |
| 446,800 | 0.184956698 | 0.17829887 |
| 446,900 | 0.184898485 | 0.17835719 |
| 447,000 | 0.184818824 | 0.17837032 |
| 447,100 | 0.184963286 | 0.17832683 |
| 447,200 | 0.184877934 | 0.17826662 |
| 447,300 | 0.184812096 | 0.1783884 |
| 447,400 | 0.184652095 | 0.17832765 |
| 447,500 | 0.18464468 | 0.17842524 |
| 447,600 | 0.184611663 | 0.17826216 |
| 447,700 | 0.184609917 | 0.17826717 |
| 447,800 | 0.184694604 | 0.17834829 |

| | | |
|---|---|---|
| 447,900 | 0.184749087 | 0.17835304 |
| 448,000 | 0.184616873 | 0.17835956 |
| 448,100 | 0.184649854 | 0.17831658 |
| 448,200 | 0.184877524 | 0.17826871 |
| 448,300 | 0.184612478 | 0.17830494 |
| 448,400 | 0.184572874 | 0.17824793 |
| 448,500 | 0.184560763 | 0.1782978 |
| 448,600 | 0.184635069 | 0.17833371 |
| 448,700 | 0.184511618 | 0.17834852 |
| 448,800 | 0.184542133 | 0.17826217 |
| 448,900 | 0.184559686 | 0.1783015 |
| 449,000 | 0.184486656 | 0.17826212 |
| 449,100 | 0.184565337 | 0.17836511 |
| 449,200 | 0.184738937 | 0.17828655 |
| 449,300 | 0.184671203 | 0.17830438 |
| 449,400 | 0.184584799 | 0.17833717 |
| 449,500 | 0.18464133 | 0.17833207 |
| 449,600 | 0.184715822 | 0.17838709 |
| 449,700 | 0.184462318 | 0.17838855 |
| 449,800 | 0.184648484 | 0.17828483 |
| 449,900 | 0.184699072 | 0.17835579 |
| 450,000 | 0.184560248 | 0.17842943 |
| 450,100 | 0.184597584 | 0.17833909 |
| 450,200 | 0.184511865 | 0.17835618 |
| 450,300 | 0.184532037 | 0.17839253 |
| 450,400 | 0.184592483 | 0.17829562 |
| 450,500 | 0.184547686 | 0.17834525 |
| 450,600 | 0.184621195 | 0.17829035 |
| 450,700 | 0.184530881 | 0.17834059 |
| 450,800 | 0.184467284 | 0.1783887 |
| 450,900 | 0.184503734 | 0.17827783 |
| 451,000 | 0.184469709 | 0.17839661 |
| 451,100 | 0.184415747 | 0.17837044 |
| 451,200 | 0.184538217 | 0.17828194 |
| 451,300 | 0.184352799 | 0.17832872 |
| 451,400 | 0.184522378 | 0.17819607 |
| 451,500 | 0.184367784 | 0.1783212 |
| 451,600 | 0.184291726 | 0.17827737 |
| 451,700 | 0.184235072 | 0.1783278 |
| 451,800 | 0.184379282 | 0.17831879 |
| 451,900 | 0.184367898 | 0.17832118 |
| 452,000 | 0.184513341 | 0.17840346 |
| 452,100 | 0.184275443 | 0.17842614 |
| 452,200 | 0.184380687 | 0.17830087 |
| 452,300 | 0.184255305 | 0.17830822 |
| 452,400 | 0.184195501 | 0.17827762 |
| 452,500 | 0.184285273 | 0.17835546 |
| 452,600 | 0.18412123 | 0.17829695 |
| 452,700 | 0.184033037 | 0.17834927 |
| 452,800 | 0.184096269 | 0.17828611 |
| 452,900 | 0.184239855 | 0.178423 |
| 453,000 | 0.18415394 | 0.17834515 |
| 453,100 | 0.184265156 | 0.17830164 |
| 453,200 | 0.184408124 | 0.17837608 |
| 453,300 | 0.184178013 | 0.17830121 |
| 453,400 | 0.184251549 | 0.17835598 |
| 453,500 | 0.184179423 | 0.17844323 |
| 453,600 | 0.184294403 | 0.17831503 |
| 453,700 | 0.183984239 | 0.17832595 |
| 453,800 | 0.184169684 | 0.17814645 |
| 453,900 | 0.18417771 | 0.17833362 |
| 454,000 | 0.18428013 | 0.17832427 |
| 454,100 | 0.184015027 | 0.17830399 |
| 454,200 | 0.183926713 | 0.17825227 |
| 454,300 | 0.184049103 | 0.17834385 |
| 454,400 | 0.18392269 | 0.17833736 |
| 454,500 | 0.184066227 | 0.17836567 |

| | | |
|---|---|---|
| 454,600 | 0.183932354 | 0.17830575 |
| 454,700 | 0.18401388 | 0.17830748 |
| 454,800 | 0.183942092 | 0.17835328 |
| 454,900 | 0.183831805 | 0.17831962 |
| 455,000 | 0.183827586 | 0.17841944 |
| 455,100 | 0.183918317 | 0.17829171 |
| 455,200 | 0.183820612 | 0.17825343 |
| 455,300 | 0.183766155 | 0.17831249 |
| 455,400 | 0.183904112 | 0.17824857 |
| 455,500 | 0.183687034 | 0.17829779 |
| 455,600 | 0.183751125 | 0.1783642 |
| 455,700 | 0.183888314 | 0.17833068 |
| 455,800 | 0.183735901 | 0.17826062 |
| 455,900 | 0.183900574 | 0.17828344 |
| 456,000 | 0.18366468 | 0.17828606 |
| 456,100 | 0.183720609 | 0.1782258 |
| 456,200 | 0.183790226 | 0.17833376 |
| 456,300 | 0.183721306 | 0.17823672 |
| 456,400 | 0.183692802 | 0.17826448 |
| 456,500 | 0.183484254 | 0.17827407 |
| 456,600 | 0.183596972 | 0.17831768 |
| 456,700 | 0.183630557 | 0.17820708 |
| 456,800 | 0.183518849 | 0.17819861 |
| 456,900 | 0.183623735 | 0.17826761 |
| 457,000 | 0.183668315 | 0.17820673 |
| 457,100 | 0.183607988 | 0.17827087 |
| 457,200 | 0.183533916 | 0.1782842 |
| 457,300 | 0.183483446 | 0.17822504 |
| 457,400 | 0.183573356 | 0.17824536 |
| 457,500 | 0.183588977 | 0.17817611 |
| 457,600 | 0.18349293 | 0.17826788 |
| 457,700 | 0.183474173 | 0.17828372 |
| 457,800 | 0.183534578 | 0.17834107 |
| 457,900 | 0.183595282 | 0.17832357 |
| 458,000 | 0.183579071 | 0.17821575 |
| 458,100 | 0.18340402 | 0.17833094 |
| 458,200 | 0.183616998 | 0.17825638 |
| 458,300 | 0.183343123 | 0.17822388 |
| 458,400 | 0.183362764 | 0.17833133 |
| 458,500 | 0.18324362 | 0.17827144 |
| 458,600 | 0.183328886 | 0.17832479 |
| 458,700 | 0.183484709 | 0.17829759 |
| 458,800 | 0.18347443 | 0.17835538 |
| 458,900 | 0.183330868 | 0.17835769 |
| 459,000 | 0.183327215 | 0.17830282 |
| 459,100 | 0.183383238 | 0.17824345 |
| 459,200 | 0.183373991 | 0.17833724 |
| 459,300 | 0.183250101 | 0.17838826 |
| 459,400 | 0.183327325 | 0.17828815 |
| 459,500 | 0.18323792 | 0.17829819 |
| 459,600 | 0.183197137 | 0.17830634 |
| 459,700 | 0.183243554 | 0.17830268 |
| 459,800 | 0.183228724 | 0.17830781 |
| 459,900 | 0.183254577 | 0.17835379 |
| 460,000 | 0.183239509 | 0.17829633 |
| 460,100 | 0.183031916 | 0.1782288 |
| 460,200 | 0.183146334 | 0.17826574 |
| 460,300 | 0.18312776 | 0.17822878 |
| 460,400 | 0.183089687 | 0.17828128 |
| 460,500 | 0.183093139 | 0.17835772 |
| 460,600 | 0.183053962 | 0.17839148 |
| 460,700 | 0.183151151 | 0.17838533 |
| 460,800 | 0.183223648 | 0.17833473 |
| 460,900 | 0.183139717 | 0.17830912 |
| 461,000 | 0.183020932 | 0.17827731 |
| 461,100 | 0.18300694 | 0.17829578 |
| 461,200 | 0.182943395 | 0.17832301 |

| | | |
|---|---|---|
| 461,300 | 0.183073099 | 0.17831267 |
| 461,400 | 0.182930664 | 0.17838157 |
| 461,500 | 0.182987605 | 0.17832524 |
| 461,600 | 0.183016596 | 0.17837627 |
| 461,700 | 0.182957485 | 0.17831482 |
| 461,800 | 0.18287197 | 0.17828835 |
| 461,900 | 0.182816873 | 0.17827871 |
| 462,000 | 0.182898735 | 0.17824421 |
| 462,100 | 0.182984372 | 0.17826425 |
| 462,200 | 0.182967389 | 0.17833277 |
| 462,300 | 0.182882573 | 0.17831818 |
| 462,400 | 0.18285902 | 0.17823821 |
| 462,500 | 0.182961278 | 0.17830872 |
| 462,600 | 0.182963518 | 0.178339 |
| 462,700 | 0.182875161 | 0.17831768 |
| 462,800 | 0.182776666 | 0.17835483 |
| 462,900 | 0.182795733 | 0.17829666 |
| 463,000 | 0.182873445 | 0.17825857 |
| 463,100 | 0.182783422 | 0.17829703 |
| 463,200 | 0.182772987 | 0.17831876 |
| 463,300 | 0.182916987 | 0.17833019 |
| 463,400 | 0.182857661 | 0.17833777 |
| 463,500 | 0.182869197 | 0.17830514 |
| 463,600 | 0.182732105 | 0.17834549 |
| 463,700 | 0.182800081 | 0.17835444 |
| 463,800 | 0.182757458 | 0.17836805 |
| 463,900 | 0.182822696 | 0.17836368 |
| 464,000 | 0.182703339 | 0.17838939 |
| 464,100 | 0.182722419 | 0.1784422 |
| 464,200 | 0.18261214 | 0.17832266 |
| 464,300 | 0.182506856 | 0.17831143 |
| 464,400 | 0.182435012 | 0.17831292 |
| 464,500 | 0.182618946 | 0.17837704 |
| 464,600 | 0.182637609 | 0.17830978 |
| 464,700 | 0.182579423 | 0.17837998 |
| 464,800 | 0.182493298 | 0.17837093 |
| 464,900 | 0.182472837 | 0.17842211 |
| 465,000 | 0.182449846 | 0.17838609 |
| 465,100 | 0.182392398 | 0.17831627 |
| 465,200 | 0.182338497 | 0.17822438 |
| 465,300 | 0.182218828 | 0.17824062 |
| 465,400 | 0.182293638 | 0.17835532 |
| 465,500 | 0.182300963 | 0.17839437 |
| 465,600 | 0.182174335 | 0.17838884 |
| 465,700 | 0.182358235 | 0.17836284 |
| 465,800 | 0.182449638 | 0.17829537 |
| 465,900 | 0.182424273 | 0.17832908 |
| 466,000 | 0.182381374 | 0.17834742 |
| 466,100 | 0.182308374 | 0.17834803 |
| 466,200 | 0.182350807 | 0.17825633 |
| 466,300 | 0.182316185 | 0.178393 |
| 466,400 | 0.18226368 | 0.17828754 |
| 466,500 | 0.182358161 | 0.17826291 |
| 466,600 | 0.182432371 | 0.17840451 |
| 466,700 | 0.182180241 | 0.17831895 |
| 466,800 | 0.18218767 | 0.17819816 |
| 466,900 | 0.182219778 | 0.17830532 |
| 467,000 | 0.182314802 | 0.17825705 |
| 467,100 | 0.182380525 | 0.17836659 |
| 467,200 | 0.182245825 | 0.17826994 |
| 467,300 | 0.182216693 | 0.17830441 |
| 467,400 | 0.18216651 | 0.17833525 |
| 467,500 | 0.182261847 | 0.17820708 |
| 467,600 | 0.18217175 | 0.17830704 |
| 467,700 | 0.182207347 | 0.17831797 |
| 467,800 | 0.182242664 | 0.17829385 |
| 467,900 | 0.182250977 | 0.17836384 |

| | | |
|---|---|---|
| 468,000 | 0.182230295 | 0.17831684 |
| 468,100 | 0.182168484 | 0.17835882 |
| 468,200 | 0.181994689 | 0.17831594 |
| 468,300 | 0.182054657 | 0.17827666 |
| 468,400 | 0.18209047 | 0.17827104 |
| 468,500 | 0.182098689 | 0.1782594 |
| 468,600 | 0.182108775 | 0.17822723 |
| 468,700 | 0.18192097 | 0.17834494 |
| 468,800 | 0.181963411 | 0.17833523 |
| 468,900 | 0.181948395 | 0.1782703 |
| 469,000 | 0.181942828 | 0.17826888 |
| 469,100 | 0.181910931 | 0.17831323 |
| 469,200 | 0.181980958 | 0.1781914 |
| 469,300 | 0.181897613 | 0.17818138 |
| 469,400 | 0.181898112 | 0.17828883 |
| 469,500 | 0.181842248 | 0.17825592 |
| 469,600 | 0.181856197 | 0.17831534 |
| 469,700 | 0.181831047 | 0.17825014 |
| 469,800 | 0.181714001 | 0.17827721 |
| 469,900 | 0.181869608 | 0.17834443 |
| 470,000 | 0.181836983 | 0.17830221 |
| 470,100 | 0.181725547 | 0.17841109 |
| 470,200 | 0.181722325 | 0.17834083 |
| 470,300 | 0.181825155 | 0.17837809 |
| 470,400 | 0.18185829 | 0.17835776 |
| 470,500 | 0.181882543 | 0.17828554 |
| 470,600 | 0.181665618 | 0.17831755 |
| 470,700 | 0.181727345 | 0.17838032 |
| 470,800 | 0.181594797 | 0.17843728 |
| 470,900 | 0.181604085 | 0.17830345 |
| 471,000 | 0.181631292 | 0.17824974 |
| 471,100 | 0.181621497 | 0.17821417 |
| 471,200 | 0.181741156 | 0.17831132 |
| 471,300 | 0.181635711 | 0.17830359 |
| 471,400 | 0.181564035 | 0.17833334 |
| 471,500 | 0.181596269 | 0.17828489 |
| 471,600 | 0.18152757 | 0.1783832 |
| 471,700 | 0.181444068 | 0.1782715 |
| 471,800 | 0.181476606 | 0.17828197 |
| 471,900 | 0.181581455 | 0.17837309 |
| 472,000 | 0.181427629 | 0.17827769 |
| 472,100 | 0.181431187 | 0.17834637 |
| 472,200 | 0.181335357 | 0.17833378 |
| 472,300 | 0.181402795 | 0.17836707 |
| 472,400 | 0.181460829 | 0.17840979 |
| 472,500 | 0.181491747 | 0.17829192 |
| 472,600 | 0.181534772 | 0.17832735 |
| 472,700 | 0.181490177 | 0.17838421 |
| 472,800 | 0.18158462 | 0.178339 |
| 472,900 | 0.181471339 | 0.17828986 |
| 473,000 | 0.181501558 | 0.17830311 |
| 473,100 | 0.181552985 | 0.17833396 |
| 473,200 | 0.181540509 | 0.17837302 |
| 473,300 | 0.181324568 | 0.17830666 |
| 473,400 | 0.181356687 | 0.17831825 |
| 473,500 | 0.181253626 | 0.17832902 |
| 473,600 | 0.181132404 | 0.17835693 |
| 473,700 | 0.181262907 | 0.17842137 |
| 473,800 | 0.181147252 | 0.17840079 |
| 473,900 | 0.181119201 | 0.178373 |
| 474,000 | 0.181170059 | 0.17839875 |
| 474,100 | 0.181308212 | 0.17836108 |
| 474,200 | 0.181241947 | 0.1783284 |
| 474,300 | 0.181230596 | 0.17833088 |
| 474,400 | 0.181222188 | 0.17838331 |
| 474,500 | 0.181189616 | 0.17837697 |
| 474,600 | 0.181063462 | 0.17841252 |

| | | |
|---|---|---|
| 474,700 | 0.181037202 | 0.17842632 |
| 474,800 | 0.181072555 | 0.17835459 |
| 474,900 | 0.181246303 | 0.17834384 |
| 475,000 | 0.181159008 | 0.17830572 |
| 475,100 | 0.181101037 | 0.17829087 |
| 475,200 | 0.180910638 | 0.17834208 |
| 475,300 | 0.181159959 | 0.17839282 |
| 475,400 | 0.181045128 | 0.1782915 |
| 475,500 | 0.181004646 | 0.17836296 |
| 475,600 | 0.180971334 | 0.17835348 |
| 475,700 | 0.181026187 | 0.17839288 |
| 475,800 | 0.18090674 | 0.17840353 |
| 475,900 | 0.180804042 | 0.17831458 |
| 476,000 | 0.18088543 | 0.17837019 |
| 476,100 | 0.18083937 | 0.17841925 |
| 476,200 | 0.180913274 | 0.17836942 |
| 476,300 | 0.18085071 | 0.1783257 |
| 476,400 | 0.180791393 | 0.1783559 |
| 476,500 | 0.180894643 | 0.17835537 |
| 476,600 | 0.180895565 | 0.17837536 |
| 476,700 | 0.180944581 | 0.17833875 |
| 476,800 | 0.180796871 | 0.17841756 |
| 476,900 | 0.180787951 | 0.17834667 |
| 477,000 | 0.180823436 | 0.17837036 |
| 477,100 | 0.180859408 | 0.17837966 |
| 477,200 | 0.180818585 | 0.17833185 |
| 477,300 | 0.180798439 | 0.17828202 |
| 477,400 | 0.180732854 | 0.17834223 |
| 477,500 | 0.180897368 | 0.17831771 |
| 477,600 | 0.180759582 | 0.17833973 |
| 477,700 | 0.1807329 | 0.17839634 |
| 477,800 | 0.180583986 | 0.17835056 |
| 477,900 | 0.180547765 | 0.17835022 |
| 478,000 | 0.180604034 | 0.17834953 |
| 478,100 | 0.180636041 | 0.17832561 |
| 478,200 | 0.180573861 | 0.17842383 |
| 478,300 | 0.180604354 | 0.17830963 |
| 478,400 | 0.180664422 | 0.17830118 |
| 478,500 | 0.180651094 | 0.17826284 |
| 478,600 | 0.180598692 | 0.17833714 |
| 478,700 | 0.180429787 | 0.1783527 |
| 478,800 | 0.180550102 | 0.17827638 |
| 478,900 | 0.180346868 | 0.17820767 |
| 479,000 | 0.18046429 | 0.17827511 |
| 479,100 | 0.180400493 | 0.17833093 |
| 479,200 | 0.180460054 | 0.1782419 |
| 479,300 | 0.180472269 | 0.17829855 |
| 479,400 | 0.180232734 | 0.17831464 |
| 479,500 | 0.180388515 | 0.17822079 |
| 479,600 | 0.180405234 | 0.1782855 |
| 479,700 | 0.18034977 | 0.17822848 |
| 479,800 | 0.180281556 | 0.17834421 |
| 479,900 | 0.18036939 | 0.17836979 |
| 480,000 | 0.180434873 | 0.178319 |
| 480,100 | 0.180311138 | 0.17825838 |
| 480,200 | 0.180314594 | 0.17827205 |
| 480,300 | 0.18022098 | 0.17825657 |
| 480,400 | 0.180257876 | 0.17831317 |
| 480,500 | 0.180387524 | 0.17829479 |
| 480,600 | 0.180154566 | 0.17830957 |
| 480,700 | 0.180172542 | 0.17829105 |
| 480,800 | 0.180112182 | 0.17837724 |
| 480,900 | 0.180086771 | 0.17830561 |
| 481,000 | 0.180119596 | 0.17834573 |
| 481,100 | 0.180139268 | 0.17834153 |
| 481,200 | 0.179950764 | 0.1782241 |
| 481,300 | 0.180011661 | 0.17825168 |

| | | |
|---|---|---|
| 481,400 | 0.180044314 | 0.1782732 |
| 481,500 | 0.180043678 | 0.17835191 |
| 481,600 | 0.179908032 | 0.17822556 |
| 481,700 | 0.180053739 | 0.17829829 |
| 481,800 | 0.180062811 | 0.17828063 |
| 481,900 | 0.17999434 | 0.17830342 |
| 482,000 | 0.179804106 | 0.17825317 |
| 482,100 | 0.179938941 | 0.17834656 |
| 482,200 | 0.179873277 | 0.17832515 |
| 482,300 | 0.179980468 | 0.17822628 |
| 482,400 | 0.179972043 | 0.17827634 |
| 482,500 | 0.179886819 | 0.17825969 |
| 482,600 | 0.179856479 | 0.17822747 |
| 482,700 | 0.179819494 | 0.1782474 |
| 482,800 | 0.179890821 | 0.1782674 |
| 482,900 | 0.179850232 | 0.17824343 |
| 483,000 | 0.179925533 | 0.17825375 |
| 483,100 | 0.179825786 | 0.17826891 |
| 483,200 | 0.179857897 | 0.17833404 |
| 483,300 | 0.179840604 | 0.17831051 |
| 483,400 | 0.179767213 | 0.17829306 |
| 483,500 | 0.179853054 | 0.1783215 |
| 483,600 | 0.179690582 | 0.17836462 |
| 483,700 | 0.179670778 | 0.17834734 |
| 483,800 | 0.179655356 | 0.1783 |
| 483,900 | 0.179694314 | 0.17828029 |
| 484,000 | 0.1797355 | 0.17826979 |
| 484,100 | 0.179910352 | 0.17830859 |
| 484,200 | 0.17979381 | 0.17833158 |
| 484,300 | 0.179758344 | 0.1782738 |
| 484,400 | 0.179738776 | 0.17820614 |
| 484,500 | 0.179657236 | 0.17830481 |
| 484,600 | 0.17969411 | 0.178359 |
| 484,700 | 0.179642155 | 0.17827869 |
| 484,800 | 0.179587051 | 0.17824946 |
| 484,900 | 0.179664415 | 0.17826906 |
| 485,000 | 0.179583551 | 0.17827148 |
| 485,100 | 0.179574602 | 0.17829154 |
| 485,200 | 0.179637065 | 0.17829641 |
| 485,300 | 0.179531933 | 0.17826524 |
| 485,400 | 0.179478779 | 0.1782708 |
| 485,500 | 0.179676139 | 0.17824356 |
| 485,600 | 0.179417643 | 0.17830549 |
| 485,700 | 0.179498676 | 0.1782975 |
| 485,800 | 0.179667279 | 0.17834125 |
| 485,900 | 0.179466843 | 0.17834099 |
| 486,000 | 0.179488589 | 0.17837895 |
| 486,100 | 0.179437735 | 0.17841859 |
| 486,200 | 0.17949679 | 0.17842818 |
| 486,300 | 0.179481277 | 0.17828387 |
| 486,400 | 0.179575052 | 0.17836637 |
| 486,500 | 0.179423792 | 0.17829718 |
| 486,600 | 0.179509202 | 0.1782842 |
| 486,700 | 0.179438342 | 0.17833744 |
| 486,800 | 0.179666137 | 0.17832237 |
| 486,900 | 0.179463185 | 0.17830999 |
| 487,000 | 0.179525753 | 0.1782392 |
| 487,100 | 0.179432722 | 0.17831019 |
| 487,200 | 0.179378151 | 0.17825559 |
| 487,300 | 0.179450324 | 0.17824611 |
| 487,400 | 0.179571992 | 0.17829378 |
| 487,500 | 0.179268248 | 0.17822219 |
| 487,600 | 0.179523057 | 0.17832631 |
| 487,700 | 0.179372434 | 0.17835185 |
| 487,800 | 0.179327624 | 0.17831639 |
| 487,900 | 0.17925204 | 0.1783036 |
| 488,000 | 0.179471654 | 0.1782024 |

| | | |
|---|---|---|
| 488,100 | 0.179340967 | 0.17829407 |
| 488,200 | 0.179219215 | 0.17831468 |
| 488,300 | 0.179195372 | 0.17824824 |
| 488,400 | 0.179094057 | 0.17822351 |
| 488,500 | 0.179166899 | 0.17832479 |
| 488,600 | 0.179092387 | 0.17832604 |
| 488,700 | 0.17920106 | 0.17822302 |
| 488,800 | 0.179186787 | 0.17820137 |
| 488,900 | 0.17918667 | 0.17827333 |
| 489,000 | 0.179185052 | 0.17832433 |
| 489,100 | 0.17921696 | 0.17826781 |
| 489,200 | 0.179047042 | 0.17828906 |
| 489,300 | 0.179297551 | 0.17831448 |
| 489,400 | 0.179185045 | 0.17831092 |
| 489,500 | 0.179137973 | 0.17823703 |
| 489,600 | 0.179189029 | 0.17832041 |
| 489,700 | 0.179271644 | 0.1782176 |
| 489,800 | 0.179154414 | 0.17823276 |
| 489,900 | 0.179131752 | 0.17835789 |
| 490,000 | 0.179139253 | 0.17824853 |
| 490,100 | 0.179201992 | 0.17821983 |
| 490,200 | 0.17911585 | 0.17822365 |
| 490,300 | 0.179142558 | 0.17830136 |
| 490,400 | 0.179022273 | 0.1782233 |
| 490,500 | 0.179048191 | 0.17836333 |
| 490,600 | 0.179100658 | 0.17837121 |
| 490,700 | 0.179078402 | 0.17836385 |
| 490,800 | 0.178967951 | 0.17822726 |
| 490,900 | 0.179007664 | 0.17822745 |
| 491,000 | 0.178938767 | 0.17825982 |
| 491,100 | 0.179086647 | 0.17826355 |
| 491,200 | 0.178792464 | 0.17826527 |
| 491,300 | 0.178880123 | 0.17826372 |
| 491,400 | 0.17883468 | 0.17830398 |
| 491,500 | 0.178975694 | 0.17823082 |
| 491,600 | 0.17880592 | 0.17831903 |
| 491,700 | 0.178777981 | 0.17829897 |
| 491,800 | 0.178843826 | 0.17825759 |
| 491,900 | 0.178703044 | 0.1783331 |
| 492,000 | 0.178904268 | 0.17826742 |
| 492,100 | 0.17884428 | 0.1783393 |
| 492,200 | 0.178911421 | 0.17836355 |
| 492,300 | 0.178779198 | 0.17835407 |
| 492,400 | 0.178672874 | 0.1783535 |
| 492,500 | 0.178757709 | 0.17828218 |
| 492,600 | 0.17886946 | 0.17832517 |
| 492,700 | 0.178851313 | 0.17835932 |
| 492,800 | 0.178789784 | 0.1783248 |
| 492,900 | 0.178815698 | 0.17828008 |
| 493,000 | 0.178747048 | 0.1782898 |
| 493,100 | 0.178815521 | 0.17836101 |
| 493,200 | 0.17861366 | 0.1784003 |
| 493,300 | 0.178817106 | 0.17833594 |
| 493,400 | 0.178786945 | 0.17838295 |
| 493,500 | 0.178724774 | 0.17824932 |
| 493,600 | 0.178828129 | 0.17833009 |
| 493,700 | 0.178834088 | 0.17831882 |
| 493,800 | 0.178616746 | 0.17832299 |
| 493,900 | 0.178544354 | 0.1783237 |
| 494,000 | 0.178513117 | 0.17824025 |
| 494,100 | 0.178512374 | 0.1783545 |
| 494,200 | 0.178598602 | 0.1783195 |
| 494,300 | 0.178580519 | 0.17844885 |
| 494,400 | 0.17862034 | 0.17834564 |
| 494,500 | 0.178610913 | 0.1782835 |
| 494,600 | 0.178557738 | 0.17829428 |
| 494,700 | 0.178480138 | 0.17833152 |

| | | |
|---|---|---|
| 494,800 | 0.17859826 | 0.17834621 |
| 494,900 | 0.17842475 | 0.17833495 |
| 495,000 | 0.17848094 | 0.17841399 |
| 495,100 | 0.178432639 | 0.17827948 |
| 495,200 | 0.178507082 | 0.17834876 |
| 495,300 | 0.17842351 | 0.17830761 |
| 495,400 | 0.178428403 | 0.178372 |
| 495,500 | 0.178450743 | 0.17837325 |
| 495,600 | 0.178497924 | 0.17833388 |
| 495,700 | 0.178356593 | 0.17832564 |
| 495,800 | 0.178298151 | 0.17827629 |
| 495,900 | 0.178202852 | 0.17829647 |
| 496,000 | 0.178366186 | 0.17838891 |
| 496,100 | 0.178209234 | 0.17830351 |
| 496,200 | 0.17834966 | 0.17828951 |
| 496,300 | 0.178352687 | 0.17837285 |
| 496,400 | 0.178296309 | 0.17830939 |
| 496,500 | 0.17842461 | 0.17823926 |
| 496,600 | 0.178226424 | 0.1783203 |
| 496,700 | 0.178311111 | 0.17831102 |
| 496,800 | 0.178457999 | 0.17831327 |
| 496,900 | 0.178235883 | 0.17830845 |
| 497,000 | 0.178248345 | 0.1784624 |
| 497,100 | 0.178285303 | 0.1784021 |
| 497,200 | 0.178092156 | 0.17835664 |
| 497,300 | 0.178151607 | 0.17837015 |
| 497,400 | 0.178242298 | 0.1783661 |
| 497,500 | 0.178220511 | 0.17838759 |
| 497,600 | 0.178086893 | 0.17831285 |
| 497,700 | 0.178153541 | 0.17832884 |
| 497,800 | 0.178018892 | 0.17835319 |
| 497,900 | 0.178027687 | 0.17837478 |
| 498,000 | 0.178122499 | 0.1783717 |
| 498,100 | 0.178158151 | 0.17834868 |
| 498,200 | 0.178207679 | 0.17831413 |
| 498,300 | 0.178041583 | 0.17836874 |
| 498,400 | 0.178141099 | 0.1784206 |
| 498,500 | 0.178110782 | 0.17836117 |
| 498,600 | 0.17800326 | 0.17820839 |
| 498,700 | 0.177965106 | 0.17829708 |
| 498,800 | 0.177889238 | 0.17824742 |
| 498,900 | 0.177895889 | 0.17830392 |
| 499,000 | 0.177917599 | 0.17835746 |
| 499,100 | 0.177936031 | 0.17832807 |
| 499,200 | 0.177819769 | 0.1783878 |
| 499,300 | 0.1777698 | 0.17829548 |
| 499,400 | 0.177741916 | 0.17832546 |
| 499,500 | 0.177709802 | 0.17833429 |
| 499,600 | 0.17778177 | 0.17827738 |
| 499,700 | 0.177790629 | 0.1784201 |
| 499,800 | 0.177895671 | 0.17838231 |
| 499,900 | 0.177858901 | 0.17836373 |
| 500,000 | 0.177644281 | 0.17834837 |
| 500,100 | 0.178308855 | 0.17803098 |

**Table S9. Genes located in the selected regions between Angora and wild rabbits**

| Gene |
|---|
| CADM1 |
| REXO2 |
| LOC108176944 |
| RBM7 |
| C1H11orf71 |
| ZBTB16 |
| LOC108176950 |
| HTR3A |
| ALG9 |
| PPP2R1B |
| SIK2 |
| SMCO4 |
| CCDC67 |
| NOX4 |
| GRM5 |
| INVS |
| ERP44 |
| STX17 |
| NR4A3 |
| TGFBR1 |
| COL15A1 |
| FAM168A |
| DENND5A |
| LOC100008770 |
| INSC |
| LOC100342427 |
| ANO3 |
| SLC5A12 |
| BBOX1 |
| KIF18A |
| METTL15 |
| WT1 |
| LOC103351180 |
| TCP11L1 |
| CSTF3 |
| RNF38 |
| LOC100344965 |
| ACCSL |
| EXT2 |
| ALX4 |
| LOC100343628 |
| LOC100343874 |
| LOC100344130 |
| LOC100340210 |
| LOC100344383 |
| ADAMTSL1 |
| PTPRD |
| SPATA6L |
| GLIS3 |
| MAMDC2 |
| TRPM3 |
| LOC103347974 |
| PRUNE2 |
| DAPK1 |
| DIRAS2 |
| PRDM10 |
| CLMP |
| LOC108176867 |
| GRIK4 |
| ARHGEF12 |
| TMEM136 |
| TRIM29 |
| FAM188B |
| INHBA |
| GLI3 |

TMEM243
DMTF1
DDX56
TMED4
OGDH
KLHL7
NUPL2
IGF2BP3
TRA2A
CCDC126
FAM221A
ADGRV1
FAM172A
FBXL7
OSMR
PRKAA1
RPL37
CARD6
C6
PLCXD3
ITGA1
ACTBL2
KIF2A
DIMT1
IPO11
LOC100354788
RNF180
LOC100355797
LOC100343903
LOC100344156
LOC100344410
LOC100344660
LOC100348578
LOC100344914
LOC100345168
LOC100345425
LOC100345683
LOC100345939
TRDN
LOC103349878
MYB
LOC108177481
LOC108177387
TNFAIP3
LOC108177388
LOC108177389
PERP
ARFGEF3
UTRN
GRM1
LOC108177494
TULP4
WTAP
ACAT2
PACRG
LOC108177393
LOC103349946
QKI
LOC108177498
T
LOC103349956
TMEM217
TBC1D22B
NT5E
SNX14
LOC103349811
LOC100344662

LOC108177446
LOC108177445
ZNF292
GJB7
SMIM8
C12H6orf163
CFAP206
SLC35A1
RARS2
ORC3
AKIRIN2
CNR1
RNGTT
PNRC1
SRSF12
PM20D2
GABRR1
GABRR2
UBE2J1
RRAGD
ANKRD6
LYRM2
LOC103349813
MDN1
CASP8AP2
GJA10
BACH2
EPHA7
FUT9
UFL1
FHL5
GPR63
NDUFAF4
KLHL32
GRIK2
LOC100346633
WASF1
FYN
C13H1orf87
TRABD2B
MKNK1
KNCN
RPS6KA1
SLC2A5
SLC2A7
CA6
LOC108177748
ENO1
RERE
MRPS14
LOC108177625
TNN
TNR
ATF6
DUSP12
FCRLB
LOC100359306
PAPPA2
ASTN1
WDR47
CLCC1
LOC100342719
LOC100343733
RNPC3
LOC103350128
COL6A5
LOC100354960

BBX
CBLB
LOC100359149
ZNF654
CGGBP1
VGLL3
LOC103350578
CADM2
CAPN7
SH3BP5
GBE1
ROBO1
LOC108178019
ROBO2
USP25
NCAM2
LOC108177928
GRIK1
LOC100339009
DIP2A
RFTN1
LOC103350352
EPHB1
LOC103350398
PPP2R3A
PLCL2
LOC103350353
MSL2
TBC1D5
SOX14
CLDN18
NME9
MRAS
LOC108177934
COPB2
RBP2
LOC108177933
RBP1
NMNAT3
CLSTN2
LOC108177916
HPS3
CP
RNF13
KCNH8
PPM1L
BCHE
TRNAV-AAC
MYNN
LRRC34
LRRIQ4
PHC3
PRKCI
NCEH1
MCRIP2
ECT2
LOC100356310
SPATA16
NLGN1
NAALADL2
TTC14
CCDC39
FXR1
LOC100341365
LOC108177978
SOX2
BCL6

LOC103350534
LOC103350471
FGF12
DLG1
ROPN1
CASR
ANKRD50
LOC103350844
FAT4
LOC108178131
LARP1B
JADE1
LOC108178134
SCLT1
LOC103350856
DCHS2
LOC100358706
NR3C2
ARHGAP10
RNF150
MGAT4D
CLGN
PABPC4L
UGT8
LOC103350741
SLC39A8
SLC18A1
ATP6V1B2
LZTS1
TSPAN5
C15H4orf22
TNIP3
QRFPR
TRPC3
SPATA5
SFMBT2
LOC103351031
LOC103351032
SIPA1L2
LOC108178160
LOC108178186
DISC1
PLXDC2
CCDC185
LOC108178200
SUSD4
HLX
1-Mar
GPATCH2
ESRRG
LOC108178206
KCNK2
LOC103350974
PPP2R5A
PLXNA2
CR1
CR2
KIAA1217
LGR6
RNPEP
TIMM17A
LOC108178224
LMOD1
BRINP3
VPS13C
SLTM
FAM63B

LOC103351108
TMOD3
LOC108178256
TMOD2
GLDN
BLOC1S6
SLC30A4
MGA
EIF2AK4
GPR176
NFATC4
LOC103351182
NYNRIN
CBLN3
LOC103351196
LOC103351201
TMEM260
LOC103351261
SLC35F4
C17H14orf37
OCA2
GABRG3
LOC103351269
UBE3A
LOC103351270
LOC103351277
LOC103351279
SNRPN
LOC108178324
LOC108178325
LOC108178365
NDN
MAGEL2
MKRN3
CHRNA7
APBA2
HERC1
SAMD8
DUSP13
KAT6B
MCU
LRRC20
NPFFR1
PPA1
SAR1A
TYSND1
AIFM2
LOC100343925
COL13A1
LOC108178478
LOC108178479
ZNF365
RTKN2
LOC103351358
HPSE2
ENTPD7
LOC100355645
CUTC
ABCC2
DNMBP
LOC103351405
LOC108178412
ERLIN1
CHUK
CWF19L1
LOC100346046
LOC100346306

WNT8B
SEC31B
NDUFB8
HIF1AN
PAX2
SLF2
SEMA4G
FBXW4
RBM20
LOC100343496
ZFAND4
8-Mar
MCMBP
PAFAH1B1
CRK
NXN
EFCAB5
AATF
ZSWIM7
TMEM92
COL1A1
LOC103351547
LOC103351548
PPP1R9B
SAMD14
PDK2
ITGA3
CDK5RAP3
LOC108178554
PRR15L
PNPO
SP2
PROKR1
APLF
WDPCP
BCL11A
LOC108176354
LOC103347677
NRXN1
C2H2orf61
TTC7A
CAMKMT
LBH
YPEL5
LOC103347812
ALK
NBAS
TBC1D19
NCOA1
DNAJC27
LOC108178981
LOC103345114
GABRG1
FZD3
LOC103345349
DOCK5
LOC103345404
RWDD4
TRAPPC11
FAM149A
FAT1
UNC5D
LOC103346906
RNF149
LOC103346939
LOC100345317
LOC108176011

LOC108176013
MALL
NPHP1
LOC103347473
BUB1
ACOXL
BCL2L11
LOC103347509
ANAPC1
MERTK
TMEM87B
FBLN7
ZC3H8
ZC3H6
TTL
POLR1B
PSD4
PAX8
RIN3
LOC103351663
SLC24A4
LOC103351700
STON2
CEP128
TTLL5
C20H14orf1
FLVCR2
LOC108178627
RAD51B
LOC108178630
RDH12
RDH11
FUT8
SPPL3
LOC103351757
CCDC60
PRR14L
LRRCC1
COMMD10
CA1
CA3
NECAB1
C3H8orf88
PGGT1B
CCDC112
SAMD12
CHSY3
SQLE
KIAA0196
NSMCE2
LOC100353158
ASAP1
KCNQ3
ZFAT
LOC103344959
LOC103348141
TRAPPC9
TGFBI
LECT2
UBE2D2
FEM1C
IL12B
LOC108176486
TTC1
PWWP2A
GABRB2
KCNIP1

GABRP
RANBP17
TLX3
NPM1
FBXW11
STK10
STC2
LOC103347964
CPEB4
C3H5orf47
LOC100343708
LOC103347973
FAM193B
LOC103348008
LOC100351987
B4GALT7
COL23A1
SPIDR
LOC108176496
PXDNL
PCMTD1
ST18
FAM150A
RB1CC1
ATP6V1H
RGS20
TCEA1
LYPLA1
LOC100356693
SOX17
LOC100355685
LYN
LOC108176501
PLAG1
CHCHD7
LOC103348029
SDCBP
NSMAF
LOC100008743
LOC108176505
LOC108176507
BHLHE22
LOC100343291
ARMC1
PDE7A
LOC100342098
LOC103348037
LOC108176508
COPS5
CSPP1
CPA6
LOC108176510
LOC103348047
SULF1
SLCO5A1
EYA1
STAU2
UBE2W
TCEB1
TMEM70
PI15
CRISPLD1
PEX2
TRMT6
MCM8
CRLS1
CFAP61

CRNKL1
NAA20
RIN2
MACROD2
JAG1
SLX4IP
MKKS
SNAP25
PAK5
PLCB4
PLCB1
LOC103348254
FMNL3
TMBIM6
NCKAP5L
RACGAP1
ASIC1
LIMA1
LOC103344945
POU6F1
DAZAP2
SMAGP
BIN2
CELA1
GALNT6
KRT5
LOC100352913
LOC100347394
LOC100347647
KRT2
KRT1
KRT77
LOC100348136
LOC100357568
LOC100357823
LOC100358083
LOC100358341
PYM1
DGKA
SPRYD4
GLS2
RBMS2
BAZ2A
C4H12orf66
C4H12orf56
E2F1
NECAB3
ACTL10
C4H20orf144
CBFA2T2
LOC100341256
LOC103348339
TMTC3
DUSP6
LOC103348378
POC1B
GALNT4
CCER1
EPYC
KERA
LOC108176669
LUM
DCN
ANKS1B
ANO4
SLC5A8
BTBD11

RIC8B
CKAP4
CCDC102A
ADGRG5
ADGRG1
ADGRG3
VAC14
LSM14A
FKBP2
MKL2
SCN3A
TTC21B
CERS6
SP5
ERICH2
GAD1
HAT1
RAPGEF4
CHCHD3
LOC108176864
LOC103348788
HNRNPA3
NFE2L2
AGPS
TTC30B
LOC100341262
PDE11A
LOC100353673
RBM45
TTN
ZNF385B
PPP1R1C
PDE1A
NCKAP1
GULP1
COL3A1
COL5A2
GLS
LOC108176936
SLC39A10
DNAH7
STK17B
LOC103348804
HECW2
GTF3C3
C7H2orf66
PGAP1
ANKRD44
LOC103348806
SF3B1
COQ10B
RFTN2
LOC103348807
BOLL
PLCL1
LOC100351153
LOC108176952
SPATS2L
KCTD18
SGO2
LOC103348854
LOC108176874
BZW1
CLK1
PPIL3
NIF3L1
ORC2

| |
|---|
| FAM126B |
| NDUFB3 |
| CFLAR |
| LOC103348916 |
| CDK15 |
| ZDBF2 |
| ADAM23 |
| CCNYL1 |
| LOC108176937 |
| PLEKHM3 |
| LOC108176881 |
| LOC100355936 |
| C7H2orf80 |
| IDH1 |
| PIKFYVE |
| ABCA12 |
| FN1 |
| XRCC5 |
| FEV |
| CFAP65 |
| NHEJ1 |
| CHPF |
| EPHA4 |
| SND1 |
| LOC108176816 |
| PAX3 |
| SGPP2 |
| FARSB |
| MOGAT1 |
| ACSL3 |
| LOC108176890 |
| PSMD1 |
| HTR2B |
| CADPS2 |
| LOC103348668 |
| RNF148 |
| RNF133 |
| SEMA3C |
| MAGI2 |
| LOC103348685 |

Table S10. Enrichment of genes located in the selected regions between Angora and wild rabbits

| Annotation Cluster | Enrichment Score: 2.43 | G | | Cou | P_Value | Benjamini |
|---|---|---|---|---|---|---|
| UP_KEYWORDS | Synapse | RT | | 11 | 1.10E-04 | 1.80E-02 |
| GOTERM_MF_DIRECT | extracellular ligand-gated ion channel activity | RT | | 7 | 1.40E-04 | 4.50E-02 |
| GOTERM_MF_DIRECT | GABA-A receptor activity | RT | | 6 | 4.60E-04 | 7.60E-02 |
| INTERPRO | Neurotransmitter-gated ion-channel, conserved site | RT | | 8 | 5.40E-04 | 1.30E-01 |
| UP_KEYWORDS | Cell junction | RT | | 13 | 6.10E-04 | 3.70E-02 |
| INTERPRO | Neurotransmitter-gated ion-channel transmembrane domain | RT | | 8 | 6.30E-04 | 1.30E-01 |
| INTERPRO | Neurotransmitter-gated ion-channel | RT | | 8 | 6.30E-04 | 1.30E-01 |
| INTERPRO | Neurotransmitter-gated ion-channel ligand-binding | RT | | 8 | 6.30E-04 | 1.30E-01 |
| INTERPRO | Gamma-aminobutyric acid A receptor | RT | | 6 | 1.40E-03 | 1.70E-01 |
| GOTERM_CC_DIRECT | synapse | RT | | 10 | 1.60E-03 | 1.60E-01 |
| GOTERM_BP_DIRECT | gamma-aminobutyric acid signaling pathway | RT | | 5 | 2.50E-03 | 1.00E+00 |
| UP_KEYWORDS | Ion channel | RT | | 16 | 4.70E-03 | 1.20E-01 |
| GOTERM_CC_DIRECT | cell junction | RT | | 12 | 5.00E-03 | 2.40E-01 |
| KEGG_PATHWAY | Retrograde endocannabinoid signaling | RT | | 10 | 5.40E-03 | 4.70E-01 |
| KEGG_PATHWAY | GABAergic synapse | RT | | 9 | 6.20E-03 | 4.70E-01 |
| KEGG_PATHWAY | Nicotine addiction | RT | | 6 | 8.80E-03 | 5.00E-01 |
| KEGG_PATHWAY | Morphine addiction | RT | | 8 | 3.00E-02 | 9.60E-01 |
| UP_KEYWORDS | Ion transport | RT | | 18 | 3.50E-02 | 5.00E-01 |
| KEGG_PATHWAY | Neuroactive ligand-receptor interaction | RT | | 14 | 1.40E-01 | 1.00E+00 |
| UP_KEYWORDS | Transport | RT | | 26 | 2.60E-01 | 1.00E+00 |
| UP_KEYWORDS | Cell membrane | RT | | 22 | 1.00E+00 | 1.00E+00 |
| **Annotation Cluster** | **Enrichment Score: 2.24** | **G** | | **Cou** | **P_Value** | **Benjamini** |
| INTERPRO | Histone core | RT | | 9 | 4.20E-04 | 1.30E-01 |
| GOTERM_CC_DIRECT | nucleosome | RT | | 9 | 7.10E-04 | 1.00E-01 |
| INTERPRO | Histone-fold | RT | | 10 | 8.30E-04 | 1.30E-01 |
| UP_KEYWORDS | Chromosome | RT | | 10 | 1.10E-03 | 4.30E-02 |
| UP_KEYWORDS | Nucleosome core | RT | | 8 | 1.40E-03 | 4.50E-02 |
| KEGG_PATHWAY | Systemic lupus erythematosus | RT | | 11 | 2.30E-02 | 8.70E-01 |
| INTERPRO | Histone H2A | RT | | 4 | 2.40E-02 | 1.00E+00 |
| SMART | H2A | RT | | 4 | 3.80E-02 | 1.00E+00 |
| KEGG_PATHWAY | Alcoholism | RT | | 12 | 4.00E-02 | 9.70E-01 |
| GOTERM_MF_DIRECT | DNA binding | RT | | 21 | 1.30E-01 | 1.00E+00 |
| **Annotation Cluster** | **Enrichment Score: 1.93** | **G** | | **Cou** | **P_Value** | **Benjamini** |
| UP_KEYWORDS | Extracellular matrix | RT | | 10 | 7.10E-04 | 3.70E-02 |
| INTERPRO | Collagen triple helix repeat | RT | | 8 | 6.80E-03 | 5.20E-01 |
| KEGG_PATHWAY | Protein digestion and absorption | RT | | 6 | 3.30E-01 | 1.00E+00 |

| Annotation Cluster | Enrichment Score: 1.84 | | | Cou | P_Value | Benjamini |
|---|---|---|---|---|---|---|
| INTERPRO | Histone core | RT | | 9 | 4.20E-04 | 1.30E-01 |
| INTERPRO | Histone H3 | RT | | 3 | 7.20E-02 | 1.00E+00 |
| SMART | H3 | RT | | 3 | 9.80E-02 | 1.00E+00 |
| **Annotation Cluster** | **Enrichment Score: 1.75** | | | **Cou** | **P_Value** | **Benjamini** |
| INTERPRO | Fibronectin, type III | RT | | 15 | 3.80E-03 | 3.40E-01 |
| SMART | FN3 | RT | | 12 | 1.70E-02 | 1.00E+00 |
| INTERPRO | Immunoglobulin-like fold | RT | | 26 | 8.80E-02 | 1.00E+00 |
| **Annotation Cluster** | **Enrichment Score: 1.63** | | | **Cou** | **P_Value** | **Benjamini** |
| UP_KEYWORDS | Extracellular matrix | RT | | 10 | 7.10E-04 | 3.70E-02 |
| GOTERM_CC_DIRECT | proteinaceous extracellular matrix | RT | | 11 | 2.10E-02 | 5.60E-01 |
| UP_KEYWORDS | Secreted | RT | | 12 | 8.90E-01 | 1.00E+00 |
| **Annotation Cluster** | **Enrichment Score: 1.6** | | | **Cou** | **P_Value** | **Benjamini** |
| INTERPRO | Type II keratin | RT | | 6 | 8.90E-04 | 1.30E-01 |
| GOTERM_CC_DIRECT | keratin filament | RT | | 6 | 5.70E-03 | 2.40E-01 |
| INTERPRO | Intermediate filament protein, conserved site | RT | | 6 | 3.80E-02 | 1.00E+00 |
| UP_KEYWORDS | Intermediate filament | RT | | 6 | 5.30E-02 | 5.90E-01 |
| GOTERM_MF_DIRECT | structural molecule activity | RT | | 9 | 6.40E-02 | 1.00E+00 |
| INTERPRO | Intermediate filament protein | RT | | 6 | 7.70E-02 | 1.00E+00 |
| SMART | SM01391 | RT | | 6 | 1.20E-01 | 1.00E+00 |
| **Annotation Cluster** | **Enrichment Score: 1.51** | | | **Cou** | **P_Value** | **Benjamini** |
| INTERPRO | Phospholipase C, phosphatidylinositol-specific , X domain | RT | | 5 | 2.50E-03 | 2.70E-01 |
| SMART | PLCXc | RT | | 5 | 3.70E-03 | 9.10E-01 |
| INTERPRO | Phospholipase C, phosphoinositol-specific, EF-hand-like | RT | | 4 | 3.70E-03 | 3.40E-01 |
| INTERPRO | PLC-like phosphodiesterase, TIM beta/alpha-barrel domain | RT | | 5 | 8.30E-03 | 5.20E-01 |
| INTERPRO | Phosphoinositide phospholipase C | RT | | 4 | 8.90E-03 | 5.20E-01 |
| INTERPRO | Phospholipase C, phosphatidylinositol-specific, Y domain | RT | | 4 | 8.90E-03 | 5.20E-01 |
| SMART | PLCYc | RT | | 4 | 1.50E-02 | 1.00E+00 |
| GOTERM_MF_DIRECT | phosphatidylinositol phospholipase C activity | RT | | 4 | 1.50E-02 | 1.00E+00 |
| UP_KEYWORDS | Lipid degradation | RT | | 5 | 4.00E-02 | 5.20E-01 |
| GOTERM_MF_DIRECT | signal transducer activity | RT | | 8 | 6.10E-02 | 1.00E+00 |
| GOTERM_BP_DIRECT | lipid catabolic process | RT | | 5 | 1.10E-01 | 1.00E+00 |
| UP_KEYWORDS | Lipid metabolism | RT | | 7 | 1.10E-01 | 1.00E+00 |
| INTERPRO | EF-hand-like domain | RT | | 11 | 3.10E-01 | 1.00E+00 |
| SMART | C2 | RT | | 5 | 6.70E-01 | 1.00E+00 |
| INTERPRO | C2 calcium-dependent membrane targeting | RT | | 5 | 7.20E-01 | 1.00E+00 |
| **Annotation Cluster** | **Enrichment Score: 1.4** | | | **Cou** | **P_Value** | **Benjamini** |
| KEGG_PATHWAY | ECM-receptor interaction | RT | | 9 | 1.10E-02 | 5.00E-01 |
| KEGG_PATHWAY | PI3K-Akt signaling pathway | RT | | 18 | 5.60E-02 | 1.00E+00 |
| KEGG_PATHWAY | Focal adhesion | RT | | 12 | 1.00E-01 | 1.00E+00 |
| **Annotation Cluster** | **Enrichment Score: 1.23** | | | **Cou** | **P_Value** | **Benjamini** |
| UP_KEYWORDS | DNA-binding | RT | | 27 | 1.90E-02 | 3.70E-01 |

| | | | | | | |
|---|---|---|---|---|---|---|
| UP_KEYWORDS | Transcription regulation | RT | | 14 | 3.00E-02 | 5.00E-01 |
| UP_KEYWORDS | Transcription | RT | | 15 | 4.70E-02 | 5.60E-01 |
| GOTERM_BP_DIRECT | transcription, DNA-templated | RT | | 11 | 4.70E-01 | 1.00E+00 |
| **Annotation Cluster** | **Enrichment Score: 1.16** | G | | **Cou** | **P_Value** | **Benjamini** |
| UP_KEYWORDS | Paired box | RT | | 3 | 3.30E-02 | 5.00E-01 |
| INTERPRO | Paired domain | RT | | 3 | 3.40E-02 | 1.00E+00 |
| SMART | PAX | RT | | 3 | 4.70E-02 | 1.00E+00 |
| UP_KEYWORDS | Developmental protein | RT | | 8 | 8.50E-02 | 8.90E-01 |
| INTERPRO | Winged helix-turn-helix DNA-binding domain | RT | | 9 | 3.40E-01 | 1.00E+00 |
| **Annotation Cluster** | **Enrichment Score: 1.07** | G | | **Cou** | **P_Value** | **Benjamini** |
| GOTERM_MF_DIRECT | cation binding | RT | | 3 | 1.40E-02 | 1.00E+00 |
| INTERPRO | Glycosyl hydrolase, family 13, catalytic domain | RT | | 3 | 2.60E-02 | 1.00E+00 |
| INTERPRO | Glycoside hydrolase, family 13 | RT | | 3 | 2.60E-02 | 1.00E+00 |
| SMART | Aamy | RT | | 3 | 3.60E-02 | 1.00E+00 |
| INTERPRO | Glycosyl hydrolase, family 13, all-beta | RT | | 3 | 1.10E-01 | 1.00E+00 |
| INTERPRO | Glycoside hydrolase, catalytic domain | RT | | 4 | 1.40E-01 | 1.00E+00 |
| INTERPRO | Glycoside hydrolase, superfamily | RT | | 4 | 2.70E-01 | 1.00E+00 |
| KEGG_PATHWAY | Starch and sucrose metabolism | RT | | 3 | 3.20E-01 | 1.00E+00 |
| GOTERM_BP_DIRECT | carbohydrate metabolic process | RT | | 5 | 5.30E-01 | 1.00E+00 |
| **Annotation Cluster** | **Enrichment Score: 1.06** | G | | **Cou** | **P_Value** | **Benjamini** |
| GOTERM_CC_DIRECT | postsynaptic membrane | RT | | 7 | 2.00E-02 | 5.60E-01 |
| KEGG_PATHWAY | Glutamatergic synapse | RT | | 9 | 3.40E-02 | 9.60E-01 |
| INTERPRO | Ionotropic glutamate receptor | RT | | 3 | 9.40E-02 | 1.00E+00 |
| INTERPRO | Glutamate receptor, L-glutamate/glycine-binding | RT | | 3 | 9.40E-02 | 1.00E+00 |
| INTERPRO | NMDA receptor | RT | | 3 | 9.40E-02 | 1.00E+00 |
| GOTERM_MF_DIRECT | extracellular-glutamate-gated ion channel activity | RT | | 3 | 9.90E-02 | 1.00E+00 |
| UP_KEYWORDS | Ligand-gated ion channel | RT | | 4 | 1.20E-01 | 1.00E+00 |
| SMART | PBPe | RT | | 3 | 1.30E-01 | 1.00E+00 |
| SMART | SM00918 | RT | | 3 | 1.30E-01 | 1.00E+00 |
| UP_KEYWORDS | Postsynaptic cell membrane | RT | | 4 | 1.30E-01 | 1.00E+00 |
| INTERPRO | Extracellular ligand-binding receptor | RT | | 5 | 1.60E-01 | 1.00E+00 |
| **Annotation Cluster** | **Enrichment Score: 1.04** | G | | **Cou** | **P_Value** | **Benjamini** |
| INTERPRO | Sterile alpha motif domain | RT | | 8 | 2.00E-02 | 1.00E+00 |
| SMART | SAM | RT | | 8 | 2.80E-02 | 1.00E+00 |
| INTERPRO | Tyrosine-protein kinase, catalytic domain | RT | | 7 | 4.40E-02 | 1.00E+00 |
| INTERPRO | Sterile alpha motif/pointed domain | RT | | 8 | 6.80E-02 | 1.00E+00 |
| INTERPRO | Tyrosine-protein kinase, active site | RT | | 7 | 7.20E-02 | 1.00E+00 |
| INTERPRO | Ephrin receptor ligand binding domain | RT | | 3 | 7.20E-02 | 1.00E+00 |
| INTERPRO | Tyrosine-protein kinase, receptor class V, conserved site | RT | | 3 | 7.20E-02 | 1.00E+00 |
| INTERPRO | Tyrosine-protein kinase, ephrin receptor | RT | | 3 | 7.20E-02 | 1.00E+00 |

| | | | | | | |
|---|---|---|---|---|---|---|
| GOTERM_BP_DIRECT | protein autophosphorylation | RT | | 9 | 8.10E-02 | 1.00E+00 |
| PIR_SUPERFAMILY | tyrosine-protein kinase, ephrin receptor type | RT | | 3 | 8.40E-02 | 1.00E+00 |
| SMART | TyrKc | RT | | 7 | 8.80E-02 | 1.00E+00 |
| UP_KEYWORDS | Tyrosine-protein kinase | RT | | 6 | 9.30E-02 | 9.10E-01 |
| SMART | EPH_lbd | RT | | 3 | 9.80E-02 | 1.00E+00 |
| INTERPRO | Tyrosine-protein kinase ephrin type A/B receptor-like | RT | | 3 | 1.70E-01 | 1.00E+00 |
| SMART | SM01411 | RT | | 3 | 2.30E-01 | 1.00E+00 |
| INTERPRO | Serine-threonine/tyrosine-protein kinase catalytic domain | RT | | 7 | 2.70E-01 | 1.00E+00 |
| INTERPRO | Galactose-binding domain-like | RT | | 3 | 7.20E-01 | 1.00E+00 |
| Annotation Cluster | Enrichment Score: 1.04 | G | | Cou | P_Value | Benjamini |
| INTERPRO | Tetratricopeptide repeat-containing domain | RT | | 9 | 3.50E-02 | 1.00E+00 |
| INTERPRO | Tetratricopeptide repeat | RT | | 8 | 1.10E-01 | 1.00E+00 |
| INTERPRO | Tetratricopeptide-like helical | RT | | 11 | 1.20E-01 | 1.00E+00 |
| SMART | TPR | RT | | 8 | 1.60E-01 | 1.00E+00 |
| Annotation Cluster | Enrichment Score: 0.93 | G | | Cou | P_Value | Benjamini |
| GOTERM_MF_DIRECT | carbonate dehydratase activity | RT | | 3 | 8.70E-02 | 1.00E+00 |
| INTERPRO | Carbonic anhydrase, alpha-class | RT | | 3 | 9.40E-02 | 1.00E+00 |
| KEGG_PATHWAY | Nitrogen metabolism | RT | | 3 | 1.10E-01 | 1.00E+00 |
| INTERPRO | Alpha carbonic anhydrase | RT | | 3 | 1.20E-01 | 1.00E+00 |
| SMART | SM01057 | RT | | 3 | 1.60E-01 | 1.00E+00 |
| GOTERM_BP_DIRECT | one-carbon metabolic process | RT | | 3 | 1.60E-01 | 1.00E+00 |
| Annotation Cluster | Enrichment Score: 0.81 | G | | Cou | P_Value | Benjamini |
| KEGG_PATHWAY | Morphine addiction | RT | | 8 | 3.00E-02 | 9.60E-01 |
| INTERPRO | 3'5'-cyclic nucleotide phosphodiesterase | RT | | 3 | 9.40E-02 | 1.00E+00 |
| INTERPRO | 3'5'-cyclic nucleotide phosphodiesterase, catalytic domain | RT | | 3 | 1.20E-01 | 1.00E+00 |
| INTERPRO | 3'5'-cyclic nucleotide phosphodiesterase, conserved site | RT | | 3 | 1.20E-01 | 1.00E+00 |
| INTERPRO | HD/PDEase domain | RT | | 3 | 1.40E-01 | 1.00E+00 |
| SMART | HDc | RT | | 3 | 1.90E-01 | 1.00E+00 |
| GOTERM_BP_DIRECT | signal transduction | RT | | 9 | 4.10E-01 | 1.00E+00 |
| KEGG_PATHWAY | Purine metabolism | RT | | 6 | 7.30E-01 | 1.00E+00 |
| Annotation Cluster | Enrichment Score: 0.78 | G | | Cou | P_Value | Benjamini |
| INTERPRO | GPCR, family 2, orphan receptor, GPR56 | RT | | 3 | 4.10E-03 | 3.40E-01 |
| INTERPRO | GPS domain | RT | | 3 | 2.80E-01 | 1.00E+00 |
| INTERPRO | GPCR, family 2-like | RT | | 4 | 2.90E-01 | 1.00E+00 |
| SMART | GPS | RT | | 3 | 3.10E-01 | 1.00E+00 |
| INTERPRO | GPCR, family 2, secretin-like | RT | | 3 | 4.30E-01 | 1.00E+00 |
| GOTERM_BP_DIRECT | cell surface receptor signaling pathway | RT | | 4 | 4.90E-01 | 1.00E+00 |
| Annotation Cluster | Enrichment Score: 0.69 | G | | Cou | P_Value | Benjamini |
| INTERPRO | RNA recognition motif domain | RT | | 11 | 1.20E-01 | 1.00E+00 |
| SMART | RRM | RT | | 11 | 2.00E-01 | 1.00E+00 |
| GOTERM_MF_DIRECT | nucleotide binding | RT | | 13 | 2.50E-01 | 1.00E+00 |
| INTERPRO | Nucleotide-binding, alpha-beta plait | RT | | 11 | 3.00E-01 | 1.00E+00 |
| Annotation Cluster | Enrichment Score: 0.68 | G | | Cou | P_Value | Benjamini |

| | | | | | | |
|---|---|---|---|---|---|---|
| UP_KEYWORDS | ANK repeat | RT | | 10 | 1.50E-01 | 1.00E+00 |
| INTERPRO | Ankyrin repeat | RT | | 10 | 1.90E-01 | 1.00E+00 |
| INTERPRO | Ankyrin repeat-containing domain | RT | | 10 | 2.30E-01 | 1.00E+00 |
| SMART | ANK | RT | | 10 | 3.20E-01 | 1.00E+00 |
| Annotation Cluster : | Enrichment Score: 0.66 | G | | Cou | P_Value | Benjamini |
| INTERPRO | Src homology-3 domain | RT | | 10 | 1.70E-01 | 1.00E+00 |
| UP_KEYWORDS | SH3 domain | RT | | 7 | 2.40E-01 | 1.00E+00 |
| SMART | SH3 | RT | | 10 | 2.60E-01 | 1.00E+00 |
| Annotation Cluster : | Enrichment Score: 0.65 | G | | Cou | P_Value | Benjamini |
| INTERPRO | Ubiquitin-conjugating enzyme/RWD-like | RT | | 5 | 1.40E-01 | 1.00E+00 |
| GOTERM_MF_DIRECT | ubiquitin conjugating enzyme activity | RT | | 3 | 1.50E-01 | 1.00E+00 |
| KEGG_PATHWAY | Protein processing in endoplasmic reticulum | RT | | 9 | 2.80E-01 | 1.00E+00 |
| INTERPRO | Ubiquitin-conjugating enzyme, E2 | RT | | 3 | 4.30E-01 | 1.00E+00 |
| Annotation Cluster : | Enrichment Score: 0.61 | G | | Cou | P_Value | Benjamini |
| INTERPRO | Protein kinase, ATP binding site | RT | | 21 | 1.40E-02 | 7.60E-01 |
| INTERPRO | Protein kinase, catalytic domain | RT | | 22 | 1.00E-01 | 1.00E+00 |
| INTERPRO | Protein kinase-like domain | RT | | 22 | 1.90E-01 | 1.00E+00 |
| INTERPRO | Serine/threonine-protein kinase, active site | RT | | 14 | 2.00E-01 | 1.00E+00 |
| UP_KEYWORDS | Kinase | RT | | 15 | 2.30E-01 | 1.00E+00 |
| UP_KEYWORDS | Transferase | RT | | 28 | 2.90E-01 | 1.00E+00 |
| UP_KEYWORDS | ATP-binding | RT | | 26 | 4.40E-01 | 1.00E+00 |
| GOTERM_MF_DIRECT | protein serine/threonine kinase activity | RT | | 8 | 4.70E-01 | 1.00E+00 |
| SMART | S_TKc | RT | | 15 | 4.70E-01 | 1.00E+00 |
| GOTERM_MF_DIRECT | ATP binding | RT | | 43 | 4.70E-01 | 1.00E+00 |
| UP_KEYWORDS | Serine/threonine-protein kinase | RT | | 7 | 5.30E-01 | 1.00E+00 |
| UP_KEYWORDS | Nucleotide-binding | RT | | 31 | 5.40E-01 | 1.00E+00 |
| Annotation Cluster : | Enrichment Score: 0.6 | G | | Cou | P_Value | Benjamini |
| INTERPRO | Plexin | RT | | 3 | 2.00E-01 | 1.00E+00 |
| INTERPRO | Plexin-like fold | RT | | 4 | 2.10E-01 | 1.00E+00 |
| SMART | PSI | RT | | 4 | 2.50E-01 | 1.00E+00 |
| INTERPRO | Semaphorin/CD100 antigen | RT | | 3 | 2.80E-01 | 1.00E+00 |
| SMART | Sema | RT | | 3 | 3.40E-01 | 1.00E+00 |
| Annotation Cluster : | Enrichment Score: 0.6 | G | | Cou | P_Value | Benjamini |
| KEGG_PATHWAY | Sphingolipid signaling pathway | RT | | 9 | 4.20E-02 | 9.70E-01 |
| KEGG_PATHWAY | Dopaminergic synapse | RT | | 6 | 4.00E-01 | 1.00E+00 |
| KEGG_PATHWAY | AMPK signaling pathway | RT | | 6 | 4.40E-01 | 1.00E+00 |
| KEGG_PATHWAY | Adrenergic signaling in cardiomyocytes | RT | | 6 | 5.50E-01 | 1.00E+00 |
| Annotation Cluster : | Enrichment Score: 0.6 | G | | Cou | P_Value | Benjamini |
| INTERPRO | Immunoglobulin-like fold | RT | | 26 | 8.80E-02 | 1.00E+00 |
| INTERPRO | Immunoglobulin subtype 2 | RT | | 11 | 1.20E-01 | 1.00E+00 |
| INTERPRO | Immunoglobulin I-set | RT | | 8 | 1.20E-01 | 1.00E+00 |
| INTERPRO | Immunoglobulin subtype | RT | | 14 | 2.10E-01 | 1.00E+00 |
| SMART | IGc2 | RT | | 11 | 2.50E-01 | 1.00E+00 |
| INTERPRO | Immunoglobulin V-set | RT | | 6 | 4.10E-01 | 1.00E+00 |
| SMART | IG | RT | | 14 | 4.30E-01 | 1.00E+00 |
| INTERPRO | Immunoglobulin-like domain | RT | | 15 | 4.30E-01 | 1.00E+00 |

| | | | | | | |
|---|---|---|---|---|---|---|
| UP_KEYWORDS | Immunoglobulin domain | RT | | 5 | 4.60E-01 | 1.00E+00 |
| SMART | IGv | RT | | 4 | 4.90E-01 | 1.00E+00 |
| Annotation Cluster : | Enrichment Score: 0.56 | G | | Cou | P_Value | Benjamini |
| KEGG_PATHWAY | Glutamatergic synapse | RT | | 9 | 3.40E-02 | 9.60E-01 |
| KEGG_PATHWAY | Long-term depression | RT | | 5 | 1.40E-01 | 1.00E+00 |
| KEGG_PATHWAY | Long-term potentiation | RT | | 5 | 1.60E-01 | 1.00E+00 |
| KEGG_PATHWAY | Gap junction | RT | | 5 | 3.50E-01 | 1.00E+00 |
| KEGG_PATHWAY | Serotonergic synapse | RT | | 6 | 3.60E-01 | 1.00E+00 |
| KEGG_PATHWAY | Calcium signaling pathway | RT | | 7 | 6.10E-01 | 1.00E+00 |
| KEGG_PATHWAY | Inflammatory mediator regulation of TRP channels | RT | | 4 | 7.00E-01 | 1.00E+00 |
| KEGG_PATHWAY | Estrogen signaling pathway | RT | | 3 | 8.50E-01 | 1.00E+00 |
| Annotation Cluster : | Enrichment Score: 0.56 | G | | Cou | P_Value | Benjamini |
| KEGG_PATHWAY | Long-term depression | RT | | 5 | 1.40E-01 | 1.00E+00 |
| KEGG_PATHWAY | Chagas disease (American trypanosomiasis) | RT | | 7 | 1.80E-01 | 1.00E+00 |
| KEGG_PATHWAY | Chemokine signaling pathway | RT | | 5 | 8.40E-01 | 1.00E+00 |
| Annotation Cluster : | Enrichment Score: 0.56 | G | | Cou | P_Value | Benjamini |
| INTERPRO | Dual specificity phosphatase, catalytic domain | RT | | 4 | 1.60E-01 | 1.00E+00 |
| GOTERM_MF_DIRECT | protein tyrosine/serine/threonine phosphatase activity | RT | | 3 | 2.50E-01 | 1.00E+00 |
| INTERPRO | Dual specificity phosphatase | RT | | 3 | 2.50E-01 | 1.00E+00 |
| INTERPRO | Protein-tyrosine/Dual specificity phosphatase | RT | | 5 | 2.90E-01 | 1.00E+00 |
| INTERPRO | Dual specificity phosphatase, subgroup, catalytic domain | RT | | 3 | 3.50E-01 | 1.00E+00 |
| SMART | DSPc | RT | | 3 | 4.40E-01 | 1.00E+00 |
| Annotation Cluster : | Enrichment Score: 0.53 | G | | Cou | P_Value | Benjamini |
| KEGG_PATHWAY | Oxidative phosphorylation | RT | | 9 | 1.20E-01 | 1.00E+00 |
| KEGG_PATHWAY | Huntington's disease | RT | | 10 | 2.90E-01 | 1.00E+00 |
| KEGG_PATHWAY | Alzheimer's disease | RT | | 9 | 3.00E-01 | 1.00E+00 |
| KEGG_PATHWAY | Parkinson's disease | RT | | 8 | 3.10E-01 | 1.00E+00 |
| KEGG_PATHWAY | Non-alcoholic fatty liver disease (NAFLD) | RT | | 6 | 7.00E-01 | 1.00E+00 |
| Annotation Cluster : | Enrichment Score: 0.47 | G | | Cou | P_Value | Benjamini |
| INTERPRO | Leucine-rich repeat-containing N-terminal | RT | | 4 | 1.90E-01 | 1.00E+00 |
| INTERPRO | Leucine-rich repeat | RT | | 11 | 2.30E-01 | 1.00E+00 |
| SMART | LRRNT | RT | | 4 | 2.70E-01 | 1.00E+00 |
| INTERPRO | Leucine-rich repeat, typical subtype | RT | | 7 | 4.50E-01 | 1.00E+00 |
| UP_KEYWORDS | Leucine-rich repeat | RT | | 5 | 5.00E-01 | 1.00E+00 |
| SMART | LRR_TYP | RT | | 7 | 6.30E-01 | 1.00E+00 |
| Annotation Cluster : | Enrichment Score: 0.45 | G | | Cou | P_Value | Benjamini |
| INTERPRO | HECT | RT | | 3 | 2.50E-01 | 1.00E+00 |
| SMART | HECTc | RT | | 3 | 3.30E-01 | 1.00E+00 |
| UP_KEYWORDS | Ubl conjugation pathway | RT | | 5 | 4.20E-01 | 1.00E+00 |
| GOTERM_MF_DIRECT | ligase activity | RT | | 3 | 4.50E-01 | 1.00E+00 |
| Annotation Cluster : | Enrichment Score: 0.37 | G | | Cou | P_Value | Benjamini |

| | | | | | | |
|---|---|---|---|---|---|---|
| INTERPRO | Protein-tyrosine/Dual specificity phosphatase | RT | | 5 | 2.90E-01 | 1.00E+00 |
| GOTERM_MF_DIRECT | protein tyrosine phosphatase activity | RT | | 5 | 3.60E-01 | 1.00E+00 |
| INTERPRO | Protein-tyrosine phosphatase, active site | RT | | 3 | 7.20E-01 | 1.00E+00 |
| Annotation Cluster | Enrichment Score: 0.36 | G | | Cou | P_Value | Benjamini |
| INTERPRO | Protease-associated domain, PA | RT | | 3 | 6.10E-02 | 1.00E+00 |
| INTERPRO | Zinc finger, RING-type | RT | | 9 | 5.70E-01 | 1.00E+00 |
| INTERPRO | Zinc finger, RING/FYVE/PHD-type | RT | | 13 | 6.60E-01 | 1.00E+00 |
| SMART | RING | RT | | 8 | 7.00E-01 | 1.00E+00 |
| INTERPRO | Zinc finger, RING-type, conserved site | RT | | 3 | 9.40E-01 | 1.00E+00 |
| Annotation Cluster | Enrichment Score: 0.35 | G | | Cou | P_Value | Benjamini |
| INTERPRO | WD40/YVTN repeat-like-containing domain | RT | | 15 | 1.80E-01 | 1.00E+00 |
| INTERPRO | WD40 repeat | RT | | 10 | 4.60E-01 | 1.00E+00 |
| INTERPRO | WD40-repeat-containing domain | RT | | 11 | 4.80E-01 | 1.00E+00 |
| INTERPRO | G-protein beta WD-40 repeat | RT | | 4 | 5.10E-01 | 1.00E+00 |
| INTERPRO | WD40 repeat, conserved site | RT | | 6 | 5.70E-01 | 1.00E+00 |
| SMART | WD40 | RT | | 10 | 6.70E-01 | 1.00E+00 |
| Annotation Cluster | Enrichment Score: 0.31 | G | | Cou | P_Value | Benjamini |
| INTERPRO | AAA+ ATPase domain | RT | | 6 | 4.00E-01 | 1.00E+00 |
| GOTERM_MF_DIRECT | ATPase activity | RT | | 4 | 5.20E-01 | 1.00E+00 |
| SMART | AAA | RT | | 6 | 5.60E-01 | 1.00E+00 |
| Annotation Cluster | Enrichment Score: 0.29 | G | | Cou | P_Value | Benjamini |
| INTERPRO | Thioredoxin domain | RT | | 3 | 3.80E-01 | 1.00E+00 |
| GOTERM_BP_DIRECT | cell redox homeostasis | RT | | 4 | 3.80E-01 | 1.00E+00 |
| INTERPRO | Thioredoxin-like fold | RT | | 3 | 9.20E-01 | 1.00E+00 |
| Annotation Cluster | Enrichment Score: 0.29 | G | | Cou | P_Value | Benjamini |
| INTERPRO | Zinc finger, nuclear hormone receptor-type | RT | | 3 | 4.40E-01 | 1.00E+00 |
| INTERPRO | Nuclear hormone receptor, ligand-binding, core | RT | | 3 | 4.80E-01 | 1.00E+00 |
| SMART | ZnF_C4 | RT | | 3 | 5.40E-01 | 1.00E+00 |
| INTERPRO | Zinc finger, NHR/GATA-type | RT | | 3 | 5.50E-01 | 1.00E+00 |
| SMART | HOLI | RT | | 3 | 5.80E-01 | 1.00E+00 |
| Annotation Cluster | Enrichment Score: 0.28 | G | | Cou | P_Value | Benjamini |
| INTERPRO | Guanine-nucleotide dissociation stimulator, CDC24, conserved site | RT | | 3 | 1.60E-01 | 1.00E+00 |
| GOTERM_MF_DIRECT | Rho guanyl-nucleotide exchange factor activity | RT | | 3 | 6.40E-01 | 1.00E+00 |
| INTERPRO | Dbl homology (DH) domain | RT | | 3 | 6.80E-01 | 1.00E+00 |
| SMART | RhoGEF | RT | | 3 | 7.40E-01 | 1.00E+00 |
| GOTERM_BP_DIRECT | regulation of Rho protein signal transduction | RT | | 3 | 7.40E-01 | 1.00E+00 |
| Annotation Cluster | Enrichment Score: 0.27 | G | | Cou | P_Value | Benjamini |
| KEGG_PATHWAY | Chronic myeloid leukemia | RT | | 5 | 2.40E-01 | 1.00E+00 |
| KEGG_PATHWAY | Hepatitis B | RT | | 5 | 7.70E-01 | 1.00E+00 |
| KEGG_PATHWAY | Pancreatic cancer | RT | | 3 | 8.00E-01 | 1.00E+00 |
| Annotation Cluster | Enrichment Score: 0.27 | G | | Cou | P_Value | Benjamini |
| INTERPRO | EF-hand-like domain | RT | | 11 | 3.10E-01 | 1.00E+00 |
| INTERPRO | EF-Hand 1, calcium-binding site | RT | | 7 | 4.20E-01 | 1.00E+00 |
| GOTERM_MF_DIRECT | calcium ion binding | RT | | 18 | 5.60E-01 | 1.00E+00 |
| INTERPRO | EF-hand domain | RT | | 7 | 6.40E-01 | 1.00E+00 |
| SMART | EFh | RT | | 4 | 9.10E-01 | 1.00E+00 |
| Annotation Cluster | Enrichment Score: 0.27 | G | | Cou | P_Value | Benjamini |

| | | | | | |
|---|---|---|---|---|---|
| INTERPRO | Pleckstrin homology-like domain | RT | 15 | 3.90E-01 | 1.00E+00 |
| INTERPRO | Pleckstrin homology domain | RT | 9 | 5.60E-01 | 1.00E+00 |
| SMART | PH | RT | 9 | 7.10E-01 | 1.00E+00 |
| Annotation Cluster | Enrichment Score: 0.23 | | Cou | P_Value | Benjamini |
| GOTERM_BP_DIRECT | homophilic cell adhesion via plasma membrane adhesion molecules | RT | 4 | 4.90E-01 | 1.00E+00 |
| INTERPRO | Cadherin | RT | 3 | 5.80E-01 | 1.00E+00 |
| INTERPRO | Cadherin-like | RT | 3 | 6.10E-01 | 1.00E+00 |
| SMART | CA | RT | 3 | 6.70E-01 | 1.00E+00 |
| Annotation Cluster | Enrichment Score: 0.21 | | Cou | P_Value | Benjamini |
| KEGG_PATHWAY | Citrate cycle (TCA cycle) | RT | 3 | 4.00E-01 | 1.00E+00 |
| KEGG_PATHWAY | Carbon metabolism | RT | 5 | 6.20E-01 | 1.00E+00 |
| KEGG_PATHWAY | Biosynthesis of antibiotics | RT | 6 | 9.10E-01 | 1.00E+00 |
| Annotation Cluster | Enrichment Score: 0.2 | | Cou | P_Value | Benjamini |
| INTERPRO | Homeodomain-like | RT | 12 | 2.40E-01 | 1.00E+00 |
| GOTERM_MF_DIRECT | sequence-specific DNA binding | RT | 9 | 6.50E-01 | 1.00E+00 |
| INTERPRO | Homeobox, conserved site | RT | 5 | 7.30E-01 | 1.00E+00 |
| UP_KEYWORDS | Homeobox | RT | 6 | 7.70E-01 | 1.00E+00 |
| INTERPRO | Homeodomain | RT | 6 | 8.00E-01 | 1.00E+00 |
| SMART | HOX | RT | 5 | 9.50E-01 | 1.00E+00 |
| Annotation Cluster | Enrichment Score: 0.18 | | Cou | P_Value | Benjamini |
| KEGG_PATHWAY | Wnt signaling pathway | RT | 7 | 3.20E-01 | 1.00E+00 |
| KEGG_PATHWAY | Oxytocin signaling pathway | RT | 4 | 9.00E-01 | 1.00E+00 |
| KEGG_PATHWAY | cGMP-PKG signaling pathway | RT | 3 | 9.80E-01 | 1.00E+00 |
| Annotation Cluster | Enrichment Score: 0.16 | | Cou | P_Value | Benjamini |
| INTERPRO | Epidermal growth factor-like domain | RT | 8 | 4.80E-01 | 1.00E+00 |
| INTERPRO | EGF-type aspartate/asparagine hydroxylation site | RT | 4 | 4.90E-01 | 1.00E+00 |
| INTERPRO | EGF-like, conserved site | RT | 6 | 6.50E-01 | 1.00E+00 |
| SMART | EGF | RT | 6 | 6.90E-01 | 1.00E+00 |
| INTERPRO | EGF-like calcium-binding, conserved site | RT | 3 | 7.10E-01 | 1.00E+00 |
| INTERPRO | EGF-like calcium-binding | RT | 3 | 8.50E-01 | 1.00E+00 |
| UP_KEYWORDS | EGF-like domain | RT | 3 | 9.10E-01 | 1.00E+00 |
| SMART | EGF_CA | RT | 3 | 9.10E-01 | 1.00E+00 |
| Annotation Cluster | Enrichment Score: 0.13 | | Cou | P_Value | Benjamini |
| INTERPRO | BTB/POZ fold | RT | 7 | 5.70E-01 | 1.00E+00 |
| INTERPRO | BTB/POZ-like | RT | 5 | 8.20E-01 | 1.00E+00 |
| SMART | BTB | RT | 5 | 9.00E-01 | 1.00E+00 |
| Annotation Cluster | Enrichment Score: 0.12 | | Cou | P_Value | Benjamini |
| INTERPRO | Zinc finger, PHD-finger | RT | 3 | 6.70E-01 | 1.00E+00 |
| INTERPRO | Zinc finger, PHD-type | RT | 3 | 7.40E-01 | 1.00E+00 |
| INTERPRO | Zinc finger, FYVE/PHD-type | RT | 4 | 8.00E-01 | 1.00E+00 |
| SMART | PHD | RT | 3 | 8.20E-01 | 1.00E+00 |
| Annotation Cluster | Enrichment Score: 0.11 | | Cou | P_Value | Benjamini |
| GOTERM_MF_DIRECT | nucleic acid binding | RT | 15 | 6.80E-01 | 1.00E+00 |
| INTERPRO | Zinc finger C2H2-type/integrase DNA-binding domain | RT | 11 | 7.50E-01 | 1.00E+00 |
| GOTERM_MF_DIRECT | metal ion binding | RT | 26 | 7.80E-01 | 1.00E+00 |
| INTERPRO | Zinc finger, C2H2-like | RT | 12 | 7.90E-01 | 1.00E+00 |
| INTERPRO | Zinc finger, C2H2 | RT | 13 | 7.90E-01 | 1.00E+00 |
| SMART | ZnF_C2H2 | RT | 12 | 9.30E-01 | 1.00E+00 |

| Annotation Cluster ! | Enrichment Score: 0.06 | G | | Cou | P_Value | Benjamini |
|---|---|---|---|---|---|---|
| UP_KEYWORDS | Zinc | RT | | 22 | 8.30E-01 | 1.00E+00 |
| UP_KEYWORDS | Zinc-finger | RT | | 14 | 8.90E-01 | 1.00E+00 |
| UP_KEYWORDS | Metal-binding | RT | | 33 | 9.00E-01 | 1.00E+00 |
| Annotation Cluster ! | Enrichment Score: 0.04 | G | | Cou | P_Value | Benjamini |
| INTERPRO | Small GTP-binding protein domain | RT | | 4 | 8.80E-01 | 1.00E+00 |
| UP_KEYWORDS | GTP-binding | RT | | 4 | 8.80E-01 | 1.00E+00 |
| GOTERM_BP_DIRECT | small GTPase mediated signal transduction | RT | | 5 | 9.10E-01 | 1.00E+00 |
| INTERPRO | Small GTPase superfamily | RT | | 3 | 9.10E-01 | 1.00E+00 |
| GOTERM_MF_DIRECT | GTP binding | RT | | 5 | 9.90E-01 | 1.00E+00 |
| Annotation Cluster ! | Enrichment Score: 0.03 | G | | Cou | P_Value | Benjamini |
| KEGG_PATHWAY | Ribosome | RT | | 6 | 8.70E-01 | 1.00E+00 |
| GOTERM_MF_DIRECT | structural constituent of ribosome | RT | | 4 | 9.70E-01 | 1.00E+00 |
| GOTERM_BP_DIRECT | translation | RT | | 4 | 9.80E-01 | 1.00E+00 |
| Annotation Cluster ! | Enrichment Score: 0.01 | G | | Cou | P_Value | Benjamini |
| UP_KEYWORDS | Transmembrane helix | RT | | # | 9.80E-01 | 1.00E+00 |
| UP_KEYWORDS | Transmembrane | RT | | # | 9.80E-01 | 1.00E+00 |
| UP_KEYWORDS | Membrane | RT | | # | 9.80E-01 | 1.00E+00 |
| GOTERM_CC_DIRECT | integral component of membrane | RT | | # | 1.00E+00 | 1.00E+00 |
| Annotation Cluster ! | Enrichment Score: 0 | G | | Cou | P_Value | Benjamini |
| SMART | SM01381 | RT | | 4 | 9.30E-01 | 1.00E+00 |
| UP_KEYWORDS | Receptor | RT | | 27 | 9.90E-01 | 1.00E+00 |
| UP_KEYWORDS | Transducer | RT | | 15 | 1.00E+00 | 1.00E+00 |
| GOTERM_MF_DIRECT | G-protein coupled receptor activity | RT | | 12 | 1.00E+00 | 1.00E+00 |
| INTERPRO | G protein-coupled receptor, rhodopsin-like | RT | | 11 | 1.00E+00 | 1.00E+00 |
| UP_KEYWORDS | G-protein coupled receptor | RT | | 11 | 1.00E+00 | 1.00E+00 |
| KEGG_PATHWAY | Olfactory transduction | RT | | 9 | 1.00E+00 | 1.00E+00 |
| INTERPRO | GPCR, rhodopsin-like, 7TM | RT | | 11 | 1.00E+00 | 1.00E+00 |
| UP_KEYWORDS | Olfaction | RT | | 4 | 1.00E+00 | 1.00E+00 |
| INTERPRO | Olfactory receptor | RT | | 4 | 1.00E+00 | 1.00E+00 |
| UP_KEYWORDS | Sensory transduction | RT | | 4 | 1.00E+00 | 1.00E+00 |
| GOTERM_MF_DIRECT | olfactory receptor activity | RT | | 4 | 1.00E+00 | 1.00E+00 |

**Table S11. Significant SNPs for DFW**

| Chr | Position | Allele0 | Allele1 | Freq | *P* | Bon |
|---|---|---|---|---|---|---|
| 11 | 64948615 | G | C | 0.928331 | 6.12E-30 | 2.40E-24 |
| 11 | 64950500 | T | C | 0.928331 | 6.12E-30 | 2.40E-24 |
| 11 | 64950776 | G | A | 0.928331 | 6.12E-30 | 2.40E-24 |
| 11 | 64950793 | T | A | 0.928331 | 6.12E-30 | 2.40E-24 |
| 11 | 64950794 | T | C | 0.928331 | 6.12E-30 | 2.40E-24 |
| 11 | 64951137 | A | G | 0.928331 | 6.12E-30 | 2.40E-24 |
| 11 | 64951174 | G | C | 0.928331 | 6.12E-30 | 2.40E-24 |
| 11 | 64951364 | T | C | 0.928331 | 6.12E-30 | 2.40E-24 |
| 11 | 64951973 | A | T | 0.928331 | 6.12E-30 | 2.40E-24 |
| 11 | 64952273 | G | A | 0.928331 | 6.12E-30 | 2.40E-24 |
| 11 | 64944841 | A | T | 0.929174 | 9.26E-30 | 3.63E-24 |
| 11 | 64366972 | A | G | 0.925801 | 6.73E-28 | 2.64E-22 |
| 11 | 64521430 | T | A | 0.930017 | 2.57E-27 | 1.01E-21 |
| 11 | 64270037 | G | C | 0.924958 | 2.66E-27 | 1.04E-21 |
| 11 | 64349603 | C | T | 0.924958 | 2.66E-27 | 1.04E-21 |
| 11 | 64350722 | T | C | 0.924958 | 2.66E-27 | 1.04E-21 |
| 11 | 64351227 | T | G | 0.924958 | 2.66E-27 | 1.04E-21 |
| 11 | 64361938 | T | C | 0.924958 | 2.66E-27 | 1.04E-21 |
| 11 | 64365353 | A | G | 0.924958 | 2.66E-27 | 1.04E-21 |
| 11 | 64461388 | A | G | 0.924958 | 2.66E-27 | 1.04E-21 |
| 11 | 64468285 | C | T | 0.924958 | 2.66E-27 | 1.04E-21 |
| 11 | 64269214 | C | A | 0.925801 | 3.16E-27 | 1.24E-21 |
| 11 | 64270630 | C | T | 0.925801 | 3.87E-27 | 1.52E-21 |
| 11 | 64270744 | T | C | 0.925801 | 3.87E-27 | 1.52E-21 |
| 11 | 64234283 | C | A | 0.925801 | 9.73E-27 | 3.81E-21 |
| 11 | 64224299 | T | C | 0.924958 | 1.06E-25 | 4.14E-20 |
| 11 | 64483464 | A | G | 0.926644 | 4.27E-25 | 1.68E-19 |
| 11 | 64220895 | T | C | 0.923272 | 7.05E-25 | 2.76E-19 |
| 11 | 64839884 | G | A | 0.91484 | 1.48E-24 | 5.79E-19 |
| 11 | 64839959 | C | T | 0.91484 | 1.48E-24 | 5.79E-19 |
| 11 | 64840001 | C | G | 0.91484 | 1.48E-24 | 5.79E-19 |
| 11 | 64840014 | A | C | 0.91484 | 1.48E-24 | 5.79E-19 |
| 11 | 64840069 | A | G | 0.91484 | 1.48E-24 | 5.79E-19 |
| 11 | 64840116 | C | G | 0.91484 | 1.48E-24 | 5.79E-19 |
| 11 | 64840353 | A | G | 0.91484 | 1.48E-24 | 5.79E-19 |
| 11 | 64840572 | A | C | 0.91484 | 1.48E-24 | 5.79E-19 |
| 11 | 64840575 | T | C | 0.91484 | 1.48E-24 | 5.79E-19 |
| 11 | 65993082 | A | G | 0.93339 | 4.02E-24 | 1.58E-18 |
| 11 | 65993312 | A | G | 0.93339 | 4.02E-24 | 1.58E-18 |
| 11 | 65993521 | T | C | 0.93339 | 4.02E-24 | 1.58E-18 |
| 11 | 65993633 | T | G | 0.93339 | 4.02E-24 | 1.58E-18 |
| 11 | 64840642 | G | A | 0.915683 | 5.10E-24 | 2.00E-18 |
| 11 | 64840760 | T | A | 0.915683 | 5.10E-24 | 2.00E-18 |
| 11 | 64841000 | C | T | 0.915683 | 5.10E-24 | 2.00E-18 |
| 11 | 64242653 | T | C | 0.928331 | 5.96E-24 | 2.34E-18 |
| 11 | 64311025 | T | C | 0.926644 | 1.06E-23 | 4.16E-18 |
| 11 | 64242993 | G | A | 0.929174 | 2.43E-23 | 9.52E-18 |
| 11 | 64831085 | T | C | 0.910624 | 3.07E-23 | 1.20E-17 |
| 11 | 64834419 | T | C | 0.910624 | 3.07E-23 | 1.20E-17 |
| 11 | 64834579 | G | A | 0.910624 | 3.07E-23 | 1.20E-17 |
| 11 | 64836885 | G | T | 0.910624 | 3.07E-23 | 1.20E-17 |
| 11 | 64837646 | A | G | 0.910624 | 3.07E-23 | 1.20E-17 |
| 11 | 65990344 | T | C | 0.931703 | 4.59E-23 | 1.80E-17 |
| 11 | 64838650 | C | A | 0.911467 | 5.38E-23 | 2.11E-17 |
| 11 | 64839268 | T | C | 0.911467 | 5.38E-23 | 2.11E-17 |
| 11 | 65451349 | T | G | 0.932546 | 6.46E-23 | 2.53E-17 |
| 11 | 65451381 | C | A | 0.932546 | 6.46E-23 | 2.53E-17 |
| 11 | 65451780 | T | C | 0.932546 | 6.46E-23 | 2.53E-17 |
| 11 | 65451868 | A | G | 0.932546 | 6.46E-23 | 2.53E-17 |
| 11 | 65452088 | G | A | 0.932546 | 6.46E-23 | 2.53E-17 |
| 11 | 65452090 | T | C | 0.932546 | 6.46E-23 | 2.53E-17 |
| 11 | 65452186 | C | T | 0.932546 | 6.46E-23 | 2.53E-17 |
| 11 | 65452273 | T | C | 0.932546 | 6.46E-23 | 2.53E-17 |
| 11 | 65452436 | T | G | 0.932546 | 6.46E-23 | 2.53E-17 |
| 11 | 65452488 | C | T | 0.932546 | 6.46E-23 | 2.53E-17 |

| | | | | | | |
|---|---|---|---|---|---|---|
| 11 | 65452978 | T | C | 0.932546 | 6.46E-23 | 2.53E-17 |
| 11 | 65453191 | C | G | 0.932546 | 6.46E-23 | 2.53E-17 |
| 11 | 65453312 | A | G | 0.932546 | 6.46E-23 | 2.53E-17 |
| 11 | 65453327 | T | A | 0.932546 | 6.46E-23 | 2.53E-17 |
| 11 | 65453351 | T | C | 0.932546 | 6.46E-23 | 2.53E-17 |
| 11 | 65453374 | G | A | 0.932546 | 6.46E-23 | 2.53E-17 |
| 11 | 65453448 | G | A | 0.932546 | 6.46E-23 | 2.53E-17 |
| 11 | 65453464 | T | C | 0.932546 | 6.46E-23 | 2.53E-17 |
| 11 | 65453608 | T | C | 0.932546 | 6.46E-23 | 2.53E-17 |
| 11 | 65453650 | T | C | 0.932546 | 6.46E-23 | 2.53E-17 |
| 11 | 65454294 | T | G | 0.932546 | 6.46E-23 | 2.53E-17 |
| 11 | 65454298 | C | T | 0.932546 | 6.46E-23 | 2.53E-17 |
| 11 | 65454336 | A | G | 0.932546 | 6.46E-23 | 2.53E-17 |
| 11 | 65454560 | A | G | 0.932546 | 6.46E-23 | 2.53E-17 |
| 11 | 65454780 | T | C | 0.932546 | 6.46E-23 | 2.53E-17 |
| 11 | 65454871 | G | T | 0.932546 | 6.46E-23 | 2.53E-17 |
| 11 | 65455133 | G | T | 0.932546 | 6.46E-23 | 2.53E-17 |
| 11 | 65455213 | A | C | 0.932546 | 6.46E-23 | 2.53E-17 |
| 11 | 65455236 | C | T | 0.932546 | 6.46E-23 | 2.53E-17 |
| 11 | 65455238 | C | T | 0.932546 | 6.46E-23 | 2.53E-17 |
| 11 | 65455275 | G | A | 0.932546 | 6.46E-23 | 2.53E-17 |
| 11 | 65455359 | C | T | 0.932546 | 6.46E-23 | 2.53E-17 |
| 11 | 65455360 | A | G | 0.932546 | 6.46E-23 | 2.53E-17 |
| 11 | 65455388 | A | G | 0.932546 | 6.46E-23 | 2.53E-17 |
| 11 | 65455483 | C | T | 0.932546 | 6.46E-23 | 2.53E-17 |
| 11 | 65455598 | T | C | 0.932546 | 6.46E-23 | 2.53E-17 |
| 11 | 65455605 | G | T | 0.932546 | 6.46E-23 | 2.53E-17 |
| 11 | 65455634 | A | G | 0.932546 | 6.46E-23 | 2.53E-17 |
| 11 | 65455696 | A | T | 0.932546 | 6.46E-23 | 2.53E-17 |
| 11 | 65455796 | T | A | 0.932546 | 6.46E-23 | 2.53E-17 |
| 11 | 65455808 | T | C | 0.932546 | 6.46E-23 | 2.53E-17 |
| 11 | 65455890 | C | G | 0.932546 | 6.46E-23 | 2.53E-17 |
| 11 | 65456009 | G | C | 0.932546 | 6.46E-23 | 2.53E-17 |
| 11 | 65456365 | T | C | 0.932546 | 6.46E-23 | 2.53E-17 |
| 11 | 65456372 | T | C | 0.932546 | 6.46E-23 | 2.53E-17 |
| 11 | 65456426 | A | G | 0.932546 | 6.46E-23 | 2.53E-17 |
| 11 | 65456645 | T | A | 0.932546 | 6.46E-23 | 2.53E-17 |
| 11 | 65456708 | A | G | 0.932546 | 6.46E-23 | 2.53E-17 |
| 11 | 65456718 | A | T | 0.932546 | 6.46E-23 | 2.53E-17 |
| 11 | 65456775 | G | A | 0.932546 | 6.46E-23 | 2.53E-17 |
| 11 | 65456811 | G | A | 0.932546 | 6.46E-23 | 2.53E-17 |
| 11 | 65457384 | T | G | 0.932546 | 6.46E-23 | 2.53E-17 |
| 11 | 65457726 | T | C | 0.932546 | 6.46E-23 | 2.53E-17 |
| 11 | 65458608 | G | A | 0.932546 | 6.46E-23 | 2.53E-17 |
| 11 | 65458611 | G | A | 0.932546 | 6.46E-23 | 2.53E-17 |
| 11 | 65458618 | G | A | 0.932546 | 6.46E-23 | 2.53E-17 |
| 11 | 65458651 | A | G | 0.932546 | 6.46E-23 | 2.53E-17 |
| 11 | 65458656 | C | T | 0.932546 | 6.46E-23 | 2.53E-17 |
| 11 | 65458789 | G | A | 0.932546 | 6.46E-23 | 2.53E-17 |
| 11 | 65458805 | T | G | 0.932546 | 6.46E-23 | 2.53E-17 |
| 11 | 65458940 | T | C | 0.932546 | 6.46E-23 | 2.53E-17 |
| 11 | 65459212 | C | A | 0.932546 | 6.46E-23 | 2.53E-17 |
| 11 | 65460225 | C | A | 0.932546 | 6.46E-23 | 2.53E-17 |
| 11 | 65460287 | C | T | 0.932546 | 6.46E-23 | 2.53E-17 |
| 11 | 65460664 | C | G | 0.932546 | 6.46E-23 | 2.53E-17 |
| 11 | 65461036 | C | T | 0.932546 | 6.46E-23 | 2.53E-17 |
| 11 | 65461248 | A | G | 0.932546 | 6.46E-23 | 2.53E-17 |
| 11 | 65461284 | T | C | 0.932546 | 6.46E-23 | 2.53E-17 |
| 11 | 65461327 | A | G | 0.932546 | 6.46E-23 | 2.53E-17 |
| 11 | 65461328 | G | T | 0.932546 | 6.46E-23 | 2.53E-17 |
| 11 | 65461419 | A | G | 0.932546 | 6.46E-23 | 2.53E-17 |
| 11 | 65462928 | T | C | 0.932546 | 6.46E-23 | 2.53E-17 |
| 11 | 65463140 | G | T | 0.932546 | 6.46E-23 | 2.53E-17 |
| 11 | 65463520 | A | G | 0.932546 | 6.46E-23 | 2.53E-17 |
| 11 | 65463905 | T | C | 0.932546 | 6.46E-23 | 2.53E-17 |
| 11 | 65463978 | C | G | 0.932546 | 6.46E-23 | 2.53E-17 |
| 11 | 65464150 | A | T | 0.932546 | 6.46E-23 | 2.53E-17 |

| | | | | | | |
|---|---|---|---|---|---|---|
| 11 | 65464177 | G | T | 0.932546 | 6.46E-23 | 2.53E-17 |
| 11 | 65464203 | C | T | 0.932546 | 6.46E-23 | 2.53E-17 |
| 11 | 65464253 | A | G | 0.932546 | 6.46E-23 | 2.53E-17 |
| 11 | 65464321 | C | A | 0.932546 | 6.46E-23 | 2.53E-17 |
| 11 | 65464329 | A | G | 0.932546 | 6.46E-23 | 2.53E-17 |
| 11 | 65465313 | C | T | 0.932546 | 6.46E-23 | 2.53E-17 |
| 11 | 65472056 | C | G | 0.932546 | 6.46E-23 | 2.53E-17 |
| 11 | 65488082 | A | G | 0.932546 | 6.46E-23 | 2.53E-17 |
| 11 | 65489950 | C | T | 0.932546 | 6.46E-23 | 2.53E-17 |
| 11 | 65490161 | C | T | 0.932546 | 6.46E-23 | 2.53E-17 |
| 11 | 65490909 | A | C | 0.932546 | 6.46E-23 | 2.53E-17 |
| 11 | 65490910 | T | C | 0.932546 | 6.46E-23 | 2.53E-17 |
| 11 | 65491251 | T | C | 0.932546 | 6.46E-23 | 2.53E-17 |
| 11 | 65492735 | A | G | 0.932546 | 6.46E-23 | 2.53E-17 |
| 11 | 65493232 | C | T | 0.932546 | 6.46E-23 | 2.53E-17 |
| 11 | 65493269 | G | A | 0.932546 | 6.46E-23 | 2.53E-17 |
| 11 | 65493278 | G | A | 0.932546 | 6.46E-23 | 2.53E-17 |
| 11 | 65493407 | A | G | 0.932546 | 6.46E-23 | 2.53E-17 |
| 11 | 65494013 | T | G | 0.932546 | 6.46E-23 | 2.53E-17 |
| 11 | 65494638 | G | A | 0.932546 | 6.46E-23 | 2.53E-17 |
| 11 | 65495101 | T | G | 0.932546 | 6.46E-23 | 2.53E-17 |
| 11 | 65496569 | T | G | 0.932546 | 6.46E-23 | 2.53E-17 |
| 11 | 65497210 | G | A | 0.932546 | 6.46E-23 | 2.53E-17 |
| 11 | 65497360 | C | T | 0.932546 | 6.46E-23 | 2.53E-17 |
| 11 | 65497398 | T | C | 0.932546 | 6.46E-23 | 2.53E-17 |
| 11 | 65497641 | G | T | 0.932546 | 6.46E-23 | 2.53E-17 |
| 11 | 65499208 | A | T | 0.932546 | 6.46E-23 | 2.53E-17 |
| 11 | 65500228 | C | A | 0.932546 | 6.46E-23 | 2.53E-17 |
| 11 | 65500300 | G | A | 0.932546 | 6.46E-23 | 2.53E-17 |
| 11 | 65500401 | T | G | 0.932546 | 6.46E-23 | 2.53E-17 |
| 11 | 65500664 | A | G | 0.932546 | 6.46E-23 | 2.53E-17 |
| 11 | 65500906 | C | T | 0.932546 | 6.46E-23 | 2.53E-17 |
| 11 | 65500917 | T | C | 0.932546 | 6.46E-23 | 2.53E-17 |
| 11 | 65500981 | C | A | 0.932546 | 6.46E-23 | 2.53E-17 |
| 11 | 65501155 | C | T | 0.932546 | 6.46E-23 | 2.53E-17 |
| 11 | 65501159 | G | A | 0.932546 | 6.46E-23 | 2.53E-17 |
| 11 | 65501185 | T | C | 0.932546 | 6.46E-23 | 2.53E-17 |
| 11 | 65501250 | T | G | 0.932546 | 6.46E-23 | 2.53E-17 |
| 11 | 65501327 | T | C | 0.932546 | 6.46E-23 | 2.53E-17 |
| 11 | 65501336 | A | G | 0.932546 | 6.46E-23 | 2.53E-17 |
| 11 | 65501339 | T | C | 0.932546 | 6.46E-23 | 2.53E-17 |
| 11 | 65501417 | C | T | 0.932546 | 6.46E-23 | 2.53E-17 |
| 11 | 65501527 | G | A | 0.932546 | 6.46E-23 | 2.53E-17 |
| 11 | 65502586 | G | A | 0.932546 | 6.46E-23 | 2.53E-17 |
| 11 | 65503370 | T | C | 0.932546 | 6.46E-23 | 2.53E-17 |
| 11 | 65503943 | C | T | 0.932546 | 6.46E-23 | 2.53E-17 |
| 11 | 65503965 | G | A | 0.932546 | 6.46E-23 | 2.53E-17 |
| 11 | 65505228 | T | C | 0.932546 | 6.46E-23 | 2.53E-17 |
| 11 | 65505261 | T | C | 0.932546 | 6.46E-23 | 2.53E-17 |
| 11 | 65505272 | C | T | 0.932546 | 6.46E-23 | 2.53E-17 |
| 11 | 65505499 | T | C | 0.932546 | 6.46E-23 | 2.53E-17 |
| 11 | 65505909 | C | T | 0.932546 | 6.46E-23 | 2.53E-17 |
| 11 | 65506074 | A | G | 0.932546 | 6.46E-23 | 2.53E-17 |
| 11 | 65506141 | G | A | 0.932546 | 6.46E-23 | 2.53E-17 |
| 11 | 65506268 | A | G | 0.932546 | 6.46E-23 | 2.53E-17 |
| 11 | 65506288 | G | A | 0.932546 | 6.46E-23 | 2.53E-17 |
| 11 | 65506948 | T | C | 0.932546 | 6.46E-23 | 2.53E-17 |
| 11 | 65507112 | C | T | 0.932546 | 6.46E-23 | 2.53E-17 |
| 11 | 65507148 | G | C | 0.932546 | 6.46E-23 | 2.53E-17 |
| 11 | 65507376 | A | G | 0.932546 | 6.46E-23 | 2.53E-17 |
| 11 | 65507810 | C | A | 0.932546 | 6.46E-23 | 2.53E-17 |
| 11 | 65507816 | C | A | 0.932546 | 6.46E-23 | 2.53E-17 |
| 11 | 65507818 | T | A | 0.932546 | 6.46E-23 | 2.53E-17 |
| 11 | 65507822 | C | A | 0.932546 | 6.46E-23 | 2.53E-17 |
| 11 | 65507828 | C | A | 0.932546 | 6.46E-23 | 2.53E-17 |
| 11 | 65507834 | C | A | 0.932546 | 6.46E-23 | 2.53E-17 |
| 11 | 65507842 | T | A | 0.932546 | 6.46E-23 | 2.53E-17 |

| | | | | | | |
|---|---|---|---|---|---|---|
| 11 | 65507939 | G | A | 0.932546 | 6.46E-23 | 2.53E-17 |
| 11 | 65507943 | G | A | 0.932546 | 6.46E-23 | 2.53E-17 |
| 11 | 65507945 | G | A | 0.932546 | 6.46E-23 | 2.53E-17 |
| 11 | 65507952 | G | T | 0.932546 | 6.46E-23 | 2.53E-17 |
| 11 | 65507967 | T | C | 0.932546 | 6.46E-23 | 2.53E-17 |
| 11 | 65507977 | T | C | 0.932546 | 6.46E-23 | 2.53E-17 |
| 11 | 65508000 | T | C | 0.932546 | 6.46E-23 | 2.53E-17 |
| 11 | 65508185 | A | T | 0.932546 | 6.46E-23 | 2.53E-17 |
| 11 | 65509629 | A | G | 0.932546 | 6.46E-23 | 2.53E-17 |
| 11 | 65509687 | C | T | 0.932546 | 6.46E-23 | 2.53E-17 |
| 11 | 65509831 | T | C | 0.932546 | 6.46E-23 | 2.53E-17 |
| 11 | 65509880 | A | G | 0.932546 | 6.46E-23 | 2.53E-17 |
| 11 | 65509986 | A | C | 0.932546 | 6.46E-23 | 2.53E-17 |
| 11 | 65510812 | T | G | 0.932546 | 6.46E-23 | 2.53E-17 |
| 11 | 65511339 | T | A | 0.932546 | 6.46E-23 | 2.53E-17 |
| 11 | 65511550 | A | G | 0.932546 | 6.46E-23 | 2.53E-17 |
| 11 | 65511971 | T | A | 0.932546 | 6.46E-23 | 2.53E-17 |
| 11 | 65512710 | G | A | 0.932546 | 6.46E-23 | 2.53E-17 |
| 11 | 65512786 | G | A | 0.932546 | 6.46E-23 | 2.53E-17 |
| 11 | 65513021 | C | G | 0.932546 | 6.46E-23 | 2.53E-17 |
| 11 | 65513833 | C | T | 0.932546 | 6.46E-23 | 2.53E-17 |
| 11 | 65514174 | A | C | 0.932546 | 6.46E-23 | 2.53E-17 |
| 11 | 65514539 | A | G | 0.932546 | 6.46E-23 | 2.53E-17 |
| 11 | 65514935 | C | T | 0.932546 | 6.46E-23 | 2.53E-17 |
| 11 | 65515200 | T | C | 0.932546 | 6.46E-23 | 2.53E-17 |
| 11 | 65515201 | C | A | 0.932546 | 6.46E-23 | 2.53E-17 |
| 11 | 65515216 | A | G | 0.932546 | 6.46E-23 | 2.53E-17 |
| 11 | 65515916 | G | A | 0.932546 | 6.46E-23 | 2.53E-17 |
| 11 | 65516791 | T | C | 0.932546 | 6.46E-23 | 2.53E-17 |
| 11 | 65516822 | C | T | 0.932546 | 6.46E-23 | 2.53E-17 |
| 11 | 65516866 | T | C | 0.932546 | 6.46E-23 | 2.53E-17 |
| 11 | 65518014 | G | A | 0.932546 | 6.46E-23 | 2.53E-17 |
| 11 | 65519665 | A | T | 0.932546 | 6.46E-23 | 2.53E-17 |
| 11 | 65519681 | A | G | 0.932546 | 6.46E-23 | 2.53E-17 |
| 11 | 65519724 | T | G | 0.932546 | 6.46E-23 | 2.53E-17 |
| 11 | 65520459 | A | G | 0.932546 | 6.46E-23 | 2.53E-17 |
| 11 | 65520690 | A | G | 0.932546 | 6.46E-23 | 2.53E-17 |
| 11 | 65521027 | C | T | 0.932546 | 6.46E-23 | 2.53E-17 |
| 11 | 65521856 | G | A | 0.932546 | 6.46E-23 | 2.53E-17 |
| 11 | 65522305 | T | C | 0.932546 | 6.46E-23 | 2.53E-17 |
| 11 | 65523091 | G | A | 0.932546 | 6.46E-23 | 2.53E-17 |
| 11 | 65524906 | C | T | 0.932546 | 6.46E-23 | 2.53E-17 |
| 11 | 65525007 | C | T | 0.932546 | 6.46E-23 | 2.53E-17 |
| 11 | 65525910 | G | T | 0.932546 | 6.46E-23 | 2.53E-17 |
| 11 | 65526124 | T | C | 0.932546 | 6.46E-23 | 2.53E-17 |
| 11 | 65526463 | C | T | 0.932546 | 6.46E-23 | 2.53E-17 |
| 11 | 65527368 | G | C | 0.932546 | 6.46E-23 | 2.53E-17 |
| 11 | 65528562 | T | G | 0.932546 | 6.46E-23 | 2.53E-17 |
| 11 | 65528569 | A | G | 0.932546 | 6.46E-23 | 2.53E-17 |
| 11 | 65530514 | A | G | 0.932546 | 6.46E-23 | 2.53E-17 |
| 11 | 65530873 | C | A | 0.932546 | 6.46E-23 | 2.53E-17 |
| 11 | 65531184 | T | C | 0.932546 | 6.46E-23 | 2.53E-17 |
| 11 | 65531489 | A | G | 0.932546 | 6.46E-23 | 2.53E-17 |
| 11 | 65531868 | T | C | 0.932546 | 6.46E-23 | 2.53E-17 |
| 11 | 65541356 | T | C | 0.932546 | 6.46E-23 | 2.53E-17 |
| 11 | 65545215 | A | G | 0.932546 | 6.46E-23 | 2.53E-17 |
| 11 | 65547854 | A | C | 0.932546 | 6.46E-23 | 2.53E-17 |
| 11 | 65550124 | T | C | 0.932546 | 6.46E-23 | 2.53E-17 |
| 11 | 65550191 | A | T | 0.932546 | 6.46E-23 | 2.53E-17 |
| 11 | 65551525 | G | A | 0.932546 | 6.46E-23 | 2.53E-17 |
| 11 | 65551627 | A | G | 0.932546 | 6.46E-23 | 2.53E-17 |
| 11 | 65552378 | G | A | 0.932546 | 6.46E-23 | 2.53E-17 |
| 11 | 65560188 | T | C | 0.932546 | 6.46E-23 | 2.53E-17 |
| 11 | 65561476 | C | T | 0.932546 | 6.46E-23 | 2.53E-17 |
| 11 | 65573200 | C | T | 0.932546 | 6.46E-23 | 2.53E-17 |
| 11 | 65576340 | C | T | 0.932546 | 6.46E-23 | 2.53E-17 |
| 11 | 65576434 | G | A | 0.932546 | 6.46E-23 | 2.53E-17 |

| | | | | | | |
|---|---|---|---|---|---|---|
| 11 | 65577062 | A | G | 0.932546 | 6.46E-23 | 2.53E-17 |
| 11 | 65578244 | A | T | 0.932546 | 6.46E-23 | 2.53E-17 |
| 11 | 65578943 | T | C | 0.932546 | 6.46E-23 | 2.53E-17 |
| 11 | 65579413 | A | G | 0.932546 | 6.46E-23 | 2.53E-17 |
| 11 | 65579780 | A | G | 0.932546 | 6.46E-23 | 2.53E-17 |
| 11 | 65580262 | T | A | 0.932546 | 6.46E-23 | 2.53E-17 |
| 11 | 65990733 | T | G | 0.932546 | 6.46E-23 | 2.53E-17 |
| 11 | 65990767 | C | T | 0.932546 | 6.46E-23 | 2.53E-17 |
| 11 | 65991066 | A | C | 0.932546 | 6.46E-23 | 2.53E-17 |
| 11 | 65992538 | A | C | 0.932546 | 6.46E-23 | 2.53E-17 |
| 11 | 65992615 | C | T | 0.932546 | 6.46E-23 | 2.53E-17 |
| 11 | 65992820 | G | A | 0.932546 | 6.46E-23 | 2.53E-17 |
| 11 | 65992921 | C | T | 0.932546 | 6.46E-23 | 2.53E-17 |
| 11 | 65993024 | G | A | 0.932546 | 6.46E-23 | 2.53E-17 |
| 11 | 65996628 | C | T | 0.932546 | 6.46E-23 | 2.53E-17 |
| 11 | 64820343 | C | G | 0.918212 | 1.60E-22 | 6.27E-17 |
| 11 | 63237429 | C | T | 0.927487 | 2.11E-22 | 8.25E-17 |
| 11 | 63255501 | C | T | 0.927487 | 2.11E-22 | 8.25E-17 |
| 11 | 65488769 | A | G | 0.931703 | 3.86E-22 | 1.51E-16 |
| 11 | 65489814 | T | A | 0.931703 | 3.86E-22 | 1.51E-16 |
| 11 | 65490249 | T | A | 0.931703 | 3.86E-22 | 1.51E-16 |
| 11 | 65493282 | G | A | 0.931703 | 3.86E-22 | 1.51E-16 |
| 11 | 65495869 | A | G | 0.931703 | 3.86E-22 | 1.51E-16 |
| 11 | 64395863 | A | C | 0.930017 | 7.12E-22 | 2.79E-16 |
| 11 | 64399779 | T | C | 0.930017 | 7.12E-22 | 2.79E-16 |
| 11 | 65532744 | G | A | 0.93086 | 7.82E-22 | 3.07E-16 |
| 11 | 65532792 | T | G | 0.93086 | 7.82E-22 | 3.07E-16 |
| 11 | 65532933 | C | T | 0.93086 | 7.82E-22 | 3.07E-16 |
| 11 | 65532942 | C | T | 0.93086 | 7.82E-22 | 3.07E-16 |
| 11 | 65533113 | T | A | 0.93086 | 7.82E-22 | 3.07E-16 |
| 11 | 65533150 | A | C | 0.93086 | 7.82E-22 | 3.07E-16 |
| 11 | 65533225 | T | A | 0.93086 | 7.82E-22 | 3.07E-16 |
| 11 | 65533226 | C | A | 0.93086 | 7.82E-22 | 3.07E-16 |
| 11 | 65533239 | G | A | 0.93086 | 7.82E-22 | 3.07E-16 |
| 11 | 65533241 | G | A | 0.93086 | 7.82E-22 | 3.07E-16 |
| 11 | 66089675 | A | G | 0.932546 | 3.40E-21 | 1.33E-15 |
| 11 | 65991238 | G | A | 0.931703 | 5.45E-21 | 2.14E-15 |
| 11 | 65991351 | A | C | 0.931703 | 5.45E-21 | 2.14E-15 |
| 11 | 65991601 | A | G | 0.931703 | 5.45E-21 | 2.14E-15 |
| 11 | 65991602 | T | C | 0.931703 | 5.45E-21 | 2.14E-15 |
| 11 | 65991833 | T | G | 0.931703 | 5.45E-21 | 2.14E-15 |
| 11 | 65991915 | T | C | 0.931703 | 5.45E-21 | 2.14E-15 |
| 11 | 65991986 | T | C | 0.931703 | 5.45E-21 | 2.14E-15 |
| 11 | 65992414 | G | A | 0.931703 | 5.45E-21 | 2.14E-15 |
| 11 | 65992466 | C | T | 0.931703 | 5.45E-21 | 2.14E-15 |
| 11 | 65994518 | A | G | 0.902192 | 6.92E-20 | 2.71E-14 |
| 11 | 64944412 | C | T | 0.902192 | 6.94E-20 | 2.72E-14 |
| 11 | 65589070 | C | T | 0.930017 | 1.09E-19 | 4.27E-14 |
| 11 | 66031752 | T | C | 0.900506 | 2.97E-19 | 1.17E-13 |
| 11 | 64965032 | T | C | 0.901349 | 3.68E-19 | 1.44E-13 |
| 11 | 65994586 | C | G | 0.901349 | 4.85E-19 | 1.90E-13 |
| 11 | 65994958 | A | T | 0.901349 | 4.85E-19 | 1.90E-13 |
| 11 | 65995114 | A | C | 0.901349 | 4.85E-19 | 1.90E-13 |
| 11 | 65996864 | A | T | 0.901349 | 4.85E-19 | 1.90E-13 |
| 11 | 65997066 | G | T | 0.901349 | 4.85E-19 | 1.90E-13 |
| 11 | 65997282 | T | G | 0.901349 | 4.85E-19 | 1.90E-13 |
| 11 | 65997443 | G | C | 0.901349 | 4.85E-19 | 1.90E-13 |
| 11 | 65997614 | C | G | 0.901349 | 4.85E-19 | 1.90E-13 |
| 11 | 65999055 | C | T | 0.901349 | 4.85E-19 | 1.90E-13 |
| 11 | 65999088 | G | T | 0.901349 | 4.85E-19 | 1.90E-13 |
| 11 | 65999379 | T | C | 0.901349 | 4.85E-19 | 1.90E-13 |
| 11 | 66000061 | T | C | 0.901349 | 4.85E-19 | 1.90E-13 |
| 11 | 66000448 | A | G | 0.901349 | 4.85E-19 | 1.90E-13 |
| 11 | 66000541 | G | T | 0.901349 | 4.85E-19 | 1.90E-13 |
| 11 | 66000669 | C | T | 0.901349 | 4.85E-19 | 1.90E-13 |
| 11 | 66000821 | C | A | 0.901349 | 4.85E-19 | 1.90E-13 |
| 11 | 66000948 | T | C | 0.901349 | 4.85E-19 | 1.90E-13 |

| | | | | | | |
|---|---|---|---|---|---|---|
| 11 | 66000962 | A | T | 0.901349 | 4.85E-19 | 1.90E-13 |
| 11 | 66001174 | G | A | 0.901349 | 4.85E-19 | 1.90E-13 |
| 11 | 66001175 | G | A | 0.901349 | 4.85E-19 | 1.90E-13 |
| 11 | 66001417 | A | C | 0.901349 | 4.85E-19 | 1.90E-13 |
| 11 | 66001787 | G | A | 0.901349 | 4.85E-19 | 1.90E-13 |
| 11 | 66001840 | G | A | 0.901349 | 4.85E-19 | 1.90E-13 |
| 11 | 66001897 | T | C | 0.901349 | 4.85E-19 | 1.90E-13 |
| 11 | 66001934 | G | A | 0.901349 | 4.85E-19 | 1.90E-13 |
| 11 | 66001952 | A | T | 0.901349 | 4.85E-19 | 1.90E-13 |
| 11 | 66001976 | C | T | 0.901349 | 4.85E-19 | 1.90E-13 |
| 11 | 66001993 | G | A | 0.901349 | 4.85E-19 | 1.90E-13 |
| 11 | 66002062 | A | G | 0.901349 | 4.85E-19 | 1.90E-13 |
| 11 | 66002103 | C | G | 0.901349 | 4.85E-19 | 1.90E-13 |
| 11 | 66002107 | C | T | 0.901349 | 4.85E-19 | 1.90E-13 |
| 11 | 66002151 | T | C | 0.901349 | 4.85E-19 | 1.90E-13 |
| 11 | 66002161 | C | T | 0.901349 | 4.85E-19 | 1.90E-13 |
| 11 | 66002320 | G | T | 0.901349 | 4.85E-19 | 1.90E-13 |
| 11 | 66002342 | A | C | 0.901349 | 4.85E-19 | 1.90E-13 |
| 11 | 66002412 | C | T | 0.901349 | 4.85E-19 | 1.90E-13 |
| 11 | 66002524 | T | G | 0.901349 | 4.85E-19 | 1.90E-13 |
| 11 | 66003083 | C | T | 0.901349 | 4.85E-19 | 1.90E-13 |
| 11 | 66003459 | C | T | 0.901349 | 4.85E-19 | 1.90E-13 |
| 11 | 66003556 | C | T | 0.901349 | 4.85E-19 | 1.90E-13 |
| 11 | 66003626 | C | T | 0.901349 | 4.85E-19 | 1.90E-13 |
| 11 | 66003650 | T | G | 0.901349 | 4.85E-19 | 1.90E-13 |
| 11 | 66003679 | C | G | 0.901349 | 4.85E-19 | 1.90E-13 |
| 11 | 66003933 | T | A | 0.901349 | 4.85E-19 | 1.90E-13 |
| 11 | 66003941 | C | T | 0.901349 | 4.85E-19 | 1.90E-13 |
| 11 | 66003942 | C | T | 0.901349 | 4.85E-19 | 1.90E-13 |
| 11 | 66004181 | G | C | 0.901349 | 4.85E-19 | 1.90E-13 |
| 11 | 66004201 | C | A | 0.901349 | 4.85E-19 | 1.90E-13 |
| 11 | 66004202 | C | A | 0.901349 | 4.85E-19 | 1.90E-13 |
| 11 | 66004220 | C | T | 0.901349 | 4.85E-19 | 1.90E-13 |
| 11 | 66004342 | G | C | 0.901349 | 4.85E-19 | 1.90E-13 |
| 11 | 66004354 | T | C | 0.901349 | 4.85E-19 | 1.90E-13 |
| 11 | 66004370 | T | G | 0.901349 | 4.85E-19 | 1.90E-13 |
| 11 | 66004399 | G | A | 0.901349 | 4.85E-19 | 1.90E-13 |
| 11 | 66004413 | C | T | 0.901349 | 4.85E-19 | 1.90E-13 |
| 11 | 66004435 | T | C | 0.901349 | 4.85E-19 | 1.90E-13 |
| 11 | 66004456 | A | T | 0.901349 | 4.85E-19 | 1.90E-13 |
| 11 | 66004533 | T | C | 0.901349 | 4.85E-19 | 1.90E-13 |
| 11 | 66004605 | C | G | 0.901349 | 4.85E-19 | 1.90E-13 |
| 11 | 66004609 | C | G | 0.901349 | 4.85E-19 | 1.90E-13 |
| 11 | 66004625 | T | C | 0.901349 | 4.85E-19 | 1.90E-13 |
| 11 | 66004626 | A | G | 0.901349 | 4.85E-19 | 1.90E-13 |
| 11 | 66004628 | T | C | 0.901349 | 4.85E-19 | 1.90E-13 |
| 11 | 66004648 | A | G | 0.901349 | 4.85E-19 | 1.90E-13 |
| 11 | 66004670 | G | A | 0.901349 | 4.85E-19 | 1.90E-13 |
| 11 | 66004673 | A | G | 0.901349 | 4.85E-19 | 1.90E-13 |
| 11 | 66004708 | G | T | 0.901349 | 4.85E-19 | 1.90E-13 |
| 11 | 66004820 | G | A | 0.901349 | 4.85E-19 | 1.90E-13 |
| 11 | 66004833 | T | A | 0.901349 | 4.85E-19 | 1.90E-13 |
| 11 | 66004847 | T | C | 0.901349 | 4.85E-19 | 1.90E-13 |
| 11 | 66004852 | T | C | 0.901349 | 4.85E-19 | 1.90E-13 |
| 11 | 66005033 | C | T | 0.901349 | 4.85E-19 | 1.90E-13 |
| 11 | 66005065 | C | T | 0.901349 | 4.85E-19 | 1.90E-13 |
| 11 | 66005142 | T | C | 0.901349 | 4.85E-19 | 1.90E-13 |
| 11 | 66005292 | C | T | 0.901349 | 4.85E-19 | 1.90E-13 |
| 11 | 66005360 | A | C | 0.901349 | 4.85E-19 | 1.90E-13 |
| 11 | 66005378 | T | G | 0.901349 | 4.85E-19 | 1.90E-13 |
| 11 | 66005451 | A | G | 0.901349 | 4.85E-19 | 1.90E-13 |
| 11 | 66005472 | G | T | 0.901349 | 4.85E-19 | 1.90E-13 |
| 11 | 66005527 | T | C | 0.901349 | 4.85E-19 | 1.90E-13 |
| 11 | 66005550 | A | G | 0.901349 | 4.85E-19 | 1.90E-13 |
| 11 | 66005576 | T | C | 0.901349 | 4.85E-19 | 1.90E-13 |
| 11 | 66005579 | C | T | 0.901349 | 4.85E-19 | 1.90E-13 |
| 11 | 66005617 | C | T | 0.901349 | 4.85E-19 | 1.90E-13 |

| | | | | | | |
|---|---|---|---|---|---|---|
| 11 | 66005627 | C | T | 0.901349 | 4.85E-19 | 1.90E-13 |
| 11 | 66005628 | C | T | 0.901349 | 4.85E-19 | 1.90E-13 |
| 11 | 66005712 | T | C | 0.901349 | 4.85E-19 | 1.90E-13 |
| 11 | 66005716 | C | G | 0.901349 | 4.85E-19 | 1.90E-13 |
| 11 | 66005719 | G | A | 0.901349 | 4.85E-19 | 1.90E-13 |
| 11 | 66005757 | T | C | 0.901349 | 4.85E-19 | 1.90E-13 |
| 11 | 66005764 | C | T | 0.901349 | 4.85E-19 | 1.90E-13 |
| 11 | 66005773 | T | C | 0.901349 | 4.85E-19 | 1.90E-13 |
| 11 | 66005774 | C | G | 0.901349 | 4.85E-19 | 1.90E-13 |
| 11 | 66005787 | A | G | 0.901349 | 4.85E-19 | 1.90E-13 |
| 11 | 66005814 | C | T | 0.901349 | 4.85E-19 | 1.90E-13 |
| 11 | 66005855 | A | G | 0.901349 | 4.85E-19 | 1.90E-13 |
| 11 | 66006375 | T | C | 0.901349 | 4.85E-19 | 1.90E-13 |
| 11 | 66006384 | T | C | 0.901349 | 4.85E-19 | 1.90E-13 |
| 11 | 66006535 | G | T | 0.901349 | 4.85E-19 | 1.90E-13 |
| 11 | 66006587 | G | T | 0.901349 | 4.85E-19 | 1.90E-13 |
| 11 | 66006669 | C | A | 0.901349 | 4.85E-19 | 1.90E-13 |
| 11 | 66006765 | T | C | 0.901349 | 4.85E-19 | 1.90E-13 |
| 11 | 66006779 | G | T | 0.901349 | 4.85E-19 | 1.90E-13 |
| 11 | 66006908 | G | A | 0.901349 | 4.85E-19 | 1.90E-13 |
| 11 | 66007178 | A | T | 0.901349 | 4.85E-19 | 1.90E-13 |
| 11 | 66007191 | C | A | 0.901349 | 4.85E-19 | 1.90E-13 |
| 11 | 66007261 | A | C | 0.901349 | 4.85E-19 | 1.90E-13 |
| 11 | 66007298 | G | T | 0.901349 | 4.85E-19 | 1.90E-13 |
| 11 | 66007304 | G | A | 0.901349 | 4.85E-19 | 1.90E-13 |
| 11 | 66007370 | C | T | 0.901349 | 4.85E-19 | 1.90E-13 |
| 11 | 66007485 | A | C | 0.901349 | 4.85E-19 | 1.90E-13 |
| 11 | 66007525 | C | T | 0.901349 | 4.85E-19 | 1.90E-13 |
| 11 | 66007958 | A | C | 0.901349 | 4.85E-19 | 1.90E-13 |
| 11 | 66008030 | C | A | 0.901349 | 4.85E-19 | 1.90E-13 |
| 11 | 66008184 | C | T | 0.901349 | 4.85E-19 | 1.90E-13 |
| 11 | 66008382 | G | A | 0.901349 | 4.85E-19 | 1.90E-13 |
| 11 | 66008387 | T | C | 0.901349 | 4.85E-19 | 1.90E-13 |
| 11 | 66008501 | G | A | 0.901349 | 4.85E-19 | 1.90E-13 |
| 11 | 66008582 | A | G | 0.901349 | 4.85E-19 | 1.90E-13 |
| 11 | 66008630 | G | A | 0.901349 | 4.85E-19 | 1.90E-13 |
| 11 | 66008696 | A | G | 0.901349 | 4.85E-19 | 1.90E-13 |
| 11 | 66008789 | A | T | 0.901349 | 4.85E-19 | 1.90E-13 |
| 11 | 66008882 | C | G | 0.901349 | 4.85E-19 | 1.90E-13 |
| 11 | 66009078 | G | C | 0.901349 | 4.85E-19 | 1.90E-13 |
| 11 | 66009106 | T | C | 0.901349 | 4.85E-19 | 1.90E-13 |
| 11 | 66009109 | C | A | 0.901349 | 4.85E-19 | 1.90E-13 |
| 11 | 66009229 | T | C | 0.901349 | 4.85E-19 | 1.90E-13 |
| 11 | 66009251 | A | C | 0.901349 | 4.85E-19 | 1.90E-13 |
| 11 | 66009477 | A | G | 0.901349 | 4.85E-19 | 1.90E-13 |
| 11 | 66009503 | A | T | 0.901349 | 4.85E-19 | 1.90E-13 |
| 11 | 66009536 | C | T | 0.901349 | 4.85E-19 | 1.90E-13 |
| 11 | 66009598 | A | G | 0.901349 | 4.85E-19 | 1.90E-13 |
| 11 | 66009715 | T | C | 0.901349 | 4.85E-19 | 1.90E-13 |
| 11 | 66009840 | C | A | 0.901349 | 4.85E-19 | 1.90E-13 |
| 11 | 66009857 | G | A | 0.901349 | 4.85E-19 | 1.90E-13 |
| 11 | 66009882 | C | T | 0.901349 | 4.85E-19 | 1.90E-13 |
| 11 | 66009912 | C | G | 0.901349 | 4.85E-19 | 1.90E-13 |
| 11 | 66009914 | A | C | 0.901349 | 4.85E-19 | 1.90E-13 |
| 11 | 66009982 | C | T | 0.901349 | 4.85E-19 | 1.90E-13 |
| 11 | 66010034 | G | C | 0.901349 | 4.85E-19 | 1.90E-13 |
| 11 | 66010041 | G | T | 0.901349 | 4.85E-19 | 1.90E-13 |
| 11 | 66010050 | T | C | 0.901349 | 4.85E-19 | 1.90E-13 |
| 11 | 66010065 | C | G | 0.901349 | 4.85E-19 | 1.90E-13 |
| 11 | 66010134 | A | T | 0.901349 | 4.85E-19 | 1.90E-13 |
| 11 | 66010139 | T | C | 0.901349 | 4.85E-19 | 1.90E-13 |
| 11 | 66010364 | C | G | 0.901349 | 4.85E-19 | 1.90E-13 |
| 11 | 66010398 | T | C | 0.901349 | 4.85E-19 | 1.90E-13 |
| 11 | 66010439 | A | G | 0.901349 | 4.85E-19 | 1.90E-13 |
| 11 | 66010483 | C | T | 0.901349 | 4.85E-19 | 1.90E-13 |
| 11 | 66010512 | G | A | 0.901349 | 4.85E-19 | 1.90E-13 |
| 11 | 66010528 | A | G | 0.901349 | 4.85E-19 | 1.90E-13 |

| | | | | | | |
|---|---|---|---|---|---|---|
| 11 | 66010621 | G | T | 0.901349 | 4.85E-19 | 1.90E-13 |
| 11 | 66010634 | G | A | 0.901349 | 4.85E-19 | 1.90E-13 |
| 11 | 66010884 | A | T | 0.901349 | 4.85E-19 | 1.90E-13 |
| 11 | 66010920 | C | T | 0.901349 | 4.85E-19 | 1.90E-13 |
| 11 | 66010921 | C | T | 0.901349 | 4.85E-19 | 1.90E-13 |
| 11 | 66010927 | A | G | 0.901349 | 4.85E-19 | 1.90E-13 |
| 11 | 66010943 | C | A | 0.901349 | 4.85E-19 | 1.90E-13 |
| 11 | 66010953 | C | T | 0.901349 | 4.85E-19 | 1.90E-13 |
| 11 | 66010983 | A | G | 0.901349 | 4.85E-19 | 1.90E-13 |
| 11 | 66011059 | T | C | 0.901349 | 4.85E-19 | 1.90E-13 |
| 11 | 66011075 | G | A | 0.901349 | 4.85E-19 | 1.90E-13 |
| 11 | 66011099 | G | A | 0.901349 | 4.85E-19 | 1.90E-13 |
| 11 | 66011204 | A | G | 0.901349 | 4.85E-19 | 1.90E-13 |
| 11 | 66011238 | G | C | 0.901349 | 4.85E-19 | 1.90E-13 |
| 11 | 66011262 | A | T | 0.901349 | 4.85E-19 | 1.90E-13 |
| 11 | 66011306 | T | C | 0.901349 | 4.85E-19 | 1.90E-13 |
| 11 | 66011334 | G | A | 0.901349 | 4.85E-19 | 1.90E-13 |
| 11 | 66011531 | C | T | 0.901349 | 4.85E-19 | 1.90E-13 |
| 11 | 66011613 | A | T | 0.901349 | 4.85E-19 | 1.90E-13 |
| 11 | 66011691 | G | C | 0.901349 | 4.85E-19 | 1.90E-13 |
| 11 | 66011709 | C | A | 0.901349 | 4.85E-19 | 1.90E-13 |
| 11 | 66011734 | C | T | 0.901349 | 4.85E-19 | 1.90E-13 |
| 11 | 66011754 | T | C | 0.901349 | 4.85E-19 | 1.90E-13 |
| 11 | 66011771 | A | G | 0.901349 | 4.85E-19 | 1.90E-13 |
| 11 | 66011776 | A | C | 0.901349 | 4.85E-19 | 1.90E-13 |
| 11 | 66011787 | A | G | 0.901349 | 4.85E-19 | 1.90E-13 |
| 11 | 66011821 | G | C | 0.901349 | 4.85E-19 | 1.90E-13 |
| 11 | 66011903 | C | T | 0.901349 | 4.85E-19 | 1.90E-13 |
| 11 | 66011931 | A | C | 0.901349 | 4.85E-19 | 1.90E-13 |
| 11 | 66011941 | T | C | 0.901349 | 4.85E-19 | 1.90E-13 |
| 11 | 66011950 | A | G | 0.901349 | 4.85E-19 | 1.90E-13 |
| 11 | 66011957 | G | A | 0.901349 | 4.85E-19 | 1.90E-13 |
| 11 | 66011969 | G | A | 0.901349 | 4.85E-19 | 1.90E-13 |
| 11 | 66011987 | C | A | 0.901349 | 4.85E-19 | 1.90E-13 |
| 11 | 66012048 | T | A | 0.901349 | 4.85E-19 | 1.90E-13 |
| 11 | 66012052 | T | A | 0.901349 | 4.85E-19 | 1.90E-13 |
| 11 | 66012218 | A | C | 0.901349 | 4.85E-19 | 1.90E-13 |
| 11 | 66012275 | T | C | 0.901349 | 4.85E-19 | 1.90E-13 |
| 11 | 66012473 | C | G | 0.901349 | 4.85E-19 | 1.90E-13 |
| 11 | 66012502 | A | G | 0.901349 | 4.85E-19 | 1.90E-13 |
| 11 | 66012507 | T | C | 0.901349 | 4.85E-19 | 1.90E-13 |
| 11 | 66012683 | C | T | 0.901349 | 4.85E-19 | 1.90E-13 |
| 11 | 66012783 | A | T | 0.901349 | 4.85E-19 | 1.90E-13 |
| 11 | 66012862 | T | A | 0.901349 | 4.85E-19 | 1.90E-13 |
| 11 | 66012895 | T | A | 0.901349 | 4.85E-19 | 1.90E-13 |
| 11 | 66012983 | T | G | 0.901349 | 4.85E-19 | 1.90E-13 |
| 11 | 66013003 | T | C | 0.901349 | 4.85E-19 | 1.90E-13 |
| 11 | 66013024 | A | G | 0.901349 | 4.85E-19 | 1.90E-13 |
| 11 | 66013053 | G | A | 0.901349 | 4.85E-19 | 1.90E-13 |
| 11 | 66013396 | T | A | 0.901349 | 4.85E-19 | 1.90E-13 |
| 11 | 66013397 | T | A | 0.901349 | 4.85E-19 | 1.90E-13 |
| 11 | 66013414 | C | T | 0.901349 | 4.85E-19 | 1.90E-13 |
| 11 | 66013441 | G | A | 0.901349 | 4.85E-19 | 1.90E-13 |
| 11 | 66013603 | T | C | 0.901349 | 4.85E-19 | 1.90E-13 |
| 11 | 66013708 | T | C | 0.901349 | 4.85E-19 | 1.90E-13 |
| 11 | 66013754 | T | C | 0.901349 | 4.85E-19 | 1.90E-13 |
| 11 | 66013759 | G | A | 0.901349 | 4.85E-19 | 1.90E-13 |
| 11 | 66013799 | A | G | 0.901349 | 4.85E-19 | 1.90E-13 |
| 11 | 66013809 | C | T | 0.901349 | 4.85E-19 | 1.90E-13 |
| 11 | 66013810 | C | G | 0.901349 | 4.85E-19 | 1.90E-13 |
| 11 | 66013855 | T | C | 0.901349 | 4.85E-19 | 1.90E-13 |
| 11 | 66013881 | A | G | 0.901349 | 4.85E-19 | 1.90E-13 |
| 11 | 66014190 | C | T | 0.901349 | 4.85E-19 | 1.90E-13 |
| 11 | 66014333 | G | C | 0.901349 | 4.85E-19 | 1.90E-13 |
| 11 | 66014830 | C | T | 0.901349 | 4.85E-19 | 1.90E-13 |
| 11 | 66014865 | G | T | 0.901349 | 4.85E-19 | 1.90E-13 |
| 11 | 66014911 | T | C | 0.901349 | 4.85E-19 | 1.90E-13 |

| | | | | | | |
|---|---|---|---|---|---|---|
| 11 | 66014972 | T | C | 0.901349 | 4.85E-19 | 1.90E-13 |
| 11 | 66015012 | G | T | 0.901349 | 4.85E-19 | 1.90E-13 |
| 11 | 66015020 | T | G | 0.901349 | 4.85E-19 | 1.90E-13 |
| 11 | 66015130 | C | T | 0.901349 | 4.85E-19 | 1.90E-13 |
| 11 | 66015194 | A | C | 0.901349 | 4.85E-19 | 1.90E-13 |
| 11 | 66016014 | C | G | 0.901349 | 4.85E-19 | 1.90E-13 |
| 11 | 66016135 | G | T | 0.901349 | 4.85E-19 | 1.90E-13 |
| 11 | 66016185 | A | G | 0.901349 | 4.85E-19 | 1.90E-13 |
| 11 | 66016553 | A | G | 0.901349 | 4.85E-19 | 1.90E-13 |
| 11 | 66016801 | C | T | 0.901349 | 4.85E-19 | 1.90E-13 |
| 11 | 66016946 | T | G | 0.901349 | 4.85E-19 | 1.90E-13 |
| 11 | 66017717 | A | T | 0.901349 | 4.85E-19 | 1.90E-13 |
| 11 | 66017744 | T | G | 0.901349 | 4.85E-19 | 1.90E-13 |
| 11 | 66017805 | G | A | 0.901349 | 4.85E-19 | 1.90E-13 |
| 11 | 66017828 | C | G | 0.901349 | 4.85E-19 | 1.90E-13 |
| 11 | 66017852 | T | A | 0.901349 | 4.85E-19 | 1.90E-13 |
| 11 | 66017895 | T | C | 0.901349 | 4.85E-19 | 1.90E-13 |
| 11 | 66017904 | G | C | 0.901349 | 4.85E-19 | 1.90E-13 |
| 11 | 66017907 | C | T | 0.901349 | 4.85E-19 | 1.90E-13 |
| 11 | 66018170 | C | T | 0.901349 | 4.85E-19 | 1.90E-13 |
| 11 | 66018256 | G | A | 0.901349 | 4.85E-19 | 1.90E-13 |
| 11 | 66018261 | T | G | 0.901349 | 4.85E-19 | 1.90E-13 |
| 11 | 66018432 | C | T | 0.901349 | 4.85E-19 | 1.90E-13 |
| 11 | 66018540 | C | A | 0.901349 | 4.85E-19 | 1.90E-13 |
| 11 | 66018598 | C | T | 0.901349 | 4.85E-19 | 1.90E-13 |
| 11 | 66018834 | A | T | 0.901349 | 4.85E-19 | 1.90E-13 |
| 11 | 66018979 | G | A | 0.901349 | 4.85E-19 | 1.90E-13 |
| 11 | 66019153 | C | T | 0.901349 | 4.85E-19 | 1.90E-13 |
| 11 | 66019292 | C | G | 0.901349 | 4.85E-19 | 1.90E-13 |
| 11 | 66019958 | T | C | 0.901349 | 4.85E-19 | 1.90E-13 |
| 11 | 66020188 | A | G | 0.901349 | 4.85E-19 | 1.90E-13 |
| 11 | 66020242 | T | C | 0.901349 | 4.85E-19 | 1.90E-13 |
| 11 | 66020519 | G | A | 0.901349 | 4.85E-19 | 1.90E-13 |
| 11 | 66020761 | T | C | 0.901349 | 4.85E-19 | 1.90E-13 |
| 11 | 66020913 | T | C | 0.901349 | 4.85E-19 | 1.90E-13 |
| 11 | 66020969 | G | A | 0.901349 | 4.85E-19 | 1.90E-13 |
| 11 | 66021128 | C | T | 0.901349 | 4.85E-19 | 1.90E-13 |
| 11 | 66021263 | C | T | 0.901349 | 4.85E-19 | 1.90E-13 |
| 11 | 66021327 | C | T | 0.901349 | 4.85E-19 | 1.90E-13 |
| 11 | 66021332 | C | T | 0.901349 | 4.85E-19 | 1.90E-13 |
| 11 | 66021347 | T | G | 0.901349 | 4.85E-19 | 1.90E-13 |
| 11 | 66021373 | C | T | 0.901349 | 4.85E-19 | 1.90E-13 |
| 11 | 66021394 | G | A | 0.901349 | 4.85E-19 | 1.90E-13 |
| 11 | 66021397 | T | C | 0.901349 | 4.85E-19 | 1.90E-13 |
| 11 | 66021511 | T | C | 0.901349 | 4.85E-19 | 1.90E-13 |
| 11 | 66021550 | A | T | 0.901349 | 4.85E-19 | 1.90E-13 |
| 11 | 66021807 | C | G | 0.901349 | 4.85E-19 | 1.90E-13 |
| 11 | 66022061 | C | T | 0.901349 | 4.85E-19 | 1.90E-13 |
| 11 | 66022178 | T | C | 0.901349 | 4.85E-19 | 1.90E-13 |
| 11 | 66022264 | T | C | 0.901349 | 4.85E-19 | 1.90E-13 |
| 11 | 66022301 | C | T | 0.901349 | 4.85E-19 | 1.90E-13 |
| 11 | 66022319 | C | A | 0.901349 | 4.85E-19 | 1.90E-13 |
| 11 | 66022376 | A | T | 0.901349 | 4.85E-19 | 1.90E-13 |
| 11 | 66022435 | G | T | 0.901349 | 4.85E-19 | 1.90E-13 |
| 11 | 66022439 | T | C | 0.901349 | 4.85E-19 | 1.90E-13 |
| 11 | 66022606 | G | C | 0.901349 | 4.85E-19 | 1.90E-13 |
| 11 | 66022708 | G | A | 0.901349 | 4.85E-19 | 1.90E-13 |
| 11 | 66022760 | G | A | 0.901349 | 4.85E-19 | 1.90E-13 |
| 11 | 66023047 | C | T | 0.901349 | 4.85E-19 | 1.90E-13 |
| 11 | 66023280 | T | C | 0.901349 | 4.85E-19 | 1.90E-13 |
| 11 | 66023323 | C | A | 0.901349 | 4.85E-19 | 1.90E-13 |
| 11 | 66023339 | C | A | 0.901349 | 4.85E-19 | 1.90E-13 |
| 11 | 66023343 | C | A | 0.901349 | 4.85E-19 | 1.90E-13 |
| 11 | 66023416 | G | A | 0.901349 | 4.85E-19 | 1.90E-13 |
| 11 | 66023505 | C | A | 0.901349 | 4.85E-19 | 1.90E-13 |
| 11 | 66023525 | C | T | 0.901349 | 4.85E-19 | 1.90E-13 |
| 11 | 66023762 | T | G | 0.901349 | 4.85E-19 | 1.90E-13 |

| | | | | | | |
|---|---|---|---|---|---|---|
| 11 | 66023947 | G | A | 0.901349 | 4.85E-19 | 1.90E-13 |
| 11 | 66024251 | T | C | 0.901349 | 4.85E-19 | 1.90E-13 |
| 11 | 66024390 | A | G | 0.901349 | 4.85E-19 | 1.90E-13 |
| 11 | 66024512 | C | G | 0.901349 | 4.85E-19 | 1.90E-13 |
| 11 | 66024761 | G | T | 0.901349 | 4.85E-19 | 1.90E-13 |
| 11 | 66024830 | A | G | 0.901349 | 4.85E-19 | 1.90E-13 |
| 11 | 66024866 | A | G | 0.901349 | 4.85E-19 | 1.90E-13 |
| 11 | 66024969 | G | A | 0.901349 | 4.85E-19 | 1.90E-13 |
| 11 | 66025084 | A | C | 0.901349 | 4.85E-19 | 1.90E-13 |
| 11 | 66025238 | A | G | 0.901349 | 4.85E-19 | 1.90E-13 |
| 11 | 66025255 | T | G | 0.901349 | 4.85E-19 | 1.90E-13 |
| 11 | 66025305 | A | G | 0.901349 | 4.85E-19 | 1.90E-13 |
| 11 | 66025347 | C | A | 0.901349 | 4.85E-19 | 1.90E-13 |
| 11 | 66025469 | C | A | 0.901349 | 4.85E-19 | 1.90E-13 |
| 11 | 66025497 | A | G | 0.901349 | 4.85E-19 | 1.90E-13 |
| 11 | 66025498 | T | C | 0.901349 | 4.85E-19 | 1.90E-13 |
| 11 | 66025555 | A | T | 0.901349 | 4.85E-19 | 1.90E-13 |
| 11 | 66025585 | T | A | 0.901349 | 4.85E-19 | 1.90E-13 |
| 11 | 66025638 | G | T | 0.901349 | 4.85E-19 | 1.90E-13 |
| 11 | 66025640 | A | C | 0.901349 | 4.85E-19 | 1.90E-13 |
| 11 | 66025642 | G | T | 0.901349 | 4.85E-19 | 1.90E-13 |
| 11 | 66025721 | T | G | 0.901349 | 4.85E-19 | 1.90E-13 |
| 11 | 66025807 | C | G | 0.901349 | 4.85E-19 | 1.90E-13 |
| 11 | 66025821 | T | G | 0.901349 | 4.85E-19 | 1.90E-13 |
| 11 | 66025955 | C | T | 0.901349 | 4.85E-19 | 1.90E-13 |
| 11 | 66025967 | T | C | 0.901349 | 4.85E-19 | 1.90E-13 |
| 11 | 66026019 | A | G | 0.901349 | 4.85E-19 | 1.90E-13 |
| 11 | 66026194 | A | T | 0.901349 | 4.85E-19 | 1.90E-13 |
| 11 | 66026208 | A | G | 0.901349 | 4.85E-19 | 1.90E-13 |
| 11 | 66026249 | C | G | 0.901349 | 4.85E-19 | 1.90E-13 |
| 11 | 66026392 | C | T | 0.901349 | 4.85E-19 | 1.90E-13 |
| 11 | 66026546 | G | A | 0.901349 | 4.85E-19 | 1.90E-13 |
| 11 | 66026617 | A | C | 0.901349 | 4.85E-19 | 1.90E-13 |
| 11 | 66026687 | C | A | 0.901349 | 4.85E-19 | 1.90E-13 |
| 11 | 66026850 | C | T | 0.901349 | 4.85E-19 | 1.90E-13 |
| 11 | 66026877 | T | C | 0.901349 | 4.85E-19 | 1.90E-13 |
| 11 | 66026948 | T | C | 0.901349 | 4.85E-19 | 1.90E-13 |
| 11 | 66027066 | A | G | 0.901349 | 4.85E-19 | 1.90E-13 |
| 11 | 66027155 | T | C | 0.901349 | 4.85E-19 | 1.90E-13 |
| 11 | 66027233 | C | T | 0.901349 | 4.85E-19 | 1.90E-13 |
| 11 | 66027393 | T | C | 0.901349 | 4.85E-19 | 1.90E-13 |
| 11 | 66027441 | G | A | 0.901349 | 4.85E-19 | 1.90E-13 |
| 11 | 66027470 | C | T | 0.901349 | 4.85E-19 | 1.90E-13 |
| 11 | 66027501 | G | A | 0.901349 | 4.85E-19 | 1.90E-13 |
| 11 | 66027554 | C | T | 0.901349 | 4.85E-19 | 1.90E-13 |
| 11 | 66027574 | T | C | 0.901349 | 4.85E-19 | 1.90E-13 |
| 11 | 66027627 | A | G | 0.901349 | 4.85E-19 | 1.90E-13 |
| 11 | 66027651 | A | C | 0.901349 | 4.85E-19 | 1.90E-13 |
| 11 | 66027655 | G | A | 0.901349 | 4.85E-19 | 1.90E-13 |
| 11 | 66027671 | C | A | 0.901349 | 4.85E-19 | 1.90E-13 |
| 11 | 66027673 | T | A | 0.901349 | 4.85E-19 | 1.90E-13 |
| 11 | 66027702 | T | G | 0.901349 | 4.85E-19 | 1.90E-13 |
| 11 | 66027771 | C | G | 0.901349 | 4.85E-19 | 1.90E-13 |
| 11 | 66027777 | A | G | 0.901349 | 4.85E-19 | 1.90E-13 |
| 11 | 66027899 | G | A | 0.901349 | 4.85E-19 | 1.90E-13 |
| 11 | 66027950 | T | C | 0.901349 | 4.85E-19 | 1.90E-13 |
| 11 | 66027958 | T | C | 0.901349 | 4.85E-19 | 1.90E-13 |
| 11 | 66027975 | T | C | 0.901349 | 4.85E-19 | 1.90E-13 |
| 11 | 66027980 | G | T | 0.901349 | 4.85E-19 | 1.90E-13 |
| 11 | 66028031 | G | A | 0.901349 | 4.85E-19 | 1.90E-13 |
| 11 | 66028043 | T | G | 0.901349 | 4.85E-19 | 1.90E-13 |
| 11 | 66028063 | T | C | 0.901349 | 4.85E-19 | 1.90E-13 |
| 11 | 66028098 | C | A | 0.901349 | 4.85E-19 | 1.90E-13 |
| 11 | 66028129 | A | C | 0.901349 | 4.85E-19 | 1.90E-13 |
| 11 | 66028144 | A | T | 0.901349 | 4.85E-19 | 1.90E-13 |
| 11 | 66028259 | C | T | 0.901349 | 4.85E-19 | 1.90E-13 |
| 11 | 66028278 | A | C | 0.901349 | 4.85E-19 | 1.90E-13 |

| | | | | | | |
|---|---|---|---|---|---|---|
| 11 | 66028339 | T | C | 0.901349 | 4.85E-19 | 1.90E-13 |
| 11 | 66028372 | T | C | 0.901349 | 4.85E-19 | 1.90E-13 |
| 11 | 66028395 | A | T | 0.901349 | 4.85E-19 | 1.90E-13 |
| 11 | 66028398 | G | T | 0.901349 | 4.85E-19 | 1.90E-13 |
| 11 | 66028418 | T | C | 0.901349 | 4.85E-19 | 1.90E-13 |
| 11 | 66028545 | A | G | 0.901349 | 4.85E-19 | 1.90E-13 |
| 11 | 66028546 | G | A | 0.901349 | 4.85E-19 | 1.90E-13 |
| 11 | 66028586 | C | A | 0.901349 | 4.85E-19 | 1.90E-13 |
| 11 | 66028625 | T | G | 0.901349 | 4.85E-19 | 1.90E-13 |
| 11 | 66028699 | A | G | 0.901349 | 4.85E-19 | 1.90E-13 |
| 11 | 66028709 | A | C | 0.901349 | 4.85E-19 | 1.90E-13 |
| 11 | 66028724 | T | C | 0.901349 | 4.85E-19 | 1.90E-13 |
| 11 | 66028838 | C | T | 0.901349 | 4.85E-19 | 1.90E-13 |
| 11 | 66028846 | T | G | 0.901349 | 4.85E-19 | 1.90E-13 |
| 11 | 66028876 | G | T | 0.901349 | 4.85E-19 | 1.90E-13 |
| 11 | 66028934 | C | A | 0.901349 | 4.85E-19 | 1.90E-13 |
| 11 | 66028958 | A | G | 0.901349 | 4.85E-19 | 1.90E-13 |
| 11 | 66028975 | G | A | 0.901349 | 4.85E-19 | 1.90E-13 |
| 11 | 66028983 | T | C | 0.901349 | 4.85E-19 | 1.90E-13 |
| 11 | 66028989 | G | C | 0.901349 | 4.85E-19 | 1.90E-13 |
| 11 | 66029030 | T | C | 0.901349 | 4.85E-19 | 1.90E-13 |
| 11 | 66029126 | C | G | 0.901349 | 4.85E-19 | 1.90E-13 |
| 11 | 66029154 | C | T | 0.901349 | 4.85E-19 | 1.90E-13 |
| 11 | 66029162 | A | G | 0.901349 | 4.85E-19 | 1.90E-13 |
| 11 | 66029177 | G | T | 0.901349 | 4.85E-19 | 1.90E-13 |
| 11 | 66029217 | C | T | 0.901349 | 4.85E-19 | 1.90E-13 |
| 11 | 66029235 | C | G | 0.901349 | 4.85E-19 | 1.90E-13 |
| 11 | 66029276 | C | G | 0.901349 | 4.85E-19 | 1.90E-13 |
| 11 | 66029281 | G | A | 0.901349 | 4.85E-19 | 1.90E-13 |
| 11 | 66029307 | G | A | 0.901349 | 4.85E-19 | 1.90E-13 |
| 11 | 66029335 | A | T | 0.901349 | 4.85E-19 | 1.90E-13 |
| 11 | 66029400 | A | G | 0.901349 | 4.85E-19 | 1.90E-13 |
| 11 | 66029445 | C | A | 0.901349 | 4.85E-19 | 1.90E-13 |
| 11 | 66029593 | T | C | 0.901349 | 4.85E-19 | 1.90E-13 |
| 11 | 66029633 | A | C | 0.901349 | 4.85E-19 | 1.90E-13 |
| 11 | 66029679 | C | T | 0.901349 | 4.85E-19 | 1.90E-13 |
| 11 | 66029697 | A | C | 0.901349 | 4.85E-19 | 1.90E-13 |
| 11 | 66029758 | C | T | 0.901349 | 4.85E-19 | 1.90E-13 |
| 11 | 66029856 | A | T | 0.901349 | 4.85E-19 | 1.90E-13 |
| 11 | 66029858 | A | T | 0.901349 | 4.85E-19 | 1.90E-13 |
| 11 | 66029931 | G | T | 0.901349 | 4.85E-19 | 1.90E-13 |
| 11 | 66029936 | G | T | 0.901349 | 4.85E-19 | 1.90E-13 |
| 11 | 66029942 | G | T | 0.901349 | 4.85E-19 | 1.90E-13 |
| 11 | 66029988 | A | C | 0.901349 | 4.85E-19 | 1.90E-13 |
| 11 | 66030059 | T | A | 0.901349 | 4.85E-19 | 1.90E-13 |
| 11 | 66030066 | T | C | 0.901349 | 4.85E-19 | 1.90E-13 |
| 11 | 66030101 | T | A | 0.901349 | 4.85E-19 | 1.90E-13 |
| 11 | 66030218 | C | T | 0.901349 | 4.85E-19 | 1.90E-13 |
| 11 | 66030257 | C | T | 0.901349 | 4.85E-19 | 1.90E-13 |
| 11 | 66030338 | C | A | 0.901349 | 4.85E-19 | 1.90E-13 |
| 11 | 66030403 | C | T | 0.901349 | 4.85E-19 | 1.90E-13 |
| 11 | 66030414 | C | T | 0.901349 | 4.85E-19 | 1.90E-13 |
| 11 | 66030549 | T | C | 0.901349 | 4.85E-19 | 1.90E-13 |
| 11 | 66030556 | G | A | 0.901349 | 4.85E-19 | 1.90E-13 |
| 11 | 66030567 | A | G | 0.901349 | 4.85E-19 | 1.90E-13 |
| 11 | 66030574 | A | G | 0.901349 | 4.85E-19 | 1.90E-13 |
| 11 | 66030575 | C | T | 0.901349 | 4.85E-19 | 1.90E-13 |
| 11 | 66030580 | C | A | 0.901349 | 4.85E-19 | 1.90E-13 |
| 11 | 66030581 | T | C | 0.901349 | 4.85E-19 | 1.90E-13 |
| 11 | 66030622 | T | C | 0.901349 | 4.85E-19 | 1.90E-13 |
| 11 | 66030644 | T | C | 0.901349 | 4.85E-19 | 1.90E-13 |
| 11 | 66030650 | C | T | 0.901349 | 4.85E-19 | 1.90E-13 |
| 11 | 66030653 | T | C | 0.901349 | 4.85E-19 | 1.90E-13 |
| 11 | 66030654 | G | C | 0.901349 | 4.85E-19 | 1.90E-13 |
| 11 | 66030662 | A | T | 0.901349 | 4.85E-19 | 1.90E-13 |
| 11 | 66030665 | T | C | 0.901349 | 4.85E-19 | 1.90E-13 |
| 11 | 66030684 | T | C | 0.901349 | 4.85E-19 | 1.90E-13 |

| | | | | | | |
|---|---|---|---|---|---|---|
| 11 | 66030685 | G | A | 0.901349 | 4.85E-19 | 1.90E-13 |
| 11 | 66030725 | G | C | 0.901349 | 4.85E-19 | 1.90E-13 |
| 11 | 66030760 | A | G | 0.901349 | 4.85E-19 | 1.90E-13 |
| 11 | 66030777 | G | C | 0.901349 | 4.85E-19 | 1.90E-13 |
| 11 | 66030867 | C | A | 0.901349 | 4.85E-19 | 1.90E-13 |
| 11 | 66030871 | G | A | 0.901349 | 4.85E-19 | 1.90E-13 |
| 11 | 66030879 | G | A | 0.901349 | 4.85E-19 | 1.90E-13 |
| 11 | 66030913 | A | G | 0.901349 | 4.85E-19 | 1.90E-13 |
| 11 | 66030925 | A | G | 0.901349 | 4.85E-19 | 1.90E-13 |
| 11 | 66030984 | C | T | 0.901349 | 4.85E-19 | 1.90E-13 |
| 11 | 66030989 | G | A | 0.901349 | 4.85E-19 | 1.90E-13 |
| 11 | 66030990 | G | A | 0.901349 | 4.85E-19 | 1.90E-13 |
| 11 | 66031000 | C | T | 0.901349 | 4.85E-19 | 1.90E-13 |
| 11 | 66031001 | A | G | 0.901349 | 4.85E-19 | 1.90E-13 |
| 11 | 66031009 | C | T | 0.901349 | 4.85E-19 | 1.90E-13 |
| 11 | 66031011 | T | C | 0.901349 | 4.85E-19 | 1.90E-13 |
| 11 | 66031016 | C | A | 0.901349 | 4.85E-19 | 1.90E-13 |
| 11 | 66031020 | T | C | 0.901349 | 4.85E-19 | 1.90E-13 |
| 11 | 66031036 | G | C | 0.901349 | 4.85E-19 | 1.90E-13 |
| 11 | 66031039 | G | A | 0.901349 | 4.85E-19 | 1.90E-13 |
| 11 | 66031040 | G | A | 0.901349 | 4.85E-19 | 1.90E-13 |
| 11 | 66031042 | A | G | 0.901349 | 4.85E-19 | 1.90E-13 |
| 11 | 66031044 | G | T | 0.901349 | 4.85E-19 | 1.90E-13 |
| 11 | 66031045 | T | C | 0.901349 | 4.85E-19 | 1.90E-13 |
| 11 | 66031047 | G | A | 0.901349 | 4.85E-19 | 1.90E-13 |
| 11 | 66031049 | C | T | 0.901349 | 4.85E-19 | 1.90E-13 |
| 11 | 66031058 | G | C | 0.901349 | 4.85E-19 | 1.90E-13 |
| 11 | 66031100 | T | A | 0.901349 | 4.85E-19 | 1.90E-13 |
| 11 | 66031113 | A | G | 0.901349 | 4.85E-19 | 1.90E-13 |
| 11 | 66031114 | T | A | 0.901349 | 4.85E-19 | 1.90E-13 |
| 11 | 66031134 | A | G | 0.901349 | 4.85E-19 | 1.90E-13 |
| 11 | 66031144 | A | C | 0.901349 | 4.85E-19 | 1.90E-13 |
| 11 | 66031193 | C | T | 0.901349 | 4.85E-19 | 1.90E-13 |
| 11 | 66031243 | C | T | 0.901349 | 4.85E-19 | 1.90E-13 |
| 11 | 66031301 | C | T | 0.901349 | 4.85E-19 | 1.90E-13 |
| 11 | 66031313 | A | C | 0.901349 | 4.85E-19 | 1.90E-13 |
| 11 | 66031325 | A | G | 0.901349 | 4.85E-19 | 1.90E-13 |
| 11 | 66031349 | A | G | 0.901349 | 4.85E-19 | 1.90E-13 |
| 11 | 66031380 | T | C | 0.901349 | 4.85E-19 | 1.90E-13 |
| 11 | 66031603 | C | T | 0.901349 | 4.85E-19 | 1.90E-13 |
| 11 | 66031604 | A | G | 0.901349 | 4.85E-19 | 1.90E-13 |
| 11 | 66031637 | G | A | 0.901349 | 4.85E-19 | 1.90E-13 |
| 11 | 66031685 | T | A | 0.901349 | 4.85E-19 | 1.90E-13 |
| 11 | 66031797 | A | C | 0.901349 | 4.85E-19 | 1.90E-13 |
| 11 | 66031846 | A | T | 0.901349 | 4.85E-19 | 1.90E-13 |
| 11 | 66031847 | T | G | 0.901349 | 4.85E-19 | 1.90E-13 |
| 11 | 66031876 | G | A | 0.901349 | 4.85E-19 | 1.90E-13 |
| 11 | 66031915 | C | T | 0.901349 | 4.85E-19 | 1.90E-13 |
| 11 | 66032001 | A | G | 0.901349 | 4.85E-19 | 1.90E-13 |
| 11 | 66032015 | C | A | 0.901349 | 4.85E-19 | 1.90E-13 |
| 11 | 66032041 | G | T | 0.901349 | 4.85E-19 | 1.90E-13 |
| 11 | 66032202 | C | T | 0.901349 | 4.85E-19 | 1.90E-13 |
| 11 | 66032205 | T | C | 0.901349 | 4.85E-19 | 1.90E-13 |
| 11 | 66032239 | T | C | 0.901349 | 4.85E-19 | 1.90E-13 |
| 11 | 66032306 | T | C | 0.901349 | 4.85E-19 | 1.90E-13 |
| 11 | 66032362 | A | G | 0.901349 | 4.85E-19 | 1.90E-13 |
| 11 | 66032365 | T | G | 0.901349 | 4.85E-19 | 1.90E-13 |
| 11 | 66032366 | C | T | 0.901349 | 4.85E-19 | 1.90E-13 |
| 11 | 66032389 | A | T | 0.901349 | 4.85E-19 | 1.90E-13 |
| 11 | 66032415 | A | G | 0.901349 | 4.85E-19 | 1.90E-13 |
| 11 | 66032449 | A | G | 0.901349 | 4.85E-19 | 1.90E-13 |
| 11 | 66032488 | T | C | 0.901349 | 4.85E-19 | 1.90E-13 |
| 11 | 66032489 | T | C | 0.901349 | 4.85E-19 | 1.90E-13 |
| 11 | 66032514 | T | C | 0.901349 | 4.85E-19 | 1.90E-13 |
| 11 | 66032537 | C | A | 0.901349 | 4.85E-19 | 1.90E-13 |
| 11 | 66032677 | G | A | 0.901349 | 4.85E-19 | 1.90E-13 |
| 11 | 66032683 | T | C | 0.901349 | 4.85E-19 | 1.90E-13 |

| | | | | | | |
|---|---|---|---|---|---|---|
| 11 | 66032689 | A | C | 0.901349 | 4.85E-19 | 1.90E-13 |
| 11 | 66032705 | C | T | 0.901349 | 4.85E-19 | 1.90E-13 |
| 11 | 66032719 | G | A | 0.901349 | 4.85E-19 | 1.90E-13 |
| 11 | 66032720 | C | T | 0.901349 | 4.85E-19 | 1.90E-13 |
| 11 | 66032725 | A | G | 0.901349 | 4.85E-19 | 1.90E-13 |
| 11 | 66032762 | C | T | 0.901349 | 4.85E-19 | 1.90E-13 |
| 11 | 66032775 | G | T | 0.901349 | 4.85E-19 | 1.90E-13 |
| 11 | 66032801 | T | C | 0.901349 | 4.85E-19 | 1.90E-13 |
| 11 | 66032802 | C | T | 0.901349 | 4.85E-19 | 1.90E-13 |
| 11 | 66032836 | G | C | 0.901349 | 4.85E-19 | 1.90E-13 |
| 11 | 66032851 | A | G | 0.901349 | 4.85E-19 | 1.90E-13 |
| 11 | 66032856 | C | T | 0.901349 | 4.85E-19 | 1.90E-13 |
| 11 | 66032872 | C | T | 0.901349 | 4.85E-19 | 1.90E-13 |
| 11 | 66032896 | A | T | 0.901349 | 4.85E-19 | 1.90E-13 |
| 11 | 66033000 | T | C | 0.901349 | 4.85E-19 | 1.90E-13 |
| 11 | 66033025 | T | C | 0.901349 | 4.85E-19 | 1.90E-13 |
| 11 | 66033053 | T | C | 0.901349 | 4.85E-19 | 1.90E-13 |
| 11 | 66033058 | T | A | 0.901349 | 4.85E-19 | 1.90E-13 |
| 11 | 66033140 | T | C | 0.901349 | 4.85E-19 | 1.90E-13 |
| 11 | 66033182 | G | A | 0.901349 | 4.85E-19 | 1.90E-13 |
| 11 | 66033215 | A | G | 0.901349 | 4.85E-19 | 1.90E-13 |
| 11 | 66033225 | A | T | 0.901349 | 4.85E-19 | 1.90E-13 |
| 11 | 66033251 | C | T | 0.901349 | 4.85E-19 | 1.90E-13 |
| 11 | 66033260 | G | A | 0.901349 | 4.85E-19 | 1.90E-13 |
| 11 | 66033283 | C | T | 0.901349 | 4.85E-19 | 1.90E-13 |
| 11 | 66033293 | C | T | 0.901349 | 4.85E-19 | 1.90E-13 |
| 11 | 66033315 | A | T | 0.901349 | 4.85E-19 | 1.90E-13 |
| 11 | 66033321 | T | C | 0.901349 | 4.85E-19 | 1.90E-13 |
| 11 | 66033373 | A | G | 0.901349 | 4.85E-19 | 1.90E-13 |
| 11 | 66033394 | T | G | 0.901349 | 4.85E-19 | 1.90E-13 |
| 11 | 66033409 | A | T | 0.901349 | 4.85E-19 | 1.90E-13 |
| 11 | 66033438 | A | G | 0.901349 | 4.85E-19 | 1.90E-13 |
| 11 | 66033440 | A | T | 0.901349 | 4.85E-19 | 1.90E-13 |
| 11 | 66033448 | G | A | 0.901349 | 4.85E-19 | 1.90E-13 |
| 11 | 66033464 | C | T | 0.901349 | 4.85E-19 | 1.90E-13 |
| 11 | 66033589 | T | C | 0.901349 | 4.85E-19 | 1.90E-13 |
| 11 | 66033590 | G | A | 0.901349 | 4.85E-19 | 1.90E-13 |
| 11 | 66033595 | T | C | 0.901349 | 4.85E-19 | 1.90E-13 |
| 11 | 66033601 | G | A | 0.901349 | 4.85E-19 | 1.90E-13 |
| 11 | 66033662 | T | A | 0.901349 | 4.85E-19 | 1.90E-13 |
| 11 | 66033696 | A | G | 0.901349 | 4.85E-19 | 1.90E-13 |
| 11 | 66033706 | T | C | 0.901349 | 4.85E-19 | 1.90E-13 |
| 11 | 66033739 | G | A | 0.901349 | 4.85E-19 | 1.90E-13 |
| 11 | 66033778 | T | C | 0.901349 | 4.85E-19 | 1.90E-13 |
| 11 | 66033800 | G | A | 0.901349 | 4.85E-19 | 1.90E-13 |
| 11 | 66033804 | G | A | 0.901349 | 4.85E-19 | 1.90E-13 |
| 11 | 66033809 | C | T | 0.901349 | 4.85E-19 | 1.90E-13 |
| 11 | 66033817 | T | C | 0.901349 | 4.85E-19 | 1.90E-13 |
| 11 | 66033848 | A | C | 0.901349 | 4.85E-19 | 1.90E-13 |
| 11 | 66033891 | G | A | 0.901349 | 4.85E-19 | 1.90E-13 |
| 11 | 66033897 | G | T | 0.901349 | 4.85E-19 | 1.90E-13 |
| 11 | 66033948 | A | G | 0.901349 | 4.85E-19 | 1.90E-13 |
| 11 | 66033953 | T | G | 0.901349 | 4.85E-19 | 1.90E-13 |
| 11 | 66034030 | A | G | 0.901349 | 4.85E-19 | 1.90E-13 |
| 11 | 66034146 | T | C | 0.901349 | 4.85E-19 | 1.90E-13 |
| 11 | 66034190 | A | G | 0.901349 | 4.85E-19 | 1.90E-13 |
| 11 | 66034242 | A | G | 0.901349 | 4.85E-19 | 1.90E-13 |
| 11 | 66034248 | C | T | 0.901349 | 4.85E-19 | 1.90E-13 |
| 11 | 66034252 | C | T | 0.901349 | 4.85E-19 | 1.90E-13 |
| 11 | 66034432 | C | T | 0.901349 | 4.85E-19 | 1.90E-13 |
| 11 | 66034457 | T | C | 0.901349 | 4.85E-19 | 1.90E-13 |
| 11 | 66034480 | A | T | 0.901349 | 4.85E-19 | 1.90E-13 |
| 11 | 66034519 | A | T | 0.901349 | 4.85E-19 | 1.90E-13 |
| 11 | 66034563 | T | C | 0.901349 | 4.85E-19 | 1.90E-13 |
| 11 | 66034606 | G | C | 0.901349 | 4.85E-19 | 1.90E-13 |
| 11 | 66034615 | C | T | 0.901349 | 4.85E-19 | 1.90E-13 |
| 11 | 66034625 | C | T | 0.901349 | 4.85E-19 | 1.90E-13 |

| | | | | | | |
|---|---|---|---|---|---|---|
| 11 | 66034753 | C | T | 0.901349 | 4.85E-19 | 1.90E-13 |
| 11 | 66034779 | T | G | 0.901349 | 4.85E-19 | 1.90E-13 |
| 11 | 66034963 | C | T | 0.901349 | 4.85E-19 | 1.90E-13 |
| 11 | 66035039 | C | T | 0.901349 | 4.85E-19 | 1.90E-13 |
| 11 | 66035101 | A | G | 0.901349 | 4.85E-19 | 1.90E-13 |
| 11 | 66035132 | G | A | 0.901349 | 4.85E-19 | 1.90E-13 |
| 11 | 66035202 | G | T | 0.901349 | 4.85E-19 | 1.90E-13 |
| 11 | 66035274 | G | T | 0.901349 | 4.85E-19 | 1.90E-13 |
| 11 | 66035283 | G | A | 0.901349 | 4.85E-19 | 1.90E-13 |
| 11 | 66035428 | A | G | 0.901349 | 4.85E-19 | 1.90E-13 |
| 11 | 66035476 | C | T | 0.901349 | 4.85E-19 | 1.90E-13 |
| 11 | 66035489 | A | G | 0.901349 | 4.85E-19 | 1.90E-13 |
| 11 | 66035496 | G | A | 0.901349 | 4.85E-19 | 1.90E-13 |
| 11 | 66035540 | G | C | 0.901349 | 4.85E-19 | 1.90E-13 |
| 11 | 66035551 | A | T | 0.901349 | 4.85E-19 | 1.90E-13 |
| 11 | 66035605 | G | A | 0.901349 | 4.85E-19 | 1.90E-13 |
| 11 | 66035610 | T | C | 0.901349 | 4.85E-19 | 1.90E-13 |
| 11 | 66035654 | T | C | 0.901349 | 4.85E-19 | 1.90E-13 |
| 11 | 66035661 | A | G | 0.901349 | 4.85E-19 | 1.90E-13 |
| 11 | 66035671 | T | C | 0.901349 | 4.85E-19 | 1.90E-13 |
| 11 | 66035692 | A | G | 0.901349 | 4.85E-19 | 1.90E-13 |
| 11 | 66035731 | A | G | 0.901349 | 4.85E-19 | 1.90E-13 |
| 11 | 66035748 | T | C | 0.901349 | 4.85E-19 | 1.90E-13 |
| 11 | 66035776 | C | A | 0.901349 | 4.85E-19 | 1.90E-13 |
| 11 | 66035845 | T | C | 0.901349 | 4.85E-19 | 1.90E-13 |
| 11 | 66036032 | G | T | 0.901349 | 4.85E-19 | 1.90E-13 |
| 11 | 66036038 | A | G | 0.901349 | 4.85E-19 | 1.90E-13 |
| 11 | 66036046 | A | G | 0.901349 | 4.85E-19 | 1.90E-13 |
| 11 | 66036049 | C | T | 0.901349 | 4.85E-19 | 1.90E-13 |
| 11 | 66036075 | G | A | 0.901349 | 4.85E-19 | 1.90E-13 |
| 11 | 66036080 | G | C | 0.901349 | 4.85E-19 | 1.90E-13 |
| 11 | 66036131 | C | T | 0.901349 | 4.85E-19 | 1.90E-13 |
| 11 | 66036202 | A | G | 0.901349 | 4.85E-19 | 1.90E-13 |
| 11 | 66036207 | T | G | 0.901349 | 4.85E-19 | 1.90E-13 |
| 11 | 66036219 | A | G | 0.901349 | 4.85E-19 | 1.90E-13 |
| 11 | 66036282 | A | G | 0.901349 | 4.85E-19 | 1.90E-13 |
| 11 | 66036327 | A | T | 0.901349 | 4.85E-19 | 1.90E-13 |
| 11 | 66036334 | T | C | 0.901349 | 4.85E-19 | 1.90E-13 |
| 11 | 66036457 | G | A | 0.901349 | 4.85E-19 | 1.90E-13 |
| 11 | 66036507 | C | T | 0.901349 | 4.85E-19 | 1.90E-13 |
| 11 | 66036551 | C | T | 0.901349 | 4.85E-19 | 1.90E-13 |
| 11 | 66036552 | G | A | 0.901349 | 4.85E-19 | 1.90E-13 |
| 11 | 66036583 | T | A | 0.901349 | 4.85E-19 | 1.90E-13 |
| 11 | 66036585 | T | A | 0.901349 | 4.85E-19 | 1.90E-13 |
| 11 | 66036642 | A | G | 0.901349 | 4.85E-19 | 1.90E-13 |
| 11 | 66036678 | G | T | 0.901349 | 4.85E-19 | 1.90E-13 |
| 11 | 66036690 | T | C | 0.901349 | 4.85E-19 | 1.90E-13 |
| 11 | 66036691 | G | A | 0.901349 | 4.85E-19 | 1.90E-13 |
| 11 | 66036699 | T | G | 0.901349 | 4.85E-19 | 1.90E-13 |
| 11 | 66036700 | T | A | 0.901349 | 4.85E-19 | 1.90E-13 |
| 11 | 66036789 | A | G | 0.901349 | 4.85E-19 | 1.90E-13 |
| 11 | 66036821 | C | G | 0.901349 | 4.85E-19 | 1.90E-13 |
| 11 | 66036832 | T | G | 0.901349 | 4.85E-19 | 1.90E-13 |
| 11 | 66036863 | A | G | 0.901349 | 4.85E-19 | 1.90E-13 |
| 11 | 66037001 | T | A | 0.901349 | 4.85E-19 | 1.90E-13 |
| 11 | 66037133 | A | C | 0.901349 | 4.85E-19 | 1.90E-13 |
| 11 | 66037207 | G | A | 0.901349 | 4.85E-19 | 1.90E-13 |
| 11 | 66037209 | G | A | 0.901349 | 4.85E-19 | 1.90E-13 |
| 11 | 66037227 | G | A | 0.901349 | 4.85E-19 | 1.90E-13 |
| 11 | 66037322 | T | C | 0.901349 | 4.85E-19 | 1.90E-13 |
| 11 | 66037335 | G | C | 0.901349 | 4.85E-19 | 1.90E-13 |
| 11 | 66037407 | C | T | 0.901349 | 4.85E-19 | 1.90E-13 |
| 11 | 66037428 | T | G | 0.901349 | 4.85E-19 | 1.90E-13 |
| 11 | 66037470 | A | C | 0.901349 | 4.85E-19 | 1.90E-13 |
| 11 | 66037472 | G | A | 0.901349 | 4.85E-19 | 1.90E-13 |
| 11 | 66037475 | G | A | 0.901349 | 4.85E-19 | 1.90E-13 |
| 11 | 66037496 | T | C | 0.901349 | 4.85E-19 | 1.90E-13 |

| | | | | | | |
|---|---|---|---|---|---|---|
| 11 | 66037517 | A | G | 0.901349 | 4.85E-19 | 1.90E-13 |
| 11 | 66037518 | C | A | 0.901349 | 4.85E-19 | 1.90E-13 |
| 11 | 66037595 | G | T | 0.901349 | 4.85E-19 | 1.90E-13 |
| 11 | 66037613 | C | A | 0.901349 | 4.85E-19 | 1.90E-13 |
| 11 | 66037666 | T | C | 0.901349 | 4.85E-19 | 1.90E-13 |
| 11 | 66037668 | C | A | 0.901349 | 4.85E-19 | 1.90E-13 |
| 11 | 66037681 | C | T | 0.901349 | 4.85E-19 | 1.90E-13 |
| 11 | 66037683 | T | C | 0.901349 | 4.85E-19 | 1.90E-13 |
| 11 | 66037705 | T | C | 0.901349 | 4.85E-19 | 1.90E-13 |
| 11 | 66037811 | T | C | 0.901349 | 4.85E-19 | 1.90E-13 |
| 11 | 66037813 | G | C | 0.901349 | 4.85E-19 | 1.90E-13 |
| 11 | 66037815 | C | A | 0.901349 | 4.85E-19 | 1.90E-13 |
| 11 | 66037816 | T | C | 0.901349 | 4.85E-19 | 1.90E-13 |
| 11 | 66037845 | C | T | 0.901349 | 4.85E-19 | 1.90E-13 |
| 11 | 66037912 | C | T | 0.901349 | 4.85E-19 | 1.90E-13 |
| 11 | 66037940 | T | C | 0.901349 | 4.85E-19 | 1.90E-13 |
| 11 | 66037955 | A | G | 0.901349 | 4.85E-19 | 1.90E-13 |
| 11 | 66038039 | C | G | 0.901349 | 4.85E-19 | 1.90E-13 |
| 11 | 66038070 | T | A | 0.901349 | 4.85E-19 | 1.90E-13 |
| 11 | 66038137 | T | A | 0.901349 | 4.85E-19 | 1.90E-13 |
| 11 | 66038197 | T | C | 0.901349 | 4.85E-19 | 1.90E-13 |
| 11 | 66038246 | A | G | 0.901349 | 4.85E-19 | 1.90E-13 |
| 11 | 66038252 | A | C | 0.901349 | 4.85E-19 | 1.90E-13 |
| 11 | 66038275 | C | T | 0.901349 | 4.85E-19 | 1.90E-13 |
| 11 | 66038401 | A | G | 0.901349 | 4.85E-19 | 1.90E-13 |
| 11 | 66038406 | A | G | 0.901349 | 4.85E-19 | 1.90E-13 |
| 11 | 66038639 | A | G | 0.901349 | 4.85E-19 | 1.90E-13 |
| 11 | 66039052 | G | A | 0.901349 | 4.85E-19 | 1.90E-13 |
| 11 | 66039085 | G | A | 0.901349 | 4.85E-19 | 1.90E-13 |
| 11 | 66039172 | A | G | 0.901349 | 4.85E-19 | 1.90E-13 |
| 11 | 66039179 | C | T | 0.901349 | 4.85E-19 | 1.90E-13 |
| 11 | 66039188 | G | A | 0.901349 | 4.85E-19 | 1.90E-13 |
| 11 | 66039194 | A | G | 0.901349 | 4.85E-19 | 1.90E-13 |
| 11 | 66039251 | T | C | 0.901349 | 4.85E-19 | 1.90E-13 |
| 11 | 66039376 | C | T | 0.901349 | 4.85E-19 | 1.90E-13 |
| 11 | 66039445 | T | A | 0.901349 | 4.85E-19 | 1.90E-13 |
| 11 | 66039471 | C | T | 0.901349 | 4.85E-19 | 1.90E-13 |
| 11 | 66039560 | T | C | 0.901349 | 4.85E-19 | 1.90E-13 |
| 11 | 66039635 | G | A | 0.901349 | 4.85E-19 | 1.90E-13 |
| 11 | 66039645 | C | T | 0.901349 | 4.85E-19 | 1.90E-13 |
| 11 | 66039765 | G | T | 0.901349 | 4.85E-19 | 1.90E-13 |
| 11 | 66039785 | G | A | 0.901349 | 4.85E-19 | 1.90E-13 |
| 11 | 66039841 | G | A | 0.901349 | 4.85E-19 | 1.90E-13 |
| 11 | 66039876 | A | G | 0.901349 | 4.85E-19 | 1.90E-13 |
| 11 | 66039886 | G | A | 0.901349 | 4.85E-19 | 1.90E-13 |
| 11 | 66039899 | G | A | 0.901349 | 4.85E-19 | 1.90E-13 |
| 11 | 66039976 | C | T | 0.901349 | 4.85E-19 | 1.90E-13 |
| 11 | 66040075 | T | C | 0.901349 | 4.85E-19 | 1.90E-13 |
| 11 | 66040083 | A | T | 0.901349 | 4.85E-19 | 1.90E-13 |
| 11 | 66040150 | C | G | 0.901349 | 4.85E-19 | 1.90E-13 |
| 11 | 66040217 | A | G | 0.901349 | 4.85E-19 | 1.90E-13 |
| 11 | 66040218 | T | C | 0.901349 | 4.85E-19 | 1.90E-13 |
| 11 | 66040227 | T | G | 0.901349 | 4.85E-19 | 1.90E-13 |
| 11 | 66040238 | C | A | 0.901349 | 4.85E-19 | 1.90E-13 |
| 11 | 66040251 | C | T | 0.901349 | 4.85E-19 | 1.90E-13 |
| 11 | 66040306 | A | G | 0.901349 | 4.85E-19 | 1.90E-13 |
| 11 | 66040309 | C | T | 0.901349 | 4.85E-19 | 1.90E-13 |
| 11 | 66040310 | T | C | 0.901349 | 4.85E-19 | 1.90E-13 |
| 11 | 66040315 | G | A | 0.901349 | 4.85E-19 | 1.90E-13 |
| 11 | 66040320 | G | A | 0.901349 | 4.85E-19 | 1.90E-13 |
| 11 | 66040323 | G | A | 0.901349 | 4.85E-19 | 1.90E-13 |
| 11 | 66040366 | A | G | 0.901349 | 4.85E-19 | 1.90E-13 |
| 11 | 66040417 | A | G | 0.901349 | 4.85E-19 | 1.90E-13 |
| 11 | 66040466 | G | A | 0.901349 | 4.85E-19 | 1.90E-13 |
| 11 | 66040470 | A | G | 0.901349 | 4.85E-19 | 1.90E-13 |
| 11 | 66040499 | C | T | 0.901349 | 4.85E-19 | 1.90E-13 |
| 11 | 66040512 | G | T | 0.901349 | 4.85E-19 | 1.90E-13 |

| | | | | | | |
|---|---|---|---|---|---|---|
| 11 | 66040585 | A | T | 0.901349 | 4.85E-19 | 1.90E-13 |
| 11 | 66040612 | A | C | 0.901349 | 4.85E-19 | 1.90E-13 |
| 11 | 66040748 | A | G | 0.901349 | 4.85E-19 | 1.90E-13 |
| 11 | 66040798 | G | C | 0.901349 | 4.85E-19 | 1.90E-13 |
| 11 | 66041249 | T | C | 0.901349 | 4.85E-19 | 1.90E-13 |
| 11 | 66041294 | C | G | 0.901349 | 4.85E-19 | 1.90E-13 |
| 11 | 66041296 | C | G | 0.901349 | 4.85E-19 | 1.90E-13 |
| 11 | 66041298 | C | G | 0.901349 | 4.85E-19 | 1.90E-13 |
| 11 | 66041300 | C | G | 0.901349 | 4.85E-19 | 1.90E-13 |
| 11 | 66041302 | G | A | 0.901349 | 4.85E-19 | 1.90E-13 |
| 11 | 66041401 | A | G | 0.901349 | 4.85E-19 | 1.90E-13 |
| 11 | 66041421 | A | C | 0.901349 | 4.85E-19 | 1.90E-13 |
| 11 | 66041481 | C | T | 0.901349 | 4.85E-19 | 1.90E-13 |
| 11 | 66041584 | A | T | 0.901349 | 4.85E-19 | 1.90E-13 |
| 11 | 66041606 | G | A | 0.901349 | 4.85E-19 | 1.90E-13 |
| 11 | 66041628 | G | A | 0.901349 | 4.85E-19 | 1.90E-13 |
| 11 | 66041676 | A | G | 0.901349 | 4.85E-19 | 1.90E-13 |
| 11 | 66041848 | C | T | 0.901349 | 4.85E-19 | 1.90E-13 |
| 11 | 66042135 | C | T | 0.901349 | 4.85E-19 | 1.90E-13 |
| 11 | 66042176 | C | A | 0.901349 | 4.85E-19 | 1.90E-13 |
| 11 | 66042192 | A | G | 0.901349 | 4.85E-19 | 1.90E-13 |
| 11 | 66042220 | A | G | 0.901349 | 4.85E-19 | 1.90E-13 |
| 11 | 66042360 | T | G | 0.901349 | 4.85E-19 | 1.90E-13 |
| 11 | 66042380 | A | G | 0.901349 | 4.85E-19 | 1.90E-13 |
| 11 | 66042436 | T | C | 0.901349 | 4.85E-19 | 1.90E-13 |
| 11 | 66042595 | C | T | 0.901349 | 4.85E-19 | 1.90E-13 |
| 11 | 66042607 | T | C | 0.901349 | 4.85E-19 | 1.90E-13 |
| 11 | 66042774 | G | C | 0.901349 | 4.85E-19 | 1.90E-13 |
| 11 | 66042787 | A | T | 0.901349 | 4.85E-19 | 1.90E-13 |
| 11 | 66042812 | A | G | 0.901349 | 4.85E-19 | 1.90E-13 |
| 11 | 66042857 | C | A | 0.901349 | 4.85E-19 | 1.90E-13 |
| 11 | 66042954 | T | A | 0.901349 | 4.85E-19 | 1.90E-13 |
| 11 | 66043012 | C | T | 0.901349 | 4.85E-19 | 1.90E-13 |
| 11 | 66043032 | T | G | 0.901349 | 4.85E-19 | 1.90E-13 |
| 11 | 66043069 | C | T | 0.901349 | 4.85E-19 | 1.90E-13 |
| 11 | 66043075 | T | C | 0.901349 | 4.85E-19 | 1.90E-13 |
| 11 | 66043078 | T | G | 0.901349 | 4.85E-19 | 1.90E-13 |
| 11 | 66043099 | A | G | 0.901349 | 4.85E-19 | 1.90E-13 |
| 11 | 66043103 | G | A | 0.901349 | 4.85E-19 | 1.90E-13 |
| 11 | 66043105 | T | C | 0.901349 | 4.85E-19 | 1.90E-13 |
| 11 | 66043109 | A | T | 0.901349 | 4.85E-19 | 1.90E-13 |
| 11 | 66043193 | T | C | 0.901349 | 4.85E-19 | 1.90E-13 |
| 11 | 66043250 | C | T | 0.901349 | 4.85E-19 | 1.90E-13 |
| 11 | 66043255 | G | A | 0.901349 | 4.85E-19 | 1.90E-13 |
| 11 | 66043335 | C | T | 0.901349 | 4.85E-19 | 1.90E-13 |
| 11 | 66043336 | T | G | 0.901349 | 4.85E-19 | 1.90E-13 |
| 11 | 66043345 | T | G | 0.901349 | 4.85E-19 | 1.90E-13 |
| 11 | 66043374 | T | A | 0.901349 | 4.85E-19 | 1.90E-13 |
| 11 | 66043497 | C | T | 0.901349 | 4.85E-19 | 1.90E-13 |
| 11 | 66043532 | A | T | 0.901349 | 4.85E-19 | 1.90E-13 |
| 11 | 66043628 | C | G | 0.901349 | 4.85E-19 | 1.90E-13 |
| 11 | 66043660 | A | T | 0.901349 | 4.85E-19 | 1.90E-13 |
| 11 | 66043760 | A | T | 0.901349 | 4.85E-19 | 1.90E-13 |
| 11 | 66043826 | T | G | 0.901349 | 4.85E-19 | 1.90E-13 |
| 11 | 66043852 | T | C | 0.901349 | 4.85E-19 | 1.90E-13 |
| 11 | 66043927 | C | T | 0.901349 | 4.85E-19 | 1.90E-13 |
| 11 | 66043931 | T | C | 0.901349 | 4.85E-19 | 1.90E-13 |
| 11 | 66043939 | G | C | 0.901349 | 4.85E-19 | 1.90E-13 |
| 11 | 66043983 | T | C | 0.901349 | 4.85E-19 | 1.90E-13 |
| 11 | 66044013 | A | G | 0.901349 | 4.85E-19 | 1.90E-13 |
| 11 | 66044034 | A | G | 0.901349 | 4.85E-19 | 1.90E-13 |
| 11 | 66044098 | T | A | 0.901349 | 4.85E-19 | 1.90E-13 |
| 11 | 66044102 | T | A | 0.901349 | 4.85E-19 | 1.90E-13 |
| 11 | 66044159 | G | A | 0.901349 | 4.85E-19 | 1.90E-13 |
| 11 | 66044350 | G | A | 0.901349 | 4.85E-19 | 1.90E-13 |
| 11 | 66044403 | C | G | 0.901349 | 4.85E-19 | 1.90E-13 |
| 11 | 66044437 | T | G | 0.901349 | 4.85E-19 | 1.90E-13 |

| | | | | | | |
|---|---|---|---|---|---|---|
| 11 | 66044526 | A | G | 0.901349 | 4.85E-19 | 1.90E-13 |
| 11 | 66044586 | C | T | 0.901349 | 4.85E-19 | 1.90E-13 |
| 11 | 66044603 | C | T | 0.901349 | 4.85E-19 | 1.90E-13 |
| 11 | 66044823 | T | G | 0.901349 | 4.85E-19 | 1.90E-13 |
| 11 | 66044888 | G | T | 0.901349 | 4.85E-19 | 1.90E-13 |
| 11 | 66045164 | T | C | 0.901349 | 4.85E-19 | 1.90E-13 |
| 11 | 66045312 | A | G | 0.901349 | 4.85E-19 | 1.90E-13 |
| 11 | 66045313 | A | C | 0.901349 | 4.85E-19 | 1.90E-13 |
| 11 | 66045337 | A | G | 0.901349 | 4.85E-19 | 1.90E-13 |
| 11 | 66045379 | T | C | 0.901349 | 4.85E-19 | 1.90E-13 |
| 11 | 66045537 | C | T | 0.901349 | 4.85E-19 | 1.90E-13 |
| 11 | 66045547 | C | T | 0.901349 | 4.85E-19 | 1.90E-13 |
| 11 | 66045789 | A | T | 0.901349 | 4.85E-19 | 1.90E-13 |
| 11 | 66045837 | C | T | 0.901349 | 4.85E-19 | 1.90E-13 |
| 11 | 66045979 | T | C | 0.901349 | 4.85E-19 | 1.90E-13 |
| 11 | 66045985 | T | C | 0.901349 | 4.85E-19 | 1.90E-13 |
| 11 | 66046052 | A | G | 0.901349 | 4.85E-19 | 1.90E-13 |
| 11 | 66046133 | G | A | 0.901349 | 4.85E-19 | 1.90E-13 |
| 11 | 66046172 | C | T | 0.901349 | 4.85E-19 | 1.90E-13 |
| 11 | 66046224 | A | G | 0.901349 | 4.85E-19 | 1.90E-13 |
| 11 | 66046256 | T | G | 0.901349 | 4.85E-19 | 1.90E-13 |
| 11 | 66046403 | C | T | 0.901349 | 4.85E-19 | 1.90E-13 |
| 11 | 66046421 | C | T | 0.901349 | 4.85E-19 | 1.90E-13 |
| 11 | 66046468 | G | A | 0.901349 | 4.85E-19 | 1.90E-13 |
| 11 | 66046489 | T | C | 0.901349 | 4.85E-19 | 1.90E-13 |
| 11 | 66046525 | G | A | 0.901349 | 4.85E-19 | 1.90E-13 |
| 11 | 66046586 | C | A | 0.901349 | 4.85E-19 | 1.90E-13 |
| 11 | 66046822 | A | G | 0.901349 | 4.85E-19 | 1.90E-13 |
| 11 | 66046836 | T | C | 0.901349 | 4.85E-19 | 1.90E-13 |
| 11 | 66046887 | A | T | 0.901349 | 4.85E-19 | 1.90E-13 |
| 11 | 66046936 | A | G | 0.901349 | 4.85E-19 | 1.90E-13 |
| 11 | 66047082 | A | C | 0.901349 | 4.85E-19 | 1.90E-13 |
| 11 | 66047241 | C | T | 0.901349 | 4.85E-19 | 1.90E-13 |
| 11 | 66047252 | C | G | 0.901349 | 4.85E-19 | 1.90E-13 |
| 11 | 66047351 | G | A | 0.901349 | 4.85E-19 | 1.90E-13 |
| 11 | 66047446 | C | A | 0.901349 | 4.85E-19 | 1.90E-13 |
| 11 | 66047541 | C | T | 0.901349 | 4.85E-19 | 1.90E-13 |
| 11 | 66047678 | G | A | 0.901349 | 4.85E-19 | 1.90E-13 |
| 11 | 66047679 | G | A | 0.901349 | 4.85E-19 | 1.90E-13 |
| 11 | 66047952 | T | G | 0.901349 | 4.85E-19 | 1.90E-13 |
| 11 | 66048126 | A | T | 0.901349 | 4.85E-19 | 1.90E-13 |
| 11 | 66048143 | G | T | 0.901349 | 4.85E-19 | 1.90E-13 |
| 11 | 66048192 | A | G | 0.901349 | 4.85E-19 | 1.90E-13 |
| 11 | 66048213 | G | A | 0.901349 | 4.85E-19 | 1.90E-13 |
| 11 | 66048237 | T | C | 0.901349 | 4.85E-19 | 1.90E-13 |
| 11 | 66048321 | T | C | 0.901349 | 4.85E-19 | 1.90E-13 |
| 11 | 66048335 | T | A | 0.901349 | 4.85E-19 | 1.90E-13 |
| 11 | 66048352 | C | A | 0.901349 | 4.85E-19 | 1.90E-13 |
| 11 | 66048381 | G | C | 0.901349 | 4.85E-19 | 1.90E-13 |
| 11 | 66048618 | T | C | 0.901349 | 4.85E-19 | 1.90E-13 |
| 11 | 66048835 | T | A | 0.901349 | 4.85E-19 | 1.90E-13 |
| 11 | 66048938 | A | G | 0.901349 | 4.85E-19 | 1.90E-13 |
| 11 | 66049016 | G | A | 0.901349 | 4.85E-19 | 1.90E-13 |
| 11 | 66049074 | G | A | 0.901349 | 4.85E-19 | 1.90E-13 |
| 11 | 66049183 | G | A | 0.901349 | 4.85E-19 | 1.90E-13 |
| 11 | 66031459 | T | C | 0.897976 | 5.07E-19 | 1.99E-13 |
| 11 | 66003717 | C | G | 0.900506 | 1.32E-18 | 5.18E-13 |
| 11 | 66003725 | C | T | 0.900506 | 1.32E-18 | 5.18E-13 |
| 11 | 66003893 | T | C | 0.900506 | 1.32E-18 | 5.18E-13 |
| 11 | 66003950 | G | A | 0.900506 | 1.32E-18 | 5.18E-13 |
| 11 | 66018028 | A | C | 0.902192 | 1.60E-18 | 6.27E-13 |
| 11 | 63283612 | G | C | 0.881113 | 2.40E-18 | 9.43E-13 |
| 11 | 63302242 | A | G | 0.881113 | 2.99E-18 | 1.17E-12 |
| 11 | 65398863 | G | A | 0.924958 | 3.04E-18 | 1.19E-12 |
| 11 | 65399615 | T | C | 0.924958 | 3.04E-18 | 1.19E-12 |
| 11 | 65400434 | T | C | 0.924958 | 3.04E-18 | 1.19E-12 |
| 11 | 65400759 | T | C | 0.924958 | 3.04E-18 | 1.19E-12 |

| | | | | | | |
|---|---|---|---|---|---|---|
| 11 | 65402208 | G | A | 0.924958 | 3.04E-18 | 1.19E-12 |
| 11 | 65402424 | T | C | 0.924958 | 3.04E-18 | 1.19E-12 |
| 11 | 65403179 | C | T | 0.924958 | 3.04E-18 | 1.19E-12 |
| 11 | 65403383 | T | G | 0.924958 | 3.04E-18 | 1.19E-12 |
| 11 | 65404633 | A | G | 0.924958 | 3.04E-18 | 1.19E-12 |
| 11 | 65404823 | G | C | 0.924958 | 3.04E-18 | 1.19E-12 |
| 11 | 65404929 | G | A | 0.924958 | 3.04E-18 | 1.19E-12 |
| 11 | 65404980 | C | T | 0.924958 | 3.04E-18 | 1.19E-12 |
| 11 | 65405087 | A | G | 0.924958 | 3.04E-18 | 1.19E-12 |
| 11 | 65405090 | A | G | 0.924958 | 3.04E-18 | 1.19E-12 |
| 11 | 65405254 | A | C | 0.924958 | 3.04E-18 | 1.19E-12 |
| 11 | 65405369 | A | G | 0.924958 | 3.04E-18 | 1.19E-12 |
| 11 | 65405580 | C | A | 0.924958 | 3.04E-18 | 1.19E-12 |
| 11 | 65405683 | C | T | 0.924958 | 3.04E-18 | 1.19E-12 |
| 11 | 65405687 | C | A | 0.924958 | 3.04E-18 | 1.19E-12 |
| 11 | 65405734 | C | T | 0.924958 | 3.04E-18 | 1.19E-12 |
| 11 | 65405940 | G | T | 0.924958 | 3.04E-18 | 1.19E-12 |
| 11 | 65405943 | G | A | 0.924958 | 3.04E-18 | 1.19E-12 |
| 11 | 65405944 | T | C | 0.924958 | 3.04E-18 | 1.19E-12 |
| 11 | 65405983 | T | C | 0.924958 | 3.04E-18 | 1.19E-12 |
| 11 | 65405987 | G | A | 0.924958 | 3.04E-18 | 1.19E-12 |
| 11 | 65405999 | C | T | 0.924958 | 3.04E-18 | 1.19E-12 |
| 11 | 65406047 | C | T | 0.924958 | 3.04E-18 | 1.19E-12 |
| 11 | 65406061 | A | C | 0.924958 | 3.04E-18 | 1.19E-12 |
| 11 | 65406104 | C | T | 0.924958 | 3.04E-18 | 1.19E-12 |
| 11 | 65406160 | G | C | 0.924958 | 3.04E-18 | 1.19E-12 |
| 11 | 65406419 | T | C | 0.924958 | 3.04E-18 | 1.19E-12 |
| 11 | 65406470 | A | G | 0.924958 | 3.04E-18 | 1.19E-12 |
| 11 | 65406667 | A | G | 0.924958 | 3.04E-18 | 1.19E-12 |
| 11 | 65406674 | T | C | 0.924958 | 3.04E-18 | 1.19E-12 |
| 11 | 65406699 | T | C | 0.924958 | 3.04E-18 | 1.19E-12 |
| 11 | 65407772 | A | G | 0.924958 | 3.04E-18 | 1.19E-12 |
| 11 | 65407808 | C | T | 0.924958 | 3.04E-18 | 1.19E-12 |
| 11 | 65407947 | C | T | 0.924958 | 3.04E-18 | 1.19E-12 |
| 11 | 65408288 | G | C | 0.924958 | 3.04E-18 | 1.19E-12 |
| 11 | 65408676 | A | G | 0.924958 | 3.04E-18 | 1.19E-12 |
| 11 | 65408874 | T | A | 0.924958 | 3.04E-18 | 1.19E-12 |
| 11 | 65408986 | C | T | 0.924958 | 3.04E-18 | 1.19E-12 |
| 11 | 65408989 | C | T | 0.924958 | 3.04E-18 | 1.19E-12 |
| 11 | 65409054 | G | A | 0.924958 | 3.04E-18 | 1.19E-12 |
| 11 | 65409059 | T | G | 0.924958 | 3.04E-18 | 1.19E-12 |
| 11 | 65409211 | C | T | 0.924958 | 3.04E-18 | 1.19E-12 |
| 11 | 65409471 | T | C | 0.924958 | 3.04E-18 | 1.19E-12 |
| 11 | 65409505 | T | G | 0.924958 | 3.04E-18 | 1.19E-12 |
| 11 | 65409672 | G | A | 0.924958 | 3.04E-18 | 1.19E-12 |
| 11 | 65410049 | A | G | 0.924958 | 3.04E-18 | 1.19E-12 |
| 11 | 65410310 | G | C | 0.924958 | 3.04E-18 | 1.19E-12 |
| 11 | 65410470 | G | A | 0.924958 | 3.04E-18 | 1.19E-12 |
| 11 | 65410524 | C | T | 0.924958 | 3.04E-18 | 1.19E-12 |
| 11 | 65410536 | A | T | 0.924958 | 3.04E-18 | 1.19E-12 |
| 11 | 65410609 | A | G | 0.924958 | 3.04E-18 | 1.19E-12 |
| 11 | 65410617 | T | G | 0.924958 | 3.04E-18 | 1.19E-12 |
| 11 | 65410629 | A | G | 0.924958 | 3.04E-18 | 1.19E-12 |
| 11 | 65410754 | C | A | 0.924958 | 3.04E-18 | 1.19E-12 |
| 11 | 65410830 | G | T | 0.924958 | 3.04E-18 | 1.19E-12 |
| 11 | 65410860 | C | T | 0.924958 | 3.04E-18 | 1.19E-12 |
| 11 | 65410986 | A | T | 0.924958 | 3.04E-18 | 1.19E-12 |
| 11 | 65410994 | C | T | 0.924958 | 3.04E-18 | 1.19E-12 |
| 11 | 65411384 | T | C | 0.924958 | 3.04E-18 | 1.19E-12 |
| 11 | 65411394 | T | C | 0.924958 | 3.04E-18 | 1.19E-12 |
| 11 | 65411456 | A | G | 0.924958 | 3.04E-18 | 1.19E-12 |
| 11 | 65411628 | A | C | 0.924958 | 3.04E-18 | 1.19E-12 |
| 11 | 65411638 | G | T | 0.924958 | 3.04E-18 | 1.19E-12 |
| 11 | 65411647 | A | T | 0.924958 | 3.04E-18 | 1.19E-12 |
| 11 | 65411654 | T | A | 0.924958 | 3.04E-18 | 1.19E-12 |
| 11 | 65411709 | T | A | 0.924958 | 3.04E-18 | 1.19E-12 |
| 11 | 65411720 | T | C | 0.924958 | 3.04E-18 | 1.19E-12 |

| | | | | | | |
|---|---|---|---|---|---|---|
| 11 | 65411745 | C | T | 0.924958 | 3.04E-18 | 1.19E-12 |
| 11 | 65411818 | T | A | 0.924958 | 3.04E-18 | 1.19E-12 |
| 11 | 65411944 | G | T | 0.924958 | 3.04E-18 | 1.19E-12 |
| 11 | 65411980 | G | A | 0.924958 | 3.04E-18 | 1.19E-12 |
| 11 | 65412034 | G | A | 0.924958 | 3.04E-18 | 1.19E-12 |
| 11 | 65412035 | G | A | 0.924958 | 3.04E-18 | 1.19E-12 |
| 11 | 65412043 | T | C | 0.924958 | 3.04E-18 | 1.19E-12 |
| 11 | 65412196 | C | G | 0.924958 | 3.04E-18 | 1.19E-12 |
| 11 | 65412242 | T | C | 0.924958 | 3.04E-18 | 1.19E-12 |
| 11 | 65412265 | C | T | 0.924958 | 3.04E-18 | 1.19E-12 |
| 11 | 65412267 | G | A | 0.924958 | 3.04E-18 | 1.19E-12 |
| 11 | 65412276 | C | T | 0.924958 | 3.04E-18 | 1.19E-12 |
| 11 | 65412279 | G | A | 0.924958 | 3.04E-18 | 1.19E-12 |
| 11 | 65412460 | T | C | 0.924958 | 3.04E-18 | 1.19E-12 |
| 11 | 65412513 | C | T | 0.924958 | 3.04E-18 | 1.19E-12 |
| 11 | 65412534 | T | C | 0.924958 | 3.04E-18 | 1.19E-12 |
| 11 | 65412598 | G | C | 0.924958 | 3.04E-18 | 1.19E-12 |
| 11 | 65412680 | C | T | 0.924958 | 3.04E-18 | 1.19E-12 |
| 11 | 65412714 | C | T | 0.924958 | 3.04E-18 | 1.19E-12 |
| 11 | 65412825 | A | G | 0.924958 | 3.04E-18 | 1.19E-12 |
| 11 | 65412984 | T | C | 0.924958 | 3.04E-18 | 1.19E-12 |
| 11 | 65413017 | C | A | 0.924958 | 3.04E-18 | 1.19E-12 |
| 11 | 65413099 | C | T | 0.924958 | 3.04E-18 | 1.19E-12 |
| 11 | 65413199 | C | T | 0.924958 | 3.04E-18 | 1.19E-12 |
| 11 | 65413345 | A | C | 0.924958 | 3.04E-18 | 1.19E-12 |
| 11 | 65413409 | G | T | 0.924958 | 3.04E-18 | 1.19E-12 |
| 11 | 65413452 | A | C | 0.924958 | 3.04E-18 | 1.19E-12 |
| 11 | 65413466 | C | T | 0.924958 | 3.04E-18 | 1.19E-12 |
| 11 | 65413467 | A | G | 0.924958 | 3.04E-18 | 1.19E-12 |
| 11 | 65413639 | G | A | 0.924958 | 3.04E-18 | 1.19E-12 |
| 11 | 65413658 | A | T | 0.924958 | 3.04E-18 | 1.19E-12 |
| 11 | 65413666 | T | C | 0.924958 | 3.04E-18 | 1.19E-12 |
| 11 | 65413677 | A | T | 0.924958 | 3.04E-18 | 1.19E-12 |
| 11 | 65413819 | A | T | 0.924958 | 3.04E-18 | 1.19E-12 |
| 11 | 65413851 | G | A | 0.924958 | 3.04E-18 | 1.19E-12 |
| 11 | 65413912 | T | A | 0.924958 | 3.04E-18 | 1.19E-12 |
| 11 | 65413962 | G | A | 0.924958 | 3.04E-18 | 1.19E-12 |
| 11 | 65414061 | C | T | 0.924958 | 3.04E-18 | 1.19E-12 |
| 11 | 65414128 | T | G | 0.924958 | 3.04E-18 | 1.19E-12 |
| 11 | 65414132 | C | T | 0.924958 | 3.04E-18 | 1.19E-12 |
| 11 | 65414161 | A | G | 0.924958 | 3.04E-18 | 1.19E-12 |
| 11 | 65414182 | C | T | 0.924958 | 3.04E-18 | 1.19E-12 |
| 11 | 65414209 | G | A | 0.924958 | 3.04E-18 | 1.19E-12 |
| 11 | 65414216 | A | C | 0.924958 | 3.04E-18 | 1.19E-12 |
| 11 | 65414220 | C | G | 0.924958 | 3.04E-18 | 1.19E-12 |
| 11 | 65414245 | A | G | 0.924958 | 3.04E-18 | 1.19E-12 |
| 11 | 65414302 | C | A | 0.924958 | 3.04E-18 | 1.19E-12 |
| 11 | 65414353 | A | G | 0.924958 | 3.04E-18 | 1.19E-12 |
| 11 | 65414381 | T | C | 0.924958 | 3.04E-18 | 1.19E-12 |
| 11 | 65414432 | T | C | 0.924958 | 3.04E-18 | 1.19E-12 |
| 11 | 65414552 | A | G | 0.924958 | 3.04E-18 | 1.19E-12 |
| 11 | 65414676 | G | A | 0.924958 | 3.04E-18 | 1.19E-12 |
| 11 | 65414694 | T | C | 0.924958 | 3.04E-18 | 1.19E-12 |
| 11 | 65414724 | T | C | 0.924958 | 3.04E-18 | 1.19E-12 |
| 11 | 65414731 | C | T | 0.924958 | 3.04E-18 | 1.19E-12 |
| 11 | 65414733 | C | G | 0.924958 | 3.04E-18 | 1.19E-12 |
| 11 | 65414745 | T | C | 0.924958 | 3.04E-18 | 1.19E-12 |
| 11 | 65414746 | G | A | 0.924958 | 3.04E-18 | 1.19E-12 |
| 11 | 65414758 | T | C | 0.924958 | 3.04E-18 | 1.19E-12 |
| 11 | 65414759 | G | A | 0.924958 | 3.04E-18 | 1.19E-12 |
| 11 | 65414768 | A | G | 0.924958 | 3.04E-18 | 1.19E-12 |
| 11 | 65414770 | T | C | 0.924958 | 3.04E-18 | 1.19E-12 |
| 11 | 65414771 | G | A | 0.924958 | 3.04E-18 | 1.19E-12 |
| 11 | 65414804 | T | G | 0.924958 | 3.04E-18 | 1.19E-12 |
| 11 | 65414820 | C | T | 0.924958 | 3.04E-18 | 1.19E-12 |
| 11 | 65414821 | A | G | 0.924958 | 3.04E-18 | 1.19E-12 |
| 11 | 65414950 | A | G | 0.924958 | 3.04E-18 | 1.19E-12 |

| | | | | | | |
|---|---|---|---|---|---|---|
| 11 | 65414962 | C | T | 0.924958 | 3.04E-18 | 1.19E-12 |
| 11 | 65414979 | G | T | 0.924958 | 3.04E-18 | 1.19E-12 |
| 11 | 65414995 | G | C | 0.924958 | 3.04E-18 | 1.19E-12 |
| 11 | 65415163 | A | T | 0.924958 | 3.04E-18 | 1.19E-12 |
| 11 | 65415317 | T | C | 0.924958 | 3.04E-18 | 1.19E-12 |
| 11 | 65415393 | T | A | 0.924958 | 3.04E-18 | 1.19E-12 |
| 11 | 65415832 | C | T | 0.924958 | 3.04E-18 | 1.19E-12 |
| 11 | 65416597 | A | C | 0.924958 | 3.04E-18 | 1.19E-12 |
| 11 | 65416605 | A | G | 0.924958 | 3.04E-18 | 1.19E-12 |
| 11 | 65416795 | G | C | 0.924958 | 3.04E-18 | 1.19E-12 |
| 11 | 65417095 | T | C | 0.924958 | 3.04E-18 | 1.19E-12 |
| 11 | 65417178 | G | A | 0.924958 | 3.04E-18 | 1.19E-12 |
| 11 | 65417201 | G | C | 0.924958 | 3.04E-18 | 1.19E-12 |
| 11 | 65417275 | A | G | 0.924958 | 3.04E-18 | 1.19E-12 |
| 11 | 65417292 | A | T | 0.924958 | 3.04E-18 | 1.19E-12 |
| 11 | 65417410 | G | A | 0.924958 | 3.04E-18 | 1.19E-12 |
| 11 | 65417426 | T | C | 0.924958 | 3.04E-18 | 1.19E-12 |
| 11 | 65417527 | G | C | 0.924958 | 3.04E-18 | 1.19E-12 |
| 11 | 65417538 | C | T | 0.924958 | 3.04E-18 | 1.19E-12 |
| 11 | 65417607 | C | T | 0.924958 | 3.04E-18 | 1.19E-12 |
| 11 | 65417723 | C | A | 0.924958 | 3.04E-18 | 1.19E-12 |
| 11 | 65432659 | C | A | 0.924958 | 3.04E-18 | 1.19E-12 |
| 11 | 64901467 | T | C | 0.900506 | 3.39E-18 | 1.33E-12 |
| 11 | 64903173 | C | T | 0.900506 | 3.39E-18 | 1.33E-12 |
| 11 | 64906329 | A | G | 0.900506 | 3.39E-18 | 1.33E-12 |
| 11 | 64907134 | T | A | 0.900506 | 3.39E-18 | 1.33E-12 |
| 11 | 64909239 | A | G | 0.900506 | 3.39E-18 | 1.33E-12 |
| 11 | 64910342 | G | C | 0.900506 | 3.39E-18 | 1.33E-12 |
| 11 | 64910472 | T | A | 0.900506 | 3.39E-18 | 1.33E-12 |
| 11 | 64910593 | G | C | 0.900506 | 3.39E-18 | 1.33E-12 |
| 11 | 64910823 | A | T | 0.900506 | 3.39E-18 | 1.33E-12 |
| 11 | 64911175 | A | G | 0.900506 | 3.39E-18 | 1.33E-12 |
| 11 | 64911316 | G | A | 0.900506 | 3.39E-18 | 1.33E-12 |
| 11 | 64911479 | G | C | 0.900506 | 3.39E-18 | 1.33E-12 |
| 11 | 64911494 | A | C | 0.900506 | 3.39E-18 | 1.33E-12 |
| 11 | 64911591 | C | G | 0.900506 | 3.39E-18 | 1.33E-12 |
| 11 | 64911700 | A | G | 0.900506 | 3.39E-18 | 1.33E-12 |
| 11 | 64911719 | G | T | 0.900506 | 3.39E-18 | 1.33E-12 |
| 11 | 64911918 | A | G | 0.900506 | 3.39E-18 | 1.33E-12 |
| 11 | 64911936 | T | G | 0.900506 | 3.39E-18 | 1.33E-12 |
| 11 | 64911964 | G | A | 0.900506 | 3.39E-18 | 1.33E-12 |
| 11 | 64911971 | C | A | 0.900506 | 3.39E-18 | 1.33E-12 |
| 11 | 64912092 | T | C | 0.900506 | 3.39E-18 | 1.33E-12 |
| 11 | 64912093 | G | A | 0.900506 | 3.39E-18 | 1.33E-12 |
| 11 | 64912310 | A | G | 0.900506 | 3.39E-18 | 1.33E-12 |
| 11 | 64912467 | C | T | 0.900506 | 3.39E-18 | 1.33E-12 |
| 11 | 64912498 | T | G | 0.900506 | 3.39E-18 | 1.33E-12 |
| 11 | 64912499 | G | C | 0.900506 | 3.39E-18 | 1.33E-12 |
| 11 | 64912652 | T | C | 0.900506 | 3.39E-18 | 1.33E-12 |
| 11 | 64912876 | T | G | 0.900506 | 3.39E-18 | 1.33E-12 |
| 11 | 64912888 | T | A | 0.900506 | 3.39E-18 | 1.33E-12 |
| 11 | 64913186 | A | G | 0.900506 | 3.39E-18 | 1.33E-12 |
| 11 | 64913262 | G | T | 0.900506 | 3.39E-18 | 1.33E-12 |
| 11 | 64913316 | A | G | 0.900506 | 3.39E-18 | 1.33E-12 |
| 11 | 64913376 | T | C | 0.900506 | 3.39E-18 | 1.33E-12 |
| 11 | 64913458 | A | G | 0.900506 | 3.39E-18 | 1.33E-12 |
| 11 | 64913619 | C | T | 0.900506 | 3.39E-18 | 1.33E-12 |
| 11 | 64913778 | G | A | 0.900506 | 3.39E-18 | 1.33E-12 |
| 11 | 64913806 | T | C | 0.900506 | 3.39E-18 | 1.33E-12 |
| 11 | 64913855 | C | T | 0.900506 | 3.39E-18 | 1.33E-12 |
| 11 | 64913878 | A | G | 0.900506 | 3.39E-18 | 1.33E-12 |
| 11 | 64913893 | A | G | 0.900506 | 3.39E-18 | 1.33E-12 |
| 11 | 64913911 | A | C | 0.900506 | 3.39E-18 | 1.33E-12 |
| 11 | 64913912 | T | A | 0.900506 | 3.39E-18 | 1.33E-12 |
| 11 | 64914108 | G | T | 0.900506 | 3.39E-18 | 1.33E-12 |
| 11 | 64914152 | T | C | 0.900506 | 3.39E-18 | 1.33E-12 |
| 11 | 64914159 | A | T | 0.900506 | 3.39E-18 | 1.33E-12 |

| | | | | | | |
|---|---|---|---|---|---|---|
| 11 | 64914161 | A | C | 0.900506 | 3.39E-18 | 1.33E-12 |
| 11 | 64914164 | C | T | 0.900506 | 3.39E-18 | 1.33E-12 |
| 11 | 64914221 | T | C | 0.900506 | 3.39E-18 | 1.33E-12 |
| 11 | 64914225 | A | G | 0.900506 | 3.39E-18 | 1.33E-12 |
| 11 | 64914245 | C | G | 0.900506 | 3.39E-18 | 1.33E-12 |
| 11 | 64914317 | A | G | 0.900506 | 3.39E-18 | 1.33E-12 |
| 11 | 64914377 | G | A | 0.900506 | 3.39E-18 | 1.33E-12 |
| 11 | 64914405 | C | T | 0.900506 | 3.39E-18 | 1.33E-12 |
| 11 | 64914419 | C | T | 0.900506 | 3.39E-18 | 1.33E-12 |
| 11 | 64914488 | A | G | 0.900506 | 3.39E-18 | 1.33E-12 |
| 11 | 64914809 | A | G | 0.900506 | 3.39E-18 | 1.33E-12 |
| 11 | 64914902 | T | A | 0.900506 | 3.39E-18 | 1.33E-12 |
| 11 | 64914948 | G | A | 0.900506 | 3.39E-18 | 1.33E-12 |
| 11 | 64915056 | C | G | 0.900506 | 3.39E-18 | 1.33E-12 |
| 11 | 64915573 | C | T | 0.900506 | 3.39E-18 | 1.33E-12 |
| 11 | 64916071 | A | C | 0.900506 | 3.39E-18 | 1.33E-12 |
| 11 | 64916884 | C | A | 0.900506 | 3.39E-18 | 1.33E-12 |
| 11 | 64917145 | C | T | 0.900506 | 3.39E-18 | 1.33E-12 |
| 11 | 64917395 | T | G | 0.900506 | 3.39E-18 | 1.33E-12 |
| 11 | 64917509 | A | G | 0.900506 | 3.39E-18 | 1.33E-12 |
| 11 | 64917908 | G | A | 0.900506 | 3.39E-18 | 1.33E-12 |
| 11 | 64920473 | A | G | 0.900506 | 3.39E-18 | 1.33E-12 |
| 11 | 64920562 | C | T | 0.900506 | 3.39E-18 | 1.33E-12 |
| 11 | 64921695 | C | T | 0.900506 | 3.39E-18 | 1.33E-12 |
| 11 | 64922592 | A | G | 0.900506 | 3.39E-18 | 1.33E-12 |
| 11 | 64922943 | A | G | 0.900506 | 3.39E-18 | 1.33E-12 |
| 11 | 64923152 | C | A | 0.900506 | 3.39E-18 | 1.33E-12 |
| 11 | 64924660 | C | G | 0.900506 | 3.39E-18 | 1.33E-12 |
| 11 | 64924702 | T | C | 0.900506 | 3.39E-18 | 1.33E-12 |
| 11 | 64928796 | A | G | 0.900506 | 3.39E-18 | 1.33E-12 |
| 11 | 64929017 | T | C | 0.900506 | 3.39E-18 | 1.33E-12 |
| 11 | 64929736 | T | C | 0.900506 | 3.39E-18 | 1.33E-12 |
| 11 | 64929740 | T | A | 0.900506 | 3.39E-18 | 1.33E-12 |
| 11 | 64929820 | G | T | 0.900506 | 3.39E-18 | 1.33E-12 |
| 11 | 64930599 | C | T | 0.900506 | 3.39E-18 | 1.33E-12 |
| 11 | 64930919 | T | C | 0.900506 | 3.39E-18 | 1.33E-12 |
| 11 | 64930977 | T | C | 0.900506 | 3.39E-18 | 1.33E-12 |
| 11 | 64931489 | T | G | 0.900506 | 3.39E-18 | 1.33E-12 |
| 11 | 64936308 | T | C | 0.900506 | 3.39E-18 | 1.33E-12 |
| 11 | 64937970 | G | A | 0.900506 | 3.39E-18 | 1.33E-12 |
| 11 | 64938838 | T | C | 0.900506 | 3.39E-18 | 1.33E-12 |
| 11 | 64954467 | A | T | 0.900506 | 3.39E-18 | 1.33E-12 |
| 11 | 64954629 | G | A | 0.900506 | 3.39E-18 | 1.33E-12 |
| 11 | 64956207 | C | T | 0.900506 | 3.39E-18 | 1.33E-12 |
| 11 | 64957282 | T | G | 0.900506 | 3.39E-18 | 1.33E-12 |
| 11 | 64957283 | A | G | 0.900506 | 3.39E-18 | 1.33E-12 |
| 11 | 64957312 | G | A | 0.900506 | 3.39E-18 | 1.33E-12 |
| 11 | 64960650 | T | A | 0.900506 | 3.39E-18 | 1.33E-12 |
| 11 | 64961195 | T | C | 0.900506 | 3.39E-18 | 1.33E-12 |
| 11 | 64961891 | T | C | 0.900506 | 3.39E-18 | 1.33E-12 |
| 11 | 64962686 | A | T | 0.900506 | 3.39E-18 | 1.33E-12 |
| 11 | 64962738 | A | G | 0.900506 | 3.39E-18 | 1.33E-12 |
| 11 | 64962794 | T | A | 0.900506 | 3.39E-18 | 1.33E-12 |
| 11 | 64962965 | C | T | 0.900506 | 3.39E-18 | 1.33E-12 |
| 11 | 64963359 | A | G | 0.900506 | 3.39E-18 | 1.33E-12 |
| 11 | 64964501 | A | G | 0.900506 | 3.39E-18 | 1.33E-12 |
| 11 | 64964530 | G | A | 0.900506 | 3.39E-18 | 1.33E-12 |
| 11 | 64964542 | T | C | 0.900506 | 3.39E-18 | 1.33E-12 |
| 11 | 64964543 | G | A | 0.900506 | 3.39E-18 | 1.33E-12 |
| 11 | 64964547 | G | A | 0.900506 | 3.39E-18 | 1.33E-12 |
| 11 | 64964569 | A | G | 0.900506 | 3.39E-18 | 1.33E-12 |
| 11 | 64964653 | A | G | 0.900506 | 3.39E-18 | 1.33E-12 |
| 11 | 64964776 | C | T | 0.900506 | 3.39E-18 | 1.33E-12 |
| 11 | 64964947 | T | C | 0.900506 | 3.39E-18 | 1.33E-12 |
| 11 | 64965206 | G | A | 0.900506 | 3.39E-18 | 1.33E-12 |
| 11 | 64965228 | T | C | 0.900506 | 3.39E-18 | 1.33E-12 |
| 11 | 64965278 | C | G | 0.900506 | 3.39E-18 | 1.33E-12 |

| | | | | | | |
|---|---|---|---|---|---|---|
| 11 | 64965302 | T | C | 0.900506 | 3.39E-18 | 1.33E-12 |
| 11 | 64965320 | T | C | 0.900506 | 3.39E-18 | 1.33E-12 |
| 11 | 64965496 | T | A | 0.900506 | 3.39E-18 | 1.33E-12 |
| 11 | 64965993 | C | T | 0.900506 | 3.39E-18 | 1.33E-12 |
| 11 | 64966833 | A | C | 0.900506 | 3.39E-18 | 1.33E-12 |
| 11 | 64970575 | A | T | 0.900506 | 3.39E-18 | 1.33E-12 |
| 11 | 64970762 | A | G | 0.900506 | 3.39E-18 | 1.33E-12 |
| 11 | 64970774 | A | G | 0.900506 | 3.39E-18 | 1.33E-12 |
| 11 | 64970990 | C | T | 0.900506 | 3.39E-18 | 1.33E-12 |
| 11 | 65082655 | G | A | 0.900506 | 3.39E-18 | 1.33E-12 |
| 11 | 65083746 | T | C | 0.900506 | 3.39E-18 | 1.33E-12 |
| 11 | 65083832 | T | C | 0.900506 | 3.39E-18 | 1.33E-12 |
| 11 | 65083856 | G | C | 0.900506 | 3.39E-18 | 1.33E-12 |
| 11 | 65084475 | A | T | 0.900506 | 3.39E-18 | 1.33E-12 |
| 11 | 65084635 | A | G | 0.900506 | 3.39E-18 | 1.33E-12 |
| 11 | 65085667 | T | C | 0.900506 | 3.39E-18 | 1.33E-12 |
| 11 | 65085970 | T | C | 0.900506 | 3.39E-18 | 1.33E-12 |
| 11 | 65098838 | A | G | 0.900506 | 3.39E-18 | 1.33E-12 |
| 11 | 64943265 | T | C | 0.904722 | 3.79E-18 | 1.48E-12 |
| 11 | 64957952 | T | C | 0.902192 | 4.34E-18 | 1.70E-12 |
| 11 | 64958345 | T | G | 0.902192 | 4.34E-18 | 1.70E-12 |
| 11 | 64958583 | C | T | 0.902192 | 4.34E-18 | 1.70E-12 |
| 11 | 64958585 | C | T | 0.902192 | 4.34E-18 | 1.70E-12 |
| 11 | 64960165 | T | A | 0.902192 | 4.34E-18 | 1.70E-12 |
| 11 | 64960413 | T | G | 0.902192 | 4.34E-18 | 1.70E-12 |
| 11 | 63290071 | A | C | 0.88027 | 5.45E-18 | 2.14E-12 |
| 11 | 64958732 | C | T | 0.901349 | 5.78E-18 | 2.27E-12 |
| 11 | 64919517 | A | G | 0.901349 | 1.06E-17 | 4.14E-12 |
| 11 | 64919566 | C | A | 0.901349 | 1.06E-17 | 4.14E-12 |
| 11 | 64919597 | C | G | 0.901349 | 1.06E-17 | 4.14E-12 |
| 11 | 64919616 | A | G | 0.901349 | 1.06E-17 | 4.14E-12 |
| 11 | 64919634 | C | A | 0.901349 | 1.06E-17 | 4.14E-12 |
| 11 | 64919643 | C | T | 0.901349 | 1.06E-17 | 4.14E-12 |
| 11 | 64919668 | G | A | 0.901349 | 1.06E-17 | 4.14E-12 |
| 11 | 64919683 | C | T | 0.901349 | 1.06E-17 | 4.14E-12 |
| 11 | 64919720 | C | T | 0.901349 | 1.06E-17 | 4.14E-12 |
| 11 | 64919791 | A | G | 0.901349 | 1.06E-17 | 4.14E-12 |
| 11 | 64919927 | G | T | 0.901349 | 1.06E-17 | 4.14E-12 |
| 11 | 65080188 | C | T | 0.901349 | 1.12E-17 | 4.39E-12 |
| 11 | 64921356 | C | G | 0.89882 | 1.98E-17 | 7.75E-12 |
| 11 | 64921376 | A | G | 0.89882 | 1.98E-17 | 7.75E-12 |
| 11 | 64921535 | T | G | 0.89882 | 1.98E-17 | 7.75E-12 |
| 11 | 64921552 | C | G | 0.89882 | 1.98E-17 | 7.75E-12 |
| 11 | 64924880 | T | A | 0.899663 | 2.05E-17 | 8.05E-12 |
| 11 | 64926542 | T | C | 0.899663 | 2.05E-17 | 8.05E-12 |
| 11 | 64926622 | A | G | 0.899663 | 2.05E-17 | 8.05E-12 |
| 11 | 66099866 | A | C | 0.900506 | 2.26E-17 | 8.87E-12 |
| 11 | 65417185 | T | C | 0.922428 | 3.10E-17 | 1.22E-11 |
| 11 | 65417210 | C | A | 0.922428 | 3.10E-17 | 1.22E-11 |
| 11 | 65417024 | G | A | 0.924115 | 3.44E-17 | 1.35E-11 |
| 11 | 64942416 | T | C | 0.901349 | 3.62E-17 | 1.42E-11 |
| 11 | 63793100 | A | C | 0.897976 | 4.04E-17 | 1.59E-11 |
| 11 | 63793113 | G | C | 0.897976 | 4.04E-17 | 1.59E-11 |
| 11 | 63793250 | T | C | 0.897976 | 4.04E-17 | 1.59E-11 |
| 11 | 63793360 | A | T | 0.897976 | 4.04E-17 | 1.59E-11 |
| 11 | 63830293 | T | C | 0.89629 | 6.34E-17 | 2.48E-11 |
| 11 | 63848519 | T | C | 0.900506 | 6.56E-17 | 2.57E-11 |
| 11 | 63848625 | G | C | 0.900506 | 6.56E-17 | 2.57E-11 |
| 11 | 63930502 | C | T | 0.865936 | 7.47E-17 | 2.93E-11 |
| 11 | 63930778 | A | G | 0.865936 | 7.47E-17 | 2.93E-11 |
| 11 | 63930804 | T | C | 0.865936 | 7.47E-17 | 2.93E-11 |
| 11 | 63931480 | A | G | 0.865936 | 7.47E-17 | 2.93E-11 |
| 11 | 63931985 | A | G | 0.865936 | 7.47E-17 | 2.93E-11 |
| 11 | 63932614 | T | C | 0.865936 | 7.47E-17 | 2.93E-11 |
| 11 | 63933053 | A | G | 0.865936 | 7.47E-17 | 2.93E-11 |
| 11 | 63933073 | G | T | 0.865936 | 7.47E-17 | 2.93E-11 |
| 11 | 63933301 | A | G | 0.865936 | 7.47E-17 | 2.93E-11 |

| | | | | | | |
|---|---|---|---|---|---|---|
| 11 | 63934043 | T | C | 0.865936 | 7.47E-17 | 2.93E-11 |
| 11 | 63934711 | G | C | 0.865936 | 7.47E-17 | 2.93E-11 |
| 11 | 63934817 | T | A | 0.865936 | 7.47E-17 | 2.93E-11 |
| 11 | 63934905 | C | T | 0.865936 | 7.47E-17 | 2.93E-11 |
| 11 | 63935009 | C | T | 0.865936 | 7.47E-17 | 2.93E-11 |
| 11 | 63936846 | T | A | 0.865936 | 7.47E-17 | 2.93E-11 |
| 11 | 63939386 | A | C | 0.865936 | 7.47E-17 | 2.93E-11 |
| 11 | 63940631 | T | C | 0.865936 | 7.47E-17 | 2.93E-11 |
| 11 | 63941785 | G | A | 0.865936 | 7.47E-17 | 2.93E-11 |
| 11 | 63941887 | A | C | 0.865936 | 7.47E-17 | 2.93E-11 |
| 11 | 63941978 | T | G | 0.865936 | 7.47E-17 | 2.93E-11 |
| 11 | 63941993 | A | T | 0.865936 | 7.47E-17 | 2.93E-11 |
| 11 | 63942843 | A | G | 0.865936 | 7.47E-17 | 2.93E-11 |
| 11 | 63943128 | G | C | 0.865936 | 7.47E-17 | 2.93E-11 |
| 11 | 63944904 | T | C | 0.865936 | 7.47E-17 | 2.93E-11 |
| 11 | 63945538 | G | A | 0.865936 | 7.47E-17 | 2.93E-11 |
| 11 | 63945715 | A | C | 0.865936 | 7.47E-17 | 2.93E-11 |
| 11 | 63945934 | G | C | 0.865936 | 7.47E-17 | 2.93E-11 |
| 11 | 63946495 | A | C | 0.865936 | 7.47E-17 | 2.93E-11 |
| 11 | 63946508 | T | C | 0.865936 | 7.47E-17 | 2.93E-11 |
| 11 | 63946569 | A | G | 0.865936 | 7.47E-17 | 2.93E-11 |
| 11 | 63946570 | G | A | 0.865936 | 7.47E-17 | 2.93E-11 |
| 11 | 63946745 | G | A | 0.865936 | 7.47E-17 | 2.93E-11 |
| 11 | 63946836 | G | A | 0.865936 | 7.47E-17 | 2.93E-11 |
| 11 | 63946868 | C | T | 0.865936 | 7.47E-17 | 2.93E-11 |
| 11 | 63946877 | A | G | 0.865936 | 7.47E-17 | 2.93E-11 |
| 11 | 63946884 | T | G | 0.865936 | 7.47E-17 | 2.93E-11 |
| 11 | 63946903 | T | C | 0.865936 | 7.47E-17 | 2.93E-11 |
| 11 | 63946928 | T | A | 0.865936 | 7.47E-17 | 2.93E-11 |
| 11 | 63947702 | A | G | 0.865936 | 7.47E-17 | 2.93E-11 |
| 11 | 63947826 | T | C | 0.865936 | 7.47E-17 | 2.93E-11 |
| 11 | 63947841 | A | G | 0.865936 | 7.47E-17 | 2.93E-11 |
| 11 | 63947880 | T | G | 0.865936 | 7.47E-17 | 2.93E-11 |
| 11 | 63947915 | T | C | 0.865936 | 7.47E-17 | 2.93E-11 |
| 11 | 63948168 | C | T | 0.865936 | 7.47E-17 | 2.93E-11 |
| 11 | 63948179 | A | G | 0.865936 | 7.47E-17 | 2.93E-11 |
| 11 | 63948203 | A | T | 0.865936 | 7.47E-17 | 2.93E-11 |
| 11 | 63948437 | C | G | 0.865936 | 7.47E-17 | 2.93E-11 |
| 11 | 63948438 | A | G | 0.865936 | 7.47E-17 | 2.93E-11 |
| 11 | 63948446 | C | T | 0.865936 | 7.47E-17 | 2.93E-11 |
| 11 | 63948557 | A | G | 0.865936 | 7.47E-17 | 2.93E-11 |
| 11 | 63948574 | A | G | 0.865936 | 7.47E-17 | 2.93E-11 |
| 11 | 63948680 | T | C | 0.865936 | 7.47E-17 | 2.93E-11 |
| 11 | 63948684 | C | T | 0.865936 | 7.47E-17 | 2.93E-11 |
| 11 | 63948727 | A | G | 0.865936 | 7.47E-17 | 2.93E-11 |
| 11 | 63948803 | C | T | 0.865936 | 7.47E-17 | 2.93E-11 |
| 11 | 63948871 | A | G | 0.865936 | 7.47E-17 | 2.93E-11 |
| 11 | 63949263 | A | G | 0.865936 | 7.47E-17 | 2.93E-11 |
| 11 | 63949355 | T | C | 0.865936 | 7.47E-17 | 2.93E-11 |
| 11 | 63949675 | G | A | 0.865936 | 7.47E-17 | 2.93E-11 |
| 11 | 63949708 | A | T | 0.865936 | 7.47E-17 | 2.93E-11 |
| 11 | 63949887 | G | A | 0.865936 | 7.47E-17 | 2.93E-11 |
| 11 | 63951088 | C | T | 0.865936 | 7.47E-17 | 2.93E-11 |
| 11 | 63951362 | C | T | 0.865936 | 7.47E-17 | 2.93E-11 |
| 11 | 63951396 | T | C | 0.865936 | 7.47E-17 | 2.93E-11 |
| 11 | 63952219 | A | G | 0.865936 | 7.47E-17 | 2.93E-11 |
| 11 | 63952554 | A | G | 0.865936 | 7.47E-17 | 2.93E-11 |
| 11 | 63955292 | A | G | 0.865936 | 7.47E-17 | 2.93E-11 |
| 11 | 63955349 | T | C | 0.865936 | 7.47E-17 | 2.93E-11 |
| 11 | 63955727 | T | A | 0.865936 | 7.47E-17 | 2.93E-11 |
| 11 | 63955910 | G | A | 0.865936 | 7.47E-17 | 2.93E-11 |
| 11 | 63831954 | A | C | 0.866779 | 8.08E-17 | 3.17E-11 |
| 11 | 63770597 | C | G | 0.900506 | 8.20E-17 | 3.22E-11 |
| 11 | 65084150 | G | A | 0.902192 | 9.17E-17 | 3.60E-11 |
| 11 | 63792512 | A | T | 0.91231 | 1.01E-16 | 3.97E-11 |
| 11 | 63930484 | G | A | 0.865093 | 1.23E-16 | 4.81E-11 |
| 11 | 63930494 | C | A | 0.865093 | 1.23E-16 | 4.81E-11 |

| | | | | | | |
|---|---|---|---|---|---|---|
| 11 | 63930495 | A | C | 0.865093 | 1.23E-16 | 4.81E-11 |
| 11 | 64899359 | C | G | 0.89882 | 1.25E-16 | 4.88E-11 |
| 11 | 64899412 | C | G | 0.89882 | 1.25E-16 | 4.88E-11 |
| 11 | 64899415 | C | A | 0.89882 | 1.25E-16 | 4.88E-11 |
| 11 | 64899425 | T | G | 0.89882 | 1.25E-16 | 4.88E-11 |
| 11 | 64899451 | A | G | 0.89882 | 1.25E-16 | 4.88E-11 |
| 11 | 63930355 | G | A | 0.867622 | 1.30E-16 | 5.11E-11 |
| 11 | 63930396 | A | T | 0.867622 | 1.30E-16 | 5.11E-11 |
| 11 | 63793862 | A | G | 0.905565 | 1.32E-16 | 5.18E-11 |
| 11 | 63792635 | A | G | 0.911467 | 1.61E-16 | 6.31E-11 |
| 11 | 63792918 | C | G | 0.911467 | 1.61E-16 | 6.31E-11 |
| 11 | 63831450 | A | C | 0.86425 | 1.62E-16 | 6.33E-11 |
| 11 | 63831454 | A | G | 0.86425 | 1.62E-16 | 6.33E-11 |
| 11 | 63831060 | G | A | 0.863406 | 1.64E-16 | 6.44E-11 |
| 11 | 63959931 | C | A | 0.866779 | 1.77E-16 | 6.93E-11 |
| 11 | 63963961 | A | G | 0.866779 | 1.77E-16 | 6.93E-11 |
| 11 | 64056828 | A | G | 0.904722 | 2.18E-16 | 8.54E-11 |
| 11 | 64057013 | T | G | 0.904722 | 2.18E-16 | 8.54E-11 |
| 11 | 64057174 | A | G | 0.904722 | 2.18E-16 | 8.54E-11 |
| 11 | 64057268 | T | C | 0.904722 | 2.18E-16 | 8.54E-11 |
| 11 | 64057328 | A | G | 0.904722 | 2.18E-16 | 8.54E-11 |
| 11 | 64057447 | G | A | 0.904722 | 2.18E-16 | 8.54E-11 |
| 11 | 64057519 | G | A | 0.904722 | 2.18E-16 | 8.54E-11 |
| 11 | 64057690 | G | A | 0.904722 | 2.18E-16 | 8.54E-11 |
| 11 | 64057711 | A | C | 0.904722 | 2.18E-16 | 8.54E-11 |
| 11 | 64057900 | A | G | 0.904722 | 2.18E-16 | 8.54E-11 |
| 11 | 64057991 | A | G | 0.904722 | 2.18E-16 | 8.54E-11 |
| 11 | 64058478 | A | G | 0.904722 | 2.18E-16 | 8.54E-11 |
| 11 | 64058511 | A | G | 0.904722 | 2.18E-16 | 8.54E-11 |
| 11 | 64060440 | A | G | 0.904722 | 2.18E-16 | 8.54E-11 |
| 11 | 64060759 | A | G | 0.904722 | 2.18E-16 | 8.54E-11 |
| 11 | 64063261 | T | C | 0.904722 | 2.18E-16 | 8.54E-11 |
| 11 | 64065481 | A | G | 0.904722 | 2.18E-16 | 8.54E-11 |
| 11 | 64065536 | A | C | 0.904722 | 2.18E-16 | 8.54E-11 |
| 11 | 64067558 | A | T | 0.904722 | 2.18E-16 | 8.54E-11 |
| 11 | 64068068 | T | C | 0.904722 | 2.18E-16 | 8.54E-11 |
| 11 | 64069340 | C | T | 0.904722 | 2.18E-16 | 8.54E-11 |
| 11 | 64069346 | G | A | 0.904722 | 2.18E-16 | 8.54E-11 |
| 11 | 64070147 | T | C | 0.904722 | 2.18E-16 | 8.54E-11 |
| 11 | 64070265 | C | T | 0.904722 | 2.18E-16 | 8.54E-11 |
| 11 | 64071438 | A | G | 0.904722 | 2.18E-16 | 8.54E-11 |
| 11 | 64073916 | G | C | 0.904722 | 2.18E-16 | 8.54E-11 |
| 11 | 64075041 | T | C | 0.904722 | 2.18E-16 | 8.54E-11 |
| 11 | 64091032 | A | G | 0.904722 | 2.18E-16 | 8.54E-11 |
| 11 | 64095398 | C | T | 0.904722 | 2.18E-16 | 8.54E-11 |
| 11 | 64096537 | C | T | 0.904722 | 2.18E-16 | 8.54E-11 |
| 11 | 64103961 | A | G | 0.904722 | 2.18E-16 | 8.54E-11 |
| 11 | 64104792 | T | G | 0.904722 | 2.18E-16 | 8.54E-11 |
| 11 | 64105774 | T | C | 0.904722 | 2.18E-16 | 8.54E-11 |
| 11 | 64106749 | A | G | 0.904722 | 2.18E-16 | 8.54E-11 |
| 11 | 64108551 | T | C | 0.904722 | 2.18E-16 | 8.54E-11 |
| 11 | 64109028 | T | C | 0.904722 | 2.18E-16 | 8.54E-11 |
| 11 | 64109391 | A | G | 0.904722 | 2.18E-16 | 8.54E-11 |
| 11 | 64109535 | A | G | 0.904722 | 2.18E-16 | 8.54E-11 |
| 11 | 64109555 | T | C | 0.904722 | 2.18E-16 | 8.54E-11 |
| 11 | 64109566 | A | G | 0.904722 | 2.18E-16 | 8.54E-11 |
| 11 | 64109761 | A | G | 0.904722 | 2.18E-16 | 8.54E-11 |
| 11 | 64110589 | A | G | 0.904722 | 2.18E-16 | 8.54E-11 |
| 11 | 64110634 | A | G | 0.904722 | 2.18E-16 | 8.54E-11 |
| 11 | 64110703 | A | G | 0.904722 | 2.18E-16 | 8.54E-11 |
| 11 | 64110731 | A | G | 0.904722 | 2.18E-16 | 8.54E-11 |
| 11 | 64111345 | A | G | 0.904722 | 2.18E-16 | 8.54E-11 |
| 11 | 64111440 | T | C | 0.904722 | 2.18E-16 | 8.54E-11 |
| 11 | 64111509 | G | A | 0.904722 | 2.18E-16 | 8.54E-11 |
| 11 | 64111535 | G | A | 0.904722 | 2.18E-16 | 8.54E-11 |
| 11 | 64111835 | G | T | 0.904722 | 2.18E-16 | 8.54E-11 |
| 11 | 64111919 | T | C | 0.904722 | 2.18E-16 | 8.54E-11 |

| | | | | | | |
|---|---|---|---|---|---|---|
| 11 | 64112179 | A | G | 0.904722 | 2.18E-16 | 8.54E-11 |
| 11 | 64112202 | T | C | 0.904722 | 2.18E-16 | 8.54E-11 |
| 11 | 64112555 | T | C | 0.904722 | 2.18E-16 | 8.54E-11 |
| 11 | 64112556 | G | A | 0.904722 | 2.18E-16 | 8.54E-11 |
| 11 | 64112633 | T | C | 0.904722 | 2.18E-16 | 8.54E-11 |
| 11 | 64114104 | T | A | 0.904722 | 2.18E-16 | 8.54E-11 |
| 11 | 64114324 | A | G | 0.904722 | 2.18E-16 | 8.54E-11 |
| 11 | 64114346 | G | A | 0.904722 | 2.18E-16 | 8.54E-11 |
| 11 | 64114542 | C | T | 0.904722 | 2.18E-16 | 8.54E-11 |
| 11 | 64114603 | C | T | 0.904722 | 2.18E-16 | 8.54E-11 |
| 11 | 64063817 | C | T | 0.901349 | 2.21E-16 | 8.66E-11 |
| 11 | 64940989 | A | G | 0.899663 | 2.50E-16 | 9.81E-11 |
| 11 | 63963505 | G | A | 0.867622 | 3.31E-16 | 1.30E-10 |
| 11 | 63830678 | C | T | 0.865093 | 3.72E-16 | 1.46E-10 |
| 11 | 63831089 | C | A | 0.865093 | 3.72E-16 | 1.46E-10 |
| 11 | 63770299 | T | C | 0.899663 | 4.35E-16 | 1.71E-10 |
| 11 | 63770379 | T | C | 0.899663 | 4.35E-16 | 1.71E-10 |
| 11 | 63770397 | G | C | 0.899663 | 4.35E-16 | 1.71E-10 |
| 11 | 63770408 | C | G | 0.899663 | 4.35E-16 | 1.71E-10 |
| 11 | 65412003 | A | T | 0.920742 | 5.20E-16 | 2.04E-10 |
| 11 | 66088675 | G | A | 0.901349 | 5.45E-16 | 2.14E-10 |
| 11 | 64055656 | T | G | 0.86172 | 6.36E-16 | 2.49E-10 |
| 11 | 64898617 | T | G | 0.887858 | 7.49E-16 | 2.94E-10 |
| 11 | 64898723 | T | C | 0.887858 | 7.49E-16 | 2.94E-10 |
| 11 | 64899026 | C | T | 0.887858 | 7.49E-16 | 2.94E-10 |
| 11 | 64170644 | G | A | 0.901349 | 9.69E-16 | 3.80E-10 |
| 11 | 64897748 | C | T | 0.887015 | 1.04E-15 | 4.07E-10 |
| 11 | 63837178 | A | G | 0.892074 | 1.25E-15 | 4.90E-10 |
| 11 | 63837190 | A | G | 0.892074 | 1.25E-15 | 4.90E-10 |
| 11 | 63908717 | C | G | 0.882799 | 1.39E-15 | 5.46E-10 |
| 11 | 63908720 | C | T | 0.882799 | 1.39E-15 | 5.46E-10 |
| 11 | 63908721 | A | G | 0.882799 | 1.39E-15 | 5.46E-10 |
| 11 | 63908734 | C | T | 0.882799 | 1.39E-15 | 5.46E-10 |
| 11 | 63908766 | C | G | 0.882799 | 1.39E-15 | 5.46E-10 |
| 11 | 63908767 | T | C | 0.882799 | 1.39E-15 | 5.46E-10 |
| 11 | 63908774 | T | G | 0.882799 | 1.39E-15 | 5.46E-10 |
| 11 | 63908803 | C | T | 0.882799 | 1.39E-15 | 5.46E-10 |
| 11 | 63908825 | C | G | 0.882799 | 1.39E-15 | 5.46E-10 |
| 11 | 63908827 | C | A | 0.882799 | 1.39E-15 | 5.46E-10 |
| 11 | 63908828 | G | A | 0.882799 | 1.39E-15 | 5.46E-10 |
| 11 | 63908841 | G | A | 0.882799 | 1.39E-15 | 5.46E-10 |
| 11 | 63908891 | A | G | 0.882799 | 1.40E-15 | 5.48E-10 |
| 11 | 63908935 | A | G | 0.882799 | 1.40E-15 | 5.48E-10 |
| 11 | 63908936 | C | T | 0.882799 | 1.40E-15 | 5.48E-10 |
| 11 | 63908937 | A | G | 0.882799 | 1.40E-15 | 5.48E-10 |
| 11 | 63908945 | C | T | 0.882799 | 1.40E-15 | 5.48E-10 |
| 11 | 63768531 | T | C | 0.900506 | 1.42E-15 | 5.55E-10 |
| 11 | 63939019 | A | G | 0.867622 | 1.42E-15 | 5.57E-10 |
| 11 | 63939145 | G | C | 0.867622 | 1.42E-15 | 5.57E-10 |
| 11 | 63939146 | G | A | 0.867622 | 1.42E-15 | 5.57E-10 |
| 11 | 63939147 | C | T | 0.867622 | 1.42E-15 | 5.57E-10 |
| 11 | 63768747 | C | G | 0.897976 | 1.60E-15 | 6.28E-10 |
| 11 | 63768748 | T | G | 0.897976 | 1.60E-15 | 6.28E-10 |
| 11 | 64171172 | C | T | 0.900506 | 1.61E-15 | 6.33E-10 |
| 11 | 63908948 | C | G | 0.883642 | 2.13E-15 | 8.36E-10 |
| 11 | 63908949 | C | A | 0.883642 | 2.13E-15 | 8.36E-10 |
| 11 | 63908671 | G | C | 0.883642 | 2.16E-15 | 8.46E-10 |
| 11 | 63908679 | C | G | 0.883642 | 2.16E-15 | 8.46E-10 |
| 11 | 63908680 | C | T | 0.883642 | 2.16E-15 | 8.46E-10 |
| 11 | 63908681 | A | G | 0.883642 | 2.16E-15 | 8.46E-10 |
| 11 | 63908682 | C | A | 0.883642 | 2.16E-15 | 8.46E-10 |
| 11 | 63908703 | G | A | 0.883642 | 2.16E-15 | 8.46E-10 |
| 11 | 63908708 | A | G | 0.883642 | 2.16E-15 | 8.46E-10 |
| 11 | 63768749 | A | T | 0.897133 | 2.28E-15 | 8.94E-10 |
| 11 | 63768788 | A | G | 0.897133 | 2.28E-15 | 8.94E-10 |
| 11 | 63768808 | G | T | 0.897133 | 2.28E-15 | 8.94E-10 |
| 11 | 63908969 | C | T | 0.884486 | 2.39E-15 | 9.36E-10 |

| | | | | | | |
|---|---|---|---|---|---|---|
| 11 | 63908976 | C | G | 0.884486 | 2.39E-15 | 9.36E-10 |
| 11 | 63908981 | C | T | 0.884486 | 2.39E-15 | 9.36E-10 |
| 11 | 64114998 | T | A | 0.903035 | 2.57E-15 | 1.01E-09 |
| 11 | 64116247 | A | C | 0.903035 | 2.57E-15 | 1.01E-09 |
| 11 | 64116319 | G | T | 0.903035 | 2.57E-15 | 1.01E-09 |
| 11 | 64116360 | A | C | 0.903035 | 2.57E-15 | 1.01E-09 |
| 11 | 64116459 | A | G | 0.903035 | 2.57E-15 | 1.01E-09 |
| 11 | 64116488 | T | C | 0.903035 | 2.57E-15 | 1.01E-09 |
| 11 | 64116738 | A | G | 0.903035 | 2.57E-15 | 1.01E-09 |
| 11 | 64117082 | G | A | 0.903035 | 2.57E-15 | 1.01E-09 |
| 11 | 64117083 | T | C | 0.903035 | 2.57E-15 | 1.01E-09 |
| 11 | 64117246 | A | G | 0.903035 | 2.57E-15 | 1.01E-09 |
| 11 | 64117971 | A | T | 0.903035 | 2.57E-15 | 1.01E-09 |
| 11 | 64118120 | T | C | 0.903035 | 2.57E-15 | 1.01E-09 |
| 11 | 64118370 | A | G | 0.903035 | 2.57E-15 | 1.01E-09 |
| 11 | 64118480 | A | G | 0.903035 | 2.57E-15 | 1.01E-09 |
| 11 | 64118697 | G | A | 0.903035 | 2.57E-15 | 1.01E-09 |
| 11 | 64118758 | T | C | 0.903035 | 2.57E-15 | 1.01E-09 |
| 11 | 64119256 | G | T | 0.903035 | 2.57E-15 | 1.01E-09 |
| 11 | 64119348 | T | C | 0.903035 | 2.57E-15 | 1.01E-09 |
| 11 | 64119354 | G | A | 0.903035 | 2.57E-15 | 1.01E-09 |
| 11 | 64119899 | G | A | 0.903035 | 2.57E-15 | 1.01E-09 |
| 11 | 64119951 | C | G | 0.903035 | 2.57E-15 | 1.01E-09 |
| 11 | 64120027 | G | A | 0.903035 | 2.57E-15 | 1.01E-09 |
| 11 | 64120179 | T | C | 0.903035 | 2.57E-15 | 1.01E-09 |
| 11 | 64120190 | A | G | 0.903035 | 2.57E-15 | 1.01E-09 |
| 11 | 64120268 | T | C | 0.903035 | 2.57E-15 | 1.01E-09 |
| 11 | 64120424 | A | T | 0.903035 | 2.57E-15 | 1.01E-09 |
| 11 | 64120442 | C | T | 0.903035 | 2.57E-15 | 1.01E-09 |
| 11 | 64120461 | A | G | 0.903035 | 2.57E-15 | 1.01E-09 |
| 11 | 64120525 | A | T | 0.903035 | 2.57E-15 | 1.01E-09 |
| 11 | 64120565 | T | G | 0.903035 | 2.57E-15 | 1.01E-09 |
| 11 | 64120637 | T | C | 0.903035 | 2.57E-15 | 1.01E-09 |
| 11 | 64120720 | T | C | 0.903035 | 2.57E-15 | 1.01E-09 |
| 11 | 64120891 | A | G | 0.903035 | 2.57E-15 | 1.01E-09 |
| 11 | 64120989 | A | G | 0.903035 | 2.57E-15 | 1.01E-09 |
| 11 | 64122381 | G | A | 0.903035 | 2.57E-15 | 1.01E-09 |
| 11 | 64122611 | T | C | 0.903035 | 2.57E-15 | 1.01E-09 |
| 11 | 64123298 | T | G | 0.903035 | 2.57E-15 | 1.01E-09 |
| 11 | 64123406 | A | G | 0.903035 | 2.57E-15 | 1.01E-09 |
| 11 | 64123431 | T | G | 0.903035 | 2.57E-15 | 1.01E-09 |
| 11 | 64123454 | G | A | 0.903035 | 2.57E-15 | 1.01E-09 |
| 11 | 64123659 | A | C | 0.903035 | 2.57E-15 | 1.01E-09 |
| 11 | 64123990 | G | A | 0.903035 | 2.57E-15 | 1.01E-09 |
| 11 | 64124122 | T | C | 0.903035 | 2.57E-15 | 1.01E-09 |
| 11 | 64124761 | G | A | 0.903035 | 2.57E-15 | 1.01E-09 |
| 11 | 64125266 | G | A | 0.903035 | 2.57E-15 | 1.01E-09 |
| 11 | 64125693 | C | T | 0.903035 | 2.57E-15 | 1.01E-09 |
| 11 | 64125748 | C | T | 0.903035 | 2.57E-15 | 1.01E-09 |
| 11 | 64125877 | A | G | 0.903035 | 2.57E-15 | 1.01E-09 |
| 11 | 64125935 | T | C | 0.903035 | 2.57E-15 | 1.01E-09 |
| 11 | 64126182 | C | T | 0.903035 | 2.57E-15 | 1.01E-09 |
| 11 | 64126413 | T | A | 0.903035 | 2.57E-15 | 1.01E-09 |
| 11 | 64126733 | T | G | 0.903035 | 2.57E-15 | 1.01E-09 |
| 11 | 64126920 | T | G | 0.903035 | 2.57E-15 | 1.01E-09 |
| 11 | 64126951 | C | T | 0.903035 | 2.57E-15 | 1.01E-09 |
| 11 | 64127943 | G | C | 0.903035 | 2.57E-15 | 1.01E-09 |
| 11 | 64129940 | A | G | 0.903035 | 2.57E-15 | 1.01E-09 |
| 11 | 64130443 | A | G | 0.903035 | 2.57E-15 | 1.01E-09 |
| 11 | 64130695 | T | A | 0.903035 | 2.57E-15 | 1.01E-09 |
| 11 | 64130802 | G | A | 0.903035 | 2.57E-15 | 1.01E-09 |
| 11 | 64130907 | C | T | 0.903035 | 2.57E-15 | 1.01E-09 |
| 11 | 64130976 | T | C | 0.903035 | 2.57E-15 | 1.01E-09 |
| 11 | 64131137 | G | A | 0.903035 | 2.57E-15 | 1.01E-09 |
| 11 | 64131245 | A | G | 0.903035 | 2.57E-15 | 1.01E-09 |
| 11 | 64131268 | T | C | 0.903035 | 2.57E-15 | 1.01E-09 |
| 11 | 64131562 | A | C | 0.903035 | 2.57E-15 | 1.01E-09 |

| | | | | | | |
|---|---|---|---|---|---|---|
| 11 | 64131566 | A | G | 0.903035 | 2.57E-15 | 1.01E-09 |
| 11 | 64131809 | A | G | 0.903035 | 2.57E-15 | 1.01E-09 |
| 11 | 64132216 | G | A | 0.903035 | 2.57E-15 | 1.01E-09 |
| 11 | 64136308 | A | G | 0.903035 | 2.57E-15 | 1.01E-09 |
| 11 | 64137195 | C | T | 0.903035 | 2.57E-15 | 1.01E-09 |
| 11 | 64139435 | T | C | 0.903035 | 2.57E-15 | 1.01E-09 |
| 11 | 64139833 | A | T | 0.903035 | 2.57E-15 | 1.01E-09 |
| 11 | 64141333 | A | T | 0.903035 | 2.57E-15 | 1.01E-09 |
| 11 | 64141411 | G | A | 0.903035 | 2.57E-15 | 1.01E-09 |
| 11 | 64141759 | A | G | 0.903035 | 2.57E-15 | 1.01E-09 |
| 11 | 64142425 | C | T | 0.903035 | 2.57E-15 | 1.01E-09 |
| 11 | 64142726 | C | T | 0.903035 | 2.57E-15 | 1.01E-09 |
| 11 | 64143207 | G | C | 0.903035 | 2.57E-15 | 1.01E-09 |
| 11 | 64143218 | G | C | 0.903035 | 2.57E-15 | 1.01E-09 |
| 11 | 64146100 | A | G | 0.903035 | 2.57E-15 | 1.01E-09 |
| 11 | 64155253 | A | G | 0.903035 | 2.57E-15 | 1.01E-09 |
| 11 | 64155586 | C | T | 0.903035 | 2.57E-15 | 1.01E-09 |
| 11 | 64155908 | A | G | 0.903035 | 2.57E-15 | 1.01E-09 |
| 11 | 64157322 | T | C | 0.903035 | 2.57E-15 | 1.01E-09 |
| 11 | 64161557 | A | G | 0.903035 | 2.57E-15 | 1.01E-09 |
| 11 | 64161662 | G | A | 0.903035 | 2.57E-15 | 1.01E-09 |
| 11 | 64162545 | G | C | 0.903035 | 2.57E-15 | 1.01E-09 |
| 11 | 64162970 | G | A | 0.903035 | 2.57E-15 | 1.01E-09 |
| 11 | 64163111 | G | A | 0.903035 | 2.57E-15 | 1.01E-09 |
| 11 | 64163408 | C | T | 0.903035 | 2.57E-15 | 1.01E-09 |
| 11 | 64163418 | C | G | 0.903035 | 2.57E-15 | 1.01E-09 |
| 11 | 64163462 | G | A | 0.903035 | 2.57E-15 | 1.01E-09 |
| 11 | 64163478 | G | A | 0.903035 | 2.57E-15 | 1.01E-09 |
| 11 | 64163597 | T | C | 0.903035 | 2.57E-15 | 1.01E-09 |
| 11 | 64164421 | A | G | 0.903035 | 2.57E-15 | 1.01E-09 |
| 11 | 64169981 | T | C | 0.903035 | 2.57E-15 | 1.01E-09 |
| 11 | 64170138 | T | G | 0.903035 | 2.57E-15 | 1.01E-09 |
| 11 | 64170278 | G | A | 0.903035 | 2.57E-15 | 1.01E-09 |
| 11 | 64170517 | A | T | 0.903035 | 2.57E-15 | 1.01E-09 |
| 11 | 64170541 | T | C | 0.903035 | 2.57E-15 | 1.01E-09 |
| 11 | 64170561 | C | T | 0.903035 | 2.57E-15 | 1.01E-09 |
| 11 | 64170613 | A | G | 0.903035 | 2.57E-15 | 1.01E-09 |
| 11 | 64170625 | A | G | 0.903035 | 2.57E-15 | 1.01E-09 |
| 11 | 64170661 | G | C | 0.903035 | 2.57E-15 | 1.01E-09 |
| 11 | 64170678 | C | T | 0.903035 | 2.57E-15 | 1.01E-09 |
| 11 | 64170809 | A | G | 0.903035 | 2.57E-15 | 1.01E-09 |
| 11 | 64170894 | T | C | 0.903035 | 2.57E-15 | 1.01E-09 |
| 11 | 64170908 | A | G | 0.903035 | 2.57E-15 | 1.01E-09 |
| 11 | 64170923 | C | T | 0.903035 | 2.57E-15 | 1.01E-09 |
| 11 | 64170948 | G | A | 0.903035 | 2.57E-15 | 1.01E-09 |
| 11 | 63836197 | T | G | 0.890388 | 2.61E-15 | 1.02E-09 |
| 11 | 64155329 | T | C | 0.899663 | 2.88E-15 | 1.13E-09 |
| 11 | 64168180 | G | C | 0.901349 | 3.74E-15 | 1.47E-09 |
| 11 | 63768273 | G | A | 0.904722 | 4.00E-15 | 1.57E-09 |
| 11 | 64167618 | G | T | 0.89629 | 4.04E-15 | 1.58E-09 |
| 11 | 64167622 | A | G | 0.89629 | 4.04E-15 | 1.58E-09 |
| 11 | 64167734 | C | T | 0.89882 | 4.34E-15 | 1.70E-09 |
| 11 | 64167793 | T | C | 0.89882 | 4.34E-15 | 1.70E-09 |
| 11 | 64167851 | A | G | 0.89882 | 4.34E-15 | 1.70E-09 |
| 11 | 63908622 | C | T | 0.884486 | 4.79E-15 | 1.88E-09 |
| 11 | 63908626 | T | C | 0.884486 | 4.79E-15 | 1.88E-09 |
| 11 | 63908651 | C | T | 0.884486 | 4.79E-15 | 1.88E-09 |
| 11 | 63768875 | C | A | 0.897976 | 5.33E-15 | 2.09E-09 |
| 11 | 63829575 | A | C | 0.889545 | 5.40E-15 | 2.12E-09 |
| 11 | 64164907 | G | A | 0.901349 | 5.72E-15 | 2.24E-09 |
| 11 | 64170671 | A | G | 0.897976 | 8.03E-15 | 3.15E-09 |
| 11 | 64170821 | T | A | 0.897976 | 8.03E-15 | 3.15E-09 |
| 11 | 64160486 | A | G | 0.903879 | 8.08E-15 | 3.17E-09 |
| 11 | 64162231 | C | G | 0.903879 | 8.08E-15 | 3.17E-09 |
| 11 | 64163746 | G | A | 0.903879 | 8.08E-15 | 3.17E-09 |
| 11 | 64163869 | A | T | 0.903879 | 8.08E-15 | 3.17E-09 |
| 11 | 64163870 | A | T | 0.903879 | 8.08E-15 | 3.17E-09 |

| | | | | | | |
|---|---|---|---|---|---|---|
| 11 | 64163871 | A | T | 0.903879 | 8.08E-15 | 3.17E-09 |
| 11 | 64163944 | A | T | 0.903879 | 8.08E-15 | 3.17E-09 |
| 11 | 64163956 | A | G | 0.903879 | 8.08E-15 | 3.17E-09 |
| 11 | 64163981 | C | T | 0.903879 | 8.08E-15 | 3.17E-09 |
| 11 | 64164022 | A | G | 0.903879 | 8.08E-15 | 3.17E-09 |
| 11 | 64164053 | C | T | 0.903879 | 8.08E-15 | 3.17E-09 |
| 11 | 64164101 | T | C | 0.903879 | 8.08E-15 | 3.17E-09 |
| 11 | 64164109 | C | T | 0.903879 | 8.08E-15 | 3.17E-09 |
| 11 | 64164281 | C | T | 0.903879 | 8.08E-15 | 3.17E-09 |
| 11 | 64164282 | G | A | 0.903879 | 8.08E-15 | 3.17E-09 |
| 11 | 64164306 | C | T | 0.903879 | 8.08E-15 | 3.17E-09 |
| 11 | 64164319 | G | A | 0.903879 | 8.08E-15 | 3.17E-09 |
| 11 | 64164520 | C | A | 0.903879 | 8.08E-15 | 3.17E-09 |
| 11 | 64164556 | G | T | 0.903879 | 8.08E-15 | 3.17E-09 |
| 11 | 64164563 | G | A | 0.903879 | 8.08E-15 | 3.17E-09 |
| 11 | 64164582 | G | T | 0.903879 | 8.08E-15 | 3.17E-09 |
| 11 | 64164594 | T | C | 0.903879 | 8.08E-15 | 3.17E-09 |
| 11 | 64164609 | T | C | 0.903879 | 8.08E-15 | 3.17E-09 |
| 11 | 64164623 | G | T | 0.903879 | 8.08E-15 | 3.17E-09 |
| 11 | 64164643 | C | T | 0.903879 | 8.08E-15 | 3.17E-09 |
| 11 | 64164662 | C | T | 0.903879 | 8.08E-15 | 3.17E-09 |
| 11 | 64164663 | A | G | 0.903879 | 8.08E-15 | 3.17E-09 |
| 11 | 64164688 | G | A | 0.903879 | 8.08E-15 | 3.17E-09 |
| 11 | 64164994 | C | T | 0.899663 | 8.56E-15 | 3.36E-09 |
| 11 | 64165010 | C | A | 0.899663 | 8.56E-15 | 3.36E-09 |
| 11 | 64165074 | C | T | 0.899663 | 8.56E-15 | 3.36E-09 |
| 11 | 63768563 | A | G | 0.89882 | 1.34E-14 | 5.26E-09 |
| 11 | 63768663 | C | T | 0.89882 | 1.34E-14 | 5.26E-09 |
| 11 | 64132623 | G | A | 0.903879 | 1.63E-14 | 6.38E-09 |
| 11 | 63913520 | A | T | 0.868465 | 1.70E-14 | 6.65E-09 |
| 11 | 63913562 | C | A | 0.868465 | 1.70E-14 | 6.65E-09 |
| 11 | 63913584 | T | C | 0.868465 | 1.70E-14 | 6.65E-09 |
| 11 | 63913593 | A | T | 0.868465 | 1.70E-14 | 6.65E-09 |
| 11 | 63913607 | G | A | 0.868465 | 1.70E-14 | 6.65E-09 |
| 11 | 63913613 | A | G | 0.868465 | 1.70E-14 | 6.65E-09 |
| 11 | 63913632 | A | G | 0.868465 | 1.70E-14 | 6.65E-09 |
| 11 | 63913682 | G | T | 0.868465 | 1.70E-14 | 6.65E-09 |
| 11 | 63913749 | C | G | 0.868465 | 1.70E-14 | 6.65E-09 |
| 11 | 63913753 | T | G | 0.868465 | 1.70E-14 | 6.65E-09 |
| 11 | 63913761 | G | A | 0.868465 | 1.70E-14 | 6.65E-09 |
| 11 | 63202868 | A | T | 0.883642 | 1.85E-14 | 7.25E-09 |
| 11 | 63352244 | C | T | 0.940978 | 2.18E-14 | 8.53E-09 |
| 11 | 64165094 | G | A | 0.897976 | 2.47E-14 | 9.67E-09 |
| 11 | 64678870 | A | C | 0.872681 | 3.27E-14 | 1.28E-08 |
| 11 | 64678871 | A | C | 0.872681 | 3.27E-14 | 1.28E-08 |
| 11 | 64169440 | A | G | 0.900506 | 3.63E-14 | 1.42E-08 |
| 11 | 64169473 | T | A | 0.900506 | 3.63E-14 | 1.42E-08 |
| 11 | 64169622 | T | C | 0.900506 | 3.63E-14 | 1.42E-08 |
| 11 | 63829594 | A | G | 0.887015 | 3.83E-14 | 1.50E-08 |
| 11 | 63829641 | G | A | 0.887015 | 3.83E-14 | 1.50E-08 |
| 11 | 64795812 | T | G | 0.867622 | 3.89E-14 | 1.52E-08 |
| 11 | 64795855 | A | G | 0.867622 | 3.89E-14 | 1.52E-08 |
| 11 | 64554057 | C | A | 0.869309 | 4.42E-14 | 1.73E-08 |
| 11 | 64554656 | C | G | 0.869309 | 4.42E-14 | 1.73E-08 |
| 11 | 64555328 | A | G | 0.869309 | 4.42E-14 | 1.73E-08 |
| 11 | 64555408 | T | A | 0.869309 | 4.42E-14 | 1.73E-08 |
| 11 | 64556940 | C | A | 0.869309 | 4.42E-14 | 1.73E-08 |
| 11 | 64556948 | C | T | 0.869309 | 4.42E-14 | 1.73E-08 |
| 11 | 64558050 | A | T | 0.869309 | 4.42E-14 | 1.73E-08 |
| 11 | 64559198 | G | T | 0.869309 | 4.42E-14 | 1.73E-08 |
| 11 | 64560245 | C | G | 0.869309 | 4.42E-14 | 1.73E-08 |
| 11 | 64560517 | T | A | 0.869309 | 4.42E-14 | 1.73E-08 |
| 11 | 64561000 | A | T | 0.869309 | 4.42E-14 | 1.73E-08 |
| 11 | 64561361 | A | G | 0.869309 | 4.42E-14 | 1.73E-08 |
| 11 | 64562828 | A | G | 0.869309 | 4.42E-14 | 1.73E-08 |
| 11 | 63193930 | A | G | 0.885329 | 4.82E-14 | 1.89E-08 |
| 11 | 64551846 | T | C | 0.867622 | 4.94E-14 | 1.93E-08 |

| | | | | | | |
|---|---|---|---|---|---|---|
| 11 | 64552222 | A | T | 0.867622 | 4.94E-14 | 1.93E-08 |
| 11 | 64553205 | C | A | 0.867622 | 4.94E-14 | 1.93E-08 |
| 11 | 64169862 | G | A | 0.899663 | 5.63E-14 | 2.21E-08 |
| 11 | 63830102 | A | G | 0.885329 | 6.94E-14 | 2.72E-08 |
| 11 | 64566419 | C | T | 0.869309 | 6.99E-14 | 2.74E-08 |
| 11 | 64566482 | T | C | 0.869309 | 6.99E-14 | 2.74E-08 |
| 11 | 64567119 | A | G | 0.869309 | 6.99E-14 | 2.74E-08 |
| 11 | 63913784 | A | T | 0.869309 | 7.11E-14 | 2.79E-08 |
| 11 | 63913790 | C | G | 0.869309 | 7.11E-14 | 2.79E-08 |
| 11 | 63913796 | T | C | 0.869309 | 7.11E-14 | 2.79E-08 |
| 11 | 61510568 | C | A | 0.921585 | 7.16E-14 | 2.81E-08 |
| 11 | 63913872 | A | C | 0.870152 | 8.31E-14 | 3.26E-08 |
| 11 | 64136783 | A | C | 0.89882 | 8.72E-14 | 3.42E-08 |
| 11 | 64685212 | C | A | 0.869309 | 9.14E-14 | 3.58E-08 |
| 11 | 64685566 | T | C | 0.869309 | 9.14E-14 | 3.58E-08 |
| 11 | 64685820 | T | C | 0.869309 | 9.14E-14 | 3.58E-08 |
| 11 | 64681741 | A | G | 0.869309 | 9.20E-14 | 3.61E-08 |
| 11 | 64681903 | A | C | 0.869309 | 9.20E-14 | 3.61E-08 |
| 11 | 64681943 | A | C | 0.869309 | 9.20E-14 | 3.61E-08 |
| 11 | 64575845 | A | G | 0.870995 | 1.02E-13 | 3.99E-08 |
| 11 | 64578100 | A | G | 0.870995 | 1.02E-13 | 3.99E-08 |
| 11 | 64578112 | T | G | 0.870995 | 1.02E-13 | 3.99E-08 |
| 11 | 64553967 | A | G | 0.870995 | 1.18E-13 | 4.62E-08 |
| 11 | 64554003 | G | C | 0.870995 | 1.18E-13 | 4.62E-08 |
| 11 | 64640880 | T | G | 0.867622 | 1.18E-13 | 4.63E-08 |
| 11 | 64641198 | A | C | 0.867622 | 1.18E-13 | 4.63E-08 |
| 11 | 64641200 | A | T | 0.867622 | 1.18E-13 | 4.63E-08 |
| 11 | 63195312 | A | G | 0.884486 | 1.25E-13 | 4.89E-08 |
| 11 | 61493008 | A | C | 0.920742 | 1.30E-13 | 5.08E-08 |
| 11 | 61493010 | A | C | 0.920742 | 1.30E-13 | 5.08E-08 |
| 11 | 61493012 | A | C | 0.920742 | 1.30E-13 | 5.08E-08 |
| 11 | 61502698 | T | G | 0.920742 | 1.30E-13 | 5.08E-08 |
| 11 | 61504250 | T | G | 0.920742 | 1.30E-13 | 5.08E-08 |
| 11 | 61505761 | A | G | 0.920742 | 1.30E-13 | 5.08E-08 |
| 11 | 61506273 | T | G | 0.920742 | 1.30E-13 | 5.08E-08 |
| 11 | 61508231 | A | G | 0.920742 | 1.30E-13 | 5.08E-08 |
| 11 | 61508232 | A | G | 0.920742 | 1.30E-13 | 5.08E-08 |
| 11 | 61508375 | T | C | 0.920742 | 1.30E-13 | 5.08E-08 |
| 11 | 61508480 | T | A | 0.920742 | 1.30E-13 | 5.08E-08 |
| 11 | 61508667 | C | G | 0.920742 | 1.30E-13 | 5.08E-08 |
| 11 | 61508753 | T | C | 0.920742 | 1.30E-13 | 5.08E-08 |
| 11 | 61508801 | A | G | 0.920742 | 1.30E-13 | 5.08E-08 |
| 11 | 61508808 | A | C | 0.920742 | 1.30E-13 | 5.08E-08 |
| 11 | 61509003 | T | C | 0.920742 | 1.30E-13 | 5.08E-08 |
| 11 | 64740523 | G | C | 0.870152 | 1.41E-13 | 5.51E-08 |
| 11 | 64741686 | T | A | 0.870152 | 1.41E-13 | 5.51E-08 |
| 11 | 64741817 | A | G | 0.870152 | 1.41E-13 | 5.51E-08 |
| 11 | 64593543 | T | G | 0.867622 | 1.42E-13 | 5.58E-08 |
| 11 | 64537375 | C | T | 0.874368 | 1.56E-13 | 6.10E-08 |
| 11 | 64537412 | T | C | 0.874368 | 1.56E-13 | 6.10E-08 |
| 11 | 64537496 | C | A | 0.874368 | 1.56E-13 | 6.10E-08 |
| 11 | 64537529 | G | T | 0.874368 | 1.56E-13 | 6.10E-08 |
| 11 | 64612744 | T | C | 0.867622 | 1.57E-13 | 6.16E-08 |
| 11 | 64611420 | A | G | 0.869309 | 1.65E-13 | 6.47E-08 |
| 11 | 64614828 | G | A | 0.869309 | 1.65E-13 | 6.47E-08 |
| 11 | 64615645 | A | C | 0.869309 | 1.65E-13 | 6.47E-08 |
| 11 | 64615859 | A | G | 0.869309 | 1.65E-13 | 6.47E-08 |
| 11 | 64616056 | T | C | 0.869309 | 1.65E-13 | 6.47E-08 |
| 11 | 64616057 | G | A | 0.869309 | 1.65E-13 | 6.47E-08 |
| 11 | 64616074 | A | T | 0.869309 | 1.65E-13 | 6.47E-08 |
| 11 | 64616075 | A | T | 0.869309 | 1.65E-13 | 6.47E-08 |
| 11 | 64617014 | T | G | 0.869309 | 1.65E-13 | 6.47E-08 |
| 11 | 64617345 | A | C | 0.869309 | 1.65E-13 | 6.47E-08 |
| 11 | 64617354 | A | C | 0.869309 | 1.65E-13 | 6.47E-08 |
| 11 | 64617369 | A | C | 0.869309 | 1.65E-13 | 6.47E-08 |
| 11 | 64617374 | A | C | 0.869309 | 1.65E-13 | 6.47E-08 |
| 11 | 64617379 | A | C | 0.869309 | 1.65E-13 | 6.47E-08 |

| | | | | | | |
|---|---|---|---|---|---|---|
| 11 | 64617589 | T | A | 0.869309 | 1.65E-13 | 6.47E-08 |
| 11 | 64617850 | G | A | 0.869309 | 1.65E-13 | 6.47E-08 |
| 11 | 64617946 | G | A | 0.869309 | 1.65E-13 | 6.47E-08 |
| 11 | 64679695 | A | G | 0.868465 | 1.74E-13 | 6.84E-08 |
| 11 | 64679783 | G | A | 0.868465 | 1.74E-13 | 6.84E-08 |
| 11 | 64679866 | A | G | 0.868465 | 1.74E-13 | 6.84E-08 |
| 11 | 64680450 | A | T | 0.868465 | 1.74E-13 | 6.84E-08 |
| 11 | 64680614 | T | C | 0.868465 | 1.74E-13 | 6.84E-08 |
| 11 | 64537679 | A | C | 0.874368 | 1.75E-13 | 6.87E-08 |
| 11 | 64528229 | A | C | 0.868465 | 1.78E-13 | 6.96E-08 |
| 11 | 64528247 | T | A | 0.868465 | 1.78E-13 | 6.96E-08 |
| 11 | 64528541 | T | G | 0.868465 | 1.78E-13 | 6.96E-08 |
| 11 | 64528546 | T | G | 0.868465 | 1.78E-13 | 6.96E-08 |
| 11 | 64529253 | T | A | 0.868465 | 1.78E-13 | 6.96E-08 |
| 11 | 64530320 | G | A | 0.868465 | 1.78E-13 | 6.96E-08 |
| 11 | 64531039 | G | A | 0.868465 | 1.78E-13 | 6.96E-08 |
| 11 | 64531492 | A | C | 0.868465 | 1.78E-13 | 6.96E-08 |
| 11 | 64531495 | C | T | 0.868465 | 1.78E-13 | 6.96E-08 |
| 11 | 64531776 | A | C | 0.868465 | 1.78E-13 | 6.96E-08 |
| 11 | 64532075 | C | T | 0.868465 | 1.78E-13 | 6.96E-08 |
| 11 | 64532266 | G | A | 0.868465 | 1.78E-13 | 6.96E-08 |
| 11 | 64532387 | C | G | 0.868465 | 1.78E-13 | 6.96E-08 |
| 11 | 64533319 | A | G | 0.868465 | 1.78E-13 | 6.96E-08 |
| 11 | 64549995 | T | C | 0.868465 | 1.78E-13 | 6.96E-08 |
| 11 | 64550106 | C | G | 0.868465 | 1.78E-13 | 6.96E-08 |
| 11 | 64550856 | A | C | 0.868465 | 1.78E-13 | 6.96E-08 |
| 11 | 64550902 | T | C | 0.868465 | 1.78E-13 | 6.96E-08 |
| 11 | 64550905 | C | T | 0.868465 | 1.78E-13 | 6.96E-08 |
| 11 | 64550907 | G | C | 0.868465 | 1.78E-13 | 6.96E-08 |
| 11 | 64550925 | A | G | 0.868465 | 1.78E-13 | 6.96E-08 |
| 11 | 64550927 | C | T | 0.868465 | 1.78E-13 | 6.96E-08 |
| 11 | 64551388 | C | T | 0.868465 | 1.78E-13 | 6.96E-08 |
| 11 | 64554159 | C | T | 0.868465 | 1.78E-13 | 6.96E-08 |
| 11 | 64554311 | T | C | 0.868465 | 1.78E-13 | 6.96E-08 |
| 11 | 64554888 | T | C | 0.868465 | 1.78E-13 | 6.96E-08 |
| 11 | 64555966 | T | C | 0.868465 | 1.78E-13 | 6.96E-08 |
| 11 | 64556156 | A | C | 0.868465 | 1.78E-13 | 6.96E-08 |
| 11 | 64556284 | A | T | 0.868465 | 1.78E-13 | 6.96E-08 |
| 11 | 64556648 | A | C | 0.868465 | 1.78E-13 | 6.96E-08 |
| 11 | 64558627 | G | C | 0.868465 | 1.78E-13 | 6.96E-08 |
| 11 | 64559003 | T | G | 0.868465 | 1.78E-13 | 6.96E-08 |
| 11 | 64563368 | T | C | 0.868465 | 1.78E-13 | 6.96E-08 |
| 11 | 64563377 | A | G | 0.868465 | 1.78E-13 | 6.96E-08 |
| 11 | 64565869 | G | A | 0.868465 | 1.78E-13 | 6.96E-08 |
| 11 | 64566055 | A | T | 0.868465 | 1.78E-13 | 6.96E-08 |
| 11 | 64566336 | A | G | 0.868465 | 1.78E-13 | 6.96E-08 |
| 11 | 64569489 | T | C | 0.868465 | 1.78E-13 | 6.96E-08 |
| 11 | 64569632 | C | T | 0.868465 | 1.78E-13 | 6.96E-08 |
| 11 | 64569895 | T | C | 0.868465 | 1.78E-13 | 6.96E-08 |
| 11 | 64569913 | A | G | 0.868465 | 1.78E-13 | 6.96E-08 |
| 11 | 64570043 | T | G | 0.868465 | 1.78E-13 | 6.96E-08 |
| 11 | 64570394 | T | G | 0.868465 | 1.78E-13 | 6.96E-08 |
| 11 | 64570475 | A | T | 0.868465 | 1.78E-13 | 6.96E-08 |
| 11 | 64571522 | A | C | 0.868465 | 1.78E-13 | 6.96E-08 |
| 11 | 64571720 | A | G | 0.868465 | 1.78E-13 | 6.96E-08 |
| 11 | 64579695 | C | A | 0.868465 | 1.78E-13 | 6.96E-08 |
| 11 | 64579699 | G | A | 0.868465 | 1.78E-13 | 6.96E-08 |
| 11 | 64580209 | A | G | 0.868465 | 1.78E-13 | 6.96E-08 |
| 11 | 64581640 | A | C | 0.868465 | 1.78E-13 | 6.96E-08 |
| 11 | 64582334 | G | T | 0.868465 | 1.78E-13 | 6.96E-08 |
| 11 | 64583591 | C | T | 0.868465 | 1.78E-13 | 6.96E-08 |
| 11 | 64583612 | G | A | 0.868465 | 1.78E-13 | 6.96E-08 |
| 11 | 64583731 | T | C | 0.868465 | 1.78E-13 | 6.96E-08 |
| 11 | 64583811 | A | T | 0.868465 | 1.78E-13 | 6.96E-08 |
| 11 | 64584200 | A | C | 0.868465 | 1.78E-13 | 6.96E-08 |
| 11 | 64594265 | G | C | 0.868465 | 1.78E-13 | 6.96E-08 |
| 11 | 64594381 | T | G | 0.868465 | 1.78E-13 | 6.96E-08 |

| | | | | | | |
|---|---|---|---|---|---|---|
| 11 | 64594384 | C | T | 0.868465 | 1.78E-13 | 6.96E-08 |
| 11 | 64594449 | T | A | 0.868465 | 1.78E-13 | 6.96E-08 |
| 11 | 64594838 | A | C | 0.868465 | 1.78E-13 | 6.96E-08 |
| 11 | 64595557 | G | A | 0.868465 | 1.78E-13 | 6.96E-08 |
| 11 | 64595690 | C | A | 0.868465 | 1.78E-13 | 6.96E-08 |
| 11 | 64596442 | C | T | 0.868465 | 1.78E-13 | 6.96E-08 |
| 11 | 64596445 | C | T | 0.868465 | 1.78E-13 | 6.96E-08 |
| 11 | 64596451 | G | A | 0.868465 | 1.78E-13 | 6.96E-08 |
| 11 | 64596595 | A | C | 0.868465 | 1.78E-13 | 6.96E-08 |
| 11 | 64597449 | T | C | 0.868465 | 1.78E-13 | 6.96E-08 |
| 11 | 64597535 | T | C | 0.868465 | 1.78E-13 | 6.96E-08 |
| 11 | 64597536 | G | T | 0.868465 | 1.78E-13 | 6.96E-08 |
| 11 | 64597538 | C | T | 0.868465 | 1.78E-13 | 6.96E-08 |
| 11 | 64597544 | T | G | 0.868465 | 1.78E-13 | 6.96E-08 |
| 11 | 64597553 | A | T | 0.868465 | 1.78E-13 | 6.96E-08 |
| 11 | 64597581 | A | G | 0.868465 | 1.78E-13 | 6.96E-08 |
| 11 | 64597585 | A | C | 0.868465 | 1.78E-13 | 6.96E-08 |
| 11 | 64597754 | T | C | 0.868465 | 1.78E-13 | 6.96E-08 |
| 11 | 64597760 | C | A | 0.868465 | 1.78E-13 | 6.96E-08 |
| 11 | 64597774 | A | T | 0.868465 | 1.78E-13 | 6.96E-08 |
| 11 | 64597796 | A | C | 0.868465 | 1.78E-13 | 6.96E-08 |
| 11 | 64598577 | G | A | 0.868465 | 1.78E-13 | 6.96E-08 |
| 11 | 64598650 | T | G | 0.868465 | 1.78E-13 | 6.96E-08 |
| 11 | 64599217 | T | A | 0.868465 | 1.78E-13 | 6.96E-08 |
| 11 | 64599231 | T | G | 0.868465 | 1.78E-13 | 6.96E-08 |
| 11 | 64600158 | G | A | 0.868465 | 1.78E-13 | 6.96E-08 |
| 11 | 64600202 | C | T | 0.868465 | 1.78E-13 | 6.96E-08 |
| 11 | 64602989 | A | G | 0.868465 | 1.78E-13 | 6.96E-08 |
| 11 | 64603845 | G | T | 0.868465 | 1.78E-13 | 6.96E-08 |
| 11 | 64604389 | G | A | 0.868465 | 1.78E-13 | 6.96E-08 |
| 11 | 64605156 | A | T | 0.868465 | 1.78E-13 | 6.96E-08 |
| 11 | 64605938 | A | G | 0.868465 | 1.78E-13 | 6.96E-08 |
| 11 | 64606305 | G | A | 0.868465 | 1.78E-13 | 6.96E-08 |
| 11 | 64608220 | G | A | 0.868465 | 1.78E-13 | 6.96E-08 |
| 11 | 64608514 | G | T | 0.868465 | 1.78E-13 | 6.96E-08 |
| 11 | 64608719 | A | G | 0.868465 | 1.78E-13 | 6.96E-08 |
| 11 | 64609150 | T | G | 0.868465 | 1.78E-13 | 6.96E-08 |
| 11 | 64614451 | G | A | 0.868465 | 1.78E-13 | 6.96E-08 |
| 11 | 64614468 | A | G | 0.868465 | 1.78E-13 | 6.96E-08 |
| 11 | 64614502 | C | T | 0.868465 | 1.78E-13 | 6.96E-08 |
| 11 | 64614868 | T | C | 0.868465 | 1.78E-13 | 6.96E-08 |
| 11 | 64614915 | A | G | 0.868465 | 1.78E-13 | 6.96E-08 |
| 11 | 64614924 | A | G | 0.868465 | 1.78E-13 | 6.96E-08 |
| 11 | 64614936 | A | G | 0.868465 | 1.78E-13 | 6.96E-08 |
| 11 | 64615037 | G | A | 0.868465 | 1.78E-13 | 6.96E-08 |
| 11 | 64615050 | A | G | 0.868465 | 1.78E-13 | 6.96E-08 |
| 11 | 64615236 | T | A | 0.868465 | 1.78E-13 | 6.96E-08 |
| 11 | 64616542 | G | A | 0.868465 | 1.78E-13 | 6.96E-08 |
| 11 | 64616739 | A | G | 0.868465 | 1.78E-13 | 6.96E-08 |
| 11 | 64616781 | A | G | 0.868465 | 1.78E-13 | 6.96E-08 |
| 11 | 64616868 | A | G | 0.868465 | 1.78E-13 | 6.96E-08 |
| 11 | 64618362 | T | A | 0.868465 | 1.78E-13 | 6.96E-08 |
| 11 | 64618467 | A | C | 0.868465 | 1.78E-13 | 6.96E-08 |
| 11 | 64618557 | T | C | 0.868465 | 1.78E-13 | 6.96E-08 |
| 11 | 64618593 | T | C | 0.868465 | 1.78E-13 | 6.96E-08 |
| 11 | 64618607 | A | G | 0.868465 | 1.78E-13 | 6.96E-08 |
| 11 | 64618668 | C | T | 0.868465 | 1.78E-13 | 6.96E-08 |
| 11 | 64618672 | A | G | 0.868465 | 1.78E-13 | 6.96E-08 |
| 11 | 64618701 | T | C | 0.868465 | 1.78E-13 | 6.96E-08 |
| 11 | 64618734 | T | G | 0.868465 | 1.78E-13 | 6.96E-08 |
| 11 | 64618784 | T | C | 0.868465 | 1.78E-13 | 6.96E-08 |
| 11 | 64618951 | T | G | 0.868465 | 1.78E-13 | 6.96E-08 |
| 11 | 64618963 | C | T | 0.868465 | 1.78E-13 | 6.96E-08 |
| 11 | 64619065 | G | A | 0.868465 | 1.78E-13 | 6.96E-08 |
| 11 | 64619483 | A | T | 0.868465 | 1.78E-13 | 6.96E-08 |
| 11 | 64619579 | A | C | 0.868465 | 1.78E-13 | 6.96E-08 |
| 11 | 64619855 | T | A | 0.868465 | 1.78E-13 | 6.96E-08 |

| | | | | | | |
|---|---|---|---|---|---|---|
| 11 | 64619884 | C | G | 0.868465 | 1.78E-13 | 6.96E-08 |
| 11 | 64619982 | A | G | 0.868465 | 1.78E-13 | 6.96E-08 |
| 11 | 64620578 | A | C | 0.868465 | 1.78E-13 | 6.96E-08 |
| 11 | 64621305 | A | G | 0.868465 | 1.78E-13 | 6.96E-08 |
| 11 | 64621369 | C | T | 0.868465 | 1.78E-13 | 6.96E-08 |
| 11 | 64622724 | A | C | 0.868465 | 1.78E-13 | 6.96E-08 |
| 11 | 64623289 | T | C | 0.868465 | 1.78E-13 | 6.96E-08 |
| 11 | 64623366 | A | G | 0.868465 | 1.78E-13 | 6.96E-08 |
| 11 | 64623730 | A | G | 0.868465 | 1.78E-13 | 6.96E-08 |
| 11 | 64625066 | T | C | 0.868465 | 1.78E-13 | 6.96E-08 |
| 11 | 64626846 | A | G | 0.868465 | 1.78E-13 | 6.96E-08 |
| 11 | 64627286 | T | C | 0.868465 | 1.78E-13 | 6.96E-08 |
| 11 | 64627316 | T | A | 0.868465 | 1.78E-13 | 6.96E-08 |
| 11 | 64627555 | T | C | 0.868465 | 1.78E-13 | 6.96E-08 |
| 11 | 64627771 | A | C | 0.868465 | 1.78E-13 | 6.96E-08 |
| 11 | 64627903 | C | G | 0.868465 | 1.78E-13 | 6.96E-08 |
| 11 | 64628205 | T | A | 0.868465 | 1.78E-13 | 6.96E-08 |
| 11 | 64628299 | T | C | 0.868465 | 1.78E-13 | 6.96E-08 |
| 11 | 64631442 | A | G | 0.868465 | 1.78E-13 | 6.96E-08 |
| 11 | 64631467 | A | G | 0.868465 | 1.78E-13 | 6.96E-08 |
| 11 | 64631595 | A | G | 0.868465 | 1.78E-13 | 6.96E-08 |
| 11 | 64634289 | G | A | 0.868465 | 1.78E-13 | 6.96E-08 |
| 11 | 64634354 | T | G | 0.868465 | 1.78E-13 | 6.96E-08 |
| 11 | 64634411 | A | G | 0.868465 | 1.78E-13 | 6.96E-08 |
| 11 | 64634528 | T | G | 0.868465 | 1.78E-13 | 6.96E-08 |
| 11 | 64634603 | A | C | 0.868465 | 1.78E-13 | 6.96E-08 |
| 11 | 64634616 | C | T | 0.868465 | 1.78E-13 | 6.96E-08 |
| 11 | 64634667 | C | T | 0.868465 | 1.78E-13 | 6.96E-08 |
| 11 | 64634838 | C | T | 0.868465 | 1.78E-13 | 6.96E-08 |
| 11 | 64634957 | A | G | 0.868465 | 1.78E-13 | 6.96E-08 |
| 11 | 64635051 | G | A | 0.868465 | 1.78E-13 | 6.96E-08 |
| 11 | 64635115 | T | G | 0.868465 | 1.78E-13 | 6.96E-08 |
| 11 | 64635285 | A | T | 0.868465 | 1.78E-13 | 6.96E-08 |
| 11 | 64635312 | G | A | 0.868465 | 1.78E-13 | 6.96E-08 |
| 11 | 64635386 | T | C | 0.868465 | 1.78E-13 | 6.96E-08 |
| 11 | 64635413 | A | G | 0.868465 | 1.78E-13 | 6.96E-08 |
| 11 | 64635434 | T | C | 0.868465 | 1.78E-13 | 6.96E-08 |
| 11 | 64635641 | G | C | 0.868465 | 1.78E-13 | 6.96E-08 |
| 11 | 64635723 | T | C | 0.868465 | 1.78E-13 | 6.96E-08 |
| 11 | 64635749 | A | G | 0.868465 | 1.78E-13 | 6.96E-08 |
| 11 | 64635757 | A | G | 0.868465 | 1.78E-13 | 6.96E-08 |
| 11 | 64635783 | A | G | 0.868465 | 1.78E-13 | 6.96E-08 |
| 11 | 64635787 | T | C | 0.868465 | 1.78E-13 | 6.96E-08 |
| 11 | 64635812 | T | A | 0.868465 | 1.78E-13 | 6.96E-08 |
| 11 | 64635823 | C | T | 0.868465 | 1.78E-13 | 6.96E-08 |
| 11 | 64635938 | T | C | 0.868465 | 1.78E-13 | 6.96E-08 |
| 11 | 64635951 | G | C | 0.868465 | 1.78E-13 | 6.96E-08 |
| 11 | 64635953 | T | A | 0.868465 | 1.78E-13 | 6.96E-08 |
| 11 | 64635956 | A | G | 0.868465 | 1.78E-13 | 6.96E-08 |
| 11 | 64635964 | A | G | 0.868465 | 1.78E-13 | 6.96E-08 |
| 11 | 64636027 | T | C | 0.868465 | 1.78E-13 | 6.96E-08 |
| 11 | 64636121 | T | C | 0.868465 | 1.78E-13 | 6.96E-08 |
| 11 | 64636255 | A | G | 0.868465 | 1.78E-13 | 6.96E-08 |
| 11 | 64636341 | C | A | 0.868465 | 1.78E-13 | 6.96E-08 |
| 11 | 64636480 | G | A | 0.868465 | 1.78E-13 | 6.96E-08 |
| 11 | 64636525 | G | A | 0.868465 | 1.78E-13 | 6.96E-08 |
| 11 | 64637160 | C | G | 0.868465 | 1.78E-13 | 6.96E-08 |
| 11 | 64637197 | G | A | 0.868465 | 1.78E-13 | 6.96E-08 |
| 11 | 64637362 | A | G | 0.868465 | 1.78E-13 | 6.96E-08 |
| 11 | 64637430 | C | G | 0.868465 | 1.78E-13 | 6.96E-08 |
| 11 | 64637667 | G | A | 0.868465 | 1.78E-13 | 6.96E-08 |
| 11 | 64637709 | C | T | 0.868465 | 1.78E-13 | 6.96E-08 |
| 11 | 64637738 | A | C | 0.868465 | 1.78E-13 | 6.96E-08 |
| 11 | 64637786 | C | G | 0.868465 | 1.78E-13 | 6.96E-08 |
| 11 | 64637850 | A | G | 0.868465 | 1.78E-13 | 6.96E-08 |
| 11 | 64637879 | T | A | 0.868465 | 1.78E-13 | 6.96E-08 |
| 11 | 64638088 | C | G | 0.868465 | 1.78E-13 | 6.96E-08 |

| | | | | | | |
|---|---|---|---|---|---|---|
| 11 | 64638122 | A | G | 0.868465 | 1.78E-13 | 6.96E-08 |
| 11 | 64638142 | A | G | 0.868465 | 1.78E-13 | 6.96E-08 |
| 11 | 64638216 | G | A | 0.868465 | 1.78E-13 | 6.96E-08 |
| 11 | 64638263 | T | C | 0.868465 | 1.78E-13 | 6.96E-08 |
| 11 | 64638399 | T | G | 0.868465 | 1.78E-13 | 6.96E-08 |
| 11 | 64638411 | G | T | 0.868465 | 1.78E-13 | 6.96E-08 |
| 11 | 64638420 | G | C | 0.868465 | 1.78E-13 | 6.96E-08 |
| 11 | 64638473 | A | T | 0.868465 | 1.78E-13 | 6.96E-08 |
| 11 | 64638546 | T | A | 0.868465 | 1.78E-13 | 6.96E-08 |
| 11 | 64638551 | T | A | 0.868465 | 1.78E-13 | 6.96E-08 |
| 11 | 64638589 | C | T | 0.868465 | 1.78E-13 | 6.96E-08 |
| 11 | 64638618 | C | A | 0.868465 | 1.78E-13 | 6.96E-08 |
| 11 | 64638636 | C | T | 0.868465 | 1.78E-13 | 6.96E-08 |
| 11 | 64638653 | T | C | 0.868465 | 1.78E-13 | 6.96E-08 |
| 11 | 64638687 | A | T | 0.868465 | 1.78E-13 | 6.96E-08 |
| 11 | 64638699 | C | T | 0.868465 | 1.78E-13 | 6.96E-08 |
| 11 | 64638700 | A | G | 0.868465 | 1.78E-13 | 6.96E-08 |
| 11 | 64640805 | G | C | 0.868465 | 1.78E-13 | 6.96E-08 |
| 11 | 64640951 | G | A | 0.868465 | 1.78E-13 | 6.96E-08 |
| 11 | 64641277 | A | T | 0.868465 | 1.78E-13 | 6.96E-08 |
| 11 | 64641281 | A | G | 0.868465 | 1.78E-13 | 6.96E-08 |
| 11 | 64641282 | G | A | 0.868465 | 1.78E-13 | 6.96E-08 |
| 11 | 64641335 | T | C | 0.868465 | 1.78E-13 | 6.96E-08 |
| 11 | 64641338 | A | C | 0.868465 | 1.78E-13 | 6.96E-08 |
| 11 | 64641345 | C | T | 0.868465 | 1.78E-13 | 6.96E-08 |
| 11 | 64641444 | G | C | 0.868465 | 1.78E-13 | 6.96E-08 |
| 11 | 64641449 | C | G | 0.868465 | 1.78E-13 | 6.96E-08 |
| 11 | 64641516 | C | T | 0.868465 | 1.78E-13 | 6.96E-08 |
| 11 | 64641534 | C | T | 0.868465 | 1.78E-13 | 6.96E-08 |
| 11 | 64641607 | T | C | 0.868465 | 1.78E-13 | 6.96E-08 |
| 11 | 64641640 | A | C | 0.868465 | 1.78E-13 | 6.96E-08 |
| 11 | 64641685 | T | G | 0.868465 | 1.78E-13 | 6.96E-08 |
| 11 | 64641692 | G | T | 0.868465 | 1.78E-13 | 6.96E-08 |
| 11 | 64641705 | C | T | 0.868465 | 1.78E-13 | 6.96E-08 |
| 11 | 64641743 | T | G | 0.868465 | 1.78E-13 | 6.96E-08 |
| 11 | 64641792 | G | A | 0.868465 | 1.78E-13 | 6.96E-08 |
| 11 | 64641887 | G | C | 0.868465 | 1.78E-13 | 6.96E-08 |
| 11 | 64641943 | C | T | 0.868465 | 1.78E-13 | 6.96E-08 |
| 11 | 64642059 | G | A | 0.868465 | 1.78E-13 | 6.96E-08 |
| 11 | 64642063 | T | C | 0.868465 | 1.78E-13 | 6.96E-08 |
| 11 | 64642076 | G | A | 0.868465 | 1.78E-13 | 6.96E-08 |
| 11 | 64642114 | A | C | 0.868465 | 1.78E-13 | 6.96E-08 |
| 11 | 64642149 | C | A | 0.868465 | 1.78E-13 | 6.96E-08 |
| 11 | 64642162 | T | A | 0.868465 | 1.78E-13 | 6.96E-08 |
| 11 | 64642250 | T | G | 0.868465 | 1.78E-13 | 6.96E-08 |
| 11 | 64642274 | G | A | 0.868465 | 1.78E-13 | 6.96E-08 |
| 11 | 64642331 | A | G | 0.868465 | 1.78E-13 | 6.96E-08 |
| 11 | 64642449 | G | A | 0.868465 | 1.78E-13 | 6.96E-08 |
| 11 | 64642613 | A | C | 0.868465 | 1.78E-13 | 6.96E-08 |
| 11 | 64642653 | A | G | 0.868465 | 1.78E-13 | 6.96E-08 |
| 11 | 64642704 | T | C | 0.868465 | 1.78E-13 | 6.96E-08 |
| 11 | 64642725 | C | T | 0.868465 | 1.78E-13 | 6.96E-08 |
| 11 | 64642906 | T | C | 0.868465 | 1.78E-13 | 6.96E-08 |
| 11 | 64642956 | G | T | 0.868465 | 1.78E-13 | 6.96E-08 |
| 11 | 64643067 | C | T | 0.868465 | 1.78E-13 | 6.96E-08 |
| 11 | 64643094 | A | G | 0.868465 | 1.78E-13 | 6.96E-08 |
| 11 | 64643186 | G | A | 0.868465 | 1.78E-13 | 6.96E-08 |
| 11 | 64643347 | A | G | 0.868465 | 1.78E-13 | 6.96E-08 |
| 11 | 64643478 | T | A | 0.868465 | 1.78E-13 | 6.96E-08 |
| 11 | 64643665 | G | T | 0.868465 | 1.78E-13 | 6.96E-08 |
| 11 | 64643984 | G | A | 0.868465 | 1.78E-13 | 6.96E-08 |
| 11 | 64644001 | A | G | 0.868465 | 1.78E-13 | 6.96E-08 |
| 11 | 64644044 | T | C | 0.868465 | 1.78E-13 | 6.96E-08 |
| 11 | 64644135 | A | G | 0.868465 | 1.78E-13 | 6.96E-08 |
| 11 | 64644392 | A | C | 0.868465 | 1.78E-13 | 6.96E-08 |
| 11 | 64644449 | T | C | 0.868465 | 1.78E-13 | 6.96E-08 |
| 11 | 64644450 | G | T | 0.868465 | 1.78E-13 | 6.96E-08 |

| | | | | | | |
|---|---|---|---|---|---|---|
| 11 | 64644518 | T | C | 0.868465 | 1.78E-13 | 6.96E-08 |
| 11 | 64644609 | C | T | 0.868465 | 1.78E-13 | 6.96E-08 |
| 11 | 64644658 | C | A | 0.868465 | 1.78E-13 | 6.96E-08 |
| 11 | 64644791 | T | C | 0.868465 | 1.78E-13 | 6.96E-08 |
| 11 | 64644801 | G | T | 0.868465 | 1.78E-13 | 6.96E-08 |
| 11 | 64644813 | C | T | 0.868465 | 1.78E-13 | 6.96E-08 |
| 11 | 64644819 | G | A | 0.868465 | 1.78E-13 | 6.96E-08 |
| 11 | 64644831 | A | G | 0.868465 | 1.78E-13 | 6.96E-08 |
| 11 | 64644833 | C | T | 0.868465 | 1.78E-13 | 6.96E-08 |
| 11 | 64644834 | G | A | 0.868465 | 1.78E-13 | 6.96E-08 |
| 11 | 64644898 | A | C | 0.868465 | 1.78E-13 | 6.96E-08 |
| 11 | 64645387 | T | C | 0.868465 | 1.78E-13 | 6.96E-08 |
| 11 | 64645441 | T | C | 0.868465 | 1.78E-13 | 6.96E-08 |
| 11 | 64645444 | C | T | 0.868465 | 1.78E-13 | 6.96E-08 |
| 11 | 64645638 | A | G | 0.868465 | 1.78E-13 | 6.96E-08 |
| 11 | 64645760 | A | G | 0.868465 | 1.78E-13 | 6.96E-08 |
| 11 | 64645800 | C | T | 0.868465 | 1.78E-13 | 6.96E-08 |
| 11 | 64645917 | T | C | 0.868465 | 1.78E-13 | 6.96E-08 |
| 11 | 64645937 | T | C | 0.868465 | 1.78E-13 | 6.96E-08 |
| 11 | 64646144 | A | T | 0.868465 | 1.78E-13 | 6.96E-08 |
| 11 | 64646188 | G | A | 0.868465 | 1.78E-13 | 6.96E-08 |
| 11 | 64646355 | C | A | 0.868465 | 1.78E-13 | 6.96E-08 |
| 11 | 64646523 | C | T | 0.868465 | 1.78E-13 | 6.96E-08 |
| 11 | 64646558 | G | A | 0.868465 | 1.78E-13 | 6.96E-08 |
| 11 | 64646852 | T | A | 0.868465 | 1.78E-13 | 6.96E-08 |
| 11 | 64646991 | C | G | 0.868465 | 1.78E-13 | 6.96E-08 |
| 11 | 64647026 | C | A | 0.868465 | 1.78E-13 | 6.96E-08 |
| 11 | 64647106 | T | A | 0.868465 | 1.78E-13 | 6.96E-08 |
| 11 | 64647138 | T | C | 0.868465 | 1.78E-13 | 6.96E-08 |
| 11 | 64647266 | T | C | 0.868465 | 1.78E-13 | 6.96E-08 |
| 11 | 64647368 | G | A | 0.868465 | 1.78E-13 | 6.96E-08 |
| 11 | 64647498 | T | C | 0.868465 | 1.78E-13 | 6.96E-08 |
| 11 | 64647681 | G | A | 0.868465 | 1.78E-13 | 6.96E-08 |
| 11 | 64647923 | C | T | 0.868465 | 1.78E-13 | 6.96E-08 |
| 11 | 64647943 | A | G | 0.868465 | 1.78E-13 | 6.96E-08 |
| 11 | 64647955 | A | C | 0.868465 | 1.78E-13 | 6.96E-08 |
| 11 | 64648110 | T | C | 0.868465 | 1.78E-13 | 6.96E-08 |
| 11 | 64648139 | C | G | 0.868465 | 1.78E-13 | 6.96E-08 |
| 11 | 64648178 | G | T | 0.868465 | 1.78E-13 | 6.96E-08 |
| 11 | 64648236 | A | G | 0.868465 | 1.78E-13 | 6.96E-08 |
| 11 | 64648335 | A | C | 0.868465 | 1.78E-13 | 6.96E-08 |
| 11 | 64648402 | G | A | 0.868465 | 1.78E-13 | 6.96E-08 |
| 11 | 64648490 | T | A | 0.868465 | 1.78E-13 | 6.96E-08 |
| 11 | 64648541 | G | A | 0.868465 | 1.78E-13 | 6.96E-08 |
| 11 | 64648588 | T | C | 0.868465 | 1.78E-13 | 6.96E-08 |
| 11 | 64648617 | A | G | 0.868465 | 1.78E-13 | 6.96E-08 |
| 11 | 64648620 | C | T | 0.868465 | 1.78E-13 | 6.96E-08 |
| 11 | 64648636 | A | G | 0.868465 | 1.78E-13 | 6.96E-08 |
| 11 | 64648653 | G | A | 0.868465 | 1.78E-13 | 6.96E-08 |
| 11 | 64648698 | A | G | 0.868465 | 1.78E-13 | 6.96E-08 |
| 11 | 64648839 | T | A | 0.868465 | 1.78E-13 | 6.96E-08 |
| 11 | 64648905 | A | G | 0.868465 | 1.78E-13 | 6.96E-08 |
| 11 | 64648979 | A | T | 0.868465 | 1.78E-13 | 6.96E-08 |
| 11 | 64648985 | A | C | 0.868465 | 1.78E-13 | 6.96E-08 |
| 11 | 64649090 | T | C | 0.868465 | 1.78E-13 | 6.96E-08 |
| 11 | 64649134 | C | A | 0.868465 | 1.78E-13 | 6.96E-08 |
| 11 | 64649268 | G | C | 0.868465 | 1.78E-13 | 6.96E-08 |
| 11 | 64649311 | C | T | 0.868465 | 1.78E-13 | 6.96E-08 |
| 11 | 64649402 | C | T | 0.868465 | 1.78E-13 | 6.96E-08 |
| 11 | 64649495 | G | A | 0.868465 | 1.78E-13 | 6.96E-08 |
| 11 | 64649517 | T | C | 0.868465 | 1.78E-13 | 6.96E-08 |
| 11 | 64649518 | G | A | 0.868465 | 1.78E-13 | 6.96E-08 |
| 11 | 64649530 | G | A | 0.868465 | 1.78E-13 | 6.96E-08 |
| 11 | 64649547 | G | T | 0.868465 | 1.78E-13 | 6.96E-08 |
| 11 | 64649660 | C | T | 0.868465 | 1.78E-13 | 6.96E-08 |
| 11 | 64649741 | T | A | 0.868465 | 1.78E-13 | 6.96E-08 |
| 11 | 64649937 | A | C | 0.868465 | 1.78E-13 | 6.96E-08 |

| | | | | | | |
|---|---|---|---|---|---|---|
| 11 | 64650017 | G | A | 0.868465 | 1.78E-13 | 6.96E-08 |
| 11 | 64651350 | A | G | 0.868465 | 1.78E-13 | 6.96E-08 |
| 11 | 64651499 | T | C | 0.868465 | 1.78E-13 | 6.96E-08 |
| 11 | 64652915 | C | T | 0.868465 | 1.78E-13 | 6.96E-08 |
| 11 | 64653065 | G | A | 0.868465 | 1.78E-13 | 6.96E-08 |
| 11 | 64653146 | T | C | 0.868465 | 1.78E-13 | 6.96E-08 |
| 11 | 64653214 | C | T | 0.868465 | 1.78E-13 | 6.96E-08 |
| 11 | 64653443 | C | A | 0.868465 | 1.78E-13 | 6.96E-08 |
| 11 | 64653446 | T | A | 0.868465 | 1.78E-13 | 6.96E-08 |
| 11 | 64653674 | C | T | 0.868465 | 1.78E-13 | 6.96E-08 |
| 11 | 64653678 | T | C | 0.868465 | 1.78E-13 | 6.96E-08 |
| 11 | 64653707 | A | G | 0.868465 | 1.78E-13 | 6.96E-08 |
| 11 | 64653806 | A | G | 0.868465 | 1.78E-13 | 6.96E-08 |
| 11 | 64654349 | G | A | 0.868465 | 1.78E-13 | 6.96E-08 |
| 11 | 64654359 | C | G | 0.868465 | 1.78E-13 | 6.96E-08 |
| 11 | 64654536 | G | A | 0.868465 | 1.78E-13 | 6.96E-08 |
| 11 | 64654584 | G | A | 0.868465 | 1.78E-13 | 6.96E-08 |
| 11 | 64654613 | A | G | 0.868465 | 1.78E-13 | 6.96E-08 |
| 11 | 64654730 | C | T | 0.868465 | 1.78E-13 | 6.96E-08 |
| 11 | 64654756 | T | C | 0.868465 | 1.78E-13 | 6.96E-08 |
| 11 | 64654842 | T | C | 0.868465 | 1.78E-13 | 6.96E-08 |
| 11 | 64655214 | A | G | 0.868465 | 1.78E-13 | 6.96E-08 |
| 11 | 64655274 | T | C | 0.868465 | 1.78E-13 | 6.96E-08 |
| 11 | 64655280 | C | T | 0.868465 | 1.78E-13 | 6.96E-08 |
| 11 | 64655407 | G | A | 0.868465 | 1.78E-13 | 6.96E-08 |
| 11 | 64655462 | A | G | 0.868465 | 1.78E-13 | 6.96E-08 |
| 11 | 64655836 | C | T | 0.868465 | 1.78E-13 | 6.96E-08 |
| 11 | 64656472 | T | C | 0.868465 | 1.78E-13 | 6.96E-08 |
| 11 | 64657046 | C | T | 0.868465 | 1.78E-13 | 6.96E-08 |
| 11 | 64657717 | G | T | 0.868465 | 1.78E-13 | 6.96E-08 |
| 11 | 64658207 | T | A | 0.868465 | 1.78E-13 | 6.96E-08 |
| 11 | 64666281 | C | T | 0.868465 | 1.78E-13 | 6.96E-08 |
| 11 | 64666393 | C | T | 0.868465 | 1.78E-13 | 6.96E-08 |
| 11 | 64666472 | C | T | 0.868465 | 1.78E-13 | 6.96E-08 |
| 11 | 64666483 | G | A | 0.868465 | 1.78E-13 | 6.96E-08 |
| 11 | 64666524 | C | T | 0.868465 | 1.78E-13 | 6.96E-08 |
| 11 | 64666657 | C | T | 0.868465 | 1.78E-13 | 6.96E-08 |
| 11 | 64666870 | A | G | 0.868465 | 1.78E-13 | 6.96E-08 |
| 11 | 64667154 | T | C | 0.868465 | 1.78E-13 | 6.96E-08 |
| 11 | 64667455 | A | G | 0.868465 | 1.78E-13 | 6.96E-08 |
| 11 | 64667590 | C | G | 0.868465 | 1.78E-13 | 6.96E-08 |
| 11 | 64667689 | A | C | 0.868465 | 1.78E-13 | 6.96E-08 |
| 11 | 64667699 | C | T | 0.868465 | 1.78E-13 | 6.96E-08 |
| 11 | 64667804 | T | C | 0.868465 | 1.78E-13 | 6.96E-08 |
| 11 | 64667805 | G | A | 0.868465 | 1.78E-13 | 6.96E-08 |
| 11 | 64667881 | T | G | 0.868465 | 1.78E-13 | 6.96E-08 |
| 11 | 64667912 | T | A | 0.868465 | 1.78E-13 | 6.96E-08 |
| 11 | 64667963 | G | T | 0.868465 | 1.78E-13 | 6.96E-08 |
| 11 | 64667965 | G | A | 0.868465 | 1.78E-13 | 6.96E-08 |
| 11 | 64667970 | C | T | 0.868465 | 1.78E-13 | 6.96E-08 |
| 11 | 64668014 | G | A | 0.868465 | 1.78E-13 | 6.96E-08 |
| 11 | 64668094 | C | T | 0.868465 | 1.78E-13 | 6.96E-08 |
| 11 | 64668333 | G | A | 0.868465 | 1.78E-13 | 6.96E-08 |
| 11 | 64668559 | A | T | 0.868465 | 1.78E-13 | 6.96E-08 |
| 11 | 64668812 | G | A | 0.868465 | 1.78E-13 | 6.96E-08 |
| 11 | 64668822 | A | G | 0.868465 | 1.78E-13 | 6.96E-08 |
| 11 | 64668847 | G | A | 0.868465 | 1.78E-13 | 6.96E-08 |
| 11 | 64668943 | T | A | 0.868465 | 1.78E-13 | 6.96E-08 |
| 11 | 64669016 | A | G | 0.868465 | 1.78E-13 | 6.96E-08 |
| 11 | 64669045 | C | T | 0.868465 | 1.78E-13 | 6.96E-08 |
| 11 | 64669927 | T | A | 0.868465 | 1.78E-13 | 6.96E-08 |
| 11 | 64670301 | T | C | 0.868465 | 1.78E-13 | 6.96E-08 |
| 11 | 64670877 | G | A | 0.868465 | 1.78E-13 | 6.96E-08 |
| 11 | 64671182 | G | A | 0.868465 | 1.78E-13 | 6.96E-08 |
| 11 | 64671737 | C | T | 0.868465 | 1.78E-13 | 6.96E-08 |
| 11 | 64672297 | A | G | 0.868465 | 1.78E-13 | 6.96E-08 |
| 11 | 64672989 | C | T | 0.868465 | 1.78E-13 | 6.96E-08 |

| | | | | | | |
|---|---|---|---|---|---|---|
| 11 | 64674051 | A | G | 0.868465 | 1.78E-13 | 6.96E-08 |
| 11 | 64674375 | C | A | 0.868465 | 1.78E-13 | 6.96E-08 |
| 11 | 64674778 | G | A | 0.868465 | 1.78E-13 | 6.96E-08 |
| 11 | 64675173 | A | G | 0.868465 | 1.78E-13 | 6.96E-08 |
| 11 | 64676102 | G | A | 0.868465 | 1.78E-13 | 6.96E-08 |
| 11 | 64677030 | G | A | 0.868465 | 1.78E-13 | 6.96E-08 |
| 11 | 64677274 | A | G | 0.868465 | 1.78E-13 | 6.96E-08 |
| 11 | 64684936 | T | G | 0.868465 | 1.78E-13 | 6.96E-08 |
| 11 | 64685100 | C | T | 0.868465 | 1.78E-13 | 6.96E-08 |
| 11 | 64685101 | A | G | 0.868465 | 1.78E-13 | 6.96E-08 |
| 11 | 64685103 | T | A | 0.868465 | 1.78E-13 | 6.96E-08 |
| 11 | 64685185 | A | C | 0.868465 | 1.78E-13 | 6.96E-08 |
| 11 | 64686102 | T | C | 0.868465 | 1.78E-13 | 6.96E-08 |
| 11 | 64686899 | T | C | 0.868465 | 1.78E-13 | 6.96E-08 |
| 11 | 64686937 | A | T | 0.868465 | 1.78E-13 | 6.96E-08 |
| 11 | 64687242 | C | T | 0.868465 | 1.78E-13 | 6.96E-08 |
| 11 | 64687433 | T | C | 0.868465 | 1.78E-13 | 6.96E-08 |
| 11 | 64687483 | G | T | 0.868465 | 1.78E-13 | 6.96E-08 |
| 11 | 64687522 | G | A | 0.868465 | 1.78E-13 | 6.96E-08 |
| 11 | 64687694 | A | G | 0.868465 | 1.78E-13 | 6.96E-08 |
| 11 | 64687725 | T | C | 0.868465 | 1.78E-13 | 6.96E-08 |
| 11 | 64687867 | C | G | 0.868465 | 1.78E-13 | 6.96E-08 |
| 11 | 64688063 | C | T | 0.868465 | 1.78E-13 | 6.96E-08 |
| 11 | 64688194 | G | A | 0.868465 | 1.78E-13 | 6.96E-08 |
| 11 | 64688226 | A | T | 0.868465 | 1.78E-13 | 6.96E-08 |
| 11 | 64688260 | A | G | 0.868465 | 1.78E-13 | 6.96E-08 |
| 11 | 64688280 | C | T | 0.868465 | 1.78E-13 | 6.96E-08 |
| 11 | 64688303 | C | G | 0.868465 | 1.78E-13 | 6.96E-08 |
| 11 | 64688307 | A | G | 0.868465 | 1.78E-13 | 6.96E-08 |
| 11 | 64688351 | T | C | 0.868465 | 1.78E-13 | 6.96E-08 |
| 11 | 64688352 | G | A | 0.868465 | 1.78E-13 | 6.96E-08 |
| 11 | 64688371 | T | A | 0.868465 | 1.78E-13 | 6.96E-08 |
| 11 | 64688372 | G | A | 0.868465 | 1.78E-13 | 6.96E-08 |
| 11 | 64688375 | C | T | 0.868465 | 1.78E-13 | 6.96E-08 |
| 11 | 64688395 | T | C | 0.868465 | 1.78E-13 | 6.96E-08 |
| 11 | 64688396 | G | A | 0.868465 | 1.78E-13 | 6.96E-08 |
| 11 | 64688398 | C | T | 0.868465 | 1.78E-13 | 6.96E-08 |
| 11 | 64688402 | A | G | 0.868465 | 1.78E-13 | 6.96E-08 |
| 11 | 64688413 | A | G | 0.868465 | 1.78E-13 | 6.96E-08 |
| 11 | 64688419 | G | T | 0.868465 | 1.78E-13 | 6.96E-08 |
| 11 | 64688580 | A | G | 0.868465 | 1.78E-13 | 6.96E-08 |
| 11 | 64688736 | T | C | 0.868465 | 1.78E-13 | 6.96E-08 |
| 11 | 64688785 | T | C | 0.868465 | 1.78E-13 | 6.96E-08 |
| 11 | 64688844 | T | C | 0.868465 | 1.78E-13 | 6.96E-08 |
| 11 | 64688893 | A | G | 0.868465 | 1.78E-13 | 6.96E-08 |
| 11 | 64689525 | G | C | 0.868465 | 1.78E-13 | 6.96E-08 |
| 11 | 64689622 | T | C | 0.868465 | 1.78E-13 | 6.96E-08 |
| 11 | 64690518 | G | A | 0.868465 | 1.78E-13 | 6.96E-08 |
| 11 | 64690564 | A | G | 0.868465 | 1.78E-13 | 6.96E-08 |
| 11 | 64690698 | T | G | 0.868465 | 1.78E-13 | 6.96E-08 |
| 11 | 64690783 | T | C | 0.868465 | 1.78E-13 | 6.96E-08 |
| 11 | 64691118 | A | G | 0.868465 | 1.78E-13 | 6.96E-08 |
| 11 | 64691153 | T | C | 0.868465 | 1.78E-13 | 6.96E-08 |
| 11 | 64691184 | T | C | 0.868465 | 1.78E-13 | 6.96E-08 |
| 11 | 64691217 | C | G | 0.868465 | 1.78E-13 | 6.96E-08 |
| 11 | 64691372 | T | C | 0.868465 | 1.78E-13 | 6.96E-08 |
| 11 | 64691598 | A | C | 0.868465 | 1.78E-13 | 6.96E-08 |
| 11 | 64692002 | C | T | 0.868465 | 1.78E-13 | 6.96E-08 |
| 11 | 64692461 | C | T | 0.868465 | 1.78E-13 | 6.96E-08 |
| 11 | 64692567 | A | G | 0.868465 | 1.78E-13 | 6.96E-08 |
| 11 | 64693099 | C | T | 0.868465 | 1.78E-13 | 6.96E-08 |
| 11 | 64693391 | C | T | 0.868465 | 1.78E-13 | 6.96E-08 |
| 11 | 64693434 | G | A | 0.868465 | 1.78E-13 | 6.96E-08 |
| 11 | 64693639 | A | G | 0.868465 | 1.78E-13 | 6.96E-08 |
| 11 | 64693826 | G | T | 0.868465 | 1.78E-13 | 6.96E-08 |
| 11 | 64693862 | T | A | 0.868465 | 1.78E-13 | 6.96E-08 |
| 11 | 64693874 | A | G | 0.868465 | 1.78E-13 | 6.96E-08 |

| | | | | | | |
|---|---|---|---|---|---|---|
| 11 | 64693875 | T | G | 0.868465 | 1.78E-13 | 6.96E-08 |
| 11 | 64693958 | T | A | 0.868465 | 1.78E-13 | 6.96E-08 |
| 11 | 64693982 | A | G | 0.868465 | 1.78E-13 | 6.96E-08 |
| 11 | 64693986 | A | G | 0.868465 | 1.78E-13 | 6.96E-08 |
| 11 | 64694022 | T | C | 0.868465 | 1.78E-13 | 6.96E-08 |
| 11 | 64694137 | G | T | 0.868465 | 1.78E-13 | 6.96E-08 |
| 11 | 64694366 | T | C | 0.868465 | 1.78E-13 | 6.96E-08 |
| 11 | 64694442 | A | G | 0.868465 | 1.78E-13 | 6.96E-08 |
| 11 | 64694478 | C | G | 0.868465 | 1.78E-13 | 6.96E-08 |
| 11 | 64694579 | G | A | 0.868465 | 1.78E-13 | 6.96E-08 |
| 11 | 64694680 | C | G | 0.868465 | 1.78E-13 | 6.96E-08 |
| 11 | 64694698 | C | T | 0.868465 | 1.78E-13 | 6.96E-08 |
| 11 | 64694837 | T | C | 0.868465 | 1.78E-13 | 6.96E-08 |
| 11 | 64694888 | G | C | 0.868465 | 1.78E-13 | 6.96E-08 |
| 11 | 64694911 | C | T | 0.868465 | 1.78E-13 | 6.96E-08 |
| 11 | 64694914 | C | T | 0.868465 | 1.78E-13 | 6.96E-08 |
| 11 | 64695292 | C | T | 0.868465 | 1.78E-13 | 6.96E-08 |
| 11 | 64695372 | C | T | 0.868465 | 1.78E-13 | 6.96E-08 |
| 11 | 64695591 | G | C | 0.868465 | 1.78E-13 | 6.96E-08 |
| 11 | 64695983 | A | T | 0.868465 | 1.78E-13 | 6.96E-08 |
| 11 | 64695984 | T | A | 0.868465 | 1.78E-13 | 6.96E-08 |
| 11 | 64696053 | A | G | 0.868465 | 1.78E-13 | 6.96E-08 |
| 11 | 64696137 | C | T | 0.868465 | 1.78E-13 | 6.96E-08 |
| 11 | 64696222 | C | A | 0.868465 | 1.78E-13 | 6.96E-08 |
| 11 | 64696238 | T | C | 0.868465 | 1.78E-13 | 6.96E-08 |
| 11 | 64696239 | G | A | 0.868465 | 1.78E-13 | 6.96E-08 |
| 11 | 64696249 | A | G | 0.868465 | 1.78E-13 | 6.96E-08 |
| 11 | 64696355 | T | G | 0.868465 | 1.78E-13 | 6.96E-08 |
| 11 | 64696414 | A | G | 0.868465 | 1.78E-13 | 6.96E-08 |
| 11 | 64696589 | T | C | 0.868465 | 1.78E-13 | 6.96E-08 |
| 11 | 64696590 | C | T | 0.868465 | 1.78E-13 | 6.96E-08 |
| 11 | 64696658 | T | G | 0.868465 | 1.78E-13 | 6.96E-08 |
| 11 | 64696685 | A | G | 0.868465 | 1.78E-13 | 6.96E-08 |
| 11 | 64696758 | C | T | 0.868465 | 1.78E-13 | 6.96E-08 |
| 11 | 64696933 | C | T | 0.868465 | 1.78E-13 | 6.96E-08 |
| 11 | 64698370 | T | A | 0.868465 | 1.78E-13 | 6.96E-08 |
| 11 | 64698491 | G | C | 0.868465 | 1.78E-13 | 6.96E-08 |
| 11 | 64698533 | A | G | 0.868465 | 1.78E-13 | 6.96E-08 |
| 11 | 64698788 | A | G | 0.868465 | 1.78E-13 | 6.96E-08 |
| 11 | 64698808 | A | G | 0.868465 | 1.78E-13 | 6.96E-08 |
| 11 | 64698843 | G | A | 0.868465 | 1.78E-13 | 6.96E-08 |
| 11 | 64698873 | T | G | 0.868465 | 1.78E-13 | 6.96E-08 |
| 11 | 64699026 | C | T | 0.868465 | 1.78E-13 | 6.96E-08 |
| 11 | 64700064 | C | T | 0.868465 | 1.78E-13 | 6.96E-08 |
| 11 | 64700380 | A | C | 0.868465 | 1.78E-13 | 6.96E-08 |
| 11 | 64701331 | A | G | 0.868465 | 1.78E-13 | 6.96E-08 |
| 11 | 64701332 | C | T | 0.868465 | 1.78E-13 | 6.96E-08 |
| 11 | 64701353 | T | C | 0.868465 | 1.78E-13 | 6.96E-08 |
| 11 | 64701379 | T | C | 0.868465 | 1.78E-13 | 6.96E-08 |
| 11 | 64701438 | T | C | 0.868465 | 1.78E-13 | 6.96E-08 |
| 11 | 64701530 | C | G | 0.868465 | 1.78E-13 | 6.96E-08 |
| 11 | 64701665 | C | T | 0.868465 | 1.78E-13 | 6.96E-08 |
| 11 | 64702111 | A | G | 0.868465 | 1.78E-13 | 6.96E-08 |
| 11 | 64702321 | T | G | 0.868465 | 1.78E-13 | 6.96E-08 |
| 11 | 64702654 | T | C | 0.868465 | 1.78E-13 | 6.96E-08 |
| 11 | 64702660 | C | T | 0.868465 | 1.78E-13 | 6.96E-08 |
| 11 | 64702917 | T | C | 0.868465 | 1.78E-13 | 6.96E-08 |
| 11 | 64703025 | T | A | 0.868465 | 1.78E-13 | 6.96E-08 |
| 11 | 64703026 | G | T | 0.868465 | 1.78E-13 | 6.96E-08 |
| 11 | 64703062 | C | T | 0.868465 | 1.78E-13 | 6.96E-08 |
| 11 | 64703071 | T | A | 0.868465 | 1.78E-13 | 6.96E-08 |
| 11 | 64703078 | C | T | 0.868465 | 1.78E-13 | 6.96E-08 |
| 11 | 64703098 | G | A | 0.868465 | 1.78E-13 | 6.96E-08 |
| 11 | 64703124 | G | A | 0.868465 | 1.78E-13 | 6.96E-08 |
| 11 | 64703160 | G | A | 0.868465 | 1.78E-13 | 6.96E-08 |
| 11 | 64703214 | C | A | 0.868465 | 1.78E-13 | 6.96E-08 |
| 11 | 64703260 | T | C | 0.868465 | 1.78E-13 | 6.96E-08 |

| | | | | | | |
|---|---|---|---|---|---|---|
| 11 | 64703363 | G | T | 0.868465 | 1.78E-13 | 6.96E-08 |
| 11 | 64703372 | G | A | 0.868465 | 1.78E-13 | 6.96E-08 |
| 11 | 64703375 | T | C | 0.868465 | 1.78E-13 | 6.96E-08 |
| 11 | 64703515 | G | T | 0.868465 | 1.78E-13 | 6.96E-08 |
| 11 | 64703586 | C | A | 0.868465 | 1.78E-13 | 6.96E-08 |
| 11 | 64703592 | T | C | 0.868465 | 1.78E-13 | 6.96E-08 |
| 11 | 64703628 | T | C | 0.868465 | 1.78E-13 | 6.96E-08 |
| 11 | 64703654 | G | T | 0.868465 | 1.78E-13 | 6.96E-08 |
| 11 | 64703677 | C | A | 0.868465 | 1.78E-13 | 6.96E-08 |
| 11 | 64703698 | T | C | 0.868465 | 1.78E-13 | 6.96E-08 |
| 11 | 64703733 | G | A | 0.868465 | 1.78E-13 | 6.96E-08 |
| 11 | 64703831 | A | G | 0.868465 | 1.78E-13 | 6.96E-08 |
| 11 | 64703833 | G | A | 0.868465 | 1.78E-13 | 6.96E-08 |
| 11 | 64703847 | A | G | 0.868465 | 1.78E-13 | 6.96E-08 |
| 11 | 64703932 | G | A | 0.868465 | 1.78E-13 | 6.96E-08 |
| 11 | 64703997 | C | T | 0.868465 | 1.78E-13 | 6.96E-08 |
| 11 | 64704103 | A | G | 0.868465 | 1.78E-13 | 6.96E-08 |
| 11 | 64704311 | T | C | 0.868465 | 1.78E-13 | 6.96E-08 |
| 11 | 64704314 | C | T | 0.868465 | 1.78E-13 | 6.96E-08 |
| 11 | 64704348 | G | A | 0.868465 | 1.78E-13 | 6.96E-08 |
| 11 | 64704932 | C | T | 0.868465 | 1.78E-13 | 6.96E-08 |
| 11 | 64704933 | A | G | 0.868465 | 1.78E-13 | 6.96E-08 |
| 11 | 64705147 | C | T | 0.868465 | 1.78E-13 | 6.96E-08 |
| 11 | 64705356 | C | T | 0.868465 | 1.78E-13 | 6.96E-08 |
| 11 | 64705361 | C | T | 0.868465 | 1.78E-13 | 6.96E-08 |
| 11 | 64705370 | C | A | 0.868465 | 1.78E-13 | 6.96E-08 |
| 11 | 64705611 | T | C | 0.868465 | 1.78E-13 | 6.96E-08 |
| 11 | 64705623 | C | T | 0.868465 | 1.78E-13 | 6.96E-08 |
| 11 | 64705631 | T | C | 0.868465 | 1.78E-13 | 6.96E-08 |
| 11 | 64705640 | T | C | 0.868465 | 1.78E-13 | 6.96E-08 |
| 11 | 64705643 | T | C | 0.868465 | 1.78E-13 | 6.96E-08 |
| 11 | 64705757 | A | T | 0.868465 | 1.78E-13 | 6.96E-08 |
| 11 | 64705770 | T | C | 0.868465 | 1.78E-13 | 6.96E-08 |
| 11 | 64705773 | A | G | 0.868465 | 1.78E-13 | 6.96E-08 |
| 11 | 64705778 | G | C | 0.868465 | 1.78E-13 | 6.96E-08 |
| 11 | 64706007 | T | A | 0.868465 | 1.78E-13 | 6.96E-08 |
| 11 | 64706083 | C | T | 0.868465 | 1.78E-13 | 6.96E-08 |
| 11 | 64706145 | C | T | 0.868465 | 1.78E-13 | 6.96E-08 |
| 11 | 64706161 | C | T | 0.868465 | 1.78E-13 | 6.96E-08 |
| 11 | 64706163 | G | A | 0.868465 | 1.78E-13 | 6.96E-08 |
| 11 | 64706177 | T | C | 0.868465 | 1.78E-13 | 6.96E-08 |
| 11 | 64706186 | C | T | 0.868465 | 1.78E-13 | 6.96E-08 |
| 11 | 64706225 | A | G | 0.868465 | 1.78E-13 | 6.96E-08 |
| 11 | 64706486 | A | G | 0.868465 | 1.78E-13 | 6.96E-08 |
| 11 | 64706851 | C | T | 0.868465 | 1.78E-13 | 6.96E-08 |
| 11 | 64707332 | T | A | 0.868465 | 1.78E-13 | 6.96E-08 |
| 11 | 64707354 | C | T | 0.868465 | 1.78E-13 | 6.96E-08 |
| 11 | 64707505 | C | T | 0.868465 | 1.78E-13 | 6.96E-08 |
| 11 | 64707518 | G | T | 0.868465 | 1.78E-13 | 6.96E-08 |
| 11 | 64707534 | C | A | 0.868465 | 1.78E-13 | 6.96E-08 |
| 11 | 64707541 | T | G | 0.868465 | 1.78E-13 | 6.96E-08 |
| 11 | 64710869 | T | C | 0.868465 | 1.78E-13 | 6.96E-08 |
| 11 | 64710934 | C | G | 0.868465 | 1.78E-13 | 6.96E-08 |
| 11 | 64712878 | T | C | 0.868465 | 1.78E-13 | 6.96E-08 |
| 11 | 64713879 | A | G | 0.868465 | 1.78E-13 | 6.96E-08 |
| 11 | 64717080 | T | A | 0.868465 | 1.78E-13 | 6.96E-08 |
| 11 | 64717251 | T | A | 0.868465 | 1.78E-13 | 6.96E-08 |
| 11 | 64717254 | T | A | 0.868465 | 1.78E-13 | 6.96E-08 |
| 11 | 64717280 | A | G | 0.868465 | 1.78E-13 | 6.96E-08 |
| 11 | 64717291 | G | C | 0.868465 | 1.78E-13 | 6.96E-08 |
| 11 | 64717303 | G | T | 0.868465 | 1.78E-13 | 6.96E-08 |
| 11 | 64717394 | C | T | 0.868465 | 1.78E-13 | 6.96E-08 |
| 11 | 64717396 | C | T | 0.868465 | 1.78E-13 | 6.96E-08 |
| 11 | 64717424 | T | C | 0.868465 | 1.78E-13 | 6.96E-08 |
| 11 | 64717428 | T | G | 0.868465 | 1.78E-13 | 6.96E-08 |
| 11 | 64717430 | T | C | 0.868465 | 1.78E-13 | 6.96E-08 |
| 11 | 64717452 | A | G | 0.868465 | 1.78E-13 | 6.96E-08 |

| | | | | | | |
|---|---|---|---|---|---|---|
| 11 | 64717456 | A | G | 0.868465 | 1.78E-13 | 6.96E-08 |
| 11 | 64717461 | A | G | 0.868465 | 1.78E-13 | 6.96E-08 |
| 11 | 64717482 | T | C | 0.868465 | 1.78E-13 | 6.96E-08 |
| 11 | 64718055 | C | A | 0.868465 | 1.78E-13 | 6.96E-08 |
| 11 | 64718068 | G | T | 0.868465 | 1.78E-13 | 6.96E-08 |
| 11 | 64718099 | T | G | 0.868465 | 1.78E-13 | 6.96E-08 |
| 11 | 64718104 | A | G | 0.868465 | 1.78E-13 | 6.96E-08 |
| 11 | 64718106 | A | G | 0.868465 | 1.78E-13 | 6.96E-08 |
| 11 | 64718108 | A | T | 0.868465 | 1.78E-13 | 6.96E-08 |
| 11 | 64718180 | C | A | 0.868465 | 1.78E-13 | 6.96E-08 |
| 11 | 64718233 | G | T | 0.868465 | 1.78E-13 | 6.96E-08 |
| 11 | 64718275 | T | A | 0.868465 | 1.78E-13 | 6.96E-08 |
| 11 | 64718276 | T | A | 0.868465 | 1.78E-13 | 6.96E-08 |
| 11 | 64718277 | T | A | 0.868465 | 1.78E-13 | 6.96E-08 |
| 11 | 64718341 | A | G | 0.868465 | 1.78E-13 | 6.96E-08 |
| 11 | 64718357 | C | T | 0.868465 | 1.78E-13 | 6.96E-08 |
| 11 | 64718358 | C | T | 0.868465 | 1.78E-13 | 6.96E-08 |
| 11 | 64718384 | A | T | 0.868465 | 1.78E-13 | 6.96E-08 |
| 11 | 64718402 | G | A | 0.868465 | 1.78E-13 | 6.96E-08 |
| 11 | 64718611 | C | A | 0.868465 | 1.78E-13 | 6.96E-08 |
| 11 | 64718619 | C | A | 0.868465 | 1.78E-13 | 6.96E-08 |
| 11 | 64718697 | A | T | 0.868465 | 1.78E-13 | 6.96E-08 |
| 11 | 64718976 | G | A | 0.868465 | 1.78E-13 | 6.96E-08 |
| 11 | 64719186 | T | G | 0.868465 | 1.78E-13 | 6.96E-08 |
| 11 | 64719316 | A | G | 0.868465 | 1.78E-13 | 6.96E-08 |
| 11 | 64719391 | A | T | 0.868465 | 1.78E-13 | 6.96E-08 |
| 11 | 64719411 | G | T | 0.868465 | 1.78E-13 | 6.96E-08 |
| 11 | 64719477 | T | C | 0.868465 | 1.78E-13 | 6.96E-08 |
| 11 | 64719858 | C | T | 0.868465 | 1.78E-13 | 6.96E-08 |
| 11 | 64719965 | C | G | 0.868465 | 1.78E-13 | 6.96E-08 |
| 11 | 64720083 | T | C | 0.868465 | 1.78E-13 | 6.96E-08 |
| 11 | 64720543 | A | C | 0.868465 | 1.78E-13 | 6.96E-08 |
| 11 | 64720566 | T | C | 0.868465 | 1.78E-13 | 6.96E-08 |
| 11 | 64720708 | G | A | 0.868465 | 1.78E-13 | 6.96E-08 |
| 11 | 64720714 | T | C | 0.868465 | 1.78E-13 | 6.96E-08 |
| 11 | 64720766 | A | T | 0.868465 | 1.78E-13 | 6.96E-08 |
| 11 | 64720815 | T | A | 0.868465 | 1.78E-13 | 6.96E-08 |
| 11 | 64720926 | C | T | 0.868465 | 1.78E-13 | 6.96E-08 |
| 11 | 64720929 | G | A | 0.868465 | 1.78E-13 | 6.96E-08 |
| 11 | 64720945 | C | G | 0.868465 | 1.78E-13 | 6.96E-08 |
| 11 | 64721002 | C | T | 0.868465 | 1.78E-13 | 6.96E-08 |
| 11 | 64721073 | G | A | 0.868465 | 1.78E-13 | 6.96E-08 |
| 11 | 64721354 | T | C | 0.868465 | 1.78E-13 | 6.96E-08 |
| 11 | 64721639 | T | C | 0.868465 | 1.78E-13 | 6.96E-08 |
| 11 | 64721717 | C | A | 0.868465 | 1.78E-13 | 6.96E-08 |
| 11 | 64721807 | G | A | 0.868465 | 1.78E-13 | 6.96E-08 |
| 11 | 64721956 | A | T | 0.868465 | 1.78E-13 | 6.96E-08 |
| 11 | 64722035 | G | C | 0.868465 | 1.78E-13 | 6.96E-08 |
| 11 | 64722094 | A | G | 0.868465 | 1.78E-13 | 6.96E-08 |
| 11 | 64722155 | T | G | 0.868465 | 1.78E-13 | 6.96E-08 |
| 11 | 64722194 | G | A | 0.868465 | 1.78E-13 | 6.96E-08 |
| 11 | 64722249 | G | A | 0.868465 | 1.78E-13 | 6.96E-08 |
| 11 | 64722252 | G | A | 0.868465 | 1.78E-13 | 6.96E-08 |
| 11 | 64722270 | A | C | 0.868465 | 1.78E-13 | 6.96E-08 |
| 11 | 64722288 | A | G | 0.868465 | 1.78E-13 | 6.96E-08 |
| 11 | 64722345 | T | C | 0.868465 | 1.78E-13 | 6.96E-08 |
| 11 | 64722346 | G | A | 0.868465 | 1.78E-13 | 6.96E-08 |
| 11 | 64722348 | A | C | 0.868465 | 1.78E-13 | 6.96E-08 |
| 11 | 64722359 | A | T | 0.868465 | 1.78E-13 | 6.96E-08 |
| 11 | 64722393 | C | T | 0.868465 | 1.78E-13 | 6.96E-08 |
| 11 | 64722407 | C | A | 0.868465 | 1.78E-13 | 6.96E-08 |
| 11 | 64722447 | A | C | 0.868465 | 1.78E-13 | 6.96E-08 |
| 11 | 64722462 | C | T | 0.868465 | 1.78E-13 | 6.96E-08 |
| 11 | 64722553 | G | A | 0.868465 | 1.78E-13 | 6.96E-08 |
| 11 | 64722556 | G | A | 0.868465 | 1.78E-13 | 6.96E-08 |
| 11 | 64722571 | A | G | 0.868465 | 1.78E-13 | 6.96E-08 |
| 11 | 64722642 | A | T | 0.868465 | 1.78E-13 | 6.96E-08 |

| | | | | | | |
|---|---|---|---|---|---|---|
| 11 | 64722780 | G | A | 0.868465 | 1.78E-13 | 6.96E-08 |
| 11 | 64722784 | C | T | 0.868465 | 1.78E-13 | 6.96E-08 |
| 11 | 64722787 | A | T | 0.868465 | 1.78E-13 | 6.96E-08 |
| 11 | 64722844 | C | T | 0.868465 | 1.78E-13 | 6.96E-08 |
| 11 | 64722845 | C | T | 0.868465 | 1.78E-13 | 6.96E-08 |
| 11 | 64722856 | G | A | 0.868465 | 1.78E-13 | 6.96E-08 |
| 11 | 64722914 | C | G | 0.868465 | 1.78E-13 | 6.96E-08 |
| 11 | 64722952 | T | C | 0.868465 | 1.78E-13 | 6.96E-08 |
| 11 | 64723189 | T | G | 0.868465 | 1.78E-13 | 6.96E-08 |
| 11 | 64723251 | G | T | 0.868465 | 1.78E-13 | 6.96E-08 |
| 11 | 64723380 | A | C | 0.868465 | 1.78E-13 | 6.96E-08 |
| 11 | 64723458 | A | T | 0.868465 | 1.78E-13 | 6.96E-08 |
| 11 | 64723505 | A | C | 0.868465 | 1.78E-13 | 6.96E-08 |
| 11 | 64723528 | A | G | 0.868465 | 1.78E-13 | 6.96E-08 |
| 11 | 64723964 | A | G | 0.868465 | 1.78E-13 | 6.96E-08 |
| 11 | 64724237 | C | T | 0.868465 | 1.78E-13 | 6.96E-08 |
| 11 | 64724271 | C | T | 0.868465 | 1.78E-13 | 6.96E-08 |
| 11 | 64724282 | G | A | 0.868465 | 1.78E-13 | 6.96E-08 |
| 11 | 64724610 | C | G | 0.868465 | 1.78E-13 | 6.96E-08 |
| 11 | 64742943 | G | A | 0.868465 | 1.78E-13 | 6.96E-08 |
| 11 | 64754218 | T | A | 0.868465 | 1.78E-13 | 6.96E-08 |
| 11 | 64754893 | T | C | 0.868465 | 1.78E-13 | 6.96E-08 |
| 11 | 64756978 | T | C | 0.868465 | 1.78E-13 | 6.96E-08 |
| 11 | 64757137 | C | G | 0.868465 | 1.78E-13 | 6.96E-08 |
| 11 | 64757260 | C | A | 0.868465 | 1.78E-13 | 6.96E-08 |
| 11 | 64757678 | A | T | 0.868465 | 1.78E-13 | 6.96E-08 |
| 11 | 64757983 | A | G | 0.868465 | 1.78E-13 | 6.96E-08 |
| 11 | 64758179 | T | C | 0.868465 | 1.78E-13 | 6.96E-08 |
| 11 | 64758309 | G | A | 0.868465 | 1.78E-13 | 6.96E-08 |
| 11 | 64758335 | C | T | 0.868465 | 1.78E-13 | 6.96E-08 |
| 11 | 64758390 | A | G | 0.868465 | 1.78E-13 | 6.96E-08 |
| 11 | 64758432 | T | C | 0.868465 | 1.78E-13 | 6.96E-08 |
| 11 | 64758455 | G | A | 0.868465 | 1.78E-13 | 6.96E-08 |
| 11 | 64758464 | C | A | 0.868465 | 1.78E-13 | 6.96E-08 |
| 11 | 64758477 | A | G | 0.868465 | 1.78E-13 | 6.96E-08 |
| 11 | 64758485 | A | T | 0.868465 | 1.78E-13 | 6.96E-08 |
| 11 | 64758492 | A | G | 0.868465 | 1.78E-13 | 6.96E-08 |
| 11 | 64759683 | G | A | 0.868465 | 1.78E-13 | 6.96E-08 |
| 11 | 64759754 | G | A | 0.868465 | 1.78E-13 | 6.96E-08 |
| 11 | 64760465 | T | G | 0.868465 | 1.78E-13 | 6.96E-08 |
| 11 | 64761125 | T | A | 0.868465 | 1.78E-13 | 6.96E-08 |
| 11 | 64761187 | A | T | 0.868465 | 1.78E-13 | 6.96E-08 |
| 11 | 64761268 | T | C | 0.868465 | 1.78E-13 | 6.96E-08 |
| 11 | 64761428 | A | G | 0.868465 | 1.78E-13 | 6.96E-08 |
| 11 | 64761791 | C | T | 0.868465 | 1.78E-13 | 6.96E-08 |
| 11 | 64761832 | T | C | 0.868465 | 1.78E-13 | 6.96E-08 |
| 11 | 64761879 | T | C | 0.868465 | 1.78E-13 | 6.96E-08 |
| 11 | 64762426 | A | G | 0.868465 | 1.78E-13 | 6.96E-08 |
| 11 | 64762520 | T | A | 0.868465 | 1.78E-13 | 6.96E-08 |
| 11 | 64764857 | C | T | 0.868465 | 1.78E-13 | 6.96E-08 |
| 11 | 64765254 | G | A | 0.868465 | 1.78E-13 | 6.96E-08 |
| 11 | 64765358 | C | T | 0.868465 | 1.78E-13 | 6.96E-08 |
| 11 | 64765569 | C | G | 0.868465 | 1.78E-13 | 6.96E-08 |
| 11 | 64765743 | T | A | 0.868465 | 1.78E-13 | 6.96E-08 |
| 11 | 64766848 | A | G | 0.868465 | 1.78E-13 | 6.96E-08 |
| 11 | 64767995 | A | G | 0.868465 | 1.78E-13 | 6.96E-08 |
| 11 | 64768083 | G | C | 0.868465 | 1.78E-13 | 6.96E-08 |
| 11 | 64768269 | A | G | 0.868465 | 1.78E-13 | 6.96E-08 |
| 11 | 64768390 | T | C | 0.868465 | 1.78E-13 | 6.96E-08 |
| 11 | 64768392 | T | C | 0.868465 | 1.78E-13 | 6.96E-08 |
| 11 | 64768577 | G | A | 0.868465 | 1.78E-13 | 6.96E-08 |
| 11 | 64769048 | T | A | 0.868465 | 1.78E-13 | 6.96E-08 |
| 11 | 64769425 | A | T | 0.868465 | 1.78E-13 | 6.96E-08 |
| 11 | 64770915 | G | A | 0.868465 | 1.78E-13 | 6.96E-08 |
| 11 | 64770949 | T | C | 0.868465 | 1.78E-13 | 6.96E-08 |
| 11 | 64770968 | G | T | 0.868465 | 1.78E-13 | 6.96E-08 |
| 11 | 64771020 | C | T | 0.868465 | 1.78E-13 | 6.96E-08 |

| | | | | | | |
|---|---|---|---|---|---|---|
| 11 | 64771298 | G | A | 0.868465 | 1.78E-13 | 6.96E-08 |
| 11 | 64771361 | G | A | 0.868465 | 1.78E-13 | 6.96E-08 |
| 11 | 64771431 | C | T | 0.868465 | 1.78E-13 | 6.96E-08 |
| 11 | 64771450 | C | G | 0.868465 | 1.78E-13 | 6.96E-08 |
| 11 | 64771654 | T | G | 0.868465 | 1.78E-13 | 6.96E-08 |
| 11 | 64771668 | G | C | 0.868465 | 1.78E-13 | 6.96E-08 |
| 11 | 64771783 | C | A | 0.868465 | 1.78E-13 | 6.96E-08 |
| 11 | 64771839 | G | A | 0.868465 | 1.78E-13 | 6.96E-08 |
| 11 | 64771865 | A | G | 0.868465 | 1.78E-13 | 6.96E-08 |
| 11 | 64772130 | G | A | 0.868465 | 1.78E-13 | 6.96E-08 |
| 11 | 64772136 | T | G | 0.868465 | 1.78E-13 | 6.96E-08 |
| 11 | 64772143 | G | A | 0.868465 | 1.78E-13 | 6.96E-08 |
| 11 | 64772151 | T | C | 0.868465 | 1.78E-13 | 6.96E-08 |
| 11 | 64772177 | G | A | 0.868465 | 1.78E-13 | 6.96E-08 |
| 11 | 64772258 | G | A | 0.868465 | 1.78E-13 | 6.96E-08 |
| 11 | 64772275 | T | C | 0.868465 | 1.78E-13 | 6.96E-08 |
| 11 | 64772316 | C | T | 0.868465 | 1.78E-13 | 6.96E-08 |
| 11 | 64772369 | G | A | 0.868465 | 1.78E-13 | 6.96E-08 |
| 11 | 64772415 | T | C | 0.868465 | 1.78E-13 | 6.96E-08 |
| 11 | 64772618 | G | A | 0.868465 | 1.78E-13 | 6.96E-08 |
| 11 | 64772656 | A | G | 0.868465 | 1.78E-13 | 6.96E-08 |
| 11 | 64772736 | C | T | 0.868465 | 1.78E-13 | 6.96E-08 |
| 11 | 64772758 | A | G | 0.868465 | 1.78E-13 | 6.96E-08 |
| 11 | 64772795 | T | C | 0.868465 | 1.78E-13 | 6.96E-08 |
| 11 | 64772806 | C | T | 0.868465 | 1.78E-13 | 6.96E-08 |
| 11 | 64772864 | A | T | 0.868465 | 1.78E-13 | 6.96E-08 |
| 11 | 64772865 | A | T | 0.868465 | 1.78E-13 | 6.96E-08 |
| 11 | 64772866 | A | T | 0.868465 | 1.78E-13 | 6.96E-08 |
| 11 | 64772906 | G | C | 0.868465 | 1.78E-13 | 6.96E-08 |
| 11 | 64772978 | G | T | 0.868465 | 1.78E-13 | 6.96E-08 |
| 11 | 64773015 | A | G | 0.868465 | 1.78E-13 | 6.96E-08 |
| 11 | 64773120 | G | A | 0.868465 | 1.78E-13 | 6.96E-08 |
| 11 | 64773281 | G | A | 0.868465 | 1.78E-13 | 6.96E-08 |
| 11 | 64773322 | A | G | 0.868465 | 1.78E-13 | 6.96E-08 |
| 11 | 64776500 | C | G | 0.868465 | 1.78E-13 | 6.96E-08 |
| 11 | 64776540 | A | C | 0.868465 | 1.78E-13 | 6.96E-08 |
| 11 | 64776590 | G | A | 0.868465 | 1.78E-13 | 6.96E-08 |
| 11 | 64778079 | T | C | 0.868465 | 1.78E-13 | 6.96E-08 |
| 11 | 64782089 | T | G | 0.868465 | 1.78E-13 | 6.96E-08 |
| 11 | 64784138 | C | A | 0.868465 | 1.78E-13 | 6.96E-08 |
| 11 | 64787283 | G | A | 0.868465 | 1.78E-13 | 6.96E-08 |
| 11 | 64787836 | C | A | 0.868465 | 1.78E-13 | 6.96E-08 |
| 11 | 64788261 | G | A | 0.868465 | 1.78E-13 | 6.96E-08 |
| 11 | 64788847 | G | A | 0.868465 | 1.78E-13 | 6.96E-08 |
| 11 | 64788977 | G | A | 0.868465 | 1.78E-13 | 6.96E-08 |
| 11 | 64789920 | T | C | 0.868465 | 1.78E-13 | 6.96E-08 |
| 11 | 64790262 | T | G | 0.868465 | 1.78E-13 | 6.96E-08 |
| 11 | 64790488 | G | A | 0.868465 | 1.78E-13 | 6.96E-08 |
| 11 | 64790879 | T | G | 0.868465 | 1.78E-13 | 6.96E-08 |
| 11 | 64790880 | T | C | 0.868465 | 1.78E-13 | 6.96E-08 |
| 11 | 64790980 | A | G | 0.868465 | 1.78E-13 | 6.96E-08 |
| 11 | 64791048 | T | C | 0.868465 | 1.78E-13 | 6.96E-08 |
| 11 | 64791065 | A | G | 0.868465 | 1.78E-13 | 6.96E-08 |
| 11 | 64791110 | G | A | 0.868465 | 1.78E-13 | 6.96E-08 |
| 11 | 64791928 | A | G | 0.868465 | 1.78E-13 | 6.96E-08 |
| 11 | 64791962 | A | G | 0.868465 | 1.78E-13 | 6.96E-08 |
| 11 | 64792836 | C | T | 0.868465 | 1.78E-13 | 6.96E-08 |
| 11 | 64793332 | C | T | 0.868465 | 1.78E-13 | 6.96E-08 |
| 11 | 64793627 | C | T | 0.868465 | 1.78E-13 | 6.96E-08 |
| 11 | 64793695 | A | C | 0.868465 | 1.78E-13 | 6.96E-08 |
| 11 | 64793879 | A | T | 0.868465 | 1.78E-13 | 6.96E-08 |
| 11 | 64793881 | A | T | 0.868465 | 1.78E-13 | 6.96E-08 |
| 11 | 64793882 | A | T | 0.868465 | 1.78E-13 | 6.96E-08 |
| 11 | 64793883 | A | T | 0.868465 | 1.78E-13 | 6.96E-08 |
| 11 | 64794121 | T | C | 0.868465 | 1.78E-13 | 6.96E-08 |
| 11 | 64794220 | A | G | 0.868465 | 1.78E-13 | 6.96E-08 |
| 11 | 64794441 | C | T | 0.868465 | 1.78E-13 | 6.96E-08 |

| | | | | | | |
|---|---|---|---|---|---|---|
| 11 | 64795170 | A | G | 0.868465 | 1.78E-13 | 6.96E-08 |
| 11 | 64795230 | A | G | 0.868465 | 1.78E-13 | 6.96E-08 |
| 11 | 64795351 | G | A | 0.868465 | 1.78E-13 | 6.96E-08 |
| 11 | 64795493 | A | G | 0.868465 | 1.78E-13 | 6.96E-08 |
| 11 | 64795540 | C | A | 0.868465 | 1.78E-13 | 6.96E-08 |
| 11 | 64795577 | T | C | 0.868465 | 1.78E-13 | 6.96E-08 |
| 11 | 64795619 | C | G | 0.868465 | 1.78E-13 | 6.96E-08 |
| 11 | 64796121 | T | C | 0.868465 | 1.78E-13 | 6.96E-08 |
| 11 | 64796172 | G | A | 0.868465 | 1.78E-13 | 6.96E-08 |
| 11 | 64796703 | T | C | 0.868465 | 1.78E-13 | 6.96E-08 |
| 11 | 64796800 | T | C | 0.868465 | 1.78E-13 | 6.96E-08 |
| 11 | 64796901 | A | G | 0.868465 | 1.78E-13 | 6.96E-08 |
| 11 | 64796922 | A | G | 0.868465 | 1.78E-13 | 6.96E-08 |
| 11 | 64796928 | T | C | 0.868465 | 1.78E-13 | 6.96E-08 |
| 11 | 63829869 | T | C | 0.891231 | 1.82E-13 | 7.15E-08 |
| 11 | 69979231 | T | C | 0.91484 | 1.88E-13 | 7.38E-08 |
| 11 | 64588267 | G | T | 0.867622 | 1.92E-13 | 7.53E-08 |
| 11 | 64588513 | T | G | 0.867622 | 1.92E-13 | 7.53E-08 |
| 11 | 63913904 | T | C | 0.872681 | 1.92E-13 | 7.53E-08 |
| 11 | 63913905 | T | G | 0.872681 | 1.92E-13 | 7.53E-08 |
| 11 | 63913907 | G | T | 0.872681 | 1.92E-13 | 7.53E-08 |
| 11 | 63913925 | C | G | 0.872681 | 1.92E-13 | 7.53E-08 |
| 11 | 63913975 | T | C | 0.872681 | 1.92E-13 | 7.53E-08 |
| 11 | 63914000 | T | G | 0.872681 | 1.92E-13 | 7.53E-08 |
| 11 | 63914002 | C | A | 0.872681 | 1.92E-13 | 7.53E-08 |
| 11 | 63914005 | A | G | 0.872681 | 1.92E-13 | 7.53E-08 |
| 11 | 63914032 | T | A | 0.872681 | 1.92E-13 | 7.53E-08 |
| 11 | 63914047 | G | C | 0.872681 | 1.92E-13 | 7.53E-08 |
| 11 | 63914055 | G | T | 0.872681 | 1.92E-13 | 7.53E-08 |
| 11 | 63914070 | G | C | 0.872681 | 1.92E-13 | 7.53E-08 |
| 11 | 63914076 | T | C | 0.872681 | 1.92E-13 | 7.53E-08 |
| 11 | 64573759 | T | A | 0.870152 | 1.93E-13 | 7.55E-08 |
| 11 | 61512451 | T | C | 0.920742 | 1.97E-13 | 7.73E-08 |
| 11 | 69982101 | T | C | 0.910624 | 2.16E-13 | 8.46E-08 |
| 11 | 69982149 | T | C | 0.910624 | 2.16E-13 | 8.46E-08 |
| 11 | 69982283 | T | C | 0.910624 | 2.16E-13 | 8.46E-08 |
| 11 | 63830046 | G | A | 0.886172 | 2.49E-13 | 9.76E-08 |
| 11 | 64578904 | C | A | 0.870995 | 2.51E-13 | 9.83E-08 |
| 11 | 64578956 | A | G | 0.870995 | 2.51E-13 | 9.83E-08 |
| 11 | 64168161 | T | A | 0.891231 | 2.79E-13 | 1.09E-07 |
| 11 | 64168162 | A | T | 0.891231 | 2.79E-13 | 1.09E-07 |
| 11 | 64578971 | T | C | 0.871838 | 2.88E-13 | 1.13E-07 |
| 11 | 64640959 | A | G | 0.867622 | 3.04E-13 | 1.19E-07 |
| 11 | 64746469 | A | G | 0.870152 | 3.19E-13 | 1.25E-07 |
| 11 | 63813678 | A | G | 0.89629 | 3.36E-13 | 1.32E-07 |
| 11 | 63789992 | C | G | 0.881113 | 3.43E-13 | 1.34E-07 |
| 11 | 63790342 | G | A | 0.881113 | 3.43E-13 | 1.34E-07 |
| 11 | 63790352 | T | C | 0.881113 | 3.43E-13 | 1.34E-07 |
| 11 | 64535525 | C | T | 0.867622 | 3.61E-13 | 1.42E-07 |
| 11 | 64527892 | C | T | 0.869309 | 3.72E-13 | 1.46E-07 |
| 11 | 64535861 | G | T | 0.868465 | 3.76E-13 | 1.48E-07 |
| 11 | 64535897 | G | A | 0.868465 | 3.76E-13 | 1.48E-07 |
| 11 | 64545354 | C | T | 0.869309 | 3.97E-13 | 1.56E-07 |
| 11 | 64545674 | G | C | 0.869309 | 3.97E-13 | 1.56E-07 |
| 11 | 64545804 | C | A | 0.869309 | 3.97E-13 | 1.56E-07 |
| 11 | 64546084 | C | T | 0.869309 | 3.97E-13 | 1.56E-07 |
| 11 | 64678417 | A | T | 0.867622 | 4.47E-13 | 1.75E-07 |
| 11 | 64678514 | C | G | 0.867622 | 4.47E-13 | 1.75E-07 |
| 11 | 64681257 | T | A | 0.867622 | 4.47E-13 | 1.75E-07 |
| 11 | 64639687 | T | A | 0.869309 | 4.52E-13 | 1.77E-07 |
| 11 | 64639703 | C | T | 0.869309 | 4.52E-13 | 1.77E-07 |
| 11 | 64639765 | G | A | 0.869309 | 4.52E-13 | 1.77E-07 |
| 11 | 64639783 | T | C | 0.869309 | 4.52E-13 | 1.77E-07 |
| 11 | 64639792 | G | A | 0.869309 | 4.52E-13 | 1.77E-07 |
| 11 | 64639812 | C | T | 0.869309 | 4.52E-13 | 1.77E-07 |
| 11 | 64639874 | C | A | 0.869309 | 4.52E-13 | 1.77E-07 |
| 11 | 64639956 | C | T | 0.869309 | 4.52E-13 | 1.77E-07 |

| | | | | | | |
|---|---|---|---|---|---|---|
| 11 | 64640082 | A | C | 0.869309 | 4.52E-13 | 1.77E-07 |
| 11 | 64640085 | G | T | 0.869309 | 4.52E-13 | 1.77E-07 |
| 11 | 64553789 | A | T | 0.870152 | 4.67E-13 | 1.83E-07 |
| 11 | 64553920 | G | T | 0.870152 | 4.67E-13 | 1.83E-07 |
| 11 | 64908588 | T | A | 0.872681 | 4.89E-13 | 1.92E-07 |
| 11 | 64908996 | A | T | 0.872681 | 4.89E-13 | 1.92E-07 |
| 11 | 64910440 | A | G | 0.872681 | 4.89E-13 | 1.92E-07 |
| 11 | 64910550 | A | G | 0.872681 | 4.89E-13 | 1.92E-07 |
| 11 | 64910574 | T | C | 0.872681 | 4.89E-13 | 1.92E-07 |
| 11 | 64910577 | G | A | 0.872681 | 4.89E-13 | 1.92E-07 |
| 11 | 64910684 | T | C | 0.872681 | 4.89E-13 | 1.92E-07 |
| 11 | 64910868 | C | A | 0.872681 | 4.89E-13 | 1.92E-07 |
| 11 | 64910870 | G | A | 0.872681 | 4.89E-13 | 1.92E-07 |
| 11 | 64910942 | C | A | 0.872681 | 4.89E-13 | 1.92E-07 |
| 11 | 64911638 | C | A | 0.872681 | 4.89E-13 | 1.92E-07 |
| 11 | 64911815 | A | G | 0.872681 | 4.89E-13 | 1.92E-07 |
| 11 | 64912779 | C | T | 0.872681 | 4.89E-13 | 1.92E-07 |
| 11 | 64913175 | C | T | 0.872681 | 4.89E-13 | 1.92E-07 |
| 11 | 64913197 | G | A | 0.872681 | 4.89E-13 | 1.92E-07 |
| 11 | 64913236 | G | A | 0.872681 | 4.89E-13 | 1.92E-07 |
| 11 | 64913510 | A | G | 0.872681 | 4.89E-13 | 1.92E-07 |
| 11 | 64917915 | A | G | 0.872681 | 4.89E-13 | 1.92E-07 |
| 11 | 64917930 | T | C | 0.872681 | 4.89E-13 | 1.92E-07 |
| 11 | 64918320 | G | A | 0.872681 | 4.89E-13 | 1.92E-07 |
| 11 | 64918528 | C | T | 0.872681 | 4.89E-13 | 1.92E-07 |
| 11 | 64918868 | G | A | 0.872681 | 4.89E-13 | 1.92E-07 |
| 11 | 64918928 | T | C | 0.872681 | 4.89E-13 | 1.92E-07 |
| 11 | 64918938 | C | T | 0.872681 | 4.89E-13 | 1.92E-07 |
| 11 | 64921974 | C | T | 0.872681 | 4.89E-13 | 1.92E-07 |
| 11 | 64935039 | T | C | 0.872681 | 4.89E-13 | 1.92E-07 |
| 11 | 64935151 | T | C | 0.872681 | 4.89E-13 | 1.92E-07 |
| 11 | 64935178 | A | C | 0.872681 | 4.89E-13 | 1.92E-07 |
| 11 | 64935300 | T | C | 0.872681 | 4.89E-13 | 1.92E-07 |
| 11 | 64935394 | T | C | 0.872681 | 4.89E-13 | 1.92E-07 |
| 11 | 64935521 | T | C | 0.872681 | 4.89E-13 | 1.92E-07 |
| 11 | 64935531 | A | G | 0.872681 | 4.89E-13 | 1.92E-07 |
| 11 | 64935536 | T | C | 0.872681 | 4.89E-13 | 1.92E-07 |
| 11 | 64935546 | A | G | 0.872681 | 4.89E-13 | 1.92E-07 |
| 11 | 64935613 | G | A | 0.872681 | 4.89E-13 | 1.92E-07 |
| 11 | 64935631 | C | T | 0.872681 | 4.89E-13 | 1.92E-07 |
| 11 | 64935643 | G | C | 0.872681 | 4.89E-13 | 1.92E-07 |
| 11 | 64935713 | C | T | 0.872681 | 4.89E-13 | 1.92E-07 |
| 11 | 64936440 | G | C | 0.872681 | 4.89E-13 | 1.92E-07 |
| 11 | 64936456 | C | T | 0.872681 | 4.89E-13 | 1.92E-07 |
| 11 | 64936457 | G | T | 0.872681 | 4.89E-13 | 1.92E-07 |
| 11 | 64936610 | G | A | 0.872681 | 4.89E-13 | 1.92E-07 |
| 11 | 64937297 | T | C | 0.872681 | 4.89E-13 | 1.92E-07 |
| 11 | 64965492 | T | C | 0.872681 | 4.89E-13 | 1.92E-07 |
| 11 | 64965505 | C | T | 0.872681 | 4.89E-13 | 1.92E-07 |
| 11 | 64590492 | A | T | 0.870995 | 4.93E-13 | 1.93E-07 |
| 11 | 64536457 | G | A | 0.870995 | 5.39E-13 | 2.11E-07 |
| 11 | 64536749 | A | C | 0.870995 | 5.39E-13 | 2.11E-07 |
| 11 | 63260255 | G | T | 0.851602 | 5.64E-13 | 2.21E-07 |
| 11 | 64597596 | T | A | 0.866779 | 5.80E-13 | 2.27E-07 |
| 11 | 64682932 | T | C | 0.86425 | 6.25E-13 | 2.45E-07 |
| 11 | 64683364 | T | G | 0.86425 | 6.25E-13 | 2.45E-07 |
| 11 | 64683372 | C | T | 0.86425 | 6.25E-13 | 2.45E-07 |
| 11 | 63255963 | T | C | 0.849916 | 6.44E-13 | 2.53E-07 |
| 11 | 64585257 | C | T | 0.871838 | 6.46E-13 | 2.53E-07 |
| 11 | 64585377 | G | C | 0.871838 | 6.46E-13 | 2.53E-07 |
| 11 | 64528597 | G | A | 0.866779 | 7.78E-13 | 3.05E-07 |
| 11 | 64528602 | A | G | 0.866779 | 7.78E-13 | 3.05E-07 |
| 11 | 64529645 | G | A | 0.866779 | 7.78E-13 | 3.05E-07 |
| 11 | 64529719 | G | A | 0.866779 | 7.78E-13 | 3.05E-07 |
| 11 | 64529856 | G | A | 0.866779 | 7.78E-13 | 3.05E-07 |
| 11 | 64529887 | C | G | 0.866779 | 7.78E-13 | 3.05E-07 |
| 11 | 64587079 | G | T | 0.866779 | 7.79E-13 | 3.06E-07 |

| | | | | | | |
|---|---|---|---|---|---|---|
| 11 | 63931753 | T | C | 0.834739 | 8.51E-13 | 3.33E-07 |
| 11 | 63931771 | T | C | 0.834739 | 8.51E-13 | 3.33E-07 |
| 11 | 63934937 | A | G | 0.834739 | 8.51E-13 | 3.33E-07 |
| 11 | 63936178 | A | G | 0.834739 | 8.51E-13 | 3.33E-07 |
| 11 | 63936337 | C | T | 0.834739 | 8.51E-13 | 3.33E-07 |
| 11 | 63936343 | T | C | 0.834739 | 8.51E-13 | 3.33E-07 |
| 11 | 63936354 | A | G | 0.834739 | 8.51E-13 | 3.33E-07 |
| 11 | 63936403 | C | T | 0.834739 | 8.51E-13 | 3.33E-07 |
| 11 | 63936416 | A | G | 0.834739 | 8.51E-13 | 3.33E-07 |
| 11 | 63936445 | A | G | 0.834739 | 8.51E-13 | 3.33E-07 |
| 11 | 63936597 | T | C | 0.834739 | 8.51E-13 | 3.33E-07 |
| 11 | 63936783 | A | G | 0.834739 | 8.51E-13 | 3.33E-07 |
| 11 | 63936941 | A | G | 0.834739 | 8.51E-13 | 3.33E-07 |
| 11 | 63937286 | C | T | 0.834739 | 8.51E-13 | 3.33E-07 |
| 11 | 63937726 | A | G | 0.834739 | 8.51E-13 | 3.33E-07 |
| 11 | 63938090 | G | A | 0.834739 | 8.51E-13 | 3.33E-07 |
| 11 | 63939295 | A | T | 0.834739 | 8.51E-13 | 3.33E-07 |
| 11 | 63939335 | A | T | 0.834739 | 8.51E-13 | 3.33E-07 |
| 11 | 63939369 | G | T | 0.834739 | 8.51E-13 | 3.33E-07 |
| 11 | 63939377 | G | A | 0.834739 | 8.51E-13 | 3.33E-07 |
| 11 | 63939475 | T | G | 0.834739 | 8.51E-13 | 3.33E-07 |
| 11 | 63939535 | A | G | 0.834739 | 8.51E-13 | 3.33E-07 |
| 11 | 63939538 | T | C | 0.834739 | 8.51E-13 | 3.33E-07 |
| 11 | 63939654 | T | G | 0.834739 | 8.51E-13 | 3.33E-07 |
| 11 | 63940111 | G | C | 0.834739 | 8.51E-13 | 3.33E-07 |
| 11 | 63940145 | A | G | 0.834739 | 8.51E-13 | 3.33E-07 |
| 11 | 63940282 | G | T | 0.834739 | 8.51E-13 | 3.33E-07 |
| 11 | 63940361 | C | G | 0.834739 | 8.51E-13 | 3.33E-07 |
| 11 | 63940362 | T | G | 0.834739 | 8.51E-13 | 3.33E-07 |
| 11 | 63940392 | G | A | 0.834739 | 8.51E-13 | 3.33E-07 |
| 11 | 63940511 | A | G | 0.834739 | 8.51E-13 | 3.33E-07 |
| 11 | 63940587 | G | A | 0.834739 | 8.51E-13 | 3.33E-07 |
| 11 | 63940607 | T | C | 0.834739 | 8.51E-13 | 3.33E-07 |
| 11 | 63940693 | C | T | 0.834739 | 8.51E-13 | 3.33E-07 |
| 11 | 63940820 | A | G | 0.834739 | 8.51E-13 | 3.33E-07 |
| 11 | 63940832 | G | C | 0.834739 | 8.51E-13 | 3.33E-07 |
| 11 | 63940838 | A | G | 0.834739 | 8.51E-13 | 3.33E-07 |
| 11 | 63940884 | C | A | 0.834739 | 8.51E-13 | 3.33E-07 |
| 11 | 63940885 | T | G | 0.834739 | 8.51E-13 | 3.33E-07 |
| 11 | 63940905 | T | C | 0.834739 | 8.51E-13 | 3.33E-07 |
| 11 | 63940943 | G | T | 0.834739 | 8.51E-13 | 3.33E-07 |
| 11 | 63941098 | A | C | 0.834739 | 8.51E-13 | 3.33E-07 |
| 11 | 63941177 | A | G | 0.834739 | 8.51E-13 | 3.33E-07 |
| 11 | 63941279 | G | A | 0.834739 | 8.51E-13 | 3.33E-07 |
| 11 | 63941385 | C | T | 0.834739 | 8.51E-13 | 3.33E-07 |
| 11 | 63941412 | A | G | 0.834739 | 8.51E-13 | 3.33E-07 |
| 11 | 63941468 | G | A | 0.834739 | 8.51E-13 | 3.33E-07 |
| 11 | 63941484 | C | T | 0.834739 | 8.51E-13 | 3.33E-07 |
| 11 | 63941486 | A | C | 0.834739 | 8.51E-13 | 3.33E-07 |
| 11 | 63941506 | G | A | 0.834739 | 8.51E-13 | 3.33E-07 |
| 11 | 63941550 | G | A | 0.834739 | 8.51E-13 | 3.33E-07 |
| 11 | 63941574 | T | C | 0.834739 | 8.51E-13 | 3.33E-07 |
| 11 | 63941708 | C | G | 0.834739 | 8.51E-13 | 3.33E-07 |
| 11 | 63941919 | A | G | 0.834739 | 8.51E-13 | 3.33E-07 |
| 11 | 63942044 | G | C | 0.834739 | 8.51E-13 | 3.33E-07 |
| 11 | 63942321 | A | G | 0.834739 | 8.51E-13 | 3.33E-07 |
| 11 | 63942381 | G | T | 0.834739 | 8.51E-13 | 3.33E-07 |
| 11 | 63942385 | C | T | 0.834739 | 8.51E-13 | 3.33E-07 |
| 11 | 63947665 | G | A | 0.834739 | 8.51E-13 | 3.33E-07 |
| 11 | 63947751 | C | T | 0.834739 | 8.51E-13 | 3.33E-07 |
| 11 | 63950328 | A | G | 0.834739 | 8.51E-13 | 3.33E-07 |
| 11 | 63950729 | A | G | 0.834739 | 8.51E-13 | 3.33E-07 |
| 11 | 63951713 | A | G | 0.834739 | 8.51E-13 | 3.33E-07 |
| 11 | 63955441 | C | T | 0.834739 | 8.51E-13 | 3.33E-07 |
| 11 | 64640358 | A | G | 0.867622 | 8.56E-13 | 3.35E-07 |
| 11 | 64536145 | G | A | 0.870152 | 8.74E-13 | 3.43E-07 |
| 11 | 64536168 | C | G | 0.870152 | 8.74E-13 | 3.43E-07 |

| | | | | | | |
|---|---|---|---|---|---|---|
| 11 | 64536171 | C | T | 0.870152 | 8.74E-13 | 3.43E-07 |
| 11 | 64536174 | C | T | 0.870152 | 8.74E-13 | 3.43E-07 |
| 11 | 64586847 | G | A | 0.865936 | 1.01E-12 | 3.98E-07 |
| 11 | 64168955 | T | C | 0.892074 | 1.01E-12 | 3.98E-07 |
| 11 | 64168975 | A | G | 0.892074 | 1.01E-12 | 3.98E-07 |
| 11 | 64168977 | C | T | 0.892074 | 1.01E-12 | 3.98E-07 |
| 11 | 69806847 | T | C | 0.872681 | 1.06E-12 | 4.14E-07 |
| 11 | 66074890 | G | C | 0.869309 | 1.15E-12 | 4.49E-07 |
| 11 | 66074891 | A | C | 0.869309 | 1.15E-12 | 4.49E-07 |
| 11 | 64549529 | C | T | 0.870152 | 1.15E-12 | 4.51E-07 |
| 11 | 64550377 | C | T | 0.870152 | 1.15E-12 | 4.51E-07 |
| 11 | 64919038 | G | T | 0.873524 | 1.23E-12 | 4.80E-07 |
| 11 | 64919062 | A | T | 0.873524 | 1.23E-12 | 4.80E-07 |
| 11 | 64919073 | T | C | 0.873524 | 1.23E-12 | 4.80E-07 |
| 11 | 64919112 | T | C | 0.873524 | 1.23E-12 | 4.80E-07 |
| 11 | 64919378 | C | T | 0.873524 | 1.23E-12 | 4.80E-07 |
| 11 | 64919609 | A | T | 0.873524 | 1.23E-12 | 4.80E-07 |
| 11 | 64919877 | C | G | 0.873524 | 1.23E-12 | 4.80E-07 |
| 11 | 64919910 | T | C | 0.873524 | 1.23E-12 | 4.80E-07 |
| 11 | 64920015 | A | G | 0.873524 | 1.23E-12 | 4.80E-07 |
| 11 | 64640626 | G | C | 0.869309 | 1.36E-12 | 5.35E-07 |
| 11 | 64640630 | A | T | 0.869309 | 1.36E-12 | 5.35E-07 |
| 11 | 64640650 | A | T | 0.869309 | 1.36E-12 | 5.35E-07 |
| 11 | 64640672 | T | A | 0.869309 | 1.36E-12 | 5.35E-07 |
| 11 | 64640680 | T | A | 0.869309 | 1.36E-12 | 5.35E-07 |
| 11 | 64640698 | G | C | 0.869309 | 1.36E-12 | 5.35E-07 |
| 11 | 64080476 | G | A | 0.848229 | 1.39E-12 | 5.44E-07 |
| 11 | 64564306 | G | T | 0.869309 | 1.39E-12 | 5.45E-07 |
| 11 | 64564652 | T | C | 0.869309 | 1.39E-12 | 5.45E-07 |
| 11 | 64565270 | G | A | 0.869309 | 1.39E-12 | 5.45E-07 |
| 11 | 64744878 | T | A | 0.869309 | 1.39E-12 | 5.45E-07 |
| 11 | 64745139 | T | G | 0.869309 | 1.39E-12 | 5.45E-07 |
| 11 | 64745224 | A | G | 0.869309 | 1.39E-12 | 5.45E-07 |
| 11 | 64745365 | A | G | 0.869309 | 1.39E-12 | 5.45E-07 |
| 11 | 64745459 | A | G | 0.869309 | 1.39E-12 | 5.45E-07 |
| 11 | 69803934 | T | C | 0.873524 | 1.46E-12 | 5.74E-07 |
| 11 | 69804558 | T | C | 0.873524 | 1.46E-12 | 5.74E-07 |
| 11 | 69804850 | A | C | 0.873524 | 1.46E-12 | 5.74E-07 |
| 11 | 69804880 | A | G | 0.873524 | 1.46E-12 | 5.74E-07 |
| 11 | 69805196 | T | G | 0.873524 | 1.46E-12 | 5.74E-07 |
| 11 | 69805848 | C | T | 0.873524 | 1.46E-12 | 5.74E-07 |
| 11 | 69805885 | A | G | 0.873524 | 1.46E-12 | 5.74E-07 |
| 11 | 69806919 | T | C | 0.873524 | 1.46E-12 | 5.74E-07 |
| 11 | 69808424 | A | G | 0.873524 | 1.46E-12 | 5.74E-07 |
| 11 | 69808428 | C | T | 0.873524 | 1.46E-12 | 5.74E-07 |
| 11 | 69808475 | A | C | 0.873524 | 1.46E-12 | 5.74E-07 |
| 11 | 69808511 | T | C | 0.873524 | 1.46E-12 | 5.74E-07 |
| 11 | 69808522 | C | T | 0.873524 | 1.46E-12 | 5.74E-07 |
| 11 | 69823021 | A | G | 0.873524 | 1.46E-12 | 5.74E-07 |
| 11 | 69823030 | C | T | 0.873524 | 1.46E-12 | 5.74E-07 |
| 11 | 69823981 | A | G | 0.873524 | 1.46E-12 | 5.74E-07 |
| 11 | 69827155 | A | C | 0.873524 | 1.46E-12 | 5.74E-07 |
| 11 | 69827234 | G | T | 0.873524 | 1.46E-12 | 5.74E-07 |
| 11 | 69827273 | T | G | 0.873524 | 1.46E-12 | 5.74E-07 |
| 11 | 69827280 | A | T | 0.873524 | 1.46E-12 | 5.74E-07 |
| 11 | 69827284 | T | C | 0.873524 | 1.46E-12 | 5.74E-07 |
| 11 | 69827346 | C | T | 0.873524 | 1.46E-12 | 5.74E-07 |
| 11 | 69827983 | T | C | 0.873524 | 1.46E-12 | 5.74E-07 |
| 11 | 69827986 | A | G | 0.873524 | 1.46E-12 | 5.74E-07 |
| 11 | 69828018 | A | G | 0.873524 | 1.46E-12 | 5.74E-07 |
| 11 | 69828140 | T | C | 0.873524 | 1.46E-12 | 5.74E-07 |
| 11 | 69828164 | G | A | 0.873524 | 1.46E-12 | 5.74E-07 |
| 11 | 69828395 | C | T | 0.873524 | 1.46E-12 | 5.74E-07 |
| 11 | 69828686 | C | T | 0.873524 | 1.46E-12 | 5.74E-07 |
| 11 | 69829111 | G | C | 0.873524 | 1.46E-12 | 5.74E-07 |
| 11 | 69829556 | C | G | 0.873524 | 1.46E-12 | 5.74E-07 |
| 11 | 69829905 | T | C | 0.873524 | 1.46E-12 | 5.74E-07 |

| | | | | | | |
|---|---|---|---|---|---|---|
| 11 | 69830252 | T | C | 0.873524 | 1.46E-12 | 5.74E-07 |
| 11 | 69830257 | C | T | 0.873524 | 1.46E-12 | 5.74E-07 |
| 11 | 69830297 | A | G | 0.873524 | 1.46E-12 | 5.74E-07 |
| 11 | 69838719 | C | T | 0.873524 | 1.46E-12 | 5.74E-07 |
| 11 | 69838738 | G | A | 0.873524 | 1.46E-12 | 5.74E-07 |
| 11 | 69838747 | A | G | 0.873524 | 1.46E-12 | 5.74E-07 |
| 11 | 69838749 | T | C | 0.873524 | 1.46E-12 | 5.74E-07 |
| 11 | 69838750 | G | A | 0.873524 | 1.46E-12 | 5.74E-07 |
| 11 | 69838820 | G | A | 0.873524 | 1.46E-12 | 5.74E-07 |
| 11 | 69838821 | G | A | 0.873524 | 1.46E-12 | 5.74E-07 |
| 11 | 69838825 | T | G | 0.873524 | 1.46E-12 | 5.74E-07 |
| 11 | 69838830 | T | C | 0.873524 | 1.46E-12 | 5.74E-07 |
| 11 | 69838831 | G | A | 0.873524 | 1.46E-12 | 5.74E-07 |
| 11 | 69838854 | T | C | 0.873524 | 1.46E-12 | 5.74E-07 |
| 11 | 69839098 | A | G | 0.873524 | 1.46E-12 | 5.74E-07 |
| 11 | 69839178 | A | G | 0.873524 | 1.46E-12 | 5.74E-07 |
| 11 | 69839179 | T | C | 0.873524 | 1.46E-12 | 5.74E-07 |
| 11 | 69839196 | A | G | 0.873524 | 1.46E-12 | 5.74E-07 |
| 11 | 69839719 | T | C | 0.873524 | 1.46E-12 | 5.74E-07 |
| 11 | 64537107 | A | T | 0.867622 | 1.59E-12 | 6.24E-07 |
| 11 | 64537108 | G | A | 0.867622 | 1.59E-12 | 6.24E-07 |
| 11 | 63294760 | A | G | 0.844013 | 1.61E-12 | 6.31E-07 |
| 11 | 63295045 | A | T | 0.844013 | 1.61E-12 | 6.31E-07 |
| 11 | 63298184 | A | G | 0.844013 | 1.61E-12 | 6.31E-07 |
| 11 | 63298230 | T | G | 0.844013 | 1.61E-12 | 6.31E-07 |
| 11 | 63299294 | T | C | 0.844013 | 1.61E-12 | 6.31E-07 |
| 11 | 69807451 | G | T | 0.860877 | 1.94E-12 | 7.59E-07 |
| 11 | 69807464 | A | G | 0.860877 | 1.94E-12 | 7.59E-07 |
| 11 | 69807476 | T | A | 0.860877 | 1.94E-12 | 7.59E-07 |
| 11 | 69807612 | T | A | 0.860877 | 1.94E-12 | 7.59E-07 |
| 11 | 69807653 | T | C | 0.860877 | 1.94E-12 | 7.59E-07 |
| 11 | 69807666 | C | T | 0.860877 | 1.94E-12 | 7.59E-07 |
| 11 | 69807667 | T | G | 0.860877 | 1.94E-12 | 7.59E-07 |
| 11 | 69807769 | A | G | 0.860877 | 1.94E-12 | 7.59E-07 |
| 11 | 69807778 | T | C | 0.860877 | 1.94E-12 | 7.59E-07 |
| 11 | 69807786 | G | T | 0.860877 | 1.94E-12 | 7.59E-07 |
| 11 | 63938324 | A | C | 0.835582 | 2.06E-12 | 8.07E-07 |
| 11 | 64082184 | G | A | 0.847386 | 2.07E-12 | 8.12E-07 |
| 11 | 64083121 | A | G | 0.847386 | 2.07E-12 | 8.12E-07 |
| 11 | 64083186 | C | G | 0.847386 | 2.07E-12 | 8.12E-07 |
| 11 | 64083574 | T | C | 0.847386 | 2.07E-12 | 8.12E-07 |
| 11 | 64083770 | G | A | 0.847386 | 2.07E-12 | 8.12E-07 |
| 11 | 64084079 | A | G | 0.847386 | 2.07E-12 | 8.12E-07 |
| 11 | 64087932 | A | T | 0.847386 | 2.07E-12 | 8.12E-07 |
| 11 | 64640578 | A | G | 0.868465 | 2.26E-12 | 8.87E-07 |
| 11 | 64073205 | A | G | 0.841484 | 2.54E-12 | 9.96E-07 |
| 11 | 64168707 | A | G | 0.893761 | 2.64E-12 | 1.04E-06 |
| 11 | 64168709 | G | A | 0.893761 | 2.64E-12 | 1.04E-06 |
| 11 | 64168727 | T | A | 0.893761 | 2.64E-12 | 1.04E-06 |
| 11 | 64168728 | T | A | 0.893761 | 2.64E-12 | 1.04E-06 |
| 11 | 64168734 | T | A | 0.893761 | 2.64E-12 | 1.04E-06 |
| 11 | 64168735 | C | A | 0.893761 | 2.64E-12 | 1.04E-06 |
| 11 | 64168737 | A | T | 0.893761 | 2.64E-12 | 1.04E-06 |
| 11 | 64168857 | A | G | 0.893761 | 2.64E-12 | 1.04E-06 |
| 11 | 64168878 | T | C | 0.893761 | 2.64E-12 | 1.04E-06 |
| 11 | 64168902 | T | A | 0.893761 | 2.64E-12 | 1.04E-06 |
| 11 | 64168928 | C | T | 0.893761 | 2.64E-12 | 1.04E-06 |
| 11 | 64639472 | C | A | 0.871838 | 2.90E-12 | 1.13E-06 |
| 11 | 64639565 | G | C | 0.871838 | 2.90E-12 | 1.13E-06 |
| 11 | 64639577 | T | C | 0.871838 | 2.90E-12 | 1.13E-06 |
| 11 | 63796117 | C | T | 0.899663 | 3.13E-12 | 1.23E-06 |
| 11 | 63796121 | A | G | 0.899663 | 3.13E-12 | 1.23E-06 |
| 11 | 63796468 | G | A | 0.899663 | 3.13E-12 | 1.23E-06 |
| 11 | 64639245 | T | C | 0.872681 | 3.22E-12 | 1.26E-06 |
| 11 | 64639254 | T | A | 0.872681 | 3.22E-12 | 1.26E-06 |
| 11 | 63799192 | A | G | 0.913153 | 3.24E-12 | 1.27E-06 |
| 11 | 69811445 | A | G | 0.859191 | 3.28E-12 | 1.28E-06 |

| | | | | | | |
|---|---|---|---|---|---|---|
| 11 | 69811490 | G | A | 0.859191 | 3.28E-12 | 1.28E-06 |
| 11 | 69811585 | G | A | 0.859191 | 3.28E-12 | 1.28E-06 |
| 11 | 69811661 | T | C | 0.859191 | 3.28E-12 | 1.28E-06 |
| 11 | 69811667 | A | G | 0.859191 | 3.28E-12 | 1.28E-06 |
| 11 | 69813397 | T | C | 0.859191 | 3.28E-12 | 1.28E-06 |
| 11 | 69814659 | A | C | 0.859191 | 3.28E-12 | 1.28E-06 |
| 11 | 69814685 | T | G | 0.859191 | 3.28E-12 | 1.28E-06 |
| 11 | 69814702 | A | T | 0.859191 | 3.28E-12 | 1.28E-06 |
| 11 | 69814793 | A | C | 0.859191 | 3.28E-12 | 1.28E-06 |
| 11 | 69814899 | C | G | 0.859191 | 3.28E-12 | 1.28E-06 |
| 11 | 69814900 | C | T | 0.859191 | 3.28E-12 | 1.28E-06 |
| 11 | 64640296 | T | C | 0.870152 | 3.34E-12 | 1.31E-06 |
| 11 | 64640298 | A | G | 0.870152 | 3.34E-12 | 1.31E-06 |
| 11 | 64640301 | G | A | 0.870152 | 3.34E-12 | 1.31E-06 |
| 11 | 64683752 | G | A | 0.862563 | 3.47E-12 | 1.36E-06 |
| 11 | 69895736 | T | C | 0.886172 | 3.52E-12 | 1.38E-06 |
| 11 | 69895957 | C | T | 0.886172 | 3.52E-12 | 1.38E-06 |
| 11 | 69897137 | T | C | 0.886172 | 3.52E-12 | 1.38E-06 |
| 11 | 69897174 | C | T | 0.886172 | 3.52E-12 | 1.38E-06 |
| 11 | 69897242 | A | G | 0.886172 | 3.52E-12 | 1.38E-06 |
| 11 | 69897253 | G | A | 0.886172 | 3.52E-12 | 1.38E-06 |
| 11 | 69897270 | C | T | 0.886172 | 3.52E-12 | 1.38E-06 |
| 11 | 69897299 | C | G | 0.886172 | 3.52E-12 | 1.38E-06 |
| 11 | 69897300 | G | A | 0.886172 | 3.52E-12 | 1.38E-06 |
| 11 | 69897474 | G | C | 0.886172 | 3.52E-12 | 1.38E-06 |
| 11 | 69897490 | T | C | 0.886172 | 3.52E-12 | 1.38E-06 |
| 11 | 69898240 | G | A | 0.886172 | 3.52E-12 | 1.38E-06 |
| 11 | 69898485 | A | T | 0.886172 | 3.52E-12 | 1.38E-06 |
| 11 | 69898526 | A | G | 0.886172 | 3.52E-12 | 1.38E-06 |
| 11 | 69898570 | T | C | 0.886172 | 3.52E-12 | 1.38E-06 |
| 11 | 69898583 | T | C | 0.886172 | 3.52E-12 | 1.38E-06 |
| 11 | 69899008 | A | G | 0.886172 | 3.52E-12 | 1.38E-06 |
| 11 | 69899393 | T | C | 0.886172 | 3.52E-12 | 1.38E-06 |
| 11 | 69899537 | A | G | 0.886172 | 3.52E-12 | 1.38E-06 |
| 11 | 69900012 | A | G | 0.886172 | 3.52E-12 | 1.38E-06 |
| 11 | 69900056 | T | A | 0.886172 | 3.52E-12 | 1.38E-06 |
| 11 | 69900109 | C | G | 0.886172 | 3.52E-12 | 1.38E-06 |
| 11 | 69900266 | C | T | 0.886172 | 3.52E-12 | 1.38E-06 |
| 11 | 69901459 | G | T | 0.886172 | 3.52E-12 | 1.38E-06 |
| 11 | 69901461 | G | A | 0.886172 | 3.52E-12 | 1.38E-06 |
| 11 | 69901736 | G | A | 0.886172 | 3.52E-12 | 1.38E-06 |
| 11 | 69901898 | T | C | 0.886172 | 3.52E-12 | 1.38E-06 |
| 11 | 69902162 | T | C | 0.886172 | 3.52E-12 | 1.38E-06 |
| 11 | 69902726 | A | C | 0.886172 | 3.52E-12 | 1.38E-06 |
| 11 | 69903508 | T | C | 0.886172 | 3.52E-12 | 1.38E-06 |
| 11 | 69903607 | A | G | 0.886172 | 3.52E-12 | 1.38E-06 |
| 11 | 69903656 | A | G | 0.886172 | 3.52E-12 | 1.38E-06 |
| 11 | 69903798 | G | C | 0.886172 | 3.52E-12 | 1.38E-06 |
| 11 | 69904093 | T | A | 0.886172 | 3.52E-12 | 1.38E-06 |
| 11 | 69904254 | T | C | 0.886172 | 3.52E-12 | 1.38E-06 |
| 11 | 69904428 | C | T | 0.886172 | 3.52E-12 | 1.38E-06 |
| 11 | 69904461 | A | C | 0.886172 | 3.52E-12 | 1.38E-06 |
| 11 | 69904778 | T | C | 0.886172 | 3.52E-12 | 1.38E-06 |
| 11 | 69904832 | T | C | 0.886172 | 3.52E-12 | 1.38E-06 |
| 11 | 69905063 | T | C | 0.886172 | 3.52E-12 | 1.38E-06 |
| 11 | 69905295 | A | G | 0.886172 | 3.52E-12 | 1.38E-06 |
| 11 | 69905316 | C | T | 0.886172 | 3.52E-12 | 1.38E-06 |
| 11 | 69905568 | A | G | 0.886172 | 3.52E-12 | 1.38E-06 |
| 11 | 69905587 | C | T | 0.886172 | 3.52E-12 | 1.38E-06 |
| 11 | 69893266 | T | C | 0.887015 | 3.60E-12 | 1.41E-06 |
| 11 | 69897651 | C | T | 0.887015 | 3.60E-12 | 1.41E-06 |
| 11 | 69897673 | C | T | 0.887015 | 3.60E-12 | 1.41E-06 |
| 11 | 69897695 | A | G | 0.887015 | 3.60E-12 | 1.41E-06 |
| 11 | 69897940 | G | A | 0.887015 | 3.60E-12 | 1.41E-06 |
| 11 | 64663140 | T | C | 0.860877 | 3.62E-12 | 1.42E-06 |
| 11 | 64683719 | C | A | 0.863406 | 3.73E-12 | 1.46E-06 |
| 11 | 69804762 | T | C | 0.860034 | 3.88E-12 | 1.52E-06 |

| | | | | | | |
|---|---|---|---|---|---|---|
| 11 | 69804786 | G | A | 0.860034 | 3.88E-12 | 1.52E-06 |
| 11 | 69805660 | T | C | 0.860034 | 3.88E-12 | 1.52E-06 |
| 11 | 69805674 | T | C | 0.860034 | 3.88E-12 | 1.52E-06 |
| 11 | 69805675 | C | T | 0.860034 | 3.88E-12 | 1.52E-06 |
| 11 | 69805676 | T | C | 0.860034 | 3.88E-12 | 1.52E-06 |
| 11 | 69805677 | T | A | 0.860034 | 3.88E-12 | 1.52E-06 |
| 11 | 69808013 | T | C | 0.860034 | 3.88E-12 | 1.52E-06 |
| 11 | 69808023 | A | G | 0.860034 | 3.88E-12 | 1.52E-06 |
| 11 | 69808080 | A | C | 0.860034 | 3.88E-12 | 1.52E-06 |
| 11 | 69808099 | A | G | 0.860034 | 3.88E-12 | 1.52E-06 |
| 11 | 69811946 | T | C | 0.860034 | 3.88E-12 | 1.52E-06 |
| 11 | 69812188 | T | C | 0.860034 | 3.88E-12 | 1.52E-06 |
| 11 | 69813784 | C | T | 0.860034 | 3.88E-12 | 1.52E-06 |
| 11 | 69813805 | C | A | 0.860034 | 3.88E-12 | 1.52E-06 |
| 11 | 69813819 | A | G | 0.860034 | 3.88E-12 | 1.52E-06 |
| 11 | 69813827 | C | A | 0.860034 | 3.88E-12 | 1.52E-06 |
| 11 | 69813831 | T | C | 0.860034 | 3.88E-12 | 1.52E-06 |
| 11 | 69813979 | C | T | 0.860034 | 3.88E-12 | 1.52E-06 |
| 11 | 69814060 | G | T | 0.860034 | 3.88E-12 | 1.52E-06 |
| 11 | 69814374 | T | C | 0.860034 | 3.88E-12 | 1.52E-06 |
| 11 | 69814377 | G | A | 0.860034 | 3.88E-12 | 1.52E-06 |
| 11 | 69814571 | G | C | 0.860034 | 3.88E-12 | 1.52E-06 |
| 11 | 69815767 | G | A | 0.860034 | 3.88E-12 | 1.52E-06 |
| 11 | 69816035 | G | A | 0.860034 | 3.88E-12 | 1.52E-06 |
| 11 | 69816146 | G | A | 0.860034 | 3.88E-12 | 1.52E-06 |
| 11 | 69816277 | A | T | 0.860034 | 3.88E-12 | 1.52E-06 |
| 11 | 69816728 | C | T | 0.860034 | 3.88E-12 | 1.52E-06 |
| 11 | 69816762 | G | T | 0.860034 | 3.88E-12 | 1.52E-06 |
| 11 | 69816959 | T | C | 0.860034 | 3.88E-12 | 1.52E-06 |
| 11 | 69816969 | A | G | 0.860034 | 3.88E-12 | 1.52E-06 |
| 11 | 69817009 | C | T | 0.860034 | 3.88E-12 | 1.52E-06 |
| 11 | 69817086 | T | G | 0.860034 | 3.88E-12 | 1.52E-06 |
| 11 | 69817310 | C | A | 0.860034 | 3.88E-12 | 1.52E-06 |
| 11 | 69817377 | A | G | 0.860034 | 3.88E-12 | 1.52E-06 |
| 11 | 69817402 | C | T | 0.860034 | 3.88E-12 | 1.52E-06 |
| 11 | 69817502 | T | C | 0.860034 | 3.88E-12 | 1.52E-06 |
| 11 | 69817548 | A | G | 0.860034 | 3.88E-12 | 1.52E-06 |
| 11 | 69817678 | C | A | 0.860034 | 3.88E-12 | 1.52E-06 |
| 11 | 69817885 | A | C | 0.860034 | 3.88E-12 | 1.52E-06 |
| 11 | 69817887 | T | C | 0.860034 | 3.88E-12 | 1.52E-06 |
| 11 | 69818004 | G | A | 0.860034 | 3.88E-12 | 1.52E-06 |
| 11 | 69818103 | A | G | 0.860034 | 3.88E-12 | 1.52E-06 |
| 11 | 69818122 | T | C | 0.860034 | 3.88E-12 | 1.52E-06 |
| 11 | 69818125 | G | A | 0.860034 | 3.88E-12 | 1.52E-06 |
| 11 | 69818142 | A | G | 0.860034 | 3.88E-12 | 1.52E-06 |
| 11 | 69818205 | A | G | 0.860034 | 3.88E-12 | 1.52E-06 |
| 11 | 69818211 | A | G | 0.860034 | 3.88E-12 | 1.52E-06 |
| 11 | 69818299 | G | A | 0.860034 | 3.88E-12 | 1.52E-06 |
| 11 | 69818600 | G | A | 0.860034 | 3.88E-12 | 1.52E-06 |
| 11 | 69818705 | C | T | 0.860034 | 3.88E-12 | 1.52E-06 |
| 11 | 69819027 | C | T | 0.860034 | 3.88E-12 | 1.52E-06 |
| 11 | 69819080 | A | C | 0.860034 | 3.88E-12 | 1.52E-06 |
| 11 | 69819089 | G | A | 0.860034 | 3.88E-12 | 1.52E-06 |
| 11 | 69819132 | T | C | 0.860034 | 3.88E-12 | 1.52E-06 |
| 11 | 69819139 | C | T | 0.860034 | 3.88E-12 | 1.52E-06 |
| 11 | 69819363 | T | C | 0.860034 | 3.88E-12 | 1.52E-06 |
| 11 | 69819545 | T | C | 0.860034 | 3.88E-12 | 1.52E-06 |
| 11 | 69820568 | A | C | 0.860034 | 3.88E-12 | 1.52E-06 |
| 11 | 69820577 | C | T | 0.860034 | 3.88E-12 | 1.52E-06 |
| 11 | 69820724 | G | A | 0.860034 | 3.88E-12 | 1.52E-06 |
| 11 | 69820925 | G | A | 0.860034 | 3.88E-12 | 1.52E-06 |
| 11 | 69821007 | G | A | 0.860034 | 3.88E-12 | 1.52E-06 |
| 11 | 69821128 | C | T | 0.860034 | 3.88E-12 | 1.52E-06 |
| 11 | 69821222 | T | C | 0.860034 | 3.88E-12 | 1.52E-06 |
| 11 | 69821232 | T | A | 0.860034 | 3.88E-12 | 1.52E-06 |
| 11 | 69821422 | A | G | 0.860034 | 3.88E-12 | 1.52E-06 |
| 11 | 69821454 | T | A | 0.860034 | 3.88E-12 | 1.52E-06 |

| | | | | | | |
|---|---|---|---|---|---|---|
| 11 | 69821619 | A | G | 0.860034 | 3.88E-12 | 1.52E-06 |
| 11 | 69821621 | G | A | 0.860034 | 3.88E-12 | 1.52E-06 |
| 11 | 69821660 | A | G | 0.860034 | 3.88E-12 | 1.52E-06 |
| 11 | 69821678 | A | G | 0.860034 | 3.88E-12 | 1.52E-06 |
| 11 | 69821751 | C | T | 0.860034 | 3.88E-12 | 1.52E-06 |
| 11 | 69821857 | G | C | 0.860034 | 3.88E-12 | 1.52E-06 |
| 11 | 69821878 | G | A | 0.860034 | 3.88E-12 | 1.52E-06 |
| 11 | 69822043 | G | T | 0.860034 | 3.88E-12 | 1.52E-06 |
| 11 | 69822242 | T | C | 0.860034 | 3.88E-12 | 1.52E-06 |
| 11 | 69822275 | G | A | 0.860034 | 3.88E-12 | 1.52E-06 |
| 11 | 69822316 | C | T | 0.860034 | 3.88E-12 | 1.52E-06 |
| 11 | 69822325 | T | C | 0.860034 | 3.88E-12 | 1.52E-06 |
| 11 | 69825548 | T | C | 0.860034 | 3.88E-12 | 1.52E-06 |
| 11 | 69825818 | A | G | 0.860034 | 3.88E-12 | 1.52E-06 |
| 11 | 69903084 | A | C | 0.884486 | 3.96E-12 | 1.55E-06 |
| 11 | 64110785 | C | T | 0.847386 | 4.18E-12 | 1.64E-06 |
| 11 | 69817122 | G | A | 0.855818 | 4.54E-12 | 1.78E-06 |
| 11 | 66095760 | T | C | 0.865936 | 5.08E-12 | 1.99E-06 |
| 11 | 64552667 | T | C | 0.869309 | 5.45E-12 | 2.13E-06 |
| 11 | 64552785 | A | G | 0.869309 | 5.45E-12 | 2.13E-06 |
| 11 | 64552866 | G | A | 0.868465 | 5.78E-12 | 2.27E-06 |
| 11 | 64057169 | A | G | 0.846543 | 5.81E-12 | 2.28E-06 |
| 11 | 64057364 | C | T | 0.846543 | 5.81E-12 | 2.28E-06 |
| 11 | 64071025 | A | G | 0.846543 | 5.81E-12 | 2.28E-06 |
| 11 | 64071318 | A | G | 0.846543 | 5.81E-12 | 2.28E-06 |
| 11 | 64071423 | G | A | 0.846543 | 5.81E-12 | 2.28E-06 |
| 11 | 64072054 | T | C | 0.846543 | 5.81E-12 | 2.28E-06 |
| 11 | 64072105 | G | A | 0.846543 | 5.81E-12 | 2.28E-06 |
| 11 | 64072157 | T | G | 0.846543 | 5.81E-12 | 2.28E-06 |
| 11 | 64072527 | G | A | 0.846543 | 5.81E-12 | 2.28E-06 |
| 11 | 64072653 | G | A | 0.846543 | 5.81E-12 | 2.28E-06 |
| 11 | 64072714 | G | A | 0.846543 | 5.81E-12 | 2.28E-06 |
| 11 | 64072783 | T | G | 0.846543 | 5.81E-12 | 2.28E-06 |
| 11 | 64072803 | C | A | 0.846543 | 5.81E-12 | 2.28E-06 |
| 11 | 64072950 | T | C | 0.846543 | 5.81E-12 | 2.28E-06 |
| 11 | 64072990 | C | A | 0.846543 | 5.81E-12 | 2.28E-06 |
| 11 | 64073003 | C | T | 0.846543 | 5.81E-12 | 2.28E-06 |
| 11 | 64073190 | A | T | 0.846543 | 5.81E-12 | 2.28E-06 |
| 11 | 64073234 | T | C | 0.846543 | 5.81E-12 | 2.28E-06 |
| 11 | 64073378 | C | T | 0.846543 | 5.81E-12 | 2.28E-06 |
| 11 | 64073954 | C | A | 0.846543 | 5.81E-12 | 2.28E-06 |
| 11 | 64074008 | T | A | 0.846543 | 5.81E-12 | 2.28E-06 |
| 11 | 64074518 | T | C | 0.846543 | 5.81E-12 | 2.28E-06 |
| 11 | 64075593 | T | C | 0.846543 | 5.81E-12 | 2.28E-06 |
| 11 | 64075635 | G | A | 0.846543 | 5.81E-12 | 2.28E-06 |
| 11 | 64076591 | T | C | 0.846543 | 5.81E-12 | 2.28E-06 |
| 11 | 64076683 | A | T | 0.846543 | 5.81E-12 | 2.28E-06 |
| 11 | 64076858 | C | T | 0.846543 | 5.81E-12 | 2.28E-06 |
| 11 | 64076950 | C | G | 0.846543 | 5.81E-12 | 2.28E-06 |
| 11 | 64077124 | G | A | 0.846543 | 5.81E-12 | 2.28E-06 |
| 11 | 64077365 | C | T | 0.846543 | 5.81E-12 | 2.28E-06 |
| 11 | 64077405 | A | G | 0.846543 | 5.81E-12 | 2.28E-06 |
| 11 | 64077517 | C | T | 0.846543 | 5.81E-12 | 2.28E-06 |
| 11 | 64077530 | G | A | 0.846543 | 5.81E-12 | 2.28E-06 |
| 11 | 64077607 | T | C | 0.846543 | 5.81E-12 | 2.28E-06 |
| 11 | 64084652 | C | T | 0.846543 | 5.81E-12 | 2.28E-06 |
| 11 | 64090748 | A | T | 0.846543 | 5.81E-12 | 2.28E-06 |
| 11 | 64096672 | T | C | 0.846543 | 5.81E-12 | 2.28E-06 |
| 11 | 64097209 | A | G | 0.846543 | 5.81E-12 | 2.28E-06 |
| 11 | 64097495 | C | A | 0.846543 | 5.81E-12 | 2.28E-06 |
| 11 | 64098445 | G | A | 0.846543 | 5.81E-12 | 2.28E-06 |
| 11 | 64100779 | A | C | 0.846543 | 5.81E-12 | 2.28E-06 |
| 11 | 66055579 | A | G | 0.86425 | 6.60E-12 | 2.59E-06 |
| 11 | 69905732 | C | T | 0.869309 | 7.05E-12 | 2.77E-06 |
| 11 | 69902762 | C | T | 0.885329 | 7.11E-12 | 2.79E-06 |
| 11 | 69902764 | T | G | 0.885329 | 7.11E-12 | 2.79E-06 |
| 11 | 69903121 | C | T | 0.885329 | 7.11E-12 | 2.79E-06 |

| | | | | | | |
|---|---|---|---|---|---|---|
| 11 | 69902983 | C | G | 0.886172 | 7.29E-12 | 2.86E-06 |
| 11 | 69903037 | A | G | 0.886172 | 7.29E-12 | 2.86E-06 |
| 11 | 69807900 | T | C | 0.860877 | 7.66E-12 | 3.00E-06 |
| 11 | 69807911 | A | G | 0.860877 | 7.66E-12 | 3.00E-06 |
| 11 | 69807966 | C | T | 0.860877 | 7.66E-12 | 3.00E-06 |
| 11 | 69807985 | T | C | 0.860877 | 7.66E-12 | 3.00E-06 |
| 11 | 69808162 | A | G | 0.860877 | 7.66E-12 | 3.00E-06 |
| 11 | 69808188 | G | A | 0.860877 | 7.66E-12 | 3.00E-06 |
| 11 | 69808198 | T | C | 0.860877 | 7.66E-12 | 3.00E-06 |
| 11 | 69808771 | G | A | 0.860877 | 7.66E-12 | 3.00E-06 |
| 11 | 69815110 | A | G | 0.860877 | 7.66E-12 | 3.00E-06 |
| 11 | 69815195 | T | A | 0.860877 | 7.66E-12 | 3.00E-06 |
| 11 | 69815264 | G | A | 0.860877 | 7.66E-12 | 3.00E-06 |
| 11 | 69815345 | T | C | 0.860877 | 7.66E-12 | 3.00E-06 |
| 11 | 69815376 | G | A | 0.860877 | 7.66E-12 | 3.00E-06 |
| 11 | 69815452 | C | T | 0.860877 | 7.66E-12 | 3.00E-06 |
| 11 | 69815485 | G | A | 0.860877 | 7.66E-12 | 3.00E-06 |
| 11 | 69815750 | A | G | 0.860877 | 7.66E-12 | 3.00E-06 |
| 11 | 63806873 | C | T | 0.915683 | 8.49E-12 | 3.33E-06 |
| 11 | 66060508 | T | C | 0.865093 | 8.66E-12 | 3.40E-06 |
| 11 | 66060989 | T | C | 0.865093 | 8.66E-12 | 3.40E-06 |
| 11 | 66061675 | T | C | 0.865093 | 8.66E-12 | 3.40E-06 |
| 11 | 66062765 | T | C | 0.865093 | 8.66E-12 | 3.40E-06 |
| 11 | 66064356 | C | T | 0.865093 | 8.66E-12 | 3.40E-06 |
| 11 | 66066018 | C | A | 0.865093 | 8.66E-12 | 3.40E-06 |
| 11 | 66067152 | T | G | 0.865093 | 8.66E-12 | 3.40E-06 |
| 11 | 66067162 | A | G | 0.865093 | 8.66E-12 | 3.40E-06 |
| 11 | 66067379 | C | T | 0.865093 | 8.66E-12 | 3.40E-06 |
| 11 | 66067449 | A | G | 0.865093 | 8.66E-12 | 3.40E-06 |
| 11 | 66067499 | C | T | 0.865093 | 8.66E-12 | 3.40E-06 |
| 11 | 66067617 | A | G | 0.865093 | 8.66E-12 | 3.40E-06 |
| 11 | 66067689 | C | T | 0.865093 | 8.66E-12 | 3.40E-06 |
| 11 | 66068337 | T | G | 0.865093 | 8.66E-12 | 3.40E-06 |
| 11 | 66103159 | A | C | 0.865093 | 8.66E-12 | 3.40E-06 |
| 11 | 66110157 | A | G | 0.865093 | 8.66E-12 | 3.40E-06 |
| 11 | 66110228 | A | C | 0.865093 | 8.66E-12 | 3.40E-06 |
| 11 | 66114258 | T | C | 0.865093 | 8.66E-12 | 3.40E-06 |
| 11 | 69862332 | G | T | 0.884486 | 8.89E-12 | 3.48E-06 |
| 11 | 69863460 | C | G | 0.884486 | 8.89E-12 | 3.48E-06 |
| 11 | 69863701 | G | A | 0.884486 | 8.89E-12 | 3.48E-06 |
| 11 | 69864861 | T | C | 0.884486 | 8.89E-12 | 3.48E-06 |
| 11 | 69865076 | G | A | 0.884486 | 8.89E-12 | 3.48E-06 |
| 11 | 69865373 | T | C | 0.884486 | 8.89E-12 | 3.48E-06 |
| 11 | 69865958 | A | G | 0.884486 | 8.89E-12 | 3.48E-06 |
| 11 | 69866041 | T | C | 0.884486 | 8.89E-12 | 3.48E-06 |
| 11 | 69866824 | C | T | 0.884486 | 8.89E-12 | 3.48E-06 |
| 11 | 69867518 | T | G | 0.884486 | 8.89E-12 | 3.48E-06 |
| 11 | 69867561 | A | G | 0.884486 | 8.89E-12 | 3.48E-06 |
| 11 | 69868146 | G | A | 0.884486 | 8.89E-12 | 3.48E-06 |
| 11 | 69868275 | C | T | 0.884486 | 8.89E-12 | 3.48E-06 |
| 11 | 69868479 | T | C | 0.884486 | 8.89E-12 | 3.48E-06 |
| 11 | 69868940 | G | T | 0.884486 | 8.89E-12 | 3.48E-06 |
| 11 | 69868950 | G | T | 0.884486 | 8.89E-12 | 3.48E-06 |
| 11 | 69869242 | T | C | 0.884486 | 8.89E-12 | 3.48E-06 |
| 11 | 69869905 | A | G | 0.884486 | 8.89E-12 | 3.48E-06 |
| 11 | 69870447 | G | T | 0.884486 | 8.89E-12 | 3.48E-06 |
| 11 | 69870618 | G | A | 0.884486 | 8.89E-12 | 3.48E-06 |
| 11 | 69870652 | T | A | 0.884486 | 8.89E-12 | 3.48E-06 |
| 11 | 69870656 | T | C | 0.884486 | 8.89E-12 | 3.48E-06 |
| 11 | 69870685 | A | C | 0.884486 | 8.89E-12 | 3.48E-06 |
| 11 | 69870705 | T | C | 0.884486 | 8.89E-12 | 3.48E-06 |
| 11 | 69870858 | G | A | 0.884486 | 8.89E-12 | 3.48E-06 |
| 11 | 69870906 | C | T | 0.884486 | 8.89E-12 | 3.48E-06 |
| 11 | 69870981 | A | C | 0.884486 | 8.89E-12 | 3.48E-06 |
| 11 | 69870997 | G | A | 0.884486 | 8.89E-12 | 3.48E-06 |
| 11 | 69871068 | C | T | 0.884486 | 8.89E-12 | 3.48E-06 |
| 11 | 69871126 | T | A | 0.884486 | 8.89E-12 | 3.48E-06 |

| | | | | | | |
|---|---|---|---|---|---|---|
| 11 | 69871156 | T | C | 0.884486 | 8.89E-12 | 3.48E-06 |
| 11 | 69871178 | T | C | 0.884486 | 8.89E-12 | 3.48E-06 |
| 11 | 69871181 | T | C | 0.884486 | 8.89E-12 | 3.48E-06 |
| 11 | 69871257 | T | C | 0.884486 | 8.89E-12 | 3.48E-06 |
| 11 | 69871310 | C | T | 0.884486 | 8.89E-12 | 3.48E-06 |
| 11 | 69871420 | C | G | 0.884486 | 8.89E-12 | 3.48E-06 |
| 11 | 69871431 | T | C | 0.884486 | 8.89E-12 | 3.48E-06 |
| 11 | 69871510 | T | C | 0.884486 | 8.89E-12 | 3.48E-06 |
| 11 | 69871715 | A | G | 0.884486 | 8.89E-12 | 3.48E-06 |
| 11 | 69873344 | C | A | 0.884486 | 8.89E-12 | 3.48E-06 |
| 11 | 69873421 | G | A | 0.884486 | 8.89E-12 | 3.48E-06 |
| 11 | 69873433 | C | T | 0.884486 | 8.89E-12 | 3.48E-06 |
| 11 | 69873444 | C | T | 0.884486 | 8.89E-12 | 3.48E-06 |
| 11 | 69873878 | T | A | 0.884486 | 8.89E-12 | 3.48E-06 |
| 11 | 69873885 | A | T | 0.884486 | 8.89E-12 | 3.48E-06 |
| 11 | 69873927 | T | A | 0.884486 | 8.89E-12 | 3.48E-06 |
| 11 | 69874067 | A | G | 0.884486 | 8.89E-12 | 3.48E-06 |
| 11 | 69874100 | T | A | 0.884486 | 8.89E-12 | 3.48E-06 |
| 11 | 69874112 | C | T | 0.884486 | 8.89E-12 | 3.48E-06 |
| 11 | 69874125 | A | G | 0.884486 | 8.89E-12 | 3.48E-06 |
| 11 | 69874187 | G | A | 0.884486 | 8.89E-12 | 3.48E-06 |
| 11 | 69874209 | G | A | 0.884486 | 8.89E-12 | 3.48E-06 |
| 11 | 69874250 | G | A | 0.884486 | 8.89E-12 | 3.48E-06 |
| 11 | 69874283 | A | G | 0.884486 | 8.89E-12 | 3.48E-06 |
| 11 | 69874414 | A | G | 0.884486 | 8.89E-12 | 3.48E-06 |
| 11 | 69875599 | A | G | 0.884486 | 8.89E-12 | 3.48E-06 |
| 11 | 69875964 | C | T | 0.884486 | 8.89E-12 | 3.48E-06 |
| 11 | 69876492 | T | A | 0.884486 | 8.89E-12 | 3.48E-06 |
| 11 | 69876496 | A | G | 0.884486 | 8.89E-12 | 3.48E-06 |
| 11 | 69876587 | C | T | 0.884486 | 8.89E-12 | 3.48E-06 |
| 11 | 69876635 | G | C | 0.884486 | 8.89E-12 | 3.48E-06 |
| 11 | 69876716 | A | G | 0.884486 | 8.89E-12 | 3.48E-06 |
| 11 | 69876828 | C | A | 0.884486 | 8.89E-12 | 3.48E-06 |
| 11 | 69877054 | G | C | 0.884486 | 8.89E-12 | 3.48E-06 |
| 11 | 69877075 | T | C | 0.884486 | 8.89E-12 | 3.48E-06 |
| 11 | 69877142 | A | G | 0.884486 | 8.89E-12 | 3.48E-06 |
| 11 | 69877146 | T | C | 0.884486 | 8.89E-12 | 3.48E-06 |
| 11 | 69877186 | C | T | 0.884486 | 8.89E-12 | 3.48E-06 |
| 11 | 69877240 | C | T | 0.884486 | 8.89E-12 | 3.48E-06 |
| 11 | 69877255 | A | G | 0.884486 | 8.89E-12 | 3.48E-06 |
| 11 | 69877277 | T | C | 0.884486 | 8.89E-12 | 3.48E-06 |
| 11 | 69877281 | G | A | 0.884486 | 8.89E-12 | 3.48E-06 |
| 11 | 69877303 | C | T | 0.884486 | 8.89E-12 | 3.48E-06 |
| 11 | 69877364 | G | A | 0.884486 | 8.89E-12 | 3.48E-06 |
| 11 | 69877429 | C | T | 0.884486 | 8.89E-12 | 3.48E-06 |
| 11 | 69877553 | G | C | 0.884486 | 8.89E-12 | 3.48E-06 |
| 11 | 69877592 | A | T | 0.884486 | 8.89E-12 | 3.48E-06 |
| 11 | 69877601 | C | T | 0.884486 | 8.89E-12 | 3.48E-06 |
| 11 | 69877612 | T | C | 0.884486 | 8.89E-12 | 3.48E-06 |
| 11 | 69877752 | A | T | 0.884486 | 8.89E-12 | 3.48E-06 |
| 11 | 69877879 | T | A | 0.884486 | 8.89E-12 | 3.48E-06 |
| 11 | 69878137 | T | G | 0.884486 | 8.89E-12 | 3.48E-06 |
| 11 | 69878413 | G | T | 0.884486 | 8.89E-12 | 3.48E-06 |
| 11 | 69878667 | T | C | 0.884486 | 8.89E-12 | 3.48E-06 |
| 11 | 69878743 | T | C | 0.884486 | 8.89E-12 | 3.48E-06 |
| 11 | 69878753 | T | C | 0.884486 | 8.89E-12 | 3.48E-06 |
| 11 | 69878768 | C | T | 0.884486 | 8.89E-12 | 3.48E-06 |
| 11 | 69878807 | C | T | 0.884486 | 8.89E-12 | 3.48E-06 |
| 11 | 69879077 | C | T | 0.884486 | 8.89E-12 | 3.48E-06 |
| 11 | 69879099 | T | A | 0.884486 | 8.89E-12 | 3.48E-06 |
| 11 | 69879101 | C | T | 0.884486 | 8.89E-12 | 3.48E-06 |
| 11 | 69879134 | C | T | 0.884486 | 8.89E-12 | 3.48E-06 |
| 11 | 69879141 | A | G | 0.884486 | 8.89E-12 | 3.48E-06 |
| 11 | 69879167 | C | T | 0.884486 | 8.89E-12 | 3.48E-06 |
| 11 | 69879224 | G | A | 0.884486 | 8.89E-12 | 3.48E-06 |
| 11 | 69879344 | T | G | 0.884486 | 8.89E-12 | 3.48E-06 |
| 11 | 69879365 | C | T | 0.884486 | 8.89E-12 | 3.48E-06 |

| | | | | | | |
|---|---|---|---|---|---|---|
| 11 | 69879477 | T | C | 0.884486 | 8.89E-12 | 3.48E-06 |
| 11 | 69879630 | A | G | 0.884486 | 8.89E-12 | 3.48E-06 |
| 11 | 69879661 | T | C | 0.884486 | 8.89E-12 | 3.48E-06 |
| 11 | 69879662 | G | A | 0.884486 | 8.89E-12 | 3.48E-06 |
| 11 | 69879674 | C | T | 0.884486 | 8.89E-12 | 3.48E-06 |
| 11 | 69879678 | G | A | 0.884486 | 8.89E-12 | 3.48E-06 |
| 11 | 69879704 | A | C | 0.884486 | 8.89E-12 | 3.48E-06 |
| 11 | 69879748 | C | T | 0.884486 | 8.89E-12 | 3.48E-06 |
| 11 | 69879888 | G | C | 0.884486 | 8.89E-12 | 3.48E-06 |
| 11 | 69879906 | T | C | 0.884486 | 8.89E-12 | 3.48E-06 |
| 11 | 69879963 | T | A | 0.884486 | 8.89E-12 | 3.48E-06 |
| 11 | 69879978 | C | T | 0.884486 | 8.89E-12 | 3.48E-06 |
| 11 | 69879994 | G | A | 0.884486 | 8.89E-12 | 3.48E-06 |
| 11 | 69880088 | A | T | 0.884486 | 8.89E-12 | 3.48E-06 |
| 11 | 69880141 | G | T | 0.884486 | 8.89E-12 | 3.48E-06 |
| 11 | 69880192 | A | G | 0.884486 | 8.89E-12 | 3.48E-06 |
| 11 | 69880231 | G | A | 0.884486 | 8.89E-12 | 3.48E-06 |
| 11 | 69880393 | T | C | 0.884486 | 8.89E-12 | 3.48E-06 |
| 11 | 69880394 | G | A | 0.884486 | 8.89E-12 | 3.48E-06 |
| 11 | 69880499 | T | C | 0.884486 | 8.89E-12 | 3.48E-06 |
| 11 | 69880743 | G | A | 0.884486 | 8.89E-12 | 3.48E-06 |
| 11 | 69881275 | G | T | 0.884486 | 8.89E-12 | 3.48E-06 |
| 11 | 69881327 | T | C | 0.884486 | 8.89E-12 | 3.48E-06 |
| 11 | 69884315 | A | G | 0.884486 | 8.89E-12 | 3.48E-06 |
| 11 | 69884348 | T | C | 0.884486 | 8.89E-12 | 3.48E-06 |
| 11 | 69884356 | A | G | 0.884486 | 8.89E-12 | 3.48E-06 |
| 11 | 69884431 | C | T | 0.884486 | 8.89E-12 | 3.48E-06 |
| 11 | 69884932 | G | A | 0.884486 | 8.89E-12 | 3.48E-06 |
| 11 | 69886248 | T | C | 0.884486 | 8.89E-12 | 3.48E-06 |
| 11 | 69886296 | T | C | 0.884486 | 8.89E-12 | 3.48E-06 |
| 11 | 69886593 | A | G | 0.884486 | 8.89E-12 | 3.48E-06 |
| 11 | 69887413 | A | G | 0.884486 | 8.89E-12 | 3.48E-06 |
| 11 | 69889265 | T | C | 0.884486 | 8.89E-12 | 3.48E-06 |
| 11 | 69889316 | A | G | 0.884486 | 8.89E-12 | 3.48E-06 |
| 11 | 69889347 | T | C | 0.884486 | 8.89E-12 | 3.48E-06 |
| 11 | 69889531 | C | T | 0.884486 | 8.89E-12 | 3.48E-06 |
| 11 | 69889727 | C | T | 0.884486 | 8.89E-12 | 3.48E-06 |
| 11 | 69889995 | A | G | 0.884486 | 8.89E-12 | 3.48E-06 |
| 11 | 69890129 | A | T | 0.884486 | 8.89E-12 | 3.48E-06 |
| 11 | 69890171 | G | A | 0.884486 | 8.89E-12 | 3.48E-06 |
| 11 | 69890319 | C | T | 0.884486 | 8.89E-12 | 3.48E-06 |
| 11 | 69890436 | T | G | 0.884486 | 8.89E-12 | 3.48E-06 |
| 11 | 69891358 | A | G | 0.884486 | 8.89E-12 | 3.48E-06 |
| 11 | 69892207 | G | A | 0.884486 | 8.89E-12 | 3.48E-06 |
| 11 | 69892313 | G | A | 0.884486 | 8.89E-12 | 3.48E-06 |
| 11 | 63939103 | G | A | 0.836425 | 8.96E-12 | 3.51E-06 |
| 11 | 63939185 | G | T | 0.836425 | 8.96E-12 | 3.51E-06 |
| 11 | 69876612 | C | T | 0.885329 | 9.02E-12 | 3.54E-06 |
| 11 | 69876613 | A | C | 0.885329 | 9.02E-12 | 3.54E-06 |
| 11 | 69876614 | T | C | 0.885329 | 9.02E-12 | 3.54E-06 |
| 11 | 69878371 | T | C | 0.885329 | 9.02E-12 | 3.54E-06 |
| 11 | 69878730 | C | T | 0.885329 | 9.02E-12 | 3.54E-06 |
| 11 | 69879305 | T | C | 0.885329 | 9.02E-12 | 3.54E-06 |
| 11 | 69879529 | T | C | 0.885329 | 9.02E-12 | 3.54E-06 |
| 11 | 69879568 | T | C | 0.885329 | 9.02E-12 | 3.54E-06 |
| 11 | 69879570 | T | A | 0.885329 | 9.02E-12 | 3.54E-06 |
| 11 | 69879571 | A | T | 0.885329 | 9.02E-12 | 3.54E-06 |
| 11 | 69879624 | G | C | 0.885329 | 9.02E-12 | 3.54E-06 |
| 11 | 69879817 | C | A | 0.885329 | 9.02E-12 | 3.54E-06 |
| 11 | 69879836 | T | A | 0.885329 | 9.02E-12 | 3.54E-06 |
| 11 | 69882682 | A | G | 0.885329 | 9.02E-12 | 3.54E-06 |
| 11 | 69885365 | G | A | 0.885329 | 9.02E-12 | 3.54E-06 |
| 11 | 69888661 | C | G | 0.885329 | 9.02E-12 | 3.54E-06 |
| 11 | 69889163 | A | G | 0.885329 | 9.02E-12 | 3.54E-06 |
| 11 | 64090908 | A | G | 0.844013 | 9.05E-12 | 3.55E-06 |
| 11 | 66073329 | A | C | 0.865936 | 9.79E-12 | 3.84E-06 |
| 11 | 66073819 | C | A | 0.865936 | 9.79E-12 | 3.84E-06 |

| | | | | | | |
|---|---|---|---|---|---|---|
| 11 | 66073857 | C | T | 0.865936 | 9.79E-12 | 3.84E-06 |
| 11 | 64078821 | A | G | 0.8457 | 1.02E-11 | 4.01E-06 |
| 11 | 64078826 | T | C | 0.8457 | 1.02E-11 | 4.01E-06 |
| 11 | 64078829 | G | A | 0.8457 | 1.02E-11 | 4.01E-06 |
| 11 | 64078911 | T | C | 0.8457 | 1.02E-11 | 4.01E-06 |
| 11 | 64079109 | C | G | 0.8457 | 1.02E-11 | 4.01E-06 |
| 11 | 64079143 | C | T | 0.8457 | 1.02E-11 | 4.01E-06 |
| 11 | 64164855 | A | C | 0.844013 | 1.07E-11 | 4.19E-06 |
| 11 | 69874320 | C | A | 0.883642 | 1.15E-11 | 4.50E-06 |
| 11 | 64108428 | C | T | 0.847386 | 1.21E-11 | 4.76E-06 |
| 11 | 69847557 | T | C | 0.870995 | 1.35E-11 | 5.27E-06 |
| 11 | 69847575 | G | T | 0.870995 | 1.35E-11 | 5.27E-06 |
| 11 | 69898760 | T | G | 0.875211 | 1.43E-11 | 5.61E-06 |
| 11 | 64120003 | T | A | 0.844857 | 2.38E-11 | 9.34E-06 |
| 11 | 64123214 | A | G | 0.844857 | 2.38E-11 | 9.34E-06 |
| 11 | 64123217 | T | A | 0.844857 | 2.38E-11 | 9.34E-06 |
| 11 | 64123241 | T | C | 0.844857 | 2.38E-11 | 9.34E-06 |
| 11 | 64123360 | G | A | 0.844857 | 2.38E-11 | 9.34E-06 |
| 11 | 64124944 | C | G | 0.844857 | 2.38E-11 | 9.34E-06 |
| 11 | 64125333 | C | T | 0.844857 | 2.38E-11 | 9.34E-06 |
| 11 | 64125335 | T | C | 0.844857 | 2.38E-11 | 9.34E-06 |
| 11 | 64125980 | A | C | 0.844857 | 2.38E-11 | 9.34E-06 |
| 11 | 64125995 | A | C | 0.844857 | 2.38E-11 | 9.34E-06 |
| 11 | 64126097 | G | T | 0.844857 | 2.38E-11 | 9.34E-06 |
| 11 | 64126414 | C | T | 0.844857 | 2.38E-11 | 9.34E-06 |
| 11 | 64126524 | T | C | 0.844857 | 2.38E-11 | 9.34E-06 |
| 11 | 64126859 | C | T | 0.844857 | 2.38E-11 | 9.34E-06 |
| 11 | 64130819 | A | G | 0.844857 | 2.38E-11 | 9.34E-06 |
| 11 | 64132214 | G | A | 0.844857 | 2.38E-11 | 9.34E-06 |
| 11 | 64139242 | C | T | 0.844857 | 2.38E-11 | 9.34E-06 |
| 11 | 64144312 | G | T | 0.844857 | 2.38E-11 | 9.34E-06 |
| 11 | 64155172 | T | C | 0.844857 | 2.38E-11 | 9.34E-06 |
| 11 | 64155739 | C | T | 0.844857 | 2.38E-11 | 9.34E-06 |
| 11 | 64156037 | T | C | 0.844857 | 2.38E-11 | 9.34E-06 |
| 11 | 64158636 | T | C | 0.844857 | 2.38E-11 | 9.34E-06 |
| 11 | 64160080 | C | G | 0.844857 | 2.38E-11 | 9.34E-06 |
| 11 | 64170575 | A | G | 0.844857 | 2.38E-11 | 9.34E-06 |
| 11 | 64062528 | T | A | 0.874368 | 2.43E-11 | 9.51E-06 |
| 11 | 64062560 | C | T | 0.874368 | 2.43E-11 | 9.51E-06 |
| 11 | 69865919 | T | C | 0.882799 | 2.65E-11 | 1.04E-05 |
| 11 | 64073275 | G | A | 0.833895 | 3.30E-11 | 1.29E-05 |
| 11 | 69825649 | A | G | 0.863406 | 3.66E-11 | 1.43E-05 |
| 11 | 66096991 | C | T | 0.870152 | 3.89E-11 | 1.52E-05 |
| 11 | 69918722 | G | A | 0.88027 | 3.89E-11 | 1.52E-05 |
| 11 | 69918755 | A | T | 0.88027 | 3.89E-11 | 1.52E-05 |
| 11 | 69918760 | G | A | 0.88027 | 3.89E-11 | 1.52E-05 |
| 11 | 69918775 | G | A | 0.88027 | 3.89E-11 | 1.52E-05 |
| 11 | 69918870 | A | C | 0.88027 | 3.89E-11 | 1.52E-05 |
| 11 | 69918914 | C | T | 0.88027 | 3.89E-11 | 1.52E-05 |
| 11 | 69919036 | T | C | 0.88027 | 3.89E-11 | 1.52E-05 |
| 11 | 69919192 | C | T | 0.88027 | 3.89E-11 | 1.52E-05 |
| 11 | 69919203 | G | A | 0.88027 | 3.89E-11 | 1.52E-05 |
| 11 | 69919224 | G | C | 0.88027 | 3.89E-11 | 1.52E-05 |
| 11 | 69919249 | A | G | 0.88027 | 3.89E-11 | 1.52E-05 |
| 11 | 69919425 | T | A | 0.88027 | 3.89E-11 | 1.52E-05 |
| 11 | 69919545 | A | G | 0.88027 | 3.89E-11 | 1.52E-05 |
| 11 | 69919556 | A | G | 0.88027 | 3.89E-11 | 1.52E-05 |
| 11 | 69919703 | C | T | 0.88027 | 3.89E-11 | 1.52E-05 |
| 11 | 69919769 | C | G | 0.88027 | 3.89E-11 | 1.52E-05 |
| 11 | 69919828 | C | A | 0.88027 | 3.89E-11 | 1.52E-05 |
| 11 | 69919918 | G | A | 0.88027 | 3.89E-11 | 1.52E-05 |
| 11 | 69919979 | C | A | 0.88027 | 3.89E-11 | 1.52E-05 |
| 11 | 69920006 | T | A | 0.88027 | 3.89E-11 | 1.52E-05 |
| 11 | 69920130 | T | C | 0.88027 | 3.89E-11 | 1.52E-05 |
| 11 | 69921443 | T | G | 0.88027 | 3.89E-11 | 1.52E-05 |
| 11 | 69922047 | C | T | 0.88027 | 3.89E-11 | 1.52E-05 |
| 11 | 69922065 | A | T | 0.88027 | 3.89E-11 | 1.52E-05 |

| | | | | | | |
|---|---|---|---|---|---|---|
| 11 | 69922430 | A | T | 0.88027 | 3.89E-11 | 1.52E-05 |
| 11 | 69922701 | C | A | 0.88027 | 3.89E-11 | 1.52E-05 |
| 11 | 69924421 | T | C | 0.88027 | 3.89E-11 | 1.52E-05 |
| 11 | 69924549 | C | T | 0.88027 | 3.89E-11 | 1.52E-05 |
| 11 | 69924944 | C | A | 0.88027 | 3.89E-11 | 1.52E-05 |
| 11 | 69925081 | A | C | 0.88027 | 3.89E-11 | 1.52E-05 |
| 11 | 69925320 | C | T | 0.88027 | 3.89E-11 | 1.52E-05 |
| 11 | 64979815 | A | G | 0.867622 | 6.39E-11 | 2.51E-05 |
| 11 | 64983289 | T | C | 0.867622 | 6.39E-11 | 2.51E-05 |
| 11 | 64987371 | T | A | 0.867622 | 6.39E-11 | 2.51E-05 |
| 11 | 64994256 | A | T | 0.867622 | 6.39E-11 | 2.51E-05 |
| 11 | 64994516 | A | G | 0.867622 | 6.39E-11 | 2.51E-05 |
| 11 | 64994811 | T | C | 0.867622 | 6.39E-11 | 2.51E-05 |
| 11 | 64994942 | T | C | 0.867622 | 6.39E-11 | 2.51E-05 |
| 11 | 64995490 | A | G | 0.867622 | 6.39E-11 | 2.51E-05 |
| 11 | 64997215 | T | C | 0.867622 | 6.39E-11 | 2.51E-05 |
| 11 | 64997471 | T | G | 0.867622 | 6.39E-11 | 2.51E-05 |
| 11 | 64998157 | A | G | 0.867622 | 6.39E-11 | 2.51E-05 |
| 11 | 64998652 | G | A | 0.867622 | 6.39E-11 | 2.51E-05 |
| 11 | 64998965 | A | G | 0.867622 | 6.39E-11 | 2.51E-05 |
| 11 | 64999074 | C | A | 0.867622 | 6.39E-11 | 2.51E-05 |
| 11 | 64999215 | G | A | 0.867622 | 6.39E-11 | 2.51E-05 |
| 11 | 64999550 | T | C | 0.867622 | 6.39E-11 | 2.51E-05 |
| 11 | 64999601 | C | A | 0.867622 | 6.39E-11 | 2.51E-05 |
| 11 | 65000704 | G | T | 0.867622 | 6.39E-11 | 2.51E-05 |
| 11 | 65026036 | T | C | 0.867622 | 6.39E-11 | 2.51E-05 |
| 11 | 65028155 | C | T | 0.867622 | 6.39E-11 | 2.51E-05 |
| 11 | 65029321 | A | G | 0.867622 | 6.39E-11 | 2.51E-05 |
| 11 | 65029326 | A | C | 0.867622 | 6.39E-11 | 2.51E-05 |
| 11 | 65030054 | T | C | 0.867622 | 6.39E-11 | 2.51E-05 |
| 11 | 65031192 | T | C | 0.867622 | 6.39E-11 | 2.51E-05 |
| 11 | 65062814 | T | C | 0.867622 | 6.39E-11 | 2.51E-05 |
| 11 | 65062995 | A | G | 0.867622 | 6.39E-11 | 2.51E-05 |
| 11 | 65064236 | A | G | 0.867622 | 6.39E-11 | 2.51E-05 |
| 11 | 65065071 | T | C | 0.867622 | 6.39E-11 | 2.51E-05 |
| 11 | 65065332 | G | A | 0.867622 | 6.39E-11 | 2.51E-05 |
| 11 | 65066203 | C | T | 0.867622 | 6.39E-11 | 2.51E-05 |
| 11 | 65066551 | T | C | 0.867622 | 6.39E-11 | 2.51E-05 |
| 11 | 65066930 | C | T | 0.867622 | 6.39E-11 | 2.51E-05 |
| 11 | 65067001 | C | G | 0.867622 | 6.39E-11 | 2.51E-05 |
| 11 | 65067906 | T | C | 0.867622 | 6.39E-11 | 2.51E-05 |
| 11 | 65069085 | C | T | 0.867622 | 6.39E-11 | 2.51E-05 |
| 11 | 65069505 | A | G | 0.867622 | 6.39E-11 | 2.51E-05 |
| 11 | 65069982 | A | G | 0.867622 | 6.39E-11 | 2.51E-05 |
| 11 | 65070402 | T | C | 0.867622 | 6.39E-11 | 2.51E-05 |
| 11 | 65070656 | C | G | 0.867622 | 6.39E-11 | 2.51E-05 |
| 11 | 69920888 | A | C | 0.878583 | 6.47E-11 | 2.54E-05 |
| 11 | 64059485 | A | G | 0.874368 | 6.78E-11 | 2.66E-05 |
| 11 | 64061791 | T | C | 0.874368 | 6.81E-11 | 2.67E-05 |
| 11 | 64059124 | G | A | 0.873524 | 7.08E-11 | 2.78E-05 |
| 11 | 64059309 | T | A | 0.873524 | 7.08E-11 | 2.78E-05 |
| 11 | 64060145 | T | C | 0.873524 | 7.08E-11 | 2.78E-05 |
| 11 | 64061128 | C | T | 0.873524 | 7.08E-11 | 2.78E-05 |
| 11 | 64061744 | A | G | 0.873524 | 7.08E-11 | 2.78E-05 |
| 11 | 64061846 | T | C | 0.873524 | 7.08E-11 | 2.78E-05 |
| 11 | 64062127 | C | T | 0.873524 | 7.08E-11 | 2.78E-05 |
| 11 | 64062278 | C | T | 0.873524 | 7.08E-11 | 2.78E-05 |
| 11 | 64062293 | C | T | 0.873524 | 7.08E-11 | 2.78E-05 |
| 11 | 64067951 | G | A | 0.873524 | 7.08E-11 | 2.78E-05 |
| 11 | 64068093 | A | G | 0.873524 | 7.08E-11 | 2.78E-05 |
| 11 | 64068299 | G | A | 0.873524 | 7.08E-11 | 2.78E-05 |
| 11 | 64068509 | A | G | 0.873524 | 7.08E-11 | 2.78E-05 |
| 11 | 64068525 | A | G | 0.873524 | 7.08E-11 | 2.78E-05 |
| 11 | 64068648 | T | C | 0.873524 | 7.08E-11 | 2.78E-05 |
| 11 | 64069028 | A | G | 0.873524 | 7.08E-11 | 2.78E-05 |
| 11 | 64069223 | A | G | 0.873524 | 7.08E-11 | 2.78E-05 |
| 11 | 64069291 | T | C | 0.873524 | 7.08E-11 | 2.78E-05 |

| | | | | | | |
|---|---|---|---|---|---|---|
| 11 | 64069302 | A | G | 0.873524 | 7.08E-11 | 2.78E-05 |
| 11 | 64069682 | G | A | 0.873524 | 7.08E-11 | 2.78E-05 |
| 11 | 64069700 | G | T | 0.873524 | 7.08E-11 | 2.78E-05 |
| 11 | 64069867 | A | G | 0.873524 | 7.08E-11 | 2.78E-05 |
| 11 | 64069929 | G | A | 0.873524 | 7.08E-11 | 2.78E-05 |
| 11 | 64069930 | A | G | 0.873524 | 7.08E-11 | 2.78E-05 |
| 11 | 64069935 | T | C | 0.873524 | 7.08E-11 | 2.78E-05 |
| 11 | 64070028 | A | G | 0.873524 | 7.08E-11 | 2.78E-05 |
| 11 | 64070080 | A | G | 0.873524 | 7.08E-11 | 2.78E-05 |
| 11 | 64070196 | G | A | 0.873524 | 7.08E-11 | 2.78E-05 |
| 11 | 64070365 | T | A | 0.873524 | 7.08E-11 | 2.78E-05 |
| 11 | 64070401 | G | A | 0.873524 | 7.08E-11 | 2.78E-05 |
| 11 | 64070512 | A | G | 0.873524 | 7.08E-11 | 2.78E-05 |
| 11 | 64070521 | A | G | 0.873524 | 7.08E-11 | 2.78E-05 |
| 11 | 69905782 | G | A | 0.879427 | 8.72E-11 | 3.42E-05 |
| 11 | 69905891 | T | A | 0.879427 | 8.72E-11 | 3.42E-05 |
| 11 | 69905957 | A | C | 0.879427 | 8.72E-11 | 3.42E-05 |
| 11 | 69906063 | T | C | 0.879427 | 8.72E-11 | 3.42E-05 |
| 11 | 69906228 | A | C | 0.879427 | 8.72E-11 | 3.42E-05 |
| 11 | 69906783 | G | A | 0.879427 | 8.72E-11 | 3.42E-05 |
| 11 | 69906831 | T | A | 0.879427 | 8.72E-11 | 3.42E-05 |
| 11 | 69906840 | T | C | 0.879427 | 8.72E-11 | 3.42E-05 |
| 11 | 69906850 | G | A | 0.879427 | 8.72E-11 | 3.42E-05 |
| 11 | 69907321 | G | A | 0.879427 | 8.72E-11 | 3.42E-05 |
| 11 | 69907376 | T | C | 0.879427 | 8.72E-11 | 3.42E-05 |
| 11 | 69907395 | C | T | 0.879427 | 8.72E-11 | 3.42E-05 |
| 11 | 69907541 | A | G | 0.879427 | 8.72E-11 | 3.42E-05 |
| 11 | 69908394 | A | G | 0.879427 | 8.72E-11 | 3.42E-05 |
| 11 | 69908414 | T | C | 0.879427 | 8.72E-11 | 3.42E-05 |
| 11 | 69908666 | A | T | 0.879427 | 8.72E-11 | 3.42E-05 |
| 11 | 69908939 | C | T | 0.879427 | 8.72E-11 | 3.42E-05 |
| 11 | 69909391 | T | C | 0.879427 | 8.72E-11 | 3.42E-05 |
| 11 | 69909575 | C | T | 0.879427 | 8.72E-11 | 3.42E-05 |
| 11 | 69910483 | T | C | 0.879427 | 8.72E-11 | 3.42E-05 |
| 11 | 69910751 | A | G | 0.879427 | 8.72E-11 | 3.42E-05 |
| 11 | 69910792 | T | C | 0.879427 | 8.72E-11 | 3.42E-05 |
| 11 | 69911037 | A | C | 0.879427 | 8.72E-11 | 3.42E-05 |
| 11 | 69911323 | G | C | 0.879427 | 8.72E-11 | 3.42E-05 |
| 11 | 69912541 | G | A | 0.879427 | 8.72E-11 | 3.42E-05 |
| 11 | 69912542 | A | G | 0.879427 | 8.72E-11 | 3.42E-05 |
| 11 | 69912546 | G | C | 0.879427 | 8.72E-11 | 3.42E-05 |
| 11 | 69912845 | T | C | 0.879427 | 8.72E-11 | 3.42E-05 |
| 11 | 69912865 | T | C | 0.879427 | 8.72E-11 | 3.42E-05 |
| 11 | 69913402 | G | T | 0.879427 | 8.72E-11 | 3.42E-05 |
| 11 | 69913444 | G | A | 0.879427 | 8.72E-11 | 3.42E-05 |
| 11 | 69913667 | A | G | 0.879427 | 8.72E-11 | 3.42E-05 |
| 11 | 69917119 | G | A | 0.879427 | 8.72E-11 | 3.42E-05 |
| 11 | 69917126 | T | C | 0.879427 | 8.72E-11 | 3.42E-05 |
| 11 | 69917164 | G | A | 0.879427 | 8.72E-11 | 3.42E-05 |
| 11 | 69917192 | G | A | 0.879427 | 8.72E-11 | 3.42E-05 |
| 11 | 69917212 | T | C | 0.879427 | 8.72E-11 | 3.42E-05 |
| 11 | 69917283 | T | C | 0.879427 | 8.72E-11 | 3.42E-05 |
| 11 | 64062171 | A | T | 0.872681 | 9.84E-11 | 3.86E-05 |
| 11 | 65018136 | C | T | 0.868465 | 1.09E-10 | 4.28E-05 |
| 11 | 65025933 | T | C | 0.868465 | 1.09E-10 | 4.28E-05 |
| 11 | 65029713 | T | C | 0.868465 | 1.09E-10 | 4.28E-05 |
| 11 | 65029886 | T | C | 0.868465 | 1.09E-10 | 4.28E-05 |
| 11 | 65031721 | A | C | 0.868465 | 1.09E-10 | 4.28E-05 |
| 11 | 66081666 | G | A | 0.867622 | 1.18E-10 | 4.62E-05 |
| 11 | 63835346 | T | A | 0.851602 | 1.18E-10 | 4.64E-05 |
| 11 | 71560147 | T | C | 0.935919 | 1.36E-10 | 5.32E-05 |
| 11 | 71627270 | T | C | 0.939292 | 1.64E-10 | 6.41E-05 |
| 11 | 71627452 | T | G | 0.939292 | 1.64E-10 | 6.41E-05 |
| 11 | 71627680 | G | A | 0.939292 | 1.64E-10 | 6.41E-05 |
| 11 | 71628347 | T | C | 0.939292 | 1.64E-10 | 6.41E-05 |
| 11 | 71628475 | G | C | 0.939292 | 1.64E-10 | 6.41E-05 |
| 11 | 71628793 | T | C | 0.939292 | 1.64E-10 | 6.41E-05 |

| | | | | | | |
|---|---|---|---|---|---|---|
| 11 | 71629057 | T | G | 0.939292 | 1.64E-10 | 6.41E-05 |
| 11 | 71629925 | C | A | 0.939292 | 1.64E-10 | 6.41E-05 |
| 11 | 71630239 | A | G | 0.939292 | 1.64E-10 | 6.41E-05 |
| 11 | 71630361 | C | T | 0.939292 | 1.64E-10 | 6.41E-05 |
| 11 | 71631069 | C | A | 0.939292 | 1.64E-10 | 6.41E-05 |
| 11 | 71631443 | C | T | 0.939292 | 1.64E-10 | 6.41E-05 |
| 11 | 69531064 | T | A | 0.814503 | 1.69E-10 | 6.62E-05 |
| 11 | 69531092 | A | G | 0.814503 | 1.69E-10 | 6.62E-05 |
| 11 | 69531101 | G | A | 0.814503 | 1.69E-10 | 6.62E-05 |
| 11 | 65013075 | C | A | 0.866779 | 1.71E-10 | 6.71E-05 |
| 11 | 65014976 | C | G | 0.866779 | 1.71E-10 | 6.71E-05 |
| 11 | 71626435 | A | T | 0.938449 | 2.06E-10 | 8.09E-05 |
| 11 | 71626529 | T | C | 0.938449 | 2.06E-10 | 8.09E-05 |
| 11 | 71626996 | T | C | 0.938449 | 2.06E-10 | 8.09E-05 |
| 11 | 69530892 | C | T | 0.815346 | 2.17E-10 | 8.49E-05 |
| 11 | 69530893 | T | G | 0.815346 | 2.17E-10 | 8.49E-05 |
| 11 | 69531057 | A | C | 0.801012 | 2.25E-10 | 8.84E-05 |
| 11 | 66080058 | A | T | 0.870152 | 2.26E-10 | 8.87E-05 |
| 11 | 69785405 | T | C | 0.813659 | 2.50E-10 | 9.81E-05 |
| 11 | 71623922 | A | G | 0.936762 | 2.53E-10 | 9.91E-05 |
| 11 | 71624560 | T | G | 0.936762 | 2.53E-10 | 9.91E-05 |
| 11 | 69793327 | A | G | 0.816189 | 2.87E-10 | 0.00011235 |
| 11 | 71507889 | C | A | 0.935076 | 2.98E-10 | 0.00011695 |
| 11 | 71507957 | G | A | 0.935076 | 2.98E-10 | 0.00011695 |
| 11 | 64567442 | A | G | 0.879427 | 3.01E-10 | 0.00011787 |
| 11 | 69530287 | G | A | 0.814503 | 3.88E-10 | 0.00015213 |
| 11 | 69530313 | C | A | 0.814503 | 3.88E-10 | 0.00015213 |
| 11 | 64969848 | T | G | 0.720067 | 4.30E-10 | 0.00016858 |
| 11 | 65072644 | C | T | 0.869309 | 4.35E-10 | 0.00017065 |
| 11 | 64169836 | G | T | 0.844013 | 4.52E-10 | 0.00017699 |
| 11 | 71498274 | C | T | 0.937605 | 5.66E-10 | 0.00022182 |
| 11 | 71498368 | G | T | 0.937605 | 5.66E-10 | 0.00022182 |
| 11 | 71498450 | T | C | 0.937605 | 5.66E-10 | 0.00022182 |
| 11 | 71498627 | A | T | 0.937605 | 5.66E-10 | 0.00022182 |
| 11 | 71499087 | C | A | 0.937605 | 5.66E-10 | 0.00022182 |
| 11 | 71499162 | A | G | 0.937605 | 5.66E-10 | 0.00022182 |
| 11 | 71499267 | G | A | 0.937605 | 5.66E-10 | 0.00022182 |
| 11 | 71499542 | G | C | 0.937605 | 5.66E-10 | 0.00022182 |
| 11 | 71499781 | T | C | 0.937605 | 5.66E-10 | 0.00022182 |
| 11 | 71499829 | A | G | 0.937605 | 5.66E-10 | 0.00022182 |
| 11 | 71500598 | T | C | 0.937605 | 5.66E-10 | 0.00022182 |
| 11 | 71501211 | A | G | 0.937605 | 5.66E-10 | 0.00022182 |
| 11 | 71501415 | C | T | 0.937605 | 5.66E-10 | 0.00022182 |
| 11 | 71502096 | T | C | 0.937605 | 5.66E-10 | 0.00022182 |
| 11 | 71502102 | G | A | 0.937605 | 5.66E-10 | 0.00022182 |
| 11 | 71502614 | G | C | 0.937605 | 5.66E-10 | 0.00022182 |
| 11 | 71503182 | T | C | 0.937605 | 5.66E-10 | 0.00022182 |
| 11 | 71503761 | G | C | 0.937605 | 5.66E-10 | 0.00022182 |
| 11 | 71505501 | T | G | 0.937605 | 5.66E-10 | 0.00022182 |
| 11 | 71507273 | A | G | 0.937605 | 5.66E-10 | 0.00022182 |
| 11 | 71507725 | G | A | 0.937605 | 5.66E-10 | 0.00022182 |
| 11 | 71507773 | C | T | 0.937605 | 5.66E-10 | 0.00022182 |
| 11 | 71507803 | T | C | 0.937605 | 5.66E-10 | 0.00022182 |
| 11 | 71509158 | A | G | 0.937605 | 5.66E-10 | 0.00022182 |
| 11 | 71509207 | C | T | 0.937605 | 5.66E-10 | 0.00022182 |
| 11 | 71509246 | T | C | 0.937605 | 5.66E-10 | 0.00022182 |
| 11 | 71509281 | T | C | 0.937605 | 5.66E-10 | 0.00022182 |
| 11 | 71509443 | C | T | 0.937605 | 5.66E-10 | 0.00022182 |
| 11 | 71509541 | A | G | 0.937605 | 5.66E-10 | 0.00022182 |
| 11 | 71509746 | T | C | 0.937605 | 5.66E-10 | 0.00022182 |
| 11 | 71509907 | A | G | 0.937605 | 5.66E-10 | 0.00022182 |
| 11 | 71511466 | T | C | 0.937605 | 5.66E-10 | 0.00022182 |
| 11 | 71511579 | T | C | 0.937605 | 5.66E-10 | 0.00022182 |
| 11 | 71511584 | T | C | 0.937605 | 5.66E-10 | 0.00022182 |
| 11 | 71511896 | T | C | 0.937605 | 5.66E-10 | 0.00022182 |
| 11 | 71512032 | C | T | 0.937605 | 5.66E-10 | 0.00022182 |
| 11 | 71512064 | G | C | 0.937605 | 5.66E-10 | 0.00022182 |

| | | | | | | |
|---|---|---|---|---|---|---|
| 11 | 71512150 | A | G | 0.937605 | 5.66E-10 | 0.00022182 |
| 11 | 71512171 | C | A | 0.937605 | 5.66E-10 | 0.00022182 |
| 11 | 71512299 | C | T | 0.937605 | 5.66E-10 | 0.00022182 |
| 11 | 71512317 | G | C | 0.937605 | 5.66E-10 | 0.00022182 |
| 11 | 71512348 | C | T | 0.937605 | 5.66E-10 | 0.00022182 |
| 11 | 71512650 | G | A | 0.937605 | 5.66E-10 | 0.00022182 |
| 11 | 71512651 | G | T | 0.937605 | 5.66E-10 | 0.00022182 |
| 11 | 71512656 | G | A | 0.937605 | 5.66E-10 | 0.00022182 |
| 11 | 71512677 | A | G | 0.937605 | 5.66E-10 | 0.00022182 |
| 11 | 71513003 | G | T | 0.937605 | 5.66E-10 | 0.00022182 |
| 11 | 71513033 | T | G | 0.937605 | 5.66E-10 | 0.00022182 |
| 11 | 71513255 | A | C | 0.937605 | 5.66E-10 | 0.00022182 |
| 11 | 71513445 | A | G | 0.937605 | 5.66E-10 | 0.00022182 |
| 11 | 71513525 | C | T | 0.937605 | 5.66E-10 | 0.00022182 |
| 11 | 71513528 | C | T | 0.937605 | 5.66E-10 | 0.00022182 |
| 11 | 71514151 | A | G | 0.937605 | 5.66E-10 | 0.00022182 |
| 11 | 71514355 | T | C | 0.937605 | 5.66E-10 | 0.00022182 |
| 11 | 71514435 | T | C | 0.937605 | 5.66E-10 | 0.00022182 |
| 11 | 71514607 | G | A | 0.937605 | 5.66E-10 | 0.00022182 |
| 11 | 71515240 | T | G | 0.937605 | 5.66E-10 | 0.00022182 |
| 11 | 71515647 | G | A | 0.937605 | 5.66E-10 | 0.00022182 |
| 11 | 71515694 | C | T | 0.937605 | 5.66E-10 | 0.00022182 |
| 11 | 71515705 | G | A | 0.937605 | 5.66E-10 | 0.00022182 |
| 11 | 71516162 | C | A | 0.937605 | 5.66E-10 | 0.00022182 |
| 11 | 71516167 | G | A | 0.937605 | 5.66E-10 | 0.00022182 |
| 11 | 71516390 | A | T | 0.937605 | 5.66E-10 | 0.00022182 |
| 11 | 71517211 | C | G | 0.937605 | 5.66E-10 | 0.00022182 |
| 11 | 71517381 | C | A | 0.937605 | 5.66E-10 | 0.00022182 |
| 11 | 71517468 | A | G | 0.937605 | 5.66E-10 | 0.00022182 |
| 11 | 71517804 | A | G | 0.937605 | 5.66E-10 | 0.00022182 |
| 11 | 71517815 | T | C | 0.937605 | 5.66E-10 | 0.00022182 |
| 11 | 71517851 | C | T | 0.937605 | 5.66E-10 | 0.00022182 |
| 11 | 71517853 | A | C | 0.937605 | 5.66E-10 | 0.00022182 |
| 11 | 71517908 | A | G | 0.937605 | 5.66E-10 | 0.00022182 |
| 11 | 71518490 | T | C | 0.937605 | 5.66E-10 | 0.00022182 |
| 11 | 71518768 | C | G | 0.937605 | 5.66E-10 | 0.00022182 |
| 11 | 71518780 | A | G | 0.937605 | 5.66E-10 | 0.00022182 |
| 11 | 71518803 | G | C | 0.937605 | 5.66E-10 | 0.00022182 |
| 11 | 71518914 | C | G | 0.937605 | 5.66E-10 | 0.00022182 |
| 11 | 71519090 | T | C | 0.937605 | 5.66E-10 | 0.00022182 |
| 11 | 71519132 | C | T | 0.937605 | 5.66E-10 | 0.00022182 |
| 11 | 71519190 | T | C | 0.937605 | 5.66E-10 | 0.00022182 |
| 11 | 71519908 | G | A | 0.937605 | 5.66E-10 | 0.00022182 |
| 11 | 71520121 | A | G | 0.937605 | 5.66E-10 | 0.00022182 |
| 11 | 71520877 | T | C | 0.937605 | 5.66E-10 | 0.00022182 |
| 11 | 71521097 | C | T | 0.937605 | 5.66E-10 | 0.00022182 |
| 11 | 71521315 | T | C | 0.937605 | 5.66E-10 | 0.00022182 |
| 11 | 71521318 | T | C | 0.937605 | 5.66E-10 | 0.00022182 |
| 11 | 71521981 | C | T | 0.937605 | 5.66E-10 | 0.00022182 |
| 11 | 71523030 | C | T | 0.937605 | 5.66E-10 | 0.00022182 |
| 11 | 71523270 | G | A | 0.937605 | 5.66E-10 | 0.00022182 |
| 11 | 71523276 | A | G | 0.937605 | 5.66E-10 | 0.00022182 |
| 11 | 71523499 | A | G | 0.937605 | 5.66E-10 | 0.00022182 |
| 11 | 71524176 | A | C | 0.937605 | 5.66E-10 | 0.00022182 |
| 11 | 71524757 | T | C | 0.937605 | 5.66E-10 | 0.00022182 |
| 11 | 71524801 | A | C | 0.937605 | 5.66E-10 | 0.00022182 |
| 11 | 71524989 | G | C | 0.937605 | 5.66E-10 | 0.00022182 |
| 11 | 71525001 | C | T | 0.937605 | 5.66E-10 | 0.00022182 |
| 11 | 71525011 | C | A | 0.937605 | 5.66E-10 | 0.00022182 |
| 11 | 71525580 | C | T | 0.937605 | 5.66E-10 | 0.00022182 |
| 11 | 71526025 | G | A | 0.937605 | 5.66E-10 | 0.00022182 |
| 11 | 71526192 | A | G | 0.937605 | 5.66E-10 | 0.00022182 |
| 11 | 71526308 | T | C | 0.937605 | 5.66E-10 | 0.00022182 |
| 11 | 71526707 | T | C | 0.937605 | 5.66E-10 | 0.00022182 |
| 11 | 71527274 | G | A | 0.937605 | 5.66E-10 | 0.00022182 |
| 11 | 71527386 | A | G | 0.937605 | 5.66E-10 | 0.00022182 |
| 11 | 71528102 | G | A | 0.937605 | 5.66E-10 | 0.00022182 |

| | | | | | | |
|---|---|---|---|---|---|---|
| 11 | 71528456 | C | G | 0.937605 | 5.66E-10 | 0.00022182 |
| 11 | 71528562 | G | A | 0.937605 | 5.66E-10 | 0.00022182 |
| 11 | 71528994 | A | G | 0.937605 | 5.66E-10 | 0.00022182 |
| 11 | 71529039 | A | C | 0.937605 | 5.66E-10 | 0.00022182 |
| 11 | 71529306 | T | C | 0.937605 | 5.66E-10 | 0.00022182 |
| 11 | 71529317 | C | T | 0.937605 | 5.66E-10 | 0.00022182 |
| 11 | 71529713 | A | G | 0.937605 | 5.66E-10 | 0.00022182 |
| 11 | 71529737 | A | G | 0.937605 | 5.66E-10 | 0.00022182 |
| 11 | 71530091 | C | T | 0.937605 | 5.66E-10 | 0.00022182 |
| 11 | 71530143 | G | T | 0.937605 | 5.66E-10 | 0.00022182 |
| 11 | 71530160 | T | C | 0.937605 | 5.66E-10 | 0.00022182 |
| 11 | 71530282 | C | A | 0.937605 | 5.66E-10 | 0.00022182 |
| 11 | 71530283 | A | C | 0.937605 | 5.66E-10 | 0.00022182 |
| 11 | 71530826 | A | G | 0.937605 | 5.66E-10 | 0.00022182 |
| 11 | 71532111 | T | G | 0.937605 | 5.66E-10 | 0.00022182 |
| 11 | 71532130 | C | G | 0.937605 | 5.66E-10 | 0.00022182 |
| 11 | 71532165 | C | A | 0.937605 | 5.66E-10 | 0.00022182 |
| 11 | 71532197 | C | T | 0.937605 | 5.66E-10 | 0.00022182 |
| 11 | 71532234 | T | C | 0.937605 | 5.66E-10 | 0.00022182 |
| 11 | 71532274 | T | G | 0.937605 | 5.66E-10 | 0.00022182 |
| 11 | 71532279 | A | C | 0.937605 | 5.66E-10 | 0.00022182 |
| 11 | 71532293 | A | G | 0.937605 | 5.66E-10 | 0.00022182 |
| 11 | 71532333 | T | C | 0.937605 | 5.66E-10 | 0.00022182 |
| 11 | 71532338 | T | C | 0.937605 | 5.66E-10 | 0.00022182 |
| 11 | 71532424 | A | T | 0.937605 | 5.66E-10 | 0.00022182 |
| 11 | 71532463 | C | T | 0.937605 | 5.66E-10 | 0.00022182 |
| 11 | 71532540 | C | G | 0.937605 | 5.66E-10 | 0.00022182 |
| 11 | 71532584 | G | C | 0.937605 | 5.66E-10 | 0.00022182 |
| 11 | 71532653 | G | C | 0.937605 | 5.66E-10 | 0.00022182 |
| 11 | 71532986 | T | G | 0.937605 | 5.66E-10 | 0.00022182 |
| 11 | 71533001 | T | A | 0.937605 | 5.66E-10 | 0.00022182 |
| 11 | 71533013 | T | C | 0.937605 | 5.66E-10 | 0.00022182 |
| 11 | 71533032 | A | T | 0.937605 | 5.66E-10 | 0.00022182 |
| 11 | 71533548 | T | C | 0.937605 | 5.66E-10 | 0.00022182 |
| 11 | 71534115 | T | A | 0.937605 | 5.66E-10 | 0.00022182 |
| 11 | 71534160 | G | A | 0.937605 | 5.66E-10 | 0.00022182 |
| 11 | 71534185 | T | C | 0.937605 | 5.66E-10 | 0.00022182 |
| 11 | 71534218 | G | A | 0.937605 | 5.66E-10 | 0.00022182 |
| 11 | 71535033 | A | G | 0.937605 | 5.66E-10 | 0.00022182 |
| 11 | 71535147 | C | T | 0.937605 | 5.66E-10 | 0.00022182 |
| 11 | 71535599 | T | G | 0.937605 | 5.66E-10 | 0.00022182 |
| 11 | 71536124 | A | T | 0.937605 | 5.66E-10 | 0.00022182 |
| 11 | 71537313 | T | C | 0.937605 | 5.66E-10 | 0.00022182 |
| 11 | 71537533 | G | A | 0.937605 | 5.66E-10 | 0.00022182 |
| 11 | 71537815 | T | C | 0.937605 | 5.66E-10 | 0.00022182 |
| 11 | 71540174 | C | A | 0.937605 | 5.66E-10 | 0.00022182 |
| 11 | 71540595 | G | A | 0.937605 | 5.66E-10 | 0.00022182 |
| 11 | 71541688 | G | A | 0.937605 | 5.66E-10 | 0.00022182 |
| 11 | 71541692 | T | C | 0.937605 | 5.66E-10 | 0.00022182 |
| 11 | 71542189 | G | A | 0.937605 | 5.66E-10 | 0.00022182 |
| 11 | 71542516 | G | A | 0.937605 | 5.66E-10 | 0.00022182 |
| 11 | 71542545 | G | A | 0.937605 | 5.66E-10 | 0.00022182 |
| 11 | 71542632 | G | T | 0.937605 | 5.66E-10 | 0.00022182 |
| 11 | 71542725 | G | C | 0.937605 | 5.66E-10 | 0.00022182 |
| 11 | 71542809 | C | T | 0.937605 | 5.66E-10 | 0.00022182 |
| 11 | 71542867 | T | C | 0.937605 | 5.66E-10 | 0.00022182 |
| 11 | 71543447 | A | T | 0.937605 | 5.66E-10 | 0.00022182 |
| 11 | 71543618 | T | C | 0.937605 | 5.66E-10 | 0.00022182 |
| 11 | 71543621 | T | C | 0.937605 | 5.66E-10 | 0.00022182 |
| 11 | 71543945 | T | C | 0.937605 | 5.66E-10 | 0.00022182 |
| 11 | 71543948 | A | G | 0.937605 | 5.66E-10 | 0.00022182 |
| 11 | 71544374 | T | C | 0.937605 | 5.66E-10 | 0.00022182 |
| 11 | 71544528 | G | A | 0.937605 | 5.66E-10 | 0.00022182 |
| 11 | 71544732 | C | G | 0.937605 | 5.66E-10 | 0.00022182 |
| 11 | 71545783 | T | C | 0.937605 | 5.66E-10 | 0.00022182 |
| 11 | 71545963 | T | C | 0.937605 | 5.66E-10 | 0.00022182 |
| 11 | 71546107 | G | C | 0.937605 | 5.66E-10 | 0.00022182 |

| | | | | | | |
|---|---|---|---|---|---|---|
| 11 | 71546313 | G | A | 0.937605 | 5.66E-10 | 0.00022182 |
| 11 | 71546371 | G | T | 0.937605 | 5.66E-10 | 0.00022182 |
| 11 | 71546383 | C | T | 0.937605 | 5.66E-10 | 0.00022182 |
| 11 | 71546390 | A | C | 0.937605 | 5.66E-10 | 0.00022182 |
| 11 | 71546485 | C | T | 0.937605 | 5.66E-10 | 0.00022182 |
| 11 | 71546486 | T | G | 0.937605 | 5.66E-10 | 0.00022182 |
| 11 | 71546488 | T | C | 0.937605 | 5.66E-10 | 0.00022182 |
| 11 | 71546662 | A | C | 0.937605 | 5.66E-10 | 0.00022182 |
| 11 | 71547092 | A | G | 0.937605 | 5.66E-10 | 0.00022182 |
| 11 | 71547163 | T | A | 0.937605 | 5.66E-10 | 0.00022182 |
| 11 | 71547267 | C | A | 0.937605 | 5.66E-10 | 0.00022182 |
| 11 | 71547329 | C | T | 0.937605 | 5.66E-10 | 0.00022182 |
| 11 | 71547504 | C | T | 0.937605 | 5.66E-10 | 0.00022182 |
| 11 | 71547510 | G | A | 0.937605 | 5.66E-10 | 0.00022182 |
| 11 | 71547619 | C | T | 0.937605 | 5.66E-10 | 0.00022182 |
| 11 | 71547650 | A | G | 0.937605 | 5.66E-10 | 0.00022182 |
| 11 | 71547714 | T | C | 0.937605 | 5.66E-10 | 0.00022182 |
| 11 | 71547834 | G | A | 0.937605 | 5.66E-10 | 0.00022182 |
| 11 | 71547912 | G | C | 0.937605 | 5.66E-10 | 0.00022182 |
| 11 | 71547936 | T | A | 0.937605 | 5.66E-10 | 0.00022182 |
| 11 | 71547945 | C | T | 0.937605 | 5.66E-10 | 0.00022182 |
| 11 | 71547962 | C | A | 0.937605 | 5.66E-10 | 0.00022182 |
| 11 | 71548058 | A | C | 0.937605 | 5.66E-10 | 0.00022182 |
| 11 | 71548185 | A | C | 0.937605 | 5.66E-10 | 0.00022182 |
| 11 | 71548229 | C | T | 0.937605 | 5.66E-10 | 0.00022182 |
| 11 | 71548542 | G | T | 0.937605 | 5.66E-10 | 0.00022182 |
| 11 | 71548585 | C | T | 0.937605 | 5.66E-10 | 0.00022182 |
| 11 | 71548614 | C | T | 0.937605 | 5.66E-10 | 0.00022182 |
| 11 | 71548651 | C | T | 0.937605 | 5.66E-10 | 0.00022182 |
| 11 | 71548780 | A | T | 0.937605 | 5.66E-10 | 0.00022182 |
| 11 | 71549136 | C | T | 0.937605 | 5.66E-10 | 0.00022182 |
| 11 | 71549623 | T | C | 0.937605 | 5.66E-10 | 0.00022182 |
| 11 | 71550817 | G | T | 0.937605 | 5.66E-10 | 0.00022182 |
| 11 | 71551027 | T | G | 0.937605 | 5.66E-10 | 0.00022182 |
| 11 | 71551036 | A | G | 0.937605 | 5.66E-10 | 0.00022182 |
| 11 | 71551037 | A | C | 0.937605 | 5.66E-10 | 0.00022182 |
| 11 | 71551069 | C | T | 0.937605 | 5.66E-10 | 0.00022182 |
| 11 | 71551077 | A | T | 0.937605 | 5.66E-10 | 0.00022182 |
| 11 | 71551199 | G | T | 0.937605 | 5.66E-10 | 0.00022182 |
| 11 | 71551389 | C | T | 0.937605 | 5.66E-10 | 0.00022182 |
| 11 | 71551390 | A | G | 0.937605 | 5.66E-10 | 0.00022182 |
| 11 | 71551471 | G | A | 0.937605 | 5.66E-10 | 0.00022182 |
| 11 | 71551493 | T | C | 0.937605 | 5.66E-10 | 0.00022182 |
| 11 | 71551520 | C | T | 0.937605 | 5.66E-10 | 0.00022182 |
| 11 | 71551562 | C | T | 0.937605 | 5.66E-10 | 0.00022182 |
| 11 | 71551584 | T | A | 0.937605 | 5.66E-10 | 0.00022182 |
| 11 | 71551598 | G | A | 0.937605 | 5.66E-10 | 0.00022182 |
| 11 | 71553104 | A | G | 0.937605 | 5.66E-10 | 0.00022182 |
| 11 | 71553318 | G | A | 0.937605 | 5.66E-10 | 0.00022182 |
| 11 | 71553461 | T | C | 0.937605 | 5.66E-10 | 0.00022182 |
| 11 | 71553759 | G | C | 0.937605 | 5.66E-10 | 0.00022182 |
| 11 | 71553767 | A | G | 0.937605 | 5.66E-10 | 0.00022182 |
| 11 | 71553880 | A | G | 0.937605 | 5.66E-10 | 0.00022182 |
| 11 | 71553906 | A | G | 0.937605 | 5.66E-10 | 0.00022182 |
| 11 | 71553910 | T | C | 0.937605 | 5.66E-10 | 0.00022182 |
| 11 | 71553977 | A | G | 0.937605 | 5.66E-10 | 0.00022182 |
| 11 | 71554207 | T | C | 0.937605 | 5.66E-10 | 0.00022182 |
| 11 | 71554317 | T | C | 0.937605 | 5.66E-10 | 0.00022182 |
| 11 | 71554732 | A | G | 0.937605 | 5.66E-10 | 0.00022182 |
| 11 | 71554735 | A | G | 0.937605 | 5.66E-10 | 0.00022182 |
| 11 | 71554763 | A | G | 0.937605 | 5.66E-10 | 0.00022182 |
| 11 | 71554802 | A | C | 0.937605 | 5.66E-10 | 0.00022182 |
| 11 | 71554814 | C | T | 0.937605 | 5.66E-10 | 0.00022182 |
| 11 | 71554991 | A | G | 0.937605 | 5.66E-10 | 0.00022182 |
| 11 | 71555004 | C | T | 0.937605 | 5.66E-10 | 0.00022182 |
| 11 | 71555056 | T | C | 0.937605 | 5.66E-10 | 0.00022182 |
| 11 | 71555178 | A | G | 0.937605 | 5.66E-10 | 0.00022182 |

| | | | | | | |
|---|---|---|---|---|---|---|
| 11 | 71555268 | T | C | 0.937605 | 5.66E-10 | 0.00022182 |
| 11 | 71555391 | C | T | 0.937605 | 5.66E-10 | 0.00022182 |
| 11 | 71555581 | A | G | 0.937605 | 5.66E-10 | 0.00022182 |
| 11 | 71555621 | T | C | 0.937605 | 5.66E-10 | 0.00022182 |
| 11 | 71555622 | G | A | 0.937605 | 5.66E-10 | 0.00022182 |
| 11 | 71555786 | T | G | 0.937605 | 5.66E-10 | 0.00022182 |
| 11 | 71555789 | C | G | 0.937605 | 5.66E-10 | 0.00022182 |
| 11 | 71555801 | A | C | 0.937605 | 5.66E-10 | 0.00022182 |
| 11 | 71555818 | T | C | 0.937605 | 5.66E-10 | 0.00022182 |
| 11 | 71555901 | G | A | 0.937605 | 5.66E-10 | 0.00022182 |
| 11 | 71555910 | A | G | 0.937605 | 5.66E-10 | 0.00022182 |
| 11 | 71555987 | A | G | 0.937605 | 5.66E-10 | 0.00022182 |
| 11 | 71555994 | A | G | 0.937605 | 5.66E-10 | 0.00022182 |
| 11 | 71556121 | G | T | 0.937605 | 5.66E-10 | 0.00022182 |
| 11 | 71556123 | C | T | 0.937605 | 5.66E-10 | 0.00022182 |
| 11 | 71556189 | G | T | 0.937605 | 5.66E-10 | 0.00022182 |
| 11 | 71556334 | G | A | 0.937605 | 5.66E-10 | 0.00022182 |
| 11 | 71556361 | G | A | 0.937605 | 5.66E-10 | 0.00022182 |
| 11 | 71556426 | C | A | 0.937605 | 5.66E-10 | 0.00022182 |
| 11 | 71556551 | T | C | 0.937605 | 5.66E-10 | 0.00022182 |
| 11 | 71556553 | T | A | 0.937605 | 5.66E-10 | 0.00022182 |
| 11 | 71556822 | G | T | 0.937605 | 5.66E-10 | 0.00022182 |
| 11 | 71556970 | G | A | 0.937605 | 5.66E-10 | 0.00022182 |
| 11 | 71556991 | T | C | 0.937605 | 5.66E-10 | 0.00022182 |
| 11 | 71557110 | C | T | 0.937605 | 5.66E-10 | 0.00022182 |
| 11 | 71557111 | A | G | 0.937605 | 5.66E-10 | 0.00022182 |
| 11 | 71557141 | A | C | 0.937605 | 5.66E-10 | 0.00022182 |
| 11 | 71557144 | C | T | 0.937605 | 5.66E-10 | 0.00022182 |
| 11 | 71557195 | T | G | 0.937605 | 5.66E-10 | 0.00022182 |
| 11 | 71557218 | A | G | 0.937605 | 5.66E-10 | 0.00022182 |
| 11 | 71557336 | G | A | 0.937605 | 5.66E-10 | 0.00022182 |
| 11 | 71557382 | C | G | 0.937605 | 5.66E-10 | 0.00022182 |
| 11 | 71557469 | T | A | 0.937605 | 5.66E-10 | 0.00022182 |
| 11 | 71557480 | G | A | 0.937605 | 5.66E-10 | 0.00022182 |
| 11 | 71557484 | G | C | 0.937605 | 5.66E-10 | 0.00022182 |
| 11 | 71557489 | T | C | 0.937605 | 5.66E-10 | 0.00022182 |
| 11 | 71557511 | A | T | 0.937605 | 5.66E-10 | 0.00022182 |
| 11 | 71557538 | C | A | 0.937605 | 5.66E-10 | 0.00022182 |
| 11 | 71557554 | A | T | 0.937605 | 5.66E-10 | 0.00022182 |
| 11 | 71557637 | G | A | 0.937605 | 5.66E-10 | 0.00022182 |
| 11 | 71557790 | G | A | 0.937605 | 5.66E-10 | 0.00022182 |
| 11 | 71557931 | G | A | 0.937605 | 5.66E-10 | 0.00022182 |
| 11 | 71557948 | T | C | 0.937605 | 5.66E-10 | 0.00022182 |
| 11 | 71557996 | T | A | 0.937605 | 5.66E-10 | 0.00022182 |
| 11 | 71558108 | A | G | 0.937605 | 5.66E-10 | 0.00022182 |
| 11 | 71558157 | A | G | 0.937605 | 5.66E-10 | 0.00022182 |
| 11 | 71558298 | G | A | 0.937605 | 5.66E-10 | 0.00022182 |
| 11 | 71558356 | A | C | 0.937605 | 5.66E-10 | 0.00022182 |
| 11 | 71558385 | A | G | 0.937605 | 5.66E-10 | 0.00022182 |
| 11 | 71558488 | G | A | 0.937605 | 5.66E-10 | 0.00022182 |
| 11 | 71558539 | A | G | 0.937605 | 5.66E-10 | 0.00022182 |
| 11 | 71558981 | A | C | 0.937605 | 5.66E-10 | 0.00022182 |
| 11 | 71559025 | C | T | 0.937605 | 5.66E-10 | 0.00022182 |
| 11 | 71559183 | T | C | 0.937605 | 5.66E-10 | 0.00022182 |
| 11 | 71560304 | A | G | 0.937605 | 5.66E-10 | 0.00022182 |
| 11 | 71560890 | T | G | 0.937605 | 5.66E-10 | 0.00022182 |
| 11 | 71560962 | C | A | 0.937605 | 5.66E-10 | 0.00022182 |
| 11 | 71561173 | T | C | 0.937605 | 5.66E-10 | 0.00022182 |
| 11 | 71561236 | A | G | 0.937605 | 5.66E-10 | 0.00022182 |
| 11 | 71561363 | T | C | 0.937605 | 5.66E-10 | 0.00022182 |
| 11 | 71561514 | C | A | 0.937605 | 5.66E-10 | 0.00022182 |
| 11 | 71561608 | G | A | 0.937605 | 5.66E-10 | 0.00022182 |
| 11 | 71561610 | G | A | 0.937605 | 5.66E-10 | 0.00022182 |
| 11 | 71561649 | T | C | 0.937605 | 5.66E-10 | 0.00022182 |
| 11 | 71561789 | G | C | 0.937605 | 5.66E-10 | 0.00022182 |
| 11 | 71561961 | T | A | 0.937605 | 5.66E-10 | 0.00022182 |
| 11 | 71562016 | A | T | 0.937605 | 5.66E-10 | 0.00022182 |

| | | | | | | |
|---|---|---|---|---|---|---|
| 11 | 71562270 | T | C | 0.937605 | 5.66E-10 | 0.00022182 |
| 11 | 71562505 | T | C | 0.937605 | 5.66E-10 | 0.00022182 |
| 11 | 71562581 | A | G | 0.937605 | 5.66E-10 | 0.00022182 |
| 11 | 71562809 | C | T | 0.937605 | 5.66E-10 | 0.00022182 |
| 11 | 71563456 | C | G | 0.937605 | 5.66E-10 | 0.00022182 |
| 11 | 71563639 | C | T | 0.937605 | 5.66E-10 | 0.00022182 |
| 11 | 71564577 | C | T | 0.937605 | 5.66E-10 | 0.00022182 |
| 11 | 71565435 | G | A | 0.937605 | 5.66E-10 | 0.00022182 |
| 11 | 71566318 | G | A | 0.937605 | 5.66E-10 | 0.00022182 |
| 11 | 71566444 | C | T | 0.937605 | 5.66E-10 | 0.00022182 |
| 11 | 71566966 | C | G | 0.937605 | 5.66E-10 | 0.00022182 |
| 11 | 71567182 | A | G | 0.937605 | 5.66E-10 | 0.00022182 |
| 11 | 71567228 | C | T | 0.937605 | 5.66E-10 | 0.00022182 |
| 11 | 71567387 | G | A | 0.937605 | 5.66E-10 | 0.00022182 |
| 11 | 71567476 | C | T | 0.937605 | 5.66E-10 | 0.00022182 |
| 11 | 71567590 | C | T | 0.937605 | 5.66E-10 | 0.00022182 |
| 11 | 71567768 | T | C | 0.937605 | 5.66E-10 | 0.00022182 |
| 11 | 71567774 | T | C | 0.937605 | 5.66E-10 | 0.00022182 |
| 11 | 71568267 | C | T | 0.937605 | 5.66E-10 | 0.00022182 |
| 11 | 71568437 | A | C | 0.937605 | 5.66E-10 | 0.00022182 |
| 11 | 71568585 | C | T | 0.937605 | 5.66E-10 | 0.00022182 |
| 11 | 71568607 | C | T | 0.937605 | 5.66E-10 | 0.00022182 |
| 11 | 71568701 | G | T | 0.937605 | 5.66E-10 | 0.00022182 |
| 11 | 71568728 | T | C | 0.937605 | 5.66E-10 | 0.00022182 |
| 11 | 71569089 | A | G | 0.937605 | 5.66E-10 | 0.00022182 |
| 11 | 71569256 | C | T | 0.937605 | 5.66E-10 | 0.00022182 |
| 11 | 71569268 | T | C | 0.937605 | 5.66E-10 | 0.00022182 |
| 11 | 71569269 | G | A | 0.937605 | 5.66E-10 | 0.00022182 |
| 11 | 71569668 | A | G | 0.937605 | 5.66E-10 | 0.00022182 |
| 11 | 71569744 | T | C | 0.937605 | 5.66E-10 | 0.00022182 |
| 11 | 71569758 | G | A | 0.937605 | 5.66E-10 | 0.00022182 |
| 11 | 71569880 | G | T | 0.937605 | 5.66E-10 | 0.00022182 |
| 11 | 71569998 | G | T | 0.937605 | 5.66E-10 | 0.00022182 |
| 11 | 71570016 | C | T | 0.937605 | 5.66E-10 | 0.00022182 |
| 11 | 71570201 | T | C | 0.937605 | 5.66E-10 | 0.00022182 |
| 11 | 71570280 | A | C | 0.937605 | 5.66E-10 | 0.00022182 |
| 11 | 71570467 | A | T | 0.937605 | 5.66E-10 | 0.00022182 |
| 11 | 71570493 | G | T | 0.937605 | 5.66E-10 | 0.00022182 |
| 11 | 71570525 | A | C | 0.937605 | 5.66E-10 | 0.00022182 |
| 11 | 71570611 | T | C | 0.937605 | 5.66E-10 | 0.00022182 |
| 11 | 71571118 | C | T | 0.937605 | 5.66E-10 | 0.00022182 |
| 11 | 71571369 | C | T | 0.937605 | 5.66E-10 | 0.00022182 |
| 11 | 71571386 | C | T | 0.937605 | 5.66E-10 | 0.00022182 |
| 11 | 71571445 | A | G | 0.937605 | 5.66E-10 | 0.00022182 |
| 11 | 71571614 | A | G | 0.937605 | 5.66E-10 | 0.00022182 |
| 11 | 71571631 | G | A | 0.937605 | 5.66E-10 | 0.00022182 |
| 11 | 71571707 | A | G | 0.937605 | 5.66E-10 | 0.00022182 |
| 11 | 71571723 | C | T | 0.937605 | 5.66E-10 | 0.00022182 |
| 11 | 71571730 | T | A | 0.937605 | 5.66E-10 | 0.00022182 |
| 11 | 71571764 | T | A | 0.937605 | 5.66E-10 | 0.00022182 |
| 11 | 71571865 | C | G | 0.937605 | 5.66E-10 | 0.00022182 |
| 11 | 71571901 | T | G | 0.937605 | 5.66E-10 | 0.00022182 |
| 11 | 71572051 | G | A | 0.937605 | 5.66E-10 | 0.00022182 |
| 11 | 71572373 | C | T | 0.937605 | 5.66E-10 | 0.00022182 |
| 11 | 71572419 | A | G | 0.937605 | 5.66E-10 | 0.00022182 |
| 11 | 71572428 | C | T | 0.937605 | 5.66E-10 | 0.00022182 |
| 11 | 71572432 | A | G | 0.937605 | 5.66E-10 | 0.00022182 |
| 11 | 71572515 | A | G | 0.937605 | 5.66E-10 | 0.00022182 |
| 11 | 71574958 | T | C | 0.937605 | 5.66E-10 | 0.00022182 |
| 11 | 71574960 | C | T | 0.937605 | 5.66E-10 | 0.00022182 |
| 11 | 71574985 | A | T | 0.937605 | 5.66E-10 | 0.00022182 |
| 11 | 71575049 | T | C | 0.937605 | 5.66E-10 | 0.00022182 |
| 11 | 71575077 | C | T | 0.937605 | 5.66E-10 | 0.00022182 |
| 11 | 71575089 | A | G | 0.937605 | 5.66E-10 | 0.00022182 |
| 11 | 71575137 | T | C | 0.937605 | 5.66E-10 | 0.00022182 |
| 11 | 71575162 | C | T | 0.937605 | 5.66E-10 | 0.00022182 |
| 11 | 71575170 | A | C | 0.937605 | 5.66E-10 | 0.00022182 |

| | | | | | | |
|---|---|---|---|---|---|---|
| 11 | 71575210 | T | C | 0.937605 | 5.66E-10 | 0.00022182 |
| 11 | 71575235 | T | C | 0.937605 | 5.66E-10 | 0.00022182 |
| 11 | 71575435 | T | C | 0.937605 | 5.66E-10 | 0.00022182 |
| 11 | 71575458 | G | A | 0.937605 | 5.66E-10 | 0.00022182 |
| 11 | 71575471 | T | C | 0.937605 | 5.66E-10 | 0.00022182 |
| 11 | 71575477 | T | C | 0.937605 | 5.66E-10 | 0.00022182 |
| 11 | 71575490 | C | T | 0.937605 | 5.66E-10 | 0.00022182 |
| 11 | 71575581 | A | G | 0.937605 | 5.66E-10 | 0.00022182 |
| 11 | 71575765 | A | C | 0.937605 | 5.66E-10 | 0.00022182 |
| 11 | 71576008 | G | A | 0.937605 | 5.66E-10 | 0.00022182 |
| 11 | 71576107 | A | G | 0.937605 | 5.66E-10 | 0.00022182 |
| 11 | 71576122 | T | C | 0.937605 | 5.66E-10 | 0.00022182 |
| 11 | 71576132 | A | G | 0.937605 | 5.66E-10 | 0.00022182 |
| 11 | 71576149 | A | T | 0.937605 | 5.66E-10 | 0.00022182 |
| 11 | 71576162 | G | A | 0.937605 | 5.66E-10 | 0.00022182 |
| 11 | 71576184 | A | G | 0.937605 | 5.66E-10 | 0.00022182 |
| 11 | 71576276 | T | C | 0.937605 | 5.66E-10 | 0.00022182 |
| 11 | 71576289 | A | T | 0.937605 | 5.66E-10 | 0.00022182 |
| 11 | 71576291 | A | G | 0.937605 | 5.66E-10 | 0.00022182 |
| 11 | 71576306 | T | C | 0.937605 | 5.66E-10 | 0.00022182 |
| 11 | 71576310 | A | C | 0.937605 | 5.66E-10 | 0.00022182 |
| 11 | 71576322 | T | C | 0.937605 | 5.66E-10 | 0.00022182 |
| 11 | 71576325 | T | G | 0.937605 | 5.66E-10 | 0.00022182 |
| 11 | 71576528 | A | C | 0.937605 | 5.66E-10 | 0.00022182 |
| 11 | 71576571 | A | C | 0.937605 | 5.66E-10 | 0.00022182 |
| 11 | 71576580 | A | G | 0.937605 | 5.66E-10 | 0.00022182 |
| 11 | 71576677 | A | G | 0.937605 | 5.66E-10 | 0.00022182 |
| 11 | 71576679 | G | A | 0.937605 | 5.66E-10 | 0.00022182 |
| 11 | 71576834 | T | A | 0.937605 | 5.66E-10 | 0.00022182 |
| 11 | 71576835 | C | T | 0.937605 | 5.66E-10 | 0.00022182 |
| 11 | 71576855 | T | C | 0.937605 | 5.66E-10 | 0.00022182 |
| 11 | 71576868 | A | G | 0.937605 | 5.66E-10 | 0.00022182 |
| 11 | 71576982 | T | C | 0.937605 | 5.66E-10 | 0.00022182 |
| 11 | 71577082 | T | C | 0.937605 | 5.66E-10 | 0.00022182 |
| 11 | 71577117 | T | C | 0.937605 | 5.66E-10 | 0.00022182 |
| 11 | 71577211 | A | G | 0.937605 | 5.66E-10 | 0.00022182 |
| 11 | 71577250 | G | T | 0.937605 | 5.66E-10 | 0.00022182 |
| 11 | 71577268 | C | T | 0.937605 | 5.66E-10 | 0.00022182 |
| 11 | 71577272 | T | A | 0.937605 | 5.66E-10 | 0.00022182 |
| 11 | 71577317 | A | G | 0.937605 | 5.66E-10 | 0.00022182 |
| 11 | 71577379 | A | T | 0.937605 | 5.66E-10 | 0.00022182 |
| 11 | 71577393 | T | C | 0.937605 | 5.66E-10 | 0.00022182 |
| 11 | 71577427 | A | G | 0.937605 | 5.66E-10 | 0.00022182 |
| 11 | 71577455 | A | G | 0.937605 | 5.66E-10 | 0.00022182 |
| 11 | 71577553 | G | A | 0.937605 | 5.66E-10 | 0.00022182 |
| 11 | 71577554 | T | C | 0.937605 | 5.66E-10 | 0.00022182 |
| 11 | 71577947 | G | T | 0.937605 | 5.66E-10 | 0.00022182 |
| 11 | 71578038 | G | A | 0.937605 | 5.66E-10 | 0.00022182 |
| 11 | 71578085 | A | T | 0.937605 | 5.66E-10 | 0.00022182 |
| 11 | 71578088 | C | T | 0.937605 | 5.66E-10 | 0.00022182 |
| 11 | 71578141 | G | T | 0.937605 | 5.66E-10 | 0.00022182 |
| 11 | 71578143 | A | G | 0.937605 | 5.66E-10 | 0.00022182 |
| 11 | 71578210 | C | T | 0.937605 | 5.66E-10 | 0.00022182 |
| 11 | 71578247 | T | C | 0.937605 | 5.66E-10 | 0.00022182 |
| 11 | 71578251 | C | T | 0.937605 | 5.66E-10 | 0.00022182 |
| 11 | 71578470 | A | G | 0.937605 | 5.66E-10 | 0.00022182 |
| 11 | 71578522 | G | A | 0.937605 | 5.66E-10 | 0.00022182 |
| 11 | 71578666 | A | T | 0.937605 | 5.66E-10 | 0.00022182 |
| 11 | 71578865 | T | C | 0.937605 | 5.66E-10 | 0.00022182 |
| 11 | 71578874 | A | G | 0.937605 | 5.66E-10 | 0.00022182 |
| 11 | 71579288 | C | T | 0.937605 | 5.66E-10 | 0.00022182 |
| 11 | 71580576 | A | G | 0.937605 | 5.66E-10 | 0.00022182 |
| 11 | 71582043 | T | C | 0.937605 | 5.66E-10 | 0.00022182 |
| 11 | 71582515 | C | T | 0.937605 | 5.66E-10 | 0.00022182 |
| 11 | 71582796 | G | C | 0.937605 | 5.66E-10 | 0.00022182 |
| 11 | 71582944 | T | C | 0.937605 | 5.66E-10 | 0.00022182 |
| 11 | 71583040 | A | T | 0.937605 | 5.66E-10 | 0.00022182 |

| | | | | | | |
|---|---|---|---|---|---|---|
| 11 | 71583041 | A | T | 0.937605 | 5.66E-10 | 0.00022182 |
| 11 | 71583056 | A | G | 0.937605 | 5.66E-10 | 0.00022182 |
| 11 | 71583376 | G | A | 0.937605 | 5.66E-10 | 0.00022182 |
| 11 | 71583400 | C | T | 0.937605 | 5.66E-10 | 0.00022182 |
| 11 | 71583587 | C | T | 0.937605 | 5.66E-10 | 0.00022182 |
| 11 | 71583695 | G | C | 0.937605 | 5.66E-10 | 0.00022182 |
| 11 | 71583703 | A | T | 0.937605 | 5.66E-10 | 0.00022182 |
| 11 | 71584233 | C | T | 0.937605 | 5.66E-10 | 0.00022182 |
| 11 | 71584247 | A | G | 0.937605 | 5.66E-10 | 0.00022182 |
| 11 | 71584248 | A | G | 0.937605 | 5.66E-10 | 0.00022182 |
| 11 | 71584405 | C | A | 0.937605 | 5.66E-10 | 0.00022182 |
| 11 | 71584467 | A | G | 0.937605 | 5.66E-10 | 0.00022182 |
| 11 | 71584610 | T | G | 0.937605 | 5.66E-10 | 0.00022182 |
| 11 | 71584646 | T | C | 0.937605 | 5.66E-10 | 0.00022182 |
| 11 | 71584698 | A | T | 0.937605 | 5.66E-10 | 0.00022182 |
| 11 | 71584701 | A | T | 0.937605 | 5.66E-10 | 0.00022182 |
| 11 | 71584757 | A | G | 0.937605 | 5.66E-10 | 0.00022182 |
| 11 | 71584764 | C | A | 0.937605 | 5.66E-10 | 0.00022182 |
| 11 | 71584855 | C | T | 0.937605 | 5.66E-10 | 0.00022182 |
| 11 | 71584934 | T | C | 0.937605 | 5.66E-10 | 0.00022182 |
| 11 | 71585016 | A | C | 0.937605 | 5.66E-10 | 0.00022182 |
| 11 | 71585157 | A | G | 0.937605 | 5.66E-10 | 0.00022182 |
| 11 | 71585381 | T | C | 0.937605 | 5.66E-10 | 0.00022182 |
| 11 | 71585521 | T | C | 0.937605 | 5.66E-10 | 0.00022182 |
| 11 | 71586075 | C | T | 0.937605 | 5.66E-10 | 0.00022182 |
| 11 | 71586176 | A | T | 0.937605 | 5.66E-10 | 0.00022182 |
| 11 | 71586450 | A | G | 0.937605 | 5.66E-10 | 0.00022182 |
| 11 | 71586830 | C | T | 0.937605 | 5.66E-10 | 0.00022182 |
| 11 | 71587015 | A | G | 0.937605 | 5.66E-10 | 0.00022182 |
| 11 | 71587178 | G | C | 0.937605 | 5.66E-10 | 0.00022182 |
| 11 | 71587261 | T | A | 0.937605 | 5.66E-10 | 0.00022182 |
| 11 | 71587270 | T | C | 0.937605 | 5.66E-10 | 0.00022182 |
| 11 | 71587403 | A | G | 0.937605 | 5.66E-10 | 0.00022182 |
| 11 | 71587446 | C | T | 0.937605 | 5.66E-10 | 0.00022182 |
| 11 | 71587449 | A | G | 0.937605 | 5.66E-10 | 0.00022182 |
| 11 | 71587548 | A | G | 0.937605 | 5.66E-10 | 0.00022182 |
| 11 | 71587566 | T | C | 0.937605 | 5.66E-10 | 0.00022182 |
| 11 | 71587575 | T | C | 0.937605 | 5.66E-10 | 0.00022182 |
| 11 | 71587598 | T | C | 0.937605 | 5.66E-10 | 0.00022182 |
| 11 | 71587675 | C | T | 0.937605 | 5.66E-10 | 0.00022182 |
| 11 | 71587849 | A | G | 0.937605 | 5.66E-10 | 0.00022182 |
| 11 | 71587895 | C | G | 0.937605 | 5.66E-10 | 0.00022182 |
| 11 | 71588452 | C | A | 0.937605 | 5.66E-10 | 0.00022182 |
| 11 | 71588494 | A | C | 0.937605 | 5.66E-10 | 0.00022182 |
| 11 | 71588562 | A | G | 0.937605 | 5.66E-10 | 0.00022182 |
| 11 | 71589773 | A | G | 0.937605 | 5.66E-10 | 0.00022182 |
| 11 | 71589795 | A | G | 0.937605 | 5.66E-10 | 0.00022182 |
| 11 | 71589982 | A | G | 0.937605 | 5.66E-10 | 0.00022182 |
| 11 | 71590055 | T | C | 0.937605 | 5.66E-10 | 0.00022182 |
| 11 | 71590489 | G | T | 0.937605 | 5.66E-10 | 0.00022182 |
| 11 | 71591189 | A | G | 0.937605 | 5.66E-10 | 0.00022182 |
| 11 | 71591317 | C | T | 0.937605 | 5.66E-10 | 0.00022182 |
| 11 | 71591366 | A | G | 0.937605 | 5.66E-10 | 0.00022182 |
| 11 | 71591477 | G | A | 0.937605 | 5.66E-10 | 0.00022182 |
| 11 | 71591692 | T | C | 0.937605 | 5.66E-10 | 0.00022182 |
| 11 | 71591931 | T | A | 0.937605 | 5.66E-10 | 0.00022182 |
| 11 | 71591976 | A | G | 0.937605 | 5.66E-10 | 0.00022182 |
| 11 | 71592163 | T | A | 0.937605 | 5.66E-10 | 0.00022182 |
| 11 | 71592266 | T | A | 0.937605 | 5.66E-10 | 0.00022182 |
| 11 | 71592296 | A | C | 0.937605 | 5.66E-10 | 0.00022182 |
| 11 | 71592357 | A | T | 0.937605 | 5.66E-10 | 0.00022182 |
| 11 | 71592428 | A | G | 0.937605 | 5.66E-10 | 0.00022182 |
| 11 | 71592444 | T | G | 0.937605 | 5.66E-10 | 0.00022182 |
| 11 | 71592744 | T | C | 0.937605 | 5.66E-10 | 0.00022182 |
| 11 | 71592855 | A | G | 0.937605 | 5.66E-10 | 0.00022182 |
| 11 | 71592934 | G | A | 0.937605 | 5.66E-10 | 0.00022182 |
| 11 | 71593182 | T | C | 0.937605 | 5.66E-10 | 0.00022182 |

| | | | | | | |
|---|---|---|---|---|---|---|
| 11 | 71593226 | G | C | 0.937605 | 5.66E-10 | 0.00022182 |
| 11 | 71593254 | C | A | 0.937605 | 5.66E-10 | 0.00022182 |
| 11 | 71593255 | A | C | 0.937605 | 5.66E-10 | 0.00022182 |
| 11 | 71593304 | T | C | 0.937605 | 5.66E-10 | 0.00022182 |
| 11 | 71593325 | C | T | 0.937605 | 5.66E-10 | 0.00022182 |
| 11 | 71593327 | G | C | 0.937605 | 5.66E-10 | 0.00022182 |
| 11 | 71593444 | A | G | 0.937605 | 5.66E-10 | 0.00022182 |
| 11 | 71593496 | A | G | 0.937605 | 5.66E-10 | 0.00022182 |
| 11 | 71593533 | A | T | 0.937605 | 5.66E-10 | 0.00022182 |
| 11 | 71593535 | T | A | 0.937605 | 5.66E-10 | 0.00022182 |
| 11 | 71594173 | C | G | 0.937605 | 5.66E-10 | 0.00022182 |
| 11 | 71594223 | C | T | 0.937605 | 5.66E-10 | 0.00022182 |
| 11 | 71594419 | T | C | 0.937605 | 5.66E-10 | 0.00022182 |
| 11 | 71594751 | G | A | 0.937605 | 5.66E-10 | 0.00022182 |
| 11 | 71594844 | T | C | 0.937605 | 5.66E-10 | 0.00022182 |
| 11 | 71594872 | A | G | 0.937605 | 5.66E-10 | 0.00022182 |
| 11 | 71594911 | A | G | 0.937605 | 5.66E-10 | 0.00022182 |
| 11 | 71595379 | A | T | 0.937605 | 5.66E-10 | 0.00022182 |
| 11 | 71595527 | A | C | 0.937605 | 5.66E-10 | 0.00022182 |
| 11 | 71595960 | T | C | 0.937605 | 5.66E-10 | 0.00022182 |
| 11 | 71596912 | A | T | 0.937605 | 5.66E-10 | 0.00022182 |
| 11 | 71597108 | T | C | 0.937605 | 5.66E-10 | 0.00022182 |
| 11 | 71597559 | A | G | 0.937605 | 5.66E-10 | 0.00022182 |
| 11 | 71598646 | T | C | 0.937605 | 5.66E-10 | 0.00022182 |
| 11 | 71598983 | A | G | 0.937605 | 5.66E-10 | 0.00022182 |
| 11 | 71600789 | T | C | 0.937605 | 5.66E-10 | 0.00022182 |
| 11 | 71601408 | C | T | 0.937605 | 5.66E-10 | 0.00022182 |
| 11 | 71601418 | A | G | 0.937605 | 5.66E-10 | 0.00022182 |
| 11 | 71601560 | G | A | 0.937605 | 5.66E-10 | 0.00022182 |
| 11 | 71601738 | A | G | 0.937605 | 5.66E-10 | 0.00022182 |
| 11 | 71601882 | A | T | 0.937605 | 5.66E-10 | 0.00022182 |
| 11 | 71601883 | A | T | 0.937605 | 5.66E-10 | 0.00022182 |
| 11 | 71602404 | C | A | 0.937605 | 5.66E-10 | 0.00022182 |
| 11 | 71602509 | T | C | 0.937605 | 5.66E-10 | 0.00022182 |
| 11 | 71602590 | T | C | 0.937605 | 5.66E-10 | 0.00022182 |
| 11 | 71602839 | A | G | 0.937605 | 5.66E-10 | 0.00022182 |
| 11 | 71602843 | A | C | 0.937605 | 5.66E-10 | 0.00022182 |
| 11 | 71602875 | G | A | 0.937605 | 5.66E-10 | 0.00022182 |
| 11 | 71602881 | T | C | 0.937605 | 5.66E-10 | 0.00022182 |
| 11 | 71602937 | A | G | 0.937605 | 5.66E-10 | 0.00022182 |
| 11 | 71603001 | C | T | 0.937605 | 5.66E-10 | 0.00022182 |
| 11 | 71603743 | C | T | 0.937605 | 5.66E-10 | 0.00022182 |
| 11 | 71604239 | T | G | 0.937605 | 5.66E-10 | 0.00022182 |
| 11 | 71604273 | T | C | 0.937605 | 5.66E-10 | 0.00022182 |
| 11 | 71604562 | C | T | 0.937605 | 5.66E-10 | 0.00022182 |
| 11 | 71604600 | G | A | 0.937605 | 5.66E-10 | 0.00022182 |
| 11 | 71604655 | G | C | 0.937605 | 5.66E-10 | 0.00022182 |
| 11 | 71604718 | C | T | 0.937605 | 5.66E-10 | 0.00022182 |
| 11 | 71605348 | T | C | 0.937605 | 5.66E-10 | 0.00022182 |
| 11 | 71605437 | A | T | 0.937605 | 5.66E-10 | 0.00022182 |
| 11 | 71605803 | G | C | 0.937605 | 5.66E-10 | 0.00022182 |
| 11 | 71605892 | T | C | 0.937605 | 5.66E-10 | 0.00022182 |
| 11 | 71605943 | A | G | 0.937605 | 5.66E-10 | 0.00022182 |
| 11 | 71606348 | C | G | 0.937605 | 5.66E-10 | 0.00022182 |
| 11 | 71606408 | C | G | 0.937605 | 5.66E-10 | 0.00022182 |
| 11 | 71606415 | A | G | 0.937605 | 5.66E-10 | 0.00022182 |
| 11 | 71606434 | C | G | 0.937605 | 5.66E-10 | 0.00022182 |
| 11 | 71606439 | T | A | 0.937605 | 5.66E-10 | 0.00022182 |
| 11 | 71606689 | C | T | 0.937605 | 5.66E-10 | 0.00022182 |
| 11 | 71606706 | G | T | 0.937605 | 5.66E-10 | 0.00022182 |
| 11 | 71606738 | A | T | 0.937605 | 5.66E-10 | 0.00022182 |
| 11 | 71606739 | A | T | 0.937605 | 5.66E-10 | 0.00022182 |
| 11 | 71606997 | G | C | 0.937605 | 5.66E-10 | 0.00022182 |
| 11 | 71607052 | A | G | 0.937605 | 5.66E-10 | 0.00022182 |
| 11 | 71607187 | A | C | 0.937605 | 5.66E-10 | 0.00022182 |
| 11 | 71607193 | G | A | 0.937605 | 5.66E-10 | 0.00022182 |
| 11 | 71607231 | T | C | 0.937605 | 5.66E-10 | 0.00022182 |

| | | | | | | |
|---|---|---|---|---|---|---|
| 11 | 71607237 | T | A | 0.937605 | 5.66E-10 | 0.00022182 |
| 11 | 71607460 | A | C | 0.937605 | 5.66E-10 | 0.00022182 |
| 11 | 71607552 | T | C | 0.937605 | 5.66E-10 | 0.00022182 |
| 11 | 71607694 | T | C | 0.937605 | 5.66E-10 | 0.00022182 |
| 11 | 71607695 | G | A | 0.937605 | 5.66E-10 | 0.00022182 |
| 11 | 71607840 | T | G | 0.937605 | 5.66E-10 | 0.00022182 |
| 11 | 71607899 | C | T | 0.937605 | 5.66E-10 | 0.00022182 |
| 11 | 71607900 | C | T | 0.937605 | 5.66E-10 | 0.00022182 |
| 11 | 71607901 | C | T | 0.937605 | 5.66E-10 | 0.00022182 |
| 11 | 71608292 | T | C | 0.937605 | 5.66E-10 | 0.00022182 |
| 11 | 71608327 | C | T | 0.937605 | 5.66E-10 | 0.00022182 |
| 11 | 71608455 | A | G | 0.937605 | 5.66E-10 | 0.00022182 |
| 11 | 71608522 | T | C | 0.937605 | 5.66E-10 | 0.00022182 |
| 11 | 71608590 | T | G | 0.937605 | 5.66E-10 | 0.00022182 |
| 11 | 71608727 | G | A | 0.937605 | 5.66E-10 | 0.00022182 |
| 11 | 71608889 | C | T | 0.937605 | 5.66E-10 | 0.00022182 |
| 11 | 71608947 | A | G | 0.937605 | 5.66E-10 | 0.00022182 |
| 11 | 71609548 | A | G | 0.937605 | 5.66E-10 | 0.00022182 |
| 11 | 71609874 | C | T | 0.937605 | 5.66E-10 | 0.00022182 |
| 11 | 71610053 | G | T | 0.937605 | 5.66E-10 | 0.00022182 |
| 11 | 71610063 | T | G | 0.937605 | 5.66E-10 | 0.00022182 |
| 11 | 71610064 | T | C | 0.937605 | 5.66E-10 | 0.00022182 |
| 11 | 71610259 | C | T | 0.937605 | 5.66E-10 | 0.00022182 |
| 11 | 71610597 | G | T | 0.937605 | 5.66E-10 | 0.00022182 |
| 11 | 71610614 | A | G | 0.937605 | 5.66E-10 | 0.00022182 |
| 11 | 71610804 | T | G | 0.937605 | 5.66E-10 | 0.00022182 |
| 11 | 71611255 | A | G | 0.937605 | 5.66E-10 | 0.00022182 |
| 11 | 71611451 | T | C | 0.937605 | 5.66E-10 | 0.00022182 |
| 11 | 71611846 | T | C | 0.937605 | 5.66E-10 | 0.00022182 |
| 11 | 71611895 | A | C | 0.937605 | 5.66E-10 | 0.00022182 |
| 11 | 71612519 | C | G | 0.937605 | 5.66E-10 | 0.00022182 |
| 11 | 71612659 | C | T | 0.937605 | 5.66E-10 | 0.00022182 |
| 11 | 71612676 | G | A | 0.937605 | 5.66E-10 | 0.00022182 |
| 11 | 71612677 | A | G | 0.937605 | 5.66E-10 | 0.00022182 |
| 11 | 71612692 | C | G | 0.937605 | 5.66E-10 | 0.00022182 |
| 11 | 71612700 | A | G | 0.937605 | 5.66E-10 | 0.00022182 |
| 11 | 71612704 | T | G | 0.937605 | 5.66E-10 | 0.00022182 |
| 11 | 71612737 | T | G | 0.937605 | 5.66E-10 | 0.00022182 |
| 11 | 71612746 | T | G | 0.937605 | 5.66E-10 | 0.00022182 |
| 11 | 71612813 | T | C | 0.937605 | 5.66E-10 | 0.00022182 |
| 11 | 71612933 | C | T | 0.937605 | 5.66E-10 | 0.00022182 |
| 11 | 71613043 | A | T | 0.937605 | 5.66E-10 | 0.00022182 |
| 11 | 71613076 | G | T | 0.937605 | 5.66E-10 | 0.00022182 |
| 11 | 71613199 | A | G | 0.937605 | 5.66E-10 | 0.00022182 |
| 11 | 71613205 | C | T | 0.937605 | 5.66E-10 | 0.00022182 |
| 11 | 71613232 | C | T | 0.937605 | 5.66E-10 | 0.00022182 |
| 11 | 71613247 | A | G | 0.937605 | 5.66E-10 | 0.00022182 |
| 11 | 71613267 | A | G | 0.937605 | 5.66E-10 | 0.00022182 |
| 11 | 71613548 | T | C | 0.937605 | 5.66E-10 | 0.00022182 |
| 11 | 71614818 | C | T | 0.937605 | 5.66E-10 | 0.00022182 |
| 11 | 71614927 | T | C | 0.937605 | 5.66E-10 | 0.00022182 |
| 11 | 71614946 | C | T | 0.937605 | 5.66E-10 | 0.00022182 |
| 11 | 71614999 | A | G | 0.937605 | 5.66E-10 | 0.00022182 |
| 11 | 71615422 | C | T | 0.937605 | 5.66E-10 | 0.00022182 |
| 11 | 71615539 | A | G | 0.937605 | 5.66E-10 | 0.00022182 |
| 11 | 71615862 | T | C | 0.937605 | 5.66E-10 | 0.00022182 |
| 11 | 71616059 | G | C | 0.937605 | 5.66E-10 | 0.00022182 |
| 11 | 71616813 | A | C | 0.937605 | 5.66E-10 | 0.00022182 |
| 11 | 71617060 | C | T | 0.937605 | 5.66E-10 | 0.00022182 |
| 11 | 71617219 | A | G | 0.937605 | 5.66E-10 | 0.00022182 |
| 11 | 71617436 | T | G | 0.937605 | 5.66E-10 | 0.00022182 |
| 11 | 71617549 | T | C | 0.937605 | 5.66E-10 | 0.00022182 |
| 11 | 71619504 | A | G | 0.937605 | 5.66E-10 | 0.00022182 |
| 11 | 71619506 | T | G | 0.937605 | 5.66E-10 | 0.00022182 |
| 11 | 71621407 | G | C | 0.937605 | 5.66E-10 | 0.00022182 |
| 11 | 71621566 | C | T | 0.937605 | 5.66E-10 | 0.00022182 |
| 11 | 71621639 | G | A | 0.937605 | 5.66E-10 | 0.00022182 |

| | | | | | | |
|---|---|---|---|---|---|---|
| 11 | 71621883 | C | A | 0.937605 | 5.66E-10 | 0.00022182 |
| 11 | 71622038 | C | T | 0.937605 | 5.66E-10 | 0.00022182 |
| 11 | 71622663 | C | T | 0.937605 | 5.66E-10 | 0.00022182 |
| 11 | 71622670 | G | A | 0.937605 | 5.66E-10 | 0.00022182 |
| 11 | 71622733 | C | T | 0.937605 | 5.66E-10 | 0.00022182 |
| 11 | 71622893 | A | C | 0.937605 | 5.66E-10 | 0.00022182 |
| 11 | 71623259 | G | A | 0.937605 | 5.66E-10 | 0.00022182 |
| 11 | 71632256 | G | A | 0.937605 | 5.66E-10 | 0.00022182 |
| 11 | 69601439 | C | T | 0.801012 | 5.90E-10 | 0.00023131 |
| 11 | 69527638 | A | G | 0.812816 | 6.68E-10 | 0.00026175 |
| 11 | 69528017 | G | A | 0.812816 | 6.68E-10 | 0.00026175 |
| 11 | 69528922 | G | T | 0.812816 | 6.68E-10 | 0.00026175 |
| 11 | 69529151 | A | G | 0.812816 | 6.68E-10 | 0.00026175 |
| 11 | 69529174 | G | T | 0.812816 | 6.68E-10 | 0.00026175 |
| 11 | 69529211 | A | G | 0.812816 | 6.68E-10 | 0.00026175 |
| 11 | 69529308 | C | A | 0.812816 | 6.68E-10 | 0.00026175 |
| 11 | 69529313 | T | A | 0.812816 | 6.68E-10 | 0.00026175 |
| 11 | 69529327 | C | T | 0.812816 | 6.68E-10 | 0.00026175 |
| 11 | 69529328 | T | G | 0.812816 | 6.68E-10 | 0.00026175 |
| 11 | 69529396 | T | C | 0.812816 | 6.68E-10 | 0.00026175 |
| 11 | 69529397 | G | C | 0.812816 | 6.68E-10 | 0.00026175 |
| 11 | 69529437 | A | G | 0.812816 | 6.68E-10 | 0.00026175 |
| 11 | 69529473 | C | T | 0.812816 | 6.68E-10 | 0.00026175 |
| 11 | 69529485 | T | C | 0.812816 | 6.68E-10 | 0.00026175 |
| 11 | 69529486 | G | A | 0.812816 | 6.68E-10 | 0.00026175 |
| 11 | 69529510 | T | C | 0.812816 | 6.68E-10 | 0.00026175 |
| 11 | 69529538 | G | C | 0.812816 | 6.68E-10 | 0.00026175 |
| 11 | 69529551 | C | A | 0.812816 | 6.68E-10 | 0.00026175 |
| 11 | 69529556 | A | G | 0.812816 | 6.68E-10 | 0.00026175 |
| 11 | 69529672 | T | G | 0.812816 | 6.68E-10 | 0.00026175 |
| 11 | 69529694 | C | T | 0.812816 | 6.68E-10 | 0.00026175 |
| 11 | 69529789 | G | A | 0.812816 | 6.68E-10 | 0.00026175 |
| 11 | 69529813 | G | T | 0.812816 | 6.68E-10 | 0.00026175 |
| 11 | 69529827 | T | A | 0.812816 | 6.68E-10 | 0.00026175 |
| 11 | 69529837 | A | T | 0.812816 | 6.68E-10 | 0.00026175 |
| 11 | 69529840 | C | T | 0.812816 | 6.68E-10 | 0.00026175 |
| 11 | 69529886 | G | C | 0.812816 | 6.68E-10 | 0.00026175 |
| 11 | 69529924 | G | C | 0.812816 | 6.68E-10 | 0.00026175 |
| 11 | 69529927 | T | C | 0.812816 | 6.68E-10 | 0.00026175 |
| 11 | 69529971 | A | G | 0.812816 | 6.68E-10 | 0.00026175 |
| 11 | 69529987 | T | C | 0.812816 | 6.68E-10 | 0.00026175 |
| 11 | 69530090 | G | A | 0.812816 | 6.68E-10 | 0.00026175 |
| 11 | 69530121 | C | A | 0.812816 | 6.68E-10 | 0.00026175 |
| 11 | 69530169 | G | T | 0.812816 | 6.68E-10 | 0.00026175 |
| 11 | 69530186 | T | C | 0.812816 | 6.68E-10 | 0.00026175 |
| 11 | 69530204 | C | T | 0.812816 | 6.68E-10 | 0.00026175 |
| 11 | 69530211 | A | T | 0.812816 | 6.68E-10 | 0.00026175 |
| 11 | 69530232 | A | G | 0.812816 | 6.68E-10 | 0.00026175 |
| 11 | 69530322 | G | A | 0.812816 | 6.68E-10 | 0.00026175 |
| 11 | 69530332 | A | C | 0.812816 | 6.68E-10 | 0.00026175 |
| 11 | 69530333 | G | A | 0.812816 | 6.68E-10 | 0.00026175 |
| 11 | 69530335 | C | T | 0.812816 | 6.68E-10 | 0.00026175 |
| 11 | 69532518 | T | C | 0.812816 | 6.68E-10 | 0.00026175 |
| 11 | 69532663 | T | A | 0.812816 | 6.68E-10 | 0.00026175 |
| 11 | 69532670 | T | C | 0.812816 | 6.68E-10 | 0.00026175 |
| 11 | 69536874 | C | T | 0.812816 | 6.68E-10 | 0.00026175 |
| 11 | 69537000 | T | C | 0.812816 | 6.68E-10 | 0.00026175 |
| 11 | 69537841 | G | T | 0.812816 | 6.68E-10 | 0.00026175 |
| 11 | 69538847 | T | A | 0.812816 | 6.68E-10 | 0.00026175 |
| 11 | 69538857 | C | T | 0.812816 | 6.68E-10 | 0.00026175 |
| 11 | 69538862 | G | A | 0.812816 | 6.68E-10 | 0.00026175 |
| 11 | 69538878 | C | A | 0.812816 | 6.68E-10 | 0.00026175 |
| 11 | 69539056 | T | C | 0.812816 | 6.68E-10 | 0.00026175 |
| 11 | 69539182 | T | C | 0.812816 | 6.68E-10 | 0.00026175 |
| 11 | 69539191 | G | A | 0.812816 | 6.68E-10 | 0.00026175 |
| 11 | 69539195 | C | T | 0.812816 | 6.68E-10 | 0.00026175 |
| 11 | 69539335 | C | A | 0.812816 | 6.68E-10 | 0.00026175 |

| | | | | | | |
|---|---|---|---|---|---|---|
| 11 | 69539342 | T | C | 0.812816 | 6.68E-10 | 0.00026175 |
| 11 | 69539349 | A | C | 0.812816 | 6.68E-10 | 0.00026175 |
| 11 | 69539359 | A | G | 0.812816 | 6.68E-10 | 0.00026175 |
| 11 | 69539388 | T | C | 0.812816 | 6.68E-10 | 0.00026175 |
| 11 | 69539430 | T | C | 0.812816 | 6.68E-10 | 0.00026175 |
| 11 | 69539721 | C | T | 0.812816 | 6.68E-10 | 0.00026175 |
| 11 | 69539722 | T | C | 0.812816 | 6.68E-10 | 0.00026175 |
| 11 | 69539904 | A | G | 0.812816 | 6.68E-10 | 0.00026175 |
| 11 | 69540713 | G | A | 0.812816 | 6.68E-10 | 0.00026175 |
| 11 | 69540776 | C | T | 0.812816 | 6.68E-10 | 0.00026175 |
| 11 | 69540832 | A | G | 0.812816 | 6.68E-10 | 0.00026175 |
| 11 | 69541073 | C | T | 0.812816 | 6.68E-10 | 0.00026175 |
| 11 | 69541138 | A | G | 0.812816 | 6.68E-10 | 0.00026175 |
| 11 | 69541223 | C | A | 0.812816 | 6.68E-10 | 0.00026175 |
| 11 | 69541237 | T | C | 0.812816 | 6.68E-10 | 0.00026175 |
| 11 | 69541355 | T | C | 0.812816 | 6.68E-10 | 0.00026175 |
| 11 | 69541419 | T | C | 0.812816 | 6.68E-10 | 0.00026175 |
| 11 | 69541431 | A | G | 0.812816 | 6.68E-10 | 0.00026175 |
| 11 | 69541448 | T | C | 0.812816 | 6.68E-10 | 0.00026175 |
| 11 | 69541540 | T | C | 0.812816 | 6.68E-10 | 0.00026175 |
| 11 | 69541664 | T | C | 0.812816 | 6.68E-10 | 0.00026175 |
| 11 | 69541684 | A | G | 0.812816 | 6.68E-10 | 0.00026175 |
| 11 | 69542146 | G | A | 0.812816 | 6.68E-10 | 0.00026175 |
| 11 | 69542160 | T | C | 0.812816 | 6.68E-10 | 0.00026175 |
| 11 | 69542172 | A | T | 0.812816 | 6.68E-10 | 0.00026175 |
| 11 | 69542194 | C | T | 0.812816 | 6.68E-10 | 0.00026175 |
| 11 | 69542207 | A | G | 0.812816 | 6.68E-10 | 0.00026175 |
| 11 | 69542231 | C | G | 0.812816 | 6.68E-10 | 0.00026175 |
| 11 | 69542241 | A | G | 0.812816 | 6.68E-10 | 0.00026175 |
| 11 | 69542402 | G | A | 0.812816 | 6.68E-10 | 0.00026175 |
| 11 | 69542583 | C | T | 0.812816 | 6.68E-10 | 0.00026175 |
| 11 | 69542595 | T | A | 0.812816 | 6.68E-10 | 0.00026175 |
| 11 | 69542680 | A | G | 0.812816 | 6.68E-10 | 0.00026175 |
| 11 | 69542728 | G | C | 0.812816 | 6.68E-10 | 0.00026175 |
| 11 | 69542821 | A | C | 0.812816 | 6.68E-10 | 0.00026175 |
| 11 | 69544344 | C | T | 0.812816 | 6.68E-10 | 0.00026175 |
| 11 | 69544989 | C | T | 0.812816 | 6.68E-10 | 0.00026175 |
| 11 | 69545196 | C | A | 0.812816 | 6.68E-10 | 0.00026175 |
| 11 | 69545398 | G | A | 0.812816 | 6.68E-10 | 0.00026175 |
| 11 | 69545442 | T | C | 0.812816 | 6.68E-10 | 0.00026175 |
| 11 | 69545475 | T | C | 0.812816 | 6.68E-10 | 0.00026175 |
| 11 | 69545582 | A | G | 0.812816 | 6.68E-10 | 0.00026175 |
| 11 | 69545601 | T | A | 0.812816 | 6.68E-10 | 0.00026175 |
| 11 | 69545742 | T | C | 0.812816 | 6.68E-10 | 0.00026175 |
| 11 | 69546351 | G | T | 0.812816 | 6.68E-10 | 0.00026175 |
| 11 | 69546384 | G | T | 0.812816 | 6.68E-10 | 0.00026175 |
| 11 | 69546418 | A | G | 0.812816 | 6.68E-10 | 0.00026175 |
| 11 | 69546435 | G | T | 0.812816 | 6.68E-10 | 0.00026175 |
| 11 | 69546455 | C | T | 0.812816 | 6.68E-10 | 0.00026175 |
| 11 | 69546683 | A | G | 0.812816 | 6.68E-10 | 0.00026175 |
| 11 | 69546842 | A | G | 0.812816 | 6.68E-10 | 0.00026175 |
| 11 | 69546912 | A | G | 0.812816 | 6.68E-10 | 0.00026175 |
| 11 | 69546913 | G | C | 0.812816 | 6.68E-10 | 0.00026175 |
| 11 | 69546933 | C | A | 0.812816 | 6.68E-10 | 0.00026175 |
| 11 | 69547089 | T | C | 0.812816 | 6.68E-10 | 0.00026175 |
| 11 | 69547101 | G | C | 0.812816 | 6.68E-10 | 0.00026175 |
| 11 | 69547109 | T | C | 0.812816 | 6.68E-10 | 0.00026175 |
| 11 | 69547126 | T | C | 0.812816 | 6.68E-10 | 0.00026175 |
| 11 | 69547148 | G | A | 0.812816 | 6.68E-10 | 0.00026175 |
| 11 | 69547196 | G | A | 0.812816 | 6.68E-10 | 0.00026175 |
| 11 | 69547447 | C | A | 0.812816 | 6.68E-10 | 0.00026175 |
| 11 | 69547448 | T | C | 0.812816 | 6.68E-10 | 0.00026175 |
| 11 | 69547453 | A | T | 0.812816 | 6.68E-10 | 0.00026175 |
| 11 | 69547461 | T | C | 0.812816 | 6.68E-10 | 0.00026175 |
| 11 | 69547466 | A | T | 0.812816 | 6.68E-10 | 0.00026175 |
| 11 | 69547469 | A | G | 0.812816 | 6.68E-10 | 0.00026175 |
| 11 | 69547639 | A | G | 0.812816 | 6.68E-10 | 0.00026175 |

| | | | | | | |
|---|---|---|---|---|---|---|
| 11 | 69547647 | A | G | 0.812816 | 6.68E-10 | 0.00026175 |
| 11 | 69547657 | G | C | 0.812816 | 6.68E-10 | 0.00026175 |
| 11 | 69547735 | C | T | 0.812816 | 6.68E-10 | 0.00026175 |
| 11 | 69547894 | G | A | 0.812816 | 6.68E-10 | 0.00026175 |
| 11 | 69547916 | G | A | 0.812816 | 6.68E-10 | 0.00026175 |
| 11 | 69548232 | T | A | 0.812816 | 6.68E-10 | 0.00026175 |
| 11 | 69548272 | T | A | 0.812816 | 6.68E-10 | 0.00026175 |
| 11 | 69548565 | A | G | 0.812816 | 6.68E-10 | 0.00026175 |
| 11 | 69548609 | G | T | 0.812816 | 6.68E-10 | 0.00026175 |
| 11 | 69549735 | G | C | 0.812816 | 6.68E-10 | 0.00026175 |
| 11 | 69550083 | G | T | 0.812816 | 6.68E-10 | 0.00026175 |
| 11 | 69550320 | C | T | 0.812816 | 6.68E-10 | 0.00026175 |
| 11 | 69550401 | A | C | 0.812816 | 6.68E-10 | 0.00026175 |
| 11 | 69550414 | C | A | 0.812816 | 6.68E-10 | 0.00026175 |
| 11 | 69550487 | T | C | 0.812816 | 6.68E-10 | 0.00026175 |
| 11 | 69550488 | A | G | 0.812816 | 6.68E-10 | 0.00026175 |
| 11 | 69551439 | C | A | 0.812816 | 6.68E-10 | 0.00026175 |
| 11 | 69551461 | C | T | 0.812816 | 6.68E-10 | 0.00026175 |
| 11 | 69551473 | C | A | 0.812816 | 6.68E-10 | 0.00026175 |
| 11 | 69551530 | T | C | 0.812816 | 6.68E-10 | 0.00026175 |
| 11 | 69551626 | T | C | 0.812816 | 6.68E-10 | 0.00026175 |
| 11 | 69551692 | C | T | 0.812816 | 6.68E-10 | 0.00026175 |
| 11 | 69551986 | T | C | 0.812816 | 6.68E-10 | 0.00026175 |
| 11 | 69552014 | C | G | 0.812816 | 6.68E-10 | 0.00026175 |
| 11 | 69552490 | T | C | 0.812816 | 6.68E-10 | 0.00026175 |
| 11 | 69552828 | T | C | 0.812816 | 6.68E-10 | 0.00026175 |
| 11 | 63812204 | C | A | 0.89629 | 6.81E-10 | 0.00026709 |
| 11 | 69531611 | C | T | 0.797639 | 6.82E-10 | 0.0002673 |
| 11 | 69531695 | G | A | 0.797639 | 6.82E-10 | 0.0002673 |
| 11 | 69531792 | G | T | 0.797639 | 6.82E-10 | 0.0002673 |
| 11 | 69531852 | T | C | 0.797639 | 6.82E-10 | 0.0002673 |
| 11 | 69918765 | G | T | 0.874368 | 7.34E-10 | 0.00028755 |
| 11 | 69918766 | A | G | 0.874368 | 7.34E-10 | 0.00028755 |
| 11 | 64965365 | T | G | 0.719224 | 7.42E-10 | 0.00029086 |
| 11 | 64966275 | A | G | 0.719224 | 7.42E-10 | 0.00029086 |
| 11 | 64970832 | T | C | 0.719224 | 7.42E-10 | 0.00029086 |
| 11 | 69530651 | A | G | 0.813659 | 8.33E-10 | 0.00032662 |
| 11 | 69528130 | A | G | 0.799325 | 8.41E-10 | 0.00032961 |
| 11 | 69528205 | A | G | 0.799325 | 8.41E-10 | 0.00032961 |
| 11 | 69528277 | C | T | 0.799325 | 8.41E-10 | 0.00032961 |
| 11 | 69532935 | G | A | 0.799325 | 8.41E-10 | 0.00032961 |
| 11 | 69533078 | T | C | 0.799325 | 8.41E-10 | 0.00032961 |
| 11 | 69536192 | C | T | 0.799325 | 8.41E-10 | 0.00032961 |
| 11 | 69539033 | C | T | 0.799325 | 8.41E-10 | 0.00032961 |
| 11 | 69539530 | A | G | 0.799325 | 8.41E-10 | 0.00032961 |
| 11 | 69539708 | C | G | 0.799325 | 8.41E-10 | 0.00032961 |
| 11 | 69539741 | G | A | 0.799325 | 8.41E-10 | 0.00032961 |
| 11 | 69540114 | T | C | 0.799325 | 8.41E-10 | 0.00032961 |
| 11 | 69540133 | G | A | 0.799325 | 8.41E-10 | 0.00032961 |
| 11 | 69540160 | A | C | 0.799325 | 8.41E-10 | 0.00032961 |
| 11 | 69540201 | T | C | 0.799325 | 8.41E-10 | 0.00032961 |
| 11 | 69540203 | G | A | 0.799325 | 8.41E-10 | 0.00032961 |
| 11 | 69540216 | A | G | 0.799325 | 8.41E-10 | 0.00032961 |
| 11 | 69540991 | C | T | 0.799325 | 8.41E-10 | 0.00032961 |
| 11 | 69542184 | C | T | 0.799325 | 8.41E-10 | 0.00032961 |
| 11 | 69542329 | C | T | 0.799325 | 8.41E-10 | 0.00032961 |
| 11 | 69542633 | C | T | 0.799325 | 8.41E-10 | 0.00032961 |
| 11 | 69542799 | G | T | 0.799325 | 8.41E-10 | 0.00032961 |
| 11 | 69542812 | T | G | 0.799325 | 8.41E-10 | 0.00032961 |
| 11 | 69544610 | A | G | 0.799325 | 8.41E-10 | 0.00032961 |
| 11 | 69544908 | T | C | 0.799325 | 8.41E-10 | 0.00032961 |
| 11 | 69545419 | C | T | 0.799325 | 8.41E-10 | 0.00032961 |
| 11 | 69545488 | G | A | 0.799325 | 8.41E-10 | 0.00032961 |
| 11 | 69546166 | T | C | 0.799325 | 8.41E-10 | 0.00032961 |
| 11 | 69546753 | G | A | 0.799325 | 8.41E-10 | 0.00032961 |
| 11 | 69547117 | A | G | 0.799325 | 8.41E-10 | 0.00032961 |
| 11 | 69547127 | G | C | 0.799325 | 8.41E-10 | 0.00032961 |

| | | | | | | |
|---|---|---|---|---|---|---|
| 11 | 69547483 | A | G | 0.799325 | 8.41E-10 | 0.00032961 |
| 11 | 69548353 | T | C | 0.799325 | 8.41E-10 | 0.00032961 |
| 11 | 69548355 | T | G | 0.799325 | 8.41E-10 | 0.00032961 |
| 11 | 69548502 | T | C | 0.799325 | 8.41E-10 | 0.00032961 |
| 11 | 69548551 | A | G | 0.799325 | 8.41E-10 | 0.00032961 |
| 11 | 69548556 | T | A | 0.799325 | 8.41E-10 | 0.00032961 |
| 11 | 69548592 | C | T | 0.799325 | 8.41E-10 | 0.00032961 |
| 11 | 69548772 | G | A | 0.799325 | 8.41E-10 | 0.00032961 |
| 11 | 69548807 | T | C | 0.799325 | 8.41E-10 | 0.00032961 |
| 11 | 69548912 | A | G | 0.799325 | 8.41E-10 | 0.00032961 |
| 11 | 69548981 | C | T | 0.799325 | 8.41E-10 | 0.00032961 |
| 11 | 69549129 | T | C | 0.799325 | 8.41E-10 | 0.00032961 |
| 11 | 69549179 | A | G | 0.799325 | 8.41E-10 | 0.00032961 |
| 11 | 69549281 | G | A | 0.799325 | 8.41E-10 | 0.00032961 |
| 11 | 69549405 | A | G | 0.799325 | 8.41E-10 | 0.00032961 |
| 11 | 69549503 | G | C | 0.799325 | 8.41E-10 | 0.00032961 |
| 11 | 69549526 | C | T | 0.799325 | 8.41E-10 | 0.00032961 |
| 11 | 69549565 | A | G | 0.799325 | 8.41E-10 | 0.00032961 |
| 11 | 69549570 | G | A | 0.799325 | 8.41E-10 | 0.00032961 |
| 11 | 69549577 | C | G | 0.799325 | 8.41E-10 | 0.00032961 |
| 11 | 69549592 | G | A | 0.799325 | 8.41E-10 | 0.00032961 |
| 11 | 69549599 | T | C | 0.799325 | 8.41E-10 | 0.00032961 |
| 11 | 69549890 | A | G | 0.799325 | 8.41E-10 | 0.00032961 |
| 11 | 69549934 | C | T | 0.799325 | 8.41E-10 | 0.00032961 |
| 11 | 69549938 | A | C | 0.799325 | 8.41E-10 | 0.00032961 |
| 11 | 69549978 | C | T | 0.799325 | 8.41E-10 | 0.00032961 |
| 11 | 69550900 | T | C | 0.799325 | 8.41E-10 | 0.00032961 |
| 11 | 69551096 | A | G | 0.799325 | 8.41E-10 | 0.00032961 |
| 11 | 71599742 | T | C | 0.935919 | 1.04E-09 | 0.00040579 |
| 11 | 71505772 | G | A | 0.938449 | 1.06E-09 | 0.00041475 |
| 11 | 64980562 | A | G | 0.866779 | 1.09E-09 | 0.00042872 |
| 11 | 66065829 | C | T | 0.756324 | 1.14E-09 | 0.00044832 |
| 11 | 71589144 | T | G | 0.936762 | 1.15E-09 | 0.00045239 |
| 11 | 65074136 | G | A | 0.870995 | 1.24E-09 | 0.00048574 |
| 11 | 69539987 | T | C | 0.798482 | 1.26E-09 | 0.00049567 |
| 11 | 69539990 | G | A | 0.798482 | 1.26E-09 | 0.00049567 |
| 11 | 69539996 | C | T | 0.798482 | 1.26E-09 | 0.00049567 |
| 11 | 69540061 | T | A | 0.798482 | 1.26E-09 | 0.00049567 |
| 11 | 69540068 | G | A | 0.798482 | 1.26E-09 | 0.00049567 |
| 11 | 69540100 | T | A | 0.798482 | 1.26E-09 | 0.00049567 |
| 11 | 69540104 | C | G | 0.798482 | 1.26E-09 | 0.00049567 |
| 11 | 64079984 | T | G | 0.816189 | 1.28E-09 | 0.00050207 |
| 11 | 64087760 | C | T | 0.816189 | 1.28E-09 | 0.00050207 |
| 11 | 71591597 | A | C | 0.928331 | 1.35E-09 | 0.00053003 |
| 11 | 69534686 | T | C | 0.811973 | 1.52E-09 | 0.00059493 |
| 11 | 69534729 | A | G | 0.811973 | 1.52E-09 | 0.00059493 |
| 11 | 69534823 | G | A | 0.811973 | 1.52E-09 | 0.00059493 |
| 11 | 69534849 | G | A | 0.811973 | 1.52E-09 | 0.00059493 |
| 11 | 69534851 | C | T | 0.811973 | 1.52E-09 | 0.00059493 |
| 11 | 69535460 | T | C | 0.811973 | 1.52E-09 | 0.00059493 |
| 11 | 64897493 | A | G | 0.714165 | 1.67E-09 | 0.00065603 |
| 11 | 71621244 | C | G | 0.935919 | 1.72E-09 | 0.00067325 |
| 11 | 71613898 | C | T | 0.939292 | 1.75E-09 | 0.00068532 |
| 11 | 71613951 | T | C | 0.939292 | 1.75E-09 | 0.00068532 |
| 11 | 70490283 | A | G | 0.91231 | 1.77E-09 | 0.00069501 |
| 11 | 69534446 | T | G | 0.799325 | 1.79E-09 | 0.00070149 |
| 11 | 69534669 | G | A | 0.799325 | 1.79E-09 | 0.00070149 |
| 11 | 69535098 | C | T | 0.799325 | 1.79E-09 | 0.00070149 |
| 11 | 69535123 | G | C | 0.799325 | 1.79E-09 | 0.00070149 |
| 11 | 69535888 | C | T | 0.798482 | 1.79E-09 | 0.00070192 |
| 11 | 69535986 | C | T | 0.798482 | 1.79E-09 | 0.00070192 |
| 11 | 69673557 | A | G | 0.815346 | 1.94E-09 | 0.00075977 |
| 11 | 69533025 | A | G | 0.800169 | 1.98E-09 | 0.00077772 |
| 11 | 71605920 | A | G | 0.935919 | 2.01E-09 | 0.0007874 |
| 11 | 66075368 | C | A | 0.758853 | 2.02E-09 | 0.00079168 |
| 11 | 66053456 | C | T | 0.757167 | 2.04E-09 | 0.00079913 |
| 11 | 69652227 | A | T | 0.81113 | 2.07E-09 | 0.00081199 |

| | | | | | | |
|---|---|---|---|---|---|---|
| 11 | 69536176 | A | C | 0.800169 | 2.13E-09 | 0.00083637 |
| 11 | 71618869 | T | C | 0.936762 | 2.19E-09 | 0.00085987 |
| 11 | 71619115 | C | T | 0.936762 | 2.19E-09 | 0.00085987 |
| 11 | 71619797 | C | T | 0.936762 | 2.19E-09 | 0.00085987 |
| 11 | 71619828 | A | G | 0.936762 | 2.19E-09 | 0.00085987 |
| 11 | 71620153 | C | T | 0.936762 | 2.19E-09 | 0.00085987 |
| 11 | 71620593 | A | C | 0.936762 | 2.19E-09 | 0.00085987 |
| 11 | 71621100 | A | G | 0.936762 | 2.19E-09 | 0.00085987 |
| 11 | 64544422 | G | A | 0.909781 | 2.21E-09 | 0.00086511 |
| 11 | 69982829 | A | G | 0.921585 | 2.28E-09 | 0.00089529 |
| 11 | 66055632 | C | A | 0.854975 | 2.40E-09 | 0.00094013 |
| 11 | 66055638 | C | T | 0.854975 | 2.40E-09 | 0.00094013 |
| 11 | 66055639 | A | G | 0.854975 | 2.40E-09 | 0.00094013 |
| 11 | 66056006 | T | C | 0.854132 | 2.55E-09 | 0.00099826 |
| 11 | 71565301 | A | G | 0.936762 | 2.75E-09 | 0.00107618 |
| 11 | 71565371 | T | G | 0.936762 | 2.75E-09 | 0.00107618 |
| 11 | 71565874 | A | G | 0.936762 | 2.75E-09 | 0.00107618 |
| 11 | 71565890 | C | T | 0.936762 | 2.75E-09 | 0.00107618 |
| 11 | 71565927 | G | A | 0.936762 | 2.75E-09 | 0.00107618 |
| 11 | 71565953 | A | G | 0.936762 | 2.75E-09 | 0.00107618 |
| 11 | 71565979 | C | T | 0.936762 | 2.75E-09 | 0.00107618 |
| 11 | 71566050 | G | T | 0.936762 | 2.75E-09 | 0.00107618 |
| 11 | 71566132 | A | G | 0.936762 | 2.75E-09 | 0.00107618 |
| 11 | 71566168 | T | C | 0.936762 | 2.75E-09 | 0.00107618 |
| 11 | 71566222 | A | G | 0.936762 | 2.75E-09 | 0.00107618 |
| 11 | 71566594 | G | A | 0.936762 | 2.75E-09 | 0.00107618 |
| 11 | 71566836 | C | T | 0.936762 | 2.75E-09 | 0.00107618 |
| 11 | 64058129 | C | A | 0.815346 | 2.82E-09 | 0.00110346 |
| 11 | 64060157 | C | T | 0.815346 | 2.82E-09 | 0.00110346 |
| 11 | 64060526 | G | A | 0.815346 | 2.82E-09 | 0.00110346 |
| 11 | 64061404 | G | A | 0.815346 | 2.82E-09 | 0.00110346 |
| 11 | 64071550 | A | G | 0.815346 | 2.82E-09 | 0.00110346 |
| 11 | 64071579 | G | T | 0.815346 | 2.82E-09 | 0.00110346 |
| 11 | 64072101 | C | T | 0.815346 | 2.82E-09 | 0.00110346 |
| 11 | 64073544 | G | C | 0.815346 | 2.82E-09 | 0.00110346 |
| 11 | 64073918 | A | G | 0.815346 | 2.82E-09 | 0.00110346 |
| 11 | 64076075 | C | T | 0.815346 | 2.82E-09 | 0.00110346 |
| 11 | 64076153 | A | G | 0.815346 | 2.82E-09 | 0.00110346 |
| 11 | 64076805 | A | G | 0.815346 | 2.82E-09 | 0.00110346 |
| 11 | 64076888 | G | A | 0.815346 | 2.82E-09 | 0.00110346 |
| 11 | 64077229 | T | C | 0.815346 | 2.82E-09 | 0.00110346 |
| 11 | 64077246 | G | A | 0.815346 | 2.82E-09 | 0.00110346 |
| 11 | 64077407 | C | T | 0.815346 | 2.82E-09 | 0.00110346 |
| 11 | 64077465 | C | A | 0.815346 | 2.82E-09 | 0.00110346 |
| 11 | 64077703 | C | T | 0.815346 | 2.82E-09 | 0.00110346 |
| 11 | 64085095 | C | T | 0.815346 | 2.82E-09 | 0.00110346 |
| 11 | 64099019 | C | T | 0.815346 | 2.82E-09 | 0.00110346 |
| 11 | 71550560 | A | T | 0.935919 | 2.87E-09 | 0.00112663 |
| 11 | 64269285 | G | T | 0.768971 | 2.97E-09 | 0.00116356 |
| 11 | 64269318 | T | C | 0.768971 | 2.97E-09 | 0.00116356 |
| 11 | 69534945 | C | G | 0.818718 | 3.07E-09 | 0.00120507 |
| 11 | 69534950 | C | T | 0.818718 | 3.07E-09 | 0.00120507 |
| 11 | 69534951 | A | G | 0.818718 | 3.07E-09 | 0.00120507 |
| 11 | 64542239 | T | A | 0.91231 | 3.26E-09 | 0.00127704 |
| 11 | 64542247 | C | A | 0.91231 | 3.26E-09 | 0.00127704 |
| 11 | 64544041 | A | G | 0.91231 | 3.26E-09 | 0.00127704 |
| 11 | 64544215 | T | C | 0.91231 | 3.26E-09 | 0.00127704 |
| 11 | 64544286 | G | A | 0.91231 | 3.26E-09 | 0.00127704 |
| 11 | 64544337 | T | C | 0.91231 | 3.26E-09 | 0.00127704 |
| 11 | 71469553 | G | T | 0.93339 | 4.28E-09 | 0.00167623 |
| 11 | 71497720 | G | C | 0.932546 | 4.77E-09 | 0.00186853 |
| 11 | 71497962 | T | C | 0.932546 | 4.77E-09 | 0.00186853 |
| 11 | 71498052 | T | C | 0.932546 | 4.77E-09 | 0.00186853 |
| 11 | 66075599 | G | A | 0.756324 | 5.37E-09 | 0.00210459 |
| 11 | 71591549 | G | C | 0.93339 | 6.55E-09 | 0.002569 |
| 11 | 66056082 | G | A | 0.853288 | 6.86E-09 | 0.00268905 |
| 11 | 66056112 | G | A | 0.853288 | 6.86E-09 | 0.00268905 |

| | | | | | | |
|---|---|---|---|---|---|---|
| 11 | 66056322 | T | C | 0.853288 | 6.86E-09 | 0.00268905 |
| 11 | 66056933 | C | A | 0.853288 | 6.86E-09 | 0.00268905 |
| 11 | 66057065 | A | T | 0.853288 | 6.86E-09 | 0.00268905 |
| 11 | 66057104 | G | A | 0.853288 | 6.86E-09 | 0.00268905 |
| 11 | 66057214 | C | A | 0.853288 | 6.86E-09 | 0.00268905 |
| 11 | 66057271 | A | C | 0.853288 | 6.86E-09 | 0.00268905 |
| 11 | 66057278 | C | A | 0.853288 | 6.86E-09 | 0.00268905 |
| 11 | 66058770 | C | T | 0.853288 | 6.86E-09 | 0.00268905 |
| 11 | 66058875 | T | C | 0.853288 | 6.86E-09 | 0.00268905 |
| 11 | 66058878 | G | C | 0.853288 | 6.86E-09 | 0.00268905 |
| 11 | 66059012 | T | A | 0.853288 | 6.86E-09 | 0.00268905 |
| 11 | 66059392 | G | C | 0.853288 | 6.86E-09 | 0.00268905 |
| 11 | 66059420 | T | C | 0.853288 | 6.86E-09 | 0.00268905 |
| 11 | 66060220 | G | A | 0.853288 | 6.86E-09 | 0.00268905 |
| 11 | 66060651 | T | C | 0.853288 | 6.86E-09 | 0.00268905 |
| 11 | 66060759 | T | C | 0.853288 | 6.86E-09 | 0.00268905 |
| 11 | 66065532 | T | G | 0.853288 | 6.86E-09 | 0.00268905 |
| 11 | 66067571 | C | A | 0.853288 | 6.86E-09 | 0.00268905 |
| 11 | 64243778 | A | G | 0.736931 | 7.82E-09 | 0.00306561 |
| 11 | 63921583 | A | G | 0.915683 | 8.17E-09 | 0.00320393 |
| 11 | 64210691 | T | G | 0.731029 | 8.36E-09 | 0.00327672 |
| 11 | 64110680 | A | G | 0.696459 | 8.75E-09 | 0.00343035 |
| 11 | 64544907 | G | A | 0.886172 | 8.87E-09 | 0.00347599 |
| 11 | 64544978 | T | C | 0.886172 | 8.87E-09 | 0.00347599 |
| 11 | 69543238 | G | A | 0.794266 | 8.99E-09 | 0.00352228 |
| 11 | 69543249 | A | G | 0.794266 | 8.99E-09 | 0.00352228 |
| 11 | 72007590 | A | G | 0.918212 | 9.25E-09 | 0.00362733 |
| 11 | 72007743 | T | C | 0.918212 | 9.25E-09 | 0.00362733 |
| 11 | 72007771 | A | C | 0.918212 | 9.25E-09 | 0.00362733 |
| 11 | 72007784 | T | C | 0.918212 | 9.25E-09 | 0.00362733 |
| 11 | 72007829 | G | T | 0.918212 | 9.25E-09 | 0.00362733 |
| 11 | 72007935 | T | C | 0.918212 | 9.25E-09 | 0.00362733 |
| 11 | 72008001 | C | T | 0.918212 | 9.25E-09 | 0.00362733 |
| 11 | 63912907 | T | C | 0.919056 | 9.41E-09 | 0.00368671 |
| 11 | 64969503 | C | T | 0.692243 | 9.96E-09 | 0.00390255 |
| 11 | 63390759 | T | C | 0.690556 | 1.03E-08 | 0.00405288 |
| 11 | 64243625 | A | G | 0.738617 | 1.03E-08 | 0.00405566 |
| 11 | 65156459 | C | T | 0.712479 | 1.06E-08 | 0.00416741 |
| 11 | 64783551 | C | T | 0.637437 | 1.07E-08 | 0.00418536 |
| 11 | 69893944 | C | T | 0.802698 | 1.09E-08 | 0.00429108 |
| 11 | 69894824 | C | T | 0.802698 | 1.09E-08 | 0.00429108 |
| 11 | 63913148 | G | T | 0.923272 | 1.13E-08 | 0.00442827 |
| 11 | 64544647 | T | C | 0.893761 | 1.20E-08 | 0.00469983 |
| 11 | 64258131 | T | C | 0.741147 | 1.24E-08 | 0.00485407 |
| 11 | 64903160 | T | C | 0.600337 | 1.28E-08 | 0.00500048 |
| 11 | 64917444 | T | C | 0.600337 | 1.28E-08 | 0.00500048 |
| 11 | 64918603 | T | C | 0.600337 | 1.28E-08 | 0.00500048 |
| 11 | 64920698 | A | G | 0.600337 | 1.28E-08 | 0.00500048 |
| 11 | 64955326 | C | A | 0.692243 | 1.39E-08 | 0.0054634 |
| 11 | 64955905 | T | C | 0.692243 | 1.39E-08 | 0.0054634 |
| 11 | 63804761 | A | T | 0.870152 | 1.40E-08 | 0.005489 |
| 11 | 64250963 | G | A | 0.738617 | 1.43E-08 | 0.00561506 |
| 11 | 64111071 | G | A | 0.694772 | 1.45E-08 | 0.00569431 |
| 11 | 64111109 | C | T | 0.694772 | 1.45E-08 | 0.00569431 |
| 11 | 72008274 | A | G | 0.917369 | 1.46E-08 | 0.00573014 |
| 11 | 72008379 | T | C | 0.917369 | 1.46E-08 | 0.00573014 |
| 11 | 72008404 | T | A | 0.917369 | 1.46E-08 | 0.00573014 |
| 11 | 72008537 | A | T | 0.917369 | 1.46E-08 | 0.00573014 |
| 11 | 72008967 | A | C | 0.917369 | 1.46E-08 | 0.00573014 |
| 11 | 72009797 | C | T | 0.917369 | 1.46E-08 | 0.00573014 |
| 11 | 72010423 | A | T | 0.917369 | 1.46E-08 | 0.00573014 |
| 11 | 72010851 | A | G | 0.917369 | 1.46E-08 | 0.00573014 |
| 11 | 72011305 | G | C | 0.917369 | 1.46E-08 | 0.00573014 |
| 11 | 72011313 | C | T | 0.917369 | 1.46E-08 | 0.00573014 |
| 11 | 72011317 | G | A | 0.917369 | 1.46E-08 | 0.00573014 |
| 11 | 72011319 | T | C | 0.917369 | 1.46E-08 | 0.00573014 |
| 11 | 72011320 | G | A | 0.917369 | 1.46E-08 | 0.00573014 |

| | | | | | | |
|---|---|---|---|---|---|---|
| 11 | 72011373 | T | C | 0.917369 | 1.46E-08 | 0.00573014 |
| 11 | 72011386 | T | G | 0.917369 | 1.46E-08 | 0.00573014 |
| 11 | 72011393 | A | G | 0.917369 | 1.46E-08 | 0.00573014 |
| 11 | 72011994 | T | C | 0.917369 | 1.46E-08 | 0.00573014 |
| 11 | 72012076 | A | G | 0.917369 | 1.46E-08 | 0.00573014 |
| 11 | 72012695 | G | A | 0.917369 | 1.46E-08 | 0.00573014 |
| 11 | 72012715 | T | A | 0.917369 | 1.46E-08 | 0.00573014 |
| 11 | 72012971 | T | A | 0.916526 | 1.52E-08 | 0.00596391 |
| 11 | 72013023 | G | T | 0.916526 | 1.52E-08 | 0.00596391 |
| 11 | 64951199 | C | T | 0.6914 | 1.60E-08 | 0.00626997 |
| 11 | 64951305 | G | A | 0.6914 | 1.60E-08 | 0.00626997 |
| 11 | 64972138 | C | A | 0.6914 | 1.60E-08 | 0.00626997 |
| 11 | 64972270 | A | C | 0.6914 | 1.60E-08 | 0.00626997 |
| 11 | 65112413 | G | A | 0.713322 | 1.60E-08 | 0.00627346 |
| 11 | 69871034 | T | A | 0.801012 | 1.61E-08 | 0.00633002 |
| 11 | 69871487 | T | C | 0.801012 | 1.61E-08 | 0.00633002 |
| 11 | 69871969 | G | C | 0.801012 | 1.61E-08 | 0.00633002 |
| 11 | 69872329 | C | T | 0.801012 | 1.61E-08 | 0.00633002 |
| 11 | 69873205 | C | G | 0.801012 | 1.61E-08 | 0.00633002 |
| 11 | 69874026 | T | C | 0.801012 | 1.61E-08 | 0.00633002 |
| 11 | 69874251 | C | T | 0.801012 | 1.61E-08 | 0.00633002 |
| 11 | 69875968 | T | C | 0.801012 | 1.61E-08 | 0.00633002 |
| 11 | 69876035 | C | T | 0.801012 | 1.61E-08 | 0.00633002 |
| 11 | 69876038 | C | T | 0.801012 | 1.61E-08 | 0.00633002 |
| 11 | 69876044 | C | T | 0.801012 | 1.61E-08 | 0.00633002 |
| 11 | 69880963 | T | C | 0.801012 | 1.61E-08 | 0.00633002 |
| 11 | 69884488 | C | G | 0.801012 | 1.61E-08 | 0.00633002 |
| 11 | 69884635 | G | A | 0.801012 | 1.61E-08 | 0.00633002 |
| 11 | 69889274 | G | A | 0.801012 | 1.61E-08 | 0.00633002 |
| 11 | 69889339 | G | T | 0.801012 | 1.61E-08 | 0.00633002 |
| 11 | 69889364 | C | A | 0.801012 | 1.61E-08 | 0.00633002 |
| 11 | 69889472 | T | A | 0.801012 | 1.61E-08 | 0.00633002 |
| 11 | 69890837 | C | T | 0.801012 | 1.61E-08 | 0.00633002 |
| 11 | 69892121 | G | A | 0.801012 | 1.61E-08 | 0.00633002 |
| 11 | 69892414 | C | G | 0.801012 | 1.61E-08 | 0.00633002 |
| 11 | 69892426 | G | T | 0.801012 | 1.61E-08 | 0.00633002 |
| 11 | 69892860 | A | T | 0.801012 | 1.61E-08 | 0.00633002 |
| 11 | 69892883 | G | C | 0.801012 | 1.61E-08 | 0.00633002 |
| 11 | 69885503 | T | C | 0.801855 | 1.63E-08 | 0.00638176 |
| 11 | 69885690 | C | T | 0.801855 | 1.63E-08 | 0.00638176 |
| 11 | 69885723 | C | T | 0.801855 | 1.63E-08 | 0.00638176 |
| 11 | 69885996 | A | G | 0.801855 | 1.63E-08 | 0.00638176 |
| 11 | 69891105 | C | T | 0.801855 | 1.63E-08 | 0.00638176 |
| 11 | 64219108 | G | A | 0.740304 | 1.63E-08 | 0.00639301 |
| 11 | 64057310 | G | A | 0.694772 | 1.64E-08 | 0.00643652 |
| 11 | 64057327 | C | T | 0.694772 | 1.64E-08 | 0.00643652 |
| 11 | 64057358 | C | T | 0.694772 | 1.64E-08 | 0.00643652 |
| 11 | 64057409 | A | G | 0.694772 | 1.64E-08 | 0.00643652 |
| 11 | 64057502 | G | T | 0.694772 | 1.64E-08 | 0.00643652 |
| 11 | 64057533 | A | G | 0.694772 | 1.64E-08 | 0.00643652 |
| 11 | 64057819 | T | G | 0.694772 | 1.64E-08 | 0.00643652 |
| 11 | 64057835 | G | A | 0.694772 | 1.64E-08 | 0.00643652 |
| 11 | 64057865 | G | A | 0.694772 | 1.64E-08 | 0.00643652 |
| 11 | 64057918 | G | A | 0.694772 | 1.64E-08 | 0.00643652 |
| 11 | 64058519 | T | G | 0.694772 | 1.64E-08 | 0.00643652 |
| 11 | 64058579 | G | A | 0.694772 | 1.64E-08 | 0.00643652 |
| 11 | 64061476 | C | G | 0.694772 | 1.64E-08 | 0.00643652 |
| 11 | 64061486 | C | T | 0.694772 | 1.64E-08 | 0.00643652 |
| 11 | 64071092 | T | C | 0.694772 | 1.64E-08 | 0.00643652 |
| 11 | 64071101 | T | A | 0.694772 | 1.64E-08 | 0.00643652 |
| 11 | 64071108 | G | A | 0.694772 | 1.64E-08 | 0.00643652 |
| 11 | 64071119 | G | A | 0.694772 | 1.64E-08 | 0.00643652 |
| 11 | 64071368 | A | G | 0.694772 | 1.64E-08 | 0.00643652 |
| 11 | 64074168 | T | C | 0.694772 | 1.64E-08 | 0.00643652 |
| 11 | 64074242 | C | G | 0.694772 | 1.64E-08 | 0.00643652 |
| 11 | 64074268 | A | G | 0.694772 | 1.64E-08 | 0.00643652 |
| 11 | 64074277 | A | G | 0.694772 | 1.64E-08 | 0.00643652 |

| | | | | | | |
|---|---|---|---|---|---|---|
| 11 | 64074378 | A | G | 0.694772 | 1.64E-08 | 0.00643652 |
| 11 | 64074385 | T | G | 0.694772 | 1.64E-08 | 0.00643652 |
| 11 | 64074477 | G | C | 0.694772 | 1.64E-08 | 0.00643652 |
| 11 | 64074817 | A | G | 0.694772 | 1.64E-08 | 0.00643652 |
| 11 | 64074839 | T | G | 0.694772 | 1.64E-08 | 0.00643652 |
| 11 | 64074975 | A | G | 0.694772 | 1.64E-08 | 0.00643652 |
| 11 | 64075035 | G | C | 0.694772 | 1.64E-08 | 0.00643652 |
| 11 | 64075047 | C | T | 0.694772 | 1.64E-08 | 0.00643652 |
| 11 | 64075058 | A | G | 0.694772 | 1.64E-08 | 0.00643652 |
| 11 | 64075064 | A | G | 0.694772 | 1.64E-08 | 0.00643652 |
| 11 | 64076216 | A | G | 0.694772 | 1.64E-08 | 0.00643652 |
| 11 | 64103156 | C | G | 0.694772 | 1.64E-08 | 0.00643652 |
| 11 | 64103233 | T | G | 0.694772 | 1.64E-08 | 0.00643652 |
| 11 | 64103612 | C | G | 0.694772 | 1.64E-08 | 0.00643652 |
| 11 | 64104679 | C | T | 0.694772 | 1.64E-08 | 0.00643652 |
| 11 | 64104807 | G | C | 0.694772 | 1.64E-08 | 0.00643652 |
| 11 | 64105107 | C | T | 0.694772 | 1.64E-08 | 0.00643652 |
| 11 | 64105221 | T | A | 0.694772 | 1.64E-08 | 0.00643652 |
| 11 | 64105222 | C | A | 0.694772 | 1.64E-08 | 0.00643652 |
| 11 | 64105268 | G | A | 0.694772 | 1.64E-08 | 0.00643652 |
| 11 | 64105355 | C | T | 0.694772 | 1.64E-08 | 0.00643652 |
| 11 | 64105941 | A | G | 0.694772 | 1.64E-08 | 0.00643652 |
| 11 | 64106047 | C | G | 0.694772 | 1.64E-08 | 0.00643652 |
| 11 | 64107388 | G | A | 0.694772 | 1.64E-08 | 0.00643652 |
| 11 | 64107436 | G | T | 0.694772 | 1.64E-08 | 0.00643652 |
| 11 | 64109102 | C | T | 0.694772 | 1.64E-08 | 0.00643652 |
| 11 | 64109134 | G | A | 0.694772 | 1.64E-08 | 0.00643652 |
| 11 | 64109296 | A | G | 0.694772 | 1.64E-08 | 0.00643652 |
| 11 | 64109364 | T | G | 0.694772 | 1.64E-08 | 0.00643652 |
| 11 | 64113492 | C | T | 0.694772 | 1.64E-08 | 0.00643652 |
| 11 | 64113504 | C | T | 0.694772 | 1.64E-08 | 0.00643652 |
| 11 | 64113757 | G | A | 0.694772 | 1.64E-08 | 0.00643652 |
| 11 | 64113975 | G | T | 0.694772 | 1.64E-08 | 0.00643652 |
| 11 | 64114009 | G | C | 0.694772 | 1.64E-08 | 0.00643652 |
| 11 | 64114044 | A | G | 0.694772 | 1.64E-08 | 0.00643652 |
| 11 | 64224165 | A | C | 0.741147 | 1.65E-08 | 0.00646372 |
| 11 | 64756045 | A | G | 0.639123 | 1.68E-08 | 0.00658931 |
| 11 | 64767180 | C | T | 0.639123 | 1.68E-08 | 0.00658931 |
| 11 | 64767184 | G | A | 0.639123 | 1.68E-08 | 0.00658931 |
| 11 | 64768176 | C | T | 0.639123 | 1.68E-08 | 0.00658931 |
| 11 | 64768295 | G | A | 0.639123 | 1.68E-08 | 0.00658931 |
| 11 | 64768435 | A | G | 0.639123 | 1.68E-08 | 0.00658931 |
| 11 | 64768656 | G | A | 0.639123 | 1.68E-08 | 0.00658931 |
| 11 | 64771708 | G | A | 0.639123 | 1.68E-08 | 0.00658931 |
| 11 | 64780069 | T | G | 0.639123 | 1.68E-08 | 0.00658931 |
| 11 | 64780124 | A | G | 0.639123 | 1.68E-08 | 0.00658931 |
| 11 | 64780382 | G | T | 0.639123 | 1.68E-08 | 0.00658931 |
| 11 | 64780459 | T | C | 0.639123 | 1.68E-08 | 0.00658931 |
| 11 | 64780523 | T | C | 0.639123 | 1.68E-08 | 0.00658931 |
| 11 | 64780535 | A | C | 0.639123 | 1.68E-08 | 0.00658931 |
| 11 | 64781001 | C | T | 0.639123 | 1.68E-08 | 0.00658931 |
| 11 | 64781578 | T | A | 0.639123 | 1.68E-08 | 0.00658931 |
| 11 | 64782630 | A | G | 0.639123 | 1.68E-08 | 0.00658931 |
| 11 | 64782954 | C | T | 0.639123 | 1.68E-08 | 0.00658931 |
| 11 | 64783030 | T | A | 0.639123 | 1.68E-08 | 0.00658931 |
| 11 | 64789290 | T | C | 0.639123 | 1.68E-08 | 0.00658931 |
| 11 | 64792293 | C | G | 0.639123 | 1.68E-08 | 0.00658931 |
| 11 | 66058063 | A | T | 0.852445 | 1.69E-08 | 0.00661346 |
| 11 | 66058368 | A | C | 0.852445 | 1.69E-08 | 0.00661346 |
| 11 | 66058598 | G | T | 0.852445 | 1.69E-08 | 0.00661346 |
| 11 | 66058663 | T | C | 0.852445 | 1.69E-08 | 0.00661346 |
| 11 | 66058711 | G | C | 0.852445 | 1.69E-08 | 0.00661346 |
| 11 | 64111400 | G | C | 0.696459 | 1.72E-08 | 0.00673415 |
| 11 | 64111401 | A | C | 0.696459 | 1.72E-08 | 0.00673415 |
| 11 | 64111449 | G | A | 0.696459 | 1.72E-08 | 0.00673415 |
| 11 | 64111632 | G | A | 0.696459 | 1.72E-08 | 0.00673415 |
| 11 | 64111648 | T | C | 0.696459 | 1.72E-08 | 0.00673415 |

| | | | | | | |
|---|---|---|---|---|---|---|
| 11 | 64111658 | G | C | 0.696459 | 1.72E-08 | 0.00673415 |
| 11 | 64111660 | G | C | 0.696459 | 1.72E-08 | 0.00673415 |
| 11 | 64111701 | T | C | 0.696459 | 1.72E-08 | 0.00673415 |
| 11 | 64111712 | T | A | 0.696459 | 1.72E-08 | 0.00673415 |
| 11 | 64111716 | C | T | 0.696459 | 1.72E-08 | 0.00673415 |
| 11 | 64112497 | T | C | 0.696459 | 1.72E-08 | 0.00673415 |
| 11 | 65090186 | G | A | 0.714165 | 1.80E-08 | 0.00707093 |
| 11 | 65090374 | A | T | 0.714165 | 1.80E-08 | 0.00707093 |
| 11 | 65090640 | T | C | 0.714165 | 1.80E-08 | 0.00707093 |
| 11 | 65090999 | C | A | 0.714165 | 1.80E-08 | 0.00707093 |
| 11 | 65091705 | A | G | 0.714165 | 1.80E-08 | 0.00707093 |
| 11 | 65092167 | G | A | 0.714165 | 1.80E-08 | 0.00707093 |
| 11 | 65092500 | A | G | 0.714165 | 1.80E-08 | 0.00707093 |
| 11 | 65092506 | A | G | 0.714165 | 1.80E-08 | 0.00707093 |
| 11 | 65092956 | A | G | 0.714165 | 1.80E-08 | 0.00707093 |
| 11 | 65093168 | T | C | 0.714165 | 1.80E-08 | 0.00707093 |
| 11 | 65093386 | C | T | 0.714165 | 1.80E-08 | 0.00707093 |
| 11 | 65093578 | A | C | 0.714165 | 1.80E-08 | 0.00707093 |
| 11 | 65094034 | C | T | 0.714165 | 1.80E-08 | 0.00707093 |
| 11 | 65095116 | T | C | 0.714165 | 1.80E-08 | 0.00707093 |
| 11 | 65095704 | C | T | 0.714165 | 1.80E-08 | 0.00707093 |
| 11 | 65096145 | T | G | 0.714165 | 1.80E-08 | 0.00707093 |
| 11 | 65098060 | G | A | 0.714165 | 1.80E-08 | 0.00707093 |
| 11 | 65098479 | A | C | 0.714165 | 1.80E-08 | 0.00707093 |
| 11 | 65099297 | G | A | 0.714165 | 1.80E-08 | 0.00707093 |
| 11 | 65099872 | G | A | 0.714165 | 1.80E-08 | 0.00707093 |
| 11 | 65101642 | A | G | 0.714165 | 1.80E-08 | 0.00707093 |
| 11 | 65101701 | G | A | 0.714165 | 1.80E-08 | 0.00707093 |
| 11 | 65101840 | C | A | 0.714165 | 1.80E-08 | 0.00707093 |
| 11 | 65101995 | A | G | 0.714165 | 1.80E-08 | 0.00707093 |
| 11 | 65102098 | C | T | 0.714165 | 1.80E-08 | 0.00707093 |
| 11 | 65102239 | C | T | 0.714165 | 1.80E-08 | 0.00707093 |
| 11 | 65103039 | A | G | 0.714165 | 1.80E-08 | 0.00707093 |
| 11 | 65103061 | G | C | 0.714165 | 1.80E-08 | 0.00707093 |
| 11 | 65103181 | T | G | 0.714165 | 1.80E-08 | 0.00707093 |
| 11 | 65103367 | C | A | 0.714165 | 1.80E-08 | 0.00707093 |
| 11 | 65103636 | C | T | 0.714165 | 1.80E-08 | 0.00707093 |
| 11 | 65103990 | T | C | 0.714165 | 1.80E-08 | 0.00707093 |
| 11 | 65105116 | A | T | 0.714165 | 1.80E-08 | 0.00707093 |
| 11 | 65105211 | C | T | 0.714165 | 1.80E-08 | 0.00707093 |
| 11 | 65106021 | T | C | 0.714165 | 1.80E-08 | 0.00707093 |
| 11 | 65106600 | A | G | 0.714165 | 1.80E-08 | 0.00707093 |
| 11 | 65108009 | T | G | 0.714165 | 1.80E-08 | 0.00707093 |
| 11 | 65109333 | G | A | 0.714165 | 1.80E-08 | 0.00707093 |
| 11 | 65109440 | T | C | 0.714165 | 1.80E-08 | 0.00707093 |
| 11 | 65110174 | C | T | 0.714165 | 1.80E-08 | 0.00707093 |
| 11 | 65110323 | C | G | 0.714165 | 1.80E-08 | 0.00707093 |
| 11 | 65110408 | C | T | 0.714165 | 1.80E-08 | 0.00707093 |
| 11 | 65110628 | A | G | 0.714165 | 1.80E-08 | 0.00707093 |
| 11 | 65111406 | T | A | 0.714165 | 1.80E-08 | 0.00707093 |
| 11 | 65111620 | G | T | 0.714165 | 1.80E-08 | 0.00707093 |
| 11 | 65111701 | G | T | 0.714165 | 1.80E-08 | 0.00707093 |
| 11 | 65111798 | A | G | 0.714165 | 1.80E-08 | 0.00707093 |
| 11 | 65111842 | A | G | 0.714165 | 1.80E-08 | 0.00707093 |
| 11 | 65111886 | A | G | 0.714165 | 1.80E-08 | 0.00707093 |
| 11 | 65112094 | T | C | 0.714165 | 1.80E-08 | 0.00707093 |
| 11 | 65112300 | C | T | 0.714165 | 1.80E-08 | 0.00707093 |
| 11 | 65112302 | C | T | 0.714165 | 1.80E-08 | 0.00707093 |
| 11 | 65112503 | T | C | 0.714165 | 1.80E-08 | 0.00707093 |
| 11 | 65113230 | C | T | 0.714165 | 1.80E-08 | 0.00707093 |
| 11 | 65114643 | T | C | 0.714165 | 1.80E-08 | 0.00707093 |
| 11 | 65119236 | A | C | 0.714165 | 1.80E-08 | 0.00707093 |
| 11 | 65119321 | T | C | 0.714165 | 1.80E-08 | 0.00707093 |
| 11 | 65119374 | A | G | 0.714165 | 1.80E-08 | 0.00707093 |
| 11 | 65121828 | A | G | 0.714165 | 1.80E-08 | 0.00707093 |
| 11 | 65124391 | A | G | 0.714165 | 1.80E-08 | 0.00707093 |
| 11 | 65125902 | A | G | 0.714165 | 1.80E-08 | 0.00707093 |

| | | | | | | |
|---|---|---|---|---|---|---|
| 11 | 65126707 | C | T | 0.714165 | 1.80E-08 | 0.00707093 |
| 11 | 65128025 | A | G | 0.714165 | 1.80E-08 | 0.00707093 |
| 11 | 65128721 | A | G | 0.714165 | 1.80E-08 | 0.00707093 |
| 11 | 65128872 | A | C | 0.714165 | 1.80E-08 | 0.00707093 |
| 11 | 65129737 | A | C | 0.714165 | 1.80E-08 | 0.00707093 |
| 11 | 65132942 | A | G | 0.714165 | 1.80E-08 | 0.00707093 |
| 11 | 65134720 | G | A | 0.714165 | 1.80E-08 | 0.00707093 |
| 11 | 65136159 | C | T | 0.714165 | 1.80E-08 | 0.00707093 |
| 11 | 65136202 | G | C | 0.714165 | 1.80E-08 | 0.00707093 |
| 11 | 65136237 | A | G | 0.714165 | 1.80E-08 | 0.00707093 |
| 11 | 65136553 | A | G | 0.714165 | 1.80E-08 | 0.00707093 |
| 11 | 65137374 | T | C | 0.714165 | 1.80E-08 | 0.00707093 |
| 11 | 65138197 | T | C | 0.714165 | 1.80E-08 | 0.00707093 |
| 11 | 65138504 | G | C | 0.714165 | 1.80E-08 | 0.00707093 |
| 11 | 65138576 | C | T | 0.714165 | 1.80E-08 | 0.00707093 |
| 11 | 65138592 | C | T | 0.714165 | 1.80E-08 | 0.00707093 |
| 11 | 65138751 | T | C | 0.714165 | 1.80E-08 | 0.00707093 |
| 11 | 65145971 | G | A | 0.714165 | 1.80E-08 | 0.00707093 |
| 11 | 65146813 | A | G | 0.714165 | 1.80E-08 | 0.00707093 |
| 11 | 64110631 | A | C | 0.753794 | 1.95E-08 | 0.00765584 |
| 11 | 64110643 | T | C | 0.753794 | 1.95E-08 | 0.00765584 |
| 11 | 64110670 | A | G | 0.753794 | 1.95E-08 | 0.00765584 |
| 11 | 64110697 | A | G | 0.753794 | 1.95E-08 | 0.00765584 |
| 11 | 64110702 | C | T | 0.753794 | 1.95E-08 | 0.00765584 |
| 11 | 64110715 | T | C | 0.753794 | 1.95E-08 | 0.00765584 |
| 11 | 64110878 | C | T | 0.753794 | 1.95E-08 | 0.00765584 |
| 11 | 64110909 | T | C | 0.753794 | 1.95E-08 | 0.00765584 |
| 11 | 64111133 | T | G | 0.753794 | 1.95E-08 | 0.00765584 |
| 11 | 64111145 | G | A | 0.753794 | 1.95E-08 | 0.00765584 |
| 11 | 64207395 | C | G | 0.676223 | 2.01E-08 | 0.00787821 |
| 11 | 64920385 | G | A | 0.81113 | 2.12E-08 | 0.0083017 |
| 11 | 64920457 | T | G | 0.81113 | 2.12E-08 | 0.0083017 |
| 11 | 64920540 | G | A | 0.81113 | 2.12E-08 | 0.0083017 |
| 11 | 64920772 | T | C | 0.81113 | 2.12E-08 | 0.0083017 |
| 11 | 64920847 | G | C | 0.81113 | 2.12E-08 | 0.0083017 |
| 11 | 64933278 | C | T | 0.81113 | 2.12E-08 | 0.0083017 |
| 11 | 64242613 | C | A | 0.73946 | 2.12E-08 | 0.00830711 |
| 11 | 64242823 | T | C | 0.73946 | 2.12E-08 | 0.00830711 |
| 11 | 64242850 | T | C | 0.73946 | 2.12E-08 | 0.00830711 |
| 11 | 64243429 | T | C | 0.73946 | 2.12E-08 | 0.00830711 |
| 11 | 64204258 | T | C | 0.674536 | 2.16E-08 | 0.00845743 |
| 11 | 64204283 | T | G | 0.674536 | 2.16E-08 | 0.00845743 |
| 11 | 64204286 | C | T | 0.674536 | 2.16E-08 | 0.00845743 |
| 11 | 64204290 | T | G | 0.674536 | 2.16E-08 | 0.00845743 |
| 11 | 64220013 | G | A | 0.73946 | 2.22E-08 | 0.00871308 |
| 11 | 64220448 | A | C | 0.73946 | 2.22E-08 | 0.00871308 |
| 11 | 64221075 | C | T | 0.73946 | 2.22E-08 | 0.00871308 |
| 11 | 64226080 | A | G | 0.73946 | 2.22E-08 | 0.00871308 |
| 11 | 64228567 | C | T | 0.73946 | 2.22E-08 | 0.00871308 |
| 11 | 64229233 | C | G | 0.73946 | 2.22E-08 | 0.00871308 |
| 11 | 64229234 | A | G | 0.73946 | 2.22E-08 | 0.00871308 |
| 11 | 64229313 | A | G | 0.73946 | 2.22E-08 | 0.00871308 |
| 11 | 64233922 | A | C | 0.73946 | 2.22E-08 | 0.00871308 |
| 11 | 64234179 | A | G | 0.73946 | 2.22E-08 | 0.00871308 |
| 11 | 64234212 | A | G | 0.73946 | 2.22E-08 | 0.00871308 |
| 11 | 64234215 | T | C | 0.73946 | 2.22E-08 | 0.00871308 |
| 11 | 64234219 | G | A | 0.73946 | 2.22E-08 | 0.00871308 |
| 11 | 64234400 | A | T | 0.73946 | 2.22E-08 | 0.00871308 |
| 11 | 64234427 | A | G | 0.73946 | 2.22E-08 | 0.00871308 |
| 11 | 64234622 | G | A | 0.73946 | 2.22E-08 | 0.00871308 |
| 11 | 64236154 | A | T | 0.73946 | 2.22E-08 | 0.00871308 |
| 11 | 64236242 | G | A | 0.73946 | 2.22E-08 | 0.00871308 |
| 11 | 64236293 | A | C | 0.73946 | 2.22E-08 | 0.00871308 |
| 11 | 64236314 | A | T | 0.73946 | 2.22E-08 | 0.00871308 |
| 11 | 64236348 | G | A | 0.73946 | 2.22E-08 | 0.00871308 |
| 11 | 64236355 | T | G | 0.73946 | 2.22E-08 | 0.00871308 |
| 11 | 64236514 | A | G | 0.73946 | 2.22E-08 | 0.00871308 |

| | | | | | | |
|---|---|---|---|---|---|---|
| 11 | 64236590 | G | A | 0.73946 | 2.22E-08 | 0.00871308 |
| 11 | 64236878 | C | A | 0.73946 | 2.22E-08 | 0.00871308 |
| 11 | 64236937 | A | G | 0.73946 | 2.22E-08 | 0.00871308 |
| 11 | 64236955 | T | C | 0.73946 | 2.22E-08 | 0.00871308 |
| 11 | 64237034 | A | T | 0.73946 | 2.22E-08 | 0.00871308 |
| 11 | 64237062 | G | A | 0.73946 | 2.22E-08 | 0.00871308 |
| 11 | 64237078 | T | G | 0.73946 | 2.22E-08 | 0.00871308 |
| 11 | 64237116 | T | C | 0.73946 | 2.22E-08 | 0.00871308 |
| 11 | 64237184 | A | G | 0.73946 | 2.22E-08 | 0.00871308 |
| 11 | 64237223 | G | C | 0.73946 | 2.22E-08 | 0.00871308 |
| 11 | 64237317 | T | C | 0.73946 | 2.22E-08 | 0.00871308 |
| 11 | 64237331 | A | G | 0.73946 | 2.22E-08 | 0.00871308 |
| 11 | 64237383 | G | A | 0.73946 | 2.22E-08 | 0.00871308 |
| 11 | 64237404 | C | G | 0.73946 | 2.22E-08 | 0.00871308 |
| 11 | 64237421 | G | T | 0.73946 | 2.22E-08 | 0.00871308 |
| 11 | 64237446 | G | A | 0.73946 | 2.22E-08 | 0.00871308 |
| 11 | 64237461 | T | C | 0.73946 | 2.22E-08 | 0.00871308 |
| 11 | 64237657 | C | T | 0.73946 | 2.22E-08 | 0.00871308 |
| 11 | 64238057 | A | G | 0.73946 | 2.22E-08 | 0.00871308 |
| 11 | 64238457 | T | C | 0.73946 | 2.22E-08 | 0.00871308 |
| 11 | 64238763 | T | C | 0.73946 | 2.22E-08 | 0.00871308 |
| 11 | 64239069 | C | A | 0.73946 | 2.22E-08 | 0.00871308 |
| 11 | 64252696 | C | T | 0.73946 | 2.22E-08 | 0.00871308 |
| 11 | 64252970 | T | C | 0.73946 | 2.22E-08 | 0.00871308 |
| 11 | 64253197 | C | G | 0.73946 | 2.22E-08 | 0.00871308 |
| 11 | 64253666 | T | C | 0.73946 | 2.22E-08 | 0.00871308 |
| 11 | 64253770 | C | A | 0.73946 | 2.22E-08 | 0.00871308 |
| 11 | 64253909 | C | G | 0.73946 | 2.22E-08 | 0.00871308 |
| 11 | 64254337 | C | T | 0.73946 | 2.22E-08 | 0.00871308 |
| 11 | 64254850 | A | G | 0.73946 | 2.22E-08 | 0.00871308 |
| 11 | 64260664 | A | C | 0.73946 | 2.22E-08 | 0.00871308 |
| 11 | 64260883 | T | C | 0.73946 | 2.22E-08 | 0.00871308 |
| 11 | 64260930 | T | C | 0.73946 | 2.22E-08 | 0.00871308 |
| 11 | 64261265 | G | A | 0.73946 | 2.22E-08 | 0.00871308 |
| 11 | 64261600 | T | C | 0.73946 | 2.22E-08 | 0.00871308 |
| 11 | 64262009 | T | G | 0.73946 | 2.22E-08 | 0.00871308 |
| 11 | 64262068 | G | A | 0.73946 | 2.22E-08 | 0.00871308 |
| 11 | 64263303 | T | G | 0.73946 | 2.22E-08 | 0.00871308 |
| 11 | 64264077 | A | G | 0.73946 | 2.22E-08 | 0.00871308 |
| 11 | 64264556 | T | C | 0.73946 | 2.22E-08 | 0.00871308 |
| 11 | 64264628 | T | C | 0.73946 | 2.22E-08 | 0.00871308 |
| 11 | 64264715 | A | G | 0.73946 | 2.22E-08 | 0.00871308 |
| 11 | 64264729 | G | T | 0.73946 | 2.22E-08 | 0.00871308 |
| 11 | 64264922 | T | G | 0.73946 | 2.22E-08 | 0.00871308 |
| 11 | 64265145 | T | C | 0.73946 | 2.22E-08 | 0.00871308 |
| 11 | 64265233 | C | T | 0.73946 | 2.22E-08 | 0.00871308 |
| 11 | 64265326 | A | G | 0.73946 | 2.22E-08 | 0.00871308 |
| 11 | 64265327 | C | G | 0.73946 | 2.22E-08 | 0.00871308 |
| 11 | 64265440 | A | G | 0.73946 | 2.22E-08 | 0.00871308 |
| 11 | 64265816 | T | C | 0.73946 | 2.22E-08 | 0.00871308 |
| 11 | 64265973 | A | G | 0.73946 | 2.22E-08 | 0.00871308 |
| 11 | 64266216 | A | C | 0.73946 | 2.22E-08 | 0.00871308 |
| 11 | 64266443 | T | G | 0.73946 | 2.22E-08 | 0.00871308 |
| 11 | 63921552 | T | C | 0.925801 | 2.25E-08 | 0.00882444 |
| 11 | 63921561 | T | C | 0.925801 | 2.25E-08 | 0.00882444 |
| 11 | 63921565 | C | A | 0.925801 | 2.25E-08 | 0.00882444 |
| 11 | 64600319 | G | T | 0.681282 | 2.27E-08 | 0.00890769 |
| 11 | 64527644 | A | G | 0.598651 | 2.34E-08 | 0.00918098 |
| 11 | 64219176 | C | T | 0.73946 | 2.35E-08 | 0.00921837 |
| 11 | 64219632 | G | A | 0.73946 | 2.35E-08 | 0.00921837 |
| 11 | 64219776 | G | A | 0.73946 | 2.35E-08 | 0.00921837 |
| 11 | 64138690 | C | A | 0.692243 | 2.44E-08 | 0.0095481 |
| 11 | 64138772 | A | C | 0.692243 | 2.44E-08 | 0.0095481 |
| 11 | 64138866 | T | C | 0.692243 | 2.44E-08 | 0.0095481 |
| 11 | 64139290 | C | T | 0.692243 | 2.44E-08 | 0.0095481 |
| 11 | 65112571 | T | C | 0.713322 | 2.45E-08 | 0.00958562 |
| 11 | 65112637 | C | T | 0.713322 | 2.45E-08 | 0.00958562 |

| | | | | | | |
|---|---|---|---|---|---|---|
| 11 | 65112675 | C | T | 0.713322 | 2.45E-08 | 0.00958562 |
| 11 | 65115334 | G | A | 0.713322 | 2.46E-08 | 0.00963183 |
| 11 | 65116361 | C | T | 0.713322 | 2.46E-08 | 0.00963183 |
| 11 | 65116998 | A | G | 0.713322 | 2.46E-08 | 0.00963183 |
| 11 | 65117891 | G | A | 0.713322 | 2.46E-08 | 0.00963183 |
| 11 | 65118018 | T | G | 0.713322 | 2.46E-08 | 0.00963183 |
| 11 | 65120022 | G | T | 0.713322 | 2.46E-08 | 0.00963183 |
| 11 | 65120726 | A | G | 0.713322 | 2.46E-08 | 0.00963183 |
| 11 | 64996165 | T | C | 0.839798 | 2.47E-08 | 0.0096978 |
| 11 | 64171157 | C | T | 0.690556 | 2.48E-08 | 0.00973367 |
| 11 | 64108026 | G | A | 0.695616 | 2.67E-08 | 0.01046795 |
| 11 | 64108509 | T | C | 0.695616 | 2.67E-08 | 0.01046795 |
| 11 | 64155572 | G | T | 0.689713 | 2.68E-08 | 0.01051574 |
| 11 | 64921173 | T | C | 0.809444 | 2.74E-08 | 0.01073873 |
| 11 | 64696140 | G | A | 0.73946 | 2.77E-08 | 0.01084668 |
| 11 | 64696165 | C | T | 0.73946 | 2.77E-08 | 0.01084668 |
| 11 | 64783635 | A | G | 0.63575 | 2.80E-08 | 0.01097454 |
| 11 | 64111664 | G | C | 0.692243 | 2.84E-08 | 0.01114266 |
| 11 | 64111666 | G | C | 0.692243 | 2.84E-08 | 0.01114266 |
| 11 | 64262717 | A | G | 0.734401 | 2.86E-08 | 0.01120126 |
| 11 | 65136020 | G | A | 0.713322 | 2.88E-08 | 0.01129804 |
| 11 | 65136053 | A | C | 0.713322 | 2.88E-08 | 0.01129804 |
| 11 | 64057331 | A | G | 0.752951 | 2.94E-08 | 0.01152649 |
| 11 | 64057378 | C | T | 0.752951 | 2.94E-08 | 0.01152649 |
| 11 | 64057382 | A | G | 0.752951 | 2.94E-08 | 0.01152649 |
| 11 | 64057401 | C | T | 0.752951 | 2.94E-08 | 0.01152649 |
| 11 | 64057500 | C | T | 0.752951 | 2.94E-08 | 0.01152649 |
| 11 | 64057839 | G | A | 0.752951 | 2.94E-08 | 0.01152649 |
| 11 | 64105324 | A | G | 0.752951 | 2.94E-08 | 0.01152649 |
| 11 | 64105764 | T | C | 0.752951 | 2.94E-08 | 0.01152649 |
| 11 | 64105906 | G | C | 0.752951 | 2.94E-08 | 0.01152649 |
| 11 | 64105957 | A | G | 0.752951 | 2.94E-08 | 0.01152649 |
| 11 | 64105976 | C | G | 0.752951 | 2.94E-08 | 0.01152649 |
| 11 | 64106374 | T | C | 0.752951 | 2.94E-08 | 0.01152649 |
| 11 | 64106630 | A | G | 0.752951 | 2.94E-08 | 0.01152649 |
| 11 | 64107596 | A | G | 0.752951 | 2.94E-08 | 0.01152649 |
| 11 | 64107809 | C | T | 0.752951 | 2.94E-08 | 0.01152649 |
| 11 | 64107873 | G | A | 0.752951 | 2.94E-08 | 0.01152649 |
| 11 | 64107905 | C | T | 0.752951 | 2.94E-08 | 0.01152649 |
| 11 | 64108587 | C | T | 0.752951 | 2.94E-08 | 0.01152649 |
| 11 | 64108614 | G | A | 0.752951 | 2.94E-08 | 0.01152649 |
| 11 | 64108619 | G | A | 0.752951 | 2.94E-08 | 0.01152649 |
| 11 | 64108621 | T | C | 0.752951 | 2.94E-08 | 0.01152649 |
| 11 | 64108626 | T | C | 0.752951 | 2.94E-08 | 0.01152649 |
| 11 | 64108644 | A | T | 0.752951 | 2.94E-08 | 0.01152649 |
| 11 | 64108656 | C | A | 0.752951 | 2.94E-08 | 0.01152649 |
| 11 | 64108731 | A | G | 0.752951 | 2.94E-08 | 0.01152649 |
| 11 | 64108751 | T | C | 0.752951 | 2.94E-08 | 0.01152649 |
| 11 | 64108862 | G | A | 0.752951 | 2.94E-08 | 0.01152649 |
| 11 | 64108901 | C | A | 0.752951 | 2.94E-08 | 0.01152649 |
| 11 | 64108937 | C | T | 0.752951 | 2.94E-08 | 0.01152649 |
| 11 | 64108965 | A | T | 0.752951 | 2.94E-08 | 0.01152649 |
| 11 | 64109060 | C | A | 0.752951 | 2.94E-08 | 0.01152649 |
| 11 | 64109081 | T | C | 0.752951 | 2.94E-08 | 0.01152649 |
| 11 | 64109108 | A | G | 0.752951 | 2.94E-08 | 0.01152649 |
| 11 | 64109140 | C | G | 0.752951 | 2.94E-08 | 0.01152649 |
| 11 | 64109176 | A | C | 0.752951 | 2.94E-08 | 0.01152649 |
| 11 | 64109337 | G | A | 0.752951 | 2.94E-08 | 0.01152649 |
| 11 | 64109390 | T | A | 0.752951 | 2.94E-08 | 0.01152649 |
| 11 | 64111539 | G | A | 0.752951 | 2.94E-08 | 0.01152649 |
| 11 | 64111586 | T | C | 0.752951 | 2.94E-08 | 0.01152649 |
| 11 | 64111862 | T | C | 0.752951 | 2.94E-08 | 0.01152649 |
| 11 | 64112065 | G | T | 0.752951 | 2.94E-08 | 0.01152649 |
| 11 | 64112185 | A | C | 0.752951 | 2.94E-08 | 0.01152649 |
| 11 | 64112229 | T | C | 0.752951 | 2.94E-08 | 0.01152649 |
| 11 | 64112252 | A | G | 0.752951 | 2.94E-08 | 0.01152649 |
| 11 | 64112289 | A | G | 0.752951 | 2.94E-08 | 0.01152649 |

| | | | | | | |
|---|---|---|---|---|---|---|
| 11 | 64112330 | A | G | 0.752951 | 2.94E-08 | 0.01152649 |
| 11 | 64242972 | T | C | 0.740304 | 2.95E-08 | 0.01157489 |
| 11 | 64139512 | T | C | 0.6914 | 2.98E-08 | 0.01168833 |
| 11 | 64266775 | G | T | 0.740304 | 3.04E-08 | 0.01190655 |
| 11 | 64266890 | T | C | 0.740304 | 3.04E-08 | 0.01190655 |
| 11 | 64267009 | A | C | 0.740304 | 3.04E-08 | 0.01190655 |
| 11 | 64267215 | T | C | 0.740304 | 3.04E-08 | 0.01190655 |
| 11 | 64267888 | A | G | 0.740304 | 3.04E-08 | 0.01190655 |
| 11 | 64267999 | C | T | 0.740304 | 3.04E-08 | 0.01190655 |
| 11 | 64185448 | A | C | 0.692243 | 3.09E-08 | 0.01211022 |
| 11 | 64231178 | T | A | 0.737774 | 3.21E-08 | 0.01260144 |
| 11 | 64231306 | G | A | 0.737774 | 3.21E-08 | 0.01260144 |
| 11 | 65162056 | T | C | 0.714165 | 3.25E-08 | 0.01273706 |
| 11 | 64114150 | C | T | 0.695616 | 3.29E-08 | 0.0128888 |
| 11 | 64114170 | C | T | 0.695616 | 3.29E-08 | 0.0128888 |
| 11 | 64114228 | G | A | 0.695616 | 3.29E-08 | 0.0128888 |
| 11 | 64114390 | A | G | 0.695616 | 3.29E-08 | 0.0128888 |
| 11 | 64110123 | C | G | 0.693086 | 3.30E-08 | 0.01292329 |
| 11 | 64595978 | G | C | 0.738617 | 3.31E-08 | 0.01296876 |
| 11 | 64116145 | T | C | 0.693086 | 3.33E-08 | 0.01304022 |
| 11 | 64118462 | T | C | 0.693086 | 3.33E-08 | 0.01304022 |
| 11 | 64118587 | A | G | 0.693086 | 3.33E-08 | 0.01304022 |
| 11 | 64118647 | G | A | 0.693086 | 3.33E-08 | 0.01304022 |
| 11 | 64118722 | T | C | 0.693086 | 3.33E-08 | 0.01304022 |
| 11 | 64118806 | A | C | 0.693086 | 3.33E-08 | 0.01304022 |
| 11 | 64118903 | G | A | 0.693086 | 3.33E-08 | 0.01304022 |
| 11 | 64120158 | C | T | 0.693086 | 3.33E-08 | 0.01304022 |
| 11 | 64121251 | G | A | 0.693086 | 3.33E-08 | 0.01304022 |
| 11 | 64121441 | C | T | 0.693086 | 3.33E-08 | 0.01304022 |
| 11 | 64122451 | A | G | 0.693086 | 3.33E-08 | 0.01304022 |
| 11 | 64122474 | G | T | 0.693086 | 3.33E-08 | 0.01304022 |
| 11 | 64122507 | C | T | 0.693086 | 3.33E-08 | 0.01304022 |
| 11 | 64122750 | T | G | 0.693086 | 3.33E-08 | 0.01304022 |
| 11 | 64122796 | C | T | 0.693086 | 3.33E-08 | 0.01304022 |
| 11 | 64122823 | T | C | 0.693086 | 3.33E-08 | 0.01304022 |
| 11 | 64122875 | C | G | 0.693086 | 3.33E-08 | 0.01304022 |
| 11 | 64123156 | G | C | 0.693086 | 3.33E-08 | 0.01304022 |
| 11 | 64123633 | G | A | 0.693086 | 3.33E-08 | 0.01304022 |
| 11 | 64123861 | G | T | 0.693086 | 3.33E-08 | 0.01304022 |
| 11 | 64124278 | C | A | 0.693086 | 3.33E-08 | 0.01304022 |
| 11 | 64124626 | G | C | 0.693086 | 3.33E-08 | 0.01304022 |
| 11 | 64124655 | G | A | 0.693086 | 3.33E-08 | 0.01304022 |
| 11 | 64124770 | C | T | 0.693086 | 3.33E-08 | 0.01304022 |
| 11 | 64124978 | T | C | 0.693086 | 3.33E-08 | 0.01304022 |
| 11 | 64125027 | G | A | 0.693086 | 3.33E-08 | 0.01304022 |
| 11 | 64125120 | G | A | 0.693086 | 3.33E-08 | 0.01304022 |
| 11 | 64125322 | T | C | 0.693086 | 3.33E-08 | 0.01304022 |
| 11 | 64125523 | G | A | 0.693086 | 3.33E-08 | 0.01304022 |
| 11 | 64126012 | G | T | 0.693086 | 3.33E-08 | 0.01304022 |
| 11 | 64126095 | G | T | 0.693086 | 3.33E-08 | 0.01304022 |
| 11 | 64126195 | C | T | 0.693086 | 3.33E-08 | 0.01304022 |
| 11 | 64126203 | C | T | 0.693086 | 3.33E-08 | 0.01304022 |
| 11 | 64127166 | A | G | 0.693086 | 3.33E-08 | 0.01304022 |
| 11 | 64127462 | A | G | 0.693086 | 3.33E-08 | 0.01304022 |
| 11 | 64127497 | T | C | 0.693086 | 3.33E-08 | 0.01304022 |
| 11 | 64127580 | G | T | 0.693086 | 3.33E-08 | 0.01304022 |
| 11 | 64127691 | T | C | 0.693086 | 3.33E-08 | 0.01304022 |
| 11 | 64128045 | G | T | 0.693086 | 3.33E-08 | 0.01304022 |
| 11 | 64129693 | T | C | 0.693086 | 3.33E-08 | 0.01304022 |
| 11 | 64129931 | T | A | 0.693086 | 3.33E-08 | 0.01304022 |
| 11 | 64129932 | G | T | 0.693086 | 3.33E-08 | 0.01304022 |
| 11 | 64130402 | A | G | 0.693086 | 3.33E-08 | 0.01304022 |
| 11 | 64136571 | T | C | 0.693086 | 3.33E-08 | 0.01304022 |
| 11 | 64139430 | C | T | 0.693086 | 3.33E-08 | 0.01304022 |
| 11 | 64139722 | T | G | 0.693086 | 3.33E-08 | 0.01304022 |
| 11 | 64139854 | G | T | 0.693086 | 3.33E-08 | 0.01304022 |
| 11 | 64139907 | A | T | 0.693086 | 3.33E-08 | 0.01304022 |

| | | | | | | |
|---|---|---|---|---|---|---|
| 11 | 64139908 | A | C | 0.693086 | 3.33E-08 | 0.01304022 |
| 11 | 64139949 | A | T | 0.693086 | 3.33E-08 | 0.01304022 |
| 11 | 64139969 | C | T | 0.693086 | 3.33E-08 | 0.01304022 |
| 11 | 64139988 | T | C | 0.693086 | 3.33E-08 | 0.01304022 |
| 11 | 64140391 | A | G | 0.693086 | 3.33E-08 | 0.01304022 |
| 11 | 64140412 | G | C | 0.693086 | 3.33E-08 | 0.01304022 |
| 11 | 64140554 | G | A | 0.693086 | 3.33E-08 | 0.01304022 |
| 11 | 64140605 | C | G | 0.693086 | 3.33E-08 | 0.01304022 |
| 11 | 64140677 | A | G | 0.693086 | 3.33E-08 | 0.01304022 |
| 11 | 64140789 | C | T | 0.693086 | 3.33E-08 | 0.01304022 |
| 11 | 64140793 | C | T | 0.693086 | 3.33E-08 | 0.01304022 |
| 11 | 64140993 | T | C | 0.693086 | 3.33E-08 | 0.01304022 |
| 11 | 64141006 | G | A | 0.693086 | 3.33E-08 | 0.01304022 |
| 11 | 64141023 | A | G | 0.693086 | 3.33E-08 | 0.01304022 |
| 11 | 64141239 | G | A | 0.693086 | 3.33E-08 | 0.01304022 |
| 11 | 64141258 | T | A | 0.693086 | 3.33E-08 | 0.01304022 |
| 11 | 64141431 | C | T | 0.693086 | 3.33E-08 | 0.01304022 |
| 11 | 64141432 | A | G | 0.693086 | 3.33E-08 | 0.01304022 |
| 11 | 64141434 | T | C | 0.693086 | 3.33E-08 | 0.01304022 |
| 11 | 64141479 | A | G | 0.693086 | 3.33E-08 | 0.01304022 |
| 11 | 64141645 | A | C | 0.693086 | 3.33E-08 | 0.01304022 |
| 11 | 64142012 | T | C | 0.693086 | 3.33E-08 | 0.01304022 |
| 11 | 64142216 | T | G | 0.693086 | 3.33E-08 | 0.01304022 |
| 11 | 64142434 | G | A | 0.693086 | 3.33E-08 | 0.01304022 |
| 11 | 64145069 | T | C | 0.693086 | 3.33E-08 | 0.01304022 |
| 11 | 64145863 | C | T | 0.693086 | 3.33E-08 | 0.01304022 |
| 11 | 64145894 | G | A | 0.693086 | 3.33E-08 | 0.01304022 |
| 11 | 64148850 | C | G | 0.693086 | 3.33E-08 | 0.01304022 |
| 11 | 64153819 | C | G | 0.693086 | 3.33E-08 | 0.01304022 |
| 11 | 64154206 | G | A | 0.693086 | 3.33E-08 | 0.01304022 |
| 11 | 64154639 | G | A | 0.693086 | 3.33E-08 | 0.01304022 |
| 11 | 64154651 | A | C | 0.693086 | 3.33E-08 | 0.01304022 |
| 11 | 64155145 | G | A | 0.693086 | 3.33E-08 | 0.01304022 |
| 11 | 64155173 | G | A | 0.693086 | 3.33E-08 | 0.01304022 |
| 11 | 64155219 | G | A | 0.693086 | 3.33E-08 | 0.01304022 |
| 11 | 64156517 | T | C | 0.693086 | 3.33E-08 | 0.01304022 |
| 11 | 64157800 | G | A | 0.693086 | 3.33E-08 | 0.01304022 |
| 11 | 64159411 | C | G | 0.693086 | 3.33E-08 | 0.01304022 |
| 11 | 64159980 | C | T | 0.693086 | 3.33E-08 | 0.01304022 |
| 11 | 64160042 | C | G | 0.693086 | 3.33E-08 | 0.01304022 |
| 11 | 64160212 | G | C | 0.693086 | 3.33E-08 | 0.01304022 |
| 11 | 64161453 | T | C | 0.693086 | 3.33E-08 | 0.01304022 |
| 11 | 64161909 | G | A | 0.693086 | 3.33E-08 | 0.01304022 |
| 11 | 64161936 | T | C | 0.693086 | 3.33E-08 | 0.01304022 |
| 11 | 64162145 | A | C | 0.693086 | 3.33E-08 | 0.01304022 |
| 11 | 64162162 | A | T | 0.693086 | 3.33E-08 | 0.01304022 |
| 11 | 64162262 | G | A | 0.693086 | 3.33E-08 | 0.01304022 |
| 11 | 64162343 | G | A | 0.693086 | 3.33E-08 | 0.01304022 |
| 11 | 64162344 | C | G | 0.693086 | 3.33E-08 | 0.01304022 |
| 11 | 64162371 | G | A | 0.693086 | 3.33E-08 | 0.01304022 |
| 11 | 64162419 | A | T | 0.693086 | 3.33E-08 | 0.01304022 |
| 11 | 64162596 | G | A | 0.693086 | 3.33E-08 | 0.01304022 |
| 11 | 64162716 | G | T | 0.693086 | 3.33E-08 | 0.01304022 |
| 11 | 64163049 | G | A | 0.693086 | 3.33E-08 | 0.01304022 |
| 11 | 64163178 | A | G | 0.693086 | 3.33E-08 | 0.01304022 |
| 11 | 64163253 | A | G | 0.693086 | 3.33E-08 | 0.01304022 |
| 11 | 64170552 | G | A | 0.693086 | 3.33E-08 | 0.01304022 |
| 11 | 64171233 | G | A | 0.693086 | 3.33E-08 | 0.01304022 |
| 11 | 64172708 | C | T | 0.693086 | 3.33E-08 | 0.01304022 |
| 11 | 64173426 | G | A | 0.693086 | 3.33E-08 | 0.01304022 |
| 11 | 64174650 | A | T | 0.693086 | 3.33E-08 | 0.01304022 |
| 11 | 64182126 | G | A | 0.693086 | 3.33E-08 | 0.01304022 |
| 11 | 64924032 | C | T | 0.810287 | 3.38E-08 | 0.01324373 |
| 11 | 64110247 | G | C | 0.752951 | 3.42E-08 | 0.01340601 |
| 11 | 64110334 | T | A | 0.752951 | 3.42E-08 | 0.01340601 |
| 11 | 64212333 | A | G | 0.729342 | 3.49E-08 | 0.01366295 |
| 7 | 138975863 | T | C | 0.946037 | 3.60E-08 | 0.01410012 |

| | | | | | | |
|---|---|---|---|---|---|---|
| 11 | 64113786 | A | C | 0.693929 | 3.75E-08 | 0.01469698 |
| 11 | 64113857 | T | C | 0.693929 | 3.75E-08 | 0.01469698 |
| 11 | 64113866 | G | A | 0.693929 | 3.75E-08 | 0.01469698 |
| 11 | 64114810 | G | A | 0.693929 | 3.75E-08 | 0.01469698 |
| 11 | 64114842 | G | A | 0.693929 | 3.75E-08 | 0.01469698 |
| 11 | 64114960 | T | G | 0.693929 | 3.75E-08 | 0.01469698 |
| 11 | 64109544 | G | A | 0.751265 | 3.78E-08 | 0.01482775 |
| 11 | 64212985 | C | T | 0.733558 | 3.91E-08 | 0.01532536 |
| 11 | 64213001 | A | G | 0.733558 | 3.91E-08 | 0.01532536 |
| 11 | 64213005 | A | G | 0.733558 | 3.91E-08 | 0.01532536 |
| 11 | 64554429 | G | T | 0.749578 | 4.40E-08 | 0.01725604 |
| 11 | 64164686 | C | A | 0.694772 | 4.44E-08 | 0.01738884 |
| 11 | 64640983 | G | C | 0.741147 | 4.48E-08 | 0.017557 |
| 11 | 64201890 | A | G | 0.734401 | 4.50E-08 | 0.0176225 |
| 11 | 64202614 | T | C | 0.734401 | 4.50E-08 | 0.0176225 |
| 11 | 64204411 | C | A | 0.734401 | 4.50E-08 | 0.0176225 |
| 11 | 64205613 | T | C | 0.734401 | 4.50E-08 | 0.0176225 |
| 11 | 64206496 | A | G | 0.734401 | 4.50E-08 | 0.0176225 |
| 11 | 64206688 | A | G | 0.734401 | 4.50E-08 | 0.0176225 |
| 11 | 64206746 | A | G | 0.734401 | 4.50E-08 | 0.0176225 |
| 11 | 64206768 | G | A | 0.734401 | 4.50E-08 | 0.0176225 |
| 11 | 64206800 | T | G | 0.734401 | 4.50E-08 | 0.0176225 |
| 11 | 64207305 | G | A | 0.734401 | 4.50E-08 | 0.0176225 |
| 11 | 64207665 | G | A | 0.734401 | 4.50E-08 | 0.0176225 |
| 11 | 64207724 | A | G | 0.734401 | 4.50E-08 | 0.0176225 |
| 11 | 64207891 | G | T | 0.734401 | 4.50E-08 | 0.0176225 |
| 11 | 64207954 | T | C | 0.734401 | 4.50E-08 | 0.0176225 |
| 11 | 64208157 | C | T | 0.734401 | 4.50E-08 | 0.0176225 |
| 11 | 64208234 | C | T | 0.734401 | 4.50E-08 | 0.0176225 |
| 11 | 64208253 | C | T | 0.734401 | 4.50E-08 | 0.0176225 |
| 11 | 64208749 | T | C | 0.734401 | 4.50E-08 | 0.0176225 |
| 11 | 64208968 | A | C | 0.734401 | 4.50E-08 | 0.0176225 |
| 11 | 64209079 | T | C | 0.734401 | 4.50E-08 | 0.0176225 |
| 11 | 64209359 | G | C | 0.734401 | 4.50E-08 | 0.0176225 |
| 11 | 64213176 | T | C | 0.734401 | 4.50E-08 | 0.0176225 |
| 11 | 64213218 | T | A | 0.734401 | 4.50E-08 | 0.0176225 |
| 11 | 64213328 | C | T | 0.734401 | 4.50E-08 | 0.0176225 |
| 11 | 64213329 | A | G | 0.734401 | 4.50E-08 | 0.0176225 |
| 11 | 64213336 | A | G | 0.734401 | 4.50E-08 | 0.0176225 |
| 11 | 64213343 | A | G | 0.734401 | 4.50E-08 | 0.0176225 |
| 11 | 64213421 | C | G | 0.734401 | 4.50E-08 | 0.0176225 |
| 11 | 64213422 | G | T | 0.734401 | 4.50E-08 | 0.0176225 |
| 11 | 64213482 | C | T | 0.734401 | 4.50E-08 | 0.0176225 |
| 11 | 64213502 | A | G | 0.734401 | 4.50E-08 | 0.0176225 |
| 11 | 64213573 | A | G | 0.734401 | 4.50E-08 | 0.0176225 |
| 11 | 64213576 | T | G | 0.734401 | 4.50E-08 | 0.0176225 |
| 11 | 64213638 | C | G | 0.734401 | 4.50E-08 | 0.0176225 |
| 11 | 64213639 | A | G | 0.734401 | 4.50E-08 | 0.0176225 |
| 11 | 64213648 | T | C | 0.734401 | 4.50E-08 | 0.0176225 |
| 11 | 64213845 | G | A | 0.734401 | 4.50E-08 | 0.0176225 |
| 11 | 64213961 | A | G | 0.734401 | 4.50E-08 | 0.0176225 |
| 11 | 64213970 | T | A | 0.734401 | 4.50E-08 | 0.0176225 |
| 11 | 64214048 | T | C | 0.734401 | 4.50E-08 | 0.0176225 |
| 11 | 64214075 | G | C | 0.734401 | 4.50E-08 | 0.0176225 |
| 11 | 64116397 | T | G | 0.692243 | 4.64E-08 | 0.0182058 |
| 11 | 64116425 | C | T | 0.692243 | 4.64E-08 | 0.0182058 |
| 11 | 64116456 | A | G | 0.692243 | 4.64E-08 | 0.0182058 |
| 11 | 64211929 | T | G | 0.732715 | 4.74E-08 | 0.01857958 |
| 11 | 65122821 | C | T | 0.710793 | 4.86E-08 | 0.01903071 |
| 11 | 65122836 | G | A | 0.710793 | 4.86E-08 | 0.01903071 |
| 11 | 64138652 | A | G | 0.750422 | 5.01E-08 | 0.01962259 |
| 11 | 64171370 | G | C | 0.750422 | 5.17E-08 | 0.02025109 |
| 11 | 64116725 | A | G | 0.693929 | 5.18E-08 | 0.0203187 |
| 11 | 64116840 | G | A | 0.693929 | 5.18E-08 | 0.0203187 |
| 11 | 64116844 | G | A | 0.693929 | 5.18E-08 | 0.0203187 |
| 11 | 64116924 | G | A | 0.693929 | 5.18E-08 | 0.0203187 |
| 11 | 64117876 | G | A | 0.693929 | 5.18E-08 | 0.0203187 |

| | | | | | | |
|---|---|---|---|---|---|---|
| 11 | 64118019 | C | G | 0.693929 | 5.18E-08 | 0.0203187 |
| 11 | 64114078 | G | A | 0.6914 | 5.28E-08 | 0.02069151 |
| 11 | 64114101 | C | G | 0.6914 | 5.28E-08 | 0.02069151 |
| 11 | 64794718 | C | T | 0.742833 | 5.54E-08 | 0.02172296 |
| 11 | 64163720 | G | T | 0.693929 | 5.70E-08 | 0.02233174 |
| 11 | 64164322 | C | T | 0.693929 | 5.70E-08 | 0.02233174 |
| 11 | 64131388 | T | C | 0.692243 | 5.70E-08 | 0.02233934 |
| 11 | 64131394 | T | C | 0.692243 | 5.70E-08 | 0.02233934 |
| 11 | 64131419 | A | G | 0.692243 | 5.70E-08 | 0.02233934 |
| 11 | 64131577 | G | A | 0.692243 | 5.70E-08 | 0.02233934 |
| 11 | 64131582 | C | T | 0.692243 | 5.70E-08 | 0.02233934 |
| 11 | 64131925 | G | T | 0.692243 | 5.70E-08 | 0.02233934 |
| 11 | 64132301 | C | A | 0.692243 | 5.70E-08 | 0.02233934 |
| 11 | 64254213 | A | G | 0.740304 | 5.83E-08 | 0.02286059 |
| 11 | 64551920 | A | G | 0.742833 | 5.86E-08 | 0.02295188 |
| 11 | 69871025 | G | T | 0.79511 | 5.90E-08 | 0.02312317 |
| 11 | 64185266 | T | G | 0.750422 | 6.27E-08 | 0.02459438 |
| 11 | 64185805 | A | G | 0.750422 | 6.27E-08 | 0.02459438 |
| 11 | 64120248 | C | T | 0.751265 | 6.47E-08 | 0.02537112 |
| 11 | 64120259 | A | T | 0.751265 | 6.47E-08 | 0.02537112 |
| 11 | 64120318 | C | T | 0.751265 | 6.47E-08 | 0.02537112 |
| 11 | 64123534 | C | G | 0.751265 | 6.47E-08 | 0.02537112 |
| 11 | 64124100 | A | C | 0.751265 | 6.47E-08 | 0.02537112 |
| 11 | 64124165 | T | C | 0.751265 | 6.47E-08 | 0.02537112 |
| 11 | 64124228 | G | A | 0.751265 | 6.47E-08 | 0.02537112 |
| 11 | 64125521 | T | C | 0.751265 | 6.47E-08 | 0.02537112 |
| 11 | 64126291 | C | T | 0.751265 | 6.47E-08 | 0.02537112 |
| 11 | 64127523 | C | A | 0.751265 | 6.47E-08 | 0.02537112 |
| 11 | 64130160 | G | C | 0.751265 | 6.47E-08 | 0.02537112 |
| 11 | 64131018 | A | T | 0.751265 | 6.47E-08 | 0.02537112 |
| 11 | 64131133 | G | A | 0.751265 | 6.47E-08 | 0.02537112 |
| 11 | 64136127 | C | T | 0.751265 | 6.47E-08 | 0.02537112 |
| 11 | 64136357 | G | A | 0.751265 | 6.47E-08 | 0.02537112 |
| 11 | 64139352 | T | C | 0.751265 | 6.47E-08 | 0.02537112 |
| 11 | 64140063 | A | G | 0.751265 | 6.47E-08 | 0.02537112 |
| 11 | 64140517 | A | C | 0.751265 | 6.47E-08 | 0.02537112 |
| 11 | 64140518 | A | C | 0.751265 | 6.47E-08 | 0.02537112 |
| 11 | 64141407 | G | A | 0.751265 | 6.47E-08 | 0.02537112 |
| 11 | 64142064 | A | G | 0.751265 | 6.47E-08 | 0.02537112 |
| 11 | 64142724 | T | C | 0.751265 | 6.47E-08 | 0.02537112 |
| 11 | 64147047 | A | G | 0.751265 | 6.47E-08 | 0.02537112 |
| 11 | 64147364 | T | C | 0.751265 | 6.47E-08 | 0.02537112 |
| 11 | 64148130 | C | G | 0.751265 | 6.47E-08 | 0.02537112 |
| 11 | 64150638 | T | C | 0.751265 | 6.47E-08 | 0.02537112 |
| 11 | 64153858 | T | C | 0.751265 | 6.47E-08 | 0.02537112 |
| 11 | 64153881 | G | A | 0.751265 | 6.47E-08 | 0.02537112 |
| 11 | 64154005 | T | C | 0.751265 | 6.47E-08 | 0.02537112 |
| 11 | 64154113 | T | C | 0.751265 | 6.47E-08 | 0.02537112 |
| 11 | 64154668 | A | C | 0.751265 | 6.47E-08 | 0.02537112 |
| 11 | 64155017 | G | C | 0.751265 | 6.47E-08 | 0.02537112 |
| 11 | 64155114 | G | A | 0.751265 | 6.47E-08 | 0.02537112 |
| 11 | 64156706 | T | C | 0.751265 | 6.47E-08 | 0.02537112 |
| 11 | 64157542 | A | G | 0.751265 | 6.47E-08 | 0.02537112 |
| 11 | 64161147 | G | A | 0.751265 | 6.47E-08 | 0.02537112 |
| 11 | 64161724 | C | A | 0.751265 | 6.47E-08 | 0.02537112 |
| 11 | 64161903 | G | A | 0.751265 | 6.47E-08 | 0.02537112 |
| 11 | 64162437 | C | G | 0.751265 | 6.47E-08 | 0.02537112 |
| 11 | 64162737 | G | A | 0.751265 | 6.47E-08 | 0.02537112 |
| 11 | 64162746 | C | A | 0.751265 | 6.47E-08 | 0.02537112 |
| 11 | 64163683 | C | G | 0.751265 | 6.47E-08 | 0.02537112 |
| 11 | 64170763 | T | C | 0.751265 | 6.47E-08 | 0.02537112 |
| 11 | 64171329 | G | A | 0.751265 | 6.47E-08 | 0.02537112 |
| 11 | 64171346 | G | A | 0.751265 | 6.47E-08 | 0.02537112 |
| 11 | 64171351 | T | G | 0.751265 | 6.47E-08 | 0.02537112 |
| 11 | 64171868 | G | T | 0.751265 | 6.47E-08 | 0.02537112 |
| 11 | 64172013 | T | G | 0.751265 | 6.47E-08 | 0.02537112 |
| 11 | 64172027 | G | A | 0.751265 | 6.47E-08 | 0.02537112 |

| | | | | | | |
|---|---|---|---|---|---|---|
| 11 | 64172170 | A | G | 0.751265 | 6.47E-08 | 0.02537112 |
| 11 | 64172278 | T | C | 0.751265 | 6.47E-08 | 0.02537112 |
| 11 | 64172383 | A | G | 0.751265 | 6.47E-08 | 0.02537112 |
| 11 | 64172413 | T | C | 0.751265 | 6.47E-08 | 0.02537112 |
| 11 | 64172444 | T | C | 0.751265 | 6.47E-08 | 0.02537112 |
| 11 | 64172493 | T | A | 0.751265 | 6.47E-08 | 0.02537112 |
| 11 | 64172647 | T | C | 0.751265 | 6.47E-08 | 0.02537112 |
| 11 | 64173210 | A | G | 0.751265 | 6.47E-08 | 0.02537112 |
| 11 | 64179550 | T | C | 0.751265 | 6.47E-08 | 0.02537112 |
| 11 | 64179569 | T | C | 0.751265 | 6.47E-08 | 0.02537112 |
| 11 | 64179576 | A | G | 0.751265 | 6.47E-08 | 0.02537112 |
| 11 | 64180854 | A | T | 0.751265 | 6.47E-08 | 0.02537112 |
| 11 | 64180866 | G | A | 0.751265 | 6.47E-08 | 0.02537112 |
| 11 | 64180993 | C | T | 0.751265 | 6.47E-08 | 0.02537112 |
| 11 | 64181184 | A | G | 0.751265 | 6.47E-08 | 0.02537112 |
| 11 | 64181556 | C | T | 0.751265 | 6.47E-08 | 0.02537112 |
| 11 | 64181870 | T | C | 0.751265 | 6.47E-08 | 0.02537112 |
| 11 | 64182084 | C | G | 0.751265 | 6.47E-08 | 0.02537112 |
| 11 | 64182144 | T | C | 0.751265 | 6.47E-08 | 0.02537112 |
| 11 | 64182172 | A | G | 0.751265 | 6.47E-08 | 0.02537112 |
| 11 | 64182304 | T | C | 0.751265 | 6.47E-08 | 0.02537112 |
| 11 | 64182402 | A | T | 0.751265 | 6.47E-08 | 0.02537112 |
| 11 | 64192454 | T | C | 0.751265 | 6.47E-08 | 0.02537112 |
| 11 | 64196945 | G | C | 0.751265 | 6.47E-08 | 0.02537112 |
| 11 | 64783767 | G | A | 0.63575 | 6.49E-08 | 0.0254296 |
| 11 | 64212023 | G | T | 0.732715 | 6.55E-08 | 0.02565847 |
| 11 | 64177759 | A | G | 0.750422 | 6.55E-08 | 0.02566267 |
| 11 | 64177918 | A | G | 0.750422 | 6.55E-08 | 0.02566267 |
| 11 | 64219218 | T | C | 0.740304 | 6.55E-08 | 0.02566878 |
| 11 | 64802484 | T | C | 0.744519 | 7.02E-08 | 0.02750264 |
| 11 | 64805150 | T | C | 0.744519 | 7.02E-08 | 0.02750264 |
| 11 | 64176719 | A | G | 0.752108 | 7.16E-08 | 0.02806893 |
| 11 | 64117131 | C | T | 0.693086 | 7.19E-08 | 0.02819938 |
| 11 | 64117201 | C | T | 0.693086 | 7.19E-08 | 0.02819938 |
| 11 | 64117226 | C | T | 0.693086 | 7.19E-08 | 0.02819938 |
| 11 | 64117258 | G | A | 0.693086 | 7.19E-08 | 0.02819938 |
| 11 | 64179506 | A | G | 0.752108 | 7.35E-08 | 0.02882058 |
| 11 | 64115276 | C | G | 0.692243 | 7.36E-08 | 0.02886437 |
| 11 | 64064718 | C | T | 0.666105 | 7.37E-08 | 0.02888824 |
| 11 | 64900263 | G | A | 0.572513 | 7.50E-08 | 0.02940792 |
| 11 | 64900281 | A | C | 0.572513 | 7.50E-08 | 0.02940792 |
| 11 | 64916765 | T | C | 0.572513 | 7.50E-08 | 0.02940792 |
| 11 | 64916801 | C | A | 0.572513 | 7.50E-08 | 0.02940792 |
| 11 | 64916974 | C | A | 0.572513 | 7.50E-08 | 0.02940792 |
| 11 | 64917033 | C | T | 0.572513 | 7.50E-08 | 0.02940792 |
| 11 | 64917085 | A | T | 0.572513 | 7.50E-08 | 0.02940792 |
| 11 | 64917306 | G | A | 0.572513 | 7.50E-08 | 0.02940792 |
| 11 | 64917332 | G | A | 0.572513 | 7.50E-08 | 0.02940792 |
| 11 | 64917367 | C | T | 0.572513 | 7.50E-08 | 0.02940792 |
| 11 | 64917398 | T | G | 0.572513 | 7.50E-08 | 0.02940792 |
| 11 | 64917403 | C | T | 0.572513 | 7.50E-08 | 0.02940792 |
| 11 | 64917423 | T | G | 0.572513 | 7.50E-08 | 0.02940792 |
| 11 | 64917475 | T | C | 0.572513 | 7.50E-08 | 0.02940792 |
| 11 | 64917480 | T | C | 0.572513 | 7.50E-08 | 0.02940792 |
| 11 | 64917513 | C | G | 0.572513 | 7.50E-08 | 0.02940792 |
| 11 | 64917518 | T | C | 0.572513 | 7.50E-08 | 0.02940792 |
| 11 | 64917542 | G | C | 0.572513 | 7.50E-08 | 0.02940792 |
| 11 | 64917590 | T | A | 0.572513 | 7.50E-08 | 0.02940792 |
| 11 | 64917607 | G | T | 0.572513 | 7.50E-08 | 0.02940792 |
| 11 | 64917708 | G | A | 0.572513 | 7.50E-08 | 0.02940792 |
| 11 | 64918576 | G | T | 0.572513 | 7.50E-08 | 0.02940792 |
| 11 | 64918776 | C | G | 0.572513 | 7.50E-08 | 0.02940792 |
| 11 | 64918824 | A | G | 0.572513 | 7.50E-08 | 0.02940792 |
| 11 | 64170762 | G | A | 0.746206 | 7.62E-08 | 0.02985963 |
| 11 | 64682035 | A | C | 0.746206 | 7.79E-08 | 0.03051576 |
| 11 | 64682036 | A | T | 0.746206 | 7.79E-08 | 0.03051576 |
| 11 | 64682066 | C | T | 0.746206 | 7.79E-08 | 0.03051576 |

| | | | | | | |
|---|---|---|---|---|---|---|
| 11 | 64590934 | A | G | 0.748735 | 7.79E-08 | 0.03052705 |
| 11 | 64555315 | A | G | 0.744519 | 7.83E-08 | 0.03069874 |
| 11 | 64556772 | G | A | 0.744519 | 7.83E-08 | 0.03069874 |
| 11 | 64556906 | T | C | 0.744519 | 7.83E-08 | 0.03069874 |
| 11 | 64559493 | G | T | 0.744519 | 7.83E-08 | 0.03069874 |
| 11 | 64560392 | A | G | 0.744519 | 7.83E-08 | 0.03069874 |
| 11 | 64562104 | C | T | 0.744519 | 7.83E-08 | 0.03069874 |
| 11 | 64562117 | C | A | 0.744519 | 7.83E-08 | 0.03069874 |
| 11 | 64562153 | G | A | 0.744519 | 7.83E-08 | 0.03069874 |
| 11 | 64180855 | A | T | 0.697302 | 8.01E-08 | 0.03139277 |
| 11 | 66065257 | G | C | 0.744519 | 8.37E-08 | 0.03280518 |
| 11 | 66065393 | C | T | 0.744519 | 8.37E-08 | 0.03280518 |
| 11 | 66067997 | T | C | 0.744519 | 8.37E-08 | 0.03280518 |
| 11 | 64591338 | C | T | 0.751265 | 8.42E-08 | 0.03302143 |
| 11 | 64591352 | G | A | 0.751265 | 8.42E-08 | 0.03302143 |
| 11 | 71455256 | A | G | 0.903035 | 8.81E-08 | 0.03451799 |
| 11 | 64114125 | T | C | 0.693929 | 8.85E-08 | 0.03467667 |
| 11 | 64679587 | G | A | 0.744519 | 8.97E-08 | 0.03517655 |
| 11 | 64679653 | G | A | 0.744519 | 8.97E-08 | 0.03517655 |
| 11 | 64679656 | A | G | 0.744519 | 8.97E-08 | 0.03517655 |
| 11 | 64679689 | G | A | 0.744519 | 8.97E-08 | 0.03517655 |
| 11 | 64679718 | G | A | 0.744519 | 8.97E-08 | 0.03517655 |
| 11 | 64679821 | G | A | 0.744519 | 8.97E-08 | 0.03517655 |
| 11 | 64679991 | A | G | 0.744519 | 8.97E-08 | 0.03517655 |
| 11 | 64680004 | C | T | 0.744519 | 8.97E-08 | 0.03517655 |
| 11 | 64680053 | C | G | 0.744519 | 8.97E-08 | 0.03517655 |
| 11 | 64680061 | C | T | 0.744519 | 8.97E-08 | 0.03517655 |
| 11 | 64680090 | A | C | 0.744519 | 8.97E-08 | 0.03517655 |
| 11 | 64680110 | A | G | 0.744519 | 8.97E-08 | 0.03517655 |
| 11 | 64680112 | A | C | 0.744519 | 8.97E-08 | 0.03517655 |
| 11 | 64680120 | G | A | 0.744519 | 8.97E-08 | 0.03517655 |
| 11 | 64680177 | C | T | 0.744519 | 8.97E-08 | 0.03517655 |
| 11 | 64680358 | A | G | 0.744519 | 8.97E-08 | 0.03517655 |
| 11 | 64680414 | G | A | 0.744519 | 8.97E-08 | 0.03517655 |
| 11 | 64680551 | A | G | 0.744519 | 8.97E-08 | 0.03517655 |
| 11 | 64680686 | T | G | 0.744519 | 8.97E-08 | 0.03517655 |
| 11 | 64680757 | A | G | 0.744519 | 8.97E-08 | 0.03517655 |
| 11 | 64680787 | T | C | 0.744519 | 8.97E-08 | 0.03517655 |
| 11 | 64680957 | A | G | 0.744519 | 8.97E-08 | 0.03517655 |
| 11 | 64680962 | T | G | 0.744519 | 8.97E-08 | 0.03517655 |
| 11 | 64681204 | C | A | 0.744519 | 8.97E-08 | 0.03517655 |
| 11 | 64681392 | G | A | 0.744519 | 8.97E-08 | 0.03517655 |
| 11 | 64681404 | G | T | 0.744519 | 8.97E-08 | 0.03517655 |
| 11 | 64681423 | T | A | 0.744519 | 8.97E-08 | 0.03517655 |
| 11 | 64234872 | T | C | 0.732715 | 8.99E-08 | 0.03522457 |
| 11 | 64592962 | G | A | 0.743676 | 9.01E-08 | 0.03532107 |
| 11 | 64593052 | T | C | 0.743676 | 9.01E-08 | 0.03532107 |
| 11 | 64593092 | C | T | 0.743676 | 9.01E-08 | 0.03532107 |
| 11 | 64593109 | A | G | 0.743676 | 9.01E-08 | 0.03532107 |
| 11 | 64215669 | T | C | 0.733558 | 9.11E-08 | 0.0357167 |
| 11 | 64591054 | G | C | 0.749578 | 9.18E-08 | 0.03599492 |
| 11 | 64591119 | A | G | 0.749578 | 9.18E-08 | 0.03599492 |
| 11 | 64682903 | G | A | 0.744519 | 9.31E-08 | 0.03647533 |
| 11 | 64682956 | C | G | 0.744519 | 9.31E-08 | 0.03647533 |
| 11 | 64667419 | G | A | 0.742833 | 9.61E-08 | 0.03765012 |
| 11 | 64776777 | T | A | 0.746206 | 9.73E-08 | 0.03813876 |
| 11 | 64776861 | T | C | 0.746206 | 9.73E-08 | 0.03813876 |
| 11 | 64684717 | C | A | 0.744519 | 9.85E-08 | 0.03860489 |
| 11 | 64685598 | G | T | 0.744519 | 9.85E-08 | 0.03860489 |
| 11 | 64685647 | G | T | 0.744519 | 9.85E-08 | 0.03860489 |
| 11 | 64685744 | G | A | 0.744519 | 9.85E-08 | 0.03860489 |
| 11 | 64685831 | T | C | 0.744519 | 9.85E-08 | 0.03860489 |
| 11 | 64707405 | T | C | 0.744519 | 9.95E-08 | 0.03900447 |
| 11 | 64707516 | A | C | 0.744519 | 9.95E-08 | 0.03900447 |
| 11 | 64171186 | C | T | 0.750422 | 1.00E-07 | 0.03922621 |
| 11 | 64061509 | C | T | 0.693086 | 1.01E-07 | 0.03945474 |
| 11 | 64116467 | A | G | 0.6914 | 1.01E-07 | 0.03961819 |

| | | | | | | |
|---|---|---|---|---|---|---|
| 11 | 64116491 | C | T | 0.6914 | 1.01E-07 | 0.03961819 |
| 11 | 64116503 | T | C | 0.6914 | 1.01E-07 | 0.03961819 |
| 11 | 64578652 | G | A | 0.744519 | 1.01E-07 | 0.03977067 |
| 11 | 64578706 | A | G | 0.744519 | 1.01E-07 | 0.03977067 |
| 11 | 65033511 | C | T | 0.684654 | 1.04E-07 | 0.04065889 |
| 11 | 65033804 | T | C | 0.686341 | 1.04E-07 | 0.04076198 |
| 11 | 65033897 | A | G | 0.686341 | 1.04E-07 | 0.04076198 |
| 11 | 65033917 | A | C | 0.686341 | 1.04E-07 | 0.04076198 |
| 11 | 65033925 | A | C | 0.686341 | 1.04E-07 | 0.04076198 |
| 11 | 65033961 | G | T | 0.686341 | 1.04E-07 | 0.04076198 |
| 11 | 65033980 | G | T | 0.686341 | 1.04E-07 | 0.04076198 |
| 11 | 65033997 | T | C | 0.686341 | 1.04E-07 | 0.04076198 |
| 11 | 65033998 | G | A | 0.686341 | 1.04E-07 | 0.04076198 |
| 11 | 65034026 | T | C | 0.686341 | 1.04E-07 | 0.04076198 |
| 11 | 65034052 | C | G | 0.686341 | 1.04E-07 | 0.04076198 |
| 11 | 65034071 | T | C | 0.686341 | 1.04E-07 | 0.04076198 |
| 11 | 65034114 | G | A | 0.686341 | 1.04E-07 | 0.04076198 |
| 11 | 65034187 | G | A | 0.686341 | 1.04E-07 | 0.04076198 |
| 11 | 65034221 | C | T | 0.686341 | 1.04E-07 | 0.04076198 |
| 11 | 65034280 | T | C | 0.686341 | 1.04E-07 | 0.04076198 |
| 11 | 64577464 | A | C | 0.745363 | 1.09E-07 | 0.04259486 |
| 11 | 64594843 | G | T | 0.742833 | 1.09E-07 | 0.04269089 |
| 11 | 64594919 | C | T | 0.742833 | 1.09E-07 | 0.04269089 |
| 11 | 64594943 | T | A | 0.742833 | 1.09E-07 | 0.04269089 |
| 11 | 64594949 | T | C | 0.742833 | 1.09E-07 | 0.04269089 |
| 11 | 64593432 | A | G | 0.742833 | 1.10E-07 | 0.043086 |
| 11 | 64593533 | T | C | 0.742833 | 1.10E-07 | 0.043086 |
| 11 | 64593662 | A | G | 0.742833 | 1.10E-07 | 0.043086 |
| 11 | 64593791 | A | G | 0.742833 | 1.10E-07 | 0.043086 |
| 11 | 64681791 | A | G | 0.743676 | 1.11E-07 | 0.04342036 |
| 11 | 64681801 | G | A | 0.743676 | 1.11E-07 | 0.04342036 |
| 11 | 64681858 | C | T | 0.743676 | 1.11E-07 | 0.04342036 |
| 11 | 64235729 | G | T | 0.731029 | 1.14E-07 | 0.04461784 |
| 11 | 64235796 | C | T | 0.731029 | 1.14E-07 | 0.04461784 |
| 11 | 64537050 | T | A | 0.748735 | 1.14E-07 | 0.04481697 |
| 11 | 64537191 | G | C | 0.748735 | 1.14E-07 | 0.04481697 |
| 11 | 64537219 | T | A | 0.748735 | 1.14E-07 | 0.04481697 |
| 11 | 64081392 | A | G | 0.664418 | 1.15E-07 | 0.04499688 |
| 11 | 64082252 | G | C | 0.664418 | 1.15E-07 | 0.04499688 |
| 11 | 64083778 | T | C | 0.664418 | 1.15E-07 | 0.04499688 |
| 11 | 64087522 | T | C | 0.664418 | 1.15E-07 | 0.04499688 |
| 11 | 64087772 | A | G | 0.664418 | 1.15E-07 | 0.04499688 |
| 11 | 64801164 | G | A | 0.744519 | 1.18E-07 | 0.04640369 |
| 11 | 64801396 | A | G | 0.744519 | 1.18E-07 | 0.04640369 |
| 11 | 64131393 | T | C | 0.750422 | 1.19E-07 | 0.04679409 |
| 11 | 64131804 | A | G | 0.750422 | 1.19E-07 | 0.04679409 |
| 11 | 64131867 | A | G | 0.750422 | 1.19E-07 | 0.04679409 |
| 11 | 64132306 | C | A | 0.750422 | 1.19E-07 | 0.04679409 |
| 11 | 64062955 | G | A | 0.663575 | 1.20E-07 | 0.0472139 |
| 11 | 64836884 | G | A | 0.552277 | 1.22E-07 | 0.04765958 |
| 11 | 64770977 | T | C | 0.512648 | 1.22E-07 | 0.04771916 |
| 11 | 64771030 | A | G | 0.512648 | 1.22E-07 | 0.04771916 |
| 11 | 63343490 | A | G | 0.674536 | 1.22E-07 | 0.04784459 |
| 11 | 64068728 | G | C | 0.662732 | 1.23E-07 | 0.04832633 |
| 11 | 64919516 | C | T | 0.573356 | 1.24E-07 | 0.04849919 |
| 11 | 64919624 | C | T | 0.573356 | 1.24E-07 | 0.04849919 |
| 11 | 64919794 | C | A | 0.573356 | 1.24E-07 | 0.04849919 |
| 11 | 64919893 | A | G | 0.573356 | 1.24E-07 | 0.04849919 |
| 11 | 64919984 | T | A | 0.573356 | 1.24E-07 | 0.04849919 |
| 11 | 64919985 | T | A | 0.573356 | 1.24E-07 | 0.04849919 |
| 11 | 64562920 | C | A | 0.745363 | 1.24E-07 | 0.0486893 |
| 11 | 64563047 | A | T | 0.745363 | 1.24E-07 | 0.0486893 |
| 11 | 64612745 | G | A | 0.742833 | 1.24E-07 | 0.04870929 |
| 11 | 64760879 | A | T | 0.744519 | 1.24E-07 | 0.04871203 |
| 11 | 64761124 | T | A | 0.744519 | 1.24E-07 | 0.04871203 |

**Table S12. Significant SNPs for CVDFW**

| Chr | Position | Allele0 | Allele1 | Freq | *P* | Bon |
|---|---|---|---|---|---|---|
| 11 | 65412003 | A | T | 0.920277 | 3.79E-13 | 1.48E-07 |
| 11 | 65398863 | G | A | 0.92461 | 5.80E-12 | 2.28E-06 |
| 11 | 65399615 | T | C | 0.92461 | 5.80E-12 | 2.28E-06 |
| 11 | 65400434 | T | C | 0.92461 | 5.80E-12 | 2.28E-06 |
| 11 | 65400759 | T | C | 0.92461 | 5.80E-12 | 2.28E-06 |
| 11 | 65402208 | G | A | 0.92461 | 5.80E-12 | 2.28E-06 |
| 11 | 65402424 | T | C | 0.92461 | 5.80E-12 | 2.28E-06 |
| 11 | 65403179 | C | T | 0.92461 | 5.80E-12 | 2.28E-06 |
| 11 | 65403383 | T | G | 0.92461 | 5.80E-12 | 2.28E-06 |
| 11 | 65404633 | A | G | 0.92461 | 5.80E-12 | 2.28E-06 |
| 11 | 65404823 | G | C | 0.92461 | 5.80E-12 | 2.28E-06 |
| 11 | 65404929 | G | A | 0.92461 | 5.80E-12 | 2.28E-06 |
| 11 | 65404980 | C | T | 0.92461 | 5.80E-12 | 2.28E-06 |
| 11 | 65405087 | A | G | 0.92461 | 5.80E-12 | 2.28E-06 |
| 11 | 65405090 | A | G | 0.92461 | 5.80E-12 | 2.28E-06 |
| 11 | 65405254 | A | C | 0.92461 | 5.80E-12 | 2.28E-06 |
| 11 | 65405369 | A | G | 0.92461 | 5.80E-12 | 2.28E-06 |
| 11 | 65405580 | C | A | 0.92461 | 5.80E-12 | 2.28E-06 |
| 11 | 65405683 | C | T | 0.92461 | 5.80E-12 | 2.28E-06 |
| 11 | 65405687 | C | A | 0.92461 | 5.80E-12 | 2.28E-06 |
| 11 | 65405734 | C | T | 0.92461 | 5.80E-12 | 2.28E-06 |
| 11 | 65405940 | G | T | 0.92461 | 5.80E-12 | 2.28E-06 |
| 11 | 65405943 | G | A | 0.92461 | 5.80E-12 | 2.28E-06 |
| 11 | 65405944 | T | C | 0.92461 | 5.80E-12 | 2.28E-06 |
| 11 | 65405983 | T | C | 0.92461 | 5.80E-12 | 2.28E-06 |
| 11 | 65405987 | G | A | 0.92461 | 5.80E-12 | 2.28E-06 |
| 11 | 65405999 | C | T | 0.92461 | 5.80E-12 | 2.28E-06 |
| 11 | 65406047 | C | T | 0.92461 | 5.80E-12 | 2.28E-06 |
| 11 | 65406061 | A | C | 0.92461 | 5.80E-12 | 2.28E-06 |
| 11 | 65406104 | C | T | 0.92461 | 5.80E-12 | 2.28E-06 |
| 11 | 65406160 | G | C | 0.92461 | 5.80E-12 | 2.28E-06 |
| 11 | 65406419 | T | C | 0.92461 | 5.80E-12 | 2.28E-06 |
| 11 | 65406470 | A | G | 0.92461 | 5.80E-12 | 2.28E-06 |
| 11 | 65406667 | A | G | 0.92461 | 5.80E-12 | 2.28E-06 |
| 11 | 65406674 | T | C | 0.92461 | 5.80E-12 | 2.28E-06 |
| 11 | 65406699 | T | C | 0.92461 | 5.80E-12 | 2.28E-06 |
| 11 | 65407772 | A | G | 0.92461 | 5.80E-12 | 2.28E-06 |
| 11 | 65407808 | C | T | 0.92461 | 5.80E-12 | 2.28E-06 |
| 11 | 65407947 | C | T | 0.92461 | 5.80E-12 | 2.28E-06 |
| 11 | 65408288 | G | C | 0.92461 | 5.80E-12 | 2.28E-06 |
| 11 | 65408676 | A | G | 0.92461 | 5.80E-12 | 2.28E-06 |
| 11 | 65408874 | T | A | 0.92461 | 5.80E-12 | 2.28E-06 |
| 11 | 65408986 | C | T | 0.92461 | 5.80E-12 | 2.28E-06 |
| 11 | 65408989 | C | T | 0.92461 | 5.80E-12 | 2.28E-06 |
| 11 | 65409054 | G | A | 0.92461 | 5.80E-12 | 2.28E-06 |
| 11 | 65409059 | T | G | 0.92461 | 5.80E-12 | 2.28E-06 |
| 11 | 65409211 | C | T | 0.92461 | 5.80E-12 | 2.28E-06 |
| 11 | 65409471 | T | C | 0.92461 | 5.80E-12 | 2.28E-06 |
| 11 | 65409505 | T | G | 0.92461 | 5.80E-12 | 2.28E-06 |
| 11 | 65409672 | G | A | 0.92461 | 5.80E-12 | 2.28E-06 |
| 11 | 65410049 | A | G | 0.92461 | 5.80E-12 | 2.28E-06 |
| 11 | 65410310 | G | C | 0.92461 | 5.80E-12 | 2.28E-06 |
| 11 | 65410470 | G | A | 0.92461 | 5.80E-12 | 2.28E-06 |
| 11 | 65410524 | C | T | 0.92461 | 5.80E-12 | 2.28E-06 |
| 11 | 65410536 | A | T | 0.92461 | 5.80E-12 | 2.28E-06 |
| 11 | 65410609 | A | G | 0.92461 | 5.80E-12 | 2.28E-06 |
| 11 | 65410617 | T | G | 0.92461 | 5.80E-12 | 2.28E-06 |
| 11 | 65410629 | A | G | 0.92461 | 5.80E-12 | 2.28E-06 |
| 11 | 65410754 | C | A | 0.92461 | 5.80E-12 | 2.28E-06 |
| 11 | 65410830 | G | T | 0.92461 | 5.80E-12 | 2.28E-06 |
| 11 | 65410860 | C | T | 0.92461 | 5.80E-12 | 2.28E-06 |
| 11 | 65410986 | A | T | 0.92461 | 5.80E-12 | 2.28E-06 |
| 11 | 65410994 | C | T | 0.92461 | 5.80E-12 | 2.28E-06 |
| 11 | 65411384 | T | C | 0.92461 | 5.80E-12 | 2.28E-06 |
| 11 | 65411394 | T | C | 0.92461 | 5.80E-12 | 2.28E-06 |

| | | | | | | |
|---|---|---|---|---|---|---|
| 11 | 65411456 | A | G | 0.92461 | 5.80E-12 | 2.28E-06 |
| 11 | 65411628 | A | C | 0.92461 | 5.80E-12 | 2.28E-06 |
| 11 | 65411638 | G | T | 0.92461 | 5.80E-12 | 2.28E-06 |
| 11 | 65411647 | A | T | 0.92461 | 5.80E-12 | 2.28E-06 |
| 11 | 65411654 | T | A | 0.92461 | 5.80E-12 | 2.28E-06 |
| 11 | 65411709 | T | A | 0.92461 | 5.80E-12 | 2.28E-06 |
| 11 | 65411720 | T | C | 0.92461 | 5.80E-12 | 2.28E-06 |
| 11 | 65411745 | C | T | 0.92461 | 5.80E-12 | 2.28E-06 |
| 11 | 65411818 | T | A | 0.92461 | 5.80E-12 | 2.28E-06 |
| 11 | 65411944 | G | T | 0.92461 | 5.80E-12 | 2.28E-06 |
| 11 | 65411980 | G | A | 0.92461 | 5.80E-12 | 2.28E-06 |
| 11 | 65412034 | G | A | 0.92461 | 5.80E-12 | 2.28E-06 |
| 11 | 65412035 | G | A | 0.92461 | 5.80E-12 | 2.28E-06 |
| 11 | 65412043 | T | C | 0.92461 | 5.80E-12 | 2.28E-06 |
| 11 | 65412196 | C | G | 0.92461 | 5.80E-12 | 2.28E-06 |
| 11 | 65412242 | T | C | 0.92461 | 5.80E-12 | 2.28E-06 |
| 11 | 65412265 | C | T | 0.92461 | 5.80E-12 | 2.28E-06 |
| 11 | 65412267 | G | A | 0.92461 | 5.80E-12 | 2.28E-06 |
| 11 | 65412276 | C | T | 0.92461 | 5.80E-12 | 2.28E-06 |
| 11 | 65412279 | G | A | 0.92461 | 5.80E-12 | 2.28E-06 |
| 11 | 65412460 | T | C | 0.92461 | 5.80E-12 | 2.28E-06 |
| 11 | 65412513 | C | T | 0.92461 | 5.80E-12 | 2.28E-06 |
| 11 | 65412534 | T | C | 0.92461 | 5.80E-12 | 2.28E-06 |
| 11 | 65412598 | G | C | 0.92461 | 5.80E-12 | 2.28E-06 |
| 11 | 65412680 | C | T | 0.92461 | 5.80E-12 | 2.28E-06 |
| 11 | 65412714 | C | T | 0.92461 | 5.80E-12 | 2.28E-06 |
| 11 | 65412825 | A | G | 0.92461 | 5.80E-12 | 2.28E-06 |
| 11 | 65412984 | T | C | 0.92461 | 5.80E-12 | 2.28E-06 |
| 11 | 65413017 | C | A | 0.92461 | 5.80E-12 | 2.28E-06 |
| 11 | 65413099 | C | T | 0.92461 | 5.80E-12 | 2.28E-06 |
| 11 | 65413199 | C | T | 0.92461 | 5.80E-12 | 2.28E-06 |
| 11 | 65413345 | A | C | 0.92461 | 5.80E-12 | 2.28E-06 |
| 11 | 65413409 | G | T | 0.92461 | 5.80E-12 | 2.28E-06 |
| 11 | 65413452 | A | C | 0.92461 | 5.80E-12 | 2.28E-06 |
| 11 | 65413466 | C | T | 0.92461 | 5.80E-12 | 2.28E-06 |
| 11 | 65413467 | A | G | 0.92461 | 5.80E-12 | 2.28E-06 |
| 11 | 65413639 | G | A | 0.92461 | 5.80E-12 | 2.28E-06 |
| 11 | 65413658 | A | T | 0.92461 | 5.80E-12 | 2.28E-06 |
| 11 | 65413666 | T | C | 0.92461 | 5.80E-12 | 2.28E-06 |
| 11 | 65413677 | A | T | 0.92461 | 5.80E-12 | 2.28E-06 |
| 11 | 65413819 | A | T | 0.92461 | 5.80E-12 | 2.28E-06 |
| 11 | 65413851 | G | A | 0.92461 | 5.80E-12 | 2.28E-06 |
| 11 | 65413912 | T | A | 0.92461 | 5.80E-12 | 2.28E-06 |
| 11 | 65413962 | G | A | 0.92461 | 5.80E-12 | 2.28E-06 |
| 11 | 65414061 | C | T | 0.92461 | 5.80E-12 | 2.28E-06 |
| 11 | 65414128 | T | G | 0.92461 | 5.80E-12 | 2.28E-06 |
| 11 | 65414132 | C | T | 0.92461 | 5.80E-12 | 2.28E-06 |
| 11 | 65414161 | A | G | 0.92461 | 5.80E-12 | 2.28E-06 |
| 11 | 65414182 | C | T | 0.92461 | 5.80E-12 | 2.28E-06 |
| 11 | 65414209 | G | A | 0.92461 | 5.80E-12 | 2.28E-06 |
| 11 | 65414216 | A | C | 0.92461 | 5.80E-12 | 2.28E-06 |
| 11 | 65414220 | C | G | 0.92461 | 5.80E-12 | 2.28E-06 |
| 11 | 65414245 | A | G | 0.92461 | 5.80E-12 | 2.28E-06 |
| 11 | 65414302 | C | A | 0.92461 | 5.80E-12 | 2.28E-06 |
| 11 | 65414353 | A | G | 0.92461 | 5.80E-12 | 2.28E-06 |
| 11 | 65414381 | T | C | 0.92461 | 5.80E-12 | 2.28E-06 |
| 11 | 65414432 | T | C | 0.92461 | 5.80E-12 | 2.28E-06 |
| 11 | 65414552 | A | G | 0.92461 | 5.80E-12 | 2.28E-06 |
| 11 | 65414676 | G | A | 0.92461 | 5.80E-12 | 2.28E-06 |
| 11 | 65414694 | T | C | 0.92461 | 5.80E-12 | 2.28E-06 |
| 11 | 65414724 | T | C | 0.92461 | 5.80E-12 | 2.28E-06 |
| 11 | 65414731 | C | T | 0.92461 | 5.80E-12 | 2.28E-06 |
| 11 | 65414733 | C | G | 0.92461 | 5.80E-12 | 2.28E-06 |
| 11 | 65414745 | T | C | 0.92461 | 5.80E-12 | 2.28E-06 |
| 11 | 65414746 | G | A | 0.92461 | 5.80E-12 | 2.28E-06 |
| 11 | 65414758 | T | C | 0.92461 | 5.80E-12 | 2.28E-06 |
| 11 | 65414759 | G | A | 0.92461 | 5.80E-12 | 2.28E-06 |

| | | | | | | |
|---|---|---|---|---|---|---|
| 11 | 65414768 | A | G | 0.92461 | 5.80E-12 | 2.28E-06 |
| 11 | 65414770 | T | C | 0.92461 | 5.80E-12 | 2.28E-06 |
| 11 | 65414771 | G | A | 0.92461 | 5.80E-12 | 2.28E-06 |
| 11 | 65414804 | T | G | 0.92461 | 5.80E-12 | 2.28E-06 |
| 11 | 65414820 | C | T | 0.92461 | 5.80E-12 | 2.28E-06 |
| 11 | 65414821 | A | G | 0.92461 | 5.80E-12 | 2.28E-06 |
| 11 | 65414950 | A | G | 0.92461 | 5.80E-12 | 2.28E-06 |
| 11 | 65414962 | C | T | 0.92461 | 5.80E-12 | 2.28E-06 |
| 11 | 65414979 | G | T | 0.92461 | 5.80E-12 | 2.28E-06 |
| 11 | 65414995 | G | C | 0.92461 | 5.80E-12 | 2.28E-06 |
| 11 | 65415163 | A | T | 0.92461 | 5.80E-12 | 2.28E-06 |
| 11 | 65415317 | T | C | 0.92461 | 5.80E-12 | 2.28E-06 |
| 11 | 65415393 | T | A | 0.92461 | 5.80E-12 | 2.28E-06 |
| 11 | 65415832 | C | T | 0.92461 | 5.80E-12 | 2.28E-06 |
| 11 | 65416597 | A | C | 0.92461 | 5.80E-12 | 2.28E-06 |
| 11 | 65416605 | A | G | 0.92461 | 5.80E-12 | 2.28E-06 |
| 11 | 65416795 | G | C | 0.92461 | 5.80E-12 | 2.28E-06 |
| 11 | 65417095 | T | C | 0.92461 | 5.80E-12 | 2.28E-06 |
| 11 | 65417178 | G | A | 0.92461 | 5.80E-12 | 2.28E-06 |
| 11 | 65417201 | G | C | 0.92461 | 5.80E-12 | 2.28E-06 |
| 11 | 65417275 | A | G | 0.92461 | 5.80E-12 | 2.28E-06 |
| 11 | 65417292 | A | T | 0.92461 | 5.80E-12 | 2.28E-06 |
| 11 | 65417410 | G | A | 0.92461 | 5.80E-12 | 2.28E-06 |
| 11 | 65417426 | T | C | 0.92461 | 5.80E-12 | 2.28E-06 |
| 11 | 65417527 | G | C | 0.92461 | 5.80E-12 | 2.28E-06 |
| 11 | 65417538 | C | T | 0.92461 | 5.80E-12 | 2.28E-06 |
| 11 | 65417607 | C | T | 0.92461 | 5.80E-12 | 2.28E-06 |
| 11 | 65417723 | C | A | 0.92461 | 5.80E-12 | 2.28E-06 |
| 11 | 65432659 | C | A | 0.92461 | 5.80E-12 | 2.28E-06 |
| 11 | 64948615 | G | C | 0.928943 | 1.25E-11 | 4.90E-06 |
| 11 | 64950500 | T | C | 0.928943 | 1.25E-11 | 4.90E-06 |
| 11 | 64950776 | G | A | 0.928943 | 1.25E-11 | 4.90E-06 |
| 11 | 64950793 | T | A | 0.928943 | 1.25E-11 | 4.90E-06 |
| 11 | 64950794 | T | C | 0.928943 | 1.25E-11 | 4.90E-06 |
| 11 | 64951137 | A | G | 0.928943 | 1.25E-11 | 4.90E-06 |
| 11 | 64951174 | G | C | 0.928943 | 1.25E-11 | 4.90E-06 |
| 11 | 64951364 | T | C | 0.928943 | 1.25E-11 | 4.90E-06 |
| 11 | 64951973 | A | T | 0.928943 | 1.25E-11 | 4.90E-06 |
| 11 | 64952273 | G | A | 0.928943 | 1.25E-11 | 4.90E-06 |
| 11 | 64270630 | C | T | 0.926343 | 1.43E-11 | 5.60E-06 |
| 11 | 64270744 | T | C | 0.926343 | 1.43E-11 | 5.60E-06 |
| 11 | 65417024 | G | A | 0.923744 | 1.96E-11 | 7.68E-06 |
| 11 | 64234283 | C | A | 0.926343 | 2.82E-11 | 1.10E-05 |
| 11 | 64944841 | A | T | 0.929809 | 5.33E-11 | 2.09E-05 |
| 11 | 64270037 | G | C | 0.925477 | 5.48E-11 | 2.15E-05 |
| 11 | 64349603 | C | T | 0.925477 | 5.48E-11 | 2.15E-05 |
| 11 | 64350722 | T | C | 0.925477 | 5.48E-11 | 2.15E-05 |
| 11 | 64351227 | T | G | 0.925477 | 5.48E-11 | 2.15E-05 |
| 11 | 64361938 | T | C | 0.925477 | 5.48E-11 | 2.15E-05 |
| 11 | 64365353 | A | G | 0.925477 | 5.48E-11 | 2.15E-05 |
| 11 | 64461388 | A | G | 0.925477 | 5.48E-11 | 2.15E-05 |
| 11 | 64468285 | C | T | 0.925477 | 5.48E-11 | 2.15E-05 |
| 11 | 65417185 | T | C | 0.92201 | 6.61E-11 | 2.59E-05 |
| 11 | 65417210 | C | A | 0.92201 | 6.61E-11 | 2.59E-05 |
| 11 | 64269214 | C | A | 0.926343 | 7.67E-11 | 3.01E-05 |
| 11 | 64224299 | T | C | 0.925477 | 8.52E-11 | 3.34E-05 |
| 11 | 64483464 | A | G | 0.92721 | 1.42E-10 | 5.58E-05 |
| 11 | 64220895 | T | C | 0.92461 | 1.65E-10 | 6.45E-05 |
| 11 | 64395863 | A | C | 0.930676 | 2.46E-10 | 9.65E-05 |
| 11 | 64399779 | T | C | 0.930676 | 2.46E-10 | 9.65E-05 |
| 11 | 63237429 | C | T | 0.92721 | 2.70E-10 | 0.00010601 |
| 11 | 63255501 | C | T | 0.92721 | 2.70E-10 | 0.00010601 |
| 11 | 64366972 | A | G | 0.926343 | 3.44E-10 | 0.00013482 |
| 11 | 65990344 | T | C | 0.931542 | 3.60E-10 | 0.00014106 |
| 11 | 65993082 | A | G | 0.933276 | 4.34E-10 | 0.00016995 |
| 11 | 65993312 | A | G | 0.933276 | 4.34E-10 | 0.00016995 |
| 11 | 65993521 | T | C | 0.933276 | 4.34E-10 | 0.00016995 |

| | | | | | | |
|---|---|---|---|---|---|---|
| 11 | 65993633 | T | G | 0.933276 | 4.34E-10 | 0.00016995 |
| 11 | 64242653 | T | C | 0.928076 | 5.47E-10 | 0.00021448 |
| 11 | 64242993 | G | A | 0.928943 | 6.96E-10 | 0.00027263 |
| 11 | 65451349 | T | G | 0.932409 | 7.85E-10 | 0.00030758 |
| 11 | 65451381 | C | A | 0.932409 | 7.85E-10 | 0.00030758 |
| 11 | 65451780 | T | C | 0.932409 | 7.85E-10 | 0.00030758 |
| 11 | 65451868 | A | G | 0.932409 | 7.85E-10 | 0.00030758 |
| 11 | 65452088 | G | A | 0.932409 | 7.85E-10 | 0.00030758 |
| 11 | 65452090 | T | C | 0.932409 | 7.85E-10 | 0.00030758 |
| 11 | 65452186 | C | T | 0.932409 | 7.85E-10 | 0.00030758 |
| 11 | 65452273 | T | C | 0.932409 | 7.85E-10 | 0.00030758 |
| 11 | 65452436 | T | G | 0.932409 | 7.85E-10 | 0.00030758 |
| 11 | 65452488 | C | T | 0.932409 | 7.85E-10 | 0.00030758 |
| 11 | 65452978 | T | C | 0.932409 | 7.85E-10 | 0.00030758 |
| 11 | 65453191 | C | G | 0.932409 | 7.85E-10 | 0.00030758 |
| 11 | 65453312 | A | G | 0.932409 | 7.85E-10 | 0.00030758 |
| 11 | 65453327 | T | A | 0.932409 | 7.85E-10 | 0.00030758 |
| 11 | 65453351 | T | C | 0.932409 | 7.85E-10 | 0.00030758 |
| 11 | 65453374 | G | A | 0.932409 | 7.85E-10 | 0.00030758 |
| 11 | 65453448 | G | A | 0.932409 | 7.85E-10 | 0.00030758 |
| 11 | 65453464 | T | C | 0.932409 | 7.85E-10 | 0.00030758 |
| 11 | 65453608 | T | C | 0.932409 | 7.85E-10 | 0.00030758 |
| 11 | 65453650 | T | C | 0.932409 | 7.85E-10 | 0.00030758 |
| 11 | 65454294 | T | G | 0.932409 | 7.85E-10 | 0.00030758 |
| 11 | 65454298 | C | T | 0.932409 | 7.85E-10 | 0.00030758 |
| 11 | 65454336 | A | G | 0.932409 | 7.85E-10 | 0.00030758 |
| 11 | 65454560 | A | G | 0.932409 | 7.85E-10 | 0.00030758 |
| 11 | 65454780 | T | C | 0.932409 | 7.85E-10 | 0.00030758 |
| 11 | 65454871 | G | T | 0.932409 | 7.85E-10 | 0.00030758 |
| 11 | 65455133 | G | T | 0.932409 | 7.85E-10 | 0.00030758 |
| 11 | 65455213 | A | C | 0.932409 | 7.85E-10 | 0.00030758 |
| 11 | 65455236 | C | T | 0.932409 | 7.85E-10 | 0.00030758 |
| 11 | 65455238 | C | T | 0.932409 | 7.85E-10 | 0.00030758 |
| 11 | 65455275 | G | A | 0.932409 | 7.85E-10 | 0.00030758 |
| 11 | 65455359 | C | T | 0.932409 | 7.85E-10 | 0.00030758 |
| 11 | 65455360 | A | G | 0.932409 | 7.85E-10 | 0.00030758 |
| 11 | 65455388 | A | G | 0.932409 | 7.85E-10 | 0.00030758 |
| 11 | 65455483 | C | T | 0.932409 | 7.85E-10 | 0.00030758 |
| 11 | 65455598 | T | C | 0.932409 | 7.85E-10 | 0.00030758 |
| 11 | 65455605 | G | T | 0.932409 | 7.85E-10 | 0.00030758 |
| 11 | 65455634 | A | G | 0.932409 | 7.85E-10 | 0.00030758 |
| 11 | 65455696 | A | T | 0.932409 | 7.85E-10 | 0.00030758 |
| 11 | 65455796 | T | A | 0.932409 | 7.85E-10 | 0.00030758 |
| 11 | 65455808 | T | C | 0.932409 | 7.85E-10 | 0.00030758 |
| 11 | 65455890 | C | G | 0.932409 | 7.85E-10 | 0.00030758 |
| 11 | 65456009 | G | C | 0.932409 | 7.85E-10 | 0.00030758 |
| 11 | 65456365 | T | C | 0.932409 | 7.85E-10 | 0.00030758 |
| 11 | 65456372 | T | C | 0.932409 | 7.85E-10 | 0.00030758 |
| 11 | 65456426 | A | G | 0.932409 | 7.85E-10 | 0.00030758 |
| 11 | 65456645 | T | A | 0.932409 | 7.85E-10 | 0.00030758 |
| 11 | 65456708 | A | G | 0.932409 | 7.85E-10 | 0.00030758 |
| 11 | 65456718 | A | T | 0.932409 | 7.85E-10 | 0.00030758 |
| 11 | 65456775 | G | A | 0.932409 | 7.85E-10 | 0.00030758 |
| 11 | 65456811 | G | A | 0.932409 | 7.85E-10 | 0.00030758 |
| 11 | 65457384 | T | G | 0.932409 | 7.85E-10 | 0.00030758 |
| 11 | 65457726 | T | C | 0.932409 | 7.85E-10 | 0.00030758 |
| 11 | 65458608 | G | A | 0.932409 | 7.85E-10 | 0.00030758 |
| 11 | 65458611 | G | A | 0.932409 | 7.85E-10 | 0.00030758 |
| 11 | 65458618 | G | A | 0.932409 | 7.85E-10 | 0.00030758 |
| 11 | 65458651 | A | G | 0.932409 | 7.85E-10 | 0.00030758 |
| 11 | 65458656 | C | T | 0.932409 | 7.85E-10 | 0.00030758 |
| 11 | 65458789 | G | A | 0.932409 | 7.85E-10 | 0.00030758 |
| 11 | 65458805 | T | G | 0.932409 | 7.85E-10 | 0.00030758 |
| 11 | 65458940 | T | C | 0.932409 | 7.85E-10 | 0.00030758 |
| 11 | 65459212 | C | A | 0.932409 | 7.85E-10 | 0.00030758 |
| 11 | 65460225 | C | A | 0.932409 | 7.85E-10 | 0.00030758 |
| 11 | 65460287 | C | T | 0.932409 | 7.85E-10 | 0.00030758 |

| | | | | | | |
|---|---|---|---|---|---|---|
| 11 | 65460664 | C | G | 0.932409 | 7.85E-10 | 0.00030758 |
| 11 | 65461036 | C | T | 0.932409 | 7.85E-10 | 0.00030758 |
| 11 | 65461248 | A | G | 0.932409 | 7.85E-10 | 0.00030758 |
| 11 | 65461284 | T | C | 0.932409 | 7.85E-10 | 0.00030758 |
| 11 | 65461327 | A | G | 0.932409 | 7.85E-10 | 0.00030758 |
| 11 | 65461328 | G | T | 0.932409 | 7.85E-10 | 0.00030758 |
| 11 | 65461419 | A | G | 0.932409 | 7.85E-10 | 0.00030758 |
| 11 | 65462928 | T | C | 0.932409 | 7.85E-10 | 0.00030758 |
| 11 | 65463140 | G | T | 0.932409 | 7.85E-10 | 0.00030758 |
| 11 | 65463520 | A | G | 0.932409 | 7.85E-10 | 0.00030758 |
| 11 | 65463905 | T | C | 0.932409 | 7.85E-10 | 0.00030758 |
| 11 | 65463978 | C | G | 0.932409 | 7.85E-10 | 0.00030758 |
| 11 | 65464150 | A | T | 0.932409 | 7.85E-10 | 0.00030758 |
| 11 | 65464177 | G | T | 0.932409 | 7.85E-10 | 0.00030758 |
| 11 | 65464203 | C | T | 0.932409 | 7.85E-10 | 0.00030758 |
| 11 | 65464253 | A | G | 0.932409 | 7.85E-10 | 0.00030758 |
| 11 | 65464321 | C | A | 0.932409 | 7.85E-10 | 0.00030758 |
| 11 | 65464329 | A | G | 0.932409 | 7.85E-10 | 0.00030758 |
| 11 | 65465313 | C | T | 0.932409 | 7.85E-10 | 0.00030758 |
| 11 | 65472056 | C | G | 0.932409 | 7.85E-10 | 0.00030758 |
| 11 | 65488082 | A | G | 0.932409 | 7.85E-10 | 0.00030758 |
| 11 | 65489950 | C | T | 0.932409 | 7.85E-10 | 0.00030758 |
| 11 | 65490161 | C | T | 0.932409 | 7.85E-10 | 0.00030758 |
| 11 | 65490909 | A | C | 0.932409 | 7.85E-10 | 0.00030758 |
| 11 | 65490910 | T | C | 0.932409 | 7.85E-10 | 0.00030758 |
| 11 | 65491251 | T | C | 0.932409 | 7.85E-10 | 0.00030758 |
| 11 | 65492735 | A | G | 0.932409 | 7.85E-10 | 0.00030758 |
| 11 | 65493232 | C | T | 0.932409 | 7.85E-10 | 0.00030758 |
| 11 | 65493269 | G | A | 0.932409 | 7.85E-10 | 0.00030758 |
| 11 | 65493278 | G | A | 0.932409 | 7.85E-10 | 0.00030758 |
| 11 | 65493407 | A | G | 0.932409 | 7.85E-10 | 0.00030758 |
| 11 | 65494013 | T | G | 0.932409 | 7.85E-10 | 0.00030758 |
| 11 | 65494638 | G | A | 0.932409 | 7.85E-10 | 0.00030758 |
| 11 | 65495101 | T | G | 0.932409 | 7.85E-10 | 0.00030758 |
| 11 | 65496569 | T | G | 0.932409 | 7.85E-10 | 0.00030758 |
| 11 | 65497210 | G | A | 0.932409 | 7.85E-10 | 0.00030758 |
| 11 | 65497360 | C | T | 0.932409 | 7.85E-10 | 0.00030758 |
| 11 | 65497398 | T | C | 0.932409 | 7.85E-10 | 0.00030758 |
| 11 | 65497641 | G | T | 0.932409 | 7.85E-10 | 0.00030758 |
| 11 | 65499208 | A | T | 0.932409 | 7.85E-10 | 0.00030758 |
| 11 | 65500228 | C | A | 0.932409 | 7.85E-10 | 0.00030758 |
| 11 | 65500300 | G | A | 0.932409 | 7.85E-10 | 0.00030758 |
| 11 | 65500401 | T | G | 0.932409 | 7.85E-10 | 0.00030758 |
| 11 | 65500664 | A | G | 0.932409 | 7.85E-10 | 0.00030758 |
| 11 | 65500906 | C | T | 0.932409 | 7.85E-10 | 0.00030758 |
| 11 | 65500917 | T | C | 0.932409 | 7.85E-10 | 0.00030758 |
| 11 | 65500981 | C | A | 0.932409 | 7.85E-10 | 0.00030758 |
| 11 | 65501155 | C | T | 0.932409 | 7.85E-10 | 0.00030758 |
| 11 | 65501159 | G | A | 0.932409 | 7.85E-10 | 0.00030758 |
| 11 | 65501185 | T | C | 0.932409 | 7.85E-10 | 0.00030758 |
| 11 | 65501250 | T | G | 0.932409 | 7.85E-10 | 0.00030758 |
| 11 | 65501327 | T | C | 0.932409 | 7.85E-10 | 0.00030758 |
| 11 | 65501336 | A | G | 0.932409 | 7.85E-10 | 0.00030758 |
| 11 | 65501339 | T | C | 0.932409 | 7.85E-10 | 0.00030758 |
| 11 | 65501417 | C | T | 0.932409 | 7.85E-10 | 0.00030758 |
| 11 | 65501527 | G | A | 0.932409 | 7.85E-10 | 0.00030758 |
| 11 | 65502586 | G | A | 0.932409 | 7.85E-10 | 0.00030758 |
| 11 | 65503370 | T | C | 0.932409 | 7.85E-10 | 0.00030758 |
| 11 | 65503943 | C | T | 0.932409 | 7.85E-10 | 0.00030758 |
| 11 | 65503965 | G | A | 0.932409 | 7.85E-10 | 0.00030758 |
| 11 | 65505228 | T | C | 0.932409 | 7.85E-10 | 0.00030758 |
| 11 | 65505261 | T | C | 0.932409 | 7.85E-10 | 0.00030758 |
| 11 | 65505272 | C | T | 0.932409 | 7.85E-10 | 0.00030758 |
| 11 | 65505499 | T | C | 0.932409 | 7.85E-10 | 0.00030758 |
| 11 | 65505909 | C | T | 0.932409 | 7.85E-10 | 0.00030758 |
| 11 | 65506074 | A | G | 0.932409 | 7.85E-10 | 0.00030758 |
| 11 | 65506141 | G | A | 0.932409 | 7.85E-10 | 0.00030758 |

| | | | | | | |
|---|---|---|---|---|---|---|
| 11 | 65506268 | A | G | 0.932409 | 7.85E-10 | 0.00030758 |
| 11 | 65506288 | G | A | 0.932409 | 7.85E-10 | 0.00030758 |
| 11 | 65506948 | T | C | 0.932409 | 7.85E-10 | 0.00030758 |
| 11 | 65507112 | C | T | 0.932409 | 7.85E-10 | 0.00030758 |
| 11 | 65507148 | G | C | 0.932409 | 7.85E-10 | 0.00030758 |
| 11 | 65507376 | A | G | 0.932409 | 7.85E-10 | 0.00030758 |
| 11 | 65507810 | C | A | 0.932409 | 7.85E-10 | 0.00030758 |
| 11 | 65507816 | C | A | 0.932409 | 7.85E-10 | 0.00030758 |
| 11 | 65507818 | T | A | 0.932409 | 7.85E-10 | 0.00030758 |
| 11 | 65507822 | C | A | 0.932409 | 7.85E-10 | 0.00030758 |
| 11 | 65507828 | C | A | 0.932409 | 7.85E-10 | 0.00030758 |
| 11 | 65507834 | C | A | 0.932409 | 7.85E-10 | 0.00030758 |
| 11 | 65507842 | T | A | 0.932409 | 7.85E-10 | 0.00030758 |
| 11 | 65507939 | G | A | 0.932409 | 7.85E-10 | 0.00030758 |
| 11 | 65507943 | G | A | 0.932409 | 7.85E-10 | 0.00030758 |
| 11 | 65507945 | G | A | 0.932409 | 7.85E-10 | 0.00030758 |
| 11 | 65507952 | G | T | 0.932409 | 7.85E-10 | 0.00030758 |
| 11 | 65507967 | T | C | 0.932409 | 7.85E-10 | 0.00030758 |
| 11 | 65507977 | T | C | 0.932409 | 7.85E-10 | 0.00030758 |
| 11 | 65508000 | T | C | 0.932409 | 7.85E-10 | 0.00030758 |
| 11 | 65508185 | A | T | 0.932409 | 7.85E-10 | 0.00030758 |
| 11 | 65509629 | A | G | 0.932409 | 7.85E-10 | 0.00030758 |
| 11 | 65509687 | C | T | 0.932409 | 7.85E-10 | 0.00030758 |
| 11 | 65509831 | T | C | 0.932409 | 7.85E-10 | 0.00030758 |
| 11 | 65509880 | A | G | 0.932409 | 7.85E-10 | 0.00030758 |
| 11 | 65509986 | A | C | 0.932409 | 7.85E-10 | 0.00030758 |
| 11 | 65510812 | T | G | 0.932409 | 7.85E-10 | 0.00030758 |
| 11 | 65511339 | T | A | 0.932409 | 7.85E-10 | 0.00030758 |
| 11 | 65511550 | A | G | 0.932409 | 7.85E-10 | 0.00030758 |
| 11 | 65511971 | T | A | 0.932409 | 7.85E-10 | 0.00030758 |
| 11 | 65512710 | G | A | 0.932409 | 7.85E-10 | 0.00030758 |
| 11 | 65512786 | G | A | 0.932409 | 7.85E-10 | 0.00030758 |
| 11 | 65513021 | C | G | 0.932409 | 7.85E-10 | 0.00030758 |
| 11 | 65513833 | C | T | 0.932409 | 7.85E-10 | 0.00030758 |
| 11 | 65514174 | A | C | 0.932409 | 7.85E-10 | 0.00030758 |
| 11 | 65514539 | A | G | 0.932409 | 7.85E-10 | 0.00030758 |
| 11 | 65514935 | C | T | 0.932409 | 7.85E-10 | 0.00030758 |
| 11 | 65515200 | T | C | 0.932409 | 7.85E-10 | 0.00030758 |
| 11 | 65515201 | C | A | 0.932409 | 7.85E-10 | 0.00030758 |
| 11 | 65515216 | A | G | 0.932409 | 7.85E-10 | 0.00030758 |
| 11 | 65515916 | G | A | 0.932409 | 7.85E-10 | 0.00030758 |
| 11 | 65516791 | T | C | 0.932409 | 7.85E-10 | 0.00030758 |
| 11 | 65516822 | C | T | 0.932409 | 7.85E-10 | 0.00030758 |
| 11 | 65516866 | T | C | 0.932409 | 7.85E-10 | 0.00030758 |
| 11 | 65518014 | G | A | 0.932409 | 7.85E-10 | 0.00030758 |
| 11 | 65519665 | A | T | 0.932409 | 7.85E-10 | 0.00030758 |
| 11 | 65519681 | A | G | 0.932409 | 7.85E-10 | 0.00030758 |
| 11 | 65519724 | T | G | 0.932409 | 7.85E-10 | 0.00030758 |
| 11 | 65520459 | A | G | 0.932409 | 7.85E-10 | 0.00030758 |
| 11 | 65520690 | A | G | 0.932409 | 7.85E-10 | 0.00030758 |
| 11 | 65521027 | C | T | 0.932409 | 7.85E-10 | 0.00030758 |
| 11 | 65521856 | G | A | 0.932409 | 7.85E-10 | 0.00030758 |
| 11 | 65522305 | T | C | 0.932409 | 7.85E-10 | 0.00030758 |
| 11 | 65523091 | G | A | 0.932409 | 7.85E-10 | 0.00030758 |
| 11 | 65524906 | C | T | 0.932409 | 7.85E-10 | 0.00030758 |
| 11 | 65525007 | C | T | 0.932409 | 7.85E-10 | 0.00030758 |
| 11 | 65525910 | G | T | 0.932409 | 7.85E-10 | 0.00030758 |
| 11 | 65526124 | T | C | 0.932409 | 7.85E-10 | 0.00030758 |
| 11 | 65526463 | C | T | 0.932409 | 7.85E-10 | 0.00030758 |
| 11 | 65527368 | G | C | 0.932409 | 7.85E-10 | 0.00030758 |
| 11 | 65528562 | T | G | 0.932409 | 7.85E-10 | 0.00030758 |
| 11 | 65528569 | A | G | 0.932409 | 7.85E-10 | 0.00030758 |
| 11 | 65530514 | A | G | 0.932409 | 7.85E-10 | 0.00030758 |
| 11 | 65530873 | C | A | 0.932409 | 7.85E-10 | 0.00030758 |
| 11 | 65531184 | T | C | 0.932409 | 7.85E-10 | 0.00030758 |
| 11 | 65531489 | A | G | 0.932409 | 7.85E-10 | 0.00030758 |
| 11 | 65531868 | T | C | 0.932409 | 7.85E-10 | 0.00030758 |

| | | | | | | |
|---|---|---|---|---|---|---|
| 11 | 65541356 | T | C | 0.932409 | 7.85E-10 | 0.00030758 |
| 11 | 65545215 | A | G | 0.932409 | 7.85E-10 | 0.00030758 |
| 11 | 65547854 | A | C | 0.932409 | 7.85E-10 | 0.00030758 |
| 11 | 65550124 | T | C | 0.932409 | 7.85E-10 | 0.00030758 |
| 11 | 65550191 | A | T | 0.932409 | 7.85E-10 | 0.00030758 |
| 11 | 65551525 | G | A | 0.932409 | 7.85E-10 | 0.00030758 |
| 11 | 65551627 | A | G | 0.932409 | 7.85E-10 | 0.00030758 |
| 11 | 65552378 | G | A | 0.932409 | 7.85E-10 | 0.00030758 |
| 11 | 65560188 | T | C | 0.932409 | 7.85E-10 | 0.00030758 |
| 11 | 65561476 | C | T | 0.932409 | 7.85E-10 | 0.00030758 |
| 11 | 65573200 | C | T | 0.932409 | 7.85E-10 | 0.00030758 |
| 11 | 65576340 | C | T | 0.932409 | 7.85E-10 | 0.00030758 |
| 11 | 65576434 | G | A | 0.932409 | 7.85E-10 | 0.00030758 |
| 11 | 65577062 | A | G | 0.932409 | 7.85E-10 | 0.00030758 |
| 11 | 65578244 | A | T | 0.932409 | 7.85E-10 | 0.00030758 |
| 11 | 65578943 | T | C | 0.932409 | 7.85E-10 | 0.00030758 |
| 11 | 65579413 | A | G | 0.932409 | 7.85E-10 | 0.00030758 |
| 11 | 65579780 | A | G | 0.932409 | 7.85E-10 | 0.00030758 |
| 11 | 65580262 | T | A | 0.932409 | 7.85E-10 | 0.00030758 |
| 11 | 65990733 | T | G | 0.932409 | 7.85E-10 | 0.00030758 |
| 11 | 65990767 | C | T | 0.932409 | 7.85E-10 | 0.00030758 |
| 11 | 65991066 | A | C | 0.932409 | 7.85E-10 | 0.00030758 |
| 11 | 65992538 | A | C | 0.932409 | 7.85E-10 | 0.00030758 |
| 11 | 65992615 | C | T | 0.932409 | 7.85E-10 | 0.00030758 |
| 11 | 65992820 | G | A | 0.932409 | 7.85E-10 | 0.00030758 |
| 11 | 65992921 | C | T | 0.932409 | 7.85E-10 | 0.00030758 |
| 11 | 65993024 | G | A | 0.932409 | 7.85E-10 | 0.00030758 |
| 11 | 65996628 | C | T | 0.932409 | 7.85E-10 | 0.00030758 |
| 11 | 65532744 | G | A | 0.930676 | 9.69E-10 | 0.00037966 |
| 11 | 65532792 | T | G | 0.930676 | 9.69E-10 | 0.00037966 |
| 11 | 65532933 | C | T | 0.930676 | 9.69E-10 | 0.00037966 |
| 11 | 65532942 | C | T | 0.930676 | 9.69E-10 | 0.00037966 |
| 11 | 65533113 | T | A | 0.930676 | 9.69E-10 | 0.00037966 |
| 11 | 65533150 | A | C | 0.930676 | 9.69E-10 | 0.00037966 |
| 11 | 65533225 | T | A | 0.930676 | 9.69E-10 | 0.00037966 |
| 11 | 65533226 | C | A | 0.930676 | 9.69E-10 | 0.00037966 |
| 11 | 65533239 | G | A | 0.930676 | 9.69E-10 | 0.00037966 |
| 11 | 65533241 | G | A | 0.930676 | 9.69E-10 | 0.00037966 |
| 11 | 63290071 | A | C | 0.880416 | 1.06E-09 | 0.00041716 |
| 11 | 63302242 | A | G | 0.881282 | 1.24E-09 | 0.00048579 |
| 11 | 64521430 | T | A | 0.930676 | 1.31E-09 | 0.00051253 |
| 11 | 64839884 | G | A | 0.915945 | 1.34E-09 | 0.0005241 |
| 11 | 64839959 | C | T | 0.915945 | 1.34E-09 | 0.0005241 |
| 11 | 64840001 | C | G | 0.915945 | 1.34E-09 | 0.0005241 |
| 11 | 64840014 | A | C | 0.915945 | 1.34E-09 | 0.0005241 |
| 11 | 64840069 | A | G | 0.915945 | 1.34E-09 | 0.0005241 |
| 11 | 64840116 | C | G | 0.915945 | 1.34E-09 | 0.0005241 |
| 11 | 64840353 | A | G | 0.915945 | 1.34E-09 | 0.0005241 |
| 11 | 64840572 | A | C | 0.915945 | 1.34E-09 | 0.0005241 |
| 11 | 64840575 | T | C | 0.915945 | 1.34E-09 | 0.0005241 |
| 11 | 64840642 | G | A | 0.916811 | 1.68E-09 | 0.00065806 |
| 11 | 64840760 | T | A | 0.916811 | 1.68E-09 | 0.00065806 |
| 11 | 64841000 | C | T | 0.916811 | 1.68E-09 | 0.00065806 |
| 11 | 65991238 | G | A | 0.931542 | 2.66E-09 | 0.00104114 |
| 11 | 65991351 | A | C | 0.931542 | 2.66E-09 | 0.00104114 |
| 11 | 65991601 | A | G | 0.931542 | 2.66E-09 | 0.00104114 |
| 11 | 65991602 | T | C | 0.931542 | 2.66E-09 | 0.00104114 |
| 11 | 65991833 | T | G | 0.931542 | 2.66E-09 | 0.00104114 |
| 11 | 65991915 | T | C | 0.931542 | 2.66E-09 | 0.00104114 |
| 11 | 65991986 | T | C | 0.931542 | 2.66E-09 | 0.00104114 |
| 11 | 65992414 | G | A | 0.931542 | 2.66E-09 | 0.00104114 |
| 11 | 65992466 | C | T | 0.931542 | 2.66E-09 | 0.00104114 |
| 11 | 63283612 | G | C | 0.881282 | 3.55E-09 | 0.0013908 |
| 11 | 66089675 | A | G | 0.932409 | 3.94E-09 | 0.00154265 |
| 11 | 65488769 | A | G | 0.931542 | 5.09E-09 | 0.00199581 |
| 11 | 65489814 | T | A | 0.931542 | 5.09E-09 | 0.00199581 |
| 11 | 65490249 | T | A | 0.931542 | 5.09E-09 | 0.00199581 |

| | | | | | | |
|---|---|---|---|---|---|---|
| 11 | 65493282 | G | A | 0.931542 | 5.09E-09 | 0.00199581 |
| 11 | 65495869 | A | G | 0.931542 | 5.09E-09 | 0.00199581 |
| 11 | 61493008 | A | C | 0.921144 | 9.24E-09 | 0.00362085 |
| 11 | 61493010 | A | C | 0.921144 | 9.24E-09 | 0.00362085 |
| 11 | 61493012 | A | C | 0.921144 | 9.24E-09 | 0.00362085 |
| 11 | 61502698 | T | G | 0.921144 | 9.24E-09 | 0.00362085 |
| 11 | 61504250 | T | G | 0.921144 | 9.24E-09 | 0.00362085 |
| 11 | 61505761 | A | G | 0.921144 | 9.24E-09 | 0.00362085 |
| 11 | 61506273 | T | G | 0.921144 | 9.24E-09 | 0.00362085 |
| 11 | 61508231 | A | G | 0.921144 | 9.24E-09 | 0.00362085 |
| 11 | 61508232 | A | G | 0.921144 | 9.24E-09 | 0.00362085 |
| 11 | 61508375 | T | C | 0.921144 | 9.24E-09 | 0.00362085 |
| 11 | 61508480 | T | A | 0.921144 | 9.24E-09 | 0.00362085 |
| 11 | 61508667 | C | G | 0.921144 | 9.24E-09 | 0.00362085 |
| 11 | 61508753 | T | C | 0.921144 | 9.24E-09 | 0.00362085 |
| 11 | 61508801 | A | G | 0.921144 | 9.24E-09 | 0.00362085 |
| 11 | 61508808 | A | C | 0.921144 | 9.24E-09 | 0.00362085 |
| 11 | 61509003 | T | C | 0.921144 | 9.24E-09 | 0.00362085 |
| 11 | 64311025 | T | C | 0.92721 | 1.03E-08 | 0.00402363 |
| 11 | 64838650 | C | A | 0.912478 | 1.18E-08 | 0.00461179 |
| 11 | 64839268 | T | C | 0.912478 | 1.18E-08 | 0.00461179 |
| 11 | 61510568 | C | A | 0.92201 | 1.25E-08 | 0.00488292 |
| 11 | 61512451 | T | C | 0.921144 | 1.33E-08 | 0.0052288 |
| 11 | 63352244 | C | T | 0.940208 | 1.44E-08 | 0.0056512 |
| 11 | 63202868 | A | T | 0.883882 | 1.44E-08 | 0.0056532 |
| 11 | 63260255 | G | T | 0.85182 | 1.84E-08 | 0.00721608 |
| 11 | 63255963 | T | C | 0.850087 | 1.85E-08 | 0.00726735 |
| 11 | 63908622 | C | T | 0.886482 | 2.20E-08 | 0.0086063 |
| 11 | 63908626 | T | C | 0.886482 | 2.20E-08 | 0.0086063 |
| 11 | 63908651 | C | T | 0.886482 | 2.20E-08 | 0.0086063 |
| 11 | 63908671 | G | C | 0.885615 | 2.61E-08 | 0.01021438 |
| 11 | 63908679 | C | G | 0.885615 | 2.61E-08 | 0.01021438 |
| 11 | 63908680 | C | T | 0.885615 | 2.61E-08 | 0.01021438 |
| 11 | 63908681 | A | G | 0.885615 | 2.61E-08 | 0.01021438 |
| 11 | 63908682 | C | A | 0.885615 | 2.61E-08 | 0.01021438 |
| 11 | 63908703 | G | A | 0.885615 | 2.61E-08 | 0.01021438 |
| 11 | 63908708 | A | G | 0.885615 | 2.61E-08 | 0.01021438 |
| 11 | 63930355 | G | A | 0.868284 | 2.77E-08 | 0.01086769 |
| 11 | 63930396 | A | T | 0.868284 | 2.77E-08 | 0.01086769 |
| 11 | 63195312 | A | G | 0.884749 | 3.00E-08 | 0.01174031 |
| 11 | 64944412 | C | T | 0.903813 | 3.07E-08 | 0.01202731 |
| 11 | 63959931 | C | A | 0.867418 | 3.24E-08 | 0.01270539 |
| 11 | 63963961 | A | G | 0.867418 | 3.24E-08 | 0.01270539 |
| 11 | 63908717 | C | G | 0.884749 | 3.34E-08 | 0.01310529 |
| 11 | 63908720 | C | T | 0.884749 | 3.34E-08 | 0.01310529 |
| 11 | 63908721 | A | G | 0.884749 | 3.34E-08 | 0.01310529 |
| 11 | 63908734 | C | T | 0.884749 | 3.34E-08 | 0.01310529 |
| 11 | 63908766 | C | G | 0.884749 | 3.34E-08 | 0.01310529 |
| 11 | 63908767 | T | C | 0.884749 | 3.34E-08 | 0.01310529 |
| 11 | 63908774 | T | G | 0.884749 | 3.34E-08 | 0.01310529 |
| 11 | 63908803 | C | T | 0.884749 | 3.34E-08 | 0.01310529 |
| 11 | 63908825 | C | G | 0.884749 | 3.34E-08 | 0.01310529 |
| 11 | 63908827 | C | A | 0.884749 | 3.34E-08 | 0.01310529 |
| 11 | 63908828 | G | A | 0.884749 | 3.34E-08 | 0.01310529 |
| 11 | 63908841 | G | A | 0.884749 | 3.34E-08 | 0.01310529 |
| 11 | 63930502 | C | T | 0.866551 | 4.03E-08 | 0.0158022 |
| 11 | 63930778 | A | G | 0.866551 | 4.03E-08 | 0.0158022 |
| 11 | 63930804 | T | C | 0.866551 | 4.03E-08 | 0.0158022 |
| 11 | 63931480 | A | G | 0.866551 | 4.03E-08 | 0.0158022 |
| 11 | 63931985 | A | G | 0.866551 | 4.03E-08 | 0.0158022 |
| 11 | 63932614 | T | C | 0.866551 | 4.03E-08 | 0.0158022 |
| 11 | 63933053 | A | G | 0.866551 | 4.03E-08 | 0.0158022 |
| 11 | 63933073 | G | T | 0.866551 | 4.03E-08 | 0.0158022 |
| 11 | 63933301 | A | G | 0.866551 | 4.03E-08 | 0.0158022 |
| 11 | 63934043 | T | C | 0.866551 | 4.03E-08 | 0.0158022 |
| 11 | 63934711 | G | C | 0.866551 | 4.03E-08 | 0.0158022 |
| 11 | 63934817 | T | A | 0.866551 | 4.03E-08 | 0.0158022 |

| | | | | | | |
|---|---|---|---|---|---|---|
| 11 | 63934905 | C | T | 0.866551 | 4.03E-08 | 0.0158022 |
| 11 | 63935009 | C | T | 0.866551 | 4.03E-08 | 0.0158022 |
| 11 | 63936846 | T | A | 0.866551 | 4.03E-08 | 0.0158022 |
| 11 | 63939386 | A | C | 0.866551 | 4.03E-08 | 0.0158022 |
| 11 | 63940631 | T | C | 0.866551 | 4.03E-08 | 0.0158022 |
| 11 | 63941785 | G | A | 0.866551 | 4.03E-08 | 0.0158022 |
| 11 | 63941887 | A | C | 0.866551 | 4.03E-08 | 0.0158022 |
| 11 | 63941978 | T | G | 0.866551 | 4.03E-08 | 0.0158022 |
| 11 | 63941993 | A | T | 0.866551 | 4.03E-08 | 0.0158022 |
| 11 | 63942843 | A | G | 0.866551 | 4.03E-08 | 0.0158022 |
| 11 | 63943128 | G | C | 0.866551 | 4.03E-08 | 0.0158022 |
| 11 | 63944904 | T | C | 0.866551 | 4.03E-08 | 0.0158022 |
| 11 | 63945538 | G | A | 0.866551 | 4.03E-08 | 0.0158022 |
| 11 | 63945715 | A | C | 0.866551 | 4.03E-08 | 0.0158022 |
| 11 | 63945934 | G | C | 0.866551 | 4.03E-08 | 0.0158022 |
| 11 | 63946495 | A | C | 0.866551 | 4.03E-08 | 0.0158022 |
| 11 | 63946508 | T | C | 0.866551 | 4.03E-08 | 0.0158022 |
| 11 | 63946569 | A | G | 0.866551 | 4.03E-08 | 0.0158022 |
| 11 | 63946570 | G | A | 0.866551 | 4.03E-08 | 0.0158022 |
| 11 | 63946745 | G | A | 0.866551 | 4.03E-08 | 0.0158022 |
| 11 | 63946836 | G | A | 0.866551 | 4.03E-08 | 0.0158022 |
| 11 | 63946868 | C | T | 0.866551 | 4.03E-08 | 0.0158022 |
| 11 | 63946877 | A | G | 0.866551 | 4.03E-08 | 0.0158022 |
| 11 | 63946884 | T | G | 0.866551 | 4.03E-08 | 0.0158022 |
| 11 | 63946903 | T | C | 0.866551 | 4.03E-08 | 0.0158022 |
| 11 | 63946928 | T | A | 0.866551 | 4.03E-08 | 0.0158022 |
| 11 | 63947702 | A | G | 0.866551 | 4.03E-08 | 0.0158022 |
| 11 | 63947826 | T | C | 0.866551 | 4.03E-08 | 0.0158022 |
| 11 | 63947841 | A | G | 0.866551 | 4.03E-08 | 0.0158022 |
| 11 | 63947880 | T | G | 0.866551 | 4.03E-08 | 0.0158022 |
| 11 | 63947915 | T | C | 0.866551 | 4.03E-08 | 0.0158022 |
| 11 | 63948168 | C | T | 0.866551 | 4.03E-08 | 0.0158022 |
| 11 | 63948179 | A | G | 0.866551 | 4.03E-08 | 0.0158022 |
| 11 | 63948203 | A | T | 0.866551 | 4.03E-08 | 0.0158022 |
| 11 | 63948437 | C | G | 0.866551 | 4.03E-08 | 0.0158022 |
| 11 | 63948438 | A | G | 0.866551 | 4.03E-08 | 0.0158022 |
| 11 | 63948446 | C | T | 0.866551 | 4.03E-08 | 0.0158022 |
| 11 | 63948557 | A | G | 0.866551 | 4.03E-08 | 0.0158022 |
| 11 | 63948574 | A | G | 0.866551 | 4.03E-08 | 0.0158022 |
| 11 | 63948680 | T | C | 0.866551 | 4.03E-08 | 0.0158022 |
| 11 | 63948684 | C | T | 0.866551 | 4.03E-08 | 0.0158022 |
| 11 | 63948727 | A | G | 0.866551 | 4.03E-08 | 0.0158022 |
| 11 | 63948803 | C | T | 0.866551 | 4.03E-08 | 0.0158022 |
| 11 | 63948871 | A | G | 0.866551 | 4.03E-08 | 0.0158022 |
| 11 | 63949263 | A | G | 0.866551 | 4.03E-08 | 0.0158022 |
| 11 | 63949355 | T | C | 0.866551 | 4.03E-08 | 0.0158022 |
| 11 | 63949675 | G | A | 0.866551 | 4.03E-08 | 0.0158022 |
| 11 | 63949708 | A | T | 0.866551 | 4.03E-08 | 0.0158022 |
| 11 | 63949887 | G | A | 0.866551 | 4.03E-08 | 0.0158022 |
| 11 | 63951088 | C | T | 0.866551 | 4.03E-08 | 0.0158022 |
| 11 | 63951362 | C | T | 0.866551 | 4.03E-08 | 0.0158022 |
| 11 | 63951396 | T | C | 0.866551 | 4.03E-08 | 0.0158022 |
| 11 | 63952219 | A | G | 0.866551 | 4.03E-08 | 0.0158022 |
| 11 | 63952554 | A | G | 0.866551 | 4.03E-08 | 0.0158022 |
| 11 | 63955292 | A | G | 0.866551 | 4.03E-08 | 0.0158022 |
| 11 | 63955349 | T | C | 0.866551 | 4.03E-08 | 0.0158022 |
| 11 | 63955727 | T | A | 0.866551 | 4.03E-08 | 0.0158022 |
| 11 | 63955910 | G | A | 0.866551 | 4.03E-08 | 0.0158022 |
| 11 | 65589070 | C | T | 0.929809 | 4.17E-08 | 0.01635904 |
| 11 | 63963505 | G | A | 0.868284 | 4.37E-08 | 0.01712061 |
| 11 | 64831085 | T | C | 0.911612 | 4.46E-08 | 0.0174971 |
| 11 | 64834419 | T | C | 0.911612 | 4.46E-08 | 0.0174971 |
| 11 | 64834579 | G | A | 0.911612 | 4.46E-08 | 0.0174971 |
| 11 | 64836885 | G | T | 0.911612 | 4.46E-08 | 0.0174971 |
| 11 | 64837646 | A | G | 0.911612 | 4.46E-08 | 0.0174971 |
| 11 | 63930484 | G | A | 0.865685 | 6.14E-08 | 0.0240776 |
| 11 | 63930494 | C | A | 0.865685 | 6.14E-08 | 0.0240776 |

| | | | | | | |
|---|---|---|---|---|---|---|
| 11 | 63930495 | A | C | 0.865685 | 6.14E-08 | 0.0240776 |
| 11 | 63908948 | C | G | 0.885615 | 8.25E-08 | 0.03235323 |
| 11 | 63908949 | C | A | 0.885615 | 8.25E-08 | 0.03235323 |
| 11 | 64063817 | C | T | 0.902946 | 8.92E-08 | 0.0349801 |
| 11 | 63294760 | A | G | 0.844021 | 1.02E-07 | 0.03989493 |
| 11 | 63295045 | A | T | 0.844021 | 1.02E-07 | 0.03989493 |
| 11 | 63298184 | A | G | 0.844021 | 1.02E-07 | 0.03989493 |
| 11 | 63298230 | T | G | 0.844021 | 1.02E-07 | 0.03989493 |
| 11 | 63299294 | T | C | 0.844021 | 1.02E-07 | 0.03989493 |
| 11 | 63908891 | A | G | 0.884749 | 1.05E-07 | 0.04125077 |
| 11 | 63908935 | A | G | 0.884749 | 1.05E-07 | 0.04125077 |
| 11 | 63908936 | C | T | 0.884749 | 1.05E-07 | 0.04125077 |
| 11 | 63908937 | A | G | 0.884749 | 1.05E-07 | 0.04125077 |
| 11 | 63908945 | C | T | 0.884749 | 1.05E-07 | 0.04125077 |
| 11 | 63770299 | T | C | 0.901213 | 1.06E-07 | 0.04137816 |
| 11 | 63770379 | T | C | 0.901213 | 1.06E-07 | 0.04137816 |
| 11 | 63770397 | G | C | 0.901213 | 1.06E-07 | 0.04137816 |
| 11 | 63770408 | C | G | 0.901213 | 1.06E-07 | 0.04137816 |
| 11 | 63770597 | C | G | 0.90208 | 1.09E-07 | 0.04281397 |
| 11 | 64055656 | T | G | 0.862218 | 1.26E-07 | 0.04951166 |

**Table S13. Significant SNPs for LFW**

| Chr | Position | *P* | Bonferroni |
|---|---|---|---|
| 1 | 20330843 | 2.80E-08 | 0.010968782 |
| 2 | 104202728 | 2.18E-09 | 0.000856424 |
| 2 | 104689105 | 9.78E-09 | 0.003832275 |

**Table S14. Significant SNPs for BW**

| Chr | Position | Allele0 | Allele1 | Freq | *P* | Bon |
|---|---|---|---|---|---|---|
| 2 | 8877739 | G | A | 0.825048 | 1.01E-13 | 3.96E-08 |
| 2 | 8812199 | T | G | 0.85086 | 5.40E-13 | 2.12E-07 |
| 2 | 8824832 | C | T | 0.85086 | 5.40E-13 | 2.12E-07 |
| 2 | 8873994 | A | G | 0.852772 | 1.22E-12 | 4.77E-07 |
| 2 | 8874135 | A | G | 0.852772 | 1.22E-12 | 4.77E-07 |
| 2 | 8874149 | G | A | 0.852772 | 1.22E-12 | 4.77E-07 |
| 2 | 8874343 | A | T | 0.852772 | 1.22E-12 | 4.77E-07 |
| 2 | 8874344 | T | A | 0.852772 | 1.22E-12 | 4.77E-07 |
| 2 | 8874735 | G | A | 0.852772 | 1.22E-12 | 4.77E-07 |
| 2 | 8867759 | A | T | 0.85086 | 1.25E-12 | 4.90E-07 |
| 2 | 8870966 | T | G | 0.85086 | 1.25E-12 | 4.90E-07 |
| 2 | 8873284 | A | T | 0.85086 | 1.25E-12 | 4.90E-07 |
| 2 | 8910937 | C | A | 0.858509 | 1.35E-12 | 5.31E-07 |
| 2 | 8911001 | G | A | 0.858509 | 1.35E-12 | 5.31E-07 |
| 2 | 8911068 | A | G | 0.858509 | 1.35E-12 | 5.31E-07 |
| 2 | 8911082 | G | A | 0.858509 | 1.35E-12 | 5.31E-07 |
| 2 | 8911108 | A | G | 0.858509 | 1.35E-12 | 5.31E-07 |
| 2 | 8911109 | G | A | 0.858509 | 1.35E-12 | 5.31E-07 |
| 2 | 8911126 | C | A | 0.858509 | 1.35E-12 | 5.31E-07 |
| 2 | 8911262 | G | T | 0.858509 | 1.35E-12 | 5.31E-07 |
| 2 | 8911264 | A | T | 0.858509 | 1.35E-12 | 5.31E-07 |
| 2 | 8844569 | A | G | 0.849904 | 1.42E-12 | 5.58E-07 |
| 2 | 8785487 | T | G | 0.852772 | 1.45E-12 | 5.67E-07 |
| 2 | 8785523 | C | T | 0.852772 | 1.45E-12 | 5.67E-07 |
| 2 | 8785622 | C | T | 0.852772 | 1.45E-12 | 5.67E-07 |
| 2 | 8785705 | T | A | 0.852772 | 1.45E-12 | 5.67E-07 |
| 2 | 8786323 | C | T | 0.852772 | 1.45E-12 | 5.67E-07 |
| 2 | 8786328 | A | G | 0.852772 | 1.45E-12 | 5.67E-07 |
| 2 | 8786516 | C | T | 0.852772 | 1.45E-12 | 5.67E-07 |
| 2 | 8787506 | C | A | 0.852772 | 1.45E-12 | 5.67E-07 |
| 2 | 8922692 | A | G | 0.853728 | 1.69E-12 | 6.64E-07 |
| 2 | 8923024 | C | A | 0.853728 | 1.69E-12 | 6.64E-07 |
| 2 | 8906778 | A | G | 0.852772 | 1.73E-12 | 6.77E-07 |
| 2 | 8906981 | G | T | 0.852772 | 1.73E-12 | 6.77E-07 |
| 2 | 8907002 | T | C | 0.852772 | 1.73E-12 | 6.77E-07 |
| 2 | 8907012 | A | G | 0.852772 | 1.73E-12 | 6.77E-07 |
| 2 | 8907095 | A | G | 0.852772 | 1.73E-12 | 6.77E-07 |
| 2 | 8764787 | C | T | 0.851816 | 1.80E-12 | 7.07E-07 |
| 2 | 8764804 | T | G | 0.851816 | 1.80E-12 | 7.07E-07 |
| 2 | 8765196 | A | G | 0.851816 | 1.80E-12 | 7.07E-07 |
| 2 | 8765722 | C | G | 0.851816 | 1.80E-12 | 7.07E-07 |
| 2 | 8766079 | A | G | 0.851816 | 1.80E-12 | 7.07E-07 |
| 2 | 8766203 | G | T | 0.851816 | 1.80E-12 | 7.07E-07 |
| 2 | 8766330 | C | T | 0.851816 | 1.80E-12 | 7.07E-07 |
| 2 | 8766350 | C | G | 0.851816 | 1.80E-12 | 7.07E-07 |
| 2 | 8766533 | A | G | 0.851816 | 1.80E-12 | 7.07E-07 |
| 2 | 8766815 | T | C | 0.851816 | 1.80E-12 | 7.07E-07 |
| 2 | 8767076 | C | A | 0.851816 | 1.80E-12 | 7.07E-07 |
| 2 | 8767453 | T | C | 0.851816 | 1.80E-12 | 7.07E-07 |
| 2 | 8767586 | T | G | 0.851816 | 1.80E-12 | 7.07E-07 |
| 2 | 8767778 | C | T | 0.851816 | 1.80E-12 | 7.07E-07 |
| 2 | 8767887 | A | G | 0.851816 | 1.80E-12 | 7.07E-07 |
| 2 | 8768088 | G | A | 0.851816 | 1.80E-12 | 7.07E-07 |
| 2 | 8768112 | T | G | 0.851816 | 1.80E-12 | 7.07E-07 |
| 2 | 8768327 | C | G | 0.851816 | 1.80E-12 | 7.07E-07 |
| 2 | 8768422 | T | C | 0.851816 | 1.80E-12 | 7.07E-07 |
| 2 | 8768471 | T | C | 0.851816 | 1.80E-12 | 7.07E-07 |
| 2 | 8769866 | T | G | 0.851816 | 1.80E-12 | 7.07E-07 |
| 2 | 8770076 | T | A | 0.851816 | 1.80E-12 | 7.07E-07 |
| 2 | 8770084 | T | A | 0.851816 | 1.80E-12 | 7.07E-07 |
| 2 | 8770208 | C | T | 0.851816 | 1.80E-12 | 7.07E-07 |
| 2 | 8770258 | A | G | 0.851816 | 1.80E-12 | 7.07E-07 |
| 2 | 8770291 | A | G | 0.851816 | 1.80E-12 | 7.07E-07 |
| 2 | 8770394 | T | C | 0.851816 | 1.80E-12 | 7.07E-07 |
| 2 | 8770796 | C | A | 0.851816 | 1.80E-12 | 7.07E-07 |

| | | | | | | |
|---|---|---|---|---|---|---|
| 2 | 8770891 | C | G | 0.851816 | 1.80E-12 | 7.07E-07 |
| 2 | 8771199 | T | A | 0.851816 | 1.80E-12 | 7.07E-07 |
| 2 | 8771201 | G | A | 0.851816 | 1.80E-12 | 7.07E-07 |
| 2 | 8771376 | T | C | 0.851816 | 1.80E-12 | 7.07E-07 |
| 2 | 8771426 | T | C | 0.851816 | 1.80E-12 | 7.07E-07 |
| 2 | 8771535 | C | G | 0.851816 | 1.80E-12 | 7.07E-07 |
| 2 | 8771769 | A | G | 0.851816 | 1.80E-12 | 7.07E-07 |
| 2 | 8771903 | T | C | 0.851816 | 1.80E-12 | 7.07E-07 |
| 2 | 8772089 | T | A | 0.851816 | 1.80E-12 | 7.07E-07 |
| 2 | 8772127 | C | T | 0.851816 | 1.80E-12 | 7.07E-07 |
| 2 | 8772204 | T | C | 0.851816 | 1.80E-12 | 7.07E-07 |
| 2 | 8772280 | A | G | 0.851816 | 1.80E-12 | 7.07E-07 |
| 2 | 8772386 | G | A | 0.851816 | 1.80E-12 | 7.07E-07 |
| 2 | 8772695 | T | C | 0.851816 | 1.80E-12 | 7.07E-07 |
| 2 | 8772827 | A | T | 0.851816 | 1.80E-12 | 7.07E-07 |
| 2 | 8772897 | A | G | 0.851816 | 1.80E-12 | 7.07E-07 |
| 2 | 8772902 | A | G | 0.851816 | 1.80E-12 | 7.07E-07 |
| 2 | 8772904 | G | A | 0.851816 | 1.80E-12 | 7.07E-07 |
| 2 | 8772944 | T | C | 0.851816 | 1.80E-12 | 7.07E-07 |
| 2 | 8773081 | C | T | 0.851816 | 1.80E-12 | 7.07E-07 |
| 2 | 8773710 | C | A | 0.851816 | 1.80E-12 | 7.07E-07 |
| 2 | 8773965 | C | A | 0.851816 | 1.80E-12 | 7.07E-07 |
| 2 | 8774010 | G | T | 0.851816 | 1.80E-12 | 7.07E-07 |
| 2 | 8774217 | T | G | 0.851816 | 1.80E-12 | 7.07E-07 |
| 2 | 8774229 | A | T | 0.851816 | 1.80E-12 | 7.07E-07 |
| 2 | 8774377 | G | A | 0.851816 | 1.80E-12 | 7.07E-07 |
| 2 | 8774451 | C | T | 0.851816 | 1.80E-12 | 7.07E-07 |
| 2 | 8774786 | A | T | 0.851816 | 1.80E-12 | 7.07E-07 |
| 2 | 8774932 | G | A | 0.851816 | 1.80E-12 | 7.07E-07 |
| 2 | 8774980 | T | G | 0.851816 | 1.80E-12 | 7.07E-07 |
| 2 | 8775066 | T | C | 0.851816 | 1.80E-12 | 7.07E-07 |
| 2 | 8775267 | C | T | 0.851816 | 1.80E-12 | 7.07E-07 |
| 2 | 8775276 | G | T | 0.851816 | 1.80E-12 | 7.07E-07 |
| 2 | 8775282 | T | C | 0.851816 | 1.80E-12 | 7.07E-07 |
| 2 | 8775284 | A | C | 0.851816 | 1.80E-12 | 7.07E-07 |
| 2 | 8775545 | C | A | 0.851816 | 1.80E-12 | 7.07E-07 |
| 2 | 8775586 | C | T | 0.851816 | 1.80E-12 | 7.07E-07 |
| 2 | 8775819 | G | T | 0.851816 | 1.80E-12 | 7.07E-07 |
| 2 | 8775826 | C | T | 0.851816 | 1.80E-12 | 7.07E-07 |
| 2 | 8775908 | C | A | 0.851816 | 1.80E-12 | 7.07E-07 |
| 2 | 8776265 | G | T | 0.851816 | 1.80E-12 | 7.07E-07 |
| 2 | 8776297 | C | T | 0.851816 | 1.80E-12 | 7.07E-07 |
| 2 | 8776364 | C | A | 0.851816 | 1.80E-12 | 7.07E-07 |
| 2 | 8776467 | G | T | 0.851816 | 1.80E-12 | 7.07E-07 |
| 2 | 8776476 | T | C | 0.851816 | 1.80E-12 | 7.07E-07 |
| 2 | 8776477 | T | C | 0.851816 | 1.80E-12 | 7.07E-07 |
| 2 | 8776522 | A | C | 0.851816 | 1.80E-12 | 7.07E-07 |
| 2 | 8776656 | G | A | 0.851816 | 1.80E-12 | 7.07E-07 |
| 2 | 8776660 | A | G | 0.851816 | 1.80E-12 | 7.07E-07 |
| 2 | 8776793 | A | G | 0.851816 | 1.80E-12 | 7.07E-07 |
| 2 | 8776955 | A | G | 0.851816 | 1.80E-12 | 7.07E-07 |
| 2 | 8776984 | T | C | 0.851816 | 1.80E-12 | 7.07E-07 |
| 2 | 8777011 | T | C | 0.851816 | 1.80E-12 | 7.07E-07 |
| 2 | 8777255 | G | A | 0.851816 | 1.80E-12 | 7.07E-07 |
| 2 | 8778186 | T | C | 0.851816 | 1.80E-12 | 7.07E-07 |
| 2 | 8778308 | T | C | 0.851816 | 1.80E-12 | 7.07E-07 |
| 2 | 8778639 | A | G | 0.851816 | 1.80E-12 | 7.07E-07 |
| 2 | 8778798 | G | A | 0.851816 | 1.80E-12 | 7.07E-07 |
| 2 | 8779360 | T | C | 0.851816 | 1.80E-12 | 7.07E-07 |
| 2 | 8780086 | C | T | 0.851816 | 1.80E-12 | 7.07E-07 |
| 2 | 8780114 | C | T | 0.851816 | 1.80E-12 | 7.07E-07 |
| 2 | 8780365 | A | T | 0.851816 | 1.80E-12 | 7.07E-07 |
| 2 | 8780480 | T | G | 0.851816 | 1.80E-12 | 7.07E-07 |
| 2 | 8780881 | C | T | 0.851816 | 1.80E-12 | 7.07E-07 |
| 2 | 8781019 | T | C | 0.851816 | 1.80E-12 | 7.07E-07 |
| 2 | 8781326 | C | A | 0.851816 | 1.80E-12 | 7.07E-07 |
| 2 | 8781862 | C | T | 0.851816 | 1.80E-12 | 7.07E-07 |

| | | | | | | |
|---|---|---|---|---|---|---|
| 2 | 8782761 | A | G | 0.851816 | 1.80E-12 | 7.07E-07 |
| 2 | 8782797 | T | C | 0.851816 | 1.80E-12 | 7.07E-07 |
| 2 | 8782813 | A | G | 0.851816 | 1.80E-12 | 7.07E-07 |
| 2 | 8782824 | C | T | 0.851816 | 1.80E-12 | 7.07E-07 |
| 2 | 8782831 | C | T | 0.851816 | 1.80E-12 | 7.07E-07 |
| 2 | 8782832 | C | T | 0.851816 | 1.80E-12 | 7.07E-07 |
| 2 | 8782919 | A | G | 0.851816 | 1.80E-12 | 7.07E-07 |
| 2 | 8782975 | C | T | 0.851816 | 1.80E-12 | 7.07E-07 |
| 2 | 8783399 | T | G | 0.851816 | 1.80E-12 | 7.07E-07 |
| 2 | 8783945 | T | C | 0.851816 | 1.80E-12 | 7.07E-07 |
| 2 | 8784183 | T | C | 0.851816 | 1.80E-12 | 7.07E-07 |
| 2 | 8784571 | T | C | 0.851816 | 1.80E-12 | 7.07E-07 |
| 2 | 8784705 | C | T | 0.851816 | 1.80E-12 | 7.07E-07 |
| 2 | 8784794 | G | A | 0.851816 | 1.80E-12 | 7.07E-07 |
| 2 | 8784992 | T | A | 0.851816 | 1.80E-12 | 7.07E-07 |
| 2 | 8785223 | A | G | 0.851816 | 1.80E-12 | 7.07E-07 |
| 2 | 8786155 | T | C | 0.851816 | 1.80E-12 | 7.07E-07 |
| 2 | 8786838 | A | G | 0.851816 | 1.80E-12 | 7.07E-07 |
| 2 | 8786951 | G | A | 0.851816 | 1.80E-12 | 7.07E-07 |
| 2 | 8786975 | T | A | 0.851816 | 1.80E-12 | 7.07E-07 |
| 2 | 8787344 | A | G | 0.851816 | 1.80E-12 | 7.07E-07 |
| 2 | 8787750 | C | T | 0.851816 | 1.80E-12 | 7.07E-07 |
| 2 | 8790044 | A | G | 0.851816 | 1.80E-12 | 7.07E-07 |
| 2 | 8790294 | G | C | 0.851816 | 1.80E-12 | 7.07E-07 |
| 2 | 8790484 | T | C | 0.851816 | 1.80E-12 | 7.07E-07 |
| 2 | 8790547 | G | A | 0.851816 | 1.80E-12 | 7.07E-07 |
| 2 | 8790605 | T | C | 0.851816 | 1.80E-12 | 7.07E-07 |
| 2 | 8790606 | G | A | 0.851816 | 1.80E-12 | 7.07E-07 |
| 2 | 8790608 | T | C | 0.851816 | 1.80E-12 | 7.07E-07 |
| 2 | 8790951 | G | T | 0.851816 | 1.80E-12 | 7.07E-07 |
| 2 | 8791080 | A | G | 0.851816 | 1.80E-12 | 7.07E-07 |
| 2 | 8791893 | T | C | 0.851816 | 1.80E-12 | 7.07E-07 |
| 2 | 8792077 | C | G | 0.851816 | 1.80E-12 | 7.07E-07 |
| 2 | 8793049 | G | T | 0.851816 | 1.80E-12 | 7.07E-07 |
| 2 | 8793114 | C | T | 0.851816 | 1.80E-12 | 7.07E-07 |
| 2 | 8793116 | G | A | 0.851816 | 1.80E-12 | 7.07E-07 |
| 2 | 8793215 | C | T | 0.851816 | 1.80E-12 | 7.07E-07 |
| 2 | 8793238 | T | A | 0.851816 | 1.80E-12 | 7.07E-07 |
| 2 | 8793277 | G | A | 0.851816 | 1.80E-12 | 7.07E-07 |
| 2 | 8793319 | C | T | 0.851816 | 1.80E-12 | 7.07E-07 |
| 2 | 8793357 | G | T | 0.851816 | 1.80E-12 | 7.07E-07 |
| 2 | 8793363 | T | C | 0.851816 | 1.80E-12 | 7.07E-07 |
| 2 | 8793368 | A | C | 0.851816 | 1.80E-12 | 7.07E-07 |
| 2 | 8793442 | G | A | 0.851816 | 1.80E-12 | 7.07E-07 |
| 2 | 8793461 | T | C | 0.851816 | 1.80E-12 | 7.07E-07 |
| 2 | 8793580 | A | C | 0.851816 | 1.80E-12 | 7.07E-07 |
| 2 | 8793678 | T | C | 0.851816 | 1.80E-12 | 7.07E-07 |
| 2 | 8793718 | G | A | 0.851816 | 1.80E-12 | 7.07E-07 |
| 2 | 8793728 | G | T | 0.851816 | 1.80E-12 | 7.07E-07 |
| 2 | 8793882 | C | T | 0.851816 | 1.80E-12 | 7.07E-07 |
| 2 | 8793898 | C | T | 0.851816 | 1.80E-12 | 7.07E-07 |
| 2 | 8793966 | G | T | 0.851816 | 1.80E-12 | 7.07E-07 |
| 2 | 8794361 | C | T | 0.851816 | 1.80E-12 | 7.07E-07 |
| 2 | 8794386 | T | C | 0.851816 | 1.80E-12 | 7.07E-07 |
| 2 | 8795016 | G | A | 0.851816 | 1.80E-12 | 7.07E-07 |
| 2 | 8795699 | C | T | 0.851816 | 1.80E-12 | 7.07E-07 |
| 2 | 8795735 | G | A | 0.851816 | 1.80E-12 | 7.07E-07 |
| 2 | 8795898 | T | G | 0.851816 | 1.80E-12 | 7.07E-07 |
| 2 | 8796589 | T | A | 0.851816 | 1.80E-12 | 7.07E-07 |
| 2 | 8797306 | C | T | 0.851816 | 1.80E-12 | 7.07E-07 |
| 2 | 8797598 | G | A | 0.851816 | 1.80E-12 | 7.07E-07 |
| 2 | 8797608 | C | T | 0.851816 | 1.80E-12 | 7.07E-07 |
| 2 | 8798403 | A | G | 0.851816 | 1.80E-12 | 7.07E-07 |
| 2 | 8798474 | G | A | 0.851816 | 1.80E-12 | 7.07E-07 |
| 2 | 8798621 | A | G | 0.851816 | 1.80E-12 | 7.07E-07 |
| 2 | 8799022 | A | G | 0.851816 | 1.80E-12 | 7.07E-07 |
| 2 | 8799071 | A | T | 0.851816 | 1.80E-12 | 7.07E-07 |

| | | | | | | |
|---|---|---|---|---|---|---|
| 2 | 8799207 | G | T | 0.851816 | 1.80E-12 | 7.07E-07 |
| 2 | 8799216 | T | C | 0.851816 | 1.80E-12 | 7.07E-07 |
| 2 | 8799257 | G | A | 0.851816 | 1.80E-12 | 7.07E-07 |
| 2 | 8799755 | T | C | 0.851816 | 1.80E-12 | 7.07E-07 |
| 2 | 8799819 | T | G | 0.851816 | 1.80E-12 | 7.07E-07 |
| 2 | 8800489 | A | G | 0.851816 | 1.80E-12 | 7.07E-07 |
| 2 | 8800505 | C | T | 0.851816 | 1.80E-12 | 7.07E-07 |
| 2 | 8801241 | A | T | 0.851816 | 1.80E-12 | 7.07E-07 |
| 2 | 8802537 | T | C | 0.851816 | 1.80E-12 | 7.07E-07 |
| 2 | 8802539 | T | G | 0.851816 | 1.80E-12 | 7.07E-07 |
| 2 | 8802856 | C | G | 0.851816 | 1.80E-12 | 7.07E-07 |
| 2 | 8802916 | T | C | 0.851816 | 1.80E-12 | 7.07E-07 |
| 2 | 8803034 | A | G | 0.851816 | 1.80E-12 | 7.07E-07 |
| 2 | 8803098 | C | T | 0.851816 | 1.80E-12 | 7.07E-07 |
| 2 | 8803200 | C | T | 0.851816 | 1.80E-12 | 7.07E-07 |
| 2 | 8803587 | G | A | 0.851816 | 1.80E-12 | 7.07E-07 |
| 2 | 8804605 | T | G | 0.851816 | 1.80E-12 | 7.07E-07 |
| 2 | 8804626 | C | T | 0.851816 | 1.80E-12 | 7.07E-07 |
| 2 | 8804657 | C | T | 0.851816 | 1.80E-12 | 7.07E-07 |
| 2 | 8804746 | T | G | 0.851816 | 1.80E-12 | 7.07E-07 |
| 2 | 8804765 | G | A | 0.851816 | 1.80E-12 | 7.07E-07 |
| 2 | 8804804 | A | G | 0.851816 | 1.80E-12 | 7.07E-07 |
| 2 | 8804829 | G | A | 0.851816 | 1.80E-12 | 7.07E-07 |
| 2 | 8804836 | T | C | 0.851816 | 1.80E-12 | 7.07E-07 |
| 2 | 8804870 | C | T | 0.851816 | 1.80E-12 | 7.07E-07 |
| 2 | 8804882 | T | C | 0.851816 | 1.80E-12 | 7.07E-07 |
| 2 | 8805797 | G | A | 0.851816 | 1.80E-12 | 7.07E-07 |
| 2 | 8806023 | A | C | 0.851816 | 1.80E-12 | 7.07E-07 |
| 2 | 8806030 | A | G | 0.851816 | 1.80E-12 | 7.07E-07 |
| 2 | 8806168 | T | G | 0.851816 | 1.80E-12 | 7.07E-07 |
| 2 | 8806290 | G | A | 0.851816 | 1.80E-12 | 7.07E-07 |
| 2 | 8806347 | G | A | 0.851816 | 1.80E-12 | 7.07E-07 |
| 2 | 8806455 | T | C | 0.851816 | 1.80E-12 | 7.07E-07 |
| 2 | 8806945 | T | G | 0.851816 | 1.80E-12 | 7.07E-07 |
| 2 | 8807165 | A | T | 0.851816 | 1.80E-12 | 7.07E-07 |
| 2 | 8807214 | T | C | 0.851816 | 1.80E-12 | 7.07E-07 |
| 2 | 8807623 | G | T | 0.851816 | 1.80E-12 | 7.07E-07 |
| 2 | 8807778 | C | T | 0.851816 | 1.80E-12 | 7.07E-07 |
| 2 | 8807828 | A | C | 0.851816 | 1.80E-12 | 7.07E-07 |
| 2 | 8807988 | A | C | 0.851816 | 1.80E-12 | 7.07E-07 |
| 2 | 8808228 | C | T | 0.851816 | 1.80E-12 | 7.07E-07 |
| 2 | 8809041 | C | A | 0.851816 | 1.80E-12 | 7.07E-07 |
| 2 | 8809410 | G | A | 0.851816 | 1.80E-12 | 7.07E-07 |
| 2 | 8809651 | G | T | 0.851816 | 1.80E-12 | 7.07E-07 |
| 2 | 8810588 | T | C | 0.851816 | 1.80E-12 | 7.07E-07 |
| 2 | 8810593 | C | T | 0.851816 | 1.80E-12 | 7.07E-07 |
| 2 | 8810608 | A | G | 0.851816 | 1.80E-12 | 7.07E-07 |
| 2 | 8811140 | G | A | 0.851816 | 1.80E-12 | 7.07E-07 |
| 2 | 8811204 | T | A | 0.851816 | 1.80E-12 | 7.07E-07 |
| 2 | 8811497 | C | T | 0.851816 | 1.80E-12 | 7.07E-07 |
| 2 | 8812372 | A | C | 0.851816 | 1.80E-12 | 7.07E-07 |
| 2 | 8812380 | A | G | 0.851816 | 1.80E-12 | 7.07E-07 |
| 2 | 8812400 | G | A | 0.851816 | 1.80E-12 | 7.07E-07 |
| 2 | 8812417 | C | T | 0.851816 | 1.80E-12 | 7.07E-07 |
| 2 | 8812419 | A | C | 0.851816 | 1.80E-12 | 7.07E-07 |
| 2 | 8812507 | G | C | 0.851816 | 1.80E-12 | 7.07E-07 |
| 2 | 8812530 | C | T | 0.851816 | 1.80E-12 | 7.07E-07 |
| 2 | 8812666 | A | T | 0.851816 | 1.80E-12 | 7.07E-07 |
| 2 | 8812984 | T | C | 0.851816 | 1.80E-12 | 7.07E-07 |
| 2 | 8812990 | C | T | 0.851816 | 1.80E-12 | 7.07E-07 |
| 2 | 8813017 | C | A | 0.851816 | 1.80E-12 | 7.07E-07 |
| 2 | 8813054 | G | A | 0.851816 | 1.80E-12 | 7.07E-07 |
| 2 | 8813180 | G | A | 0.851816 | 1.80E-12 | 7.07E-07 |
| 2 | 8813244 | T | G | 0.851816 | 1.80E-12 | 7.07E-07 |
| 2 | 8813435 | T | A | 0.851816 | 1.80E-12 | 7.07E-07 |
| 2 | 8813436 | C | T | 0.851816 | 1.80E-12 | 7.07E-07 |
| 2 | 8813651 | G | C | 0.851816 | 1.80E-12 | 7.07E-07 |

| | | | | | | |
|---|---|---|---|---|---|---|
| 2 | 8813686 | A | G | 0.851816 | 1.80E-12 | 7.07E-07 |
| 2 | 8813899 | C | T | 0.851816 | 1.80E-12 | 7.07E-07 |
| 2 | 8813934 | C | T | 0.851816 | 1.80E-12 | 7.07E-07 |
| 2 | 8813963 | A | T | 0.851816 | 1.80E-12 | 7.07E-07 |
| 2 | 8814009 | A | G | 0.851816 | 1.80E-12 | 7.07E-07 |
| 2 | 8814232 | G | A | 0.851816 | 1.80E-12 | 7.07E-07 |
| 2 | 8814446 | A | T | 0.851816 | 1.80E-12 | 7.07E-07 |
| 2 | 8814702 | G | A | 0.851816 | 1.80E-12 | 7.07E-07 |
| 2 | 8815866 | A | G | 0.851816 | 1.80E-12 | 7.07E-07 |
| 2 | 8816380 | T | A | 0.851816 | 1.80E-12 | 7.07E-07 |
| 2 | 8816381 | T | A | 0.851816 | 1.80E-12 | 7.07E-07 |
| 2 | 8816382 | T | A | 0.851816 | 1.80E-12 | 7.07E-07 |
| 2 | 8816525 | T | C | 0.851816 | 1.80E-12 | 7.07E-07 |
| 2 | 8816534 | T | G | 0.851816 | 1.80E-12 | 7.07E-07 |
| 2 | 8817228 | T | C | 0.851816 | 1.80E-12 | 7.07E-07 |
| 2 | 8817280 | A | T | 0.851816 | 1.80E-12 | 7.07E-07 |
| 2 | 8817300 | A | G | 0.851816 | 1.80E-12 | 7.07E-07 |
| 2 | 8817309 | C | T | 0.851816 | 1.80E-12 | 7.07E-07 |
| 2 | 8817553 | G | A | 0.851816 | 1.80E-12 | 7.07E-07 |
| 2 | 8817825 | C | T | 0.851816 | 1.80E-12 | 7.07E-07 |
| 2 | 8817866 | G | C | 0.851816 | 1.80E-12 | 7.07E-07 |
| 2 | 8818140 | C | T | 0.851816 | 1.80E-12 | 7.07E-07 |
| 2 | 8818401 | T | C | 0.851816 | 1.80E-12 | 7.07E-07 |
| 2 | 8818407 | A | G | 0.851816 | 1.80E-12 | 7.07E-07 |
| 2 | 8818462 | C | T | 0.851816 | 1.80E-12 | 7.07E-07 |
| 2 | 8818773 | A | G | 0.851816 | 1.80E-12 | 7.07E-07 |
| 2 | 8818873 | G | T | 0.851816 | 1.80E-12 | 7.07E-07 |
| 2 | 8819102 | A | G | 0.851816 | 1.80E-12 | 7.07E-07 |
| 2 | 8819852 | T | C | 0.851816 | 1.80E-12 | 7.07E-07 |
| 2 | 8820112 | G | A | 0.851816 | 1.80E-12 | 7.07E-07 |
| 2 | 8820214 | G | A | 0.851816 | 1.80E-12 | 7.07E-07 |
| 2 | 8820486 | T | A | 0.851816 | 1.80E-12 | 7.07E-07 |
| 2 | 8820544 | C | A | 0.851816 | 1.80E-12 | 7.07E-07 |
| 2 | 8820860 | C | T | 0.851816 | 1.80E-12 | 7.07E-07 |
| 2 | 8820915 | T | A | 0.851816 | 1.80E-12 | 7.07E-07 |
| 2 | 8821035 | T | C | 0.851816 | 1.80E-12 | 7.07E-07 |
| 2 | 8821109 | A | G | 0.851816 | 1.80E-12 | 7.07E-07 |
| 2 | 8821129 | T | C | 0.851816 | 1.80E-12 | 7.07E-07 |
| 2 | 8821219 | T | C | 0.851816 | 1.80E-12 | 7.07E-07 |
| 2 | 8821370 | A | G | 0.851816 | 1.80E-12 | 7.07E-07 |
| 2 | 8821619 | A | C | 0.851816 | 1.80E-12 | 7.07E-07 |
| 2 | 8821698 | A | T | 0.851816 | 1.80E-12 | 7.07E-07 |
| 2 | 8821837 | T | C | 0.851816 | 1.80E-12 | 7.07E-07 |
| 2 | 8821990 | A | G | 0.851816 | 1.80E-12 | 7.07E-07 |
| 2 | 8822011 | T | C | 0.851816 | 1.80E-12 | 7.07E-07 |
| 2 | 8822155 | T | C | 0.851816 | 1.80E-12 | 7.07E-07 |
| 2 | 8822970 | A | G | 0.851816 | 1.80E-12 | 7.07E-07 |
| 2 | 8823144 | G | A | 0.851816 | 1.80E-12 | 7.07E-07 |
| 2 | 8823842 | A | C | 0.851816 | 1.80E-12 | 7.07E-07 |
| 2 | 8823998 | A | T | 0.851816 | 1.80E-12 | 7.07E-07 |
| 2 | 8824191 | T | C | 0.851816 | 1.80E-12 | 7.07E-07 |
| 2 | 8824238 | A | T | 0.851816 | 1.80E-12 | 7.07E-07 |
| 2 | 8824248 | G | A | 0.851816 | 1.80E-12 | 7.07E-07 |
| 2 | 8824251 | G | A | 0.851816 | 1.80E-12 | 7.07E-07 |
| 2 | 8824281 | C | A | 0.851816 | 1.80E-12 | 7.07E-07 |
| 2 | 8824443 | T | A | 0.851816 | 1.80E-12 | 7.07E-07 |
| 2 | 8824543 | A | T | 0.851816 | 1.80E-12 | 7.07E-07 |
| 2 | 8824629 | T | C | 0.851816 | 1.80E-12 | 7.07E-07 |
| 2 | 8824631 | C | T | 0.851816 | 1.80E-12 | 7.07E-07 |
| 2 | 8824853 | T | C | 0.851816 | 1.80E-12 | 7.07E-07 |
| 2 | 8824914 | T | A | 0.851816 | 1.80E-12 | 7.07E-07 |
| 2 | 8824949 | A | T | 0.851816 | 1.80E-12 | 7.07E-07 |
| 2 | 8825053 | C | G | 0.851816 | 1.80E-12 | 7.07E-07 |
| 2 | 8825138 | T | C | 0.851816 | 1.80E-12 | 7.07E-07 |
| 2 | 8825190 | G | A | 0.851816 | 1.80E-12 | 7.07E-07 |
| 2 | 8825218 | G | A | 0.851816 | 1.80E-12 | 7.07E-07 |
| 2 | 8825270 | G | A | 0.851816 | 1.80E-12 | 7.07E-07 |

| | | | | | | |
|---|---|---|---|---|---|---|
| 2 | 8825477 | G | A | 0.851816 | 1.80E-12 | 7.07E-07 |
| 2 | 8825561 | A | G | 0.851816 | 1.80E-12 | 7.07E-07 |
| 2 | 8825596 | A | G | 0.851816 | 1.80E-12 | 7.07E-07 |
| 2 | 8825605 | A | G | 0.851816 | 1.80E-12 | 7.07E-07 |
| 2 | 8825766 | T | G | 0.851816 | 1.80E-12 | 7.07E-07 |
| 2 | 8825785 | C | T | 0.851816 | 1.80E-12 | 7.07E-07 |
| 2 | 8825920 | T | C | 0.851816 | 1.80E-12 | 7.07E-07 |
| 2 | 8825929 | G | A | 0.851816 | 1.80E-12 | 7.07E-07 |
| 2 | 8826161 | C | T | 0.851816 | 1.80E-12 | 7.07E-07 |
| 2 | 8826242 | G | T | 0.851816 | 1.80E-12 | 7.07E-07 |
| 2 | 8826332 | C | T | 0.851816 | 1.80E-12 | 7.07E-07 |
| 2 | 8826362 | C | T | 0.851816 | 1.80E-12 | 7.07E-07 |
| 2 | 8826378 | G | C | 0.851816 | 1.80E-12 | 7.07E-07 |
| 2 | 8826494 | T | G | 0.851816 | 1.80E-12 | 7.07E-07 |
| 2 | 8827266 | A | G | 0.851816 | 1.80E-12 | 7.07E-07 |
| 2 | 8827831 | G | A | 0.851816 | 1.80E-12 | 7.07E-07 |
| 2 | 8828095 | G | A | 0.851816 | 1.80E-12 | 7.07E-07 |
| 2 | 8828129 | T | C | 0.851816 | 1.80E-12 | 7.07E-07 |
| 2 | 8828232 | G | A | 0.851816 | 1.80E-12 | 7.07E-07 |
| 2 | 8828321 | C | T | 0.851816 | 1.80E-12 | 7.07E-07 |
| 2 | 8828322 | A | G | 0.851816 | 1.80E-12 | 7.07E-07 |
| 2 | 8828376 | C | T | 0.851816 | 1.80E-12 | 7.07E-07 |
| 2 | 8828390 | C | T | 0.851816 | 1.80E-12 | 7.07E-07 |
| 2 | 8828553 | A | G | 0.851816 | 1.80E-12 | 7.07E-07 |
| 2 | 8828747 | C | A | 0.851816 | 1.80E-12 | 7.07E-07 |
| 2 | 8828850 | G | A | 0.851816 | 1.80E-12 | 7.07E-07 |
| 2 | 8828875 | A | G | 0.851816 | 1.80E-12 | 7.07E-07 |
| 2 | 8828917 | C | T | 0.851816 | 1.80E-12 | 7.07E-07 |
| 2 | 8829260 | T | C | 0.851816 | 1.80E-12 | 7.07E-07 |
| 2 | 8829360 | A | G | 0.851816 | 1.80E-12 | 7.07E-07 |
| 2 | 8829513 | C | T | 0.851816 | 1.80E-12 | 7.07E-07 |
| 2 | 8829811 | G | T | 0.851816 | 1.80E-12 | 7.07E-07 |
| 2 | 8830134 | T | C | 0.851816 | 1.80E-12 | 7.07E-07 |
| 2 | 8830163 | A | G | 0.851816 | 1.80E-12 | 7.07E-07 |
| 2 | 8830241 | C | T | 0.851816 | 1.80E-12 | 7.07E-07 |
| 2 | 8830242 | C | T | 0.851816 | 1.80E-12 | 7.07E-07 |
| 2 | 8830546 | T | A | 0.851816 | 1.80E-12 | 7.07E-07 |
| 2 | 8830892 | T | C | 0.851816 | 1.80E-12 | 7.07E-07 |
| 2 | 8831094 | C | T | 0.851816 | 1.80E-12 | 7.07E-07 |
| 2 | 8831185 | T | C | 0.851816 | 1.80E-12 | 7.07E-07 |
| 2 | 8831249 | T | C | 0.851816 | 1.80E-12 | 7.07E-07 |
| 2 | 8831700 | A | G | 0.851816 | 1.80E-12 | 7.07E-07 |
| 2 | 8831771 | C | A | 0.851816 | 1.80E-12 | 7.07E-07 |
| 2 | 8831772 | A | G | 0.851816 | 1.80E-12 | 7.07E-07 |
| 2 | 8832020 | G | A | 0.851816 | 1.80E-12 | 7.07E-07 |
| 2 | 8832022 | C | T | 0.851816 | 1.80E-12 | 7.07E-07 |
| 2 | 8832217 | A | T | 0.851816 | 1.80E-12 | 7.07E-07 |
| 2 | 8832260 | A | C | 0.851816 | 1.80E-12 | 7.07E-07 |
| 2 | 8832279 | C | T | 0.851816 | 1.80E-12 | 7.07E-07 |
| 2 | 8832623 | A | T | 0.851816 | 1.80E-12 | 7.07E-07 |
| 2 | 8832699 | A | G | 0.851816 | 1.80E-12 | 7.07E-07 |
| 2 | 8832770 | C | T | 0.851816 | 1.80E-12 | 7.07E-07 |
| 2 | 8832865 | T | C | 0.851816 | 1.80E-12 | 7.07E-07 |
| 2 | 8833021 | T | G | 0.851816 | 1.80E-12 | 7.07E-07 |
| 2 | 8833168 | A | G | 0.851816 | 1.80E-12 | 7.07E-07 |
| 2 | 8833263 | T | C | 0.851816 | 1.80E-12 | 7.07E-07 |
| 2 | 8833324 | G | A | 0.851816 | 1.80E-12 | 7.07E-07 |
| 2 | 8833345 | T | A | 0.851816 | 1.80E-12 | 7.07E-07 |
| 2 | 8833565 | A | C | 0.851816 | 1.80E-12 | 7.07E-07 |
| 2 | 8833651 | C | T | 0.851816 | 1.80E-12 | 7.07E-07 |
| 2 | 8835473 | C | T | 0.851816 | 1.80E-12 | 7.07E-07 |
| 2 | 8835514 | C | A | 0.851816 | 1.80E-12 | 7.07E-07 |
| 2 | 8837873 | C | T | 0.851816 | 1.80E-12 | 7.07E-07 |
| 2 | 8837889 | C | T | 0.851816 | 1.80E-12 | 7.07E-07 |
| 2 | 8837920 | T | C | 0.851816 | 1.80E-12 | 7.07E-07 |
| 2 | 8839892 | A | T | 0.851816 | 1.80E-12 | 7.07E-07 |
| 2 | 8841022 | T | C | 0.851816 | 1.80E-12 | 7.07E-07 |

| | | | | | | |
|---|---|---|---|---|---|---|
| 2 | 8841782 | C | T | 0.851816 | 1.80E-12 | 7.07E-07 |
| 2 | 8842092 | T | C | 0.851816 | 1.80E-12 | 7.07E-07 |
| 2 | 8842187 | T | G | 0.851816 | 1.80E-12 | 7.07E-07 |
| 2 | 8842559 | A | G | 0.851816 | 1.80E-12 | 7.07E-07 |
| 2 | 8842706 | C | T | 0.851816 | 1.80E-12 | 7.07E-07 |
| 2 | 8842806 | C | T | 0.851816 | 1.80E-12 | 7.07E-07 |
| 2 | 8842882 | C | T | 0.851816 | 1.80E-12 | 7.07E-07 |
| 2 | 8843070 | G | A | 0.851816 | 1.80E-12 | 7.07E-07 |
| 2 | 8843075 | C | G | 0.851816 | 1.80E-12 | 7.07E-07 |
| 2 | 8843144 | G | C | 0.851816 | 1.80E-12 | 7.07E-07 |
| 2 | 8843202 | C | T | 0.851816 | 1.80E-12 | 7.07E-07 |
| 2 | 8843229 | C | T | 0.851816 | 1.80E-12 | 7.07E-07 |
| 2 | 8843340 | A | G | 0.851816 | 1.80E-12 | 7.07E-07 |
| 2 | 8843500 | T | C | 0.851816 | 1.80E-12 | 7.07E-07 |
| 2 | 8843555 | T | C | 0.851816 | 1.80E-12 | 7.07E-07 |
| 2 | 8843614 | T | A | 0.851816 | 1.80E-12 | 7.07E-07 |
| 2 | 8843616 | G | A | 0.851816 | 1.80E-12 | 7.07E-07 |
| 2 | 8843816 | A | G | 0.851816 | 1.80E-12 | 7.07E-07 |
| 2 | 8844095 | T | A | 0.851816 | 1.80E-12 | 7.07E-07 |
| 2 | 8844107 | G | A | 0.851816 | 1.80E-12 | 7.07E-07 |
| 2 | 8844133 | T | G | 0.851816 | 1.80E-12 | 7.07E-07 |
| 2 | 8844162 | C | A | 0.851816 | 1.80E-12 | 7.07E-07 |
| 2 | 8844295 | G | A | 0.851816 | 1.80E-12 | 7.07E-07 |
| 2 | 8844387 | G | T | 0.851816 | 1.80E-12 | 7.07E-07 |
| 2 | 8844503 | C | A | 0.851816 | 1.80E-12 | 7.07E-07 |
| 2 | 8845494 | T | C | 0.851816 | 1.80E-12 | 7.07E-07 |
| 2 | 8845499 | C | T | 0.851816 | 1.80E-12 | 7.07E-07 |
| 2 | 8845694 | T | G | 0.851816 | 1.80E-12 | 7.07E-07 |
| 2 | 8845743 | C | T | 0.851816 | 1.80E-12 | 7.07E-07 |
| 2 | 8845946 | C | T | 0.851816 | 1.80E-12 | 7.07E-07 |
| 2 | 8846073 | T | C | 0.851816 | 1.80E-12 | 7.07E-07 |
| 2 | 8846301 | C | T | 0.851816 | 1.80E-12 | 7.07E-07 |
| 2 | 8846302 | C | A | 0.851816 | 1.80E-12 | 7.07E-07 |
| 2 | 8846303 | C | G | 0.851816 | 1.80E-12 | 7.07E-07 |
| 2 | 8846352 | C | T | 0.851816 | 1.80E-12 | 7.07E-07 |
| 2 | 8846465 | A | G | 0.851816 | 1.80E-12 | 7.07E-07 |
| 2 | 8846578 | A | T | 0.851816 | 1.80E-12 | 7.07E-07 |
| 2 | 8846610 | T | C | 0.851816 | 1.80E-12 | 7.07E-07 |
| 2 | 8846653 | A | C | 0.851816 | 1.80E-12 | 7.07E-07 |
| 2 | 8846743 | C | A | 0.851816 | 1.80E-12 | 7.07E-07 |
| 2 | 8846773 | T | C | 0.851816 | 1.80E-12 | 7.07E-07 |
| 2 | 8846816 | C | A | 0.851816 | 1.80E-12 | 7.07E-07 |
| 2 | 8847140 | G | T | 0.851816 | 1.80E-12 | 7.07E-07 |
| 2 | 8847147 | T | G | 0.851816 | 1.80E-12 | 7.07E-07 |
| 2 | 8847216 | T | C | 0.851816 | 1.80E-12 | 7.07E-07 |
| 2 | 8847336 | C | T | 0.851816 | 1.80E-12 | 7.07E-07 |
| 2 | 8847375 | T | C | 0.851816 | 1.80E-12 | 7.07E-07 |
| 2 | 8847388 | T | C | 0.851816 | 1.80E-12 | 7.07E-07 |
| 2 | 8847406 | T | C | 0.851816 | 1.80E-12 | 7.07E-07 |
| 2 | 8847433 | C | T | 0.851816 | 1.80E-12 | 7.07E-07 |
| 2 | 8847594 | A | G | 0.851816 | 1.80E-12 | 7.07E-07 |
| 2 | 8847782 | T | C | 0.851816 | 1.80E-12 | 7.07E-07 |
| 2 | 8847876 | T | A | 0.851816 | 1.80E-12 | 7.07E-07 |
| 2 | 8848459 | G | A | 0.851816 | 1.80E-12 | 7.07E-07 |
| 2 | 8848590 | C | T | 0.851816 | 1.80E-12 | 7.07E-07 |
| 2 | 8848640 | T | C | 0.851816 | 1.80E-12 | 7.07E-07 |
| 2 | 8848808 | A | G | 0.851816 | 1.80E-12 | 7.07E-07 |
| 2 | 8848828 | T | G | 0.851816 | 1.80E-12 | 7.07E-07 |
| 2 | 8848914 | A | C | 0.851816 | 1.80E-12 | 7.07E-07 |
| 2 | 8848980 | A | G | 0.851816 | 1.80E-12 | 7.07E-07 |
| 2 | 8848991 | C | A | 0.851816 | 1.80E-12 | 7.07E-07 |
| 2 | 8849070 | C | T | 0.851816 | 1.80E-12 | 7.07E-07 |
| 2 | 8849187 | C | T | 0.851816 | 1.80E-12 | 7.07E-07 |
| 2 | 8849226 | T | G | 0.851816 | 1.80E-12 | 7.07E-07 |
| 2 | 8849253 | C | A | 0.851816 | 1.80E-12 | 7.07E-07 |
| 2 | 8849410 | G | A | 0.851816 | 1.80E-12 | 7.07E-07 |
| 2 | 8849470 | T | C | 0.851816 | 1.80E-12 | 7.07E-07 |

| | | | | | | |
|---|---|---|---|---|---|---|
| 2 | 8849558 | C | T | 0.851816 | 1.80E-12 | 7.07E-07 |
| 2 | 8849569 | A | C | 0.851816 | 1.80E-12 | 7.07E-07 |
| 2 | 8849673 | A | G | 0.851816 | 1.80E-12 | 7.07E-07 |
| 2 | 8849796 | T | G | 0.851816 | 1.80E-12 | 7.07E-07 |
| 2 | 8849832 | G | A | 0.851816 | 1.80E-12 | 7.07E-07 |
| 2 | 8849896 | T | C | 0.851816 | 1.80E-12 | 7.07E-07 |
| 2 | 8850061 | G | A | 0.851816 | 1.80E-12 | 7.07E-07 |
| 2 | 8850213 | A | C | 0.851816 | 1.80E-12 | 7.07E-07 |
| 2 | 8851179 | C | T | 0.851816 | 1.80E-12 | 7.07E-07 |
| 2 | 8851199 | A | T | 0.851816 | 1.80E-12 | 7.07E-07 |
| 2 | 8851312 | A | G | 0.851816 | 1.80E-12 | 7.07E-07 |
| 2 | 8851457 | G | A | 0.851816 | 1.80E-12 | 7.07E-07 |
| 2 | 8851626 | C | G | 0.851816 | 1.80E-12 | 7.07E-07 |
| 2 | 8851639 | T | A | 0.851816 | 1.80E-12 | 7.07E-07 |
| 2 | 8851684 | G | A | 0.851816 | 1.80E-12 | 7.07E-07 |
| 2 | 8851760 | A | T | 0.851816 | 1.80E-12 | 7.07E-07 |
| 2 | 8851781 | C | G | 0.851816 | 1.80E-12 | 7.07E-07 |
| 2 | 8851893 | G | A | 0.851816 | 1.80E-12 | 7.07E-07 |
| 2 | 8851991 | G | A | 0.851816 | 1.80E-12 | 7.07E-07 |
| 2 | 8852059 | G | A | 0.851816 | 1.80E-12 | 7.07E-07 |
| 2 | 8852200 | T | C | 0.851816 | 1.80E-12 | 7.07E-07 |
| 2 | 8852429 | T | C | 0.851816 | 1.80E-12 | 7.07E-07 |
| 2 | 8852528 | C | A | 0.851816 | 1.80E-12 | 7.07E-07 |
| 2 | 8852702 | A | G | 0.851816 | 1.80E-12 | 7.07E-07 |
| 2 | 8852719 | G | A | 0.851816 | 1.80E-12 | 7.07E-07 |
| 2 | 8852806 | G | C | 0.851816 | 1.80E-12 | 7.07E-07 |
| 2 | 8852839 | C | T | 0.851816 | 1.80E-12 | 7.07E-07 |
| 2 | 8852854 | G | A | 0.851816 | 1.80E-12 | 7.07E-07 |
| 2 | 8853049 | C | A | 0.851816 | 1.80E-12 | 7.07E-07 |
| 2 | 8853197 | T | C | 0.851816 | 1.80E-12 | 7.07E-07 |
| 2 | 8853238 | G | T | 0.851816 | 1.80E-12 | 7.07E-07 |
| 2 | 8853797 | G | T | 0.851816 | 1.80E-12 | 7.07E-07 |
| 2 | 8853810 | T | C | 0.851816 | 1.80E-12 | 7.07E-07 |
| 2 | 8854068 | T | C | 0.851816 | 1.80E-12 | 7.07E-07 |
| 2 | 8854143 | C | T | 0.851816 | 1.80E-12 | 7.07E-07 |
| 2 | 8854173 | A | G | 0.851816 | 1.80E-12 | 7.07E-07 |
| 2 | 8854634 | C | T | 0.851816 | 1.80E-12 | 7.07E-07 |
| 2 | 8854882 | G | A | 0.851816 | 1.80E-12 | 7.07E-07 |
| 2 | 8855011 | T | C | 0.851816 | 1.80E-12 | 7.07E-07 |
| 2 | 8855061 | T | C | 0.851816 | 1.80E-12 | 7.07E-07 |
| 2 | 8855104 | A | G | 0.851816 | 1.80E-12 | 7.07E-07 |
| 2 | 8855219 | C | T | 0.851816 | 1.80E-12 | 7.07E-07 |
| 2 | 8855591 | C | T | 0.851816 | 1.80E-12 | 7.07E-07 |
| 2 | 8855698 | T | C | 0.851816 | 1.80E-12 | 7.07E-07 |
| 2 | 8855746 | T | C | 0.851816 | 1.80E-12 | 7.07E-07 |
| 2 | 8856018 | T | A | 0.851816 | 1.80E-12 | 7.07E-07 |
| 2 | 8856168 | T | G | 0.851816 | 1.80E-12 | 7.07E-07 |
| 2 | 8856208 | T | G | 0.851816 | 1.80E-12 | 7.07E-07 |
| 2 | 8856542 | C | T | 0.851816 | 1.80E-12 | 7.07E-07 |
| 2 | 8856545 | G | T | 0.851816 | 1.80E-12 | 7.07E-07 |
| 2 | 8856546 | G | T | 0.851816 | 1.80E-12 | 7.07E-07 |
| 2 | 8856576 | C | A | 0.851816 | 1.80E-12 | 7.07E-07 |
| 2 | 8856612 | T | C | 0.851816 | 1.80E-12 | 7.07E-07 |
| 2 | 8856754 | T | C | 0.851816 | 1.80E-12 | 7.07E-07 |
| 2 | 8856799 | C | T | 0.851816 | 1.80E-12 | 7.07E-07 |
| 2 | 8856884 | A | C | 0.851816 | 1.80E-12 | 7.07E-07 |
| 2 | 8857661 | C | T | 0.851816 | 1.80E-12 | 7.07E-07 |
| 2 | 8857787 | C | T | 0.851816 | 1.80E-12 | 7.07E-07 |
| 2 | 8857891 | T | C | 0.851816 | 1.80E-12 | 7.07E-07 |
| 2 | 8857915 | C | T | 0.851816 | 1.80E-12 | 7.07E-07 |
| 2 | 8857934 | T | C | 0.851816 | 1.80E-12 | 7.07E-07 |
| 2 | 8858080 | C | A | 0.851816 | 1.80E-12 | 7.07E-07 |
| 2 | 8858119 | A | G | 0.851816 | 1.80E-12 | 7.07E-07 |
| 2 | 8858433 | T | A | 0.851816 | 1.80E-12 | 7.07E-07 |
| 2 | 8858529 | T | C | 0.851816 | 1.80E-12 | 7.07E-07 |
| 2 | 8859201 | A | G | 0.851816 | 1.80E-12 | 7.07E-07 |
| 2 | 8859276 | A | G | 0.851816 | 1.80E-12 | 7.07E-07 |

| | | | | | | |
|---|---|---|---|---|---|---|
| 2 | 8859569 | C | T | 0.851816 | 1.80E-12 | 7.07E-07 |
| 2 | 8859630 | A | T | 0.851816 | 1.80E-12 | 7.07E-07 |
| 2 | 8859675 | A | G | 0.851816 | 1.80E-12 | 7.07E-07 |
| 2 | 8859796 | T | C | 0.851816 | 1.80E-12 | 7.07E-07 |
| 2 | 8859811 | C | A | 0.851816 | 1.80E-12 | 7.07E-07 |
| 2 | 8860295 | G | T | 0.851816 | 1.80E-12 | 7.07E-07 |
| 2 | 8860369 | G | A | 0.851816 | 1.80E-12 | 7.07E-07 |
| 2 | 8860811 | T | G | 0.851816 | 1.80E-12 | 7.07E-07 |
| 2 | 8860889 | T | G | 0.851816 | 1.80E-12 | 7.07E-07 |
| 2 | 8860993 | C | T | 0.851816 | 1.80E-12 | 7.07E-07 |
| 2 | 8861131 | T | C | 0.851816 | 1.80E-12 | 7.07E-07 |
| 2 | 8861363 | G | A | 0.851816 | 1.80E-12 | 7.07E-07 |
| 2 | 8861427 | A | G | 0.851816 | 1.80E-12 | 7.07E-07 |
| 2 | 8861692 | A | T | 0.851816 | 1.80E-12 | 7.07E-07 |
| 2 | 8862613 | A | T | 0.851816 | 1.80E-12 | 7.07E-07 |
| 2 | 8864134 | G | C | 0.851816 | 1.80E-12 | 7.07E-07 |
| 2 | 8864176 | G | T | 0.851816 | 1.80E-12 | 7.07E-07 |
| 2 | 8864194 | T | C | 0.851816 | 1.80E-12 | 7.07E-07 |
| 2 | 8864199 | G | A | 0.851816 | 1.80E-12 | 7.07E-07 |
| 2 | 8864227 | A | G | 0.851816 | 1.80E-12 | 7.07E-07 |
| 2 | 8864272 | T | C | 0.851816 | 1.80E-12 | 7.07E-07 |
| 2 | 8864506 | T | A | 0.851816 | 1.80E-12 | 7.07E-07 |
| 2 | 8865254 | A | G | 0.851816 | 1.80E-12 | 7.07E-07 |
| 2 | 8865375 | A | G | 0.851816 | 1.80E-12 | 7.07E-07 |
| 2 | 8865504 | C | T | 0.851816 | 1.80E-12 | 7.07E-07 |
| 2 | 8865672 | C | T | 0.851816 | 1.80E-12 | 7.07E-07 |
| 2 | 8865700 | C | T | 0.851816 | 1.80E-12 | 7.07E-07 |
| 2 | 8865997 | C | T | 0.851816 | 1.80E-12 | 7.07E-07 |
| 2 | 8866200 | A | G | 0.851816 | 1.80E-12 | 7.07E-07 |
| 2 | 8866362 | C | T | 0.851816 | 1.80E-12 | 7.07E-07 |
| 2 | 8866876 | T | C | 0.851816 | 1.80E-12 | 7.07E-07 |
| 2 | 8866979 | G | A | 0.851816 | 1.80E-12 | 7.07E-07 |
| 2 | 8867293 | A | C | 0.851816 | 1.80E-12 | 7.07E-07 |
| 2 | 8867978 | A | G | 0.851816 | 1.80E-12 | 7.07E-07 |
| 2 | 8868100 | G | A | 0.851816 | 1.80E-12 | 7.07E-07 |
| 2 | 8868557 | A | T | 0.851816 | 1.80E-12 | 7.07E-07 |
| 2 | 8868718 | C | T | 0.851816 | 1.80E-12 | 7.07E-07 |
| 2 | 8868892 | G | C | 0.851816 | 1.80E-12 | 7.07E-07 |
| 2 | 8868896 | C | T | 0.851816 | 1.80E-12 | 7.07E-07 |
| 2 | 8869036 | A | G | 0.851816 | 1.80E-12 | 7.07E-07 |
| 2 | 8869988 | T | C | 0.851816 | 1.80E-12 | 7.07E-07 |
| 2 | 8870127 | T | G | 0.851816 | 1.80E-12 | 7.07E-07 |
| 2 | 8870290 | G | A | 0.851816 | 1.80E-12 | 7.07E-07 |
| 2 | 8870420 | C | A | 0.851816 | 1.80E-12 | 7.07E-07 |
| 2 | 8870421 | A | T | 0.851816 | 1.80E-12 | 7.07E-07 |
| 2 | 8870542 | T | A | 0.851816 | 1.80E-12 | 7.07E-07 |
| 2 | 8871178 | G | A | 0.851816 | 1.80E-12 | 7.07E-07 |
| 2 | 8871596 | C | T | 0.851816 | 1.80E-12 | 7.07E-07 |
| 2 | 8871699 | C | T | 0.851816 | 1.80E-12 | 7.07E-07 |
| 2 | 8871700 | A | G | 0.851816 | 1.80E-12 | 7.07E-07 |
| 2 | 8871773 | G | C | 0.851816 | 1.80E-12 | 7.07E-07 |
| 2 | 8871848 | C | A | 0.851816 | 1.80E-12 | 7.07E-07 |
| 2 | 8871903 | A | T | 0.851816 | 1.80E-12 | 7.07E-07 |
| 2 | 8871905 | A | T | 0.851816 | 1.80E-12 | 7.07E-07 |
| 2 | 8871996 | C | T | 0.851816 | 1.80E-12 | 7.07E-07 |
| 2 | 8872162 | T | C | 0.851816 | 1.80E-12 | 7.07E-07 |
| 2 | 8872167 | G | A | 0.851816 | 1.80E-12 | 7.07E-07 |
| 2 | 8872190 | T | C | 0.851816 | 1.80E-12 | 7.07E-07 |
| 2 | 8872625 | G | A | 0.851816 | 1.80E-12 | 7.07E-07 |
| 2 | 8872771 | A | G | 0.851816 | 1.80E-12 | 7.07E-07 |
| 2 | 8872850 | C | A | 0.851816 | 1.80E-12 | 7.07E-07 |
| 2 | 8873146 | G | A | 0.851816 | 1.80E-12 | 7.07E-07 |
| 2 | 8873558 | C | T | 0.851816 | 1.80E-12 | 7.07E-07 |
| 2 | 8873747 | T | C | 0.851816 | 1.80E-12 | 7.07E-07 |
| 2 | 8875417 | G | A | 0.851816 | 1.80E-12 | 7.07E-07 |
| 2 | 8875501 | A | G | 0.851816 | 1.80E-12 | 7.07E-07 |
| 2 | 8875551 | G | A | 0.851816 | 1.80E-12 | 7.07E-07 |

| | | | | | | |
|---|---|---|---|---|---|---|
| 2 | 8875556 | A | G | 0.851816 | 1.80E-12 | 7.07E-07 |
| 2 | 8875637 | A | C | 0.851816 | 1.80E-12 | 7.07E-07 |
| 2 | 8875871 | T | G | 0.851816 | 1.80E-12 | 7.07E-07 |
| 2 | 8876132 | C | A | 0.851816 | 1.80E-12 | 7.07E-07 |
| 2 | 8876949 | C | T | 0.851816 | 1.80E-12 | 7.07E-07 |
| 2 | 8877654 | G | A | 0.851816 | 1.80E-12 | 7.07E-07 |
| 2 | 8877825 | G | C | 0.851816 | 1.80E-12 | 7.07E-07 |
| 2 | 8877905 | A | G | 0.851816 | 1.80E-12 | 7.07E-07 |
| 2 | 8878070 | T | C | 0.851816 | 1.80E-12 | 7.07E-07 |
| 2 | 8878657 | C | T | 0.851816 | 1.80E-12 | 7.07E-07 |
| 2 | 8880199 | T | C | 0.851816 | 1.80E-12 | 7.07E-07 |
| 2 | 8880361 | G | A | 0.851816 | 1.80E-12 | 7.07E-07 |
| 2 | 8880638 | G | A | 0.851816 | 1.80E-12 | 7.07E-07 |
| 2 | 8881296 | G | C | 0.851816 | 1.80E-12 | 7.07E-07 |
| 2 | 8881424 | C | T | 0.851816 | 1.80E-12 | 7.07E-07 |
| 2 | 8881504 | G | A | 0.851816 | 1.80E-12 | 7.07E-07 |
| 2 | 8881508 | G | A | 0.851816 | 1.80E-12 | 7.07E-07 |
| 2 | 8881613 | G | A | 0.851816 | 1.80E-12 | 7.07E-07 |
| 2 | 8881766 | A | G | 0.851816 | 1.80E-12 | 7.07E-07 |
| 2 | 8881768 | C | A | 0.851816 | 1.80E-12 | 7.07E-07 |
| 2 | 8882606 | A | C | 0.851816 | 1.80E-12 | 7.07E-07 |
| 2 | 8883141 | T | C | 0.851816 | 1.80E-12 | 7.07E-07 |
| 2 | 8883333 | G | A | 0.851816 | 1.80E-12 | 7.07E-07 |
| 2 | 8883793 | C | A | 0.851816 | 1.80E-12 | 7.07E-07 |
| 2 | 8883983 | G | T | 0.851816 | 1.80E-12 | 7.07E-07 |
| 2 | 8884766 | A | G | 0.851816 | 1.80E-12 | 7.07E-07 |
| 2 | 8884809 | T | C | 0.851816 | 1.80E-12 | 7.07E-07 |
| 2 | 8886216 | A | G | 0.851816 | 1.80E-12 | 7.07E-07 |
| 2 | 8886379 | A | G | 0.851816 | 1.80E-12 | 7.07E-07 |
| 2 | 8886415 | G | A | 0.851816 | 1.80E-12 | 7.07E-07 |
| 2 | 8886418 | C | T | 0.851816 | 1.80E-12 | 7.07E-07 |
| 2 | 8886776 | T | C | 0.851816 | 1.80E-12 | 7.07E-07 |
| 2 | 8887350 | A | G | 0.851816 | 1.80E-12 | 7.07E-07 |
| 2 | 8888135 | A | G | 0.851816 | 1.80E-12 | 7.07E-07 |
| 2 | 8888178 | A | C | 0.851816 | 1.80E-12 | 7.07E-07 |
| 2 | 8888842 | G | A | 0.851816 | 1.80E-12 | 7.07E-07 |
| 2 | 8888892 | C | T | 0.851816 | 1.80E-12 | 7.07E-07 |
| 2 | 8889257 | A | G | 0.851816 | 1.80E-12 | 7.07E-07 |
| 2 | 8889359 | C | T | 0.851816 | 1.80E-12 | 7.07E-07 |
| 2 | 8889668 | G | A | 0.851816 | 1.80E-12 | 7.07E-07 |
| 2 | 8889768 | T | C | 0.851816 | 1.80E-12 | 7.07E-07 |
| 2 | 8891548 | C | A | 0.851816 | 1.80E-12 | 7.07E-07 |
| 2 | 8891647 | T | C | 0.851816 | 1.80E-12 | 7.07E-07 |
| 2 | 8891805 | A | G | 0.851816 | 1.80E-12 | 7.07E-07 |
| 2 | 8891889 | T | A | 0.851816 | 1.80E-12 | 7.07E-07 |
| 2 | 8892133 | T | C | 0.851816 | 1.80E-12 | 7.07E-07 |
| 2 | 8892279 | G | A | 0.851816 | 1.80E-12 | 7.07E-07 |
| 2 | 8892385 | T | C | 0.851816 | 1.80E-12 | 7.07E-07 |
| 2 | 8892525 | C | T | 0.851816 | 1.80E-12 | 7.07E-07 |
| 2 | 8892757 | C | T | 0.851816 | 1.80E-12 | 7.07E-07 |
| 2 | 8892828 | T | C | 0.851816 | 1.80E-12 | 7.07E-07 |
| 2 | 8892959 | G | A | 0.851816 | 1.80E-12 | 7.07E-07 |
| 2 | 8894140 | C | A | 0.851816 | 1.80E-12 | 7.07E-07 |
| 2 | 8894170 | C | T | 0.851816 | 1.80E-12 | 7.07E-07 |
| 2 | 8894177 | C | A | 0.851816 | 1.80E-12 | 7.07E-07 |
| 2 | 8894218 | T | G | 0.851816 | 1.80E-12 | 7.07E-07 |
| 2 | 8894706 | G | A | 0.851816 | 1.80E-12 | 7.07E-07 |
| 2 | 8894779 | T | C | 0.851816 | 1.80E-12 | 7.07E-07 |
| 2 | 8895204 | C | G | 0.851816 | 1.80E-12 | 7.07E-07 |
| 2 | 8895267 | T | C | 0.851816 | 1.80E-12 | 7.07E-07 |
| 2 | 8896384 | A | T | 0.851816 | 1.80E-12 | 7.07E-07 |
| 2 | 8896488 | T | C | 0.851816 | 1.80E-12 | 7.07E-07 |
| 2 | 8896581 | G | A | 0.851816 | 1.80E-12 | 7.07E-07 |
| 2 | 8896603 | A | T | 0.851816 | 1.80E-12 | 7.07E-07 |
| 2 | 8896604 | G | A | 0.851816 | 1.80E-12 | 7.07E-07 |
| 2 | 8896605 | C | A | 0.851816 | 1.80E-12 | 7.07E-07 |
| 2 | 8896840 | T | C | 0.851816 | 1.80E-12 | 7.07E-07 |

| | | | | | | |
|---|---|---|---|---|---|---|
| 2 | 8897000 | A | G | 0.851816 | 1.80E-12 | 7.07E-07 |
| 2 | 8897046 | T | C | 0.851816 | 1.80E-12 | 7.07E-07 |
| 2 | 8897124 | T | G | 0.851816 | 1.80E-12 | 7.07E-07 |
| 2 | 8897176 | A | G | 0.851816 | 1.80E-12 | 7.07E-07 |
| 2 | 8897891 | G | A | 0.851816 | 1.80E-12 | 7.07E-07 |
| 2 | 8897912 | A | C | 0.851816 | 1.80E-12 | 7.07E-07 |
| 2 | 8898001 | T | A | 0.851816 | 1.80E-12 | 7.07E-07 |
| 2 | 8898004 | A | C | 0.851816 | 1.80E-12 | 7.07E-07 |
| 2 | 8898060 | C | T | 0.851816 | 1.80E-12 | 7.07E-07 |
| 2 | 8899001 | A | G | 0.851816 | 1.80E-12 | 7.07E-07 |
| 2 | 8899054 | T | C | 0.851816 | 1.80E-12 | 7.07E-07 |
| 2 | 8899055 | G | A | 0.851816 | 1.80E-12 | 7.07E-07 |
| 2 | 8899110 | G | A | 0.851816 | 1.80E-12 | 7.07E-07 |
| 2 | 8899165 | T | C | 0.851816 | 1.80E-12 | 7.07E-07 |
| 2 | 8899213 | C | T | 0.851816 | 1.80E-12 | 7.07E-07 |
| 2 | 8899217 | G | A | 0.851816 | 1.80E-12 | 7.07E-07 |
| 2 | 8899271 | A | G | 0.851816 | 1.80E-12 | 7.07E-07 |
| 2 | 8900029 | C | A | 0.851816 | 1.80E-12 | 7.07E-07 |
| 2 | 8900081 | G | A | 0.851816 | 1.80E-12 | 7.07E-07 |
| 2 | 8900100 | G | A | 0.851816 | 1.80E-12 | 7.07E-07 |
| 2 | 8900146 | G | C | 0.851816 | 1.80E-12 | 7.07E-07 |
| 2 | 8900152 | T | C | 0.851816 | 1.80E-12 | 7.07E-07 |
| 2 | 8900168 | A | G | 0.851816 | 1.80E-12 | 7.07E-07 |
| 2 | 8900194 | C | G | 0.851816 | 1.80E-12 | 7.07E-07 |
| 2 | 8900215 | G | A | 0.851816 | 1.80E-12 | 7.07E-07 |
| 2 | 8900246 | C | T | 0.851816 | 1.80E-12 | 7.07E-07 |
| 2 | 8900276 | A | G | 0.851816 | 1.80E-12 | 7.07E-07 |
| 2 | 8900310 | A | T | 0.851816 | 1.80E-12 | 7.07E-07 |
| 2 | 8900328 | C | A | 0.851816 | 1.80E-12 | 7.07E-07 |
| 2 | 8900349 | C | T | 0.851816 | 1.80E-12 | 7.07E-07 |
| 2 | 8900372 | T | C | 0.851816 | 1.80E-12 | 7.07E-07 |
| 2 | 8900412 | G | A | 0.851816 | 1.80E-12 | 7.07E-07 |
| 2 | 8900442 | G | A | 0.851816 | 1.80E-12 | 7.07E-07 |
| 2 | 8900498 | G | A | 0.851816 | 1.80E-12 | 7.07E-07 |
| 2 | 8900531 | C | T | 0.851816 | 1.80E-12 | 7.07E-07 |
| 2 | 8900573 | C | T | 0.851816 | 1.80E-12 | 7.07E-07 |
| 2 | 8900580 | C | T | 0.851816 | 1.80E-12 | 7.07E-07 |
| 2 | 8900643 | A | G | 0.851816 | 1.80E-12 | 7.07E-07 |
| 2 | 8900713 | C | T | 0.851816 | 1.80E-12 | 7.07E-07 |
| 2 | 8900973 | A | G | 0.851816 | 1.80E-12 | 7.07E-07 |
| 2 | 8901022 | G | A | 0.851816 | 1.80E-12 | 7.07E-07 |
| 2 | 8901033 | A | C | 0.851816 | 1.80E-12 | 7.07E-07 |
| 2 | 8901262 | G | T | 0.851816 | 1.80E-12 | 7.07E-07 |
| 2 | 8901447 | A | G | 0.851816 | 1.80E-12 | 7.07E-07 |
| 2 | 8901459 | T | C | 0.851816 | 1.80E-12 | 7.07E-07 |
| 2 | 8901618 | C | T | 0.851816 | 1.80E-12 | 7.07E-07 |
| 2 | 8901689 | C | T | 0.851816 | 1.80E-12 | 7.07E-07 |
| 2 | 8901694 | G | C | 0.851816 | 1.80E-12 | 7.07E-07 |
| 2 | 8901752 | C | T | 0.851816 | 1.80E-12 | 7.07E-07 |
| 2 | 8901851 | C | T | 0.851816 | 1.80E-12 | 7.07E-07 |
| 2 | 8901926 | G | T | 0.851816 | 1.80E-12 | 7.07E-07 |
| 2 | 8901959 | G | A | 0.851816 | 1.80E-12 | 7.07E-07 |
| 2 | 8902495 | A | C | 0.851816 | 1.80E-12 | 7.07E-07 |
| 2 | 8902512 | G | T | 0.851816 | 1.80E-12 | 7.07E-07 |
| 2 | 8902524 | T | C | 0.851816 | 1.80E-12 | 7.07E-07 |
| 2 | 8902525 | A | T | 0.851816 | 1.80E-12 | 7.07E-07 |
| 2 | 8902652 | G | A | 0.851816 | 1.80E-12 | 7.07E-07 |
| 2 | 8902718 | G | A | 0.851816 | 1.80E-12 | 7.07E-07 |
| 2 | 8902787 | G | C | 0.851816 | 1.80E-12 | 7.07E-07 |
| 2 | 8902804 | G | A | 0.851816 | 1.80E-12 | 7.07E-07 |
| 2 | 8903227 | C | A | 0.851816 | 1.80E-12 | 7.07E-07 |
| 2 | 8903316 | G | C | 0.851816 | 1.80E-12 | 7.07E-07 |
| 2 | 8903379 | C | A | 0.851816 | 1.80E-12 | 7.07E-07 |
| 2 | 8903407 | T | A | 0.851816 | 1.80E-12 | 7.07E-07 |
| 2 | 8903544 | A | T | 0.851816 | 1.80E-12 | 7.07E-07 |
| 2 | 8903696 | T | G | 0.851816 | 1.80E-12 | 7.07E-07 |
| 2 | 8903783 | G | A | 0.851816 | 1.80E-12 | 7.07E-07 |

| | | | | | | |
|---|---|---|---|---|---|---|
| 2 | 8903980 | A | G | 0.851816 | 1.80E-12 | 7.07E-07 |
| 2 | 8904612 | A | C | 0.851816 | 1.80E-12 | 7.07E-07 |
| 2 | 8904904 | C | T | 0.851816 | 1.80E-12 | 7.07E-07 |
| 2 | 8905125 | C | T | 0.851816 | 1.80E-12 | 7.07E-07 |
| 2 | 8905174 | A | G | 0.851816 | 1.80E-12 | 7.07E-07 |
| 2 | 8905226 | G | C | 0.851816 | 1.80E-12 | 7.07E-07 |
| 2 | 8905322 | C | T | 0.851816 | 1.80E-12 | 7.07E-07 |
| 2 | 8905343 | C | T | 0.851816 | 1.80E-12 | 7.07E-07 |
| 2 | 8905388 | G | C | 0.851816 | 1.80E-12 | 7.07E-07 |
| 2 | 8905730 | T | C | 0.851816 | 1.80E-12 | 7.07E-07 |
| 2 | 8905763 | G | A | 0.851816 | 1.80E-12 | 7.07E-07 |
| 2 | 8905765 | T | C | 0.851816 | 1.80E-12 | 7.07E-07 |
| 2 | 8905861 | T | C | 0.851816 | 1.80E-12 | 7.07E-07 |
| 2 | 8905925 | G | A | 0.851816 | 1.80E-12 | 7.07E-07 |
| 2 | 8905969 | G | A | 0.851816 | 1.80E-12 | 7.07E-07 |
| 2 | 8906036 | G | T | 0.851816 | 1.80E-12 | 7.07E-07 |
| 2 | 8906060 | T | C | 0.851816 | 1.80E-12 | 7.07E-07 |
| 2 | 8906121 | T | C | 0.851816 | 1.80E-12 | 7.07E-07 |
| 2 | 8906530 | C | T | 0.851816 | 1.80E-12 | 7.07E-07 |
| 2 | 8911310 | A | G | 0.851816 | 1.80E-12 | 7.07E-07 |
| 2 | 8911387 | G | T | 0.851816 | 1.80E-12 | 7.07E-07 |
| 2 | 8911449 | T | C | 0.851816 | 1.80E-12 | 7.07E-07 |
| 2 | 8911498 | C | G | 0.851816 | 1.80E-12 | 7.07E-07 |
| 2 | 8911520 | T | C | 0.851816 | 1.80E-12 | 7.07E-07 |
| 2 | 8911522 | T | C | 0.851816 | 1.80E-12 | 7.07E-07 |
| 2 | 8911659 | A | T | 0.851816 | 1.80E-12 | 7.07E-07 |
| 2 | 8911666 | C | T | 0.851816 | 1.80E-12 | 7.07E-07 |
| 2 | 8911667 | G | T | 0.851816 | 1.80E-12 | 7.07E-07 |
| 2 | 8911748 | G | T | 0.851816 | 1.80E-12 | 7.07E-07 |
| 2 | 8911773 | T | C | 0.851816 | 1.80E-12 | 7.07E-07 |
| 2 | 8911819 | A | G | 0.851816 | 1.80E-12 | 7.07E-07 |
| 2 | 8911836 | T | A | 0.851816 | 1.80E-12 | 7.07E-07 |
| 2 | 8911852 | A | T | 0.851816 | 1.80E-12 | 7.07E-07 |
| 2 | 8911859 | A | C | 0.851816 | 1.80E-12 | 7.07E-07 |
| 2 | 8912236 | T | C | 0.851816 | 1.80E-12 | 7.07E-07 |
| 2 | 8912340 | G | A | 0.851816 | 1.80E-12 | 7.07E-07 |
| 2 | 8912481 | T | C | 0.851816 | 1.80E-12 | 7.07E-07 |
| 2 | 8912529 | C | T | 0.851816 | 1.80E-12 | 7.07E-07 |
| 2 | 8912588 | C | T | 0.851816 | 1.80E-12 | 7.07E-07 |
| 2 | 8912664 | C | T | 0.851816 | 1.80E-12 | 7.07E-07 |
| 2 | 8912665 | A | G | 0.851816 | 1.80E-12 | 7.07E-07 |
| 2 | 8912703 | T | C | 0.851816 | 1.80E-12 | 7.07E-07 |
| 2 | 8912789 | A | G | 0.851816 | 1.80E-12 | 7.07E-07 |
| 2 | 8912808 | A | T | 0.851816 | 1.80E-12 | 7.07E-07 |
| 2 | 8912868 | G | A | 0.851816 | 1.80E-12 | 7.07E-07 |
| 2 | 8912920 | T | A | 0.851816 | 1.80E-12 | 7.07E-07 |
| 2 | 8913013 | T | C | 0.851816 | 1.80E-12 | 7.07E-07 |
| 2 | 8913229 | T | C | 0.851816 | 1.80E-12 | 7.07E-07 |
| 2 | 8913231 | C | T | 0.851816 | 1.80E-12 | 7.07E-07 |
| 2 | 8913587 | C | T | 0.851816 | 1.80E-12 | 7.07E-07 |
| 2 | 8913665 | T | C | 0.851816 | 1.80E-12 | 7.07E-07 |
| 2 | 8913780 | T | A | 0.851816 | 1.80E-12 | 7.07E-07 |
| 2 | 8913876 | T | C | 0.851816 | 1.80E-12 | 7.07E-07 |
| 2 | 8914083 | A | C | 0.851816 | 1.80E-12 | 7.07E-07 |
| 2 | 8914097 | C | A | 0.851816 | 1.80E-12 | 7.07E-07 |
| 2 | 8914106 | G | A | 0.851816 | 1.80E-12 | 7.07E-07 |
| 2 | 8914189 | C | T | 0.851816 | 1.80E-12 | 7.07E-07 |
| 2 | 8914218 | C | T | 0.851816 | 1.80E-12 | 7.07E-07 |
| 2 | 8914302 | G | A | 0.851816 | 1.80E-12 | 7.07E-07 |
| 2 | 8914328 | G | A | 0.851816 | 1.80E-12 | 7.07E-07 |
| 2 | 8914371 | G | T | 0.851816 | 1.80E-12 | 7.07E-07 |
| 2 | 8914450 | A | T | 0.851816 | 1.80E-12 | 7.07E-07 |
| 2 | 8914601 | G | A | 0.851816 | 1.80E-12 | 7.07E-07 |
| 2 | 8914731 | A | G | 0.851816 | 1.80E-12 | 7.07E-07 |
| 2 | 8914845 | A | G | 0.851816 | 1.80E-12 | 7.07E-07 |
| 2 | 8914846 | C | T | 0.851816 | 1.80E-12 | 7.07E-07 |
| 2 | 8915026 | G | A | 0.851816 | 1.80E-12 | 7.07E-07 |

| | | | | | | |
|---|---|---|---|---|---|---|
| 2 | 8915110 | A | G | 0.851816 | 1.80E-12 | 7.07E-07 |
| 2 | 8915311 | T | C | 0.851816 | 1.80E-12 | 7.07E-07 |
| 2 | 8915350 | C | T | 0.851816 | 1.80E-12 | 7.07E-07 |
| 2 | 8915359 | T | A | 0.851816 | 1.80E-12 | 7.07E-07 |
| 2 | 8915774 | C | T | 0.851816 | 1.80E-12 | 7.07E-07 |
| 2 | 8915798 | G | A | 0.851816 | 1.80E-12 | 7.07E-07 |
| 2 | 8915814 | A | T | 0.851816 | 1.80E-12 | 7.07E-07 |
| 2 | 8915849 | C | T | 0.851816 | 1.80E-12 | 7.07E-07 |
| 2 | 8915851 | C | T | 0.851816 | 1.80E-12 | 7.07E-07 |
| 2 | 8916284 | G | A | 0.851816 | 1.80E-12 | 7.07E-07 |
| 2 | 8916322 | C | A | 0.851816 | 1.80E-12 | 7.07E-07 |
| 2 | 8916323 | A | G | 0.851816 | 1.80E-12 | 7.07E-07 |
| 2 | 8916328 | A | G | 0.851816 | 1.80E-12 | 7.07E-07 |
| 2 | 8916450 | T | C | 0.851816 | 1.80E-12 | 7.07E-07 |
| 2 | 8916832 | T | C | 0.851816 | 1.80E-12 | 7.07E-07 |
| 2 | 8916839 | A | G | 0.851816 | 1.80E-12 | 7.07E-07 |
| 2 | 8917106 | G | A | 0.851816 | 1.80E-12 | 7.07E-07 |
| 2 | 8917147 | T | C | 0.851816 | 1.80E-12 | 7.07E-07 |
| 2 | 8917149 | G | A | 0.851816 | 1.80E-12 | 7.07E-07 |
| 2 | 8917384 | T | A | 0.851816 | 1.80E-12 | 7.07E-07 |
| 2 | 8917446 | A | G | 0.851816 | 1.80E-12 | 7.07E-07 |
| 2 | 8917468 | A | G | 0.851816 | 1.80E-12 | 7.07E-07 |
| 2 | 8917481 | C | T | 0.851816 | 1.80E-12 | 7.07E-07 |
| 2 | 8917509 | G | A | 0.851816 | 1.80E-12 | 7.07E-07 |
| 2 | 8917530 | C | A | 0.851816 | 1.80E-12 | 7.07E-07 |
| 2 | 8917546 | A | G | 0.851816 | 1.80E-12 | 7.07E-07 |
| 2 | 8917579 | G | A | 0.851816 | 1.80E-12 | 7.07E-07 |
| 2 | 8917634 | G | A | 0.851816 | 1.80E-12 | 7.07E-07 |
| 2 | 8917650 | C | A | 0.851816 | 1.80E-12 | 7.07E-07 |
| 2 | 8917651 | T | C | 0.851816 | 1.80E-12 | 7.07E-07 |
| 2 | 8917701 | C | T | 0.851816 | 1.80E-12 | 7.07E-07 |
| 2 | 8917706 | C | T | 0.851816 | 1.80E-12 | 7.07E-07 |
| 2 | 8917715 | A | G | 0.851816 | 1.80E-12 | 7.07E-07 |
| 2 | 8917791 | C | G | 0.851816 | 1.80E-12 | 7.07E-07 |
| 2 | 8917911 | C | T | 0.851816 | 1.80E-12 | 7.07E-07 |
| 2 | 8918135 | C | T | 0.851816 | 1.80E-12 | 7.07E-07 |
| 2 | 8918156 | C | T | 0.851816 | 1.80E-12 | 7.07E-07 |
| 2 | 8918217 | A | T | 0.851816 | 1.80E-12 | 7.07E-07 |
| 2 | 8918303 | G | A | 0.851816 | 1.80E-12 | 7.07E-07 |
| 2 | 8918324 | T | C | 0.851816 | 1.80E-12 | 7.07E-07 |
| 2 | 8918881 | A | G | 0.851816 | 1.80E-12 | 7.07E-07 |
| 2 | 8918890 | C | T | 0.851816 | 1.80E-12 | 7.07E-07 |
| 2 | 8918916 | A | C | 0.851816 | 1.80E-12 | 7.07E-07 |
| 2 | 8918953 | A | T | 0.851816 | 1.80E-12 | 7.07E-07 |
| 2 | 8918964 | A | T | 0.851816 | 1.80E-12 | 7.07E-07 |
| 2 | 8919035 | G | A | 0.851816 | 1.80E-12 | 7.07E-07 |
| 2 | 8919318 | A | G | 0.851816 | 1.80E-12 | 7.07E-07 |
| 2 | 8919324 | G | C | 0.851816 | 1.80E-12 | 7.07E-07 |
| 2 | 8919360 | G | A | 0.851816 | 1.80E-12 | 7.07E-07 |
| 2 | 8919452 | G | T | 0.851816 | 1.80E-12 | 7.07E-07 |
| 2 | 8919618 | G | C | 0.851816 | 1.80E-12 | 7.07E-07 |
| 2 | 8919723 | G | A | 0.851816 | 1.80E-12 | 7.07E-07 |
| 2 | 8919769 | T | A | 0.851816 | 1.80E-12 | 7.07E-07 |
| 2 | 8919824 | A | T | 0.851816 | 1.80E-12 | 7.07E-07 |
| 2 | 8919950 | G | A | 0.851816 | 1.80E-12 | 7.07E-07 |
| 2 | 8920003 | A | C | 0.851816 | 1.80E-12 | 7.07E-07 |
| 2 | 8920036 | G | T | 0.851816 | 1.80E-12 | 7.07E-07 |
| 2 | 8920644 | A | G | 0.851816 | 1.80E-12 | 7.07E-07 |
| 2 | 8920795 | T | C | 0.851816 | 1.80E-12 | 7.07E-07 |
| 2 | 8920813 | T | C | 0.851816 | 1.80E-12 | 7.07E-07 |
| 2 | 8920901 | A | G | 0.851816 | 1.80E-12 | 7.07E-07 |
| 2 | 8921008 | A | C | 0.851816 | 1.80E-12 | 7.07E-07 |
| 2 | 8921069 | T | C | 0.851816 | 1.80E-12 | 7.07E-07 |
| 2 | 8921071 | T | C | 0.851816 | 1.80E-12 | 7.07E-07 |
| 2 | 8921103 | T | C | 0.851816 | 1.80E-12 | 7.07E-07 |
| 2 | 8921108 | T | C | 0.851816 | 1.80E-12 | 7.07E-07 |
| 2 | 8921128 | C | T | 0.851816 | 1.80E-12 | 7.07E-07 |

| | | | | | | |
|---|---|---|---|---|---|---|
| 2 | 8921218 | G | C | 0.851816 | 1.80E-12 | 7.07E-07 |
| 2 | 8921271 | C | T | 0.851816 | 1.80E-12 | 7.07E-07 |
| 2 | 8921300 | A | G | 0.851816 | 1.80E-12 | 7.07E-07 |
| 2 | 8921386 | A | T | 0.851816 | 1.80E-12 | 7.07E-07 |
| 2 | 8921464 | T | G | 0.851816 | 1.80E-12 | 7.07E-07 |
| 2 | 8921481 | C | G | 0.851816 | 1.80E-12 | 7.07E-07 |
| 2 | 8921537 | G | A | 0.851816 | 1.80E-12 | 7.07E-07 |
| 2 | 8921577 | C | T | 0.851816 | 1.80E-12 | 7.07E-07 |
| 2 | 8921627 | A | G | 0.851816 | 1.80E-12 | 7.07E-07 |
| 2 | 8921842 | T | C | 0.851816 | 1.80E-12 | 7.07E-07 |
| 2 | 8921866 | G | A | 0.851816 | 1.80E-12 | 7.07E-07 |
| 2 | 8921882 | C | T | 0.851816 | 1.80E-12 | 7.07E-07 |
| 2 | 8921926 | G | A | 0.851816 | 1.80E-12 | 7.07E-07 |
| 2 | 8921957 | A | G | 0.851816 | 1.80E-12 | 7.07E-07 |
| 2 | 8922036 | G | C | 0.851816 | 1.80E-12 | 7.07E-07 |
| 2 | 8922099 | A | C | 0.851816 | 1.80E-12 | 7.07E-07 |
| 2 | 8922293 | A | T | 0.851816 | 1.80E-12 | 7.07E-07 |
| 2 | 8850504 | T | G | 0.852772 | 2.19E-12 | 8.59E-07 |
| 2 | 8850542 | G | T | 0.852772 | 2.19E-12 | 8.59E-07 |
| 2 | 8850794 | A | T | 0.852772 | 2.19E-12 | 8.59E-07 |
| 2 | 8833968 | A | G | 0.85086 | 2.22E-12 | 8.69E-07 |
| 2 | 8816257 | C | T | 0.849904 | 2.88E-12 | 1.13E-06 |
| 2 | 8907176 | G | C | 0.853728 | 3.22E-12 | 1.26E-06 |
| 2 | 8907260 | T | C | 0.853728 | 3.22E-12 | 1.26E-06 |
| 2 | 8907277 | C | T | 0.853728 | 3.22E-12 | 1.26E-06 |
| 2 | 8893117 | A | T | 0.852772 | 3.36E-12 | 1.32E-06 |
| 2 | 8893155 | C | A | 0.852772 | 3.36E-12 | 1.32E-06 |
| 2 | 8893245 | A | G | 0.852772 | 3.36E-12 | 1.32E-06 |
| 2 | 8893440 | A | T | 0.852772 | 3.36E-12 | 1.32E-06 |
| 2 | 8893875 | C | T | 0.852772 | 3.36E-12 | 1.32E-06 |
| 2 | 8893883 | A | C | 0.852772 | 3.36E-12 | 1.32E-06 |
| 2 | 8893922 | A | C | 0.852772 | 3.36E-12 | 1.32E-06 |
| 2 | 8893978 | A | C | 0.852772 | 3.36E-12 | 1.32E-06 |
| 2 | 8849175 | C | T | 0.84608 | 3.39E-12 | 1.33E-06 |
| 2 | 8890755 | A | G | 0.852772 | 3.49E-12 | 1.37E-06 |
| 2 | 8890916 | C | A | 0.852772 | 3.49E-12 | 1.37E-06 |
| 2 | 8890934 | T | C | 0.852772 | 3.49E-12 | 1.37E-06 |
| 2 | 8890945 | T | C | 0.852772 | 3.49E-12 | 1.37E-06 |
| 2 | 8891092 | A | G | 0.852772 | 3.49E-12 | 1.37E-06 |
| 2 | 8891460 | T | A | 0.852772 | 3.49E-12 | 1.37E-06 |
| 2 | 8787865 | G | A | 0.85086 | 3.58E-12 | 1.40E-06 |
| 2 | 8788847 | C | T | 0.85086 | 3.58E-12 | 1.40E-06 |
| 2 | 8789498 | G | A | 0.85086 | 3.58E-12 | 1.40E-06 |
| 2 | 8789518 | G | A | 0.85086 | 3.58E-12 | 1.40E-06 |
| 2 | 8789582 | A | C | 0.85086 | 3.58E-12 | 1.40E-06 |
| 2 | 8794423 | T | G | 0.85086 | 3.58E-12 | 1.40E-06 |
| 2 | 8794526 | T | C | 0.85086 | 3.58E-12 | 1.40E-06 |
| 2 | 8794532 | G | C | 0.85086 | 3.58E-12 | 1.40E-06 |
| 2 | 8794587 | A | G | 0.85086 | 3.58E-12 | 1.40E-06 |
| 2 | 8797780 | C | A | 0.85086 | 3.58E-12 | 1.40E-06 |
| 2 | 8797784 | A | G | 0.85086 | 3.58E-12 | 1.40E-06 |
| 2 | 8797821 | G | A | 0.85086 | 3.58E-12 | 1.40E-06 |
| 2 | 8797982 | G | A | 0.85086 | 3.58E-12 | 1.40E-06 |
| 2 | 8798830 | T | C | 0.85086 | 3.58E-12 | 1.40E-06 |
| 2 | 8798896 | C | T | 0.85086 | 3.58E-12 | 1.40E-06 |
| 2 | 8800772 | T | A | 0.85086 | 3.58E-12 | 1.40E-06 |
| 2 | 8805109 | T | A | 0.85086 | 3.58E-12 | 1.40E-06 |
| 2 | 8911272 | C | T | 0.857553 | 3.59E-12 | 1.41E-06 |
| 2 | 8911273 | A | G | 0.857553 | 3.59E-12 | 1.41E-06 |
| 2 | 8882157 | C | T | 0.849904 | 4.12E-12 | 1.61E-06 |
| 2 | 8776552 | G | A | 0.848948 | 4.33E-12 | 1.70E-06 |
| 2 | 8776650 | G | A | 0.848948 | 4.33E-12 | 1.70E-06 |
| 2 | 8776658 | A | G | 0.848948 | 4.33E-12 | 1.70E-06 |
| 2 | 8776666 | A | G | 0.848948 | 4.33E-12 | 1.70E-06 |
| 2 | 8896276 | G | C | 0.852772 | 4.58E-12 | 1.80E-06 |
| 2 | 8896680 | A | G | 0.852772 | 4.58E-12 | 1.80E-06 |
| 2 | 8897527 | G | C | 0.852772 | 4.58E-12 | 1.80E-06 |

| | | | | | | |
|---|---|---|---|---|---|---|
| 2 | 8898291 | A | T | 0.852772 | 4.58E-12 | 1.80E-06 |
| 2 | 8898335 | A | G | 0.852772 | 4.58E-12 | 1.80E-06 |
| 2 | 8898363 | G | A | 0.852772 | 4.58E-12 | 1.80E-06 |
| 2 | 8898387 | A | G | 0.852772 | 4.58E-12 | 1.80E-06 |
| 2 | 8898584 | C | T | 0.852772 | 4.58E-12 | 1.80E-06 |
| 2 | 8898616 | C | A | 0.852772 | 4.58E-12 | 1.80E-06 |
| 2 | 8898642 | G | A | 0.852772 | 4.58E-12 | 1.80E-06 |
| 2 | 8898649 | C | T | 0.852772 | 4.58E-12 | 1.80E-06 |
| 2 | 8898664 | A | G | 0.852772 | 4.58E-12 | 1.80E-06 |
| 2 | 8898675 | T | C | 0.852772 | 4.58E-12 | 1.80E-06 |
| 2 | 8898708 | C | A | 0.852772 | 4.58E-12 | 1.80E-06 |
| 2 | 8898900 | T | C | 0.852772 | 4.58E-12 | 1.80E-06 |
| 2 | 8898925 | T | G | 0.852772 | 4.58E-12 | 1.80E-06 |
| 2 | 8898939 | T | C | 0.852772 | 4.58E-12 | 1.80E-06 |
| 2 | 8898947 | C | T | 0.852772 | 4.58E-12 | 1.80E-06 |
| 2 | 8899273 | A | T | 0.852772 | 4.58E-12 | 1.80E-06 |
| 2 | 8899327 | G | A | 0.852772 | 4.58E-12 | 1.80E-06 |
| 2 | 8899441 | A | C | 0.852772 | 4.58E-12 | 1.80E-06 |
| 2 | 8899491 | A | G | 0.852772 | 4.58E-12 | 1.80E-06 |
| 2 | 8899876 | A | G | 0.852772 | 4.58E-12 | 1.80E-06 |
| 2 | 8899915 | C | T | 0.852772 | 4.58E-12 | 1.80E-06 |
| 2 | 8899916 | A | G | 0.852772 | 4.58E-12 | 1.80E-06 |
| 2 | 8899956 | A | G | 0.852772 | 4.58E-12 | 1.80E-06 |
| 2 | 8895333 | G | C | 0.852772 | 5.04E-12 | 1.97E-06 |
| 2 | 8895493 | A | G | 0.852772 | 5.04E-12 | 1.97E-06 |
| 2 | 8910847 | C | A | 0.859465 | 5.13E-12 | 2.01E-06 |
| 2 | 8910867 | C | T | 0.859465 | 5.13E-12 | 2.01E-06 |
| 2 | 8910868 | A | G | 0.859465 | 5.13E-12 | 2.01E-06 |
| 2 | 8910918 | G | C | 0.859465 | 5.13E-12 | 2.01E-06 |
| 2 | 8875246 | C | A | 0.853728 | 6.47E-12 | 2.54E-06 |
| 2 | 8841688 | G | C | 0.85086 | 8.69E-12 | 3.41E-06 |
| 2 | 8841753 | G | A | 0.85086 | 8.69E-12 | 3.41E-06 |
| 2 | 8864700 | G | A | 0.85086 | 8.76E-12 | 3.43E-06 |
| 2 | 11262235 | C | T | 0.840344 | 1.12E-11 | 4.38E-06 |
| 2 | 11262261 | A | G | 0.840344 | 1.12E-11 | 4.38E-06 |
| 2 | 11262316 | T | C | 0.840344 | 1.12E-11 | 4.38E-06 |
| 2 | 11262346 | C | G | 0.840344 | 1.12E-11 | 4.38E-06 |
| 2 | 11262370 | A | G | 0.840344 | 1.12E-11 | 4.38E-06 |
| 2 | 11262379 | G | A | 0.840344 | 1.12E-11 | 4.38E-06 |
| 2 | 11262399 | C | T | 0.840344 | 1.12E-11 | 4.38E-06 |
| 2 | 11262408 | G | A | 0.840344 | 1.12E-11 | 4.38E-06 |
| 2 | 8907302 | G | T | 0.854685 | 1.25E-11 | 4.90E-06 |
| 2 | 8907310 | G | A | 0.854685 | 1.25E-11 | 4.90E-06 |
| 2 | 8907518 | T | C | 0.854685 | 1.25E-11 | 4.90E-06 |
| 2 | 8907682 | C | T | 0.854685 | 1.25E-11 | 4.90E-06 |
| 2 | 8907874 | G | A | 0.854685 | 1.25E-11 | 4.90E-06 |
| 2 | 8907892 | C | A | 0.854685 | 1.25E-11 | 4.90E-06 |
| 2 | 8907910 | C | T | 0.854685 | 1.25E-11 | 4.90E-06 |
| 2 | 8907917 | T | G | 0.854685 | 1.25E-11 | 4.90E-06 |
| 2 | 11262435 | G | A | 0.83652 | 1.36E-11 | 5.33E-06 |
| 2 | 8890029 | A | T | 0.855641 | 1.82E-11 | 7.14E-06 |
| 2 | 11269194 | A | G | 0.866157 | 2.06E-11 | 8.08E-06 |
| 2 | 11269614 | A | G | 0.866157 | 2.06E-11 | 8.08E-06 |
| 2 | 8257045 | G | T | 0.881453 | 2.47E-11 | 9.68E-06 |
| 2 | 11282426 | T | C | 0.864245 | 3.87E-11 | 1.52E-05 |
| 2 | 11282433 | G | A | 0.864245 | 3.87E-11 | 1.52E-05 |
| 2 | 8894966 | T | C | 0.853728 | 4.28E-11 | 1.68E-05 |
| 2 | 8130114 | A | G | 0.90153 | 4.32E-11 | 1.69E-05 |
| 2 | 8130116 | G | A | 0.90153 | 4.32E-11 | 1.69E-05 |
| 2 | 8130425 | A | G | 0.90153 | 4.32E-11 | 1.69E-05 |
| 2 | 8130640 | G | A | 0.90153 | 4.32E-11 | 1.69E-05 |
| 2 | 8130886 | T | C | 0.90153 | 4.32E-11 | 1.69E-05 |
| 2 | 8131825 | C | T | 0.90153 | 4.32E-11 | 1.69E-05 |
| 2 | 8131855 | A | G | 0.90153 | 4.32E-11 | 1.69E-05 |
| 2 | 8131880 | C | T | 0.90153 | 4.32E-11 | 1.69E-05 |
| 2 | 8131932 | A | T | 0.90153 | 4.32E-11 | 1.69E-05 |
| 2 | 8132026 | A | G | 0.90153 | 4.32E-11 | 1.69E-05 |

| | | | | | | |
|---|---|---|---|---|---|---|
| 2 | 8132055 | G | A | 0.90153 | 4.32E-11 | 1.69E-05 |
| 2 | 8132067 | C | T | 0.90153 | 4.32E-11 | 1.69E-05 |
| 2 | 8132076 | A | G | 0.90153 | 4.32E-11 | 1.69E-05 |
| 2 | 8132528 | T | G | 0.90153 | 4.32E-11 | 1.69E-05 |
| 2 | 8132838 | G | T | 0.90153 | 4.32E-11 | 1.69E-05 |
| 2 | 8133769 | T | C | 0.90153 | 4.32E-11 | 1.69E-05 |
| 2 | 8133771 | C | A | 0.90153 | 4.32E-11 | 1.69E-05 |
| 2 | 8134793 | C | T | 0.90153 | 4.32E-11 | 1.69E-05 |
| 2 | 8134971 | C | T | 0.90153 | 4.32E-11 | 1.69E-05 |
| 2 | 8138597 | G | A | 0.90153 | 4.32E-11 | 1.69E-05 |
| 2 | 8138913 | C | G | 0.90153 | 4.32E-11 | 1.69E-05 |
| 2 | 8139240 | C | T | 0.90153 | 4.32E-11 | 1.69E-05 |
| 2 | 8139303 | G | A | 0.90153 | 4.32E-11 | 1.69E-05 |
| 2 | 8139314 | C | G | 0.90153 | 4.32E-11 | 1.69E-05 |
| 2 | 8139322 | A | G | 0.90153 | 4.32E-11 | 1.69E-05 |
| 2 | 8139344 | T | C | 0.90153 | 4.32E-11 | 1.69E-05 |
| 2 | 8139520 | C | T | 0.90153 | 4.32E-11 | 1.69E-05 |
| 2 | 8139584 | A | G | 0.90153 | 4.32E-11 | 1.69E-05 |
| 2 | 8139585 | C | A | 0.90153 | 4.32E-11 | 1.69E-05 |
| 2 | 8142291 | C | G | 0.90153 | 4.32E-11 | 1.69E-05 |
| 2 | 8142350 | G | A | 0.90153 | 4.32E-11 | 1.69E-05 |
| 2 | 8142597 | A | G | 0.90153 | 4.32E-11 | 1.69E-05 |
| 2 | 8142625 | T | A | 0.90153 | 4.32E-11 | 1.69E-05 |
| 2 | 8142628 | A | G | 0.90153 | 4.32E-11 | 1.69E-05 |
| 2 | 8142776 | C | T | 0.90153 | 4.32E-11 | 1.69E-05 |
| 2 | 8142805 | T | C | 0.90153 | 4.32E-11 | 1.69E-05 |
| 2 | 8142821 | G | T | 0.90153 | 4.32E-11 | 1.69E-05 |
| 2 | 8142823 | G | T | 0.90153 | 4.32E-11 | 1.69E-05 |
| 2 | 8145940 | A | G | 0.90153 | 4.32E-11 | 1.69E-05 |
| 2 | 8146597 | C | T | 0.90153 | 4.32E-11 | 1.69E-05 |
| 2 | 8147631 | T | C | 0.90153 | 4.32E-11 | 1.69E-05 |
| 2 | 8148310 | G | A | 0.90153 | 4.32E-11 | 1.69E-05 |
| 2 | 8149725 | G | A | 0.90153 | 4.32E-11 | 1.69E-05 |
| 2 | 8149749 | C | T | 0.90153 | 4.32E-11 | 1.69E-05 |
| 2 | 8149823 | A | G | 0.90153 | 4.32E-11 | 1.69E-05 |
| 2 | 8150177 | T | C | 0.90153 | 4.32E-11 | 1.69E-05 |
| 2 | 8151152 | C | G | 0.90153 | 4.32E-11 | 1.69E-05 |
| 2 | 8151244 | A | C | 0.90153 | 4.32E-11 | 1.69E-05 |
| 2 | 8151369 | T | C | 0.90153 | 4.32E-11 | 1.69E-05 |
| 2 | 8151478 | G | A | 0.90153 | 4.32E-11 | 1.69E-05 |
| 2 | 8151960 | T | A | 0.90153 | 4.32E-11 | 1.69E-05 |
| 2 | 8153509 | G | A | 0.90153 | 4.32E-11 | 1.69E-05 |
| 2 | 8153617 | C | T | 0.90153 | 4.32E-11 | 1.69E-05 |
| 2 | 8153656 | G | A | 0.90153 | 4.32E-11 | 1.69E-05 |
| 2 | 8153764 | A | G | 0.90153 | 4.32E-11 | 1.69E-05 |
| 2 | 8153766 | A | G | 0.90153 | 4.32E-11 | 1.69E-05 |
| 2 | 8153770 | C | T | 0.90153 | 4.32E-11 | 1.69E-05 |
| 2 | 8153776 | G | A | 0.90153 | 4.32E-11 | 1.69E-05 |
| 2 | 8154003 | A | G | 0.90153 | 4.32E-11 | 1.69E-05 |
| 2 | 8154099 | C | T | 0.90153 | 4.32E-11 | 1.69E-05 |
| 2 | 8154225 | T | C | 0.90153 | 4.32E-11 | 1.69E-05 |
| 2 | 8154230 | C | A | 0.90153 | 4.32E-11 | 1.69E-05 |
| 2 | 8154416 | C | T | 0.90153 | 4.32E-11 | 1.69E-05 |
| 2 | 8154453 | T | G | 0.90153 | 4.32E-11 | 1.69E-05 |
| 2 | 8154477 | T | C | 0.90153 | 4.32E-11 | 1.69E-05 |
| 2 | 8154512 | G | A | 0.90153 | 4.32E-11 | 1.69E-05 |
| 2 | 8154544 | C | G | 0.90153 | 4.32E-11 | 1.69E-05 |
| 2 | 8154546 | G | T | 0.90153 | 4.32E-11 | 1.69E-05 |
| 2 | 8154581 | C | T | 0.90153 | 4.32E-11 | 1.69E-05 |
| 2 | 8154591 | A | G | 0.90153 | 4.32E-11 | 1.69E-05 |
| 2 | 8154602 | G | A | 0.90153 | 4.32E-11 | 1.69E-05 |
| 2 | 8154801 | T | C | 0.90153 | 4.32E-11 | 1.69E-05 |
| 2 | 8155021 | C | G | 0.90153 | 4.32E-11 | 1.69E-05 |
| 2 | 8155023 | A | G | 0.90153 | 4.32E-11 | 1.69E-05 |
| 2 | 8155056 | T | C | 0.90153 | 4.32E-11 | 1.69E-05 |
| 2 | 8155128 | G | A | 0.90153 | 4.32E-11 | 1.69E-05 |
| 2 | 8155183 | T | C | 0.90153 | 4.32E-11 | 1.69E-05 |

| | | | | | | |
|---|---|---|---|---|---|---|
| 2 | 8155328 | T | G | 0.90153 | 4.32E-11 | 1.69E-05 |
| 2 | 8155332 | A | C | 0.90153 | 4.32E-11 | 1.69E-05 |
| 2 | 8155377 | C | T | 0.90153 | 4.32E-11 | 1.69E-05 |
| 2 | 8155443 | T | G | 0.90153 | 4.32E-11 | 1.69E-05 |
| 2 | 8155498 | T | C | 0.90153 | 4.32E-11 | 1.69E-05 |
| 2 | 8155609 | T | A | 0.90153 | 4.32E-11 | 1.69E-05 |
| 2 | 8155628 | T | A | 0.90153 | 4.32E-11 | 1.69E-05 |
| 2 | 8155731 | T | G | 0.90153 | 4.32E-11 | 1.69E-05 |
| 2 | 8155737 | G | A | 0.90153 | 4.32E-11 | 1.69E-05 |
| 2 | 8155781 | G | A | 0.90153 | 4.32E-11 | 1.69E-05 |
| 2 | 8155905 | C | A | 0.90153 | 4.32E-11 | 1.69E-05 |
| 2 | 8155913 | C | T | 0.90153 | 4.32E-11 | 1.69E-05 |
| 2 | 8155969 | T | C | 0.90153 | 4.32E-11 | 1.69E-05 |
| 2 | 8156141 | A | G | 0.90153 | 4.32E-11 | 1.69E-05 |
| 2 | 8156247 | G | T | 0.90153 | 4.32E-11 | 1.69E-05 |
| 2 | 8156308 | A | G | 0.90153 | 4.32E-11 | 1.69E-05 |
| 2 | 8156378 | A | C | 0.90153 | 4.32E-11 | 1.69E-05 |
| 2 | 8156422 | C | T | 0.90153 | 4.32E-11 | 1.69E-05 |
| 2 | 8156481 | C | T | 0.90153 | 4.32E-11 | 1.69E-05 |
| 2 | 8156514 | A | G | 0.90153 | 4.32E-11 | 1.69E-05 |
| 2 | 8156590 | T | C | 0.90153 | 4.32E-11 | 1.69E-05 |
| 2 | 8156699 | G | C | 0.90153 | 4.32E-11 | 1.69E-05 |
| 2 | 8156728 | G | T | 0.90153 | 4.32E-11 | 1.69E-05 |
| 2 | 8156794 | C | A | 0.90153 | 4.32E-11 | 1.69E-05 |
| 2 | 8156944 | G | A | 0.90153 | 4.32E-11 | 1.69E-05 |
| 2 | 8157039 | G | A | 0.90153 | 4.32E-11 | 1.69E-05 |
| 2 | 8157066 | G | A | 0.90153 | 4.32E-11 | 1.69E-05 |
| 2 | 8157189 | C | T | 0.90153 | 4.32E-11 | 1.69E-05 |
| 2 | 8157214 | A | T | 0.90153 | 4.32E-11 | 1.69E-05 |
| 2 | 8157222 | A | G | 0.90153 | 4.32E-11 | 1.69E-05 |
| 2 | 8157244 | C | A | 0.90153 | 4.32E-11 | 1.69E-05 |
| 2 | 8157271 | C | A | 0.90153 | 4.32E-11 | 1.69E-05 |
| 2 | 8157408 | A | G | 0.90153 | 4.32E-11 | 1.69E-05 |
| 2 | 8157467 | A | G | 0.90153 | 4.32E-11 | 1.69E-05 |
| 2 | 8157741 | T | G | 0.90153 | 4.32E-11 | 1.69E-05 |
| 2 | 8157770 | T | C | 0.90153 | 4.32E-11 | 1.69E-05 |
| 2 | 8157799 | T | C | 0.90153 | 4.32E-11 | 1.69E-05 |
| 2 | 8157800 | G | A | 0.90153 | 4.32E-11 | 1.69E-05 |
| 2 | 8157805 | C | T | 0.90153 | 4.32E-11 | 1.69E-05 |
| 2 | 8157806 | A | G | 0.90153 | 4.32E-11 | 1.69E-05 |
| 2 | 8157843 | T | C | 0.90153 | 4.32E-11 | 1.69E-05 |
| 2 | 8157880 | G | T | 0.90153 | 4.32E-11 | 1.69E-05 |
| 2 | 8157888 | T | A | 0.90153 | 4.32E-11 | 1.69E-05 |
| 2 | 8157921 | G | A | 0.90153 | 4.32E-11 | 1.69E-05 |
| 2 | 8157929 | C | T | 0.90153 | 4.32E-11 | 1.69E-05 |
| 2 | 8157935 | A | G | 0.90153 | 4.32E-11 | 1.69E-05 |
| 2 | 8157937 | G | A | 0.90153 | 4.32E-11 | 1.69E-05 |
| 2 | 8157938 | G | A | 0.90153 | 4.32E-11 | 1.69E-05 |
| 2 | 8157964 | G | A | 0.90153 | 4.32E-11 | 1.69E-05 |
| 2 | 8157966 | A | G | 0.90153 | 4.32E-11 | 1.69E-05 |
| 2 | 8157982 | A | C | 0.90153 | 4.32E-11 | 1.69E-05 |
| 2 | 8160874 | A | G | 0.90153 | 4.32E-11 | 1.69E-05 |
| 2 | 8160878 | T | C | 0.90153 | 4.32E-11 | 1.69E-05 |
| 2 | 8160890 | A | G | 0.90153 | 4.32E-11 | 1.69E-05 |
| 2 | 8160901 | T | C | 0.90153 | 4.32E-11 | 1.69E-05 |
| 2 | 8160965 | C | T | 0.90153 | 4.32E-11 | 1.69E-05 |
| 2 | 8161127 | C | T | 0.90153 | 4.32E-11 | 1.69E-05 |
| 2 | 8161316 | G | A | 0.90153 | 4.32E-11 | 1.69E-05 |
| 2 | 8161477 | T | C | 0.90153 | 4.32E-11 | 1.69E-05 |
| 2 | 8161731 | C | T | 0.90153 | 4.32E-11 | 1.69E-05 |
| 2 | 8161963 | C | T | 0.90153 | 4.32E-11 | 1.69E-05 |
| 2 | 8162044 | T | A | 0.90153 | 4.32E-11 | 1.69E-05 |
| 2 | 8162071 | T | C | 0.90153 | 4.32E-11 | 1.69E-05 |
| 2 | 8162137 | T | C | 0.90153 | 4.32E-11 | 1.69E-05 |
| 2 | 8162240 | T | C | 0.90153 | 4.32E-11 | 1.69E-05 |
| 2 | 8162302 | T | C | 0.90153 | 4.32E-11 | 1.69E-05 |
| 2 | 8162321 | G | A | 0.90153 | 4.32E-11 | 1.69E-05 |

| | | | | | | |
|---|---|---|---|---|---|---|
| 2 | 8162334 | A | T | 0.90153 | 4.32E-11 | 1.69E-05 |
| 2 | 8162416 | T | C | 0.90153 | 4.32E-11 | 1.69E-05 |
| 2 | 8162467 | T | C | 0.90153 | 4.32E-11 | 1.69E-05 |
| 2 | 8162479 | A | G | 0.90153 | 4.32E-11 | 1.69E-05 |
| 2 | 8162481 | G | A | 0.90153 | 4.32E-11 | 1.69E-05 |
| 2 | 8162533 | C | T | 0.90153 | 4.32E-11 | 1.69E-05 |
| 2 | 8162539 | T | C | 0.90153 | 4.32E-11 | 1.69E-05 |
| 2 | 8162547 | T | C | 0.90153 | 4.32E-11 | 1.69E-05 |
| 2 | 8162557 | A | G | 0.90153 | 4.32E-11 | 1.69E-05 |
| 2 | 8162685 | C | G | 0.90153 | 4.32E-11 | 1.69E-05 |
| 2 | 8162833 | C | T | 0.90153 | 4.32E-11 | 1.69E-05 |
| 2 | 8162842 | A | G | 0.90153 | 4.32E-11 | 1.69E-05 |
| 2 | 8163012 | G | A | 0.90153 | 4.32E-11 | 1.69E-05 |
| 2 | 8163085 | G | A | 0.90153 | 4.32E-11 | 1.69E-05 |
| 2 | 8163101 | T | A | 0.90153 | 4.32E-11 | 1.69E-05 |
| 2 | 8163458 | C | A | 0.90153 | 4.32E-11 | 1.69E-05 |
| 2 | 8163460 | C | A | 0.90153 | 4.32E-11 | 1.69E-05 |
| 2 | 8163911 | G | T | 0.90153 | 4.32E-11 | 1.69E-05 |
| 2 | 8163951 | T | C | 0.90153 | 4.32E-11 | 1.69E-05 |
| 2 | 8163958 | T | C | 0.90153 | 4.32E-11 | 1.69E-05 |
| 2 | 8163967 | C | T | 0.90153 | 4.32E-11 | 1.69E-05 |
| 2 | 8164217 | G | T | 0.90153 | 4.32E-11 | 1.69E-05 |
| 2 | 8164430 | C | T | 0.90153 | 4.32E-11 | 1.69E-05 |
| 2 | 8164510 | T | C | 0.90153 | 4.32E-11 | 1.69E-05 |
| 2 | 8164553 | T | C | 0.90153 | 4.32E-11 | 1.69E-05 |
| 2 | 8167109 | A | G | 0.90153 | 4.32E-11 | 1.69E-05 |
| 2 | 8167121 | C | T | 0.90153 | 4.32E-11 | 1.69E-05 |
| 2 | 8167171 | C | T | 0.90153 | 4.32E-11 | 1.69E-05 |
| 2 | 8167222 | T | C | 0.90153 | 4.32E-11 | 1.69E-05 |
| 2 | 8167225 | G | A | 0.90153 | 4.32E-11 | 1.69E-05 |
| 2 | 8167229 | A | G | 0.90153 | 4.32E-11 | 1.69E-05 |
| 2 | 8167333 | A | G | 0.90153 | 4.32E-11 | 1.69E-05 |
| 2 | 8167431 | C | T | 0.90153 | 4.32E-11 | 1.69E-05 |
| 2 | 8167674 | G | C | 0.90153 | 4.32E-11 | 1.69E-05 |
| 2 | 8167777 | A | G | 0.90153 | 4.32E-11 | 1.69E-05 |
| 2 | 8167825 | G | A | 0.90153 | 4.32E-11 | 1.69E-05 |
| 2 | 8168082 | A | G | 0.90153 | 4.32E-11 | 1.69E-05 |
| 2 | 8168114 | A | G | 0.90153 | 4.32E-11 | 1.69E-05 |
| 2 | 8168487 | A | G | 0.90153 | 4.32E-11 | 1.69E-05 |
| 2 | 8168706 | A | G | 0.90153 | 4.32E-11 | 1.69E-05 |
| 2 | 8168713 | A | G | 0.90153 | 4.32E-11 | 1.69E-05 |
| 2 | 8169038 | C | T | 0.90153 | 4.32E-11 | 1.69E-05 |
| 2 | 8196769 | A | G | 0.90153 | 4.32E-11 | 1.69E-05 |
| 2 | 8196772 | A | G | 0.90153 | 4.32E-11 | 1.69E-05 |
| 2 | 11126937 | A | C | 0.839388 | 4.72E-11 | 1.85E-05 |
| 2 | 8164401 | A | G | 0.902486 | 4.82E-11 | 1.89E-05 |
| 2 | 8164998 | A | C | 0.902486 | 4.86E-11 | 1.91E-05 |
| 2 | 8166075 | T | C | 0.903442 | 5.13E-11 | 2.01E-05 |
| 2 | 8166211 | A | C | 0.903442 | 5.13E-11 | 2.01E-05 |
| 2 | 8166271 | T | C | 0.903442 | 5.13E-11 | 2.01E-05 |
| 2 | 11268009 | G | C | 0.863289 | 5.18E-11 | 2.03E-05 |
| 2 | 11268035 | A | G | 0.863289 | 5.18E-11 | 2.03E-05 |
| 2 | 11268036 | T | G | 0.863289 | 5.18E-11 | 2.03E-05 |
| 2 | 11268046 | T | C | 0.863289 | 5.18E-11 | 2.03E-05 |
| 2 | 11268166 | A | G | 0.863289 | 5.18E-11 | 2.03E-05 |
| 2 | 11268207 | A | G | 0.863289 | 5.18E-11 | 2.03E-05 |
| 2 | 11268215 | A | G | 0.863289 | 5.18E-11 | 2.03E-05 |
| 2 | 11270931 | G | A | 0.863289 | 5.18E-11 | 2.03E-05 |
| 2 | 11270950 | T | G | 0.863289 | 5.18E-11 | 2.03E-05 |
| 2 | 11271951 | T | G | 0.863289 | 5.18E-11 | 2.03E-05 |
| 2 | 11273024 | G | C | 0.863289 | 5.18E-11 | 2.03E-05 |
| 2 | 11274679 | T | C | 0.863289 | 5.18E-11 | 2.03E-05 |
| 2 | 11274737 | G | T | 0.863289 | 5.18E-11 | 2.03E-05 |
| 2 | 11276407 | C | T | 0.863289 | 5.18E-11 | 2.03E-05 |
| 2 | 11277927 | T | C | 0.863289 | 5.18E-11 | 2.03E-05 |
| 2 | 11279908 | T | C | 0.863289 | 5.18E-11 | 2.03E-05 |
| 2 | 11280867 | C | T | 0.863289 | 5.18E-11 | 2.03E-05 |

| | | | | | | |
|---|---|---|---|---|---|---|
| 2 | 11282123 | C | G | 0.863289 | 5.18E-11 | 2.03E-05 |
| 2 | 11282774 | T | C | 0.863289 | 5.18E-11 | 2.03E-05 |
| 2 | 11283078 | C | T | 0.863289 | 5.18E-11 | 2.03E-05 |
| 2 | 11283102 | T | C | 0.863289 | 5.18E-11 | 2.03E-05 |
| 2 | 11283169 | T | A | 0.863289 | 5.18E-11 | 2.03E-05 |
| 2 | 11283482 | A | G | 0.863289 | 5.18E-11 | 2.03E-05 |
| 2 | 11283569 | G | C | 0.863289 | 5.18E-11 | 2.03E-05 |
| 2 | 11283853 | T | A | 0.863289 | 5.18E-11 | 2.03E-05 |
| 2 | 11283874 | G | C | 0.863289 | 5.18E-11 | 2.03E-05 |
| 2 | 11284150 | G | A | 0.863289 | 5.18E-11 | 2.03E-05 |
| 2 | 11284254 | A | G | 0.863289 | 5.18E-11 | 2.03E-05 |
| 2 | 8257100 | A | C | 0.882409 | 5.28E-11 | 2.07E-05 |
| 2 | 11263247 | T | C | 0.838432 | 5.61E-11 | 2.20E-05 |
| 2 | 11263286 | C | T | 0.838432 | 5.61E-11 | 2.20E-05 |
| 2 | 8140650 | T | A | 0.902486 | 5.65E-11 | 2.21E-05 |
| 2 | 8140658 | A | G | 0.902486 | 5.65E-11 | 2.21E-05 |
| 2 | 8143378 | C | A | 0.902486 | 5.65E-11 | 2.21E-05 |
| 2 | 11116078 | A | G | 0.837476 | 5.88E-11 | 2.31E-05 |
| 2 | 11116092 | T | C | 0.837476 | 5.88E-11 | 2.31E-05 |
| 2 | 11116172 | C | T | 0.837476 | 5.88E-11 | 2.31E-05 |
| 2 | 11116537 | G | T | 0.837476 | 5.88E-11 | 2.31E-05 |
| 2 | 11117221 | T | G | 0.837476 | 5.88E-11 | 2.31E-05 |
| 2 | 11117704 | A | T | 0.837476 | 5.88E-11 | 2.31E-05 |
| 2 | 11117844 | A | G | 0.837476 | 5.88E-11 | 2.31E-05 |
| 2 | 11119603 | C | T | 0.837476 | 5.88E-11 | 2.31E-05 |
| 2 | 11122005 | T | G | 0.837476 | 5.88E-11 | 2.31E-05 |
| 2 | 11237198 | A | G | 0.839388 | 6.09E-11 | 2.39E-05 |
| 2 | 11237260 | A | G | 0.839388 | 6.09E-11 | 2.39E-05 |
| 2 | 11239488 | T | C | 0.839388 | 6.09E-11 | 2.39E-05 |
| 2 | 11128031 | A | T | 0.837476 | 6.53E-11 | 2.56E-05 |
| 2 | 11270336 | G | A | 0.867113 | 6.77E-11 | 2.65E-05 |
| 2 | 11270650 | T | C | 0.867113 | 6.77E-11 | 2.65E-05 |
| 2 | 11263122 | G | A | 0.840344 | 7.33E-11 | 2.87E-05 |
| 2 | 11263141 | A | G | 0.840344 | 7.33E-11 | 2.87E-05 |
| 2 | 11263155 | G | C | 0.840344 | 7.33E-11 | 2.87E-05 |
| 2 | 11263267 | T | C | 0.840344 | 7.33E-11 | 2.87E-05 |
| 2 | 8166407 | T | C | 0.899618 | 7.83E-11 | 3.07E-05 |
| 2 | 8166498 | C | T | 0.904398 | 8.03E-11 | 3.15E-05 |
| 2 | 8166540 | T | C | 0.904398 | 8.03E-11 | 3.15E-05 |
| 2 | 8166554 | A | G | 0.904398 | 8.03E-11 | 3.15E-05 |
| 2 | 8166566 | A | C | 0.904398 | 8.03E-11 | 3.15E-05 |
| 2 | 8166581 | A | G | 0.904398 | 8.03E-11 | 3.15E-05 |
| 2 | 8166615 | T | C | 0.904398 | 8.03E-11 | 3.15E-05 |
| 2 | 8165898 | A | G | 0.902486 | 8.17E-11 | 3.20E-05 |
| 2 | 8850999 | C | T | 0.84608 | 8.22E-11 | 3.22E-05 |
| 2 | 11271426 | C | T | 0.862333 | 8.40E-11 | 3.29E-05 |
| 2 | 11271500 | A | G | 0.862333 | 8.40E-11 | 3.29E-05 |
| 2 | 11277318 | G | A | 0.862333 | 8.40E-11 | 3.29E-05 |
| 2 | 11281415 | T | C | 0.862333 | 8.40E-11 | 3.29E-05 |
| 2 | 11281538 | G | C | 0.862333 | 8.40E-11 | 3.29E-05 |
| 2 | 11281597 | T | C | 0.862333 | 8.40E-11 | 3.29E-05 |
| 2 | 11281620 | G | A | 0.862333 | 8.40E-11 | 3.29E-05 |
| 2 | 11281865 | T | C | 0.862333 | 8.40E-11 | 3.29E-05 |
| 2 | 11281882 | T | C | 0.862333 | 8.40E-11 | 3.29E-05 |
| 2 | 11281947 | C | A | 0.862333 | 8.40E-11 | 3.29E-05 |
| 2 | 11282064 | T | C | 0.862333 | 8.40E-11 | 3.29E-05 |
| 2 | 11310994 | C | G | 0.870937 | 8.78E-11 | 3.44E-05 |
| 2 | 11312400 | C | T | 0.870937 | 8.78E-11 | 3.44E-05 |
| 2 | 11313403 | A | G | 0.870937 | 8.78E-11 | 3.44E-05 |
| 2 | 11314035 | A | G | 0.870937 | 8.78E-11 | 3.44E-05 |
| 2 | 11314823 | T | C | 0.870937 | 8.78E-11 | 3.44E-05 |
| 2 | 11314941 | C | T | 0.870937 | 8.78E-11 | 3.44E-05 |
| 2 | 11315405 | A | G | 0.870937 | 8.78E-11 | 3.44E-05 |
| 2 | 11316924 | T | C | 0.870937 | 8.78E-11 | 3.44E-05 |
| 2 | 11317572 | A | G | 0.870937 | 8.78E-11 | 3.44E-05 |
| 2 | 11318053 | T | A | 0.870937 | 8.78E-11 | 3.44E-05 |
| 2 | 11318673 | A | G | 0.870937 | 8.78E-11 | 3.44E-05 |

| | | | | | | |
|---|---|---|---|---|---|---|
| 2 | 11322880 | C | T | 0.870937 | 8.78E-11 | 3.44E-05 |
| 2 | 11341451 | T | G | 0.871893 | 9.18E-11 | 3.60E-05 |
| 2 | 11352418 | G | A | 0.871893 | 9.18E-11 | 3.60E-05 |
| 2 | 10791629 | C | T | 0.83652 | 9.80E-11 | 3.84E-05 |
| 2 | 11109441 | T | G | 0.835564 | 1.05E-10 | 4.12E-05 |
| 2 | 11109455 | G | A | 0.835564 | 1.05E-10 | 4.12E-05 |
| 2 | 8260191 | A | G | 0.879541 | 1.06E-10 | 4.16E-05 |
| 2 | 8260192 | A | G | 0.879541 | 1.06E-10 | 4.16E-05 |
| 2 | 8260218 | G | A | 0.879541 | 1.06E-10 | 4.16E-05 |
| 2 | 8895686 | C | T | 0.854685 | 1.09E-10 | 4.27E-05 |
| 2 | 8895690 | T | A | 0.854685 | 1.09E-10 | 4.27E-05 |
| 2 | 11096781 | T | C | 0.838432 | 1.16E-10 | 4.55E-05 |
| 2 | 11096806 | G | A | 0.838432 | 1.16E-10 | 4.55E-05 |
| 2 | 11097349 | A | G | 0.838432 | 1.16E-10 | 4.55E-05 |
| 2 | 11098133 | T | G | 0.838432 | 1.16E-10 | 4.55E-05 |
| 2 | 11098323 | A | T | 0.838432 | 1.16E-10 | 4.55E-05 |
| 2 | 11099114 | T | C | 0.838432 | 1.16E-10 | 4.55E-05 |
| 2 | 11100004 | T | C | 0.838432 | 1.16E-10 | 4.55E-05 |
| 2 | 11100873 | T | G | 0.838432 | 1.16E-10 | 4.55E-05 |
| 2 | 11100898 | T | A | 0.838432 | 1.16E-10 | 4.55E-05 |
| 2 | 11101259 | T | C | 0.838432 | 1.16E-10 | 4.55E-05 |
| 2 | 11101566 | G | A | 0.838432 | 1.16E-10 | 4.55E-05 |
| 2 | 11101972 | C | A | 0.838432 | 1.16E-10 | 4.55E-05 |
| 2 | 11102129 | G | A | 0.838432 | 1.16E-10 | 4.55E-05 |
| 2 | 11102236 | A | G | 0.838432 | 1.16E-10 | 4.55E-05 |
| 2 | 11102459 | A | G | 0.838432 | 1.16E-10 | 4.55E-05 |
| 2 | 11103053 | A | G | 0.838432 | 1.16E-10 | 4.55E-05 |
| 2 | 11103364 | T | A | 0.838432 | 1.16E-10 | 4.55E-05 |
| 2 | 11103398 | G | A | 0.838432 | 1.16E-10 | 4.55E-05 |
| 2 | 11104207 | G | A | 0.838432 | 1.16E-10 | 4.55E-05 |
| 2 | 11104652 | T | C | 0.838432 | 1.16E-10 | 4.55E-05 |
| 2 | 11107470 | G | A | 0.838432 | 1.16E-10 | 4.55E-05 |
| 2 | 11107938 | G | T | 0.838432 | 1.16E-10 | 4.55E-05 |
| 2 | 11111710 | G | A | 0.838432 | 1.16E-10 | 4.55E-05 |
| 2 | 11115338 | G | T | 0.838432 | 1.16E-10 | 4.55E-05 |
| 2 | 11115649 | A | G | 0.838432 | 1.16E-10 | 4.55E-05 |
| 2 | 11115723 | A | T | 0.838432 | 1.16E-10 | 4.55E-05 |
| 2 | 11117095 | T | C | 0.838432 | 1.16E-10 | 4.55E-05 |
| 2 | 11118023 | A | G | 0.838432 | 1.16E-10 | 4.55E-05 |
| 2 | 11118756 | A | G | 0.838432 | 1.16E-10 | 4.55E-05 |
| 2 | 11118995 | G | A | 0.838432 | 1.16E-10 | 4.55E-05 |
| 2 | 11119809 | C | G | 0.838432 | 1.16E-10 | 4.55E-05 |
| 2 | 11119814 | C | A | 0.838432 | 1.16E-10 | 4.55E-05 |
| 2 | 11120009 | C | G | 0.838432 | 1.16E-10 | 4.55E-05 |
| 2 | 11121904 | A | G | 0.838432 | 1.16E-10 | 4.55E-05 |
| 2 | 11123599 | T | C | 0.838432 | 1.16E-10 | 4.55E-05 |
| 2 | 11124417 | A | G | 0.838432 | 1.16E-10 | 4.55E-05 |
| 2 | 11127568 | A | T | 0.838432 | 1.16E-10 | 4.55E-05 |
| 2 | 11127570 | T | C | 0.838432 | 1.16E-10 | 4.55E-05 |
| 2 | 11127645 | C | G | 0.838432 | 1.16E-10 | 4.55E-05 |
| 2 | 11128032 | A | T | 0.838432 | 1.16E-10 | 4.55E-05 |
| 2 | 11128137 | C | T | 0.838432 | 1.16E-10 | 4.55E-05 |
| 2 | 11128180 | A | G | 0.838432 | 1.16E-10 | 4.55E-05 |
| 2 | 11128481 | T | G | 0.838432 | 1.16E-10 | 4.55E-05 |
| 2 | 11128615 | A | C | 0.838432 | 1.16E-10 | 4.55E-05 |
| 2 | 11129003 | C | T | 0.838432 | 1.16E-10 | 4.55E-05 |
| 2 | 11129082 | A | C | 0.838432 | 1.16E-10 | 4.55E-05 |
| 2 | 11129128 | A | C | 0.838432 | 1.16E-10 | 4.55E-05 |
| 2 | 11129175 | C | T | 0.838432 | 1.16E-10 | 4.55E-05 |
| 2 | 11129274 | A | G | 0.838432 | 1.16E-10 | 4.55E-05 |
| 2 | 11129336 | A | G | 0.838432 | 1.16E-10 | 4.55E-05 |
| 2 | 11129980 | A | G | 0.838432 | 1.16E-10 | 4.55E-05 |
| 2 | 11130012 | C | A | 0.838432 | 1.16E-10 | 4.55E-05 |
| 2 | 11130140 | A | C | 0.838432 | 1.16E-10 | 4.55E-05 |
| 2 | 11130437 | T | G | 0.838432 | 1.16E-10 | 4.55E-05 |
| 2 | 11130655 | A | C | 0.838432 | 1.16E-10 | 4.55E-05 |
| 2 | 11130872 | C | A | 0.838432 | 1.16E-10 | 4.55E-05 |

| | | | | | | |
|---|---|---|---|---|---|---|
| 2 | 11131579 | T | C | 0.838432 | 1.16E-10 | 4.55E-05 |
| 2 | 11131653 | A | G | 0.838432 | 1.16E-10 | 4.55E-05 |
| 2 | 11132203 | A | G | 0.838432 | 1.16E-10 | 4.55E-05 |
| 2 | 11135008 | G | A | 0.838432 | 1.16E-10 | 4.55E-05 |
| 2 | 11135116 | A | G | 0.838432 | 1.16E-10 | 4.55E-05 |
| 2 | 11135219 | T | C | 0.838432 | 1.16E-10 | 4.55E-05 |
| 2 | 11135343 | C | T | 0.838432 | 1.16E-10 | 4.55E-05 |
| 2 | 11135527 | T | G | 0.838432 | 1.16E-10 | 4.55E-05 |
| 2 | 11135570 | G | C | 0.838432 | 1.16E-10 | 4.55E-05 |
| 2 | 11135589 | G | A | 0.838432 | 1.16E-10 | 4.55E-05 |
| 2 | 11135727 | G | A | 0.838432 | 1.16E-10 | 4.55E-05 |
| 2 | 11135754 | A | G | 0.838432 | 1.16E-10 | 4.55E-05 |
| 2 | 11135780 | G | A | 0.838432 | 1.16E-10 | 4.55E-05 |
| 2 | 11135896 | T | C | 0.838432 | 1.16E-10 | 4.55E-05 |
| 2 | 11135911 | C | G | 0.838432 | 1.16E-10 | 4.55E-05 |
| 2 | 11136007 | T | G | 0.838432 | 1.16E-10 | 4.55E-05 |
| 2 | 11136033 | T | C | 0.838432 | 1.16E-10 | 4.55E-05 |
| 2 | 11136123 | C | T | 0.838432 | 1.16E-10 | 4.55E-05 |
| 2 | 11136189 | G | T | 0.838432 | 1.16E-10 | 4.55E-05 |
| 2 | 11136312 | C | G | 0.838432 | 1.16E-10 | 4.55E-05 |
| 2 | 11136387 | G | A | 0.838432 | 1.16E-10 | 4.55E-05 |
| 2 | 11136393 | G | T | 0.838432 | 1.16E-10 | 4.55E-05 |
| 2 | 11136414 | T | C | 0.838432 | 1.16E-10 | 4.55E-05 |
| 2 | 11136472 | A | G | 0.838432 | 1.16E-10 | 4.55E-05 |
| 2 | 11136522 | G | C | 0.838432 | 1.16E-10 | 4.55E-05 |
| 2 | 11136535 | C | T | 0.838432 | 1.16E-10 | 4.55E-05 |
| 2 | 11136655 | C | T | 0.838432 | 1.16E-10 | 4.55E-05 |
| 2 | 11137886 | C | T | 0.838432 | 1.16E-10 | 4.55E-05 |
| 2 | 11138170 | T | C | 0.838432 | 1.16E-10 | 4.55E-05 |
| 2 | 11138276 | T | C | 0.838432 | 1.16E-10 | 4.55E-05 |
| 2 | 11138698 | G | A | 0.838432 | 1.16E-10 | 4.55E-05 |
| 2 | 11140882 | G | T | 0.838432 | 1.16E-10 | 4.55E-05 |
| 2 | 11141935 | G | A | 0.838432 | 1.16E-10 | 4.55E-05 |
| 2 | 11148994 | A | G | 0.838432 | 1.16E-10 | 4.55E-05 |
| 2 | 11149671 | T | C | 0.838432 | 1.16E-10 | 4.55E-05 |
| 2 | 11150886 | T | C | 0.838432 | 1.16E-10 | 4.55E-05 |
| 2 | 11150976 | C | A | 0.838432 | 1.16E-10 | 4.55E-05 |
| 2 | 11151112 | G | A | 0.838432 | 1.16E-10 | 4.55E-05 |
| 2 | 11151207 | C | G | 0.838432 | 1.16E-10 | 4.55E-05 |
| 2 | 11151258 | A | G | 0.838432 | 1.16E-10 | 4.55E-05 |
| 2 | 11151275 | A | G | 0.838432 | 1.16E-10 | 4.55E-05 |
| 2 | 11151721 | A | G | 0.838432 | 1.16E-10 | 4.55E-05 |
| 2 | 11151724 | C | T | 0.838432 | 1.16E-10 | 4.55E-05 |
| 2 | 11153215 | C | G | 0.838432 | 1.16E-10 | 4.55E-05 |
| 2 | 11154436 | G | A | 0.838432 | 1.16E-10 | 4.55E-05 |
| 2 | 11154582 | A | G | 0.838432 | 1.16E-10 | 4.55E-05 |
| 2 | 11155944 | G | A | 0.838432 | 1.16E-10 | 4.55E-05 |
| 2 | 11263408 | G | C | 0.837476 | 1.20E-10 | 4.69E-05 |
| 2 | 11270845 | A | G | 0.866157 | 1.20E-10 | 4.69E-05 |
| 2 | 11270932 | T | C | 0.866157 | 1.20E-10 | 4.69E-05 |
| 2 | 11270962 | T | A | 0.866157 | 1.20E-10 | 4.69E-05 |
| 2 | 11272339 | T | C | 0.866157 | 1.20E-10 | 4.69E-05 |
| 2 | 11272357 | T | G | 0.866157 | 1.20E-10 | 4.69E-05 |
| 2 | 11272774 | A | G | 0.866157 | 1.20E-10 | 4.69E-05 |
| 2 | 11273008 | T | C | 0.866157 | 1.20E-10 | 4.69E-05 |
| 2 | 11273320 | A | C | 0.866157 | 1.20E-10 | 4.69E-05 |
| 2 | 11274671 | T | C | 0.866157 | 1.20E-10 | 4.69E-05 |
| 2 | 11275420 | A | G | 0.866157 | 1.20E-10 | 4.69E-05 |
| 2 | 11275474 | C | G | 0.866157 | 1.20E-10 | 4.69E-05 |
| 2 | 11275669 | A | G | 0.866157 | 1.20E-10 | 4.69E-05 |
| 2 | 11276101 | T | G | 0.866157 | 1.20E-10 | 4.69E-05 |
| 2 | 11276288 | T | C | 0.866157 | 1.20E-10 | 4.69E-05 |
| 2 | 11276585 | G | A | 0.866157 | 1.20E-10 | 4.69E-05 |
| 2 | 11276606 | G | T | 0.866157 | 1.20E-10 | 4.69E-05 |
| 2 | 11277056 | T | C | 0.866157 | 1.20E-10 | 4.69E-05 |
| 2 | 11277696 | G | C | 0.866157 | 1.20E-10 | 4.69E-05 |
| 2 | 11277782 | G | A | 0.866157 | 1.20E-10 | 4.69E-05 |

| | | | | | | |
|---|---|---|---|---|---|---|
| 2 | 11277834 | G | A | 0.866157 | 1.20E-10 | 4.69E-05 |
| 2 | 11282867 | C | T | 0.866157 | 1.20E-10 | 4.69E-05 |
| 2 | 11282974 | C | T | 0.866157 | 1.20E-10 | 4.69E-05 |
| 2 | 11283126 | A | G | 0.866157 | 1.20E-10 | 4.69E-05 |
| 2 | 11283211 | G | A | 0.866157 | 1.20E-10 | 4.69E-05 |
| 2 | 11283722 | T | A | 0.866157 | 1.20E-10 | 4.69E-05 |
| 2 | 11283829 | C | G | 0.866157 | 1.20E-10 | 4.69E-05 |
| 2 | 11283873 | T | C | 0.866157 | 1.20E-10 | 4.69E-05 |
| 2 | 11283970 | A | G | 0.866157 | 1.20E-10 | 4.69E-05 |
| 2 | 11284024 | G | A | 0.866157 | 1.20E-10 | 4.69E-05 |
| 2 | 11284025 | T | A | 0.866157 | 1.20E-10 | 4.69E-05 |
| 2 | 11284322 | G | T | 0.866157 | 1.20E-10 | 4.69E-05 |
| 2 | 11284532 | A | G | 0.866157 | 1.20E-10 | 4.69E-05 |
| 2 | 11284732 | T | C | 0.866157 | 1.20E-10 | 4.69E-05 |
| 2 | 11284757 | G | C | 0.866157 | 1.20E-10 | 4.69E-05 |
| 2 | 11285043 | A | T | 0.866157 | 1.20E-10 | 4.69E-05 |
| 2 | 11285666 | G | A | 0.866157 | 1.20E-10 | 4.69E-05 |
| 2 | 11285671 | A | C | 0.866157 | 1.20E-10 | 4.69E-05 |
| 2 | 11285681 | G | A | 0.866157 | 1.20E-10 | 4.69E-05 |
| 2 | 11285841 | C | T | 0.866157 | 1.20E-10 | 4.69E-05 |
| 2 | 11285848 | A | G | 0.866157 | 1.20E-10 | 4.69E-05 |
| 2 | 11285919 | T | C | 0.866157 | 1.20E-10 | 4.69E-05 |
| 2 | 11286177 | A | G | 0.866157 | 1.20E-10 | 4.69E-05 |
| 2 | 11286415 | G | A | 0.866157 | 1.20E-10 | 4.69E-05 |
| 2 | 11286418 | C | T | 0.866157 | 1.20E-10 | 4.69E-05 |
| 2 | 11286458 | A | G | 0.866157 | 1.20E-10 | 4.69E-05 |
| 2 | 11286612 | T | A | 0.866157 | 1.20E-10 | 4.69E-05 |
| 2 | 11286774 | G | A | 0.866157 | 1.20E-10 | 4.69E-05 |
| 2 | 11286900 | T | C | 0.866157 | 1.20E-10 | 4.69E-05 |
| 2 | 11286916 | C | A | 0.866157 | 1.20E-10 | 4.69E-05 |
| 2 | 11286970 | G | A | 0.866157 | 1.20E-10 | 4.69E-05 |
| 2 | 11287106 | T | C | 0.866157 | 1.20E-10 | 4.69E-05 |
| 2 | 11287909 | G | A | 0.866157 | 1.20E-10 | 4.69E-05 |
| 2 | 11288056 | A | G | 0.866157 | 1.20E-10 | 4.69E-05 |
| 2 | 11288248 | C | A | 0.866157 | 1.20E-10 | 4.69E-05 |
| 2 | 11288569 | T | C | 0.866157 | 1.20E-10 | 4.69E-05 |
| 2 | 11288995 | G | A | 0.866157 | 1.20E-10 | 4.69E-05 |
| 2 | 11289158 | G | A | 0.866157 | 1.20E-10 | 4.69E-05 |
| 2 | 11289369 | C | T | 0.866157 | 1.20E-10 | 4.69E-05 |
| 2 | 11289412 | G | T | 0.866157 | 1.20E-10 | 4.69E-05 |
| 2 | 11289467 | T | C | 0.866157 | 1.20E-10 | 4.69E-05 |
| 2 | 11289574 | A | G | 0.866157 | 1.20E-10 | 4.69E-05 |
| 2 | 11289609 | A | T | 0.866157 | 1.20E-10 | 4.69E-05 |
| 2 | 11289672 | T | C | 0.866157 | 1.20E-10 | 4.69E-05 |
| 2 | 11289694 | T | C | 0.866157 | 1.20E-10 | 4.69E-05 |
| 2 | 11289766 | A | G | 0.866157 | 1.20E-10 | 4.69E-05 |
| 2 | 11289772 | C | G | 0.866157 | 1.20E-10 | 4.69E-05 |
| 2 | 11289812 | G | A | 0.866157 | 1.20E-10 | 4.69E-05 |
| 2 | 11289853 | T | C | 0.866157 | 1.20E-10 | 4.69E-05 |
| 2 | 11289860 | A | G | 0.866157 | 1.20E-10 | 4.69E-05 |
| 2 | 11289861 | T | C | 0.866157 | 1.20E-10 | 4.69E-05 |
| 2 | 11289889 | G | A | 0.866157 | 1.20E-10 | 4.69E-05 |
| 2 | 11289935 | T | C | 0.866157 | 1.20E-10 | 4.69E-05 |
| 2 | 11290103 | G | A | 0.866157 | 1.20E-10 | 4.69E-05 |
| 2 | 11290160 | C | T | 0.866157 | 1.20E-10 | 4.69E-05 |
| 2 | 11290212 | T | C | 0.866157 | 1.20E-10 | 4.69E-05 |
| 2 | 11290253 | G | A | 0.866157 | 1.20E-10 | 4.69E-05 |
| 2 | 11290471 | T | A | 0.866157 | 1.20E-10 | 4.69E-05 |
| 2 | 11290846 | G | A | 0.866157 | 1.20E-10 | 4.69E-05 |
| 2 | 11291059 | G | A | 0.866157 | 1.20E-10 | 4.69E-05 |
| 2 | 11291142 | C | A | 0.866157 | 1.20E-10 | 4.69E-05 |
| 2 | 11291853 | A | G | 0.866157 | 1.20E-10 | 4.69E-05 |
| 2 | 11291873 | C | T | 0.866157 | 1.20E-10 | 4.69E-05 |
| 2 | 11292081 | A | G | 0.866157 | 1.20E-10 | 4.69E-05 |
| 2 | 11292553 | T | C | 0.866157 | 1.20E-10 | 4.69E-05 |
| 2 | 11292762 | T | C | 0.866157 | 1.20E-10 | 4.69E-05 |
| 2 | 11293239 | T | C | 0.866157 | 1.20E-10 | 4.69E-05 |

| | | | | | | |
|---|---|---|---|---|---|---|
| 2 | 11293844 | T | G | 0.866157 | 1.20E-10 | 4.69E-05 |
| 2 | 11293848 | G | A | 0.866157 | 1.20E-10 | 4.69E-05 |
| 2 | 11294999 | T | C | 0.866157 | 1.20E-10 | 4.69E-05 |
| 2 | 11295776 | T | A | 0.866157 | 1.20E-10 | 4.69E-05 |
| 2 | 11295906 | G | A | 0.866157 | 1.20E-10 | 4.69E-05 |
| 2 | 11295940 | A | G | 0.866157 | 1.20E-10 | 4.69E-05 |
| 2 | 11295964 | A | G | 0.866157 | 1.20E-10 | 4.69E-05 |
| 2 | 11296147 | T | C | 0.866157 | 1.20E-10 | 4.69E-05 |
| 2 | 11296148 | C | T | 0.866157 | 1.20E-10 | 4.69E-05 |
| 2 | 11296567 | G | A | 0.866157 | 1.20E-10 | 4.69E-05 |
| 2 | 11297146 | C | T | 0.866157 | 1.20E-10 | 4.69E-05 |
| 2 | 11297256 | C | T | 0.866157 | 1.20E-10 | 4.69E-05 |
| 2 | 11297761 | A | T | 0.866157 | 1.20E-10 | 4.69E-05 |
| 2 | 11297762 | T | G | 0.866157 | 1.20E-10 | 4.69E-05 |
| 2 | 11297763 | G | C | 0.866157 | 1.20E-10 | 4.69E-05 |
| 2 | 11297892 | A | T | 0.866157 | 1.20E-10 | 4.69E-05 |
| 2 | 11297975 | A | C | 0.866157 | 1.20E-10 | 4.69E-05 |
| 2 | 11298030 | A | G | 0.866157 | 1.20E-10 | 4.69E-05 |
| 2 | 11298051 | A | C | 0.866157 | 1.20E-10 | 4.69E-05 |
| 2 | 11298112 | C | T | 0.866157 | 1.20E-10 | 4.69E-05 |
| 2 | 11299192 | G | A | 0.866157 | 1.20E-10 | 4.69E-05 |
| 2 | 11299355 | C | T | 0.866157 | 1.20E-10 | 4.69E-05 |
| 2 | 11299405 | G | A | 0.866157 | 1.20E-10 | 4.69E-05 |
| 2 | 11299588 | C | T | 0.866157 | 1.20E-10 | 4.69E-05 |
| 2 | 11301070 | T | C | 0.866157 | 1.20E-10 | 4.69E-05 |
| 2 | 11301108 | A | G | 0.866157 | 1.20E-10 | 4.69E-05 |
| 2 | 11301138 | G | T | 0.866157 | 1.20E-10 | 4.69E-05 |
| 2 | 11301174 | T | C | 0.866157 | 1.20E-10 | 4.69E-05 |
| 2 | 11301186 | T | C | 0.866157 | 1.20E-10 | 4.69E-05 |
| 2 | 11301344 | T | C | 0.866157 | 1.20E-10 | 4.69E-05 |
| 2 | 11301456 | G | A | 0.866157 | 1.20E-10 | 4.69E-05 |
| 2 | 11301693 | A | C | 0.866157 | 1.20E-10 | 4.69E-05 |
| 2 | 11302048 | G | A | 0.866157 | 1.20E-10 | 4.69E-05 |
| 2 | 11304529 | A | C | 0.866157 | 1.20E-10 | 4.69E-05 |
| 2 | 11305851 | G | A | 0.866157 | 1.20E-10 | 4.69E-05 |
| 2 | 11309397 | A | G | 0.866157 | 1.20E-10 | 4.69E-05 |
| 2 | 11309724 | G | A | 0.866157 | 1.20E-10 | 4.69E-05 |
| 2 | 11310236 | C | T | 0.866157 | 1.20E-10 | 4.69E-05 |
| 2 | 11165001 | C | T | 0.83652 | 1.22E-10 | 4.78E-05 |
| 2 | 11282669 | G | A | 0.865201 | 1.23E-10 | 4.81E-05 |
| 2 | 11282698 | A | G | 0.865201 | 1.23E-10 | 4.81E-05 |
| 2 | 11125619 | A | T | 0.835564 | 1.27E-10 | 4.98E-05 |
| 2 | 8666854 | T | C | 0.878585 | 1.27E-10 | 5.00E-05 |
| 2 | 8668480 | C | T | 0.878585 | 1.27E-10 | 5.00E-05 |
| 2 | 8668481 | A | G | 0.878585 | 1.27E-10 | 5.00E-05 |
| 2 | 8668493 | G | C | 0.878585 | 1.27E-10 | 5.00E-05 |
| 2 | 8668764 | C | G | 0.878585 | 1.27E-10 | 5.00E-05 |
| 2 | 8668845 | C | T | 0.878585 | 1.27E-10 | 5.00E-05 |
| 2 | 8668877 | C | G | 0.878585 | 1.27E-10 | 5.00E-05 |
| 2 | 8669040 | T | C | 0.878585 | 1.27E-10 | 5.00E-05 |
| 2 | 8672459 | T | G | 0.878585 | 1.27E-10 | 5.00E-05 |
| 2 | 8674858 | T | C | 0.878585 | 1.27E-10 | 5.00E-05 |
| 2 | 8674986 | A | G | 0.878585 | 1.27E-10 | 5.00E-05 |
| 2 | 8686283 | A | G | 0.878585 | 1.27E-10 | 5.00E-05 |
| 2 | 11099797 | T | C | 0.837476 | 1.28E-10 | 5.03E-05 |
| 2 | 11104939 | T | C | 0.837476 | 1.28E-10 | 5.03E-05 |
| 2 | 11266174 | T | G | 0.867113 | 1.29E-10 | 5.06E-05 |
| 2 | 11214376 | T | C | 0.83652 | 1.34E-10 | 5.25E-05 |
| 2 | 11215211 | C | A | 0.83652 | 1.34E-10 | 5.25E-05 |
| 2 | 11227800 | T | C | 0.83652 | 1.34E-10 | 5.25E-05 |
| 2 | 11232073 | G | A | 0.83652 | 1.34E-10 | 5.25E-05 |
| 2 | 11232119 | C | T | 0.83652 | 1.34E-10 | 5.25E-05 |
| 2 | 11232125 | G | A | 0.83652 | 1.34E-10 | 5.25E-05 |
| 2 | 11232145 | A | G | 0.83652 | 1.34E-10 | 5.25E-05 |
| 2 | 11235115 | T | C | 0.83652 | 1.34E-10 | 5.25E-05 |
| 2 | 11235125 | G | A | 0.83652 | 1.34E-10 | 5.25E-05 |
| 2 | 11245144 | A | G | 0.83652 | 1.34E-10 | 5.25E-05 |

| | | | | | | |
|---|---|---|---|---|---|---|
| 2 | 11247787 | A | G | 0.83652 | 1.34E-10 | 5.25E-05 |
| 2 | 11247907 | T | C | 0.83652 | 1.34E-10 | 5.25E-05 |
| 2 | 11247917 | G | C | 0.83652 | 1.34E-10 | 5.25E-05 |
| 2 | 11248059 | C | T | 0.83652 | 1.34E-10 | 5.25E-05 |
| 2 | 11248130 | A | G | 0.83652 | 1.34E-10 | 5.25E-05 |
| 2 | 11248352 | G | A | 0.83652 | 1.34E-10 | 5.25E-05 |
| 2 | 11248568 | C | T | 0.83652 | 1.34E-10 | 5.25E-05 |
| 2 | 11248574 | C | T | 0.83652 | 1.34E-10 | 5.25E-05 |
| 2 | 11248983 | A | G | 0.83652 | 1.34E-10 | 5.25E-05 |
| 2 | 11253357 | T | C | 0.83652 | 1.34E-10 | 5.25E-05 |
| 2 | 11253485 | G | T | 0.83652 | 1.34E-10 | 5.25E-05 |
| 2 | 11254561 | T | C | 0.83652 | 1.34E-10 | 5.25E-05 |
| 2 | 11254609 | T | A | 0.83652 | 1.34E-10 | 5.25E-05 |
| 2 | 11254654 | C | T | 0.83652 | 1.34E-10 | 5.25E-05 |
| 2 | 11254708 | T | C | 0.83652 | 1.34E-10 | 5.25E-05 |
| 2 | 11254712 | C | G | 0.83652 | 1.34E-10 | 5.25E-05 |
| 2 | 11254720 | A | G | 0.83652 | 1.34E-10 | 5.25E-05 |
| 2 | 11254736 | C | T | 0.83652 | 1.34E-10 | 5.25E-05 |
| 2 | 11254779 | C | G | 0.83652 | 1.34E-10 | 5.25E-05 |
| 2 | 11254841 | T | C | 0.83652 | 1.34E-10 | 5.25E-05 |
| 2 | 11254900 | A | G | 0.83652 | 1.34E-10 | 5.25E-05 |
| 2 | 11254917 | G | T | 0.83652 | 1.34E-10 | 5.25E-05 |
| 2 | 11254931 | T | C | 0.83652 | 1.34E-10 | 5.25E-05 |
| 2 | 11254942 | A | T | 0.83652 | 1.34E-10 | 5.25E-05 |
| 2 | 11254959 | A | C | 0.83652 | 1.34E-10 | 5.25E-05 |
| 2 | 11255110 | C | T | 0.83652 | 1.34E-10 | 5.25E-05 |
| 2 | 11255358 | T | C | 0.83652 | 1.34E-10 | 5.25E-05 |
| 2 | 11255475 | A | G | 0.83652 | 1.34E-10 | 5.25E-05 |
| 2 | 11255601 | A | G | 0.83652 | 1.34E-10 | 5.25E-05 |
| 2 | 11255611 | C | G | 0.83652 | 1.34E-10 | 5.25E-05 |
| 2 | 11255678 | A | G | 0.83652 | 1.34E-10 | 5.25E-05 |
| 2 | 11255710 | A | G | 0.83652 | 1.34E-10 | 5.25E-05 |
| 2 | 11255777 | C | G | 0.83652 | 1.34E-10 | 5.25E-05 |
| 2 | 11255802 | T | G | 0.83652 | 1.34E-10 | 5.25E-05 |
| 2 | 11256419 | A | G | 0.83652 | 1.34E-10 | 5.25E-05 |
| 2 | 11256490 | A | C | 0.83652 | 1.34E-10 | 5.25E-05 |
| 2 | 11256522 | T | C | 0.83652 | 1.34E-10 | 5.25E-05 |
| 2 | 11256587 | C | A | 0.83652 | 1.34E-10 | 5.25E-05 |
| 2 | 11256594 | G | A | 0.83652 | 1.34E-10 | 5.25E-05 |
| 2 | 11256598 | G | A | 0.83652 | 1.34E-10 | 5.25E-05 |
| 2 | 11256643 | T | A | 0.83652 | 1.34E-10 | 5.25E-05 |
| 2 | 11256646 | G | A | 0.83652 | 1.34E-10 | 5.25E-05 |
| 2 | 11256648 | C | T | 0.83652 | 1.34E-10 | 5.25E-05 |
| 2 | 11256661 | A | G | 0.83652 | 1.34E-10 | 5.25E-05 |
| 2 | 11256670 | A | G | 0.83652 | 1.34E-10 | 5.25E-05 |
| 2 | 11256677 | G | T | 0.83652 | 1.34E-10 | 5.25E-05 |
| 2 | 11256681 | C | T | 0.83652 | 1.34E-10 | 5.25E-05 |
| 2 | 11256685 | G | A | 0.83652 | 1.34E-10 | 5.25E-05 |
| 2 | 11256690 | A | G | 0.83652 | 1.34E-10 | 5.25E-05 |
| 2 | 11256691 | T | G | 0.83652 | 1.34E-10 | 5.25E-05 |
| 2 | 11256724 | G | A | 0.83652 | 1.34E-10 | 5.25E-05 |
| 2 | 11256737 | G | A | 0.83652 | 1.34E-10 | 5.25E-05 |
| 2 | 11256767 | C | A | 0.83652 | 1.34E-10 | 5.25E-05 |
| 2 | 11256823 | C | T | 0.83652 | 1.34E-10 | 5.25E-05 |
| 2 | 11256835 | C | A | 0.83652 | 1.34E-10 | 5.25E-05 |
| 2 | 11257071 | A | C | 0.83652 | 1.34E-10 | 5.25E-05 |
| 2 | 11257142 | C | T | 0.83652 | 1.34E-10 | 5.25E-05 |
| 2 | 11257285 | C | A | 0.83652 | 1.34E-10 | 5.25E-05 |
| 2 | 11257288 | G | A | 0.83652 | 1.34E-10 | 5.25E-05 |
| 2 | 11257465 | A | G | 0.83652 | 1.34E-10 | 5.25E-05 |
| 2 | 11263746 | C | T | 0.83652 | 1.34E-10 | 5.25E-05 |
| 2 | 11263772 | G | T | 0.83652 | 1.34E-10 | 5.25E-05 |
| 2 | 11263913 | C | A | 0.83652 | 1.34E-10 | 5.25E-05 |
| 2 | 11263991 | A | G | 0.83652 | 1.34E-10 | 5.25E-05 |
| 2 | 11264138 | G | A | 0.83652 | 1.34E-10 | 5.25E-05 |
| 2 | 11264160 | A | G | 0.83652 | 1.34E-10 | 5.25E-05 |
| 2 | 11264184 | G | A | 0.83652 | 1.34E-10 | 5.25E-05 |

| | | | | | | |
|---|---|---|---|---|---|---|
| 2 | 11264521 | T | C | 0.83652 | 1.34E-10 | 5.25E-05 |
| 2 | 11264651 | C | T | 0.83652 | 1.34E-10 | 5.25E-05 |
| 2 | 11264861 | C | T | 0.83652 | 1.34E-10 | 5.25E-05 |
| 2 | 11265698 | A | G | 0.83652 | 1.34E-10 | 5.25E-05 |
| 2 | 10639215 | C | G | 0.823136 | 1.45E-10 | 5.69E-05 |
| 2 | 10639245 | A | G | 0.823136 | 1.45E-10 | 5.69E-05 |
| 2 | 10639300 | A | C | 0.823136 | 1.45E-10 | 5.69E-05 |
| 2 | 10639313 | G | A | 0.823136 | 1.45E-10 | 5.69E-05 |
| 2 | 10639391 | A | G | 0.823136 | 1.45E-10 | 5.69E-05 |
| 2 | 11129699 | T | G | 0.83652 | 1.52E-10 | 5.96E-05 |
| 2 | 11129709 | A | G | 0.83652 | 1.52E-10 | 5.96E-05 |
| 2 | 11129714 | C | A | 0.83652 | 1.52E-10 | 5.96E-05 |
| 2 | 11129747 | T | C | 0.83652 | 1.52E-10 | 5.96E-05 |
| 2 | 11263428 | T | G | 0.839388 | 1.53E-10 | 5.98E-05 |
| 2 | 11261156 | T | C | 0.838432 | 1.63E-10 | 6.40E-05 |
| 2 | 11279809 | C | T | 0.864245 | 1.68E-10 | 6.57E-05 |
| 2 | 11283632 | G | C | 0.864245 | 1.68E-10 | 6.57E-05 |
| 2 | 8278925 | A | G | 0.877629 | 1.70E-10 | 6.68E-05 |
| 2 | 11214103 | T | C | 0.838432 | 1.71E-10 | 6.71E-05 |
| 2 | 11214265 | C | A | 0.838432 | 1.71E-10 | 6.71E-05 |
| 2 | 11215235 | C | G | 0.838432 | 1.71E-10 | 6.71E-05 |
| 2 | 11216076 | G | A | 0.838432 | 1.71E-10 | 6.71E-05 |
| 2 | 11216218 | C | T | 0.838432 | 1.71E-10 | 6.71E-05 |
| 2 | 11216247 | T | C | 0.838432 | 1.71E-10 | 6.71E-05 |
| 2 | 11216264 | T | C | 0.838432 | 1.71E-10 | 6.71E-05 |
| 2 | 11216848 | C | G | 0.838432 | 1.71E-10 | 6.71E-05 |
| 2 | 11216857 | G | A | 0.838432 | 1.71E-10 | 6.71E-05 |
| 2 | 11217261 | G | A | 0.838432 | 1.71E-10 | 6.71E-05 |
| 2 | 11217503 | G | C | 0.838432 | 1.71E-10 | 6.71E-05 |
| 2 | 11217837 | C | T | 0.838432 | 1.71E-10 | 6.71E-05 |
| 2 | 11217838 | T | G | 0.838432 | 1.71E-10 | 6.71E-05 |
| 2 | 11218299 | T | C | 0.838432 | 1.71E-10 | 6.71E-05 |
| 2 | 11219438 | A | G | 0.838432 | 1.71E-10 | 6.71E-05 |
| 2 | 11219994 | C | A | 0.838432 | 1.71E-10 | 6.71E-05 |
| 2 | 11220152 | A | C | 0.838432 | 1.71E-10 | 6.71E-05 |
| 2 | 11220221 | T | C | 0.838432 | 1.71E-10 | 6.71E-05 |
| 2 | 11220872 | G | A | 0.838432 | 1.71E-10 | 6.71E-05 |
| 2 | 11221526 | A | G | 0.838432 | 1.71E-10 | 6.71E-05 |
| 2 | 11221727 | A | C | 0.838432 | 1.71E-10 | 6.71E-05 |
| 2 | 11222182 | A | C | 0.838432 | 1.71E-10 | 6.71E-05 |
| 2 | 11222696 | A | G | 0.838432 | 1.71E-10 | 6.71E-05 |
| 2 | 11222698 | A | G | 0.838432 | 1.71E-10 | 6.71E-05 |
| 2 | 11222887 | C | T | 0.838432 | 1.71E-10 | 6.71E-05 |
| 2 | 11222932 | G | A | 0.838432 | 1.71E-10 | 6.71E-05 |
| 2 | 11223331 | T | C | 0.838432 | 1.71E-10 | 6.71E-05 |
| 2 | 11223749 | A | G | 0.838432 | 1.71E-10 | 6.71E-05 |
| 2 | 11223978 | G | A | 0.838432 | 1.71E-10 | 6.71E-05 |
| 2 | 11224077 | T | G | 0.838432 | 1.71E-10 | 6.71E-05 |
| 2 | 11224774 | A | T | 0.838432 | 1.71E-10 | 6.71E-05 |
| 2 | 11225193 | T | C | 0.838432 | 1.71E-10 | 6.71E-05 |
| 2 | 11225420 | A | G | 0.838432 | 1.71E-10 | 6.71E-05 |
| 2 | 11225521 | A | T | 0.838432 | 1.71E-10 | 6.71E-05 |
| 2 | 11226129 | A | G | 0.838432 | 1.71E-10 | 6.71E-05 |
| 2 | 11226647 | A | T | 0.838432 | 1.71E-10 | 6.71E-05 |
| 2 | 11226711 | C | T | 0.838432 | 1.71E-10 | 6.71E-05 |
| 2 | 11226872 | C | T | 0.838432 | 1.71E-10 | 6.71E-05 |
| 2 | 11227079 | G | A | 0.838432 | 1.71E-10 | 6.71E-05 |
| 2 | 11227327 | C | A | 0.838432 | 1.71E-10 | 6.71E-05 |
| 2 | 11227339 | G | A | 0.838432 | 1.71E-10 | 6.71E-05 |
| 2 | 11227371 | A | G | 0.838432 | 1.71E-10 | 6.71E-05 |
| 2 | 11227591 | C | T | 0.838432 | 1.71E-10 | 6.71E-05 |
| 2 | 11227678 | A | G | 0.838432 | 1.71E-10 | 6.71E-05 |
| 2 | 11227909 | C | T | 0.838432 | 1.71E-10 | 6.71E-05 |
| 2 | 11227910 | A | G | 0.838432 | 1.71E-10 | 6.71E-05 |
| 2 | 11227982 | C | T | 0.838432 | 1.71E-10 | 6.71E-05 |
| 2 | 11228061 | C | T | 0.838432 | 1.71E-10 | 6.71E-05 |
| 2 | 11228185 | G | A | 0.838432 | 1.71E-10 | 6.71E-05 |

| | | | | | | |
|---|---|---|---|---|---|---|
| 2 | 11228301 | T | A | 0.838432 | 1.71E-10 | 6.71E-05 |
| 2 | 11228463 | C | G | 0.838432 | 1.71E-10 | 6.71E-05 |
| 2 | 11228644 | A | T | 0.838432 | 1.71E-10 | 6.71E-05 |
| 2 | 11228729 | A | G | 0.838432 | 1.71E-10 | 6.71E-05 |
| 2 | 11228773 | A | G | 0.838432 | 1.71E-10 | 6.71E-05 |
| 2 | 11228848 | C | T | 0.838432 | 1.71E-10 | 6.71E-05 |
| 2 | 11229139 | A | G | 0.838432 | 1.71E-10 | 6.71E-05 |
| 2 | 11229342 | G | C | 0.838432 | 1.71E-10 | 6.71E-05 |
| 2 | 11229675 | A | G | 0.838432 | 1.71E-10 | 6.71E-05 |
| 2 | 11231304 | G | C | 0.838432 | 1.71E-10 | 6.71E-05 |
| 2 | 11231435 | C | T | 0.838432 | 1.71E-10 | 6.71E-05 |
| 2 | 11232446 | A | G | 0.838432 | 1.71E-10 | 6.71E-05 |
| 2 | 11232448 | C | T | 0.838432 | 1.71E-10 | 6.71E-05 |
| 2 | 11232451 | G | A | 0.838432 | 1.71E-10 | 6.71E-05 |
| 2 | 11232460 | C | T | 0.838432 | 1.71E-10 | 6.71E-05 |
| 2 | 11232535 | T | G | 0.838432 | 1.71E-10 | 6.71E-05 |
| 2 | 11233486 | T | G | 0.838432 | 1.71E-10 | 6.71E-05 |
| 2 | 11233497 | A | G | 0.838432 | 1.71E-10 | 6.71E-05 |
| 2 | 11234163 | G | A | 0.838432 | 1.71E-10 | 6.71E-05 |
| 2 | 11236087 | C | T | 0.838432 | 1.71E-10 | 6.71E-05 |
| 2 | 11236123 | C | T | 0.838432 | 1.71E-10 | 6.71E-05 |
| 2 | 11236360 | A | C | 0.838432 | 1.71E-10 | 6.71E-05 |
| 2 | 11241430 | C | T | 0.838432 | 1.71E-10 | 6.71E-05 |
| 2 | 11241551 | A | G | 0.838432 | 1.71E-10 | 6.71E-05 |
| 2 | 11241591 | A | T | 0.838432 | 1.71E-10 | 6.71E-05 |
| 2 | 11241661 | T | A | 0.838432 | 1.71E-10 | 6.71E-05 |
| 2 | 11241757 | G | A | 0.838432 | 1.71E-10 | 6.71E-05 |
| 2 | 11242056 | G | A | 0.838432 | 1.71E-10 | 6.71E-05 |
| 2 | 11242182 | G | C | 0.838432 | 1.71E-10 | 6.71E-05 |
| 2 | 11242257 | G | A | 0.838432 | 1.71E-10 | 6.71E-05 |
| 2 | 11242258 | G | A | 0.838432 | 1.71E-10 | 6.71E-05 |
| 2 | 11242463 | T | C | 0.838432 | 1.71E-10 | 6.71E-05 |
| 2 | 11242767 | T | G | 0.838432 | 1.71E-10 | 6.71E-05 |
| 2 | 11243091 | C | T | 0.838432 | 1.71E-10 | 6.71E-05 |
| 2 | 11243101 | C | A | 0.838432 | 1.71E-10 | 6.71E-05 |
| 2 | 11243205 | C | T | 0.838432 | 1.71E-10 | 6.71E-05 |
| 2 | 11243207 | A | G | 0.838432 | 1.71E-10 | 6.71E-05 |
| 2 | 11243561 | A | G | 0.838432 | 1.71E-10 | 6.71E-05 |
| 2 | 11243593 | T | C | 0.838432 | 1.71E-10 | 6.71E-05 |
| 2 | 11243761 | G | A | 0.838432 | 1.71E-10 | 6.71E-05 |
| 2 | 11243782 | A | G | 0.838432 | 1.71E-10 | 6.71E-05 |
| 2 | 11243812 | A | G | 0.838432 | 1.71E-10 | 6.71E-05 |
| 2 | 11245733 | A | G | 0.838432 | 1.71E-10 | 6.71E-05 |
| 2 | 11247131 | G | T | 0.838432 | 1.71E-10 | 6.71E-05 |
| 2 | 11247180 | G | A | 0.838432 | 1.71E-10 | 6.71E-05 |
| 2 | 11247242 | A | G | 0.838432 | 1.71E-10 | 6.71E-05 |
| 2 | 11247259 | T | C | 0.838432 | 1.71E-10 | 6.71E-05 |
| 2 | 11247967 | A | G | 0.838432 | 1.71E-10 | 6.71E-05 |
| 2 | 11248711 | G | A | 0.838432 | 1.71E-10 | 6.71E-05 |
| 2 | 11249669 | C | A | 0.838432 | 1.71E-10 | 6.71E-05 |
| 2 | 11249769 | T | G | 0.838432 | 1.71E-10 | 6.71E-05 |
| 2 | 11250407 | G | A | 0.838432 | 1.71E-10 | 6.71E-05 |
| 2 | 11251054 | T | C | 0.838432 | 1.71E-10 | 6.71E-05 |
| 2 | 11251097 | A | T | 0.838432 | 1.71E-10 | 6.71E-05 |
| 2 | 11251117 | T | C | 0.838432 | 1.71E-10 | 6.71E-05 |
| 2 | 11253021 | A | G | 0.838432 | 1.71E-10 | 6.71E-05 |
| 2 | 11253075 | C | G | 0.838432 | 1.71E-10 | 6.71E-05 |
| 2 | 11253195 | C | T | 0.838432 | 1.71E-10 | 6.71E-05 |
| 2 | 11253285 | A | G | 0.838432 | 1.71E-10 | 6.71E-05 |
| 2 | 11253562 | T | C | 0.838432 | 1.71E-10 | 6.71E-05 |
| 2 | 11253813 | A | G | 0.838432 | 1.71E-10 | 6.71E-05 |
| 2 | 11260065 | C | A | 0.838432 | 1.71E-10 | 6.71E-05 |
| 2 | 11260246 | A | G | 0.838432 | 1.71E-10 | 6.71E-05 |
| 2 | 11260288 | T | A | 0.838432 | 1.71E-10 | 6.71E-05 |
| 2 | 11260291 | A | G | 0.838432 | 1.71E-10 | 6.71E-05 |
| 2 | 11260323 | G | A | 0.838432 | 1.71E-10 | 6.71E-05 |
| 2 | 11260327 | T | C | 0.838432 | 1.71E-10 | 6.71E-05 |

| | | | | | | |
|---|---|---|---|---|---|---|
| 2 | 11260328 | G | A | 0.838432 | 1.71E-10 | 6.71E-05 |
| 2 | 11260602 | T | C | 0.838432 | 1.71E-10 | 6.71E-05 |
| 2 | 11260615 | T | C | 0.838432 | 1.71E-10 | 6.71E-05 |
| 2 | 11260736 | T | G | 0.838432 | 1.71E-10 | 6.71E-05 |
| 2 | 11263711 | G | T | 0.838432 | 1.71E-10 | 6.71E-05 |
| 2 | 11263737 | C | T | 0.838432 | 1.71E-10 | 6.71E-05 |
| 2 | 11263873 | T | C | 0.838432 | 1.71E-10 | 6.71E-05 |
| 2 | 11264058 | G | A | 0.838432 | 1.71E-10 | 6.71E-05 |
| 2 | 11264070 | T | G | 0.838432 | 1.71E-10 | 6.71E-05 |
| 2 | 11264180 | A | G | 0.838432 | 1.71E-10 | 6.71E-05 |
| 2 | 11264195 | C | T | 0.838432 | 1.71E-10 | 6.71E-05 |
| 2 | 11264446 | A | G | 0.838432 | 1.71E-10 | 6.71E-05 |
| 2 | 11264488 | G | T | 0.838432 | 1.71E-10 | 6.71E-05 |
| 2 | 11264493 | T | G | 0.838432 | 1.71E-10 | 6.71E-05 |
| 2 | 11264851 | G | A | 0.838432 | 1.71E-10 | 6.71E-05 |
| 2 | 11264869 | T | C | 0.838432 | 1.71E-10 | 6.71E-05 |
| 2 | 11265117 | C | T | 0.838432 | 1.71E-10 | 6.71E-05 |
| 2 | 11265159 | A | T | 0.838432 | 1.71E-10 | 6.71E-05 |
| 2 | 11265160 | T | G | 0.838432 | 1.71E-10 | 6.71E-05 |
| 2 | 11265161 | T | A | 0.838432 | 1.71E-10 | 6.71E-05 |
| 2 | 11265268 | T | G | 0.838432 | 1.71E-10 | 6.71E-05 |
| 2 | 11265563 | A | G | 0.838432 | 1.71E-10 | 6.71E-05 |
| 2 | 11265798 | T | G | 0.838432 | 1.71E-10 | 6.71E-05 |
| 2 | 11265906 | C | T | 0.838432 | 1.71E-10 | 6.71E-05 |
| 2 | 11265907 | C | A | 0.838432 | 1.71E-10 | 6.71E-05 |
| 2 | 11265923 | A | G | 0.838432 | 1.71E-10 | 6.71E-05 |
| 2 | 11265991 | C | T | 0.838432 | 1.71E-10 | 6.71E-05 |
| 2 | 11271311 | T | C | 0.865201 | 1.82E-10 | 7.14E-05 |
| 2 | 11271490 | T | C | 0.865201 | 1.82E-10 | 7.14E-05 |
| 2 | 11271529 | T | G | 0.865201 | 1.82E-10 | 7.14E-05 |
| 2 | 11271769 | G | C | 0.865201 | 1.82E-10 | 7.14E-05 |
| 2 | 11274055 | T | C | 0.865201 | 1.82E-10 | 7.14E-05 |
| 2 | 11277631 | A | T | 0.865201 | 1.82E-10 | 7.14E-05 |
| 2 | 8291608 | C | T | 0.874761 | 1.91E-10 | 7.49E-05 |
| 2 | 8292630 | A | G | 0.874761 | 1.91E-10 | 7.49E-05 |
| 2 | 8292940 | C | T | 0.874761 | 1.91E-10 | 7.49E-05 |
| 2 | 11298084 | C | T | 0.868069 | 1.94E-10 | 7.61E-05 |
| 2 | 10639205 | C | T | 0.824092 | 1.94E-10 | 7.62E-05 |
| 2 | 8366545 | A | G | 0.878585 | 1.99E-10 | 7.79E-05 |
| 2 | 8272795 | A | G | 0.876673 | 1.99E-10 | 7.81E-05 |
| 2 | 8281598 | A | G | 0.876673 | 1.99E-10 | 7.81E-05 |
| 2 | 8281862 | A | G | 0.876673 | 1.99E-10 | 7.81E-05 |
| 2 | 8311509 | A | G | 0.876673 | 1.99E-10 | 7.81E-05 |
| 2 | 8166009 | C | T | 0.905354 | 2.05E-10 | 8.04E-05 |
| 2 | 8166010 | A | G | 0.905354 | 2.05E-10 | 8.04E-05 |
| 2 | 8205823 | G | A | 0.871893 | 2.23E-10 | 8.76E-05 |
| 2 | 11097265 | T | G | 0.83652 | 2.25E-10 | 8.81E-05 |
| 2 | 8281924 | G | T | 0.874761 | 2.29E-10 | 8.98E-05 |
| 2 | 8272731 | G | A | 0.873805 | 2.29E-10 | 8.99E-05 |
| 2 | 8893567 | A | G | 0.858509 | 2.40E-10 | 9.42E-05 |
| 2 | 8893697 | T | C | 0.858509 | 2.40E-10 | 9.42E-05 |
| 2 | 11130249 | A | G | 0.834608 | 2.42E-10 | 9.49E-05 |
| 2 | 11130260 | G | T | 0.834608 | 2.42E-10 | 9.49E-05 |
| 2 | 11130385 | A | G | 0.834608 | 2.42E-10 | 9.49E-05 |
| 2 | 11130588 | G | A | 0.834608 | 2.42E-10 | 9.49E-05 |
| 2 | 11130642 | G | A | 0.834608 | 2.42E-10 | 9.49E-05 |
| 2 | 11130689 | T | G | 0.834608 | 2.42E-10 | 9.49E-05 |
| 2 | 11130693 | T | C | 0.834608 | 2.42E-10 | 9.49E-05 |
| 2 | 11135703 | A | G | 0.834608 | 2.42E-10 | 9.49E-05 |
| 2 | 11136497 | T | C | 0.834608 | 2.42E-10 | 9.49E-05 |
| 2 | 11136843 | G | T | 0.834608 | 2.42E-10 | 9.49E-05 |
| 2 | 11136845 | A | T | 0.834608 | 2.42E-10 | 9.49E-05 |
| 2 | 11136978 | C | T | 0.834608 | 2.42E-10 | 9.49E-05 |
| 2 | 11129546 | C | T | 0.839388 | 2.48E-10 | 9.73E-05 |
| 2 | 8895735 | T | G | 0.855641 | 2.49E-10 | 9.78E-05 |
| 2 | 8169879 | T | C | 0.903442 | 2.56E-10 | 0.000100371 |
| 2 | 8287077 | C | T | 0.875717 | 2.58E-10 | 0.000101271 |

| | | | | | | |
|---|---|---|---|---|---|---|
| 2 | 8287103 | C | T | 0.875717 | 2.58E-10 | 0.000101271 |
| 2 | 8287140 | G | A | 0.875717 | 2.58E-10 | 0.000101271 |
| 2 | 8287163 | G | T | 0.875717 | 2.58E-10 | 0.000101271 |
| 2 | 8287306 | A | G | 0.875717 | 2.58E-10 | 0.000101271 |
| 2 | 8287387 | G | A | 0.875717 | 2.58E-10 | 0.000101271 |
| 2 | 8287397 | T | C | 0.875717 | 2.58E-10 | 0.000101271 |
| 2 | 8287407 | T | G | 0.875717 | 2.58E-10 | 0.000101271 |
| 2 | 8287579 | A | G | 0.875717 | 2.58E-10 | 0.000101271 |
| 2 | 8287622 | C | T | 0.875717 | 2.58E-10 | 0.000101271 |
| 2 | 8287635 | C | T | 0.875717 | 2.58E-10 | 0.000101271 |
| 2 | 8287637 | C | A | 0.875717 | 2.58E-10 | 0.000101271 |
| 2 | 8287726 | C | A | 0.875717 | 2.58E-10 | 0.000101271 |
| 2 | 8287753 | C | T | 0.875717 | 2.58E-10 | 0.000101271 |
| 2 | 8287802 | A | G | 0.875717 | 2.58E-10 | 0.000101271 |
| 2 | 8287906 | G | A | 0.875717 | 2.58E-10 | 0.000101271 |
| 2 | 8287920 | C | A | 0.875717 | 2.58E-10 | 0.000101271 |
| 2 | 8339994 | C | A | 0.875717 | 2.58E-10 | 0.000101271 |
| 2 | 8468422 | G | A | 0.875717 | 2.58E-10 | 0.000101271 |
| 2 | 10607390 | G | A | 0.824092 | 2.64E-10 | 0.000103309 |
| 2 | 10607487 | T | C | 0.824092 | 2.64E-10 | 0.000103309 |
| 2 | 10607500 | G | A | 0.824092 | 2.64E-10 | 0.000103309 |
| 2 | 10607702 | A | T | 0.824092 | 2.64E-10 | 0.000103309 |
| 2 | 8366442 | G | T | 0.874761 | 2.76E-10 | 0.000108097 |
| 2 | 8158086 | T | C | 0.902486 | 2.90E-10 | 0.000113822 |
| 2 | 8158143 | A | C | 0.902486 | 2.90E-10 | 0.000113822 |
| 2 | 8158168 | T | C | 0.902486 | 2.90E-10 | 0.000113822 |
| 2 | 11164815 | G | A | 0.835564 | 2.93E-10 | 0.000114661 |
| 2 | 11164868 | T | C | 0.835564 | 2.93E-10 | 0.000114661 |
| 2 | 11164959 | T | C | 0.835564 | 2.93E-10 | 0.000114661 |
| 2 | 11131970 | A | C | 0.833652 | 2.99E-10 | 0.000117281 |
| 2 | 11301798 | G | A | 0.865201 | 3.02E-10 | 0.000118396 |
| 2 | 10638868 | C | T | 0.825048 | 3.12E-10 | 0.000122175 |
| 2 | 10638948 | A | G | 0.825048 | 3.12E-10 | 0.000122175 |
| 2 | 10638988 | T | A | 0.825048 | 3.12E-10 | 0.000122175 |
| 2 | 10639110 | T | C | 0.825048 | 3.12E-10 | 0.000122175 |
| 2 | 10639113 | T | A | 0.825048 | 3.12E-10 | 0.000122175 |
| 2 | 10639128 | A | G | 0.825048 | 3.12E-10 | 0.000122175 |
| 2 | 10639147 | A | G | 0.825048 | 3.12E-10 | 0.000122175 |
| 2 | 8152717 | A | G | 0.900574 | 3.28E-10 | 0.00012861 |
| 2 | 8152780 | C | T | 0.900574 | 3.28E-10 | 0.00012861 |
| 2 | 8152787 | C | T | 0.900574 | 3.28E-10 | 0.00012861 |
| 2 | 8152836 | C | A | 0.900574 | 3.28E-10 | 0.00012861 |
| 2 | 8152923 | A | C | 0.900574 | 3.28E-10 | 0.00012861 |
| 2 | 8152989 | G | A | 0.900574 | 3.28E-10 | 0.00012861 |
| 2 | 8153038 | G | A | 0.900574 | 3.28E-10 | 0.00012861 |
| 2 | 8153148 | A | G | 0.900574 | 3.28E-10 | 0.00012861 |
| 2 | 8153400 | G | A | 0.900574 | 3.28E-10 | 0.00012861 |
| 2 | 10802685 | A | G | 0.838432 | 3.30E-10 | 0.000129262 |
| 2 | 10814805 | T | G | 0.838432 | 3.30E-10 | 0.000129262 |
| 2 | 11126921 | A | C | 0.834608 | 3.33E-10 | 0.00013034 |
| 2 | 11126990 | G | A | 0.834608 | 3.33E-10 | 0.00013034 |
| 2 | 11157504 | T | C | 0.837476 | 3.46E-10 | 0.000135778 |
| 2 | 11157523 | A | G | 0.837476 | 3.46E-10 | 0.000135778 |
| 2 | 11157647 | A | C | 0.837476 | 3.46E-10 | 0.000135778 |
| 2 | 11157816 | G | C | 0.837476 | 3.46E-10 | 0.000135778 |
| 2 | 11157817 | C | T | 0.837476 | 3.46E-10 | 0.000135778 |
| 2 | 11157824 | C | A | 0.837476 | 3.46E-10 | 0.000135778 |
| 2 | 11157892 | C | G | 0.837476 | 3.46E-10 | 0.000135778 |
| 2 | 11157943 | T | A | 0.837476 | 3.46E-10 | 0.000135778 |
| 2 | 11158005 | C | T | 0.837476 | 3.46E-10 | 0.000135778 |
| 2 | 11158182 | A | G | 0.837476 | 3.46E-10 | 0.000135778 |
| 2 | 11158225 | C | T | 0.837476 | 3.46E-10 | 0.000135778 |
| 2 | 11158328 | C | G | 0.837476 | 3.46E-10 | 0.000135778 |
| 2 | 11158350 | A | G | 0.837476 | 3.46E-10 | 0.000135778 |
| 2 | 11158781 | T | C | 0.837476 | 3.46E-10 | 0.000135778 |
| 2 | 11158835 | G | A | 0.837476 | 3.46E-10 | 0.000135778 |
| 2 | 11159023 | A | G | 0.837476 | 3.46E-10 | 0.000135778 |

| | | | | | | |
|---|---|---|---|---|---|---|
| 2 | 11159059 | G | A | 0.837476 | 3.46E-10 | 0.000135778 |
| 2 | 11159182 | G | T | 0.837476 | 3.46E-10 | 0.000135778 |
| 2 | 11159185 | C | T | 0.837476 | 3.46E-10 | 0.000135778 |
| 2 | 11159197 | G | A | 0.837476 | 3.46E-10 | 0.000135778 |
| 2 | 11159454 | T | A | 0.837476 | 3.46E-10 | 0.000135778 |
| 2 | 11159497 | A | T | 0.837476 | 3.46E-10 | 0.000135778 |
| 2 | 11159529 | C | T | 0.837476 | 3.46E-10 | 0.000135778 |
| 2 | 11159553 | T | C | 0.837476 | 3.46E-10 | 0.000135778 |
| 2 | 11161799 | A | G | 0.837476 | 3.46E-10 | 0.000135778 |
| 2 | 11161865 | A | C | 0.837476 | 3.46E-10 | 0.000135778 |
| 2 | 11162134 | T | C | 0.837476 | 3.46E-10 | 0.000135778 |
| 2 | 11162217 | A | C | 0.837476 | 3.46E-10 | 0.000135778 |
| 2 | 11162242 | A | G | 0.837476 | 3.46E-10 | 0.000135778 |
| 2 | 11162269 | G | A | 0.837476 | 3.46E-10 | 0.000135778 |
| 2 | 11162305 | A | G | 0.837476 | 3.46E-10 | 0.000135778 |
| 2 | 11162307 | A | G | 0.837476 | 3.46E-10 | 0.000135778 |
| 2 | 11162346 | T | A | 0.837476 | 3.46E-10 | 0.000135778 |
| 2 | 11163063 | G | C | 0.837476 | 3.46E-10 | 0.000135778 |
| 2 | 11163189 | C | T | 0.837476 | 3.46E-10 | 0.000135778 |
| 2 | 11163218 | C | T | 0.837476 | 3.46E-10 | 0.000135778 |
| 2 | 11163271 | C | T | 0.837476 | 3.46E-10 | 0.000135778 |
| 2 | 11163278 | T | C | 0.837476 | 3.46E-10 | 0.000135778 |
| 2 | 11163981 | T | A | 0.837476 | 3.46E-10 | 0.000135778 |
| 2 | 11164046 | G | T | 0.837476 | 3.46E-10 | 0.000135778 |
| 2 | 11164074 | T | C | 0.837476 | 3.46E-10 | 0.000135778 |
| 2 | 11164151 | T | C | 0.837476 | 3.46E-10 | 0.000135778 |
| 2 | 11164395 | T | A | 0.837476 | 3.46E-10 | 0.000135778 |
| 2 | 11164412 | G | T | 0.837476 | 3.46E-10 | 0.000135778 |
| 2 | 11164422 | G | T | 0.837476 | 3.46E-10 | 0.000135778 |
| 2 | 11164469 | A | G | 0.837476 | 3.46E-10 | 0.000135778 |
| 2 | 11164478 | C | T | 0.837476 | 3.46E-10 | 0.000135778 |
| 2 | 11164486 | T | C | 0.837476 | 3.46E-10 | 0.000135778 |
| 2 | 11164505 | T | G | 0.837476 | 3.46E-10 | 0.000135778 |
| 2 | 11164601 | G | A | 0.837476 | 3.46E-10 | 0.000135778 |
| 2 | 11164770 | C | T | 0.837476 | 3.46E-10 | 0.000135778 |
| 2 | 11164779 | T | C | 0.837476 | 3.46E-10 | 0.000135778 |
| 2 | 11164987 | A | C | 0.837476 | 3.46E-10 | 0.000135778 |
| 2 | 11165103 | C | T | 0.837476 | 3.46E-10 | 0.000135778 |
| 2 | 11165208 | G | C | 0.837476 | 3.46E-10 | 0.000135778 |
| 2 | 11165210 | A | C | 0.837476 | 3.46E-10 | 0.000135778 |
| 2 | 11294187 | C | T | 0.865201 | 3.47E-10 | 0.00013582 |
| 2 | 8664868 | A | G | 0.876673 | 3.53E-10 | 0.00013853 |
| 2 | 8664931 | G | A | 0.876673 | 3.53E-10 | 0.00013853 |
| 2 | 8664934 | T | C | 0.876673 | 3.53E-10 | 0.00013853 |
| 2 | 8664976 | T | C | 0.876673 | 3.53E-10 | 0.00013853 |
| 2 | 11159777 | A | G | 0.83652 | 3.58E-10 | 0.0001405 |
| 2 | 11159802 | G | A | 0.83652 | 3.58E-10 | 0.0001405 |
| 2 | 11159843 | C | T | 0.83652 | 3.58E-10 | 0.0001405 |
| 2 | 11159892 | G | C | 0.83652 | 3.58E-10 | 0.0001405 |
| 2 | 11159990 | A | C | 0.83652 | 3.58E-10 | 0.0001405 |
| 2 | 11160341 | A | G | 0.83652 | 3.58E-10 | 0.0001405 |
| 2 | 11160356 | T | C | 0.83652 | 3.58E-10 | 0.0001405 |
| 2 | 11160441 | T | G | 0.83652 | 3.58E-10 | 0.0001405 |
| 2 | 11160840 | A | G | 0.83652 | 3.58E-10 | 0.0001405 |
| 2 | 11160975 | T | C | 0.83652 | 3.58E-10 | 0.0001405 |
| 2 | 11161083 | G | T | 0.83652 | 3.58E-10 | 0.0001405 |
| 2 | 8278930 | C | T | 0.880497 | 3.69E-10 | 0.00014481 |
| 2 | 11116164 | G | C | 0.832696 | 3.78E-10 | 0.000148187 |
| 2 | 11117140 | C | T | 0.832696 | 3.78E-10 | 0.000148187 |
| 2 | 11117395 | G | A | 0.832696 | 3.78E-10 | 0.000148187 |
| 2 | 11117499 | G | A | 0.832696 | 3.78E-10 | 0.000148187 |
| 2 | 11117731 | A | G | 0.832696 | 3.78E-10 | 0.000148187 |
| 2 | 11119431 | T | G | 0.832696 | 3.78E-10 | 0.000148187 |
| 2 | 11119486 | G | A | 0.832696 | 3.78E-10 | 0.000148187 |
| 2 | 11124031 | T | C | 0.832696 | 3.78E-10 | 0.000148187 |
| 2 | 8365110 | G | A | 0.878585 | 3.82E-10 | 0.000149647 |
| 2 | 8272083 | A | G | 0.874761 | 3.96E-10 | 0.000155206 |

| | | | | | | |
|---|---|---|---|---|---|---|
| 2 | 8272087 | G | A | 0.874761 | 3.96E-10 | 0.000155206 |
| 2 | 8272099 | T | C | 0.874761 | 3.96E-10 | 0.000155206 |
| 2 | 8272121 | G | A | 0.874761 | 3.96E-10 | 0.000155206 |
| 2 | 8273363 | G | A | 0.874761 | 3.96E-10 | 0.000155206 |
| 2 | 8273369 | A | C | 0.874761 | 3.96E-10 | 0.000155206 |
| 2 | 8273384 | G | C | 0.874761 | 3.96E-10 | 0.000155206 |
| 2 | 8275249 | G | A | 0.874761 | 3.96E-10 | 0.000155206 |
| 2 | 8275354 | A | G | 0.874761 | 3.96E-10 | 0.000155206 |
| 2 | 8275467 | T | C | 0.874761 | 3.96E-10 | 0.000155206 |
| 2 | 8276831 | A | T | 0.874761 | 3.96E-10 | 0.000155206 |
| 2 | 8282851 | C | A | 0.874761 | 3.96E-10 | 0.000155206 |
| 2 | 8282852 | T | C | 0.874761 | 3.96E-10 | 0.000155206 |
| 2 | 8282957 | C | A | 0.874761 | 3.96E-10 | 0.000155206 |
| 2 | 8283030 | C | T | 0.874761 | 3.96E-10 | 0.000155206 |
| 2 | 8283038 | A | G | 0.874761 | 3.96E-10 | 0.000155206 |
| 2 | 8283128 | C | T | 0.874761 | 3.96E-10 | 0.000155206 |
| 2 | 8283166 | G | C | 0.874761 | 3.96E-10 | 0.000155206 |
| 2 | 8285254 | C | T | 0.874761 | 3.96E-10 | 0.000155206 |
| 2 | 8285315 | G | A | 0.874761 | 3.96E-10 | 0.000155206 |
| 2 | 8285333 | T | C | 0.874761 | 3.96E-10 | 0.000155206 |
| 2 | 8285352 | G | A | 0.874761 | 3.96E-10 | 0.000155206 |
| 2 | 8285439 | T | G | 0.874761 | 3.96E-10 | 0.000155206 |
| 2 | 8285466 | A | G | 0.874761 | 3.96E-10 | 0.000155206 |
| 2 | 8285554 | C | T | 0.874761 | 3.96E-10 | 0.000155206 |
| 2 | 8286082 | T | A | 0.874761 | 3.96E-10 | 0.000155206 |
| 2 | 8286123 | G | T | 0.874761 | 3.96E-10 | 0.000155206 |
| 2 | 8286219 | C | T | 0.874761 | 3.96E-10 | 0.000155206 |
| 2 | 8286236 | T | C | 0.874761 | 3.96E-10 | 0.000155206 |
| 2 | 8286861 | T | C | 0.874761 | 3.96E-10 | 0.000155206 |
| 2 | 8286869 | C | T | 0.874761 | 3.96E-10 | 0.000155206 |
| 2 | 8286986 | C | A | 0.874761 | 3.96E-10 | 0.000155206 |
| 2 | 8286987 | T | C | 0.874761 | 3.96E-10 | 0.000155206 |
| 2 | 8286991 | C | T | 0.874761 | 3.96E-10 | 0.000155206 |
| 2 | 8287034 | T | C | 0.874761 | 3.96E-10 | 0.000155206 |
| 2 | 8670474 | G | A | 0.874761 | 3.96E-10 | 0.000155206 |
| 2 | 8670590 | A | C | 0.874761 | 3.96E-10 | 0.000155206 |
| 2 | 8670591 | G | T | 0.874761 | 3.96E-10 | 0.000155206 |
| 2 | 8670872 | G | A | 0.874761 | 3.96E-10 | 0.000155206 |
| 2 | 8671352 | C | T | 0.874761 | 3.96E-10 | 0.000155206 |
| 2 | 8672197 | A | G | 0.874761 | 3.96E-10 | 0.000155206 |
| 2 | 8674185 | T | G | 0.874761 | 3.96E-10 | 0.000155206 |
| 2 | 8674684 | A | G | 0.874761 | 3.96E-10 | 0.000155206 |
| 2 | 8675445 | G | A | 0.874761 | 3.96E-10 | 0.000155206 |
| 2 | 8676265 | A | G | 0.874761 | 3.96E-10 | 0.000155206 |
| 2 | 8677633 | A | G | 0.874761 | 3.96E-10 | 0.000155206 |
| 2 | 8678124 | T | G | 0.874761 | 3.96E-10 | 0.000155206 |
| 2 | 8680506 | T | C | 0.874761 | 3.96E-10 | 0.000155206 |
| 2 | 8680514 | A | T | 0.874761 | 3.96E-10 | 0.000155206 |
| 2 | 8680622 | T | A | 0.874761 | 3.96E-10 | 0.000155206 |
| 2 | 8681124 | A | G | 0.874761 | 3.96E-10 | 0.000155206 |
| 2 | 8682786 | C | A | 0.874761 | 3.96E-10 | 0.000155206 |
| 2 | 8683228 | C | T | 0.874761 | 3.96E-10 | 0.000155206 |
| 2 | 8683453 | T | C | 0.874761 | 3.96E-10 | 0.000155206 |
| 2 | 8683967 | T | C | 0.874761 | 3.96E-10 | 0.000155206 |
| 2 | 8684990 | T | C | 0.874761 | 3.96E-10 | 0.000155206 |
| 2 | 8686963 | T | C | 0.874761 | 3.96E-10 | 0.000155206 |
| 2 | 8687026 | T | C | 0.874761 | 3.96E-10 | 0.000155206 |
| 2 | 11321517 | G | A | 0.865201 | 4.04E-10 | 0.000158364 |
| 2 | 11163361 | A | T | 0.83652 | 4.04E-10 | 0.00015848 |
| 2 | 11163667 | A | G | 0.83652 | 4.04E-10 | 0.00015848 |
| 2 | 11232154 | A | C | 0.837476 | 4.14E-10 | 0.000162436 |
| 2 | 11232223 | C | A | 0.837476 | 4.14E-10 | 0.000162436 |
| 2 | 11232233 | G | A | 0.837476 | 4.14E-10 | 0.000162436 |
| 2 | 11232248 | G | A | 0.837476 | 4.14E-10 | 0.000162436 |
| 2 | 11245936 | C | T | 0.837476 | 4.14E-10 | 0.000162436 |
| 2 | 11162496 | A | G | 0.835564 | 4.16E-10 | 0.000162895 |
| 2 | 10455395 | A | G | 0.823136 | 4.25E-10 | 0.000166594 |

| | | | | | | |
|---|---|---|---|---|---|---|
| 2 | 11125632 | A | C | 0.832696 | 4.27E-10 | 0.000167265 |
| 2 | 11125634 | A | T | 0.832696 | 4.27E-10 | 0.000167265 |
| 2 | 11125653 | C | T | 0.832696 | 4.27E-10 | 0.000167265 |
| 2 | 11125801 | A | T | 0.832696 | 4.27E-10 | 0.000167265 |
| 2 | 11125850 | T | C | 0.832696 | 4.27E-10 | 0.000167265 |
| 2 | 11125890 | T | C | 0.832696 | 4.27E-10 | 0.000167265 |
| 2 | 8666274 | T | C | 0.879541 | 4.38E-10 | 0.00017177 |
| 2 | 8666519 | C | T | 0.879541 | 4.38E-10 | 0.00017177 |
| 2 | 8277475 | T | G | 0.878585 | 4.44E-10 | 0.000174138 |
| 2 | 8277840 | A | C | 0.878585 | 4.44E-10 | 0.000174138 |
| 2 | 8278579 | G | C | 0.878585 | 4.44E-10 | 0.000174138 |
| 2 | 8278668 | T | G | 0.878585 | 4.44E-10 | 0.000174138 |
| 2 | 10453413 | T | C | 0.820268 | 4.54E-10 | 0.000178066 |
| 2 | 10453456 | G | A | 0.820268 | 4.54E-10 | 0.000178066 |
| 2 | 8207779 | T | C | 0.877629 | 4.63E-10 | 0.000181367 |
| 2 | 8215397 | C | A | 0.877629 | 4.63E-10 | 0.000181367 |
| 2 | 8215400 | A | G | 0.877629 | 4.63E-10 | 0.000181367 |
| 2 | 8220034 | C | T | 0.877629 | 4.63E-10 | 0.000181367 |
| 2 | 8222532 | C | T | 0.877629 | 4.63E-10 | 0.000181367 |
| 2 | 8222554 | G | A | 0.877629 | 4.63E-10 | 0.000181367 |
| 2 | 8222560 | G | A | 0.877629 | 4.63E-10 | 0.000181367 |
| 2 | 8222571 | G | A | 0.877629 | 4.63E-10 | 0.000181367 |
| 2 | 8222798 | T | A | 0.877629 | 4.63E-10 | 0.000181367 |
| 2 | 8223006 | A | G | 0.877629 | 4.63E-10 | 0.000181367 |
| 2 | 8223459 | G | A | 0.877629 | 4.63E-10 | 0.000181367 |
| 2 | 8223473 | C | A | 0.877629 | 4.63E-10 | 0.000181367 |
| 2 | 8224156 | C | T | 0.877629 | 4.63E-10 | 0.000181367 |
| 2 | 8226900 | G | A | 0.877629 | 4.63E-10 | 0.000181367 |
| 2 | 8226901 | A | G | 0.877629 | 4.63E-10 | 0.000181367 |
| 2 | 8233841 | T | C | 0.877629 | 4.63E-10 | 0.000181367 |
| 2 | 8233895 | T | A | 0.877629 | 4.63E-10 | 0.000181367 |
| 2 | 8233905 | C | A | 0.877629 | 4.63E-10 | 0.000181367 |
| 2 | 8233986 | G | A | 0.877629 | 4.63E-10 | 0.000181367 |
| 2 | 8234083 | A | G | 0.877629 | 4.63E-10 | 0.000181367 |
| 2 | 8234450 | G | T | 0.877629 | 4.63E-10 | 0.000181367 |
| 2 | 8234575 | A | G | 0.877629 | 4.63E-10 | 0.000181367 |
| 2 | 8234579 | G | T | 0.877629 | 4.63E-10 | 0.000181367 |
| 2 | 8234625 | G | T | 0.877629 | 4.63E-10 | 0.000181367 |
| 2 | 8234637 | G | A | 0.877629 | 4.63E-10 | 0.000181367 |
| 2 | 8234823 | C | T | 0.877629 | 4.63E-10 | 0.000181367 |
| 2 | 8234932 | G | A | 0.877629 | 4.63E-10 | 0.000181367 |
| 2 | 8235260 | T | C | 0.877629 | 4.63E-10 | 0.000181367 |
| 2 | 8235329 | A | C | 0.877629 | 4.63E-10 | 0.000181367 |
| 2 | 8235405 | C | T | 0.877629 | 4.63E-10 | 0.000181367 |
| 2 | 8235503 | G | T | 0.877629 | 4.63E-10 | 0.000181367 |
| 2 | 8235533 | C | T | 0.877629 | 4.63E-10 | 0.000181367 |
| 2 | 8235566 | C | T | 0.877629 | 4.63E-10 | 0.000181367 |
| 2 | 8236756 | T | C | 0.877629 | 4.63E-10 | 0.000181367 |
| 2 | 8236843 | G | T | 0.877629 | 4.63E-10 | 0.000181367 |
| 2 | 8237085 | G | T | 0.877629 | 4.63E-10 | 0.000181367 |
| 2 | 8237210 | C | T | 0.877629 | 4.63E-10 | 0.000181367 |
| 2 | 8237983 | G | T | 0.877629 | 4.63E-10 | 0.000181367 |
| 2 | 8238144 | A | T | 0.877629 | 4.63E-10 | 0.000181367 |
| 2 | 8238178 | G | A | 0.877629 | 4.63E-10 | 0.000181367 |
| 2 | 8238191 | C | T | 0.877629 | 4.63E-10 | 0.000181367 |
| 2 | 8238568 | T | A | 0.877629 | 4.63E-10 | 0.000181367 |
| 2 | 8239217 | G | A | 0.877629 | 4.63E-10 | 0.000181367 |
| 2 | 8239719 | G | A | 0.877629 | 4.63E-10 | 0.000181367 |
| 2 | 8240115 | G | T | 0.877629 | 4.63E-10 | 0.000181367 |
| 2 | 8240453 | T | C | 0.877629 | 4.63E-10 | 0.000181367 |
| 2 | 8240619 | A | G | 0.877629 | 4.63E-10 | 0.000181367 |
| 2 | 8241369 | A | G | 0.877629 | 4.63E-10 | 0.000181367 |
| 2 | 8254042 | T | C | 0.877629 | 4.63E-10 | 0.000181367 |
| 2 | 8255015 | G | A | 0.877629 | 4.63E-10 | 0.000181367 |
| 2 | 8256029 | C | T | 0.877629 | 4.63E-10 | 0.000181367 |
| 2 | 8260413 | G | A | 0.877629 | 4.63E-10 | 0.000181367 |
| 2 | 8260451 | A | C | 0.877629 | 4.63E-10 | 0.000181367 |

| | | | | | | |
|---|---|---|---|---|---|---|
| 2 | 8260591 | T | A | 0.877629 | 4.63E-10 | 0.000181367 |
| 2 | 8260667 | A | G | 0.877629 | 4.63E-10 | 0.000181367 |
| 2 | 8260682 | A | G | 0.877629 | 4.63E-10 | 0.000181367 |
| 2 | 8260706 | G | C | 0.877629 | 4.63E-10 | 0.000181367 |
| 2 | 8260873 | G | C | 0.877629 | 4.63E-10 | 0.000181367 |
| 2 | 8261062 | C | A | 0.877629 | 4.63E-10 | 0.000181367 |
| 2 | 8261093 | A | C | 0.877629 | 4.63E-10 | 0.000181367 |
| 2 | 8261236 | T | G | 0.877629 | 4.63E-10 | 0.000181367 |
| 2 | 8261265 | G | C | 0.877629 | 4.63E-10 | 0.000181367 |
| 2 | 8261332 | T | G | 0.877629 | 4.63E-10 | 0.000181367 |
| 2 | 8261712 | G | A | 0.877629 | 4.63E-10 | 0.000181367 |
| 2 | 8261983 | A | G | 0.877629 | 4.63E-10 | 0.000181367 |
| 2 | 8262440 | C | T | 0.877629 | 4.63E-10 | 0.000181367 |
| 2 | 8262684 | C | T | 0.877629 | 4.63E-10 | 0.000181367 |
| 2 | 8262796 | C | T | 0.877629 | 4.63E-10 | 0.000181367 |
| 2 | 8262838 | C | T | 0.877629 | 4.63E-10 | 0.000181367 |
| 2 | 8263088 | C | T | 0.877629 | 4.63E-10 | 0.000181367 |
| 2 | 8267140 | G | A | 0.877629 | 4.63E-10 | 0.000181367 |
| 2 | 8268384 | G | A | 0.877629 | 4.63E-10 | 0.000181367 |
| 2 | 8271629 | T | C | 0.877629 | 4.63E-10 | 0.000181367 |
| 2 | 8271661 | C | A | 0.877629 | 4.63E-10 | 0.000181367 |
| 2 | 8272097 | T | C | 0.877629 | 4.63E-10 | 0.000181367 |
| 2 | 8272144 | A | G | 0.877629 | 4.63E-10 | 0.000181367 |
| 2 | 8272253 | T | C | 0.877629 | 4.63E-10 | 0.000181367 |
| 2 | 8273316 | T | C | 0.877629 | 4.63E-10 | 0.000181367 |
| 2 | 8273948 | C | T | 0.877629 | 4.63E-10 | 0.000181367 |
| 2 | 8274456 | A | G | 0.877629 | 4.63E-10 | 0.000181367 |
| 2 | 8274683 | T | G | 0.877629 | 4.63E-10 | 0.000181367 |
| 2 | 8275038 | A | G | 0.877629 | 4.63E-10 | 0.000181367 |
| 2 | 8275387 | A | G | 0.877629 | 4.63E-10 | 0.000181367 |
| 2 | 8275871 | T | G | 0.877629 | 4.63E-10 | 0.000181367 |
| 2 | 8276152 | A | G | 0.877629 | 4.63E-10 | 0.000181367 |
| 2 | 8276764 | T | G | 0.877629 | 4.63E-10 | 0.000181367 |
| 2 | 8281600 | T | C | 0.877629 | 4.63E-10 | 0.000181367 |
| 2 | 8282356 | T | A | 0.877629 | 4.63E-10 | 0.000181367 |
| 2 | 8282381 | C | T | 0.877629 | 4.63E-10 | 0.000181367 |
| 2 | 8282650 | A | G | 0.877629 | 4.63E-10 | 0.000181367 |
| 2 | 8282724 | G | T | 0.877629 | 4.63E-10 | 0.000181367 |
| 2 | 8283359 | T | C | 0.877629 | 4.63E-10 | 0.000181367 |
| 2 | 8285343 | A | C | 0.877629 | 4.63E-10 | 0.000181367 |
| 2 | 8285518 | T | C | 0.877629 | 4.63E-10 | 0.000181367 |
| 2 | 8285578 | G | A | 0.877629 | 4.63E-10 | 0.000181367 |
| 2 | 8285600 | A | G | 0.877629 | 4.63E-10 | 0.000181367 |
| 2 | 8285604 | A | G | 0.877629 | 4.63E-10 | 0.000181367 |
| 2 | 8285609 | A | G | 0.877629 | 4.63E-10 | 0.000181367 |
| 2 | 8285656 | G | C | 0.877629 | 4.63E-10 | 0.000181367 |
| 2 | 8285698 | A | T | 0.877629 | 4.63E-10 | 0.000181367 |
| 2 | 8286102 | A | C | 0.877629 | 4.63E-10 | 0.000181367 |
| 2 | 8286144 | G | A | 0.877629 | 4.63E-10 | 0.000181367 |
| 2 | 8286206 | A | G | 0.877629 | 4.63E-10 | 0.000181367 |
| 2 | 8286220 | A | G | 0.877629 | 4.63E-10 | 0.000181367 |
| 2 | 8286246 | A | G | 0.877629 | 4.63E-10 | 0.000181367 |
| 2 | 8286382 | A | G | 0.877629 | 4.63E-10 | 0.000181367 |
| 2 | 8286522 | A | G | 0.877629 | 4.63E-10 | 0.000181367 |
| 2 | 8286585 | G | A | 0.877629 | 4.63E-10 | 0.000181367 |
| 2 | 8286645 | G | A | 0.877629 | 4.63E-10 | 0.000181367 |
| 2 | 8286879 | G | A | 0.877629 | 4.63E-10 | 0.000181367 |
| 2 | 8286992 | A | G | 0.877629 | 4.63E-10 | 0.000181367 |
| 2 | 8287049 | T | C | 0.877629 | 4.63E-10 | 0.000181367 |
| 2 | 8287106 | C | T | 0.877629 | 4.63E-10 | 0.000181367 |
| 2 | 8287156 | C | T | 0.877629 | 4.63E-10 | 0.000181367 |
| 2 | 8287222 | T | C | 0.877629 | 4.63E-10 | 0.000181367 |
| 2 | 8287347 | C | A | 0.877629 | 4.63E-10 | 0.000181367 |
| 2 | 8287820 | A | G | 0.877629 | 4.63E-10 | 0.000181367 |
| 2 | 8288168 | C | T | 0.877629 | 4.63E-10 | 0.000181367 |
| 2 | 8288169 | C | T | 0.877629 | 4.63E-10 | 0.000181367 |
| 2 | 8288265 | G | T | 0.877629 | 4.63E-10 | 0.000181367 |

| | | | | | | |
|---|---|---|---|---|---|---|
| 2 | 8289363 | C | T | 0.877629 | 4.63E-10 | 0.000181367 |
| 2 | 8289452 | T | C | 0.877629 | 4.63E-10 | 0.000181367 |
| 2 | 8289596 | T | C | 0.877629 | 4.63E-10 | 0.000181367 |
| 2 | 8290631 | G | A | 0.877629 | 4.63E-10 | 0.000181367 |
| 2 | 8290654 | G | A | 0.877629 | 4.63E-10 | 0.000181367 |
| 2 | 8290824 | C | T | 0.877629 | 4.63E-10 | 0.000181367 |
| 2 | 8291135 | C | T | 0.877629 | 4.63E-10 | 0.000181367 |
| 2 | 8291686 | G | A | 0.877629 | 4.63E-10 | 0.000181367 |
| 2 | 8292354 | T | G | 0.877629 | 4.63E-10 | 0.000181367 |
| 2 | 8292462 | G | T | 0.877629 | 4.63E-10 | 0.000181367 |
| 2 | 8293280 | C | T | 0.877629 | 4.63E-10 | 0.000181367 |
| 2 | 8293282 | T | G | 0.877629 | 4.63E-10 | 0.000181367 |
| 2 | 8293293 | G | A | 0.877629 | 4.63E-10 | 0.000181367 |
| 2 | 8294881 | G | T | 0.877629 | 4.63E-10 | 0.000181367 |
| 2 | 8294923 | T | A | 0.877629 | 4.63E-10 | 0.000181367 |
| 2 | 8310025 | C | T | 0.877629 | 4.63E-10 | 0.000181367 |
| 2 | 8312780 | G | A | 0.877629 | 4.63E-10 | 0.000181367 |
| 2 | 8317705 | C | T | 0.877629 | 4.63E-10 | 0.000181367 |
| 2 | 8320115 | G | A | 0.877629 | 4.63E-10 | 0.000181367 |
| 2 | 8328300 | C | T | 0.877629 | 4.63E-10 | 0.000181367 |
| 2 | 8329242 | C | T | 0.877629 | 4.63E-10 | 0.000181367 |
| 2 | 8337626 | A | T | 0.877629 | 4.63E-10 | 0.000181367 |
| 2 | 8344826 | G | T | 0.877629 | 4.63E-10 | 0.000181367 |
| 2 | 8357851 | A | T | 0.877629 | 4.63E-10 | 0.000181367 |
| 2 | 8374533 | C | T | 0.877629 | 4.63E-10 | 0.000181367 |
| 2 | 8375246 | T | G | 0.877629 | 4.63E-10 | 0.000181367 |
| 2 | 8378059 | C | T | 0.877629 | 4.63E-10 | 0.000181367 |
| 2 | 8378378 | G | A | 0.877629 | 4.63E-10 | 0.000181367 |
| 2 | 8379642 | T | C | 0.877629 | 4.63E-10 | 0.000181367 |
| 2 | 8384496 | G | A | 0.877629 | 4.63E-10 | 0.000181367 |
| 2 | 8385640 | T | G | 0.877629 | 4.63E-10 | 0.000181367 |
| 2 | 8386229 | A | C | 0.877629 | 4.63E-10 | 0.000181367 |
| 2 | 8389300 | C | A | 0.877629 | 4.63E-10 | 0.000181367 |
| 2 | 8389404 | G | A | 0.877629 | 4.63E-10 | 0.000181367 |
| 2 | 8389696 | G | A | 0.877629 | 4.63E-10 | 0.000181367 |
| 2 | 8391863 | C | T | 0.877629 | 4.63E-10 | 0.000181367 |
| 2 | 8395207 | G | C | 0.877629 | 4.63E-10 | 0.000181367 |
| 2 | 8401510 | T | G | 0.877629 | 4.63E-10 | 0.000181367 |
| 2 | 8401514 | C | G | 0.877629 | 4.63E-10 | 0.000181367 |
| 2 | 8413192 | C | T | 0.877629 | 4.63E-10 | 0.000181367 |
| 2 | 8415152 | G | C | 0.877629 | 4.63E-10 | 0.000181367 |
| 2 | 8415209 | A | T | 0.877629 | 4.63E-10 | 0.000181367 |
| 2 | 8415616 | A | G | 0.877629 | 4.63E-10 | 0.000181367 |
| 2 | 8416420 | G | C | 0.877629 | 4.63E-10 | 0.000181367 |
| 2 | 8419669 | T | A | 0.877629 | 4.63E-10 | 0.000181367 |
| 2 | 8422818 | G | T | 0.877629 | 4.63E-10 | 0.000181367 |
| 2 | 8424408 | G | T | 0.877629 | 4.63E-10 | 0.000181367 |
| 2 | 8424496 | A | T | 0.877629 | 4.63E-10 | 0.000181367 |
| 2 | 8426323 | G | A | 0.877629 | 4.63E-10 | 0.000181367 |
| 2 | 8426665 | C | A | 0.877629 | 4.63E-10 | 0.000181367 |
| 2 | 8429774 | A | T | 0.877629 | 4.63E-10 | 0.000181367 |
| 2 | 8432790 | T | C | 0.877629 | 4.63E-10 | 0.000181367 |
| 2 | 8433849 | A | G | 0.877629 | 4.63E-10 | 0.000181367 |
| 2 | 8434068 | A | G | 0.877629 | 4.63E-10 | 0.000181367 |
| 2 | 8436224 | G | A | 0.877629 | 4.63E-10 | 0.000181367 |
| 2 | 8443562 | C | T | 0.877629 | 4.63E-10 | 0.000181367 |
| 2 | 8444232 | G | T | 0.877629 | 4.63E-10 | 0.000181367 |
| 2 | 8445110 | T | C | 0.877629 | 4.63E-10 | 0.000181367 |
| 2 | 8446845 | A | G | 0.877629 | 4.63E-10 | 0.000181367 |
| 2 | 8447829 | C | T | 0.877629 | 4.63E-10 | 0.000181367 |
| 2 | 8449770 | C | T | 0.877629 | 4.63E-10 | 0.000181367 |
| 2 | 8449905 | G | A | 0.877629 | 4.63E-10 | 0.000181367 |
| 2 | 8450963 | T | C | 0.877629 | 4.63E-10 | 0.000181367 |
| 2 | 8451612 | T | A | 0.877629 | 4.63E-10 | 0.000181367 |
| 2 | 8456479 | C | T | 0.877629 | 4.63E-10 | 0.000181367 |
| 2 | 8459402 | C | T | 0.877629 | 4.63E-10 | 0.000181367 |
| 2 | 8460962 | C | T | 0.877629 | 4.63E-10 | 0.000181367 |

| | | | | | | |
|---|---|---|---|---|---|---|
| 2 | 8461101 | T | C | 0.877629 | 4.63E-10 | 0.000181367 |
| 2 | 8467685 | C | T | 0.877629 | 4.63E-10 | 0.000181367 |
| 2 | 8666900 | G | C | 0.877629 | 4.63E-10 | 0.000181367 |
| 2 | 8666912 | A | G | 0.877629 | 4.63E-10 | 0.000181367 |
| 2 | 8667030 | T | C | 0.877629 | 4.63E-10 | 0.000181367 |
| 2 | 8667234 | C | A | 0.877629 | 4.63E-10 | 0.000181367 |
| 2 | 8667387 | T | C | 0.877629 | 4.63E-10 | 0.000181367 |
| 2 | 8667954 | T | G | 0.877629 | 4.63E-10 | 0.000181367 |
| 2 | 8667964 | C | T | 0.877629 | 4.63E-10 | 0.000181367 |
| 2 | 8668007 | T | C | 0.877629 | 4.63E-10 | 0.000181367 |
| 2 | 8668168 | T | G | 0.877629 | 4.63E-10 | 0.000181367 |
| 2 | 8668169 | C | A | 0.877629 | 4.63E-10 | 0.000181367 |
| 2 | 8669143 | C | T | 0.877629 | 4.63E-10 | 0.000181367 |
| 2 | 8669147 | C | T | 0.877629 | 4.63E-10 | 0.000181367 |
| 2 | 8669149 | T | C | 0.877629 | 4.63E-10 | 0.000181367 |
| 2 | 10455503 | G | A | 0.82218 | 4.84E-10 | 0.000189601 |
| 2 | 10455536 | C | T | 0.82218 | 4.84E-10 | 0.000189601 |
| 2 | 10456075 | C | T | 0.82218 | 4.84E-10 | 0.000189601 |
| 2 | 10605186 | T | C | 0.82218 | 4.84E-10 | 0.000189601 |
| 2 | 10610764 | C | T | 0.82218 | 4.84E-10 | 0.000189601 |
| 2 | 10611150 | A | G | 0.82218 | 4.84E-10 | 0.000189601 |
| 2 | 10611186 | T | A | 0.82218 | 4.84E-10 | 0.000189601 |
| 2 | 10611301 | G | A | 0.82218 | 4.84E-10 | 0.000189601 |
| 2 | 10611430 | A | T | 0.82218 | 4.84E-10 | 0.000189601 |
| 2 | 10611490 | C | T | 0.82218 | 4.84E-10 | 0.000189601 |
| 2 | 10611767 | G | A | 0.82218 | 4.84E-10 | 0.000189601 |
| 2 | 10611941 | A | C | 0.82218 | 4.84E-10 | 0.000189601 |
| 2 | 10612202 | T | A | 0.82218 | 4.84E-10 | 0.000189601 |
| 2 | 10612252 | T | C | 0.82218 | 4.84E-10 | 0.000189601 |
| 2 | 10612996 | G | T | 0.82218 | 4.84E-10 | 0.000189601 |
| 2 | 10613226 | C | T | 0.82218 | 4.84E-10 | 0.000189601 |
| 2 | 10613477 | C | T | 0.82218 | 4.84E-10 | 0.000189601 |
| 2 | 10613680 | G | T | 0.82218 | 4.84E-10 | 0.000189601 |
| 2 | 10613693 | A | T | 0.82218 | 4.84E-10 | 0.000189601 |
| 2 | 10613809 | A | G | 0.82218 | 4.84E-10 | 0.000189601 |
| 2 | 10613922 | G | A | 0.82218 | 4.84E-10 | 0.000189601 |
| 2 | 10614053 | T | G | 0.82218 | 4.84E-10 | 0.000189601 |
| 2 | 10614093 | C | T | 0.82218 | 4.84E-10 | 0.000189601 |
| 2 | 10614863 | T | G | 0.82218 | 4.84E-10 | 0.000189601 |
| 2 | 10614995 | G | A | 0.82218 | 4.84E-10 | 0.000189601 |
| 2 | 10615774 | T | C | 0.82218 | 4.84E-10 | 0.000189601 |
| 2 | 10615859 | T | C | 0.82218 | 4.84E-10 | 0.000189601 |
| 2 | 10616583 | C | T | 0.82218 | 4.84E-10 | 0.000189601 |
| 2 | 10616942 | C | T | 0.82218 | 4.84E-10 | 0.000189601 |
| 2 | 10617081 | C | G | 0.82218 | 4.84E-10 | 0.000189601 |
| 2 | 10617941 | G | T | 0.82218 | 4.84E-10 | 0.000189601 |
| 2 | 10618507 | T | C | 0.82218 | 4.84E-10 | 0.000189601 |
| 2 | 10618748 | A | C | 0.82218 | 4.84E-10 | 0.000189601 |
| 2 | 10618756 | T | C | 0.82218 | 4.84E-10 | 0.000189601 |
| 2 | 10618854 | T | C | 0.82218 | 4.84E-10 | 0.000189601 |
| 2 | 10619095 | T | C | 0.82218 | 4.84E-10 | 0.000189601 |
| 2 | 10619407 | A | G | 0.82218 | 4.84E-10 | 0.000189601 |
| 2 | 10619502 | C | T | 0.82218 | 4.84E-10 | 0.000189601 |
| 2 | 10619609 | C | A | 0.82218 | 4.84E-10 | 0.000189601 |
| 2 | 10619622 | C | T | 0.82218 | 4.84E-10 | 0.000189601 |
| 2 | 10619768 | T | G | 0.82218 | 4.84E-10 | 0.000189601 |
| 2 | 10620837 | A | G | 0.82218 | 4.84E-10 | 0.000189601 |
| 2 | 10626284 | C | T | 0.82218 | 4.84E-10 | 0.000189601 |
| 2 | 10626677 | A | G | 0.82218 | 4.84E-10 | 0.000189601 |
| 2 | 10627006 | G | A | 0.82218 | 4.84E-10 | 0.000189601 |
| 2 | 10627619 | A | G | 0.82218 | 4.84E-10 | 0.000189601 |
| 2 | 10627846 | G | C | 0.82218 | 4.84E-10 | 0.000189601 |
| 2 | 10628042 | T | G | 0.82218 | 4.84E-10 | 0.000189601 |
| 2 | 10628933 | A | G | 0.82218 | 4.84E-10 | 0.000189601 |
| 2 | 10628988 | T | A | 0.82218 | 4.84E-10 | 0.000189601 |
| 2 | 10629174 | T | C | 0.82218 | 4.84E-10 | 0.000189601 |
| 2 | 10629772 | T | C | 0.82218 | 4.84E-10 | 0.000189601 |

| | | | | | | |
|---|---|---|---|---|---|---|
| 2 | 10629942 | G | A | 0.82218 | 4.84E-10 | 0.000189601 |
| 2 | 10630240 | A | T | 0.82218 | 4.84E-10 | 0.000189601 |
| 2 | 10630332 | C | A | 0.82218 | 4.84E-10 | 0.000189601 |
| 2 | 10630345 | G | A | 0.82218 | 4.84E-10 | 0.000189601 |
| 2 | 10630447 | T | A | 0.82218 | 4.84E-10 | 0.000189601 |
| 2 | 10630721 | A | G | 0.82218 | 4.84E-10 | 0.000189601 |
| 2 | 10630979 | G | A | 0.82218 | 4.84E-10 | 0.000189601 |
| 2 | 10631121 | A | C | 0.82218 | 4.84E-10 | 0.000189601 |
| 2 | 10631193 | T | A | 0.82218 | 4.84E-10 | 0.000189601 |
| 2 | 10631197 | T | C | 0.82218 | 4.84E-10 | 0.000189601 |
| 2 | 10631248 | T | C | 0.82218 | 4.84E-10 | 0.000189601 |
| 2 | 10631290 | T | A | 0.82218 | 4.84E-10 | 0.000189601 |
| 2 | 10631298 | T | C | 0.82218 | 4.84E-10 | 0.000189601 |
| 2 | 10631302 | G | C | 0.82218 | 4.84E-10 | 0.000189601 |
| 2 | 10631396 | C | T | 0.82218 | 4.84E-10 | 0.000189601 |
| 2 | 10631541 | C | A | 0.82218 | 4.84E-10 | 0.000189601 |
| 2 | 10631570 | A | G | 0.82218 | 4.84E-10 | 0.000189601 |
| 2 | 10631832 | G | A | 0.82218 | 4.84E-10 | 0.000189601 |
| 2 | 10631997 | A | G | 0.82218 | 4.84E-10 | 0.000189601 |
| 2 | 10632038 | C | G | 0.82218 | 4.84E-10 | 0.000189601 |
| 2 | 10632048 | C | T | 0.82218 | 4.84E-10 | 0.000189601 |
| 2 | 10632193 | A | C | 0.82218 | 4.84E-10 | 0.000189601 |
| 2 | 10632325 | A | G | 0.82218 | 4.84E-10 | 0.000189601 |
| 2 | 10632357 | T | C | 0.82218 | 4.84E-10 | 0.000189601 |
| 2 | 10632367 | T | C | 0.82218 | 4.84E-10 | 0.000189601 |
| 2 | 10632677 | T | G | 0.82218 | 4.84E-10 | 0.000189601 |
| 2 | 10632705 | T | G | 0.82218 | 4.84E-10 | 0.000189601 |
| 2 | 10632851 | T | C | 0.82218 | 4.84E-10 | 0.000189601 |
| 2 | 10632899 | C | G | 0.82218 | 4.84E-10 | 0.000189601 |
| 2 | 10632955 | A | G | 0.82218 | 4.84E-10 | 0.000189601 |
| 2 | 10633038 | A | C | 0.82218 | 4.84E-10 | 0.000189601 |
| 2 | 10633069 | A | C | 0.82218 | 4.84E-10 | 0.000189601 |
| 2 | 10633579 | A | G | 0.82218 | 4.84E-10 | 0.000189601 |
| 2 | 10633676 | G | T | 0.82218 | 4.84E-10 | 0.000189601 |
| 2 | 10633689 | T | C | 0.82218 | 4.84E-10 | 0.000189601 |
| 2 | 10633773 | C | T | 0.82218 | 4.84E-10 | 0.000189601 |
| 2 | 10633820 | A | G | 0.82218 | 4.84E-10 | 0.000189601 |
| 2 | 10633832 | G | A | 0.82218 | 4.84E-10 | 0.000189601 |
| 2 | 10633861 | A | G | 0.82218 | 4.84E-10 | 0.000189601 |
| 2 | 10633918 | A | G | 0.82218 | 4.84E-10 | 0.000189601 |
| 2 | 10633982 | A | G | 0.82218 | 4.84E-10 | 0.000189601 |
| 2 | 10634226 | T | C | 0.82218 | 4.84E-10 | 0.000189601 |
| 2 | 10634242 | T | C | 0.82218 | 4.84E-10 | 0.000189601 |
| 2 | 10634360 | T | C | 0.82218 | 4.84E-10 | 0.000189601 |
| 2 | 10634374 | A | G | 0.82218 | 4.84E-10 | 0.000189601 |
| 2 | 10634407 | A | G | 0.82218 | 4.84E-10 | 0.000189601 |
| 2 | 10634413 | T | C | 0.82218 | 4.84E-10 | 0.000189601 |
| 2 | 10634471 | G | A | 0.82218 | 4.84E-10 | 0.000189601 |
| 2 | 10634606 | G | A | 0.82218 | 4.84E-10 | 0.000189601 |
| 2 | 10634655 | G | A | 0.82218 | 4.84E-10 | 0.000189601 |
| 2 | 10638694 | T | C | 0.82218 | 4.84E-10 | 0.000189601 |
| 2 | 10639488 | C | T | 0.82218 | 4.84E-10 | 0.000189601 |
| 2 | 10639632 | A | G | 0.82218 | 4.84E-10 | 0.000189601 |
| 2 | 10639633 | T | A | 0.82218 | 4.84E-10 | 0.000189601 |
| 2 | 10639672 | C | T | 0.82218 | 4.84E-10 | 0.000189601 |
| 2 | 10639753 | C | T | 0.82218 | 4.84E-10 | 0.000189601 |
| 2 | 10639771 | A | G | 0.82218 | 4.84E-10 | 0.000189601 |
| 2 | 10639773 | G | T | 0.82218 | 4.84E-10 | 0.000189601 |
| 2 | 10639848 | A | T | 0.82218 | 4.84E-10 | 0.000189601 |
| 2 | 10640297 | A | C | 0.82218 | 4.84E-10 | 0.000189601 |
| 2 | 10640303 | A | C | 0.82218 | 4.84E-10 | 0.000189601 |
| 2 | 10640309 | A | C | 0.82218 | 4.84E-10 | 0.000189601 |
| 2 | 10640335 | A | G | 0.82218 | 4.84E-10 | 0.000189601 |
| 2 | 10640544 | C | A | 0.82218 | 4.84E-10 | 0.000189601 |
| 2 | 10640655 | A | T | 0.82218 | 4.84E-10 | 0.000189601 |
| 2 | 10640822 | G | A | 0.82218 | 4.84E-10 | 0.000189601 |
| 2 | 10640867 | G | A | 0.82218 | 4.84E-10 | 0.000189601 |

| | | | | | | |
|---|---|---|---|---|---|---|
| 2 | 10640987 | T | C | 0.82218 | 4.84E-10 | 0.000189601 |
| 2 | 10641166 | T | C | 0.82218 | 4.84E-10 | 0.000189601 |
| 2 | 10641260 | G | A | 0.82218 | 4.84E-10 | 0.000189601 |
| 2 | 10641327 | A | C | 0.82218 | 4.84E-10 | 0.000189601 |
| 2 | 10641855 | C | A | 0.82218 | 4.84E-10 | 0.000189601 |
| 2 | 10641892 | A | T | 0.82218 | 4.84E-10 | 0.000189601 |
| 2 | 10641893 | A | T | 0.82218 | 4.84E-10 | 0.000189601 |
| 2 | 10641925 | G | C | 0.82218 | 4.84E-10 | 0.000189601 |
| 2 | 10642208 | C | T | 0.82218 | 4.84E-10 | 0.000189601 |
| 2 | 10642317 | G | T | 0.82218 | 4.84E-10 | 0.000189601 |
| 2 | 10642463 | G | C | 0.82218 | 4.84E-10 | 0.000189601 |
| 2 | 10642558 | G | A | 0.82218 | 4.84E-10 | 0.000189601 |
| 2 | 10642591 | A | G | 0.82218 | 4.84E-10 | 0.000189601 |
| 2 | 10642596 | A | G | 0.82218 | 4.84E-10 | 0.000189601 |
| 2 | 10642604 | A | T | 0.82218 | 4.84E-10 | 0.000189601 |
| 2 | 10642701 | C | T | 0.82218 | 4.84E-10 | 0.000189601 |
| 2 | 10642822 | G | T | 0.82218 | 4.84E-10 | 0.000189601 |
| 2 | 10642836 | T | C | 0.82218 | 4.84E-10 | 0.000189601 |
| 2 | 10642982 | C | T | 0.82218 | 4.84E-10 | 0.000189601 |
| 2 | 10642986 | G | A | 0.82218 | 4.84E-10 | 0.000189601 |
| 2 | 10643549 | C | T | 0.82218 | 4.84E-10 | 0.000189601 |
| 2 | 10643676 | G | C | 0.82218 | 4.84E-10 | 0.000189601 |
| 2 | 10643777 | A | G | 0.82218 | 4.84E-10 | 0.000189601 |
| 2 | 10643871 | A | G | 0.82218 | 4.84E-10 | 0.000189601 |
| 2 | 10643909 | A | C | 0.82218 | 4.84E-10 | 0.000189601 |
| 2 | 10644272 | G | A | 0.82218 | 4.84E-10 | 0.000189601 |
| 2 | 10644363 | T | C | 0.82218 | 4.84E-10 | 0.000189601 |
| 2 | 10644365 | T | C | 0.82218 | 4.84E-10 | 0.000189601 |
| 2 | 10644803 | G | T | 0.82218 | 4.84E-10 | 0.000189601 |
| 2 | 10645083 | T | C | 0.82218 | 4.84E-10 | 0.000189601 |
| 2 | 8158246 | A | G | 0.903442 | 4.86E-10 | 0.000190361 |
| 2 | 8158258 | T | C | 0.903442 | 4.86E-10 | 0.000190361 |
| 2 | 8158276 | G | A | 0.903442 | 4.86E-10 | 0.000190361 |
| 2 | 11260834 | C | A | 0.839388 | 4.96E-10 | 0.000194278 |
| 2 | 11261027 | G | A | 0.839388 | 4.96E-10 | 0.000194278 |
| 2 | 11261061 | G | C | 0.839388 | 4.96E-10 | 0.000194278 |
| 2 | 11261122 | T | C | 0.839388 | 4.96E-10 | 0.000194278 |
| 2 | 11261174 | T | C | 0.839388 | 4.96E-10 | 0.000194278 |
| 2 | 11261399 | C | T | 0.839388 | 4.96E-10 | 0.000194278 |
| 2 | 10473150 | A | G | 0.82696 | 4.99E-10 | 0.000195675 |
| 2 | 8206174 | A | G | 0.873805 | 5.00E-10 | 0.000195817 |
| 2 | 11167719 | A | G | 0.838432 | 5.10E-10 | 0.000199961 |
| 2 | 11161968 | A | G | 0.838432 | 5.16E-10 | 0.000202098 |
| 2 | 8366971 | T | G | 0.870937 | 5.19E-10 | 0.000203301 |
| 2 | 8283861 | G | A | 0.873805 | 5.20E-10 | 0.000203937 |
| 2 | 8283910 | G | A | 0.873805 | 5.20E-10 | 0.000203937 |
| 2 | 10619881 | A | G | 0.837476 | 5.29E-10 | 0.000207492 |
| 2 | 8166051 | G | A | 0.908222 | 5.32E-10 | 0.000208503 |
| 2 | 10621965 | A | T | 0.823136 | 5.36E-10 | 0.00021004 |
| 2 | 11136695 | G | A | 0.835564 | 5.54E-10 | 0.00021727 |
| 2 | 11136708 | A | G | 0.835564 | 5.54E-10 | 0.00021727 |
| 2 | 11136711 | G | A | 0.835564 | 5.54E-10 | 0.00021727 |
| 2 | 10645588 | C | G | 0.823136 | 5.55E-10 | 0.0002174 |
| 2 | 10645589 | A | G | 0.823136 | 5.55E-10 | 0.0002174 |
| 2 | 10645688 | A | G | 0.823136 | 5.55E-10 | 0.0002174 |
| 2 | 11159348 | A | T | 0.835564 | 5.62E-10 | 0.000220276 |
| 2 | 11159608 | A | G | 0.835564 | 5.62E-10 | 0.000220276 |
| 2 | 8295473 | C | T | 0.875717 | 5.75E-10 | 0.000225241 |
| 2 | 8205999 | C | G | 0.867113 | 5.77E-10 | 0.000226235 |
| 2 | 8206093 | T | C | 0.867113 | 5.77E-10 | 0.000226235 |
| 2 | 8283692 | T | C | 0.876673 | 5.88E-10 | 0.00023044 |
| 2 | 8283758 | T | A | 0.876673 | 5.88E-10 | 0.00023044 |
| 2 | 8283759 | A | T | 0.876673 | 5.88E-10 | 0.00023044 |
| 2 | 8283953 | G | A | 0.876673 | 5.88E-10 | 0.00023044 |
| 2 | 8283959 | T | A | 0.876673 | 5.88E-10 | 0.00023044 |
| 2 | 8283971 | C | A | 0.876673 | 5.88E-10 | 0.00023044 |
| 2 | 8284265 | T | C | 0.876673 | 5.88E-10 | 0.00023044 |

| | | | | | | |
|---|---|---|---|---|---|---|
| 2 | 8284416 | C | T | 0.876673 | 5.88E-10 | 0.00023044 |
| 2 | 8284532 | G | T | 0.876673 | 5.88E-10 | 0.00023044 |
| 2 | 11233599 | A | G | 0.837476 | 5.99E-10 | 0.000234901 |
| 2 | 11233606 | A | G | 0.837476 | 5.99E-10 | 0.000234901 |
| 2 | 8408993 | C | T | 0.876673 | 6.05E-10 | 0.00023706 |
| 2 | 8410258 | A | G | 0.876673 | 6.05E-10 | 0.00023706 |
| 2 | 8410989 | T | G | 0.876673 | 6.05E-10 | 0.00023706 |
| 2 | 8411013 | T | G | 0.876673 | 6.05E-10 | 0.00023706 |
| 2 | 8424699 | A | T | 0.876673 | 6.05E-10 | 0.00023706 |
| 2 | 11136913 | T | A | 0.83652 | 6.40E-10 | 0.0002509 |
| 2 | 11137011 | T | C | 0.83652 | 6.40E-10 | 0.0002509 |
| 2 | 11137052 | T | C | 0.83652 | 6.40E-10 | 0.0002509 |
| 2 | 11137068 | A | G | 0.83652 | 6.40E-10 | 0.0002509 |
| 2 | 11137137 | C | T | 0.83652 | 6.40E-10 | 0.0002509 |
| 2 | 11137146 | G | A | 0.83652 | 6.40E-10 | 0.0002509 |
| 2 | 11137148 | G | C | 0.83652 | 6.40E-10 | 0.0002509 |
| 2 | 11137376 | T | G | 0.83652 | 6.40E-10 | 0.0002509 |
| 2 | 11137477 | G | C | 0.83652 | 6.40E-10 | 0.0002509 |
| 2 | 11244812 | T | C | 0.840344 | 6.40E-10 | 0.000250949 |
| 2 | 11244911 | T | C | 0.840344 | 6.40E-10 | 0.000250949 |
| 2 | 11141673 | T | C | 0.828872 | 6.66E-10 | 0.000260899 |
| 2 | 10795656 | A | G | 0.830784 | 6.70E-10 | 0.000262511 |
| 2 | 8366979 | C | T | 0.873805 | 6.84E-10 | 0.000268108 |
| 2 | 11134518 | C | T | 0.834608 | 7.27E-10 | 0.00028506 |
| 2 | 11162681 | G | T | 0.83652 | 7.34E-10 | 0.000287889 |
| 2 | 11162783 | G | A | 0.83652 | 7.34E-10 | 0.000287889 |
| 2 | 11162799 | T | C | 0.83652 | 7.34E-10 | 0.000287889 |
| 2 | 11095390 | C | T | 0.833652 | 7.46E-10 | 0.000292254 |
| 2 | 11095478 | A | G | 0.833652 | 7.46E-10 | 0.000292254 |
| 2 | 11095483 | T | C | 0.833652 | 7.46E-10 | 0.000292254 |
| 2 | 11095492 | C | T | 0.833652 | 7.46E-10 | 0.000292254 |
| 2 | 11095502 | C | T | 0.833652 | 7.46E-10 | 0.000292254 |
| 2 | 11095587 | C | G | 0.833652 | 7.46E-10 | 0.000292254 |
| 2 | 11095591 | A | G | 0.833652 | 7.46E-10 | 0.000292254 |
| 2 | 11095675 | G | A | 0.833652 | 7.46E-10 | 0.000292254 |
| 2 | 11095681 | A | C | 0.833652 | 7.46E-10 | 0.000292254 |
| 2 | 11095899 | C | A | 0.833652 | 7.46E-10 | 0.000292254 |
| 2 | 11096950 | T | C | 0.833652 | 7.46E-10 | 0.000292254 |
| 2 | 11097121 | A | G | 0.833652 | 7.46E-10 | 0.000292254 |
| 2 | 11097188 | G | A | 0.833652 | 7.46E-10 | 0.000292254 |
| 2 | 11098238 | T | G | 0.833652 | 7.46E-10 | 0.000292254 |
| 2 | 11098516 | C | T | 0.833652 | 7.46E-10 | 0.000292254 |
| 2 | 11099215 | G | A | 0.833652 | 7.46E-10 | 0.000292254 |
| 2 | 11099308 | T | C | 0.833652 | 7.46E-10 | 0.000292254 |
| 2 | 11100139 | C | T | 0.833652 | 7.46E-10 | 0.000292254 |
| 2 | 11105164 | T | C | 0.833652 | 7.46E-10 | 0.000292254 |
| 2 | 11105766 | G | A | 0.833652 | 7.46E-10 | 0.000292254 |
| 2 | 11106136 | A | C | 0.833652 | 7.46E-10 | 0.000292254 |
| 2 | 11111370 | A | G | 0.833652 | 7.46E-10 | 0.000292254 |
| 2 | 11111712 | C | G | 0.833652 | 7.46E-10 | 0.000292254 |
| 2 | 11111833 | G | A | 0.833652 | 7.46E-10 | 0.000292254 |
| 2 | 11111890 | T | A | 0.833652 | 7.46E-10 | 0.000292254 |
| 2 | 11112221 | C | T | 0.833652 | 7.46E-10 | 0.000292254 |
| 2 | 11113909 | G | A | 0.833652 | 7.46E-10 | 0.000292254 |
| 2 | 11114714 | T | C | 0.833652 | 7.46E-10 | 0.000292254 |
| 2 | 11115420 | C | T | 0.833652 | 7.46E-10 | 0.000292254 |
| 2 | 11115846 | G | A | 0.833652 | 7.46E-10 | 0.000292254 |
| 2 | 11115945 | T | G | 0.833652 | 7.46E-10 | 0.000292254 |
| 2 | 11116010 | A | G | 0.833652 | 7.46E-10 | 0.000292254 |
| 2 | 11118446 | T | C | 0.833652 | 7.46E-10 | 0.000292254 |
| 2 | 11118480 | A | G | 0.833652 | 7.46E-10 | 0.000292254 |
| 2 | 11118989 | T | C | 0.833652 | 7.46E-10 | 0.000292254 |
| 2 | 11119036 | G | A | 0.833652 | 7.46E-10 | 0.000292254 |
| 2 | 11119259 | T | C | 0.833652 | 7.46E-10 | 0.000292254 |
| 2 | 11119894 | G | A | 0.833652 | 7.46E-10 | 0.000292254 |
| 2 | 11119921 | A | C | 0.833652 | 7.46E-10 | 0.000292254 |
| 2 | 11119931 | C | T | 0.833652 | 7.46E-10 | 0.000292254 |

| | | | | | | |
|---|---|---|---|---|---|---|
| 2 | 11120721 | G | A | 0.833652 | 7.46E-10 | 0.000292254 |
| 2 | 11121188 | C | T | 0.833652 | 7.46E-10 | 0.000292254 |
| 2 | 11122164 | C | T | 0.833652 | 7.46E-10 | 0.000292254 |
| 2 | 11122293 | G | A | 0.833652 | 7.46E-10 | 0.000292254 |
| 2 | 11122448 | G | T | 0.833652 | 7.46E-10 | 0.000292254 |
| 2 | 11122595 | T | A | 0.833652 | 7.46E-10 | 0.000292254 |
| 2 | 11122910 | T | G | 0.833652 | 7.46E-10 | 0.000292254 |
| 2 | 11123156 | G | A | 0.833652 | 7.46E-10 | 0.000292254 |
| 2 | 11123480 | C | T | 0.833652 | 7.46E-10 | 0.000292254 |
| 2 | 11123710 | C | T | 0.833652 | 7.46E-10 | 0.000292254 |
| 2 | 11123760 | T | G | 0.833652 | 7.46E-10 | 0.000292254 |
| 2 | 11124410 | C | T | 0.833652 | 7.46E-10 | 0.000292254 |
| 2 | 11124524 | A | G | 0.833652 | 7.46E-10 | 0.000292254 |
| 2 | 11124914 | A | G | 0.833652 | 7.46E-10 | 0.000292254 |
| 2 | 11127178 | C | A | 0.833652 | 7.46E-10 | 0.000292254 |
| 2 | 11127402 | G | A | 0.833652 | 7.46E-10 | 0.000292254 |
| 2 | 11127404 | C | T | 0.833652 | 7.46E-10 | 0.000292254 |
| 2 | 11127549 | G | T | 0.833652 | 7.46E-10 | 0.000292254 |
| 2 | 11127566 | C | G | 0.833652 | 7.46E-10 | 0.000292254 |
| 2 | 11128389 | C | A | 0.833652 | 7.46E-10 | 0.000292254 |
| 2 | 11128537 | A | C | 0.833652 | 7.46E-10 | 0.000292254 |
| 2 | 11128565 | T | C | 0.833652 | 7.46E-10 | 0.000292254 |
| 2 | 11128582 | C | A | 0.833652 | 7.46E-10 | 0.000292254 |
| 2 | 11128618 | G | T | 0.833652 | 7.46E-10 | 0.000292254 |
| 2 | 11129024 | G | A | 0.833652 | 7.46E-10 | 0.000292254 |
| 2 | 11129072 | C | T | 0.833652 | 7.46E-10 | 0.000292254 |
| 2 | 11129317 | G | A | 0.833652 | 7.46E-10 | 0.000292254 |
| 2 | 11129455 | G | A | 0.833652 | 7.46E-10 | 0.000292254 |
| 2 | 8198268 | C | T | 0.889101 | 7.83E-10 | 0.000307045 |
| 2 | 8675414 | A | G | 0.872849 | 7.99E-10 | 0.000313345 |
| 2 | 8686213 | C | A | 0.872849 | 7.99E-10 | 0.000313345 |
| 2 | 8396366 | C | T | 0.876673 | 7.99E-10 | 0.000313351 |
| 2 | 8396569 | G | A | 0.876673 | 7.99E-10 | 0.000313351 |
| 2 | 8256449 | G | A | 0.873805 | 8.00E-10 | 0.000313473 |
| 2 | 6080771 | C | A | 0.875717 | 6.26E-08 | 0.024538325 |
| 2 | 6081373 | C | T | 0.875717 | 6.26E-08 | 0.024538325 |
| 2 | 6082272 | A | C | 0.877629 | 9.39E-08 | 0.036787732 |
| 2 | 6082531 | A | C | 0.877629 | 9.39E-08 | 0.036787732 |
| 2 | 6082620 | C | T | 0.877629 | 9.39E-08 | 0.036787732 |
| 2 | 6082653 | G | A | 0.877629 | 9.39E-08 | 0.036787732 |
| 2 | 6083064 | A | G | 0.877629 | 9.39E-08 | 0.036787732 |
| 2 | 6083645 | A | G | 0.877629 | 9.39E-08 | 0.036787732 |
| 2 | 6083715 | C | G | 0.877629 | 9.39E-08 | 0.036787732 |
| 2 | 6083794 | G | T | 0.877629 | 9.39E-08 | 0.036787732 |
| 2 | 6083895 | A | T | 0.877629 | 9.39E-08 | 0.036787732 |
| 2 | 6083961 | G | A | 0.877629 | 9.39E-08 | 0.036787732 |
| 2 | 6083976 | C | A | 0.877629 | 9.39E-08 | 0.036787732 |
| 2 | 6084005 | G | A | 0.877629 | 9.39E-08 | 0.036787732 |
| 2 | 6084045 | C | A | 0.877629 | 9.39E-08 | 0.036787732 |
| 2 | 6084122 | A | C | 0.877629 | 9.39E-08 | 0.036787732 |
| 2 | 6084123 | T | C | 0.877629 | 9.39E-08 | 0.036787732 |
| 2 | 6084180 | A | G | 0.875717 | 1.13E-07 | 0.044292112 |
| 2 | 6084193 | T | A | 0.877629 | 9.39E-08 | 0.036787732 |
| 2 | 6084219 | T | A | 0.877629 | 9.39E-08 | 0.036787732 |
| 2 | 6084238 | A | G | 0.877629 | 9.39E-08 | 0.036787732 |
| 2 | 6084480 | C | T | 0.877629 | 9.39E-08 | 0.036787732 |
| 2 | 6084493 | C | A | 0.877629 | 9.39E-08 | 0.036787732 |
| 2 | 6084549 | T | G | 0.877629 | 9.39E-08 | 0.036787732 |
| 2 | 6085088 | C | T | 0.877629 | 9.39E-08 | 0.036787732 |
| 2 | 6085089 | A | G | 0.877629 | 9.39E-08 | 0.036787732 |
| 2 | 6085201 | A | G | 0.877629 | 9.39E-08 | 0.036787732 |
| 2 | 6085482 | G | A | 0.877629 | 9.39E-08 | 0.036787732 |
| 2 | 6085611 | T | C | 0.877629 | 9.39E-08 | 0.036787732 |
| 2 | 6085618 | A | T | 0.877629 | 9.39E-08 | 0.036787732 |
| 2 | 6085619 | C | T | 0.877629 | 9.39E-08 | 0.036787732 |
| 2 | 6085630 | T | G | 0.877629 | 9.39E-08 | 0.036787732 |
| 2 | 6085691 | G | A | 0.877629 | 9.39E-08 | 0.036787732 |

| | | | | | | |
|---|---|---|---|---|---|---|
| 2 | 6085700 | A | T | 0.877629 | 9.39E-08 | 0.036787732 |
| 2 | 6085730 | C | T | 0.877629 | 9.39E-08 | 0.036787732 |
| 2 | 6085856 | A | C | 0.877629 | 9.39E-08 | 0.036787732 |
| 2 | 6085882 | G | T | 0.876673 | 4.54E-08 | 0.017813193 |
| 2 | 6085937 | T | C | 0.876673 | 4.54E-08 | 0.017813193 |
| 2 | 6085952 | C | T | 0.876673 | 4.54E-08 | 0.017813193 |
| 2 | 6085964 | A | G | 0.876673 | 4.54E-08 | 0.017813193 |
| 2 | 6086907 | C | T | 0.877629 | 9.39E-08 | 0.036787732 |
| 2 | 6086928 | T | C | 0.877629 | 9.39E-08 | 0.036787732 |
| 2 | 6086941 | A | G | 0.877629 | 9.39E-08 | 0.036787732 |
| 2 | 6087014 | G | A | 0.877629 | 9.39E-08 | 0.036787732 |
| 2 | 6087018 | A | G | 0.877629 | 9.39E-08 | 0.036787732 |
| 2 | 6087304 | T | C | 0.877629 | 9.39E-08 | 0.036787732 |
| 2 | 6087423 | C | T | 0.877629 | 9.39E-08 | 0.036787732 |
| 2 | 6087499 | C | T | 0.877629 | 9.39E-08 | 0.036787732 |
| 2 | 6087766 | G | C | 0.877629 | 9.39E-08 | 0.036787732 |
| 2 | 6087817 | G | A | 0.877629 | 9.39E-08 | 0.036787732 |
| 2 | 6087947 | A | G | 0.874761 | 1.16E-07 | 0.045561722 |
| 2 | 6088104 | A | G | 0.877629 | 9.39E-08 | 0.036787732 |
| 2 | 6088111 | C | G | 0.877629 | 9.39E-08 | 0.036787732 |
| 2 | 6088305 | T | C | 0.877629 | 9.39E-08 | 0.036787732 |
| 2 | 6088345 | G | A | 0.877629 | 9.39E-08 | 0.036787732 |
| 2 | 6088347 | A | G | 0.877629 | 9.39E-08 | 0.036787732 |
| 2 | 6088453 | T | C | 0.877629 | 9.39E-08 | 0.036787732 |
| 2 | 6088564 | C | T | 0.877629 | 9.39E-08 | 0.036787732 |
| 2 | 6088581 | C | T | 0.877629 | 9.39E-08 | 0.036787732 |
| 2 | 6088693 | A | G | 0.877629 | 9.39E-08 | 0.036787732 |
| 2 | 6088766 | T | A | 0.877629 | 9.39E-08 | 0.036787732 |
| 2 | 6088775 | C | T | 0.877629 | 9.39E-08 | 0.036787732 |
| 2 | 6088778 | A | G | 0.877629 | 9.39E-08 | 0.036787732 |
| 2 | 6088779 | C | T | 0.877629 | 9.39E-08 | 0.036787732 |
| 2 | 6088818 | C | T | 0.877629 | 9.39E-08 | 0.036787732 |
| 2 | 6088945 | C | T | 0.877629 | 9.39E-08 | 0.036787732 |
| 2 | 6088952 | G | A | 0.877629 | 9.39E-08 | 0.036787732 |
| 2 | 6088967 | A | T | 0.877629 | 9.39E-08 | 0.036787732 |
| 2 | 6089076 | C | G | 0.877629 | 9.39E-08 | 0.036787732 |
| 2 | 6089089 | A | G | 0.877629 | 9.39E-08 | 0.036787732 |
| 2 | 6089172 | A | G | 0.877629 | 9.39E-08 | 0.036787732 |
| 2 | 6089228 | G | A | 0.877629 | 9.39E-08 | 0.036787732 |
| 2 | 6089271 | A | T | 0.877629 | 9.39E-08 | 0.036787732 |
| 2 | 6089433 | A | C | 0.877629 | 9.39E-08 | 0.036787732 |
| 2 | 6089463 | G | A | 0.877629 | 9.39E-08 | 0.036787732 |
| 2 | 6089486 | C | T | 0.877629 | 9.39E-08 | 0.036787732 |
| 2 | 6089539 | A | T | 0.877629 | 9.39E-08 | 0.036787732 |
| 2 | 6089622 | C | T | 0.877629 | 9.39E-08 | 0.036787732 |
| 2 | 6089734 | C | A | 0.877629 | 9.39E-08 | 0.036787732 |
| 2 | 6089887 | A | G | 0.877629 | 9.39E-08 | 0.036787732 |
| 2 | 6090335 | T | C | 0.877629 | 9.39E-08 | 0.036787732 |
| 2 | 6090620 | T | A | 0.877629 | 9.39E-08 | 0.036787732 |
| 2 | 6091515 | A | G | 0.877629 | 9.39E-08 | 0.036787732 |
| 2 | 6091776 | G | T | 0.877629 | 9.39E-08 | 0.036787732 |
| 2 | 6092033 | T | C | 0.877629 | 9.39E-08 | 0.036787732 |
| 2 | 6092117 | G | C | 0.877629 | 9.39E-08 | 0.036787732 |
| 2 | 6092376 | T | C | 0.877629 | 9.39E-08 | 0.036787732 |
| 2 | 6092378 | A | G | 0.877629 | 9.39E-08 | 0.036787732 |
| 2 | 6092498 | A | G | 0.877629 | 9.39E-08 | 0.036787732 |
| 2 | 6092521 | A | T | 0.877629 | 9.39E-08 | 0.036787732 |
| 2 | 6092859 | A | G | 0.877629 | 9.39E-08 | 0.036787732 |
| 2 | 6095107 | T | C | 0.877629 | 9.39E-08 | 0.036787732 |
| 2 | 6095162 | A | G | 0.877629 | 9.39E-08 | 0.036787732 |
| 2 | 6096532 | T | C | 0.877629 | 9.39E-08 | 0.036787732 |
| 2 | 6096722 | T | C | 0.877629 | 9.39E-08 | 0.036787732 |
| 2 | 6096766 | T | A | 0.877629 | 9.39E-08 | 0.036787732 |
| 2 | 6096880 | G | A | 0.877629 | 9.39E-08 | 0.036787732 |
| 2 | 6097194 | T | C | 0.877629 | 9.39E-08 | 0.036787732 |
| 2 | 6097267 | A | G | 0.877629 | 9.39E-08 | 0.036787732 |
| 2 | 6097511 | T | C | 0.877629 | 9.39E-08 | 0.036787732 |

| | | | | | | |
|---|---|---|---|---|---|---|
| 2 | 6097685 | G | C | 0.877629 | 9.39E-08 | 0.036787732 |
| 2 | 6097698 | T | C | 0.877629 | 9.39E-08 | 0.036787732 |
| 2 | 6097730 | T | C | 0.877629 | 9.39E-08 | 0.036787732 |
| 2 | 6097732 | A | G | 0.877629 | 9.39E-08 | 0.036787732 |
| 2 | 6098879 | A | G | 0.877629 | 9.39E-08 | 0.036787732 |
| 2 | 6099212 | T | C | 0.877629 | 9.39E-08 | 0.036787732 |
| 2 | 6099676 | T | C | 0.877629 | 9.39E-08 | 0.036787732 |
| 2 | 6099706 | T | C | 0.877629 | 9.39E-08 | 0.036787732 |
| 2 | 6100469 | A | G | 0.877629 | 9.39E-08 | 0.036787732 |
| 2 | 6100470 | G | C | 0.877629 | 9.39E-08 | 0.036787732 |
| 2 | 6100743 | A | G | 0.877629 | 9.39E-08 | 0.036787732 |
| 2 | 6100974 | T | C | 0.877629 | 9.39E-08 | 0.036787732 |
| 2 | 6101065 | C | T | 0.877629 | 9.39E-08 | 0.036787732 |
| 2 | 6101946 | A | C | 0.877629 | 9.39E-08 | 0.036787732 |
| 2 | 6103121 | A | T | 0.877629 | 9.39E-08 | 0.036787732 |
| 2 | 6104398 | A | G | 0.877629 | 9.39E-08 | 0.036787732 |
| 2 | 6104611 | C | A | 0.877629 | 9.39E-08 | 0.036787732 |
| 2 | 6105387 | A | G | 0.877629 | 9.39E-08 | 0.036787732 |
| 2 | 6105417 | G | A | 0.877629 | 9.39E-08 | 0.036787732 |
| 2 | 6105580 | C | T | 0.877629 | 9.39E-08 | 0.036787732 |
| 2 | 6105677 | G | C | 0.877629 | 9.39E-08 | 0.036787732 |
| 2 | 6105826 | T | G | 0.877629 | 9.39E-08 | 0.036787732 |
| 2 | 6106031 | T | C | 0.877629 | 9.39E-08 | 0.036787732 |
| 2 | 6107396 | C | T | 0.877629 | 9.39E-08 | 0.036787732 |
| 2 | 6107583 | C | T | 0.877629 | 9.39E-08 | 0.036787732 |
| 2 | 6107663 | G | A | 0.877629 | 9.39E-08 | 0.036787732 |
| 2 | 6107833 | A | C | 0.877629 | 9.39E-08 | 0.036787732 |
| 2 | 6109698 | C | G | 0.877629 | 9.39E-08 | 0.036787732 |
| 2 | 6109798 | A | G | 0.877629 | 9.39E-08 | 0.036787732 |
| 2 | 6110061 | G | T | 0.877629 | 9.39E-08 | 0.036787732 |
| 2 | 6110141 | C | G | 0.877629 | 9.39E-08 | 0.036787732 |
| 2 | 6110175 | A | G | 0.877629 | 9.39E-08 | 0.036787732 |
| 2 | 6110303 | A | T | 0.877629 | 9.39E-08 | 0.036787732 |
| 2 | 6110707 | C | A | 0.877629 | 9.39E-08 | 0.036787732 |
| 2 | 6110736 | G | A | 0.877629 | 9.39E-08 | 0.036787732 |
| 2 | 6111085 | T | C | 0.877629 | 9.39E-08 | 0.036787732 |
| 2 | 6111239 | A | T | 0.877629 | 9.39E-08 | 0.036787732 |
| 2 | 6111854 | T | C | 0.877629 | 9.39E-08 | 0.036787732 |
| 2 | 6112064 | G | A | 0.877629 | 9.39E-08 | 0.036787732 |
| 2 | 6112158 | A | C | 0.877629 | 9.39E-08 | 0.036787732 |
| 2 | 6112318 | T | C | 0.877629 | 9.39E-08 | 0.036787732 |
| 2 | 6112321 | C | T | 0.877629 | 9.39E-08 | 0.036787732 |
| 2 | 6113533 | C | T | 0.877629 | 9.39E-08 | 0.036787732 |
| 2 | 6115079 | C | T | 0.888145 | 9.41E-08 | 0.036891801 |
| 2 | 6115160 | G | T | 0.888145 | 9.41E-08 | 0.036891801 |
| 2 | 6115272 | T | C | 0.888145 | 9.41E-08 | 0.036891801 |
| 2 | 6115306 | A | G | 0.888145 | 9.41E-08 | 0.036891801 |
| 2 | 6116273 | C | T | 0.877629 | 9.39E-08 | 0.036787732 |
| 2 | 6118491 | A | G | 0.877629 | 9.39E-08 | 0.036787732 |
| 2 | 6166613 | T | C | 0.881453 | 1.15E-07 | 0.045172882 |
| 2 | 6169273 | T | C | 0.881453 | 1.15E-07 | 0.045172882 |
| 2 | 6194084 | G | A | 0.870937 | 1.21E-07 | 0.047405969 |
| 2 | 6194089 | C | T | 0.870937 | 1.21E-07 | 0.047405969 |
| 2 | 6216501 | T | C | 0.877629 | 9.39E-08 | 0.036787732 |
| 2 | 6217056 | A | G | 0.877629 | 9.39E-08 | 0.036787732 |
| 2 | 6224002 | A | C | 0.877629 | 9.39E-08 | 0.036787732 |
| 2 | 6241160 | T | C | 0.877629 | 9.39E-08 | 0.036787732 |
| 2 | 6241280 | A | G | 0.877629 | 9.39E-08 | 0.036787732 |
| 2 | 6241490 | A | G | 0.877629 | 9.39E-08 | 0.036787732 |
| 2 | 6243115 | A | G | 0.877629 | 9.39E-08 | 0.036787732 |
| 2 | 6243127 | G | T | 0.877629 | 9.39E-08 | 0.036787732 |
| 2 | 6243407 | T | C | 0.877629 | 9.39E-08 | 0.036787732 |
| 2 | 6244382 | A | C | 0.877629 | 9.39E-08 | 0.036787732 |
| 2 | 6244650 | G | T | 0.877629 | 9.39E-08 | 0.036787732 |
| 2 | 6244673 | C | T | 0.877629 | 9.39E-08 | 0.036787732 |
| 2 | 6244899 | G | C | 0.877629 | 9.39E-08 | 0.036787732 |
| 2 | 6245037 | A | G | 0.877629 | 9.39E-08 | 0.036787732 |

| | | | | | | |
|---|---|---|---|---|---|---|
| 2 | 6245221 | T | C | 0.877629 | 9.39E-08 | 0.036787732 |
| 2 | 6245250 | G | A | 0.877629 | 9.39E-08 | 0.036787732 |
| 2 | 6245668 | A | G | 0.877629 | 9.39E-08 | 0.036787732 |
| 2 | 6245759 | C | T | 0.877629 | 9.39E-08 | 0.036787732 |
| 2 | 6245862 | G | C | 0.877629 | 9.39E-08 | 0.036787732 |
| 2 | 6245907 | T | C | 0.877629 | 9.39E-08 | 0.036787732 |
| 2 | 6246099 | C | G | 0.877629 | 9.39E-08 | 0.036787732 |
| 2 | 6246415 | A | G | 0.877629 | 9.39E-08 | 0.036787732 |
| 2 | 6246620 | A | G | 0.877629 | 9.39E-08 | 0.036787732 |
| 2 | 6247181 | T | C | 0.877629 | 9.39E-08 | 0.036787732 |
| 2 | 6247528 | T | A | 0.877629 | 9.39E-08 | 0.036787732 |
| 2 | 6247820 | G | A | 0.877629 | 9.39E-08 | 0.036787732 |
| 2 | 6247877 | C | A | 0.877629 | 9.39E-08 | 0.036787732 |
| 2 | 6247909 | C | T | 0.877629 | 9.39E-08 | 0.036787732 |
| 2 | 6248166 | G | A | 0.877629 | 9.39E-08 | 0.036787732 |
| 2 | 6248237 | T | C | 0.877629 | 9.39E-08 | 0.036787732 |
| 2 | 6248243 | T | C | 0.877629 | 9.39E-08 | 0.036787732 |
| 2 | 6248253 | C | T | 0.877629 | 9.39E-08 | 0.036787732 |
| 2 | 6248500 | T | C | 0.877629 | 9.39E-08 | 0.036787732 |
| 2 | 6248837 | C | G | 0.877629 | 9.39E-08 | 0.036787732 |
| 2 | 6249235 | T | C | 0.877629 | 9.39E-08 | 0.036787732 |
| 2 | 6249324 | A | G | 0.877629 | 9.39E-08 | 0.036787732 |
| 2 | 8198892 | G | T | 0.869981 | 1.04E-09 | 0.000408525 |
| 2 | 8199035 | A | C | 0.872849 | 1.13E-09 | 0.000442894 |
| 2 | 8199039 | C | T | 0.872849 | 1.13E-09 | 0.000442894 |
| 2 | 8199201 | T | G | 0.872849 | 1.13E-09 | 0.000442894 |
| 2 | 8199675 | G | T | 0.872849 | 1.13E-09 | 0.000442894 |
| 2 | 8199878 | T | A | 0.872849 | 1.13E-09 | 0.000442894 |
| 2 | 8200049 | T | G | 0.867113 | 6.10E-09 | 0.002390125 |
| 2 | 8200578 | T | C | 0.872849 | 1.13E-09 | 0.000442894 |
| 2 | 8201365 | A | C | 0.872849 | 1.13E-09 | 0.000442894 |
| 2 | 8201400 | G | A | 0.872849 | 1.13E-09 | 0.000442894 |
| 2 | 8201757 | A | G | 0.866157 | 4.06E-09 | 0.001590595 |
| 2 | 8202570 | A | G | 0.872849 | 1.13E-09 | 0.000442894 |
| 2 | 8202778 | C | T | 0.867113 | 6.10E-09 | 0.002390125 |
| 2 | 8202840 | C | T | 0.867113 | 6.10E-09 | 0.002390125 |
| 2 | 8202897 | C | A | 0.872849 | 1.13E-09 | 0.000442894 |
| 2 | 8202907 | C | T | 0.872849 | 1.13E-09 | 0.000442894 |
| 2 | 8202964 | C | T | 0.872849 | 1.13E-09 | 0.000442894 |
| 2 | 8203266 | C | T | 0.872849 | 1.13E-09 | 0.000442894 |
| 2 | 8203331 | C | T | 0.872849 | 1.13E-09 | 0.000442894 |
| 2 | 8203394 | C | A | 0.872849 | 1.13E-09 | 0.000442894 |
| 2 | 8205569 | C | T | 0.872849 | 1.13E-09 | 0.000442894 |
| 2 | 8205665 | T | C | 0.866157 | 1.33E-09 | 0.000520525 |
| 2 | 8206869 | G | C | 0.869981 | 7.90E-09 | 0.003097739 |
| 2 | 8206986 | A | G | 0.875717 | 1.43E-09 | 0.000559977 |
| 2 | 8207701 | C | G | 0.869981 | 7.90E-09 | 0.003097739 |
| 2 | 8208144 | C | T | 0.871893 | 2.66E-09 | 0.001044036 |
| 2 | 8208179 | A | G | 0.871893 | 2.66E-09 | 0.001044036 |
| 2 | 8212353 | G | C | 0.871893 | 2.66E-09 | 0.001044036 |
| 2 | 8215349 | A | G | 0.871893 | 2.66E-09 | 0.001044036 |
| 2 | 8215459 | G | T | 0.871893 | 2.66E-09 | 0.001044036 |
| 2 | 8215803 | G | C | 0.871893 | 2.66E-09 | 0.001044036 |
| 2 | 8215821 | T | G | 0.871893 | 2.66E-09 | 0.001044036 |
| 2 | 8216070 | C | T | 0.871893 | 2.66E-09 | 0.001044036 |
| 2 | 8216212 | C | T | 0.871893 | 2.66E-09 | 0.001044036 |
| 2 | 8216494 | G | A | 0.871893 | 2.66E-09 | 0.001044036 |
| 2 | 8216573 | C | G | 0.871893 | 2.66E-09 | 0.001044036 |
| 2 | 8216833 | A | G | 0.871893 | 2.66E-09 | 0.001044036 |
| 2 | 8216948 | C | T | 0.871893 | 2.66E-09 | 0.001044036 |
| 2 | 8217040 | T | C | 0.871893 | 2.66E-09 | 0.001044036 |
| 2 | 8217064 | A | G | 0.871893 | 2.66E-09 | 0.001044036 |
| 2 | 8217429 | T | C | 0.868069 | 1.63E-09 | 0.000637486 |
| 2 | 8217456 | G | A | 0.868069 | 1.63E-09 | 0.000637486 |
| 2 | 8217476 | T | G | 0.868069 | 1.63E-09 | 0.000637486 |
| 2 | 8217510 | C | T | 0.868069 | 1.63E-09 | 0.000637486 |
| 2 | 8217817 | C | A | 0.868069 | 1.63E-09 | 0.000637486 |

| | | | | | | |
|---|---|---|---|---|---|---|
| 2 | 8219339 | A | C | 0.871893 | 2.66E-09 | 0.001044036 |
| 2 | 8219768 | T | G | 0.871893 | 2.66E-09 | 0.001044036 |
| 2 | 8220015 | C | T | 0.871893 | 2.66E-09 | 0.001044036 |
| 2 | 8220186 | G | A | 0.871893 | 2.66E-09 | 0.001044036 |
| 2 | 8220199 | G | A | 0.871893 | 2.66E-09 | 0.001044036 |
| 2 | 8221333 | A | T | 0.871893 | 2.66E-09 | 0.001044036 |
| 2 | 8221709 | C | T | 0.871893 | 2.66E-09 | 0.001044036 |
| 2 | 8224044 | G | A | 0.871893 | 2.66E-09 | 0.001044036 |
| 2 | 8224138 | G | A | 0.871893 | 2.66E-09 | 0.001044036 |
| 2 | 8224383 | C | T | 0.871893 | 2.66E-09 | 0.001044036 |
| 2 | 8224482 | T | C | 0.871893 | 2.66E-09 | 0.001044036 |
| 2 | 8224523 | C | T | 0.871893 | 2.66E-09 | 0.001044036 |
| 2 | 8224703 | G | T | 0.871893 | 2.66E-09 | 0.001044036 |
| 2 | 8224865 | C | T | 0.871893 | 2.66E-09 | 0.001044036 |
| 2 | 8224949 | A | G | 0.871893 | 2.66E-09 | 0.001044036 |
| 2 | 8224965 | C | T | 0.871893 | 2.66E-09 | 0.001044036 |
| 2 | 8225304 | C | T | 0.873805 | 1.23E-09 | 0.000482511 |
| 2 | 8225338 | C | T | 0.873805 | 1.23E-09 | 0.000482511 |
| 2 | 8225418 | C | T | 0.873805 | 1.23E-09 | 0.000482511 |
| 2 | 8225427 | G | A | 0.873805 | 1.23E-09 | 0.000482511 |
| 2 | 8225794 | C | A | 0.873805 | 1.23E-09 | 0.000482511 |
| 2 | 8225856 | C | T | 0.873805 | 1.23E-09 | 0.000482511 |
| 2 | 8225918 | G | A | 0.873805 | 1.23E-09 | 0.000482511 |
| 2 | 8225978 | G | A | 0.871893 | 2.66E-09 | 0.001044036 |
| 2 | 8225994 | G | A | 0.871893 | 2.66E-09 | 0.001044036 |
| 2 | 8226090 | C | T | 0.871893 | 2.66E-09 | 0.001044036 |
| 2 | 8226130 | T | G | 0.871893 | 2.66E-09 | 0.001044036 |
| 2 | 8227204 | C | A | 0.871893 | 2.66E-09 | 0.001044036 |
| 2 | 8227215 | C | T | 0.871893 | 2.66E-09 | 0.001044036 |
| 2 | 8227316 | C | T | 0.871893 | 2.66E-09 | 0.001044036 |
| 2 | 8227416 | T | C | 0.871893 | 2.66E-09 | 0.001044036 |
| 2 | 8227518 | C | T | 0.871893 | 2.66E-09 | 0.001044036 |
| 2 | 8227555 | C | T | 0.871893 | 2.66E-09 | 0.001044036 |
| 2 | 8227568 | T | C | 0.871893 | 2.66E-09 | 0.001044036 |
| 2 | 8227696 | A | G | 0.871893 | 2.66E-09 | 0.001044036 |
| 2 | 8227890 | G | A | 0.871893 | 2.66E-09 | 0.001044036 |
| 2 | 8228052 | C | T | 0.871893 | 2.66E-09 | 0.001044036 |
| 2 | 8228388 | T | G | 0.871893 | 2.66E-09 | 0.001044036 |
| 2 | 8228427 | A | T | 0.871893 | 2.66E-09 | 0.001044036 |
| 2 | 8228428 | A | C | 0.871893 | 2.66E-09 | 0.001044036 |
| 2 | 8228455 | T | G | 0.871893 | 2.66E-09 | 0.001044036 |
| 2 | 8228493 | G | A | 0.871893 | 2.66E-09 | 0.001044036 |
| 2 | 8228503 | T | C | 0.871893 | 2.66E-09 | 0.001044036 |
| 2 | 8228560 | C | T | 0.871893 | 2.66E-09 | 0.001044036 |
| 2 | 8228580 | G | A | 0.871893 | 2.66E-09 | 0.001044036 |
| 2 | 8228791 | C | T | 0.871893 | 2.66E-09 | 0.001044036 |
| 2 | 8228915 | T | C | 0.871893 | 2.66E-09 | 0.001044036 |
| 2 | 8228959 | A | G | 0.871893 | 2.66E-09 | 0.001044036 |
| 2 | 8229035 | A | G | 0.871893 | 2.66E-09 | 0.001044036 |
| 2 | 8229356 | C | T | 0.871893 | 2.66E-09 | 0.001044036 |
| 2 | 8230005 | C | T | 0.871893 | 2.66E-09 | 0.001044036 |
| 2 | 8230150 | C | T | 0.871893 | 2.66E-09 | 0.001044036 |
| 2 | 8230160 | G | A | 0.871893 | 2.66E-09 | 0.001044036 |
| 2 | 8230226 | T | C | 0.871893 | 2.66E-09 | 0.001044036 |
| 2 | 8230288 | G | A | 0.871893 | 2.66E-09 | 0.001044036 |
| 2 | 8230660 | T | C | 0.871893 | 2.66E-09 | 0.001044036 |
| 2 | 8231099 | T | A | 0.871893 | 2.66E-09 | 0.001044036 |
| 2 | 8231344 | G | A | 0.871893 | 2.66E-09 | 0.001044036 |
| 2 | 8231404 | A | G | 0.871893 | 2.66E-09 | 0.001044036 |
| 2 | 8231464 | C | G | 0.871893 | 2.66E-09 | 0.001044036 |
| 2 | 8231591 | T | A | 0.871893 | 2.66E-09 | 0.001044036 |
| 2 | 8231592 | C | A | 0.871893 | 2.66E-09 | 0.001044036 |
| 2 | 8231734 | T | C | 0.871893 | 2.66E-09 | 0.001044036 |
| 2 | 8231774 | T | C | 0.871893 | 2.66E-09 | 0.001044036 |
| 2 | 8231840 | C | T | 0.871893 | 2.66E-09 | 0.001044036 |
| 2 | 8231841 | G | A | 0.871893 | 2.66E-09 | 0.001044036 |
| 2 | 8231880 | T | C | 0.871893 | 2.66E-09 | 0.001044036 |

| | | | | | | |
|---|---|---|---|---|---|---|
| 2 | 8231897 | C | G | 0.871893 | 2.66E-09 | 0.001044036 |
| 2 | 8232606 | C | T | 0.871893 | 2.66E-09 | 0.001044036 |
| 2 | 8232774 | C | T | 0.871893 | 2.66E-09 | 0.001044036 |
| 2 | 8233486 | T | C | 0.871893 | 2.66E-09 | 0.001044036 |
| 2 | 8233724 | G | A | 0.871893 | 2.66E-09 | 0.001044036 |
| 2 | 8235593 | T | C | 0.871893 | 2.66E-09 | 0.001044036 |
| 2 | 8249520 | G | A | 0.872849 | 5.45E-09 | 0.002135975 |
| 2 | 8249549 | C | G | 0.872849 | 5.45E-09 | 0.002135975 |
| 2 | 8250687 | G | T | 0.872849 | 5.45E-09 | 0.002135975 |
| 2 | 8250809 | T | C | 0.872849 | 5.45E-09 | 0.002135975 |
| 2 | 8251036 | G | T | 0.872849 | 5.45E-09 | 0.002135975 |
| 2 | 8252669 | A | G | 0.871893 | 2.66E-09 | 0.001044036 |
| 2 | 8252682 | A | G | 0.871893 | 2.66E-09 | 0.001044036 |
| 2 | 8252863 | T | C | 0.871893 | 2.66E-09 | 0.001044036 |
| 2 | 8252868 | A | T | 0.871893 | 2.66E-09 | 0.001044036 |
| 2 | 8252886 | G | A | 0.871893 | 2.66E-09 | 0.001044036 |
| 2 | 8252942 | T | A | 0.871893 | 2.66E-09 | 0.001044036 |
| 2 | 8253034 | C | A | 0.871893 | 2.66E-09 | 0.001044036 |
| 2 | 8253395 | T | C | 0.871893 | 2.66E-09 | 0.001044036 |
| 2 | 8253570 | T | C | 0.871893 | 2.66E-09 | 0.001044036 |
| 2 | 8253914 | T | C | 0.871893 | 2.66E-09 | 0.001044036 |
| 2 | 8254376 | A | C | 0.871893 | 2.66E-09 | 0.001044036 |
| 2 | 8255179 | T | C | 0.871893 | 2.66E-09 | 0.001044036 |
| 2 | 8255201 | G | C | 0.871893 | 2.66E-09 | 0.001044036 |
| 2 | 8261703 | A | G | 0.871893 | 2.66E-09 | 0.001044036 |
| 2 | 8262264 | C | T | 0.871893 | 2.66E-09 | 0.001044036 |
| 2 | 8262272 | C | T | 0.871893 | 2.66E-09 | 0.001044036 |
| 2 | 8264009 | A | G | 0.871893 | 2.66E-09 | 0.001044036 |
| 2 | 8264238 | A | C | 0.871893 | 2.66E-09 | 0.001044036 |
| 2 | 8264383 | G | A | 0.871893 | 2.66E-09 | 0.001044036 |
| 2 | 8264835 | G | A | 0.871893 | 2.66E-09 | 0.001044036 |
| 2 | 8264895 | A | G | 0.871893 | 2.66E-09 | 0.001044036 |
| 2 | 8265003 | G | A | 0.871893 | 2.66E-09 | 0.001044036 |
| 2 | 8265359 | T | C | 0.871893 | 2.66E-09 | 0.001044036 |
| 2 | 8265501 | C | T | 0.871893 | 2.66E-09 | 0.001044036 |
| 2 | 8265567 | C | T | 0.871893 | 2.66E-09 | 0.001044036 |
| 2 | 8265575 | C | T | 0.871893 | 2.66E-09 | 0.001044036 |
| 2 | 8265606 | A | T | 0.871893 | 2.66E-09 | 0.001044036 |
| 2 | 8265611 | G | A | 0.871893 | 2.66E-09 | 0.001044036 |
| 2 | 8265613 | A | G | 0.871893 | 2.66E-09 | 0.001044036 |
| 2 | 8265648 | G | T | 0.871893 | 2.66E-09 | 0.001044036 |
| 2 | 8266047 | C | T | 0.871893 | 2.66E-09 | 0.001044036 |
| 2 | 8266193 | T | C | 0.871893 | 2.66E-09 | 0.001044036 |
| 2 | 8266263 | G | A | 0.871893 | 2.66E-09 | 0.001044036 |
| 2 | 8266387 | T | A | 0.871893 | 2.66E-09 | 0.001044036 |
| 2 | 8266396 | A | G | 0.871893 | 2.66E-09 | 0.001044036 |
| 2 | 8266418 | A | G | 0.871893 | 2.66E-09 | 0.001044036 |
| 2 | 8266453 | T | A | 0.871893 | 2.66E-09 | 0.001044036 |
| 2 | 8266535 | A | G | 0.871893 | 2.66E-09 | 0.001044036 |
| 2 | 8267024 | G | A | 0.871893 | 2.66E-09 | 0.001044036 |
| 2 | 8268180 | C | T | 0.871893 | 2.66E-09 | 0.001044036 |
| 2 | 8268192 | G | A | 0.871893 | 2.66E-09 | 0.001044036 |
| 2 | 8268276 | A | C | 0.871893 | 2.66E-09 | 0.001044036 |
| 2 | 8268449 | C | A | 0.871893 | 2.66E-09 | 0.001044036 |
| 2 | 8269293 | A | G | 0.871893 | 2.66E-09 | 0.001044036 |
| 2 | 8269341 | C | T | 0.871893 | 2.66E-09 | 0.001044036 |
| 2 | 8269385 | G | A | 0.871893 | 2.66E-09 | 0.001044036 |
| 2 | 8269422 | C | T | 0.870937 | 3.69E-09 | 0.00144489 |
| 2 | 8269423 | A | G | 0.871893 | 2.66E-09 | 0.001044036 |
| 2 | 8269427 | G | A | 0.870937 | 3.69E-09 | 0.00144489 |
| 2 | 8269536 | T | C | 0.870937 | 3.69E-09 | 0.00144489 |
| 2 | 8269692 | A | C | 0.870937 | 7.71E-09 | 0.003022468 |
| 2 | 8269953 | G | A | 0.869025 | 1.95E-08 | 0.007640435 |
| 2 | 8270029 | C | T | 0.869025 | 1.95E-08 | 0.007640435 |
| 2 | 8270061 | G | T | 0.869981 | 1.20E-08 | 0.004687288 |
| 2 | 8270090 | C | G | 0.869981 | 1.20E-08 | 0.004687288 |
| 2 | 8270100 | G | A | 0.869981 | 1.20E-08 | 0.004687288 |

| | | | | | | |
|---|---|---|---|---|---|---|
| 2 | 8270238 | C | T | 0.869981 | 1.20E-08 | 0.004687288 |
| 2 | 8270266 | G | A | 0.869981 | 1.20E-08 | 0.004687288 |
| 2 | 8270270 | T | A | 0.870937 | 7.71E-09 | 0.003022468 |
| 2 | 8270310 | C | A | 0.872849 | 1.10E-08 | 0.00432424 |
| 2 | 8270345 | C | A | 0.872849 | 1.10E-08 | 0.00432424 |
| 2 | 8270396 | T | C | 0.872849 | 1.10E-08 | 0.00432424 |
| 2 | 8270398 | T | C | 0.872849 | 1.10E-08 | 0.00432424 |
| 2 | 8270399 | G | A | 0.872849 | 1.10E-08 | 0.00432424 |
| 2 | 8270487 | T | A | 0.872849 | 1.10E-08 | 0.00432424 |
| 2 | 8270512 | A | G | 0.872849 | 1.10E-08 | 0.00432424 |
| 2 | 8270534 | G | T | 0.872849 | 1.10E-08 | 0.00432424 |
| 2 | 8270599 | T | G | 0.871893 | 6.42E-09 | 0.002518371 |
| 2 | 8270702 | T | C | 0.870937 | 7.71E-09 | 0.003022468 |
| 2 | 8270743 | T | C | 0.869981 | 1.54E-08 | 0.006027611 |
| 2 | 8270805 | G | C | 0.869981 | 1.54E-08 | 0.006027611 |
| 2 | 8270822 | G | T | 0.870937 | 5.72E-09 | 0.002243729 |
| 2 | 8270952 | C | A | 0.872849 | 5.45E-09 | 0.002135975 |
| 2 | 8270953 | A | G | 0.872849 | 5.45E-09 | 0.002135975 |
| 2 | 8270954 | C | A | 0.872849 | 5.45E-09 | 0.002135975 |
| 2 | 8270982 | A | G | 0.872849 | 5.45E-09 | 0.002135975 |
| 2 | 8271009 | G | A | 0.872849 | 5.45E-09 | 0.002135975 |
| 2 | 8271017 | C | T | 0.869981 | 4.50E-09 | 0.001763998 |
| 2 | 8271168 | T | C | 0.872849 | 5.45E-09 | 0.002135975 |
| 2 | 8271185 | T | C | 0.872849 | 5.45E-09 | 0.002135975 |
| 2 | 8271596 | C | T | 0.872849 | 5.45E-09 | 0.002135975 |
| 2 | 8271813 | A | T | 0.867113 | 9.76E-08 | 0.03823965 |
| 2 | 8278135 | C | T | 0.872849 | 1.54E-09 | 0.000602173 |
| 2 | 8278298 | A | G | 0.872849 | 1.54E-09 | 0.000602173 |
| 2 | 8282022 | A | T | 0.883365 | 5.70E-09 | 0.002236047 |
| 2 | 8282030 | T | A | 0.885277 | 1.03E-08 | 0.004023085 |
| 2 | 8282035 | T | A | 0.883365 | 5.70E-09 | 0.002236047 |
| 2 | 8282036 | T | A | 0.883365 | 5.70E-09 | 0.002236047 |
| 2 | 8282840 | T | C | 0.875717 | 1.02E-09 | 0.000401579 |
| 2 | 8285723 | C | T | 0.873805 | 8.42E-10 | 0.000329914 |
| 2 | 8285777 | A | G | 0.876673 | 1.03E-09 | 0.000403124 |
| 2 | 8285923 | T | G | 0.873805 | 8.42E-10 | 0.000329914 |
| 2 | 8286062 | G | T | 0.873805 | 8.42E-10 | 0.000329914 |
| 2 | 8295076 | A | G | 0.878585 | 1.41E-09 | 0.000552584 |
| 2 | 8295082 | A | G | 0.878585 | 1.41E-09 | 0.000552584 |
| 2 | 8295166 | C | G | 0.878585 | 1.41E-09 | 0.000552584 |
| 2 | 8366548 | G | T | 0.872849 | 8.60E-10 | 0.000337047 |
| 2 | 8411937 | T | C | 0.876673 | 1.39E-09 | 0.000544094 |
| 2 | 8411938 | T | A | 0.876673 | 1.39E-09 | 0.000544094 |
| 2 | 8411940 | A | C | 0.876673 | 1.39E-09 | 0.000544094 |
| 2 | 8461224 | G | A | 0.873805 | 6.16E-09 | 0.00241547 |
| 2 | 8501036 | C | T | 0.885277 | 1.21E-07 | 0.047457318 |
| 2 | 8543871 | A | C | 0.909178 | 1.26E-08 | 0.004930039 |
| 2 | 8583946 | G | A | 0.876673 | 1.03E-09 | 0.000402238 |
| 2 | 8595072 | C | T | 0.869981 | 7.78E-09 | 0.003051521 |
| 2 | 8595998 | C | G | 0.869981 | 7.78E-09 | 0.003051521 |
| 2 | 8596822 | A | C | 0.869025 | 1.63E-08 | 0.006402536 |
| 2 | 8641012 | G | A | 0.869981 | 4.50E-09 | 0.001763998 |
| 2 | 8642356 | G | A | 0.869981 | 4.50E-09 | 0.001763998 |
| 2 | 8663773 | C | T | 0.875717 | 1.29E-09 | 0.00050597 |
| 2 | 8663789 | G | A | 0.875717 | 1.29E-09 | 0.00050597 |
| 2 | 8664693 | C | T | 0.875717 | 1.29E-09 | 0.00050597 |
| 2 | 8665135 | C | T | 0.875717 | 1.29E-09 | 0.00050597 |
| 2 | 8665218 | A | G | 0.875717 | 1.29E-09 | 0.00050597 |
| 2 | 8665277 | G | A | 0.875717 | 1.29E-09 | 0.00050597 |
| 2 | 8665347 | T | C | 0.875717 | 1.29E-09 | 0.00050597 |
| 2 | 8665474 | G | A | 0.875717 | 1.29E-09 | 0.00050597 |
| 2 | 8665945 | C | T | 0.870937 | 1.47E-09 | 0.000574417 |
| 2 | 8667036 | C | G | 0.869025 | 2.20E-09 | 0.000861457 |
| 2 | 8667258 | G | A | 0.869025 | 2.20E-09 | 0.000861457 |
| 2 | 8669404 | G | A | 0.872849 | 9.91E-10 | 0.000388366 |
| 2 | 8669641 | G | A | 0.869025 | 2.20E-09 | 0.000861457 |
| 2 | 8670125 | T | G | 0.869025 | 2.20E-09 | 0.000861457 |

| | | | | | | |
|---|---|---|---|---|---|---|
| 2 | 8670169 | T | C | 0.869025 | 2.20E-09 | 0.000861457 |
| 2 | 8670177 | T | C | 0.869025 | 2.20E-09 | 0.000861457 |
| 2 | 8670241 | C | T | 0.869025 | 2.20E-09 | 0.000861457 |
| 2 | 8670260 | T | C | 0.869025 | 2.20E-09 | 0.000861457 |
| 2 | 8670282 | G | A | 0.869025 | 2.20E-09 | 0.000861457 |
| 2 | 8670332 | C | T | 0.869025 | 2.20E-09 | 0.000861457 |
| 2 | 8670347 | A | C | 0.869025 | 2.20E-09 | 0.000861457 |
| 2 | 8670541 | G | A | 0.869025 | 2.20E-09 | 0.000861457 |
| 2 | 8670559 | C | T | 0.869025 | 2.20E-09 | 0.000861457 |
| 2 | 8670577 | C | T | 0.869025 | 2.20E-09 | 0.000861457 |
| 2 | 8670903 | T | C | 0.869025 | 2.20E-09 | 0.000861457 |
| 2 | 8671068 | A | G | 0.869025 | 2.20E-09 | 0.000861457 |
| 2 | 8671069 | C | T | 0.869025 | 2.20E-09 | 0.000861457 |
| 2 | 8671138 | C | T | 0.869025 | 2.20E-09 | 0.000861457 |
| 2 | 8672720 | C | T | 0.869025 | 2.20E-09 | 0.000861457 |
| 2 | 8672982 | A | G | 0.869025 | 2.20E-09 | 0.000861457 |
| 2 | 8673895 | A | C | 0.869025 | 2.20E-09 | 0.000861457 |
| 2 | 8674218 | C | T | 0.869025 | 2.20E-09 | 0.000861457 |
| 2 | 8674747 | C | T | 0.869025 | 2.20E-09 | 0.000861457 |
| 2 | 8676724 | G | T | 0.869025 | 2.20E-09 | 0.000861457 |
| 2 | 8676948 | A | G | 0.869025 | 2.20E-09 | 0.000861457 |
| 2 | 8677596 | G | A | 0.869025 | 2.20E-09 | 0.000861457 |
| 2 | 8677757 | A | C | 0.869025 | 2.20E-09 | 0.000861457 |
| 2 | 8679691 | A | G | 0.869025 | 2.20E-09 | 0.000861457 |
| 2 | 8680188 | G | A | 0.869025 | 2.20E-09 | 0.000861457 |
| 2 | 8680556 | C | T | 0.869025 | 2.20E-09 | 0.000861457 |
| 2 | 8680763 | C | T | 0.869025 | 2.20E-09 | 0.000861457 |
| 2 | 8681818 | G | A | 0.869025 | 2.20E-09 | 0.000861457 |
| 2 | 8682798 | G | A | 0.869025 | 2.20E-09 | 0.000861457 |
| 2 | 8683888 | A | G | 0.869025 | 2.20E-09 | 0.000861457 |
| 2 | 8684412 | G | C | 0.869025 | 2.20E-09 | 0.000861457 |
| 2 | 8684619 | T | C | 0.869025 | 2.20E-09 | 0.000861457 |
| 2 | 8685383 | G | C | 0.869025 | 2.20E-09 | 0.000861457 |
| 2 | 8685395 | T | G | 0.869025 | 2.20E-09 | 0.000861457 |
| 2 | 8685490 | A | G | 0.869025 | 2.20E-09 | 0.000861457 |
| 2 | 8685526 | C | A | 0.869025 | 2.20E-09 | 0.000861457 |
| 2 | 8686926 | G | A | 0.869025 | 2.20E-09 | 0.000861457 |
| 2 | 8687028 | T | G | 0.869025 | 2.20E-09 | 0.000861457 |
| 2 | 8687502 | C | A | 0.869025 | 2.20E-09 | 0.000861457 |
| 2 | 8687543 | G | A | 0.869025 | 2.20E-09 | 0.000861457 |
| 2 | 8687588 | A | C | 0.869025 | 2.20E-09 | 0.000861457 |
| 2 | 8687919 | G | A | 0.869025 | 2.20E-09 | 0.000861457 |
| 2 | 8688600 | T | A | 0.869025 | 2.20E-09 | 0.000861457 |
| 2 | 8688857 | T | C | 0.871893 | 2.66E-09 | 0.001044036 |
| 2 | 8688863 | C | T | 0.869025 | 2.20E-09 | 0.000861457 |
| 2 | 8688985 | T | C | 0.870937 | 4.26E-09 | 0.001670327 |
| 2 | 8689423 | G | A | 0.870937 | 4.26E-09 | 0.001670327 |
| 2 | 8689915 | C | T | 0.869025 | 2.20E-09 | 0.000861457 |
| 2 | 8690126 | T | G | 0.869025 | 2.20E-09 | 0.000861457 |
| 2 | 8692340 | G | A | 0.869025 | 2.20E-09 | 0.000861457 |
| 2 | 8692695 | G | C | 0.869025 | 2.20E-09 | 0.000861457 |
| 2 | 8692961 | A | G | 0.869025 | 2.20E-09 | 0.000861457 |
| 2 | 8693187 | G | A | 0.869025 | 2.20E-09 | 0.000861457 |
| 2 | 8693581 | G | A | 0.871893 | 2.66E-09 | 0.001044036 |
| 2 | 8693870 | A | C | 0.869025 | 2.20E-09 | 0.000861457 |
| 2 | 8693980 | A | G | 0.870937 | 4.26E-09 | 0.001670327 |
| 2 | 8694135 | A | C | 0.869981 | 1.52E-09 | 0.000595988 |
| 2 | 8697079 | C | T | 0.871893 | 2.66E-09 | 0.001044036 |
| 2 | 8699102 | A | G | 0.869025 | 2.20E-09 | 0.000861457 |
| 2 | 8700581 | C | T | 0.869025 | 2.20E-09 | 0.000861457 |
| 2 | 8701137 | C | T | 0.869025 | 2.20E-09 | 0.000861457 |
| 2 | 8701613 | T | C | 0.869025 | 2.20E-09 | 0.000861457 |
| 2 | 8701831 | G | A | 0.869025 | 2.20E-09 | 0.000861457 |
| 2 | 8701994 | T | C | 0.869981 | 1.18E-09 | 0.000463049 |
| 2 | 8702817 | C | T | 0.869025 | 2.20E-09 | 0.000861457 |
| 2 | 8702951 | G | T | 0.869025 | 2.20E-09 | 0.000861457 |
| 2 | 8703618 | C | T | 0.869025 | 2.20E-09 | 0.000861457 |

| | | | | | | |
|---|---|---|---|---|---|---|
| 2 | 8703759 | A | G | 0.869025 | 2.20E-09 | 0.000861457 |
| 2 | 8703847 | G | C | 0.869025 | 2.20E-09 | 0.000861457 |
| 2 | 8704002 | C | T | 0.869025 | 2.20E-09 | 0.000861457 |
| 2 | 8704106 | G | A | 0.869025 | 2.20E-09 | 0.000861457 |
| 2 | 8704909 | T | C | 0.869025 | 2.20E-09 | 0.000861457 |
| 2 | 8704948 | G | A | 0.869025 | 2.20E-09 | 0.000861457 |
| 2 | 8706431 | T | A | 0.869025 | 2.20E-09 | 0.000861457 |
| 2 | 8706670 | G | A | 0.869025 | 2.20E-09 | 0.000861457 |
| 2 | 8707086 | G | C | 0.867113 | 8.95E-09 | 0.003506794 |
| 2 | 8708757 | A | T | 0.876673 | 3.98E-09 | 0.001561785 |
| 2 | 8709090 | T | C | 0.873805 | 8.47E-09 | 0.003321385 |
| 2 | 8713757 | G | T | 0.869025 | 2.20E-09 | 0.000861457 |
| 2 | 8720155 | A | T | 0.869025 | 2.20E-09 | 0.000861457 |
| 2 | 8722898 | C | T | 0.869025 | 2.20E-09 | 0.000861457 |
| 2 | 8727247 | T | A | 0.869025 | 2.20E-09 | 0.000861457 |
| 2 | 8727673 | T | C | 0.869025 | 2.20E-09 | 0.000861457 |
| 2 | 8728049 | A | G | 0.869025 | 2.20E-09 | 0.000861457 |
| 2 | 8737916 | G | A | 0.871893 | 2.66E-09 | 0.001044036 |
| 2 | 8742223 | G | A | 0.869025 | 2.20E-09 | 0.000861457 |
| 2 | 8744871 | C | T | 0.869025 | 2.20E-09 | 0.000861457 |
| 2 | 8747154 | A | G | 0.869025 | 2.20E-09 | 0.000861457 |
| 2 | 8750304 | G | A | 0.869025 | 2.20E-09 | 0.000861457 |
| 2 | 8750838 | G | A | 0.869025 | 2.20E-09 | 0.000861457 |
| 2 | 8753432 | T | C | 0.869025 | 2.20E-09 | 0.000861457 |
| 2 | 8754263 | T | G | 0.869981 | 1.52E-09 | 0.000595988 |
| 2 | 8759136 | C | A | 0.869025 | 2.20E-09 | 0.000861457 |
| 2 | 8759654 | G | A | 0.869025 | 2.20E-09 | 0.000861457 |
| 2 | 8759902 | A | G | 0.869025 | 2.20E-09 | 0.000861457 |
| 2 | 8760010 | A | G | 0.871893 | 2.66E-09 | 0.001044036 |
| 2 | 8760974 | C | T | 0.869981 | 1.52E-09 | 0.000595988 |
| 2 | 8761017 | C | G | 0.869981 | 1.52E-09 | 0.000595988 |
| 2 | 8761920 | G | A | 0.870937 | 1.29E-09 | 0.000504991 |
| 2 | 8762211 | C | T | 0.871893 | 2.66E-09 | 0.001044036 |
| 2 | 8763207 | C | T | 0.871893 | 2.66E-09 | 0.001044036 |
| 2 | 8763452 | C | A | 0.871893 | 2.66E-09 | 0.001044036 |
| 2 | 8900300 | A | T | 0.916826 | 9.20E-08 | 0.036061557 |
| 2 | 8915463 | C | A | 0.914914 | 6.09E-08 | 0.023863969 |
| 2 | 10452262 | T | C | 0.803059 | 5.37E-08 | 0.021046916 |
| 2 | 10452573 | G | C | 0.82218 | 1.22E-09 | 0.000477192 |
| 2 | 10452723 | C | T | 0.82218 | 1.22E-09 | 0.000477192 |
| 2 | 10453071 | G | A | 0.82218 | 1.22E-09 | 0.000477192 |
| 2 | 10453200 | A | G | 0.821224 | 8.36E-10 | 0.000327835 |
| 2 | 10453232 | C | T | 0.821224 | 8.36E-10 | 0.000327835 |
| 2 | 10453938 | A | T | 0.825048 | 7.50E-09 | 0.002941125 |
| 2 | 10454021 | A | T | 0.825048 | 7.50E-09 | 0.002941125 |
| 2 | 10454107 | C | T | 0.825048 | 7.50E-09 | 0.002941125 |
| 2 | 10463060 | C | A | 0.819312 | 8.10E-09 | 0.003175155 |
| 2 | 10635097 | C | T | 0.808795 | 1.05E-07 | 0.041275073 |
| 2 | 10635132 | T | A | 0.808795 | 1.05E-07 | 0.041275073 |
| 2 | 10638113 | A | T | 0.823136 | 9.07E-10 | 0.000355465 |
| 2 | 10638114 | C | A | 0.823136 | 9.07E-10 | 0.000355465 |
| 2 | 10638143 | A | G | 0.823136 | 9.07E-10 | 0.000355465 |
| 2 | 10638252 | T | C | 0.823136 | 9.07E-10 | 0.000355465 |
| 2 | 10638278 | G | T | 0.823136 | 9.07E-10 | 0.000355465 |
| 2 | 10638365 | T | C | 0.823136 | 9.07E-10 | 0.000355465 |
| 2 | 10638466 | T | C | 0.823136 | 9.07E-10 | 0.000355465 |
| 2 | 10646029 | T | C | 0.830784 | 2.86E-09 | 0.001121224 |
| 2 | 10646031 | C | T | 0.830784 | 2.86E-09 | 0.001121224 |
| 2 | 10646089 | T | C | 0.830784 | 2.86E-09 | 0.001121224 |
| 2 | 10646210 | T | A | 0.830784 | 2.86E-09 | 0.001121224 |
| 2 | 10646235 | T | A | 0.830784 | 2.86E-09 | 0.001121224 |
| 2 | 10646303 | G | T | 0.830784 | 2.86E-09 | 0.001121224 |
| 2 | 10646404 | T | C | 0.830784 | 2.86E-09 | 0.001121224 |
| 2 | 10646703 | A | T | 0.830784 | 2.86E-09 | 0.001121224 |
| 2 | 10646781 | C | T | 0.830784 | 2.86E-09 | 0.001121224 |
| 2 | 10646866 | G | A | 0.830784 | 2.86E-09 | 0.001121224 |
| 2 | 10647074 | C | T | 0.830784 | 2.86E-09 | 0.001121224 |

| | | | | | | |
|---|---|---|---|---|---|---|
| 2 | 10647175 | C | T | 0.830784 | 2.86E-09 | 0.001121224 |
| 2 | 10647215 | A | T | 0.830784 | 2.86E-09 | 0.001121224 |
| 2 | 10647311 | T | C | 0.830784 | 2.86E-09 | 0.001121224 |
| 2 | 10647522 | G | A | 0.830784 | 2.86E-09 | 0.001121224 |
| 2 | 10647737 | C | T | 0.830784 | 2.86E-09 | 0.001121224 |
| 2 | 10647775 | T | A | 0.830784 | 2.86E-09 | 0.001121224 |
| 2 | 10647788 | G | A | 0.830784 | 2.86E-09 | 0.001121224 |
| 2 | 10647790 | G | A | 0.830784 | 2.86E-09 | 0.001121224 |
| 2 | 10647807 | G | A | 0.830784 | 2.86E-09 | 0.001121224 |
| 2 | 10647820 | G | A | 0.830784 | 2.86E-09 | 0.001121224 |
| 2 | 10648019 | A | G | 0.830784 | 2.86E-09 | 0.001121224 |
| 2 | 10648113 | C | A | 0.830784 | 2.86E-09 | 0.001121224 |
| 2 | 10648134 | C | A | 0.830784 | 2.86E-09 | 0.001121224 |
| 2 | 10648271 | G | A | 0.830784 | 2.86E-09 | 0.001121224 |
| 2 | 10648361 | T | A | 0.830784 | 2.86E-09 | 0.001121224 |
| 2 | 10649065 | T | C | 0.830784 | 2.86E-09 | 0.001121224 |
| 2 | 10649385 | T | C | 0.830784 | 2.86E-09 | 0.001121224 |
| 2 | 10649481 | C | T | 0.830784 | 2.86E-09 | 0.001121224 |
| 2 | 10649482 | C | T | 0.830784 | 2.86E-09 | 0.001121224 |
| 2 | 10649764 | T | C | 0.830784 | 2.86E-09 | 0.001121224 |
| 2 | 10649806 | G | C | 0.830784 | 2.86E-09 | 0.001121224 |
| 2 | 10649864 | T | C | 0.830784 | 2.86E-09 | 0.001121224 |
| 2 | 10649958 | A | G | 0.830784 | 2.86E-09 | 0.001121224 |
| 2 | 10649966 | G | A | 0.830784 | 2.86E-09 | 0.001121224 |
| 2 | 10649972 | C | T | 0.830784 | 2.86E-09 | 0.001121224 |
| 2 | 10650133 | A | G | 0.830784 | 2.86E-09 | 0.001121224 |
| 2 | 10650218 | A | G | 0.830784 | 2.86E-09 | 0.001121224 |
| 2 | 10650403 | A | G | 0.830784 | 2.86E-09 | 0.001121224 |
| 2 | 10650430 | A | G | 0.830784 | 2.86E-09 | 0.001121224 |
| 2 | 10650468 | G | C | 0.830784 | 2.86E-09 | 0.001121224 |
| 2 | 10650469 | T | A | 0.830784 | 2.86E-09 | 0.001121224 |
| 2 | 10650645 | C | T | 0.830784 | 2.86E-09 | 0.001121224 |
| 2 | 10650667 | C | A | 0.830784 | 2.86E-09 | 0.001121224 |
| 2 | 10650823 | A | T | 0.830784 | 2.86E-09 | 0.001121224 |
| 2 | 10651366 | C | G | 0.830784 | 2.86E-09 | 0.001121224 |
| 2 | 10652262 | A | G | 0.830784 | 2.86E-09 | 0.001121224 |
| 2 | 10711376 | C | T | 0.83174 | 2.43E-09 | 0.00095071 |
| 2 | 10747664 | C | A | 0.832696 | 3.62E-09 | 0.001420686 |
| 2 | 10747872 | T | A | 0.832696 | 3.62E-09 | 0.001420686 |
| 2 | 10765742 | G | A | 0.805927 | 6.40E-08 | 0.025080741 |
| 2 | 10765804 | G | A | 0.805927 | 6.40E-08 | 0.025080741 |
| 2 | 10769439 | G | A | 0.83174 | 1.03E-09 | 0.000402187 |
| 2 | 10775435 | A | G | 0.830784 | 2.86E-09 | 0.001121224 |
| 2 | 10775442 | T | A | 0.830784 | 2.86E-09 | 0.001121224 |
| 2 | 10775452 | A | G | 0.830784 | 2.86E-09 | 0.001121224 |
| 2 | 10775467 | A | G | 0.830784 | 2.86E-09 | 0.001121224 |
| 2 | 10775509 | C | T | 0.830784 | 2.86E-09 | 0.001121224 |
| 2 | 10775651 | T | C | 0.830784 | 2.86E-09 | 0.001121224 |
| 2 | 10775654 | A | G | 0.830784 | 2.86E-09 | 0.001121224 |
| 2 | 10775666 | G | T | 0.830784 | 2.86E-09 | 0.001121224 |
| 2 | 10775668 | A | G | 0.830784 | 2.86E-09 | 0.001121224 |
| 2 | 10775689 | G | C | 0.830784 | 2.86E-09 | 0.001121224 |
| 2 | 10775723 | G | T | 0.830784 | 2.86E-09 | 0.001121224 |
| 2 | 10776109 | G | A | 0.830784 | 2.86E-09 | 0.001121224 |
| 2 | 10776199 | T | C | 0.830784 | 2.86E-09 | 0.001121224 |
| 2 | 10776277 | C | T | 0.830784 | 2.86E-09 | 0.001121224 |
| 2 | 10776349 | A | G | 0.830784 | 2.86E-09 | 0.001121224 |
| 2 | 10776482 | A | G | 0.830784 | 2.86E-09 | 0.001121224 |
| 2 | 10776671 | A | G | 0.830784 | 2.86E-09 | 0.001121224 |
| 2 | 10776947 | A | G | 0.830784 | 2.86E-09 | 0.001121224 |
| 2 | 10776997 | G | T | 0.830784 | 2.86E-09 | 0.001121224 |
| 2 | 10777152 | G | A | 0.830784 | 2.86E-09 | 0.001121224 |
| 2 | 10777160 | C | T | 0.830784 | 2.86E-09 | 0.001121224 |
| 2 | 10777165 | A | T | 0.830784 | 2.86E-09 | 0.001121224 |
| 2 | 10777285 | T | C | 0.830784 | 2.86E-09 | 0.001121224 |
| 2 | 10777306 | T | A | 0.830784 | 2.86E-09 | 0.001121224 |
| 2 | 10777340 | T | C | 0.830784 | 2.86E-09 | 0.001121224 |

| | | | | | | |
|---|---|---|---|---|---|---|
| 2 | 10777387 | G | A | 0.830784 | 2.86E-09 | 0.001121224 |
| 2 | 10777442 | C | G | 0.830784 | 2.86E-09 | 0.001121224 |
| 2 | 10777798 | T | C | 0.830784 | 2.86E-09 | 0.001121224 |
| 2 | 10777875 | C | T | 0.830784 | 2.86E-09 | 0.001121224 |
| 2 | 10778379 | C | A | 0.83174 | 3.87E-09 | 0.00151592 |
| 2 | 10778773 | A | G | 0.830784 | 2.86E-09 | 0.001121224 |
| 2 | 10778893 | C | T | 0.830784 | 2.86E-09 | 0.001121224 |
| 2 | 10780482 | G | A | 0.830784 | 2.86E-09 | 0.001121224 |
| 2 | 10780853 | C | A | 0.830784 | 2.86E-09 | 0.001121224 |
| 2 | 10780868 | A | G | 0.830784 | 2.86E-09 | 0.001121224 |
| 2 | 10781028 | T | A | 0.830784 | 2.86E-09 | 0.001121224 |
| 2 | 10782155 | G | C | 0.830784 | 2.86E-09 | 0.001121224 |
| 2 | 10782284 | T | C | 0.830784 | 2.86E-09 | 0.001121224 |
| 2 | 10782361 | A | G | 0.830784 | 2.86E-09 | 0.001121224 |
| 2 | 10782468 | A | G | 0.830784 | 2.86E-09 | 0.001121224 |
| 2 | 10782623 | A | T | 0.830784 | 2.86E-09 | 0.001121224 |
| 2 | 10782635 | T | C | 0.830784 | 2.86E-09 | 0.001121224 |
| 2 | 10782646 | A | C | 0.830784 | 2.86E-09 | 0.001121224 |
| 2 | 10782790 | A | T | 0.830784 | 2.86E-09 | 0.001121224 |
| 2 | 10782983 | G | A | 0.830784 | 2.86E-09 | 0.001121224 |
| 2 | 10782991 | T | C | 0.830784 | 2.86E-09 | 0.001121224 |
| 2 | 10782996 | T | C | 0.830784 | 2.86E-09 | 0.001121224 |
| 2 | 10783002 | G | A | 0.830784 | 2.86E-09 | 0.001121224 |
| 2 | 10783009 | G | A | 0.830784 | 2.86E-09 | 0.001121224 |
| 2 | 10783077 | G | T | 0.830784 | 2.86E-09 | 0.001121224 |
| 2 | 10783140 | A | T | 0.830784 | 2.86E-09 | 0.001121224 |
| 2 | 10783155 | A | G | 0.830784 | 2.86E-09 | 0.001121224 |
| 2 | 10783178 | A | C | 0.830784 | 2.86E-09 | 0.001121224 |
| 2 | 10783949 | C | A | 0.830784 | 2.86E-09 | 0.001121224 |
| 2 | 10784724 | A | G | 0.830784 | 2.86E-09 | 0.001121224 |
| 2 | 10784944 | C | T | 0.830784 | 2.86E-09 | 0.001121224 |
| 2 | 10793615 | T | C | 0.83174 | 1.59E-09 | 0.000624771 |
| 2 | 10794747 | A | G | 0.83174 | 1.03E-09 | 0.000402187 |
| 2 | 10795310 | C | T | 0.83174 | 1.03E-09 | 0.000402187 |
| 2 | 10795314 | G | A | 0.83174 | 1.03E-09 | 0.000402187 |
| 2 | 10800031 | A | G | 0.839388 | 9.92E-10 | 0.00038871 |
| 2 | 10812499 | G | A | 0.838432 | 9.20E-10 | 0.000360746 |
| 2 | 10812592 | A | G | 0.838432 | 9.20E-10 | 0.000360746 |
| 2 | 10813531 | G | A | 0.837476 | 9.60E-10 | 0.000376405 |
| 2 | 10813544 | C | G | 0.837476 | 9.60E-10 | 0.000376405 |
| 2 | 10814282 | G | A | 0.837476 | 9.60E-10 | 0.000376405 |
| 2 | 10814319 | G | A | 0.837476 | 9.60E-10 | 0.000376405 |
| 2 | 10818838 | C | T | 0.825048 | 1.09E-09 | 0.000427622 |
| 2 | 10826862 | A | G | 0.802103 | 1.14E-07 | 0.044756212 |
| 2 | 10826934 | T | C | 0.802103 | 1.14E-07 | 0.044756212 |
| 2 | 10827141 | A | G | 0.802103 | 1.14E-07 | 0.044756212 |
| 2 | 11136646 | C | G | 0.8413 | 1.19E-09 | 0.00046584 |
| 2 | 11136656 | T | G | 0.8413 | 1.19E-09 | 0.00046584 |
| 2 | 11136658 | A | C | 0.8413 | 1.19E-09 | 0.00046584 |
| 2 | 11136701 | G | A | 0.8413 | 1.19E-09 | 0.00046584 |
| 2 | 11136706 | C | T | 0.8413 | 1.19E-09 | 0.00046584 |
| 2 | 11136716 | A | G | 0.8413 | 1.19E-09 | 0.00046584 |
| 2 | 11136720 | C | T | 0.8413 | 1.19E-09 | 0.00046584 |
| 2 | 11136725 | T | G | 0.8413 | 1.19E-09 | 0.00046584 |
| 2 | 11136726 | C | T | 0.8413 | 1.19E-09 | 0.00046584 |
| 2 | 11136727 | A | G | 0.8413 | 1.19E-09 | 0.00046584 |
| 2 | 11138284 | A | G | 0.829828 | 8.77E-10 | 0.000343919 |
| 2 | 11148449 | C | T | 0.843212 | 1.24E-09 | 0.000487555 |
| 2 | 11151869 | A | G | 0.826004 | 1.41E-09 | 0.000550746 |
| 2 | 11155339 | A | C | 0.843212 | 1.24E-09 | 0.000487555 |
| 2 | 11155773 | A | G | 0.826004 | 1.41E-09 | 0.000550746 |
| 2 | 11157503 | C | G | 0.827916 | 1.78E-09 | 0.000698964 |
| 2 | 11157881 | G | A | 0.827916 | 1.78E-09 | 0.000698964 |
| 2 | 11157928 | C | T | 0.827916 | 1.78E-09 | 0.000698964 |
| 2 | 11158022 | T | C | 0.842256 | 3.60E-09 | 0.001409542 |
| 2 | 11158132 | C | G | 0.827916 | 1.78E-09 | 0.000698964 |
| 2 | 11158282 | G | A | 0.827916 | 1.78E-09 | 0.000698964 |

| | | | | | | |
|---|---|---|---|---|---|---|
| 2 | 11158351 | A | G | 0.827916 | 1.78E-09 | 0.000698964 |
| 2 | 11158366 | C | T | 0.827916 | 1.78E-09 | 0.000698964 |
| 2 | 11158376 | A | G | 0.827916 | 1.78E-09 | 0.000698964 |
| 2 | 11158627 | G | A | 0.827916 | 1.78E-09 | 0.000698964 |
| 2 | 11159414 | G | T | 0.82696 | 3.93E-09 | 0.001538757 |
| 2 | 11159415 | C | T | 0.82696 | 3.93E-09 | 0.001538757 |
| 2 | 11159887 | C | G | 0.826004 | 2.60E-09 | 0.001020133 |
| 2 | 11159923 | G | T | 0.826004 | 2.60E-09 | 0.001020133 |
| 2 | 11160104 | A | T | 0.826004 | 2.60E-09 | 0.001020133 |
| 2 | 11161298 | T | C | 0.825048 | 1.45E-09 | 0.000566539 |
| 2 | 11161409 | A | G | 0.825048 | 1.45E-09 | 0.000566539 |
| 2 | 11161846 | G | A | 0.826004 | 1.41E-09 | 0.000550746 |
| 2 | 11162131 | C | T | 0.826004 | 1.41E-09 | 0.000550746 |
| 2 | 11162273 | A | T | 0.826004 | 1.41E-09 | 0.000550746 |
| 2 | 11162549 | T | C | 0.835564 | 1.27E-09 | 0.000496892 |
| 2 | 11162589 | A | T | 0.834608 | 1.47E-09 | 0.000575691 |
| 2 | 11162655 | A | G | 0.824092 | 4.15E-09 | 0.001626285 |
| 2 | 11163295 | G | A | 0.826004 | 1.41E-09 | 0.000550746 |
| 2 | 11163602 | C | T | 0.825048 | 1.67E-09 | 0.000655835 |
| 2 | 11163638 | T | C | 0.82696 | 1.18E-09 | 0.000463692 |
| 2 | 11163749 | C | G | 0.825048 | 1.67E-09 | 0.000655835 |
| 2 | 11163889 | G | C | 0.826004 | 1.41E-09 | 0.000550746 |
| 2 | 11164230 | T | C | 0.827916 | 1.78E-09 | 0.000698964 |
| 2 | 11164390 | A | G | 0.827916 | 1.78E-09 | 0.000698964 |
| 2 | 11164555 | G | A | 0.827916 | 1.78E-09 | 0.000698964 |
| 2 | 11164587 | C | A | 0.827916 | 1.78E-09 | 0.000698964 |
| 2 | 11164725 | T | C | 0.827916 | 1.78E-09 | 0.000698964 |
| 2 | 11164932 | C | T | 0.834608 | 1.33E-09 | 0.000523072 |
| 2 | 11189749 | G | T | 0.85086 | 3.58E-09 | 0.001402533 |
| 2 | 11204893 | C | T | 0.847036 | 1.61E-09 | 0.000632618 |
| 2 | 11214446 | A | G | 0.833652 | 1.08E-09 | 0.000424302 |
| 2 | 11214527 | T | C | 0.833652 | 1.08E-09 | 0.000424302 |
| 2 | 11215180 | C | T | 0.83174 | 8.45E-10 | 0.000331308 |
| 2 | 11218394 | A | G | 0.833652 | 1.08E-09 | 0.000424302 |
| 2 | 11220151 | C | T | 0.833652 | 1.08E-09 | 0.000424302 |
| 2 | 11222044 | A | G | 0.832696 | 2.48E-09 | 0.000973053 |
| 2 | 11222725 | C | T | 0.83174 | 8.45E-10 | 0.000331308 |
| 2 | 11246227 | G | A | 0.833652 | 1.08E-09 | 0.000424302 |
| 2 | 11260789 | A | C | 0.838432 | 1.17E-09 | 0.000459408 |
| 2 | 11261042 | A | C | 0.838432 | 1.17E-09 | 0.000459408 |
| 2 | 11261054 | A | G | 0.838432 | 1.17E-09 | 0.000459408 |
| 2 | 11295769 | A | G | 0.861377 | 1.10E-09 | 0.000431428 |
| 2 | 11295963 | C | T | 0.861377 | 1.10E-09 | 0.000431428 |
| 2 | 11296169 | C | T | 0.861377 | 1.10E-09 | 0.000431428 |
| 2 | 11296234 | A | G | 0.861377 | 1.10E-09 | 0.000431428 |
| 2 | 11296312 | G | A | 0.861377 | 1.10E-09 | 0.000431428 |
| 2 | 11296339 | A | T | 0.861377 | 1.10E-09 | 0.000431428 |
| 2 | 11296366 | T | C | 0.861377 | 1.10E-09 | 0.000431428 |
| 2 | 11296503 | A | C | 0.861377 | 1.10E-09 | 0.000431428 |
| 2 | 11296511 | G | C | 0.861377 | 1.10E-09 | 0.000431428 |
| 2 | 11296512 | G | A | 0.861377 | 1.10E-09 | 0.000431428 |
| 2 | 11296544 | C | A | 0.861377 | 1.10E-09 | 0.000431428 |
| 2 | 11296550 | A | G | 0.861377 | 1.10E-09 | 0.000431428 |
| 2 | 11296582 | T | A | 0.861377 | 1.10E-09 | 0.000431428 |
| 2 | 11296658 | G | A | 0.861377 | 1.10E-09 | 0.000431428 |
| 2 | 11296781 | G | T | 0.861377 | 1.10E-09 | 0.000431428 |
| 2 | 11297076 | G | A | 0.861377 | 1.10E-09 | 0.000431428 |
| 2 | 11297187 | C | A | 0.861377 | 1.10E-09 | 0.000431428 |
| 2 | 11297214 | T | A | 0.861377 | 1.10E-09 | 0.000431428 |
| 2 | 11297339 | T | C | 0.861377 | 1.10E-09 | 0.000431428 |
| 2 | 11297495 | T | A | 0.861377 | 1.10E-09 | 0.000431428 |
| 2 | 11297543 | G | A | 0.861377 | 1.10E-09 | 0.000431428 |
| 2 | 11297642 | C | A | 0.861377 | 1.10E-09 | 0.000431428 |
| 2 | 11297912 | T | A | 0.861377 | 1.10E-09 | 0.000431428 |
| 2 | 11298495 | A | G | 0.865201 | 3.17E-09 | 0.001243752 |
| 2 | 11298528 | G | A | 0.865201 | 3.17E-09 | 0.001243752 |
| 2 | 11298534 | A | C | 0.865201 | 3.17E-09 | 0.001243752 |

| | | | | | | |
|---|---|---|---|---|---|---|
| 2 | 11299449 | T | C | 0.861377 | 1.10E-09 | 0.000431428 |
| 2 | 11299478 | T | G | 0.861377 | 1.10E-09 | 0.000431428 |
| 2 | 11299727 | T | C | 0.861377 | 1.10E-09 | 0.000431428 |
| 2 | 11299757 | C | A | 0.861377 | 1.10E-09 | 0.000431428 |
| 2 | 11299825 | C | T | 0.861377 | 1.10E-09 | 0.000431428 |
| 2 | 11299875 | C | T | 0.861377 | 1.10E-09 | 0.000431428 |
| 2 | 11299971 | A | C | 0.861377 | 1.10E-09 | 0.000431428 |
| 2 | 11300018 | C | T | 0.861377 | 1.10E-09 | 0.000431428 |
| 2 | 11300067 | T | C | 0.861377 | 1.10E-09 | 0.000431428 |
| 2 | 11300170 | C | T | 0.861377 | 1.10E-09 | 0.000431428 |
| 2 | 11300237 | C | T | 0.861377 | 1.10E-09 | 0.000431428 |
| 2 | 11300239 | C | G | 0.861377 | 1.10E-09 | 0.000431428 |
| 2 | 11300532 | A | G | 0.861377 | 1.10E-09 | 0.000431428 |
| 2 | 11300558 | C | T | 0.861377 | 1.10E-09 | 0.000431428 |
| 2 | 11364052 | C | T | 0.863289 | 4.37E-09 | 0.00171259 |
| 2 | 11364130 | C | T | 0.875717 | 5.39E-08 | 0.021107947 |
| 2 | 11364179 | A | G | 0.876673 | 2.46E-09 | 0.00096283 |
| 2 | 13590275 | T | C | 0.885277 | 4.70E-08 | 0.018403352 |
| 2 | 13590873 | T | A | 0.885277 | 4.70E-08 | 0.018403352 |
| 2 | 13603173 | G | A | 0.885277 | 4.70E-08 | 0.018403352 |
| 2 | 13603320 | G | C | 0.885277 | 4.70E-08 | 0.018403352 |
| 2 | 13603540 | G | A | 0.885277 | 4.70E-08 | 0.018403352 |
| 2 | 13603555 | G | A | 0.885277 | 4.70E-08 | 0.018403352 |
| 2 | 13603559 | T | C | 0.885277 | 4.70E-08 | 0.018403352 |
| 4 | 38127528 | A | G | 0.916826 | 1.09E-07 | 0.042842977 |
| 12 | 148369145 | G | A | 0.916826 | 8.02E-08 | 0.031453252 |

**Table S15. SNP-based heritability**

| SNP-based heritability | BW | CVDFW | DFW | LFW | LCW | RCW |
|---|---|---|---|---|---|---|
| **D35** [1] | 53.87% | — | — | — | — | — |
| **D70** | 23.95% | 1.37% | 27.01% | 21.18% | 23.06% | 5.27% |
| **D140** | 39.01% | 10.20% | 30.52% | 2.82% | 5.96% | 0.23% |
| **D210** | 39.35% | 10.79% | 33.73% | 11.20% | 19.46% | 21.73% |
| **Mean** | 39.05% | 7.45% | 30.42% | 11.74% | 16.16% | 9.07% |
| **Standard Error** | 5.29% | 2.49% | 1.59% | 4.33% | 4.25% | 5.30% |

1. The trait was measured at 35 days of age.

**Table S16. QTL contribution to heritability**

| Trait | QTL No. | Gene No. in QTLs | Variance explained [1] | Variance explained by QTL1 | Variance explained by QTL2 | Variance explained by QTL3 |
|---|---|---|---|---|---|---|
| **CVDFW** | 1 | 6 | 2.28% | 2.28% | — | — |
| **DFW** | 2 | 0 | 8.52% | 1.10% | 7.50% | — |
| **BW** | 3 | 68 | 5.49% | 2.32% | 0.42% | 3.15% |

1. Total phenotypic variance explained by QTLs.

**Table S17. Genes overlapped by QTLs**

| Chr | QTLstart | QTLend | ID | Trait | Gene |
|---|---|---|---|---|---|
| 2 | 8257045 | 8923024 | QTL1_95%CI | BW | ID=gene-FAM184B;Dbxref=GeneID:100347213;Name=FAM184B;gbkey=Gene;gene=FAM184B;gene_biotype=protein_coding |
| 2 | 8257045 | 8923024 | QTL1_95%CI | BW | ID=gene-DCAF16;Dbxref=GeneID:100347465;Name=DCAF16;gbkey=Gene;gene=DCAF16;gene_biotype=protein_coding |
| 2 | 8257045 | 8923024 | QTL1_95%CI | BW | ID=gene-NCAPG;Dbxref=GeneID:100347720;Name=NCAPG;gbkey=Gene;gene=NCAPG;gene_biotype=protein_coding |
| 2 | 8257045 | 8923024 | QTL1_95%CI | BW | ID=gene-LCORL;Dbxref=GeneID:100347964;Name=LCORL;gbkey=Gene;gene=LCORL;gene_biotype=protein_coding |
| 4 | 36704911 | 38161753 | QTL2_95%CI | BW | ID=gene-LOC100348136;Dbxref=GeneID:100348136;Name=LOC100348136;gbkey=Gene;gene=LOC100348136;gene_biotype=protein_coding |
| 4 | 36704911 | 38161753 | QTL2_95%CI | BW | ID=gene-KRT3;Dbxref=GeneID:100353668;Name=KRT3;gbkey=Gene;gene=KRT3;gene_biotype=protein_coding |
| 4 | 36704911 | 38161753 | QTL2_95%CI | BW | ID=gene-KRT4;Dbxref=GeneID:100348388;Name=KRT4;gbkey=Gene;gene=KRT4;gene_biotype=protein_coding |
| 4 | 36704911 | 38161753 | QTL2_95%CI | BW | ID=gene-LOC100348640;Dbxref=GeneID:100348640;Name=LOC100348640;gbkey=Gene;gene=LOC100348640;gene_biotype=protein_coding |
| 4 | 36704911 | 38161753 | QTL2_95%CI | BW | ID=gene-KRT78;Dbxref=GeneID:100353913;Name=KRT78;gbkey=Gene;gene=KRT78;gene_biotype=protein_coding |
| 4 | 36704911 | 38161753 | QTL2_95%CI | BW | ID=gene-KRT8;Dbxref=GeneID:100348891;Name=KRT8;gbkey=Gene;gene=KRT8;gene_biotype=protein_coding |
| 4 | 36704911 | 38161753 | QTL2_95%CI | BW | ID=gene-LOC103348274;Dbxref=GeneID:103348274;Name=LOC103348274;gbkey=Gene;gene=LOC103348274;gene_biotype=lncRNA |
| 4 | 36704911 | 38161753 | QTL2_95%CI | BW | ID=gene-KRT18;Dbxref=GeneID:100354174;Name=KRT18;gbkey=Gene;gene=KRT18;gene_biotype=protein_coding;partial=true |
| 4 | 36704911 | 38161753 | QTL2_95%CI | BW | ID=gene-EIF4B;Dbxref=GeneID:100354421;Name=EIF4B;gbkey=Gene;gene=EIF4B;gene_biotype=protein_coding |
| 4 | 36704911 | 38161753 | QTL2_95%CI | BW | ID=gene-TNS2;Dbxref=GeneID:103348305;Name=TNS2;gbkey=Gene;gene=TNS2;gene_biotype=protein_coding;partial=true |
| 4 | 36704911 | 38161753 | QTL2_95%CI | BW | ID=gene-LOC103348275;Dbxref=GeneID:103348275;Name=LOC103348275;gbkey=Gene;gene=LOC103348275;gene_biotype=protein_coding;partial=true;start_range=.,36925711 |
| 4 | 36704911 | 38161753 | QTL2_95%CI | BW | ID=gene-SPRYD3;Dbxref=GeneID:100354677;Name=SPRYD3;gbkey=Gene;gene=SPRYD3;gene_biotype=protein_coding |

| | | | | | |
|---|---|---|---|---|---|
| 4 | 36704911 | 38161753 | QTL2_95%CI | BW | ID=gene-LOC108176645;Dbxref=GeneID:108176645;Name=LOC108176645;gbkey=Gene;gene=LOC108176645;gene_biotype=lncRNA |
| 4 | 36704911 | 38161753 | QTL2_95%CI | BW | ID=gene-IGFBP6;Dbxref=GeneID:100355364;Name=IGFBP6;gbkey=Gene;gene=IGFBP6;gene_biotype=protein_coding |
| 4 | 36704911 | 38161753 | QTL2_95%CI | BW | ID=gene-SOAT2;Dbxref=GeneID:100349393;Name=SOAT2;gbkey=Gene;gene=SOAT2;gene_biotype=protein_coding |
| 4 | 36704911 | 38161753 | QTL2_95%CI | BW | ID=gene-CSAD;Dbxref=GeneID:100355612;Name=CSAD;gbkey=Gene;gene=CSAD;gene_biotype=protein_coding |
| 4 | 36704911 | 38161753 | QTL2_95%CI | BW | ID=gene-ZNF740;Dbxref=GeneID:100349908;Name=ZNF740;gbkey=Gene;gene=ZNF740;gene_biotype=protein_coding |
| 4 | 36704911 | 38161753 | QTL2_95%CI | BW | ID=gene-ITGB7;Dbxref=GeneID:100350557;Name=ITGB7;gbkey=Gene;gene=ITGB7;gene_biotype=protein_coding |
| 4 | 36704911 | 38161753 | QTL2_95%CI | BW | ID=gene-RARG;Dbxref=GeneID:100355857;Name=RARG;end_range=37071079,.;gbkey=Gene;gene=RARG;gene_biotype=protein_coding;partial=true |
| 4 | 36704911 | 38161753 | QTL2_95%CI | BW | ID=gene-MFSD5;Dbxref=GeneID:100350813;Name=MFSD5;gbkey=Gene;gene=MFSD5;gene_biotype=protein_coding |
| 4 | 36704911 | 38161753 | QTL2_95%CI | BW | ID=gene-ESPL1;Dbxref=GeneID:100356105;Name=ESPL1;gbkey=Gene;gene=ESPL1;gene_biotype=protein_coding |
| 4 | 36704911 | 38161753 | QTL2_95%CI | BW | ID=gene-PFDN5;Dbxref=GeneID:100356360;Name=PFDN5;gbkey=Gene;gene=PFDN5;gene_biotype=protein_coding |
| 4 | 36704911 | 38161753 | QTL2_95%CI | BW | ID=gene-C4H12orf10;Dbxref=GeneID:100351065;Name=C4H12orf10;gbkey=Gene;gene=C4H12orf10;gene_biotype=protein_coding |
| 4 | 36704911 | 38161753 | QTL2_95%CI | BW | ID=gene-AAAS;Dbxref=GeneID:100356621;Name=AAAS;gbkey=Gene;gene=AAAS;gene_biotype=protein_coding |
| 4 | 36704911 | 38161753 | QTL2_95%CI | BW | ID=gene-SP7;Dbxref=GeneID:100356871;Name=SP7;gbkey=Gene;gene=SP7;gene_biotype=protein_coding |
| 4 | 36704911 | 38161753 | QTL2_95%CI | BW | ID=gene-SP1;Dbxref=GeneID:100351317;Name=SP1;gbkey=Gene;gene=SP1;gene_biotype=protein_coding |
| 4 | 36704911 | 38161753 | QTL2_95%CI | BW | ID=gene-AMHR2;Dbxref=GeneID:100009599;Name=AMHR2;description=anti-Mullerian |
| 4 | 36704911 | 38161753 | QTL2_95%CI | BW | ID=gene-PRR13;Dbxref=GeneID:100357129;Name=PRR13;gbkey=Gene;gene=PRR13;gene_biotype=protein_coding |

| | | | | | |
|---|---|---|---|---|---|
| 4 | 36704911 | 38161753 | QTL2_95%CI | BW | ID=gene-PCBP2;Dbxref=GeneID:100357382;Name=PCBP2;gbkey=Gene;gene=PCBP2;gene_biotype=protein_coding |
| 4 | 36704911 | 38161753 | QTL2_95%CI | BW | ID=gene-MAP3K12;Dbxref=GeneID:100358158;Name=MAP3K12;gbkey=Gene;gene=MAP3K12;gene_biotype=protein_coding |
| 4 | 36704911 | 38161753 | QTL2_95%CI | BW | ID=gene-TARBP2;Dbxref=GeneID:100358598;Name=TARBP2;gbkey=Gene;gene=TARBP2;gene_biotype=protein_coding |
| 4 | 36704911 | 38161753 | QTL2_95%CI | BW | ID=gene-NPFF;Dbxref=GeneID:100358860;Name=NPFF;gbkey=Gene;gene=NPFF;gene_biotype=protein_coding |
| 4 | 36704911 | 38161753 | QTL2_95%CI | BW | ID=gene-ATF7;Dbxref=GeneID:100351569;Name=ATF7;gbkey=Gene;gene=ATF7;gene_biotype=protein_coding |
| 4 | 36704911 | 38161753 | QTL2_95%CI | BW | ID=gene-LOC100359118;Dbxref=GeneID:100359118;Name=LOC100359118;gbkey=Gene;gene=LOC100359118;gene_biotype=protein_coding |
| 4 | 36704911 | 38161753 | QTL2_95%CI | BW | ID=gene-LOC108176647;Dbxref=GeneID:108176647;Name=LOC108176647;gbkey=Gene;gene=LOC108176647;gene_biotype=lncRNA |
| 4 | 36704911 | 38161753 | QTL2_95%CI | BW | ID=gene-ATP5G2;Dbxref=GeneID:100351820;Name=ATP5G2;gbkey=Gene;gene=ATP5G2;gene_biotype=protein_coding |
| 4 | 36704911 | 38161753 | QTL2_95%CI | BW | ID=gene-LOC108176646;Dbxref=GeneID:108176646;Name=LOC108176646;gbkey=Gene;gene=LOC108176646;gene_biotype=lncRNA |
| 4 | 36704911 | 38161753 | QTL2_95%CI | BW | ID=gene-CALCOCO1;Dbxref=GeneID:100337884;Name=CALCOCO1;gbkey=Gene;gene=CALCOCO1;gene_biotype=protein_coding;partial=true |
| 4 | 36704911 | 38161753 | QTL2_95%CI | BW | ID=gene-LOC103348277;Dbxref=GeneID:103348277;Name=LOC103348277;gbkey=Gene;gene=LOC103348277;gene_biotype=lncRNA |
| 4 | 36704911 | 38161753 | QTL2_95%CI | BW | ID=gene-HOXC13;Dbxref=GeneID:100352321;Name=HOXC13;gbkey=Gene;gene=HOXC13;gene_biotype=protein_coding |
| 4 | 36704911 | 38161753 | QTL2_95%CI | BW | ID=gene-HOXC12;Dbxref=GeneID:100338310;Name=HOXC12;gbkey=Gene;gene=HOXC12;gene_biotype=protein_coding |
| 4 | 36704911 | 38161753 | QTL2_95%CI | BW | ID=gene-HOXC11;Dbxref=GeneID:100338559;Name=HOXC11;gbkey=Gene;gene=HOXC11;gene_biotype=protein_coding |
| 4 | 36704911 | 38161753 | QTL2_95%CI | BW | ID=gene-HOXC10;Dbxref=GeneID:100338808;Name=HOXC10;gbkey=Gene;gene=HOXC10;gene_biotype=protein_coding |
| 4 | 36704911 | 38161753 | QTL2_95%CI | BW | ID=gene-LOC103348279;Dbxref=GeneID:103348279;Name=LOC103348279;gbkey=Gene;gene=LOC103348279;gene_biotype=lncRNA |

| | | | | | |
|---|---|---|---|---|---|
| 4 | 36704911 | 38161753 | QTL2_95%CI | BW | ID=gene-HOXC9;Dbxref=GeneID:103348280;Name=HOXC9;gbkey=Gene;gene=HOXC9;gene_biotype=protein_coding |
| 4 | 36704911 | 38161753 | QTL2_95%CI | BW | ID=gene-HOXC8;Dbxref=GeneID:103348281;Name=HOXC8;gbkey=Gene;gene=HOXC8;gene_biotype=protein_coding |
| 4 | 36704911 | 38161753 | QTL2_95%CI | BW | ID=gene-HOXC6;Dbxref=GeneID:100339058;Name=HOXC6;gbkey=Gene;gene=HOXC6;gene_biotype=protein_coding |
| 4 | 36704911 | 38161753 | QTL2_95%CI | BW | ID=gene-HOXC5;Dbxref=GeneID:103348282;Name=HOXC5;end_range=37817354,.;gbkey=Gene;gene=HOXC5;gene_biotype=protein_coding;partial=true |
| 4 | 36704911 | 38161753 | QTL2_95%CI | BW | ID=gene-HOXC4;Dbxref=GeneID:100339303;Name=HOXC4;gbkey=Gene;gene=HOXC4;gene_biotype=protein_coding;partial=true |
| 4 | 36704911 | 38161753 | QTL2_95%CI | BW | ID=gene-LOC103348283;Dbxref=GeneID:103348283;Name=LOC103348283;gbkey=Gene;gene=LOC103348283;gene_biotype=lncRNA |
| 4 | 36704911 | 38161753 | QTL2_95%CI | BW | ID=gene-LOC103348284;Dbxref=GeneID:103348284;Name=LOC103348284;gbkey=Gene;gene=LOC103348284;gene_biotype=lncRNA |
| 4 | 36704911 | 38161753 | QTL2_95%CI | BW | ID=gene-LOC108176648;Dbxref=GeneID:108176648;Name=LOC108176648;gbkey=Gene;gene=LOC108176648;gene_biotype=lncRNA |
| 4 | 36704911 | 38161753 | QTL2_95%CI | BW | ID=gene-SMUG1;Dbxref=GeneID:100352573;Name=SMUG1;gbkey=Gene;gene=SMUG1;gene_biotype=protein_coding |
| 4 | 36704911 | 38161753 | QTL2_95%CI | BW | ID=gene-TRNAK-CUU-2;Dbxref=GeneID:108176713;Name=TRNAK-CUU;gbkey=Gene;gene=TRNAK-CUU;gene_biotype=tRNA |
| 4 | 36704911 | 38161753 | QTL2_95%CI | BW | ID=gene-CBX5;Dbxref=GeneID:100339807;Name=CBX5;gbkey=Gene;gene=CBX5;gene_biotype=protein_coding |
| 4 | 36704911 | 38161753 | QTL2_95%CI | BW | ID=gene-HNRNPA1;Dbxref=GeneID:100340058;Name=HNRNPA1;gbkey=Gene;gene=HNRNPA1;gene_biotype=protein_coding |
| 4 | 36704911 | 38161753 | QTL2_95%CI | BW | ID=gene-NFE2;Dbxref=GeneID:100340311;Name=NFE2;gbkey=Gene;gene=NFE2;gene_biotype=protein_coding |
| 4 | 36704911 | 38161753 | QTL2_95%CI | BW | ID=gene-COPZ1;Dbxref=GeneID:100352831;Name=COPZ1;gbkey=Gene;gene=COPZ1;gene_biotype=protein_coding |
| 4 | 36704911 | 38161753 | QTL2_95%CI | BW | ID=gene-GPR84;Dbxref=GeneID:100353087;Name=GPR84;gbkey=Gene;gene=GPR84;gene_biotype=protein_coding |
| 4 | 36704911 | 38161753 | QTL2_95%CI | BW | ID=gene-ZNF385A;Dbxref=GeneID:100353338;Name=ZNF385A;gbkey=Gene;gene=ZNF385A;gene_biotype=protein_coding |

| | | | | | |
|---|---|---|---|---|---|
| 4 | 36704911 | 38161753 | QTL2_95%CI | BW | ID=gene-ITGA5;Dbxref=GeneID:100009172;Name=ITGA5;gbkey=Gene;gene=ITGA5;gene_biotype=protein_coding |
| 12 | 148360582 | 148642693 | QTL3_95%CI | BW | ID=gene-LOC108177494;Dbxref=GeneID:108177494;Name=LOC108177494;gbkey=Gene;gene=LOC108177494;gene_biotype=protein_coding;partial=true;start_range=.,148368830 |
| 12 | 148360582 | 148642693 | QTL3_95%CI | BW | ID=gene-TULP4;Dbxref=GeneID:100339243;Name=TULP4;gbkey=Gene;gene=TULP4;gene_biotype=protein_coding |
| 12 | 148360582 | 148642693 | QTL3_95%CI | BW | ID=gene-TMEM181;Dbxref=GeneID:100355801;Name=TMEM181;gbkey=Gene;gene=TMEM181;gene_biotype=protein_coding;partial=true |
| 11 | 64270630 | 65432659 | QTL1_95%CI | CVDFW | ID=gene-TMEM267;Dbxref=GeneID:100339987;Name=TMEM267;gbkey=Gene;gene=TMEM267;gene_biotype=protein_coding |
| 11 | 64270630 | 65432659 | QTL1_95%CI | CVDFW | ID=gene-C11H5orf34;Dbxref=GeneID:100342453;Name=C11H5orf34;gbkey=Gene;gene=C11H5orf34;gene_biotype=protein_coding;partial=true |
| 11 | 64270630 | 65432659 | QTL1_95%CI | CVDFW | ID=gene-PAIP1;Dbxref=GeneID:100340240;Name=PAIP1;gbkey=Gene;gene=PAIP1;gene_biotype=protein_coding |
| 11 | 64270630 | 65432659 | QTL1_95%CI | CVDFW | ID=gene-LOC103349617;Dbxref=GeneID:103349617;Name=LOC103349617;gbkey=Gene;gene=LOC103349617;gene_biotype=lncRNA |
| 11 | 64270630 | 65432659 | QTL1_95%CI | CVDFW | ID=gene-NNT;Dbxref=GeneID:100341011;Name=NNT;gbkey=Gene;gene=NNT;gene_biotype=protein_coding |
| 11 | 64270630 | 65432659 | QTL1_95%CI | CVDFW | ID=gene-FGF10;Dbxref=GeneID:100342802;Name=FGF10;gbkey=Gene;gene=FGF10;gene_biotype=protein_coding |
| 7 | 138910411 | 139598335 | QTL1_95%CI | DFW | ID=gene-LOC100351153;Dbxref=GeneID:100351153;Name=LOC100351153;gbkey=Gene;gene=LOC100351153;gene_biotype=protein_coding |
| 7 | 138910411 | 139598335 | QTL1_95%CI | DFW | ID=gene-SATB2;Dbxref=GeneID:100351405;Name=SATB2;gbkey=Gene;gene=SATB2;gene_biotype=protein_coding |
| 11 | 64944841 | 64952273 | QTL2_95%CI | DFW | ID=gene-NNT;Dbxref=GeneID:100341011;Name=NNT;gbkey=Gene;gene=NNT;gene_biotype=protein_coding |
| 11 | 64944841 | 64952273 | QTL2_95%CI | DFW | ID=gene-FGF10;Dbxref=GeneID:100342802;Name=FGF10;gbkey=Gene;gene=FGF10;gene_biotype=protein_coding |
